# Sleep duration and brain structure – phenotypic associations and genotypic covariance

**DOI:** 10.1101/2022.02.15.480501

**Authors:** Anders M. Fjell, Øystein Sørensen, Yunpeng Wang, Inge K. Amlien, William F.C. Baaré, David Bartrés-Faz, Lars Bertram, Carl-Johan Boraxbekk, Andreas M. Brandmaier, Ilja Demuth, Christian A Drevon, Klaus P. Ebmeier, Paolo Ghisletta, Rogier Kievit, Simone Kühn, Kathrine Skak Madsen, Athanasia M Mowinckel, Lars Nyberg, Claire E. Sexton, Cristina Solé-Padullés, Didac Vidal-Piñeiro, Gerd Wagner, Leiv Otto Watne, Kristine B. Walhovd

## Abstract

The question of how much sleep is best for the brain attracts scientific and public interest, and there is concern that insuficient sleep leads to poorer brain health. However, it is unknown how much sleep is sufficient and how much is too much. We analyzed 51,295 brain magnetic resonnance images from 47,039 participants, and calculated the self-reported sleep duration associated with the largest regional volumes and smallest ventricles relative to intracranial volume (ICV) and thickest cortex. 6.8 hours of sleep was associated with the most favorable brain outcome overall. Critical values, defined by 95% confidence intervals, were 5.7 and 7.9 hours. There was regional variation, with for instance the hippocampus showing largest volume at 6.3 hours. Moderately long sleep (> 8 hours) was more strongly associated with smaller relative volumes, thinner cortex and larger ventricles than even very short sleep (< 5 hours), but effect sizes were modest. People with larger ICV reported longer sleep (7.5 hours), so not correcting for ICV yielded longer durations associated with maximal volume. Controlling for socioeconomic status, body mass index and depression symptoms did not alter the associations. Genetic analyses showed that genes related to longer sleep in short sleepers were related to shorter sleep in long sleepers. This may indicate a genetically controlled homeostatic regulation of sleep duration. Mendelian randomization analyses did not suggest sleep duration to have a causal impact on brain structure in the analyzed datasets. The findings challenge the notion that habitual short sleep is negatively related to brain structure.

**Significance statement:** According to consensus recommendations, adults should sleep between 7 and 9 hours to optimize their health. We found that sleeping less than the recommended amount was associated with greater regional brain volumes relative to intracranial volume, and very short sleep was only weakly related to smaller volumes. Genetic analyses did not show causal effects of sleep duration on brain structure. Taken together, the results suggest that habitual short sleep is not an important contributor to lower brain volumes in adults on a group level, and that large individual dfferences in sleep need likely exist.

## Introduction

It is a worry that insufficient sleep could be a pervasive negative factor for physical, mental and cognitive health^1–4^. Lack of sleep is also a concern for brain health, and older adults with sleep problems have higher probability of developing Alzheimer’s Disease (AD) and other dementias^5–10^. It has been argued that 15% of AD in the population may be attributed to sleep problems^11^, and studies have reported that long^12^ and short^13^ sleep duration is associated with increased risk of cognitive decline. Still, sleep was not included among 12 potentially modifiable risk factors for dementia in the Lancet Commission on dementia prevention^14^. It is not clear whether a causal relationship between variations in habitual sleep duration and brain health exists. Even if sleep *is* essential, we do not know how much sleep is necessary for optimal brain health. Expert recommendations exist for sleep duration and overall health^15–19^ but not for brain health specifically. The aim of the present study was to provide estimates of self-reported sleep duration associated with the largest regional brain volumes, thickest cortex and smallest ventricles across adult life. Further, we used genetic information to address causality of associations between sleep duration and brain structure.

Important aspects of brain health can be measured by structural MRI, which is sensitive to brain changes in aging^20^ and a range of clinical conditions such as AD^21^. Global brain volume is consistently related to higher general cognitive function^22^, and volumetric loss in specific regions has been shown to be associated with reduction of functions such as memory^23^. Effect sizes are modest, but it is established that in adulthood, larger regional brain volumes, thicker cortex and smaller ventricles are associated with higher cognitive function and less risk for cognitive decline^24^. Hence, if insufficient everyday sleep has detrimental effects on the brain, it is likely that we will detect these by brain morphometric analyses. Still, previous research has not yet convincingly established that self-reported sleep duration is related to brain morphometry in healthy adults. We reviewed 18 studies reporting relationships between self-reported sleep duration and at least one morphometric brain measure in adult participants without specific sleep disorders (Table 1). One cross-sectional study found shorter sleep duration to be related to smaller brain volume ^25^, one found longer duration to be related to small volume ^26^, and a third reported a quadratic relationship where shorter and longer durations were related to smaller hippocampal volumes ^27^. The remaining studies reported no significant relationship ^27–34^. Of six longitudinal studies, one found sleep duration shorter and longer than seven hours to be associated with more frontotemporal gray matter loss^35^, one found duration > 9 hours to be associated with smaller hippocampal subfields^36^, and four studies did not report any significant relationships ^27,37,38^. Hence, despite a growing body of literature, the relationship between sleep duration and brain structure remains unclear. Furthermore, no attempts have yet been made to calculate which sleep duration is associated with the largest regional brain volumes, a proxy for brain health^21^. This lack of evidence is probably partly due to insufficient statistical power and too scarce sampling of very short and long sleep durations. For instance, in a study of 28,000 participants, faster cognitive decline was observed in individuals sleeping 4 hours or less or 10 hours or more each night, compared to a reference group sleeping 7 hours^39^. However, no relationship between sleep duration and global cognitive decline was observed between these extreme intervals. We previously reported that in 21,000 participants from the UK Biobank, variations within the range of 5–9 hours of sleep were not related to hippocampal volume, whereas shorter and longer durations were associated with smaller volumes ^27^. Thus, it is possible that associations between sleep duration and brain structure exist only outside the normal ranges of sleep, but addressing this question requires very large samples.

**Table 1.** Reviewed studies of sleep duration – brain structure. X: This sample is a subsample of Fjell et al. 2019. Y: This is a longitudinal subsample of the Fjell et al. 2019 study above UKB: UK Biobank imaging study. LBC: Lothian Birth Cohort study. VBM: Voxel-based morphometry. ROI: Regions of interest. SPM: Statistical parametric mapping. TBV: Total Brain Volume. TGMV: Total Gray Matter Volume. Fjell et al. 2019 is cited twice for the cross-sectional results, since the study reports analyses from two independent samples. The sample in Fjell et al. 2020 is largely overlapping the longitudinal part of Fjell et al. 2019.

| Study | N | Sample | Main result and method |
| --- | --- | --- | --- |
| **Cross-sectional** |  |  |  |
| Alperin et al., 2019 | 69 | Older | Duration correlated negatively with hippocampal and cortical volume and thickness reductions. ROI, FreeSurfer. |
| Branger et al., 2016 | 51 | Older | No relationship between sleep duration and GM volume. VBM, SPM |
| Grau-Rivera et al., 2020 | 366 | Middle-aged/ older | Insomnia symptoms associated with lower GM volume, not surviving corrections. VBM, SPM. |
| Kim et al., 2021 | 1353 | Middle-aged/ older | No relationship in absence of sleep apnea. ROI, custom. |
| Lim et al., 2016 | 141 | Older | No relationship to volume in any of 68 cortical regions. ROI, FreeSurfer. |
| Stoffers et al. 2012 | 65 | Young/ middle-aged | Early morning awakenings negatively related to orbitofrontal volumes measured, not for initiating or maintain sleep. VBM, SPM. |
| Zitser et al., 2020 | 613 | Middle-aged/ older | No relationship between global GM volume and sleep duration over several years. VBM, FSL. |
| Fjell et al., 2019 | 3105 | Adult lifespan | No relationship between duration and hippocampal volume. ROI, FreeSurfer. |
| Fjell et al., 2019 | 21000 | Middle-aged/ older (UKB) | ≤ 5 hours and ≥ 9 hours related to smaller hippocampal volume. ROI, FreeSurfer. |
| Aribisala et al., 2020 | 457 | Older (LBC) | Nighttime sleep duration not associated with total brain volume. ROI, custom. |
| Sabeti et al. 2018 | 79 | Older (90+) | No relationship between sleep duration and hippocampal volume. ROI, custom. |
| Westwood et al., 2017 | 2457 | Older (incl dementia) | Long sleep (> 9 hours) negatively associated with total cerebral brain volume. ROI, custom. |
| **Longitudinal** |  |  |  |
| Sexton et al. 2014* | 147 | Adult lifespan | No relationship between duration and volume reduction. Vertex-wise, ROI, FreeSurfer. |
| Lo et al., 2014 | 119 | Older | No relationship between duration and change in total cerebral, <u>inferior</u> or superior frontal volume, pos relationship with ventricular expansion. ROI, FreeSurfer. |
| Spira et al., 2016 | 122 | Older | Sleep < 7 hours and > 7 hours associated with frontotemporal loss. Vertex-wise, FreeSurfer. |
| Fjell et al., 2019 | 1299 | Adult lifespan | No relationship with hippocampal volume loss. ROI, FreeSurfer. |
| Fjell et al., 2020<sup>y</sup> | 2205 | Adult lifespan | No relationship with thickness change. Vertex-wise, FreeSurfer. |
| De Looze et al., 2022 | 417 | Older | ≥ sleep associated with smaller hippocampal subfields |

In the present study we combined cross-sectional and longitudinal data from the Lifebrain consortium^40^ and tested the relationships between self-reported sleep duration and brain structure using 51,295 magnetic resonance imaging (MRI) brain scans from 47,029 participants across the adult life (20–89 years). We calculated the sleep duration associated with the most favorable brain structural properties across the observed age range and estimated how much brain structural deviations could be expected as a function of less or more sleep than this duration. In line with previous research on brain structure and cognitive function in adulthood and aging, larger regional brain volumes, smaller ventricles and thicker cortex were seen as favorable^24^. We systematically tested the influence of relevant somatic, mental and societal variables. Further, twin and genome wide association studies (GWAS) have shown heritability and polygenic influences on sleep duration^41–47^. Pleiotropy between genes for sleep duration and other conditions that may directly or indirectly be related to brain health is possible^41,43,48^. To test this hypothesis, we used Mendelian randomization analyses to assess the bidirectional causality of sleep duration – brain structure relationships.

## Results

Associations were tested by use of Generalized Additive Mixed Models (GAMM) run in R^49^. Code, detailed model statistics, complementary results and exact sample size for each sub-analysis are presented in supplemental information (SI). Mean self-reported sleep duration per night as a function of age is shown in Figure 1, imposed on the US National Sleep Foundation (NSF) recommendations^15^. The average sleep duration was relatively stable around 7 hours across the lifespan, and while significantly related to age (F = 33.1, p < 2e^16^), age explained minute variance (R^2^ = 0.006). The average reported sleep durations were at or below the lower recommended limits at most ages.

**Figure 1.**
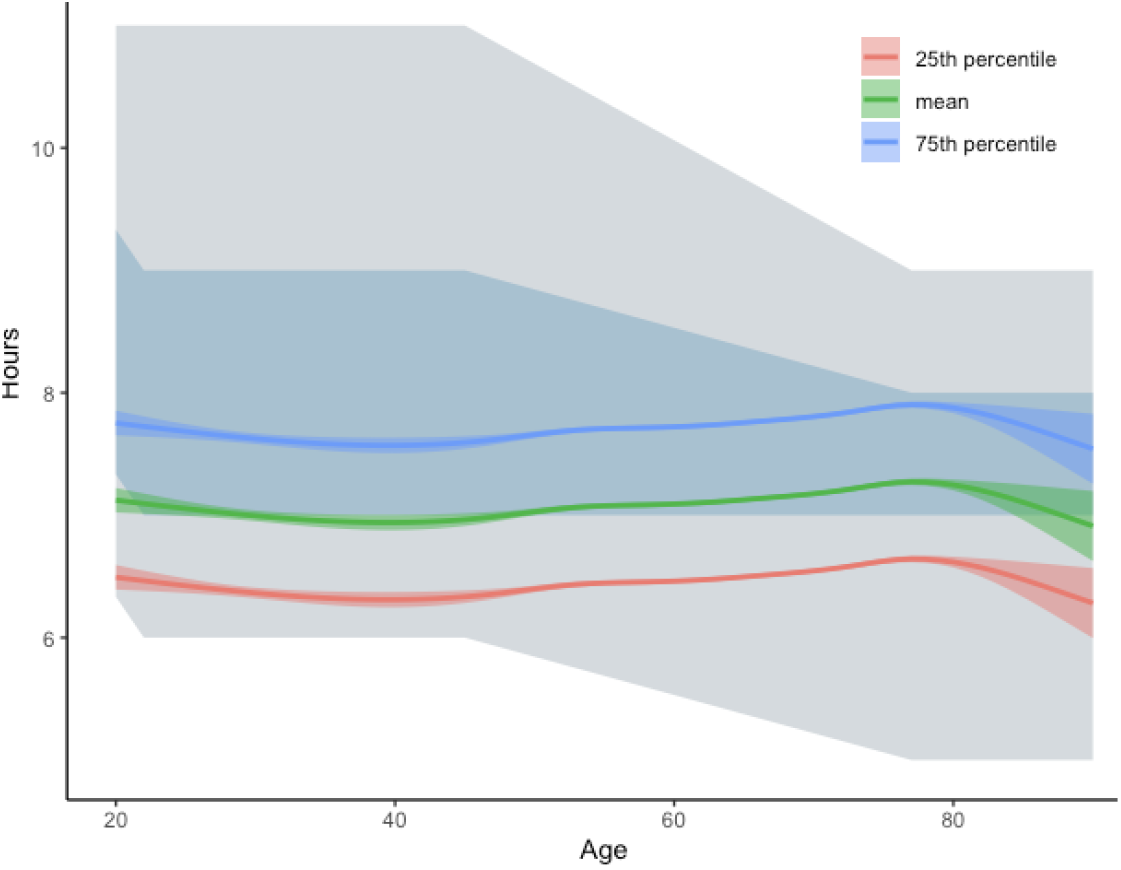
Self-reported sleep duration superimposed on the recommended sleep intervals from the National Sleep Foundation. Blue/ gray area depicts recommended sleep interval (blue: “recommended”, gray: “may be appropriate”). The green line shows average self-reported sleep in this study, the blue and red show 75^th^ and 25^th^ percentiles, respectively. The shaded area around the curves shows 95% confidence interval.

### Estimation of sleep duration - brain structure associations

To estimate the sleep duration associated with maximum subcortical volume and cortical thickness, and smallest ventricles, we ran three GAMM models: 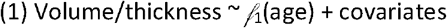 (sex, site, estimated intracranial volume [ICV]), 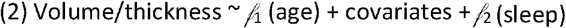, 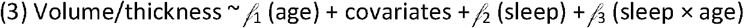. Smooth functions 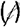 were used for age, sleep and sleep × age to allow non-linear relationships between the predictors and brain volume. A likelihood ratio test was used to select between models. The models provided the best fits taking both cross-sectional and longitudinal observations into account. We have previously reported on change relationships specifically^50^. 11 subcortical regions, four segments of the corpus callosum (CC), total ventricular volume and total gray matter volume, which is the sum of cortical, subcortical and cerebellar gray matter, were included. ICV was tested in a separate analysis and was additionally included as covariate of no interest in all volumetric analyses. The cortical analyses were focused on thickness, which changes considerably with age^50–52^. 32 cortical regions from the Desikan-Killiany parcellation scheme^53^ were included, visualized using ggseg^54^. The results were entered into a to a K-means cluster analysis. Inspection of the scree plot showed that three clusters represented a reasonable solution, with the regions within each cluster showing more similar sleep-thickness relationships than regions outside the cluster. The clusters are shown in Figure 2, left panel (see SI Cluster analysis for details). One cluster (#1) covered posterior medial cortices, superior parietal and caudal anterior cingulate cortex. The second and largest cluster (#2) included most of the lateral surface and superior frontal cortex. The third cluster (#3) included the rest of the lateral cortex, i.e. insula and pars opercularis, and medial regions such as medial orbitofrontal cortex, rostral anterior cingulate, posterior cingulate, and parahippocampal gyrus.

**Figure 2.**
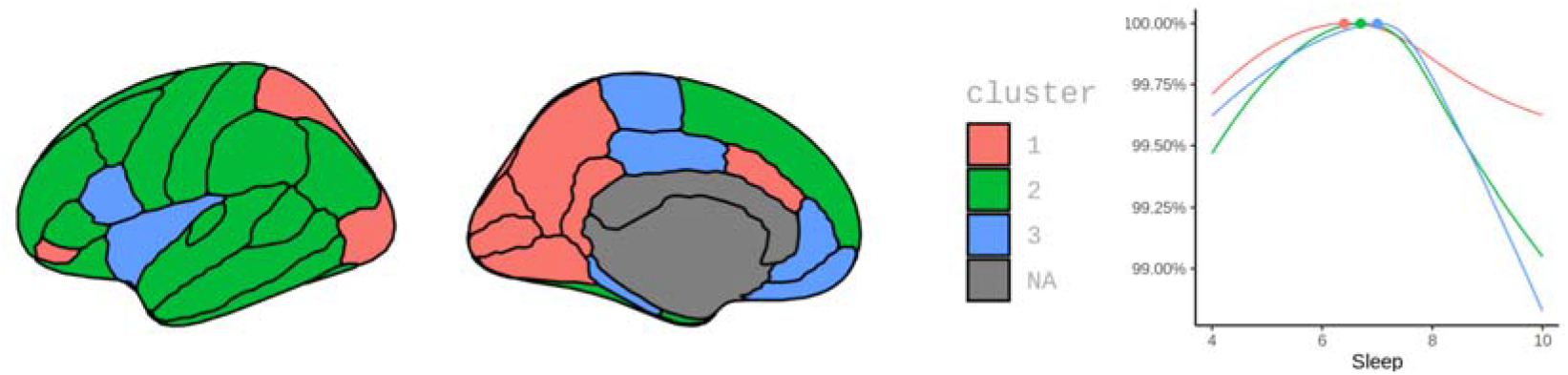
Cortical clusters. Left panel: Clusters of regions showing similar thickness – sleep duration relationships. Right panel: Thickness in each cluster as a function of sleep duration. 100% is the maximum thickness, as illustrated by the colored dots. X-axis indicates reported number of hours of sleep.

For each variable, the analyses yielded a sleep duration value associated with the maximum (minimum for ventricles) volume or thickness. The results are presented in Table 2. Together with the corresponding confidence intervals (CI), these were entered into a meta-analysis to find the sleep duration associated with the most favorable outcome across all the brain regions. We excluded total gray matter volume since this variable is a summary of other included variables. Corpus callosum structures were excluded because, with one exception, their estimated sleep duration at maximum volume could not be defined since the sleep-volume relationship was monotonous. The three cortical clusters and the subcortical regions were weighted such that cortex and subcortex contributed equally to the meta-analytic fit. For the ventricles, we estimated number of hours of sleep corresponding to minimal volume, and for all other regions we estimated the sleep duration that was associated with maximum volumes/ thickness.

**Table 2.**
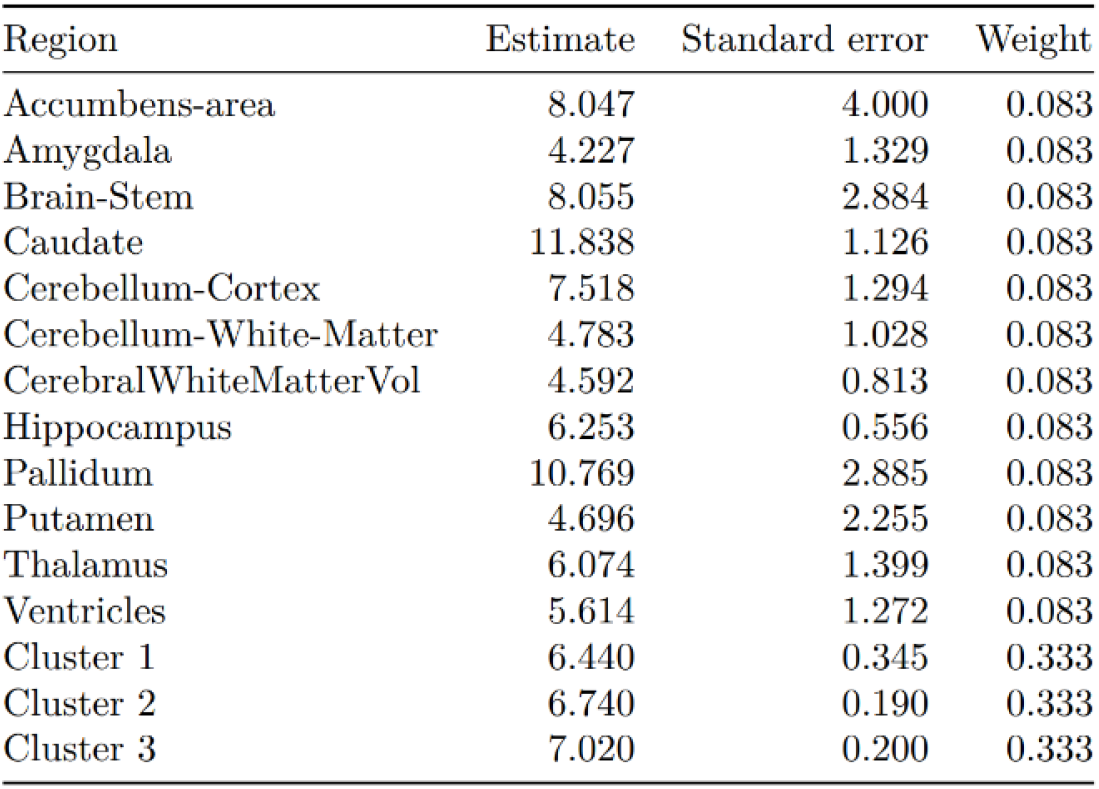
Estimated sleep duration associated with maximum (minimum for ventricles) volume or thickness. Cluster 1, 2 & 3 refers to the cortical thickness clusters in Figure 2.

| Region | Estimate | Standard error | Weight |
| --- | --- | --- | --- |
| Accumbens-area | 8.047 | 4.000 | 0.083 |
| Amygdala | 4.227 | 1.329 | 0.083 |
| Brain-Stem | 8.055 | 2.884 | 0.083 |
| Caudate | 11.838 | 1.126 | 0.083 |
| Cerebellum-Cortex | 7.518 | 1.294 | 0.083 |
| Cerebellum-White-Matter | 4.783 | 1.028 | 0.083 |
| CerebralWhiteMatterVol | 4.592 | 0.813 | 0.083 |
| Hippocampus | 6.253 | 0.556 | 0.083 |
| Pallidum | 10.769 | 2.885 | 0.083 |
| Putamen | 4.696 | 2.255 | 0.083 |
| Thalamus | 6.074 | 1.399 | 0.083 |
| Ventricles | 5.614 | 1.272 | 0.083 |
| Cluster 1 | 6.440 | 0.345 | 0.333 |
| Cluster 2 | 6.740 | 0.190 | 0.333 |
| Cluster 3 | 7.020 | 0.200 | 0.333 |

**Table 3.**
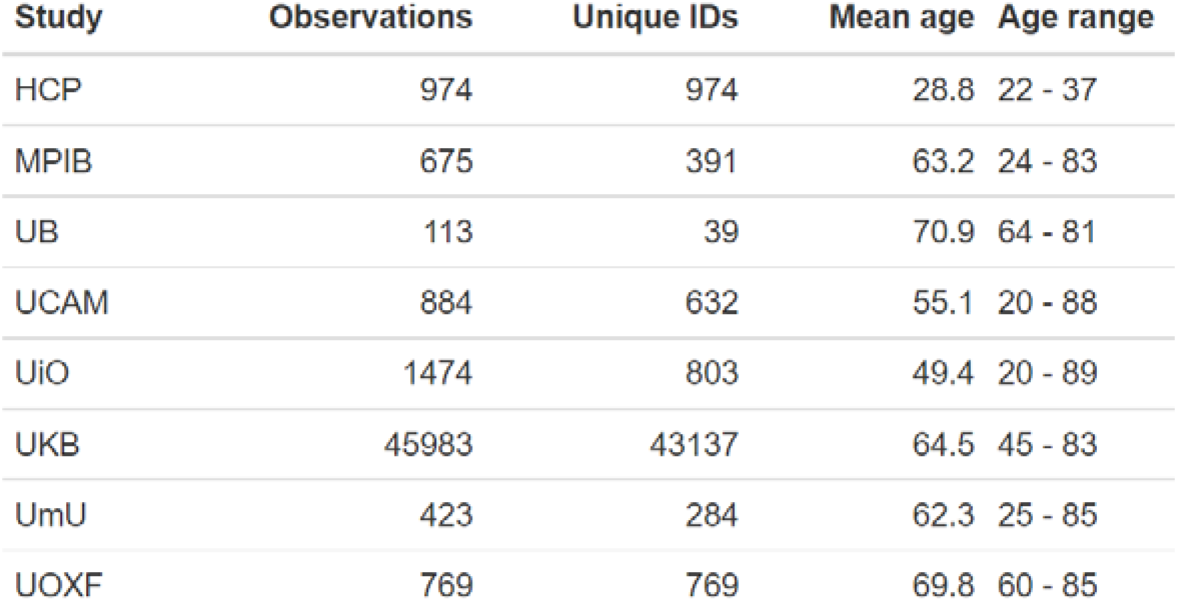
Sample origins. HCP: Human Connectome Project, MPIB: Max Planck Institute Berlin (BASE-II), UB: University of Barcelona, UCAM: University of Cambridge (CamCAN), UiO: University of Oslo, UKB: UK Biobank, UmU: University of Umea (Betula), UOXF: Oxford University (Whitehall-II)

Random effects analysis was used, as we did not expect each region to have the same sleep duration associated with maximum volume/ thickness. To be able to compare estimates across all regions, we used the model with a smooth term for sleep and no age interaction for all subcortical regions, since a significant age-interaction was found for the ventricles only. The estimates and standard errors were computed by sampling 5000 Monte Carlo samples from the empirical Bayes posterior distribution of the model for each region, constraining the number of hours of sleep to be between 4 and 12.

Detailed results are presented in SI Meta-analysis. The analysis showed that sleep duration of 6.8 hours was associated with maximum subcortical volume, smallest ventricles and thickest cortex. The critical values were 5.7 and 7.9 hours, as defined by the 95% CI. The variability across age was small while variability across regions was considerable. Thus, in the following we present results for different regions, additional covariates and genetic influences.

### Cortical thickness

The cortical analyses focused on thickness, and ICV was not included as covariate. For completeness, results for area and volume are presented in SI Cluster analysis. For 27 of the 32 regions, a significant relationship between sleep duration and thickness was found after false discovery rate (FDR)-correction (critical p-value = .0041), the exceptions being pericalcarine sulcus, lingual gyrus, lateral occipital cortex, isthmus cingulate and cuneus. Maximum cortical thickness as a function of sleep duration for each cluster is plotted in Figure 2, right panel. 6.4 (CI 5.7, 7.1), 6.7 (CI 6.3, 7.1) and 7.0 (CI 6.4, 7.2) hours were associated with maximum thickness for clusters 1, 2 and 3, respectively. Cluster #1 was only weakly related to sleep duration, with 4 and 10 hours of sleep being associated with about 0.25% thinner cortex. For clusters #2 and #3, shorter and longer sleep was associated with thinner cortex, but the negative associations of longer sleep were numerically larger than the negative associations of shorter sleep: about 1% thinner cortex at 10 hours vs. about 0.5% at 4 hours.

In addition, we tested associations between sleep duration and cortical thickness by vertex-wise spatiotemporal linear mixed-effect models (LME)^55,56^. The hierarchical nature of the data - repeated measurements nested within participants - was accounted for using a random intercept term for subject ID. Surface results were corrected for multiple testing by 10,000 Z Monte Carlo simulations using p < .01 as cluster generation threshold and p < .05 as cluster threshold^57^. As we previously have seen that sleep-cortex relationships tend to be stronger with higher age, we ran separate analyses in young/ middle-aged (20-60 years) and older adults (>60 years), and for participants with below vs. above average reported sleep duration. Analyses were thus run in four groups: young/ middle-aged with below (n = 9464) vs. above (n = 6421) average sleep, and older adults with below (n = 17119) vs. above (n = 15053) average sleep. The main results are shown in Figure 3 (SI Cortical surface analyses for area and volume). The spatially strongest effects were found in the older adults sleeping above average, where longer sleep duration was associated with thinner cortex across most of the brain surface. There were positive associations between sleep duration and thickness in the younger and older below average sleep groups, and no effects in the young group sleeping above average.

**Figure 3.**
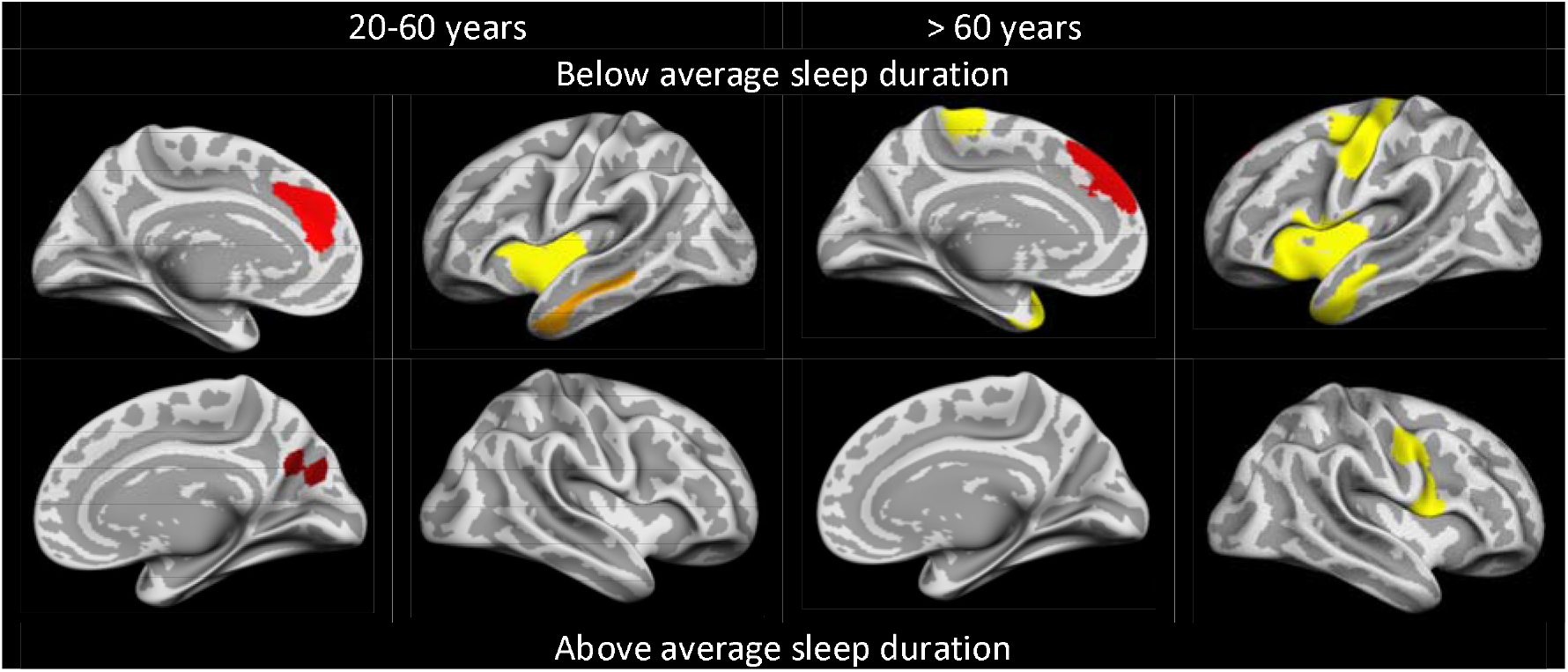
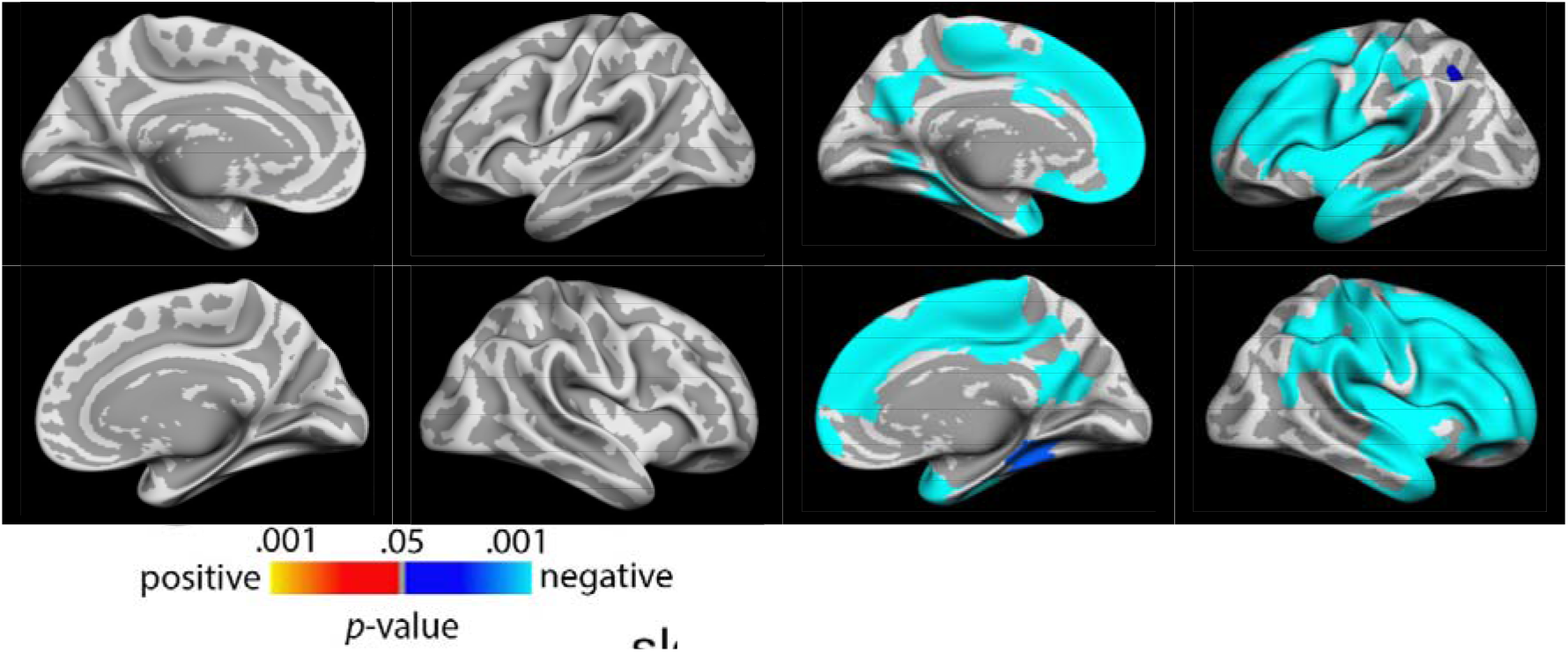
Associations between sleep duration and cortical thickness. Spatiotemporal LMEs were used to take advantage of both the longitudinal and cross-sectional data. Only clusters surviving corrections for multiple comparisons are shown. Warm colors indicate positive and cold colors indicate negative relationships between sleep duration and cortical thickness.

### Volumetric and subcortical regions

GAMMs as described above were run for each region. See SI Subcortical results for scatterplots, age trajectories and detailed model statistics. Correction for multiple comparisons was done by the Benjamini-Hochberg procedure for 19 regions. Main effects of sleep on volume were observed for brain stem (p < 5e^−7^), cerebellum cortex (p < 3.3e^−5^), cerebellum WM (p < 8.5^e-5^), cerebral WM (p < 1.2e^−5^), hippocampus (p < 2e^−16^), thalamus (p < 1.2e^−5^), total gray matter volume (p < 2e^−16^), ventricles (p < 1.5e^−3^) and ICV (p < 2e^−16^). Significant effects were also observed for segments of the corpus callosum: anterior (p < .005), central (p < 7.6e^−5^), middle anterior (p < 8.9e^−06^) and posterior (p = .016). Only for the ventricles was the sleep × age interaction model preferred (p = .0012). No significant main effect of sleep or sleep × age interactions were seen for accumbens, amygdala, caudate, pallidum and putamen.

Duration associated with maximal volume ranged from 6.3 hours for the hippocampus to 7.3 hours for posterior corpus callosum (Figures 4 and 5). More than 7.6 hours of sleep was associated with significantly smaller volume for all regions, and critical values for short sleep ranged from 4.0 hours (brain stem) to 7.1 hours (total gray matter volume). For all WM structures - cerebral WM (max volume at 4.0 hours [CI 4.0, 6.1]), cerebellar WM (4.5 hours [CI 4.0, 6.1]), and the corpus callosum (4.0 hours [CI 4, 4], except the posterior CC) - the relationships were close to monotonous negative, i.e., shorter sleep was related to larger volume. ICV was the MRI-derived measure for which maximal volume was associated with the longest sleep duration (7.5 hours, [CI 7.4, 7.6]). 10 hours of sleep was typically associated with about 1% smaller volume, with higher values for the hippocampus (1.5%) and the segments of the corpus callosum (1.5-3%). Sleeping 4 hours was associated with smaller deviations in volume, from 0% to 0.8%.

**Figure 4.**
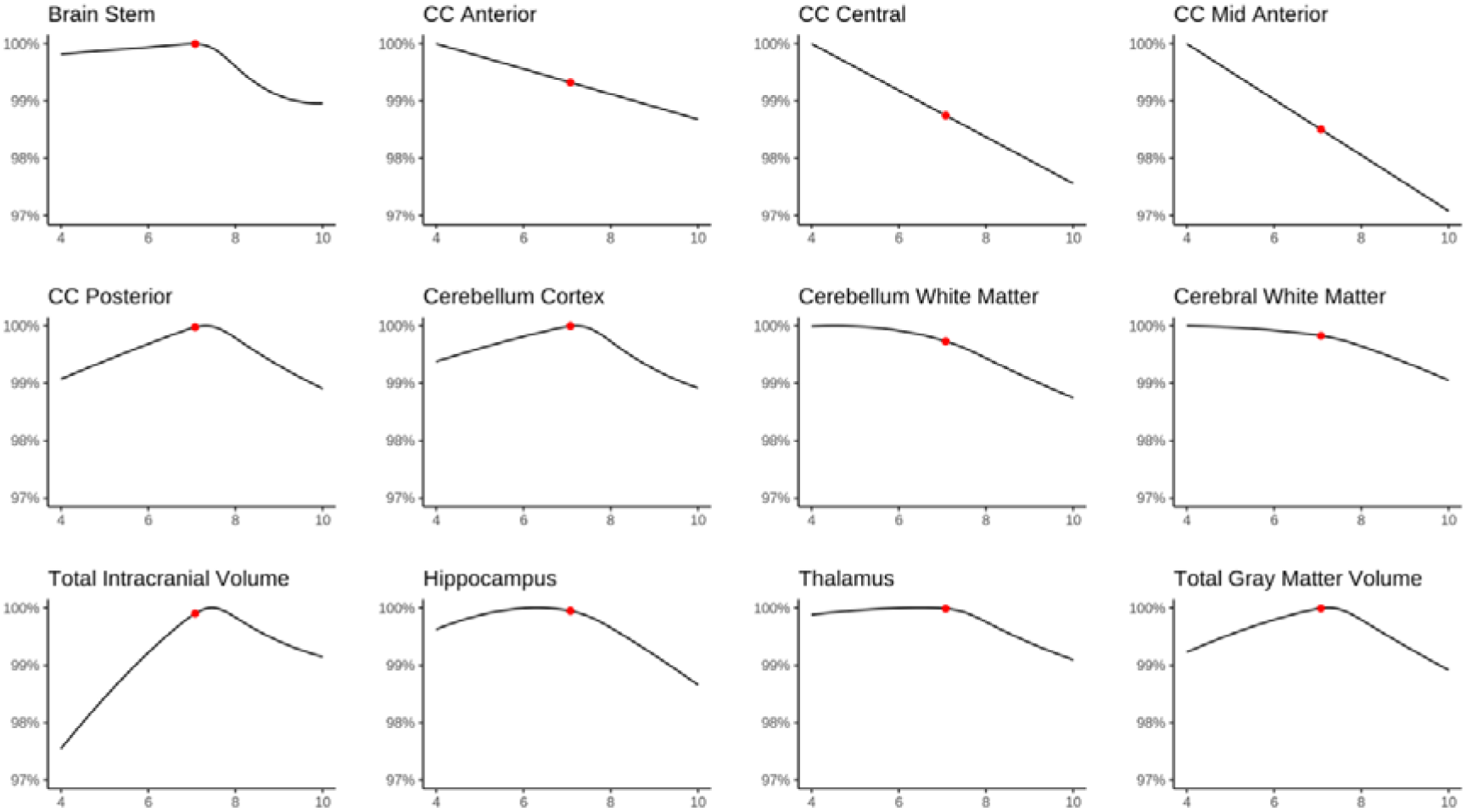
Subcortical and global volumes as a function of sleep duration. 100% is the maximum volume, and the scale shows volumetric deviations from the maximum volume as a function of sleep duration. The red dots show average reported sleep duration. Only regions significantly related to sleep duration are shown. All plots are corrected for age, sex and site and ICV. CC: Corpus callosum.

**Figure 5.**
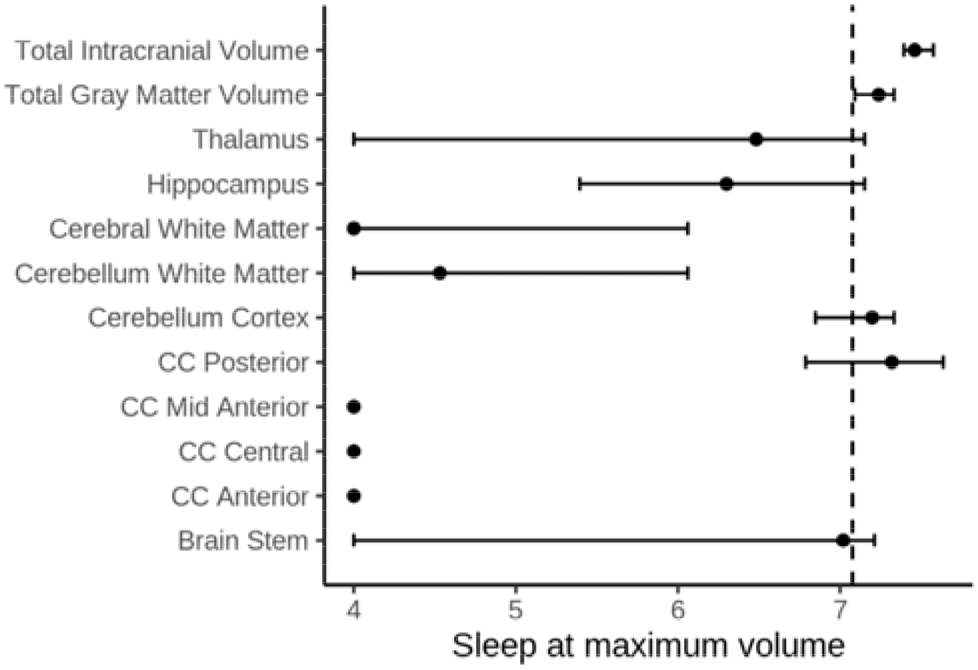
Sleep duration associated with maximum volume and 95% confidence intervals. The dotted line shows the average reported sleep duration. Only regions significantly related to sleep duration are shown.

For the ventricles, sleep duration and age interacted, as illustrated in Figure 6. In young and older adults, 7.0 hours or less was associated with the largest volume, and 10 hours was associated with the largest volumes during middle age. As large ventricular volume when controlling for ICV is considered to be negative, this suggests that short sleep was associated with beneficial outcome for middle aged participants, whereas long sleep was associated with beneficial outcome for young and older participants. This complex relationship was not expected and not easily interpretable.

**Figure 6.**
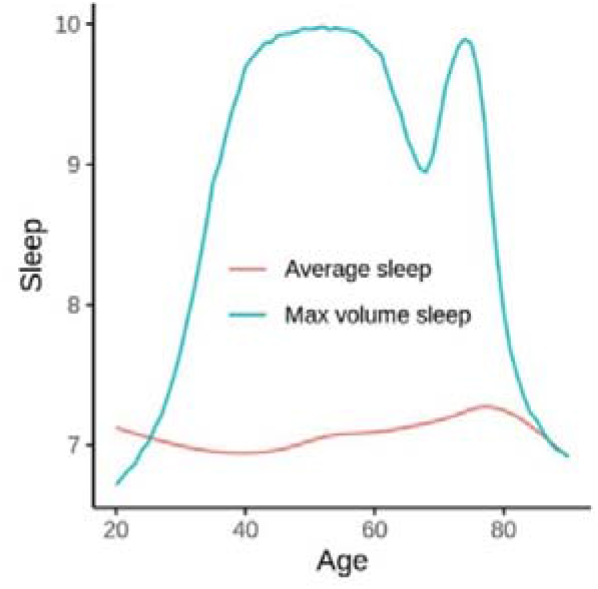
Sleep duration associated with maximum ventricular volume. The blue line shows how many hours of sleep (y-axis) was associated with the largest volume continuously across the age-range (x-axis). Average reported sleep across age is illustrated by the red line.

### Post hoc analyses – modelling covariates

We re-ran the volumetric analyses controlling for socioeconomic status (SES; income and education), body mass index (BMI) and depression symptoms (see SI Subcortical results). In the first set of analyses, we tested whether inclusion of each covariate affected the sleep-brain relationships. In a second set of analyses, we tested for interactions between sleep and each covariate. Controlling for the main effects of SES, no sleep duration-brain relationships changed from significant to not significant. Sleep-SES interaction effects did not survive corrections for multiple comparisons. Inclusion of BMI and depression symptoms as covariates had no notable effects on the results, and no significant interaction effects were found (See SI Subcortical results for details).

As seen above, ICV was the MRI-derived measure for which maximum volume was associated with the longest sleep duration, meaning that people with larger heads on average report longer sleep. Although there is no ground truth for what the most appropriate procedure is, ICV is usually controlled for in volumetric studies because regional structures scale with head size. Hence, we included ICV as covariate in our main volumetric analyses. However, since ICV showed a relationship with sleep duration, we re-ran the meta-analysis without controlling for ICV. This affected the results, yielding 7.38 hours (CI 6.43, 8.33) as the duration associated with maximal regional volume and cortical thickness and smallest ventricles. Thus, maximal relative volume is associated with shorter sleep duration than maximal absolute volume, since head size is associated with longer reported sleep duration.

### Polygenic scores, genetic correlations and Mendelian randomization

To determine the plausible direction of causality between brain structure and sleep duration, we performed a series of genetic analysis using cross-sectional data from UK Biobank. Hippocampus, total gray matter volume and ICV were chosen as regions of interest for the genetic analyses as they showed the typical inverted U-shaped relationship to sleep duration. For selection of participant, quality control procedures and genetic analyses, see SI Genetic analyses.

Samples were stratified into shorter (≤ 7 hours) and longer (>7 hours) sleepers, since different relationships was expected in these two groups. Three independent samples were used for GWAS: (1) participants with MRI (n=29,155), (2) participants sleeping < 7 hours without MRI (n=197,137), and (3) participants sleeping ≤ 7 hours without MRI (n=112,839). GWAS were performed independently for each trait in the corresponding sample. Manhattan and QQplots showing GWAS results are presented in SI Genetic analyses.

Single nucleotide polymorphism heritability (SNP-h^2^) was estimated by the linkage disequilibrium regression models^58^ for sleep duration in the short sleepers (h^2^= 0.045, se=0.0035) and long sleepers (h^2^ = 0.021, se=0.0047), hippocampal volume (h^2^ = 0.29, se=0.03), total gray matter volume (h^2^ = 0.22, se=0.03) and ICV (h^2^ = 0.35, se=0.03). For the further genetic analyses, age, sex and the top 10 genetic principal components were included as covariates. For hippocampal volume and total gray matter volume, ICV was included as an additional covariate. The genetic correlation for sleep duration was negative for the short vs. long sleepers (r_g_ = −.40, se = 0.10, p = 9.65e^−5^), showing that the genes related to longer sleep in the below average sleep duration group are related to shorter sleep in the above average sleep group.

Corresponding polygenic scores (PGS) were calculated for each variable in each group separately. PGSs for ICV (PGS-ICV) and total gray matter volume (PGS-TGV) were significantly associated with sleep duration in the short sleepers (PGS-ICV, t = 8.47, p_*fdr*_ = 2.4×10^−15^; PGS-TGV, t = 4.65, p_*fdr*_ = 3.28×10^−5^; Figure 7a, c and SI). PGS for sleep duration in the short sleepers was significantly related to ICV (t = 6.99, p*_fdr_* = 3.03×10^−11^, Figure 7b) and to a lesser extent with total gray matter volume (t = 2.69, p_*fdr*_ = 6.42×10^−2^, Figure 7d). No significant associations for other pairs of traits were identified.

**Figure 7.**
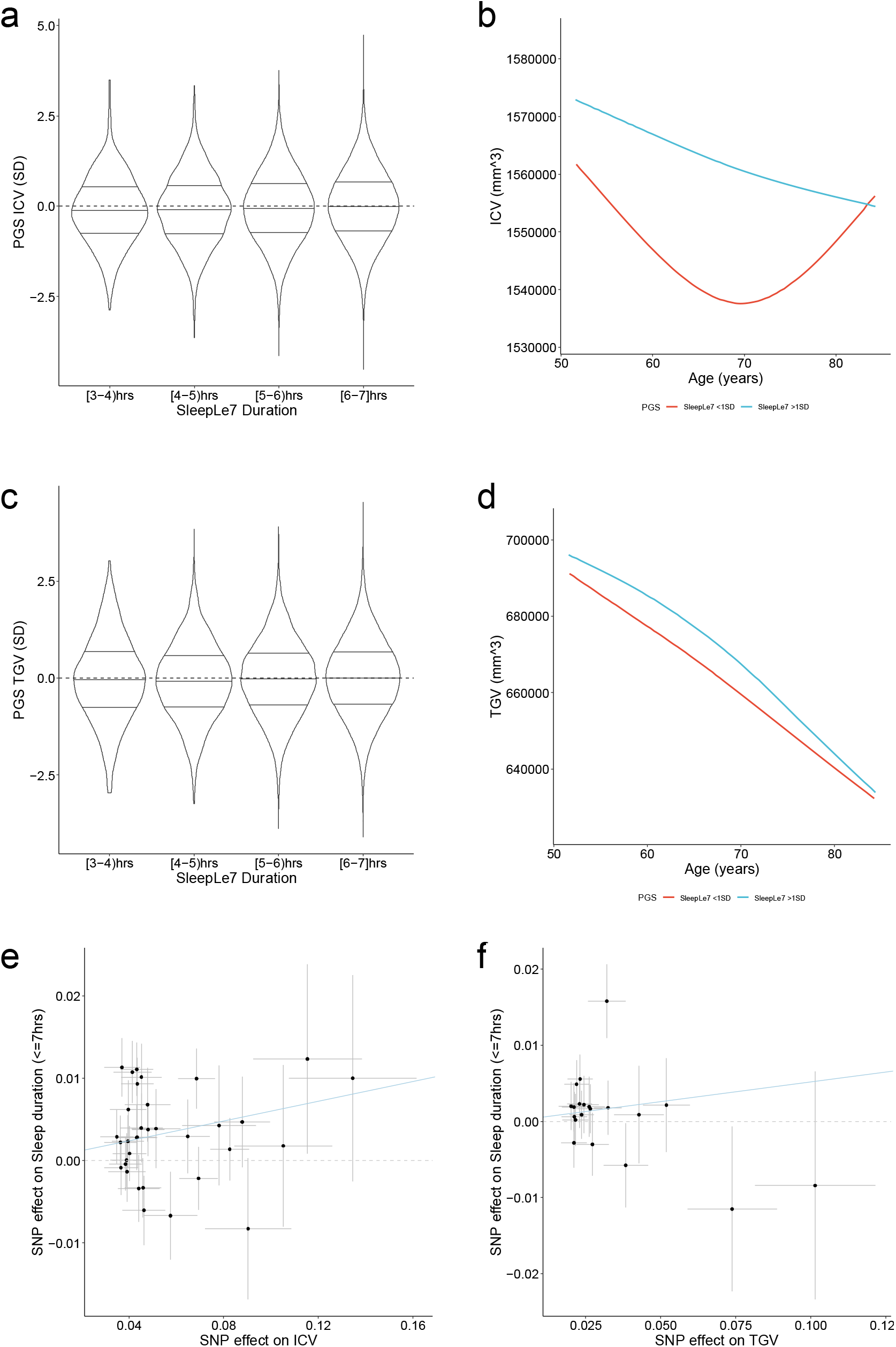
Genetic relations between sleep duration and brain structure. a. Distribution of PGS-ICV in different sleep duration strata among short sleepers. b. ICV for short sleepers (≤ 7 hours) with one standard deviation above (blue) and below (red) average PGS for sleep duration c. Distribution of PGS-TGV (total gray matter volume) in different sleep duration strata among short sleepers. d. Total gray matter volume (TGV) for short sleepers with one standard deviation above (blue) and below (red) average PGS for sleep duration. e. SNP effects on ICV (x axis) and sleep duration (y axis) for the short sleepers in units of sample standard deviation. f. SNP effects on total gray matter volume (x axis) and sleep duration (y axis) for the short sleepers in units of sample standard deviation.

We performed bidirectional Mendelian randomization analysis for each pair of traits. Among the 12 pairs, ICV showed a significant causal effect (inverse-variance weighted beta = 0.060, se = 0.017, p = 5.36×10^−4^) on sleep duration for the short sleepers (Figure 7e), and total gray matter volume showed a nominally significant negative effect for the long sleepers (beta = −0.35, se = 0.14, p = 0.012, uncorrected) (Figure 7f). Due to low heritability for sleep in the study, resulting in a weaker genetic instrument, we performed a robust MR analysis using robust adjusted profile score ^59^ for the direction from sleep to brain traits, but also did not to detect significant causal relationships for this directionality of effects. For details, see SI Genetic analyses, SI Genetics supplementary notes and SI Genetics supplementary tables.

## Discussion

Shorter sleep than currently recommended was not a consistent factor associated with brain structure across the adult lifespan, at least not as measured by relative regional brain volume, cortical thickness and ventricular volume. Rather, sleeping less than the recommended amount was associated with greater volumes and thickness. 6.8 hours of sleep was associated with the most favorable brain outcome, which is just below the lower limit of the current international recommendations^16–19^. Interestingly, this number was well aligned with the average reported sleep duration of about 7 hours, in line with the genetic analyses that indicated a possible homeostatic drive for sleep duration. The critical values for short and long sleep were 5.7 and 7.9 hours, respectively. This suggests that self-reported sleep up to 1.3 hours less than the lower limit of the recommendations is not necessarily associated with smaller regional volume or thinner cortex. The upper limit of 7.9 hours demonstrates that sleep durations well within the recommended range is clearly associated with less favorable volumetric brain outcomes. In fact, long sleep was more strongly associated with lower brain volumes and thinner cortex than short sleep. Although definite claims about causality cannot be made based on observational studies, we believe the present results make a strong case that short habitual sleep is not a prevalent cause of poorer brain health as indicated by structural brain measures in the samples studied here. In line with this conclusion, the Mendelian randomization analysis did not show causality from sleep to brain volume. These results are in line with findings of great individual differences in sleep need^60^.

### Sleep duration and the brain through adult life

Sleep duration is the most widely studied, best supported, and most straightforward sleep measure to address in relation to health^3^. It is also an aspect of sleep that to some degree may be modified by lifestyle choices. Average reported sleep was around 7 hours across the age range, which fits well with a recent meta-analysis of more than 1 million participants^61^. Seven hours represent the lower limits of the recommendations from both the NSF and the American Academy of Sleep Medicine and Sleep Research (AASM-SRS)^16–19^. It is noteworthy that according to our results, sleeping just below the recommended interval is associated with the most favorable brain outcomes. Even very short sleep was associated with larger regional volume and thicker cortex than moderately long sleep. The volumetric results, but not the thickness results, were obtained controlling for ICV. The genetic analyses showed that overlapping genes had opposite effects on sleep duration in below vs. above average sleepers. One interpretation is that there is a propensity towards optimal sleep duration for the brain. Still, there are substantial individual differences in sleep duration, which partly can be explained by genetic differences^41–47^. Nevertheless, the results suggest that optimal and average sleep are relatively well aligned in the present samples from mostly European countries.

Regarding the hypothesis that short sleep may exert a causal negative effect on brain health, the association between ICV and sleep duration is interesting. ICV was the MRI-derived measure that was most positively associated with sleep duration. As sleep has no causal effect on ICV in adults, this relationship must reflect other factors, and demonstrates that associations between sleep duration and MRI-derived volumes may reflect non-causal effects. The Mendelian randomization analyses also showed causal effects of ICV on sleep duration but not the inverse. Hence, people with larger heads on average report to sleep longer. This effect was removed from the estimated sleep duration – brain volume relationships by covarying for ICV. This was not done in the cortical thickness analyses, as thickness and ICV usually are weakly related^62^. Controlling for ICV removes the effect of global scaling, i.e. that regional brain volumes scale with head size. Since ICV is sometimes regarded as a proxy for maximal brain size, controlling for ICV yields regional volumes reflecting deviations from the expected based on head size. This will to a larger extent reflect volumetric alterations, not only offset effects. About 4000 longitudinal MRIs were included, which further causes brain changes to affect the results. Regardless, as most of the data were cross-sectional, we cannot readily distinguish between offset effects – such as those seen for ICV – and effects representing brain change. Still, we believe the combined results represent evidence against the notion that habitual short sleep causes smaller regional volumes and thinner cortex as detectable by MRI in the presently studied populations. It is still worth noting that when absolute volumes were used in the analyses, sleep duration associated with maximal volume was substantially longer. Hence, maximal relative volume – partly reflecting deviations from the individual’s largest life-time volume – is associated with shorter sleep duration than absolute volume.

Even moderately long sleep was more strongly related to smaller relative regional volumes, thinner cortex and larger ventricles than short sleep. Sleeping 4 hours was associated with at most 0.8% smaller regional brain volume. In contrast, sleeping for more than 7.6 hours was associated with significantly smaller volume or thinner cortex for all regions, with even shorter critical values for e.g. the hippocampus and the cerebral cortex. AASM-SRS proposed no upper limit^3^ on sleep duration whereas NSF recommended a maximum of 9 hours through most of adulthood, and 8 hours in older adults^15^. Our results show that 8 hours of sleep is associated with smaller relative regional brain volumes and thinner cortex throughout adult life. Previous research has established associations between long sleep and poorer brain^36^, cognitive^26,27,35,39,63^ and somatic health^64^, which often are ascribed to underlying comorbidities^3,15,64^. In the present analyses we attempted to address this by controlling for somatic (BMI), mental (symptoms of depression) and social (SES) factors. However, these had negligible effects on the relationships. Whether long sleep itself has negative impacts on the brain is not known, but the present observational results do show an association between moderately long sleep and smaller volumes and thinner cortex.

An important qualification, however, is that individual differences in sleep need likely exist, due to factors such as genotype and previous sleep history^60^. If deviations from an individuals’ sleep need leads to poorer brain health for that individual, this may not be picked up in our group analyses. Thus, we cannot from our results conclude that people should try to sleep 6.8 hours each night. What we can say is that people who report to sleep 6.8 hours tend to have the largest regional brain volumes and smallest ventricles relative to ICV, as well as the thickest cortex. Optimal sleep for an individual may deviate from this number in both directions. Intra-individual effects on the brain of changing sleep duration can be assessed in sleep deprivation studies. Unfortunately, experimental sleep deprivation does not resemble habitual variations in sleep duration, and the long-term consequences on the brain from of sleep deprivation, taking into account adaptations^60^, are not known.

### Caveats and limitations

First, self-reported sleep duration is not accurate, and may reflect several other aspects of sleep than duration only. There is no perfect way to measure sleep duration without disrupting routine^65^. Self-reports are only moderately correlated with actigraph measures^65,66^. However, although actigraph results often correlate highly with polysomnography^67^ (PSG), they tend to over-estimate duration^67–71^, and it is not known how well actigraphs perform outside a sleep lab setting. One study reported that the same genetic loci were related to sleep duration whether it was measured by actigraphs or self-reports^41^. The international recommendations for sleep duration were mostly based on studies involving self-report^3,15^, and self-reported sleep is the most relevant variable for clinical, public health and policy recommendations^3^. While acknowledging the limitations of self reports, we also believe it to be the most relevant measure in the present context. Second, we have not considered other potentially important aspects of self-reported sleep, such as subjective sleep quality. This can be relevant in relation to brain health, but was not considered in the present analyses as the focus of the study was sleep duration. Third, we studied morphometric brain measures only. Whereas other measures could show different sensitivity to sleep duration, such as white matter microstructure ^72^ or Aβ accumulation ^73^, brain morphometry is sensitive to normal and pathological brain changes^21^, and atrophy has consistently been identified as a factor governing age-related sleep changes^74^. Fourth, we have not considered cognitive function, for which different ranges of sleep duration may possibly be optimal. It is still likely that associations between sleep duration and ith cognitive performance or mental health are transient and reversable after restoritative sleep, whereas associations with brain structure may be more permanent. Fifth, the samples were not thoroughly screened for sleep disorders such as sleep apnea^75^. If individuals with sleep problems were included, this would probably not attenuate the relationships, and is therefore unlikely to explain the weak sleep-brain associations observed in the study. Finally, although some of the samples are population based, no MRI study is fully representative of the population from which it us sampled. Despite including studies from multiple European countries and the US, we can not exclude the possibility that other sleep-brain patterns exist in other populations from Europe or elsewhere.

### Conclusion

Short sleep does not seem to be associated with smaller regional brain volumes, thinner cortex or smaller ventricles in the present samples. Rather, sleeping less than the recommended amount was associated with greater regional brain volumes relative to intracranial volume and thicker cortex, and moderately long sleep showed a more negative association than even very short sleep.

## Supporting information

SI Genetic analyses

SI Genetics supplementary notes

SI Genetics supplementary tables

Si Meta analysis

SI Subcortical results

SI MRI Methods

SI Sample characteristics

SI Cortical clustering

SI Cortical surface analyses

## Acknowledgement

The Lifebrain project is funded by the EU Horizon 2020 Grant agreement number 732592 (Lifebrain). In addition, the different sub-studies are supported by different sources: LCBC: The European Research Council under grant agreements 283634, 725025 (to A.M.F.) and 313440 (to K.B.W.), as well as the Norwegian Research Council (to A.M.F., K.B.W.), The National Association for Public Health’s dementia research program, Norway (to A.M.F). Betula: a scholar grant from the Knut and Alice Wallenberg (KAW) foundation to L.N. Barcelona: Partially supported by a Spanish Ministry of Economy and Competitiveness (MINECO) grant to D-BF [grant number PSI2015-64227-R (AEI/FEDER, UE)]; by the Walnuts and Healthy Aging study (http://www.clinicaltrials.gov; Grant NCT01634841) funded by the California Walnut Commission, Sacramento, California; and an ICREA Academia 2019 award. BASE-II has been supported by the German Federal Ministry of Education and Research under grant numbers 16SV5537/ 16SV5837/ 16SV5538/ 16SV5536K/ 01UW0808/ 01UW0706/ 01GL1716A/ 01GL1716B, the European Research Council under grant agreement 677804 (to S.K.). Work on the Whitehall II Imaging Substudy was mainly funded by Lifelong Health and Well-being Programme Grant G1001354 from the UK Medical Research Council (“Predicting MRI Abnormalities with Longitudinal Data of the Whitehall II Substudy”) to K.E. The Wellcome Centre for Integrative Neuroimaging is supported by core funding from award 203139/Z/16/Z from the Wellcome Trust. Data were provided [in part] by the Human Connectome Project, WU-Minn Consortium (Principal Investigators: David Van Essen and Kamil Ugurbil; 1U54MH091657) funded by the 16 NIH Institutes and Centers that support the NIH Blueprint for Neuroscience Research; and by the McDonnell Center for Systems Neuroscience at Washington University. Part of the research was conducted using the UK Biobank resource under application number 32048.

## Materials and Methods

### Sample

Community-dwelling participants were recruited from multiple countries in Europe and the US. Some were convenience samples, whereas others were contacted on the basis of population registry information. All participants at age of majority gave written informed consent. All procedures were approved by a relevant ethical review board. For Lifebrain, approval was given by the Regional Ethical Committee for South Norway, and all sub-studies were approved by the relevant national review boards. For UKB, ethical approval was obtained from the National Health Service National Research Ethics Service (Ref 11/NW/0382).

In total, data from 47,039 participants (20.0-89.4 years) with information about sleep duration and MRI of the brain were included. For 3,910, two or more MRI examinations were available, yielding a total of 51,320 MRIs (mean follow-up interval 2.5 years, range 0.005-11.2, 26,811 female/ 24,509 male observations). Demographics of the samples are given in Table 1 and a brief description of each is given below (see SI Sample characteristics for details).

### Statistical analyses

ROI analyses were run in R version 4.0.0 ^76^, by use of Generalized Additive Mixed Models (GAMM) using the packages “gamm4” version 0.2-26^77^ and “mgcv” version 1.8-28 ^49^. The Desikan-Killiany parcellation included in FreeSurfer yields 34 regions, but the temporal and the frontal poles were excluded from analysis due to substantial noise in these regions. Volumetric outliers were defined by having a residual more than four times the magnitude of the residuals standard error in an analysis of age effects and removed from the analyses. Computer code can be found in SI in the relevant sections.

### Data availability

Data supporting the results of the current study are available from the PI of each sub-study on request, given appropriate ethical and data protection approvals. Contact information can be obtained from the corresponding authors. UK Biobank data requests can be submitted to http://www.ukbiobank.ac.uk.

