## Supplementary material for "Sleep duration and brain structure – phenotypic associations and genotypic covariance": SI Genetic analyses

### **Genome-Wide Association studies (GWAS) Using UK Biobank data**

We accessed the UK Biobank genetic data (<https://www.ukbiobank.ac.uk/about-biobank-uk/> released in February 2020 through the application no. 32048. Ethical approval of UK Biobank project was obtained from the National Health Service National Research Ethics Service (Ref 11/NW/0382) and all participants provided written informed consent. Among the 502,507 participants, 40,682 had undergone MRI and 487,409 had DNA genotypes. The detailed information on genotyping, imputation and quality controls was published by^1^. Before performing GWAS, we excluded participants who are not self-reported white-British (n=92 900), have relatives in the biobank (n=148 689), had been labelled as outliers in missingness or heterozygosity(n=968) or had conflicting self-reported vs. genetic sex (n=378) by UK Biobank team. Among the remaining 339696 participants, 125 800 reported sleep duration longer than 7 hours, and 213 331 reported sleep duration less or equal to 7 hours. 29 155 of these participants had undergone MRI. Genotypes of the UK Biobank participants were filtered by removing variants that are non-SNP, had a minor allele frequency less than 0.01, failed the Hardy-Weinberg equilibrium test (p<1e^-6^), or had large missingness (>0.05). In total, 9 153 887 autosomal SNPs were analyzed.

#### *GWAS for sleep duration among participants sleeping <= 7hours*

We performed GWAS for sleep durations for UKB participants who reported less than or equal to 7 hours of sleep and had not undergone MRI (n=197,137). The sleep duration was scaled by subtracting the mean and divided by the standard deviations in this subsample. A linear regression model was used to identify SNPs that were associated with scaled sleep durations. Age at baseline, sex and the top 10 genetic principal components (PC, obtained from UK Biobank) were included as covariates in the model. The PLINK2 command –glm was used for this analysis. A genomic inflation factor was estimated to be 1.03 based on the GWAS summary statistics, i.e., no discernable inflation of the associations was observed. The association quantile-quantile plot, and the Manhattan plot are shown below.

| 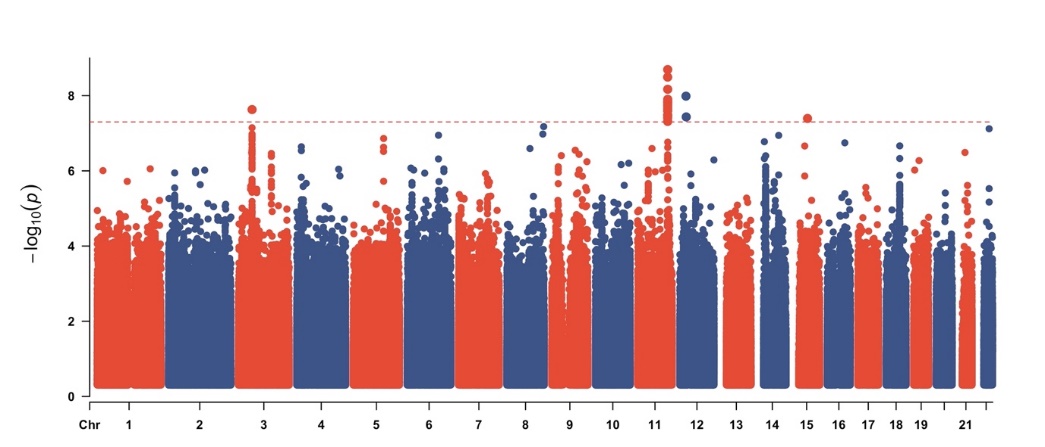 | 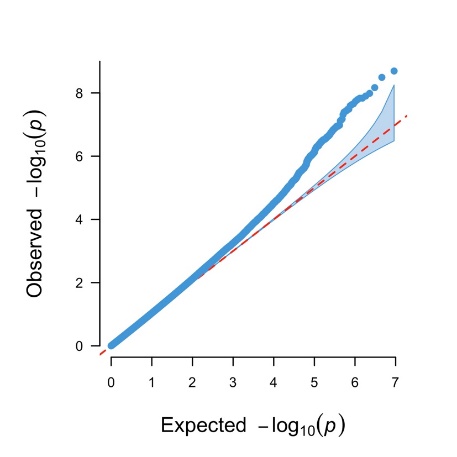 |
| --- | --- |

*Left panel: Manhattan plot of GWAS for sleep duration in participants sleeping ≤ 7 hours, with the dotted red line showing the genome-wide significance level (p < 5×10^-8^). Right panel: QQ plot of GWAS for sleep duration.*

#### *GWAS for sleep duration among participants sleeping > 7 hours*

We also performed GWAS for sleep durations for UKB participants who reported more than 7 hours of sleep and had not undergone MRI (n=112,839). The sleep duration was scaled by subtracting the mean and divided by the standard deviations in this subsample. A linear regression model was used for identified SNPs that associated with scaled sleep durations. Age at baseline, sex and the top 10 genetic principal components (obtained from UK Biobank) were included as covariates in the model. The PLINK2^2^ command –glm was used for this analysis. A genomic inflation factor was estimated to 1.02 based on the GWAS summary statistics, i.e., no discernable inflation of the associations was observed. The association quantile-quantile plot, and the Manhattan plot are shown below.

| 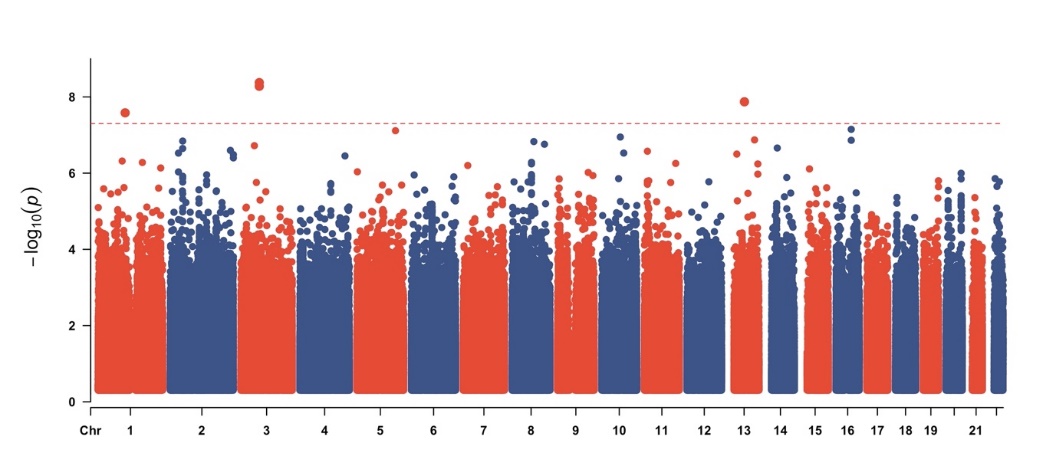 | 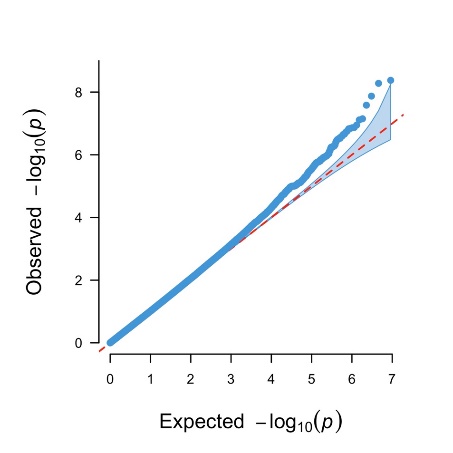 |
| --- | --- |

*Left panel: Manhattan plot of GWAS for sleep duration in participants sleeping > 7 hours, with the dotted red line showing the genome-wide significance level (p < 5×10^-8^). Right panel: QQ plot of GWAS for sleep duration.*

#### *GWAS for Hippocampal volume, total grey matter volume and intracranial volume*

For the 29 155 participants, GWAS for total hippocampal volume (HippV), total grey matter volume (TGV) and estimated intracranial volume (ICV) were performed using the –glm function from PLINK2, separately for each MRI derived measure. In each GWAS, the phenotype was first scaled as for sleep duration. In addition to age at MRI, sex and the top 10 genetic PCs, ICV was included as a covariate for the GWAS of HippV and TGV. Genomic inflation factors were estimated to 1.05, 1.06 and 1.02 for the GWAS summary statistics for HippV, ICV and TGV, respectively. The association quantile-quantile plots, and the Manhattan plots are shown below.

| 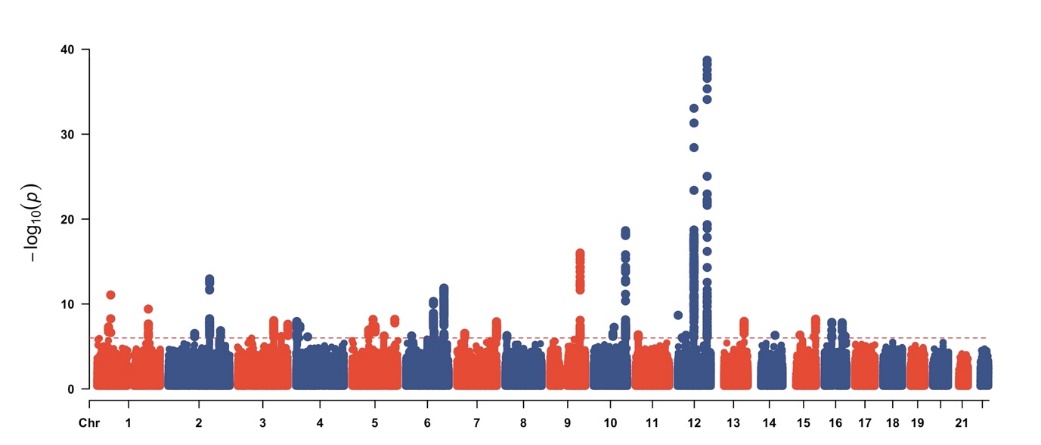 | 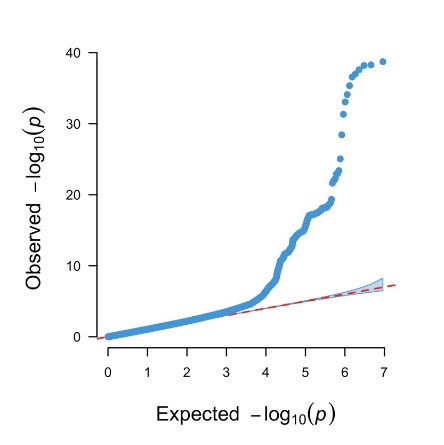 |
| --- | --- |

*Left panel: Manhattan plot of GWAS for hippocampal volume, with the dotted red line showing the genome-wide significance level (p < 5×10^-8^). Right panel: QQ plot of GWAS for hippocampal volume.*

| 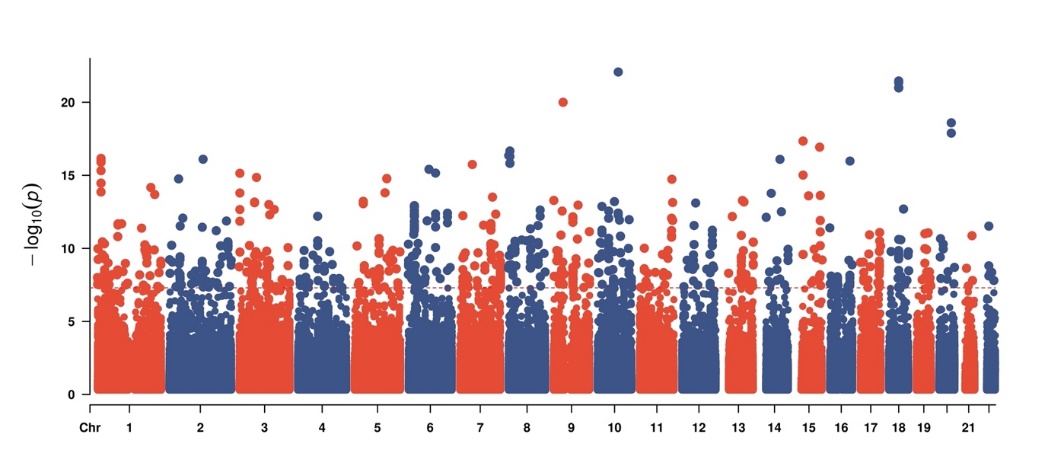 | 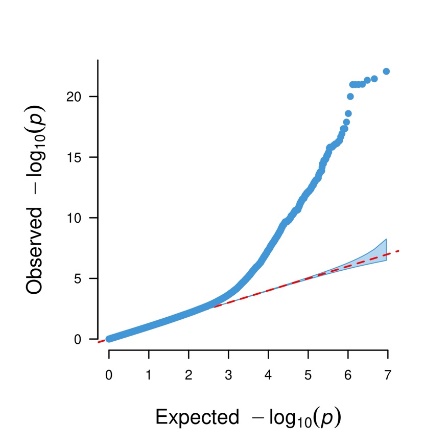 |
| --- | --- |

*Left panel: Manhattan plot of GWAS for total gray matter volume, with the dotted red line showing the genome-wide significance level (p < 5×10^-8^). Right panel: QQ plot of GWAS for total gray matter volume.*

| 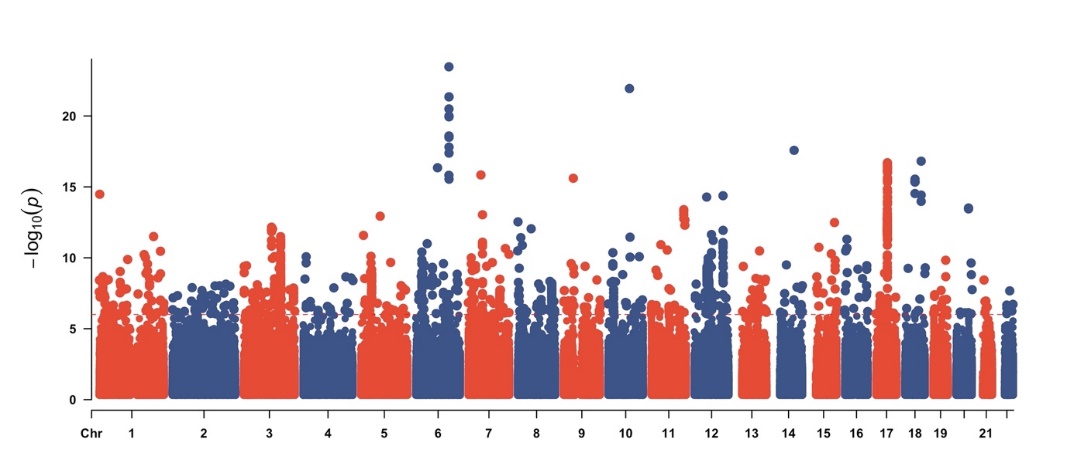 | 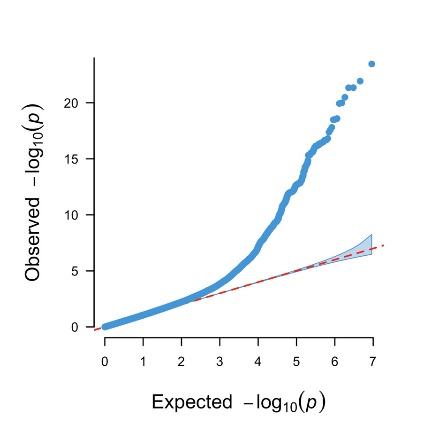 |
| --- | --- |

*Left panel: Manhattan plot of GWAS for intracranial volume, with the dotted red line showing the genome-wide significance level (p < 5×10^-8^). Right panel: QQ plot of GWAS for intracranial volume.*

**SNP heritability and genetic correlations**

The Linkage Disequilibrium score regression (LD)^3^ model was used for estimating SNP heritability and between-traits genetic correlations using GWAS summary statistics. Association statistics were first aligned with high quality HapMap 3 SNPs, and then the LD scores for these set SNPs were used. The signed-sumstats were assigned to regression beta, 0. The SNP heritabilities for hippocampal volume, ICV and total gray matter volume were estimated to 0.29 (se=0.03), 0.35 (se=0.03) and 0.22 (se=0.02), respectively. The numbers for Sleep durations were 0.04 (se=0.003) and 0.02 (se=0.005) for participants sleeping less than or equal to versus longer than 7 hours, respectively. The genetic correlations between sleep duration (<=7 hours) and hippocampal volume, ICV and total gray matter volume were estimated to be 0.04 (se=0.05, p=0.47), 0.1 (se=0.05, p=0.05) and 0.09 (se=0.06, p=0.10), respectively. The genetic correlations between sleep duration (>7 hours) and hippocampal volume, ICV and total gray matter volume were estimated to be -0.08 (se=0.10, p=0.43), -0.07 (se=0.08, p=0.44) and 0.04 (se=0.10, p=0.70), respectively.

### **Polygenic scores**

To accurately estimate the polygenic scores for a trait, we first computed the posterior effect size per SNP using the Bayesian mixture model implemented in PRS-CS^4^. We only performed analysis for SNPs that exist in the high quality HapMap3 SNP set. The polygenic parameter (phi) was set to 0.01 for all analysis in the study, and the default values for other parameters from PRS-CS were used. These computed posterior effects were used as weights in the computation of PGSs for a trait by using the score function from PLINK2. To examine the associations between PGS for a trait with a second trait, linear regression models were used. The same covariates included in the GWAS analysis were included as covariates in addition to PGSs in these models.

| PGS | Trait | Beta | Se | t-score | P | FDR |
| --- | --- | --- | --- | --- | --- | --- |
| ICV | SleepLe7 | 1.91x10^-2^ | 2.56x10^-3^ | 8.47 | <2x10^-16^ | 2.40x10^-15^ |
|  | SleepGt7 | -6.23x10^-3^ | 2.98x10^-3^ | -2.09 | 0.04 | 0.32 |
| HippV | SleepLe7 | 2.43x10^-3^ | 2.26x10^-3^ | 1.07 | 0.28 | 0.66 |
|  | SleepGt7 | -4.16x10^-3^ | 2.98x10^-3^ | -1.40 | 0.16 | 0.66 |
| TGV | SleepLe7 | 1.05x10^-2^ | 2.56x10^-3^ | 4.65 | 3.28x10^-6^ | 3.28x10^-5^ |
|  | SleepGt7 | -4.23x10^-3^ | 2.97x10^-3^ | -1.42 | 0.16 | 0.66 |
| SleepLe7 | ICV | 3.33x10^-2^ | 4.77x10^-3^ | 6.99 | 2.75x10^-12^ | 3.03x10^-11^ |
|  | HippV | 3.25x10^-3^ | 4.56x10^-3^ | 0.713 | 0.48 | 0.66 |
|  | TGV | 7.84x10^-3^ | 2.91x10^-3^ | 2.69 | 7.13x10^-3^ | 6.42x10^-2^ |
| SleepGt7 | ICV | -8.42x10^-3^ | 4.78x10^-3^ | -1.76 | 7.81x10^-2^ | 0.51 |
|  | HippV | -7.30x10^-3^ | 4.56x10^-3^ | -1.60 | 0.11 | 0.66 |
|  | TGV | -4.50x10^-3^ | 2.90x10^-3^ | -1.55 | 0.12 | 0.66 |

*Polygenic score (PGS) associations between sleep duration and HippV, ICV and TGV. PGS were tested for associations with measured traits (Traits) in independent samples. Linear regression models were used to estimate the effect sizes of PGS to sleep duration for those sleeping less than or equal 7 hours per day (SleepLe7) and those sleeping more than 7 hours per day (SleepGt7). Generalized additive models were used for estimating effects of PGS to total Hippocampal volume (HippV), total grey matter volume (TGV) and estimated intracranial volume (ICV), taking age at scan as the smoothed variable. Age, sex and the top 10 genetic principal components were included as covariates in all models. For HippV and TGV, ICV was included as an additional covariate. Note, effect sizes (beta) indicate the amount of change in standard deviation of a trait by one standard deviate increase of a PGS.*

### **Mendelian randomization**

*Instrumental variants selection*

To determine the SNP-phenotype associations, the subset of SNPs associated with the phenotype having p <1x10^-6^ and also existing in the GWAS results were used. In addition, SNPs with minor allele frequency (MAF) < 0.05 in the GWAS sample, or having ambiguous allelic coding, i.e., A/T or C/G, were excluded from the instrument set. To remove correlated SNPs, the linkage disequilibrium (LD) clumping method implemented by PLINK was used. The following parameters in PLINK were set, clump-kb 10,000 kb, and --clump-r2 0.1. The LD structure from the corresponding GWAS sample were used for clumping. The full list of SNPs selected as instruments for each analysis are presented in SI Genetics supplementary tables. The strength of the instruments to an outcome was evaluated using the F statistic. As a sensitivity analysis, we repeated the analysis using p<10^-5^ as the threshold to select instrumental SNPs (SI Genetic supplementary tables). No qualitative differences in the results were detected.

| Exposure | outcome | Beta | Se | P |
| --- | --- | --- | --- | --- |
| ICV | SleepLe7 | 5.99x10^-2^ | 1.73x10^-2^ | 5.36x10^-4^ |
|  | SleepGt7 | -3.67x10^-3^ | 2.57x10^-2^ | 0.87 |
| HippV | SleepLe7 | 1.13x10^-2^ | 1.45x10^-2^ | 0.44 |
|  | SleepGt7 | 9.61x10^-3^ | 2.18x10^-2^ | 0.66 |
| TGV | SleepLe7 | 5.18x10^-2^ | 3.22x10^-2^ | 0.11 |
|  | SleepGt7 | -8.99x10^-3^ | 6.59x10^-2^ | 0.89 |
| SleepLe7 | ICV | 4.38x10-2 | 0.14 | 0.75 |
|  | HippV | 0.16 | 0.15 | 0.30 |
|  | TGV | 2.66x10^-2^ | 8.35x10^-2^ | 0.75 |
| SleepGt7 | ICV | 6.14x10^-2^ | 0.38 | 0.87 |
|  | HippV | -0.16 | 0.22 | 0.48 |
|  | TGV | -0.35 | 0.14 | 1.22x10^-2^ |

*Inverse variance weighted Mendelian randomization results between sleep duration and HippV, ICV and TGV.*

Finally, we performed two-sample Mendelian randomization analysis using several popular methods, the inverse variance weighted (IVW), the Egger regression, the weighted median and the robust adjusted profile score (RAPS) methods. The RAPs was only applied to weaker instruments, i.e. P <10^-5^. Full report for each pair of traits and each method were presented in SI Genetics supplementary notes.

| Exposure | outcome | Beta | Se | P |
| --- | --- | --- | --- | --- |
| SleepLe7 | ICV | -7.90x10-3 | 8.47x10-2 | 0.93 |
|  | HippV | 2.72x10-3 | 8.10x10-2 | 0.97 |
|  | TGV | 6.96x10-3 | 5.15x10-2 | 0.89 |
| SleepGt7 | ICV | 5.29x10^-3^ | 7.29x10-2 | 0.94 |
|  | HippV | -6.95x10^-2^ | 6.89x10^-2^ | 0.31 |
|  | TGV | -2.10x10^-2^ | 4.36x10^-2^ | 0.63 |

*Mendelian randomization by RAPS between sleep duration and HippV, ICV and TGV.*
