## Supplementary material for "Sleep duration and brain structure – phenotypic associations and genotypic covariance": SI Genetics supplementary notes

### Two sample MR report

---

#### HippV against SleepGt7

Date: **24 september, 2021**

---

##### Results from two sample MR:

| method | nsnp | b | se | pval |
|---|---|---|---|---|
| MR Egger | 76 | 0.0675178 | 0.0370249 | 0.0722541 |
| Weighted median | 76 | -0.0061928 | 0.0211743 | 0.7699293 |
| Inverse variance weighted | 76 | -0.0057491 | 0.0149908 | 0.7013456 |
| Simple mode | 76 | 0.0223998 | 0.0475225 | 0.6387583 |
| Weighted mode | 76 | 0.0163095 | 0.0347411 | 0.6401034 |

---

##### Heterogeneity tests

| method | Q | Q_df | Q_pval |
|---|---|---|---|
| MR Egger | 89.81645 | 74 | 0.1017810 |
| Inverse variance weighted | 95.45002 | 75 | 0.0557322 |

---

##### Test for directional horizontal pleiotropy

| egger_intercept | se | pval |
|---|---|---|
| -0.0038145 | 0.0017706 | 0.0344626 |

---

##### Test that the exposure is upstream of the outcome

| snp_r2.exposure | snp_r2.outcome | correct_causal_direction | steiger_pval |
|---|---|---|---|
| 0.0937249 | 0.0011498 | TRUE | 0 |

Note - R^2 values are approximate

---

**Forest plot of single SNP MR**

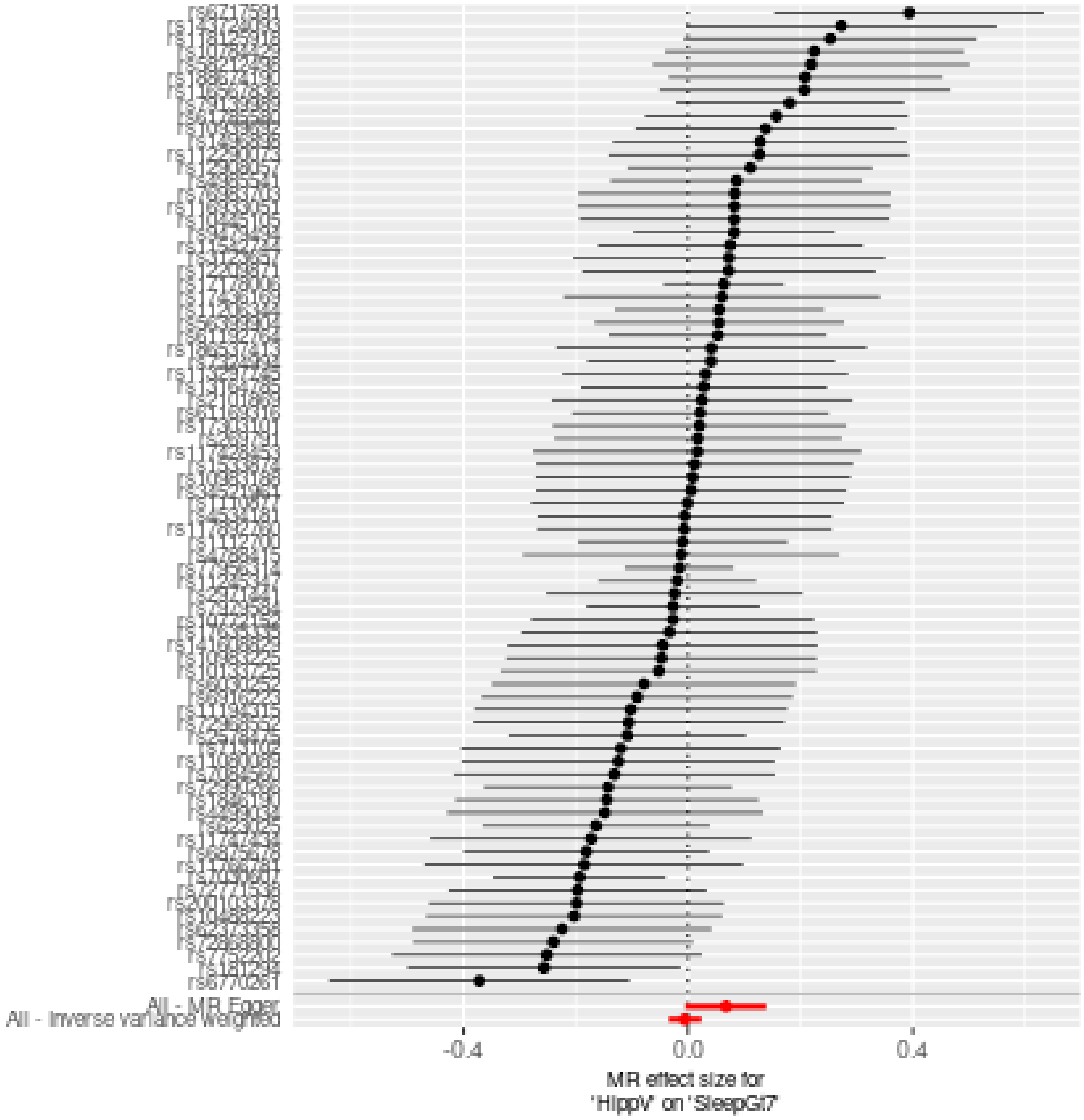

**Comparison of results using different MR methods**

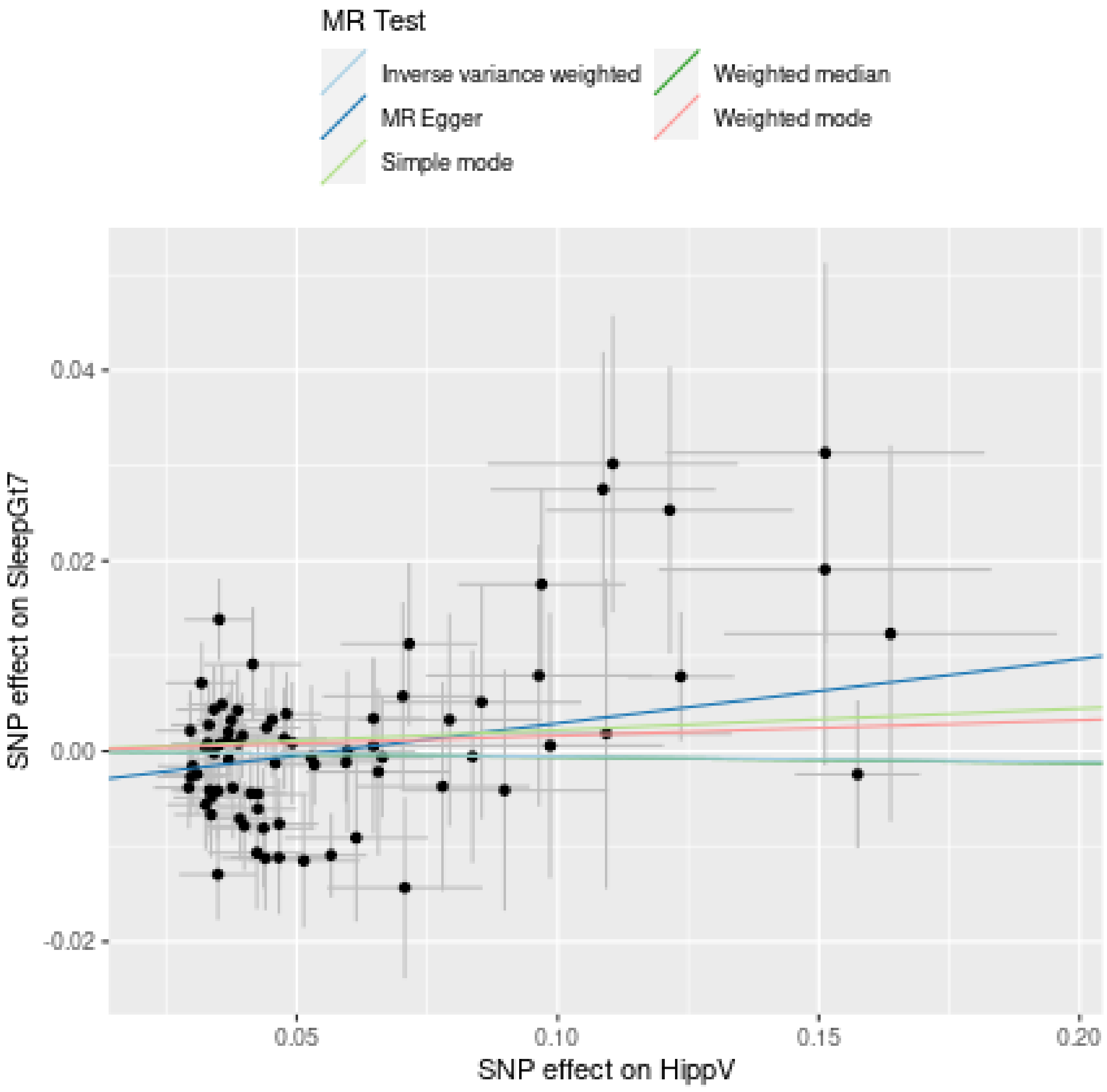

**Funnel plot**

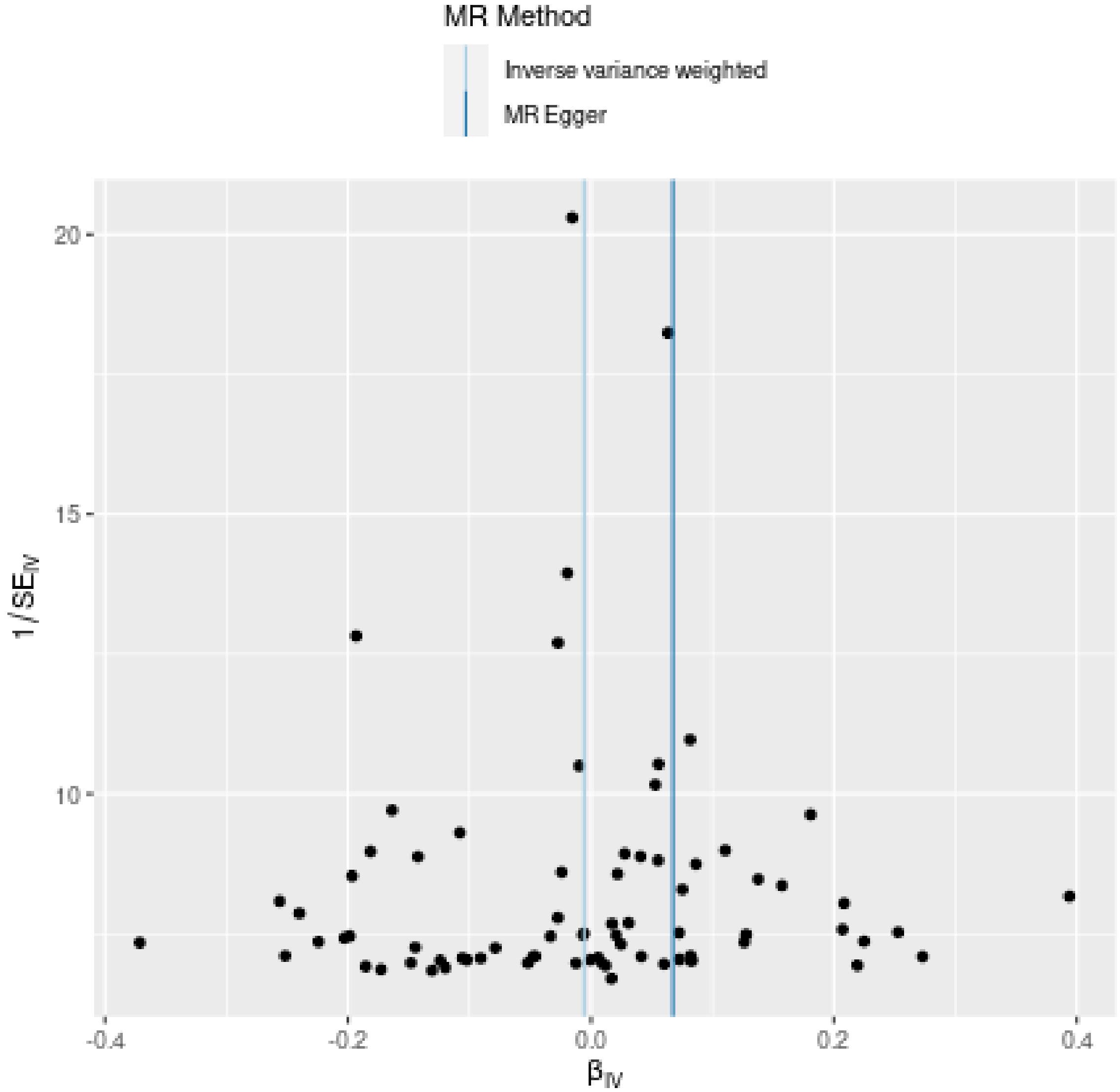

MR Method
Inverse variance weighted
MR Egger
20
15
10
1/SE$_{IV}$
-0.4
-0.2
0.0
0.2
0.4
β$_{IV}$

**Leave-one-out sensitivity analysis**

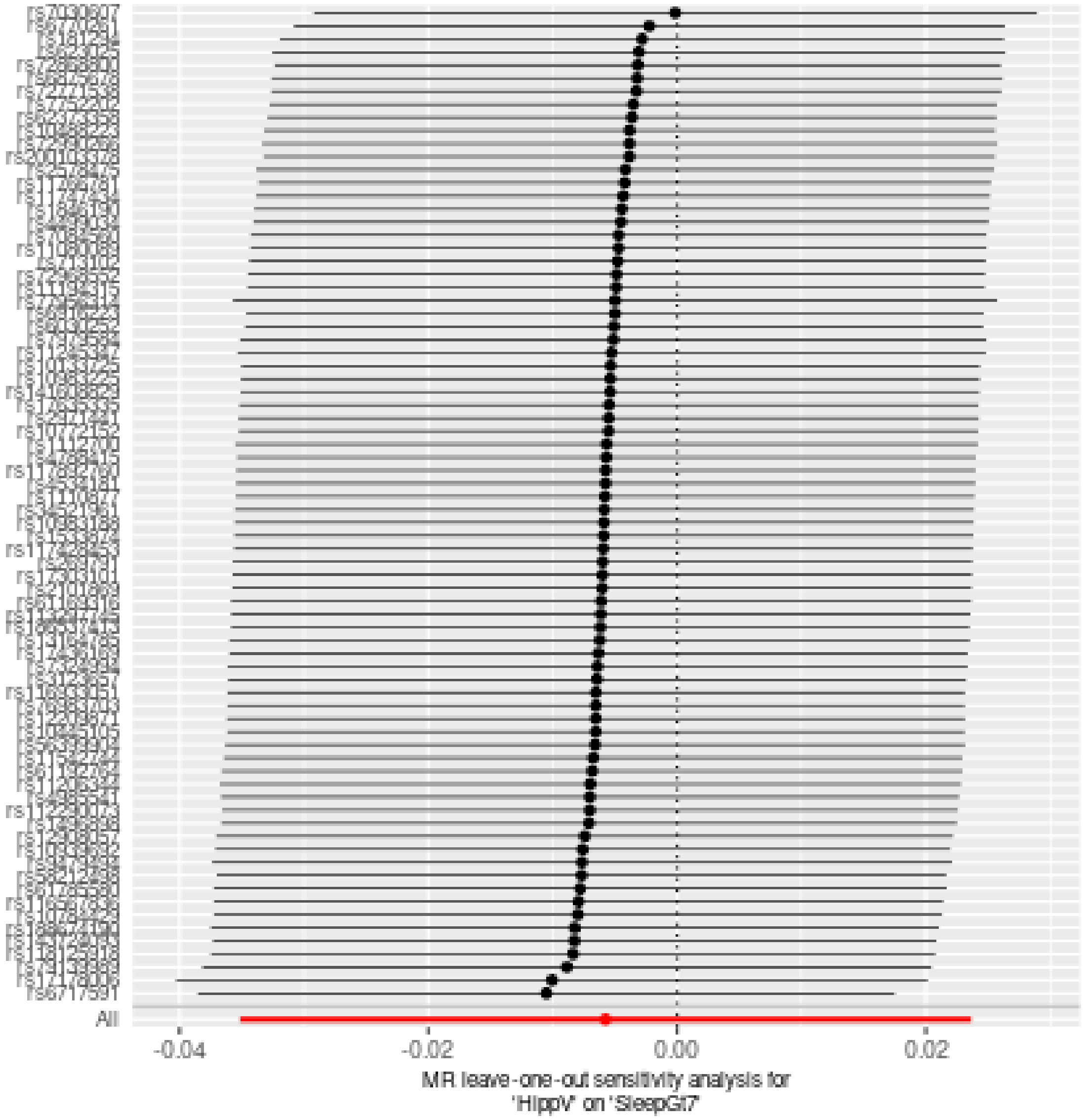

### Two sample MR report

---

#### HippV against SleepLe7

Date: **24 september, 2021**

---

##### Results from two sample MR:

| method | nsnp | b | se | pval |
|---|---|---|---|---|
| MR Egger | 76 | 0.0169608 | 0.0265944 | 0.5255981 |
| Weighted median | 76 | 0.0014014 | 0.0156517 | 0.9286559 |
| Inverse variance weighted | 76 | 0.0105060 | 0.0104622 | 0.3152862 |
| Simple mode | 76 | -0.0103722 | 0.0392096 | 0.7920952 |
| Weighted mode | 76 | 0.0097953 | 0.0251273 | 0.6977701 |

---

##### Heterogeneity tests

| method | Q | Q_df | Q_pval |
|---|---|---|---|
| MR Egger | 80.97263 | 74 | 0.2707679 |
| Inverse variance weighted | 81.04907 | 75 | 0.2962110 |

---

##### Test for directional horizontal pleiotropy

| egger_intercept | se | pval |
|---|---|---|
| -0.0003362 | 0.0012719 | 0.79228 |

---

##### Test that the exposure is upstream of the outcome

| snp_r2.exposure | snp_r2.outcome | correct_causal_direction | steiger_pval |
|---|---|---|---|
| 0.0937249 | 0.0004956 | TRUE | 0 |

Note - R^2 values are approximate

---

**Forest plot of single SNP MR**

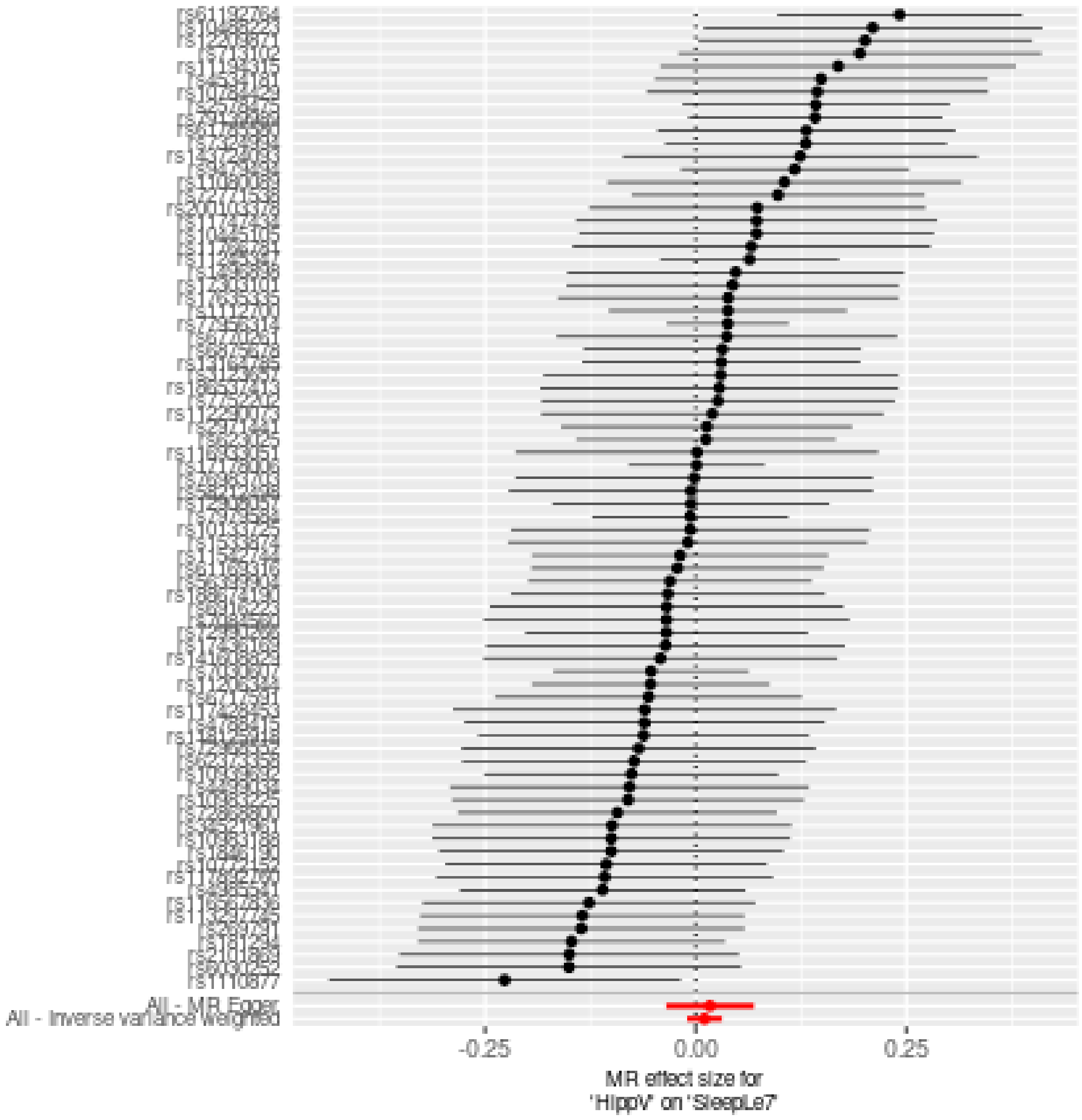

**Comparison of results using different MR methods**

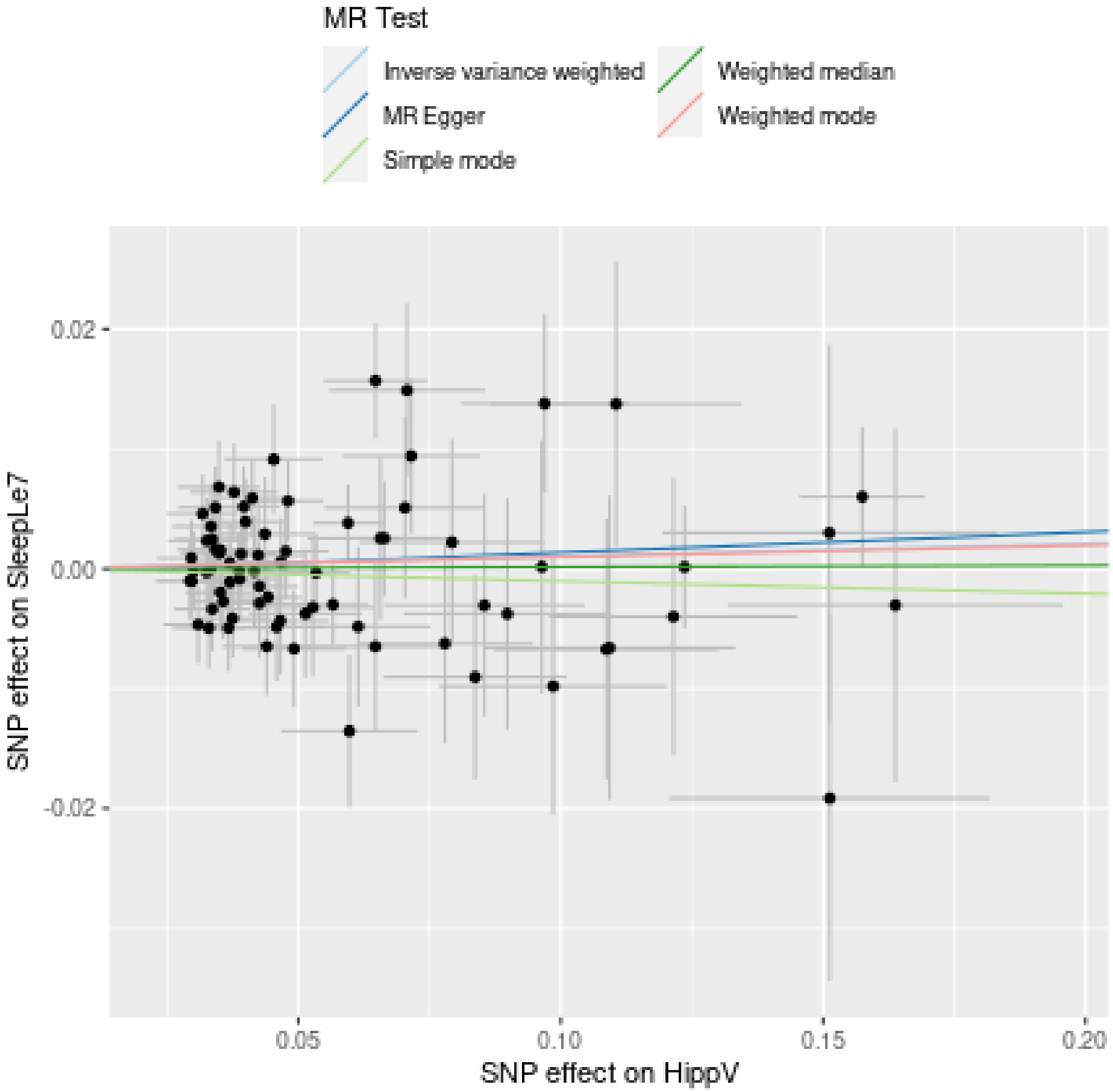

**Funnel plot**

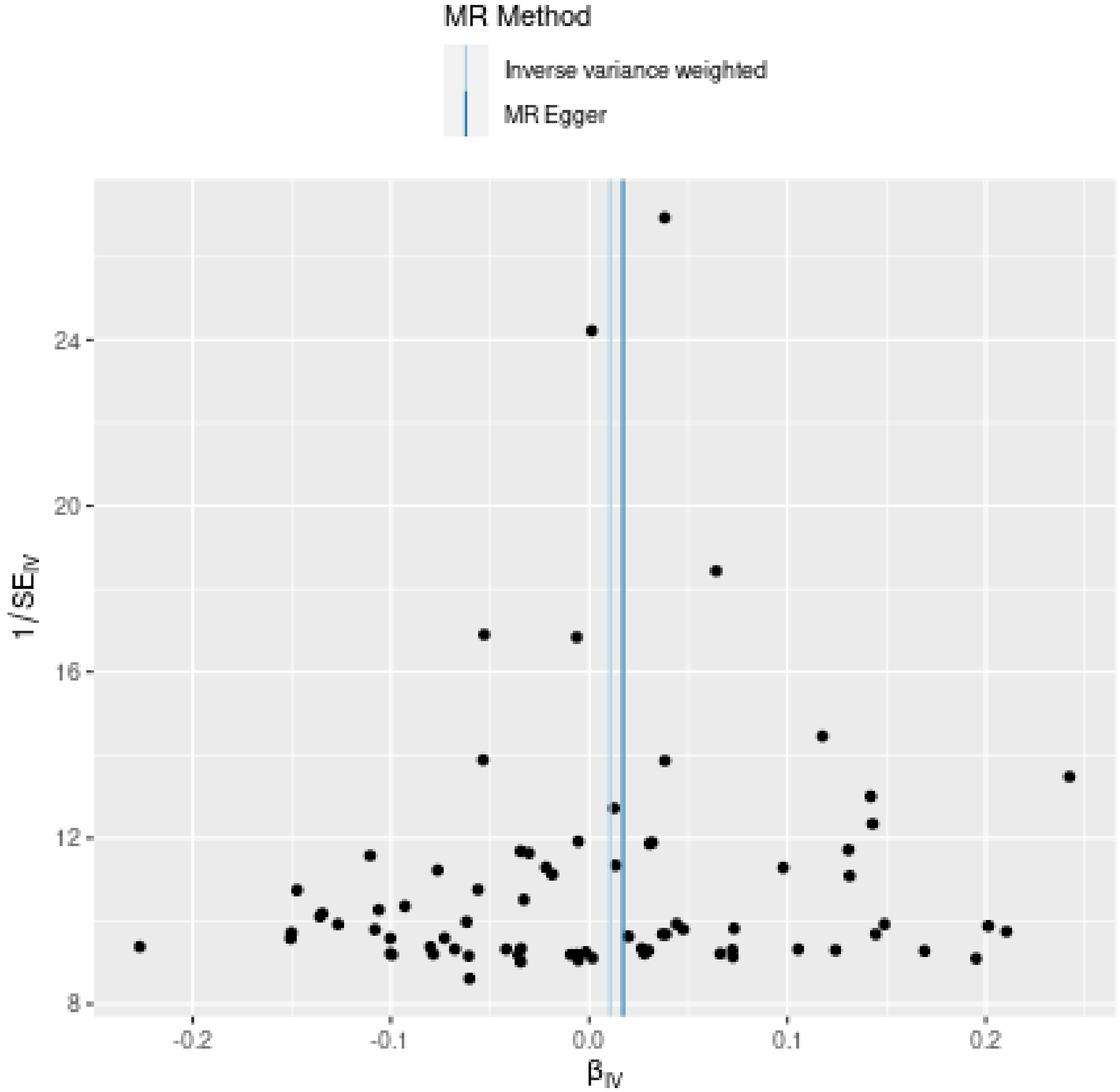
MR Method
Inverse variance weighted
MR Egger
24
20
16
12
8
-0.2
-0.1
0.0
0.1
0.2
$\beta_{IV}$
$1/SE_{IV}$

**Leave-one-out sensitivity analysis**

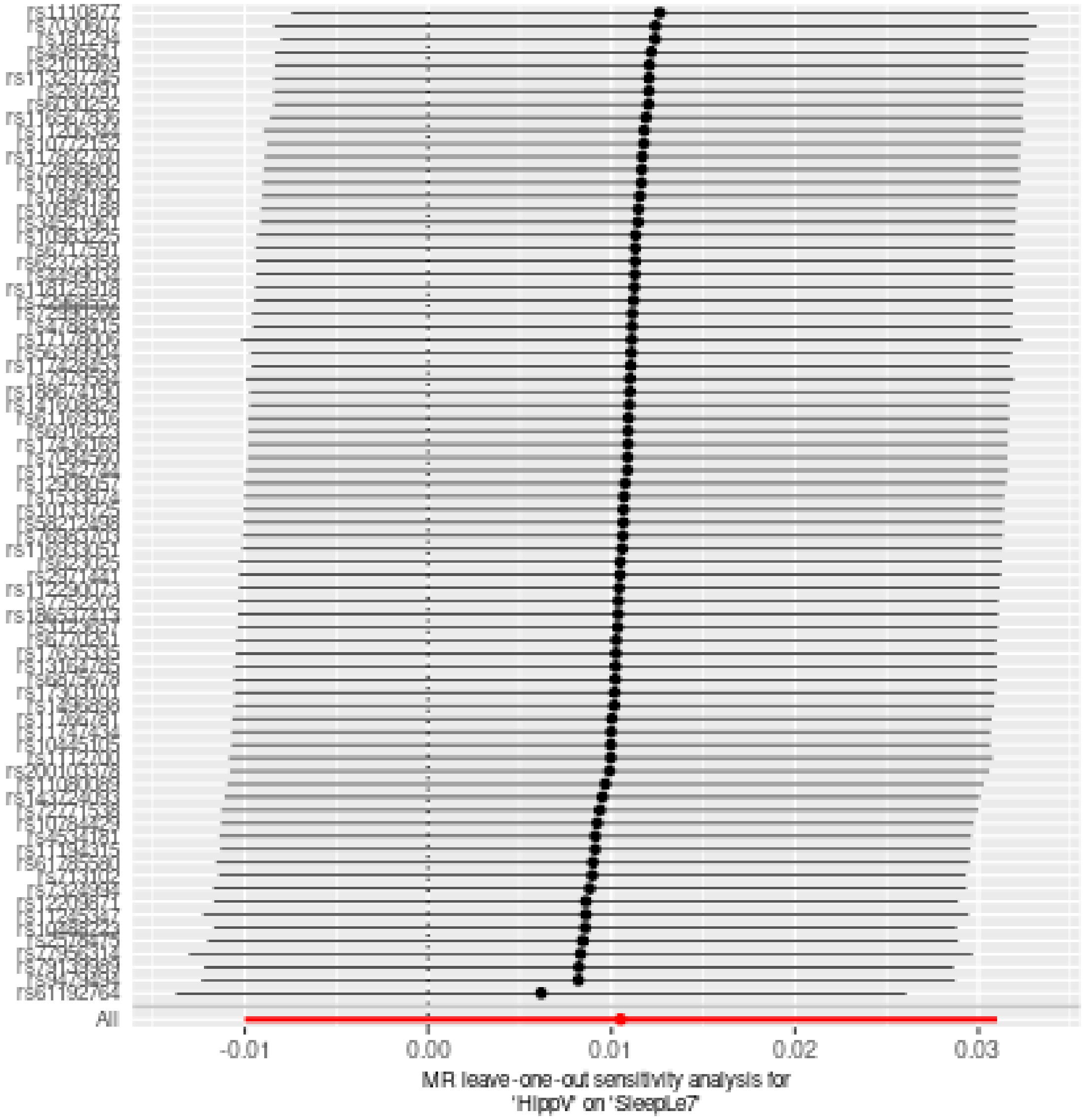

### Two sample MR report

#### ICV against SleepGt7

Date: **24 september, 2021**

##### Results from two sample MR:

| method | nsnp | b | se | pval |
|---|---|---|---|---|
| MR Egger | 88 | 0.0339083 | 0.0396457 | 0.3947706 |
| Weighted median | 88 | -0.0173681 | 0.0195531 | 0.3744033 |
| Inverse variance weighted | 88 | -0.0070838 | 0.0147017 | 0.6299251 |
| Simple mode | 88 | -0.0274712 | 0.0485554 | 0.5730056 |
| Weighted mode | 88 | -0.0150815 | 0.0406290 | 0.7113908 |

##### Heterogeneity tests

| method | Q | Q_df | Q_pval |
|---|---|---|---|
| MR Egger | 113.5486 | 86 | 0.0249814 |
| Inverse variance weighted | 115.1845 | 87 | 0.0232427 |

##### Test for directional horizontal pleiotropy

| egger_intercept | se | pval |
|---|---|---|
| -0.0021613 | 0.0019417 | 0.2687687 |

##### Test that the exposure is upstream of the outcome

| snp_r2.exposure | snp_r2.outcome | correct_causal_direction | steiger_pval |
|---|---|---|---|
| 0.097292 | 0.001197 | TRUE | 0 |

Note - R^2 values are approximate

**Forest plot of single SNP MR**

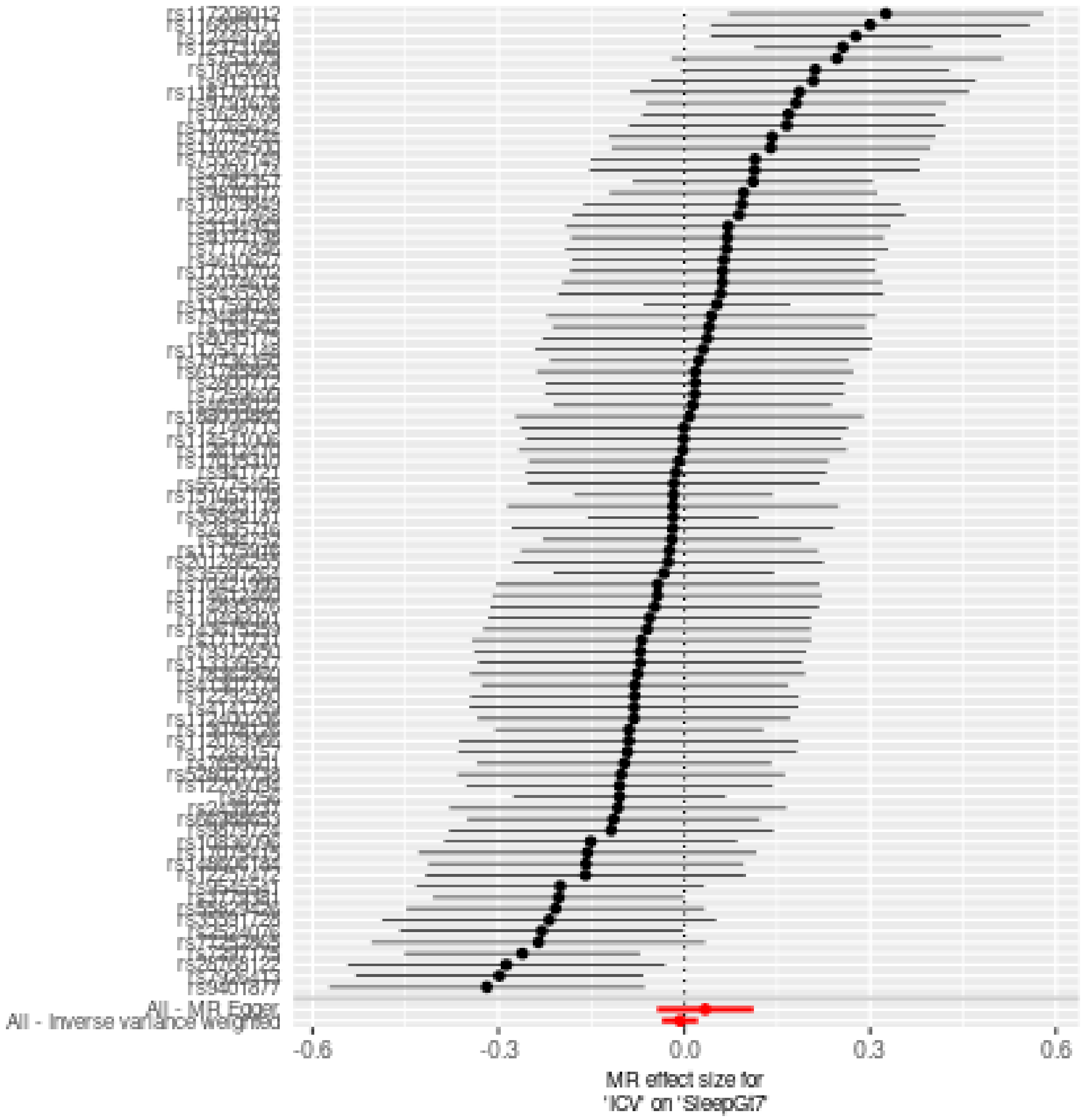

**Comparison of results using different MR methods**

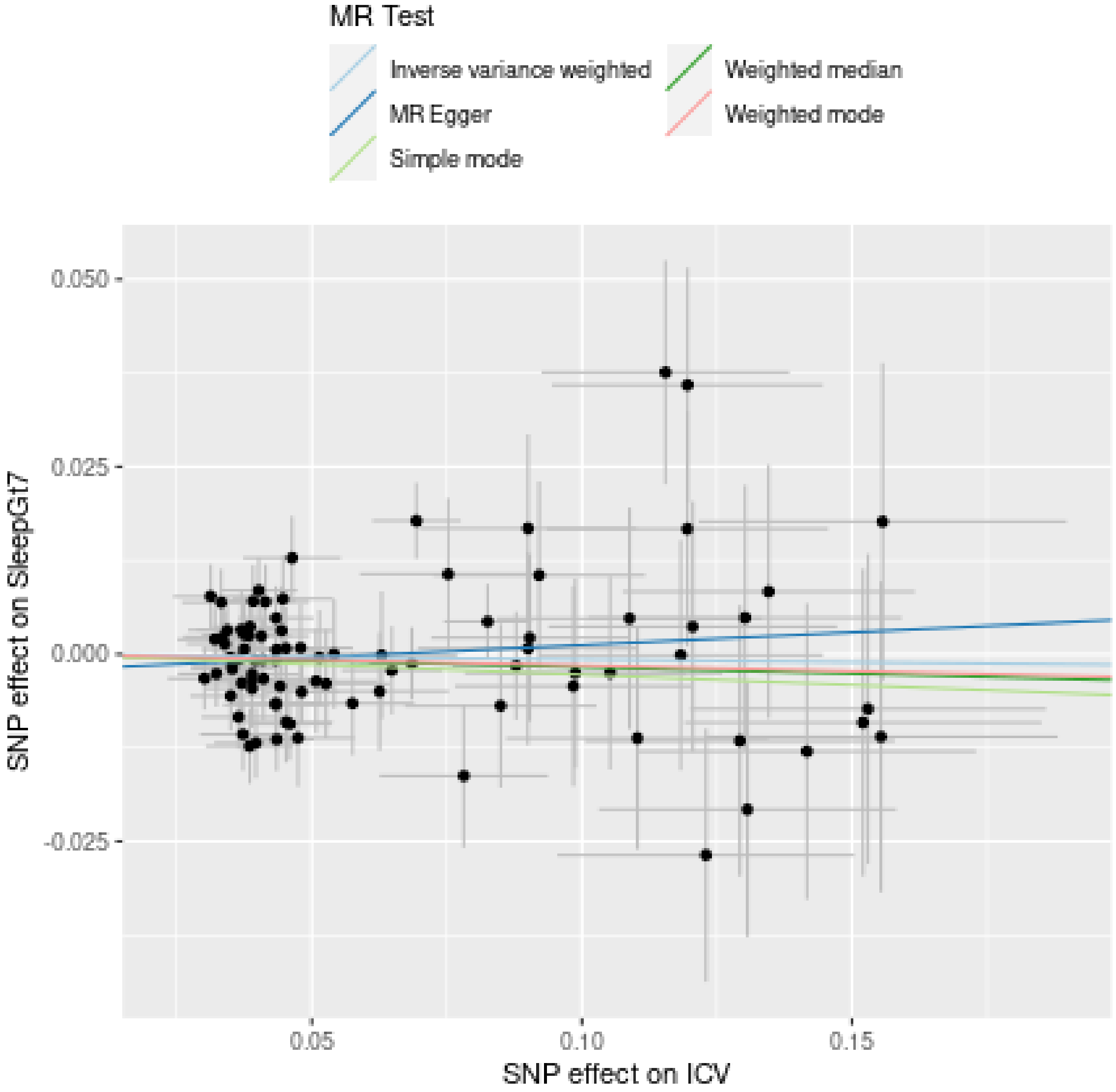

**Funnel plot**

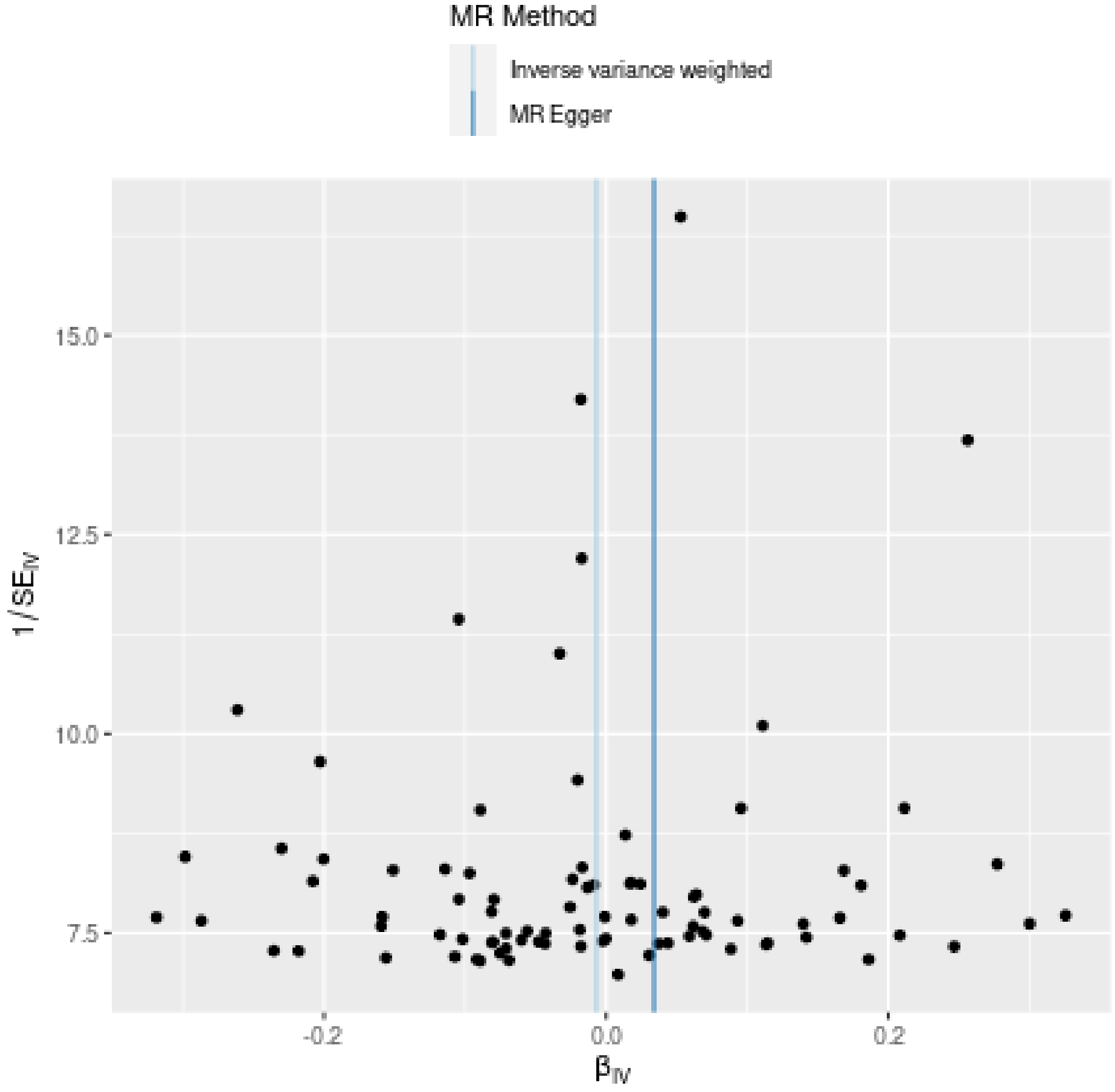
MR Method
Inverse variance weighted
MR Egger
15.0
12.5
10.0
7.5
$1/SE_{IV}$
-0.2
0.0
0.2
$\beta_{IV}$

**Leave-one-out sensitivity analysis**

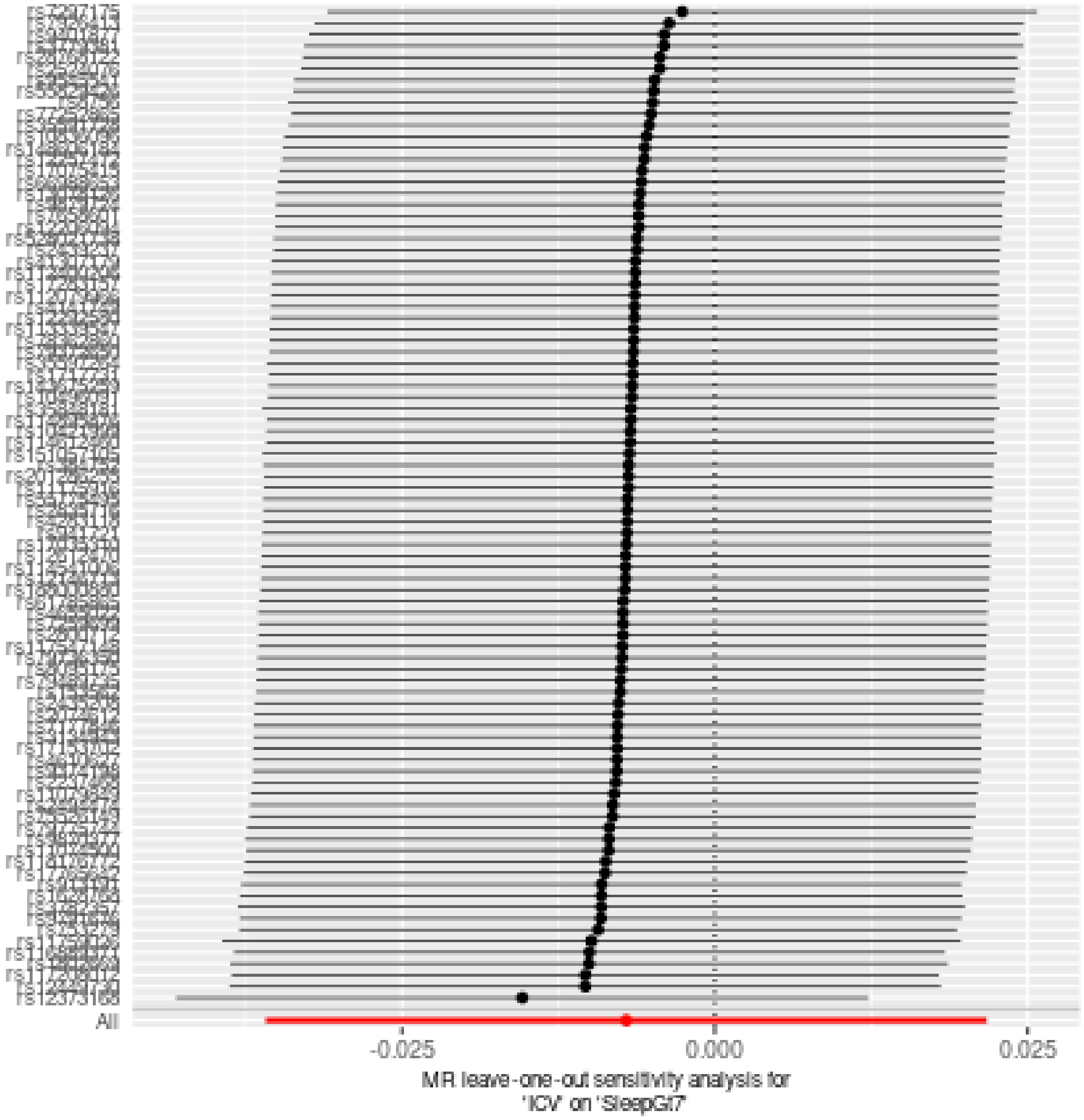

### Two sample MR report

---

#### ICV against SleepLe7

Date: **24 september, 2021**

---

##### Results from two sample MR:

| method | nsnp | b | se | pval |
|---|---|---|---|---|
| MR Egger | 88 | -0.0023802 | 0.0318884 | 0.9406727 |
| Weighted median | 88 | 0.0427685 | 0.0150103 | 0.0043818 |
| Inverse variance weighted | 88 | 0.0408021 | 0.0118785 | 0.0005926 |
| Simple mode | 88 | 0.0482814 | 0.0384868 | 0.2130235 |
| Weighted mode | 88 | 0.0414786 | 0.0337732 | 0.2227018 |

---

##### Heterogeneity tests

| method | Q | Q_df | Q_pval |
|---|---|---|---|
| MR Egger | 127.3433 | 86 | 0.0025303 |
| Inverse variance weighted | 130.4896 | 87 | 0.0017729 |

---

##### Test for directional horizontal pleiotropy

| egger_intercept | se | pval |
|---|---|---|
| 0.0022751 | 0.0015608 | 0.1485691 |

---

##### Test that the exposure is upstream of the outcome

| snp_r2.exposure | snp_r2.outcome | correct_causal_direction | steiger_pval |
|---|---|---|---|
| 0.097292 | 0.0008324 | TRUE | 0 |

Note - R^2 values are approximate

---

**Forest plot of single SNP MR**

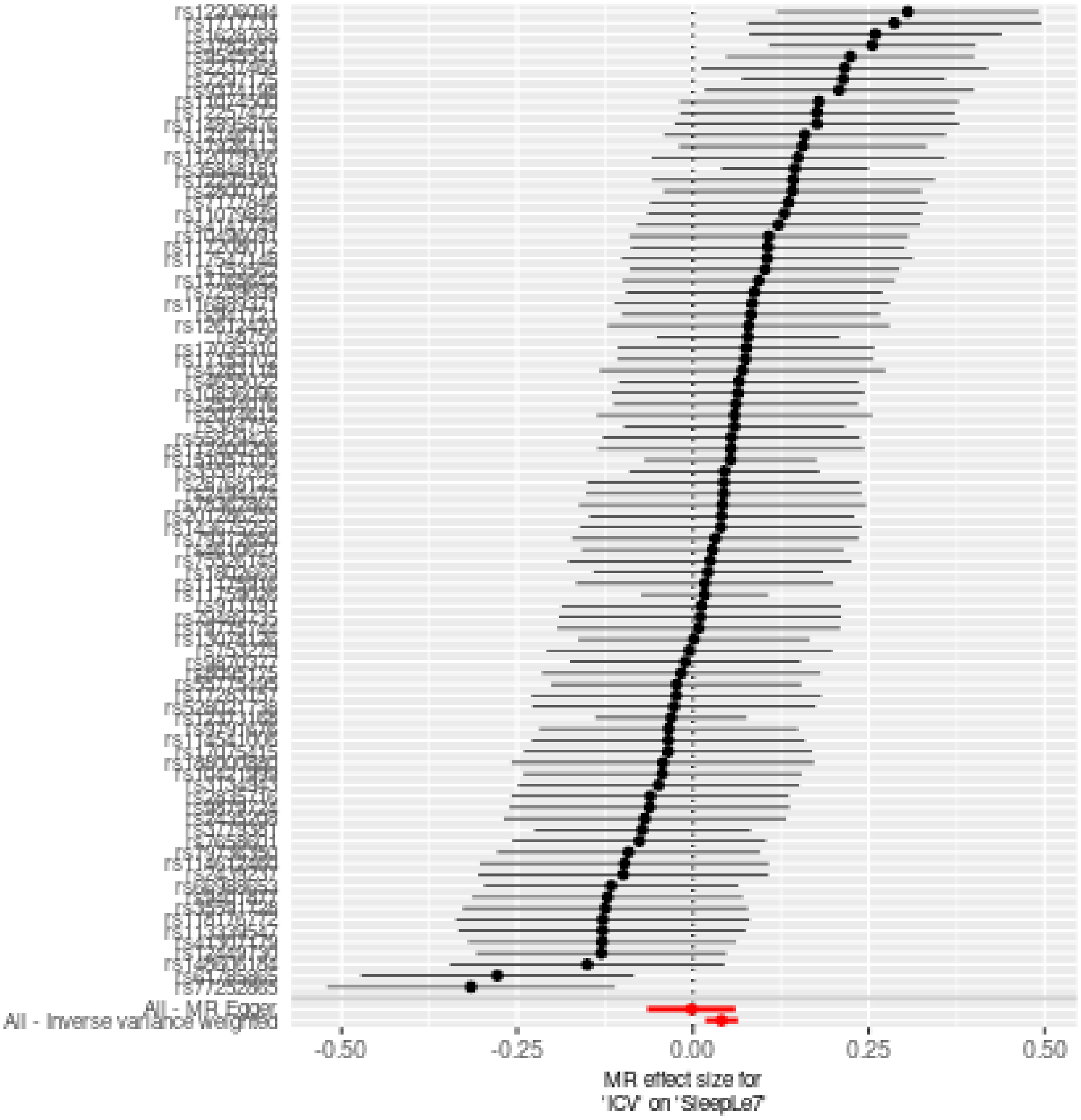

**Comparison of results using different MR methods**

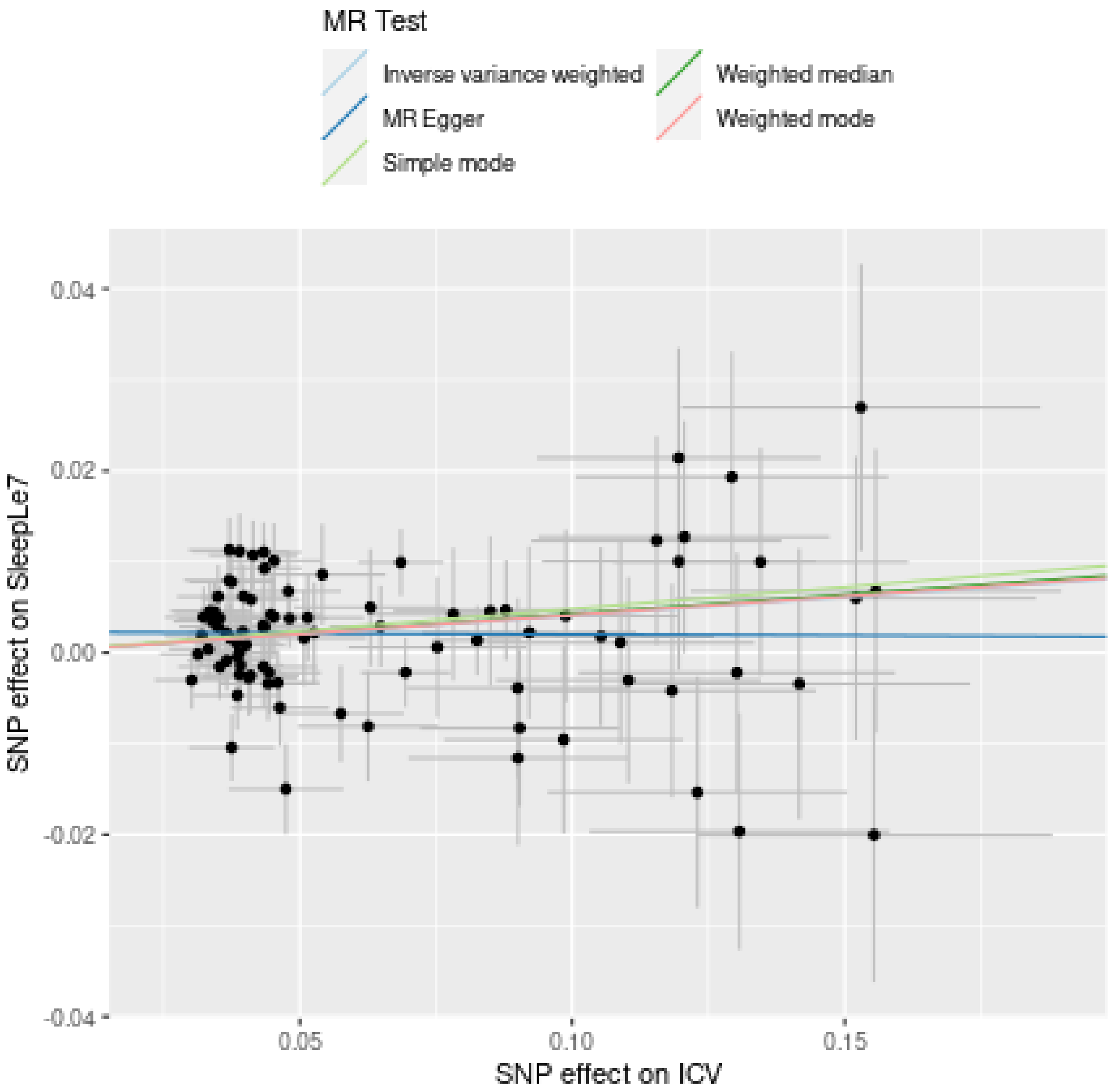

**Funnel plot**

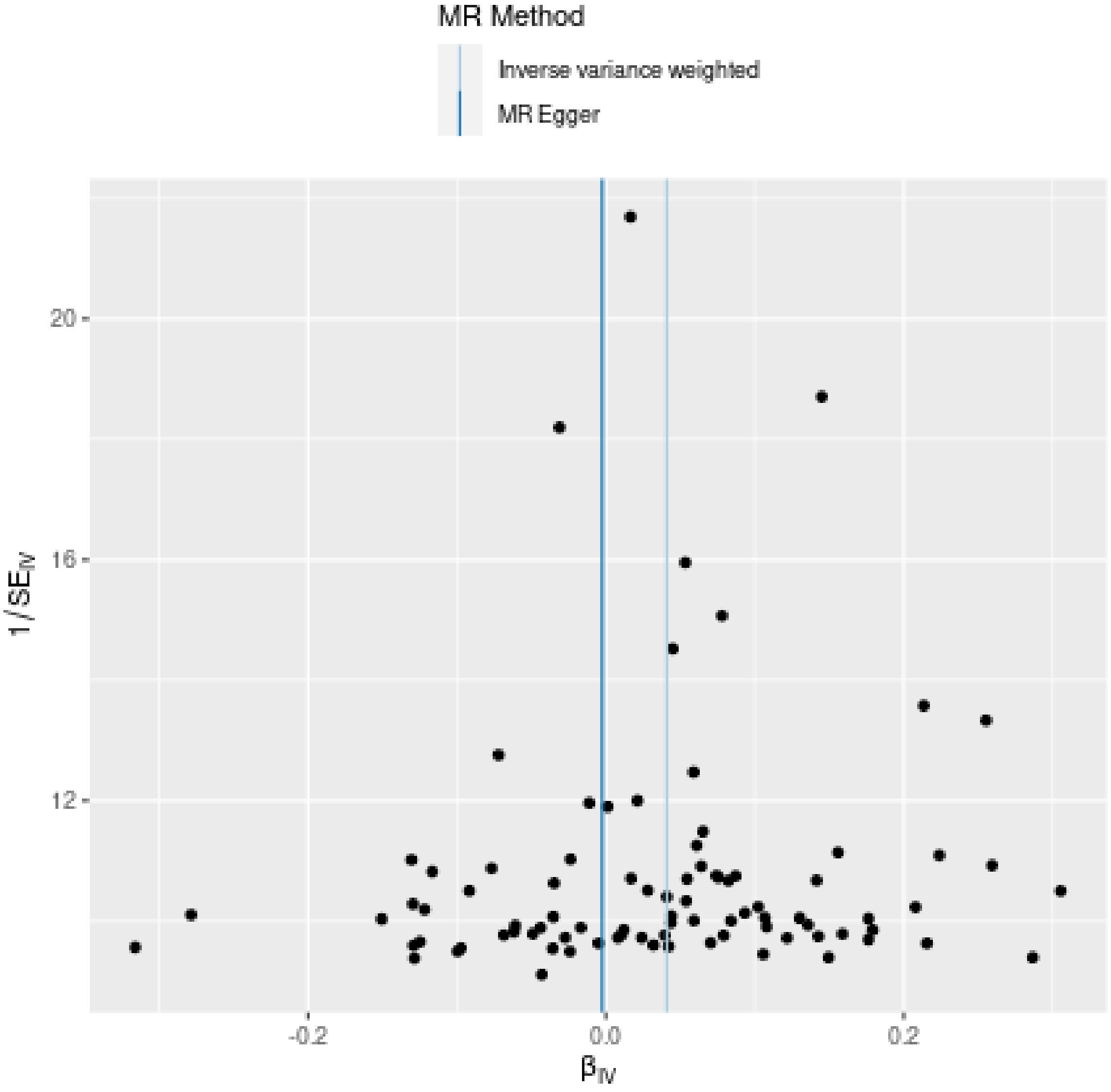

MR Method
Inverse variance weighted
MR Egger
20
16
12
1/SE$_{IV}$
-0.2
0.0
0.2
β$_{IV}$

**Leave-one-out sensitivity analysis**

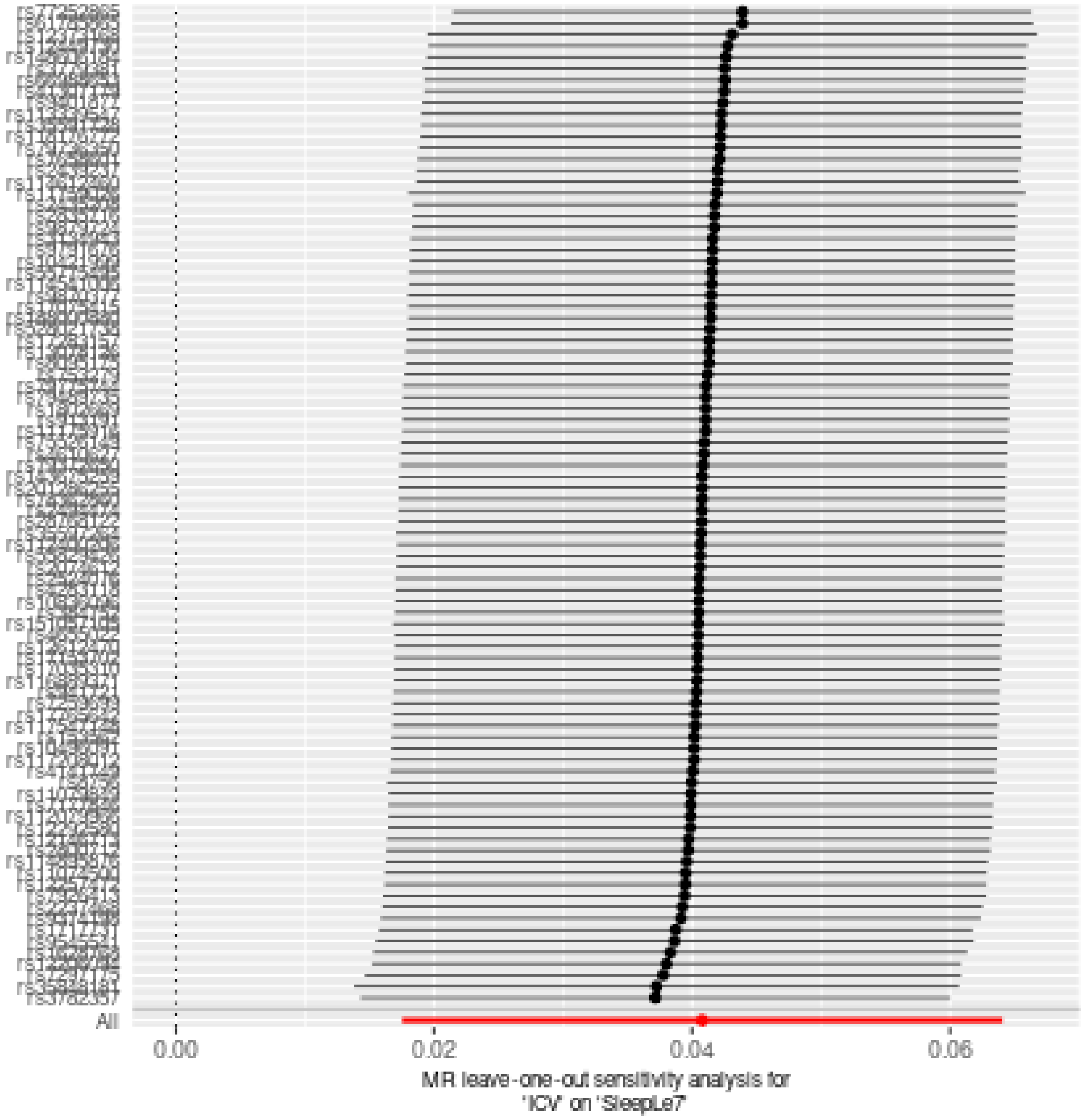

### Two sample MR report

---

#### SleepGt7 against HippV

Date: **24 september, 2021**

---

##### Results from two sample MR:

| method | nsnp | b | se | pval |
|---|---|---|---|---|
| MR Egger | 21 | -0.0925011 | 0.1512298 | 0.5480129 |
| Weighted median | 21 | -0.0458650 | 0.1029470 | 0.6559435 |
| Inverse variance weighted | 21 | -0.0746943 | 0.0728970 | 0.3055257 |
| Simple mode | 21 | -0.0566075 | 0.1817977 | 0.7587370 |
| Weighted mode | 21 | -0.0606288 | 0.1772525 | 0.7358833 |

---

##### Heterogeneity tests

| method | Q | Q_df | Q_pval |
|---|---|---|---|
| MR Egger | 16.35114 | 19 | 0.6337366 |
| Inverse variance weighted | 16.36920 | 20 | 0.6934719 |

---

##### Test for directional horizontal pleiotropy

| egger_intercept | se | pval |
|---|---|---|
| 0.0006559 | 0.0048807 | 0.8945073 |

---

##### Test that the exposure is upstream of the outcome

| snp_r2.exposure | snp_r2.outcome | correct_causal_direction | steiger_pval |
|---|---|---|---|
| 0.0052712 | 0.0010812 | TRUE | 0 |

Note - R^2 values are approximate

---

**Forest plot of single SNP MR**

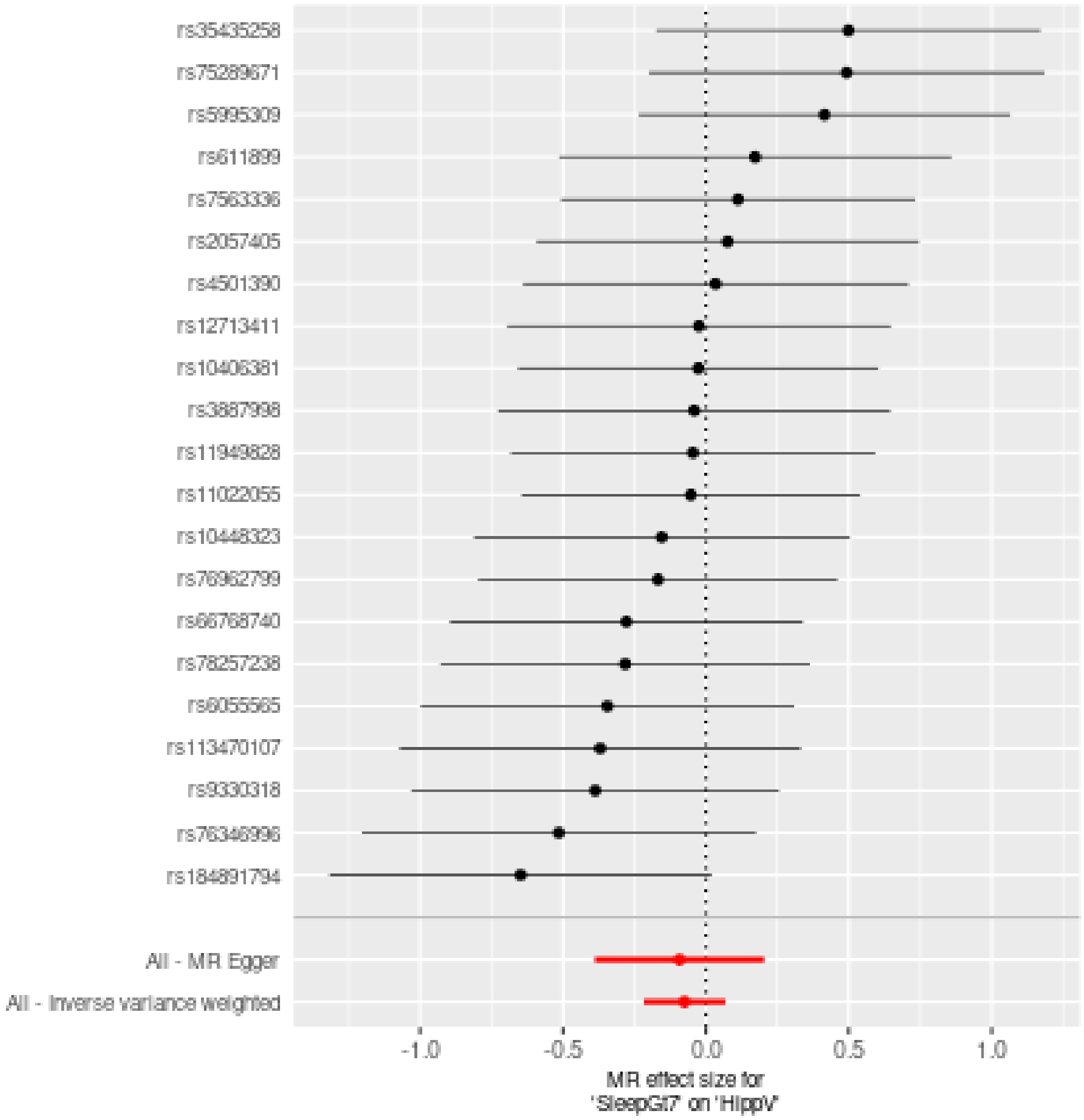

**Comparison of results using different MR methods**

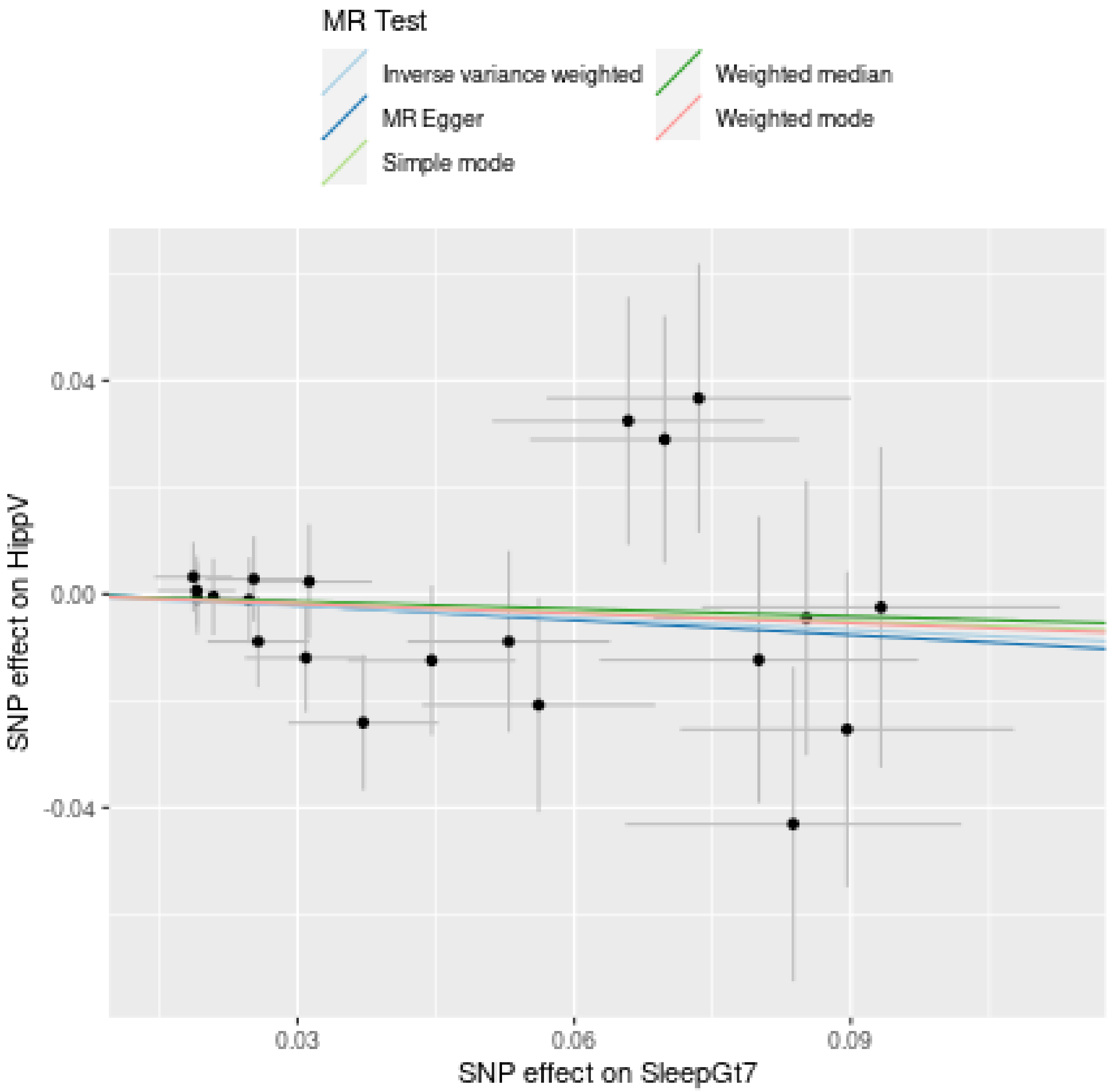

**Funnel plot**

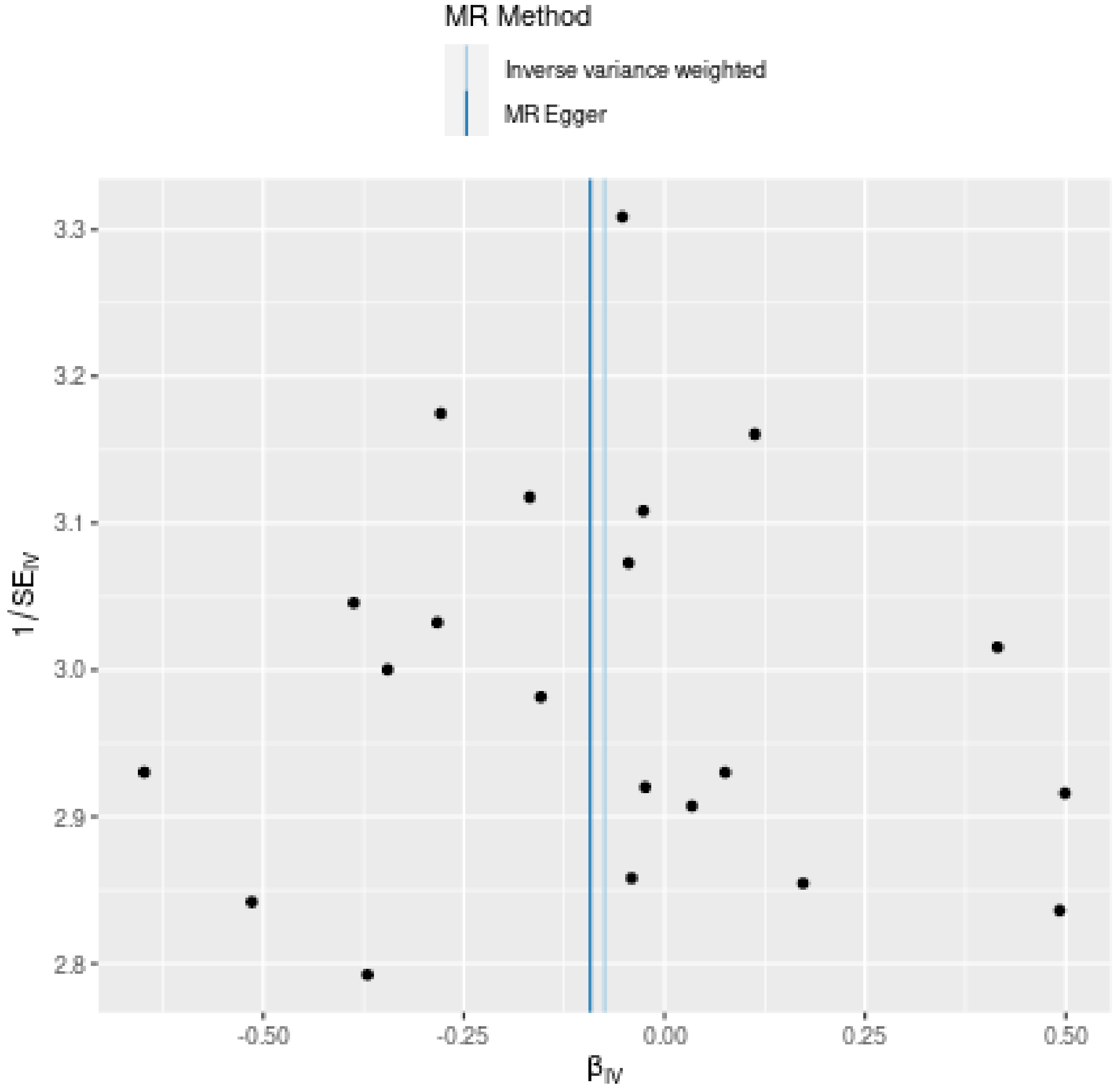

MR Method
Inverse variance weighted
MR Egger
3.3
3.2
3.1
3.0
2.9
2.8
1/SE$_{IV}$
-0.50
-0.25
0.00
0.25
0.50
β$_{IV}$

**Leave-one-out sensitivity analysis**

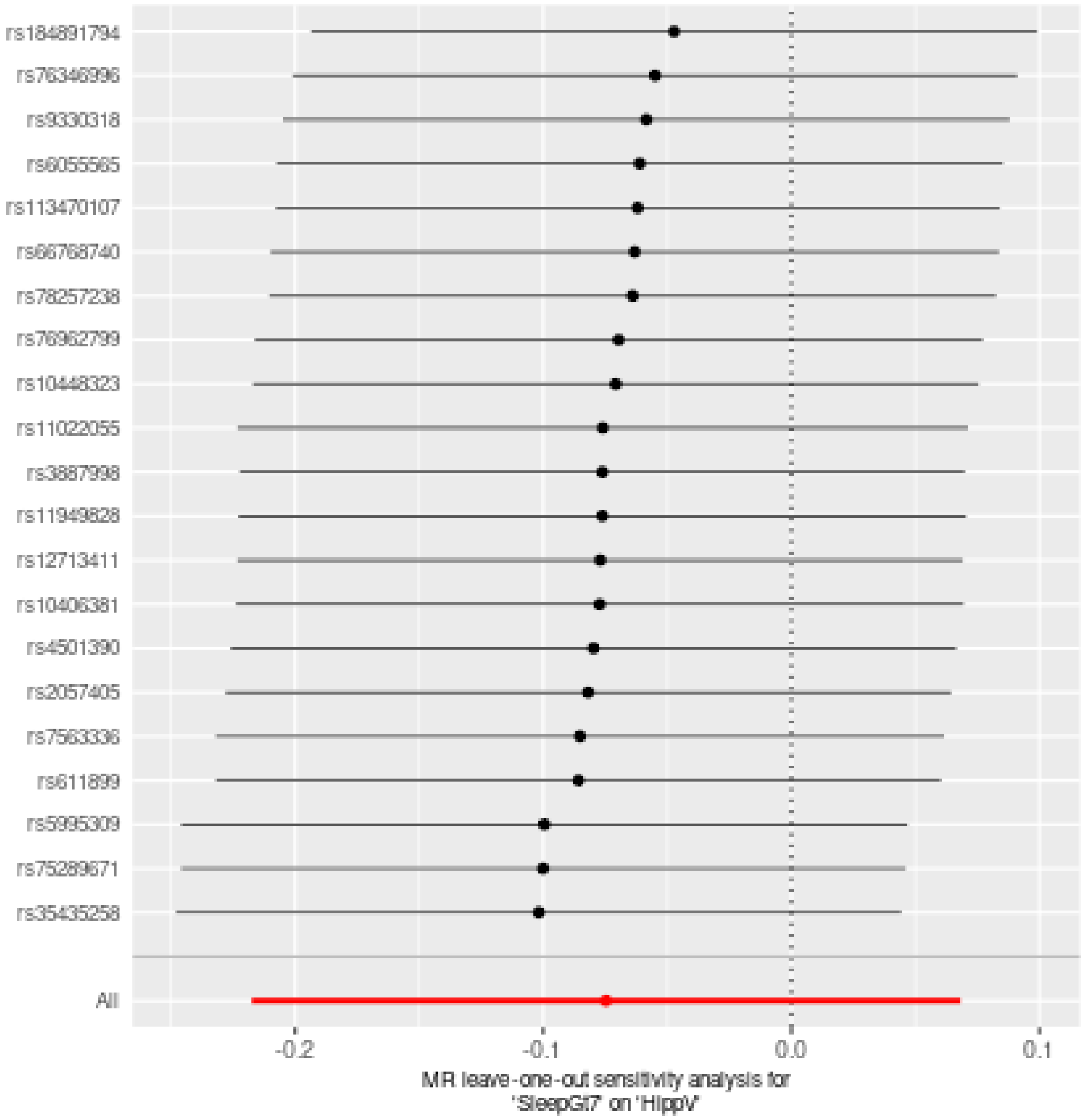

### Two sample MR report

---

#### SleepGt7 against ICV

Date: **24 september, 2021**

---

##### Results from two sample MR:

| method | nsnp | b | se | pval |
|---|---|---|---|---|
| MR Egger | 21 | -0.0290303 | 0.1581733 | 0.8563230 |
| Weighted median | 21 | -0.0036594 | 0.1063354 | 0.9725472 |
| Inverse variance weighted | 21 | 0.0243909 | 0.0760816 | 0.7485224 |
| Simple mode | 21 | 0.0004752 | 0.1914223 | 0.9980439 |
| Weighted mode | 21 | -0.0031448 | 0.1987966 | 0.9875353 |

---

##### Heterogeneity tests

| method | Q | Q_df | Q_pval |
|---|---|---|---|
| MR Egger | 12.19716 | 19 | 0.8770258 |
| Inverse variance weighted | 12.34556 | 20 | 0.9036687 |

---

##### Test for directional horizontal pleiotropy

| egger_intercept | se | pval |
|---|---|---|
| 0.0019644 | 0.0050993 | 0.7043465 |

---

##### Test that the exposure is upstream of the outcome

| snp_r2.exposure | snp_r2.outcome | correct_causal_direction | steiger_pval |
|---|---|---|---|
| 0.0052712 | 0.0005336 | TRUE | 0 |

Note - R^2 values are approximate

---

**Forest plot of single SNP MR**

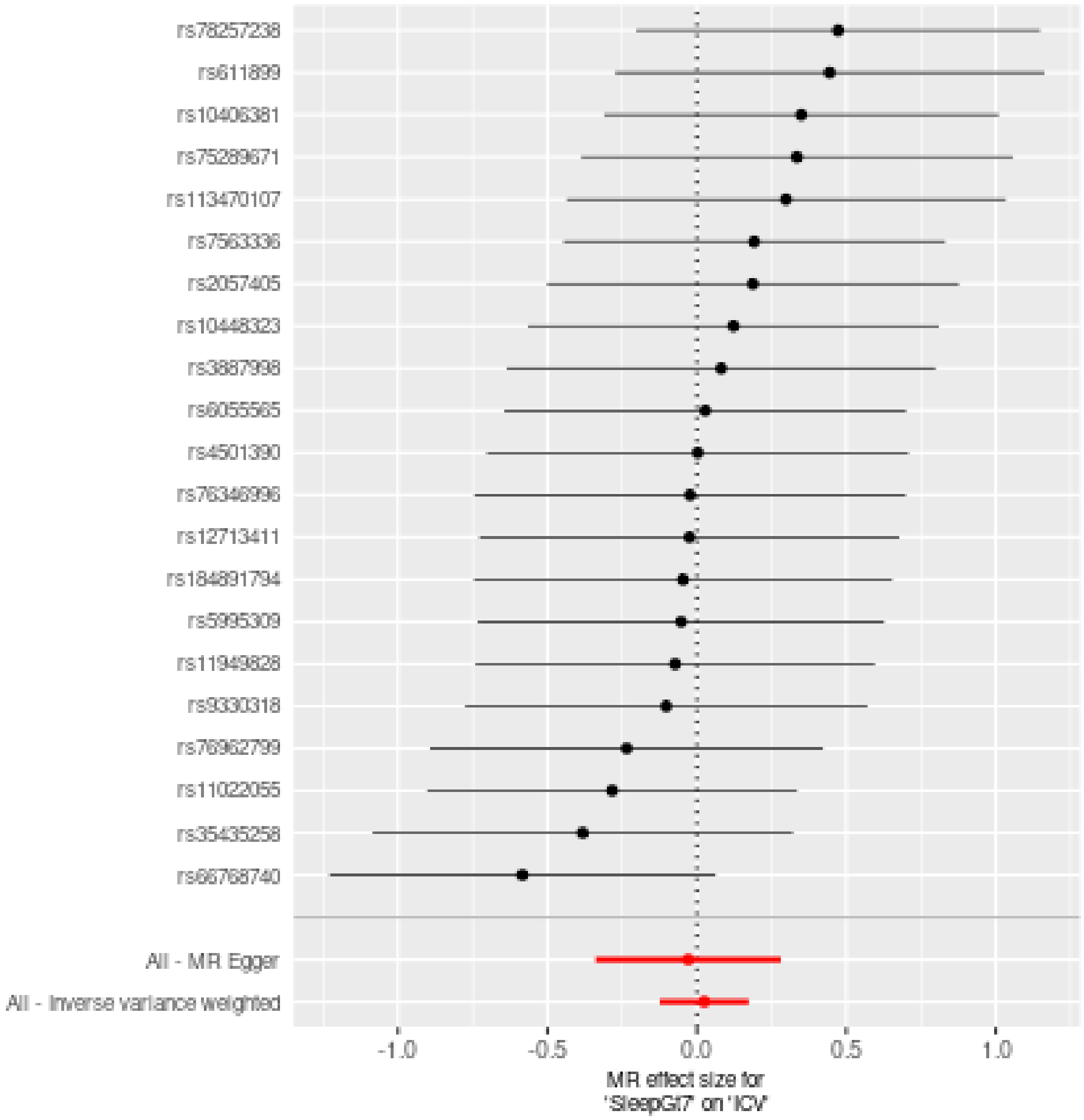

**Comparison of results using different MR methods**

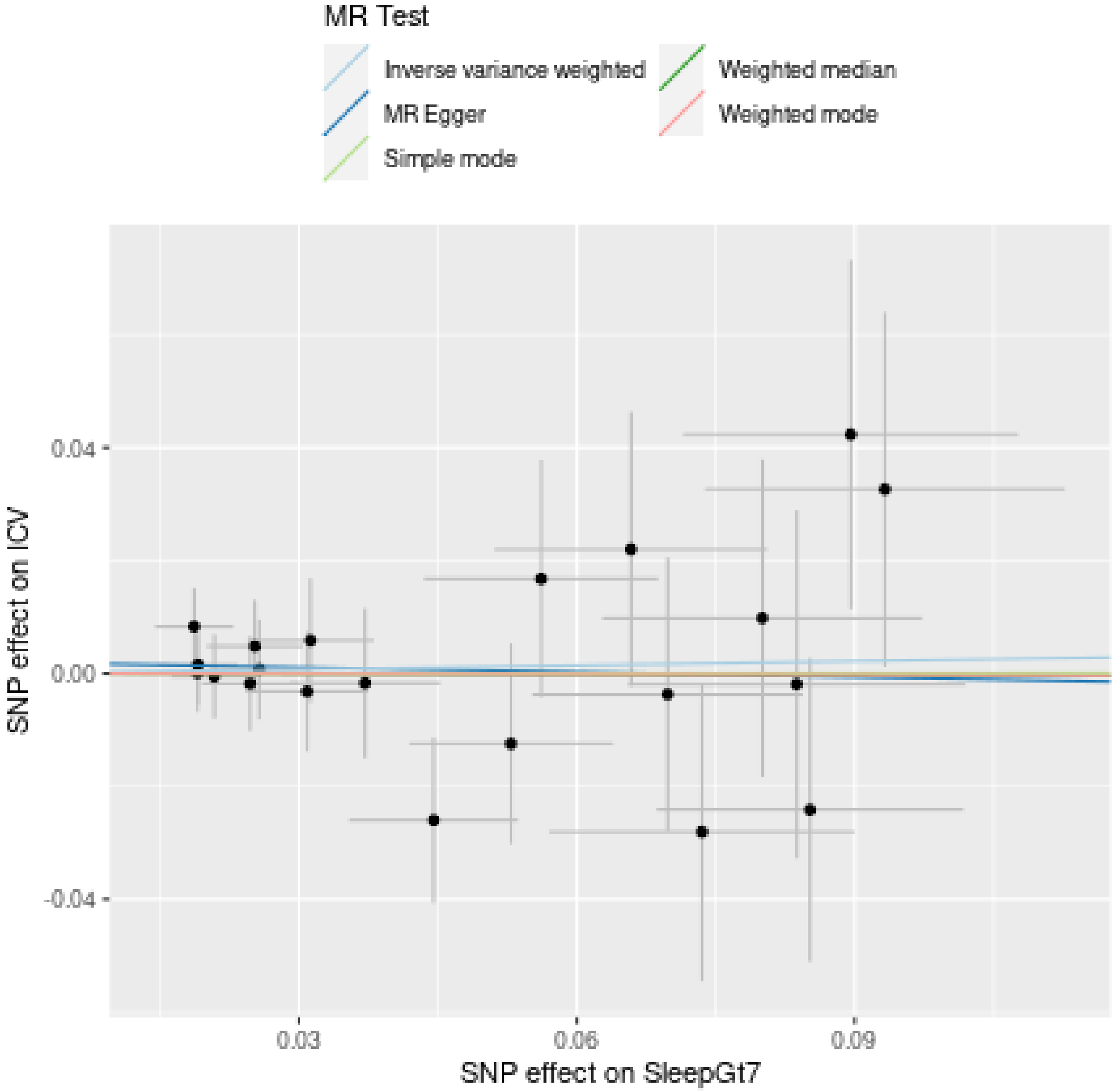

**Funnel plot**

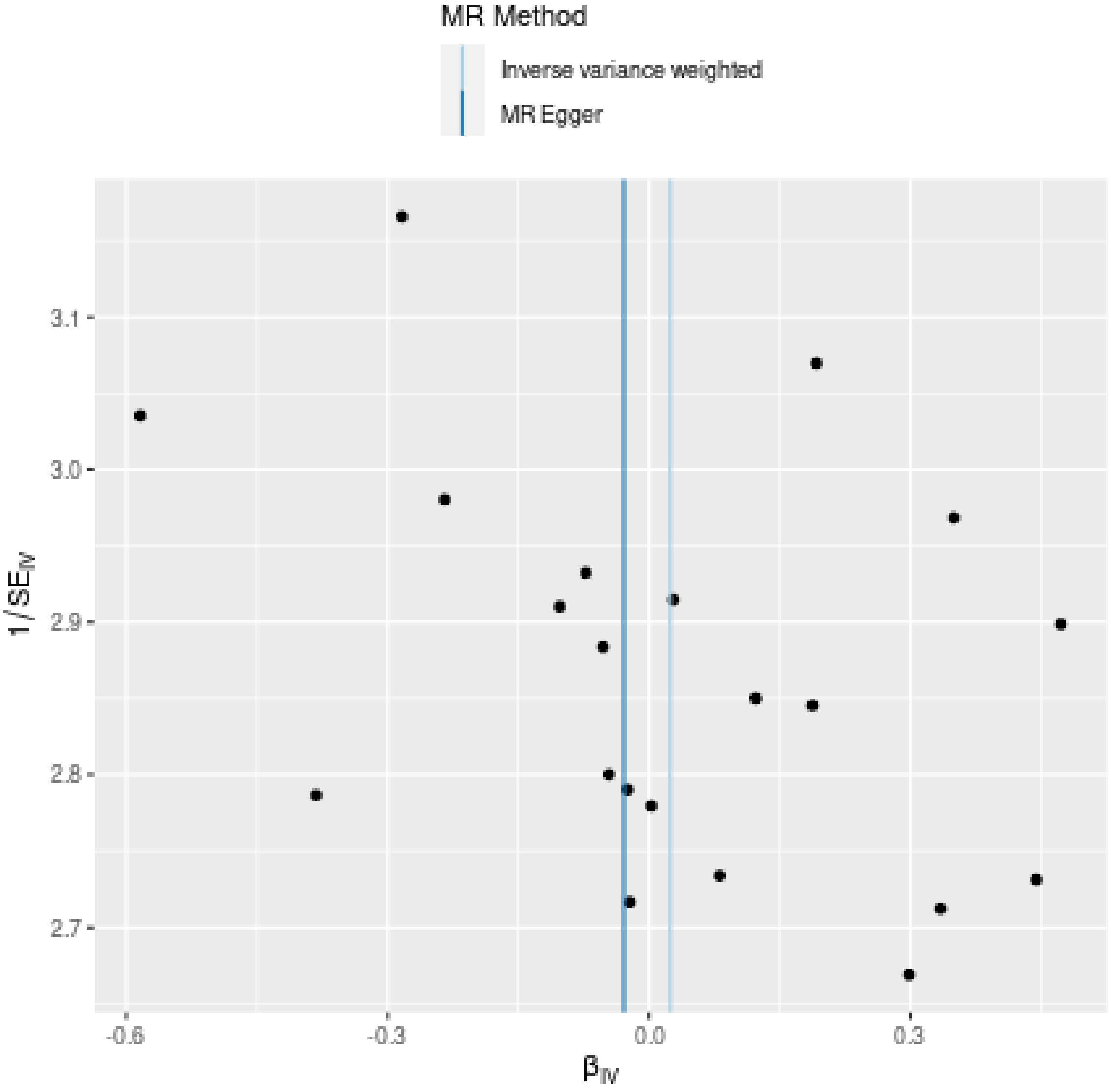
MR Method
Inverse variance weighted
MR Egger
3.1
3.0
2.9
2.8
2.7
1/SE$_{IV}$
-0.6
-0.3
0.0
0.3
β$_{IV}$

**Leave-one-out sensitivity analysis**

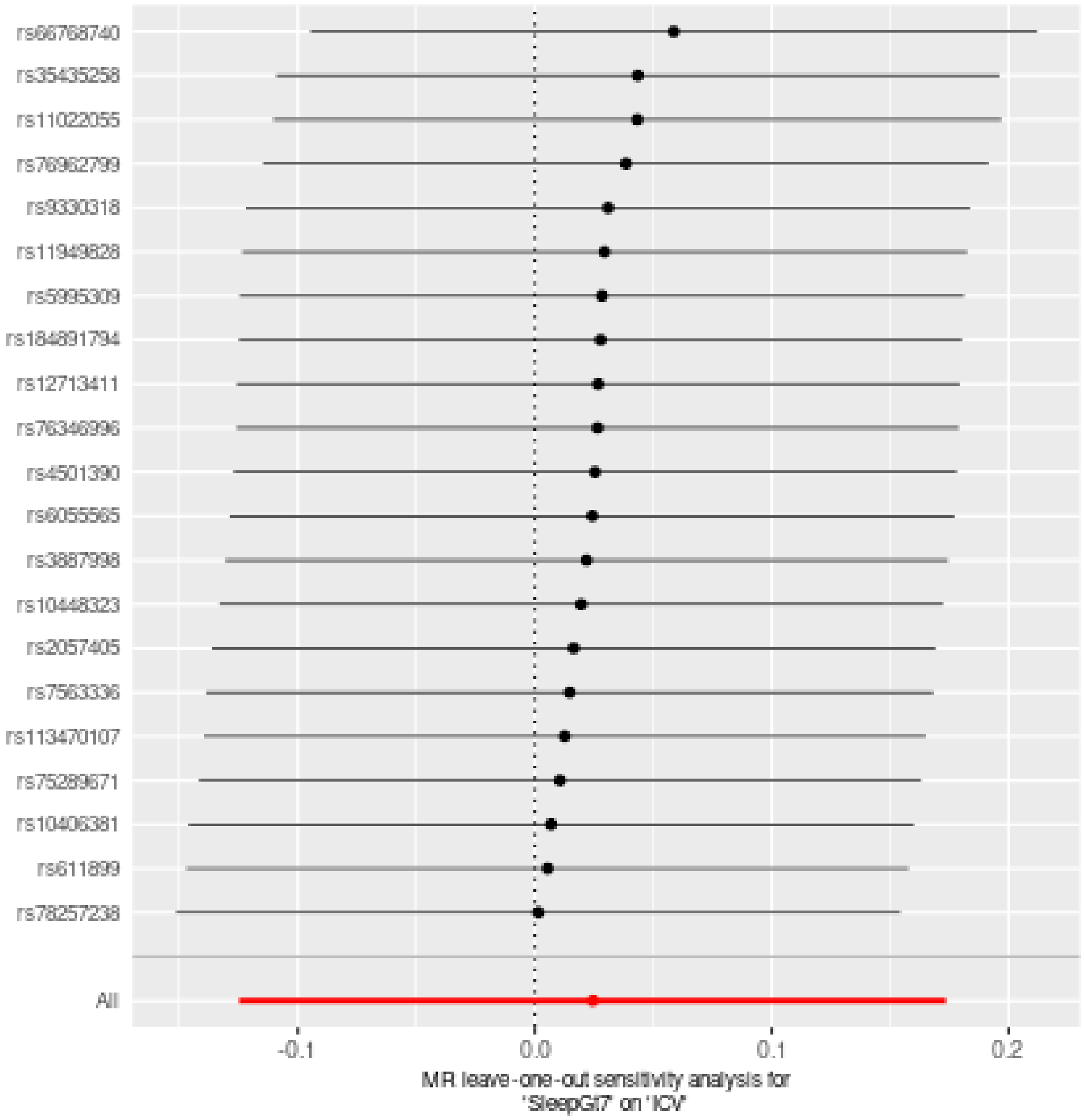

### Two sample MR report

---

#### SleepGt7 against TGV

Date: **24 september, 2021**

---

##### Results from two sample MR:

| method | nsnp | b | se | pval |
|---|---|---|---|---|
| MR Egger | 21 | -0.1411426 | 0.0960387 | 0.1580233 |
| Weighted median | 21 | -0.0374634 | 0.0657610 | 0.5688877 |
| Inverse variance weighted | 21 | -0.0540507 | 0.0461293 | 0.2413081 |
| Simple mode | 21 | 0.0416703 | 0.1285359 | 0.7491586 |
| Weighted mode | 21 | 0.0566952 | 0.1393968 | 0.6885346 |

---

##### Heterogeneity tests

| method | Q | Q_df | Q_pval |
|---|---|---|---|
| MR Egger | 16.74293 | 19 | 0.6072791 |
| Inverse variance weighted | 17.81191 | 20 | 0.5997972 |

---

##### Test for directional horizontal pleiotropy

| egger_intercept | se | pval |
|---|---|---|
| 0.0031989 | 0.0030939 | 0.3141566 |

---

##### Test that the exposure is upstream of the outcome

| snp_r2.exposure | snp_r2.outcome | correct_causal_direction | steiger_pval |
|---|---|---|---|
| 0.0052712 | 0.0010403 | TRUE | 0 |

Note - R^2 values are approximate

---

**Forest plot of single SNP MR**

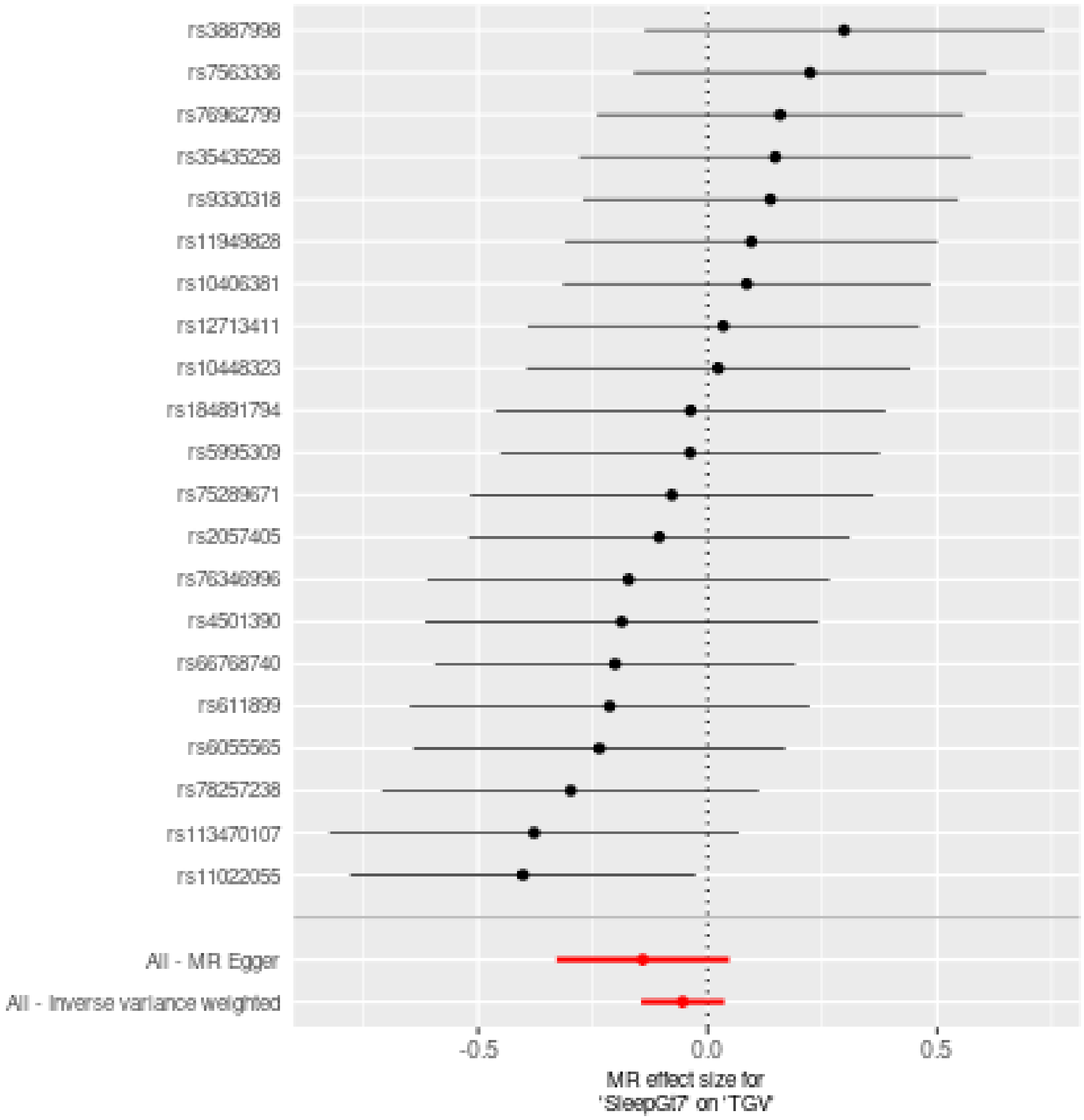

**Comparison of results using different MR methods**

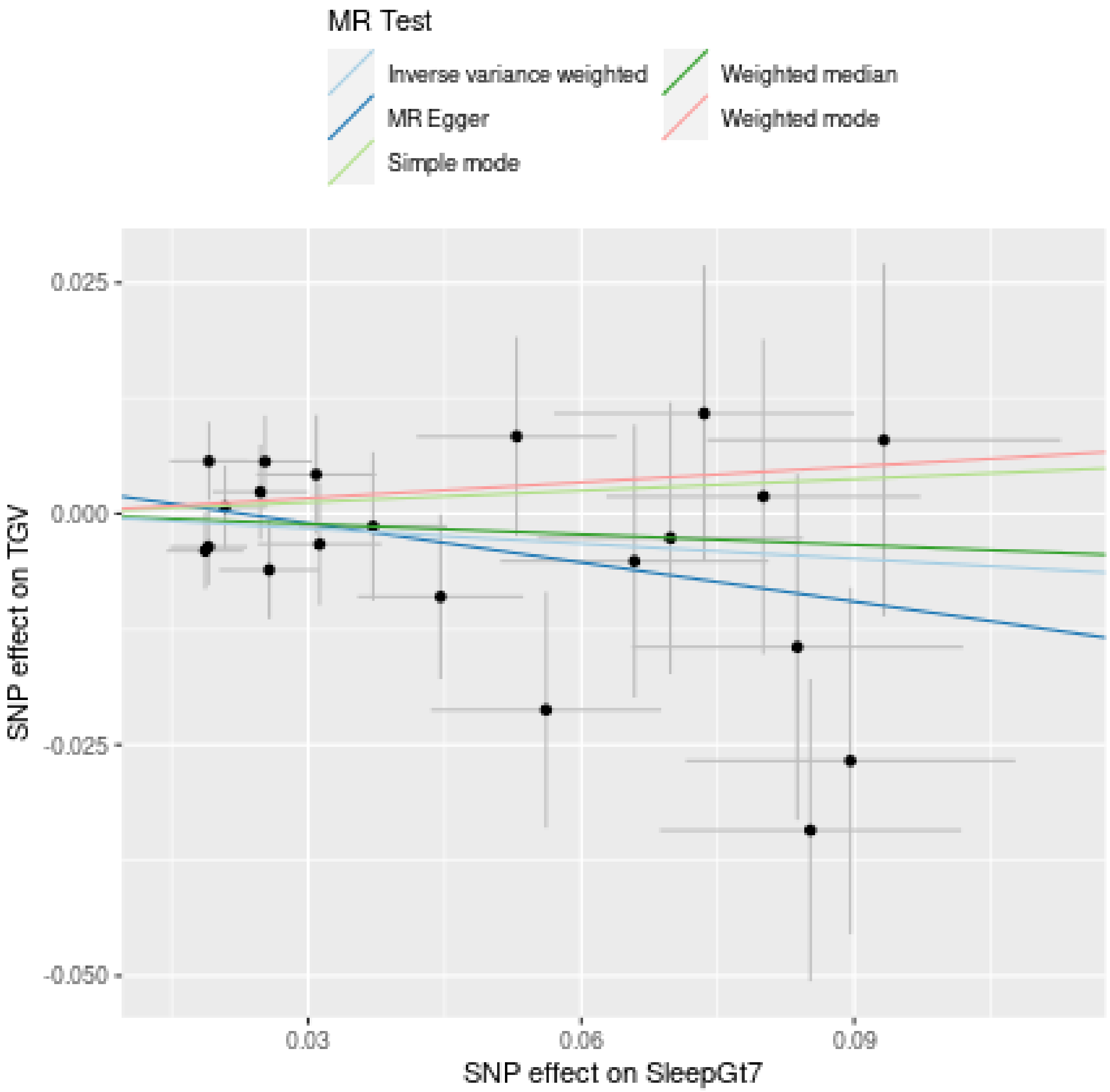

**Funnel plot**

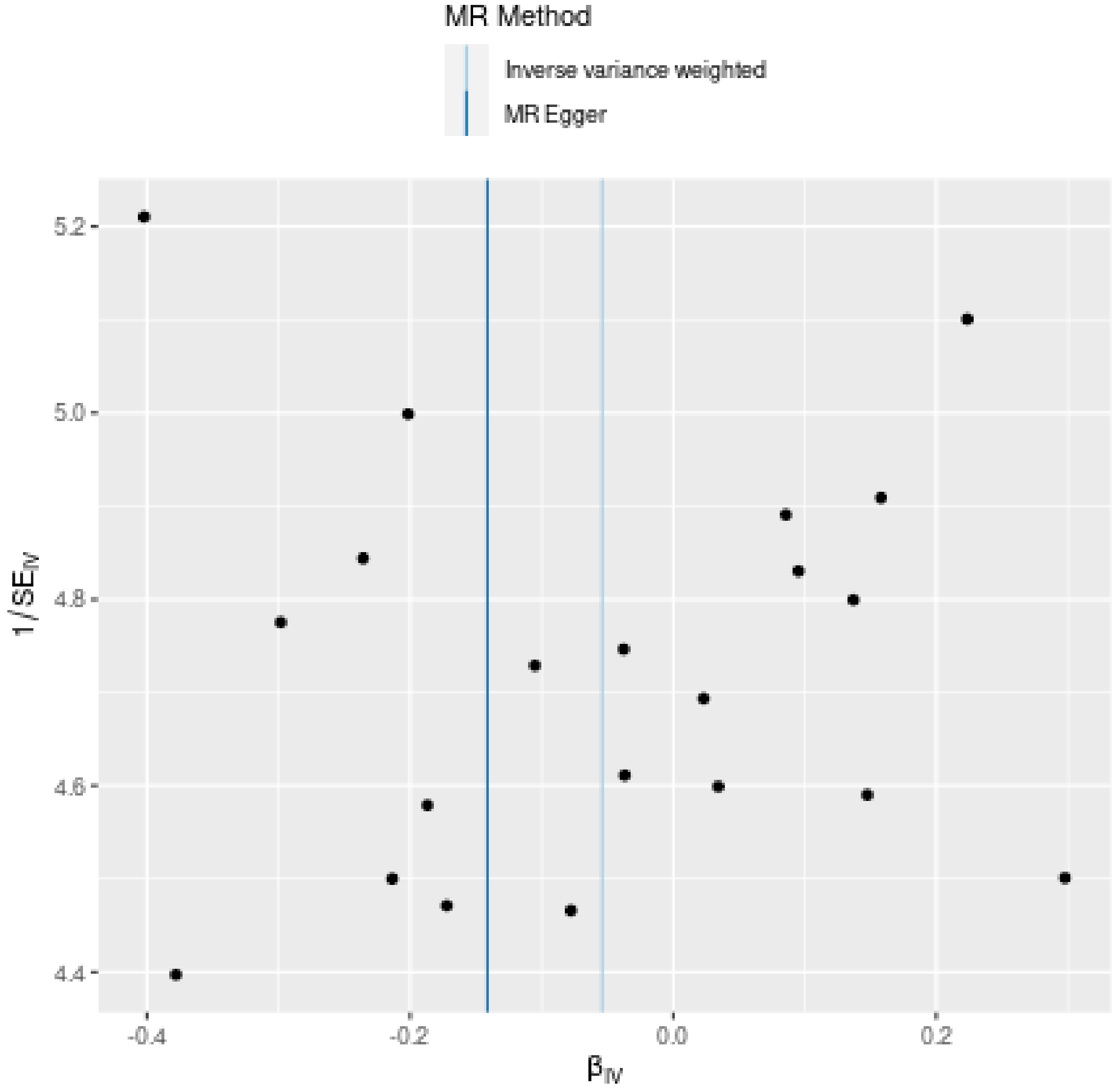
MR Method
Inverse variance weighted
MR Egger
5.2
5.0
4.8
4.6
4.4
1/SE$_{IV}$
-0.4
-0.2
0.0
0.2
β$_{IV}$

**Leave-one-out sensitivity analysis**

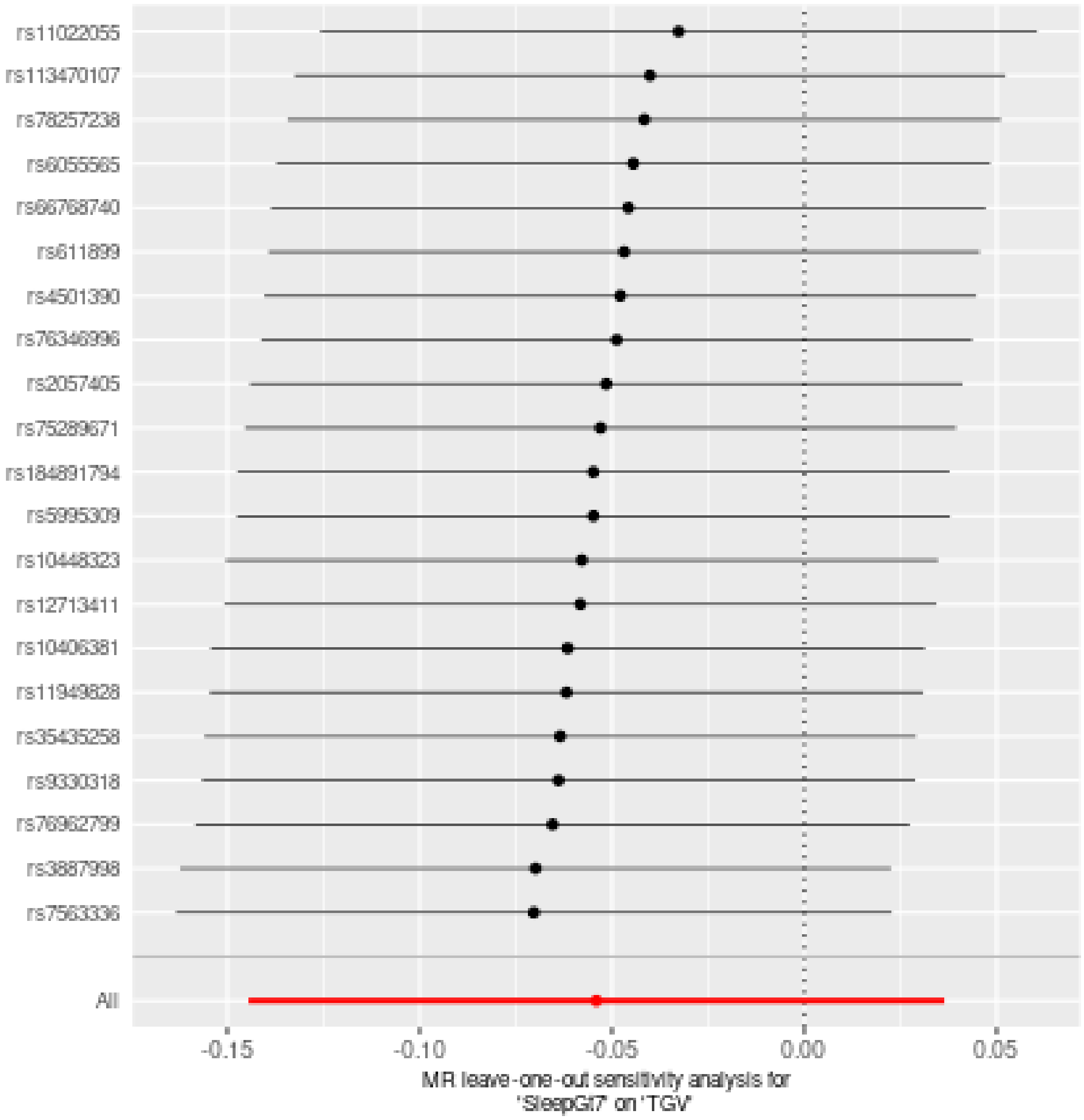

### Two sample MR report

---

#### SleepLe7 against HippV

Date: **24 september, 2021**

---

##### Results from two sample MR:

| method | nsnp | b | se | pval |
|---|---|---|---|---|
| MR Egger | 30 | -0.0678012 | 0.2229604 | 0.7633039 |
| Weighted median | 30 | -0.0519938 | 0.1143050 | 0.6492035 |
| Inverse variance weighted | 30 | 0.0022258 | 0.0790990 | 0.9775514 |
| Simple mode | 30 | -0.1871948 | 0.2508017 | 0.4614427 |
| Weighted mode | 30 | -0.2103634 | 0.2740287 | 0.4488894 |

---

##### Heterogeneity tests

| method | Q | Q_df | Q_pval |
|---|---|---|---|
| MR Egger | 29.85086 | 28 | 0.3703822 |
| Inverse variance weighted | 29.97172 | 29 | 0.4154165 |

---

##### Test for directional horizontal pleiotropy

| egger_intercept | se | pval |
|---|---|---|
| 0.0015696 | 0.0046618 | 0.7388584 |

---

##### Test that the exposure is upstream of the outcome

| snp_r2.exposure | snp_r2.outcome | correct_causal_direction | steiger_pval |
|---|---|---|---|
| 0.0037888 | 0.0010377 | TRUE | 3.2e-06 |

Note - R^2 values are approximate

---

**Forest plot of single SNP MR**

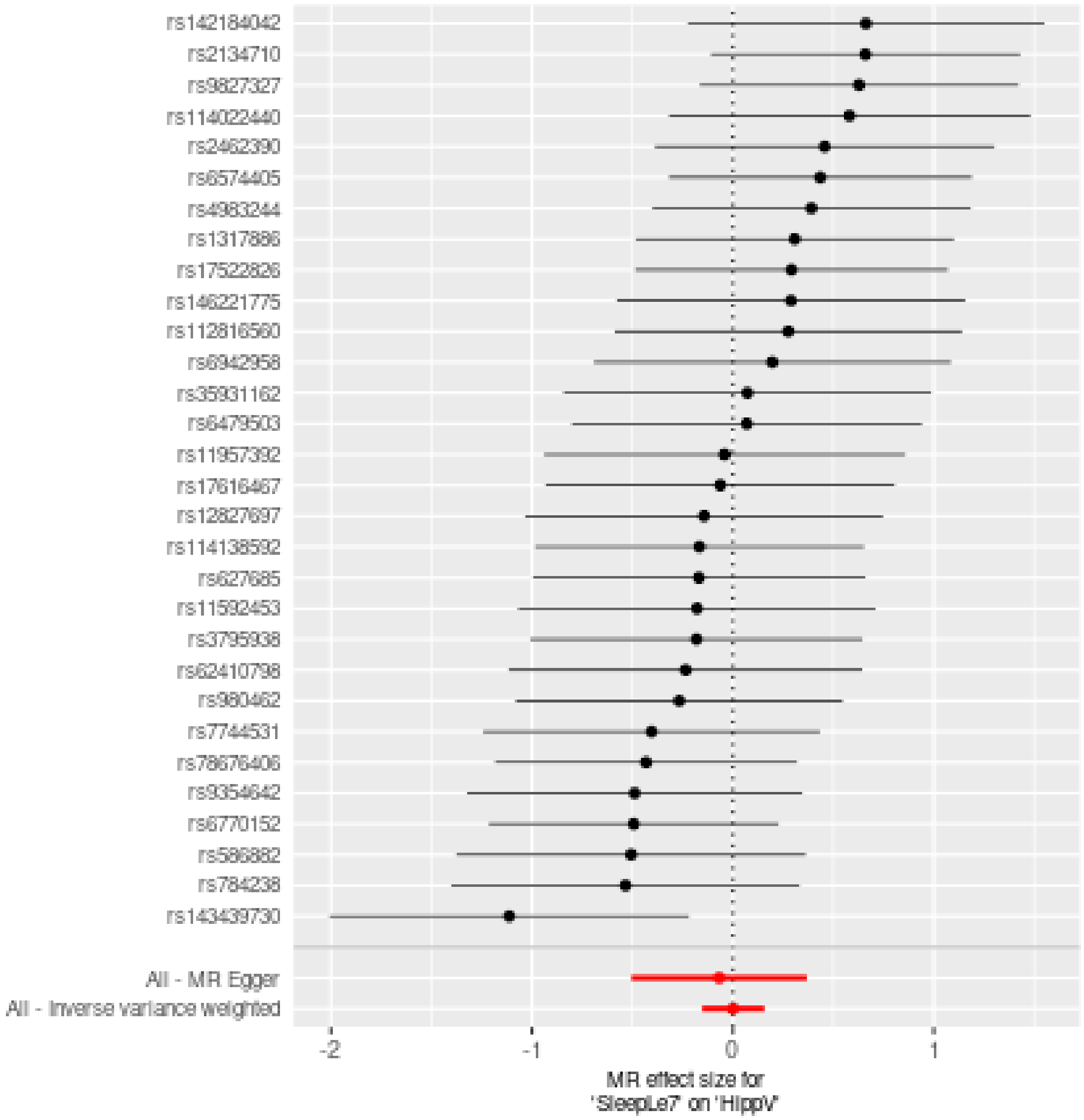

**Comparison of results using different MR methods**

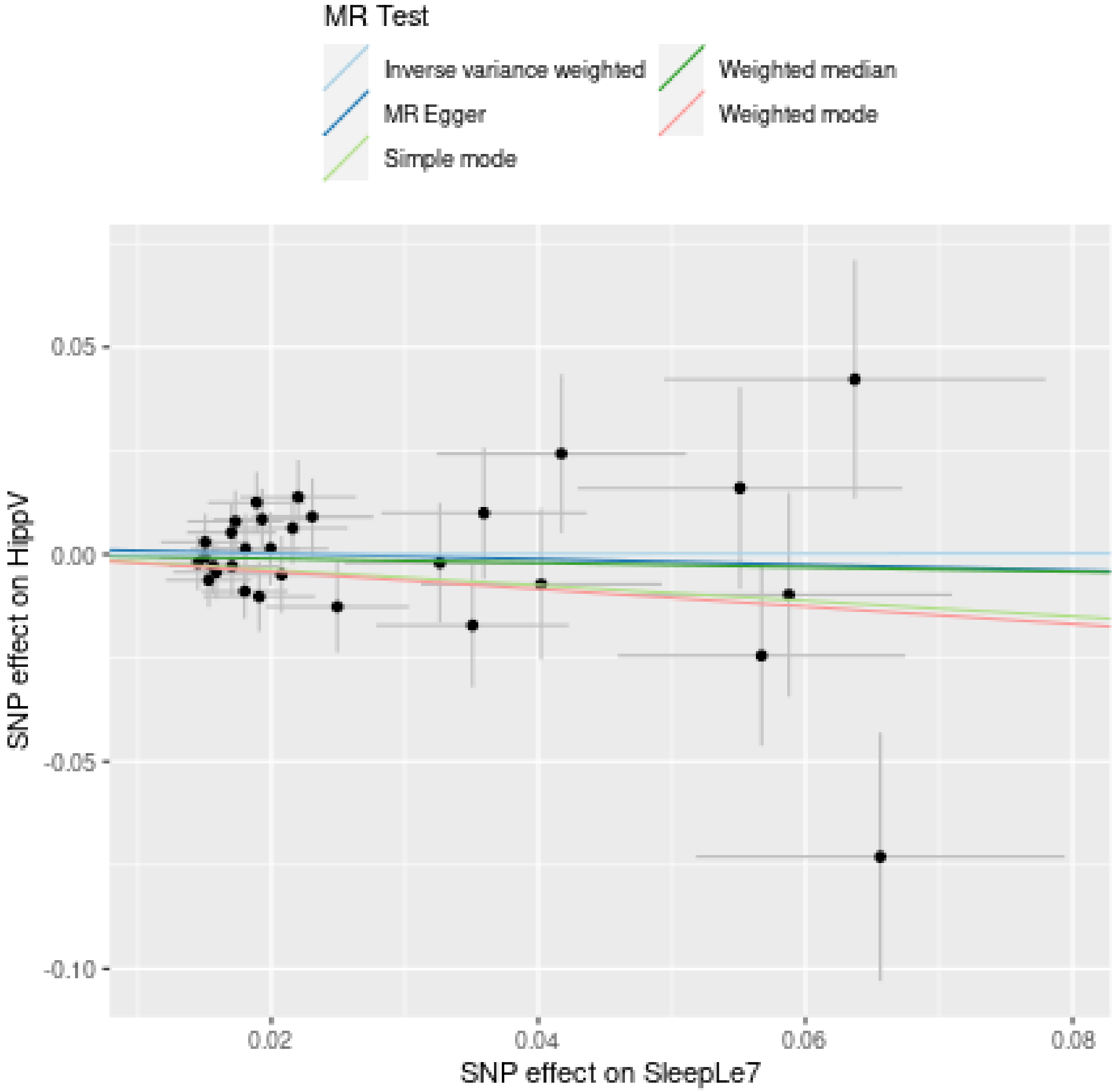

**Funnel plot**

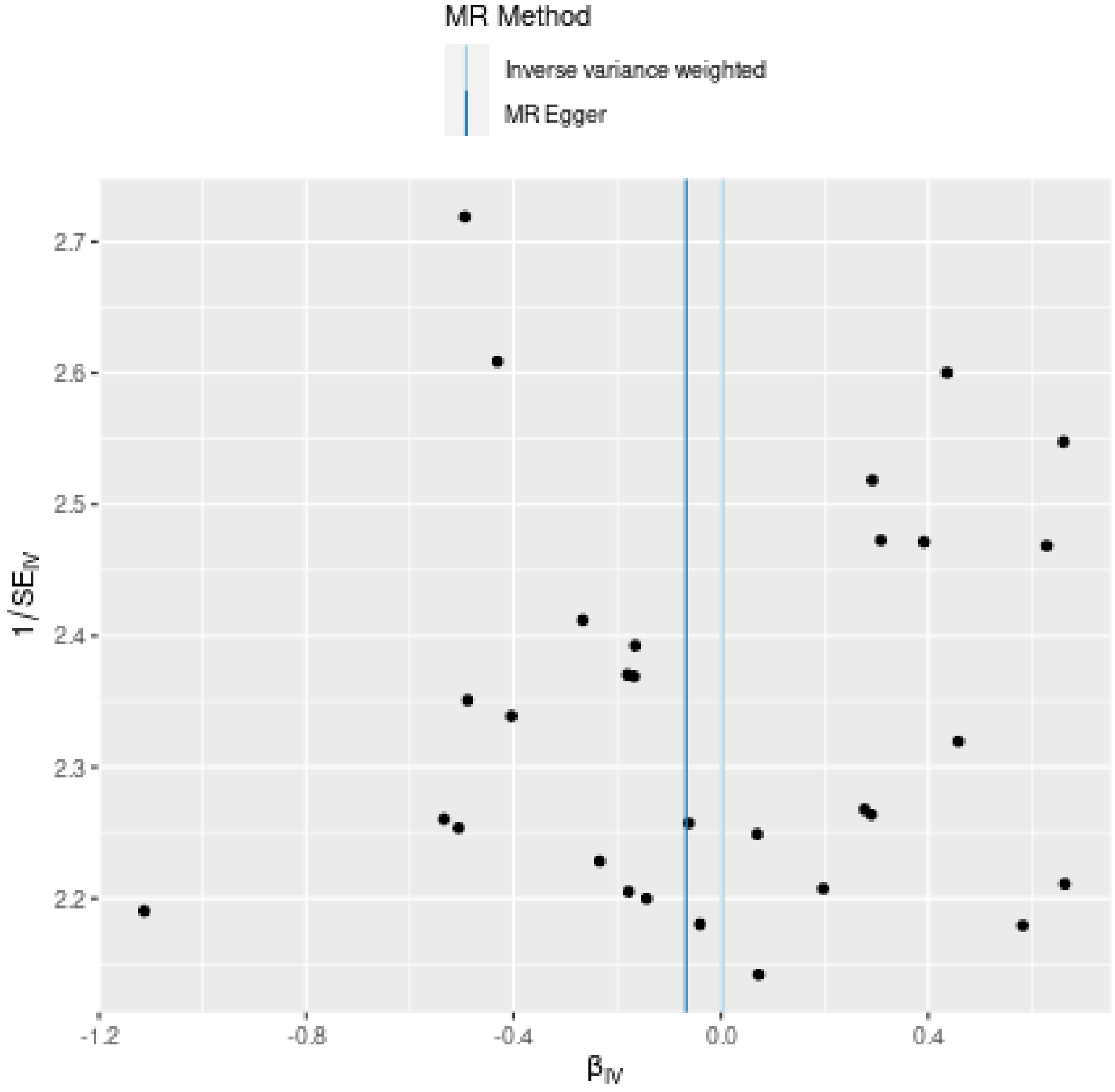
MR Method
Inverse variance weighted
MR Egger
2.7
2.6
2.5
2.4
2.3
2.2
1/SE$_{IV}$
-1.2
-0.8
-0.4
0.0
0.4
β$_{IV}$

**Leave-one-out sensitivity analysis**

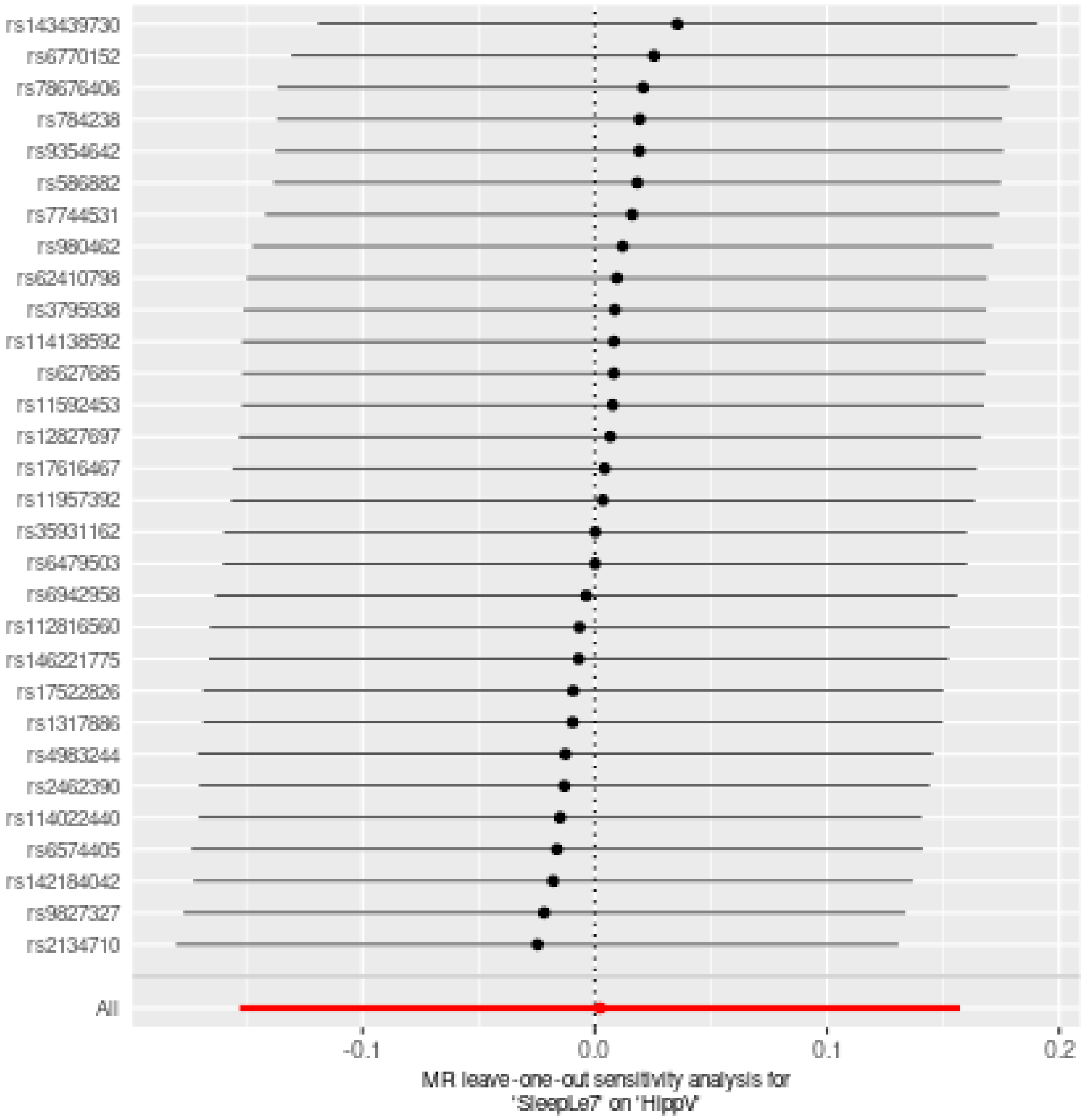

### Two sample MR report

---

#### SleepLe7 against ICV

Date: **24 september, 2021**

---

##### Results from two sample MR:

| method | nsnp | b | se | pval |
|---|---|---|---|---|
| MR Egger | 30 | 0.0849180 | 0.2261827 | 0.7101625 |
| Weighted median | 30 | -0.0461790 | 0.1143781 | 0.6864042 |
| Inverse variance weighted | 30 | 0.0318054 | 0.0813939 | 0.6959755 |
| Simple mode | 30 | -0.0890807 | 0.2332230 | 0.7052794 |
| Weighted mode | 30 | -0.0890807 | 0.2263548 | 0.6967960 |

---

##### Heterogeneity tests

| method | Q | Q_df | Q_pval |
|---|---|---|---|
| MR Egger | 28.07049 | 28 | 0.4607169 |
| Inverse variance weighted | 28.13401 | 29 | 0.5107562 |

---

##### Test for directional horizontal pleiotropy

| egger_intercept | se | pval |
|---|---|---|
| -0.0011905 | 0.0047292 | 0.8030881 |

---

##### Test that the exposure is upstream of the outcome

| snp_r2.exposure | snp_r2.outcome | correct_causal_direction | steiger_pval |
|---|---|---|---|
| 0.0037888 | 0.0010717 | TRUE | 4.8e-06 |

Note - R^2 values are approximate

---

**Forest plot of single SNP MR**

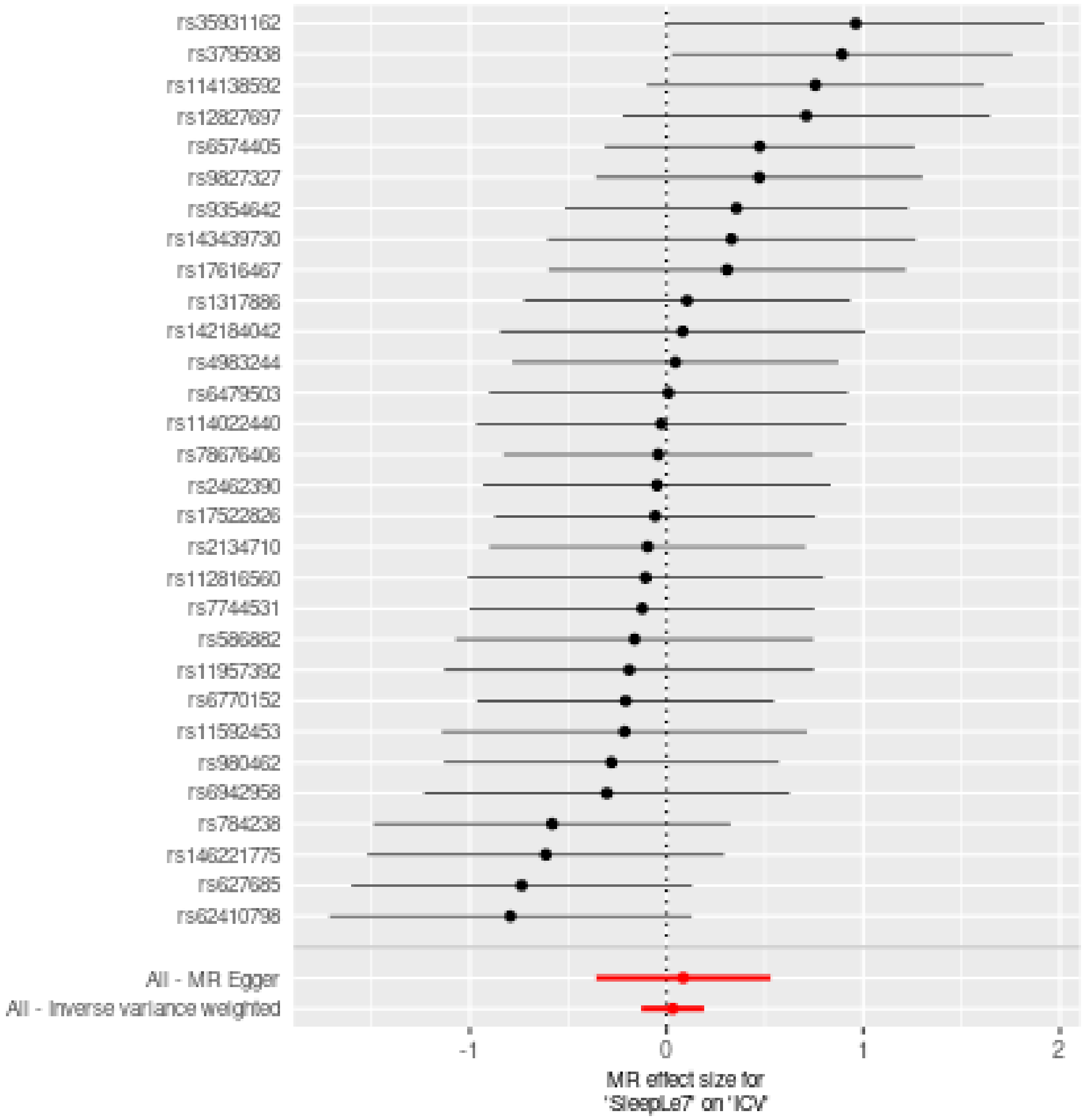

**Comparison of results using different MR methods**

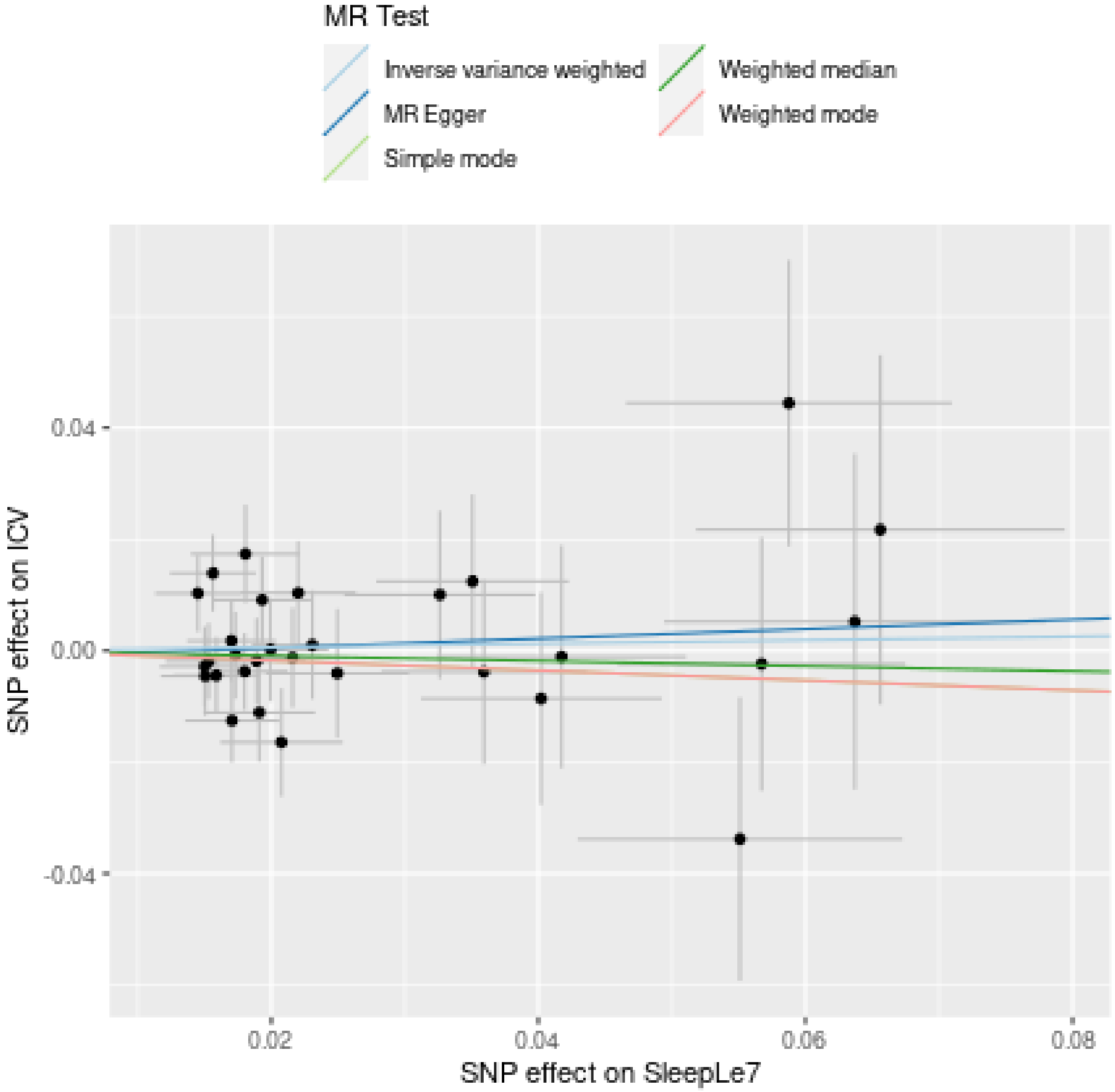

**Funnel plot**

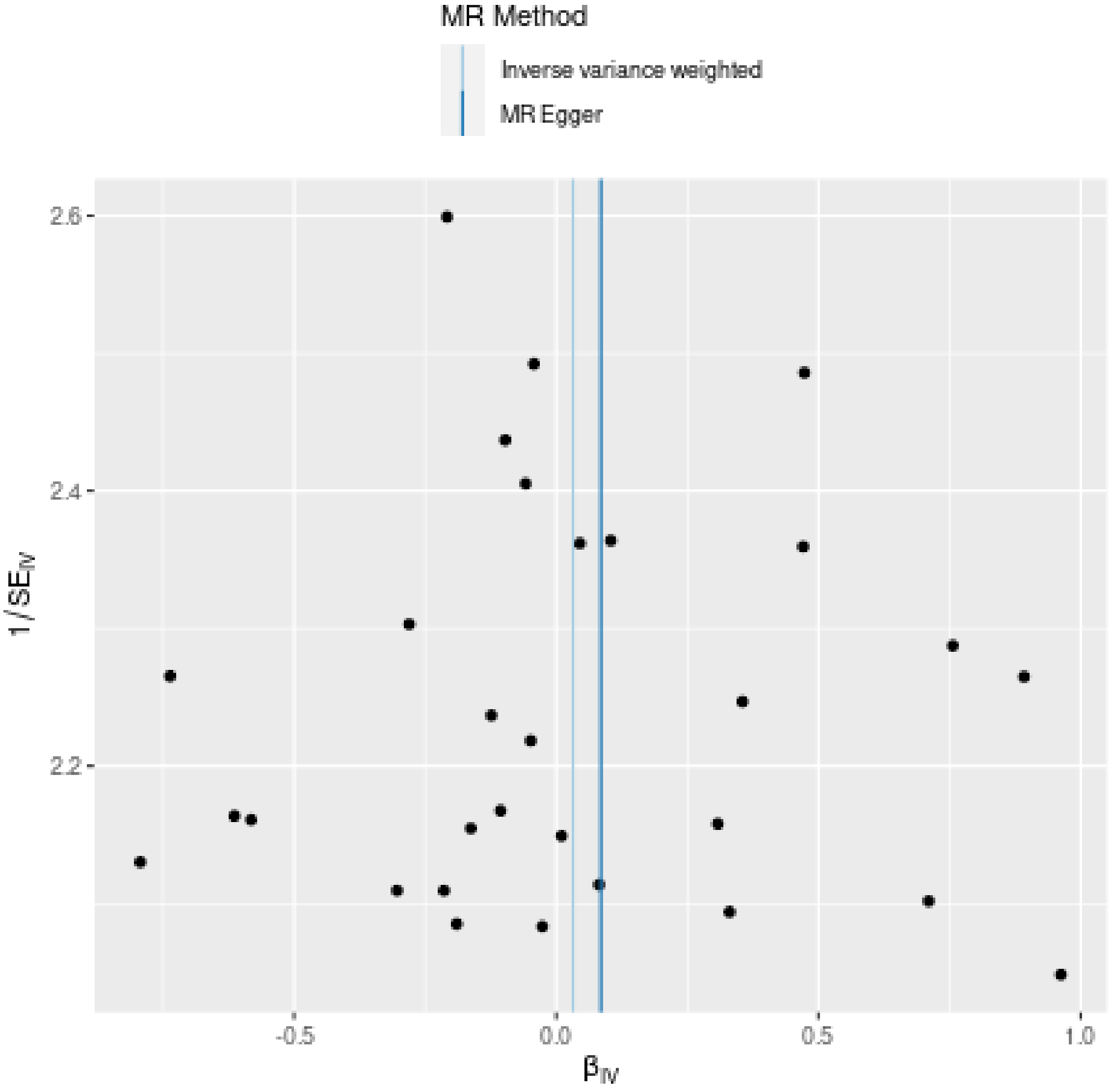
MR Method
Inverse variance weighted
MR Egger
2.6
2.4
2.2
1/SE_IV
-0.5
0.0
0.5
1.0
β_IV

**Leave-one-out sensitivity analysis**

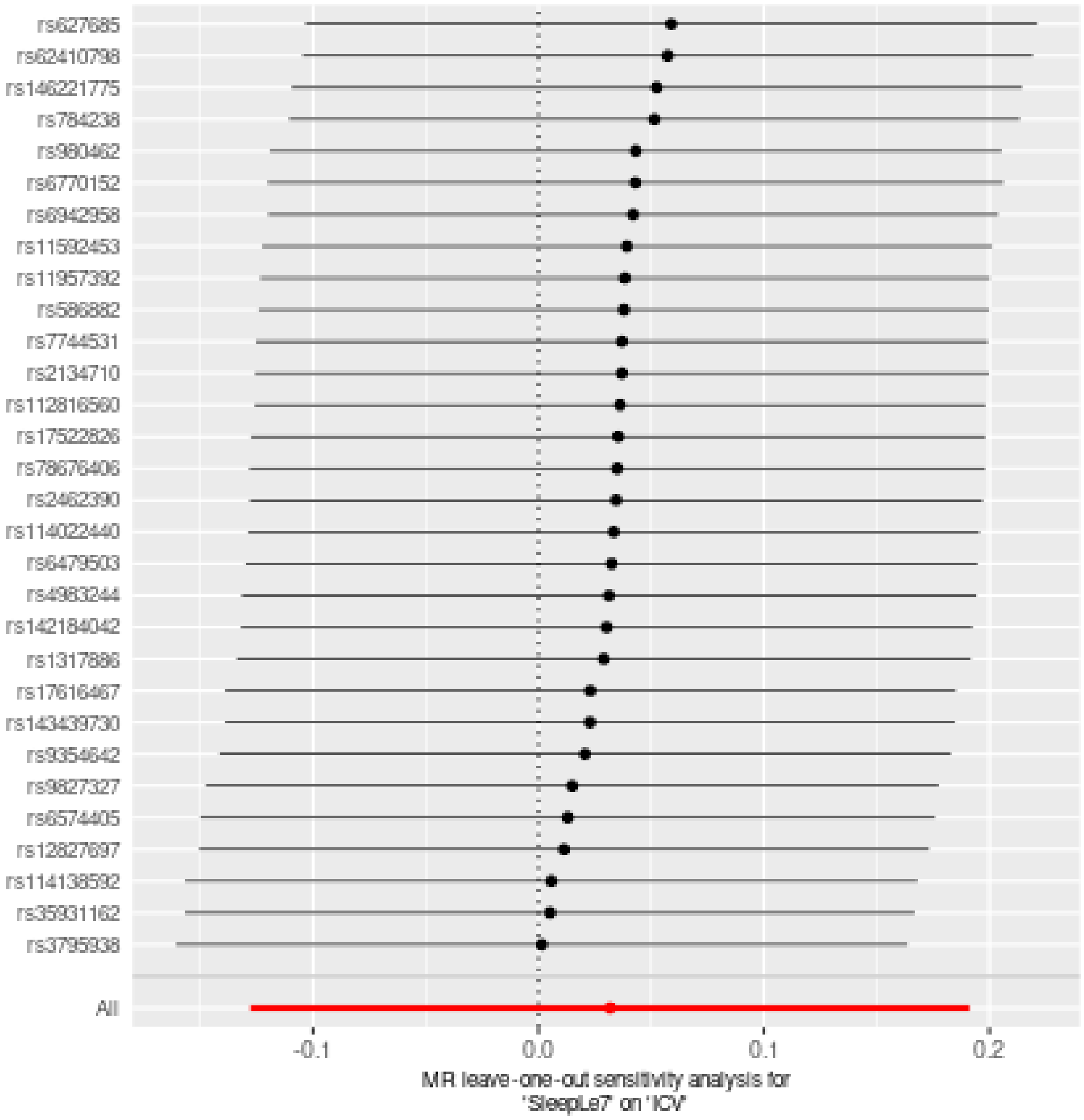

### Two sample MR report

---

#### SleepLe7 against TGV

Date: **24 september, 2021**

---

##### Results from two sample MR:

| method | nsnp | b | se | pval |
|---|---|---|---|---|
| MR Egger | 30 | -0.0961532 | 0.1379010 | 0.4913878 |
| Weighted median | 30 | -0.0319099 | 0.0698793 | 0.6479281 |
| Inverse variance weighted | 30 | -0.0103353 | 0.0494239 | 0.8343585 |
| Simple mode | 30 | -0.0908140 | 0.1403078 | 0.5225623 |
| Weighted mode | 30 | -0.0835624 | 0.1423746 | 0.5617996 |

---

##### Heterogeneity tests

| method | Q | Q_df | Q_pval |
|---|---|---|---|
| MR Egger | 28.29881 | 28 | 0.4487004 |
| Inverse variance weighted | 28.74861 | 29 | 0.4782141 |

---

##### Test for directional horizontal pleiotropy

| egger_intercept | se | pval |
|---|---|---|
| 0.0019235 | 0.0028832 | 0.5101537 |

---

##### Test that the exposure is upstream of the outcome

| snp_r2.exposure | snp_r2.outcome | correct_causal_direction | steiger_pval |
|---|---|---|---|
| 0.0037888 | 0.0010039 | TRUE | 2.1e-06 |

Note - R^2 values are approximate

---

**Forest plot of single SNP MR**

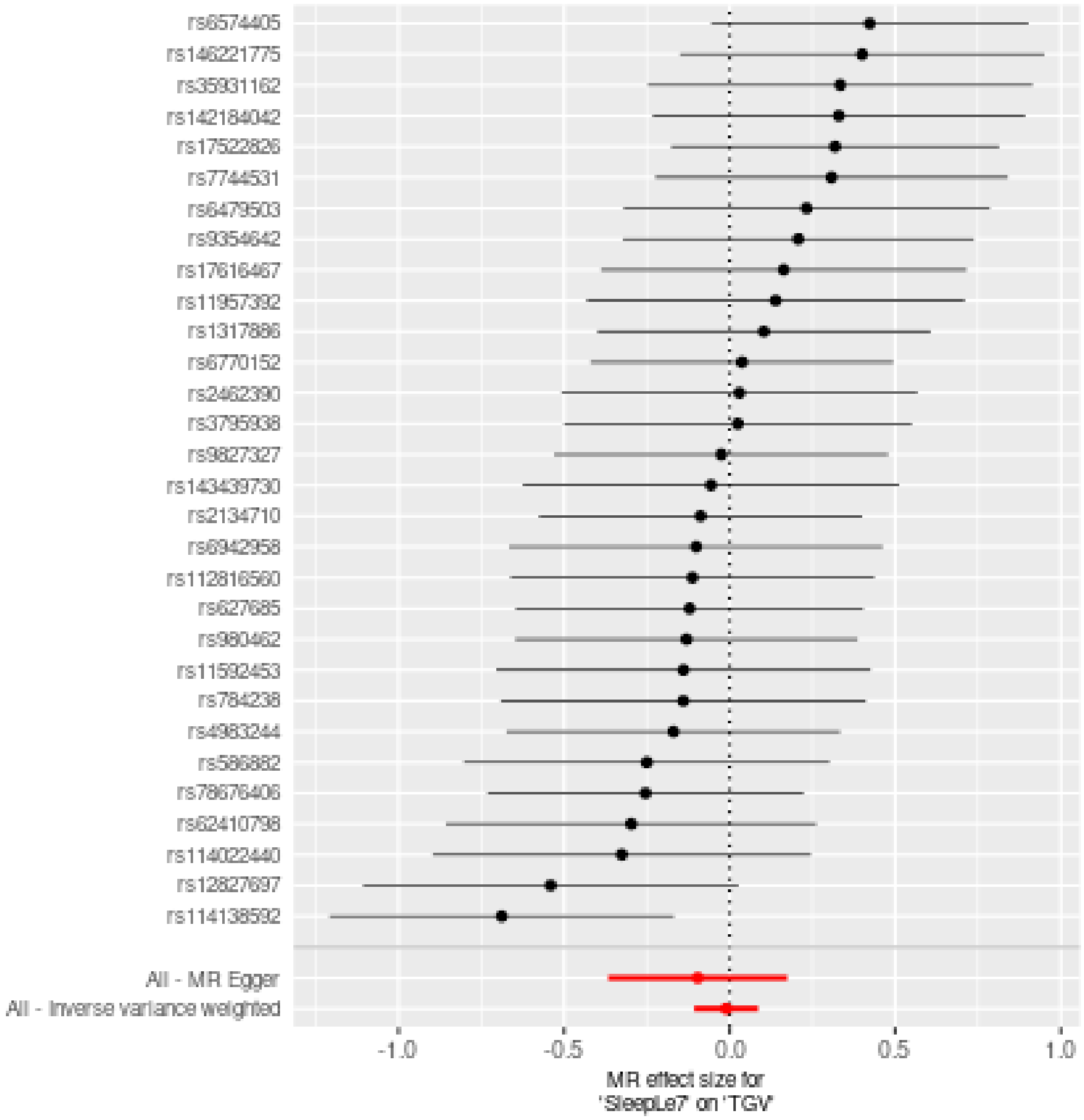

**Comparison of results using different MR methods**

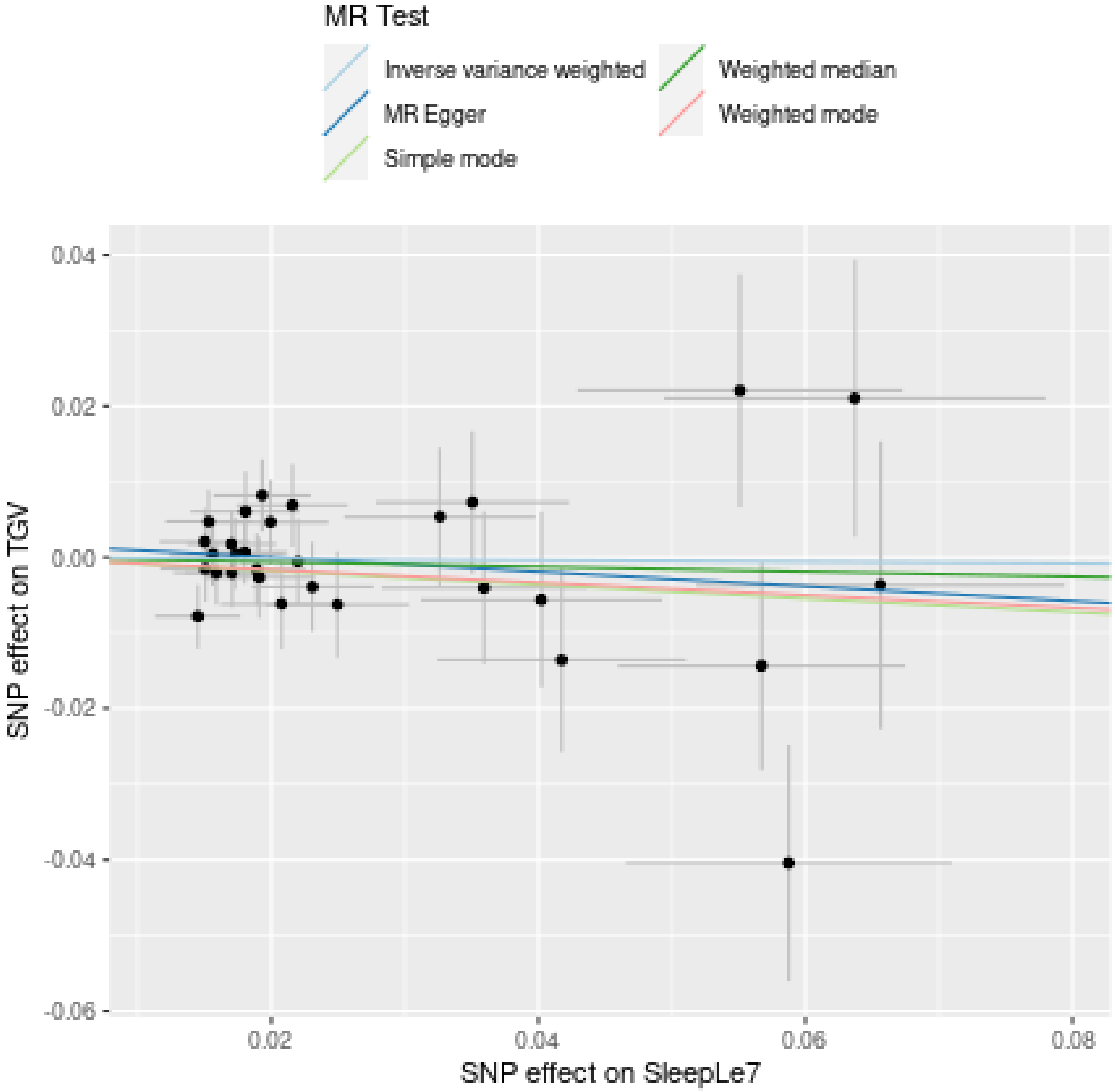

**Funnel plot**

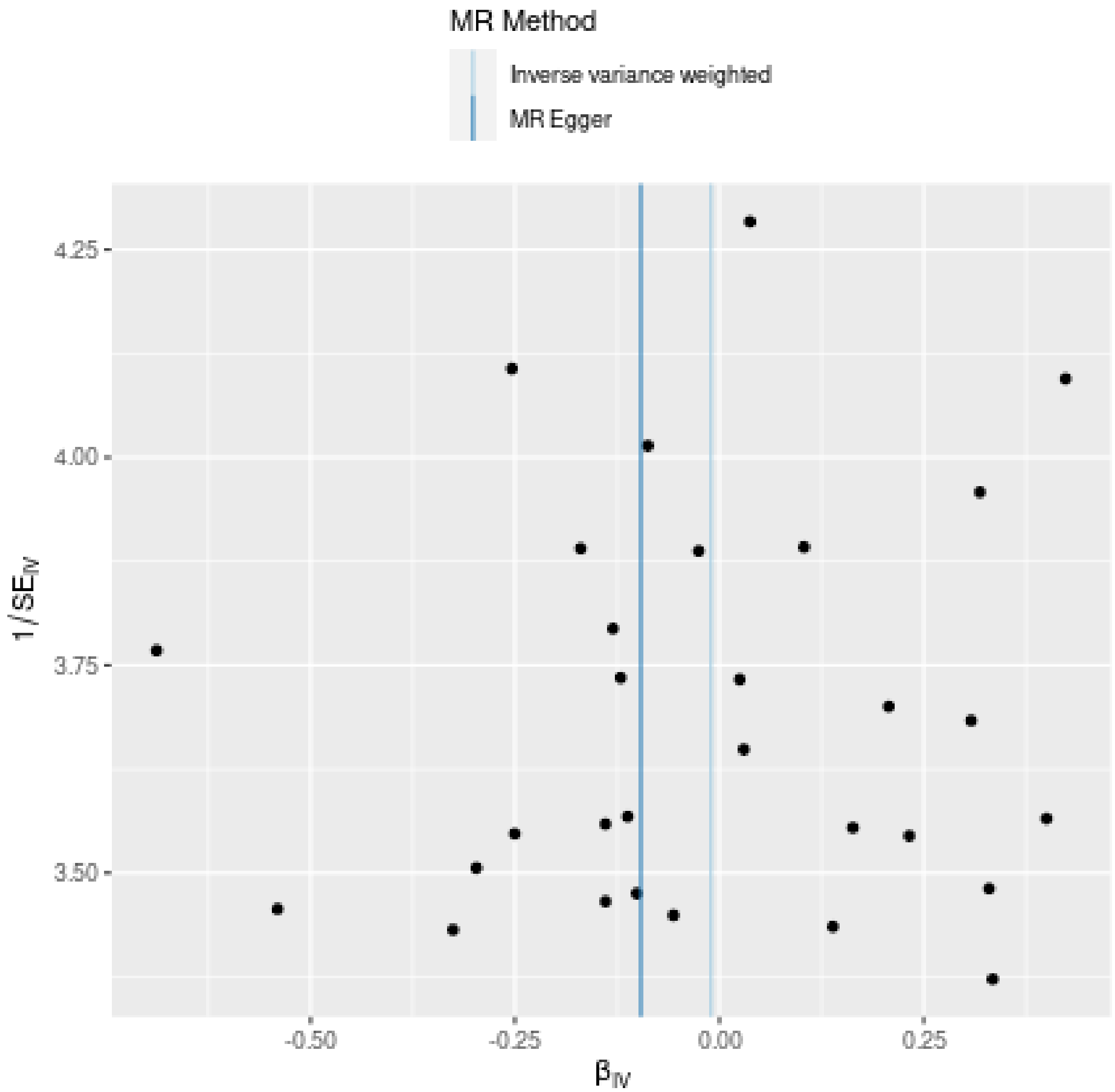

MR Method
Inverse variance weighted
MR Egger
4.25
4.00
3.75
3.50
1/SE_IV
-0.50
-0.25
0.00
0.25
β_IV

**Leave-one-out sensitivity analysis**

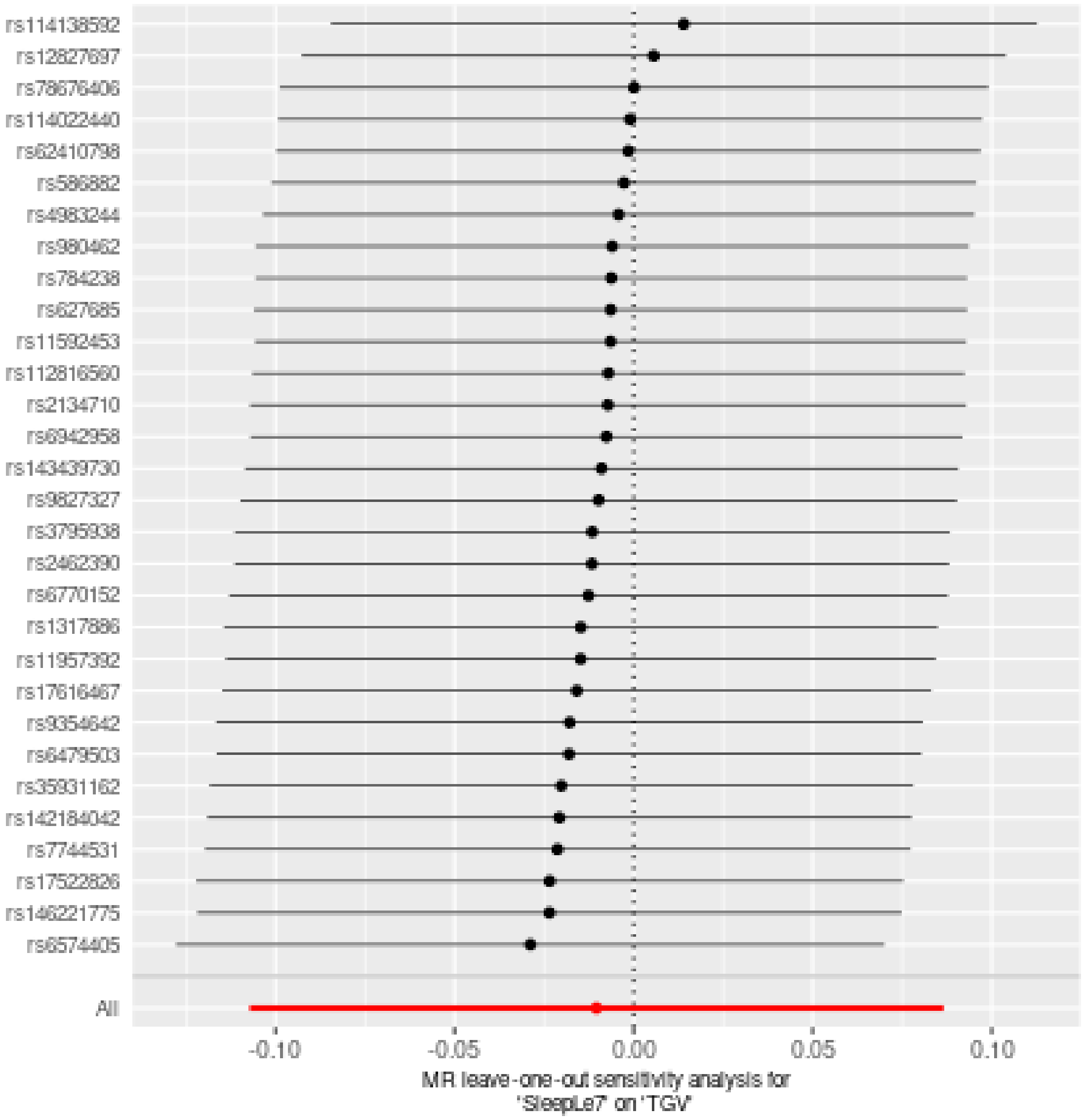

### Two sample MR report

---

#### TGV against SleepGt7

Date: **24 september, 2021**

---

##### Results from two sample MR:

| method | nsnp | b | se | pval |
|---|---|---|---|---|
| MR Egger | 67 | 0.1054601 | 0.0837270 | 0.2123280 |
| Weighted median | 67 | -0.0008322 | 0.0383481 | 0.9826867 |
| Inverse variance weighted | 67 | -0.0001759 | 0.0331697 | 0.9957690 |
| Simple mode | 67 | 0.0111666 | 0.0984704 | 0.9100569 |
| Weighted mode | 67 | 0.0147731 | 0.0977878 | 0.8803788 |

---

##### Heterogeneity tests

| method | Q | Q_df | Q_pval |
|---|---|---|---|
| MR Egger | 109.1096 | 65 | 0.0005061 |
| Inverse variance weighted | 112.2713 | 66 | 0.0003332 |

---

##### Test for directional horizontal pleiotropy

| egger_intercept | se | pval |
|---|---|---|
| -0.0032295 | 0.0023531 | 0.1746514 |

---

##### Test that the exposure is upstream of the outcome

| snp_r2.exposure | snp_r2.outcome | correct_causal_direction | steiger_pval |
|---|---|---|---|
| 0.0620032 | 0.0011137 | TRUE | 0 |

Note - R^2 values are approximate

---

**Forest plot of single SNP MR**

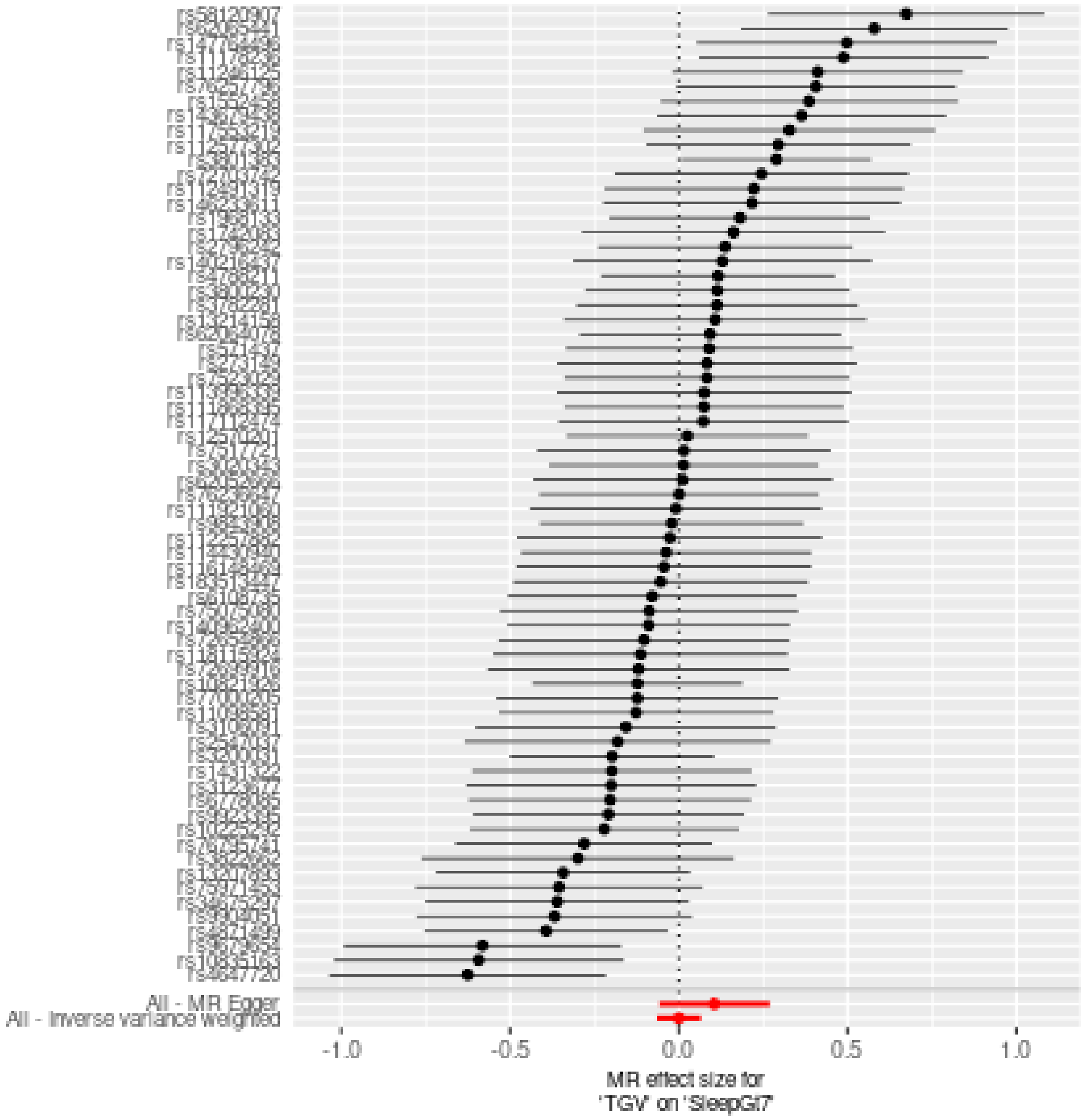

**Comparison of results using different MR methods**

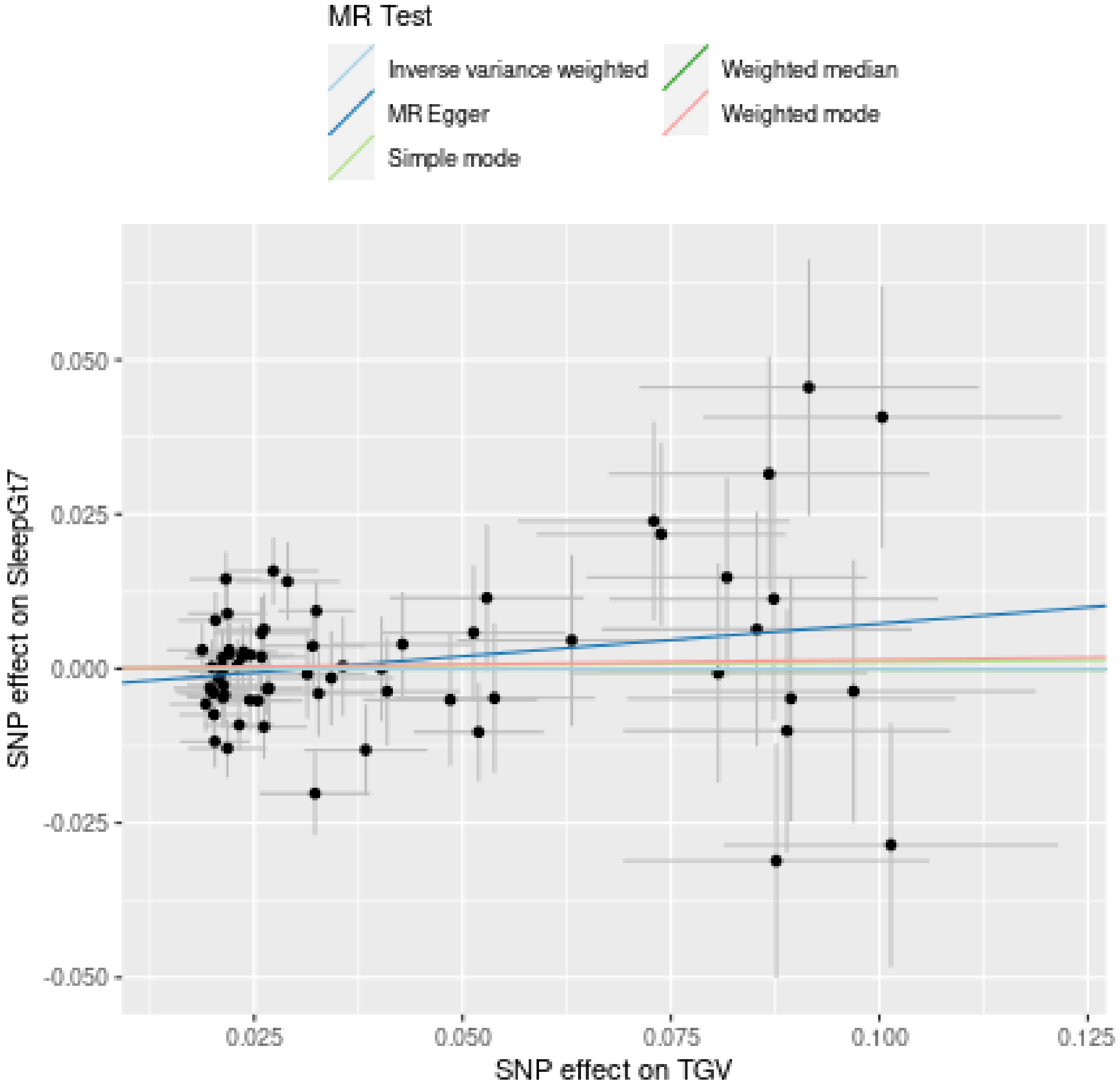

**Funnel plot**

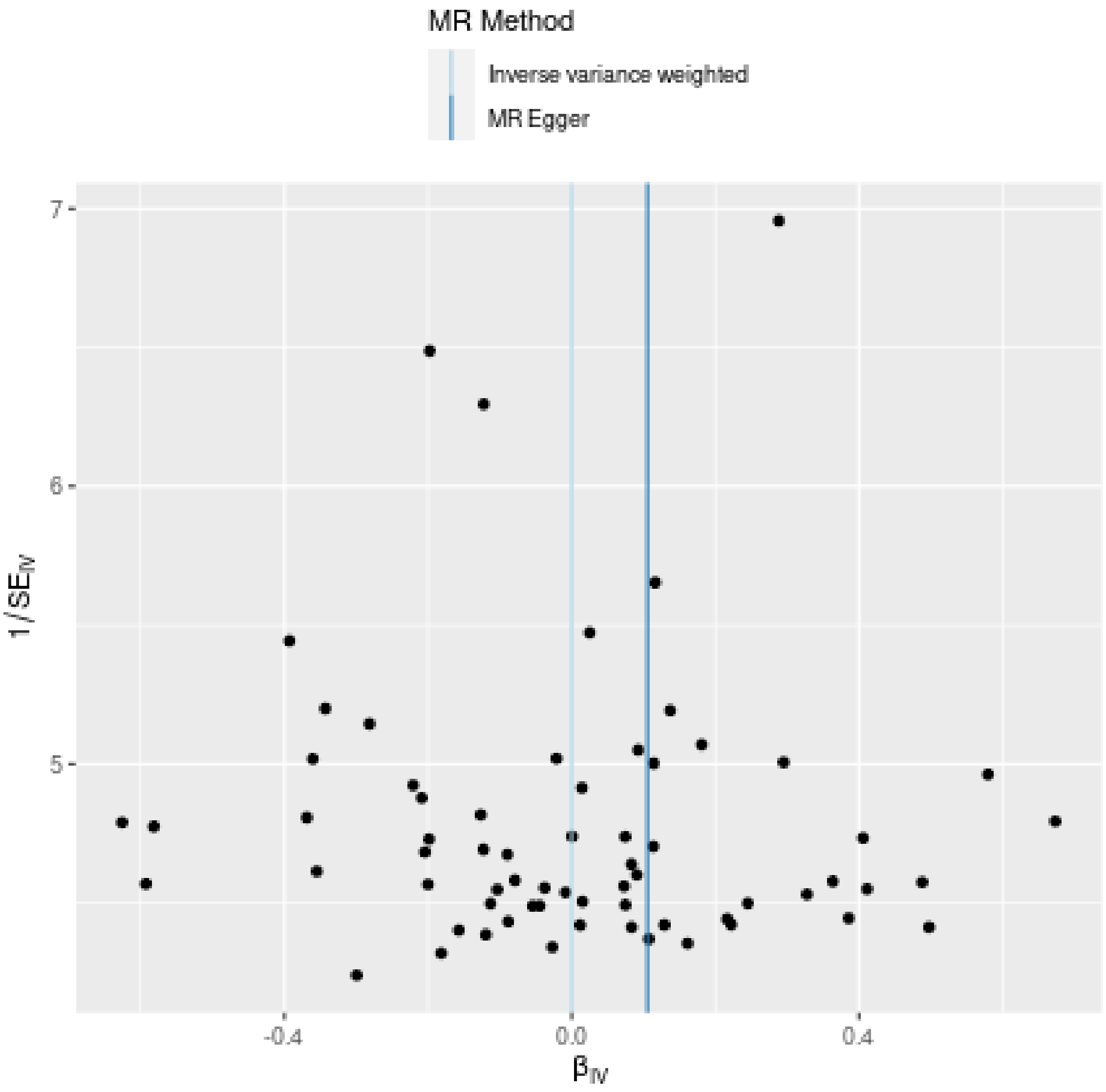
MR Method
Inverse variance weighted
MR Egger
7
6
5
1/$SE_{IV}$
-0.4
0.0
0.4
$\beta_{IV}$

**Leave-one-out sensitivity analysis**

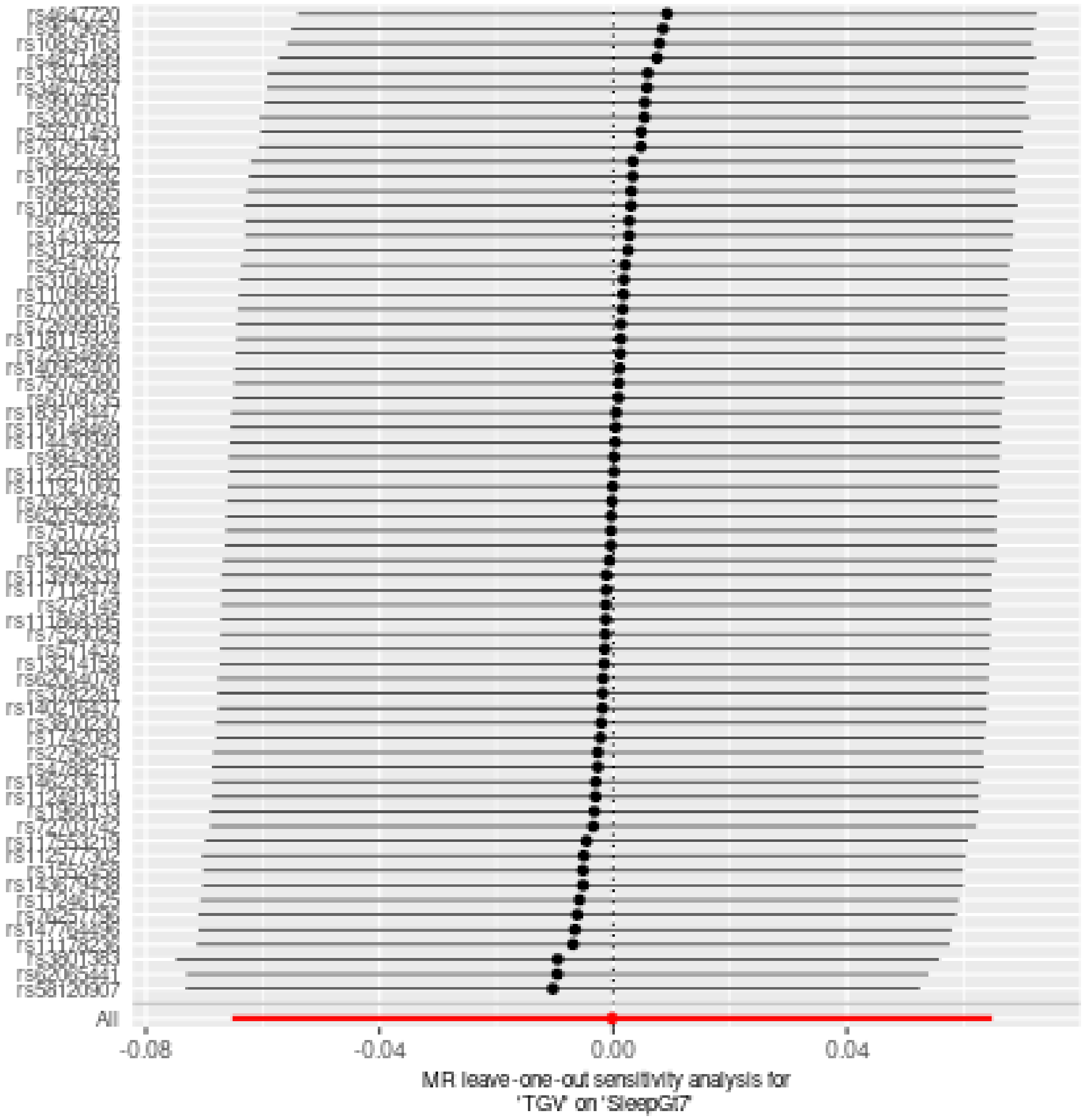

### Two sample MR report

---

#### TGV against SleepLe7

Date: **24 september, 2021**

---

###### Results from two sample MR:

| method | nsnp | b | se | pval |
|---|---|---|---|---|
| MR Egger | 67 | -0.0178969 | 0.0540929 | 0.7418183 |
| Weighted median | 67 | 0.0223517 | 0.0285948 | 0.4344087 |
| Inverse variance weighted | 67 | 0.0361167 | 0.0213874 | 0.0912793 |
| Simple mode | 67 | -0.0029460 | 0.0643265 | 0.9636094 |
| Weighted mode | 67 | 0.0199181 | 0.0593531 | 0.7382476 |

---

###### Heterogeneity tests

| method | Q | Q_df | Q_pval |
|---|---|---|---|
| MR Egger | 80.03294 | 65 | 0.0991988 |
| Inverse variance weighted | 81.48735 | 66 | 0.0947592 |

---

###### Test for directional horizontal pleiotropy

| egger_intercept | se | pval |
|---|---|---|
| 0.0016544 | 0.0015222 | 0.2811208 |

---

###### Test that the exposure is upstream of the outcome

| snp_r2.exposure | snp_r2.outcome | correct_causal_direction | steiger_pval |
|---|---|---|---|
| 0.0620032 | 0.0004911 | TRUE | 0 |

Note - R^2 values are approximate

---

**Forest plot of single SNP MR**

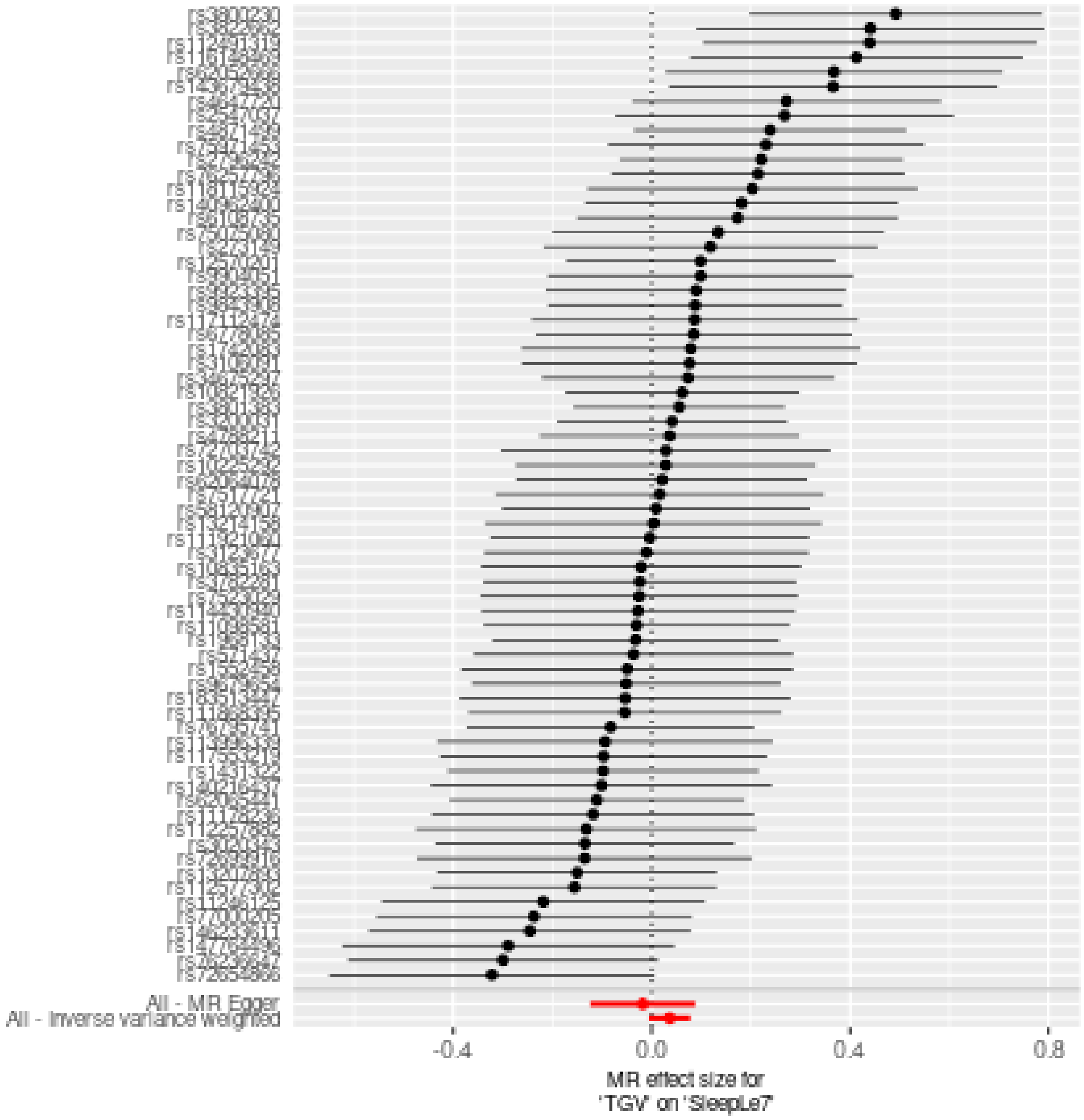

**Comparison of results using different MR methods**

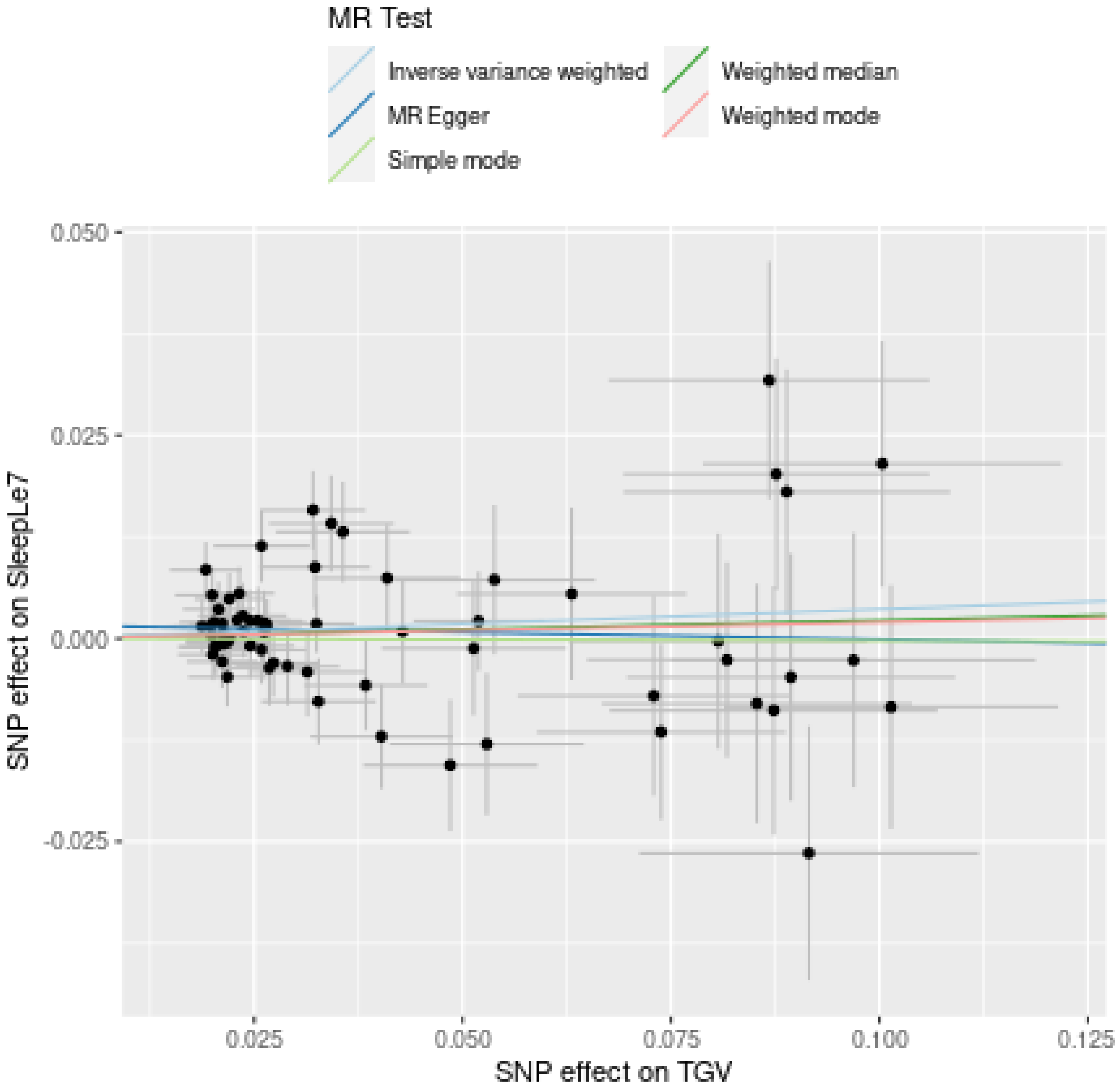

**Funnel plot**

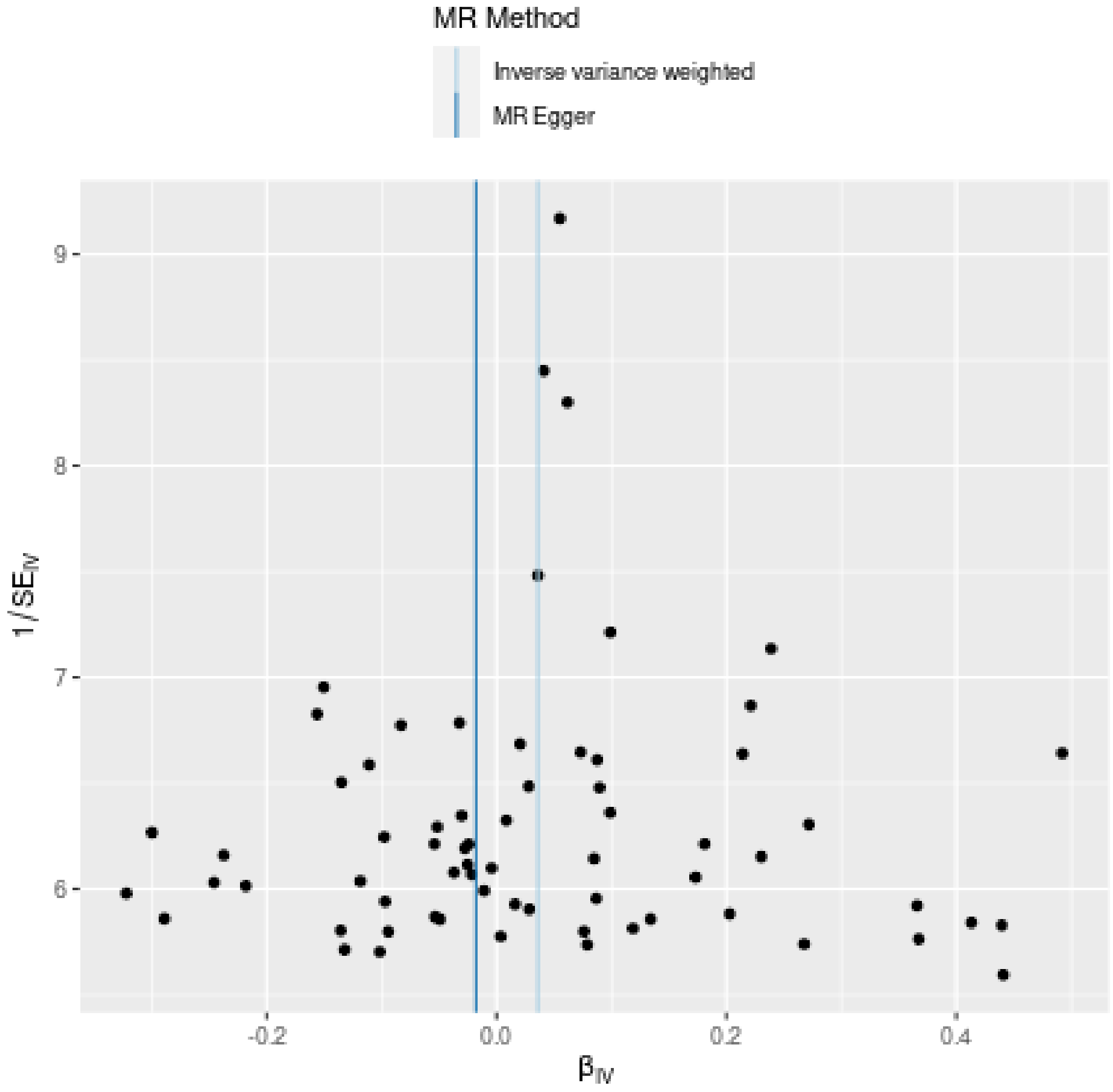

MR Method
Inverse variance weighted
MR Egger
9
8
7
6
1/SE_IV
-0.2
0.0
0.2
0.4
β_IV

**Leave-one-out sensitivity analysis**

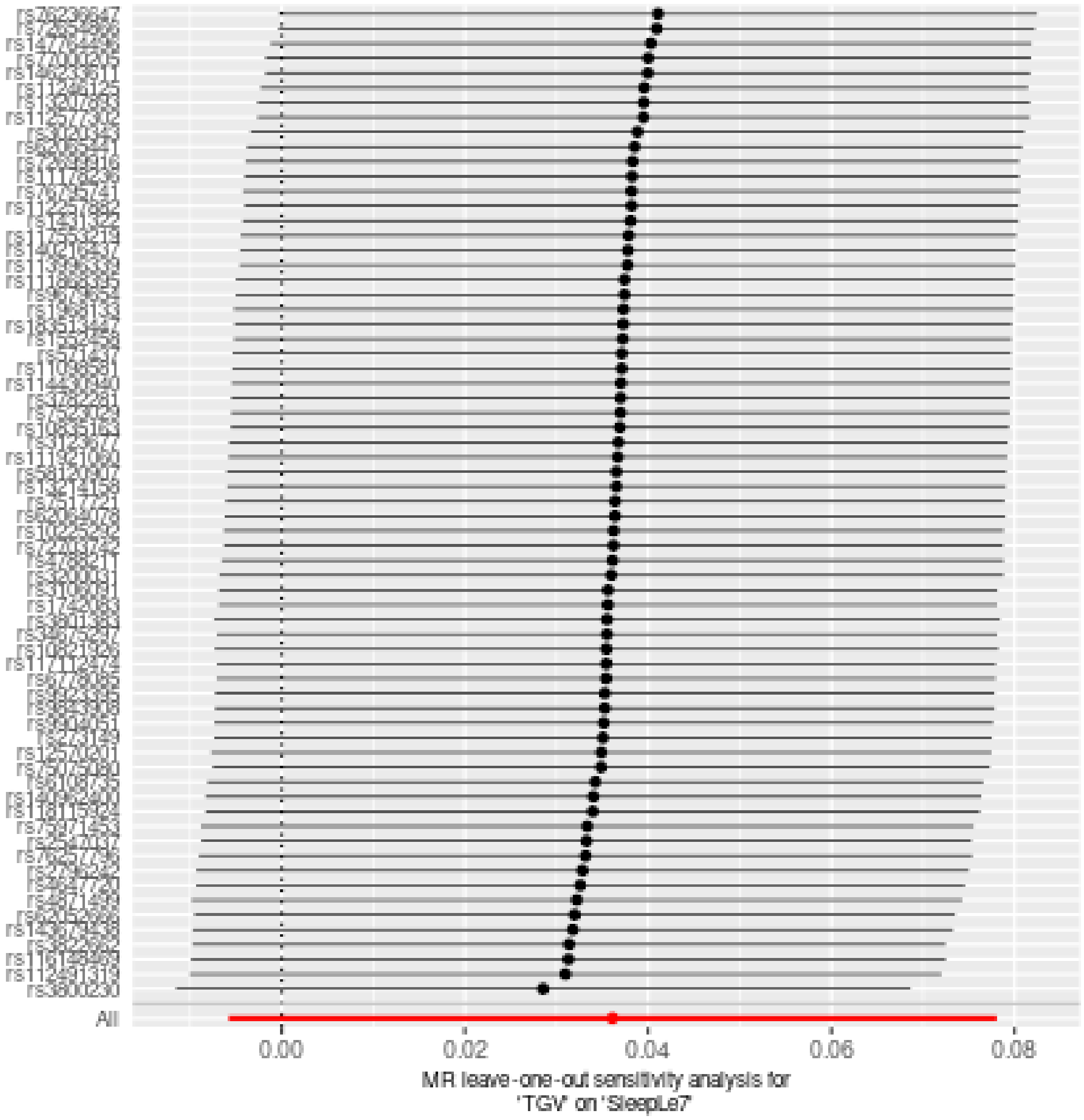
