## Supplementary material for "Sleep duration and brain structure – phenotypic associations and genotypic covariance": Si Meta analysis

The analyses to be described were conducted in R version 4.0.0 (R Core Team 2020). We used the `metafor` package (Viechtbauer 2010) for the meta analysis and `tidyverse` (Wickham et al. 2019) for data manipulation and visualization.

```
library(tidyverse)
library(metafor)
```

### Controling for ICV

We start the meta analysis using the models for subcortical volumes which controled for intracranial volume.

We performed a meta-analytic fit in order to find the average across both cortical and subcortical regions of the sleep associated with maximum cortical thickness and subcortical volume. Corpos callosum structures were excluded because with one exception their estimated maximum volume sleep were fixed at the lower boundary of four hours. Including them would hence lead to division by zero in the meta analysis. This can also be seen in the supplementary material for subcortical volumes. We also excluded total gray matter volume and intracranial volume, and weighted the three cortical clusters and the seven subcortical regions such that cortex and subcortex contributed equally. Random effects meta analysis was used, since we did not expect each region to have the same sleep duration associated with maximum thickness/volume.

To be able to compare estimates across all regions, we used the model with a smooth term for sleep and no age interaction for all subcortical regions. In the supplementary material for subcortical volumes, this model specification is titled `mod_no_interaction`. For the ventricles, we searched for the number of hours of sleep corresponding to minimal volume, while for all other regions we search for the sleep duration that maximized the volumes. The estimates and standard errors were computed by sampling 5000 Monte Carlo samples from the empirical Bayes posterior distribution of the model for each region, constraining the number of hours of sleep to be between 4 and 12.

The complete data are shown below, where Cluster 1, 2, and 3 refer to the cortical clusters.

| Region | Estimate | Standard error | Weight |
|---|---|---|---|
| Accumbens-area | 8.061 | 4.000 | 0.083 |
| Amygdala | 4.222 | 1.315 | 0.083 |
| Brain-Stem | 8.069 | 2.879 | 0.083 |
| Caudate | 11.821 | 1.184 | 0.083 |
| Cerebellum-Cortex | 7.509 | 1.291 | 0.083 |
| Cerebellum-White-Matter | 4.793 | 1.018 | 0.083 |
| CerebralWhiteMatterVol | 4.596 | 0.804 | 0.083 |
| Hippocampus | 6.268 | 0.530 | 0.083 |
| Pallidum | 10.742 | 2.912 | 0.083 |
| Putamen | 4.709 | 2.272 | 0.083 |
| Thalamus | 6.075 | 1.406 | 0.083 |
| Ventricles | 5.622 | 1.302 | 0.083 |
| Cluster 1 | 6.440 | 0.345 | 0.333 |
| Cluster 2 | 6.740 | 0.190 | 0.333 |
| Cluster 3 | 7.020 | 0.200 | 0.333 |

The following call was used to fit the model.

```
meta_mod <- rma(yi = estimate, sei = standard_error, weights = weight,
                method = "REML", data = meta_dat)
```

And next is the model summary. The estimate refers to the average of the sleep associated with maximum subcortical volume and cortical thickness. The p-value is with respect to a null hypothesis that zero hours of sleep is optimal, and is hence not relevant. The confidence intervals, on the other hand, can be interpreted as usual.

```
summary(meta_mod)
```

```
##
## Random-Effects Model (k = 15; tau^2 estimator: REML)
##
##   logLik  deviance       AIC       BIC      AICc
## -30.5789   61.1577   65.1577   66.4358   66.2486
##
## tau^2 (estimated amount of total heterogeneity): 2.1738 (SE = 1.2755)
## tau (square root of estimated tau^2 value):      1.4744
## I^2 (total heterogeneity / total variability):   88.53%
## H^2 (total variability / sampling variability):  8.72
##
## Test for Heterogeneity:
## Q(df = 14) = 40.6873, p-val = 0.0002
##
## Model Results:
##
## estimate      se     zval    pval   ci.lb   ci.ub
##   6.8036  0.5620  12.1064  <.0001  5.7022  7.9051  ***
##
## ---
## Signif. codes:  0 '***' 0.001 '**' 0.01 '*' 0.05 '.' 0.1 ' ' 1
```

#### Additional sensitivity check

Since we use a random effects meta analysis it is also possible to include the corpus callosum structures, despite them having zero standard errors. We show this below, as a check of the robustness of the conclusions.

We weighted the corpus callosum structures such that they in total contributed as much as a single other subcortical region. The complete data are shown below.

```
meta_dat %>%
  rename(
    Region = region,
    Estimate = estimate,
    `Standard error` = standard_error,
    Weight = weight
  ) %>%
  knitr::kable(digits = 3)
```

| Region | Estimate | Standard error | weight0 | Weight |
|---|---|---|---|---|
| Accumbens-area | 8.061 | 4.000 | 1.0 | 0.059 |
| Amygdala | 4.222 | 1.315 | 1.0 | 0.059 |
| Brain-Stem | 8.069 | 2.879 | 1.0 | 0.059 |

| Region | Estimate | Standard error | weight0 | Weight |
|---|---|---|---|---|
| Caudate | 11.821 | 1.184 | 1.0 | 0.059 |
| CC_Anterior | 4.010 | 0.277 | 0.2 | 0.012 |
| CC_Central | 4.000 | 0.000 | 0.2 | 0.012 |
| CC_Mid_Anterior | 4.000 | 0.000 | 0.2 | 0.012 |
| CC_Mid_Posterior | 5.981 | 1.927 | 0.2 | 0.012 |
| CC_Posterior | 7.779 | 1.537 | 0.2 | 0.012 |
| Cerebellum-Cortex | 7.509 | 1.291 | 1.0 | 0.059 |
| Cerebellum-White-Matter | 4.793 | 1.018 | 1.0 | 0.059 |
| CerebralWhiteMatterVol | 4.596 | 0.804 | 1.0 | 0.059 |
| Hippocampus | 6.268 | 0.530 | 1.0 | 0.059 |
| Pallidum | 10.742 | 2.912 | 1.0 | 0.059 |
| Putamen | 4.709 | 2.272 | 1.0 | 0.059 |
| Thalamus | 6.075 | 1.406 | 1.0 | 0.059 |
| Ventricles | 5.622 | 1.302 | 1.0 | 0.059 |
| Cluster 1 | 6.440 | 0.345 | 1.0 | 0.333 |
| Cluster 2 | 6.740 | 0.190 | 1.0 | 0.333 |
| Cluster 3 | 7.020 | 0.200 | 1.0 | 0.333 |

Next we fit the model. Note the warnings, which are expected.

```
meta_mod <- rma(yi = estimate, sei = standard_error, weights = weight,
                method = "REML", data = meta_dat)
```

```
## Warning: There are outcomes with non-positive sampling variances.
```

```
## Warning: Cannot compute Q-test, I^2, or H^2 when there are non-positive sampling
## variances in the data.
```

Here is the summary, which shows estimates close to what we got when excluding corpus callosum.

```
summary(meta_mod)
```

```
##
## Random-Effects Model (k = 20; tau^2 estimator: REML)
##
##   logLik  deviance       AIC       BIC      AICc
## -42.6055   85.2111   89.2111   91.1000   89.9611
##
## tau^2 (estimated amount of total heterogeneity): 2.8415 (SE = 1.2795)
## tau (square root of estimated tau^2 value):      1.6857
##
## Model Results:
##
## estimate      se     zval    pval   ci.lb   ci.ub
##   6.7369  0.6353  10.6040  <.0001  5.4917  7.9821  ***
##
## ---
## Signif. codes:  0 '***' 0.001 '**' 0.01 '*' 0.05 '.' 0.1 ' ' 1
```

### Not Controling for ICV

We next use the subcortical models which did not contain ICV. In this case only the ventricles had zero standard error, but to make the results comparable to above, we performed one analysis excluding the corpos callosum and one analysis including corpos callosum.

The complete data are shown below.

| Region | Estimate | Standard error | Weight |
|---|---|---|---|
| Accumbens-area | 7.890 | 1.237 | 0.083 |
| Amygdala | 8.584 | 2.152 | 0.083 |
| Brain-Stem | 8.766 | 2.200 | 0.083 |
| Caudate | 10.594 | 1.609 | 0.083 |
| Cerebellum-Cortex | 7.718 | 1.323 | 0.083 |
| Cerebellum-White-Matter | 7.935 | 1.845 | 0.083 |
| CerebralWhiteMatterVol | 7.412 | 0.526 | 0.083 |
| Hippocampus | 7.135 | 0.223 | 0.083 |
| Pallidum | 9.570 | 2.216 | 0.083 |
| Putamen | 9.125 | 2.245 | 0.083 |
| Thalamus | 7.644 | 1.235 | 0.083 |
| Ventricles | 4.000 | 0.000 | 0.083 |
| Cluster 1 | 6.440 | 0.345 | 0.333 |
| Cluster 2 | 6.740 | 0.190 | 0.333 |
| Cluster 3 | 7.020 | 0.200 | 0.333 |

The following call was used to fit the model.

```
meta_mod <- rma(yi = estimate, sei = standard_error, weights = weight,
                method = "REML", data = meta_dat)
```

```
## Warning: There are outcomes with non-positive sampling variances.
```

```
## Warning: Cannot compute Q-test, I^2, or H^2 when there are non-positive sampling
## variances in the data.
```

And next is the model summary.

```
summary(meta_mod)
```

```
##
## Random-Effects Model (k = 15; tau^2 estimator: REML)
##
##   logLik  deviance       AIC       BIC      AICc
## -27.1104   54.2208   58.2208   59.4989   59.3117
##
## tau^2 (estimated amount of total heterogeneity): 1.6863 (SE = 0.9681)
## tau (square root of estimated tau^2 value):      1.2986
##
## Model Results:
##
## estimate      se     zval    pval   ci.lb   ci.ub
##   7.3822  0.4838  15.2572  <.0001  6.4338  8.3305  ***
##
## ---
## Signif. codes:  0 '***' 0.001 '**' 0.01 '*' 0.05 '.' 0.1 ' ' 1
```

Next, we included the corpos callosum in the same way as in the sensitivity check above.

We weighted the corpus callosum structures such that they in total contributed as much as a single other subcortical region. The complete data are shown below.

```r
meta_dat %>%
  rename(
    Region = region,
    Estimate = estimate,
    `Standard error` = standard_error,
    Weight = weight
  ) %>%
  knitr::kable(digits = 3)
```

| Region | Estimate | Standard error | weight0 | Weight |
|---|---|---|---|---|
| Accumbens-area | 7.890 | 1.237 | 1.0 | 0.059 |
| Amygdala | 8.584 | 2.152 | 1.0 | 0.059 |
| Brain-Stem | 8.766 | 2.200 | 1.0 | 0.059 |
| Caudate | 10.594 | 1.609 | 1.0 | 0.059 |
| CC_Anterior | 7.769 | 1.510 | 0.2 | 0.012 |
| CC_Central | 4.833 | 1.029 | 0.2 | 0.012 |
| CC_Mid_Anterior | 4.782 | 1.151 | 0.2 | 0.012 |
| CC_Mid_Posterior | 7.468 | 1.025 | 0.2 | 0.012 |
| CC_Posterior | 8.108 | 1.681 | 0.2 | 0.012 |
| Cerebellum-Cortex | 7.718 | 1.323 | 1.0 | 0.059 |
| Cerebellum-White-Matter | 7.935 | 1.845 | 1.0 | 0.059 |
| CerebralWhiteMatterVol | 7.412 | 0.526 | 1.0 | 0.059 |
| Hippocampus | 7.135 | 0.223 | 1.0 | 0.059 |
| Pallidum | 9.570 | 2.216 | 1.0 | 0.059 |
| Putamen | 9.125 | 2.245 | 1.0 | 0.059 |
| Thalamus | 7.644 | 1.235 | 1.0 | 0.059 |
| Ventricles | 4.000 | 0.000 | 1.0 | 0.059 |
| Cluster 1 | 6.440 | 0.345 | 1.0 | 0.333 |
| Cluster 2 | 6.740 | 0.190 | 1.0 | 0.333 |
| Cluster 3 | 7.020 | 0.200 | 1.0 | 0.333 |

Next we fit the model.

```r
meta_mod <- rma(yi = estimate, sei = standard_error, weights = weight,
                method = "REML", data = meta_dat)
```

```
## Warning: There are outcomes with non-positive sampling variances.

## Warning: Cannot compute Q-test, I^2, or H^2 when there are non-positive sampling
## variances in the data.
```

Here is the summary, which shows estimates close to what we got when excluding corpus callosum.

```r
summary(meta_mod)
```

```
##
## Random-Effects Model (k = 20; tau^2 estimator: REML)
##
##   logLik  deviance       AIC       BIC      AICc
## -36.8311   73.6622   77.6622   79.5510   78.4122
##
## tau^2 (estimated amount of total heterogeneity): 1.5113 (SE = 0.7965)
## tau (square root of estimated tau^2 value):      1.2293
##
```

```
## Model Results:
##
## estimate      se     zval    pval   ci.lb   ci.ub
##   7.2477  0.4728  15.3296  <.0001  6.3210  8.1743  ***
##
## ---
## Signif. codes:  0 '***' 0.001 '**' 0.01 '*' 0.05 '.' 0.1 ' ' 1
```
