## Supplementary material for "Sleep duration and brain structure – phenotypic associations and genotypic covariance": SI Subcortical results

### Subcortical volumes

#### Data overview

The total number of observations was 51320, from 47039 unique participants. The age range was from 20 to 89.4. The sex count is shown below.

| sex | n |
|---|---|
| female | 26811 |
| male | 24509 |

Follow-up intervals are shown below.

| Participants with follow-up | Mean follow-up | Max follow-up | Min follow-up |
|---|---|---|---|
| 3910 | 2.511982 | 11.22656 | 0.0054795 |

In order to remove outliers, the following model was fit to each region:

```
mod <- gamm4(value ~ s(age, bs = "cr") + sex + site,
             random = ~(1|id), data = data)
```

Outliers were defined by having a residual more than four times the magnitude of the residuals standard error. Technically, all values which returned `TRUE` on the following test were retained, all others were removed as outliers.

```
abs(residuals(mod$mer)) < 4 * sigma(mod$mer)
```

The table below shows the number of outliers removed per region.

| region | initial_rows | retained_rows | outliers_removed |
|---|---|---|---|
| Accumbens-area | 51320 | 51315 | 5 |
| Amygdala | 51320 | 51314 | 6 |
| Brain-Stem | 51320 | 51297 | 23 |
| Caudate | 51320 | 51306 | 14 |
| CC_Anterior | 51320 | 51304 | 16 |
| CC_Central | 51320 | 51307 | 13 |
| CC_Mid_Anterior | 51320 | 51298 | 22 |
| CC_Mid_Posterior | 51320 | 51279 | 41 |
| CC_Posterior | 51320 | 51266 | 54 |
| Cerebellum-Cortex | 51320 | 51292 | 28 |
| Cerebellum-White-Matter | 51320 | 51285 | 35 |
| CerebralWhiteMatterVol | 51320 | 51311 | 9 |
| EstimatedTotalIntraCranialVol | 51320 | 51300 | 20 |
| Hippocampus | 51320 | 51310 | 10 |
| Pallidum | 51320 | 51311 | 9 |
| Putamen | 51320 | 51298 | 22 |

| region | initial_rows | retained_rows | outliers_removed |
|---|---|---|---|
| Thalamus | 51319 | 51299 | 20 |
| TotalGrayVol | 51320 | 51312 | 8 |
| Ventricles | 51320 | 51299 | 21 |

The plot below shows the average sleep duration as a function of age, fit with a GAM. This model will be used for comparisons later. All sleep durations in this document are confined to lie between 4 and 10 hours, to avoid nonsensical answers.

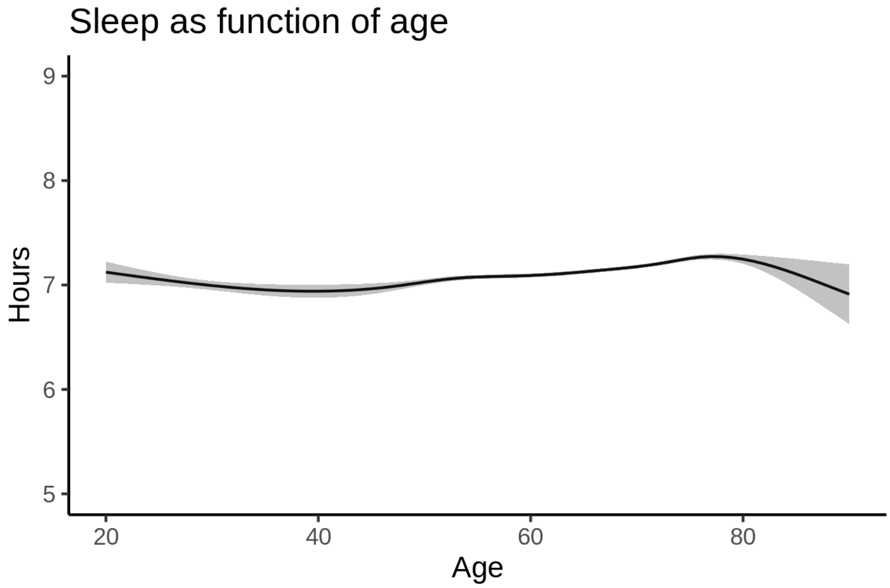


The model statistics is shown next.

```
##
## Family: gaussian
## Link function: identity
##
## Formula:
## sleep ~ s(age, bs = "cr")
##
## Parametric coefficients:
##             Estimate Std. Error t value Pr(>|t|)
## (Intercept) 7.134122   0.004306    1657   <2e-16 ***
## ---
## Signif. codes:  0 ‘***’ 0.001 ‘**’ 0.01 ‘*’ 0.05 ‘.’ 0.1 ‘ ’ 1
##
## Approximate significance of smooth terms:
##         edf Ref.df     F p-value
## s(age) 6.91  7.902 33.12  <2e-16 ***
## ---
## Signif. codes:  0 ‘***’ 0.001 ‘**’ 0.01 ‘*’ 0.05 ‘.’ 0.1 ‘ ’ 1
##
## R-sq.(adj) =  0.00558   Deviance explained = 0.572%
## -REML =  63545  Scale est. = 0.87216   n = 47039
```

#### Accumbens-area

##### Descriptive statistics

| Study | Observations | Unique IDs | Mean age | Age range |
|---|---|---|---|---|
| HCP | 974 | 974 | 28.8 | 22 - 37 |
| MPIB | 675 | 391 | 63.2 | 24 - 83 |
| UB | 113 | 39 | 70.9 | 64 - 81 |
| UCAM | 884 | 632 | 55.1 | 20 - 88 |
| UiO | 1474 | 803 | 49.4 | 20 - 89 |
| UKB | 45983 | 43137 | 64.5 | 45 - 83 |
| UmU | 423 | 284 | 62.3 | 25 - 85 |
| UOXF | 769 | 769 | 69.8 | 60 - 85 |

##### Spaghetti plot

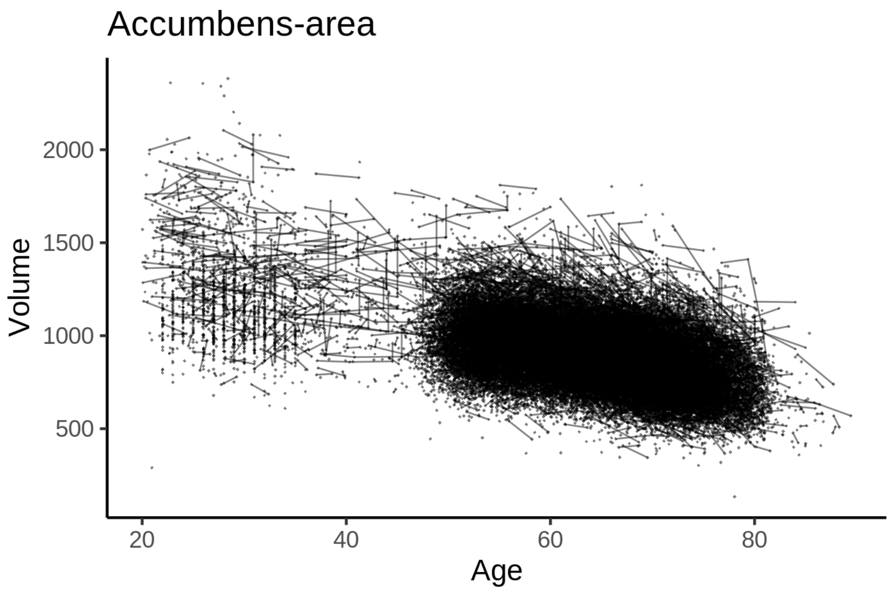


##### Model outputs

Model without sleep term

```
##
## Family: gaussian
## Link function: identity
##
## Formula:
## value ~ sex + site + icv + s(age_z, k = 10, bs = "cr")
## <environment: 0x55cf3fd47368>
##
## Parametric coefficients:
##                Estimate Std. Error t value Pr(>|t|)
## (Intercept)   804.7854     7.7562 103.761  < 2e-16 ***
```

```
## sexmale        19.7698     1.5399  12.838  < 2e-16 ***
## siteMPIB       93.3082     9.6688   9.650  < 2e-16 ***
## siteousAvanto 238.5742     8.2501  28.918  < 2e-16 ***
## siteousPrisma 101.7528    13.6266   7.467 8.32e-14 ***
## siteousSkyra  407.8480     7.4142  55.009  < 2e-16 ***
## siteUB         64.8784    22.1438   2.930  0.00339 **
## siteUCAM        4.6942     8.4308   0.557  0.57767
## siteUKB        82.5192     7.9615  10.365  < 2e-16 ***
## siteUmU       367.0385    10.8002  33.984  < 2e-16 ***
## siteUOXF       59.8483     9.3456   6.404 1.53e-10 ***
## icv            59.5677     0.7755  76.808  < 2e-16 ***
## ---
## Signif. codes:  0 '***' 0.001 '**' 0.01 '*' 0.05 '.' 0.1 ' ' 1
##
## Approximate significance of smooth terms:
##            edf Ref.df    F p-value
## s(age_z) 8.142  8.142 1888  <2e-16 ***
## ---
## Signif. codes:  0 '***' 0.001 '**' 0.01 '*' 0.05 '.' 0.1 ' ' 1
##
## R-sq.(adj) =  0.468
## lmer.REML = 6.443e+05  Scale est. = 2922.1    n = 51295
```

Model with only main effects of age and sleep

```
##
## Family: gaussian
## Link function: identity
##
## Formula:
## value ~ sex + site + icv + s(age_z, k = 10, bs = "cr") + s(sleep_z,
##     k = 5, bs = "cr")
## <environment: 0x55cf3fd47368>
##
## Parametric coefficients:
##               Estimate Std. Error t value Pr(>|t|)
## (Intercept)   804.7917     7.7638 103.659  < 2e-16 ***
## sexmale        19.7701     1.5399  12.838  < 2e-16 ***
## siteMPIB       93.3009     9.6760   9.643  < 2e-16 ***
## siteousAvanto 238.5709     8.2521  28.910  < 2e-16 ***
## siteousPrisma 101.7509    13.6271   7.467 8.35e-14 ***
## siteousSkyra  407.8446     7.4166  54.991  < 2e-16 ***
## siteUB         64.8772    22.1441   2.930  0.00339 **
## siteUCAM        4.6913     8.4328   0.556  0.57800
## siteUKB        82.5124     7.9700  10.353  < 2e-16 ***
## siteUmU       367.0232    10.8261  33.902  < 2e-16 ***
## siteUOXF       59.8456     9.3474   6.402 1.54e-10 ***
## icv            59.5672     0.7759  76.772  < 2e-16 ***
## ---
## Signif. codes:  0 '***' 0.001 '**' 0.01 '*' 0.05 '.' 0.1 ' ' 1
##
## Approximate significance of smooth terms:
##              edf Ref.df    F p-value
## s(age_z)   8.142  8.142 1883  <2e-16 ***
## s(sleep_z) 1.000  1.000    0   0.982
```

```
## ---
## Signif. codes:  0 '***' 0.001 '**' 0.01 '*' 0.05 '.' 0.1 ' ' 1
##
## R-sq.(adj) =  0.468
## lmer.REML = 6.443e+05  Scale est. = 2922.2    n = 51295
```

Model with full interaction between age and sleep

```
##
## Family: gaussian
## Link function: identity
##
## Formula:
## value ~ sex + site + icv + t2(age_z, sleep_z, k = c(10, 4), bs = "cr")
## <environment: 0x55cf3fd47368>
##
## Parametric coefficients:
##                Estimate Std. Error t value Pr(>|t|)
## (Intercept)    805.9183     7.8135 103.144  < 2e-16 ***
## sexmale         19.7654     1.5433  12.807  < 2e-16 ***
## siteMPIB        92.4267     9.7396   9.490  < 2e-16 ***
## siteousAvanto  237.6135     8.3000  28.628  < 2e-16 ***
## siteousPrisma  100.7471    13.6595   7.376 1.66e-13 ***
## siteousSkyra   406.8795     7.4728  54.448  < 2e-16 ***
## siteUB          63.8033    22.1684   2.878    0.004 **
## siteUCAM         3.4953     8.4921   0.412    0.681
## siteUKB         81.3575     8.0206  10.144  < 2e-16 ***
## siteUmU        366.0625    10.8761  33.658  < 2e-16 ***
## siteUOXF        58.6861     9.3987   6.244 4.30e-10 ***
## icv             59.5497     0.7761  76.728  < 2e-16 ***
## ---
## Signif. codes:  0 '***' 0.001 '**' 0.01 '*' 0.05 '.' 0.1 ' ' 1
##
## Approximate significance of smooth terms:
##                      edf Ref.df     F p-value
## t2(age_z,sleep_z) 13.55  13.55 33.24  <2e-16 ***
## ---
## Signif. codes:  0 '***' 0.001 '**' 0.01 '*' 0.05 '.' 0.1 ' ' 1
##
## R-sq.(adj) =  0.468
## lmer.REML = 6.4431e+05  Scale est. = 2921.5    n = 51295
```

#### Model comparison

`mod_no_sleep` refers to model without sleep term, `mod_no_interaction` refers to model with only main effect of sleep, and `mod_full` refers to model with a full interaction between age and sleep. This is a nested model comparison, and the p-value at a given line refers to comparing the model at the line to the model on the line above. Hence, significance implies that the more complicated model is supported on statistical grounds.

To be even more specific, the p-value on the second row tests whether there is an association between sleep and volume. The p-value on the third row tests whether this association depends on age.

```
## Data: NULL
## Models:
```

```
## mod_list$mod_no_sleep$mer: NULL
## mod_list$mod_no_interaction$mer: NULL
## mod_list$mod_full$mer: NULL
##                                npar    AIC    BIC  logLik deviance Chisq Df Pr(>Chisq)
## mod_list$mod_no_sleep$mer         16 644334 644475 -322151   644302
## mod_list$mod_no_interaction$mer   18 644338 644497 -322151   644302 5e-04  2     0.9998
## mod_list$mod_full$mer             20 644354 644531 -322157   644314 0e+00  2     1.0000
```

We chose the model based on the likelihood ratio test with 5 % significance level, which was `mod_no_sleep`.

##### Lifespan brain trajectory

The trajectory shown is from the chosen model `mod_no_sleep`.

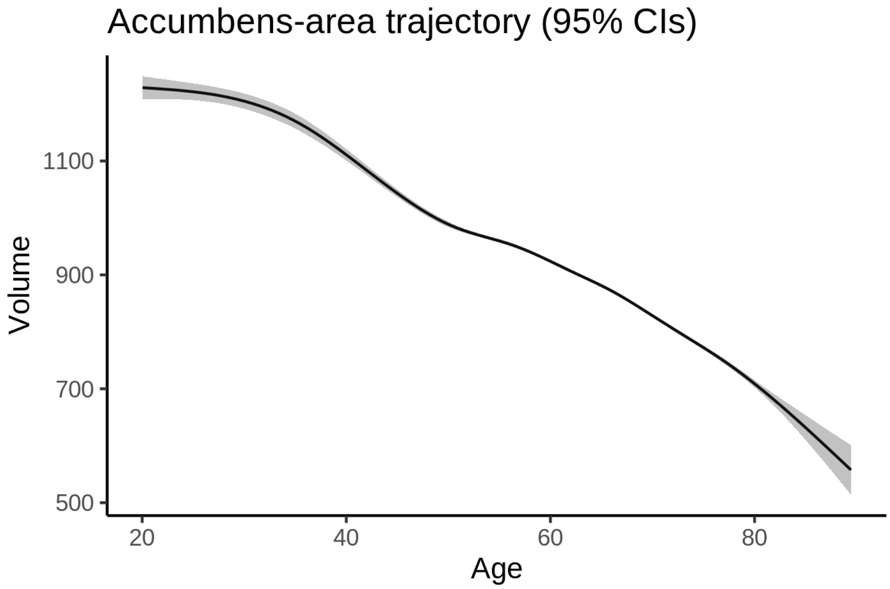


##### Effect of sleep

The chosen model did not include a sleep term, and hence we don't have any estimated effect of sleep.

We show the full interaction model for completeness, although it was not selected.

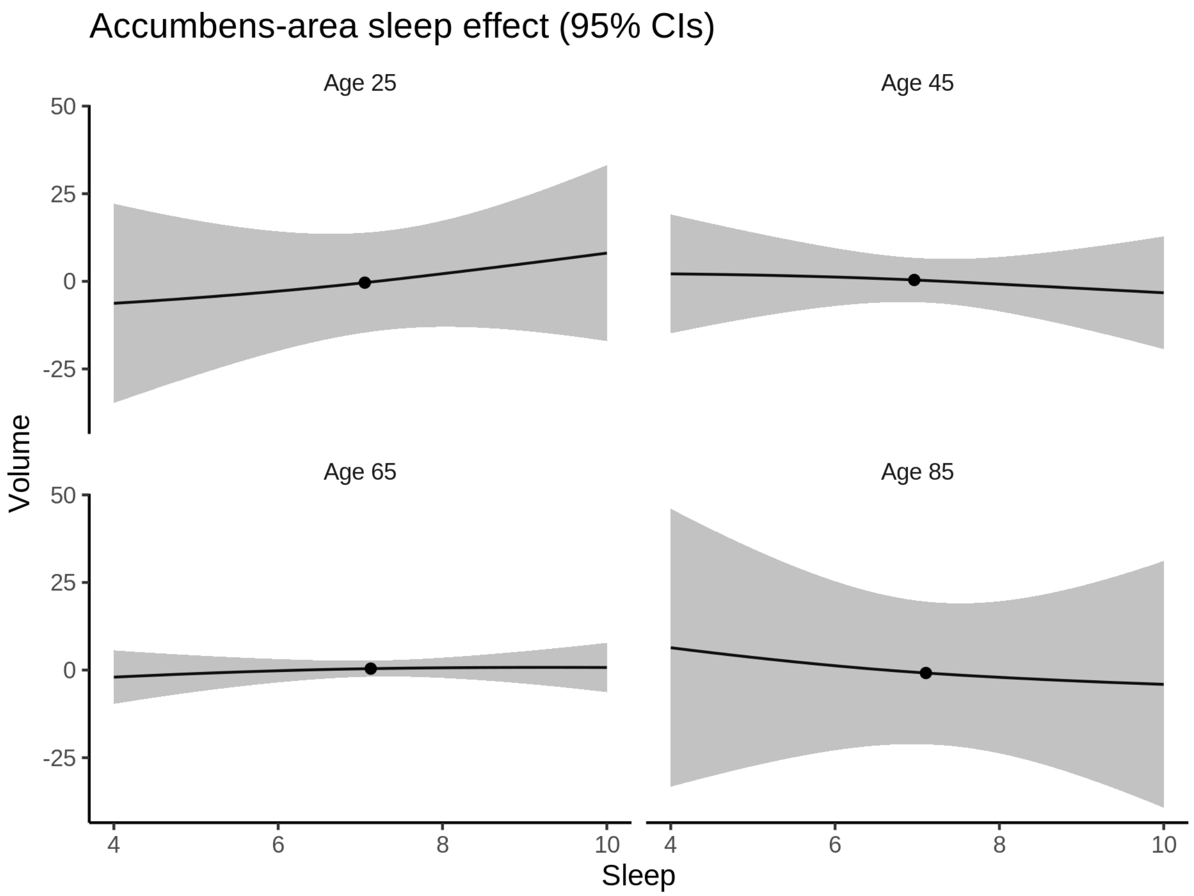


##### Deviation from sleep associated with maximal volume

Model with no sleep term was selected. No plots to show. (Although we can of course dig up the plots, which will be pretty flat).

##### Comparison of mean sleep and sleep associated with maximum volume

Nothing to show, as we did not find an association between sleep and volume.

### Amygdala

#### Descriptive statistics

| Study | Observations | Unique IDs | Mean age | Age range |
|---|---|---|---|---|
| HCP | 974 | 974 | 28.8 | 22 - 37 |
| MPIB | 677 | 391 | 63.1 | 24 - 83 |
| UB | 113 | 39 | 70.9 | 64 - 81 |
| UCAM | 884 | 632 | 55.1 | 20 - 88 |
| UiO | 1472 | 803 | 49.3 | 20 - 89 |
| UKB | 45982 | 43138 | 64.5 | 45 - 83 |

| Study | Observations | Unique IDs | Mean age | Age range |
|---|---|---|---|---|
| UmU | 423 | 284 | 62.3 | 25 - 85 |
| UOXF | 769 | 769 | 69.8 | 60 - 85 |

##### Spaghetti plot

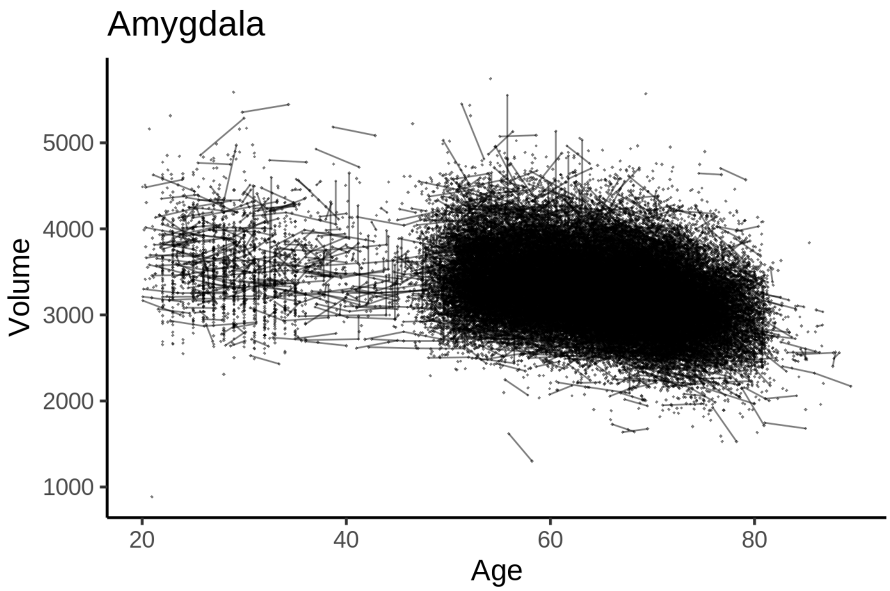


##### Model outputs

Model without sleep term

```
##
## Family: gaussian
## Link function: identity
##
## Formula:
## value ~ sex + site + icv + s(age_z, k = 10, bs = "cr")
## <environment: 0x55cf39151140>
##
## Parametric coefficients:
##                Estimate Std. Error t value Pr(>|t|)
## (Intercept)   3032.751     18.118 167.386  < 2e-16 ***
## sexmale        147.607      3.645  40.494  < 2e-16 ***
## siteMPIB       342.390     22.852  14.983  < 2e-16 ***
## siteousAvanto -100.927     18.894  -5.342 9.24e-08 ***
## siteousPrisma  188.671     31.730   5.946 2.76e-09 ***
## siteousSkyra   508.714     17.451  29.151  < 2e-16 ***
## siteUB          81.988     53.036   1.546   0.1221
## siteUCAM       102.681     19.862   5.170 2.35e-07 ***
## siteUKB        190.839     18.591  10.265  < 2e-16 ***
## siteUmU        157.735     25.501   6.185 6.24e-10 ***
## siteUOXF        46.149     21.908   2.106   0.0352 *
## icv            202.559      1.834 110.417  < 2e-16 ***
```

```
## ---
## Signif. codes:  0 ‘***’ 0.001 ‘**’ 0.01 ‘*’ 0.05 ‘.’ 0.1 ‘ ’ 1
##
## Approximate significance of smooth terms:
##            edf Ref.df    F p-value
## s(age_z) 7.503  7.503 1303  <2e-16 ***
## ---
## Signif. codes:  0 ‘***’ 0.001 ‘**’ 0.01 ‘*’ 0.05 ‘.’ 0.1 ‘ ’ 1
##
## R-sq.(adj) =  0.457
## lmer.REML = 7.3126e+05  Scale est. = 11634     n = 51294
```

Model with only main effects of age and sleep

```
##
## Family: gaussian
## Link function: identity
##
## Formula:
## value ~ sex + site + icv + s(age_z, k = 10, bs = "cr") + s(sleep_z,
##     k = 5, bs = "cr")
## <environment: 0x55cf39151140>
##
## Parametric coefficients:
##              Estimate Std. Error t value Pr(>|t|)
## (Intercept)   3031.209     18.136 167.138  < 2e-16 ***
## sexmale        147.552      3.645  40.480  < 2e-16 ***
## siteMPIB       344.063     22.868  15.046  < 2e-16 ***
## siteousAvanto -100.111     18.898  -5.297 1.18e-07 ***
## siteousPrisma  189.217     31.730   5.963 2.49e-09 ***
## siteousSkyra   509.561     17.456  29.191  < 2e-16 ***
## siteUB          82.478     53.035   1.555   0.1199
## siteUCAM       103.484     19.866   5.209 1.91e-07 ***
## siteUKB        192.476     18.610  10.343  < 2e-16 ***
## siteUmU        161.103     25.561   6.303 2.95e-10 ***
## siteUOXF        46.959     21.912   2.143   0.0321 *
## icv            202.666      1.835 110.428  < 2e-16 ***
## ---
## Signif. codes:  0 ‘***’ 0.001 ‘**’ 0.01 ‘*’ 0.05 ‘.’ 0.1 ‘ ’ 1
##
## Approximate significance of smooth terms:
##              edf Ref.df        F p-value
## s(age_z)   7.507  7.507 1294.646  <2e-16 ***
## s(sleep_z) 1.000  1.000    3.651  0.0561 .
## ---
## Signif. codes:  0 ‘***’ 0.001 ‘**’ 0.01 ‘*’ 0.05 ‘.’ 0.1 ‘ ’ 1
##
## R-sq.(adj) =  0.457
## lmer.REML = 7.3125e+05  Scale est. = 11635     n = 51294
```

Model with full interaction between age and sleep

```
##
## Family: gaussian
## Link function: identity
##
```

```
## Formula:
## value ~ sex + site + icv + t2(age_z, sleep_z, k = c(10, 4), bs = "cr")
## <environment: 0x55cf39151140>
##
## Parametric coefficients:
##                Estimate Std. Error t value Pr(>|t|)    
## (Intercept)    3034.166     18.214 166.583  < 2e-16 ***
## sexmale         147.649      3.653  40.418  < 2e-16 ***
## siteMPIB        341.225     22.996  14.838  < 2e-16 ***
## siteousAvanto -102.655     18.999  -5.403 6.57e-08 ***
## siteousPrisma  186.506     31.799   5.865 4.51e-09 ***
## siteousSkyra   506.860     17.572  28.844  < 2e-16 ***
## siteUB          79.479     53.074   1.498   0.1343    
## siteUCAM       100.339     19.984   5.021 5.16e-07 ***
## siteUKB        189.407     18.688  10.135  < 2e-16 ***
## siteUmU        158.176     25.653   6.166 7.05e-10 ***
## siteUOXF        43.759     21.998   1.989   0.0467 *  
## icv            202.575      1.836 110.342  < 2e-16 ***
## ---
## Signif. codes:  0 ‘***’ 0.001 ‘**’ 0.01 ‘*’ 0.05 ‘.’ 0.1 ‘ ’ 1
##
## Approximate significance of smooth terms:
##                     edf Ref.df    F p-value    
## t2(age_z,sleep_z) 12.89  12.89 21.8  <2e-16 ***
## ---
## Signif. codes:  0 ‘***’ 0.001 ‘**’ 0.01 ‘*’ 0.05 ‘.’ 0.1 ‘ ’ 1
##
## R-sq.(adj) =  0.457   
## lmer.REML = 7.3126e+05  Scale est. = 11634     n = 51294
```

#### Model comparison

`mod_no_sleep` refers to model without sleep term, `mod_no_interaction` refers to model with only main effect of sleep, and `mod_full` refers to model with a full interaction between age and sleep. This is a nested model comparison, and the p-value at a given line refers to comparing the model at the line to the model on the line above. Hence, significance implies that the more complicated model is supported on statistical grounds.

To be even more specific, the p-value on the second row tests whether there is an association between sleep and volume. The p-value on the third row tests whether this association depends on age.

```
## Data: NULL
## Models:
## mod_list$mod_no_sleep$mer: NULL
## mod_list$mod_no_interaction$mer: NULL
## mod_list$mod_full$mer: NULL
##                                 npar    AIC    BIC  logLik deviance  Chisq Df Pr(>Chisq)
## mod_list$mod_no_sleep$mer         16 731291 731432 -365629   731259                     
## mod_list$mod_no_interaction$mer   18 731291 731450 -365627   731255 3.6504  2     0.1612
## mod_list$mod_full$mer             20 731302 731479 -365631   731262 0.0000  2     1.0000
```

We chose the model based on the likelihood ratio test with 5 % significance level, which was `mod_no_sleep`.

##### Lifespan brain trajectory

The trajectory shown is from the chosen model `mod_no_sleep`.

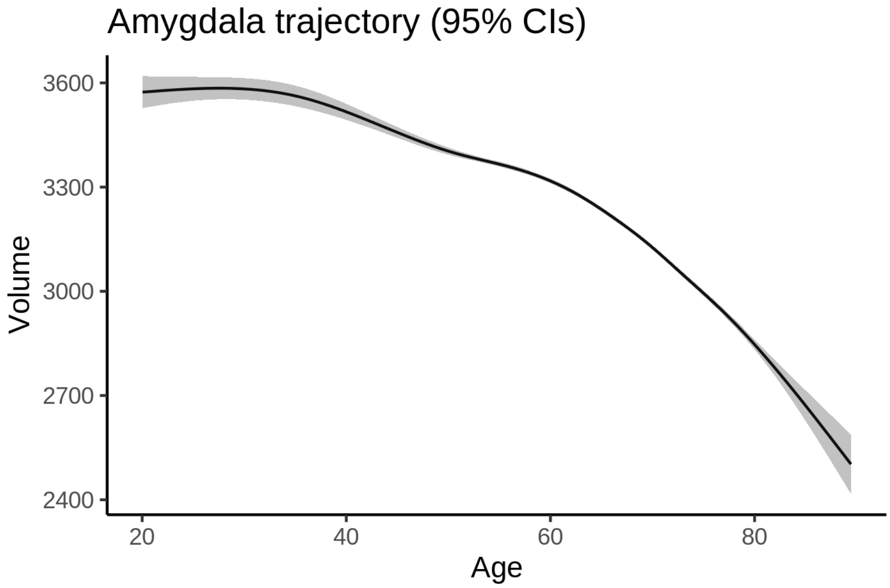


##### Effect of sleep

The chosen model did not include a sleep term, and hence we don't have any estimated effect of sleep.

We show the full interaction model for completeness, although it was not selected.

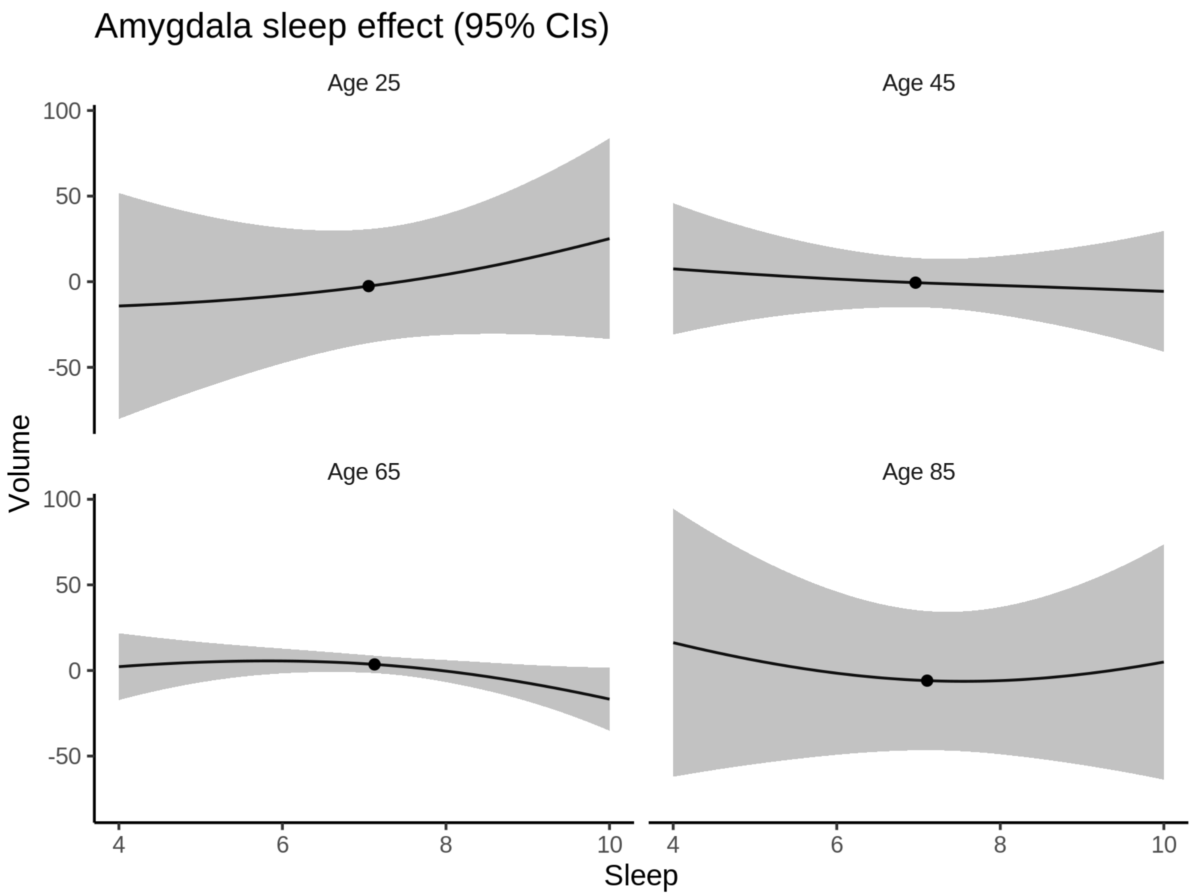


**Deviation from sleep associated with maximal volume**

Model with no sleep term was selected. No plots to show. (Although we can of course dig up the plots, which will be pretty flat).

**Comparison of mean sleep and sleep associated with maximum volume**

Nothing to show, as we did not find an association between sleep and volume.

### Brain-Stem

#### Descriptive statistics

| Study | Observations | Unique IDs | Mean age | Age range |
|---|---|---|---|---|
| HCP | 974 | 974 | 28.8 | 22 - 37 |
| MPIB | 677 | 391 | 63.1 | 24 - 83 |
| UB | 113 | 39 | 70.9 | 64 - 81 |
| UCAM | 884 | 632 | 55.1 | 20 - 88 |
| UiO | 1474 | 803 | 49.3 | 20 - 88 |
| UKB | 45963 | 43127 | 64.5 | 45 - 83 |

| Study | Observations | Unique IDs | Mean age | Age range |
|---|---|---|---|---|
| UmU | 423 | 284 | 62.3 | 25 - 85 |
| UOXF | 769 | 769 | 69.8 | 60 - 85 |

##### Spaghetti plot

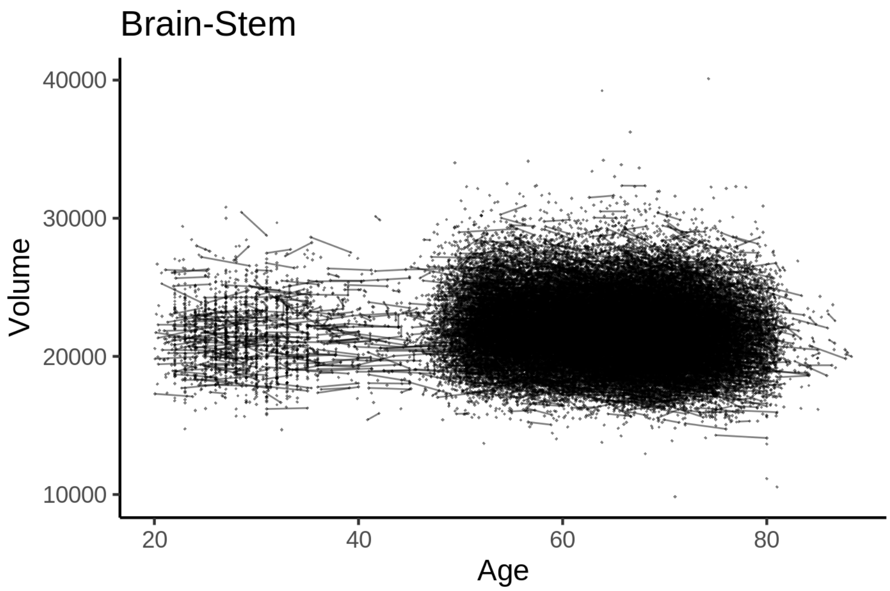


##### Model outputs

Model without sleep term

```
##
## Family: gaussian
## Link function: identity
##
## Formula:
## value ~ sex + site + icv + s(age_z, k = 10, bs = "cr")
## <environment: 0x55cffba7dac0>
##
## Parametric coefficients:
##                 Estimate Std. Error t value Pr(>|t|)
## (Intercept)    20850.985     93.007 224.188  < 2e-16 ***
## sexmale          799.227     20.272  39.424  < 2e-16 ***
## siteMPIB        2479.723    124.152  19.973  < 2e-16 ***
## siteousAvanto     -6.604     96.087  -0.069   0.9452
## siteousPrisma    256.268    158.127   1.621   0.1051
## siteousSkyra     670.917     94.273   7.117 1.12e-12 ***
## siteUB          -292.806    299.802  -0.977   0.3287
## siteUCAM        -116.359    106.881  -1.089   0.2763
## siteUKB          750.462     95.155   7.887 3.16e-15 ***
## siteUmU        -1590.851    138.225 -11.509  < 2e-16 ***
## siteUOXF         317.696    115.158   2.759   0.0058 **
## icv             1480.207     10.143 145.930  < 2e-16 ***
```

```
## ---
## Signif. codes:  0 ‘***’ 0.001 ‘**’ 0.01 ‘*’ 0.05 ‘.’ 0.1 ‘ ’ 1
##
## Approximate significance of smooth terms:
##             edf Ref.df     F p-value
## s(age_z) 6.792  6.792 130.2  <2e-16 ***
## ---
## Signif. codes:  0 ‘***’ 0.001 ‘**’ 0.01 ‘*’ 0.05 ‘.’ 0.1 ‘ ’ 1
##
## R-sq.(adj) =  0.486
## lmer.REML = 9.0188e+05  Scale est. = 1.013e+05  n = 51277
```

Model with only main effects of age and sleep

```
##
## Family: gaussian
## Link function: identity
##
## Formula:
## value ~ sex + site + icv + s(age_z, k = 10, bs = "cr") + s(sleep_z,
##     k = 5, bs = "cr")
## <environment: 0x55cffba7dac0>
##
## Parametric coefficients:
##                Estimate Std. Error t value Pr(>|t|)
## (Intercept)   20843.218     93.074 223.943  < 2e-16 ***
## sexmale         798.629     20.266  39.408  < 2e-16 ***
## siteMPIB       2497.217    124.176  20.110  < 2e-16 ***
## siteousAvanto    -3.718     96.088  -0.039  0.96914
## siteousPrisma   257.932    158.099   1.631  0.10280
## siteousSkyra    672.492     94.276   7.133 9.93e-13 ***
## siteUB         -281.535    299.670  -0.939  0.34749
## siteUCAM       -107.309    106.855  -1.004  0.31526
## siteUKB         758.722     95.233   7.967 1.66e-15 ***
## siteUmU       -1550.020    138.482 -11.193  < 2e-16 ***
## siteUOXF        321.189    115.131   2.790  0.00528 **
## icv            1479.841     10.151 145.788  < 2e-16 ***
## ---
## Signif. codes:  0 ‘***’ 0.001 ‘**’ 0.01 ‘*’ 0.05 ‘.’ 0.1 ‘ ’ 1
##
## Approximate significance of smooth terms:
##               edf Ref.df       F  p-value
## s(age_z)   6.787  6.787 125.050  < 2e-16 ***
## s(sleep_z) 3.585  3.585   9.959 1.03e-07 ***
## ---
## Signif. codes:  0 ‘***’ 0.001 ‘**’ 0.01 ‘*’ 0.05 ‘.’ 0.1 ‘ ’ 1
##
## R-sq.(adj) =  0.487
## lmer.REML = 9.0185e+05  Scale est. = 1.0137e+05  n = 51277
```

Model with full interaction between age and sleep

```
##
## Family: gaussian
## Link function: identity
##
```

```
## Formula:
## value ~ sex + site + icv + t2(age_z, sleep_z, k = c(10, 4), bs = "cr")
## <environment: 0x55cffba7dac0>
##
## Parametric coefficients:
##                Estimate Std. Error t value Pr(>|t|)
## (Intercept)    20819.17      92.62 224.788  < 2e-16 ***
## sexmale          797.91      20.31  39.295  < 2e-16 ***
## siteMPIB        2518.88     124.33  20.260  < 2e-16 ***
## siteousAvanto     16.97      96.37   0.176  0.86020
## siteousPrisma    277.04     158.42   1.749  0.08033 .
## siteousSkyra     695.75      94.58   7.356 1.92e-13 ***
## siteUB          -268.04     299.61  -0.895  0.37099
## siteUCAM         -89.37     106.95  -0.836  0.40336
## siteUKB          783.81      94.72   8.275  < 2e-16 ***
## siteUmU        -1531.87     138.37 -11.071  < 2e-16 ***
## siteUOXF         343.02     114.89   2.986  0.00283 **
## icv             1480.92      10.15 145.957  < 2e-16 ***
## ---
## Signif. codes:  0 ‘***’ 0.001 ‘**’ 0.01 ‘*’ 0.05 ‘.’ 0.1 ‘ ’ 1
##
## Approximate significance of smooth terms:
##                    edf Ref.df     F p-value
## t2(age_z,sleep_z)  11     11 12.47  <2e-16 ***
## ---
## Signif. codes:  0 ‘***’ 0.001 ‘**’ 0.01 ‘*’ 0.05 ‘.’ 0.1 ‘ ’ 1
##
## R-sq.(adj) =  0.487
## lmer.REML = 9.0186e+05  Scale est. = 1.0137e+05  n = 51277
```

#### Model comparison

`mod_no_sleep` refers to model without sleep term, `mod_no_interaction` refers to model with only main effect of sleep, and `mod_full` refers to model with a full interaction between age and sleep. This is a nested model comparison, and the p-value at a given line refers to comparing the model at the line to the model on the line above. Hence, significance implies that the more complicated model is supported on statistical grounds.

To be even more specific, the p-value on the second row tests whether there is an association between sleep and volume. The p-value on the third row tests whether this association depends on age.

```
## Data: NULL
## Models:
## mod_list$mod_no_sleep$mer: NULL
## mod_list$mod_no_interaction$mer: NULL
## mod_list$mod_full$mer: NULL
##                                 npar    AIC    BIC  logLik deviance  Chisq Df Pr(>Chisq)
## mod_list$mod_no_sleep$mer         16 901914 902056 -450941   901882
## mod_list$mod_no_interaction$mer   18 901888 902048 -450926   901852 29.867  2  3.269e-07 ***
## mod_list$mod_full$mer             20 901896 902073 -450928   901856  0.000  2          1
## ---
## Signif. codes:  0 ‘***’ 0.001 ‘**’ 0.01 ‘*’ 0.05 ‘.’ 0.1 ‘ ’ 1
```

We chose the model based on the likelihood ratio test with 5 % significance level, which was `mod_no_interaction`.

#### Lifespan brain trajectory

The trajectory shown is from the chosen model `mod_no_interaction`.

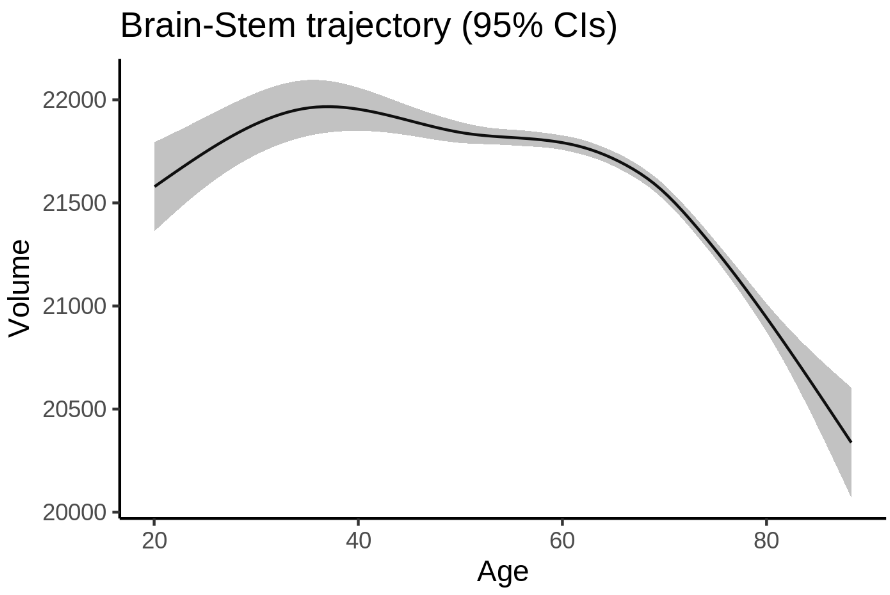


#### Effect of sleep

The chosen model only included the main effect of sleep, and hence the effect does not vary with age. The black dot shows the average sleep duration across all ages in the sample.

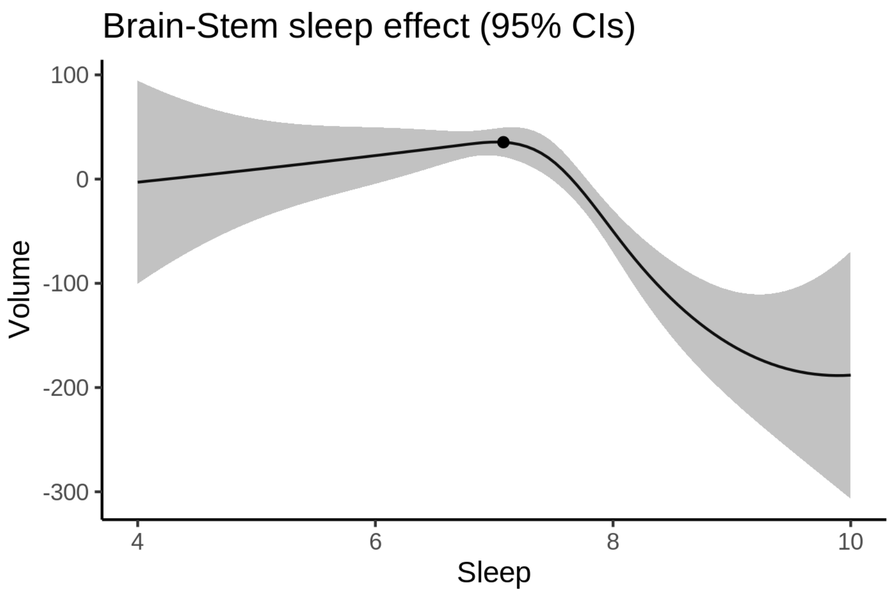


We also show the full interaction model for completeness, although it was not selected.

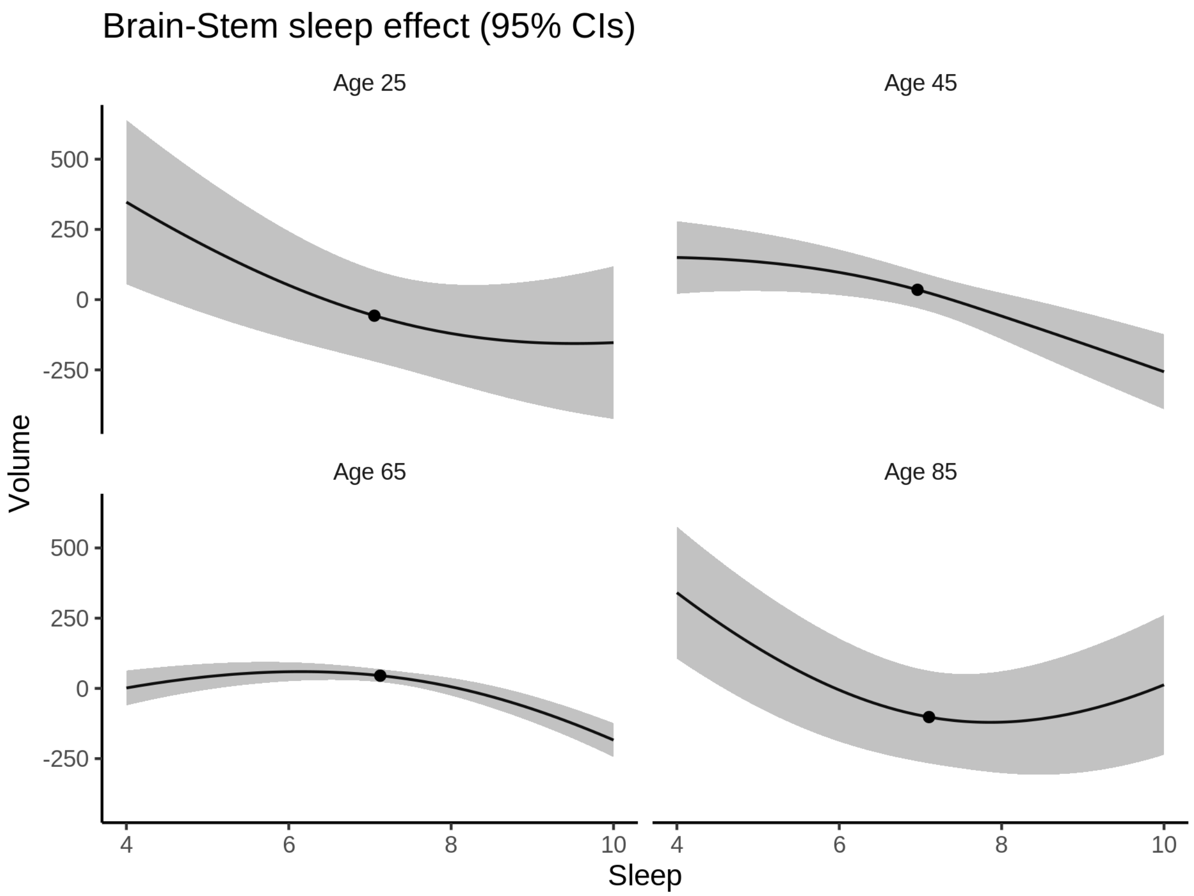


**Deviation from sleep associated with maximal volume**

Model with only main effect of sleep was chosen, so we show it for all ages at once. Maximum volume is attained at 7 hours of sleep. The percentage values in the plot are calculated as follows: The maximum at 100 % refers to a person at an arbitrary age with a sleep duration associated with maximum volume. For a female, this volume is 21554 and for a male it is 22353. The other percentage values show how large the expected volume is for someone with other sleep durations. For example, 99 % implies a 1 % reduction.

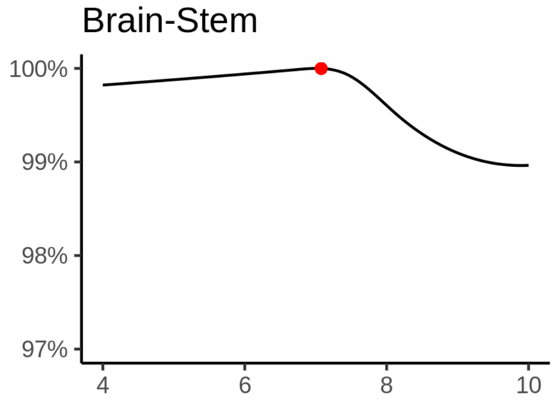

**Comparison of mean sleep and sleep associated with maximum volume**

A 95 % confidence interval for the sleep associated with maximum volume is [4, 7.21].

The plot below compares average sleep to the sleep associated with maximum volume.

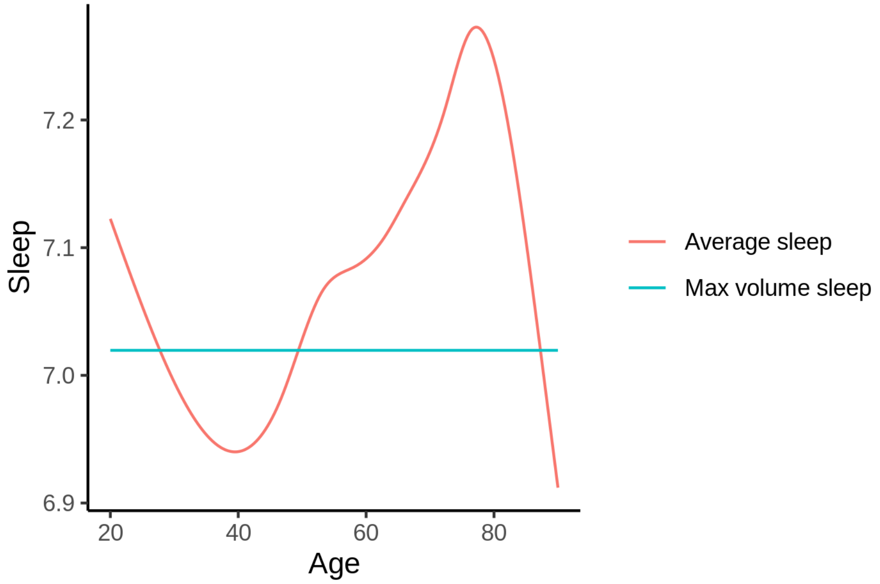


The next plot shows the difference between average sleep and sleep associated with maximum volume. The shaded region is a 95 % confidence interval.

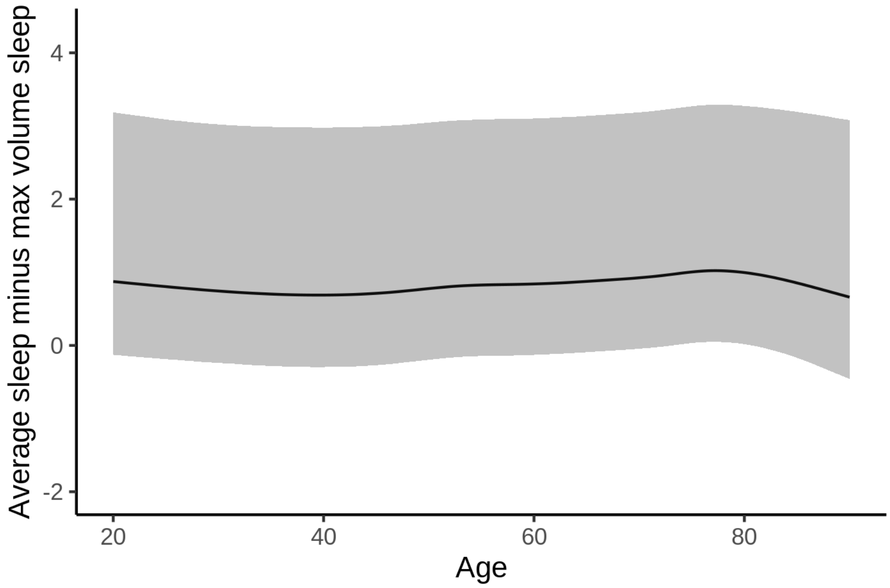


The next plot shows the probability that the sleep duration associated with maximum volume is longer than the average sleep duration, as a function of age. Probility below .05 can be interpreted as evidence that the sleep associated with maximum volume is shorter than the mean sleep, and probability above .95 can be interpreted the opposite way.

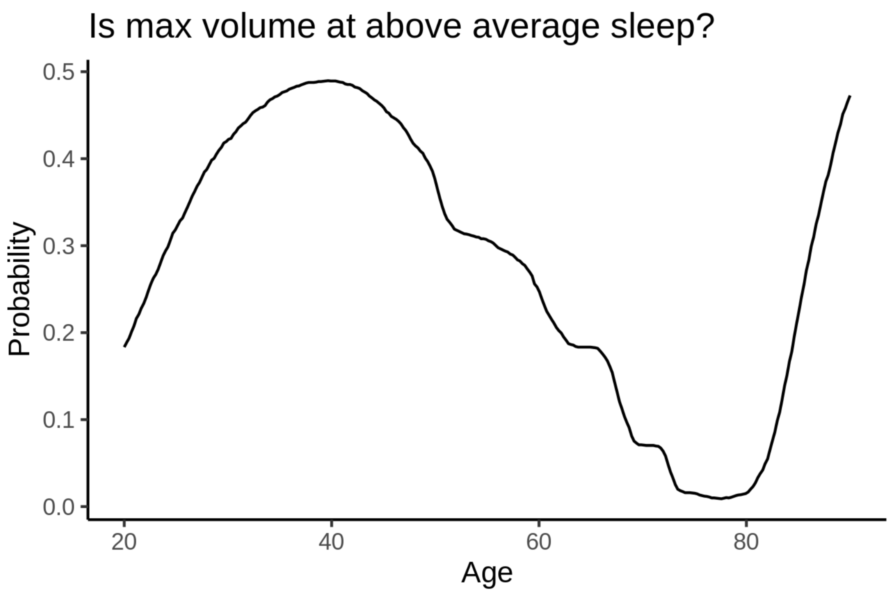


##### Controling for covariates

Below is the output for a model in which we only include data with income and education.

```
##
## Family: gaussian
## Link function: identity
##
## Formula:
## value ~ sex + site + s(age_z, k = 10, bs = "cr") + s(sleep_z,
##     k = 5, bs = "cr") + icv
## <environment: 0x55cffaf4b048>
##
## Parametric coefficients:
##               Estimate Std. Error t value Pr(>|t|)
## (Intercept)   23651.60     235.72 100.338  < 2e-16 ***
## sexmale         745.12      25.82  28.856  < 2e-16 ***
## siteousAvanto -3029.84     268.46 -11.286  < 2e-16 ***
## siteousPrisma -2324.05     284.19  -8.178 3.00e-16 ***
## siteousSkyra  -2235.92     254.91  -8.772  < 2e-16 ***
## siteUKB       -2024.83     235.21  -8.609  < 2e-16 ***
## siteUOXF      -2140.67     264.66  -8.088 6.27e-16 ***
## icv            1558.00      13.21 117.978  < 2e-16 ***
## ---
## Signif. codes:  0 ‘***’ 0.001 ‘**’ 0.01 ‘*’ 0.05 ‘.’ 0.1 ‘ ’ 1
##
## Approximate significance of smooth terms:
##              edf Ref.df      F  p-value
## s(age_z)   6.341  6.341 54.422  < 2e-16 ***
## s(sleep_z) 3.124  3.124  8.596 7.43e-06 ***
## ---
## Signif. codes:  0 ‘***’ 0.001 ‘**’ 0.01 ‘*’ 0.05 ‘.’ 0.1 ‘ ’ 1
##
```

```
## R-sq.(adj) =  0.489
## lmer.REML = 5.4865e+05  Scale est. = 1.0297e+05  n = 31180
```

Below is the output for a model in which we control for the main effects of income and education.

```
##
## Family: gaussian
## Link function: identity
##
## Formula:
## value ~ sex + site + s(age_z, k = 10, bs = "cr") + s(sleep_z,
##     k = 5, bs = "cr") + icv + income_scaled + education_scaled
## <environment: 0x55cffaf4b048>
##
## Parametric coefficients:
##                  Estimate Std. Error t value Pr(>|t|)
## (Intercept)      23423.07     236.97  98.844  < 2e-16 ***
## sexmale            745.41      25.86  28.823  < 2e-16 ***
## siteousAvanto    -3025.99     268.24 -11.281  < 2e-16 ***
## siteousPrisma    -2339.71     283.89  -8.242  < 2e-16 ***
## siteousSkyra     -2237.19     254.66  -8.785  < 2e-16 ***
## siteUKB          -2025.90     235.14  -8.616  < 2e-16 ***
## siteUOXF         -2069.34     264.54  -7.822 5.35e-15 ***
## icv               1543.52      13.30 116.082  < 2e-16 ***
## income_scaled      154.01      30.72   5.014 5.36e-07 ***
## education_scaled   200.91      36.41   5.518 3.46e-08 ***
## ---
## Signif. codes:  0 ‘***’ 0.001 ‘**’ 0.01 ‘*’ 0.05 ‘.’ 0.1 ‘ ’ 1
##
## Approximate significance of smooth terms:
##              edf Ref.df      F  p-value
## s(age_z)   6.485  6.485 43.181  < 2e-16 ***
## s(sleep_z) 2.866  2.866  7.319 0.000102 ***
## ---
## Signif. codes:  0 ‘***’ 0.001 ‘**’ 0.01 ‘*’ 0.05 ‘.’ 0.1 ‘ ’ 1
##
## R-sq.(adj) =   0.49
## lmer.REML = 5.4856e+05  Scale est. = 1.0308e+05  n = 31180
```

We also included interaction effects between sleep duration and education and income, in another model. The output is shown below, and the interaction terms are `income_scaled:sleep_z` and `education_scaled:sleep_z`.

```
##
## Family: gaussian
## Link function: identity
##
## Formula:
## value ~ sex + site + s(age_z, k = 10, bs = "cr") + s(sleep_z,
##     k = 5, bs = "cr") + icv + income_scaled + education_scaled +
##     income_scaled:sleep_z + education_scaled:sleep_z
## <environment: 0x55cffaf4b048>
##
## Parametric coefficients:
##                          Estimate Std. Error t value Pr(>|t|)
## (Intercept)              23425.02     236.98  98.848  < 2e-16 ***
```

```
## sexmale                     743.13      25.88  28.717  < 2e-16 ***
## siteousAvanto            -3030.57     268.24 -11.298  < 2e-16 ***
## siteousPrisma            -2348.82     283.90  -8.273  < 2e-16 ***
## siteousSkyra             -2241.56     254.65  -8.802  < 2e-16 ***
## siteUKB                  -2028.06     235.16  -8.624  < 2e-16 ***
## siteUOXF                 -2070.95     264.61  -7.826 5.18e-15 ***
## icv                       1543.83      13.30 116.102  < 2e-16 ***
## income_scaled              156.22      30.73   5.084 3.72e-07 ***
## education_scaled           199.46      36.42   5.477 4.35e-08 ***
## income_scaled:sleep_z      -62.14      30.71  -2.024   0.0430 *
## education_scaled:sleep_z    60.43      35.34   1.710   0.0873 .
## ---
## Signif. codes:  0 ‘***’ 0.001 ‘**’ 0.01 ‘*’ 0.05 ‘.’ 0.1 ‘ ’ 1
##
## Approximate significance of smooth terms:
##              edf Ref.df      F p-value
## s(age_z)   6.498  6.498 42.991 < 2e-16 ***
## s(sleep_z) 2.850  2.850  4.668 0.00385 **
## ---
## Signif. codes:  0 ‘***’ 0.001 ‘**’ 0.01 ‘*’ 0.05 ‘.’ 0.1 ‘ ’ 1
##
## R-sq.(adj) =   0.49
## lmer.REML = 5.4853e+05  Scale est. = 1.0307e+05  n = 31180
```

We did the same controling for BMI. Below is the model with no covariates but only keeping data with BMI.

```
##
## Family: gaussian
## Link function: identity
##
## Formula:
## value ~ sex + site + s(age_z, k = 10, bs = "cr") + s(sleep_z,
##     k = 5, bs = "cr") + icv
## <environment: 0x55cf3be23950>
##
## Parametric coefficients:
##               Estimate Std. Error t value Pr(>|t|)
## (Intercept)   20966.08     129.42 161.997  < 2e-16 ***
## sexmale         734.31      25.06  29.307  < 2e-16 ***
## siteousPrisma   259.38     171.25   1.515    0.130
## siteousSkyra    760.07      80.95   9.389  < 2e-16 ***
## siteUCAM       -149.77     140.94  -1.063    0.288
## siteUKB         661.69     130.93   5.054 4.35e-07 ***
## siteUmU       -1499.67     166.48  -9.008  < 2e-16 ***
## icv            1560.44      12.85 121.416  < 2e-16 ***
## ---
## Signif. codes:  0 ‘***’ 0.001 ‘**’ 0.01 ‘*’ 0.05 ‘.’ 0.1 ‘ ’ 1
##
## Approximate significance of smooth terms:
##              edf Ref.df     F  p-value
## s(age_z)   6.291  6.291 69.69  < 2e-16 ***
## s(sleep_z) 2.773  2.773  8.38 3.55e-05 ***
## ---
## Signif. codes:  0 ‘***’ 0.001 ‘**’ 0.01 ‘*’ 0.05 ‘.’ 0.1 ‘ ’ 1
##
```

```
## R-sq.(adj) =  0.495
## lmer.REML = 5.8762e+05  Scale est. = 99124     n = 33435
```

Below is the model output with main effect.

```
##
## Family: gaussian
## Link function: identity
##
## Formula:
## value ~ sex + site + s(age_z, k = 10, bs = "cr") + s(sleep_z,
##     k = 5, bs = "cr") + icv + bmi
## <environment: 0x55cf3be23950>
##
## Parametric coefficients:
##                 Estimate Std. Error t value Pr(>|t|)
## (Intercept)   21422.114    143.278 149.515  < 2e-16 ***
## sexmale         748.728     25.110  29.818  < 2e-16 ***
## siteousPrisma   271.451    171.189   1.586    0.113
## siteousSkyra    760.818     80.955   9.398  < 2e-16 ***
## siteUCAM       -139.823    140.873  -0.993    0.321
## siteUKB         677.335    130.892   5.175 2.30e-07 ***
## siteUmU       -1479.725    166.404  -8.892  < 2e-16 ***
## icv            1563.260     12.846 121.692  < 2e-16 ***
## bmi             -18.139      2.448  -7.410 1.29e-13 ***
## ---
## Signif. codes:  0 '***' 0.001 '**' 0.01 '*' 0.05 '.' 0.1 ' ' 1
##
## Approximate significance of smooth terms:
##               edf Ref.df      F  p-value
## s(age_z)   6.310  6.310 70.563  < 2e-16 ***
## s(sleep_z) 2.625  2.625  9.495 3.16e-05 ***
## ---
## Signif. codes:  0 '***' 0.001 '**' 0.01 '*' 0.05 '.' 0.1 ' ' 1
##
## R-sq.(adj) =  0.496
## lmer.REML = 5.8757e+05  Scale est. = 99128     n = 33435
```

Next is the model with BMI-sleep interaction.

```
##
## Family: gaussian
## Link function: identity
##
## Formula:
## value ~ sex + site + s(age_z, k = 10, bs = "cr") + s(sleep_z,
##     k = 5, bs = "cr") + icv + bmi + bmi:sleep_z
## <environment: 0x55cf3be23950>
##
## Parametric coefficients:
##                 Estimate Std. Error t value Pr(>|t|)
## (Intercept)   21418.116    143.363 149.398  < 2e-16 ***
## sexmale         748.806     25.110  29.821  < 2e-16 ***
## siteousPrisma   271.425    171.188   1.586    0.113
## siteousSkyra    761.074     80.954   9.401  < 2e-16 ***
## siteUCAM       -138.552    140.881  -0.983    0.325
```

```
## siteUKB         678.711    130.900   5.185 2.17e-07 ***
## siteUmU       -1478.667    166.409  -8.886  < 2e-16 ***
## icv            1563.355     12.846 121.696  < 2e-16 ***
## bmi             -18.075      2.449  -7.380 1.63e-13 ***
## bmi:sleep_z       1.873      2.322   0.807    0.420
## ---
## Signif. codes:  0 '***' 0.001 '**' 0.01 '*' 0.05 '.' 0.1 ' ' 1
##
## Approximate significance of smooth terms:
##              edf Ref.df      F p-value
## s(age_z)   6.307  6.307 70.662  <2e-16 ***
## s(sleep_z) 2.578  2.578  3.756  0.0284 *
## ---
## Signif. codes:  0 '***' 0.001 '**' 0.01 '*' 0.05 '.' 0.1 ' ' 1
##
## R-sq.(adj) =  0.496
## lmer.REML = 5.8756e+05  Scale est. = 99124     n = 33435
```

We did the same controling for depression. Below is the model with no covariates but only keeping data with depression.

```
##
## Family: gaussian
## Link function: identity
##
## Formula:
## value ~ sex + site + s(age_z, k = 10, bs = "cr") + s(sleep_z,
##     k = 5, bs = "cr") + icv
## <environment: 0x55cf3fe20f08>
##
## Parametric coefficients:
##               Estimate Std. Error t value Pr(>|t|)
## (Intercept)   23161.49     120.87 191.626  < 2e-16 ***
## sexmale         739.43      25.17  29.372  < 2e-16 ***
## siteousAvanto -2316.00     214.47 -10.799  < 2e-16 ***
## siteousPrisma -1773.87     218.92  -8.103 5.56e-16 ***
## siteousSkyra  -1343.77     176.54  -7.612 2.78e-14 ***
## siteUCAM      -2487.74     164.91 -15.086  < 2e-16 ***
## siteUKB       -1536.00     119.56 -12.848  < 2e-16 ***
## siteUmU       -3701.94     161.71 -22.893  < 2e-16 ***
## icv            1551.00      12.89 120.284  < 2e-16 ***
## ---
## Signif. codes:  0 '***' 0.001 '**' 0.01 '*' 0.05 '.' 0.1 ' ' 1
##
## Approximate significance of smooth terms:
##              edf Ref.df      F p-value
## s(age_z)   6.521  6.521 72.335 < 2e-16 ***
## s(sleep_z) 2.714  2.714  8.015 6.7e-05 ***
## ---
## Signif. codes:  0 '***' 0.001 '**' 0.01 '*' 0.05 '.' 0.1 ' ' 1
##
## R-sq.(adj) =  0.491
## lmer.REML = 5.8578e+05  Scale est. = 98306     n = 33347
```

Below is the model output with main effect.

```
##
## Family: gaussian
## Link function: identity
##
## Formula:
## value ~ sex + site + s(age_z, k = 10, bs = "cr") + s(sleep_z,
##     k = 5, bs = "cr") + icv + depression
## <environment: 0x55cf3fe20f08>
##
## Parametric coefficients:
##                Estimate Std. Error t value Pr(>|t|)
## (Intercept)    23209.23     121.34 191.270  < 2e-16 ***
## sexmale          734.39      25.19  29.149  < 2e-16 ***
## siteousAvanto -2290.77     214.46 -10.682  < 2e-16 ***
## siteousPrisma -1750.31     218.90  -7.996 1.33e-15 ***
## siteousSkyra  -1315.69     176.58  -7.451 9.49e-14 ***
## siteUCAM      -2476.05     164.88 -15.017  < 2e-16 ***
## siteUKB       -1555.49     119.61 -13.004  < 2e-16 ***
## siteUmU       -3556.25     165.11 -21.539  < 2e-16 ***
## icv            1551.73      12.89 120.366  < 2e-16 ***
## depression     -391.89      90.18  -4.345 1.39e-05 ***
## ---
## Signif. codes:  0 '***' 0.001 '**' 0.01 '*' 0.05 '.' 0.1 ' ' 1
##
## Approximate significance of smooth terms:
##              edf Ref.df      F  p-value
## s(age_z)   6.496  6.496 74.864  < 2e-16 ***
## s(sleep_z) 2.553  2.553  8.151 7.53e-05 ***
## ---
## Signif. codes:  0 '***' 0.001 '**' 0.01 '*' 0.05 '.' 0.1 ' ' 1
##
## R-sq.(adj) =  0.491
## lmer.REML = 5.8575e+05  Scale est. = 98228     n = 33347
```

Next is the model with depression-sleep interaction.

```
##
## Family: gaussian
## Link function: identity
##
## Formula:
## value ~ sex + site + s(age_z, k = 10, bs = "cr") + s(sleep_z,
##     k = 5, bs = "cr") + icv + depression + depression:sleep_z
## <environment: 0x55cf3fe20f08>
##
## Parametric coefficients:
##                     Estimate Std. Error t value Pr(>|t|)
## (Intercept)         23209.22     121.34 191.267  < 2e-16 ***
## sexmale               734.52      25.20  29.149  < 2e-16 ***
## siteousAvanto       -2289.93     214.48 -10.677  < 2e-16 ***
## siteousPrisma       -1749.47     218.93  -7.991 1.38e-15 ***
## siteousSkyra        -1314.89     176.61  -7.445 9.90e-14 ***
## siteUCAM            -2475.14     164.92 -15.008  < 2e-16 ***
## siteUKB             -1555.51     119.61 -13.004  < 2e-16 ***
## siteUmU             -3562.32     166.52 -21.392  < 2e-16 ***
```

```
## icv                  1551.72      12.89 120.363  < 2e-16 ***
## depression           -390.42      90.35  -4.321 1.56e-05 ***
## depression:sleep_z     18.66      66.46   0.281    0.779
## ---
## Signif. codes:  0 '***' 0.001 '**' 0.01 '*' 0.05 '.' 0.1 ' ' 1
##
## Approximate significance of smooth terms:
##               edf Ref.df      F  p-value
## s(age_z)    6.498  6.498 74.854  < 2e-16 ***
## s(sleep_z) 2.523  2.523  6.582 0.000394 ***
## ---
## Signif. codes:  0 '***' 0.001 '**' 0.01 '*' 0.05 '.' 0.1 ' ' 1
##
## R-sq.(adj) =  0.491
## lmer.REML = 5.8574e+05  Scale est. = 98228     n = 33347
```

The plot below shows the sleep-volume curve for the original model and for the model with main effects of SES.

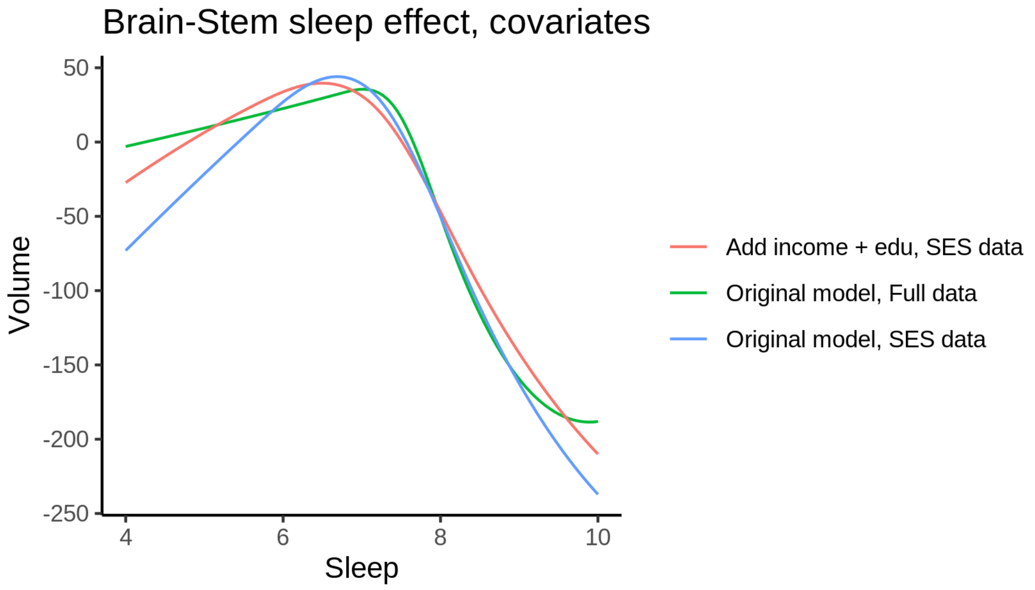


The plot below shows the sleep-volume curve for the original model and for the model with main effects of BMI.

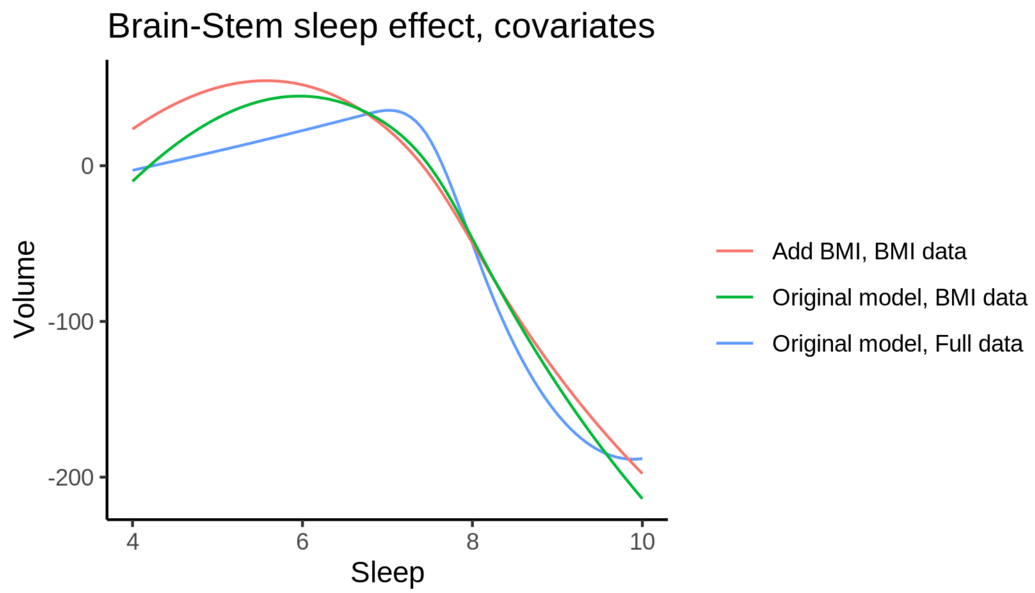


The plot below shows the sleep-volume curve for the original model and for the model with main effects of depression.

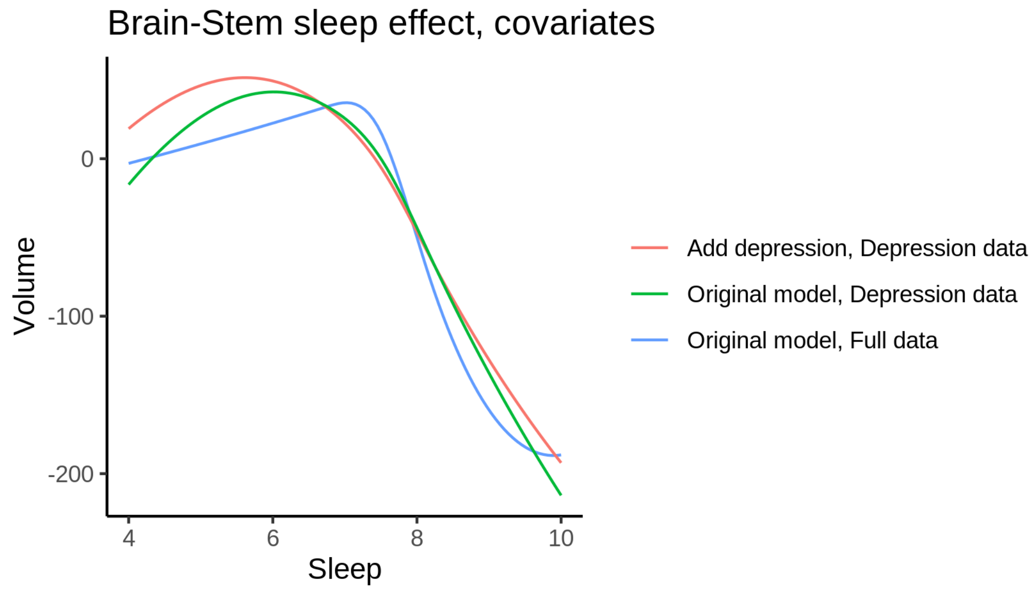


### Caudate

#### Descriptive statistics

| Study | Observations | Unique IDs | Mean age | Age range |
|---|---|---|---|---|
| HCP | 974 | 974 | 28.8 | 22 - 37 |
| MPIB | 675 | 390 | 63.2 | 24 - 83 |
| UB | 113 | 39 | 70.9 | 64 - 81 |
| UCAM | 884 | 632 | 55.1 | 20 - 88 |

| Study | Observations | Unique IDs | Mean age | Age range |
|---|---|---|---|---|
| UiO | 1473 | 803 | 49.3 | 20 - 88 |
| UKB | 45976 | 43134 | 64.5 | 45 - 83 |
| UmU | 423 | 284 | 62.3 | 25 - 85 |
| UOXF | 769 | 769 | 69.8 | 60 - 85 |

##### Spaghetti plot

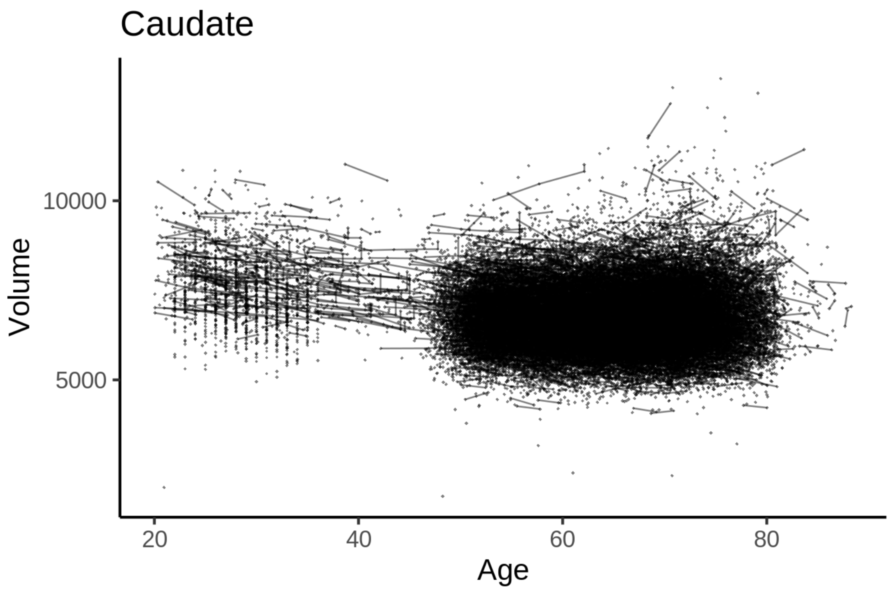


##### Model outputs

Model without sleep term

```
##
## Family: gaussian
## Link function: identity
##
## Formula:
## value ~ sex + site + icv + s(age_z, k = 10, bs = "cr")
## <environment: 0x55cf3d115068>
##
## Parametric coefficients:
##                Estimate Std. Error t value Pr(>|t|)
## (Intercept)   6551.365     39.661 165.182  < 2e-16 ***
## sexmale         18.801      8.356   2.250 0.024446 *
## siteMPIB      1253.335     51.828  24.183  < 2e-16 ***
## siteousAvanto  497.065     40.412  12.300  < 2e-16 ***
## siteousPrisma  830.657     67.572  12.293  < 2e-16 ***
## siteousSkyra   836.695     39.298  21.291  < 2e-16 ***
## siteUB         554.003    123.505   4.486 7.28e-06 ***
## siteUCAM       618.390     44.688  13.838  < 2e-16 ***
## siteUKB        149.431     40.629   3.678 0.000235 ***
## siteUmU        798.836     57.655  13.856  < 2e-16 ***
```

```
## siteUOXF       631.752     48.599  12.999  < 2e-16 ***
## icv            462.244      4.189 110.344  < 2e-16 ***
## ---
## Signif. codes:  0 ‘***’ 0.001 ‘**’ 0.01 ‘*’ 0.05 ‘.’ 0.1 ‘ ’ 1
##
## Approximate significance of smooth terms:
##            edf Ref.df     F p-value
## s(age_z) 7.795  7.795 76.84  <2e-16 ***
## ---
## Signif. codes:  0 ‘***’ 0.001 ‘**’ 0.01 ‘*’ 0.05 ‘.’ 0.1 ‘ ’ 1
##
## R-sq.(adj) =   0.33
## lmer.REML = 8.1237e+05  Scale est. = 23485     n = 51287
```

Model with only main effects of age and sleep

```
##
## Family: gaussian
## Link function: identity
##
## Formula:
## value ~ sex + site + icv + s(age_z, k = 10, bs = "cr") + s(sleep_z,
##     k = 5, bs = "cr")
## <environment: 0x55cf3d115068>
##
## Parametric coefficients:
##                Estimate Std. Error t value Pr(>|t|)
## (Intercept)    6554.791     39.693 165.138  < 2e-16 ***
## sexmale          18.925      8.356   2.265 0.023522 *
## siteMPIB       1249.564     51.858  24.096  < 2e-16 ***
## siteousAvanto   495.199     40.420  12.251  < 2e-16 ***
## siteousPrisma   829.348     67.571  12.274  < 2e-16 ***
## siteousSkyra    834.795     39.307  21.238  < 2e-16 ***
## siteUB          553.092    123.501   4.478 7.54e-06 ***
## siteUCAM        616.697     44.693  13.799  < 2e-16 ***
## siteUKB         145.789     40.664   3.585 0.000337 ***
## siteUmU         790.969     57.782  13.689  < 2e-16 ***
## siteUOXF        630.095     48.602  12.964  < 2e-16 ***
## icv             461.988      4.191 110.236  < 2e-16 ***
## ---
## Signif. codes:  0 ‘***’ 0.001 ‘**’ 0.01 ‘*’ 0.05 ‘.’ 0.1 ‘ ’ 1
##
## Approximate significance of smooth terms:
##              edf Ref.df      F p-value
## s(age_z)   7.788  7.788 76.387  <2e-16 ***
## s(sleep_z) 1.000  1.000  4.043  0.0444 *
## ---
## Signif. codes:  0 ‘***’ 0.001 ‘**’ 0.01 ‘*’ 0.05 ‘.’ 0.1 ‘ ’ 1
##
## R-sq.(adj) =   0.33
## lmer.REML = 8.1236e+05  Scale est. = 23479     n = 51287
```

Model with full interaction between age and sleep

```
##
## Family: gaussian
```

```
## Link function: identity
##
## Formula:
## value ~ sex + site + icv + t2(age_z, sleep_z, k = c(10, 4), bs = "cr")
## <environment: 0x55cf3d115068>
##
## Parametric coefficients:
##                Estimate Std. Error t value Pr(>|t|)
## (Intercept)   6557.691     39.940 164.188  < 2e-16 ***
## sexmale         19.850      8.372   2.371   0.0177 *
## siteMPIB      1246.088     52.157  23.891  < 2e-16 ***
## siteousAvanto  493.928     40.689  12.139  < 2e-16 ***
## siteousPrisma  827.509     67.732  12.217  < 2e-16 ***
## siteousSkyra   832.673     39.597  21.029  < 2e-16 ***
## siteUB         548.279    123.611   4.436  9.2e-06 ***
## siteUCAM       610.893     44.983  13.581  < 2e-16 ***
## siteUKB        142.420     40.917   3.481   0.0005 ***
## siteUmU        790.605     58.012  13.628  < 2e-16 ***
## siteUOXF       624.235     48.858  12.777  < 2e-16 ***
## icv            461.840      4.192 110.183  < 2e-16 ***
## ---
## Signif. codes:  0 ‘***’ 0.001 ‘**’ 0.01 ‘*’ 0.05 ‘.’ 0.1 ‘ ’ 1
##
## Approximate significance of smooth terms:
##                      edf Ref.df     F p-value
## t2(age_z,sleep_z) 12.38  12.38 10.34  <2e-16 ***
## ---
## Signif. codes:  0 ‘***’ 0.001 ‘**’ 0.01 ‘*’ 0.05 ‘.’ 0.1 ‘ ’ 1
##
## R-sq.(adj) =   0.33
## lmer.REML = 8.1237e+05  Scale est. = 23455     n = 51287
```

#### Model comparison

`mod_no_sleep` refers to model without sleep term, `mod_no_interaction` refers to model with only main effect of sleep, and `mod_full` refers to model with a full interaction between age and sleep. This is a nested model comparison, and the p-value at a given line refers to comparing the model at the line to the model on the line above. Hence, significance implies that the more complicated model is supported on statistical grounds.

To be even more specific, the p-value on the second row tests whether there is an association between sleep and volume. The p-value on the third row tests whether this association depends on age.

```
## Data: NULL
## Models:
## mod_list$mod_no_sleep$mer: NULL
## mod_list$mod_no_interaction$mer: NULL
## mod_list$mod_full$mer: NULL
##                                 npar    AIC    BIC  logLik deviance  Chisq Df Pr(>Chisq)
## mod_list$mod_no_sleep$mer         16 812399 812540 -406183   812367
## mod_list$mod_no_interaction$mer   18 812399 812558 -406181   812363 4.0426  2     0.1325
## mod_list$mod_full$mer             20 812407 812584 -406184   812367 0.0000  2     1.0000
```

We chose the model based on the likelihood ratio test with 5 % significance level, which was `mod_no_sleep`.

##### Lifespan brain trajectory

The trajectory shown is from the chosen model `mod_no_sleep`.

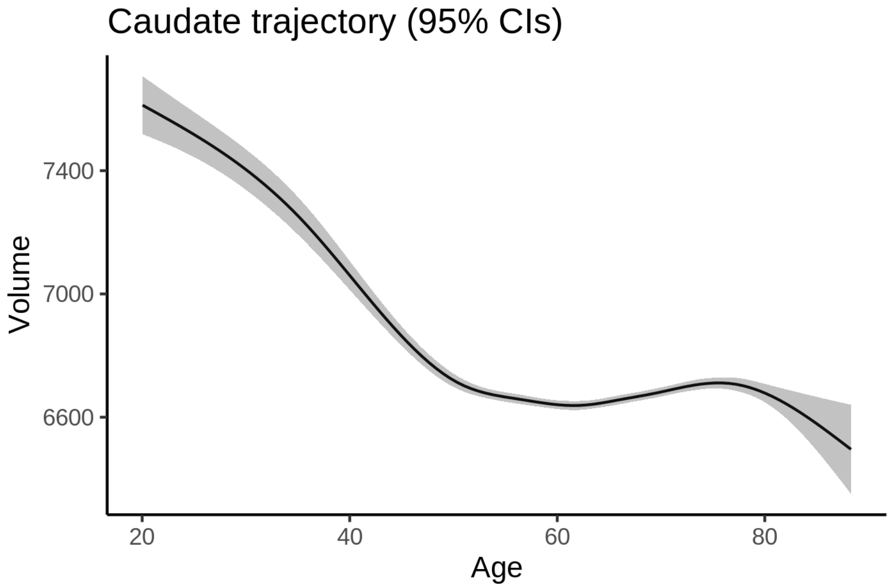


##### Effect of sleep

The chosen model did not include a sleep term, and hence we don't have any estimated effect of sleep.

We show the full interaction model for completeness, although it was not selected.

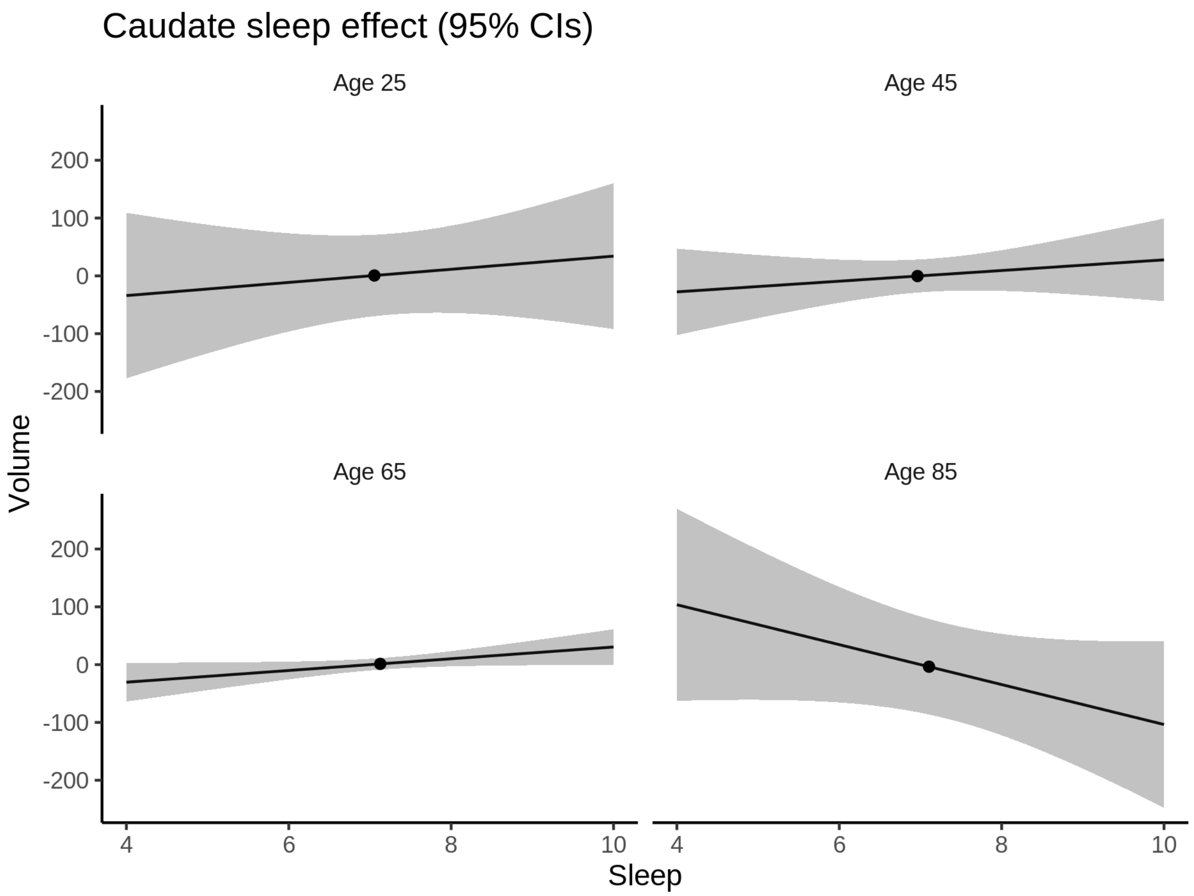


**Deviation from sleep associated with maximal volume**

Model with no sleep term was selected. No plots to show. (Although we can of course dig up the plots, which will be pretty flat).

**Comparison of mean sleep and sleep associated with maximum volume**

Nothing to show, as we did not find an association between sleep and volume.

### CC_Anterior

#### Descriptive statistics

| Study | Observations | Unique IDs | Mean age | Age range |
|---|---|---|---|---|
| HCP | 974 | 974 | 28.8 | 22 - 37 |
| MPIB | 677 | 391 | 63.1 | 24 - 83 |
| UB | 113 | 39 | 70.9 | 64 - 81 |
| UCAM | 884 | 632 | 55.1 | 20 - 88 |
| UiO | 1475 | 803 | 49.4 | 20 - 89 |
| UKB | 45969 | 43132 | 64.5 | 45 - 83 |

| Study | Observations | Unique IDs | Mean age | Age range |
|---|---|---|---|---|
| UmU | 423 | 284 | 62.3 | 25 - 85 |
| UOXF | 769 | 769 | 69.8 | 60 - 85 |

##### Spaghetti plot

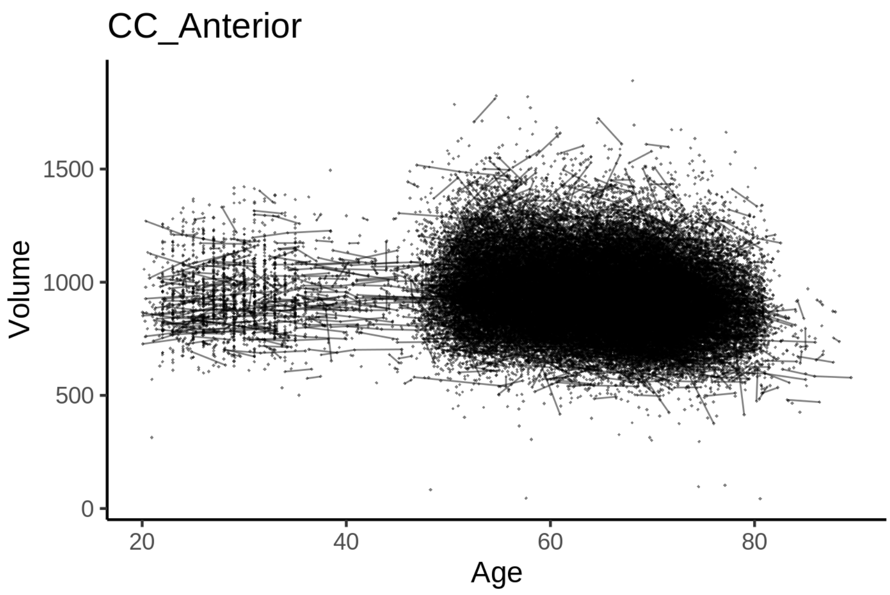


##### Model outputs

Model without sleep term

```
##
## Family: gaussian
## Link function: identity
##
## Formula:
## value ~ sex + site + icv + s(age_z, k = 10, bs = "cr")
## <environment: 0x55cffd3bfd10>
##
## Parametric coefficients:
##                Estimate Std. Error t value Pr(>|t|)
## (Intercept)    938.3374     8.0469 116.609  < 2e-16 ***
## sexmale        -40.6055     1.6406 -24.750  < 2e-16 ***
## siteMPIB        94.3007    10.2573   9.194  < 2e-16 ***
## siteousAvanto -71.3019     8.2722  -8.619  < 2e-16 ***
## siteousPrisma -31.7953    14.0082  -2.270 0.023226 *
## siteousSkyra  -32.8286     7.8115  -4.203 2.64e-05 ***
## siteUB        -59.3537    24.0465  -2.468 0.013580 *
## siteUCAM      -62.1113     8.8934  -6.984 2.90e-12 ***
## siteUKB        22.3476     8.2530   2.708 0.006775 **
## siteUmU       -10.4134    11.4377  -0.910 0.362592
## siteUOXF      -32.5601     9.7659  -3.334 0.000856 ***
## icv            80.9732     0.8249  98.159  < 2e-16 ***
```

```
## ---
## Signif. codes:  0 '***' 0.001 '**' 0.01 '*' 0.05 '.' 0.1 ' ' 1
##
## Approximate significance of smooth terms:
##            edf Ref.df     F p-value
## s(age_z) 7.374  7.374 130.6  <2e-16 ***
## ---
## Signif. codes:  0 '***' 0.001 '**' 0.01 '*' 0.05 '.' 0.1 ' ' 1
##
## R-sq.(adj) =  0.215
## lmer.REML = 6.4793e+05  Scale est. = 1716.7    n = 51284
```

Model with only main effects of age and sleep

```
##
## Family: gaussian
## Link function: identity
##
## Formula:
## value ~ sex + site + icv + s(age_z, k = 10, bs = "cr") + s(sleep_z,
##     k = 5, bs = "cr")
## <environment: 0x55cffd3bfd10>
##
## Parametric coefficients:
##              Estimate Std. Error t value Pr(>|t|)
## (Intercept)   937.2788     8.0545 116.367  < 2e-16 ***
## sexmale       -40.6425     1.6405 -24.774  < 2e-16 ***
## siteMPIB       95.4518    10.2638   9.300  < 2e-16 ***
## siteousAvanto -70.7360     8.2739  -8.549  < 2e-16 ***
## siteousPrisma -31.4120    14.0076  -2.242  0.02493 *
## siteousSkyra  -32.2450     7.8135  -4.127 3.68e-05 ***
## siteUB        -59.0261    24.0447  -2.455  0.01410 *
## siteUCAM      -61.5673     8.8947  -6.922 4.51e-12 ***
## siteUKB        23.4721     8.2614   2.841  0.00450 **
## siteUmU        -8.0777    11.4638  -0.705  0.48104
## siteUOXF      -32.0125     9.7672  -3.278  0.00105 **
## icv            81.0477     0.8252  98.213  < 2e-16 ***
## ---
## Signif. codes:  0 '***' 0.001 '**' 0.01 '*' 0.05 '.' 0.1 ' ' 1
##
## Approximate significance of smooth terms:
##              edf Ref.df       F p-value
## s(age_z)   7.383  7.383 128.422 < 2e-16 ***
## s(sleep_z) 1.000  1.000   8.783 0.00304 **
## ---
## Signif. codes:  0 '***' 0.001 '**' 0.01 '*' 0.05 '.' 0.1 ' ' 1
##
## R-sq.(adj) =  0.215
## lmer.REML = 6.4792e+05  Scale est. = 1716.7    n = 51284
```

Model with full interaction between age and sleep

```
##
## Family: gaussian
## Link function: identity
##
```

```
## Formula:
## value ~ sex + site + icv + t2(age_z, sleep_z, k = c(10, 4), bs = "cr")
## <environment: 0x55cffd3bfd10>
##
## Parametric coefficients:
##                Estimate Std. Error t value Pr(>|t|)
## (Intercept)   937.1934     7.9720 117.561  < 2e-16 ***
## sexmale       -40.6119     1.6440 -24.703  < 2e-16 ***
## siteMPIB       95.7500    10.2565   9.336  < 2e-16 ***
## siteousAvanto -70.6534     8.2809  -8.532  < 2e-16 ***
## siteousPrisma -31.6230    14.0311  -2.254 0.024215 *
## siteousSkyra  -32.2046     7.8248  -4.116 3.87e-05 ***
## siteUB        -58.9158    24.0198  -2.453 0.014178 *
## siteUCAM      -61.2326     8.8823  -6.894 5.50e-12 ***
## siteUKB        23.5381     8.1713   2.881 0.003971 **
## siteUmU        -7.7029    11.4294  -0.674 0.500345
## siteUOXF      -32.0463     9.7069  -3.301 0.000963 ***
## icv            80.9810     0.8254  98.116  < 2e-16 ***
## ---
## Signif. codes:  0 '***' 0.001 '**' 0.01 '*' 0.05 '.' 0.1 ' ' 1
##
## Approximate significance of smooth terms:
##                      edf Ref.df     F p-value
## t2(age_z,sleep_z) 11.57  11.57 10.79  <2e-16 ***
## ---
## Signif. codes:  0 '***' 0.001 '**' 0.01 '*' 0.05 '.' 0.1 ' ' 1
##
## R-sq.(adj) =  0.215
## lmer.REML = 6.4792e+05  Scale est. = 1716.7    n = 51284
```

#### Model comparison

`mod_no_sleep` refers to model without sleep term, `mod_no_interaction` refers to model with only main effect of sleep, and `mod_full` refers to model with a full interaction between age and sleep. This is a nested model comparison, and the p-value at a given line refers to comparing the model at the line to the model on the line above. Hence, significance implies that the more complicated model is supported on statistical grounds.

To be even more specific, the p-value on the second row tests whether there is an association between sleep and volume. The p-value on the third row tests whether this association depends on age.

```
## Data: NULL
## Models:
## mod_list$mod_no_sleep$mer: NULL
## mod_list$mod_no_interaction$mer: NULL
## mod_list$mod_full$mer: NULL
##                                 npar    AIC    BIC  logLik deviance  Chisq Df Pr(>Chisq)
## mod_list$mod_no_sleep$mer         16 647958 648100 -323963   647926
## mod_list$mod_no_interaction$mer   18 647953 648113 -323959   647917 8.7818  2    0.01239 *
## mod_list$mod_full$mer             20 647960 648137 -323960   647920 0.0000  2    1.00000
## ---
## Signif. codes:  0 '***' 0.001 '**' 0.01 '*' 0.05 '.' 0.1 ' ' 1
```

We chose the model based on the likelihood ratio test with 5 % significance level, which was `mod_no_interaction`.

##### Lifespan brain trajectory

The trajectory shown is from the chosen model `mod_no_interaction`.

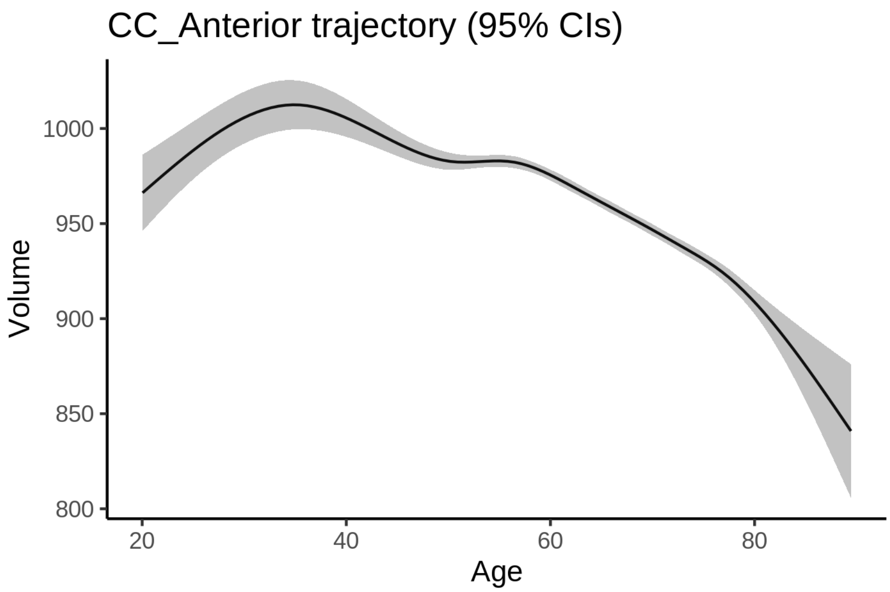


##### Effect of sleep

The chosen model only included the main effect of sleep, and hence the effect does not vary with age. The black dot shows the average sleep duration across all ages in the sample.

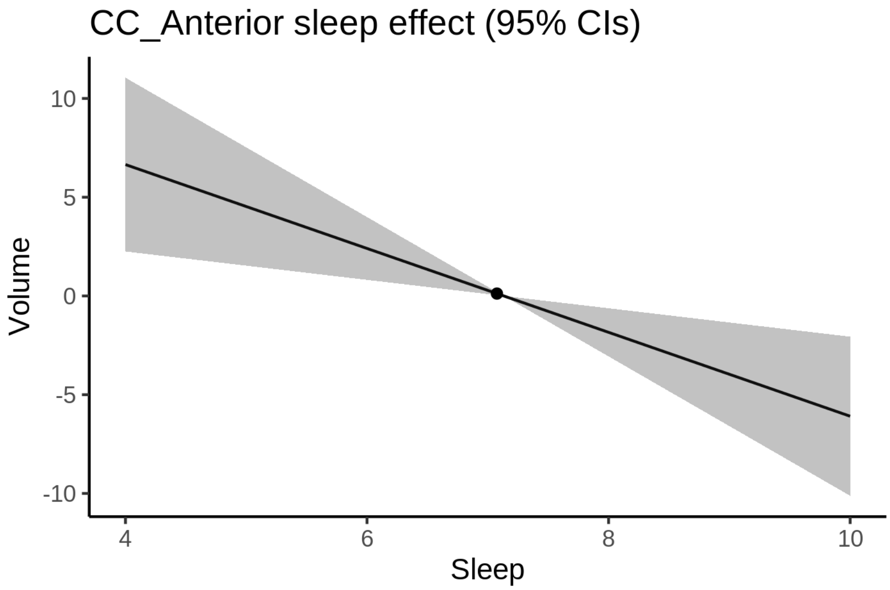


We also show the full interaction model for completeness, although it was not selected.

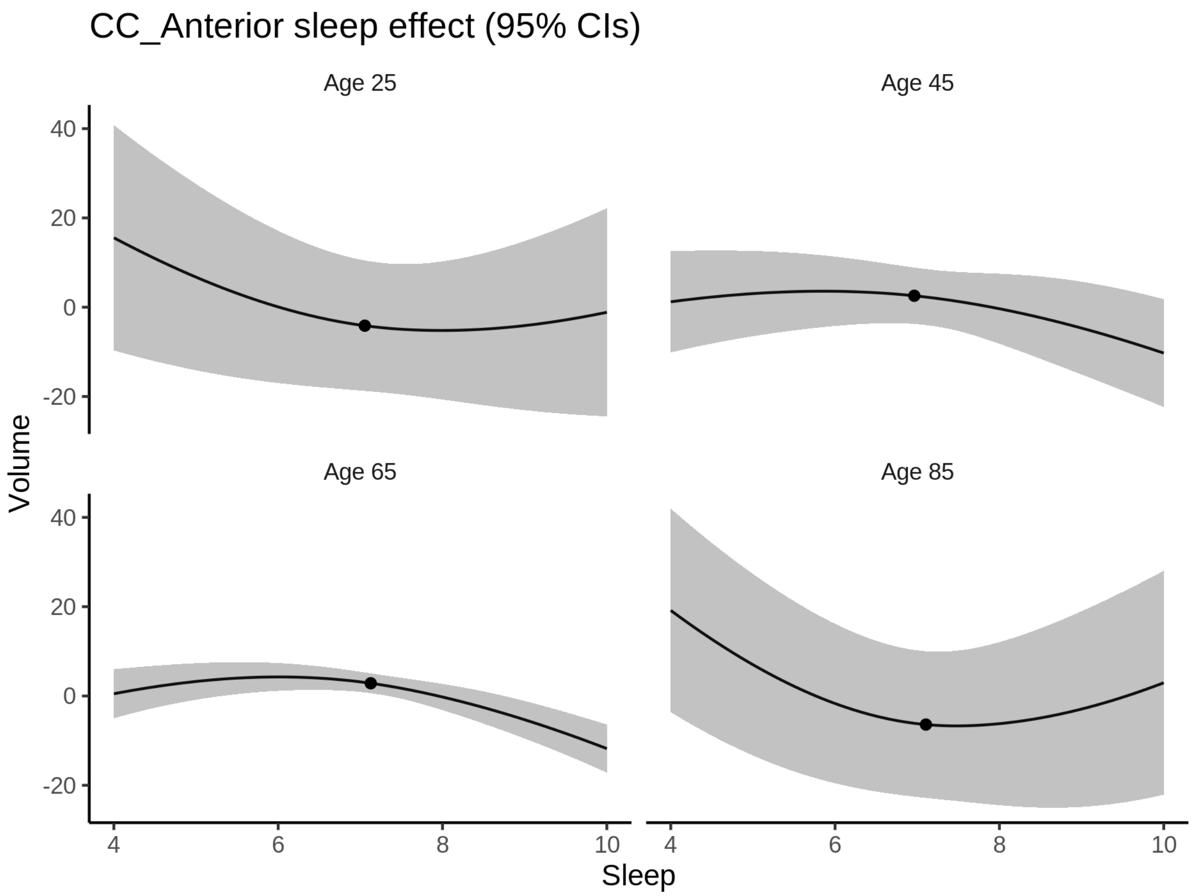


**Deviation from sleep associated with maximal volume**

Model with only main effect of sleep was chosen, so we show it for all ages at once. Maximum volume is attained at 4 hours of sleep. The percentage values in the plot are calculated as follows: The maximum at 100 % refers to a person at an arbitrary age with a sleep duration associated with maximum volume. For a female, this volume is 962 and for a male it is 921. The other percentage values show how large the expected volume is for someone with other sleep durations. For example, 99 % implies a 1 % reduction.

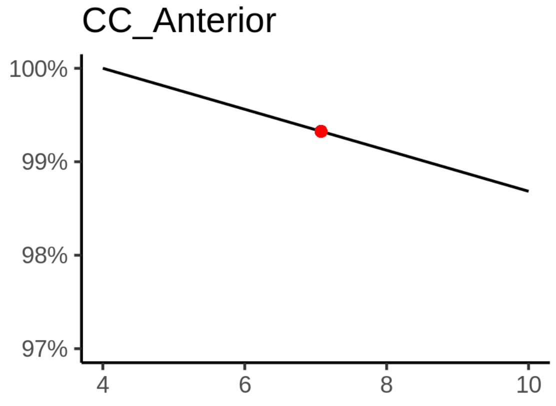

**Comparison of mean sleep and sleep associated with maximum volume**

A 95 % confidence interval for the sleep associated with maximum volume is [4, 4].

The plot below compares average sleep to the sleep associated with maximum volume.

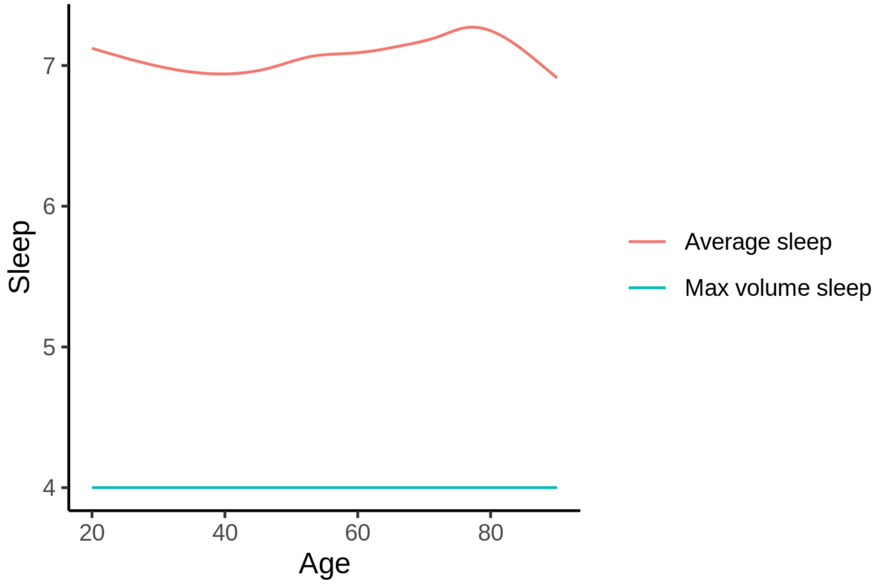


The next plot shows the difference between average sleep and sleep associated with maximum volume. The shaded region is a 95 % confidence interval.

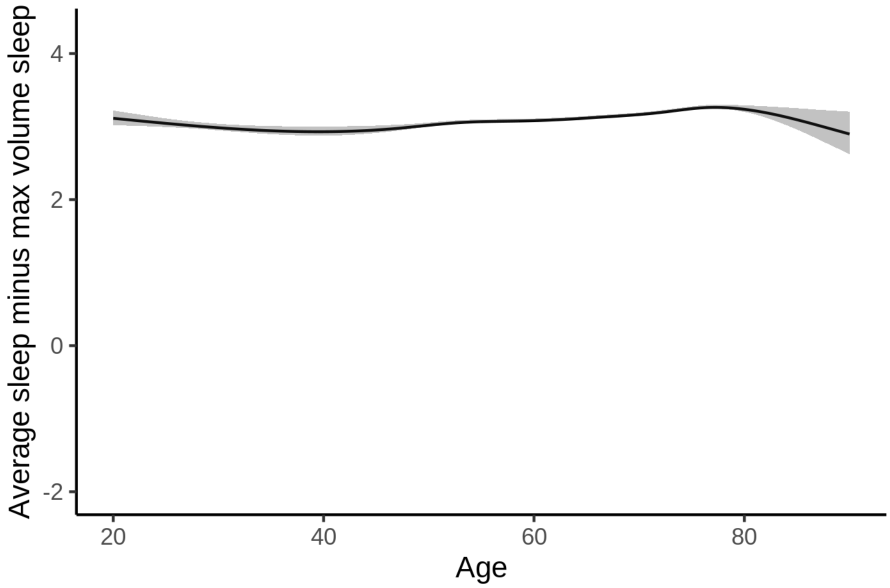


The next plot shows the probability that the sleep duration associated with maximum volume is longer than the average sleep duration, as a function of age. Probility below .05 can be interpreted as evidence that the sleep associated with maximum volume is shorter than the mean sleep, and probability above .95 can be interpreted the opposite way.

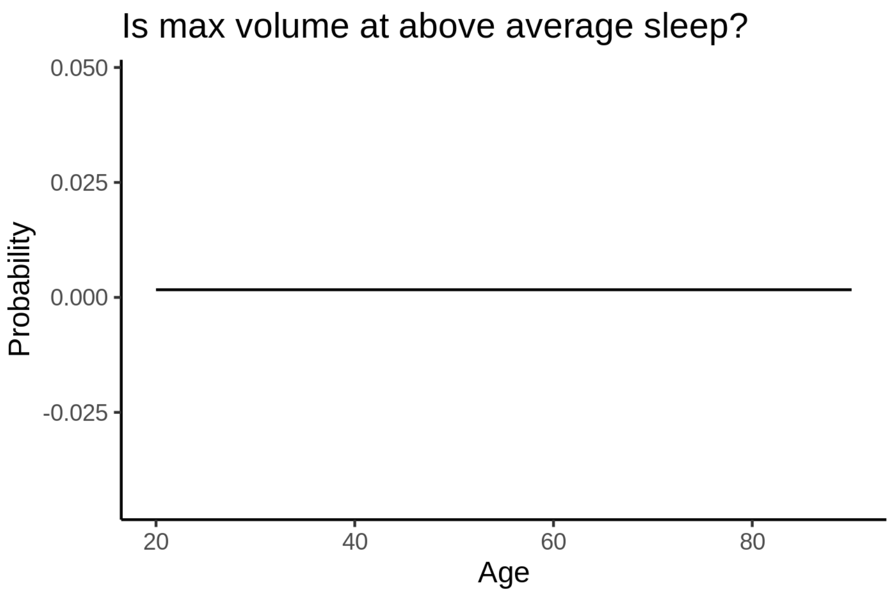


##### Controling for covariates

Below is the output for a model in which we only include data with income and education.

```
##
## Family: gaussian
## Link function: identity
##
## Formula:
## value ~ sex + site + s(age_z, k = 10, bs = "cr") + s(sleep_z,
##     k = 5, bs = "cr") + icv
## <environment: 0x55cf3b9d6820>
##
## Parametric coefficients:
##               Estimate Std. Error t value Pr(>|t|)
## (Intercept)   1070.002     19.035  56.214  < 2e-16 ***
## sexmale        -43.547      2.102 -20.719  < 2e-16 ***
## siteousAvanto -202.615     23.314  -8.691  < 2e-16 ***
## siteousPrisma -143.408     23.506  -6.101 1.07e-09 ***
## siteousSkyra  -147.718     20.647  -7.154 8.58e-13 ***
## siteUKB       -109.953     18.998  -5.788 7.21e-09 ***
## siteUOXF      -149.823     21.405  -7.000 2.62e-12 ***
## icv             84.900      1.080  78.609  < 2e-16 ***
## ---
## Signif. codes:  0 ‘***’ 0.001 ‘**’ 0.01 ‘*’ 0.05 ‘.’ 0.1 ‘ ’ 1
##
## Approximate significance of smooth terms:
##              edf Ref.df     F p-value
## s(age_z)   4.989  4.989 83.70  <2e-16 ***
## s(sleep_z) 1.705  1.705  2.23  0.0679 .
## ---
## Signif. codes:  0 ‘***’ 0.001 ‘**’ 0.01 ‘*’ 0.05 ‘.’ 0.1 ‘ ’ 1
##
```

```
## R-sq.(adj) =  0.211
## lmer.REML = 3.9487e+05  Scale est. = 2119.2    n = 31188
```

Below is the output for a model in which we control for the main effects of income and education.

```
##
## Family: gaussian
## Link function: identity
##
## Formula:
## value ~ sex + site + s(age_z, k = 10, bs = "cr") + s(sleep_z,
##     k = 5, bs = "cr") + icv + income_scaled + education_scaled
## <environment: 0x55cf3b9d6820>
##
## Parametric coefficients:
##                   Estimate Std. Error t value Pr(>|t|)
## (Intercept)      1069.8103    19.1589  55.839  < 2e-16 ***
## sexmale           -43.5166     2.1080 -20.643  < 2e-16 ***
## siteousAvanto    -202.7119    23.3200  -8.693  < 2e-16 ***
## siteousPrisma    -143.4437    23.5081  -6.102 1.06e-09 ***
## siteousSkyra     -147.8117    20.6533  -7.157 8.44e-13 ***
## siteUKB          -110.0777    19.0100  -5.791 7.08e-09 ***
## siteUOXF         -149.8168    21.4174  -6.995 2.70e-12 ***
## icv                84.8901     1.0894  77.922  < 2e-16 ***
## income_scaled      -0.2847     2.5045  -0.114    0.909
## education_scaled    0.5576     2.9587   0.188    0.851
## ---
## Signif. codes:  0 '***' 0.001 '**' 0.01 '*' 0.05 '.' 0.1 ' ' 1
##
## Approximate significance of smooth terms:
##              edf Ref.df      F p-value
## s(age_z)   4.992  4.992 76.609  <2e-16 ***
## s(sleep_z) 1.702  1.702  2.239  0.0674 .
## ---
## Signif. codes:  0 '***' 0.001 '**' 0.01 '*' 0.05 '.' 0.1 ' ' 1
##
## R-sq.(adj) =  0.211
## lmer.REML = 3.9486e+05  Scale est. = 2119.2    n = 31188
```

We also included interaction effects between sleep duration and education and income, in another model. The output is shown below, and the interaction terms are `income_scaled:sleep_z` and `education_scaled:sleep_z`.

```
##
## Family: gaussian
## Link function: identity
##
## Formula:
## value ~ sex + site + s(age_z, k = 10, bs = "cr") + s(sleep_z,
##     k = 5, bs = "cr") + icv + income_scaled + education_scaled +
##     income_scaled:sleep_z + education_scaled:sleep_z
## <environment: 0x55cf3b9d6820>
##
## Parametric coefficients:
##                           Estimate Std. Error t value Pr(>|t|)
## (Intercept)              1069.7614    19.1603  55.832  < 2e-16 ***
```

```
## sexmale                    -43.4054     2.1095 -20.576  < 2e-16 ***
## siteousAvanto            -202.5313    23.3209  -8.685  < 2e-16 ***
## siteousPrisma            -142.9874    23.5108  -6.082 1.20e-09 ***
## siteousSkyra             -147.6296    20.6540  -7.148 9.01e-13 ***
## siteUKB                  -110.0329    19.0123  -5.787 7.21e-09 ***
## siteUOXF                 -149.8448    21.4237  -6.994 2.72e-12 ***
## icv                        84.8782     1.0895  77.904  < 2e-16 ***
## income_scaled              -0.3923     2.5056  -0.157    0.876
## education_scaled            0.6293     2.9591   0.213    0.832
## income_scaled:sleep_z       3.2233     2.4816   1.299    0.194
## education_scaled:sleep_z   -2.5040     2.8678  -0.873    0.383
## ---
## Signif. codes:  0 '***' 0.001 '**' 0.01 '*' 0.05 '.' 0.1 ' ' 1
##
## Approximate significance of smooth terms:
##              edf Ref.df      F p-value
## s(age_z)   4.997  4.997 76.710  <2e-16 ***
## s(sleep_z) 1.645  1.645  0.355   0.543
## ---
## Signif. codes:  0 '***' 0.001 '**' 0.01 '*' 0.05 '.' 0.1 ' ' 1
##
## R-sq.(adj) =  0.211
## lmer.REML = 3.9485e+05  Scale est. = 2119.3    n = 31188
```

We did the same controling for BMI. Below is the model with no covariates but only keeping data with BMI.

```
##
## Family: gaussian
## Link function: identity
##
## Formula:
## value ~ sex + site + s(age_z, k = 10, bs = "cr") + s(sleep_z,
##     k = 5, bs = "cr") + icv
## <environment: 0x55cf3ae0d9c8>
##
## Parametric coefficients:
##               Estimate Std. Error t value Pr(>|t|)
## (Intercept)    876.935     13.616  64.404  < 2e-16 ***
## sexmale        -43.188      2.035 -21.228  < 2e-16 ***
## siteousPrisma   42.892     17.912   2.395 0.016642 *
## siteousSkyra    46.640     11.064   4.215 2.50e-05 ***
## siteUCAM         7.295     14.319   0.509 0.610432
## siteUKB         83.003     13.717   6.051 1.45e-09 ***
## siteUmU         56.936     16.007   3.557 0.000376 ***
## icv             84.890      1.048  80.972  < 2e-16 ***
## ---
## Signif. codes:  0 '***' 0.001 '**' 0.01 '*' 0.05 '.' 0.1 ' ' 1
##
## Approximate significance of smooth terms:
##              edf Ref.df     F p-value
## s(age_z)   5.022  5.022 98.70  <2e-16 ***
## s(sleep_z) 1.304  1.304  4.15  0.0532 .
## ---
## Signif. codes:  0 '***' 0.001 '**' 0.01 '*' 0.05 '.' 0.1 ' ' 1
##
```

```
## R-sq.(adj) =  0.214
## lmer.REML = 4.2272e+05  Scale est. = 1979.2    n = 33443
```

Below is the model output with main effect.

```
##
## Family: gaussian
## Link function: identity
##
## Formula:
## value ~ sex + site + s(age_z, k = 10, bs = "cr") + s(sleep_z,
##     k = 5, bs = "cr") + icv + bmi
## <environment: 0x55cf3ae0d9c8>
##
## Parametric coefficients:
##               Estimate Std. Error t value Pr(>|t|)
## (Intercept)   926.4403    14.4914  63.930  < 2e-16 ***
## sexmale       -41.6206     2.0374 -20.428  < 2e-16 ***
## siteousPrisma  44.9093    17.8959   2.509 0.012096 *
## siteousSkyra   46.9702    11.0616   4.246 2.18e-05 ***
## siteUCAM        8.5601    14.3090   0.598 0.549690
## siteUKB        84.6479    13.7107   6.174 6.74e-10 ***
## siteUmU        59.1880    15.9952   3.700 0.000216 ***
## icv            85.1817     1.0469  81.364  < 2e-16 ***
## bmi            -1.9676     0.1977  -9.954  < 2e-16 ***
## ---
## Signif. codes:  0 ‘***’ 0.001 ‘**’ 0.01 ‘*’ 0.05 ‘.’ 0.1 ‘ ’ 1
##
## Approximate significance of smooth terms:
##              edf Ref.df      F p-value
## s(age_z)   5.083  5.083 99.024  <2e-16 ***
## s(sleep_z) 1.000  1.000  6.585  0.0103 *
## ---
## Signif. codes:  0 ‘***’ 0.001 ‘**’ 0.01 ‘*’ 0.05 ‘.’ 0.1 ‘ ’ 1
##
## R-sq.(adj) =  0.217
## lmer.REML = 4.2262e+05  Scale est. = 1978.6    n = 33443
```

Next is the model with BMI-sleep interaction.

```
##
## Family: gaussian
## Link function: identity
##
## Formula:
## value ~ sex + site + s(age_z, k = 10, bs = "cr") + s(sleep_z,
##     k = 5, bs = "cr") + icv + bmi + bmi:sleep_z
## <environment: 0x55cf3ae0d9c8>
##
## Parametric coefficients:
##               Estimate Std. Error t value Pr(>|t|)
## (Intercept)   927.0218    14.4979  63.942  < 2e-16 ***
## sexmale       -41.6312     2.0374 -20.434  < 2e-16 ***
## siteousPrisma  44.8992    17.8957   2.509 0.012114 *
## siteousSkyra   46.9075    11.0616   4.241 2.24e-05 ***
## siteUCAM        8.3327    14.3100   0.582 0.560371
```

```
## siteUKB        84.4259    13.7117   6.157 7.49e-10 ***
## siteUmU        58.9868    15.9958   3.688 0.000227 ***
## icv            85.1723     1.0469  81.354  < 2e-16 ***
## bmi            -1.9765     0.1978  -9.994  < 2e-16 ***
## bmi:sleep_z    -0.2503     0.1872  -1.337 0.181133
## ---
## Signif. codes:  0 '***' 0.001 '**' 0.01 '*' 0.05 '.' 0.1 ' ' 1
##
## Approximate significance of smooth terms:
##              edf Ref.df      F p-value
## s(age_z)   5.084  5.084 98.807  <2e-16 ***
## s(sleep_z) 1.000  1.000  0.793   0.373
## ---
## Signif. codes:  0 '***' 0.001 '**' 0.01 '*' 0.05 '.' 0.1 ' ' 1
##
## R-sq.(adj) =  0.217
## lmer.REML = 4.2262e+05  Scale est. = 1978.5    n = 33443
```

We did the same controling for depression. Below is the model with no covariates but only keeping data with depression.

```
##
## Family: gaussian
## Link function: identity
##
## Formula:
## value ~ sex + site + s(age_z, k = 10, bs = "cr") + s(sleep_z,
##     k = 5, bs = "cr") + icv
## <environment: 0x55cf39cd3f08>
##
## Parametric coefficients:
##               Estimate Std. Error t value Pr(>|t|)
## (Intercept)   1012.170      9.677 104.595  < 2e-16 ***
## sexmale        -43.409      2.047 -21.202  < 2e-16 ***
## siteousAvanto -129.226     21.545  -5.998 2.02e-09 ***
## siteousPrisma  -85.649     18.649  -4.593 4.39e-06 ***
## siteousSkyra   -78.756     14.531  -5.420 6.01e-08 ***
## siteUCAM      -129.567     13.222  -9.799  < 2e-16 ***
## siteUKB        -52.453      9.568  -5.482 4.24e-08 ***
## siteUmU        -79.068     12.981  -6.091 1.13e-09 ***
## icv             84.672      1.054  80.366  < 2e-16 ***
## ---
## Signif. codes:  0 '***' 0.001 '**' 0.01 '*' 0.05 '.' 0.1 ' ' 1
##
## Approximate significance of smooth terms:
##              edf Ref.df      F p-value
## s(age_z)   5.194  5.194 89.045  <2e-16 ***
## s(sleep_z) 1.576  1.576  4.961  0.0369 *
## ---
## Signif. codes:  0 '***' 0.001 '**' 0.01 '*' 0.05 '.' 0.1 ' ' 1
##
## R-sq.(adj) =  0.212
## lmer.REML = 4.2149e+05  Scale est. = 1926.4    n = 33355
```

Below is the model output with main effect.

```
##
## Family: gaussian
## Link function: identity
##
## Formula:
## value ~ sex + site + s(age_z, k = 10, bs = "cr") + s(sleep_z,
##     k = 5, bs = "cr") + icv + depression
## <environment: 0x55cf39cd3f08>
##
## Parametric coefficients:
##                Estimate Std. Error t value Pr(>|t|)
## (Intercept)    1011.093      9.720 104.023  < 2e-16 ***
## sexmale         -43.297      2.049 -21.126  < 2e-16 ***
## siteousAvanto -129.763     21.547  -6.022 1.74e-09 ***
## siteousPrisma  -86.124     18.651  -4.618 3.90e-06 ***
## siteousSkyra   -79.371     14.537  -5.460 4.80e-08 ***
## siteUCAM      -129.803     13.223  -9.816  < 2e-16 ***
## siteUKB        -52.017      9.576  -5.432 5.61e-08 ***
## siteUmU        -82.351     13.259  -6.211 5.32e-10 ***
## icv             84.652      1.054  80.336  < 2e-16 ***
## depression       8.880      7.295   1.217    0.224
## ---
## Signif. codes:  0 ‘***’ 0.001 ‘**’ 0.01 ‘*’ 0.05 ‘.’ 0.1 ‘ ’ 1
##
## Approximate significance of smooth terms:
##              edf Ref.df      F p-value
## s(age_z)   5.177  5.177 86.298  <2e-16 ***
## s(sleep_z) 1.757  1.757  4.552  0.0438 *
## ---
## Signif. codes:  0 ‘***’ 0.001 ‘**’ 0.01 ‘*’ 0.05 ‘.’ 0.1 ‘ ’ 1
##
## R-sq.(adj) =  0.212
## lmer.REML = 4.2148e+05  Scale est. = 1926.2    n = 33355
```

Next is the model with depression-sleep interaction.

```
##
## Family: gaussian
## Link function: identity
##
## Formula:
## value ~ sex + site + s(age_z, k = 10, bs = "cr") + s(sleep_z,
##     k = 5, bs = "cr") + icv + depression + depression:sleep_z
## <environment: 0x55cf39cd3f08>
##
## Parametric coefficients:
##                     Estimate Std. Error t value Pr(>|t|)
## (Intercept)         1011.092      9.720 104.023  < 2e-16 ***
## sexmale              -43.330      2.050 -21.139  < 2e-16 ***
## siteousAvanto       -130.015     21.549  -6.033 1.62e-09 ***
## siteousPrisma        -86.371     18.653  -4.631 3.66e-06 ***
## siteousSkyra         -79.612     14.539  -5.476 4.39e-08 ***
## siteUCAM            -130.047     13.226  -9.833  < 2e-16 ***
## siteUKB              -52.003      9.576  -5.431 5.66e-08 ***
## siteUmU              -80.809     13.369  -6.045 1.51e-09 ***
```

```
## icv                   84.655      1.054  80.339  < 2e-16 ***
## depression             8.506      7.307   1.164    0.244
## depression:sleep_z    -4.795      5.337  -0.898    0.369
## ---
## Signif. codes:  0 '***' 0.001 '**' 0.01 '*' 0.05 '.' 0.1 ' ' 1
##
## Approximate significance of smooth terms:
##              edf Ref.df      F p-value
## s(age_z)   5.169  5.169 86.463  <2e-16 ***
## s(sleep_z) 1.748  1.748  2.458   0.188
## ---
## Signif. codes:  0 '***' 0.001 '**' 0.01 '*' 0.05 '.' 0.1 ' ' 1
##
## R-sq.(adj) =  0.212
## lmer.REML = 4.2147e+05  Scale est. = 1926.3    n = 33355
```

The plot below shows the sleep-volume curve for the original model and for the model with main effects of SES.

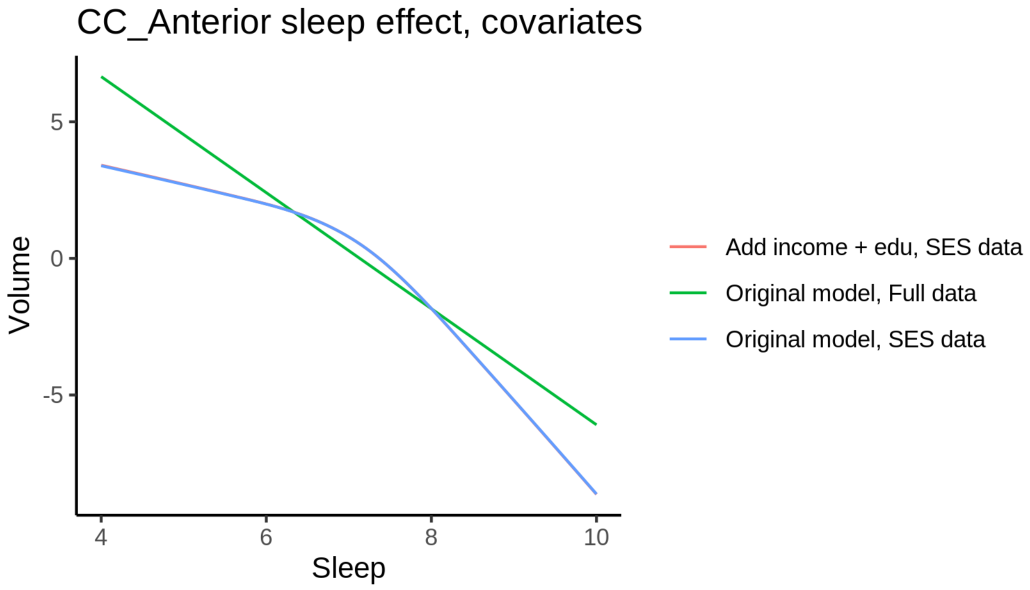


The plot below shows the sleep-volume curve for the original model and for the model with main effects of BMI.

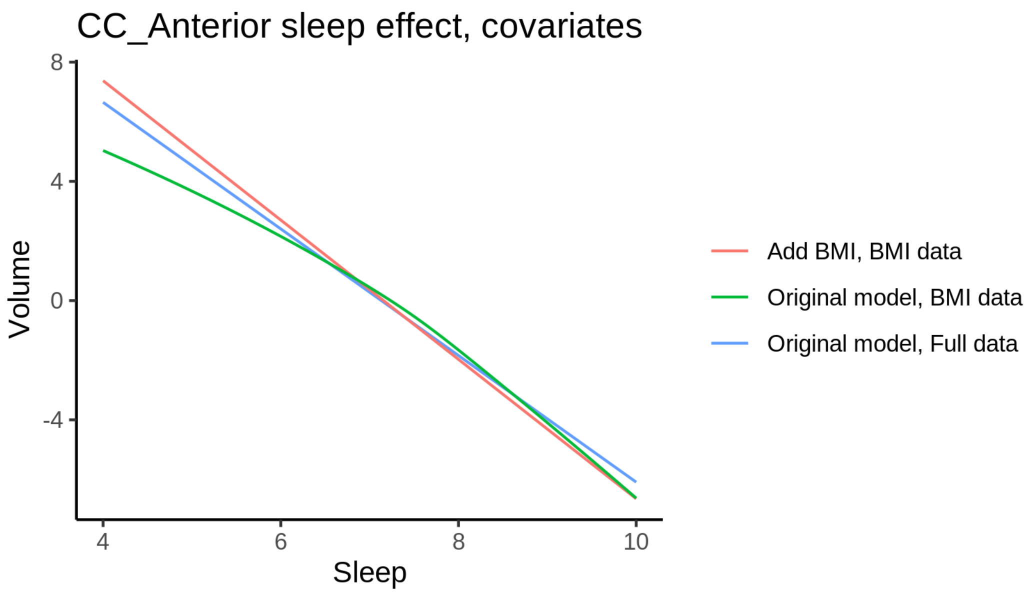


The plot below shows the sleep-volume curve for the original model and for the model with main effects of depression.

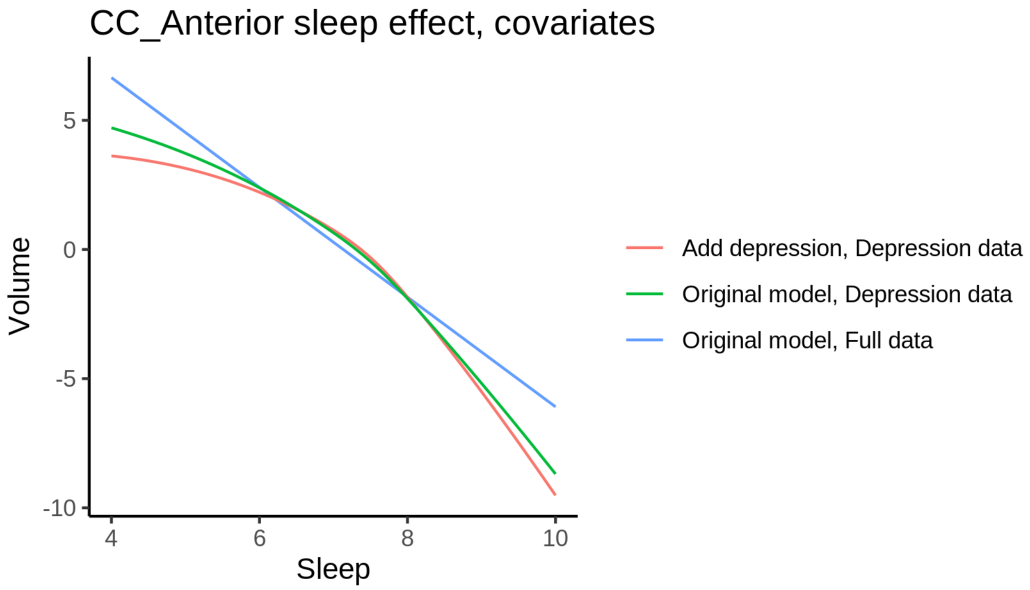


### CC_Central

#### Descriptive statistics

| Study | Observations | Unique IDs | Mean age | Age range |
|---|---|---|---|---|
| HCP | 974 | 974 | 28.8 | 22 - 37 |
| MPIB | 677 | 391 | 63.1 | 24 - 83 |
| UB | 113 | 39 | 70.9 | 64 - 81 |
| UCAM | 883 | 632 | 55.1 | 20 - 88 |

| Study | Observations | Unique IDs | Mean age | Age range |
|---|---|---|---|---|
| UiO | 1475 | 803 | 49.4 | 20 - 89 |
| UKB | 45975 | 43135 | 64.5 | 45 - 83 |
| UmU | 423 | 284 | 62.3 | 25 - 85 |
| UOXF | 769 | 769 | 69.8 | 60 - 85 |

##### Spaghetti plot

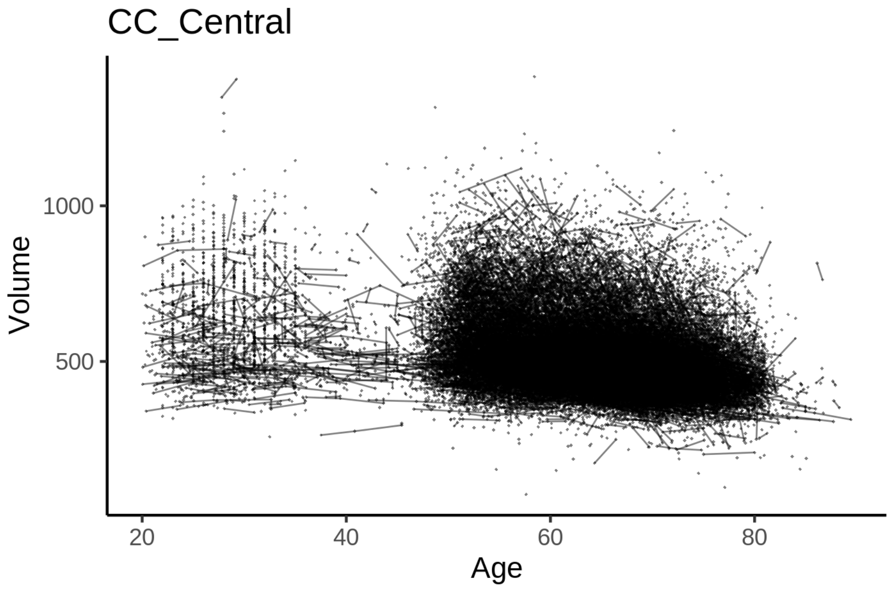


##### Model outputs

Model without sleep term

```
##
## Family: gaussian
## Link function: identity
##
## Formula:
## value ~ sex + site + icv + s(age_z, k = 10, bs = "cr")
## <environment: 0x55cffc0d4a88>
##
## Parametric coefficients:
##                 Estimate Std. Error t value Pr(>|t|)
## (Intercept)     591.0919     6.1098  96.745  < 2e-16 ***
## sexmale         -18.7683     1.2655 -14.831  < 2e-16 ***
## siteMPIB        -66.4966     7.8605  -8.460  < 2e-16 ***
## siteousAvanto -149.2461     6.4162 -23.261  < 2e-16 ***
## siteousPrisma  -68.1815    10.9081  -6.251 4.12e-10 ***
## siteousSkyra  -156.4902     5.9914 -26.119  < 2e-16 ***
## siteUB        -115.3724    18.4347  -6.258 3.92e-10 ***
## siteUCAM       -84.1117     6.8109 -12.350  < 2e-16 ***
## siteUKB        -58.5092     6.2627  -9.343  < 2e-16 ***
## siteUmU        -79.4964     8.7623  -9.073  < 2e-16 ***
```

```
## siteUOXF       -45.4166     7.4483  -6.098 1.08e-09 ***
## icv             20.4333     0.6366  32.100  < 2e-16 ***
## ---
## Signif. codes:  0 ‘***’ 0.001 ‘**’ 0.01 ‘*’ 0.05 ‘.’ 0.1 ‘ ’ 1
##
## Approximate significance of smooth terms:
##            edf Ref.df     F p-value
## s(age_z) 5.612  5.612 812.1  <2e-16 ***
## ---
## Signif. codes:  0 ‘***’ 0.001 ‘**’ 0.01 ‘*’ 0.05 ‘.’ 0.1 ‘ ’ 1
##
## R-sq.(adj) =  0.146
## lmer.REML = 6.2202e+05  Scale est. = 1202.5    n = 51289
```

Model with only main effects of age and sleep

```
##
## Family: gaussian
## Link function: identity
##
## Formula:
## value ~ sex + site + icv + s(age_z, k = 10, bs = "cr") + s(sleep_z,
##     k = 5, bs = "cr")
## <environment: 0x55cffc0d4a88>
##
## Parametric coefficients:
##                 Estimate Std. Error t value Pr(>|t|)
## (Intercept)    589.9592     6.1122  96.522  < 2e-16 ***
## sexmale        -18.8082     1.2653 -14.865  < 2e-16 ***
## siteMPIB       -65.2609     7.8634  -8.299  < 2e-16 ***
## siteousAvanto -148.6462     6.4160 -23.168  < 2e-16 ***
## siteousPrisma  -67.7805    10.9066  -6.215 5.18e-10 ***
## siteousSkyra  -155.8716     5.9912 -26.017  < 2e-16 ***
## siteUB        -115.0301    18.4305  -6.241 4.37e-10 ***
## siteUCAM       -83.5356     6.8098 -12.267  < 2e-16 ***
## siteUKB        -57.3055     6.2655  -9.146  < 2e-16 ***
## siteUmU        -76.9895     8.7800  -8.769  < 2e-16 ***
## siteUOXF       -44.8365     7.4460  -6.022 1.74e-09 ***
## icv             20.5131     0.6367  32.216  < 2e-16 ***
## ---
## Signif. codes:  0 ‘***’ 0.001 ‘**’ 0.01 ‘*’ 0.05 ‘.’ 0.1 ‘ ’ 1
##
## Approximate significance of smooth terms:
##              edf Ref.df      F  p-value
## s(age_z)   5.586  5.586 807.61  < 2e-16 ***
## s(sleep_z) 1.000  1.000  17.08 3.59e-05 ***
## ---
## Signif. codes:  0 ‘***’ 0.001 ‘**’ 0.01 ‘*’ 0.05 ‘.’ 0.1 ‘ ’ 1
##
## R-sq.(adj) =  0.146
## lmer.REML = 6.2201e+05  Scale est. = 1202.5    n = 51289
```

Model with full interaction between age and sleep

```
##
## Family: gaussian
```

```
## Link function: identity
##
## Formula:
## value ~ sex + site + icv + t2(age_z, sleep_z, k = c(10, 4), bs = "cr")
## <environment: 0x55cffc0d4a88>
##
## Parametric coefficients:
##                 Estimate Std. Error t value Pr(>|t|)
## (Intercept)     589.8465     6.1378  96.100  < 2e-16 ***
## sexmale         -19.0312     1.2677 -15.013  < 2e-16 ***
## siteMPIB        -64.8241     7.9046  -8.201 2.44e-16 ***
## siteousAvanto -148.7267     6.4490 -23.062  < 2e-16 ***
## siteousPrisma  -67.8894    10.9271  -6.213 5.24e-10 ***
## siteousSkyra  -155.7878     6.0296 -25.837  < 2e-16 ***
## siteUB        -114.3415    18.4415  -6.200 5.68e-10 ***
## siteUCAM       -83.0414     6.8478 -12.127  < 2e-16 ***
## siteUKB        -57.0914     6.2912  -9.075  < 2e-16 ***
## siteUmU        -77.1219     8.8082  -8.756  < 2e-16 ***
## siteUOXF       -44.1593     7.4750  -5.908 3.49e-09 ***
## icv             20.5251     0.6368  32.231  < 2e-16 ***
## ---
## Signif. codes:  0 ‘***’ 0.001 ‘**’ 0.01 ‘*’ 0.05 ‘.’ 0.1 ‘ ’ 1
##
## Approximate significance of smooth terms:
##                      edf Ref.df     F p-value
## t2(age_z,sleep_z) 9.226  9.226 27.14  <2e-16 ***
## ---
## Signif. codes:  0 ‘***’ 0.001 ‘**’ 0.01 ‘*’ 0.05 ‘.’ 0.1 ‘ ’ 1
##
## R-sq.(adj) =  0.146
## lmer.REML = 6.22e+05  Scale est. = 1202.4    n = 51289
```

#### Model comparison

`mod_no_sleep` refers to model without sleep term, `mod_no_interaction` refers to model with only main effect of sleep, and `mod_full` refers to model with a full interaction between age and sleep. This is a nested model comparison, and the p-value at a given line refers to comparing the model at the line to the model on the line above. Hence, significance implies that the more complicated model is supported on statistical grounds.

To be even more specific, the p-value on the second row tests whether there is an association between sleep and volume. The p-value on the third row tests whether this association depends on age.

```
## Data: NULL
## Models:
## mod_list$mod_no_sleep$mer: NULL
## mod_list$mod_no_interaction$mer: NULL
## mod_list$mod_full$mer: NULL
##                                 npar    AIC    BIC  logLik deviance   Chisq Df Pr(>Chisq)
## mod_list$mod_no_sleep$mer         16 622056 622198 -311012   622024
## mod_list$mod_no_interaction$mer   18 622043 622203 -311004   622007 17.0736  2  0.0001961 ***
## mod_list$mod_full$mer             20 622043 622220 -311002   622003  4.1849  2  0.1233863
## ---
## Signif. codes:  0 ‘***’ 0.001 ‘**’ 0.01 ‘*’ 0.05 ‘.’ 0.1 ‘ ’ 1
```

We chose the model based on the likelihood ratio test with 5 % significance level, which was `mod_no_interaction`.

#### Lifespan brain trajectory

The trajectory shown is from the chosen model `mod_no_interaction`.

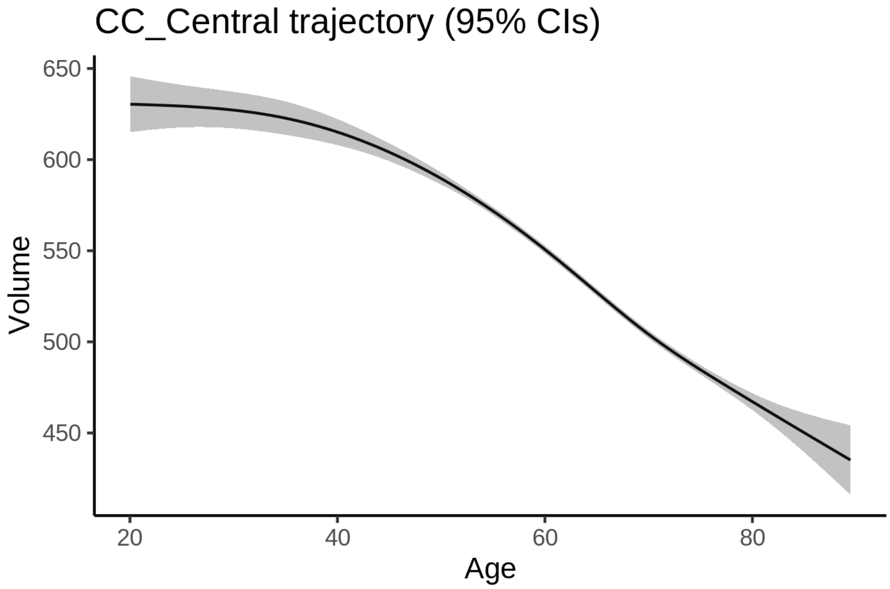


#### Effect of sleep

The chosen model only included the main effect of sleep, and hence the effect does not vary with age. The black dot shows the average sleep duration across all ages in the sample.

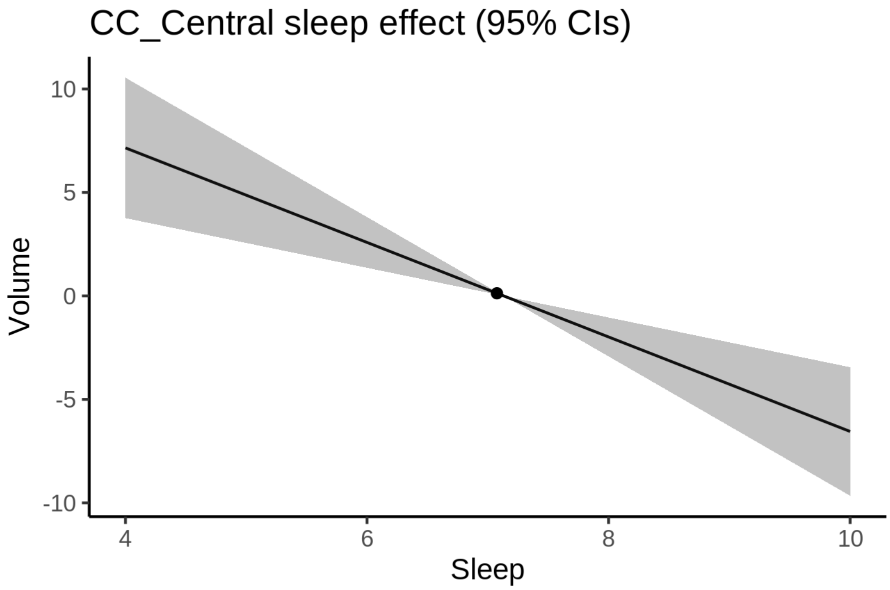


We also show the full interaction model for completeness, although it was not selected.

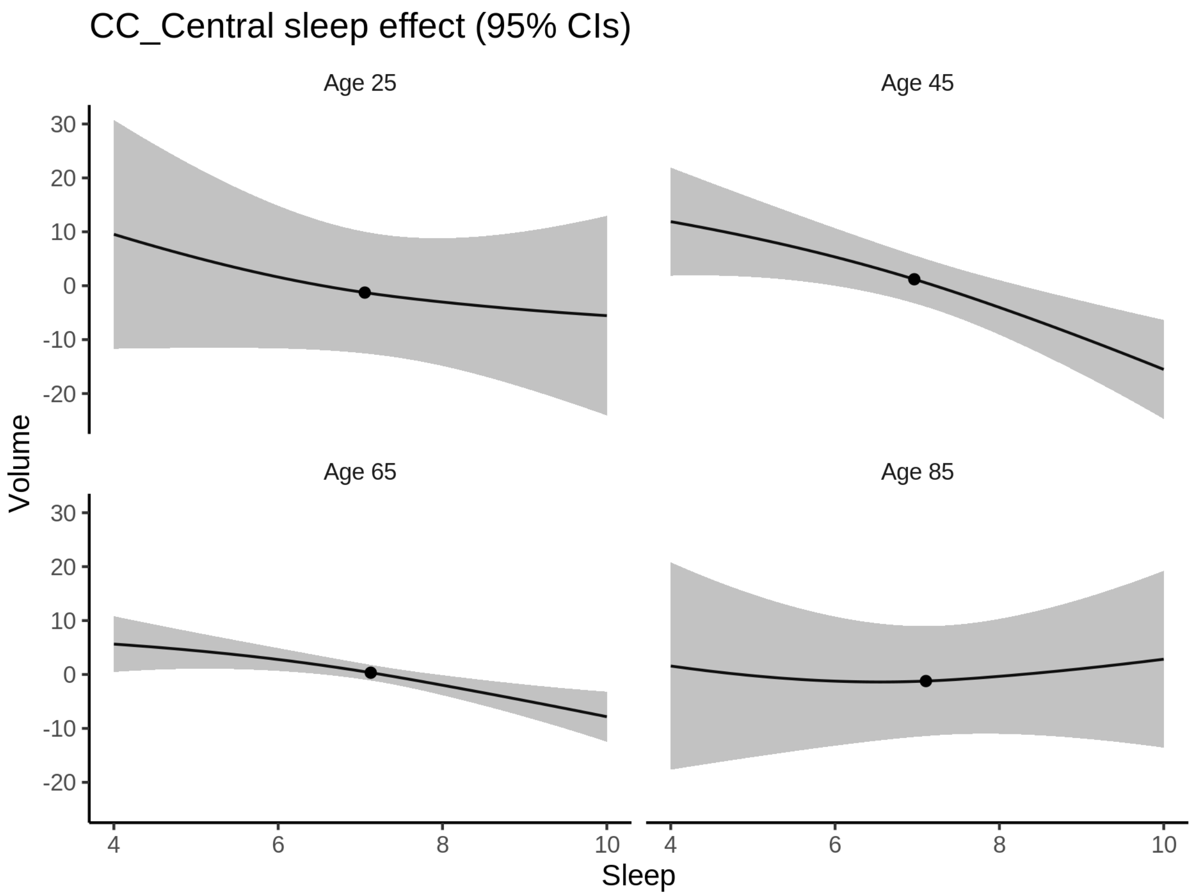


**Deviation from sleep associated with maximal volume**

Model with only main effect of sleep was chosen, so we show it for all ages at once. Maximum volume is attained at 4 hours of sleep. The percentage values in the plot are calculated as follows: The maximum at 100 % refers to a person at an arbitrary age with a sleep duration associated with maximum volume. For a female, this volume is 555 and for a male it is 536. The other percentage values show how large the expected volume is for someone with other sleep durations. For example, 99 % implies a 1 % reduction.

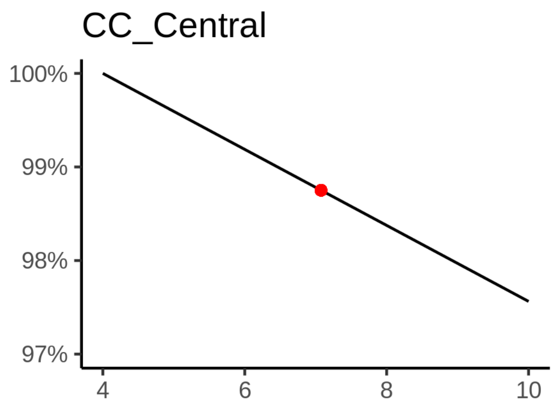

**Comparison of mean sleep and sleep associated with maximum volume**

A 95 % confidence interval for the sleep associated with maximum volume is [4, 4].

The plot below compares average sleep to the sleep associated with maximum volume.

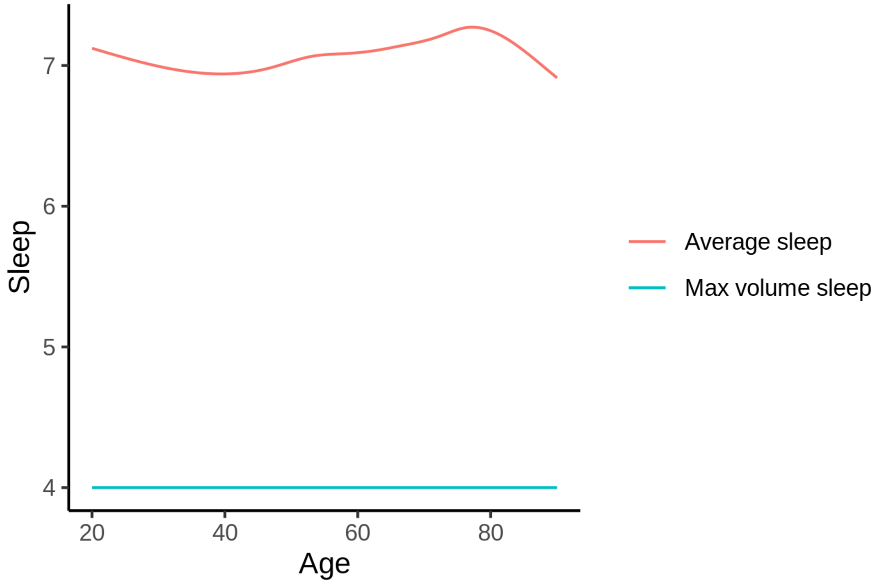


The next plot shows the difference between average sleep and sleep associated with maximum volume. The shaded region is a 95 % confidence interval.

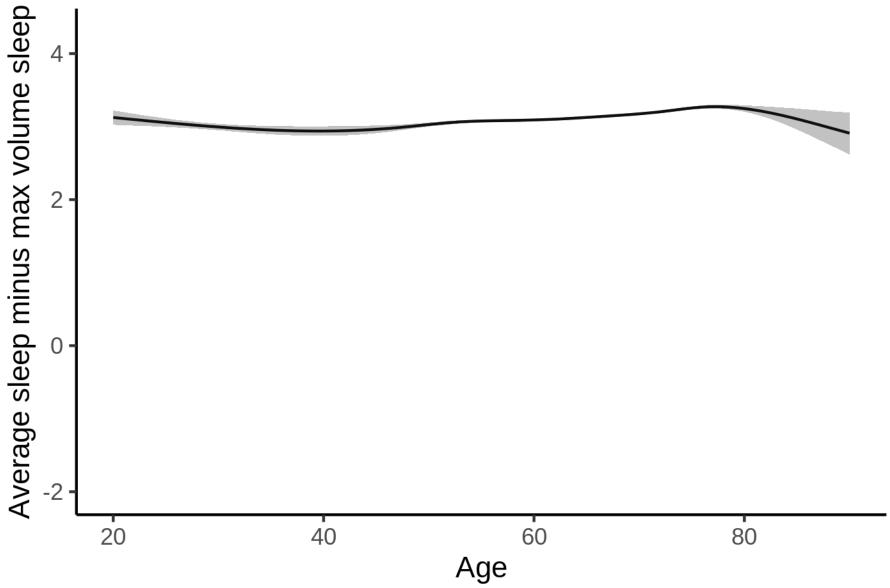


The next plot shows the probability that the sleep duration associated with maximum volume is longer than the average sleep duration, as a function of age. Probility below .05 can be interpreted as evidence that the sleep associated with maximum volume is shorter than the mean sleep, and probability above .95 can be interpreted the opposite way.

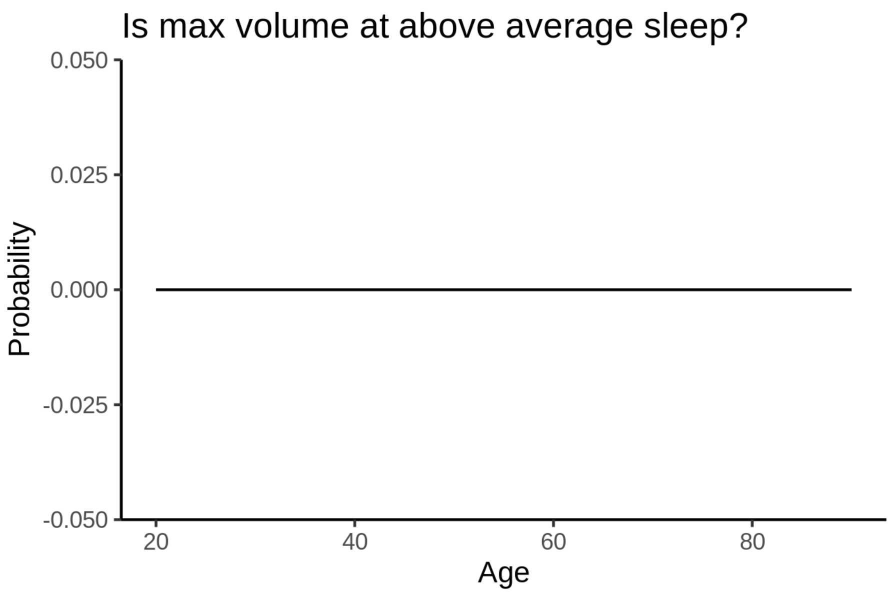


##### Controling for covariates

Below is the output for a model in which we only include data with income and education.

```
##
## Family: gaussian
## Link function: identity
##
## Formula:
## value ~ sex + site + s(age_z, k = 10, bs = "cr") + s(sleep_z,
##     k = 5, bs = "cr") + icv
## <environment: 0x55cf401af668>
##
## Parametric coefficients:
##               Estimate Std. Error t value Pr(>|t|)
## (Intercept)   516.2376    14.4730  35.669  < 2e-16 ***
## sexmale       -18.7158     1.6012 -11.689  < 2e-16 ***
## siteousAvanto -88.3558    17.9518  -4.922 8.62e-07 ***
## siteousPrisma  12.4867    17.9160   0.697    0.486
## siteousSkyra  -89.0267    15.7033  -5.669 1.45e-08 ***
## siteUKB        14.0811    14.4453   0.975    0.330
## siteUOXF       22.1866    16.2830   1.363    0.173
## icv            21.1422     0.8228  25.697  < 2e-16 ***
## ---
## Signif. codes:  0 ‘***’ 0.001 ‘**’ 0.01 ‘*’ 0.05 ‘.’ 0.1 ‘ ’ 1
##
## Approximate significance of smooth terms:
##              edf Ref.df       F p-value
## s(age_z)   4.899  4.899 548.764  <2e-16 ***
## s(sleep_z) 1.000  1.000   5.131  0.0235 *
## ---
## Signif. codes:  0 ‘***’ 0.001 ‘**’ 0.01 ‘*’ 0.05 ‘.’ 0.1 ‘ ’ 1
##
```

```
## R-sq.(adj) =  0.114
## lmer.REML = 3.7828e+05  Scale est. = 1410.2    n = 31193
```

Below is the output for a model in which we control for the main effects of income and education.

```
##
## Family: gaussian
## Link function: identity
##
## Formula:
## value ~ sex + site + s(age_z, k = 10, bs = "cr") + s(sleep_z,
##     k = 5, bs = "cr") + icv + income_scaled + education_scaled
## <environment: 0x55cf401af668>
##
## Parametric coefficients:
##                  Estimate Std. Error t value Pr(>|t|)
## (Intercept)       521.426     14.568  35.792  < 2e-16 ***
## sexmale           -18.767      1.606 -11.688  < 2e-16 ***
## siteousAvanto     -88.079     17.956  -4.905 9.37e-07 ***
## siteousPrisma      13.172     17.916   0.735   0.4622
## siteousSkyra      -88.725     15.706  -5.649 1.63e-08 ***
## siteUKB            14.159     14.456   0.979   0.3274
## siteUOXF           20.547     16.293   1.261   0.2073
## icv                21.477      0.830  25.876  < 2e-16 ***
## income_scaled      -2.856      1.906  -1.498   0.1341
## education_scaled   -4.998      2.252  -2.219   0.0265 *
## ---
## Signif. codes:  0 ‘***’ 0.001 ‘**’ 0.01 ‘*’ 0.05 ‘.’ 0.1 ‘ ’ 1
##
## Approximate significance of smooth terms:
##              edf Ref.df       F p-value
## s(age_z)   4.966  4.966 507.207  <2e-16 ***
## s(sleep_z) 1.075  1.075   4.387  0.0298 *
## ---
## Signif. codes:  0 ‘***’ 0.001 ‘**’ 0.01 ‘*’ 0.05 ‘.’ 0.1 ‘ ’ 1
##
## R-sq.(adj) =  0.115
## lmer.REML = 3.7826e+05  Scale est. = 1409.9    n = 31193
```

We also included interaction effects between sleep duration and education and income, in another model. The output is shown below, and the interaction terms are `income_scaled:sleep_z` and `education_scaled:sleep_z`.

```
##
## Family: gaussian
## Link function: identity
##
## Formula:
## value ~ sex + site + s(age_z, k = 10, bs = "cr") + s(sleep_z,
##     k = 5, bs = "cr") + icv + income_scaled + education_scaled +
##     income_scaled:sleep_z + education_scaled:sleep_z
## <environment: 0x55cf401af668>
##
## Parametric coefficients:
##                          Estimate Std. Error t value Pr(>|t|)
## (Intercept)               521.302     14.569  35.782  < 2e-16 ***
```

```
## sexmale                   -18.651      1.607 -11.607  < 2e-16 ***
## siteousAvanto             -87.833     17.956  -4.892 1.00e-06 ***
## siteousPrisma              13.723     17.918   0.766   0.4438
## siteousSkyra              -88.486     15.706  -5.634 1.78e-08 ***
## siteUKB                    14.311     14.457   0.990   0.3222
## siteUOXF                   20.741     16.298   1.273   0.2032
## icv                        21.457      0.830  25.851  < 2e-16 ***
## income_scaled              -2.973      1.907  -1.559   0.1189
## education_scaled           -4.922      2.253  -2.185   0.0289 *
## income_scaled:sleep_z       2.915      1.886   1.545   0.1223
## education_scaled:sleep_z   -3.701      2.181  -1.697   0.0897 .
## ---
## Signif. codes:  0 ‘***’ 0.001 ‘**’ 0.01 ‘*’ 0.05 ‘.’ 0.1 ‘ ’ 1
##
## Approximate significance of smooth terms:
##              edf Ref.df       F p-value
## s(age_z)   4.976  4.976 506.822  <2e-16 ***
## s(sleep_z) 1.000  1.000   0.001   0.977
## ---
## Signif. codes:  0 ‘***’ 0.001 ‘**’ 0.01 ‘*’ 0.05 ‘.’ 0.1 ‘ ’ 1
##
## R-sq.(adj) =  0.115
## lmer.REML = 3.7825e+05  Scale est. = 1409.8    n = 31193
```

We did the same controling for BMI. Below is the model with no covariates but only keeping data with BMI.

```
##
## Family: gaussian
## Link function: identity
##
## Formula:
## value ~ sex + site + s(age_z, k = 10, bs = "cr") + s(sleep_z,
##     k = 5, bs = "cr") + icv
## <environment: 0x55cf3f88a628>
##
## Parametric coefficients:
##               Estimate Std. Error t value Pr(>|t|)
## (Intercept)   429.6670    10.8772  39.501  < 2e-16 ***
## sexmale       -18.1909     1.5542 -11.704  < 2e-16 ***
## siteousPrisma  91.7286    14.1516   6.482 9.19e-11 ***
## siteousSkyra    0.4329     9.0640   0.048    0.962
## siteUCAM       77.1885    11.3820   6.782 1.21e-11 ***
## siteUKB       101.4436    10.9521   9.262  < 2e-16 ***
## siteUmU        84.1346    12.6246   6.664 2.70e-11 ***
## icv            20.9038     0.8010  26.096  < 2e-16 ***
## ---
## Signif. codes:  0 ‘***’ 0.001 ‘**’ 0.01 ‘*’ 0.05 ‘.’ 0.1 ‘ ’ 1
##
## Approximate significance of smooth terms:
##             edf Ref.df       F p-value
## s(age_z)   5.41   5.41 558.567  <2e-16 ***
## s(sleep_z) 1.00   1.00   5.666  0.0173 *
## ---
## Signif. codes:  0 ‘***’ 0.001 ‘**’ 0.01 ‘*’ 0.05 ‘.’ 0.1 ‘ ’ 1
##
```

```
## R-sq.(adj) =  0.119
## lmer.REML = 4.0518e+05  Scale est. = 1345.6    n = 33448
```

Below is the model output with main effect.

```
##
## Family: gaussian
## Link function: identity
##
## Formula:
## value ~ sex + site + s(age_z, k = 10, bs = "cr") + s(sleep_z,
##     k = 5, bs = "cr") + icv + bmi
## <environment: 0x55cf3f88a628>
##
## Parametric coefficients:
##               Estimate Std. Error t value Pr(>|t|)
## (Intercept)   461.2566    11.5189  40.044  < 2e-16 ***
## sexmale       -17.1918     1.5572 -11.041  < 2e-16 ***
## siteousPrisma  93.0805    14.1433   6.581 4.73e-11 ***
## siteousSkyra    0.6502     9.0623   0.072    0.943
## siteUCAM       78.0101    11.3763   6.857 7.14e-12 ***
## siteUKB       102.4378    10.9482   9.357  < 2e-16 ***
## siteUmU        85.5654    12.6178   6.781 1.21e-11 ***
## icv            21.0822     0.8005  26.338  < 2e-16 ***
## bmi            -1.2536     0.1510  -8.299  < 2e-16 ***
## ---
## Signif. codes:  0 '***' 0.001 '**' 0.01 '*' 0.05 '.' 0.1 ' ' 1
##
## Approximate significance of smooth terms:
##              edf Ref.df      F p-value
## s(age_z)   5.439  5.439 556.78 < 2e-16 ***
## s(sleep_z) 1.000  1.000   7.58 0.00591 **
## ---
## Signif. codes:  0 '***' 0.001 '**' 0.01 '*' 0.05 '.' 0.1 ' ' 1
##
## R-sq.(adj) =  0.121
## lmer.REML = 4.0511e+05  Scale est. = 1345.3    n = 33448
```

Next is the model with BMI-sleep interaction.

```
##
## Family: gaussian
## Link function: identity
##
## Formula:
## value ~ sex + site + s(age_z, k = 10, bs = "cr") + s(sleep_z,
##     k = 5, bs = "cr") + icv + bmi + bmi:sleep_z
## <environment: 0x55cf3f88a628>
##
## Parametric coefficients:
##               Estimate Std. Error t value Pr(>|t|)
## (Intercept)   460.9914    11.5239  40.003  < 2e-16 ***
## sexmale       -17.1869     1.5572 -11.037  < 2e-16 ***
## siteousPrisma  93.0877    14.1433   6.582 4.72e-11 ***
## siteousSkyra    0.6819     9.0623   0.075    0.940
## siteUCAM       78.1154    11.3771   6.866 6.72e-12 ***
```

```
## siteUKB       102.5407    10.9490   9.365  < 2e-16 ***
## siteUmU        85.6589    12.6185   6.788 1.15e-11 ***
## icv            21.0864     0.8005  26.342  < 2e-16 ***
## bmi            -1.2495     0.1511  -8.268  < 2e-16 ***
## bmi:sleep_z     0.1131     0.1430   0.791    0.429
## ---
## Signif. codes:  0 '***' 0.001 '**' 0.01 '*' 0.05 '.' 0.1 ' ' 1
##
## Approximate significance of smooth terms:
##              edf Ref.df       F p-value
## s(age_z)   5.441  5.441 556.730  <2e-16 ***
## s(sleep_z) 1.000  1.000   1.535   0.215
## ---
## Signif. codes:  0 '***' 0.001 '**' 0.01 '*' 0.05 '.' 0.1 ' ' 1
##
## R-sq.(adj) =  0.121
## lmer.REML = 4.0511e+05  Scale est. = 1345.3    n = 33448
```

We did the same controling for depression. Below is the model with no covariates but only keeping data with depression.

```
##
## Family: gaussian
## Link function: identity
##
## Formula:
## value ~ sex + site + s(age_z, k = 10, bs = "cr") + s(sleep_z,
##     k = 5, bs = "cr") + icv
## <environment: 0x55cf3d0705b8>
##
## Parametric coefficients:
##               Estimate Std. Error t value Pr(>|t|)
## (Intercept)    517.792      7.336  70.578  < 2e-16 ***
## sexmale        -17.977      1.558 -11.536  < 2e-16 ***
## siteousAvanto  -91.668     17.090  -5.364  8.2e-08 ***
## siteousPrisma   -1.249     14.263  -0.088   0.9302
## siteousSkyra   -97.644     11.076  -8.816  < 2e-16 ***
## siteUCAM       -18.108     10.033  -1.805   0.0711 .
## siteUKB         12.553      7.253   1.731   0.0835 .
## siteUmU         -6.165      9.850  -0.626   0.5314
## icv             20.474      0.802  25.530  < 2e-16 ***
## ---
## Signif. codes:  0 '***' 0.001 '**' 0.01 '*' 0.05 '.' 0.1 ' ' 1
##
## Approximate significance of smooth terms:
##              edf Ref.df       F p-value
## s(age_z)   5.274  5.274 550.287  <2e-16 ***
## s(sleep_z) 1.000  1.000   5.385  0.0203 *
## ---
## Signif. codes:  0 '***' 0.001 '**' 0.01 '*' 0.05 '.' 0.1 ' ' 1
##
## R-sq.(adj) =  0.117
## lmer.REML = 4.038e+05  Scale est. = 1319.2    n = 33360
```

Below is the model output with main effect.

```
##
## Family: gaussian
## Link function: identity
##
## Formula:
## value ~ sex + site + s(age_z, k = 10, bs = "cr") + s(sleep_z,
##     k = 5, bs = "cr") + icv + depression
## <environment: 0x55cf3d0705b8>
##
## Parametric coefficients:
##                Estimate Std. Error t value Pr(>|t|)
## (Intercept)   519.0433     7.3694  70.432  < 2e-16 ***
## sexmale       -18.0998     1.5598 -11.604  < 2e-16 ***
## siteousAvanto -91.0391    17.0945  -5.326 1.01e-07 ***
## siteousPrisma  -0.6967    14.2677  -0.049   0.9611
## siteousSkyra  -96.9344    11.0842  -8.745  < 2e-16 ***
## siteUCAM      -17.8312    10.0342  -1.777   0.0756 .
## siteUKB        12.0105     7.2593   1.655   0.0980 .
## siteUmU        -2.5116    10.0600  -0.250   0.8028
## icv            20.4850     0.8020  25.543  < 2e-16 ***
## depression     -9.8820     5.5266  -1.788   0.0738 .
## ---
## Signif. codes:  0 ‘***’ 0.001 ‘**’ 0.01 ‘*’ 0.05 ‘.’ 0.1 ‘ ’ 1
##
## Approximate significance of smooth terms:
##              edf Ref.df       F p-value
## s(age_z)   5.301  5.301 542.003  <2e-16 ***
## s(sleep_z) 1.000  1.000   5.956  0.0147 *
## ---
## Signif. codes:  0 ‘***’ 0.001 ‘**’ 0.01 ‘*’ 0.05 ‘.’ 0.1 ‘ ’ 1
##
## R-sq.(adj) =  0.117
## lmer.REML = 4.038e+05  Scale est. = 1319.1    n = 33360
```

Next is the model with depression-sleep interaction.

```
##
## Family: gaussian
## Link function: identity
##
## Formula:
## value ~ sex + site + s(age_z, k = 10, bs = "cr") + s(sleep_z,
##     k = 5, bs = "cr") + icv + depression + depression:sleep_z
## <environment: 0x55cf3d0705b8>
##
## Parametric coefficients:
##                     Estimate Std. Error t value Pr(>|t|)
## (Intercept)          519.038      7.369  70.434  < 2e-16 ***
## sexmale              -18.153      1.560 -11.637  < 2e-16 ***
## siteousAvanto        -91.462     17.095  -5.350 8.84e-08 ***
## siteousPrisma         -1.105     14.268  -0.077   0.9383
## siteousSkyra         -97.339     11.085  -8.781  < 2e-16 ***
## siteUCAM             -18.228     10.036  -1.816   0.0693 .
## siteUKB               12.035      7.259   1.658   0.0973 .
## siteUmU               -0.038     10.142  -0.004   0.9970
```

```
## icv                   20.492      0.802  25.552  < 2e-16 ***
## depression           -10.471      5.535  -1.892   0.0585 .
## depression:sleep_z    -7.741      4.037  -1.918   0.0552 .
## ---
## Signif. codes:  0 '***' 0.001 '**' 0.01 '*' 0.05 '.' 0.1 ' ' 1
##
## Approximate significance of smooth terms:
##               edf Ref.df       F p-value
## s(age_z)    5.291  5.291 543.411  <2e-16 ***
## s(sleep_z) 1.000  1.000   1.067   0.302
## ---
## Signif. codes:  0 '***' 0.001 '**' 0.01 '*' 0.05 '.' 0.1 ' ' 1
##
## R-sq.(adj) =  0.117
## lmer.REML = 4.0379e+05  Scale est. = 1319.2    n = 33360
```

The plot below shows the sleep-volume curve for the original model and for the model with main effects of SES.

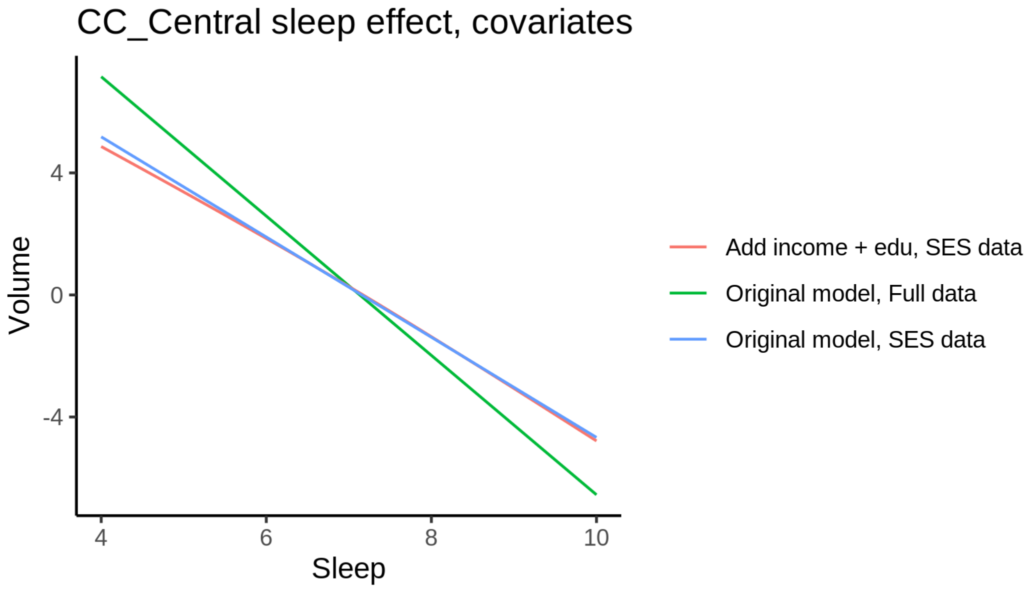


The plot below shows the sleep-volume curve for the original model and for the model with main effects of BMI.

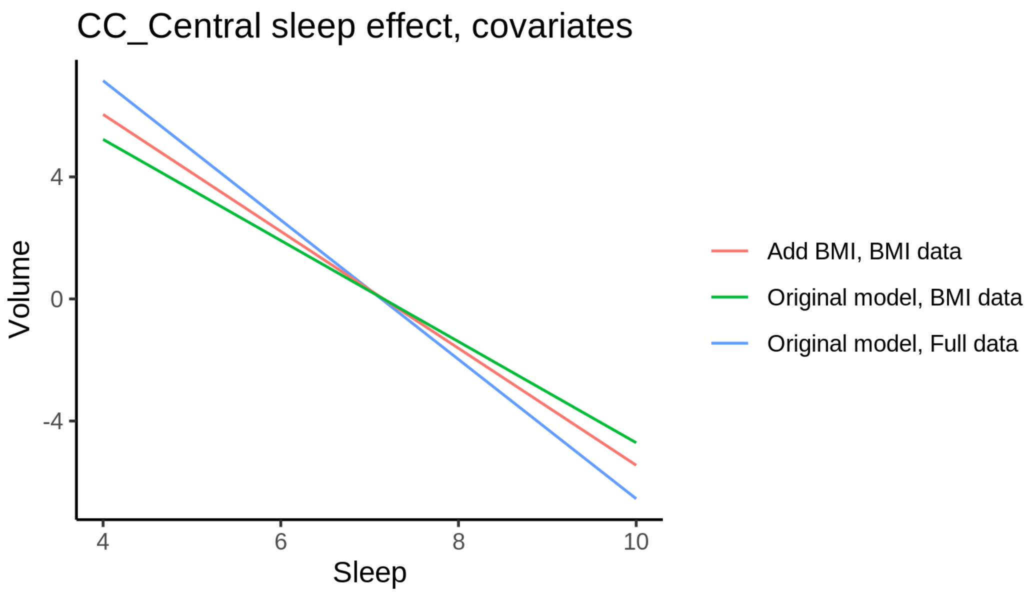


The plot below shows the sleep-volume curve for the original model and for the model with main effects of depression.

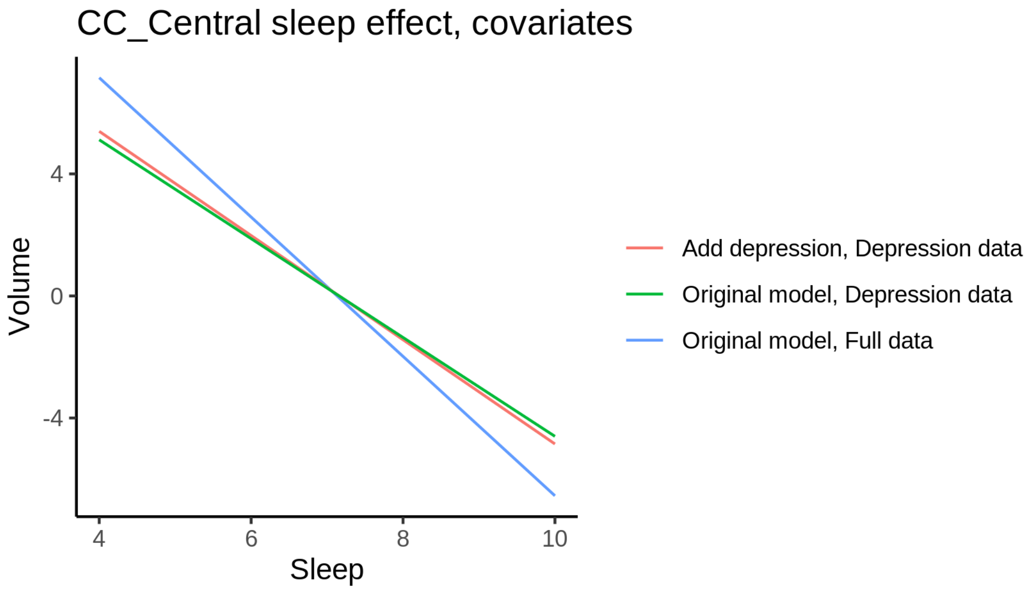


### CC_Mid_Anterior

#### Descriptive statistics

| Study | Observations | Unique IDs | Mean age | Age range |
|---|---|---|---|---|
| HCP | 974 | 974 | 28.8 | 22 - 37 |
| MPIB | 677 | 391 | 63.1 | 24 - 83 |
| UB | 113 | 39 | 70.9 | 64 - 81 |
| UCAM | 881 | 631 | 55.1 | 20 - 88 |

| Study | Observations | Unique IDs | Mean age | Age range |
|---|---|---|---|---|
| UiO | 1475 | 803 | 49.4 | 20 - 89 |
| UKB | 45966 | 43131 | 64.5 | 45 - 83 |
| UmU | 423 | 284 | 62.3 | 25 - 85 |
| UOXF | 769 | 769 | 69.8 | 60 - 85 |

##### Spaghetti plot

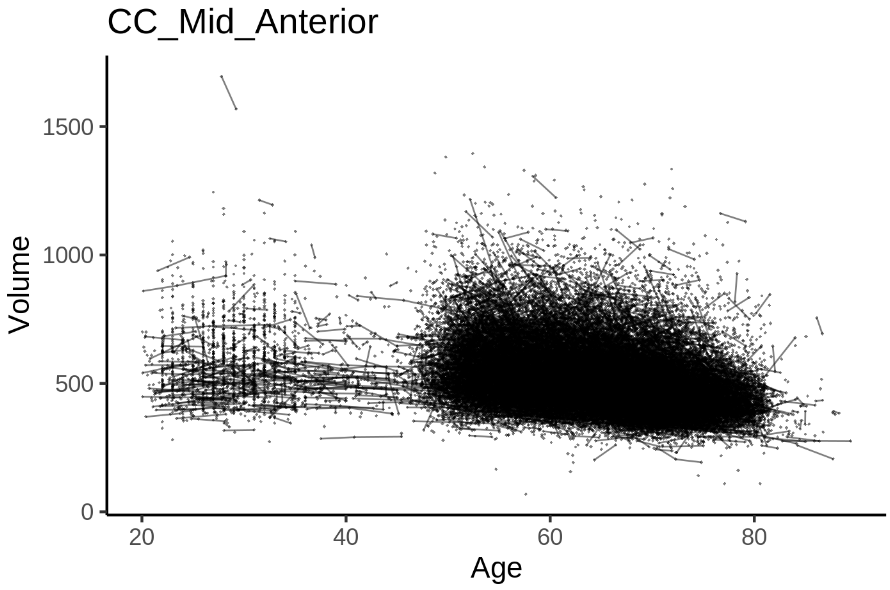


##### Model outputs

Model without sleep term

```
##
## Family: gaussian
## Link function: identity
##
## Formula:
## value ~ sex + site + icv + s(age_z, k = 10, bs = "cr")
## <environment: 0x55cf42eee758>
##
## Parametric coefficients:
##                 Estimate Std. Error t value Pr(>|t|)
## (Intercept)     552.3647     6.3874  86.477  < 2e-16 ***
## sexmale         -17.0689     1.3101 -13.029  < 2e-16 ***
## siteMPIB         -0.8302     8.1646  -0.102 0.919004
## siteousAvanto -104.7971     6.6723 -15.706  < 2e-16 ***
## siteousPrisma  -38.5515    11.3100  -3.409 0.000653 ***
## siteousSkyra  -101.7467     6.2245 -16.346  < 2e-16 ***
## siteUB         -75.3073    19.0945  -3.944 8.03e-05 ***
## siteUCAM       -52.9758     7.0815  -7.481 7.50e-14 ***
## siteUKB        -18.4330     6.5496  -2.814 0.004889 **
## siteUmU        -39.3966     9.1040  -4.327 1.51e-05 ***
```

```
## siteUOXF        -8.6763     7.7648  -1.117 0.263831
## icv             30.8259     0.6592  46.766  < 2e-16 ***
## ---
## Signif. codes:  0 '***' 0.001 '**' 0.01 '*' 0.05 '.' 0.1 ' ' 1
##
## Approximate significance of smooth terms:
##            edf Ref.df     F p-value
## s(age_z) 6.189  6.189 844.6  <2e-16 ***
## ---
## Signif. codes:  0 '***' 0.001 '**' 0.01 '*' 0.05 '.' 0.1 ' ' 1
##
## R-sq.(adj) =   0.16
## lmer.REML = 6.2553e+05  Scale est. = 1315.2    n = 51278
```

Model with only main effects of age and sleep

```
##
## Family: gaussian
## Link function: identity
##
## Formula:
## value ~ sex + site + icv + s(age_z, k = 10, bs = "cr") + s(sleep_z,
##     k = 5, bs = "cr")
## <environment: 0x55cf42eee758>
##
## Parametric coefficients:
##                  Estimate Std. Error t value Pr(>|t|)
## (Intercept)      551.0236     6.3900  86.232  < 2e-16 ***
## sexmale          -17.1166     1.3098 -13.068  < 2e-16 ***
## siteMPIB           0.6327     8.1675   0.077  0.93825
## siteousAvanto  -104.0882     6.6719 -15.601  < 2e-16 ***
## siteousPrisma   -38.0773    11.3079  -3.367  0.00076 ***
## siteousSkyra   -101.0144     6.2242 -16.229  < 2e-16 ***
## siteUB          -74.9009    19.0893  -3.924 8.73e-05 ***
## siteUCAM        -52.2898     7.0802  -7.385 1.54e-13 ***
## siteUKB         -17.0075     6.5527  -2.596  0.00945 **
## siteUmU         -36.4298     9.1222  -3.994 6.52e-05 ***
## siteUOXF         -7.9887     7.7624  -1.029  0.30342
## icv              30.9206     0.6593  46.899  < 2e-16 ***
## ---
## Signif. codes:  0 '***' 0.001 '**' 0.01 '*' 0.05 '.' 0.1 ' ' 1
##
## Approximate significance of smooth terms:
##              edf Ref.df      F  p-value
## s(age_z)   6.164  6.164 838.38  < 2e-16 ***
## s(sleep_z) 1.000  1.000  22.32 2.34e-06 ***
## ---
## Signif. codes:  0 '***' 0.001 '**' 0.01 '*' 0.05 '.' 0.1 ' ' 1
##
## R-sq.(adj) =   0.16
## lmer.REML = 6.2551e+05  Scale est. = 1315.1    n = 51278
```

Model with full interaction between age and sleep

```
##
## Family: gaussian
```

```
## Link function: identity
##
## Formula:
## value ~ sex + site + icv + t2(age_z, sleep_z, k = c(10, 4), bs = "cr")
## <environment: 0x55cf42eee758>
##
## Parametric coefficients:
##                Estimate Std. Error t value Pr(>|t|)
## (Intercept)    550.7137     6.4179  85.810  < 2e-16 ***
## sexmale        -17.2953     1.3124 -13.178  < 2e-16 ***
## siteMPIB         1.1718     8.2120   0.143 0.886534
## siteousAvanto -104.0577     6.7076 -15.513  < 2e-16 ***
## siteousPrisma  -38.1061    11.3302  -3.363 0.000771 ***
## siteousSkyra  -100.8759     6.2654 -16.100  < 2e-16 ***
## siteUB         -74.0957    19.1023  -3.879 0.000105 ***
## siteUCAM       -51.7512     7.1221  -7.266 3.75e-13 ***
## siteUKB        -16.6063     6.5806  -2.524 0.011621 *
## siteUmU        -36.3852     9.1534  -3.975 7.05e-05 ***
## siteUOXF        -7.2257     7.7934  -0.927 0.353850
## icv             30.9409     0.6594  46.925  < 2e-16 ***
## ---
## Signif. codes:  0 ‘***’ 0.001 ‘**’ 0.01 ‘*’ 0.05 ‘.’ 0.1 ‘ ’ 1
##
## Approximate significance of smooth terms:
##                     edf Ref.df     F p-value
## t2(age_z,sleep_z) 9.538  9.538 17.09  <2e-16 ***
## ---
## Signif. codes:  0 ‘***’ 0.001 ‘**’ 0.01 ‘*’ 0.05 ‘.’ 0.1 ‘ ’ 1
##
## R-sq.(adj) =   0.16
## lmer.REML = 6.2551e+05  Scale est. = 1314.7    n = 51278
```

#### Model comparison

`mod_no_sleep` refers to model without sleep term, `mod_no_interaction` refers to model with only main effect of sleep, and `mod_full` refers to model with a full interaction between age and sleep. This is a nested model comparison, and the p-value at a given line refers to comparing the model at the line to the model on the line above. Hence, significance implies that the more complicated model is supported on statistical grounds.

To be even more specific, the p-value on the second row tests whether there is an association between sleep and volume. The p-value on the third row tests whether this association depends on age.

```
## Data: NULL
## Models:
## mod_list$mod_no_sleep$mer: NULL
## mod_list$mod_no_interaction$mer: NULL
## mod_list$mod_full$mer: NULL
##                                 npar    AIC    BIC  logLik deviance  Chisq Df Pr(>Chisq)
## mod_list$mod_no_sleep$mer         16 625564 625706 -312766   625532
## mod_list$mod_no_interaction$mer   18 625546 625705 -312755   625510 22.312  2  1.429e-05 ***
## mod_list$mod_full$mer             20 625551 625728 -312755   625511  0.000  2          1
## ---
## Signif. codes:  0 ‘***’ 0.001 ‘**’ 0.01 ‘*’ 0.05 ‘.’ 0.1 ‘ ’ 1
```

We chose the model based on the likelihood ratio test with 5 % significance level, which was `mod_no_interaction`.

#### Lifespan brain trajectory

The trajectory shown is from the chosen model `mod_no_interaction`.

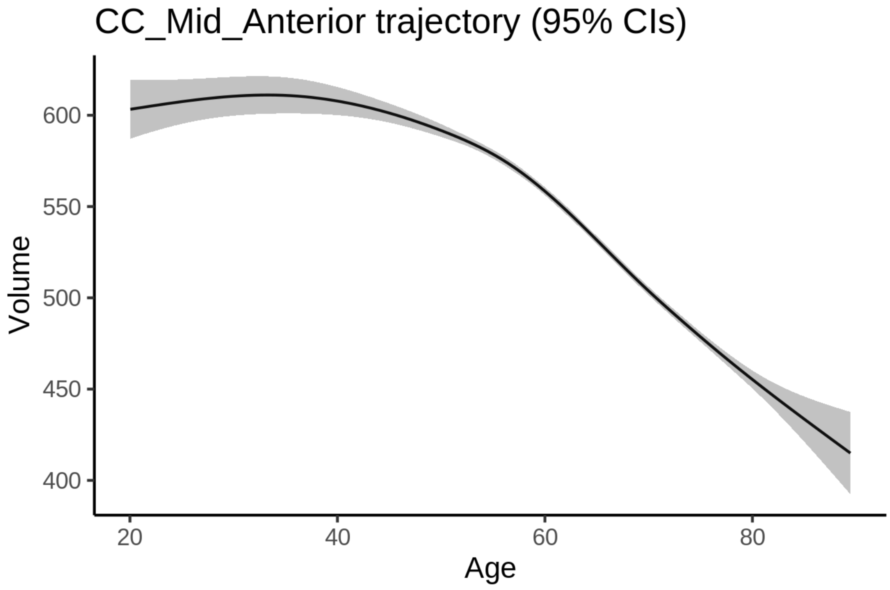


#### Effect of sleep

The chosen model only included the main effect of sleep, and hence the effect does not vary with age. The black dot shows the average sleep duration across all ages in the sample.

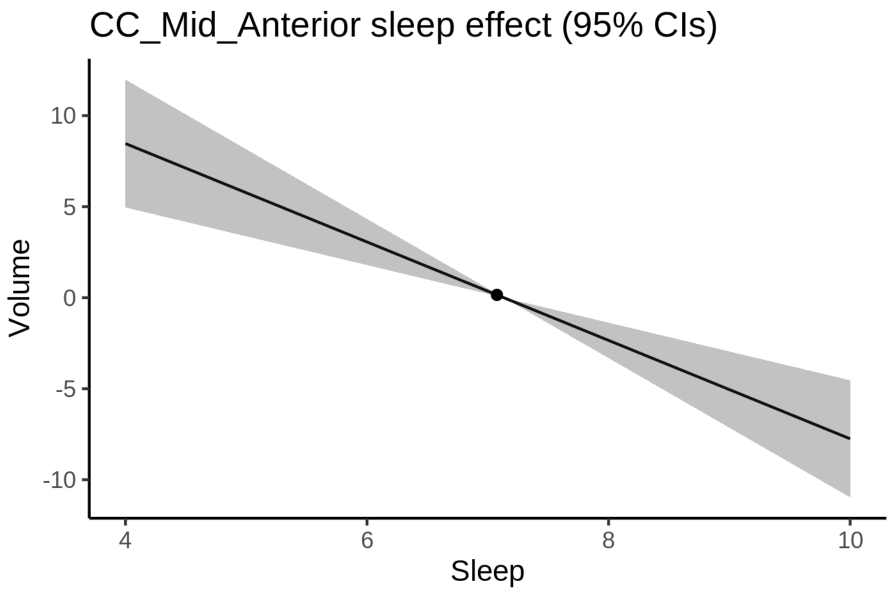


We also show the full interaction model for completeness, although it was not selected.

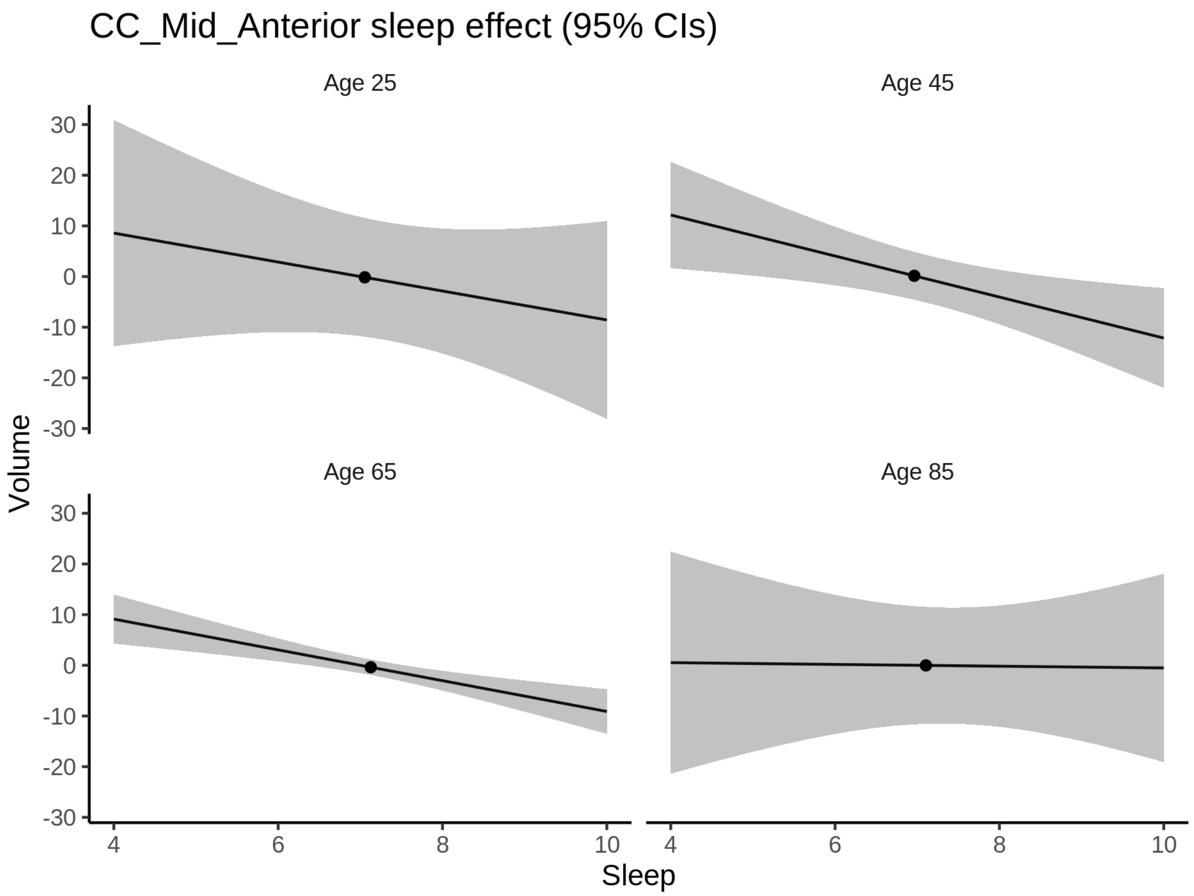


**Deviation from sleep associated with maximal volume**

Model with only main effect of sleep was chosen, so we show it for all ages at once. Maximum volume is attained at 4 hours of sleep. The percentage values in the plot are calculated as follows: The maximum at 100 % refers to a person at an arbitrary age with a sleep duration associated with maximum volume. For a female, this volume is 548 and for a male it is 531. The other percentage values show how large the expected volume is for someone with other sleep durations. For example, 99 % implies a 1 % reduction.

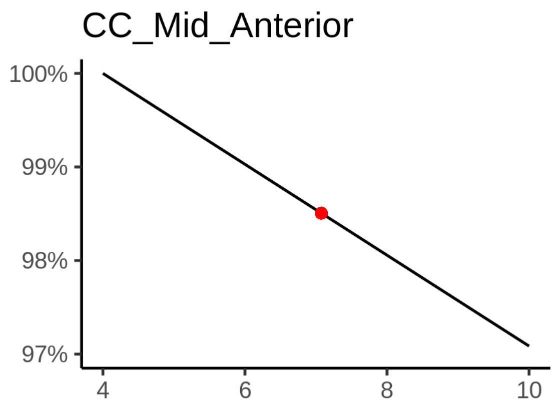

**Comparison of mean sleep and sleep associated with maximum volume**

A 95 % confidence interval for the sleep associated with maximum volume is [4, 4].

The plot below compares average sleep to the sleep associated with maximum volume.

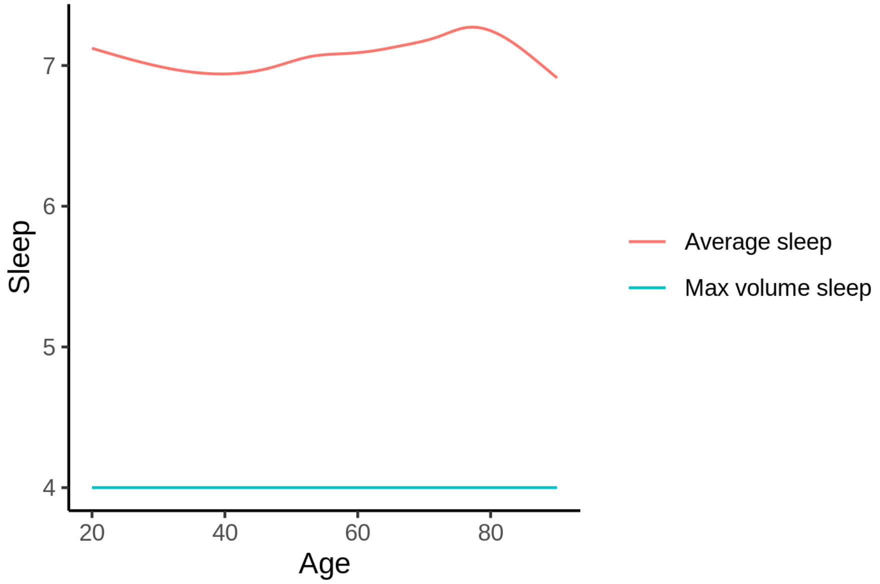


The next plot shows the difference between average sleep and sleep associated with maximum volume. The shaded region is a 95 % confidence interval.

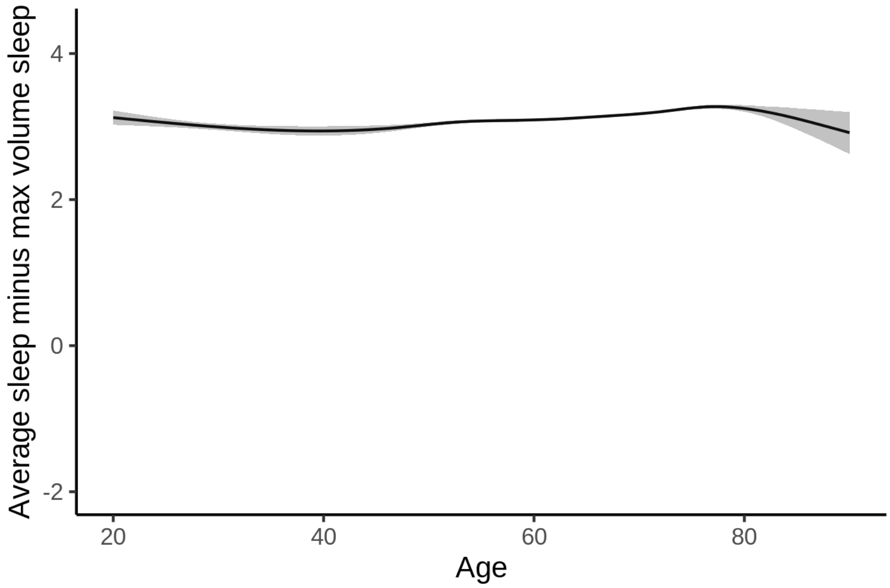


The next plot shows the probability that the sleep duration associated with maximum volume is longer than the average sleep duration, as a function of age. Probility below .05 can be interpreted as evidence that the sleep associated with maximum volume is shorter than the mean sleep, and probability above .95 can be interpreted the opposite way.

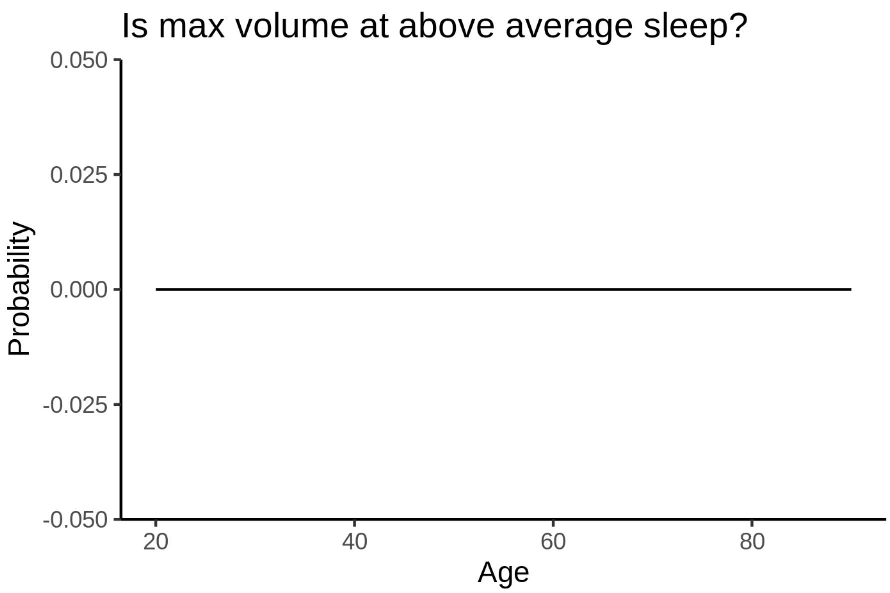


##### Controling for covariates

Below is the output for a model in which we only include data with income and education.

```
##
## Family: gaussian
## Link function: identity
##
## Formula:
## value ~ sex + site + s(age_z, k = 10, bs = "cr") + s(sleep_z,
##     k = 5, bs = "cr") + icv
## <environment: 0x55cf3df85088>
##
## Parametric coefficients:
##                Estimate Std. Error t value Pr(>|t|)
## (Intercept)    555.7995    14.9399  37.202  < 2e-16 ***
## sexmale        -18.4100     1.6524 -11.141  < 2e-16 ***
## siteousAvanto -119.1874    18.6147  -6.403 1.55e-10 ***
## siteousPrisma  -41.8377    18.4940  -2.262   0.0237 *
## siteousSkyra  -111.3991    16.1943  -6.879 6.15e-12 ***
## siteUKB        -23.5826    14.9125  -1.581   0.1138
## siteUOXF       -15.0958    16.8079  -0.898   0.3691
## icv             32.3789     0.8492  38.128  < 2e-16 ***
## ---
## Signif. codes:  0 '***' 0.001 '**' 0.01 '*' 0.05 '.' 0.1 ' ' 1
##
## Approximate significance of smooth terms:
##              edf Ref.df       F p-value
## s(age_z)   5.218  5.218 608.281 < 2e-16 ***
## s(sleep_z) 1.000  1.000   9.265 0.00234 **
## ---
## Signif. codes:  0 '***' 0.001 '**' 0.01 '*' 0.05 '.' 0.1 ' ' 1
##
```

```
## R-sq.(adj) =   0.15
## lmer.REML = 3.8026e+05  Scale est. = 1585.3    n = 31185
```

Below is the output for a model in which we control for the main effects of income and education.

```
##
## Family: gaussian
## Link function: identity
##
## Formula:
## value ~ sex + site + s(age_z, k = 10, bs = "cr") + s(sleep_z,
##     k = 5, bs = "cr") + icv + income_scaled + education_scaled
## <environment: 0x55cf3df85088>
##
## Parametric coefficients:
##                   Estimate Std. Error t value Pr(>|t|)
## (Intercept)       561.7247    15.0377  37.354  < 2e-16 ***
## sexmale           -18.4897     1.6570 -11.159  < 2e-16 ***
## siteousAvanto    -118.8309    18.6176  -6.383 1.76e-10 ***
## siteousPrisma     -41.0324    18.4934  -2.219   0.0265 *
## siteousSkyra     -110.9974    16.1967  -6.853 7.36e-12 ***
## siteUKB           -23.4352    14.9228  -1.570   0.1163
## siteUOXF          -16.9538    16.8180  -1.008   0.3134
## icv                32.7604     0.8566  38.243  < 2e-16 ***
## income_scaled      -2.9597     1.9667  -1.505   0.1324
## education_scaled   -5.9524     2.3242  -2.561   0.0104 *
## ---
## Signif. codes:  0 ‘***’ 0.001 ‘**’ 0.01 ‘*’ 0.05 ‘.’ 0.1 ‘ ’ 1
##
## Approximate significance of smooth terms:
##              edf Ref.df       F p-value
## s(age_z)   5.279  5.279 561.243 < 2e-16 ***
## s(sleep_z) 1.000  1.000   8.956 0.00277 **
## ---
## Signif. codes:  0 ‘***’ 0.001 ‘**’ 0.01 ‘*’ 0.05 ‘.’ 0.1 ‘ ’ 1
##
## R-sq.(adj) =  0.151
## lmer.REML = 3.8024e+05  Scale est. = 1584.9    n = 31185
```

We also included interaction effects between sleep duration and education and income, in another model. The output is shown below, and the interaction terms are `income_scaled:sleep_z` and `education_scaled:sleep_z`.

```
##
## Family: gaussian
## Link function: identity
##
## Formula:
## value ~ sex + site + s(age_z, k = 10, bs = "cr") + s(sleep_z,
##     k = 5, bs = "cr") + icv + income_scaled + education_scaled +
##     income_scaled:sleep_z + education_scaled:sleep_z
## <environment: 0x55cf3df85088>
##
## Parametric coefficients:
##                           Estimate Std. Error t value Pr(>|t|)
## (Intercept)               561.8578    15.0389  37.360  < 2e-16 ***
```

```
## sexmale                   -18.3803     1.6581 -11.085  < 2e-16 ***
## siteousAvanto            -118.7398    18.6180  -6.378 1.82e-10 ***
## siteousPrisma             -40.6529    18.4951  -2.198   0.0280 *
## siteousSkyra             -110.8987    16.1968  -6.847 7.68e-12 ***
## siteUKB                   -23.5949    14.9246  -1.581   0.1139
## siteUOXF                  -17.3012    16.8229  -1.028   0.3038
## icv                        32.7541     0.8567  38.232  < 2e-16 ***
## income_scaled              -3.0758     1.9678  -1.563   0.1180
## education_scaled           -5.8957     2.3244  -2.536   0.0112 *
## income_scaled:sleep_z       3.5910     1.9462   1.845   0.0650 .
## education_scaled:sleep_z   -1.1372     2.2498  -0.505   0.6132
## ---
## Signif. codes:  0 ‘***’ 0.001 ‘**’ 0.01 ‘*’ 0.05 ‘.’ 0.1 ‘ ’ 1
##
## Approximate significance of smooth terms:
##              edf Ref.df       F p-value
## s(age_z)   5.293  5.293 560.349  <2e-16 ***
## s(sleep_z) 1.000  1.000   2.303   0.129
## ---
## Signif. codes:  0 ‘***’ 0.001 ‘**’ 0.01 ‘*’ 0.05 ‘.’ 0.1 ‘ ’ 1
##
## R-sq.(adj) =  0.151
## lmer.REML = 3.8023e+05  Scale est. = 1584.9    n = 31185
```

We did the same controling for BMI. Below is the model with no covariates but only keeping data with BMI.

```
##
## Family: gaussian
## Link function: identity
##
## Formula:
## value ~ sex + site + s(age_z, k = 10, bs = "cr") + s(sleep_z,
##     k = 5, bs = "cr") + icv
## <environment: 0x55cf418e0640>
##
## Parametric coefficients:
##               Estimate Std. Error t value Pr(>|t|)
## (Intercept)   449.2700    11.3258  39.668  < 2e-16 ***
## sexmale       -17.4284     1.5993 -10.897  < 2e-16 ***
## siteousPrisma  64.7800    14.6743   4.415 1.02e-05 ***
## siteousSkyra    1.8415     9.4780   0.194    0.846
## siteUCAM       51.2029    11.8289   4.329 1.50e-05 ***
## siteUKB        83.2341    11.4036   7.299 2.97e-13 ***
## siteUmU        69.5968    13.0973   5.314 1.08e-07 ***
## icv            32.1221     0.8246  38.957  < 2e-16 ***
## ---
## Signif. codes:  0 ‘***’ 0.001 ‘**’ 0.01 ‘*’ 0.05 ‘.’ 0.1 ‘ ’ 1
##
## Approximate significance of smooth terms:
##              edf Ref.df       F p-value
## s(age_z)   5.957  5.957 576.135 < 2e-16 ***
## s(sleep_z) 1.043  1.043   8.879 0.00244 **
## ---
## Signif. codes:  0 ‘***’ 0.001 ‘**’ 0.01 ‘*’ 0.05 ‘.’ 0.1 ‘ ’ 1
##
```

```
## R-sq.(adj) =  0.153
## lmer.REML = 4.0706e+05  Scale est. = 1475.2    n = 33438
```

Below is the model output with main effect.

```
##
## Family: gaussian
## Link function: identity
##
## Formula:
## value ~ sex + site + s(age_z, k = 10, bs = "cr") + s(sleep_z,
##     k = 5, bs = "cr") + icv + bmi
## <environment: 0x55cf418e0640>
##
## Parametric coefficients:
##               Estimate Std. Error t value Pr(>|t|)
## (Intercept)   470.7728    11.9842  39.283  < 2e-16 ***
## sexmale       -16.7470     1.6033 -10.445  < 2e-16 ***
## siteousPrisma  65.7213    14.6713   4.480 7.50e-06 ***
## siteousSkyra    2.0154     9.4777   0.213    0.832
## siteUCAM       51.7124    11.8273   4.372 1.23e-05 ***
## siteUKB        83.9202    11.4032   7.359 1.89e-13 ***
## siteUmU        70.5143    13.0955   5.385 7.31e-08 ***
## icv            32.2604     0.8248  39.115  < 2e-16 ***
## bmi            -0.8536     0.1557  -5.484 4.18e-08 ***
## ---
## Signif. codes:  0 '***' 0.001 '**' 0.01 '*' 0.05 '.' 0.1 ' ' 1
##
## Approximate significance of smooth terms:
##              edf Ref.df       F p-value
## s(age_z)   5.990  5.990 573.810 < 2e-16 ***
## s(sleep_z) 1.579  1.579   5.984 0.00364 **
## ---
## Signif. codes:  0 '***' 0.001 '**' 0.01 '*' 0.05 '.' 0.1 ' ' 1
##
## R-sq.(adj) =  0.154
## lmer.REML = 4.0703e+05  Scale est. = 1475.2    n = 33438
```

Next is the model with BMI-sleep interaction.

```
##
## Family: gaussian
## Link function: identity
##
## Formula:
## value ~ sex + site + s(age_z, k = 10, bs = "cr") + s(sleep_z,
##     k = 5, bs = "cr") + icv + bmi + bmi:sleep_z
## <environment: 0x55cf418e0640>
##
## Parametric coefficients:
##                Estimate Std. Error t value Pr(>|t|)
## (Intercept)   470.85388   11.98922  39.273  < 2e-16 ***
## sexmale       -16.74877    1.60337 -10.446  < 2e-16 ***
## siteousPrisma  65.71758   14.67139   4.479 7.51e-06 ***
## siteousSkyra    2.00447    9.47776   0.211    0.833
## siteUCAM       51.68047   11.82813   4.369 1.25e-05 ***
```

```
## siteUKB        83.88702   11.40404   7.356 1.94e-13 ***
## siteUmU        70.48640   13.09617   5.382 7.41e-08 ***
## icv            32.25854    0.82479  39.111  < 2e-16 ***
## bmi            -0.85478    0.15574  -5.489 4.08e-08 ***
## bmi:sleep_z    -0.03542    0.14739  -0.240    0.810
## ---
## Signif. codes:  0 '***' 0.001 '**' 0.01 '*' 0.05 '.' 0.1 ' ' 1
##
## Approximate significance of smooth terms:
##              edf Ref.df       F p-value
## s(age_z)   5.989  5.989 573.611  <2e-16 ***
## s(sleep_z) 1.561  1.561   0.182    0.72
## ---
## Signif. codes:  0 '***' 0.001 '**' 0.01 '*' 0.05 '.' 0.1 ' ' 1
##
## R-sq.(adj) =  0.154
## lmer.REML = 4.0703e+05  Scale est. = 1475.2    n = 33438
```

We did the same controling for depression. Below is the model with no covariates but only keeping data with depression.

```
##
## Family: gaussian
## Link function: identity
##
## Formula:
## value ~ sex + site + s(age_z, k = 10, bs = "cr") + s(sleep_z,
##     k = 5, bs = "cr") + icv
## <environment: 0x55cffba79778>
##
## Parametric coefficients:
##                Estimate Std. Error t value Pr(>|t|)
## (Intercept)    542.0368     7.5570  71.727  < 2e-16 ***
## sexmale        -17.5091     1.6062 -10.901  < 2e-16 ***
## siteousAvanto -101.3754    17.7220  -5.720 1.07e-08 ***
## siteousPrisma  -30.1393    14.7297  -2.046   0.0407 *
## siteousSkyra  -101.5808    11.4403  -8.879  < 2e-16 ***
## siteUCAM       -47.6493    10.3531  -4.602 4.19e-06 ***
## siteUKB        -10.0097     7.4712  -1.340   0.1803
## siteUmU        -24.4606    10.1487  -2.410   0.0159 *
## icv             31.7893     0.8269  38.446  < 2e-16 ***
## ---
## Signif. codes:  0 '***' 0.001 '**' 0.01 '*' 0.05 '.' 0.1 ' ' 1
##
## Approximate significance of smooth terms:
##              edf Ref.df       F p-value
## s(age_z)   5.554  5.554 602.201 < 2e-16 ***
## s(sleep_z) 1.000  1.000   9.173 0.00246 **
## ---
## Signif. codes:  0 '***' 0.001 '**' 0.01 '*' 0.05 '.' 0.1 ' ' 1
##
## R-sq.(adj) =   0.15
## lmer.REML = 4.0576e+05  Scale est. = 1431      n = 33350
```

Below is the model output with main effect.

```
##
## Family: gaussian
## Link function: identity
##
## Formula:
## value ~ sex + site + s(age_z, k = 10, bs = "cr") + s(sleep_z,
##     k = 5, bs = "cr") + icv + depression
## <environment: 0x55cffba79778>
##
## Parametric coefficients:
##                Estimate Std. Error t value Pr(>|t|)
## (Intercept)    542.3941     7.5915  71.448  < 2e-16 ***
## sexmale        -17.5442     1.6078 -10.912  < 2e-16 ***
## siteousAvanto -101.1926    17.7258  -5.709 1.15e-08 ***
## siteousPrisma  -29.9785    14.7336  -2.035   0.0419 *
## siteousSkyra  -101.3751    11.4478  -8.855  < 2e-16 ***
## siteUCAM       -47.5671    10.3548  -4.594 4.37e-06 ***
## siteUKB        -10.1649     7.4779  -1.359   0.1741
## siteUmU        -23.4174    10.3653  -2.259   0.0239 *
## icv             31.7924     0.8269  38.448  < 2e-16 ***
## depression      -2.8188     5.6960  -0.495   0.6207
## ---
## Signif. codes:  0 ‘***’ 0.001 ‘**’ 0.01 ‘*’ 0.05 ‘.’ 0.1 ‘ ’ 1
##
## Approximate significance of smooth terms:
##              edf Ref.df       F p-value
## s(age_z)   5.561  5.561 591.503 < 2e-16 ***
## s(sleep_z) 1.000  1.000   9.339 0.00225 **
## ---
## Signif. codes:  0 ‘***’ 0.001 ‘**’ 0.01 ‘*’ 0.05 ‘.’ 0.1 ‘ ’ 1
##
## R-sq.(adj) =   0.15
## lmer.REML = 4.0575e+05  Scale est. = 1431      n = 33350
```

Next is the model with depression-sleep interaction.

```
##
## Family: gaussian
## Link function: identity
##
## Formula:
## value ~ sex + site + s(age_z, k = 10, bs = "cr") + s(sleep_z,
##     k = 5, bs = "cr") + icv + depression + depression:sleep_z
## <environment: 0x55cffba79778>
##
## Parametric coefficients:
##                     Estimate Std. Error t value Pr(>|t|)
## (Intercept)         542.3891     7.5911  71.451  < 2e-16 ***
## sexmale             -17.5966     1.6080 -10.943  < 2e-16 ***
## siteousAvanto      -101.6011    17.7269  -5.731 1.00e-08 ***
## siteousPrisma       -30.3718    14.7342  -2.061   0.0393 *
## siteousSkyra       -101.7655    11.4490  -8.889  < 2e-16 ***
## siteUCAM            -47.9564    10.3564  -4.631 3.66e-06 ***
## siteUKB             -10.1418     7.4776  -1.356   0.1750
## siteUmU             -20.9872    10.4498  -2.008   0.0446 *
```

```
## icv                  31.7990     0.8269  38.457  < 2e-16 ***
## depression           -3.3986     5.7046  -0.596   0.5513
## depression:sleep_z   -7.6053     4.1600  -1.828   0.0675 .
## ---
## Signif. codes:  0 '***' 0.001 '**' 0.01 '*' 0.05 '.' 0.1 ' ' 1
##
## Approximate significance of smooth terms:
##              edf Ref.df       F p-value
## s(age_z)   5.557  5.557 592.251  <2e-16 ***
## s(sleep_z) 1.000  1.000   2.561    0.11
## ---
## Signif. codes:  0 '***' 0.001 '**' 0.01 '*' 0.05 '.' 0.1 ' ' 1
##
## R-sq.(adj) =   0.15
## lmer.REML = 4.0574e+05  Scale est. = 1431.1    n = 33350
```

The plot below shows the sleep-volume curve for the original model and for the model with main effects of SES.

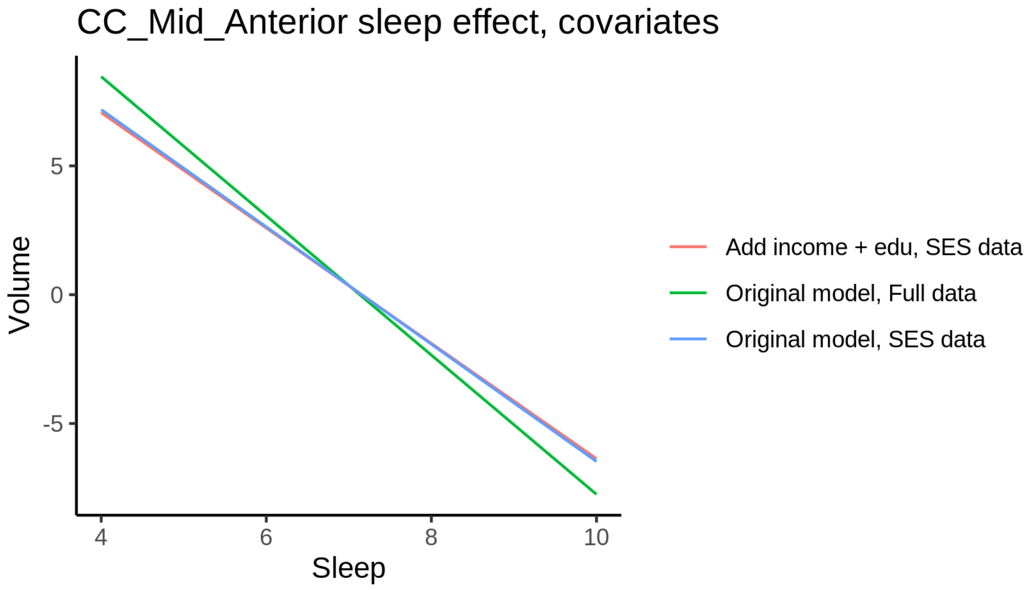


The plot below shows the sleep-volume curve for the original model and for the model with main effects of BMI.

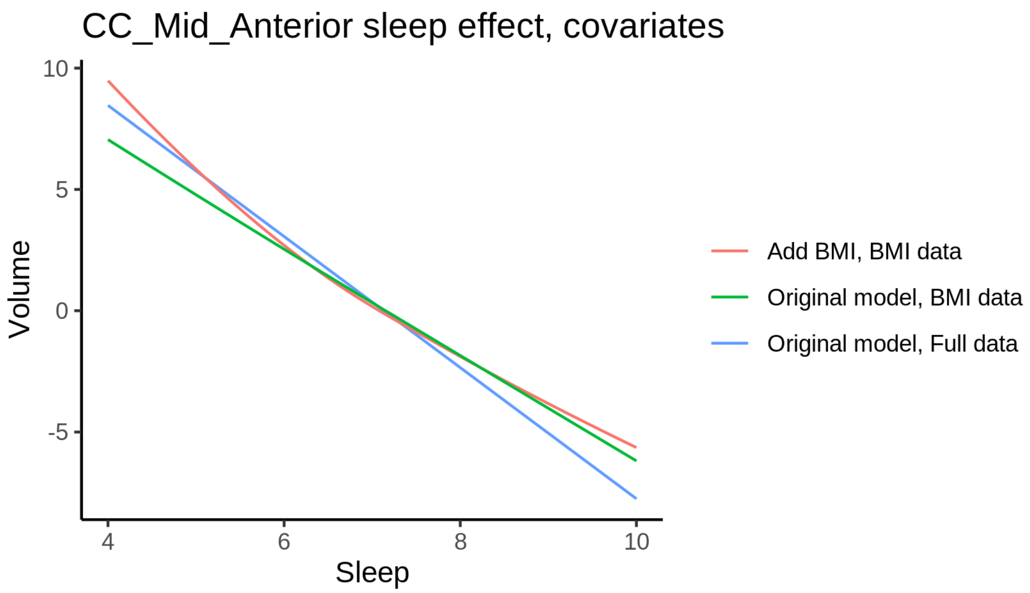


The plot below shows the sleep-volume curve for the original model and for the model with main effects of depression.

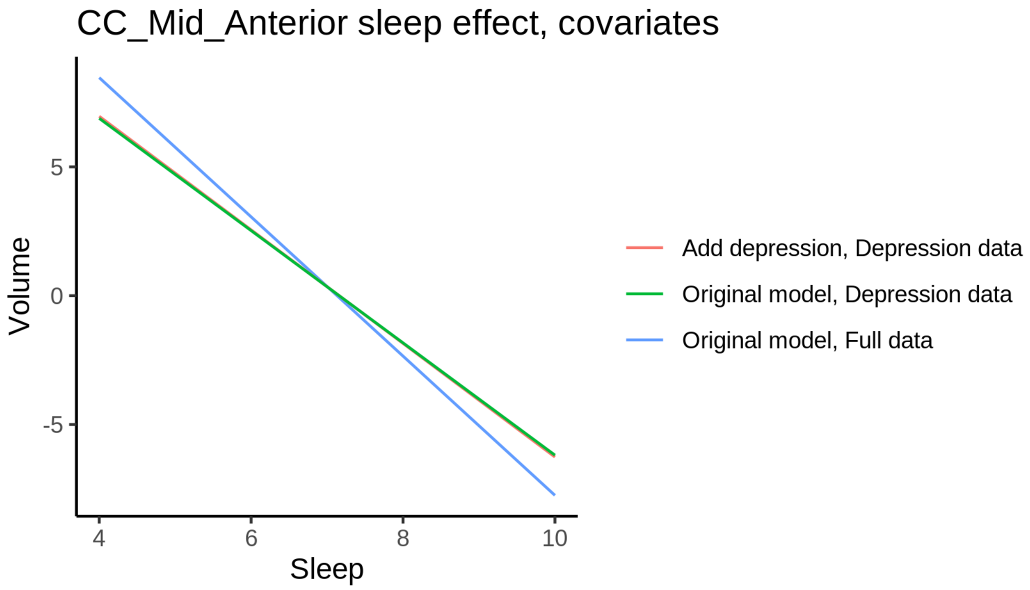


### CC_Mid_Posterior

#### Descriptive statistics

| Study | Observations | Unique IDs | Mean age | Age range |
|---|---|---|---|---|
| HCP | 974 | 974 | 28.8 | 22 - 37 |
| MPIB | 677 | 391 | 63.1 | 24 - 83 |
| UB | 113 | 39 | 70.9 | 64 - 81 |
| UCAM | 884 | 632 | 55.1 | 20 - 88 |

| Study | Observations | Unique IDs | Mean age | Age range |
|---|---|---|---|---|
| UiO | 1475 | 803 | 49.4 | 20 - 89 |
| UKB | 45944 | 43120 | 64.5 | 45 - 83 |
| UmU | 423 | 284 | 62.3 | 25 - 85 |
| UOXF | 769 | 769 | 69.8 | 60 - 85 |

##### Spaghetti plot

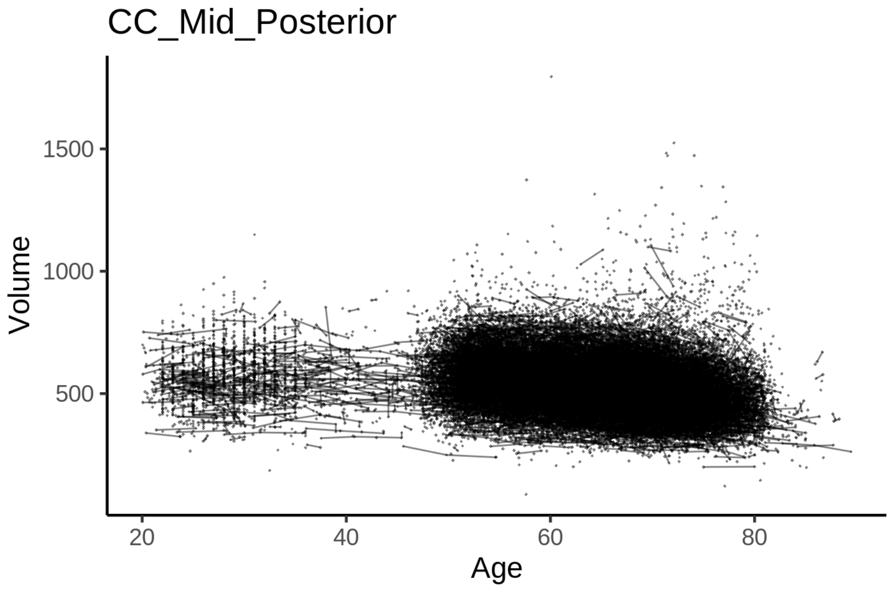


##### Model outputs

Model without sleep term

```
##
## Family: gaussian
## Link function: identity
##
## Formula:
## value ~ sex + site + icv + s(age_z, k = 10, bs = "cr")
## <environment: 0x55cf3c511450>
##
## Parametric coefficients:
##                Estimate Std. Error t value Pr(>|t|)
## (Intercept)   526.4686     5.6751  92.768  < 2e-16 ***
## sexmale       -18.4574     1.1772 -15.679  < 2e-16 ***
## siteMPIB       14.6600     7.3209   2.002   0.0452 *
## siteousAvanto -60.7977     5.8401 -10.410  < 2e-16 ***
## siteousPrisma -21.6656     9.9150  -2.185   0.0289 *
## siteousSkyra  -76.2374     5.5681 -13.692  < 2e-16 ***
## siteUB        -44.2611    17.2887  -2.560   0.0105 *
## siteUCAM      -29.2657     6.3345  -4.620 3.85e-06 ***
## siteUKB        23.5889     5.8171   4.055 5.02e-05 ***
## siteUmU       -14.3191     8.1584  -1.755   0.0792 .
```

```
## siteUOXF       -6.7178     6.9212  -0.971   0.3317
## icv            27.3316     0.5915  46.204  < 2e-16 ***
## ---
## Signif. codes:  0 '***' 0.001 '**' 0.01 '*' 0.05 '.' 0.1 ' ' 1
##
## Approximate significance of smooth terms:
##            edf Ref.df     F p-value
## s(age_z) 6.555  6.555 461.6  <2e-16 ***
## ---
## Signif. codes:  0 '***' 0.001 '**' 0.01 '*' 0.05 '.' 0.1 ' ' 1
##
## R-sq.(adj) =  0.124
## lmer.REML = 6.1285e+05  Scale est. = 737.18    n = 51259
```

Model with only main effects of age and sleep

```
##
## Family: gaussian
## Link function: identity
##
## Formula:
## value ~ sex + site + icv + s(age_z, k = 10, bs = "cr") + s(sleep_z,
##     k = 5, bs = "cr")
## <environment: 0x55cf3c511450>
##
## Parametric coefficients:
##               Estimate Std. Error t value Pr(>|t|)
## (Intercept)    526.279      5.680  92.652  < 2e-16 ***
## sexmale        -18.473      1.177 -15.692  < 2e-16 ***
## siteMPIB        14.983      7.325   2.045   0.0408 *
## siteousAvanto  -60.735      5.842 -10.397  < 2e-16 ***
## siteousPrisma  -21.627      9.915  -2.181   0.0292 *
## siteousSkyra   -76.186      5.570 -13.678  < 2e-16 ***
## siteUB         -44.088     17.288  -2.550   0.0108 *
## siteUCAM       -29.113      6.335  -4.596 4.33e-06 ***
## siteUKB         23.794      5.823   4.086 4.39e-05 ***
## siteUmU        -13.633      8.177  -1.667   0.0955 .
## siteUOXF        -6.634      6.922  -0.958   0.3378
## icv             27.331      0.592  46.168  < 2e-16 ***
## ---
## Signif. codes:  0 '***' 0.001 '**' 0.01 '*' 0.05 '.' 0.1 ' ' 1
##
## Approximate significance of smooth terms:
##              edf Ref.df       F p-value
## s(age_z)   6.548  6.548 457.976  <2e-16 ***
## s(sleep_z) 1.715  1.715   2.383   0.212
## ---
## Signif. codes:  0 '***' 0.001 '**' 0.01 '*' 0.05 '.' 0.1 ' ' 1
##
## R-sq.(adj) =  0.124
## lmer.REML = 6.1285e+05  Scale est. = 737.23    n = 51259
```

Model with full interaction between age and sleep

```
##
## Family: gaussian
```

```
## Link function: identity
##
## Formula:
## value ~ sex + site + icv + t2(age_z, sleep_z, k = c(10, 4), bs = "cr")
## <environment: 0x55cf3c511450>
##
## Parametric coefficients:
##                Estimate Std. Error t value Pr(>|t|)
## (Intercept)     526.782      5.696  92.487  < 2e-16 ***
## sexmale         -18.513      1.180 -15.694  < 2e-16 ***
## siteMPIB         14.569      7.360   1.979  0.04777 *
## siteousAvanto   -61.119      5.871 -10.410  < 2e-16 ***
## siteousPrisma   -21.977      9.935  -2.212  0.02697 *
## siteousSkyra    -76.519      5.603 -13.657  < 2e-16 ***
## siteUB          -44.630     17.297  -2.580  0.00988 **
## siteUCAM        -29.563      6.367  -4.644 3.43e-06 ***
## siteUKB          23.294      5.838   3.990 6.61e-05 ***
## siteUmU         -14.124      8.199  -1.723  0.08496 .
## siteUOXF         -7.116      6.941  -1.025  0.30533
## icv              27.335      0.592  46.178  < 2e-16 ***
## ---
## Signif. codes:  0 ‘***’ 0.001 ‘**’ 0.01 ‘*’ 0.05 ‘.’ 0.1 ‘ ’ 1
##
## Approximate significance of smooth terms:
##                      edf Ref.df     F p-value
## t2(age_z,sleep_z) 10.94  10.94 10.27  <2e-16 ***
## ---
## Signif. codes:  0 ‘***’ 0.001 ‘**’ 0.01 ‘*’ 0.05 ‘.’ 0.1 ‘ ’ 1
##
## R-sq.(adj) =  0.124
## lmer.REML = 6.1285e+05  Scale est. = 737.09    n = 51259
```

#### Model comparison

`mod_no_sleep` refers to model without sleep term, `mod_no_interaction` refers to model with only main effect of sleep, and `mod_full` refers to model with a full interaction between age and sleep. This is a nested model comparison, and the p-value at a given line refers to comparing the model at the line to the model on the line above. Hence, significance implies that the more complicated model is supported on statistical grounds.

To be even more specific, the p-value on the second row tests whether there is an association between sleep and volume. The p-value on the third row tests whether this association depends on age.

```
## Data: NULL
## Models:
## mod_list$mod_no_sleep$mer: NULL
## mod_list$mod_no_interaction$mer: NULL
## mod_list$mod_full$mer: NULL
##                                 npar    AIC    BIC  logLik deviance  Chisq Df Pr(>Chisq)
## mod_list$mod_no_sleep$mer         16 612882 613024 -306425   612850
## mod_list$mod_no_interaction$mer   18 612885 613044 -306424   612849 1.6366  2     0.4412
## mod_list$mod_full$mer             20 612892 613069 -306426   612852 0.0000  2     1.0000
```

We chose the model based on the likelihood ratio test with 5 % significance level, which was `mod_no_sleep`.

##### Lifespan brain trajectory

The trajectory shown is from the chosen model `mod_no_sleep`.

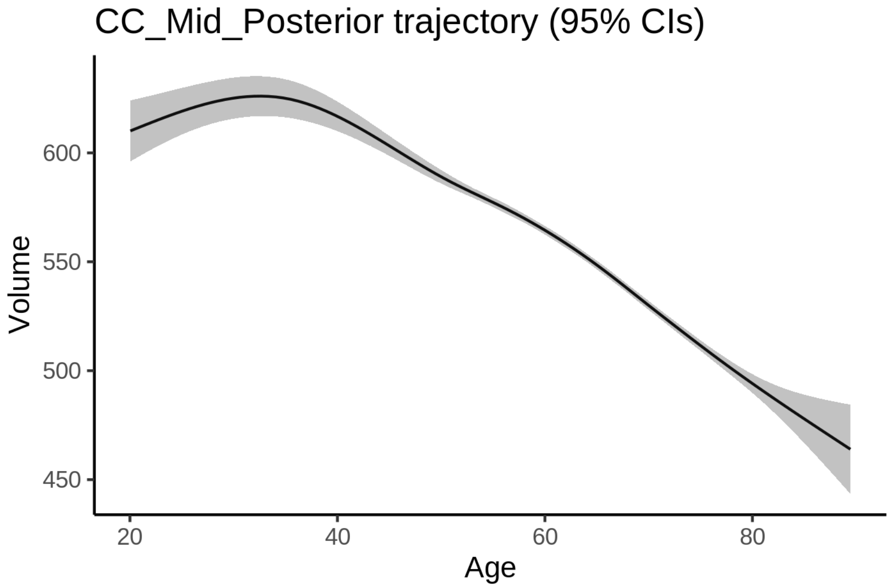


##### Effect of sleep

The chosen model did not include a sleep term, and hence we don't have any estimated effect of sleep.

We show the full interaction model for completeness, although it was not selected.

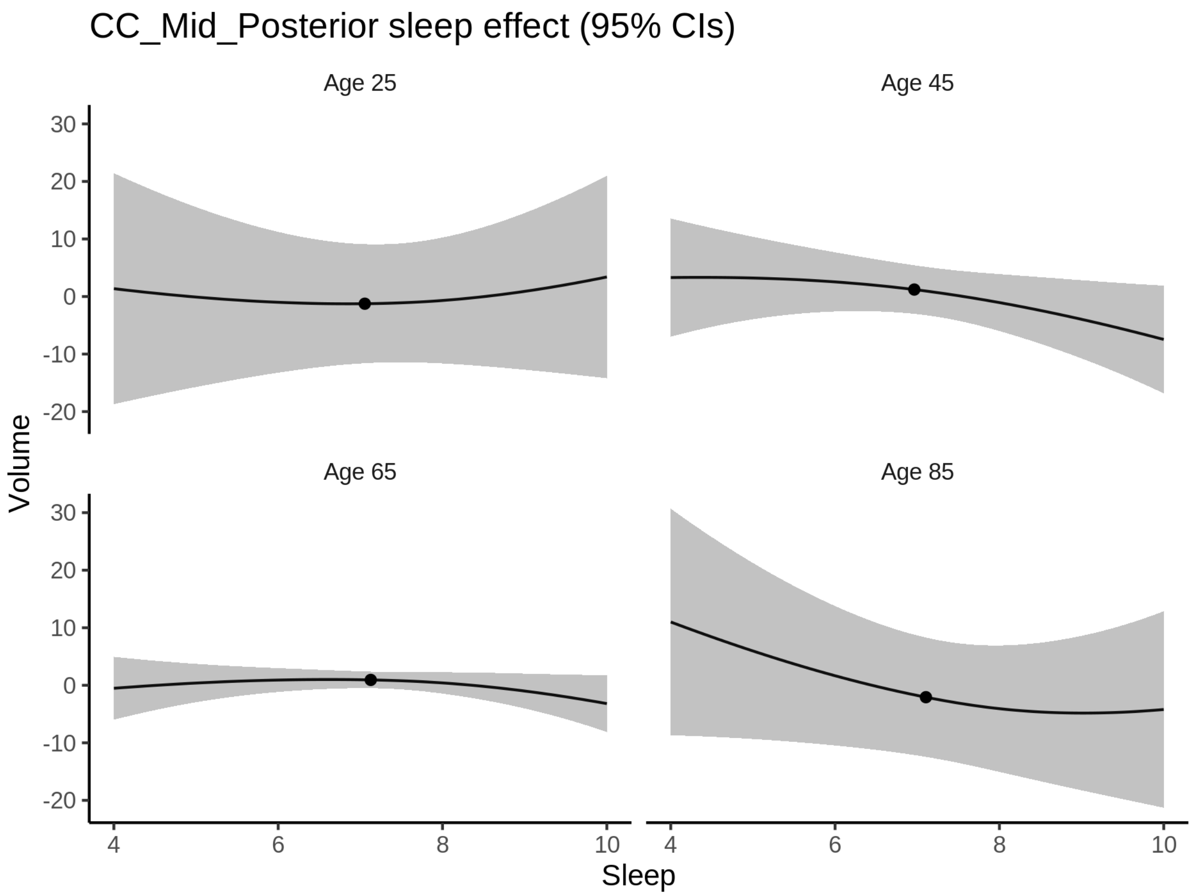


**Deviation from sleep associated with maximal volume**

Model with no sleep term was selected. No plots to show. (Although we can of course dig up the plots, which will be pretty flat).

**Comparison of mean sleep and sleep associated with maximum volume**

Nothing to show, as we did not find an association between sleep and volume.

### CC_Posterior

#### Descriptive statistics

| Study | Observations | Unique IDs | Mean age | Age range |
|---|---|---|---|---|
| HCP | 974 | 974 | 28.8 | 22 - 37 |
| MPIB | 675 | 390 | 63.1 | 24 - 83 |
| UB | 113 | 39 | 70.9 | 64 - 81 |
| UCAM | 878 | 630 | 54.9 | 20 - 88 |
| UiO | 1475 | 803 | 49.4 | 20 - 89 |
| UKB | 45941 | 43118 | 64.5 | 45 - 83 |

| Study | Observations | Unique IDs | Mean age | Age range |
|---|---|---|---|---|
| UmU | 421 | 283 | 62.3 | 25 - 85 |
| UOXF | 769 | 769 | 69.8 | 60 - 85 |

##### Spaghetti plot

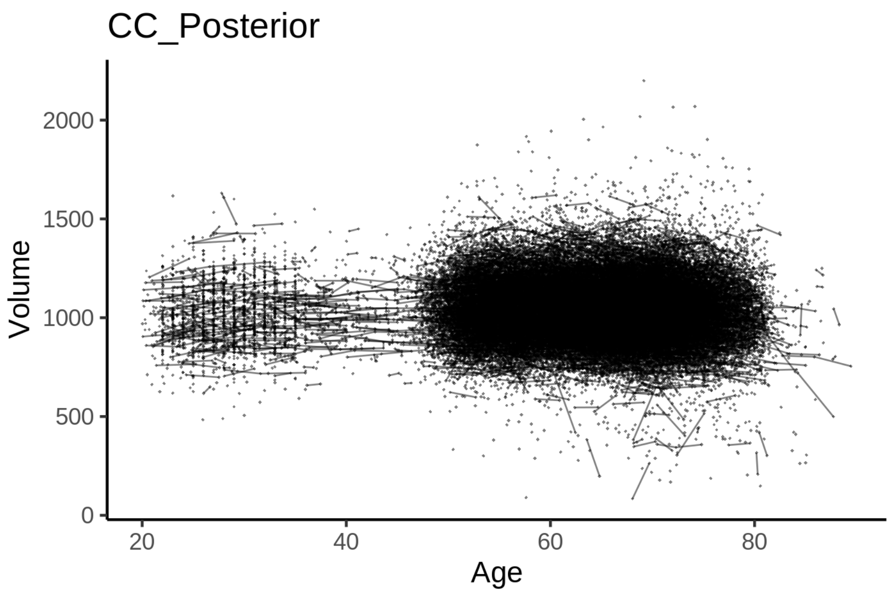


##### Model outputs

Model without sleep term

```
##
## Family: gaussian
## Link function: identity
##
## Formula:
## value ~ sex + site + icv + s(age_z, k = 10, bs = "cr")
## <environment: 0x55cf3be1b918>
##
## Parametric coefficients:
##                Estimate Std. Error t value Pr(>|t|)
## (Intercept)   1048.3788     8.0564 130.130  < 2e-16 ***
## sexmale        -36.4997     1.6833 -21.684  < 2e-16 ***
## siteMPIB        30.4711    10.4565   2.914  0.00357 **
## siteousAvanto  -60.0208     8.2027  -7.317 2.57e-13 ***
## siteousPrisma  -15.6531    13.7881  -1.135  0.25627
## siteousSkyra   -45.6734     7.9393  -5.753 8.83e-09 ***
## siteUB         -74.8581    24.8484  -3.013  0.00259 **
## siteUCAM       -46.9145     9.0358  -5.192 2.09e-07 ***
## siteUKB         14.2660     8.2558   1.728  0.08399 .
## siteUmU         -3.8288    11.6562  -0.328  0.74255
## siteUOXF       -25.0382     9.8471  -2.543  0.01100 *
## icv             65.5434     0.8446  77.604  < 2e-16 ***
```

```
## ---
## Signif. codes:  0 ‘***’ 0.001 ‘**’ 0.01 ‘*’ 0.05 ‘.’ 0.1 ‘ ’ 1
##
## Approximate significance of smooth terms:
##            edf Ref.df     F p-value
## s(age_z) 7.792  7.792 15.94  <2e-16 ***
## ---
## Signif. codes:  0 ‘***’ 0.001 ‘**’ 0.01 ‘*’ 0.05 ‘.’ 0.1 ‘ ’ 1
##
## R-sq.(adj) =  0.136
## lmer.REML = 6.4804e+05  Scale est. = 1085.1    n = 51246
```

Model with only main effects of age and sleep

```
##
## Family: gaussian
## Link function: identity
##
## Formula:
## value ~ sex + site + icv + s(age_z, k = 10, bs = "cr") + s(sleep_z,
##     k = 5, bs = "cr")
## <environment: 0x55cf3be1b918>
##
## Parametric coefficients:
##                Estimate Std. Error t value Pr(>|t|)
## (Intercept)   1048.9766     8.0639 130.083  < 2e-16 ***
## sexmale        -36.5026     1.6832 -21.686  < 2e-16 ***
## siteMPIB        30.3587    10.4620   2.902  0.00371 **
## siteousAvanto  -60.4880     8.2048  -7.372 1.70e-13 ***
## siteousPrisma  -15.9920    13.7876  -1.160  0.24610
## siteousSkyra   -46.2223     7.9418  -5.820 5.92e-09 ***
## siteUB         -74.4149    24.8452  -2.995  0.00274 **
## siteUCAM       -46.9243     9.0363  -5.193 2.08e-07 ***
## siteUKB         13.6367     8.2641   1.650  0.09893 .
## siteUmU         -3.9638    11.6816  -0.339  0.73437
## siteUOXF       -25.3487     9.8474  -2.574  0.01005 *
## icv             65.4342     0.8454  77.400  < 2e-16 ***
## ---
## Signif. codes:  0 ‘***’ 0.001 ‘**’ 0.01 ‘*’ 0.05 ‘.’ 0.1 ‘ ’ 1
##
## Approximate significance of smooth terms:
##              edf Ref.df      F p-value
## s(age_z)   7.785  7.785 15.632  <2e-16 ***
## s(sleep_z) 2.974  2.974  3.484  0.0112 *
## ---
## Signif. codes:  0 ‘***’ 0.001 ‘**’ 0.01 ‘*’ 0.05 ‘.’ 0.1 ‘ ’ 1
##
## R-sq.(adj) =  0.136
## lmer.REML = 6.4804e+05  Scale est. = 1085.3    n = 51246
```

Model with full interaction between age and sleep

```
##
## Family: gaussian
## Link function: identity
##
```

```
## Formula:
## value ~ sex + site + icv + t2(age_z, sleep_z, k = c(10, 4), bs = "cr")
## <environment: 0x55cf3be1b918>
##
## Parametric coefficients:
##                Estimate Std. Error t value Pr(>|t|)
## (Intercept)   1048.9976     8.0080 130.994  < 2e-16 ***
## sexmale        -36.5047     1.6868 -21.641  < 2e-16 ***
## siteMPIB        30.1511    10.4688   2.880  0.00398 **
## siteousAvanto  -60.5762     8.2239  -7.366 1.79e-13 ***
## siteousPrisma  -16.3358    13.8146  -1.183  0.23701
## siteousSkyra   -46.2287     7.9634  -5.805 6.47e-09 ***
## siteUB         -75.0788    24.8325  -3.023  0.00250 **
## siteUCAM       -47.2699     9.0384  -5.230 1.70e-07 ***
## siteUKB         13.6255     8.2020   1.661  0.09667 .
## siteUmU         -4.2010    11.6631  -0.360  0.71870
## siteUOXF       -25.6016     9.8113  -2.609  0.00907 **
## icv             65.4731     0.8451  77.475  < 2e-16 ***
## ---
## Signif. codes:  0 ‘***’ 0.001 ‘**’ 0.01 ‘*’ 0.05 ‘.’ 0.1 ‘ ’ 1
##
## Approximate significance of smooth terms:
##                     edf Ref.df    F p-value
## t2(age_z,sleep_z) 12.83  12.83 9.28  <2e-16 ***
## ---
## Signif. codes:  0 ‘***’ 0.001 ‘**’ 0.01 ‘*’ 0.05 ‘.’ 0.1 ‘ ’ 1
##
## R-sq.(adj) =  0.136
## lmer.REML = 6.4804e+05  Scale est. = 1084.8    n = 51246
```

#### Model comparison

`mod_no_sleep` refers to model without sleep term, `mod_no_interaction` refers to model with only main effect of sleep, and `mod_full` refers to model with a full interaction between age and sleep. This is a nested model comparison, and the p-value at a given line refers to comparing the model at the line to the model on the line above. Hence, significance implies that the more complicated model is supported on statistical grounds.

To be even more specific, the p-value on the second row tests whether there is an association between sleep and volume. The p-value on the third row tests whether this association depends on age.

```
## Data: NULL
## Models:
## mod_list$mod_no_sleep$mer: NULL
## mod_list$mod_no_interaction$mer: NULL
## mod_list$mod_full$mer: NULL
##                                 npar    AIC    BIC  logLik deviance  Chisq Df Pr(>Chisq)
## mod_list$mod_no_sleep$mer         16 648074 648215 -324021   648042
## mod_list$mod_no_interaction$mer   18 648071 648231 -324018   648035 6.1887  2     0.0453 *
## mod_list$mod_full$mer             20 648080 648257 -324020   648040 0.0000  2     1.0000
## ---
## Signif. codes:  0 ‘***’ 0.001 ‘**’ 0.01 ‘*’ 0.05 ‘.’ 0.1 ‘ ’ 1
```

We chose the model based on the likelihood ratio test with 5 % significance level, which was `mod_no_interaction`.

##### Lifespan brain trajectory

The trajectory shown is from the chosen model `mod_no_interaction`.

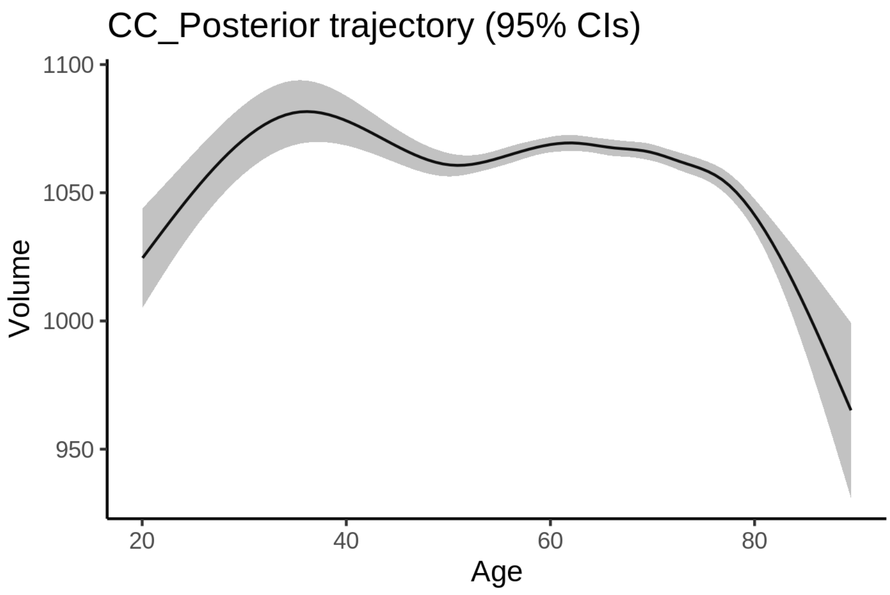


##### Effect of sleep

The chosen model only included the main effect of sleep, and hence the effect does not vary with age. The black dot shows the average sleep duration across all ages in the sample.

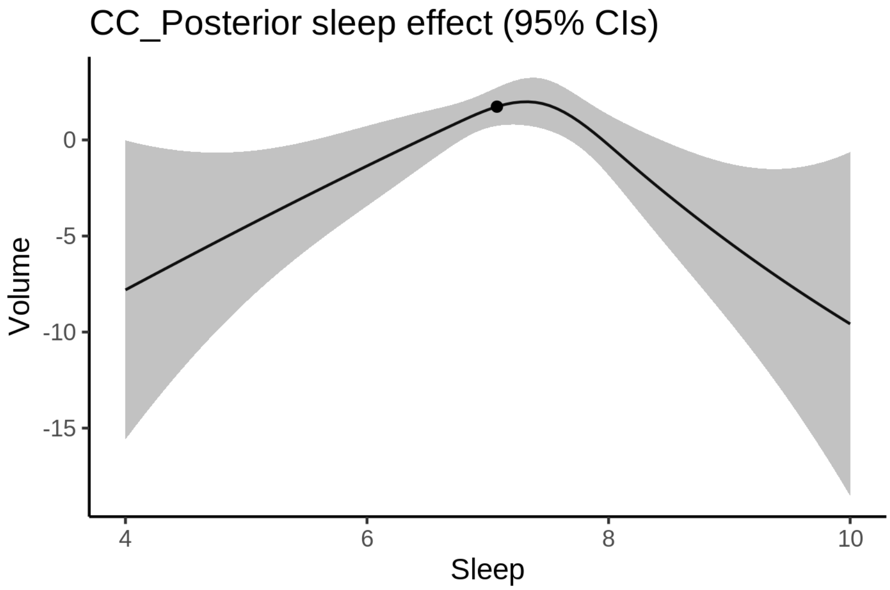


We also show the full interaction model for completeness, although it was not selected.

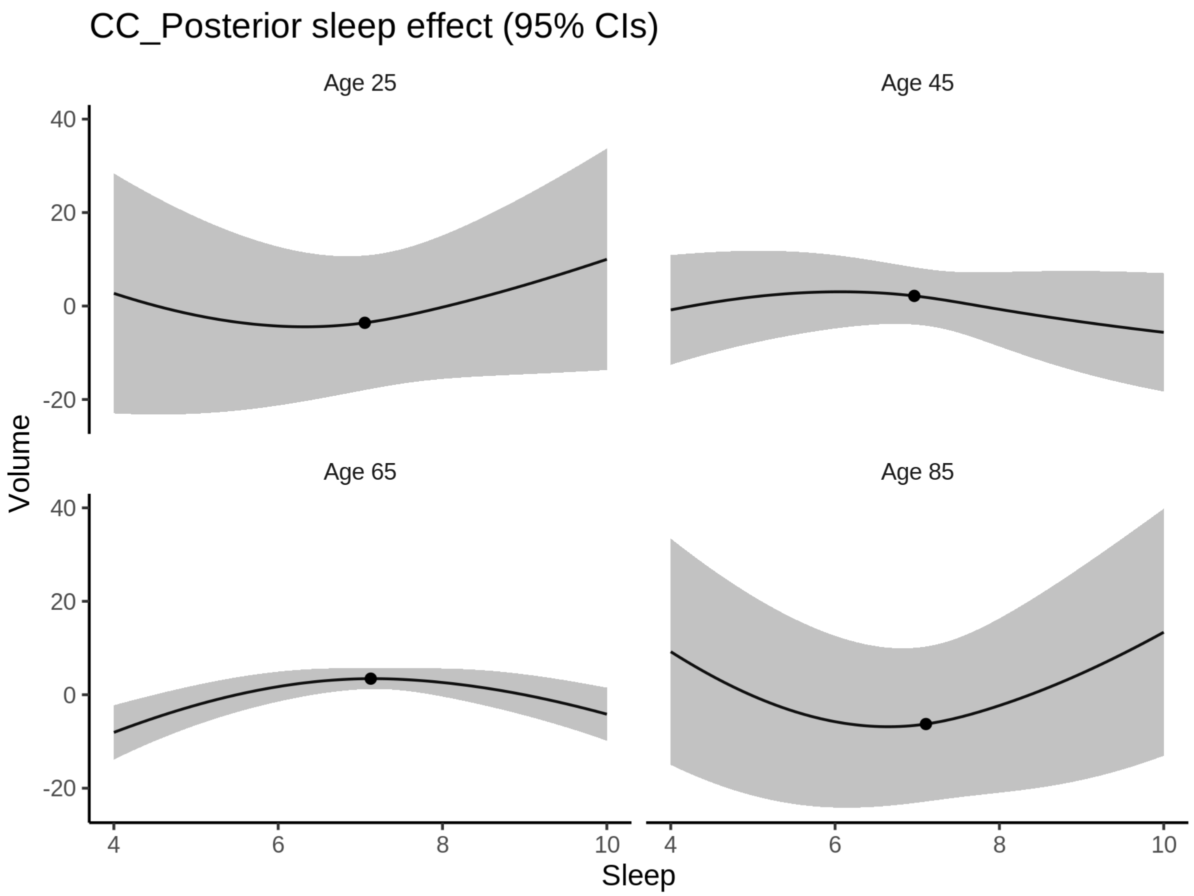


**Deviation from sleep associated with maximal volume**

Model with only main effect of sleep was chosen, so we show it for all ages at once. Maximum volume is attained at 7.3 hours of sleep. The percentage values in the plot are calculated as follows: The maximum at 100 % refers to a person at an arbitrary age with a sleep duration associated with maximum volume. For a female, this volume is 1055 and for a male it is 1018. The other percentage values show how large the expected volume is for someone with other sleep durations. For example, 99 % implies a 1 % reduction.

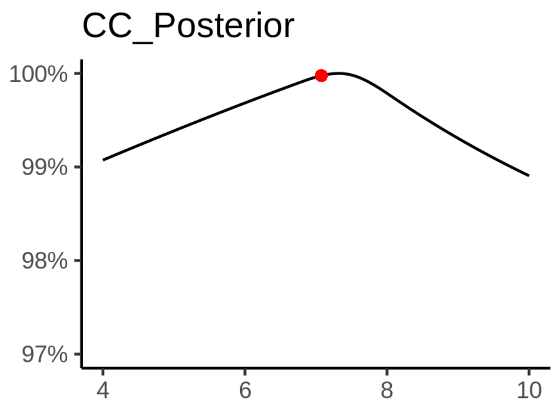

**Comparison of mean sleep and sleep associated with maximum volume**

A 95 % confidence interval for the sleep associated with maximum volume is [6.85, 7.58].

The plot below compares average sleep to the sleep associated with maximum volume.

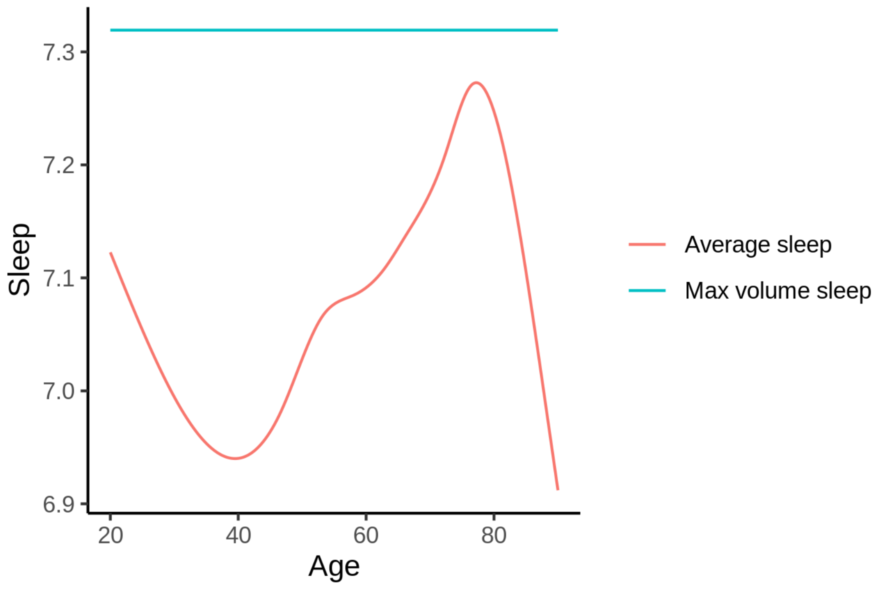


The next plot shows the difference between average sleep and sleep associated with maximum volume. The shaded region is a 95 % confidence interval.

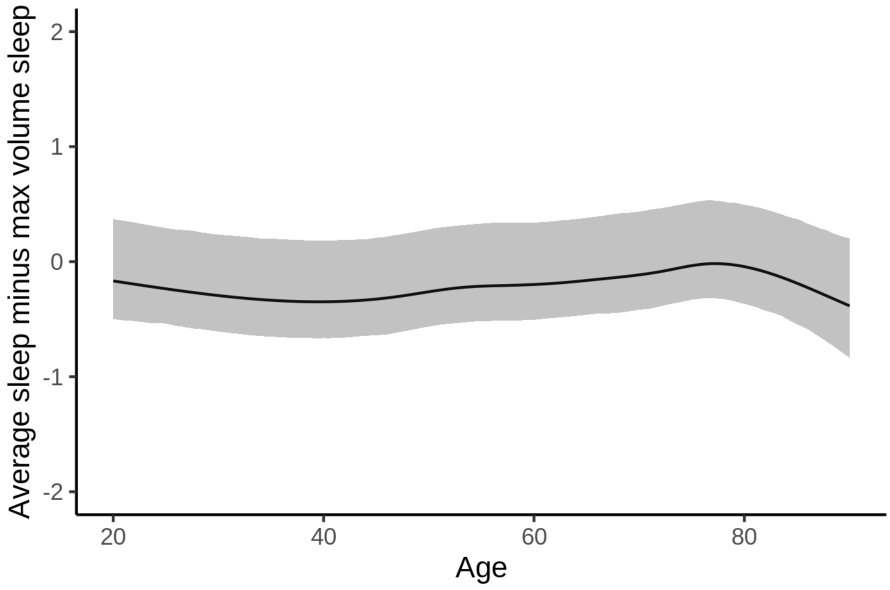


The next plot shows the probability that the sleep duration associated with maximum volume is longer than the average sleep duration, as a function of age. Probility below .05 can be interpreted as evidence that the sleep associated with maximum volume is shorter than the mean sleep, and probability above .95 can be interpreted the opposite way.

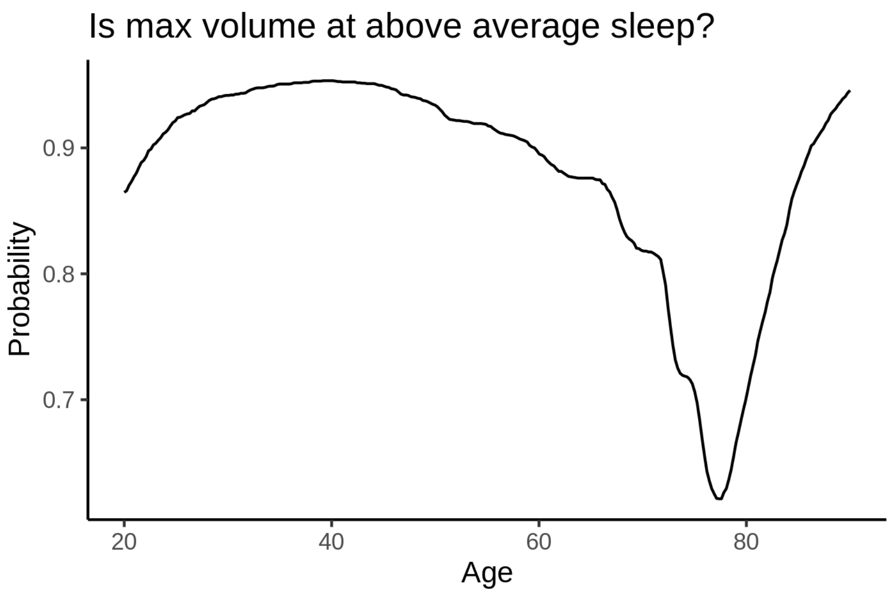


##### Controling for covariates

Below is the output for a model in which we only include data with income and education.

```
##
## Family: gaussian
## Link function: identity
##
## Formula:
## value ~ sex + site + s(age_z, k = 10, bs = "cr") + s(sleep_z,
##     k = 5, bs = "cr") + icv
## <environment: 0x55cf38300600>
##
## Parametric coefficients:
##                Estimate Std. Error t value Pr(>|t|)
## (Intercept)    1114.748     19.322  57.693  < 2e-16 ***
## sexmale         -40.777      2.130 -19.148  < 2e-16 ***
## siteousAvanto -122.154     22.688  -5.384 7.33e-08 ***
## siteousPrisma  -57.543     23.653  -2.433  0.01499 *
## siteousSkyra  -102.896     20.993  -4.901 9.56e-07 ***
## siteUKB         -49.960     19.279  -2.591  0.00956 **
## siteUOXF        -71.296     21.715  -3.283  0.00103 **
## icv              69.076      1.092  63.253  < 2e-16 ***
## ---
## Signif. codes:  0 ‘***’ 0.001 ‘**’ 0.01 ‘*’ 0.05 ‘.’ 0.1 ‘ ’ 1
##
## Approximate significance of smooth terms:
##              edf Ref.df     F  p-value
## s(age_z)   4.507  4.507 4.923 0.000424 ***
## s(sleep_z) 2.472  2.472 2.181 0.210885
## ---
## Signif. codes:  0 ‘***’ 0.001 ‘**’ 0.01 ‘*’ 0.05 ‘.’ 0.1 ‘ ’ 1
##
```

```
## R-sq.(adj) =  0.137
## lmer.REML = 3.9407e+05  Scale est. = 1199.7    n = 31165
```

Below is the output for a model in which we control for the main effects of income and education.

```
##
## Family: gaussian
## Link function: identity
##
## Formula:
## value ~ sex + site + s(age_z, k = 10, bs = "cr") + s(sleep_z,
##     k = 5, bs = "cr") + icv + income_scaled + education_scaled
## <environment: 0x55cf38300600>
##
## Parametric coefficients:
##                  Estimate Std. Error t value Pr(>|t|)
## (Intercept)      1109.676     19.443  57.072  < 2e-16 ***
## sexmale           -40.670      2.135 -19.045  < 2e-16 ***
## siteousAvanto    -122.396     22.694  -5.393 6.97e-08 ***
## siteousPrisma     -58.053     23.653  -2.454  0.01412 *
## siteousSkyra     -103.217     20.998  -4.916 8.90e-07 ***
## siteUKB           -50.363     19.289  -2.611  0.00903 **
## siteUOXF          -69.902     21.726  -3.217  0.00129 **
## icv                68.761      1.101  62.444  < 2e-16 ***
## income_scaled       2.012      2.536   0.793  0.42755
## education_scaled    5.734      3.001   1.910  0.05611 .
## ---
## Signif. codes:  0 ‘***’ 0.001 ‘**’ 0.01 ‘*’ 0.05 ‘.’ 0.1 ‘ ’ 1
##
## Approximate significance of smooth terms:
##              edf Ref.df     F  p-value
## s(age_z)   4.531  4.531 4.798 0.000527 ***
## s(sleep_z) 2.338  2.338 1.803 0.272701
## ---
## Signif. codes:  0 ‘***’ 0.001 ‘**’ 0.01 ‘*’ 0.05 ‘.’ 0.1 ‘ ’ 1
##
## R-sq.(adj) =  0.137
## lmer.REML = 3.9406e+05  Scale est. = 1200.2    n = 31165
```

We also included interaction effects between sleep duration and education and income, in another model. The output is shown below, and the interaction terms are `income_scaled:sleep_z` and `education_scaled:sleep_z`.

```
##
## Family: gaussian
## Link function: identity
##
## Formula:
## value ~ sex + site + s(age_z, k = 10, bs = "cr") + s(sleep_z,
##     k = 5, bs = "cr") + icv + income_scaled + education_scaled +
##     income_scaled:sleep_z + education_scaled:sleep_z
## <environment: 0x55cf38300600>
##
## Parametric coefficients:
##                          Estimate Std. Error t value Pr(>|t|)
## (Intercept)              1109.514     19.445  57.060  < 2e-16 ***
```

```
## sexmale                   -40.602      2.137 -19.000  < 2e-16 ***
## siteousAvanto            -122.209     22.695  -5.385 7.30e-08 ***
## siteousPrisma             -57.703     23.656  -2.439  0.01472 *
## siteousSkyra             -103.036     20.999  -4.907 9.31e-07 ***
## siteUKB                   -50.177     19.291  -2.601  0.00930 **
## siteUOXF                  -69.638     21.732  -3.204  0.00135 **
## icv                        68.747      1.101  62.424  < 2e-16 ***
## income_scaled               1.951      2.537   0.769  0.44194
## education_scaled            5.788      3.002   1.928  0.05384 .
## income_scaled:sleep_z       1.514      2.524   0.600  0.54854
## education_scaled:sleep_z   -2.824      2.912  -0.970  0.33212
## ---
## Signif. codes:  0 ‘***’ 0.001 ‘**’ 0.01 ‘*’ 0.05 ‘.’ 0.1 ‘ ’ 1
##
## Approximate significance of smooth terms:
##              edf Ref.df     F  p-value
## s(age_z)   4.524  4.524 4.806 0.000522 ***
## s(sleep_z) 2.291  2.291 1.540 0.336913
## ---
## Signif. codes:  0 ‘***’ 0.001 ‘**’ 0.01 ‘*’ 0.05 ‘.’ 0.1 ‘ ’ 1
##
## R-sq.(adj) =  0.137
## lmer.REML = 3.9405e+05  Scale est. = 1200.2    n = 31165
```

We did the same controling for BMI. Below is the model with no covariates but only keeping data with BMI.

```
##
## Family: gaussian
## Link function: identity
##
## Formula:
## value ~ sex + site + s(age_z, k = 10, bs = "cr") + s(sleep_z,
##     k = 5, bs = "cr") + icv
## <environment: 0x55cf356dbea8>
##
## Parametric coefficients:
##               Estimate Std. Error t value Pr(>|t|)
## (Intercept)   1002.621     11.899  84.260  < 2e-16 ***
## sexmale        -40.008      2.071 -19.318  < 2e-16 ***
## siteousPrisma   39.250     15.941   2.462 0.013813 *
## siteousSkyra    12.307      8.487   1.450 0.147064
## siteUCAM         8.293     12.728   0.652 0.514713
## siteUKB         61.681     12.017   5.133 2.87e-07 ***
## siteUmU         49.935     14.695   3.398 0.000679 ***
## icv             68.913      1.065  64.693  < 2e-16 ***
## ---
## Signif. codes:  0 ‘***’ 0.001 ‘**’ 0.01 ‘*’ 0.05 ‘.’ 0.1 ‘ ’ 1
##
## Approximate significance of smooth terms:
##              edf Ref.df     F  p-value
## s(age_z)   5.760  5.760 7.172 1.42e-06 ***
## s(sleep_z) 2.487  2.487 2.198     0.24
## ---
## Signif. codes:  0 ‘***’ 0.001 ‘**’ 0.01 ‘*’ 0.05 ‘.’ 0.1 ‘ ’ 1
##
```

```
## R-sq.(adj) =  0.138
## lmer.REML = 4.2197e+05  Scale est. = 1117.3    n = 33414
```

Below is the model output with main effect.

```
##
## Family: gaussian
## Link function: identity
##
## Formula:
## value ~ sex + site + s(age_z, k = 10, bs = "cr") + s(sleep_z,
##     k = 5, bs = "cr") + icv + bmi
## <environment: 0x55cf356dbea8>
##
## Parametric coefficients:
##                Estimate Std. Error t value Pr(>|t|)
## (Intercept)   1044.5577    12.9321  80.772  < 2e-16 ***
## sexmale        -38.6815     2.0750 -18.641  < 2e-16 ***
## siteousPrisma   40.5781    15.9315   2.547 0.010869 *
## siteousSkyra    12.4422     8.4856   1.466 0.142585
## siteUCAM         9.1921    12.7191   0.723 0.469868
## siteUKB         63.0705    12.0090   5.252 1.51e-07 ***
## siteUmU         51.6763    14.6845   3.519 0.000434 ***
## icv             69.1821     1.0643  65.000  < 2e-16 ***
## bmi             -1.6664     0.2018  -8.257  < 2e-16 ***
## ---
## Signif. codes:  0 ‘***’ 0.001 ‘**’ 0.01 ‘*’ 0.05 ‘.’ 0.1 ‘ ’ 1
##
## Approximate significance of smooth terms:
##              edf Ref.df     F  p-value
## s(age_z)   5.739  5.739 7.742 1.14e-06 ***
## s(sleep_z) 1.927  1.927 0.924    0.452
## ---
## Signif. codes:  0 ‘***’ 0.001 ‘**’ 0.01 ‘*’ 0.05 ‘.’ 0.1 ‘ ’ 1
##
## R-sq.(adj) =   0.14
## lmer.REML = 4.219e+05  Scale est. = 1117.1    n = 33414
```

Next is the model with BMI-sleep interaction.

```
##
## Family: gaussian
## Link function: identity
##
## Formula:
## value ~ sex + site + s(age_z, k = 10, bs = "cr") + s(sleep_z,
##     k = 5, bs = "cr") + icv + bmi + bmi:sleep_z
## <environment: 0x55cf356dbea8>
##
## Parametric coefficients:
##                Estimate Std. Error t value Pr(>|t|)
## (Intercept)   1044.8182    12.9394  80.747  < 2e-16 ***
## sexmale        -38.6864     2.0751 -18.643  < 2e-16 ***
## siteousPrisma   40.5780    15.9316   2.547 0.010870 *
## siteousSkyra    12.4247     8.4857   1.464 0.143153
## siteUCAM         9.0982    12.7202   0.715 0.474455
```

```
## siteUKB          62.9864    12.0103   5.244 1.58e-07 ***
## siteUmU          51.5948    14.6854   3.513 0.000443 ***
## icv              69.1778     1.0644  64.994  < 2e-16 ***
## bmi              -1.6709     0.2019  -8.274  < 2e-16 ***
## bmi:sleep_z      -0.1183     0.1912  -0.619 0.535940
## ---
## Signif. codes:  0 ‘***’ 0.001 ‘**’ 0.01 ‘*’ 0.05 ‘.’ 0.1 ‘ ’ 1
##
## Approximate significance of smooth terms:
##               edf Ref.df     F  p-value
## s(age_z)    5.744  5.744 7.757 1.12e-06 ***
## s(sleep_z) 1.907  1.907 0.600    0.484
## ---
## Signif. codes:  0 ‘***’ 0.001 ‘**’ 0.01 ‘*’ 0.05 ‘.’ 0.1 ‘ ’ 1
##
## R-sq.(adj) =   0.14
## lmer.REML = 4.219e+05  Scale est. = 1117.1    n = 33414
```

We did the same controling for depression. Below is the model with no covariates but only keeping data with depression.

```
##
## Family: gaussian
## Link function: identity
##
## Formula:
## value ~ sex + site + s(age_z, k = 10, bs = "cr") + s(sleep_z,
##     k = 5, bs = "cr") + icv
## <environment: 0x55cffafa0890>
##
## Parametric coefficients:
##                Estimate Std. Error t value Pr(>|t|)
## (Intercept)    1058.021      9.955 106.278  < 2e-16 ***
## sexmale         -39.435      2.081 -18.950  < 2e-16 ***
## siteousAvanto  -59.323     19.223  -3.086 0.002030 **
## siteousPrisma   -7.190     18.555  -0.388 0.698374
## siteousSkyra   -35.224     14.700  -2.396 0.016576 *
## siteUCAM       -45.863     13.587  -3.376 0.000738 ***
## siteUKB          6.095      9.846   0.619 0.535889
## siteUmU         -7.296     13.328  -0.547 0.584078
## icv             68.492      1.069  64.085  < 2e-16 ***
## ---
## Signif. codes:  0 ‘***’ 0.001 ‘**’ 0.01 ‘*’ 0.05 ‘.’ 0.1 ‘ ’ 1
##
## Approximate significance of smooth terms:
##               edf Ref.df     F  p-value
## s(age_z)    5.599  5.599 6.141 3.21e-05 ***
## s(sleep_z) 2.597  2.597 2.587    0.174
## ---
## Signif. codes:  0 ‘***’ 0.001 ‘**’ 0.01 ‘*’ 0.05 ‘.’ 0.1 ‘ ’ 1
##
## R-sq.(adj) =  0.138
## lmer.REML = 4.2055e+05  Scale est. = 1083.2    n = 33324
```

Below is the model output with main effect.

```
##
## Family: gaussian
## Link function: identity
##
## Formula:
## value ~ sex + site + s(age_z, k = 10, bs = "cr") + s(sleep_z,
##     k = 5, bs = "cr") + icv + depression
## <environment: 0x55cffafa0890>
##
## Parametric coefficients:
##               Estimate Std. Error t value Pr(>|t|)
## (Intercept)   1058.712      9.998 105.895  < 2e-16 ***
## sexmale        -39.508      2.083 -18.965  < 2e-16 ***
## siteousAvanto  -58.948     19.227  -3.066 0.002172 **
## siteousPrisma   -6.852     18.559  -0.369 0.711955
## siteousSkyra   -34.806     14.708  -2.366 0.017965 *
## siteUCAM       -45.692     13.588  -3.363 0.000773 ***
## siteUKB          5.813      9.854   0.590 0.555241
## siteUmU         -5.184     13.611  -0.381 0.703309
## icv             68.503      1.069  64.091  < 2e-16 ***
## depression      -5.674      7.446  -0.762 0.446026
## ---
## Signif. codes:  0 '***' 0.001 '**' 0.01 '*' 0.05 '.' 0.1 ' ' 1
##
## Approximate significance of smooth terms:
##              edf Ref.df     F  p-value
## s(age_z)   5.586  5.586 6.112 3.45e-05 ***
## s(sleep_z) 2.530  2.530 2.347    0.207
## ---
## Signif. codes:  0 '***' 0.001 '**' 0.01 '*' 0.05 '.' 0.1 ' ' 1
##
## R-sq.(adj) =  0.138
## lmer.REML = 4.2054e+05  Scale est. = 1083.2    n = 33324
```

Next is the model with depression-sleep interaction.

```
##
## Family: gaussian
## Link function: identity
##
## Formula:
## value ~ sex + site + s(age_z, k = 10, bs = "cr") + s(sleep_z,
##     k = 5, bs = "cr") + icv + depression + depression:sleep_z
## <environment: 0x55cffafa0890>
##
## Parametric coefficients:
##                    Estimate Std. Error t value Pr(>|t|)
## (Intercept)        1058.708      9.998 105.895  < 2e-16 ***
## sexmale             -39.548      2.084 -18.981  < 2e-16 ***
## siteousAvanto       -59.180     19.227  -3.078 0.002086 **
## siteousPrisma        -7.087     18.559  -0.382 0.702545
## siteousSkyra        -35.027     14.708  -2.381 0.017251 *
## siteUCAM            -45.954     13.591  -3.381 0.000722 ***
## siteUKB               5.837      9.854   0.592 0.553588
## siteUmU              -3.342     13.726  -0.243 0.807627
```

```
## icv                    68.508      1.069  64.097  < 2e-16 ***
## depression             -6.202      7.458  -0.831 0.405702
## depression:sleep_z     -5.567      5.479  -1.016 0.309584
## ---
## Signif. codes:  0 '***' 0.001 '**' 0.01 '*' 0.05 '.' 0.1 ' ' 1
##
## Approximate significance of smooth terms:
##               edf Ref.df     F  p-value
## s(age_z)    5.569  5.569 6.089 3.63e-05 ***
## s(sleep_z) 2.447  2.447 2.123    0.231
## ---
## Signif. codes:  0 '***' 0.001 '**' 0.01 '*' 0.05 '.' 0.1 ' ' 1
##
## R-sq.(adj) =  0.138
## lmer.REML = 4.2053e+05  Scale est. = 1083.2    n = 33324
```

The plot below shows the sleep-volume curve for the original model and for the model with main effects of SES.

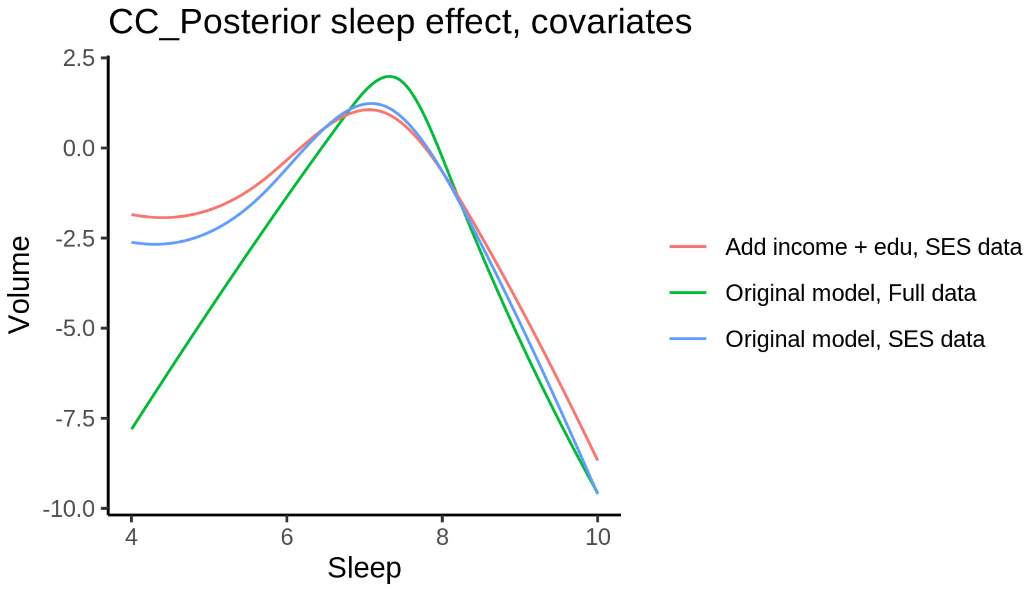


The plot below shows the sleep-volume curve for the original model and for the model with main effects of BMI.

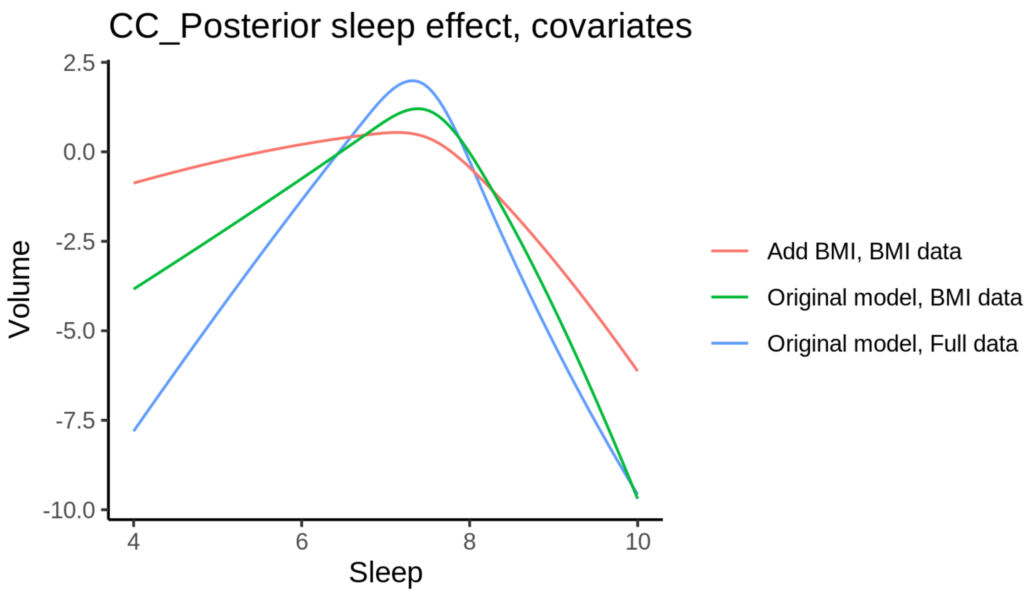


The plot below shows the sleep-volume curve for the original model and for the model with main effects of depression.

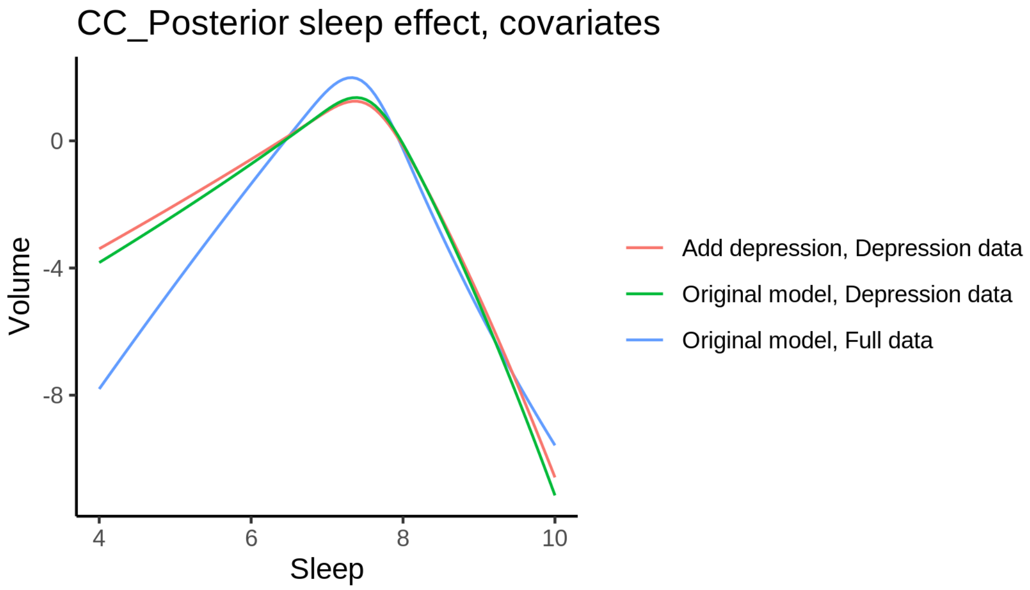


### Cerebellum-Cortex

#### Descriptive statistics

| Study | Observations | Unique IDs | Mean age | Age range |
|---|---|---|---|---|
| HCP | 974 | 974 | 28.8 | 22 - 37 |
| MPIB | 677 | 391 | 63.1 | 24 - 83 |
| UB | 113 | 39 | 70.9 | 64 - 81 |
| UCAM | 884 | 632 | 55.1 | 20 - 88 |

| Study | Observations | Unique IDs | Mean age | Age range |
|---|---|---|---|---|
| UiO | 1474 | 803 | 49.3 | 20 - 89 |
| UKB | 45964 | 43128 | 64.5 | 45 - 83 |
| UmU | 417 | 282 | 62.2 | 25 - 85 |
| UOXF | 769 | 769 | 69.8 | 60 - 85 |

##### Spaghetti plot

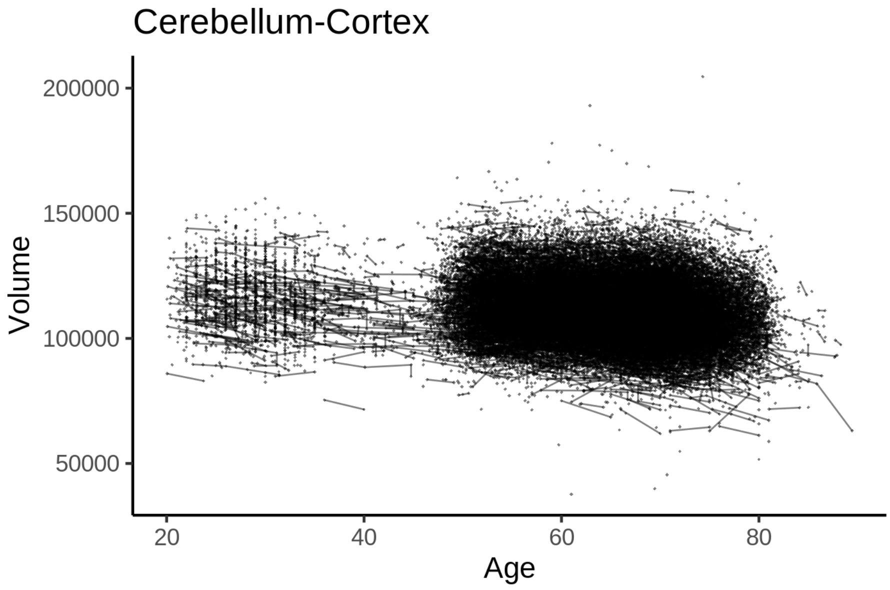


##### Model outputs

Model without sleep term

```
##
## Family: gaussian
## Link function: identity
##
## Formula:
## value ~ sex + site + icv + s(age_z, k = 10, bs = "cr")
## <environment: 0x55cffd315db0>
##
## Parametric coefficients:
##                 Estimate Std. Error t value Pr(>|t|)
## (Intercept)   104987.87     492.72 213.079  < 2e-16 ***
## sexmale         5654.68     103.50  54.635  < 2e-16 ***
## siteMPIB        2235.72     641.51   3.485 0.000492 ***
## siteousAvanto  -4736.69     502.95  -9.418  < 2e-16 ***
## siteousPrisma   2602.29     845.14   3.079 0.002077 **
## siteousSkyra   -6903.58     487.26 -14.168  < 2e-16 ***
## siteUB         -5650.17    1527.78  -3.698 0.000217 ***
## siteUCAM        -584.48     554.13  -1.055 0.291531
## siteUKB         4629.09     504.81   9.170  < 2e-16 ***
## siteUmU       -12200.00     716.02 -17.039  < 2e-16 ***
```

```
## siteUOXF       -4862.83     603.18  -8.062 7.67e-16 ***
## icv             5473.49      51.93 105.394  < 2e-16 ***
## ---
## Signif. codes:  0 ‘***’ 0.001 ‘**’ 0.01 ‘*’ 0.05 ‘.’ 0.1 ‘ ’ 1
##
## Approximate significance of smooth terms:
##            edf Ref.df     F p-value
## s(age_z) 7.381  7.381 345.5  <2e-16 ***
## ---
## Signif. codes:  0 ‘***’ 0.001 ‘**’ 0.01 ‘*’ 0.05 ‘.’ 0.1 ‘ ’ 1
##
## R-sq.(adj) =  0.436
## lmer.REML = 1.0706e+06  Scale est. = 3.9784e+06  n = 51272
```

Model with only main effects of age and sleep

```
##
## Family: gaussian
## Link function: identity
##
## Formula:
## value ~ sex + site + icv + s(age_z, k = 10, bs = "cr") + s(sleep_z,
##     k = 5, bs = "cr")
## <environment: 0x55cffd315db0>
##
## Parametric coefficients:
##                Estimate Std. Error t value Pr(>|t|)
## (Intercept)   105003.09     493.14 212.928  < 2e-16 ***
## sexmale         5652.91     103.48  54.629  < 2e-16 ***
## siteMPIB        2270.02     641.74   3.537 0.000405 ***
## siteousAvanto  -4757.64     503.00  -9.458  < 2e-16 ***
## siteousPrisma   2586.14     844.99   3.061 0.002210 **
## siteousSkyra   -6932.00     487.35 -14.224  < 2e-16 ***
## siteUB         -5595.29    1527.30  -3.664 0.000249 ***
## siteUCAM        -562.50     554.08  -1.015 0.310014
## siteUKB         4613.43     505.29   9.130  < 2e-16 ***
## siteUmU       -12117.80     717.46 -16.890  < 2e-16 ***
## siteUOXF       -4870.34     603.12  -8.075 6.88e-16 ***
## icv             5465.92      51.98 105.160  < 2e-16 ***
## ---
## Signif. codes:  0 ‘***’ 0.001 ‘**’ 0.01 ‘*’ 0.05 ‘.’ 0.1 ‘ ’ 1
##
## Approximate significance of smooth terms:
##              edf Ref.df       F p-value
## s(age_z)   7.382  7.382 339.984 < 2e-16 ***
## s(sleep_z) 3.420  3.420   7.662 1.4e-05 ***
## ---
## Signif. codes:  0 ‘***’ 0.001 ‘**’ 0.01 ‘*’ 0.05 ‘.’ 0.1 ‘ ’ 1
##
## R-sq.(adj) =  0.436
## lmer.REML = 1.0706e+06  Scale est. = 3.9792e+06  n = 51272
```

Model with full interaction between age and sleep

```
##
## Family: gaussian
```

```
## Link function: identity
##
## Formula:
## value ~ sex + site + icv + t2(age_z, sleep_z, k = c(10, 4), bs = "cr")
## <environment: 0x55cffd315db0>
##
## Parametric coefficients:
##                Estimate Std. Error t value Pr(>|t|)
## (Intercept)   105088.51     494.52 212.507  < 2e-16 ***
## sexmale         5646.17     103.70  54.448  < 2e-16 ***
## siteMPIB        2174.87     644.86   3.373 0.000745 ***
## siteousAvanto  -4848.44     505.95  -9.583  < 2e-16 ***
## siteousPrisma   2510.26     847.04   2.964 0.003042 **
## siteousSkyra   -7014.03     490.46 -14.301  < 2e-16 ***
## siteUB         -5720.84    1528.21  -3.743 0.000182 ***
## siteUCAM        -671.38     556.94  -1.205 0.228022
## siteUKB         4530.60     506.63   8.943  < 2e-16 ***
## siteUmU       -12231.94     719.52 -17.000  < 2e-16 ***
## siteUOXF       -4962.99     604.97  -8.204 2.39e-16 ***
## icv             5468.72      51.98 105.217  < 2e-16 ***
## ---
## Signif. codes:  0 ‘***’ 0.001 ‘**’ 0.01 ‘*’ 0.05 ‘.’ 0.1 ‘ ’ 1
##
## Approximate significance of smooth terms:
##                    edf Ref.df     F p-value
## t2(age_z,sleep_z) 14.8   14.8 9.041  <2e-16 ***
## ---
## Signif. codes:  0 ‘***’ 0.001 ‘**’ 0.01 ‘*’ 0.05 ‘.’ 0.1 ‘ ’ 1
##
## R-sq.(adj) =  0.436
## lmer.REML = 1.0706e+06  Scale est. = 3.9815e+06  n = 51272
```

#### Model comparison

`mod_no_sleep` refers to model without sleep term, `mod_no_interaction` refers to model with only main effect of sleep, and `mod_full` refers to model with a full interaction between age and sleep. This is a nested model comparison, and the p-value at a given line refers to comparing the model at the line to the model on the line above. Hence, significance implies that the more complicated model is supported on statistical grounds.

To be even more specific, the p-value on the second row tests whether there is an association between sleep and volume. The p-value on the third row tests whether this association depends on age.

```
## Data: NULL
## Models:
## mod_list$mod_no_sleep$mer: NULL
## mod_list$mod_no_interaction$mer: NULL
## mod_list$mod_full$mer: NULL
##                                 npar     AIC     BIC  logLik deviance  Chisq Df Pr(>Chisq)
## mod_list$mod_no_sleep$mer         16 1070617 1070758 -535292  1070585
## mod_list$mod_no_interaction$mer   18 1070600 1070759 -535282  1070564 21.137  2  2.571e-05 ***
## mod_list$mod_full$mer             20 1070619 1070796 -535290  1070579  0.000  2          1
## ---
## Signif. codes:  0 ‘***’ 0.001 ‘**’ 0.01 ‘*’ 0.05 ‘.’ 0.1 ‘ ’ 1
```

We chose the model based on the likelihood ratio test with 5 % significance level, which was `mod_no_interaction`.

##### Lifespan brain trajectory

The trajectory shown is from the chosen model `mod_no_interaction`.

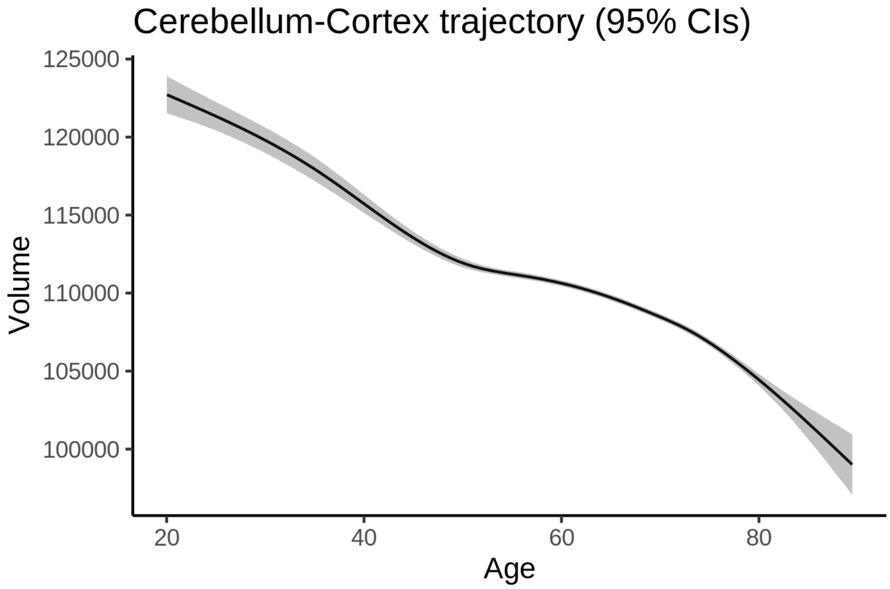


##### Effect of sleep

The chosen model only included the main effect of sleep, and hence the effect does not vary with age. The black dot shows the average sleep duration across all ages in the sample.

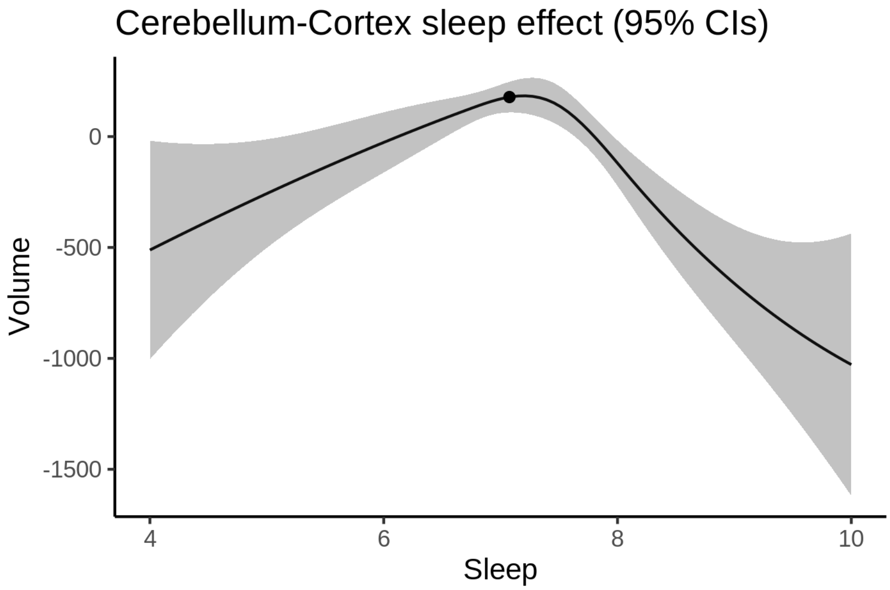


We also show the full interaction model for completeness, although it was not selected.

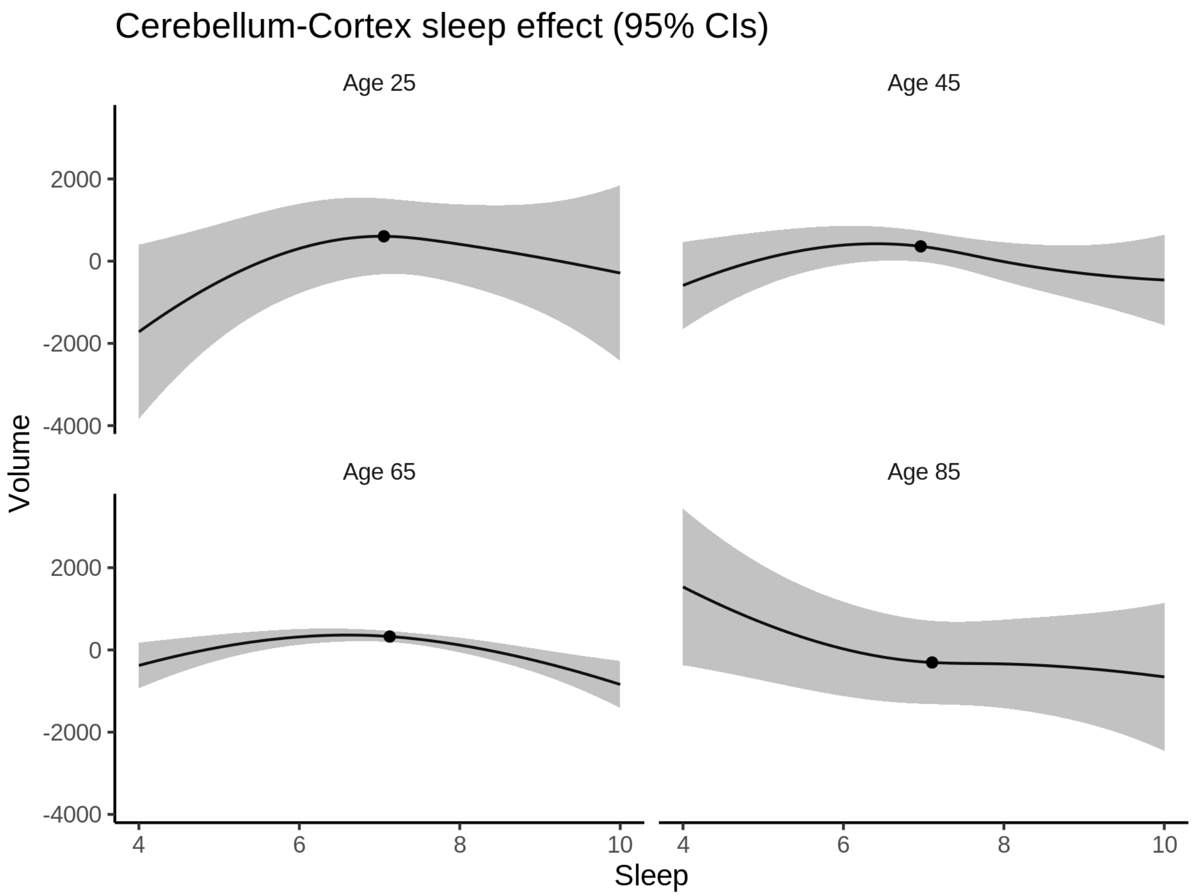


**Deviation from sleep associated with maximal volume**

Model with only main effect of sleep was chosen, so we show it for all ages at once. Maximum volume is attained at 7.2 hours of sleep. The percentage values in the plot are calculated as follows: The maximum at 100 % refers to a person at an arbitrary age with a sleep duration associated with maximum volume. For a female, this volume is 111703 and for a male it is 117356. The other percentage values show how large the expected volume is for someone with other sleep durations. For example, 99 % implies a 1 % reduction.

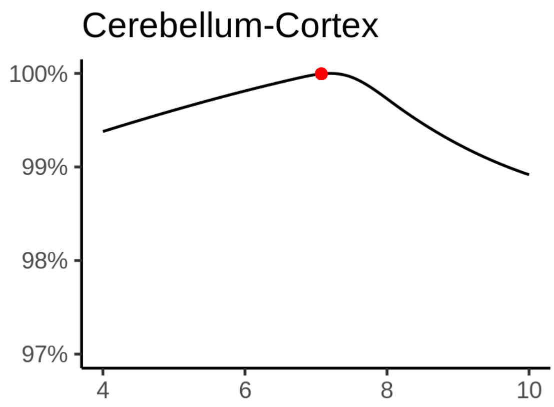

**Comparison of mean sleep and sleep associated with maximum volume**

A 95 % confidence interval for the sleep associated with maximum volume is [6.91, 7.39].

The plot below compares average sleep to the sleep associated with maximum volume.

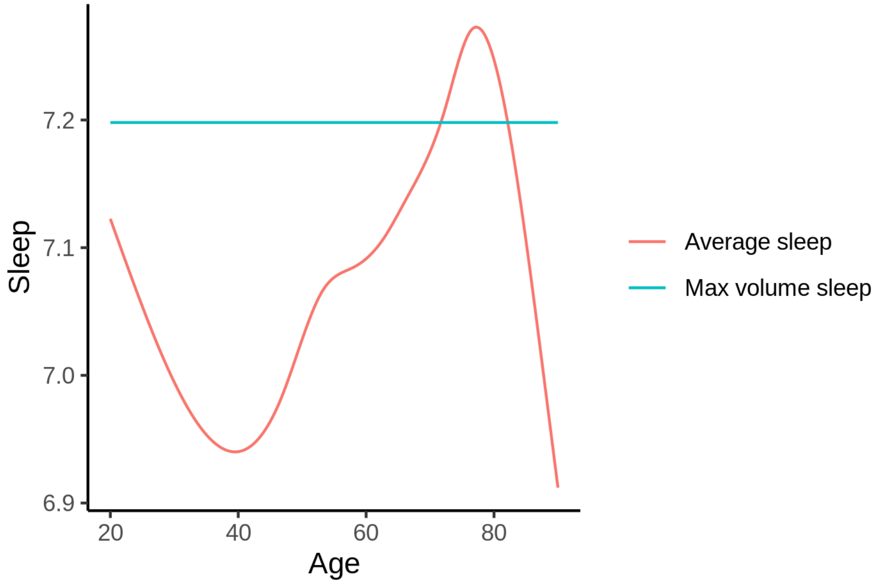


The next plot shows the difference between average sleep and sleep associated with maximum volume. The shaded region is a 95 % confidence interval.

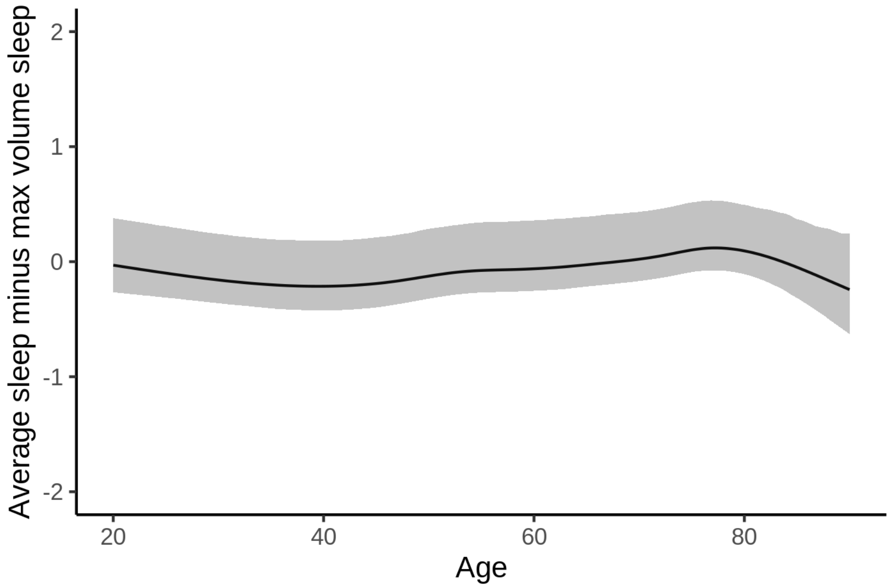


The next plot shows the probability that the sleep duration associated with maximum volume is longer than the average sleep duration, as a function of age. Probility below .05 can be interpreted as evidence that the sleep associated with maximum volume is shorter than the mean sleep, and probability above .95 can be interpreted the opposite way.

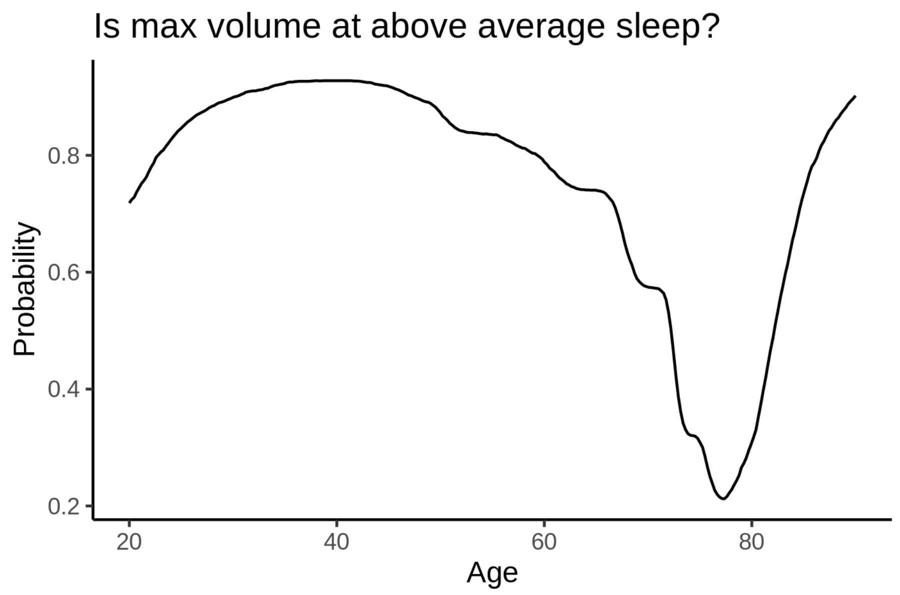


##### Controling for covariates

Below is the output for a model in which we only include data with income and education.

```
##
## Family: gaussian
## Link function: identity
##
## Formula:
## value ~ sex + site + s(age_z, k = 10, bs = "cr") + s(sleep_z,
##     k = 5, bs = "cr") + icv
## <environment: 0x55d072eb1a58>
##
## Parametric coefficients:
##                 Estimate Std. Error t value Pr(>|t|)
## (Intercept)   108960.76    1207.77  90.217  < 2e-16 ***
## sexmale         5582.76     132.41  42.163  < 2e-16 ***
## siteousAvanto -10148.35    1398.56  -7.256 4.07e-13 ***
## siteousPrisma    -14.98    1467.78  -0.010    0.992
## siteousSkyra  -11244.89    1306.60  -8.606  < 2e-16 ***
## siteUKB          534.83    1205.32   0.444    0.657
## siteUOXF       -7639.12    1356.11  -5.633 1.78e-08 ***
## icv             5663.68      67.88  83.433  < 2e-16 ***
## ---
## Signif. codes:  0 ‘***’ 0.001 ‘**’ 0.01 ‘*’ 0.05 ‘.’ 0.1 ‘ ’ 1
##
## Approximate significance of smooth terms:
##              edf Ref.df       F p-value
## s(age_z)   6.036  6.036 168.237  <2e-16 ***
## s(sleep_z) 2.728  2.728   5.459  0.0019 **
## ---
## Signif. codes:  0 ‘***’ 0.001 ‘**’ 0.01 ‘*’ 0.05 ‘.’ 0.1 ‘ ’ 1
##
```

```
## R-sq.(adj) =  0.408
## lmer.REML = 6.513e+05  Scale est. = 3.8427e+06  n = 31180
```

Below is the output for a model in which we control for the main effects of income and education.

```
##
## Family: gaussian
## Link function: identity
##
## Formula:
## value ~ sex + site + s(age_z, k = 10, bs = "cr") + s(sleep_z,
##     k = 5, bs = "cr") + icv + income_scaled + education_scaled
## <environment: 0x55d072eb1a58>
##
## Parametric coefficients:
##                   Estimate Std. Error t value Pr(>|t|)
## (Intercept)      107614.93    1213.67  88.669  < 2e-16 ***
## sexmale            5576.85     132.56  42.071  < 2e-16 ***
## siteousAvanto    -10113.57    1396.91  -7.240 4.59e-13 ***
## siteousPrisma      -121.76    1465.54  -0.083    0.934
## siteousSkyra     -11232.68    1304.71  -8.609  < 2e-16 ***
## siteUKB             567.24    1204.37   0.471    0.638
## siteUOXF          -7203.91    1354.87  -5.317 1.06e-07 ***
## icv                5576.85      68.33  81.620  < 2e-16 ***
## income_scaled      1017.96     157.45   6.465 1.03e-10 ***
## education_scaled   1071.70     186.38   5.750 9.00e-09 ***
## ---
## Signif. codes:  0 ‘***’ 0.001 ‘**’ 0.01 ‘*’ 0.05 ‘.’ 0.1 ‘ ’ 1
##
## Approximate significance of smooth terms:
##              edf Ref.df       F p-value
## s(age_z)   6.156  6.156 140.661  <2e-16 ***
## s(sleep_z) 2.284  2.284   4.245  0.0115 *
## ---
## Signif. codes:  0 ‘***’ 0.001 ‘**’ 0.01 ‘*’ 0.05 ‘.’ 0.1 ‘ ’ 1
##
## R-sq.(adj) =   0.41
## lmer.REML = 6.5117e+05  Scale est. = 3.8472e+06  n = 31180
```

We also included interaction effects between sleep duration and education and income, in another model. The output is shown below, and the interaction terms are `income_scaled:sleep_z` and `education_scaled:sleep_z`.

```
##
## Family: gaussian
## Link function: identity
##
## Formula:
## value ~ sex + site + s(age_z, k = 10, bs = "cr") + s(sleep_z,
##     k = 5, bs = "cr") + icv + income_scaled + education_scaled +
##     income_scaled:sleep_z + education_scaled:sleep_z
## <environment: 0x55d072eb1a58>
##
## Parametric coefficients:
##                           Estimate Std. Error t value Pr(>|t|)
## (Intercept)              107628.57    1213.72  88.677  < 2e-16 ***
```

```
## sexmale                    5565.78     132.64  41.960  < 2e-16 ***
## siteousAvanto            -10137.58    1396.88  -7.257 4.04e-13 ***
## siteousPrisma              -169.96    1465.61  -0.116   0.9077
## siteousSkyra             -11255.57    1304.68  -8.627  < 2e-16 ***
## siteUKB                     552.29    1204.45   0.459   0.6466
## siteUOXF                  -7219.16    1355.20  -5.327 1.01e-07 ***
## icv                        5578.45      68.33  81.639  < 2e-16 ***
## income_scaled              1028.34     157.53   6.528 6.77e-11 ***
## education_scaled           1063.98     186.40   5.708 1.15e-08 ***
## income_scaled:sleep_z      -291.52     156.73  -1.860   0.0629 .
## education_scaled:sleep_z    328.17     180.79   1.815   0.0695 .
## ---
## Signif. codes:  0 '***' 0.001 '**' 0.01 '*' 0.05 '.' 0.1 ' ' 1
##
## Approximate significance of smooth terms:
##              edf Ref.df       F p-value
## s(age_z)   6.162  6.162 140.298  <2e-16 ***
## s(sleep_z) 2.332  2.332   3.349  0.0342 *
## ---
## Signif. codes:  0 '***' 0.001 '**' 0.01 '*' 0.05 '.' 0.1 ' ' 1
##
## R-sq.(adj) =   0.41
## lmer.REML = 6.5114e+05  Scale est. = 3.8472e+06  n = 31180
```

We did the same controling for BMI. Below is the model with no covariates but only keeping data with BMI.

```
##
## Family: gaussian
## Link function: identity
##
## Formula:
## value ~ sex + site + s(age_z, k = 10, bs = "cr") + s(sleep_z,
##     k = 5, bs = "cr") + icv
## <environment: 0x55cffbc3c740>
##
## Parametric coefficients:
##               Estimate Std. Error t value Pr(>|t|)
## (Intercept)   98809.64     730.46 135.270  < 2e-16 ***
## sexmale        5572.67     128.36  43.414  < 2e-16 ***
## siteousPrisma  8219.31     974.87   8.431  < 2e-16 ***
## siteousSkyra   -966.00     511.87  -1.887   0.0591 .
## siteUCAM       5482.29     780.91   7.020 2.25e-12 ***
## siteUKB       10771.24     738.00  14.595  < 2e-16 ***
## siteUmU       -5757.06     905.16  -6.360 2.04e-10 ***
## icv            5643.96      66.04  85.466  < 2e-16 ***
## ---
## Signif. codes:  0 '***' 0.001 '**' 0.01 '*' 0.05 '.' 0.1 ' ' 1
##
## Approximate significance of smooth terms:
##              edf Ref.df       F p-value
## s(age_z)   6.646  6.646 225.333 < 2e-16 ***
## s(sleep_z) 2.742  2.742   5.497 0.00116 **
## ---
## Signif. codes:  0 '***' 0.001 '**' 0.01 '*' 0.05 '.' 0.1 ' ' 1
##
```

```
## R-sq.(adj) =  0.428
## lmer.REML = 6.9782e+05  Scale est. = 4.0433e+06  n = 33429
```

Below is the model output with main effect.

```
##
## Family: gaussian
## Link function: identity
##
## Formula:
## value ~ sex + site + s(age_z, k = 10, bs = "cr") + s(sleep_z,
##     k = 5, bs = "cr") + icv + bmi
## <environment: 0x55cffbc3c740>
##
## Parametric coefficients:
##                Estimate Std. Error t value Pr(>|t|)
## (Intercept)   102585.45     794.68 129.090  < 2e-16 ***
## sexmale         5691.92     128.46  44.311  < 2e-16 ***
## siteousPrisma   8342.91     973.66   8.569  < 2e-16 ***
## siteousSkyra    -955.31     511.75  -1.867   0.0619 .
## siteUCAM        5562.12     779.80   7.133 1.00e-12 ***
## siteUKB        10892.82     737.06  14.779  < 2e-16 ***
## siteUmU        -5600.19     903.74  -6.197 5.83e-10 ***
## icv             5667.54      65.91  85.989  < 2e-16 ***
## bmi             -149.90      12.50 -11.990  < 2e-16 ***
## ---
## Signif. codes:  0 ‘***’ 0.001 ‘**’ 0.01 ‘*’ 0.05 ‘.’ 0.1 ‘ ’ 1
##
## Approximate significance of smooth terms:
##              edf Ref.df       F p-value
## s(age_z)   6.634  6.634 225.845 < 2e-16 ***
## s(sleep_z) 2.382  2.382   5.905 0.00161 **
## ---
## Signif. codes:  0 ‘***’ 0.001 ‘**’ 0.01 ‘*’ 0.05 ‘.’ 0.1 ‘ ’ 1
##
## R-sq.(adj) =   0.43
## lmer.REML = 6.9767e+05  Scale est. = 4.0424e+06  n = 33429
```

Next is the model with BMI-sleep interaction.

```
##
## Family: gaussian
## Link function: identity
##
## Formula:
## value ~ sex + site + s(age_z, k = 10, bs = "cr") + s(sleep_z,
##     k = 5, bs = "cr") + icv + bmi + bmi:sleep_z
## <environment: 0x55cffbc3c740>
##
## Parametric coefficients:
##                 Estimate Std. Error t value Pr(>|t|)
## (Intercept)    1.026e+05  7.951e+02 129.019  < 2e-16 ***
## sexmale        5.692e+03  1.285e+02  44.309  < 2e-16 ***
## siteousPrisma  8.343e+03  9.737e+02   8.568  < 2e-16 ***
## siteousSkyra  -9.553e+02  5.118e+02  -1.867   0.0619 .
## siteUCAM       5.562e+03  7.799e+02   7.132 1.01e-12 ***
```

```
## siteUKB        1.089e+04  7.371e+02  14.777  < 2e-16 ***
## siteUmU       -5.600e+03  9.038e+02  -6.196 5.84e-10 ***
## icv            5.668e+03  6.591e+01  85.985  < 2e-16 ***
## bmi           -1.499e+02  1.251e+01 -11.983  < 2e-16 ***
## bmi:sleep_z   -1.392e-01  1.185e+01  -0.012   0.9906
## ---
## Signif. codes:  0 ‘***’ 0.001 ‘**’ 0.01 ‘*’ 0.05 ‘.’ 0.1 ‘ ’ 1
##
## Approximate significance of smooth terms:
##              edf Ref.df      F p-value
## s(age_z)   6.634  6.634 225.74  <2e-16 ***
## s(sleep_z) 2.381  2.381   2.22   0.095 .
## ---
## Signif. codes:  0 ‘***’ 0.001 ‘**’ 0.01 ‘*’ 0.05 ‘.’ 0.1 ‘ ’ 1
##
## R-sq.(adj) =   0.43
## lmer.REML = 6.9767e+05  Scale est. = 4.0424e+06  n = 33429
```

We did the same controling for depression. Below is the model with no covariates but only keeping data with depression.

```
##
## Family: gaussian
## Link function: identity
##
## Formula:
## value ~ sex + site + s(age_z, k = 10, bs = "cr") + s(sleep_z,
##     k = 5, bs = "cr") + icv
## <environment: 0x55cffd20b258>
##
## Parametric coefficients:
##                 Estimate Std. Error t value Pr(>|t|)
## (Intercept)    106215.39     615.75 172.498  < 2e-16 ***
## sexmale          5603.57     128.83  43.498  < 2e-16 ***
## siteousAvanto   -5079.60    1175.60  -4.321 1.56e-05 ***
## siteousPrisma    1630.21    1149.66   1.418  0.15620
## siteousSkyra    -7300.10     915.11  -7.977 1.54e-15 ***
## siteUCAM        -2195.05     842.51  -2.605  0.00918 **
## siteUKB          3279.97     609.01   5.386 7.26e-08 ***
## siteUmU        -13196.02     826.07 -15.974  < 2e-16 ***
## icv              5623.01      66.16  84.986  < 2e-16 ***
## ---
## Signif. codes:  0 ‘***’ 0.001 ‘**’ 0.01 ‘*’ 0.05 ‘.’ 0.1 ‘ ’ 1
##
## Approximate significance of smooth terms:
##              edf Ref.df      F  p-value
## s(age_z)   6.882  6.882 173.15  < 2e-16 ***
## s(sleep_z) 2.782  2.782   6.43 0.000825 ***
## ---
## Signif. codes:  0 ‘***’ 0.001 ‘**’ 0.01 ‘*’ 0.05 ‘.’ 0.1 ‘ ’ 1
##
## R-sq.(adj) =  0.425
## lmer.REML = 6.9552e+05  Scale est. = 3.795e+06  n = 33341
```

Below is the model output with main effect.

```
##
## Family: gaussian
## Link function: identity
##
## Formula:
## value ~ sex + site + s(age_z, k = 10, bs = "cr") + s(sleep_z,
##     k = 5, bs = "cr") + icv + depression
## <environment: 0x55cffd20b258>
##
## Parametric coefficients:
##                Estimate Std. Error t value Pr(>|t|)
## (Intercept)   106436.61     618.22 172.167  < 2e-16 ***
## sexmale         5580.15     128.93  43.280  < 2e-16 ***
## siteousAvanto  -4961.26    1175.70  -4.220 2.45e-05 ***
## siteousPrisma   1738.82    1149.65   1.512    0.130
## siteousSkyra   -7167.82     915.40  -7.830 5.01e-15 ***
## siteUCAM       -2141.92     842.39  -2.543    0.011 *
## siteUKB         3189.83     609.32   5.235 1.66e-07 ***
## siteUmU       -12517.50     843.59 -14.838  < 2e-16 ***
## icv             5626.59      66.15  85.057  < 2e-16 ***
## depression     -1817.93     461.13  -3.942 8.09e-05 ***
## ---
## Signif. codes:  0 ‘***’ 0.001 ‘**’ 0.01 ‘*’ 0.05 ‘.’ 0.1 ‘ ’ 1
##
## Approximate significance of smooth terms:
##              edf Ref.df      F p-value
## s(age_z)   6.864  6.864 176.06 < 2e-16 ***
## s(sleep_z) 2.566  2.566   6.14 0.00132 **
## ---
## Signif. codes:  0 ‘***’ 0.001 ‘**’ 0.01 ‘*’ 0.05 ‘.’ 0.1 ‘ ’ 1
##
## R-sq.(adj) =  0.426
## lmer.REML = 6.9549e+05  Scale est. = 3.7947e+06  n = 33341
```

Next is the model with depression-sleep interaction.

```
##
## Family: gaussian
## Link function: identity
##
## Formula:
## value ~ sex + site + s(age_z, k = 10, bs = "cr") + s(sleep_z,
##     k = 5, bs = "cr") + icv + depression + depression:sleep_z
## <environment: 0x55cffd20b258>
##
## Parametric coefficients:
##                     Estimate Std. Error t value Pr(>|t|)
## (Intercept)        106436.96     618.21 172.169  < 2e-16 ***
## sexmale              5577.38     128.95  43.252  < 2e-16 ***
## siteousAvanto       -4980.18    1175.80  -4.236 2.29e-05 ***
## siteousPrisma        1719.59    1149.75   1.496   0.1348
## siteousSkyra        -7186.06     915.52  -7.849 4.31e-15 ***
## siteUCAM            -2161.59     842.55  -2.566   0.0103 *
## siteUKB              3190.41     609.32   5.236 1.65e-07 ***
## siteUmU            -12385.89     850.95 -14.555  < 2e-16 ***
```

```
## icv                   5626.74      66.15  85.058  < 2e-16 ***
## depression           -1849.86     462.10  -4.003 6.26e-05 ***
## depression:sleep_z   -403.02     339.93  -1.186   0.2358
## ---
## Signif. codes:  0 '***' 0.001 '**' 0.01 '*' 0.05 '.' 0.1 ' ' 1
##
## Approximate significance of smooth terms:
##              edf Ref.df      F p-value
## s(age_z)   6.862  6.862 176.19  <2e-16 ***
## s(sleep_z) 2.597  2.597   4.05  0.0126 *
## ---
## Signif. codes:  0 '***' 0.001 '**' 0.01 '*' 0.05 '.' 0.1 ' ' 1
##
## R-sq.(adj) =  0.426
## lmer.REML = 6.9547e+05  Scale est. = 3.7946e+06  n = 33341
```

The plot below shows the sleep-volume curve for the original model and for the model with main effects of SES.

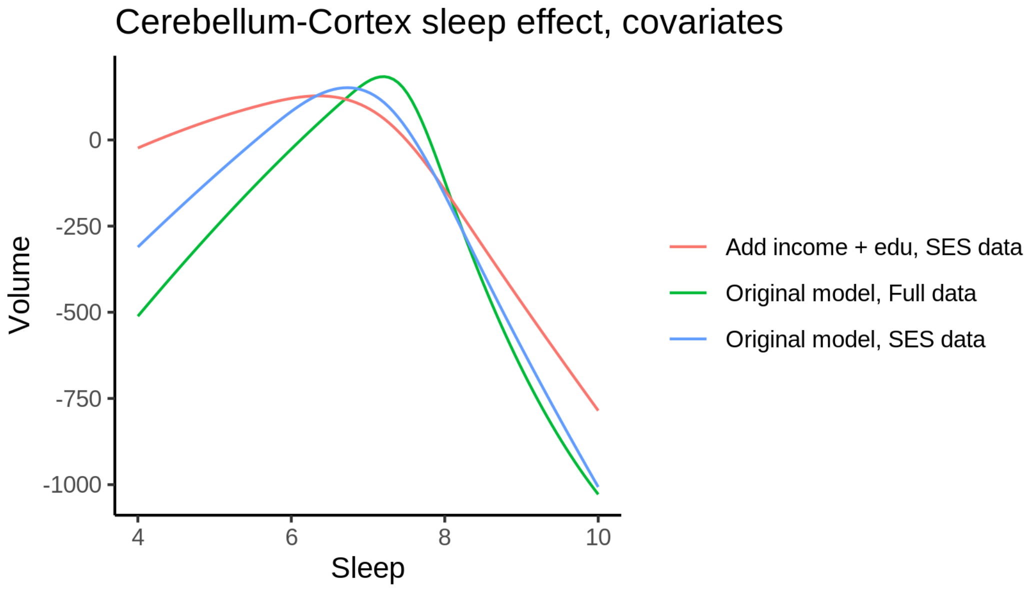


The plot below shows the sleep-volume curve for the original model and for the model with main effects of BMI.

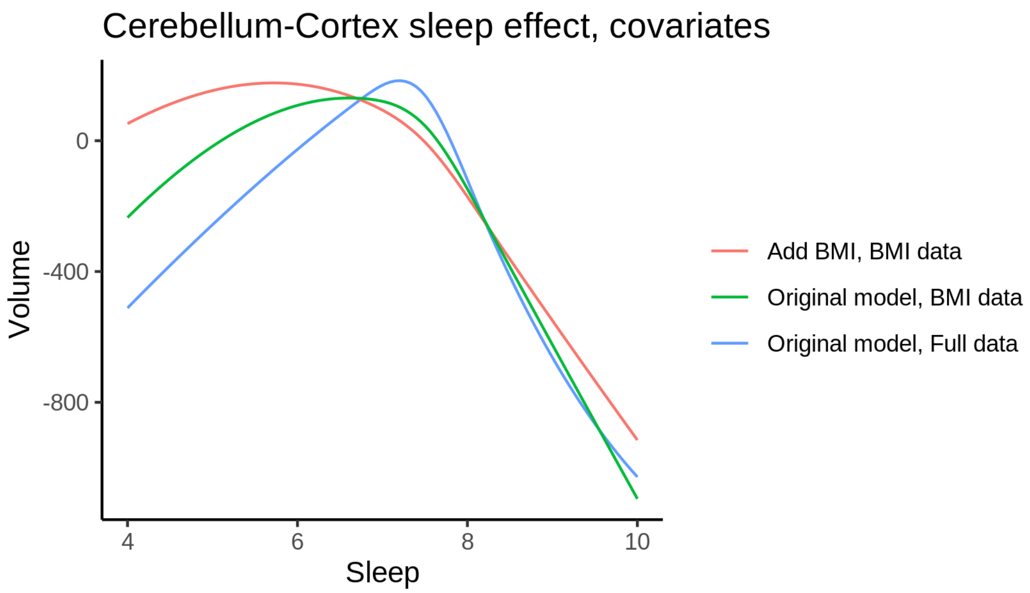


The plot below shows the sleep-volume curve for the original model and for the model with main effects of depression.

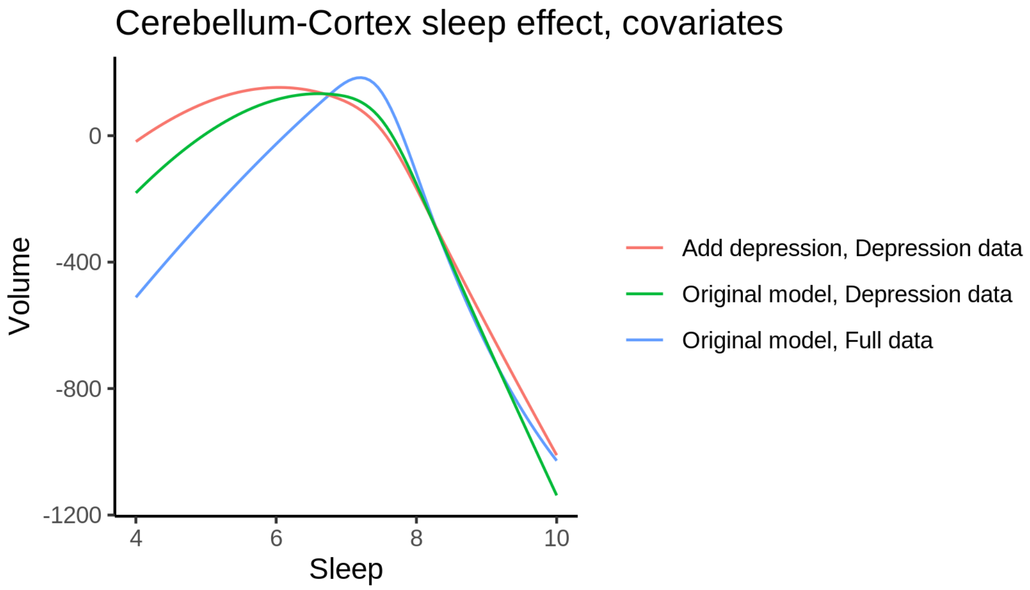


### Cerebellum-White-Matter

#### Descriptive statistics

| Study | Observations | Unique IDs | Mean age | Age range |
|---|---|---|---|---|
| HCP | 974 | 974 | 28.8 | 22 - 37 |
| MPIB | 676 | 391 | 63.2 | 24 - 83 |
| UB | 113 | 39 | 70.9 | 64 - 81 |
| UCAM | 884 | 632 | 55.1 | 20 - 88 |

| Study | Observations | Unique IDs | Mean age | Age range |
|---|---|---|---|---|
| UiO | 1475 | 803 | 49.4 | 20 - 89 |
| UKB | 45957 | 43127 | 64.5 | 45 - 83 |
| UmU | 417 | 283 | 62.3 | 25 - 85 |
| UOXF | 769 | 769 | 69.8 | 60 - 85 |

##### Spaghetti plot

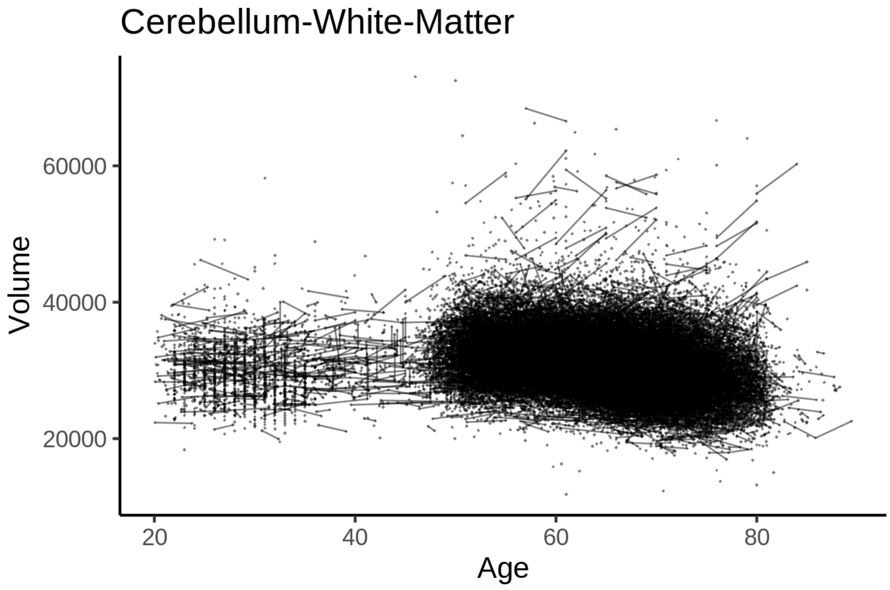


##### Model outputs

Model without sleep term

```
##
## Family: gaussian
## Link function: identity
##
## Formula:
## value ~ sex + site + icv + s(age_z, k = 10, bs = "cr")
## <environment: 0x55cf3feca068>
##
## Parametric coefficients:
##                Estimate Std. Error t value Pr(>|t|)
## (Intercept)   28027.15     200.58 139.733  < 2e-16 ***
## sexmale        -325.54      40.27  -8.085 6.36e-16 ***
## siteMPIB       2435.65     252.34   9.652  < 2e-16 ***
## siteousAvanto   758.12     210.83   3.596 0.000323 ***
## siteousPrisma  -910.44     352.84  -2.580 0.009873 **
## siteousSkyra   3578.81     192.95  18.548  < 2e-16 ***
## siteUB         -306.67     583.21  -0.526 0.599010
## siteUCAM        454.21     219.51   2.069 0.038535 *
## siteUKB        3245.89     205.82  15.771  < 2e-16 ***
## siteUmU       10760.81     281.95  38.166  < 2e-16 ***
```

```
## siteUOXF        765.03     242.39   3.156 0.001599 **
## icv            2066.72      20.27 101.952  < 2e-16 ***
## ---
## Signif. codes:  0 ‘***’ 0.001 ‘**’ 0.01 ‘*’ 0.05 ‘.’ 0.1 ‘ ’ 1
##
## Approximate significance of smooth terms:
##            edf Ref.df     F p-value
## s(age_z) 7.328  7.328 610.1  <2e-16 ***
## ---
## Signif. codes:  0 ‘***’ 0.001 ‘**’ 0.01 ‘*’ 0.05 ‘.’ 0.1 ‘ ’ 1
##
## R-sq.(adj) =  0.324
## lmer.REML = 9.777e+05  Scale est. = 1.6224e+06  n = 51265
```

Model with only main effects of age and sleep

```
##
## Family: gaussian
## Link function: identity
##
## Formula:
## value ~ sex + site + icv + s(age_z, k = 10, bs = "cr") + s(sleep_z,
##     k = 5, bs = "cr")
## <environment: 0x55cf3feca068>
##
## Parametric coefficients:
##                Estimate Std. Error t value Pr(>|t|)
## (Intercept)   27995.42     200.72 139.478  < 2e-16 ***
## sexmale        -327.26      40.26  -8.129 4.44e-16 ***
## siteMPIB       2475.43     252.46   9.805  < 2e-16 ***
## siteousAvanto   772.77     210.84   3.665 0.000247 ***
## siteousPrisma  -901.03     352.78  -2.554 0.010650 *
## siteousSkyra   3593.49     192.98  18.621  < 2e-16 ***
## siteUB         -290.35     583.07  -0.498 0.618509
## siteUCAM        474.65     219.50   2.162 0.030592 *
## siteUKB        3279.85     205.98  15.923  < 2e-16 ***
## siteUmU       10842.12     282.54  38.373  < 2e-16 ***
## siteUOXF        781.77     242.36   3.226 0.001258 **
## icv            2068.04      20.29 101.947  < 2e-16 ***
## ---
## Signif. codes:  0 ‘***’ 0.001 ‘**’ 0.01 ‘*’ 0.05 ‘.’ 0.1 ‘ ’ 1
##
## Approximate significance of smooth terms:
##              edf Ref.df       F  p-value
## s(age_z)   7.308  7.308 601.487  < 2e-16 ***
## s(sleep_z) 2.058  2.058   9.973 4.48e-05 ***
## ---
## Signif. codes:  0 ‘***’ 0.001 ‘**’ 0.01 ‘*’ 0.05 ‘.’ 0.1 ‘ ’ 1
##
## R-sq.(adj) =  0.324
## lmer.REML = 9.7768e+05  Scale est. = 1.6222e+06  n = 51265
```

Model with full interaction between age and sleep

```
##
## Family: gaussian
```

```
## Link function: identity
##
## Formula:
## value ~ sex + site + icv + t2(age_z, sleep_z, k = c(10, 4), bs = "cr")
## <environment: 0x55cf3feca068>
##
## Parametric coefficients:
##                Estimate Std. Error t value Pr(>|t|)
## (Intercept)   27965.91     201.56 138.748  < 2e-16 ***
## sexmale        -329.75      40.35  -8.173 3.09e-16 ***
## siteMPIB       2505.11     253.88   9.867  < 2e-16 ***
## siteousAvanto   798.44     211.92   3.768 0.000165 ***
## siteousPrisma  -879.76     353.54  -2.488 0.012833 *
## siteousSkyra   3623.43     194.23  18.655  < 2e-16 ***
## siteUB         -265.86     583.51  -0.456 0.648661
## siteUCAM        508.10     220.79   2.301 0.021381 *
## siteUKB        3311.07     206.82  16.009  < 2e-16 ***
## siteUmU       10867.45     283.54  38.327  < 2e-16 ***
## siteUOXF        817.66     243.31   3.361 0.000778 ***
## icv            2068.68      20.28 101.987  < 2e-16 ***
## ---
## Signif. codes:  0 ‘***’ 0.001 ‘**’ 0.01 ‘*’ 0.05 ‘.’ 0.1 ‘ ’ 1
##
## Approximate significance of smooth terms:
##                      edf Ref.df     F p-value
## t2(age_z,sleep_z) 12.87  12.87 8.321  <2e-16 ***
## ---
## Signif. codes:  0 ‘***’ 0.001 ‘**’ 0.01 ‘*’ 0.05 ‘.’ 0.1 ‘ ’ 1
##
## R-sq.(adj) =  0.324
## lmer.REML = 9.7768e+05  Scale est. = 1.6219e+06  n = 51265
```

#### Model comparison

`mod_no_sleep` refers to model without sleep term, `mod_no_interaction` refers to model with only main effect of sleep, and `mod_full` refers to model with a full interaction between age and sleep. This is a nested model comparison, and the p-value at a given line refers to comparing the model at the line to the model on the line above. Hence, significance implies that the more complicated model is supported on statistical grounds.

To be even more specific, the p-value on the second row tests whether there is an association between sleep and volume. The p-value on the third row tests whether this association depends on age.

```
## Data: NULL
## Models:
## mod_list$mod_no_sleep$mer: NULL
## mod_list$mod_no_interaction$mer: NULL
## mod_list$mod_full$mer: NULL
##                                 npar    AIC    BIC  logLik deviance  Chisq Df Pr(>Chisq)
## mod_list$mod_no_sleep$mer         16 977728 977869 -488848   977696
## mod_list$mod_no_interaction$mer   18 977714 977873 -488839   977678 17.448  2  0.0001626 ***
## mod_list$mod_full$mer             20 977722 977899 -488841   977682  0.000  2  1.0000000
## ---
## Signif. codes:  0 ‘***’ 0.001 ‘**’ 0.01 ‘*’ 0.05 ‘.’ 0.1 ‘ ’ 1
```

We chose the model based on the likelihood ratio test with 5 % significance level, which was `mod_no_interaction`.

#### Lifespan brain trajectory

The trajectory shown is from the chosen model `mod_no_interaction`.

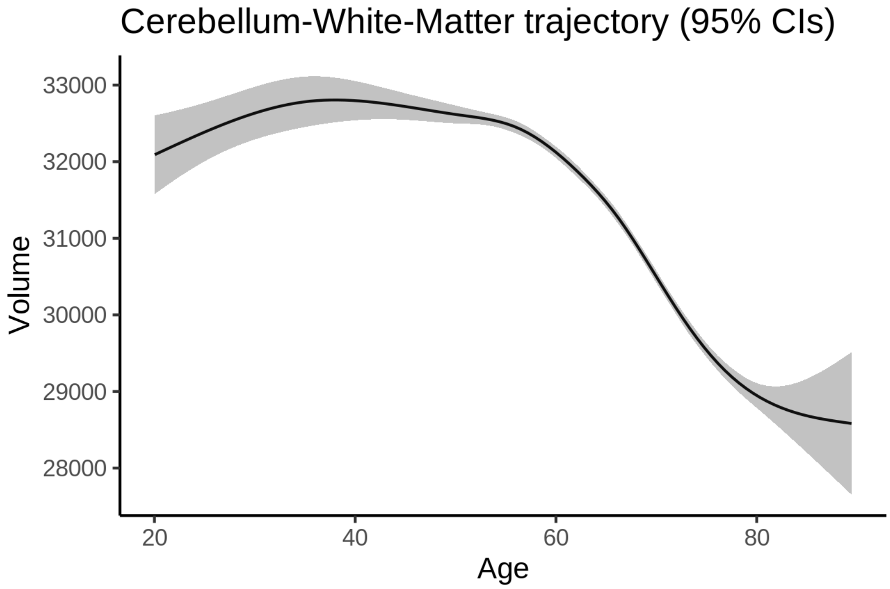


#### Effect of sleep

The chosen model only included the main effect of sleep, and hence the effect does not vary with age. The black dot shows the average sleep duration across all ages in the sample.

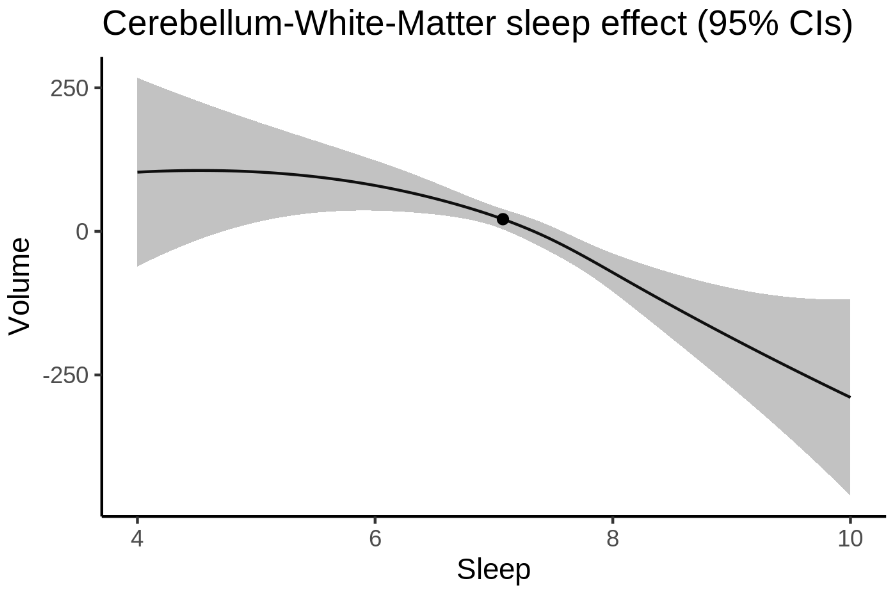


We also show the full interaction model for completeness, although it was not selected.

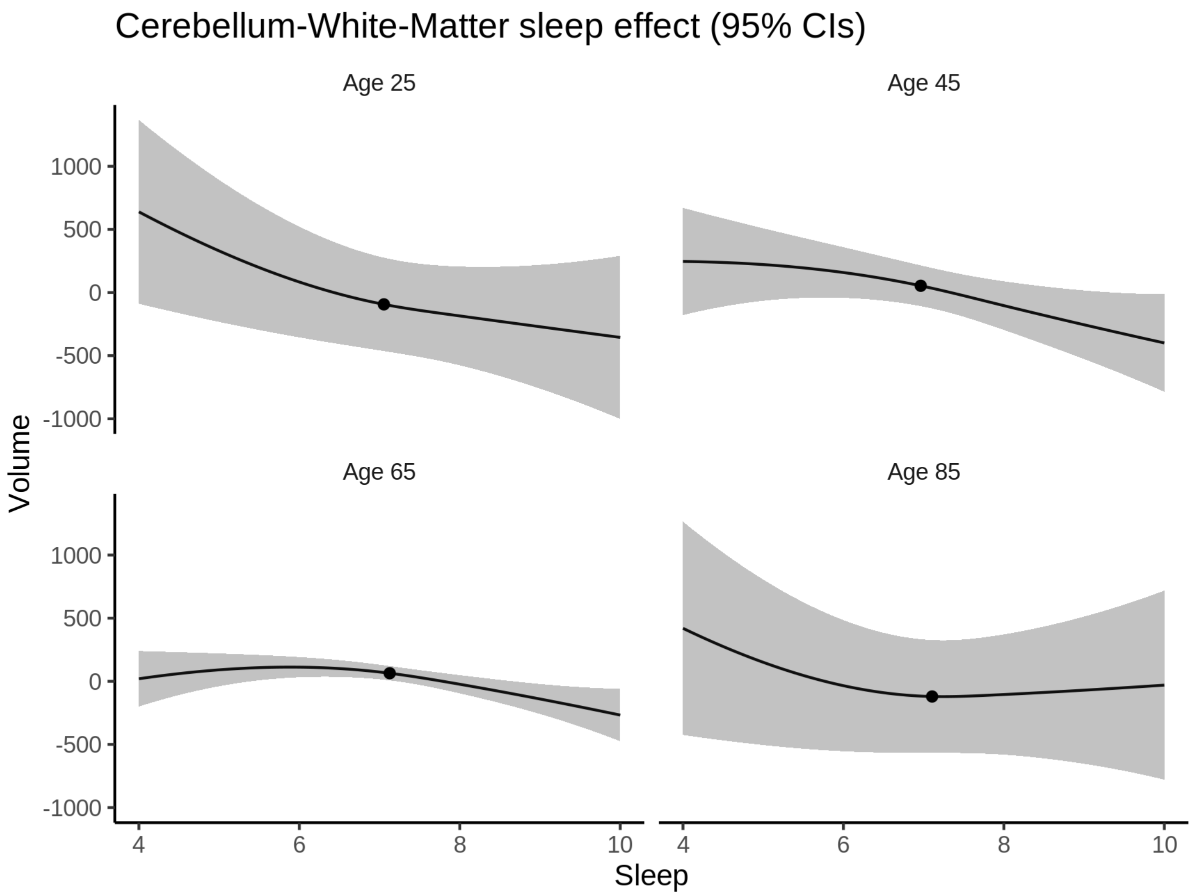


**Deviation from sleep associated with maximal volume**

Model with only main effect of sleep was chosen, so we show it for all ages at once. Maximum volume is attained at 4.5 hours of sleep. The percentage values in the plot are calculated as follows: The maximum at 100 % refers to a person at an arbitrary age with a sleep duration associated with maximum volume. For a female, this volume is 31417 and for a male it is 31090. The other percentage values show how large the expected volume is for someone with other sleep durations. For example, 99 % implies a 1 % reduction.

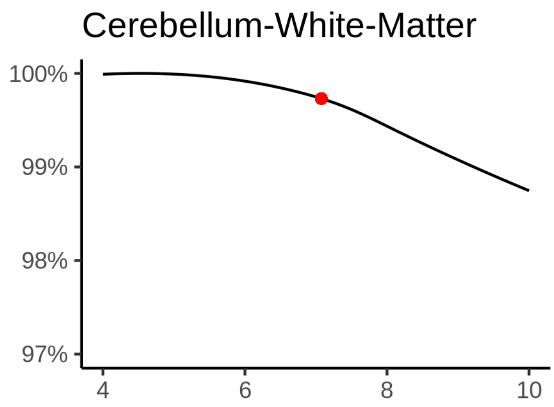

**Comparison of mean sleep and sleep associated with maximum volume**

A 95 % confidence interval for the sleep associated with maximum volume is [4, 6.06].

The plot below compares average sleep to the sleep associated with maximum volume.

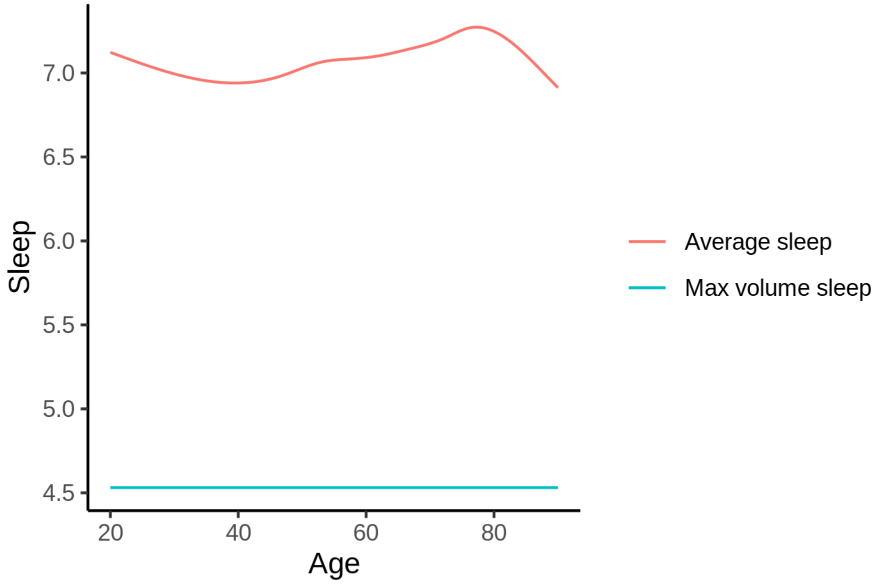


The next plot shows the difference between average sleep and sleep associated with maximum volume. The shaded region is a 95 % confidence interval.

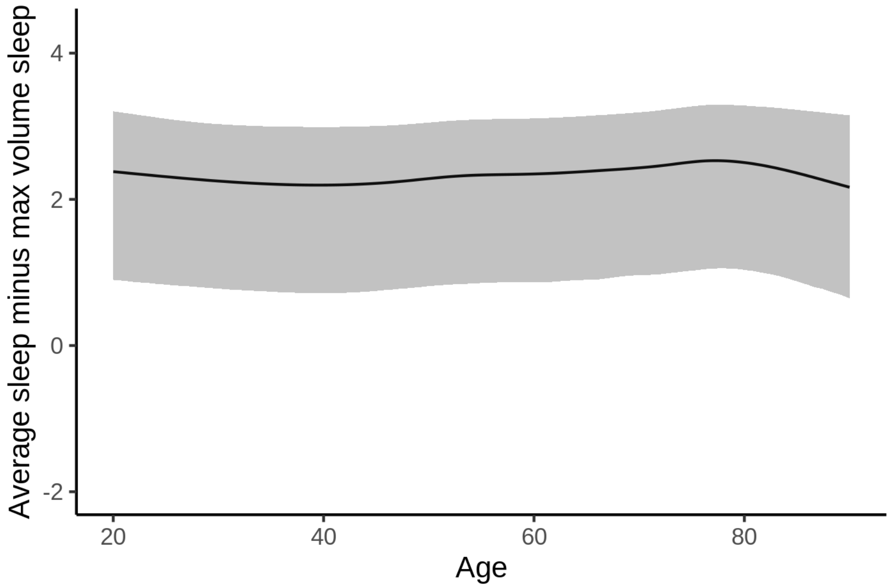


The next plot shows the probability that the sleep duration associated with maximum volume is longer than the average sleep duration, as a function of age. Probility below .05 can be interpreted as evidence that the sleep associated with maximum volume is shorter than the mean sleep, and probability above .95 can be interpreted the opposite way.

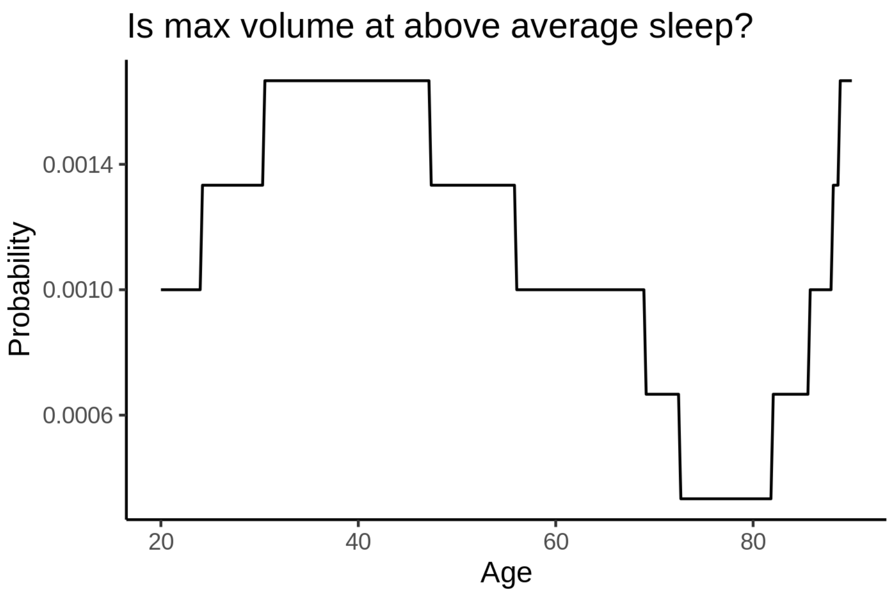


##### Controling for covariates

Below is the output for a model in which we only include data with income and education.

```
##
## Family: gaussian
## Link function: identity
##
## Formula:
## value ~ sex + site + s(age_z, k = 10, bs = "cr") + s(sleep_z,
##     k = 5, bs = "cr") + icv
## <environment: 0x55d07603ad80>
##
## Parametric coefficients:
##               Estimate Std. Error t value Pr(>|t|)
## (Intercept)   30837.74     454.76  67.811  < 2e-16 ***
## sexmale        -409.89      50.22  -8.163 3.40e-16 ***
## siteousAvanto -3033.27     573.54  -5.289 1.24e-07 ***
## siteousPrisma -3764.78     562.38  -6.694 2.20e-11 ***
## siteousSkyra    547.36     491.22   1.114  0.26517
## siteUKB         462.96     454.05   1.020  0.30791
## siteUOXF      -1362.15     511.55  -2.663  0.00775 **
## icv            2150.91      25.83  83.270  < 2e-16 ***
## ---
## Signif. codes:  0 '***' 0.001 '**' 0.01 '*' 0.05 '.' 0.1 ' ' 1
##
## Approximate significance of smooth terms:
##              edf Ref.df       F  p-value
## s(age_z)   6.400  6.400 423.807  < 2e-16 ***
## s(sleep_z) 2.694  2.694   6.072 0.000664 ***
## ---
## Signif. codes:  0 '***' 0.001 '**' 0.01 '*' 0.05 '.' 0.1 ' ' 1
##
```

```
## R-sq.(adj) =  0.303
## lmer.REML = 5.9336e+05  Scale est. = 1.6984e+06  n = 31179
```

Below is the output for a model in which we control for the main effects of income and education.

```
##
## Family: gaussian
## Link function: identity
##
## Formula:
## value ~ sex + site + s(age_z, k = 10, bs = "cr") + s(sleep_z,
##     k = 5, bs = "cr") + icv + income_scaled + education_scaled
## <environment: 0x55d07603ad80>
##
## Parametric coefficients:
##                  Estimate Std. Error t value Pr(>|t|)
## (Intercept)      30635.97     457.64  66.943  < 2e-16 ***
## sexmale           -407.59      50.35  -8.095 5.94e-16 ***
## siteousAvanto    -3044.18     573.56  -5.308 1.12e-07 ***
## siteousPrisma    -3790.45     562.29  -6.741 1.60e-11 ***
## siteousSkyra       535.21     491.23   1.090  0.27593
## siteUKB            457.44     454.17   1.007  0.31385
## siteUOXF         -1301.09     511.70  -2.543  0.01100 *
## icv               2138.26      26.04  82.101  < 2e-16 ***
## income_scaled      103.74      59.91   1.732  0.08333 .
## education_scaled   201.71      70.66   2.855  0.00431 **
## ---
## Signif. codes:  0 '***' 0.001 '**' 0.01 '*' 0.05 '.' 0.1 ' ' 1
##
## Approximate significance of smooth terms:
##              edf Ref.df       F  p-value
## s(age_z)   6.355  6.355 380.965  < 2e-16 ***
## s(sleep_z) 2.449  2.449   6.306 0.000783 ***
## ---
## Signif. codes:  0 '***' 0.001 '**' 0.01 '*' 0.05 '.' 0.1 ' ' 1
##
## R-sq.(adj) =  0.304
## lmer.REML = 5.9333e+05  Scale est. = 1.6988e+06  n = 31179
```

We also included interaction effects between sleep duration and education and income, in another model. The output is shown below, and the interaction terms are `income_scaled:sleep_z` and `education_scaled:sleep_z`.

```
##
## Family: gaussian
## Link function: identity
##
## Formula:
## value ~ sex + site + s(age_z, k = 10, bs = "cr") + s(sleep_z,
##     k = 5, bs = "cr") + icv + income_scaled + education_scaled +
##     income_scaled:sleep_z + education_scaled:sleep_z
## <environment: 0x55d07603ad80>
##
## Parametric coefficients:
##                          Estimate Std. Error t value Pr(>|t|)
## (Intercept)              30644.43     457.66  66.959  < 2e-16 ***
```

```
## sexmale                  -410.16      50.39  -8.140 4.09e-16 ***
## siteousAvanto           -3052.49     573.55  -5.322 1.03e-07 ***
## siteousPrisma           -3805.89     562.33  -6.768 1.33e-11 ***
## siteousSkyra              527.13     491.23   1.073  0.28325
## siteUKB                   447.61     454.20   0.985  0.32440
## siteUOXF                -1316.26     511.82  -2.572  0.01012 *
## icv                      2138.89      26.05  82.119  < 2e-16 ***
## income_scaled             105.96      59.94   1.768  0.07713 .
## education_scaled          199.38      70.67   2.821  0.00478 **
## income_scaled:sleep_z     -47.60      59.50  -0.800  0.42373
## education_scaled:sleep_z  131.74      68.57   1.921  0.05470 .
## ---
## Signif. codes:  0 '***' 0.001 '**' 0.01 '*' 0.05 '.' 0.1 ' ' 1
##
## Approximate significance of smooth terms:
##              edf Ref.df       F p-value
## s(age_z)   6.349  6.349 380.679 < 2e-16 ***
## s(sleep_z) 2.536  2.536   4.063 0.00947 **
## ---
## Signif. codes:  0 '***' 0.001 '**' 0.01 '*' 0.05 '.' 0.1 ' ' 1
##
## R-sq.(adj) =  0.304
## lmer.REML = 5.933e+05  Scale est. = 1.6988e+06  n = 31179
```

We did the same controling for BMI. Below is the model with no covariates but only keeping data with BMI.

```
##
## Family: gaussian
## Link function: identity
##
## Formula:
## value ~ sex + site + s(age_z, k = 10, bs = "cr") + s(sleep_z,
##     k = 5, bs = "cr") + icv
## <environment: 0x55cffbe978f0>
##
## Parametric coefficients:
##               Estimate Std. Error t value Pr(>|t|)
## (Intercept)   29123.37     376.69  77.313  < 2e-16 ***
## sexmale        -456.22      50.32  -9.066  < 2e-16 ***
## siteousPrisma -1686.88     480.76  -3.509 0.000451 ***
## siteousSkyra   2851.11     323.58   8.811  < 2e-16 ***
## siteUCAM       -429.40     391.11  -1.098 0.272260
## siteUKB        2190.09     379.10   5.777 7.67e-09 ***
## siteUmU        9969.42     429.41  23.216  < 2e-16 ***
## icv            2182.75      25.97  84.063  < 2e-16 ***
## ---
## Signif. codes:  0 '***' 0.001 '**' 0.01 '*' 0.05 '.' 0.1 ' ' 1
##
## Approximate significance of smooth terms:
##              edf Ref.df       F  p-value
## s(age_z)   7.237  7.237 367.765  < 2e-16 ***
## s(sleep_z) 2.037  2.037   6.878 0.000926 ***
## ---
## Signif. codes:  0 '***' 0.001 '**' 0.01 '*' 0.05 '.' 0.1 ' ' 1
##
```

```
## R-sq.(adj) =  0.309
## lmer.REML = 6.3794e+05  Scale est. = 1.7505e+06  n = 33428
```

Below is the model output with main effect.

```
##
## Family: gaussian
## Link function: identity
##
## Formula:
## value ~ sex + site + s(age_z, k = 10, bs = "cr") + s(sleep_z,
##     k = 5, bs = "cr") + icv + bmi
## <environment: 0x55cffbe978f0>
##
## Parametric coefficients:
##               Estimate Std. Error t value Pr(>|t|)
## (Intercept)   30342.89     396.07  76.610  < 2e-16 ***
## sexmale        -417.76      50.40  -8.290  < 2e-16 ***
## siteousPrisma -1629.25     480.36  -3.392 0.000695 ***
## siteousSkyra   2862.95     323.50   8.850  < 2e-16 ***
## siteUCAM       -398.20     390.82  -1.019 0.308265
## siteUKB        2231.40     378.85   5.890  3.9e-09 ***
## siteUmU       10025.16     429.06  23.365  < 2e-16 ***
## icv            2190.64      25.93  84.491  < 2e-16 ***
## bmi             -48.50       4.89  -9.918  < 2e-16 ***
## ---
## Signif. codes:  0 ‘***’ 0.001 ‘**’ 0.01 ‘*’ 0.05 ‘.’ 0.1 ‘ ’ 1
##
## Approximate significance of smooth terms:
##              edf Ref.df      F  p-value
## s(age_z)   7.242  7.242 371.46  < 2e-16 ***
## s(sleep_z) 1.402  1.402  12.25 0.000383 ***
## ---
## Signif. codes:  0 ‘***’ 0.001 ‘**’ 0.01 ‘*’ 0.05 ‘.’ 0.1 ‘ ’ 1
##
## R-sq.(adj) =  0.311
## lmer.REML = 6.3784e+05  Scale est. = 1.7502e+06  n = 33428
```

Next is the model with BMI-sleep interaction.

```
##
## Family: gaussian
## Link function: identity
##
## Formula:
## value ~ sex + site + s(age_z, k = 10, bs = "cr") + s(sleep_z,
##     k = 5, bs = "cr") + icv + bmi + bmi:sleep_z
## <environment: 0x55cffbe978f0>
##
## Parametric coefficients:
##                Estimate Std. Error t value Pr(>|t|)
## (Intercept)   30337.329    396.221  76.567  < 2e-16 ***
## sexmale        -417.583     50.397  -8.286  < 2e-16 ***
## siteousPrisma -1628.690    480.366  -3.391 0.000698 ***
## siteousSkyra   2864.400    323.509   8.854  < 2e-16 ***
## siteUCAM       -396.501    390.853  -1.014 0.310375
```

```
## siteUKB        2235.141    378.880   5.899 3.68e-09 ***
## siteUmU       10026.875    429.087  23.368  < 2e-16 ***
## icv            2191.119     25.921  84.529  < 2e-16 ***
## bmi             -48.488      4.890  -9.916  < 2e-16 ***
## bmi:sleep_z       2.963      4.628   0.640 0.522039
## ---
## Signif. codes:  0 '***' 0.001 '**' 0.01 '*' 0.05 '.' 0.1 ' ' 1
##
## Approximate significance of smooth terms:
##              edf Ref.df      F p-value
## s(age_z)   7.245  7.245 372.06  <2e-16 ***
## s(sleep_z) 1.000  1.000   1.65   0.199
## ---
## Signif. codes:  0 '***' 0.001 '**' 0.01 '*' 0.05 '.' 0.1 ' ' 1
##
## R-sq.(adj) =  0.311
## lmer.REML = 6.3783e+05  Scale est. = 1.7502e+06  n = 33428
```

We did the same controling for depression. Below is the model with no covariates but only keeping data with depression.

```
##
## Family: gaussian
## Link function: identity
##
## Formula:
## value ~ sex + site + s(age_z, k = 10, bs = "cr") + s(sleep_z,
##     k = 5, bs = "cr") + icv
## <environment: 0x55cffcf831d0>
##
## Parametric coefficients:
##               Estimate Std. Error t value Pr(>|t|)
## (Intercept)   29743.85     236.40 125.818  < 2e-16 ***
## sexmale        -439.89      50.51  -8.710  < 2e-16 ***
## siteousAvanto  -269.42     585.92  -0.460 0.645650
## siteousPrisma -1610.45     467.68  -3.443 0.000575 ***
## siteousSkyra   2972.65     362.61   8.198 2.53e-16 ***
## siteUCAM      -1350.52     325.14  -4.154 3.28e-05 ***
## siteUKB        1540.93     233.71   6.593 4.37e-11 ***
## siteUmU        9346.59     318.37  29.358  < 2e-16 ***
## icv            2159.80      26.02  83.006  < 2e-16 ***
## ---
## Signif. codes:  0 '***' 0.001 '**' 0.01 '*' 0.05 '.' 0.1 ' ' 1
##
## Approximate significance of smooth terms:
##              edf Ref.df       F p-value
## s(age_z)   7.046  7.046 368.904 < 2e-16 ***
## s(sleep_z) 2.068  2.068   6.073 0.00193 **
## ---
## Signif. codes:  0 '***' 0.001 '**' 0.01 '*' 0.05 '.' 0.1 ' ' 1
##
## R-sq.(adj) =  0.319
## lmer.REML = 6.3601e+05  Scale est. = 1.702e+06  n = 33340
```

Below is the model output with main effect.

```
##
## Family: gaussian
## Link function: identity
##
## Formula:
## value ~ sex + site + s(age_z, k = 10, bs = "cr") + s(sleep_z,
##     k = 5, bs = "cr") + icv + depression
## <environment: 0x55cffcf831d0>
##
## Parametric coefficients:
##               Estimate Std. Error t value Pr(>|t|)
## (Intercept)   29839.51     237.39 125.699  < 2e-16 ***
## sexmale        -449.62      50.54  -8.896  < 2e-16 ***
## siteousAvanto  -221.06     585.99  -0.377 0.705991
## siteousPrisma -1566.76     467.68  -3.350 0.000809 ***
## siteousSkyra   3028.34     362.76   8.348  < 2e-16 ***
## siteUCAM      -1329.01     325.10  -4.088 4.36e-05 ***
## siteUKB        1501.22     233.83   6.420 1.38e-10 ***
## siteUmU        9634.00     325.15  29.630  < 2e-16 ***
## icv            2161.30      26.01  83.097  < 2e-16 ***
## depression     -777.89     179.77  -4.327 1.51e-05 ***
## ---
## Signif. codes:  0 ‘***’ 0.001 ‘**’ 0.01 ‘*’ 0.05 ‘.’ 0.1 ‘ ’ 1
##
## Approximate significance of smooth terms:
##              edf Ref.df       F p-value
## s(age_z)   7.062  7.062 370.757 < 2e-16 ***
## s(sleep_z) 1.661  1.661   9.159 0.00191 **
## ---
## Signif. codes:  0 ‘***’ 0.001 ‘**’ 0.01 ‘*’ 0.05 ‘.’ 0.1 ‘ ’ 1
##
## R-sq.(adj) =  0.319
## lmer.REML = 6.3598e+05  Scale est. = 1.7023e+06  n = 33340
```

Next is the model with depression-sleep interaction.

```
##
## Family: gaussian
## Link function: identity
##
## Formula:
## value ~ sex + site + s(age_z, k = 10, bs = "cr") + s(sleep_z,
##     k = 5, bs = "cr") + icv + depression + depression:sleep_z
## <environment: 0x55cffcf831d0>
##
## Parametric coefficients:
##                    Estimate Std. Error t value Pr(>|t|)
## (Intercept)        29839.14     237.38 125.700  < 2e-16 ***
## sexmale             -448.42      50.55  -8.871  < 2e-16 ***
## siteousAvanto       -211.74     586.03  -0.361 0.717866
## siteousPrisma      -1557.13     467.71  -3.329 0.000872 ***
## siteousSkyra        3037.66     362.80   8.373  < 2e-16 ***
## siteUCAM           -1320.15     325.15  -4.060 4.92e-05 ***
## siteUKB             1501.48     233.82   6.422 1.37e-10 ***
## siteUmU             9576.41     327.81  29.214  < 2e-16 ***
```

```
## icv                  2161.38       26.01  83.111  < 2e-16 ***
## depression           -767.19      179.79  -4.267 1.99e-05 ***
## depression:sleep_z    183.62      131.12   1.400 0.161400
## ---
## Signif. codes:  0 '***' 0.001 '**' 0.01 '*' 0.05 '.' 0.1 ' ' 1
##
## Approximate significance of smooth terms:
##               edf Ref.df      F  p-value
## s(age_z)    7.057  7.057 371.40  < 2e-16 ***
## s(sleep_z) 1.406  1.406  11.47 0.000745 ***
## ---
## Signif. codes:  0 '***' 0.001 '**' 0.01 '*' 0.05 '.' 0.1 ' ' 1
##
## R-sq.(adj) =  0.319
## lmer.REML = 6.3597e+05  Scale est. = 1.7024e+06  n = 33340
```

The plot below shows the sleep-volume curve for the original model and for the model with main effects of SES.

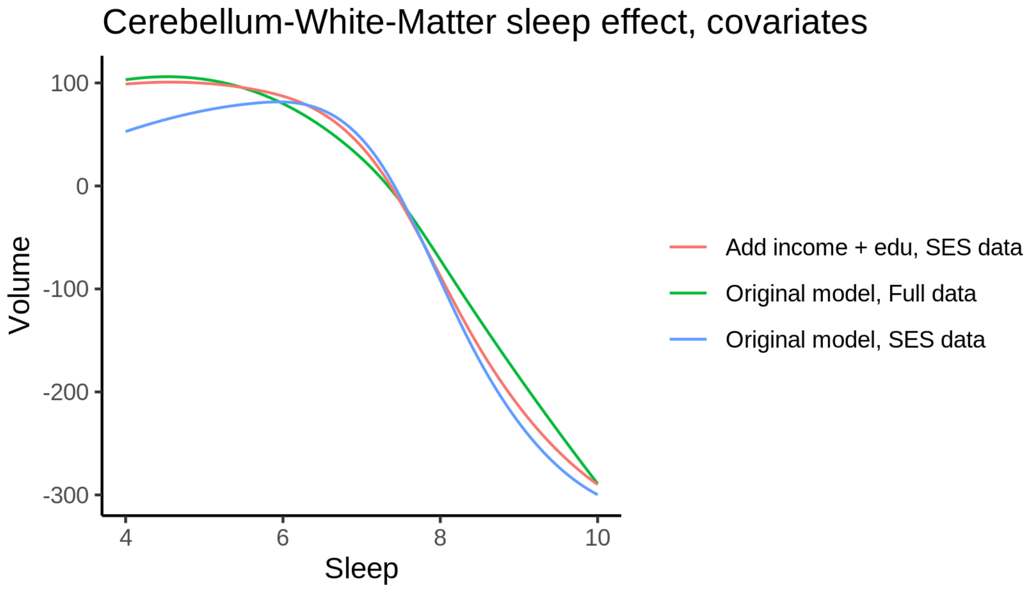


The plot below shows the sleep-volume curve for the original model and for the model with main effects of BMI.

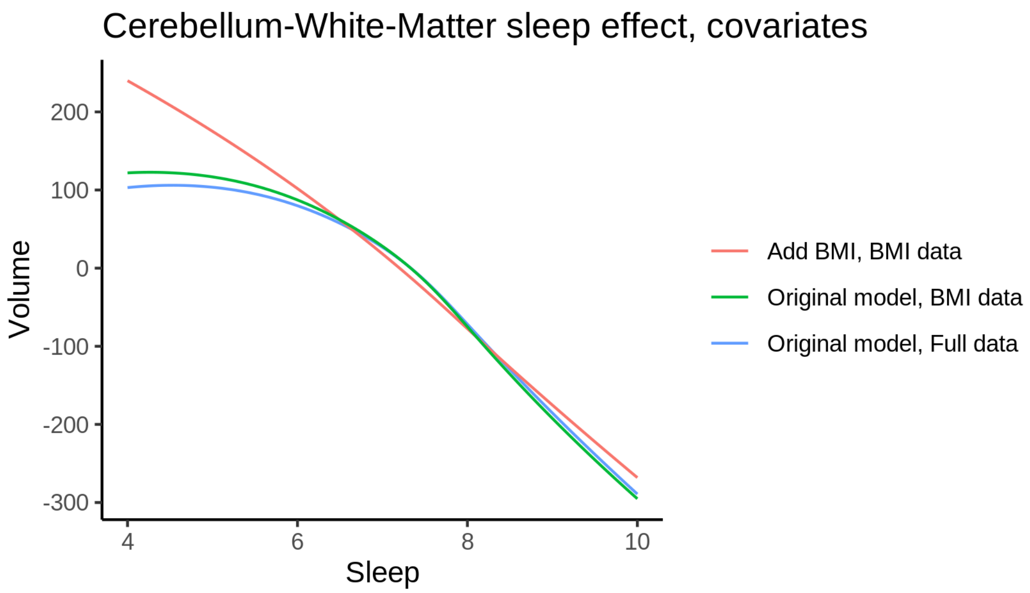


The plot below shows the sleep-volume curve for the original model and for the model with main effects of depression.

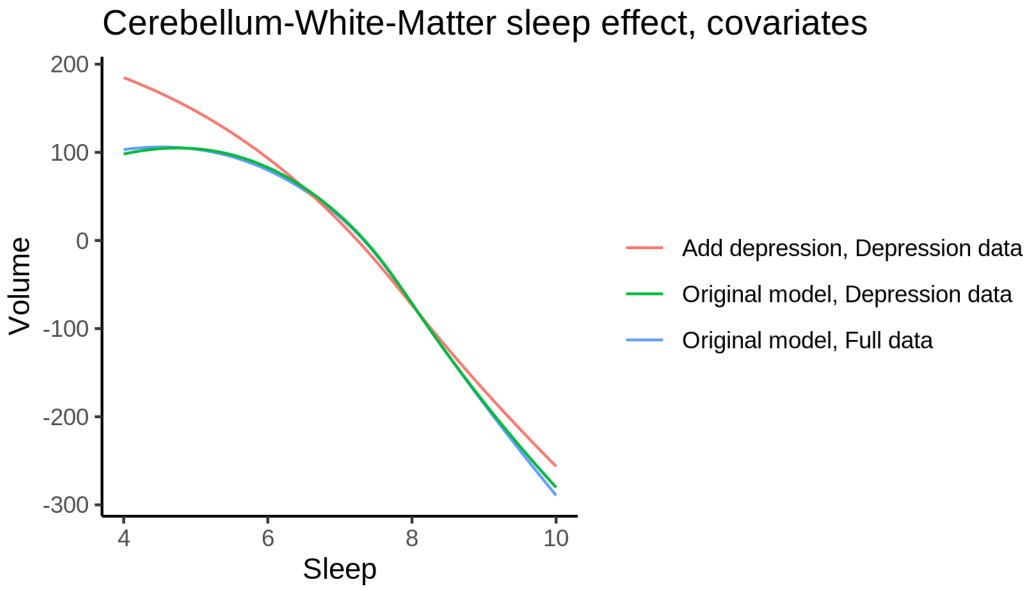


### CerebralWhiteMatterVol

#### Descriptive statistics

| Study | Observations | Unique IDs | Mean age | Age range |
|---|---|---|---|---|
| HCP | 974 | 974 | 28.8 | 22 - 37 |
| MPIB | 677 | 391 | 63.1 | 24 - 83 |
| UB | 113 | 39 | 70.9 | 64 - 81 |
| UCAM | 881 | 631 | 55.1 | 20 - 88 |

| Study | Observations | Unique IDs | Mean age | Age range |
|---|---|---|---|---|
| UiO | 1475 | 803 | 49.4 | 20 - 89 |
| UKB | 45979 | 43135 | 64.5 | 45 - 83 |
| UmU | 423 | 284 | 62.3 | 25 - 85 |
| UOXF | 769 | 769 | 69.8 | 60 - 85 |

#### Spaghetti plot

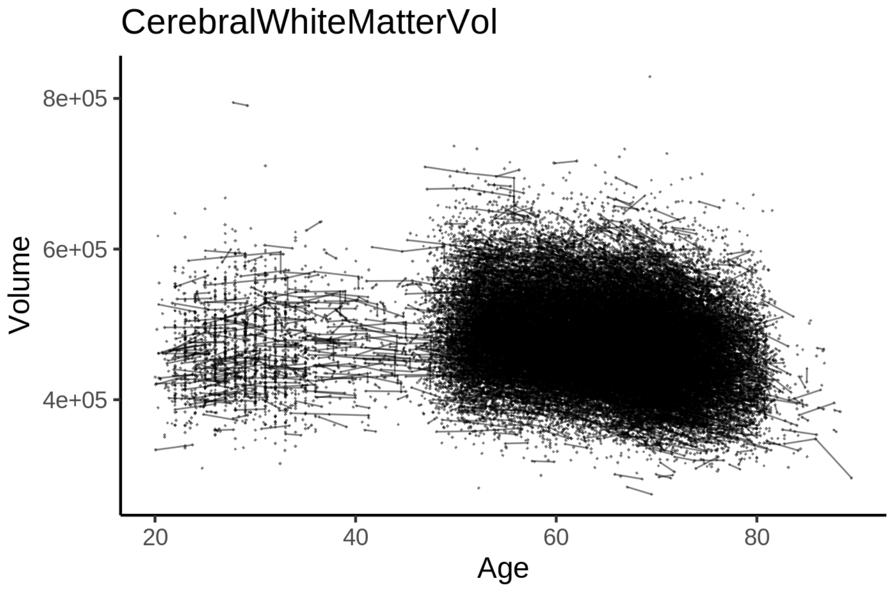


#### Model outputs

Model without sleep term

```
##
## Family: gaussian
## Link function: identity
##
## Formula:
## value ~ sex + site + icv + s(age_z, k = 10, bs = "cr")
## <environment: 0x55cf3e6fa040>
##
## Parametric coefficients:
##                 Estimate Std. Error t value Pr(>|t|)
## (Intercept)   449338.5     1636.3 274.614  < 2e-16 ***
## sexmale         8623.8      340.4  25.332  < 2e-16 ***
## siteMPIB       61328.7     2117.1  28.968  < 2e-16 ***
## siteousAvanto   2159.4     1662.2   1.299   0.1939
## siteousPrisma  -6244.7     2792.4  -2.236   0.0253 *
## siteousSkyra   -1602.0     1608.4  -0.996   0.3192
## siteUB        -11018.2     5028.0  -2.191   0.0284 *
## siteUCAM       -9203.8     1830.9  -5.027 5.00e-07 ***
## siteUKB        23318.6     1676.9  13.905  < 2e-16 ***
## siteUmU        13111.1     2359.3   5.557 2.75e-08 ***
```

```
## siteUOXF        1281.0     1997.6   0.641   0.5213
## icv            45591.8      170.8 266.939  < 2e-16 ***
## ---
## Signif. codes:  0 '***' 0.001 '**' 0.01 '*' 0.05 '.' 0.1 ' ' 1
##
## Approximate significance of smooth terms:
##            edf Ref.df     F p-value
## s(age_z) 8.333  8.333 879.2  <2e-16 ***
## ---
## Signif. codes:  0 '***' 0.001 '**' 0.01 '*' 0.05 '.' 0.1 ' ' 1
##
## R-sq.(adj) =  0.729
## lmer.REML = 1.1933e+06  Scale est. = 4.4856e+07  n = 51291
```

Model with only main effects of age and sleep

```
##
## Family: gaussian
## Link function: identity
##
## Formula:
## value ~ sex + site + icv + s(age_z, k = 10, bs = "cr") + s(sleep_z,
##     k = 5, bs = "cr")
## <environment: 0x55cf3e6fa040>
##
## Parametric coefficients:
##                 Estimate Std. Error t value Pr(>|t|)
## (Intercept)   449079.9     1637.6 274.238  < 2e-16 ***
## sexmale         8609.1      340.3  25.295  < 2e-16 ***
## siteMPIB       61682.7     2117.9  29.125  < 2e-16 ***
## siteousAvanto   2276.0     1662.3   1.369   0.1710
## siteousPrisma  -6156.0     2792.1  -2.205   0.0275 *
## siteousSkyra   -1490.6     1608.6  -0.927   0.3541
## siteUB        -10875.8     5026.4  -2.164   0.0305 *
## siteUCAM       -9043.8     1830.7  -4.940 7.83e-07 ***
## siteUKB        23595.7     1678.4  14.058  < 2e-16 ***
## siteUmU        13854.1     2364.0   5.860 4.65e-09 ***
## siteUOXF        1401.4     1997.4   0.702   0.4829
## icv            45602.9      170.9 266.828  < 2e-16 ***
## ---
## Signif. codes:  0 '***' 0.001 '**' 0.01 '*' 0.05 '.' 0.1 ' ' 1
##
## Approximate significance of smooth terms:
##              edf Ref.df      F  p-value
## s(age_z)   8.332  8.332 866.04  < 2e-16 ***
## s(sleep_z) 2.298  2.298  11.24 4.43e-06 ***
## ---
## Signif. codes:  0 '***' 0.001 '**' 0.01 '*' 0.05 '.' 0.1 ' ' 1
##
## R-sq.(adj) =  0.729
## lmer.REML = 1.1933e+06  Scale est. = 4.4883e+07  n = 51291
```

Model with full interaction between age and sleep

```
##
## Family: gaussian
```

```
## Link function: identity
##
## Formula:
## value ~ sex + site + icv + t2(age_z, sleep_z, k = c(10, 4), bs = "cr")
## <environment: 0x55cf3e6fa040>
##
## Parametric coefficients:
##                Estimate Std. Error t value Pr(>|t|)
## (Intercept)    448964.2     1648.7 272.306  < 2e-16 ***
## sexmale          8608.2      341.1  25.239  < 2e-16 ***
## siteMPIB        61800.1     2131.1  28.999  < 2e-16 ***
## siteousAvanto    2401.9     1673.8   1.435   0.1513
## siteousPrisma   -6066.4     2799.4  -2.167   0.0302 *
## siteousSkyra    -1344.6     1620.8  -0.830   0.4068
## siteUB         -10856.8     5031.3  -2.158   0.0309 *
## siteUCAM        -8967.2     1843.4  -4.865 1.15e-06 ***
## siteUKB         23713.9     1689.9  14.033  < 2e-16 ***
## siteUmU         13989.0     2374.3   5.892 3.84e-09 ***
## siteUOXF         1485.2     2008.9   0.739   0.4597
## icv             45607.4      170.9 266.881  < 2e-16 ***
## ---
## Signif. codes:  0 '***' 0.001 '**' 0.01 '*' 0.05 '.' 0.1 ' ' 1
##
## Approximate significance of smooth terms:
##                     edf Ref.df     F p-value
## t2(age_z,sleep_z) 15.51  15.51 42.11  <2e-16 ***
## ---
## Signif. codes:  0 '***' 0.001 '**' 0.01 '*' 0.05 '.' 0.1 ' ' 1
##
## R-sq.(adj) =  0.729
## lmer.REML = 1.1933e+06  Scale est. = 4.4888e+07  n = 51291
```

#### Model comparison

`mod_no_sleep` refers to model without sleep term, `mod_no_interaction` refers to model with only main effect of sleep, and `mod_full` refers to model with a full interaction between age and sleep. This is a nested model comparison, and the p-value at a given line refers to comparing the model at the line to the model on the line above. Hence, significance implies that the more complicated model is supported on statistical grounds.

To be even more specific, the p-value on the second row tests whether there is an association between sleep and volume. The p-value on the third row tests whether this association depends on age.

```
## Data: NULL
## Models:
## mod_list$mod_no_sleep$mer: NULL
## mod_list$mod_no_interaction$mer: NULL
## mod_list$mod_full$mer: NULL
##                                 npar     AIC     BIC  logLik deviance  Chisq Df Pr(>Chisq)
## mod_list$mod_no_sleep$mer         16 1193318 1193460 -596643  1193286
## mod_list$mod_no_interaction$mer   18 1193299 1193458 -596631  1193263 23.551  2  7.689e-06 ***
## mod_list$mod_full$mer             20 1193312 1193489 -596636  1193272  0.000  2          1
## ---
## Signif. codes:  0 '***' 0.001 '**' 0.01 '*' 0.05 '.' 0.1 ' ' 1
```

We chose the model based on the likelihood ratio test with 5 % significance level, which was `mod_no_interaction`.

##### Lifespan brain trajectory

The trajectory shown is from the chosen model `mod_no_interaction`.

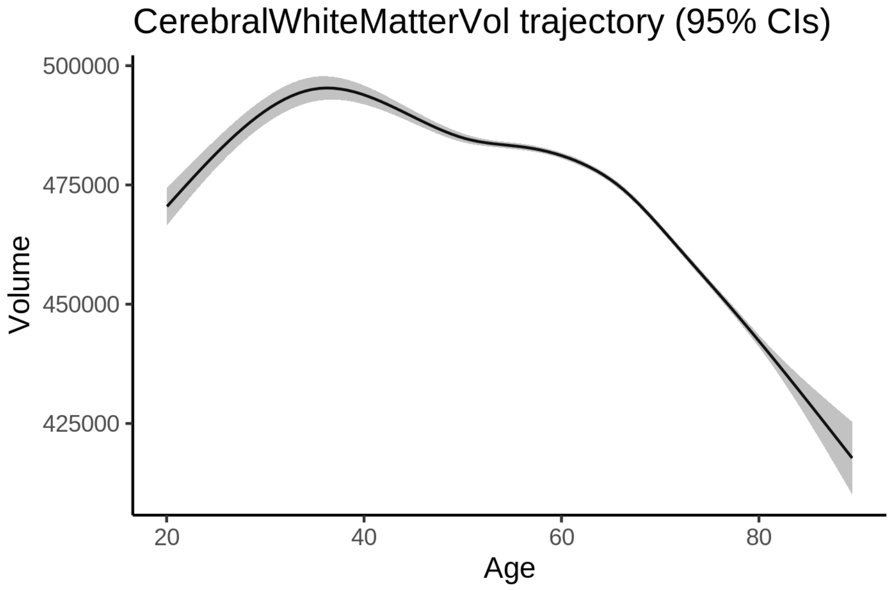


##### Effect of sleep

The chosen model only included the main effect of sleep, and hence the effect does not vary with age. The black dot shows the average sleep duration across all ages in the sample.

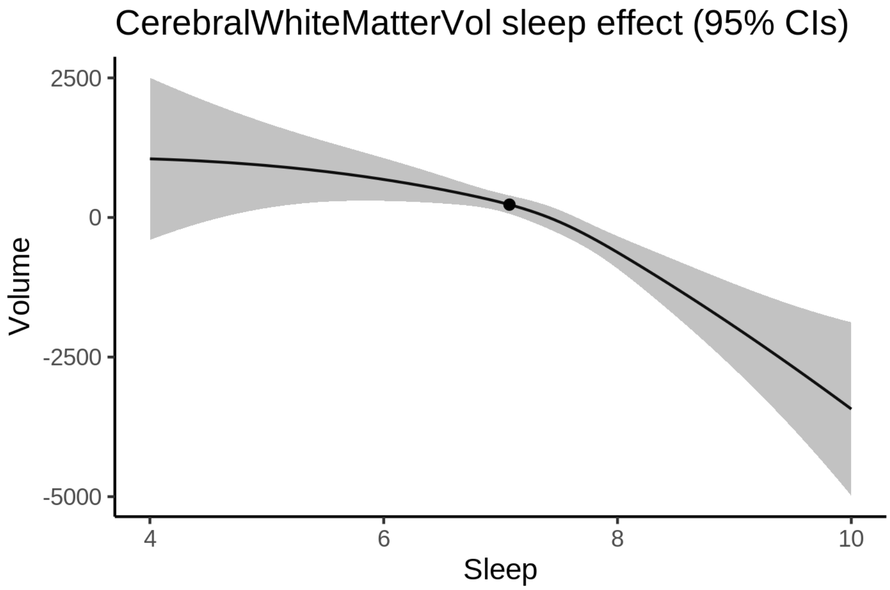


We also show the full interaction model for completeness, although it was not selected.

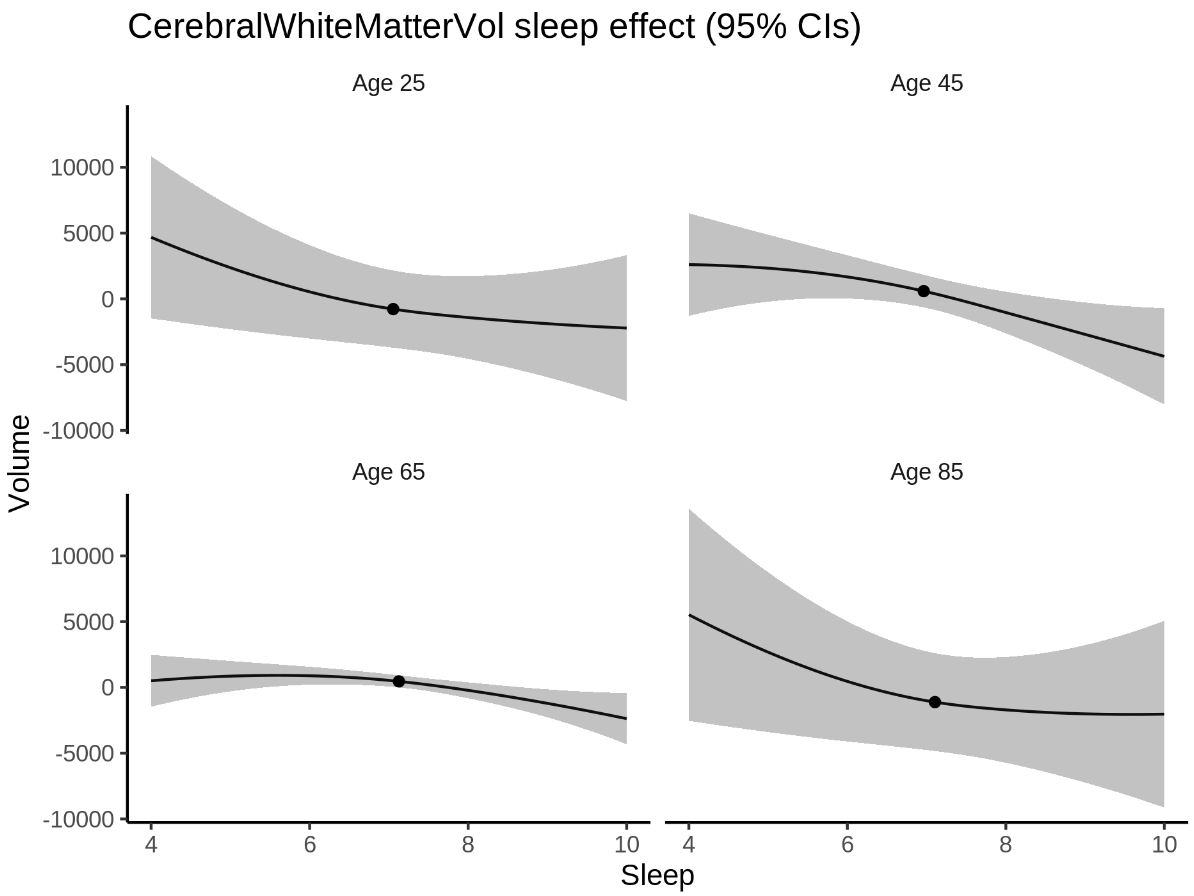


**Deviation from sleep associated with maximal volume**

Model with only main effect of sleep was chosen, so we show it for all ages at once. Maximum volume is attained at 4 hours of sleep. The percentage values in the plot are calculated as follows: The maximum at 100 % refers to a person at an arbitrary age with a sleep duration associated with maximum volume. For a female, this volume is 471970 and for a male it is 480579. The other percentage values show how large the expected volume is for someone with other sleep durations. For example, 99 % implies a 1 % reduction.

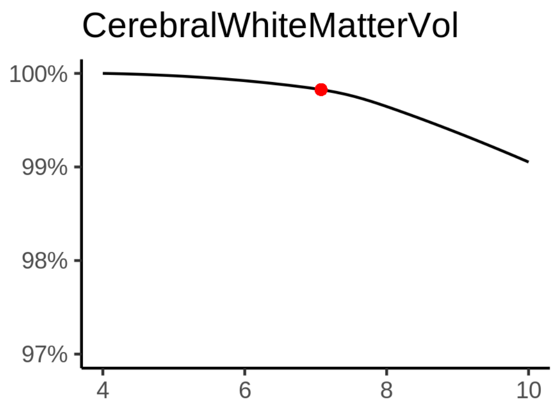

###### Comparison of mean sleep and sleep associated with maximum volume

A 95 % confidence interval for the sleep associated with maximum volume is [4, 6.06].

The plot below compares average sleep to the sleep associated with maximum volume.

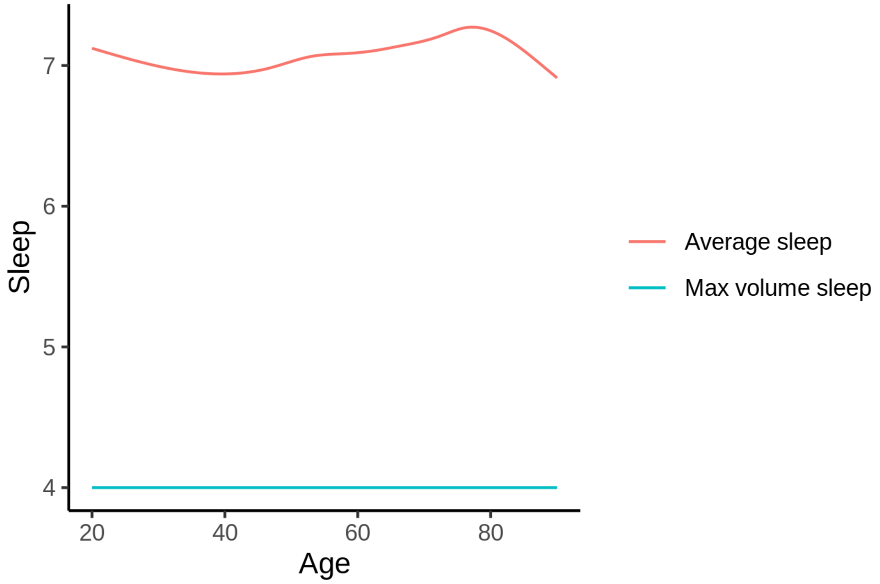


The next plot shows the difference between average sleep and sleep associated with maximum volume. The shaded region is a 95 % confidence interval.

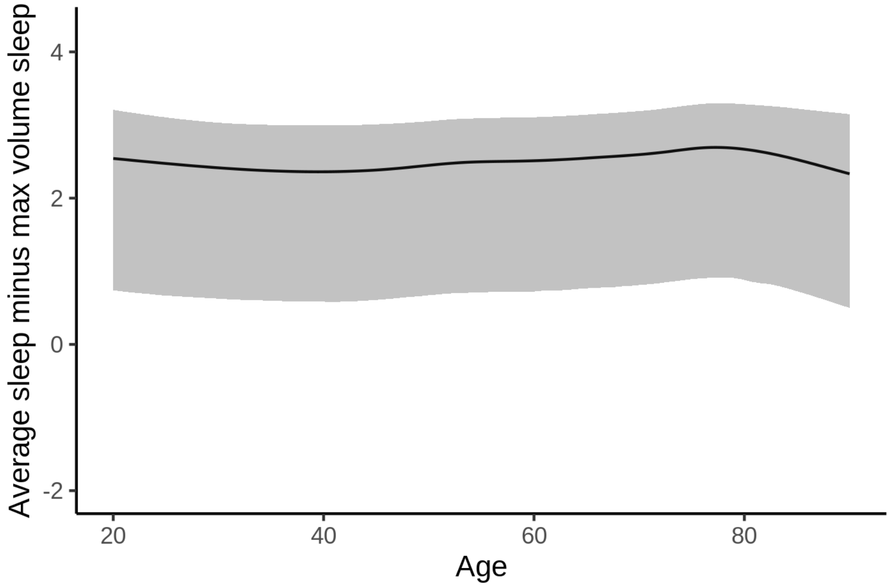


The next plot shows the probability that the sleep duration associated with maximum volume is longer than the average sleep duration, as a function of age. Probility below .05 can be interpreted as evidence that the sleep associated with maximum volume is shorter than the mean sleep, and probability above .95 can be interpreted the opposite way.

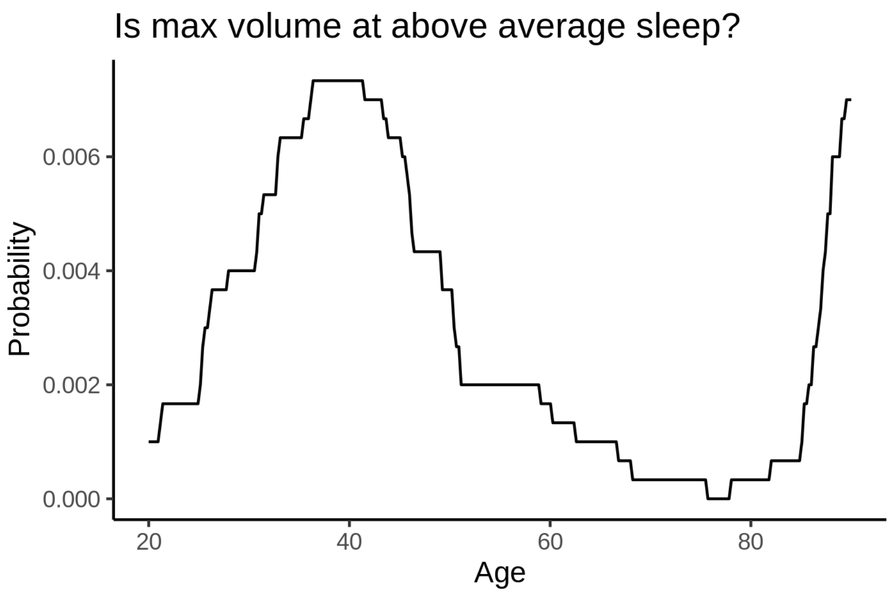


##### Controling for covariates

Below is the output for a model in which we only include data with income and education.

```
##
## Family: gaussian
## Link function: identity
##
## Formula:
## value ~ sex + site + s(age_z, k = 10, bs = "cr") + s(sleep_z,
##     k = 5, bs = "cr") + icv
## <environment: 0x55cf3c0f1868>
##
## Parametric coefficients:
##                Estimate Std. Error t value Pr(>|t|)
## (Intercept)    521949.5     3900.2  133.83   <2e-16 ***
## sexmale          7792.6      425.4   18.32   <2e-16 ***
## siteousAvanto  -77988.9     4526.8  -17.23   <2e-16 ***
## siteousPrisma  -74182.5     4731.4  -15.68   <2e-16 ***
## siteousSkyra   -75120.2     4198.2  -17.89   <2e-16 ***
## siteUKB        -48770.3     3893.6  -12.53   <2e-16 ***
## siteUOXF       -59368.6     4376.1  -13.57   <2e-16 ***
## icv             47017.2      218.0  215.66   <2e-16 ***
## ---
## Signif. codes:  0 ‘***’ 0.001 ‘**’ 0.01 ‘*’ 0.05 ‘.’ 0.1 ‘ ’ 1
##
## Approximate significance of smooth terms:
##              edf Ref.df       F p-value
## s(age_z)   7.754  7.754 475.792  <2e-16 ***
## s(sleep_z) 1.478  1.478   2.587  0.0555 .
## ---
## Signif. codes:  0 ‘***’ 0.001 ‘**’ 0.01 ‘*’ 0.05 ‘.’ 0.1 ‘ ’ 1
##
```

```
## R-sq.(adj) =  0.735
## lmer.REML = 7.2471e+05  Scale est. = 4.5032e+07  n = 31196
```

Below is the output for a model in which we control for the main effects of income and education.

```
##
## Family: gaussian
## Link function: identity
##
## Formula:
## value ~ sex + site + s(age_z, k = 10, bs = "cr") + s(sleep_z,
##     k = 5, bs = "cr") + icv + income_scaled + education_scaled
## <environment: 0x55cf3c0f1868>
##
## Parametric coefficients:
##                   Estimate Std. Error t value Pr(>|t|)
## (Intercept)       522573.0     3924.4 133.160   <2e-16 ***
## sexmale             7727.1      426.6  18.114   <2e-16 ***
## siteousAvanto     -77788.1     4527.8 -17.180   <2e-16 ***
## siteousPrisma     -74081.3     4731.3 -15.658   <2e-16 ***
## siteousSkyra      -74918.5     4199.1 -17.841   <2e-16 ***
## siteUKB           -48498.7     3895.8 -12.449   <2e-16 ***
## siteUOXF          -59441.5     4378.2 -13.577   <2e-16 ***
## icv                47052.2      219.9 214.002   <2e-16 ***
## income_scaled        464.4      506.5   0.917   0.3592
## education_scaled   -1386.4      599.5  -2.313   0.0207 *
## ---
## Signif. codes:  0 ‘***’ 0.001 ‘**’ 0.01 ‘*’ 0.05 ‘.’ 0.1 ‘ ’ 1
##
## Approximate significance of smooth terms:
##              edf Ref.df     F p-value
## s(age_z)   7.761  7.761 437.3  <2e-16 ***
## s(sleep_z) 1.563  1.563   2.3  0.0667 .
## ---
## Signif. codes:  0 ‘***’ 0.001 ‘**’ 0.01 ‘*’ 0.05 ‘.’ 0.1 ‘ ’ 1
##
## R-sq.(adj) =  0.735
## lmer.REML = 7.2468e+05  Scale est. = 4.5048e+07  n = 31196
```

We also included interaction effects between sleep duration and education and income, in another model. The output is shown below, and the interaction terms are `income_scaled:sleep_z` and `education_scaled:sleep_z`.

```
##
## Family: gaussian
## Link function: identity
##
## Formula:
## value ~ sex + site + s(age_z, k = 10, bs = "cr") + s(sleep_z,
##     k = 5, bs = "cr") + icv + income_scaled + education_scaled +
##     income_scaled:sleep_z + education_scaled:sleep_z
## <environment: 0x55cf3c0f1868>
##
## Parametric coefficients:
##                           Estimate Std. Error t value Pr(>|t|)
## (Intercept)               522642.1     3924.6 133.170   <2e-16 ***
```

```
## sexmale                    7743.5      426.9  18.140   <2e-16 ***
## siteousAvanto            -77799.4     4527.8 -17.182   <2e-16 ***
## siteousPrisma            -74054.8     4731.7 -15.651   <2e-16 ***
## siteousSkyra             -74929.1     4199.2 -17.844   <2e-16 ***
## siteUKB                  -48579.2     3896.1 -12.469   <2e-16 ***
## siteUOXF                 -59591.4     4379.4 -13.607   <2e-16 ***
## icv                       47053.1      219.9 213.983   <2e-16 ***
## income_scaled               443.8      506.9   0.876   0.3812
## education_scaled          -1382.7      599.6  -2.306   0.0211 *
## income_scaled:sleep_z       702.0      502.7   1.396   0.1626
## education_scaled:sleep_z    269.9      581.0   0.464   0.6423
## ---
## Signif. codes:  0 ‘***’ 0.001 ‘**’ 0.01 ‘*’ 0.05 ‘.’ 0.1 ‘ ’ 1
##
## Approximate significance of smooth terms:
##              edf Ref.df       F p-value
## s(age_z)   7.757  7.757 437.496  <2e-16 ***
## s(sleep_z) 1.649  1.649   1.772   0.108
## ---
## Signif. codes:  0 ‘***’ 0.001 ‘**’ 0.01 ‘*’ 0.05 ‘.’ 0.1 ‘ ’ 1
##
## R-sq.(adj) =  0.735
## lmer.REML = 7.2465e+05  Scale est. = 4.5049e+07  n = 31196
```

We did the same controling for BMI. Below is the model with no covariates but only keeping data with BMI.

```
##
## Family: gaussian
## Link function: identity
##
## Formula:
## value ~ sex + site + s(age_z, k = 10, bs = "cr") + s(sleep_z,
##     k = 5, bs = "cr") + icv
## <environment: 0x55cffd5ecaf0>
##
## Parametric coefficients:
##               Estimate Std. Error t value Pr(>|t|)
## (Intercept)   452757.0     2365.7 191.384  < 2e-16 ***
## sexmale         7085.3      408.9  17.329  < 2e-16 ***
## siteousPrisma  -6114.8     3145.7  -1.944   0.0519 .
## siteousSkyra    -470.0     1675.2  -0.281   0.7791
## siteUCAM      -10345.7     2519.3  -4.107 4.02e-05 ***
## siteUKB        20720.8     2390.0   8.670  < 2e-16 ***
## siteUmU        13205.8     2908.5   4.540 5.63e-06 ***
## icv            47511.1      210.1 226.096  < 2e-16 ***
## ---
## Signif. codes:  0 ‘***’ 0.001 ‘**’ 0.01 ‘*’ 0.05 ‘.’ 0.1 ‘ ’ 1
##
## Approximate significance of smooth terms:
##              edf Ref.df       F p-value
## s(age_z)   7.697  7.697 555.050  <2e-16 ***
## s(sleep_z) 1.002  1.002   5.707  0.0169 *
## ---
## Signif. codes:  0 ‘***’ 0.001 ‘**’ 0.01 ‘*’ 0.05 ‘.’ 0.1 ‘ ’ 1
##
```

```
## R-sq.(adj) =  0.742
## lmer.REML = 7.7586e+05  Scale est. = 4.3344e+07  n = 33449
```

Below is the model output with main effect.

```
##
## Family: gaussian
## Link function: identity
##
## Formula:
## value ~ sex + site + s(age_z, k = 10, bs = "cr") + s(sleep_z,
##     k = 5, bs = "cr") + icv + bmi
## <environment: 0x55cffd5ecaf0>
##
## Parametric coefficients:
##                Estimate Std. Error t value Pr(>|t|)
## (Intercept)   469696.67    2564.10 183.182  < 2e-16 ***
## sexmale         7615.37     408.21  18.655  < 2e-16 ***
## siteousPrisma  -5479.31    3138.56  -1.746   0.0809 .
## siteousSkyra    -426.20    1675.55  -0.254   0.7992
## siteUCAM       -9935.84    2512.80  -3.954 7.70e-05 ***
## siteUKB        21227.30    2384.67   8.902  < 2e-16 ***
## siteUmU        13945.37    2899.77   4.809 1.52e-06 ***
## icv            47609.94     209.25 227.528  < 2e-16 ***
## bmi             -671.05      39.65 -16.926  < 2e-16 ***
## ---
## Signif. codes:  0 ‘***’ 0.001 ‘**’ 0.01 ‘*’ 0.05 ‘.’ 0.1 ‘ ’ 1
##
## Approximate significance of smooth terms:
##              edf Ref.df      F p-value
## s(age_z)   7.695  7.695 563.32 < 2e-16 ***
## s(sleep_z) 1.000  1.000  10.01 0.00156 **
## ---
## Signif. codes:  0 ‘***’ 0.001 ‘**’ 0.01 ‘*’ 0.05 ‘.’ 0.1 ‘ ’ 1
##
## R-sq.(adj) =  0.745
## lmer.REML = 7.7557e+05  Scale est. = 4.3385e+07  n = 33449
```

Next is the model with BMI-sleep interaction.

```
##
## Family: gaussian
## Link function: identity
##
## Formula:
## value ~ sex + site + s(age_z, k = 10, bs = "cr") + s(sleep_z,
##     k = 5, bs = "cr") + icv + bmi + bmi:sleep_z
## <environment: 0x55cffd5ecaf0>
##
## Parametric coefficients:
##                Estimate Std. Error t value Pr(>|t|)
## (Intercept)   469730.54    2565.47 183.097  < 2e-16 ***
## sexmale         7614.71     408.22  18.653  < 2e-16 ***
## siteousPrisma  -5479.25    3138.58  -1.746   0.0809 .
## siteousSkyra    -428.75    1675.57  -0.256   0.7980
## siteUCAM       -9948.24    2513.01  -3.959 7.55e-05 ***
```

```
## siteUKB        21215.35    2384.88   8.896  < 2e-16 ***
## siteUmU        13934.63    2899.92   4.805 1.55e-06 ***
## icv            47609.37     209.26 227.517  < 2e-16 ***
## bmi             -671.59      39.67 -16.930  < 2e-16 ***
## bmi:sleep_z      -15.28      37.54  -0.407   0.6839
## ---
## Signif. codes:  0 '***' 0.001 '**' 0.01 '*' 0.05 '.' 0.1 ' ' 1
##
## Approximate significance of smooth terms:
##              edf Ref.df       F p-value
## s(age_z)   7.695  7.695 563.147  <2e-16 ***
## s(sleep_z) 1.000  1.000   0.016   0.899
## ---
## Signif. codes:  0 '***' 0.001 '**' 0.01 '*' 0.05 '.' 0.1 ' ' 1
##
## R-sq.(adj) =  0.745
## lmer.REML = 7.7556e+05  Scale est. = 4.3385e+07  n = 33449
```

We did the same controling for depression. Below is the model with no covariates but only keeping data with depression.

```
##
## Family: gaussian
## Link function: identity
##
## Formula:
## value ~ sex + site + s(age_z, k = 10, bs = "cr") + s(sleep_z,
##     k = 5, bs = "cr") + icv
## <environment: 0x55cf3f91e498>
##
## Parametric coefficients:
##                Estimate Std. Error t value Pr(>|t|)
## (Intercept)   502308.9     1973.3  254.55   <2e-16 ***
## sexmale         7244.3      413.3   17.53   <2e-16 ***
## siteousAvanto -46978.7     3847.4  -12.21   <2e-16 ***
## siteousPrisma -51554.6     3717.9  -13.87   <2e-16 ***
## siteousSkyra  -49103.8     2954.4  -16.62   <2e-16 ***
## siteUCAM      -61968.8     2709.1  -22.88   <2e-16 ***
## siteUKB       -28983.2     1951.7  -14.85   <2e-16 ***
## siteUmU       -36652.0     2644.1  -13.86   <2e-16 ***
## icv            47241.6      212.1  222.70   <2e-16 ***
## ---
## Signif. codes:  0 '***' 0.001 '**' 0.01 '*' 0.05 '.' 0.1 ' ' 1
##
## Approximate significance of smooth terms:
##              edf Ref.df     F p-value
## s(age_z)   7.703  7.703 545.5  <2e-16 ***
## s(sleep_z) 1.408  1.408   5.2  0.0301 *
## ---
## Signif. codes:  0 '***' 0.001 '**' 0.01 '*' 0.05 '.' 0.1 ' ' 1
##
## R-sq.(adj) =  0.736
## lmer.REML = 7.7393e+05  Scale est. = 4.2768e+07  n = 33361
```

Below is the model output with main effect.

```
##
## Family: gaussian
## Link function: identity
##
## Formula:
## value ~ sex + site + s(age_z, k = 10, bs = "cr") + s(sleep_z,
##     k = 5, bs = "cr") + icv + depression
## <environment: 0x55cf3f91e498>
##
## Parametric coefficients:
##               Estimate Std. Error t value Pr(>|t|)
## (Intercept)   503027.8     1981.5 253.865  < 2e-16 ***
## sexmale         7170.9      413.6  17.338  < 2e-16 ***
## siteousAvanto -46607.7     3847.5 -12.114  < 2e-16 ***
## siteousPrisma -51218.5     3718.0 -13.776  < 2e-16 ***
## siteousSkyra  -48688.6     2955.6 -16.473  < 2e-16 ***
## siteUCAM      -61794.6     2708.9 -22.812  < 2e-16 ***
## siteUKB       -29282.9     1952.8 -14.995  < 2e-16 ***
## siteUmU       -34490.8     2699.0 -12.779  < 2e-16 ***
## icv            47251.0      212.0 222.837  < 2e-16 ***
## depression     -5826.8     1467.3  -3.971 7.17e-05 ***
## ---
## Signif. codes:  0 ‘***’ 0.001 ‘**’ 0.01 ‘*’ 0.05 ‘.’ 0.1 ‘ ’ 1
##
## Approximate significance of smooth terms:
##            edf Ref.df       F p-value
## s(age_z)   7.7    7.7 547.274 < 2e-16 ***
## s(sleep_z) 1.0    1.0   7.296 0.00692 **
## ---
## Signif. codes:  0 ‘***’ 0.001 ‘**’ 0.01 ‘*’ 0.05 ‘.’ 0.1 ‘ ’ 1
##
## R-sq.(adj) =  0.736
## lmer.REML = 7.7389e+05  Scale est. = 4.2739e+07  n = 33361
```

Next is the model with depression-sleep interaction.

```
##
## Family: gaussian
## Link function: identity
##
## Formula:
## value ~ sex + site + s(age_z, k = 10, bs = "cr") + s(sleep_z,
##     k = 5, bs = "cr") + icv + depression + depression:sleep_z
## <environment: 0x55cf3f91e498>
##
## Parametric coefficients:
##                    Estimate Std. Error t value Pr(>|t|)
## (Intercept)        503027.6     1981.5 253.860  < 2e-16 ***
## sexmale              7169.3      413.7  17.331  < 2e-16 ***
## siteousAvanto      -46618.2     3847.8 -12.115  < 2e-16 ***
## siteousPrisma      -51228.9     3718.4 -13.777  < 2e-16 ***
## siteousSkyra       -48698.7     2956.0 -16.474  < 2e-16 ***
## siteUCAM           -61805.8     2709.4 -22.811  < 2e-16 ***
## siteUKB            -29282.2     1952.9 -14.994  < 2e-16 ***
## siteUmU            -34415.5     2721.0 -12.648  < 2e-16 ***
```

```
## icv                  47251.2      212.0 222.832  < 2e-16 ***
## depression           -5844.7     1469.6  -3.977 6.99e-05 ***
## depression:sleep_z    -233.8     1072.4  -0.218    0.827
## ---
## Signif. codes:  0 '***' 0.001 '**' 0.01 '*' 0.05 '.' 0.1 ' ' 1
##
## Approximate significance of smooth terms:
##               edf Ref.df       F p-value
## s(age_z)    7.699  7.699 547.349  <2e-16 ***
## s(sleep_z) 1.000  1.000   4.682  0.0305 *
## ---
## Signif. codes:  0 '***' 0.001 '**' 0.01 '*' 0.05 '.' 0.1 ' ' 1
##
## R-sq.(adj) =  0.736
## lmer.REML = 7.7388e+05  Scale est. = 4.2737e+07  n = 33361
```

The plot below shows the sleep-volume curve for the original model and for the model with main effects of SES.

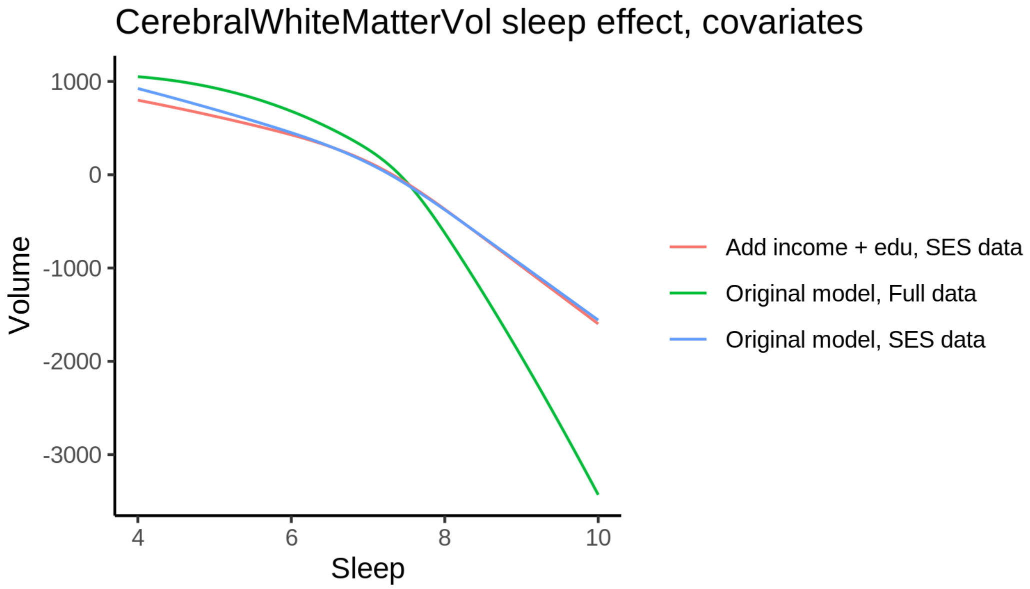


The plot below shows the sleep-volume curve for the original model and for the model with main effects of BMI.

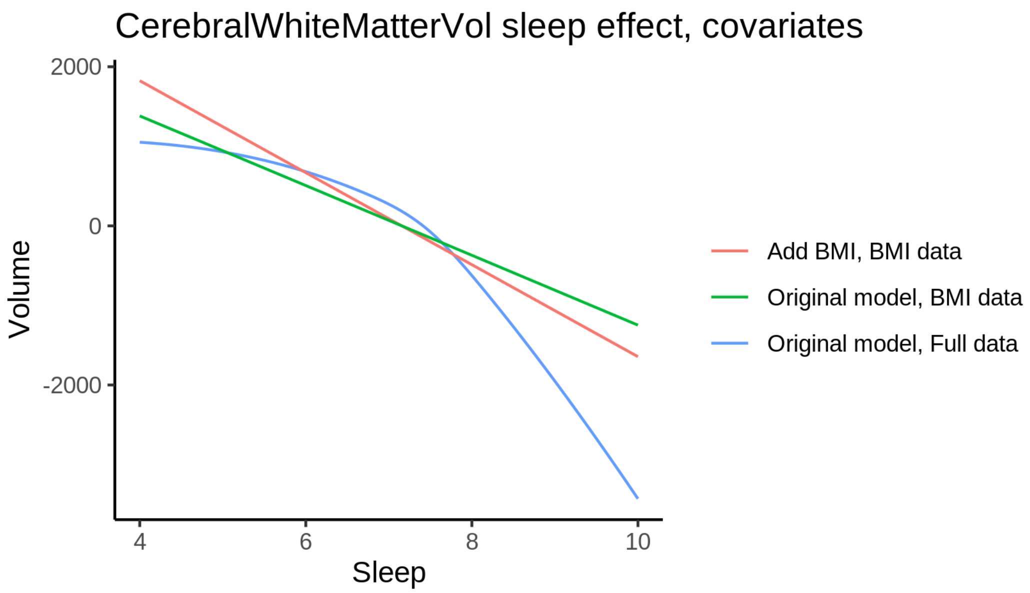


The plot below shows the sleep-volume curve for the original model and for the model with main effects of depression.

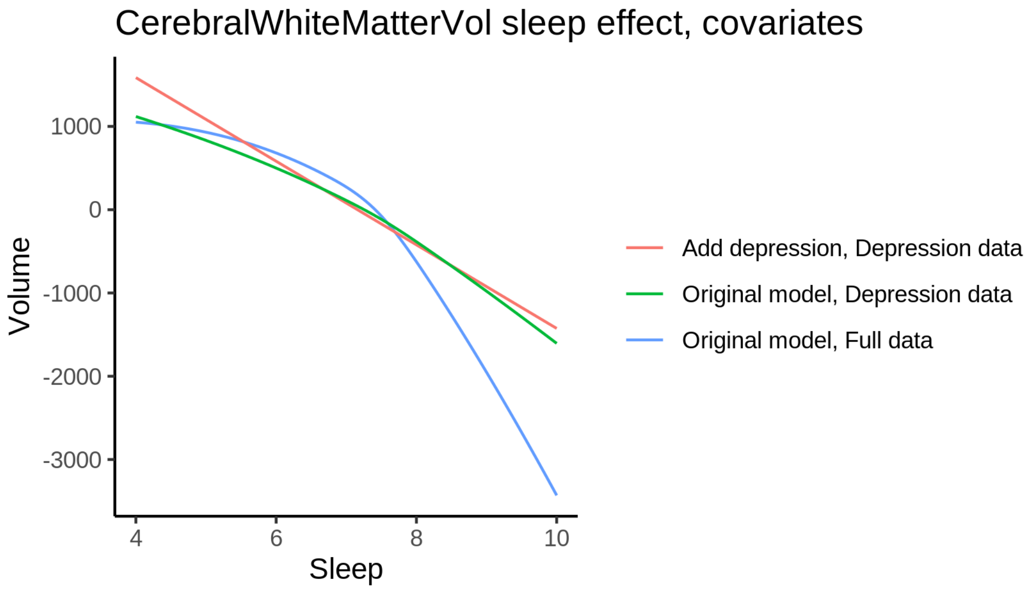


#### EstimatedTotalIntraCranialVol

##### Descriptive statistics

| Study | Observations | Unique IDs | Mean age | Age range |
|---|---|---|---|---|
| HCP | 974 | 974 | 28.8 | 22 - 37 |
| MPIB | 677 | 391 | 63.1 | 24 - 83 |
| UB | 113 | 39 | 70.9 | 64 - 81 |
| UCAM | 884 | 632 | 55.1 | 20 - 88 |

| Study | Observations | Unique IDs | Mean age | Age range |
|---|---|---|---|---|
| UiO | 1475 | 803 | 49.4 | 20 - 89 |
| UKB | 45985 | 43138 | 64.5 | 45 - 83 |
| UmU | 423 | 284 | 62.3 | 25 - 85 |
| UOXF | 769 | 769 | 69.8 | 60 - 85 |

#### Spaghetti plot

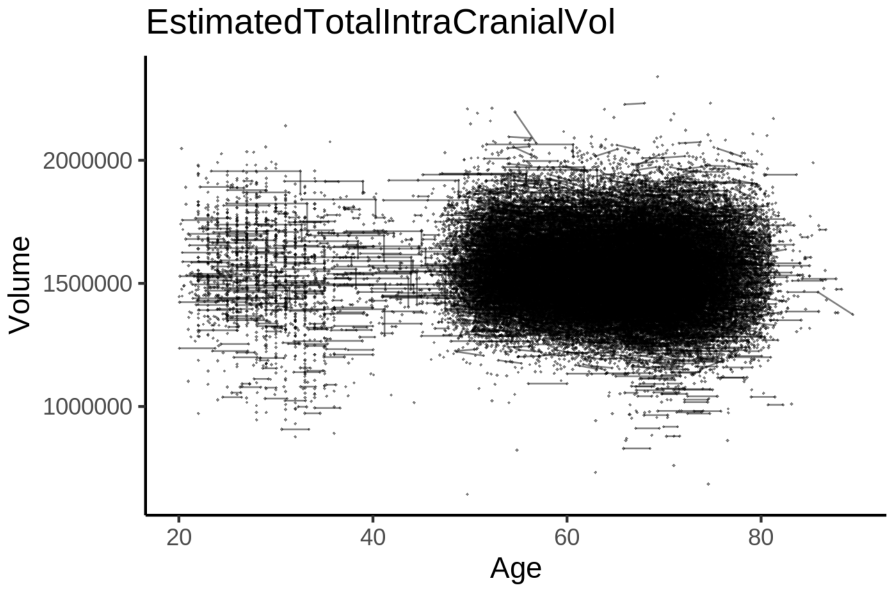


#### Model outputs

Model without sleep term

```
##
## Family: gaussian
## Link function: identity
##
## Formula:
## value ~ sex + site + s(age_z, k = 10, bs = "cr")
## <environment: 0x55cf39dd06a0>
##
## Parametric coefficients:
##                Estimate Std. Error t value Pr(>|t|)
## (Intercept)     1443926       5748 251.225  < 2e-16 ***
## sexmale          180501       1191 151.590  < 2e-16 ***
## siteMPIB        -245302       8373 -29.297  < 2e-16 ***
## siteousAvanto     66330       6468  10.255  < 2e-16 ***
## siteousPrisma     97934       9270  10.565  < 2e-16 ***
## siteousSkyra      10193       6459   1.578  0.11455
## siteUB           -33150      21269  -1.559  0.11910
## siteUCAM          19052       7207   2.644  0.00821 **
## siteUKB           19270       5862   3.287  0.00101 **
## siteUmU          -49797       9395  -5.301 1.16e-07 ***
```

```
## siteUOXF        -22773       7479  -3.045  0.00233 **
## ---
## Signif. codes:  0 ‘***’ 0.001 ‘**’ 0.01 ‘*’ 0.05 ‘.’ 0.1 ‘ ’ 1
##
## Approximate significance of smooth terms:
##            edf Ref.df     F p-value
## s(age_z) 6.833  6.833 78.33  <2e-16 ***
## ---
## Signif. codes:  0 ‘***’ 0.001 ‘**’ 0.01 ‘*’ 0.05 ‘.’ 0.1 ‘ ’ 1
##
## R-sq.(adj) =   0.35
## lmer.REML = 1.335e+06  Scale est. = 1.5233e+08  n = 51300
```

Model with only main effects of age and sleep

```
##
## Family: gaussian
## Link function: identity
##
## Formula:
## value ~ sex + site + s(age_z, k = 10, bs = "cr") + s(sleep_z,
##     k = 5, bs = "cr")
## <environment: 0x55cf39dd06a0>
##
## Parametric coefficients:
##               Estimate Std. Error t value Pr(>|t|)
## (Intercept)    1446829       5751 251.582  < 2e-16 ***
## sexmale         180118       1190 151.356  < 2e-16 ***
## siteMPIB       -246617       8368 -29.472  < 2e-16 ***
## siteousAvanto    64422       6464   9.966  < 2e-16 ***
## siteousPrisma    96164       9264  10.380  < 2e-16 ***
## siteousSkyra      8246       6455   1.277  0.20144
## siteUB          -32215      21245  -1.516  0.12943
## siteUCAM         18476       7200   2.566  0.01029 *
## siteUKB          16403       5866   2.796  0.00517 **
## siteUmU         -53634       9404  -5.703 1.18e-08 ***
## siteUOXF        -23837       7473  -3.190  0.00143 **
## ---
## Signif. codes:  0 ‘***’ 0.001 ‘**’ 0.01 ‘*’ 0.05 ‘.’ 0.1 ‘ ’ 1
##
## Approximate significance of smooth terms:
##              edf Ref.df     F p-value
## s(age_z)   6.860  6.860 77.46  <2e-16 ***
## s(sleep_z) 3.615  3.615 30.13  <2e-16 ***
## ---
## Signif. codes:  0 ‘***’ 0.001 ‘**’ 0.01 ‘*’ 0.05 ‘.’ 0.1 ‘ ’ 1
##
## R-sq.(adj) =  0.352
## lmer.REML = 1.3349e+06  Scale est. = 1.5234e+08  n = 51300
```

Model with full interaction between age and sleep

```
##
## Family: gaussian
## Link function: identity
##
```

```
## Formula:
## value ~ sex + site + t2(age_z, sleep_z, k = c(10, 4), bs = "cr")
## <environment: 0x55cf39dd06a0>
##
## Parametric coefficients:
##               Estimate Std. Error t value Pr(>|t|)
## (Intercept)    1447957       5818 248.868  < 2e-16 ***
## sexmale         180559       1192 151.472  < 2e-16 ***
## siteMPIB       -249581       8421 -29.637  < 2e-16 ***
## siteousAvanto    63757       6510   9.793  < 2e-16 ***
## siteousPrisma    95052       9301  10.219  < 2e-16 ***
## siteousSkyra      7419       6504   1.141 0.253957
## siteUB          -36174      21274  -1.700 0.089068 .
## siteUCAM         15824       7255   2.181 0.029181 *
## siteUKB          15083       5936   2.541 0.011052 *
## siteUmU         -55780       9450  -5.902 3.61e-09 ***
## siteUOXF        -26301       7537  -3.490 0.000484 ***
## ---
## Signif. codes:  0 '***' 0.001 '**' 0.01 '*' 0.05 '.' 0.1 ' ' 1
##
## Approximate significance of smooth terms:
##                     edf Ref.df     F  p-value
## t2(age_z,sleep_z) 13.2   13.2 3.727 2.06e-05 ***
## ---
## Signif. codes:  0 '***' 0.001 '**' 0.01 '*' 0.05 '.' 0.1 ' ' 1
##
## R-sq.(adj) =  0.351
## lmer.REML = 1.3349e+06  Scale est. = 1.5246e+08  n = 51300
```

#### Model comparison

`mod_no_sleep` refers to model without sleep term, `mod_no_interaction` refers to model with only main effect of sleep, and `mod_full` refers to model with a full interaction between age and sleep. This is a nested model comparison, and the p-value at a given line refers to comparing the model at the line to the model on the line above. Hence, significance implies that the more complicated model is supported on statistical grounds.

To be even more specific, the p-value on the second row tests whether there is an association between sleep and volume. The p-value on the third row tests whether this association depends on age.

```
## Data: NULL
## Models:
## mod_list$mod_no_sleep$mer: NULL
## mod_list$mod_no_interaction$mer: NULL
## mod_list$mod_full$mer: NULL
##                                 npar     AIC     BIC  logLik deviance  Chisq Df Pr(>Chisq)
## mod_list$mod_no_sleep$mer         15 1335031 1335164 -667501  1335001
## mod_list$mod_no_interaction$mer   17 1334934 1335084 -667450  1334900 101.77  2     <2e-16 ***
## mod_list$mod_full$mer             19 1334976 1335144 -667469  1334938   0.00  2          1
## ---
## Signif. codes:  0 '***' 0.001 '**' 0.01 '*' 0.05 '.' 0.1 ' ' 1
```

We chose the model based on the likelihood ratio test with 5 % significance level, which was `mod_no_interaction`.

##### Lifespan brain trajectory

The trajectory shown is from the chosen model `mod_no_interaction`.

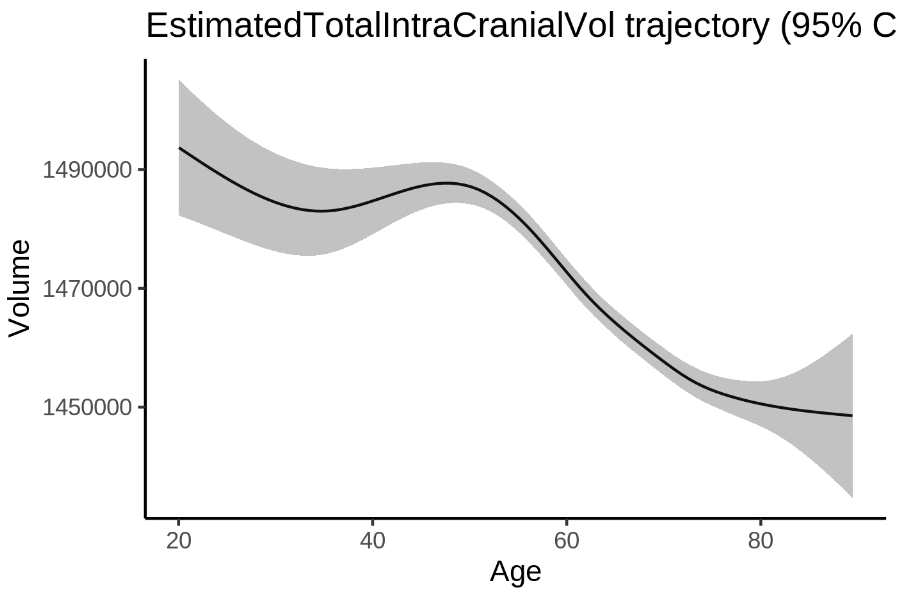


##### Effect of sleep

The chosen model only included the main effect of sleep, and hence the effect does not vary with age. The black dot shows the average sleep duration across all ages in the sample.

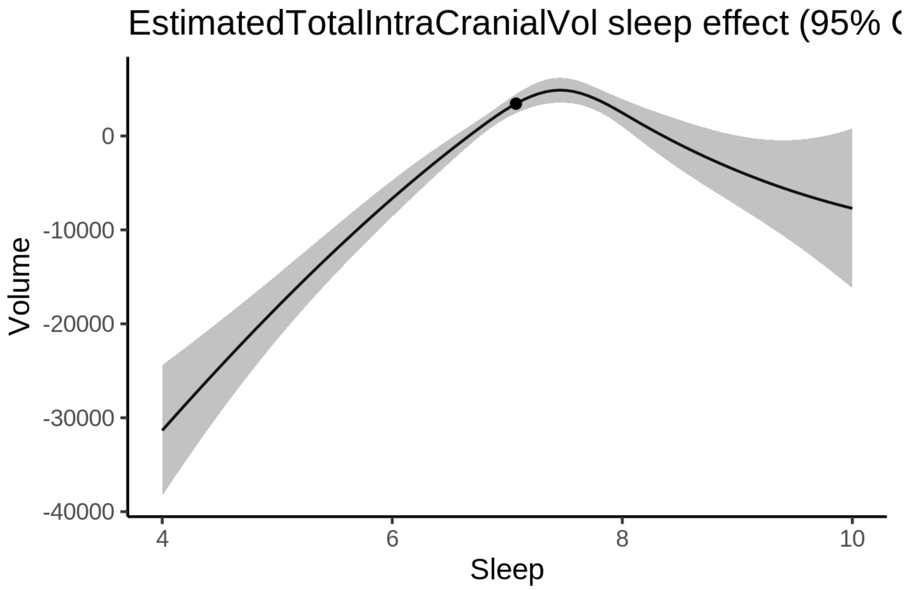


We also show the full interaction model for completeness, although it was not selected.

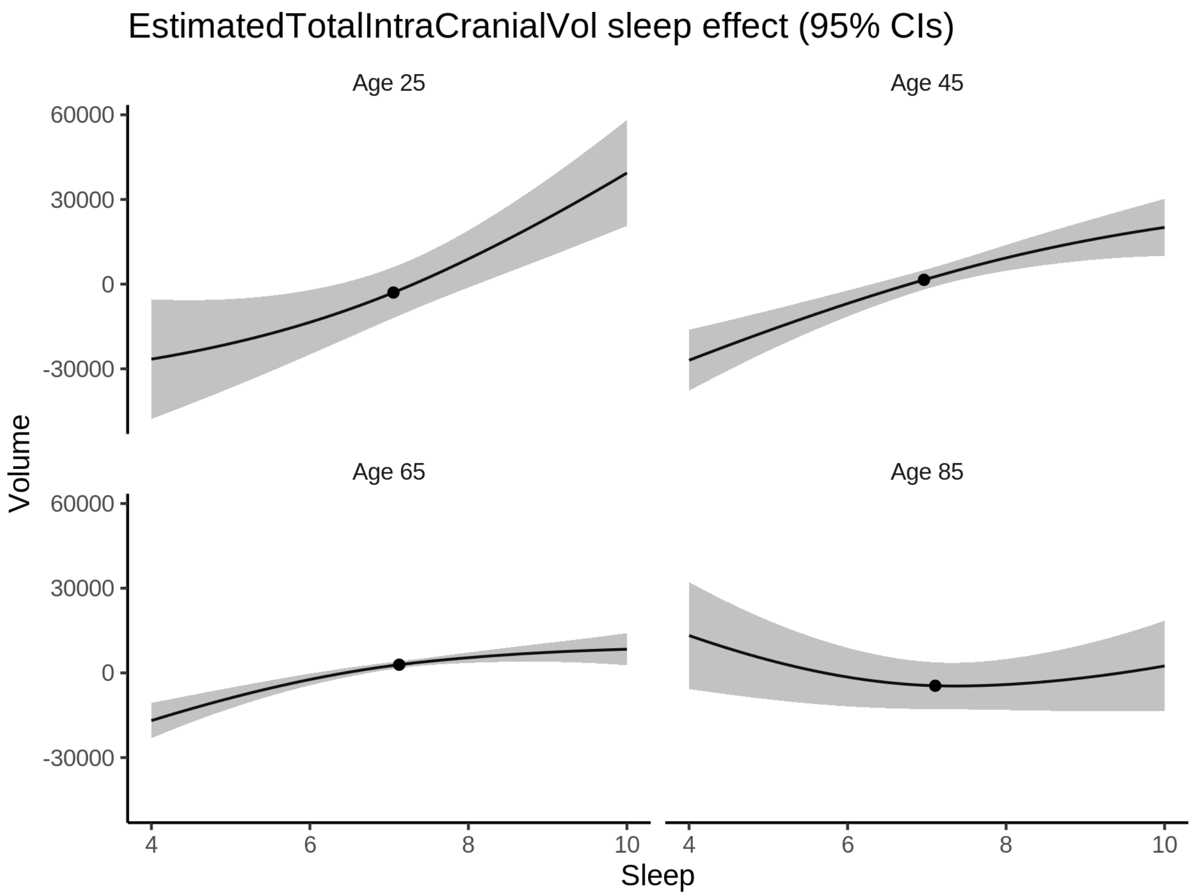


**Deviation from sleep associated with maximal volume**

Model with only main effect of sleep was chosen, so we show it for all ages at once. Maximum volume is attained at 7.5 hours of sleep. The percentage values in the plot are calculated as follows: The maximum at 100 % refers to a person at an arbitrary age with a sleep duration associated with maximum volume. For a female, this volume is 1472474 and for a male it is 1652592. The other percentage values show how large the expected volume is for someone with other sleep durations. For example, 99 % implies a 1 % reduction.

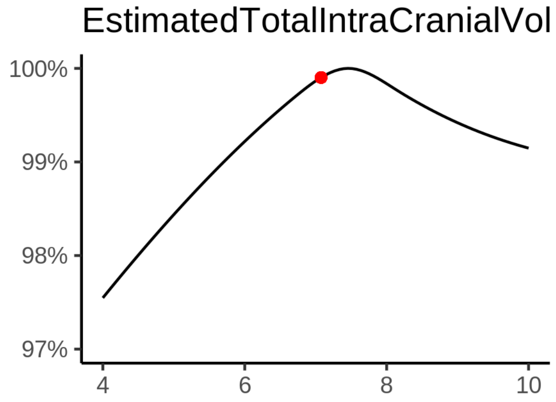

**Comparison of mean sleep and sleep associated with maximum volume**

A 95 % confidence interval for the sleep associated with maximum volume is [7.39, 7.58].

The plot below compares average sleep to the sleep associated with maximum volume.

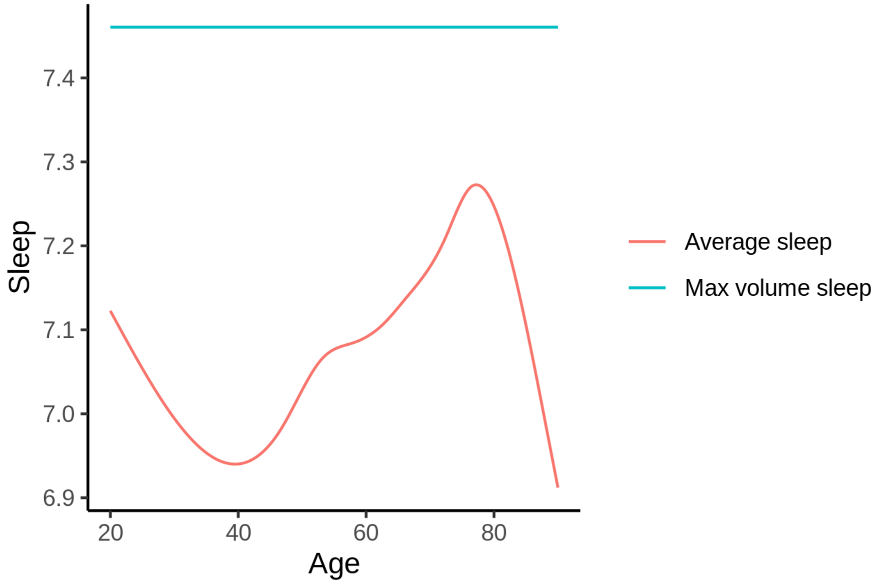


The next plot shows the difference between average sleep and sleep associated with maximum volume. The shaded region is a 95 % confidence interval.

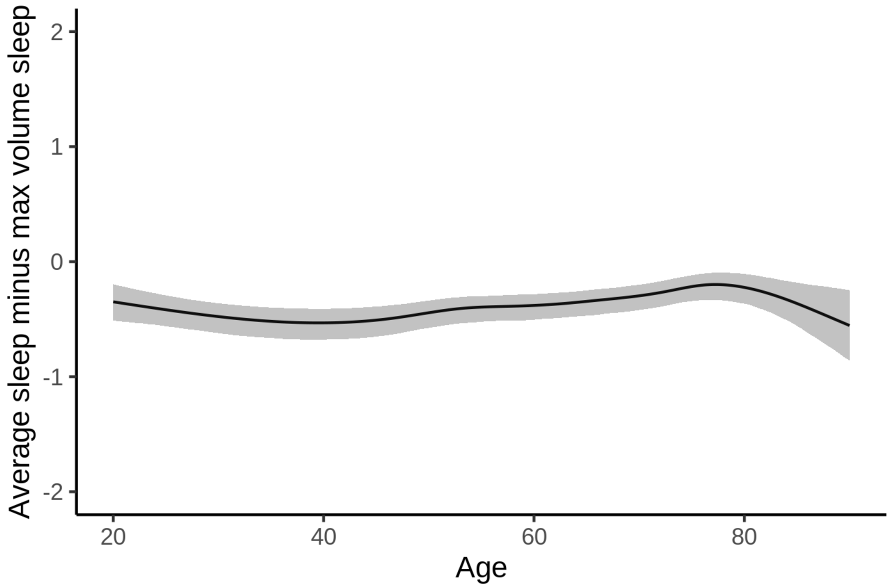


The next plot shows the probability that the sleep duration associated with maximum volume is longer than the average sleep duration, as a function of age. Probility below .05 can be interpreted as evidence that the sleep associated with maximum volume is shorter than the mean sleep, and probability above .95 can be interpreted the opposite way.

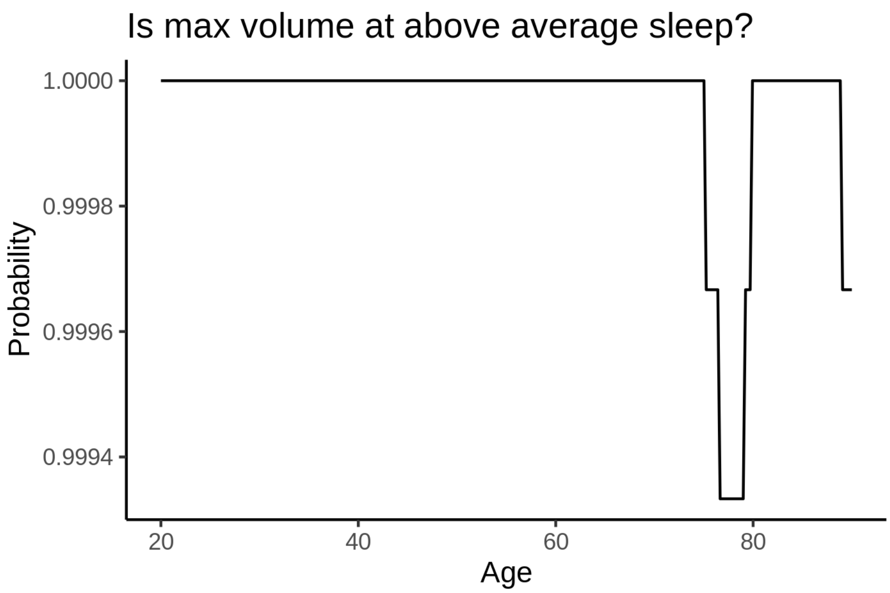


##### Controling for covariates

Below is the output for a model in which we only include data with income and education.

```
##
## Family: gaussian
## Link function: identity
##
## Formula:
## value ~ sex + site + s(age_z, k = 10, bs = "cr") + s(sleep_z,
##     k = 5, bs = "cr")
## <environment: 0x55cff9aaee40>
##
## Parametric coefficients:
##                Estimate Std. Error t value Pr(>|t|)
## (Intercept)     1170189      16243   72.04   <2e-16 ***
## sexmale          177837       1484  119.86   <2e-16 ***
## siteousAvanto    330643      18015   18.35   <2e-16 ***
## siteousPrisma    338578      19048   17.77   <2e-16 ***
## siteousSkyra     266856      17662   15.11   <2e-16 ***
## siteUKB          297548      16250   18.31   <2e-16 ***
## siteUOXF         217283      18377   11.82   <2e-16 ***
## ---
## Signif. codes:  0 ‘***’ 0.001 ‘**’ 0.01 ‘*’ 0.05 ‘.’ 0.1 ‘ ’ 1
##
## Approximate significance of smooth terms:
##              edf Ref.df     F p-value
## s(age_z)   6.580  6.580 63.34  <2e-16 ***
## s(sleep_z) 3.619  3.619 19.82  <2e-16 ***
## ---
## Signif. codes:  0 ‘***’ 0.001 ‘**’ 0.01 ‘*’ 0.05 ‘.’ 0.1 ‘ ’ 1
##
## R-sq.(adj) =  0.339
```

```
## lmer.REML = 8.1142e+05  Scale est. = 1.6633e+08  n = 31198
```

Below is the output for a model in which we control for the main effects of income and education.

```
##
## Family: gaussian
## Link function: identity
##
## Formula:
## value ~ sex + site + s(age_z, k = 10, bs = "cr") + s(sleep_z,
##     k = 5, bs = "cr") + income_scaled + education_scaled
## <environment: 0x55cff9aaee40>
##
## Parametric coefficients:
##                  Estimate Std. Error t value Pr(>|t|)
## (Intercept)       1137576      16159   70.40   <2e-16 ***
## sexmale            175337       1482  118.34   <2e-16 ***
## siteousAvanto      326634      17882   18.27   <2e-16 ***
## siteousPrisma      331656      18908   17.54   <2e-16 ***
## siteousSkyra       262689      17524   14.99   <2e-16 ***
## siteUKB            291247      16107   18.08   <2e-16 ***
## siteUOXF           225307      18212   12.37   <2e-16 ***
## income_scaled       24950       2114   11.80   <2e-16 ***
## education_scaled    36120       2525   14.31   <2e-16 ***
## ---
## Signif. codes:  0 '***' 0.001 '**' 0.01 '*' 0.05 '.' 0.1 ' ' 1
##
## Approximate significance of smooth terms:
##              edf Ref.df     F p-value
## s(age_z)   5.906  5.906 42.96  <2e-16 ***
## s(sleep_z) 3.349  3.349 12.77  <2e-16 ***
## ---
## Signif. codes:  0 '***' 0.001 '**' 0.01 '*' 0.05 '.' 0.1 ' ' 1
##
## R-sq.(adj) =   0.35
## lmer.REML = 8.1092e+05  Scale est. = 1.6785e+08  n = 31198
```

We also included interaction effects between sleep duration and education and income, in another model. The output is shown below, and the interaction terms are `income_scaled:sleep_z` and `education_scaled:sleep_z`.

```
##
## Family: gaussian
## Link function: identity
##
## Formula:
## value ~ sex + site + s(age_z, k = 10, bs = "cr") + s(sleep_z,
##     k = 5, bs = "cr") + income_scaled + education_scaled + income_scaled:sleep_z +
##     education_scaled:sleep_z
## <environment: 0x55cff9aaee40>
##
## Parametric coefficients:
##                          Estimate Std. Error t value Pr(>|t|)
## (Intercept)               1137216      16159  70.378   <2e-16 ***
## sexmale                    175397       1483 118.288   <2e-16 ***
## siteousAvanto              326930      17881  18.283   <2e-16 ***
```

```
## siteousPrisma              332110      18908  17.564   <2e-16 ***
## siteousSkyra               262985      17524  15.007   <2e-16 ***
## siteUKB                    291707      16107  18.110   <2e-16 ***
## siteUOXF                   226094      18216  12.412   <2e-16 ***
## income_scaled               24872       2115  11.761   <2e-16 ***
## education_scaled            36199       2525  14.336   <2e-16 ***
## income_scaled:sleep_z        1292       2142   0.603   0.5465
## education_scaled:sleep_z    -5869       2460  -2.386   0.0171 *
## ---
## Signif. codes:  0 '***' 0.001 '**' 0.01 '*' 0.05 '.' 0.1 ' ' 1
##
## Approximate significance of smooth terms:
##              edf Ref.df     F  p-value
## s(age_z)   5.914  5.914 43.03  < 2e-16 ***
## s(sleep_z) 3.342  3.342 10.82 5.99e-08 ***
## ---
## Signif. codes:  0 '***' 0.001 '**' 0.01 '*' 0.05 '.' 0.1 ' ' 1
##
## R-sq.(adj) =   0.35
## lmer.REML = 8.1088e+05  Scale est. = 1.6783e+08  n = 31198
```

We did the same controling for BMI. Below is the model with no covariates but only keeping data with BMI.

```
##
## Family: gaussian
## Link function: identity
##
## Formula:
## value ~ sex + site + s(age_z, k = 10, bs = "cr") + s(sleep_z,
##     k = 5, bs = "cr")
## <environment: 0x55cf42c46d98>
##
## Parametric coefficients:
##                Estimate Std. Error t value Pr(>|t|)
## (Intercept)    1503984       7265 207.027  < 2e-16 ***
## sexmale         178117       1430 124.570  < 2e-16 ***
## siteousPrisma    28500       8342   3.416 0.000636 ***
## siteousSkyra    -61544       3176 -19.378  < 2e-16 ***
## siteUCAM        -39016       8475  -4.604 4.16e-06 ***
## siteUKB         -36357       7359  -4.941 7.83e-07 ***
## siteUmU        -112001      10365 -10.805  < 2e-16 ***
## ---
## Signif. codes:  0 '***' 0.001 '**' 0.01 '*' 0.05 '.' 0.1 ' ' 1
##
## Approximate significance of smooth terms:
##              edf Ref.df     F p-value
## s(age_z)   6.208  6.208 73.14  <2e-16 ***
## s(sleep_z) 3.507  3.507 23.24  <2e-16 ***
## ---
## Signif. codes:  0 '***' 0.001 '**' 0.01 '*' 0.05 '.' 0.1 ' ' 1
##
## R-sq.(adj) =  0.338
## lmer.REML = 8.6864e+05  Scale est. = 1.4743e+08  n = 33453
```

Below is the model output with main effect.

```
##
## Family: gaussian
## Link function: identity
##
## Formula:
## value ~ sex + site + s(age_z, k = 10, bs = "cr") + s(sleep_z,
##     k = 5, bs = "cr") + bmi
## <environment: 0x55cf42c46d98>
##
## Parametric coefficients:
##                Estimate Std. Error t value Pr(>|t|)
## (Intercept)   1482076.0     8408.4 176.261  < 2e-16 ***
## sexmale        177266.3     1438.7 123.214  < 2e-16 ***
## siteousPrisma   28222.8     8341.9   3.383 0.000717 ***
## siteousSkyra   -61511.1     3176.0 -19.367  < 2e-16 ***
## siteUCAM       -39578.2     8472.2  -4.672 3.00e-06 ***
## siteUKB        -37344.3     7358.1  -5.075 3.89e-07 ***
## siteUmU       -113078.3    10363.2 -10.912  < 2e-16 ***
## bmi               882.5      170.8   5.167 2.39e-07 ***
## ---
## Signif. codes:  0 '***' 0.001 '**' 0.01 '*' 0.05 '.' 0.1 ' ' 1
##
## Approximate significance of smooth terms:
##              edf Ref.df     F p-value
## s(age_z)   6.187  6.187 73.18  <2e-16 ***
## s(sleep_z) 3.539  3.539 25.12  <2e-16 ***
## ---
## Signif. codes:  0 '***' 0.001 '**' 0.01 '*' 0.05 '.' 0.1 ' ' 1
##
## R-sq.(adj) =  0.338
## lmer.REML = 8.686e+05  Scale est. = 1.4745e+08  n = 33453
```

Next is the model with BMI-sleep interaction.

```
##
## Family: gaussian
## Link function: identity
##
## Formula:
## value ~ sex + site + s(age_z, k = 10, bs = "cr") + s(sleep_z,
##     k = 5, bs = "cr") + bmi + bmi:sleep_z
## <environment: 0x55cf42c46d98>
##
## Parametric coefficients:
##                Estimate Std. Error t value Pr(>|t|)
## (Intercept)   1482526.6     8414.7 176.184  < 2e-16 ***
## sexmale        177247.2     1438.7 123.198  < 2e-16 ***
## siteousPrisma   28217.7     8342.0   3.383 0.000719 ***
## siteousSkyra   -61524.0     3176.0 -19.371  < 2e-16 ***
## siteUCAM       -39701.0     8472.5  -4.686 2.80e-06 ***
## siteUKB        -37461.8     7358.3  -5.091 3.58e-07 ***
## siteUmU       -113161.1    10363.1 -10.920  < 2e-16 ***
## bmi               874.3      170.9   5.116 3.15e-07 ***
## bmi:sleep_z      -223.3      162.6  -1.373 0.169745
## ---
```

```
## Signif. codes:  0 '***' 0.001 '**' 0.01 '*' 0.05 '.' 0.1 ' ' 1
##
## Approximate significance of smooth terms:
##               edf Ref.df    F p-value
## s(age_z)    6.182  6.182 73.0  <2e-16 ***
## s(sleep_z) 3.549  3.549 17.5  <2e-16 ***
## ---
## Signif. codes:  0 '***' 0.001 '**' 0.01 '*' 0.05 '.' 0.1 ' ' 1
##
## R-sq.(adj) =  0.338
## lmer.REML = 8.6858e+05  Scale est. = 1.4745e+08  n = 33453
```

We did the same controling for depression. Below is the model with no covariates but only keeping data with depression.

```
##
## Family: gaussian
## Link function: identity
##
## Formula:
## value ~ sex + site + s(age_z, k = 10, bs = "cr") + s(sleep_z,
##     k = 5, bs = "cr")
## <environment: 0x55cffb41e390>
##
## Parametric coefficients:
##                Estimate Std. Error t value Pr(>|t|)
## (Intercept)     1226926       8272  148.33   <2e-16 ***
## sexmale          178444       1436  124.30   <2e-16 ***
## siteousAvanto    272632      12544   21.73   <2e-16 ***
## siteousPrisma    285879      13410   21.32   <2e-16 ***
## siteousSkyra     210583      11514   18.29   <2e-16 ***
## siteUCAM         247976      11372   21.81   <2e-16 ***
## siteUKB          240213       8262   29.08   <2e-16 ***
## siteUmU          167224      11266   14.84   <2e-16 ***
## ---
## Signif. codes:  0 '***' 0.001 '**' 0.01 '*' 0.05 '.' 0.1 ' ' 1
##
## Approximate significance of smooth terms:
##               edf Ref.df     F p-value
## s(age_z)    6.397  6.397 71.05  <2e-16 ***
## s(sleep_z) 3.438  3.438 21.74  <2e-16 ***
## ---
## Signif. codes:  0 '***' 0.001 '**' 0.01 '*' 0.05 '.' 0.1 ' ' 1
##
## R-sq.(adj) =  0.357
## lmer.REML = 8.658e+05  Scale est. = 1.4235e+08  n = 33365
```

Below is the model output with main effect.

```
##
## Family: gaussian
## Link function: identity
##
## Formula:
## value ~ sex + site + s(age_z, k = 10, bs = "cr") + s(sleep_z,
##     k = 5, bs = "cr") + depression
```

```
## <environment: 0x55cffb41e390>
##
## Parametric coefficients:
##               Estimate Std. Error t value Pr(>|t|)
## (Intercept)    1225404       8303 147.579   <2e-16 ***
## sexmale         178594       1437 124.250   <2e-16 ***
## siteousAvanto   271638      12549  21.645   <2e-16 ***
## siteousPrisma   284868      13416  21.234   <2e-16 ***
## siteousSkyra    209525      11521  18.187   <2e-16 ***
## siteUCAM        247479      11373  21.761   <2e-16 ***
## siteUKB         240796       8266  29.132   <2e-16 ***
## siteUmU         162259      11511  14.096   <2e-16 ***
## depression       13159       6289   2.092   0.0364 *
## ---
## Signif. codes:  0 ‘***’ 0.001 ‘**’ 0.01 ‘*’ 0.05 ‘.’ 0.1 ‘ ’ 1
##
## Approximate significance of smooth terms:
##              edf Ref.df     F p-value
## s(age_z)   6.369  6.369 68.46  <2e-16 ***
## s(sleep_z) 3.462  3.462 22.55  <2e-16 ***
## ---
## Signif. codes:  0 ‘***’ 0.001 ‘**’ 0.01 ‘*’ 0.05 ‘.’ 0.1 ‘ ’ 1
##
## R-sq.(adj) =  0.357
## lmer.REML = 8.6578e+05  Scale est. = 1.4238e+08  n = 33365
```

Next is the model with depression-sleep interaction.

```
##
## Family: gaussian
## Link function: identity
##
## Formula:
## value ~ sex + site + s(age_z, k = 10, bs = "cr") + s(sleep_z,
##     k = 5, bs = "cr") + depression + depression:sleep_z
## <environment: 0x55cffb41e390>
##
## Parametric coefficients:
##                    Estimate Std. Error t value Pr(>|t|)
## (Intercept)         1225403       8304 147.577   <2e-16 ***
## sexmale              178609       1438 124.232   <2e-16 ***
## siteousAvanto        271716      12550  21.650   <2e-16 ***
## siteousPrisma        284949      13417  21.238   <2e-16 ***
## siteousSkyra         209600      11522  18.191   <2e-16 ***
## siteUCAM             247585      11375  21.766   <2e-16 ***
## siteUKB              240790       8266  29.131   <2e-16 ***
## siteUmU              161488      11616  13.902   <2e-16 ***
## depression            13368       6303   2.121   0.0339 *
## depression:sleep_z     2324       4695   0.495   0.6206
## ---
## Signif. codes:  0 ‘***’ 0.001 ‘**’ 0.01 ‘*’ 0.05 ‘.’ 0.1 ‘ ’ 1
##
## Approximate significance of smooth terms:
##              edf Ref.df     F p-value
## s(age_z)   6.368  6.368 68.46  <2e-16 ***
```

```
## s(sleep_z) 3.447  3.447 19.25  <2e-16 ***
## ---
## Signif. codes:  0 '***' 0.001 '**' 0.01 '*' 0.05 '.' 0.1 ' ' 1
##
## R-sq.(adj) =  0.357
## lmer.REML = 8.6576e+05  Scale est. = 1.4238e+08   n = 33365
```

The plot below shows the sleep-volume curve for the original model and for the model with main effects of SES.

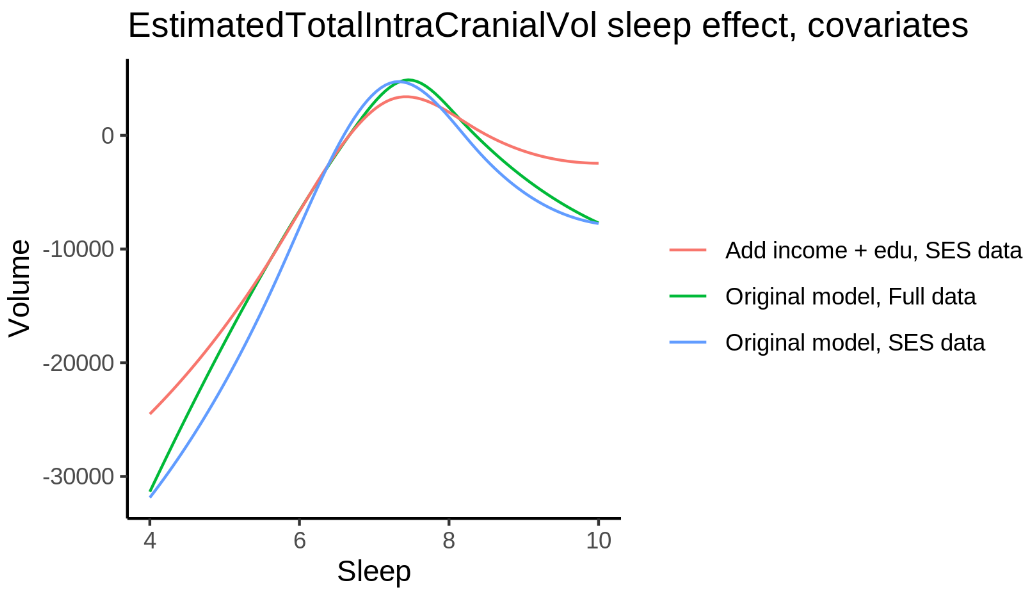


The plot below shows the sleep-volume curve for the original model and for the model with main effects of BMI.

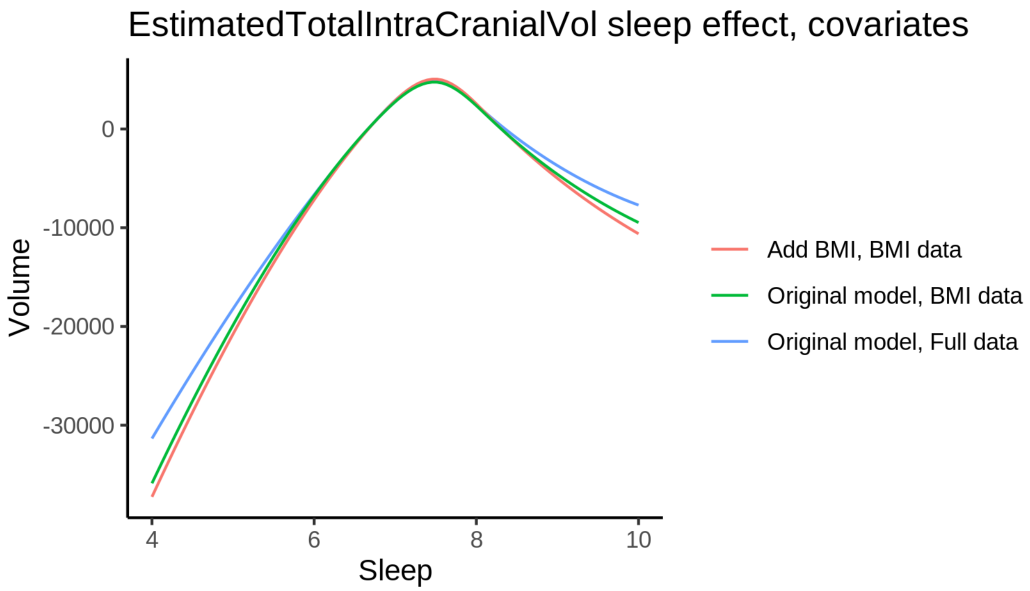


The plot below shows the sleep-volume curve for the original model and for the model with main effects of depression.

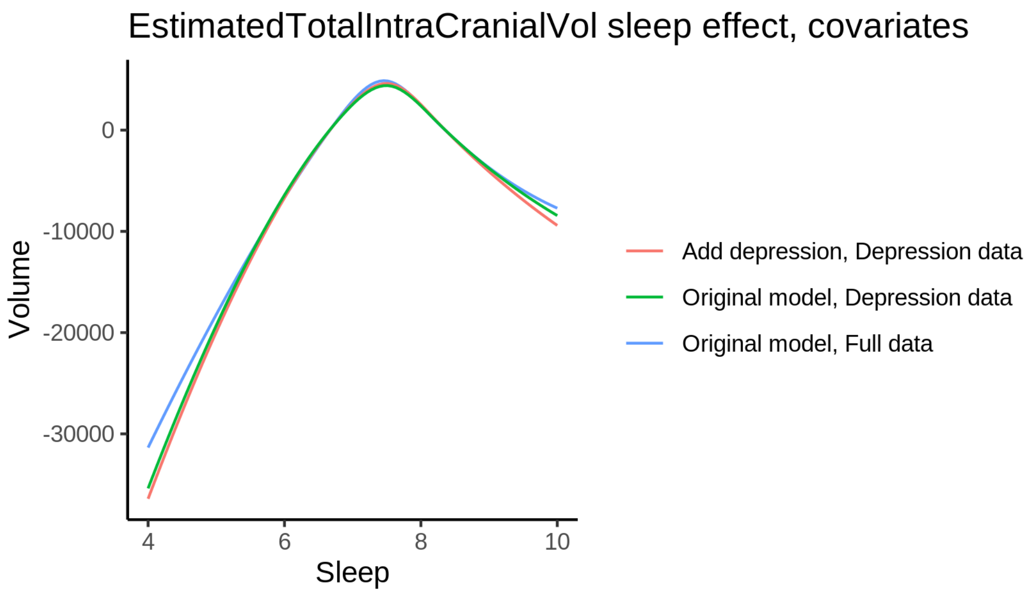


### Hippocampus

#### Descriptive statistics

| Study | Observations | Unique IDs | Mean age | Age range |
|---|---|---|---|---|
| HCP | 974 | 974 | 28.8 | 22 - 37 |
| MPIB | 677 | 391 | 63.1 | 24 - 83 |
| UB | 113 | 39 | 70.9 | 64 - 81 |
| UCAM | 884 | 632 | 55.1 | 20 - 88 |
| UiO | 1475 | 803 | 49.4 | 20 - 89 |
| UKB | 45975 | 43133 | 64.5 | 45 - 83 |
| UmU | 423 | 284 | 62.3 | 25 - 85 |
| UOXF | 769 | 769 | 69.8 | 60 - 85 |

##### Spaghetti plot

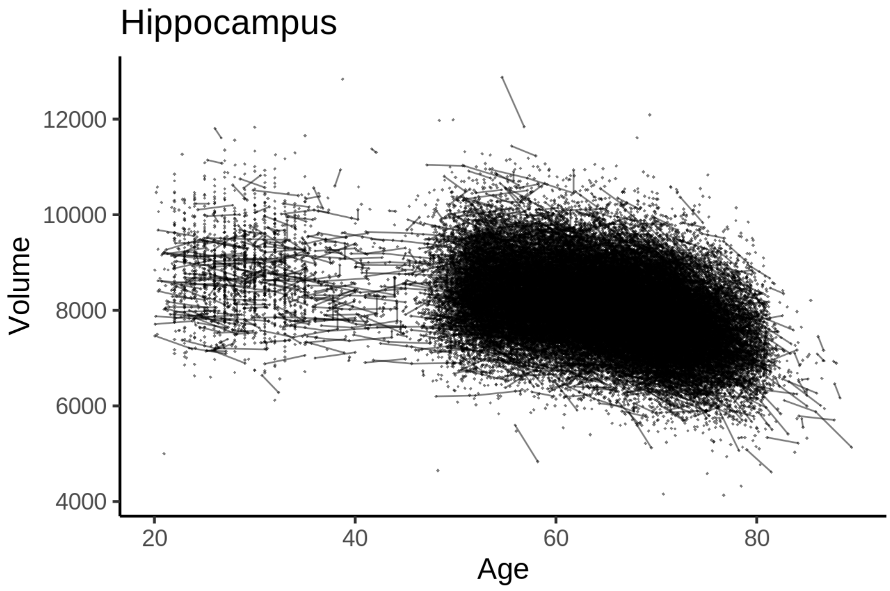


##### Model outputs

Model without sleep term

```
##
## Family: gaussian
## Link function: identity
##
## Formula:
## value ~ sex + site + icv + s(age_z, k = 10, bs = "cr")
## <environment: 0x55cf408eb8e0>
##
## Parametric coefficients:
##               Estimate Std. Error t value Pr(>|t|)
## (Intercept)   8112.510     34.441 235.551  < 2e-16 ***
## sexmale        123.282      7.125  17.304  < 2e-16 ***
## siteMPIB       274.524     44.380   6.186 6.23e-10 ***
## siteousAvanto -490.342     35.051 -13.990  < 2e-16 ***
## siteousPrisma  -92.627     59.135  -1.566  0.11727
## siteousSkyra  -193.006     33.731  -5.722 1.06e-08 ***
## siteUB        -139.267    105.056  -1.326  0.18496
## siteUCAM      -104.010     38.397  -2.709  0.00676 **
## siteUKB       -102.714     35.305  -2.909  0.00362 **
## siteUmU       -119.373     49.462  -2.413  0.01581 *
## siteUOXF      -303.527     41.974  -7.231 4.85e-13 ***
## icv            395.665      3.577 110.608  < 2e-16 ***
## ---
## Signif. codes:  0 ‘***’ 0.001 ‘**’ 0.01 ‘*’ 0.05 ‘.’ 0.1 ‘ ’ 1
##
## Approximate significance of smooth terms:
##            edf Ref.df    F p-value
```

```
## s(age_z) 7.952  7.952 1518  <2e-16 ***
## ---
## Signif. codes:  0 ‘***’ 0.001 ‘**’ 0.01 ‘*’ 0.05 ‘.’ 0.1 ‘ ’ 1
##
## R-sq.(adj) =  0.423
## lmer.REML = 7.9714e+05  Scale est. = 22392     n = 51290
```

Model with only main effects of age and sleep

```
##
## Family: gaussian
## Link function: identity
##
## Formula:
## value ~ sex + site + icv + s(age_z, k = 10, bs = "cr") + s(sleep_z,
##     k = 5, bs = "cr")
## <environment: 0x55cf408eb8e0>
##
## Parametric coefficients:
##               Estimate Std. Error t value Pr(>|t|)
## (Intercept)   8108.801     34.466 235.272  < 2e-16 ***
## sexmale        122.872      7.122  17.252  < 2e-16 ***
## siteMPIB       281.505     44.391   6.341 2.29e-10 ***
## siteousAvanto -489.345     35.051 -13.961  < 2e-16 ***
## siteousPrisma  -91.985     59.119  -1.556  0.11973
## siteousSkyra  -192.363     33.733  -5.703 1.19e-08 ***
## siteUB        -135.001    105.009  -1.286  0.19858
## siteUCAM      -100.500     38.390  -2.618  0.00885 **
## siteUKB        -98.657     35.333  -2.792  0.00524 **
## siteUmU       -104.394     49.558  -2.107  0.03516 *
## siteUOXF      -301.875     41.965  -7.193 6.40e-13 ***
## icv            395.516      3.579 110.495  < 2e-16 ***
## ---
## Signif. codes:  0 ‘***’ 0.001 ‘**’ 0.01 ‘*’ 0.05 ‘.’ 0.1 ‘ ’ 1
##
## Approximate significance of smooth terms:
##              edf Ref.df       F p-value
## s(age_z)   7.953  7.953 1495.45  <2e-16 ***
## s(sleep_z) 2.960  2.960   13.72  <2e-16 ***
## ---
## Signif. codes:  0 ‘***’ 0.001 ‘**’ 0.01 ‘*’ 0.05 ‘.’ 0.1 ‘ ’ 1
##
## R-sq.(adj) =  0.423
## lmer.REML = 7.971e+05  Scale est. = 22404     n = 51290
```

Model with full interaction between age and sleep

```
##
## Family: gaussian
## Link function: identity
##
## Formula:
## value ~ sex + site + icv + t2(age_z, sleep_z, k = c(10, 4), bs = "cr")
## <environment: 0x55cf408eb8e0>
##
## Parametric coefficients:
```

```
##                Estimate Std. Error t value Pr(>|t|)
## (Intercept)   8108.596     34.545 234.724  < 2e-16 ***
## sexmale        122.606      7.137  17.178  < 2e-16 ***
## siteMPIB       280.965     44.596   6.300 3.00e-10 ***
## siteousAvanto -488.582     35.231 -13.868  < 2e-16 ***
## siteousPrisma  -93.586     59.251  -1.579  0.11423
## siteousSkyra  -191.333     33.928  -5.639 1.72e-08 ***
## siteUB        -135.920    105.058  -1.294  0.19575
## siteUCAM      -101.849     38.566  -2.641  0.00827 **
## siteUKB        -98.296     35.409  -2.776  0.00551 **
## siteUmU       -107.478     49.679  -2.163  0.03051 *
## siteUOXF      -300.924     42.077  -7.152 8.69e-13 ***
## icv            395.567      3.578 110.542  < 2e-16 ***
## ---
## Signif. codes:  0 '***' 0.001 '**' 0.01 '*' 0.05 '.' 0.1 ' ' 1
##
## Approximate significance of smooth terms:
##                     edf Ref.df     F p-value
## t2(age_z,sleep_z) 14.22  14.22 37.66  <2e-16 ***
## ---
## Signif. codes:  0 '***' 0.001 '**' 0.01 '*' 0.05 '.' 0.1 ' ' 1
##
## R-sq.(adj) =  0.423
## lmer.REML = 7.971e+05  Scale est. = 22392     n = 51290
```

#### Model comparison

mod_no_sleep refers to model without sleep term, mod_no_interaction refers to model with only main effect of sleep, and mod_full refers to model with a full interaction between age and sleep. This is a nested model comparison, and the p-value at a given line refers to comparing the model at the line to the model on the line above. Hence, significance implies that the more complicated model is supported on statistical grounds.

To be even more specific, the p-value on the second row tests whether there is an association between sleep and volume. The p-value on the third row tests whether this association depends on age.

```
## Data: NULL
## Models:
## mod_list$mod_no_sleep$mer: NULL
## mod_list$mod_no_interaction$mer: NULL
## mod_list$mod_full$mer: NULL
##                                npar    AIC    BIC  logLik deviance   Chisq Df Pr(>Chisq)
## mod_list$mod_no_sleep$mer         16 797170 797311 -398569   797138
## mod_list$mod_no_interaction$mer   18 797139 797298 -398551   797103 34.6416  2  3.004e-08 ***
## mod_list$mod_full$mer             20 797142 797319 -398551   797102  1.0135  2     0.6025
## ---
## Signif. codes:  0 '***' 0.001 '**' 0.01 '*' 0.05 '.' 0.1 ' ' 1
```

We chose the model based on the likelihood ratio test with 5 % significance level, which was mod_no_interaction.

#### Lifespan brain trajectory

The trajectory shown is from the chosen model mod_no_interaction.

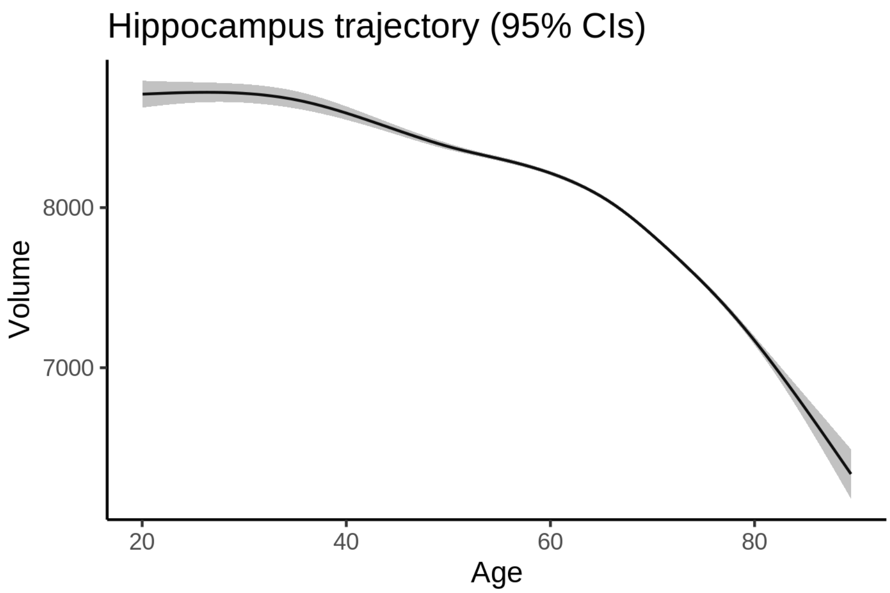


##### Effect of sleep

The chosen model only included the main effect of sleep, and hence the effect does not vary with age. The black dot shows the average sleep duration across all ages in the sample.

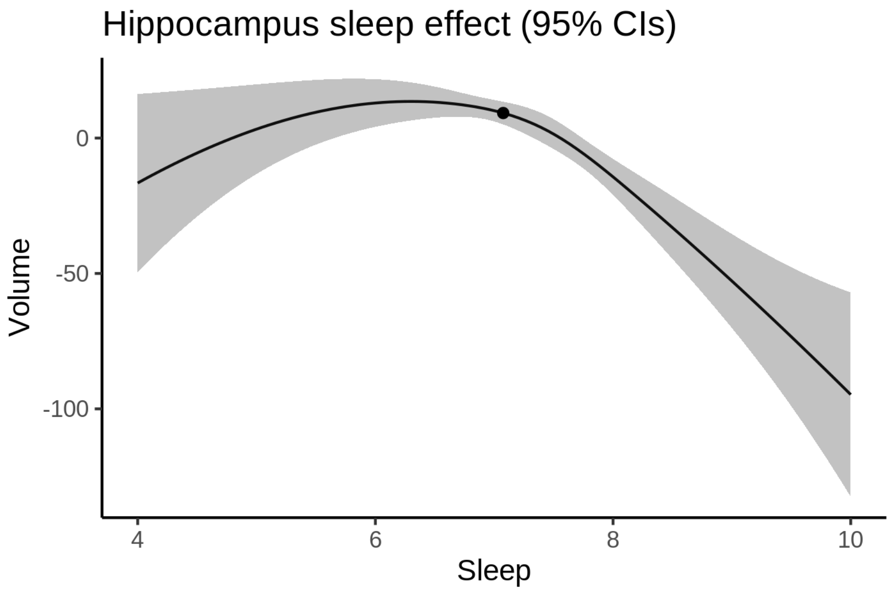


We also show the full interaction model for completeness, although it was not selected.

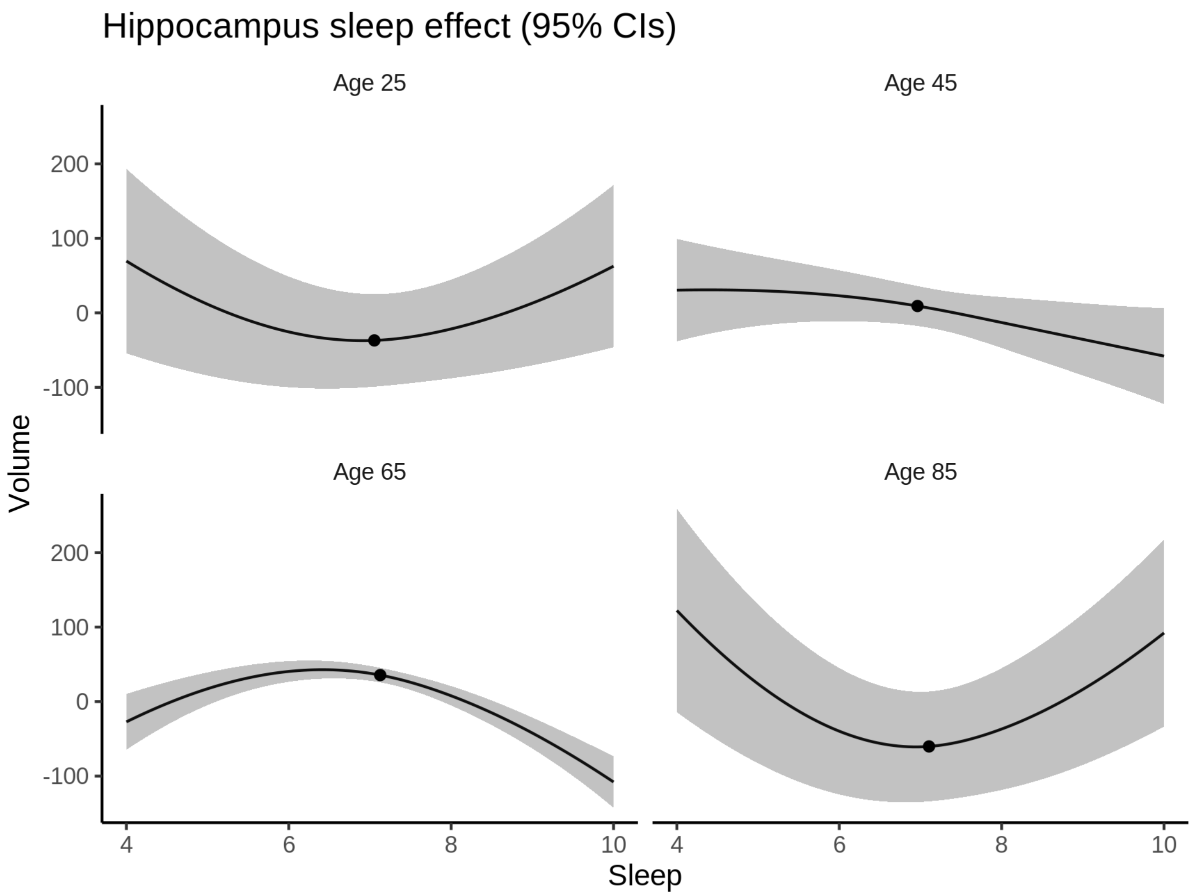


**Deviation from sleep associated with maximal volume**

Model with only main effect of sleep was chosen, so we show it for all ages at once. Maximum volume is attained at 6.3 hours of sleep. The percentage values in the plot are calculated as follows: The maximum at 100 % refers to a person at an arbitrary age with a sleep duration associated with maximum volume. For a female, this volume is 8060 and for a male it is 8183. The other percentage values show how large the expected volume is for someone with other sleep durations. For example, 99 % implies a 1 % reduction.

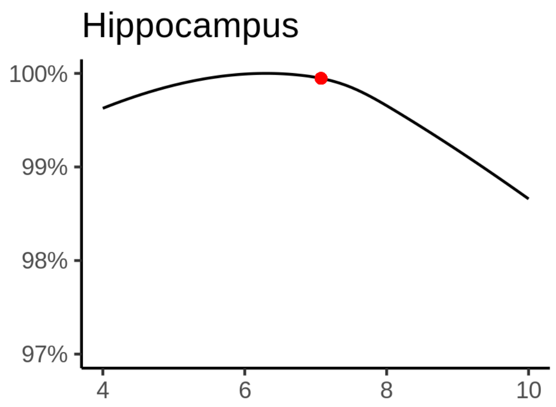

**Comparison of mean sleep and sleep associated with maximum volume**

A 95 % confidence interval for the sleep associated with maximum volume is [5.21, 7.15].

The plot below compares average sleep to the sleep associated with maximum volume.

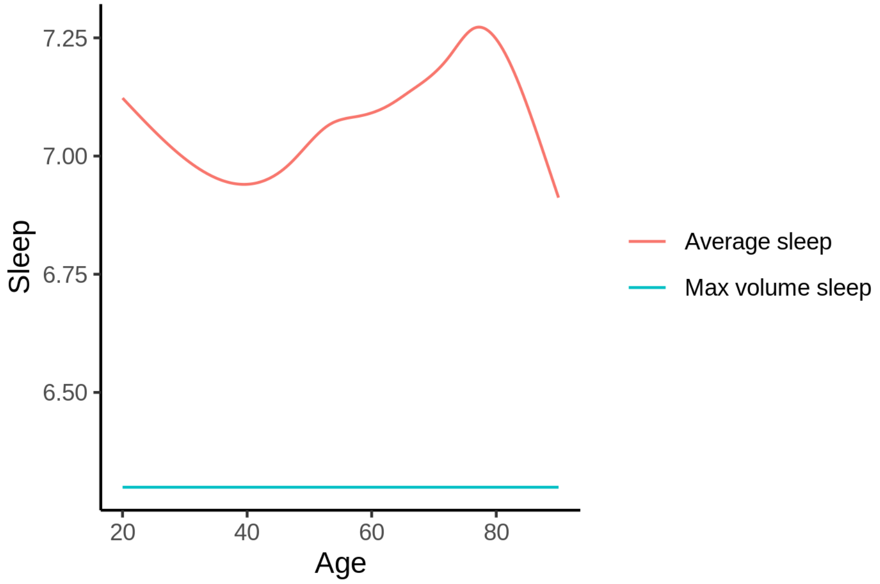


The next plot shows the difference between average sleep and sleep associated with maximum volume. The shaded region is a 95 % confidence interval.

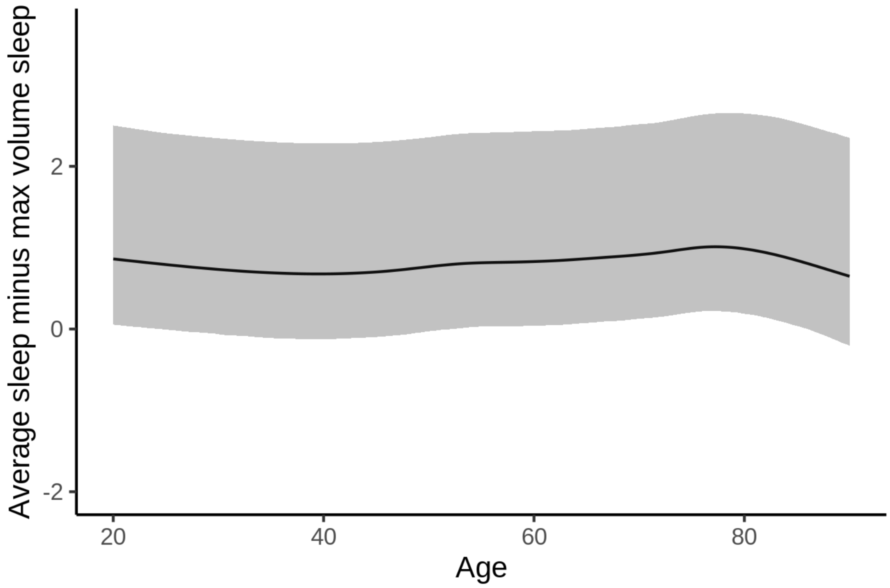


The next plot shows the probability that the sleep duration associated with maximum volume is longer than the average sleep duration, as a function of age. Probility below .05 can be interpreted as evidence that the sleep associated with maximum volume is shorter than the mean sleep, and probability above .95 can be interpreted the opposite way.

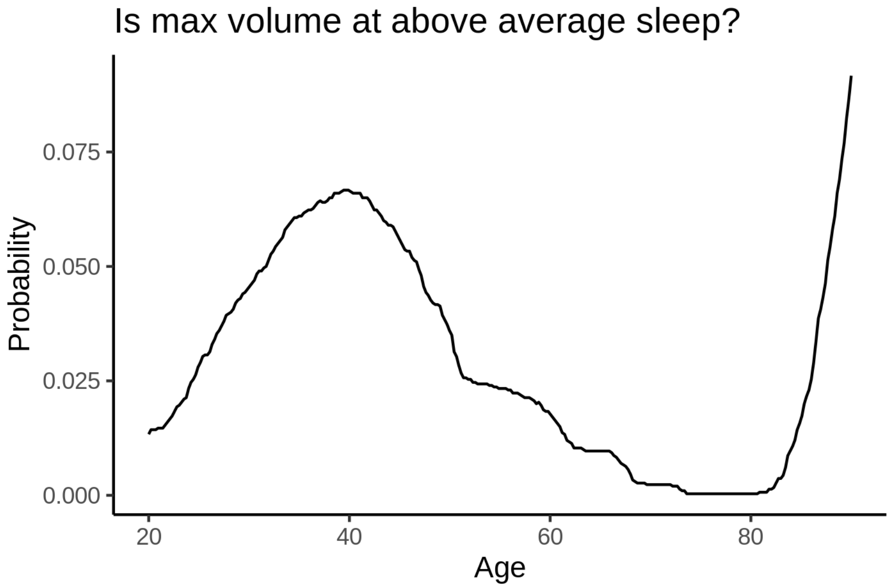


##### Controling for covariates

Below is the output for a model in which we only include data with income and education.

```
##
## Family: gaussian
## Link function: identity
##
## Formula:
## value ~ sex + site + s(age_z, k = 10, bs = "cr") + s(sleep_z,
##     k = 5, bs = "cr") + icv
## <environment: 0x55d075f74ee8>
##
## Parametric coefficients:
##                Estimate Std. Error t value Pr(>|t|)
## (Intercept)    8469.311     82.446 102.725  < 2e-16 ***
## sexmale         102.521      9.018  11.368  < 2e-16 ***
## siteousAvanto -898.627     96.829  -9.281  < 2e-16 ***
## siteousPrisma -344.268    100.492  -3.426 0.000614 ***
## siteousSkyra  -556.398     88.938  -6.256 4.00e-10 ***
## siteUKB       -457.516     82.303  -5.559 2.74e-08 ***
## siteUOXF      -543.695     92.544  -5.875 4.27e-09 ***
## icv            412.033      4.628  89.035  < 2e-16 ***
## ---
## Signif. codes:  0 ‘***’ 0.001 ‘**’ 0.01 ‘*’ 0.05 ‘.’ 0.1 ‘ ’ 1
##
## Approximate significance of smooth terms:
##              edf Ref.df       F  p-value
## s(age_z)   6.898  6.898 906.250  < 2e-16 ***
## s(sleep_z) 2.701  2.701   7.544 0.000231 ***
## ---
## Signif. codes:  0 ‘***’ 0.001 ‘**’ 0.01 ‘*’ 0.05 ‘.’ 0.1 ‘ ’ 1
##
```

```
## R-sq.(adj) =  0.401
## lmer.REML = 4.846e+05  Scale est. = 23769     n = 31190
```

Below is the output for a model in which we control for the main effects of income and education.

```
##
## Family: gaussian
## Link function: identity
##
## Formula:
## value ~ sex + site + s(age_z, k = 10, bs = "cr") + s(sleep_z,
##     k = 5, bs = "cr") + icv + income_scaled + education_scaled
## <environment: 0x55d075f74ee8>
##
## Parametric coefficients:
##                  Estimate Std. Error t value Pr(>|t|)
## (Intercept)      8438.563     82.959 101.720  < 2e-16 ***
## sexmale           102.719      9.043  11.359  < 2e-16 ***
## siteousAvanto    -898.983     96.844  -9.283  < 2e-16 ***
## siteousPrisma    -347.177    100.481  -3.455 0.000551 ***
## siteousSkyra     -557.210     88.947  -6.265 3.79e-10 ***
## siteUKB          -457.907     82.338  -5.561 2.70e-08 ***
## siteUOXF         -534.197     92.579  -5.770 7.99e-09 ***
## icv               410.078      4.666  87.893  < 2e-16 ***
## income_scaled      18.482     10.750   1.719 0.085569 .
## education_scaled   28.624     12.707   2.253 0.024284 *
## ---
## Signif. codes:  0 '***' 0.001 '**' 0.01 '*' 0.05 '.' 0.1 ' ' 1
##
## Approximate significance of smooth terms:
##              edf Ref.df       F  p-value
## s(age_z)   6.897  6.897 825.692  < 2e-16 ***
## s(sleep_z) 2.595  2.595   7.106 0.000452 ***
## ---
## Signif. codes:  0 '***' 0.001 '**' 0.01 '*' 0.05 '.' 0.1 ' ' 1
##
## R-sq.(adj) =  0.402
## lmer.REML = 4.8458e+05  Scale est. = 23791     n = 31190
```

We also included interaction effects between sleep duration and education and income, in another model. The output is shown below, and the interaction terms are `income_scaled:sleep_z` and `education_scaled:sleep_z`.

```
##
## Family: gaussian
## Link function: identity
##
## Formula:
## value ~ sex + site + s(age_z, k = 10, bs = "cr") + s(sleep_z,
##     k = 5, bs = "cr") + icv + income_scaled + education_scaled +
##     income_scaled:sleep_z + education_scaled:sleep_z
## <environment: 0x55d075f74ee8>
##
## Parametric coefficients:
##                          Estimate Std. Error t value Pr(>|t|)
## (Intercept)              8438.866     82.963 101.718  < 2e-16 ***
```

```
## sexmale                    102.082      9.049  11.281  < 2e-16 ***
## siteousAvanto             -900.126     96.845  -9.295  < 2e-16 ***
## siteousPrisma             -349.758    100.489  -3.481 0.000501 ***
## siteousSkyra              -558.326     88.947  -6.277 3.50e-10 ***
## siteUKB                   -458.180     82.345  -5.564 2.66e-08 ***
## siteUOXF                  -534.034     92.603  -5.767 8.15e-09 ***
## icv                        410.146      4.666  87.901  < 2e-16 ***
## income_scaled               19.087     10.755   1.775 0.075968 .
## education_scaled            28.215     12.708   2.220 0.026413 *
## income_scaled:sleep_z      -18.477     10.703  -1.726 0.084315 .
## education_scaled:sleep_z    14.270     12.331   1.157 0.247173
## ---
## Signif. codes:  0 ‘***’ 0.001 ‘**’ 0.01 ‘*’ 0.05 ‘.’ 0.1 ‘ ’ 1
##
## Approximate significance of smooth terms:
##              edf Ref.df       F p-value
## s(age_z)   6.897  6.897 824.909  <2e-16 ***
## s(sleep_z) 2.635  2.635   4.576  0.0111 *
## ---
## Signif. codes:  0 ‘***’ 0.001 ‘**’ 0.01 ‘*’ 0.05 ‘.’ 0.1 ‘ ’ 1
##
## R-sq.(adj) =  0.402
## lmer.REML = 4.8456e+05  Scale est. = 23791     n = 31190
```

We did the same controling for BMI. Below is the model with no covariates but only keeping data with BMI.

```
##
## Family: gaussian
## Link function: identity
##
## Formula:
## value ~ sex + site + s(age_z, k = 10, bs = "cr") + s(sleep_z,
##     k = 5, bs = "cr") + icv
## <environment: 0x55cffd511c48>
##
## Parametric coefficients:
##               Estimate Std. Error t value Pr(>|t|)
## (Intercept)   7606.759     52.116 145.959  < 2e-16 ***
## sexmale         96.281      8.758  10.994  < 2e-16 ***
## siteousPrisma  471.568     69.283   6.806 1.02e-11 ***
## siteousSkyra   387.221     38.042  10.179  < 2e-16 ***
## siteUCAM       430.543     55.307   7.785 7.20e-15 ***
## siteUKB        411.476     52.624   7.819 5.47e-15 ***
## siteUmU        435.593     63.458   6.864 6.80e-12 ***
## icv            417.266      4.507  92.578  < 2e-16 ***
## ---
## Signif. codes:  0 ‘***’ 0.001 ‘**’ 0.01 ‘*’ 0.05 ‘.’ 0.1 ‘ ’ 1
##
## Approximate significance of smooth terms:
##              edf Ref.df      F  p-value
## s(age_z)   7.289  7.289 987.33  < 2e-16 ***
## s(sleep_z) 2.596  2.596   7.91 0.000148 ***
## ---
## Signif. codes:  0 ‘***’ 0.001 ‘**’ 0.01 ‘*’ 0.05 ‘.’ 0.1 ‘ ’ 1
##
```

```
## R-sq.(adj) =  0.412
## lmer.REML = 5.1907e+05  Scale est. = 22530     n = 33445
```

Below is the model output with main effect.

```
##
## Family: gaussian
## Link function: identity
##
## Formula:
## value ~ sex + site + s(age_z, k = 10, bs = "cr") + s(sleep_z,
##     k = 5, bs = "cr") + icv + bmi
## <environment: 0x55cffd511c48>
##
## Parametric coefficients:
##                 Estimate Std. Error t value Pr(>|t|)
## (Intercept)    7589.5638    56.4029 134.560  < 2e-16 ***
## sexmale          95.7381     8.7840  10.899  < 2e-16 ***
## siteousPrisma   471.0061    69.2870   6.798 1.08e-11 ***
## siteousSkyra    387.1637    38.0431  10.177  < 2e-16 ***
## siteUCAM        430.1804    55.3100   7.778 7.60e-15 ***
## siteUKB         410.9495    52.6289   7.808 5.96e-15 ***
## siteUmU         434.8825    63.4645   6.852 7.39e-12 ***
## icv             417.1578     4.5092  92.514  < 2e-16 ***
## bmi               0.6817     0.8548   0.797    0.425
## ---
## Signif. codes:  0 ‘***’ 0.001 ‘**’ 0.01 ‘*’ 0.05 ‘.’ 0.1 ‘ ’ 1
##
## Approximate significance of smooth terms:
##              edf Ref.df      F  p-value
## s(age_z)   7.288  7.288 987.09  < 2e-16 ***
## s(sleep_z) 2.614  2.614   7.61 0.000144 ***
## ---
## Signif. codes:  0 ‘***’ 0.001 ‘**’ 0.01 ‘*’ 0.05 ‘.’ 0.1 ‘ ’ 1
##
## R-sq.(adj) =  0.412
## lmer.REML = 5.1907e+05  Scale est. = 22531     n = 33445
```

Next is the model with BMI-sleep interaction.

```
##
## Family: gaussian
## Link function: identity
##
## Formula:
## value ~ sex + site + s(age_z, k = 10, bs = "cr") + s(sleep_z,
##     k = 5, bs = "cr") + icv + bmi + bmi:sleep_z
## <environment: 0x55cffd511c48>
##
## Parametric coefficients:
##                 Estimate Std. Error t value Pr(>|t|)
## (Intercept)    7587.9785    56.4324 134.461  < 2e-16 ***
## sexmale          95.7691     8.7841  10.903  < 2e-16 ***
## siteousPrisma   471.0122    69.2870   6.798 1.08e-11 ***
## siteousSkyra    387.2929    38.0431  10.180  < 2e-16 ***
## siteUCAM        430.7383    55.3136   7.787 7.05e-15 ***
```

```
## siteUKB        411.5115    52.6327   7.819 5.50e-15 ***
## siteUmU        435.3608    63.4669   6.860 7.02e-12 ***
## icv            417.1870     4.5093  92.517  < 2e-16 ***
## bmi              0.7073     0.8553   0.827    0.408
## bmi:sleep_z      0.7032     0.8108   0.867    0.386
## ---
## Signif. codes:  0 '***' 0.001 '**' 0.01 '*' 0.05 '.' 0.1 ' ' 1
##
## Approximate significance of smooth terms:
##              edf Ref.df       F p-value
## s(age_z)   7.287  7.287 987.291 < 2e-16 ***
## s(sleep_z) 2.610  2.610   4.226 0.00772 **
## ---
## Signif. codes:  0 '***' 0.001 '**' 0.01 '*' 0.05 '.' 0.1 ' ' 1
##
## R-sq.(adj) =  0.412
## lmer.REML = 5.1907e+05  Scale est. = 22530     n = 33445
```

We did the same controling for depression. Below is the model with no covariates but only keeping data with depression.

```
##
## Family: gaussian
## Link function: identity
##
## Formula:
## value ~ sex + site + s(age_z, k = 10, bs = "cr") + s(sleep_z,
##     k = 5, bs = "cr") + icv
## <environment: 0x55cffb399a28>
##
## Parametric coefficients:
##               Estimate Std. Error t value Pr(>|t|)
## (Intercept)   8352.007     41.973 198.984  < 2e-16 ***
## sexmale         95.391      8.806  10.833  < 2e-16 ***
## siteousAvanto -710.959     83.921  -8.472  < 2e-16 ***
## siteousPrisma -194.873     79.554  -2.450   0.0143 *
## siteousSkyra  -296.464     62.943  -4.710 2.49e-06 ***
## siteUCAM      -351.896     57.537  -6.116 9.70e-10 ***
## siteUKB       -338.155     41.512  -8.146 3.89e-16 ***
## siteUmU       -305.071     56.245  -5.424 5.87e-08 ***
## icv            414.420      4.525  91.578  < 2e-16 ***
## ---
## Signif. codes:  0 '***' 0.001 '**' 0.01 '*' 0.05 '.' 0.1 ' ' 1
##
## Approximate significance of smooth terms:
##              edf Ref.df       F  p-value
## s(age_z)   7.219  7.219 954.439  < 2e-16 ***
## s(sleep_z) 2.670  2.670   9.018 4.83e-05 ***
## ---
## Signif. codes:  0 '***' 0.001 '**' 0.01 '*' 0.05 '.' 0.1 ' ' 1
##
## R-sq.(adj) =  0.406
## lmer.REML = 5.1749e+05  Scale est. = 22060     n = 33357
```

Below is the model output with main effect.

```
##
## Family: gaussian
## Link function: identity
##
## Formula:
## value ~ sex + site + s(age_z, k = 10, bs = "cr") + s(sleep_z,
##     k = 5, bs = "cr") + icv + depression
## <environment: 0x55cffb399a28>
##
## Parametric coefficients:
##                Estimate Std. Error t value Pr(>|t|)
## (Intercept)    8368.205     42.142 198.572  < 2e-16 ***
## sexmale          93.712      8.813  10.634  < 2e-16 ***
## siteousAvanto -702.588     83.923  -8.372  < 2e-16 ***
## siteousPrisma -187.324     79.553  -2.355   0.0185 *
## siteousSkyra  -287.038     62.965  -4.559 5.17e-06 ***
## siteUCAM      -348.098     57.529  -6.051 1.46e-09 ***
## siteUKB       -344.803     41.533  -8.302  < 2e-16 ***
## siteUmU       -255.878     57.431  -4.455 8.40e-06 ***
## icv            414.674      4.524  91.655  < 2e-16 ***
## depression    -132.626     31.489  -4.212 2.54e-05 ***
## ---
## Signif. codes:  0 ‘***’ 0.001 ‘**’ 0.01 ‘*’ 0.05 ‘.’ 0.1 ‘ ’ 1
##
## Approximate significance of smooth terms:
##              edf Ref.df       F  p-value
## s(age_z)   7.210  7.210 953.699  < 2e-16 ***
## s(sleep_z) 2.447  2.447   9.146 5.27e-05 ***
## ---
## Signif. codes:  0 ‘***’ 0.001 ‘**’ 0.01 ‘*’ 0.05 ‘.’ 0.1 ‘ ’ 1
##
## R-sq.(adj) =  0.406
## lmer.REML = 5.1747e+05  Scale est. = 22051     n = 33357
```

Next is the model with depression-sleep interaction.

```
##
## Family: gaussian
## Link function: identity
##
## Formula:
## value ~ sex + site + s(age_z, k = 10, bs = "cr") + s(sleep_z,
##     k = 5, bs = "cr") + icv + depression + depression:sleep_z
## <environment: 0x55cffb399a28>
##
## Parametric coefficients:
##                     Estimate Std. Error t value Pr(>|t|)
## (Intercept)         8368.208     42.142 198.572  < 2e-16 ***
## sexmale               93.554      8.814  10.614  < 2e-16 ***
## siteousAvanto       -703.682     83.930  -8.384  < 2e-16 ***
## siteousPrisma       -188.428     79.560  -2.368   0.0179 *
## siteousSkyra        -288.090     62.973  -4.575 4.78e-06 ***
## siteUCAM            -349.229     57.540  -6.069 1.30e-09 ***
## siteUKB             -344.746     41.533  -8.301  < 2e-16 ***
## siteUmU             -248.433     57.923  -4.289 1.80e-05 ***
```

```
## icv                 414.688      4.524  91.657  < 2e-16 ***
## depression         -134.501     31.548  -4.263 2.02e-05 ***
## depression:sleep_z  -22.906     23.170  -0.989   0.3229
## ---
## Signif. codes:  0 '***' 0.001 '**' 0.01 '*' 0.05 '.' 0.1 ' ' 1
##
## Approximate significance of smooth terms:
##               edf Ref.df       F p-value
## s(age_z)    7.208  7.208 954.084 < 2e-16 ***
## s(sleep_z) 2.437  2.437   5.718 0.00154 **
## ---
## Signif. codes:  0 '***' 0.001 '**' 0.01 '*' 0.05 '.' 0.1 ' ' 1
##
## R-sq.(adj) =  0.406
## lmer.REML = 5.1746e+05  Scale est. = 22050     n = 33357
```

The plot below shows the sleep-volume curve for the original model and for the model with main effects of SES.

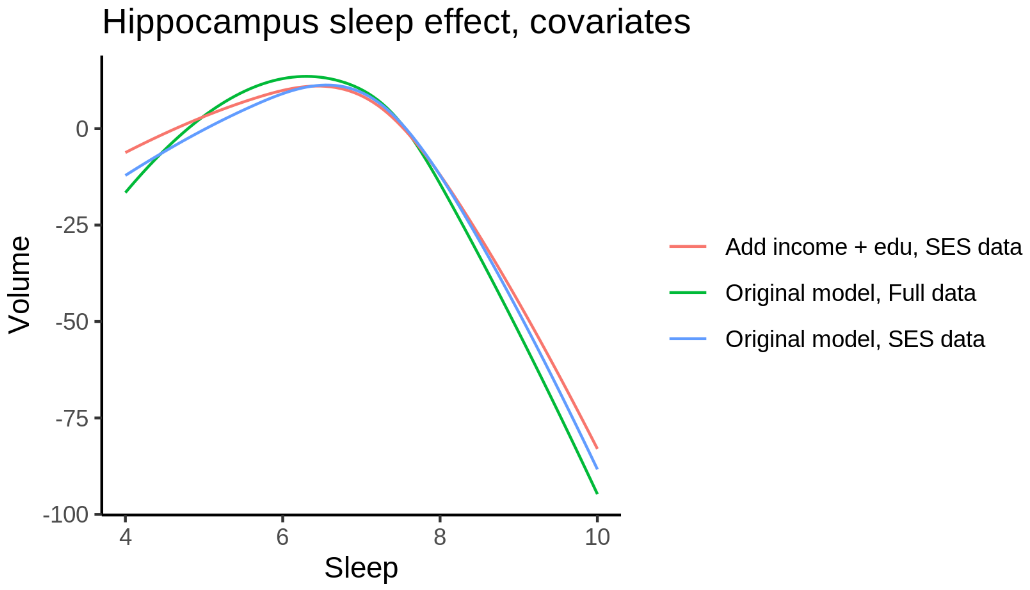


The plot below shows the sleep-volume curve for the original model and for the model with main effects of BMI.

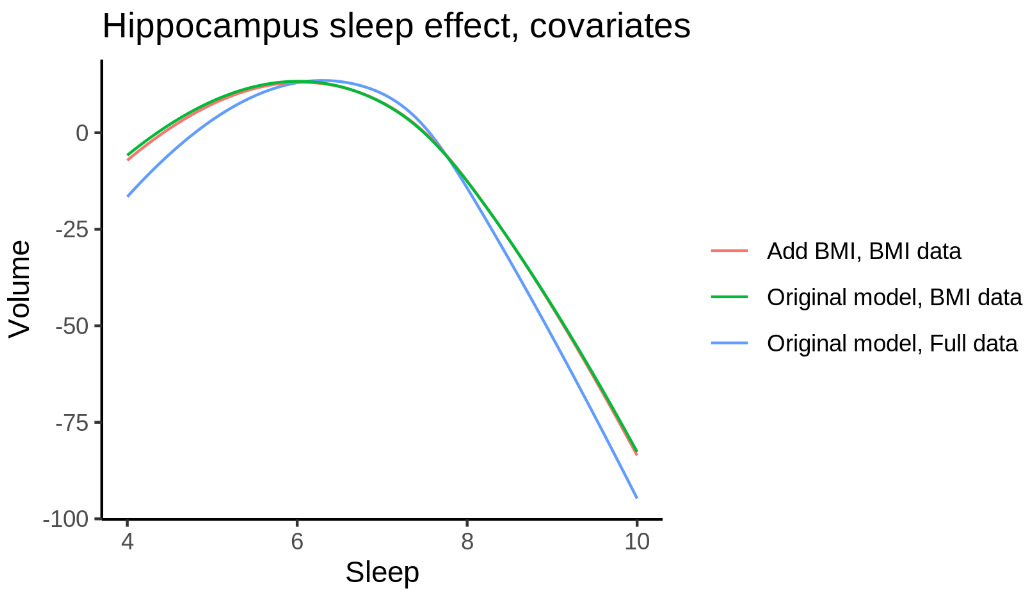


The plot below shows the sleep-volume curve for the original model and for the model with main effects of depression.

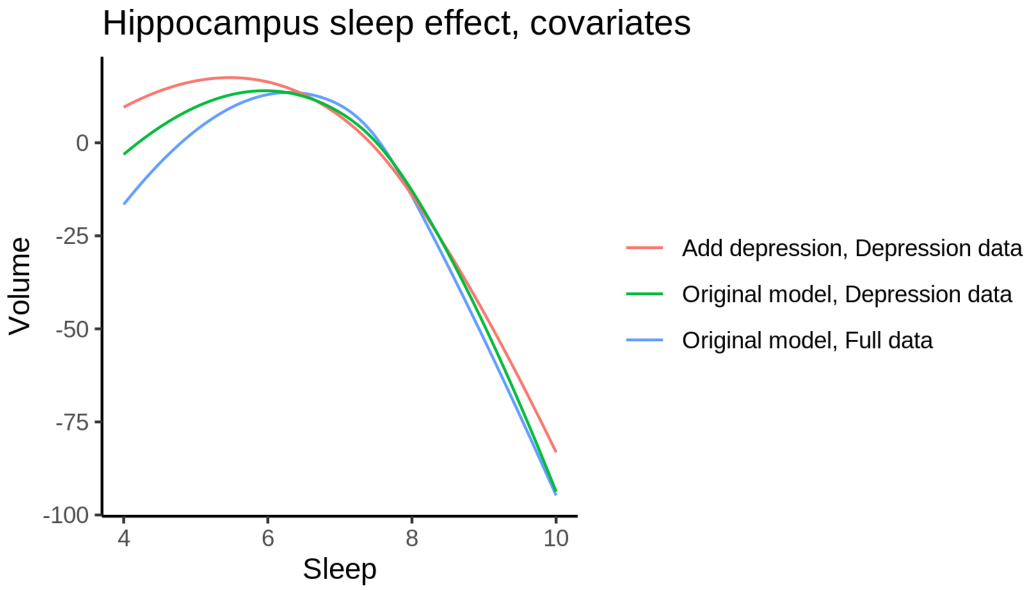


### Pallidum

#### Descriptive statistics

| Study | Observations | Unique IDs | Mean age | Age range |
|---|---|---|---|---|
| HCP | 974 | 974 | 28.8 | 22 - 37 |
| MPIB | 677 | 391 | 63.1 | 24 - 83 |
| UB | 113 | 39 | 70.9 | 64 - 81 |
| UCAM | 884 | 632 | 55.1 | 20 - 88 |

| Study | Observations | Unique IDs | Mean age | Age range |
|---|---|---|---|---|
| UiO | 1474 | 803 | 49.3 | 20 - 89 |
| UKB | 45983 | 43137 | 64.5 | 45 - 83 |
| UmU | 423 | 284 | 62.3 | 25 - 85 |
| UOXF | 769 | 769 | 69.8 | 60 - 85 |

##### Spaghetti plot

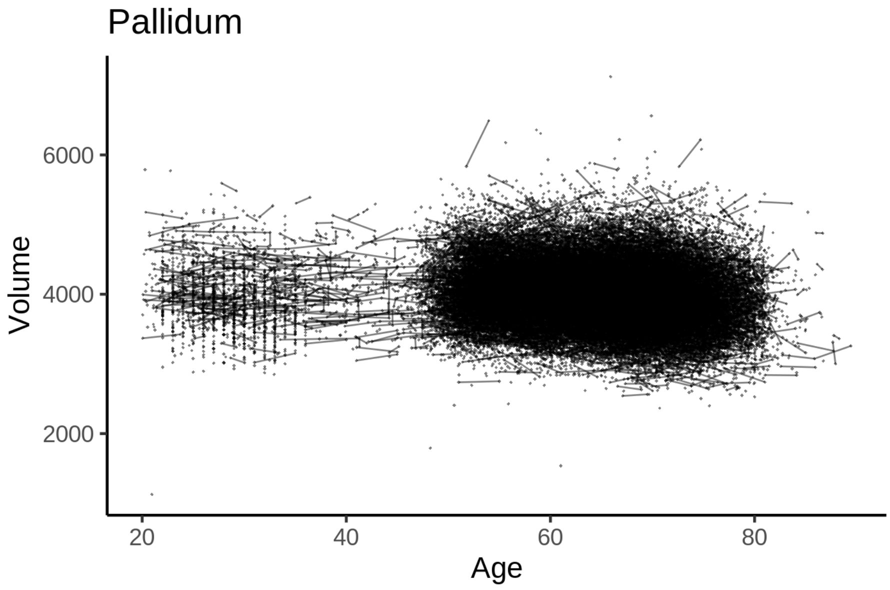


##### Model outputs

Model without sleep term

```
##
## Family: gaussian
## Link function: identity
##
## Formula:
## value ~ sex + site + icv + s(age_z, k = 10, bs = "cr")
## <environment: 0x55cf3c0125d8>
##
## Parametric coefficients:
##                Estimate Std. Error t value Pr(>|t|)
## (Intercept)    3623.601     19.029 190.425  < 2e-16 ***
## sexmale          84.926      3.870  21.947  < 2e-16 ***
## siteMPIB        516.109     24.191  21.335  < 2e-16 ***
## siteousAvanto   113.306     19.767   5.732 9.97e-09 ***
## siteousPrisma   229.801     33.418   6.877 6.20e-12 ***
## siteousSkyra    231.643     18.447  12.557  < 2e-16 ***
## siteUB           78.720     56.465   1.394    0.163
## siteUCAM        260.560     20.993  12.412  < 2e-16 ***
## siteUKB         346.464     19.518  17.751  < 2e-16 ***
## siteUmU         140.290     26.981   5.200 2.01e-07 ***
```

```
## siteUOXF       104.509     23.078   4.529 5.95e-06 ***
## icv            260.146      1.947 133.635  < 2e-16 ***
## ---
## Signif. codes:  0 '***' 0.001 '**' 0.01 '*' 0.05 '.' 0.1 ' ' 1
##
## Approximate significance of smooth terms:
##            edf Ref.df     F p-value
## s(age_z) 6.887  6.887 198.8  <2e-16 ***
## ---
## Signif. codes:  0 '***' 0.001 '**' 0.01 '*' 0.05 '.' 0.1 ' ' 1
##
## R-sq.(adj) =  0.423
## lmer.REML = 7.3686e+05  Scale est. = 11451     n = 51297
```

Model with only main effects of age and sleep

```
##
## Family: gaussian
## Link function: identity
##
## Formula:
## value ~ sex + site + icv + s(age_z, k = 10, bs = "cr") + s(sleep_z,
##     k = 5, bs = "cr")
## <environment: 0x55cf3c0125d8>
##
## Parametric coefficients:
##                Estimate Std. Error t value Pr(>|t|)
## (Intercept)    3624.456     19.047 190.285  < 2e-16 ***
## sexmale          84.956      3.870  21.954  < 2e-16 ***
## siteMPIB        515.177     24.208  21.281  < 2e-16 ***
## siteousAvanto   112.853     19.772   5.708 1.15e-08 ***
## siteousPrisma   229.497     33.419   6.867 6.61e-12 ***
## siteousSkyra    231.174     18.452  12.528  < 2e-16 ***
## siteUB           78.459     56.465   1.390    0.165
## siteUCAM        260.125     20.997  12.389  < 2e-16 ***
## siteUKB         345.556     19.538  17.686  < 2e-16 ***
## siteUmU         138.399     27.045   5.117 3.11e-07 ***
## siteUOXF        104.070     23.081   4.509 6.53e-06 ***
## icv             260.085      1.948 133.543  < 2e-16 ***
## ---
## Signif. codes:  0 '***' 0.001 '**' 0.01 '*' 0.05 '.' 0.1 ' ' 1
##
## Approximate significance of smooth terms:
##              edf Ref.df       F p-value
## s(age_z)   6.888  6.888 198.822  <2e-16 ***
## s(sleep_z) 1.000  1.000   1.038   0.308
## ---
## Signif. codes:  0 '***' 0.001 '**' 0.01 '*' 0.05 '.' 0.1 ' ' 1
##
## R-sq.(adj) =  0.423
## lmer.REML = 7.3686e+05  Scale est. = 11450     n = 51297
```

Model with full interaction between age and sleep

```
##
## Family: gaussian
```

```
## Link function: identity
##
## Formula:
## value ~ sex + site + icv + t2(age_z, sleep_z, k = c(10, 4), bs = "cr")
## <environment: 0x55cf3c0125d8>
##
## Parametric coefficients:
##                Estimate Std. Error t value Pr(>|t|)
## (Intercept)   3627.991     18.878 192.176  < 2e-16 ***
## sexmale         85.471      3.878  22.041  < 2e-16 ***
## siteMPIB       511.657     24.201  21.142  < 2e-16 ***
## siteousAvanto  110.654     19.793   5.591 2.28e-08 ***
## siteousPrisma  227.351     33.473   6.792 1.12e-11 ***
## siteousSkyra   228.620     18.488  12.366  < 2e-16 ***
## siteUB          74.304     56.415   1.317    0.188
## siteUCAM       255.798     20.980  12.193  < 2e-16 ***
## siteUKB        341.685     19.353  17.655  < 2e-16 ***
## siteUmU        136.112     26.977   5.046 4.54e-07 ***
## siteUOXF        99.123     22.962   4.317 1.59e-05 ***
## icv            259.959      1.948 133.461  < 2e-16 ***
## ---
## Signif. codes:  0 '***' 0.001 '**' 0.01 '*' 0.05 '.' 0.1 ' ' 1
##
## Approximate significance of smooth terms:
##                       edf Ref.df     F p-value
## t2(age_z,sleep_z) 11.06  11.06 25.84  <2e-16 ***
## ---
## Signif. codes:  0 '***' 0.001 '**' 0.01 '*' 0.05 '.' 0.1 ' ' 1
##
## R-sq.(adj) =  0.423
## lmer.REML = 7.3686e+05  Scale est. = 11441     n = 51297
```

#### Model comparison

`mod_no_sleep` refers to model without sleep term, `mod_no_interaction` refers to model with only main effect of sleep, and `mod_full` refers to model with a full interaction between age and sleep. This is a nested model comparison, and the p-value at a given line refers to comparing the model at the line to the model on the line above. Hence, significance implies that the more complicated model is supported on statistical grounds.

To be even more specific, the p-value on the second row tests whether there is an association between sleep and volume. The p-value on the third row tests whether this association depends on age.

```
## Data: NULL
## Models:
## mod_list$mod_no_sleep$mer: NULL
## mod_list$mod_no_interaction$mer: NULL
## mod_list$mod_full$mer: NULL
##                                 npar    AIC    BIC  logLik deviance  Chisq Df Pr(>Chisq)
## mod_list$mod_no_sleep$mer         16 736896 737037 -368432   736864
## mod_list$mod_no_interaction$mer   18 736899 737058 -368431   736863 1.0385  2     0.5950
## mod_list$mod_full$mer             20 736899 737076 -368429   736859 3.6731  2     0.1594
```

We chose the model based on the likelihood ratio test with 5 % significance level, which was `mod_no_sleep`.

##### Lifespan brain trajectory

The trajectory shown is from the chosen model `mod_no_sleep`.

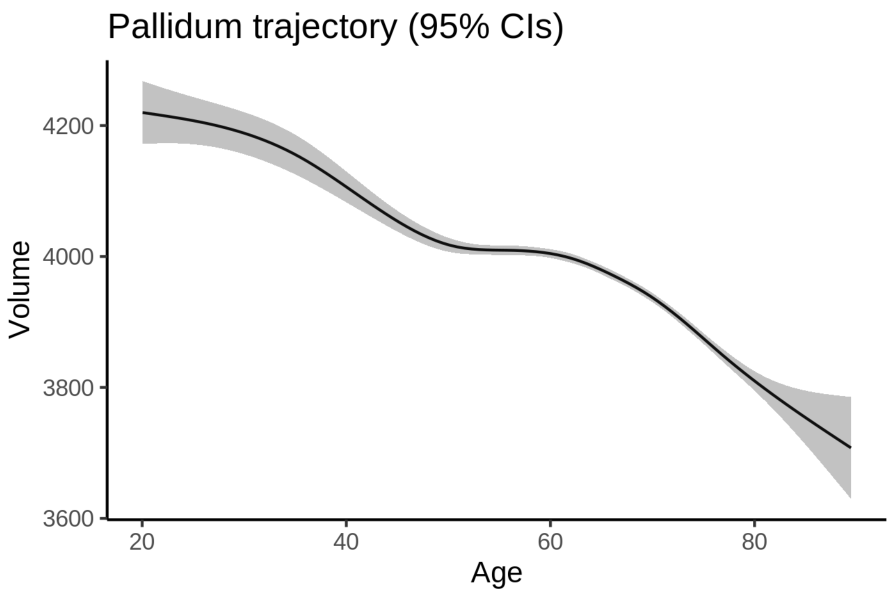


##### Effect of sleep

The chosen model did not include a sleep term, and hence we don't have any estimated effect of sleep.

We show the full interaction model for completeness, although it was not selected.

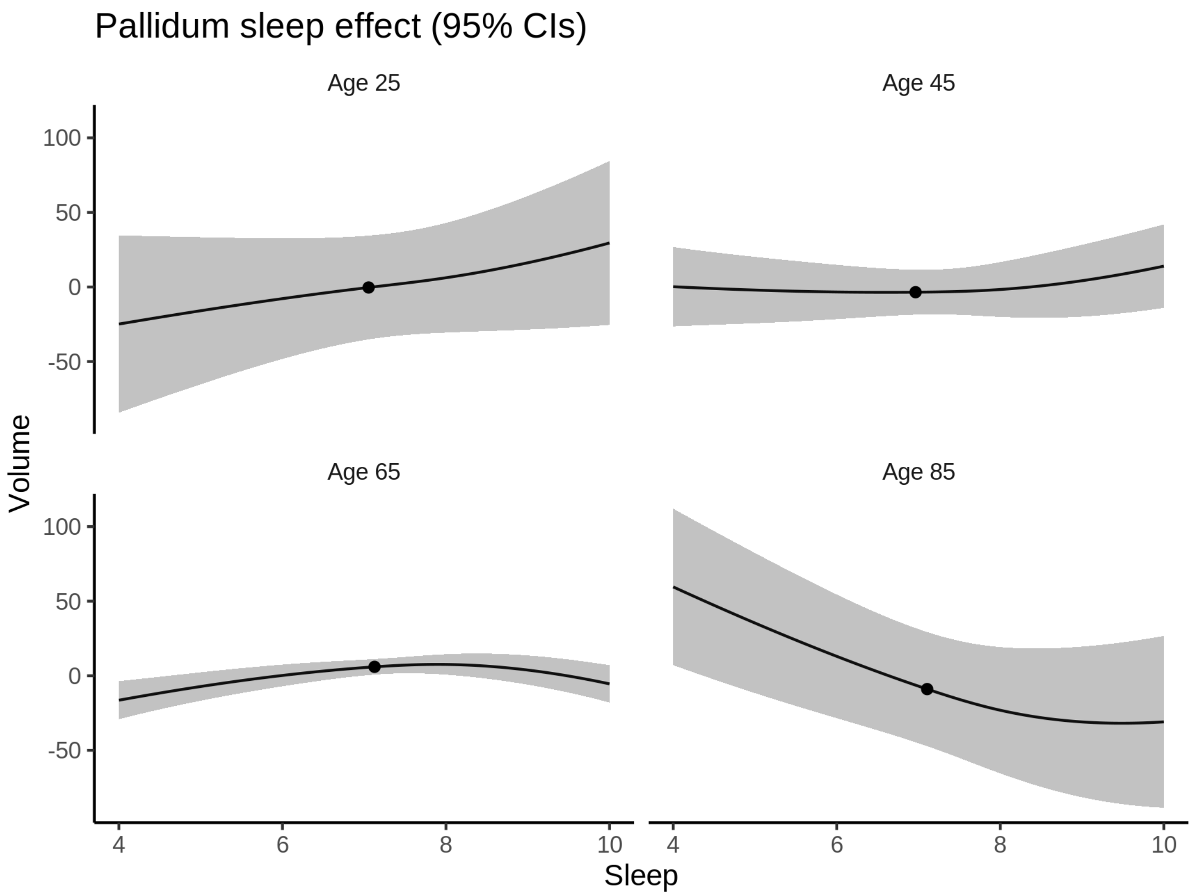


###### Deviation from sleep associated with maximal volume

Model with no sleep term was selected. No plots to show. (Although we can of course dig up the plots, which will be pretty flat).

###### Comparison of mean sleep and sleep associated with maximum volume

Nothing to show, as we did not find an association between sleep and volume.

#### Putamen

##### Descriptive statistics

| Study | Observations | Unique IDs | Mean age | Age range |
|---|---|---|---|---|
| HCP | 974 | 974 | 28.8 | 22 - 37 |
| MPIB | 677 | 391 | 63.1 | 24 - 83 |
| UB | 113 | 39 | 70.9 | 64 - 81 |
| UCAM | 884 | 632 | 55.1 | 20 - 88 |
| UiO | 1469 | 802 | 49.3 | 20 - 88 |
| UKB | 45969 | 43130 | 64.5 | 45 - 83 |

| Study | Observations | Unique IDs | Mean age | Age range |
|---|---|---|---|---|
| UmU | 423 | 284 | 62.3 | 25 - 85 |
| UOXF | 769 | 769 | 69.8 | 60 - 85 |

##### Spaghetti plot

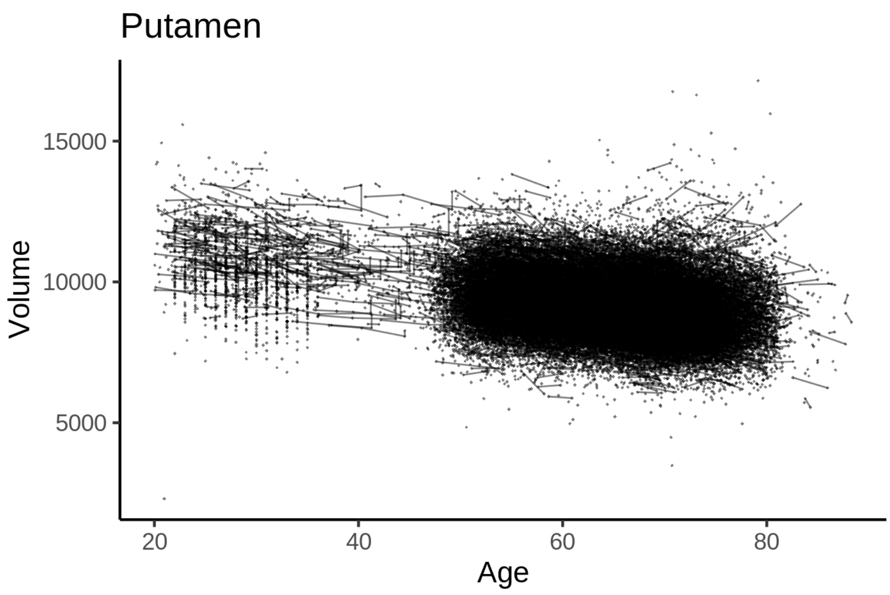


##### Model outputs

Model without sleep term

```
##
## Family: gaussian
## Link function: identity
##
## Formula:
## value ~ sex + site + icv + s(age_z, k = 10, bs = "cr")
## <environment: 0x55cf36e17858>
##
## Parametric coefficients:
##                Estimate Std. Error t value Pr(>|t|)
## (Intercept)   8661.443     45.829 188.994  < 2e-16 ***
## sexmale        356.849      9.804  36.399  < 2e-16 ***
## siteMPIB      1260.091     60.387  20.867  < 2e-16 ***
## siteousAvanto  601.066     47.397  12.681  < 2e-16 ***
## siteousPrisma  768.506     79.982   9.608  < 2e-16 ***
## siteousSkyra  1240.209     45.879  27.032  < 2e-16 ***
## siteUB         777.051    144.441   5.380 7.49e-08 ***
## siteUCAM       666.852     52.083  12.804  < 2e-16 ***
## siteUKB        349.492     46.921   7.449 9.59e-14 ***
## siteUmU        723.407     67.241  10.758  < 2e-16 ***
## siteUOXF       864.405     56.408  15.324  < 2e-16 ***
## icv            494.164      4.917 100.495  < 2e-16 ***
```

```
## ---
## Signif. codes:  0 ‘***’ 0.001 ‘**’ 0.01 ‘*’ 0.05 ‘.’ 0.1 ‘ ’ 1
##
## Approximate significance of smooth terms:
##             edf Ref.df     F p-value
## s(age_z) 5.824  5.824 886.5  <2e-16 ***
## ---
## Signif. codes:  0 ‘***’ 0.001 ‘**’ 0.01 ‘*’ 0.05 ‘.’ 0.1 ‘ ’ 1
##
## R-sq.(adj) =  0.408
## lmer.REML = 8.29e+05  Scale est. = 35692     n = 51278
```

Model with only main effects of age and sleep

```
##
## Family: gaussian
## Link function: identity
##
## Formula:
## value ~ sex + site + icv + s(age_z, k = 10, bs = "cr") + s(sleep_z,
##     k = 5, bs = "cr")
## <environment: 0x55cf36e17858>
##
## Parametric coefficients:
##               Estimate Std. Error t value Pr(>|t|)
## (Intercept)   8658.548     45.879 188.725  < 2e-16 ***
## sexmale        356.754      9.804  36.390  < 2e-16 ***
## siteMPIB      1263.236     60.429  20.905  < 2e-16 ***
## siteousAvanto  602.625     47.411  12.711  < 2e-16 ***
## siteousPrisma  769.601     79.986   9.622  < 2e-16 ***
## siteousSkyra  1241.801     45.893  27.058  < 2e-16 ***
## siteUB         777.924    144.441   5.386 7.25e-08 ***
## siteUCAM       668.294     52.095  12.828  < 2e-16 ***
## siteUKB        352.565     46.976   7.505 6.23e-14 ***
## siteUmU        729.847     67.398  10.829  < 2e-16 ***
## siteUOXF       865.874     56.423  15.346  < 2e-16 ***
## icv            494.370      4.919 100.494  < 2e-16 ***
## ---
## Signif. codes:  0 ‘***’ 0.001 ‘**’ 0.01 ‘*’ 0.05 ‘.’ 0.1 ‘ ’ 1
##
## Approximate significance of smooth terms:
##               edf Ref.df       F p-value
## s(age_z)   5.836  5.836 881.518  <2e-16 ***
## s(sleep_z) 1.000  1.000   1.893   0.169
## ---
## Signif. codes:  0 ‘***’ 0.001 ‘**’ 0.01 ‘*’ 0.05 ‘.’ 0.1 ‘ ’ 1
##
## R-sq.(adj) =  0.408
## lmer.REML = 8.29e+05  Scale est. = 35695     n = 51278
```

Model with full interaction between age and sleep

```
##
## Family: gaussian
## Link function: identity
##
```

```
## Formula:
## value ~ sex + site + icv + t2(age_z, sleep_z, k = c(10, 4), bs = "cr")
## <environment: 0x55cf36e17858>
##
## Parametric coefficients:
##                Estimate Std. Error t value Pr(>|t|)
## (Intercept)    8666.581     46.056 188.175  < 2e-16 ***
## sexmale         356.637      9.822  36.310  < 2e-16 ***
## siteMPIB       1256.471     60.726  20.691  < 2e-16 ***
## siteousAvanto   596.990     47.673  12.523  < 2e-16 ***
## siteousPrisma   764.751     80.152   9.541  < 2e-16 ***
## siteousSkyra   1236.082     46.184  26.764  < 2e-16 ***
## siteUB          770.514    144.527   5.331 9.79e-08 ***
## siteUCAM        661.042     52.374  12.622  < 2e-16 ***
## siteUKB         344.332     47.153   7.302 2.87e-13 ***
## siteUmU         721.813     67.595  10.678  < 2e-16 ***
## siteUOXF        857.948     56.629  15.150  < 2e-16 ***
## icv             494.317      4.920 100.468  < 2e-16 ***
## ---
## Signif. codes:  0 '***' 0.001 '**' 0.01 '*' 0.05 '.' 0.1 ' ' 1
##
## Approximate significance of smooth terms:
##                      edf Ref.df     F p-value
## t2(age_z,sleep_z) 8.953  8.953 50.65  <2e-16 ***
## ---
## Signif. codes:  0 '***' 0.001 '**' 0.01 '*' 0.05 '.' 0.1 ' ' 1
##
## R-sq.(adj) =  0.408
## lmer.REML = 8.29e+05  Scale est. = 35691     n = 51278
```

#### Model comparison

`mod_no_sleep` refers to model without sleep term, `mod_no_interaction` refers to model with only main effect of sleep, and `mod_full` refers to model with a full interaction between age and sleep. This is a nested model comparison, and the p-value at a given line refers to comparing the model at the line to the model on the line above. Hence, significance implies that the more complicated model is supported on statistical grounds.

To be even more specific, the p-value on the second row tests whether there is an association between sleep and volume. The p-value on the third row tests whether this association depends on age.

```
## Data: NULL
## Models:
## mod_list$mod_no_sleep$mer: NULL
## mod_list$mod_no_interaction$mer: NULL
## mod_list$mod_full$mer: NULL
##                                 npar    AIC    BIC  logLik deviance  Chisq Df Pr(>Chisq)
## mod_list$mod_no_sleep$mer         16 829035 829177 -414502   829003
## mod_list$mod_no_interaction$mer   18 829037 829196 -414501   829001 1.8926  2     0.3882
## mod_list$mod_full$mer             20 829045 829222 -414502   829005 0.0000  2     1.0000
```

We chose the model based on the likelihood ratio test with 5 % significance level, which was `mod_no_sleep`.

#### Lifespan brain trajectory

The trajectory shown is from the chosen model `mod_no_sleep`.

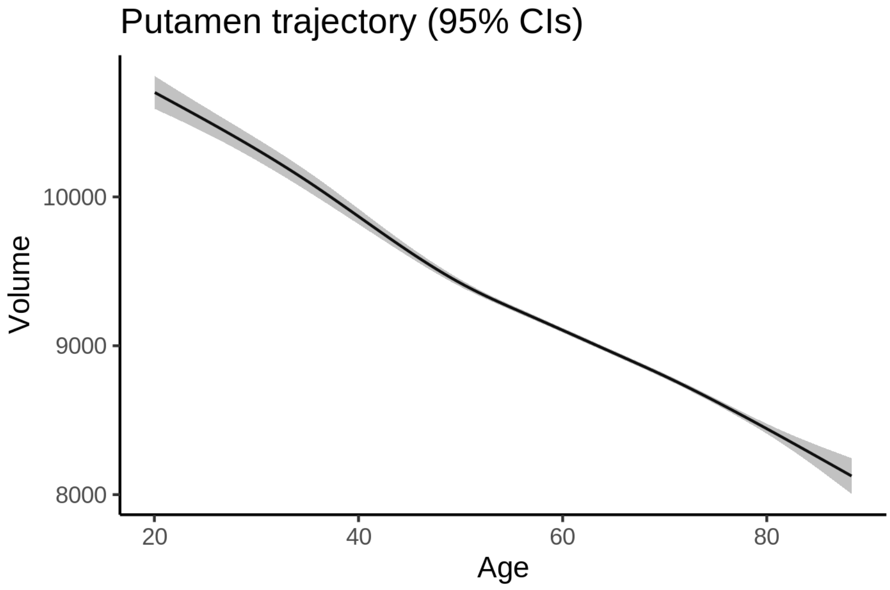


#### Effect of sleep

The chosen model did not include a sleep term, and hence we don't have any estimated effect of sleep.

We show the full interaction model for completeness, although it was not selected.

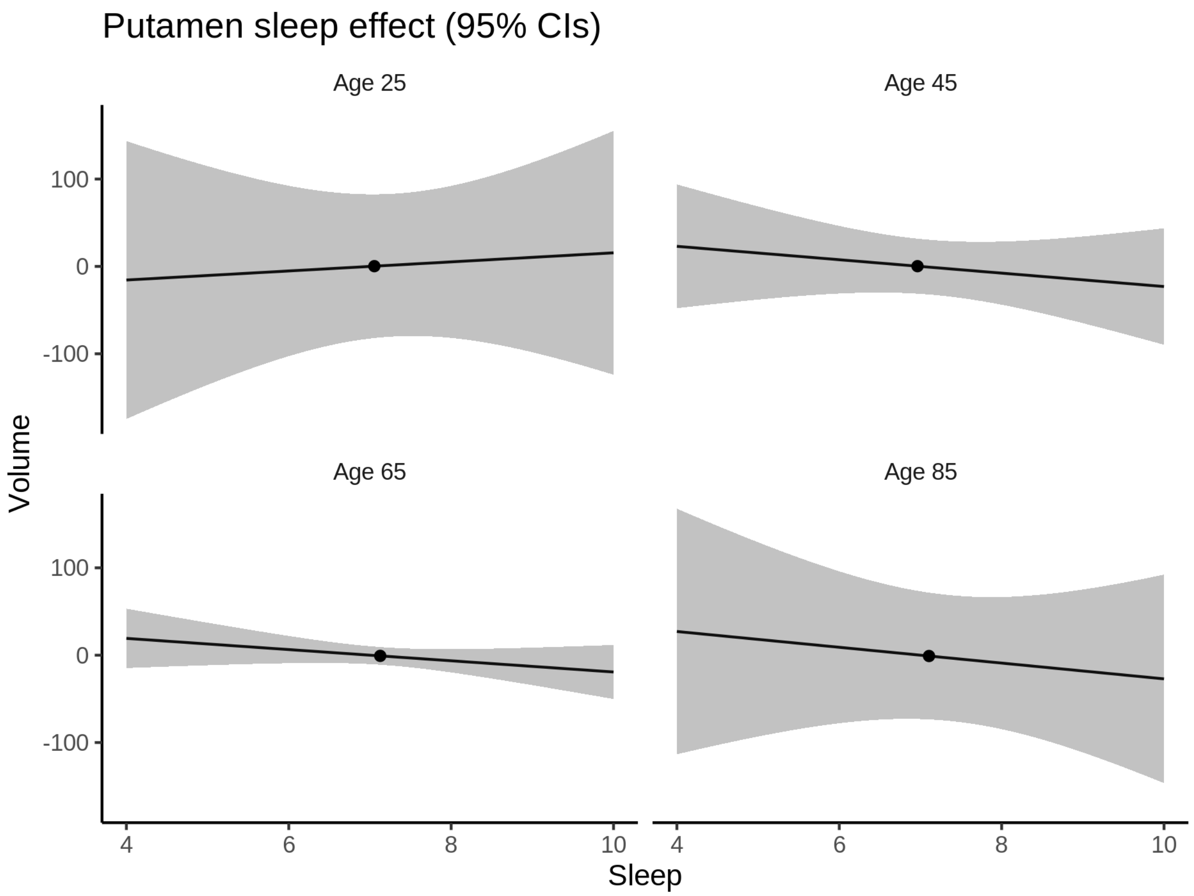


**Deviation from sleep associated with maximal volume**

Model with no sleep term was selected. No plots to show. (Although we can of course dig up the plots, which will be pretty flat).

**Comparison of mean sleep and sleep associated with maximum volume**

Nothing to show, as we did not find an association between sleep and volume.

### Thalamus

#### Descriptive statistics

| Study | Observations | Unique IDs | Mean age | Age range |
|---|---|---|---|---|
| HCP | 974 | 974 | 28.8 | 22 - 37 |
| MPIB | 677 | 391 | 63.1 | 24 - 83 |
| UB | 113 | 39 | 70.9 | 64 - 81 |
| UCAM | 884 | 632 | 55.1 | 20 - 88 |
| UiO | 1475 | 803 | 49.4 | 20 - 89 |
| UKB | 45975 | 43133 | 64.5 | 45 - 83 |

| Study | Observations | Unique IDs | Mean age | Age range |
|---|---|---|---|---|
| UmU | 423 | 284 | 62.3 | 25 - 85 |
| UOXF | 769 | 769 | 69.8 | 60 - 85 |

##### Spaghetti plot

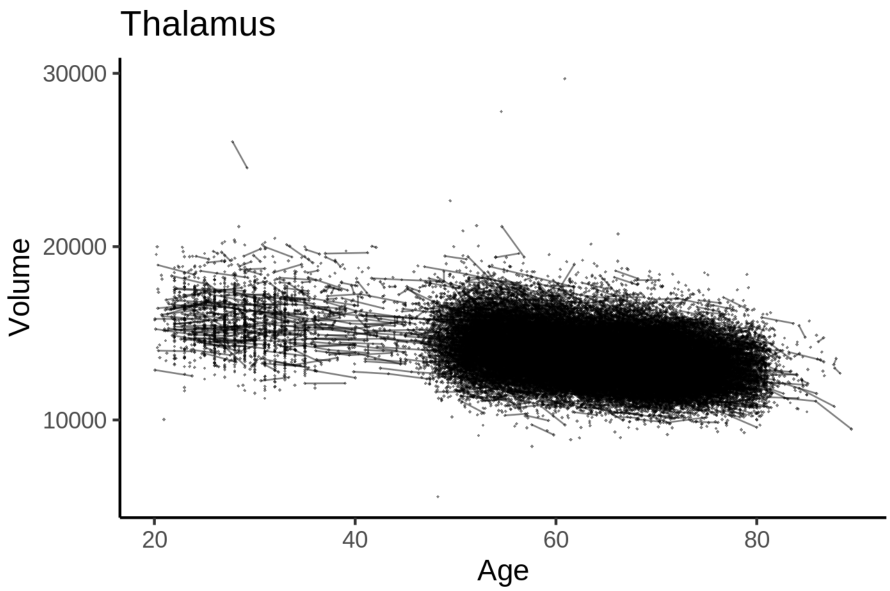


##### Model outputs

Model without sleep term

```
##
## Family: gaussian
## Link function: identity
##
## Formula:
## value ~ sex + site + icv + s(age_z, k = 10, bs = "cr")
## <environment: 0x55cffa104428>
##
## Parametric coefficients:
##                 Estimate Std. Error t value Pr(>|t|)
## (Intercept)   13375.389     52.667 253.963  < 2e-16 ***
## sexmale         197.026     10.656  18.490  < 2e-16 ***
## siteMPIB       1613.341     66.767  24.164  < 2e-16 ***
## siteousAvanto   306.103     54.184   5.649 1.62e-08 ***
## siteousPrisma  1061.610     91.549  11.596  < 2e-16 ***
## siteousSkyra    408.481     50.891   8.027 1.02e-15 ***
## siteUB         1052.544    155.959   6.749 1.51e-11 ***
## siteUCAM       1052.585     57.951  18.163  < 2e-16 ***
## siteUKB         204.487     54.030   3.785 0.000154 ***
## siteUmU         -29.398     74.476  -0.395 0.693039
## siteUOXF        910.068     63.781  14.269  < 2e-16 ***
## icv             880.265      5.359 164.247  < 2e-16 ***
```

```
## ---
## Signif. codes:  0 ‘***’ 0.001 ‘**’ 0.01 ‘*’ 0.05 ‘.’ 0.1 ‘ ’ 1
##
## Approximate significance of smooth terms:
##            edf Ref.df    F p-value
## s(age_z) 7.867  7.867 1555  <2e-16 ***
## ---
## Signif. codes:  0 ‘***’ 0.001 ‘**’ 0.01 ‘*’ 0.05 ‘.’ 0.1 ‘ ’ 1
##
## R-sq.(adj) =  0.581
## lmer.REML = 8.403e+05  Scale est. = 79373     n = 51290
```

Model with only main effects of age and sleep

```
##
## Family: gaussian
## Link function: identity
##
## Formula:
## value ~ sex + site + icv + s(age_z, k = 10, bs = "cr") + s(sleep_z,
##     k = 5, bs = "cr")
## <environment: 0x55cffa104428>
##
## Parametric coefficients:
##                Estimate Std. Error t value Pr(>|t|)
## (Intercept)   13369.602     52.712 253.633  < 2e-16 ***
## sexmale         196.605     10.653  18.455  < 2e-16 ***
## siteMPIB       1623.150     66.793  24.301  < 2e-16 ***
## siteousAvanto   308.055     54.192   5.684 1.32e-08 ***
## siteousPrisma  1062.925     91.531  11.613  < 2e-16 ***
## siteousSkyra    410.070     50.899   8.056 8.02e-16 ***
## siteUB         1058.758    155.903   6.791 1.12e-11 ***
## siteUCAM       1057.814     57.947  18.255  < 2e-16 ***
## siteUKB         210.697     54.081   3.896 9.79e-05 ***
## siteUmU          -8.803     74.631  -0.118    0.906
## siteUOXF        913.212     63.776  14.319  < 2e-16 ***
## icv             880.163      5.363 164.103  < 2e-16 ***
## ---
## Signif. codes:  0 ‘***’ 0.001 ‘**’ 0.01 ‘*’ 0.05 ‘.’ 0.1 ‘ ’ 1
##
## Approximate significance of smooth terms:
##              edf Ref.df        F  p-value
## s(age_z)   7.871  7.871 1537.252  < 2e-16 ***
## s(sleep_z) 3.017  3.017    9.202 4.49e-06 ***
## ---
## Signif. codes:  0 ‘***’ 0.001 ‘**’ 0.01 ‘*’ 0.05 ‘.’ 0.1 ‘ ’ 1
##
## R-sq.(adj) =  0.582
## lmer.REML = 8.4028e+05  Scale est. = 79443     n = 51290
```

Model with full interaction between age and sleep

```
##
## Family: gaussian
## Link function: identity
##
```

```
## Formula:
## value ~ sex + site + icv + t2(age_z, sleep_z, k = c(10, 4), bs = "cr")
## <environment: 0x55cffa104428>
##
## Parametric coefficients:
##                 Estimate Std. Error t value Pr(>|t|)
## (Intercept)   13372.186     52.910 252.737  < 2e-16 ***
## sexmale         196.301     10.677  18.385  < 2e-16 ***
## siteMPIB       1620.768     67.155  24.135  < 2e-16 ***
## siteousAvanto   308.453     54.481   5.662 1.51e-08 ***
## siteousPrisma  1062.274     91.748  11.578  < 2e-16 ***
## siteousSkyra    411.341     51.224   8.030 9.93e-16 ***
## siteUB         1052.036    156.019   6.743 1.57e-11 ***
## siteUCAM       1055.088     58.265  18.108  < 2e-16 ***
## siteUKB         208.134     54.277   3.835 0.000126 ***
## siteUmU         -13.106     74.871  -0.175 0.861047
## siteUOXF        909.819     64.013  14.213  < 2e-16 ***
## icv             880.439      5.362 164.185  < 2e-16 ***
## ---
## Signif. codes:  0 '***' 0.001 '**' 0.01 '*' 0.05 '.' 0.1 ' ' 1
##
## Approximate significance of smooth terms:
##                     edf Ref.df    F p-value
## t2(age_z,sleep_z) 14.53  14.53 28.8  <2e-16 ***
## ---
## Signif. codes:  0 '***' 0.001 '**' 0.01 '*' 0.05 '.' 0.1 ' ' 1
##
## R-sq.(adj) =  0.582
## lmer.REML = 8.4029e+05  Scale est. = 79414     n = 51290
```

#### Model comparison

`mod_no_sleep` refers to model without sleep term, `mod_no_interaction` refers to model with only main effect of sleep, and `mod_full` refers to model with a full interaction between age and sleep. This is a nested model comparison, and the p-value at a given line refers to comparing the model at the line to the model on the line above. Hence, significance implies that the more complicated model is supported on statistical grounds.

To be even more specific, the p-value on the second row tests whether there is an association between sleep and volume. The p-value on the third row tests whether this association depends on age.

```
## Data: NULL
## Models:
## mod_list$mod_no_sleep$mer: NULL
## mod_list$mod_no_interaction$mer: NULL
## mod_list$mod_full$mer: NULL
##                                 npar    AIC    BIC  logLik deviance Chisq Df Pr(>Chisq)
## mod_list$mod_no_sleep$mer         16 840335 840476 -420151   840303
## mod_list$mod_no_interaction$mer   18 840316 840475 -420140   840280  22.9  2  1.065e-05 ***
## mod_list$mod_full$mer             20 840330 840507 -420145   840290   0.0  2          1
## ---
## Signif. codes:  0 '***' 0.001 '**' 0.01 '*' 0.05 '.' 0.1 ' ' 1
```

We chose the model based on the likelihood ratio test with 5 % significance level, which was `mod_no_interaction`.

##### Lifespan brain trajectory

The trajectory shown is from the chosen model `mod_no_interaction`.

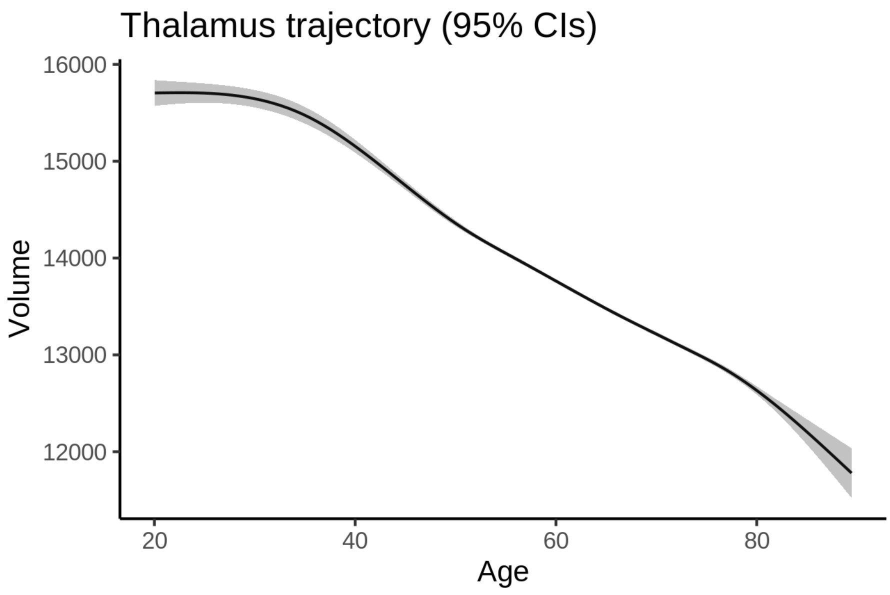


##### Effect of sleep

The chosen model only included the main effect of sleep, and hence the effect does not vary with age. The black dot shows the average sleep duration across all ages in the sample.

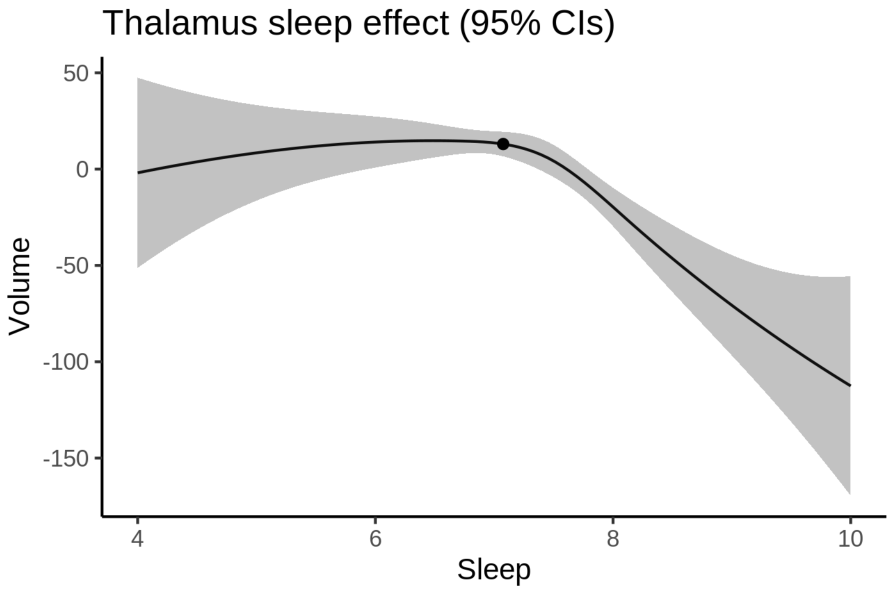


We also show the full interaction model for completeness, although it was not selected.

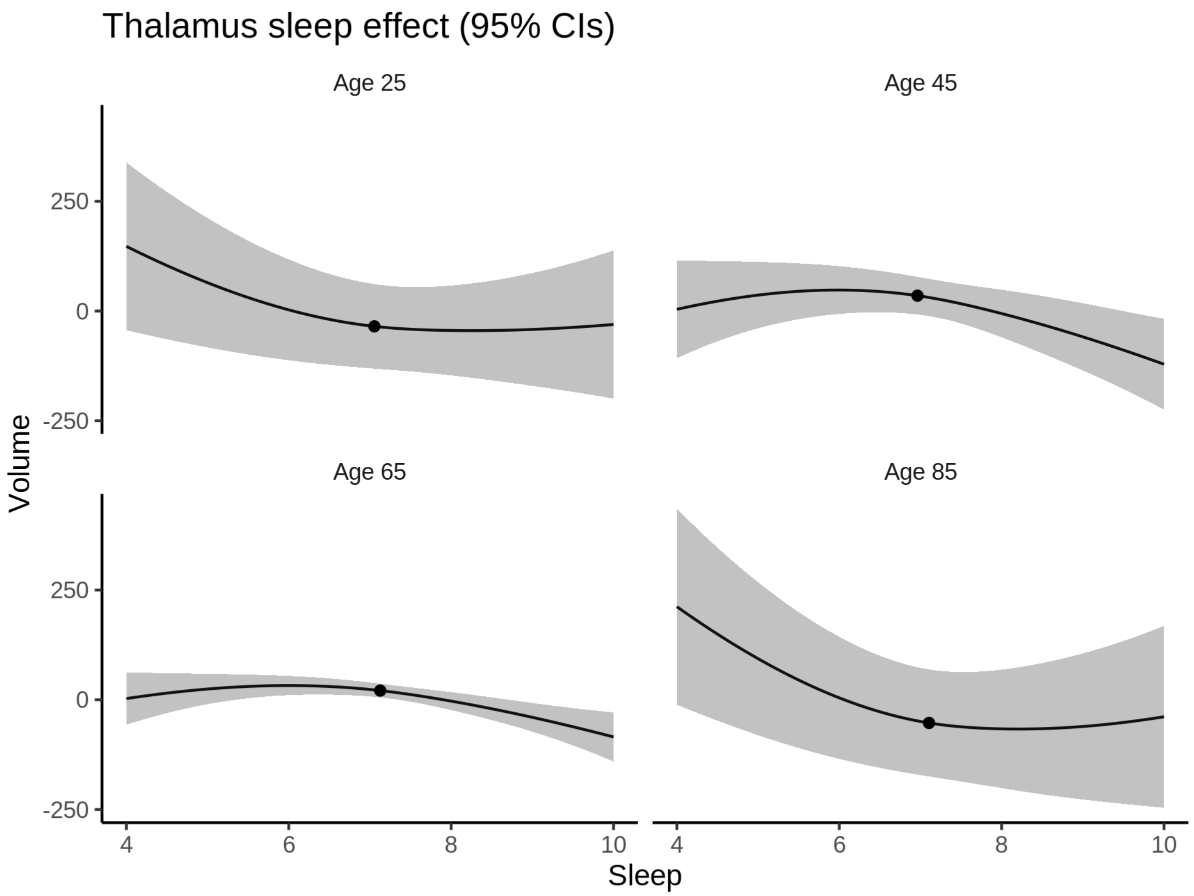


**Deviation from sleep associated with maximal volume**

Model with only main effect of sleep was chosen, so we show it for all ages at once. Maximum volume is attained at 6.5 hours of sleep. The percentage values in the plot are calculated as follows: The maximum at 100 % refers to a person at an arbitrary age with a sleep duration associated with maximum volume. For a female, this volume is 14076 and for a male it is 14273. The other percentage values show how large the expected volume is for someone with other sleep durations. For example, 99 % implies a 1 % reduction.

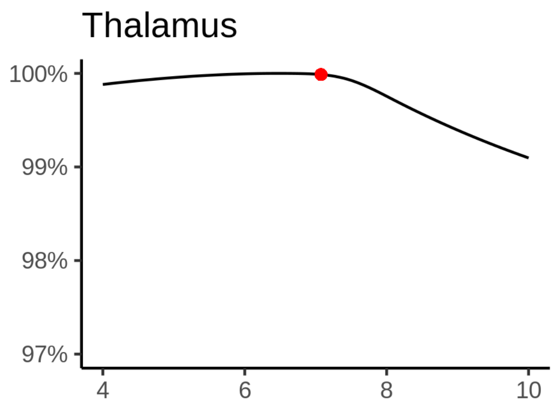

**Comparison of mean sleep and sleep associated with maximum volume**

A 95 % confidence interval for the sleep associated with maximum volume is [4, 7.15].

The plot below compares average sleep to the sleep associated with maximum volume.

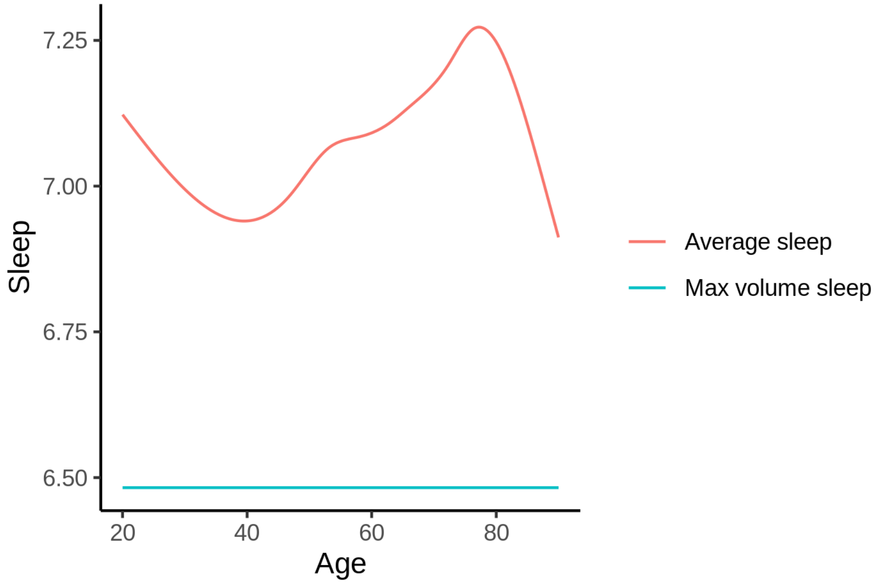


The next plot shows the difference between average sleep and sleep associated with maximum volume. The shaded region is a 95 % confidence interval.

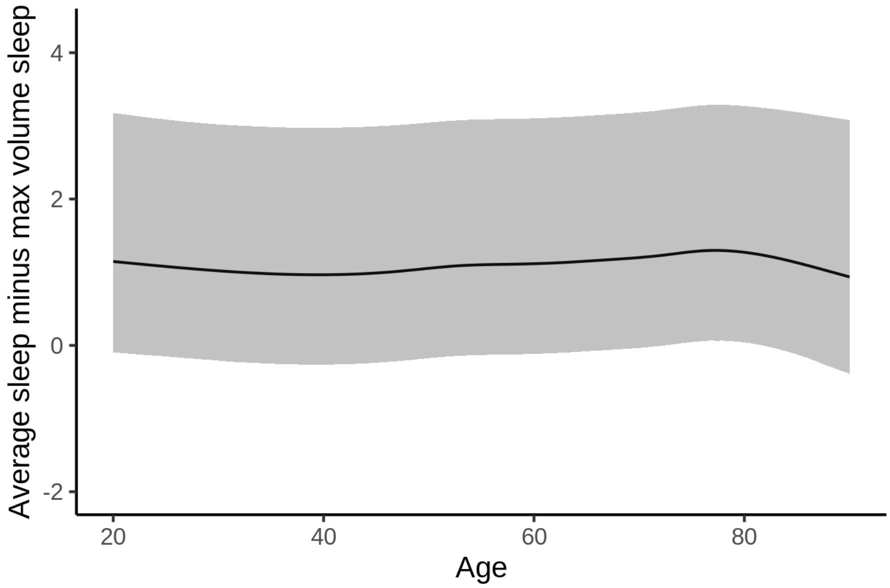


The next plot shows the probability that the sleep duration associated with maximum volume is longer than the average sleep duration, as a function of age. Probility below .05 can be interpreted as evidence that the sleep associated with maximum volume is shorter than the mean sleep, and probability above .95 can be interpreted the opposite way.

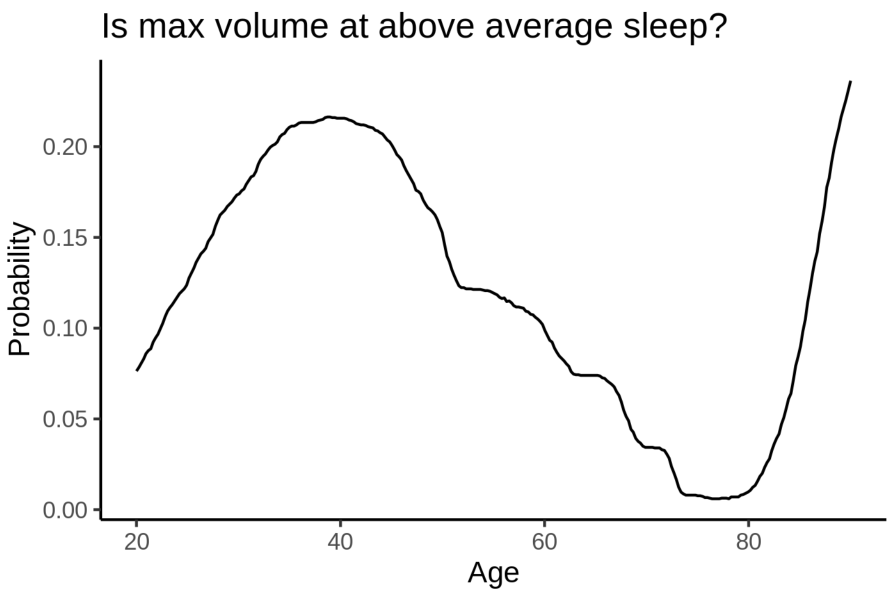


##### Controling for covariates

Below is the output for a model in which we only include data with income and education.

```
##
## Family: gaussian
## Link function: identity
##
## Formula:
## value ~ sex + site + s(age_z, k = 10, bs = "cr") + s(sleep_z,
##     k = 5, bs = "cr") + icv
## <environment: 0x55cf411e55b0>
##
## Parametric coefficients:
##                Estimate Std. Error t value Pr(>|t|)
## (Intercept)   15076.901    120.778 124.832  < 2e-16 ***
## sexmale         174.596     13.418  13.012  < 2e-16 ***
## siteousAvanto -1709.261    147.616 -11.579  < 2e-16 ***
## siteousPrisma  -406.556    149.583  -2.718  0.00657 **
## siteousSkyra  -1297.356    131.679  -9.852  < 2e-16 ***
## siteUKB       -1536.755    120.485 -12.755  < 2e-16 ***
## siteUOXF       -711.377    135.975  -5.232 1.69e-07 ***
## icv             898.979      6.896 130.364  < 2e-16 ***
## ---
## Signif. codes:  0 ‘***’ 0.001 ‘**’ 0.01 ‘*’ 0.05 ‘.’ 0.1 ‘ ’ 1
##
## Approximate significance of smooth terms:
##              edf Ref.df        F  p-value
## s(age_z)   3.331  3.331 1791.515  < 2e-16 ***
## s(sleep_z) 2.771  2.771    7.336 0.000221 ***
## ---
## Signif. codes:  0 ‘***’ 0.001 ‘**’ 0.01 ‘*’ 0.05 ‘.’ 0.1 ‘ ’ 1
##
```

```
## R-sq.(adj) =  0.555
## lmer.REML = 5.1044e+05  Scale est. = 81739     n = 31194
```

Below is the output for a model in which we control for the main effects of income and education.

```
##
## Family: gaussian
## Link function: identity
##
## Formula:
## value ~ sex + site + s(age_z, k = 10, bs = "cr") + s(sleep_z,
##     k = 5, bs = "cr") + icv + income_scaled + education_scaled
## <environment: 0x55cf411e55b0>
##
## Parametric coefficients:
##                   Estimate Std. Error t value Pr(>|t|)
## (Intercept)      15046.075    121.464 123.873  < 2e-16 ***
## sexmale            174.729     13.455  12.986  < 2e-16 ***
## siteousAvanto    -1709.996    147.605 -11.585  < 2e-16 ***
## siteousPrisma     -409.784    149.538  -2.740  0.00614 **
## siteousSkyra     -1298.148    131.688  -9.858  < 2e-16 ***
## siteUKB          -1537.457    120.467 -12.763  < 2e-16 ***
## siteUOXF          -702.322    135.969  -5.165 2.42e-07 ***
## icv                896.973      6.954 128.982  < 2e-16 ***
## income_scaled       19.899     15.974   1.246  0.21288
## education_scaled    28.297     18.897   1.497  0.13429
## ---
## Signif. codes:  0 ‘***’ 0.001 ‘**’ 0.01 ‘*’ 0.05 ‘.’ 0.1 ‘ ’ 1
##
## Approximate significance of smooth terms:
##              edf Ref.df        F  p-value
## s(age_z)   3.241  3.241 1688.365  < 2e-16 ***
## s(sleep_z) 2.665  2.665    7.206 0.000326 ***
## ---
## Signif. codes:  0 ‘***’ 0.001 ‘**’ 0.01 ‘*’ 0.05 ‘.’ 0.1 ‘ ’ 1
##
## R-sq.(adj) =  0.555
## lmer.REML = 5.1042e+05  Scale est. = 81800     n = 31194
```

We also included interaction effects between sleep duration and education and income, in another model. The output is shown below, and the interaction terms are `income_scaled:sleep_z` and `education_scaled:sleep_z`.

```
##
## Family: gaussian
## Link function: identity
##
## Formula:
## value ~ sex + site + s(age_z, k = 10, bs = "cr") + s(sleep_z,
##     k = 5, bs = "cr") + icv + income_scaled + education_scaled +
##     income_scaled:sleep_z + education_scaled:sleep_z
## <environment: 0x55cf411e55b0>
##
## Parametric coefficients:
##                           Estimate Std. Error t value Pr(>|t|)
## (Intercept)              15045.297    121.455 123.875  < 2e-16 ***
```

```
## sexmale                    174.148     13.464  12.934  < 2e-16 ***
## siteousAvanto            -1710.595    147.596 -11.590  < 2e-16 ***
## siteousPrisma             -411.496    149.542  -2.752  0.00593 **
## siteousSkyra             -1298.685    131.686  -9.862  < 2e-16 ***
## siteUKB                  -1536.414    120.462 -12.754  < 2e-16 ***
## siteUOXF                  -699.674    135.993  -5.145 2.69e-07 ***
## icv                        896.974      6.955 128.971  < 2e-16 ***
## income_scaled               20.515     15.981   1.284  0.19926
## education_scaled            28.010     18.900   1.482  0.13834
## income_scaled:sleep_z      -21.628     15.926  -1.358  0.17445
## education_scaled:sleep_z     1.352     18.344   0.074  0.94124
## ---
## Signif. codes:  0 '***' 0.001 '**' 0.01 '*' 0.05 '.' 0.1 ' ' 1
##
## Approximate significance of smooth terms:
##              edf Ref.df        F p-value
## s(age_z)   3.217  3.217 1699.202  <2e-16 ***
## s(sleep_z) 2.673  2.673    2.868   0.069 .
## ---
## Signif. codes:  0 '***' 0.001 '**' 0.01 '*' 0.05 '.' 0.1 ' ' 1
##
## R-sq.(adj) =  0.555
## lmer.REML = 5.104e+05  Scale est. = 81806     n = 31194
```

We did the same controling for BMI. Below is the model with no covariates but only keeping data with BMI.

```
##
## Family: gaussian
## Link function: identity
##
## Formula:
## value ~ sex + site + s(age_z, k = 10, bs = "cr") + s(sleep_z,
##     k = 5, bs = "cr") + icv
## <environment: 0x55cff9b36638>
##
## Parametric coefficients:
##                Estimate Std. Error t value Pr(>|t|)
## (Intercept)   13635.869     85.714 159.085  < 2e-16 ***
## sexmale         163.769     13.030  12.569  < 2e-16 ***
## siteousPrisma   822.343    112.901   7.284 3.32e-13 ***
## siteousSkyra    232.174     68.558   3.387 0.000709 ***
## siteUCAM        814.823     90.175   9.036  < 2e-16 ***
## siteUKB         -67.346     86.384  -0.780 0.435624
## siteUmU        -233.709    101.216  -2.309 0.020949 *
## icv             909.969      6.716 135.495  < 2e-16 ***
## ---
## Signif. codes:  0 '***' 0.001 '**' 0.01 '*' 0.05 '.' 0.1 ' ' 1
##
## Approximate significance of smooth terms:
##              edf Ref.df        F  p-value
## s(age_z)   5.651  5.651 1366.485  < 2e-16 ***
## s(sleep_z) 2.589  2.589    6.966 0.000428 ***
## ---
## Signif. codes:  0 '***' 0.001 '**' 0.01 '*' 0.05 '.' 0.1 ' ' 1
##
```

#### R-sq.(adj) =  0.582
## lmer.REML = 5.4679e+05  Scale est. = 75474     n = 33449

Below is the model output with main effect.

##
#### Family: gaussian
#### Link function: identity
##
#### Formula:
#### value ~ sex + site + s(age_z, k = 10, bs = "cr") + s(sleep_z,
##     k = 5, bs = "cr") + icv + bmi
#### <environment: 0x55cff9b36638>
##
#### Parametric coefficients:
##                 Estimate Std. Error t value Pr(>|t|)
## (Intercept)   13614.2523    91.4995 148.790  < 2e-16 ***
## sexmale         163.0917    13.0687  12.480  < 2e-16 ***
## siteousPrisma   821.5215   112.9095   7.276 3.52e-13 ***
## siteousSkyra    232.0673    68.5592   3.385 0.000713 ***
## siteUCAM        814.3570    90.1799   9.030  < 2e-16 ***
## siteUKB         -67.9988    86.3937  -0.787 0.431240
## siteUmU        -234.5920   101.2282  -2.317 0.020485 *
## icv             909.8307     6.7189 135.414  < 2e-16 ***
## bmi               0.8566     1.2711   0.674 0.500377
## ---
#### Signif. codes:  0 ‘***’ 0.001 ‘**’ 0.01 ‘*’ 0.05 ‘.’ 0.1 ‘ ’ 1
##
#### Approximate significance of smooth terms:
##              edf Ref.df        F  p-value
## s(age_z)   5.656  5.656 1365.113  < 2e-16 ***
## s(sleep_z) 2.617  2.617    6.669 0.000438 ***
## ---
#### Signif. codes:  0 ‘***’ 0.001 ‘**’ 0.01 ‘*’ 0.05 ‘.’ 0.1 ‘ ’ 1
##
#### R-sq.(adj) =  0.582
## lmer.REML = 5.4678e+05  Scale est. = 75476     n = 33449

Next is the model with BMI-sleep interaction.

##
#### Family: gaussian
#### Link function: identity
##
#### Formula:
#### value ~ sex + site + s(age_z, k = 10, bs = "cr") + s(sleep_z,
##     k = 5, bs = "cr") + icv + bmi + bmi:sleep_z
#### <environment: 0x55cff9b36638>
##
#### Parametric coefficients:
##                Estimate Std. Error t value Pr(>|t|)
## (Intercept)   13610.616     91.537 148.690  < 2e-16 ***
## sexmale         163.160     13.069  12.485  < 2e-16 ***
## siteousPrisma   821.591    112.907   7.277  3.5e-13 ***
## siteousSkyra    232.435     68.558   3.390 0.000699 ***
## siteUCAM        815.697     90.183   9.045  < 2e-16 ***

```
## siteUKB         -66.647     86.394  -0.771 0.440462
## siteUmU        -233.461    101.229  -2.306 0.021101 *
## icv             909.910      6.719 135.425  < 2e-16 ***
## bmi               0.912      1.272   0.717 0.473281
## bmi:sleep_z       1.640      1.206   1.360 0.173772
## ---
## Signif. codes:  0 ‘***’ 0.001 ‘**’ 0.01 ‘*’ 0.05 ‘.’ 0.1 ‘ ’ 1
##
## Approximate significance of smooth terms:
##              edf Ref.df        F p-value
## s(age_z)   5.646  5.646 1368.029  <2e-16 ***
## s(sleep_z) 2.565  2.565    3.456    0.02 *
## ---
## Signif. codes:  0 ‘***’ 0.001 ‘**’ 0.01 ‘*’ 0.05 ‘.’ 0.1 ‘ ’ 1
##
## R-sq.(adj) =  0.582
## lmer.REML = 5.4678e+05  Scale est. = 75472     n = 33449
```

We did the same controling for depression. Below is the model with no covariates but only keeping data with depression.

```
##
## Family: gaussian
## Link function: identity
##
## Formula:
## value ~ sex + site + s(age_z, k = 10, bs = "cr") + s(sleep_z,
##     k = 5, bs = "cr") + icv
## <environment: 0x55cf40fe2cc8>
##
## Parametric coefficients:
##                Estimate Std. Error t value Pr(>|t|)
## (Intercept)   14864.399     61.807 240.498  < 2e-16 ***
## sexmale         165.624     13.056  12.686  < 2e-16 ***
## siteousAvanto -1109.929    135.769  -8.175 3.06e-16 ***
## siteousPrisma  -324.664    119.706  -2.712  0.00669 **
## siteousSkyra   -755.799     93.726  -8.064 7.64e-16 ***
## siteUCAM       -448.938     84.864  -5.290 1.23e-07 ***
## siteUKB       -1311.541     61.119 -21.459  < 2e-16 ***
## siteUmU       -1469.083     82.941 -17.712  < 2e-16 ***
## icv             902.973      6.719 134.387  < 2e-16 ***
## ---
## Signif. codes:  0 ‘***’ 0.001 ‘**’ 0.01 ‘*’ 0.05 ‘.’ 0.1 ‘ ’ 1
##
## Approximate significance of smooth terms:
##              edf Ref.df       F  p-value
## s(age_z)   7.099  7.099 951.524  < 2e-16 ***
## s(sleep_z) 2.594  2.594   7.873 0.000155 ***
## ---
## Signif. codes:  0 ‘***’ 0.001 ‘**’ 0.01 ‘*’ 0.05 ‘.’ 0.1 ‘ ’ 1
##
## R-sq.(adj) =   0.57
## lmer.REML = 5.4493e+05  Scale est. = 72753     n = 33361
```

Below is the model output with main effect.

```
##
## Family: gaussian
## Link function: identity
##
## Formula:
## value ~ sex + site + s(age_z, k = 10, bs = "cr") + s(sleep_z,
##     k = 5, bs = "cr") + icv + depression
## <environment: 0x55cf40fe2cc8>
##
## Parametric coefficients:
##                 Estimate Std. Error t value Pr(>|t|)
## (Intercept)   14886.219     62.064 239.853  < 2e-16 ***
## sexmale         163.360     13.067  12.502  < 2e-16 ***
## siteousAvanto -1098.740    135.767  -8.093 6.02e-16 ***
## siteousPrisma  -314.774    119.712  -2.629 0.008557 **
## siteousSkyra   -743.116     93.769  -7.925 2.35e-15 ***
## siteUCAM       -444.007     84.860  -5.232 1.68e-07 ***
## siteUKB       -1320.492     61.155 -21.593  < 2e-16 ***
## siteUmU       -1402.974     84.707 -16.563  < 2e-16 ***
## icv             903.327      6.718 134.464  < 2e-16 ***
## depression     -178.587     46.642  -3.829 0.000129 ***
## ---
## Signif. codes:  0 '***' 0.001 '**' 0.01 '*' 0.05 '.' 0.1 ' ' 1
##
## Approximate significance of smooth terms:
##              edf Ref.df       F p-value
## s(age_z)   7.102  7.102 945.970 < 2e-16 ***
## s(sleep_z) 2.347  2.347   8.394 0.00012 ***
## ---
## Signif. codes:  0 '***' 0.001 '**' 0.01 '*' 0.05 '.' 0.1 ' ' 1
##
## R-sq.(adj) =   0.57
## lmer.REML = 5.4491e+05  Scale est. = 72703     n = 33361
```

Next is the model with depression-sleep interaction.

```
##
## Family: gaussian
## Link function: identity
##
## Formula:
## value ~ sex + site + s(age_z, k = 10, bs = "cr") + s(sleep_z,
##     k = 5, bs = "cr") + icv + depression + depression:sleep_z
## <environment: 0x55cf40fe2cc8>
##
## Parametric coefficients:
##                      Estimate Std. Error t value Pr(>|t|)
## (Intercept)        14886.264     62.065 239.851  < 2e-16 ***
## sexmale              163.187     13.069  12.486  < 2e-16 ***
## siteousAvanto      -1099.863    135.774  -8.101 5.65e-16 ***
## siteousPrisma       -315.911    119.720  -2.639 0.008325 **
## siteousSkyra        -744.221     93.779  -7.936 2.15e-15 ***
## siteUCAM            -445.164     84.875  -5.245 1.57e-07 ***
## siteUKB            -1320.498     61.155 -21.592  < 2e-16 ***
## siteUmU            -1394.521     85.437 -16.322  < 2e-16 ***
```

```
## icv                   903.328      6.718 134.460  < 2e-16 ***
## depression           -180.478     46.739  -3.861 0.000113 ***
## depression:sleep_z   -26.174     34.302  -0.763 0.445446
## ---
## Signif. codes:  0 '***' 0.001 '**' 0.01 '*' 0.05 '.' 0.1 ' ' 1
##
## Approximate significance of smooth terms:
##               edf Ref.df       F p-value
## s(age_z)    7.091  7.091 947.297  <2e-16 ***
## s(sleep_z) 2.396  2.396   5.078  0.0031 **
## ---
## Signif. codes:  0 '***' 0.001 '**' 0.01 '*' 0.05 '.' 0.1 ' ' 1
##
## R-sq.(adj) =   0.57
## lmer.REML = 5.449e+05  Scale est. = 72702     n = 33361
```

The plot below shows the sleep-volume curve for the original model and for the model with main effects of SES.

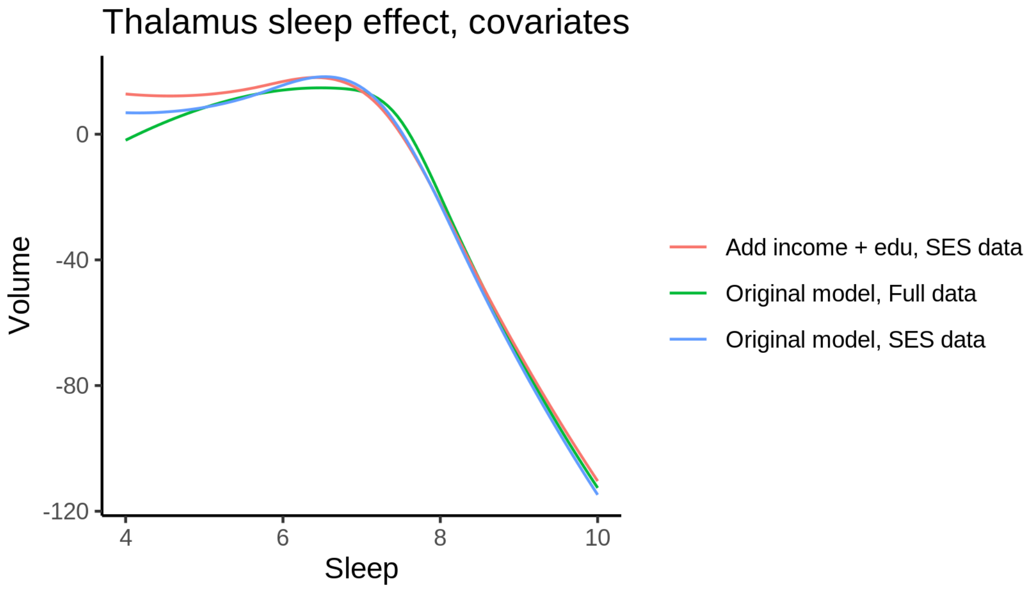


The plot below shows the sleep-volume curve for the original model and for the model with main effects of BMI.

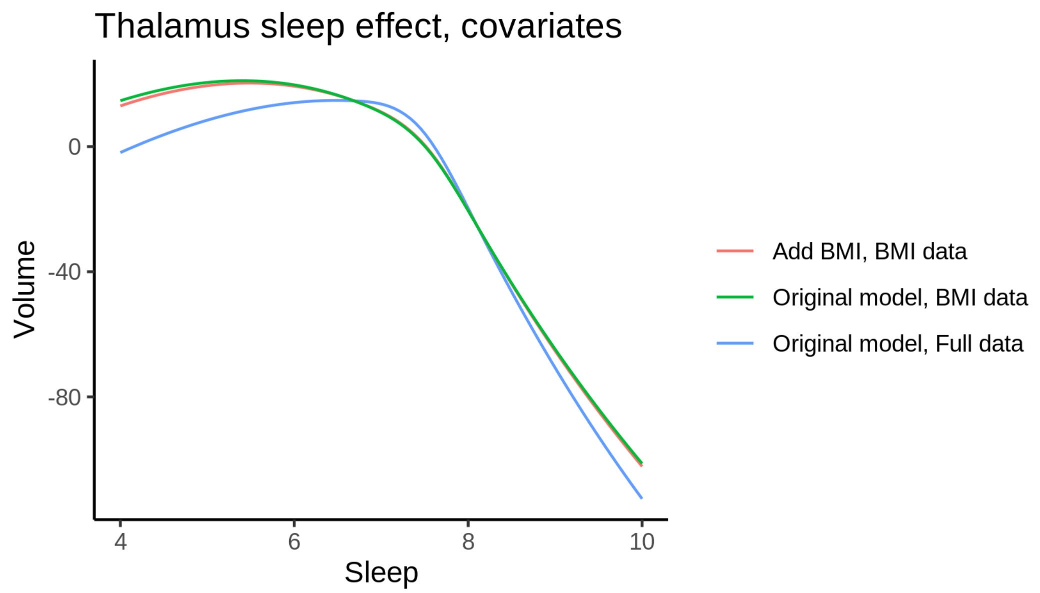


The plot below shows the sleep-volume curve for the original model and for the model with main effects of depression.

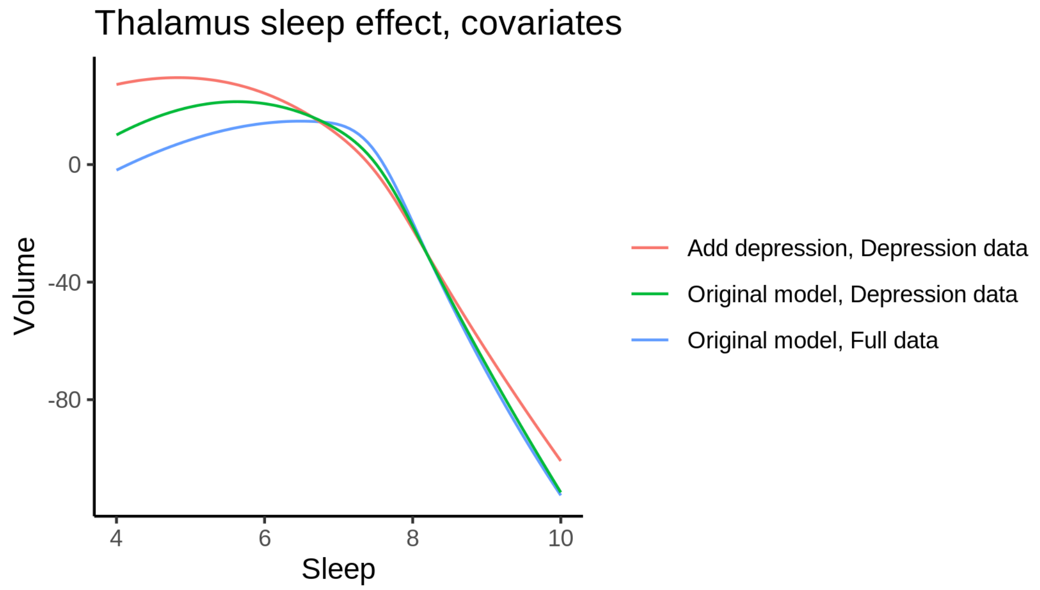


### TotalGrayVol

#### Descriptive statistics

| Study | Observations | Unique IDs | Mean age | Age range |
|---|---|---|---|---|
| HCP | 974 | 974 | 28.8 | 22 - 37 |
| MPIB | 677 | 391 | 63.1 | 24 - 83 |
| UB | 113 | 39 | 70.9 | 64 - 81 |
| UCAM | 884 | 632 | 55.1 | 20 - 88 |

| Study | Observations | Unique IDs | Mean age | Age range |
|---|---|---|---|---|
| UiO | 1475 | 803 | 49.4 | 20 - 89 |
| UKB | 45977 | 43134 | 64.5 | 45 - 83 |
| UmU | 423 | 284 | 62.3 | 25 - 85 |
| UOXF | 769 | 769 | 69.8 | 60 - 85 |

##### Spaghetti plot

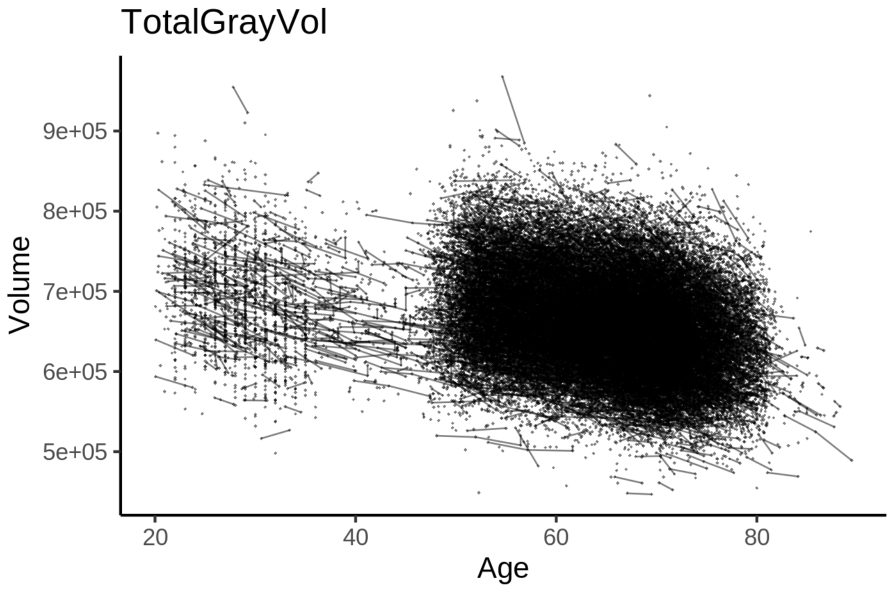


##### Model outputs

Model without sleep term

```
##
## Family: gaussian
## Link function: identity
##
## Formula:
## value ~ sex + site + icv + s(age_z, k = 10, bs = "cr")
## <environment: 0x55cf3a5f66f8>
##
## Parametric coefficients:
##                Estimate Std. Error t value Pr(>|t|)
## (Intercept)    606904.9     1672.3 362.922  < 2e-16 ***
## sexmale         15975.2      334.8  47.718  < 2e-16 ***
## siteMPIB        56062.0     2101.2  26.680  < 2e-16 ***
## siteousAvanto   -5008.2     1745.2  -2.870  0.00411 **
## siteousPrisma   31161.0     2924.7  10.654  < 2e-16 ***
## siteousSkyra    20115.4     1606.1  12.524  < 2e-16 ***
## siteUB          13348.1     4862.9   2.745  0.00606 **
## siteUCAM        14852.7     1828.0   8.125 4.57e-16 ***
## siteUKB         51638.1     1716.2  30.089  < 2e-16 ***
## siteUmU         25127.9     2345.6  10.713  < 2e-16 ***
```

```
## siteUOXF       10093.5     2019.4   4.998 5.80e-07 ***
## icv            45480.7      168.5 269.886  < 2e-16 ***
## ---
## Signif. codes:  0 '***' 0.001 '**' 0.01 '*' 0.05 '.' 0.1 ' ' 1
##
## Approximate significance of smooth terms:
##            edf Ref.df    F p-value
## s(age_z) 7.935  7.935 1799  <2e-16 ***
## ---
## Signif. codes:  0 '***' 0.001 '**' 0.01 '*' 0.05 '.' 0.1 ' ' 1
##
## R-sq.(adj) =  0.766
## lmer.REML = 1.1952e+06  Scale est. = 1.0487e+08  n = 51292
```

Model with only main effects of age and sleep

```
##
## Family: gaussian
## Link function: identity
##
## Formula:
## value ~ sex + site + icv + s(age_z, k = 10, bs = "cr") + s(sleep_z,
##     k = 5, bs = "cr")
## <environment: 0x55cf3a5f66f8>
##
## Parametric coefficients:
##               Estimate Std. Error t value Pr(>|t|)
## (Intercept)   607105.1     1672.6 362.962  < 2e-16 ***
## sexmale        15964.2      334.5  47.727  < 2e-16 ***
## siteMPIB       56144.5     2100.7  26.727  < 2e-16 ***
## siteousAvanto  -5217.4     1744.4  -2.991  0.00278 **
## siteousPrisma  31029.6     2922.3  10.618  < 2e-16 ***
## siteousSkyra   19865.6     1605.3  12.375  < 2e-16 ***
## siteUB         13643.5     4857.8   2.809  0.00498 **
## siteUCAM       14921.0     1826.7   8.168 3.20e-16 ***
## siteUKB        51431.1     1716.7  29.960  < 2e-16 ***
## siteUmU        25345.1     2349.0  10.790  < 2e-16 ***
## siteUOXF        9990.0     2017.9   4.951 7.41e-07 ***
## icv            45425.4      168.6 269.505  < 2e-16 ***
## ---
## Signif. codes:  0 '***' 0.001 '**' 0.01 '*' 0.05 '.' 0.1 ' ' 1
##
## Approximate significance of smooth terms:
##              edf Ref.df       F p-value
## s(age_z)   7.931  7.931 1777.75  <2e-16 ***
## s(sleep_z) 3.673  3.673   24.24  <2e-16 ***
## ---
## Signif. codes:  0 '***' 0.001 '**' 0.01 '*' 0.05 '.' 0.1 ' ' 1
##
## R-sq.(adj) =  0.767
## lmer.REML = 1.1951e+06  Scale est. = 1.049e+08  n = 51292
```

Model with full interaction between age and sleep

```
##
## Family: gaussian
```

```
## Link function: identity
##
## Formula:
## value ~ sex + site + icv + t2(age_z, sleep_z, k = c(10, 4), bs = "cr")
## <environment: 0x55cf3a5f66f8>
##
## Parametric coefficients:
##                Estimate Std. Error t value Pr(>|t|)
## (Intercept)    607516.4     1680.2 361.577  < 2e-16 ***
## sexmale         15991.1      335.2  47.700  < 2e-16 ***
## siteMPIB        55623.9     2113.5  26.319  < 2e-16 ***
## siteousAvanto   -5589.9     1755.0  -3.185  0.00145 **
## siteousPrisma   30618.6     2929.5  10.452  < 2e-16 ***
## siteousSkyra    19509.1     1617.4  12.062  < 2e-16 ***
## siteUB          12915.9     4861.9   2.657  0.00790 **
## siteUCAM        14352.9     1838.8   7.806 6.03e-15 ***
## siteUKB         51003.1     1724.2  29.580  < 2e-16 ***
## siteUmU         24892.2     2358.4  10.555  < 2e-16 ***
## siteUOXF         9408.4     2026.6   4.642 3.45e-06 ***
## icv             45428.8      168.6 269.523  < 2e-16 ***
## ---
## Signif. codes:  0 '***' 0.001 '**' 0.01 '*' 0.05 '.' 0.1 ' ' 1
##
## Approximate significance of smooth terms:
##                      edf Ref.df     F p-value
## t2(age_z,sleep_z) 17.55  17.55 24.68  <2e-16 ***
## ---
## Signif. codes:  0 '***' 0.001 '**' 0.01 '*' 0.05 '.' 0.1 ' ' 1
##
## R-sq.(adj) =  0.767
## lmer.REML = 1.1951e+06  Scale est. = 1.049e+08  n = 51292
```

#### Model comparison

`mod_no_sleep` refers to model without sleep term, `mod_no_interaction` refers to model with only main effect of sleep, and `mod_full` refers to model with a full interaction between age and sleep. This is a nested model comparison, and the p-value at a given line refers to comparing the model at the line to the model on the line above. Hence, significance implies that the more complicated model is supported on statistical grounds.

To be even more specific, the p-value on the second row tests whether there is an association between sleep and volume. The p-value on the third row tests whether this association depends on age.

```
## Data: NULL
## Models:
## mod_list$mod_no_sleep$mer: NULL
## mod_list$mod_no_interaction$mer: NULL
## mod_list$mod_full$mer: NULL
##                                 npar     AIC     BIC  logLik deviance  Chisq Df Pr(>Chisq)
## mod_list$mod_no_sleep$mer         16 1195222 1195364 -597595  1195190
## mod_list$mod_no_interaction$mer   18 1195144 1195304 -597554  1195108 81.651  2     <2e-16 ***
## mod_list$mod_full$mer             20 1195163 1195340 -597562  1195123  0.000  2          1
## ---
## Signif. codes:  0 '***' 0.001 '**' 0.01 '*' 0.05 '.' 0.1 ' ' 1
```

We chose the model based on the likelihood ratio test with 5 % significance level, which was `mod_no_interaction`.

#### Lifespan brain trajectory

The trajectory shown is from the chosen model `mod_no_interaction`.

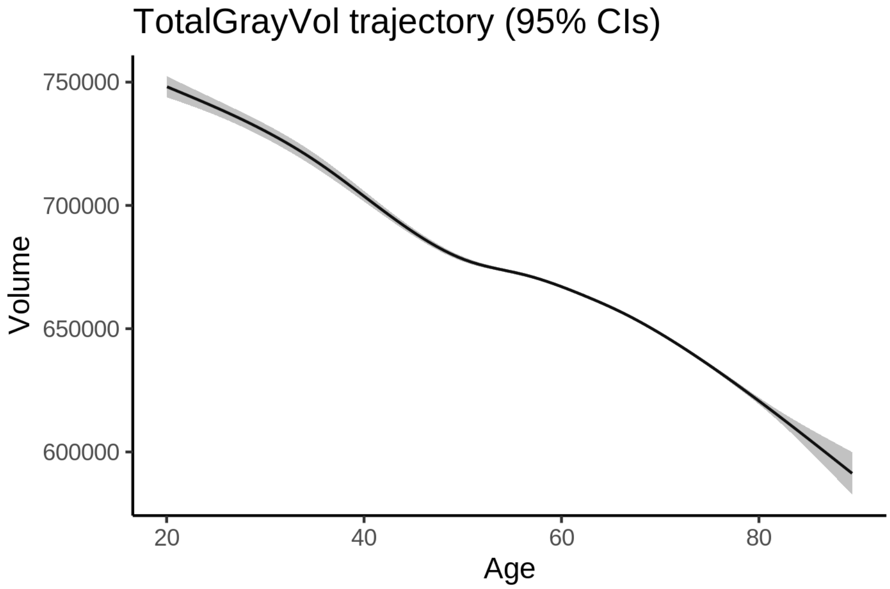


#### Effect of sleep

The chosen model only included the main effect of sleep, and hence the effect does not vary with age. The black dot shows the average sleep duration across all ages in the sample.

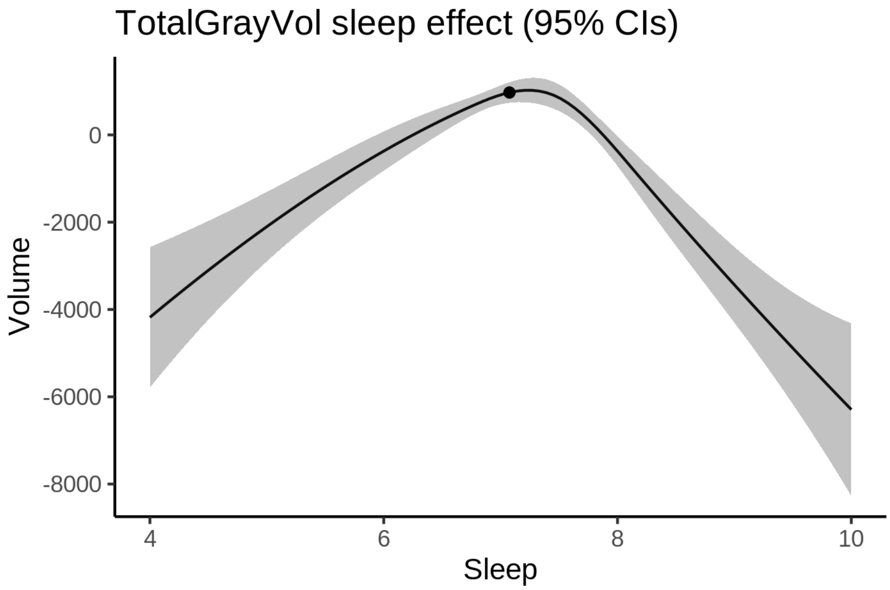


We also show the full interaction model for completeness, although it was not selected.

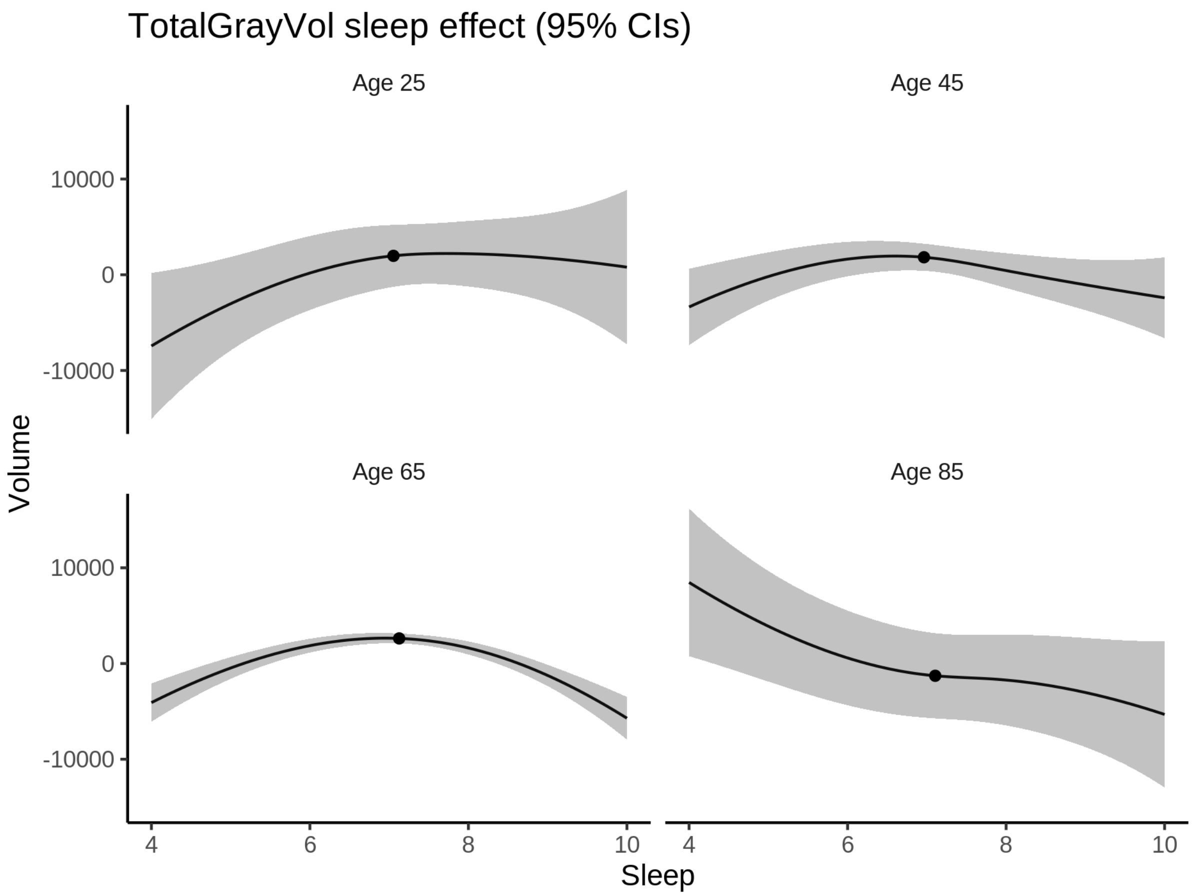


**Deviation from sleep associated with maximal volume**

Model with only main effect of sleep was chosen, so we show it for all ages at once. Maximum volume is attained at 7.2 hours of sleep. The percentage values in the plot are calculated as follows: The maximum at 100 % refers to a person at an arbitrary age with a sleep duration associated with maximum volume. For a female, this volume is 673977 and for a male it is 689942. The other percentage values show how large the expected volume is for someone with other sleep durations. For example, 99 % implies a 1 % reduction.

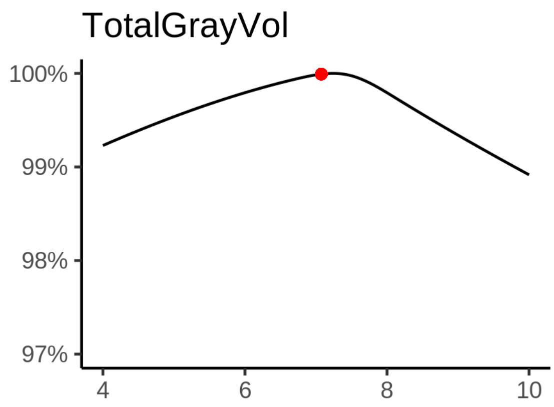

**Comparison of mean sleep and sleep associated with maximum volume**

A 95 % confidence interval for the sleep associated with maximum volume is [7.09, 7.33].

The plot below compares average sleep to the sleep associated with maximum volume.

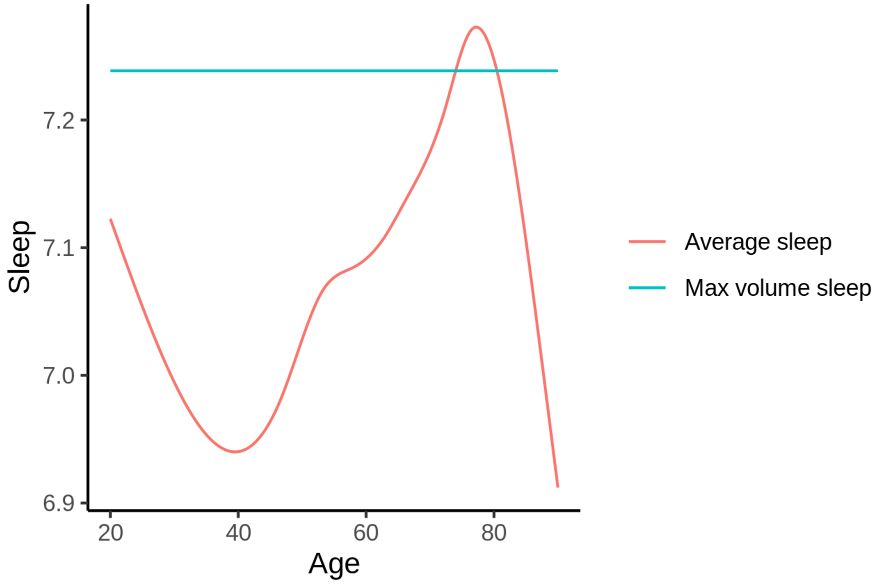


The next plot shows the difference between average sleep and sleep associated with maximum volume. The shaded region is a 95 % confidence interval.

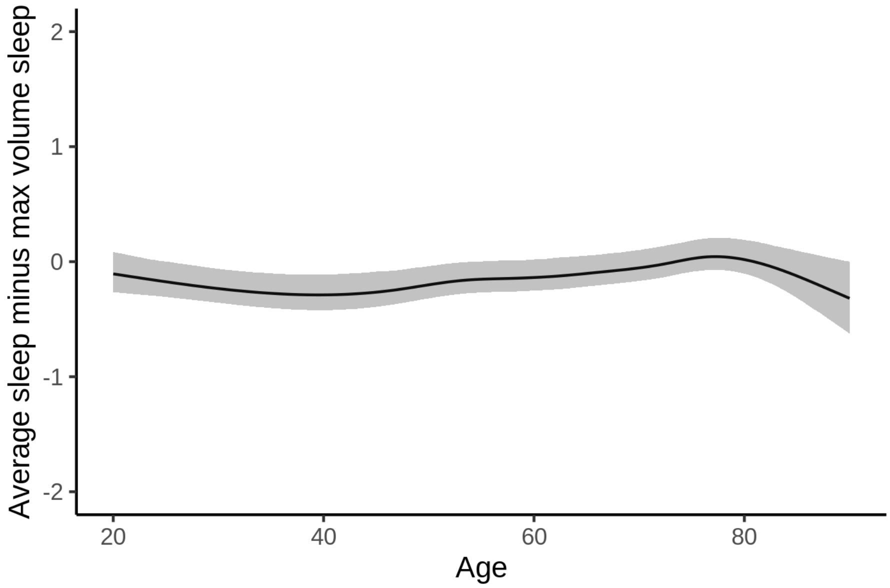


The next plot shows the probability that the sleep duration associated with maximum volume is longer than the average sleep duration, as a function of age. Probility below .05 can be interpreted as evidence that the sleep associated with maximum volume is shorter than the mean sleep, and probability above .95 can be interpreted the opposite way.

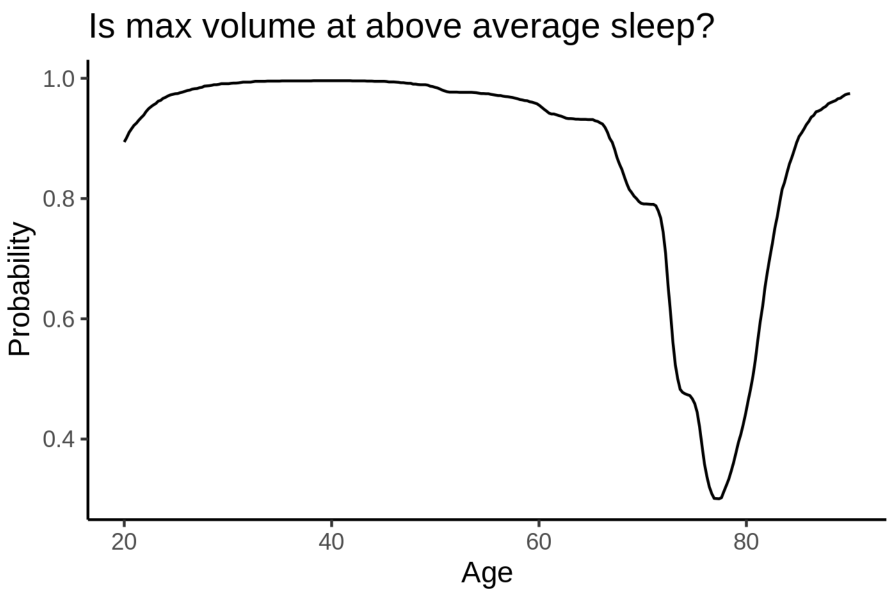


##### Controling for covariates

Below is the output for a model in which we only include data with income and education.

```
##
## Family: gaussian
## Link function: identity
##
## Formula:
## value ~ sex + site + s(age_z, k = 10, bs = "cr") + s(sleep_z,
##     k = 5, bs = "cr") + icv
## <environment: 0x55cffc024d68>
##
## Parametric coefficients:
##                Estimate Std. Error t value Pr(>|t|)
## (Intercept)    675222.2     3798.7 177.751  < 2e-16 ***
## sexmale         15652.0      419.1  37.350  < 2e-16 ***
## siteousAvanto  -80433.2     4797.6 -16.765  < 2e-16 ***
## siteousPrisma  -32342.0     4693.8  -6.890 5.67e-12 ***
## siteousSkyra   -49323.3     4098.2 -12.035  < 2e-16 ***
## siteUKB        -17877.4     3793.0  -4.713 2.45e-06 ***
## siteUOXF       -50294.6     4272.6 -11.771  < 2e-16 ***
## icv             46477.7      215.6 215.572  < 2e-16 ***
## ---
## Signif. codes:  0 ‘***’ 0.001 ‘**’ 0.01 ‘*’ 0.05 ‘.’ 0.1 ‘ ’ 1
##
## Approximate significance of smooth terms:
##              edf Ref.df       F p-value
## s(age_z)   6.812  6.812 1046.94  <2e-16 ***
## s(sleep_z) 3.511  3.511   15.32  <2e-16 ***
## ---
## Signif. codes:  0 ‘***’ 0.001 ‘**’ 0.01 ‘*’ 0.05 ‘.’ 0.1 ‘ ’ 1
##
```

```
## R-sq.(adj) =  0.763
## lmer.REML = 7.2597e+05  Scale est. = 1.2113e+08  n = 31192
```

Below is the output for a model in which we control for the main effects of income and education.

```
##
## Family: gaussian
## Link function: identity
##
## Formula:
## value ~ sex + site + s(age_z, k = 10, bs = "cr") + s(sleep_z,
##     k = 5, bs = "cr") + icv + income_scaled + education_scaled
## <environment: 0x55cffc024d68>
##
## Parametric coefficients:
##                  Estimate Std. Error t value Pr(>|t|)
## (Intercept)      672500.4     3821.9 175.958  < 2e-16 ***
## sexmale           15615.2      420.0  37.179  < 2e-16 ***
## siteousAvanto    -80362.9     4796.3 -16.755  < 2e-16 ***
## siteousPrisma    -32672.6     4690.9  -6.965 3.35e-12 ***
## siteousSkyra     -49284.0     4096.4 -12.031  < 2e-16 ***
## siteUKB          -17634.0     3793.2  -4.649 3.35e-06 ***
## siteUOXF         -49321.5     4272.6 -11.544  < 2e-16 ***
## icv               46302.5      217.3 213.104  < 2e-16 ***
## income_scaled      2316.7      499.9   4.634 3.60e-06 ***
## education_scaled   1803.3      589.4   3.059  0.00222 **
## ---
## Signif. codes:  0 ‘***’ 0.001 ‘**’ 0.01 ‘*’ 0.05 ‘.’ 0.1 ‘ ’ 1
##
## Approximate significance of smooth terms:
##              edf Ref.df      F p-value
## s(age_z)   6.835  6.835 940.52  <2e-16 ***
## s(sleep_z) 3.440  3.440  13.04  <2e-16 ***
## ---
## Signif. codes:  0 ‘***’ 0.001 ‘**’ 0.01 ‘*’ 0.05 ‘.’ 0.1 ‘ ’ 1
##
## R-sq.(adj) =  0.763
## lmer.REML = 7.259e+05  Scale est. = 1.212e+08  n = 31192
```

We also included interaction effects between sleep duration and education and income, in another model. The output is shown below, and the interaction terms are `income_scaled:sleep_z` and `education_scaled:sleep_z`.

```
##
## Family: gaussian
## Link function: identity
##
## Formula:
## value ~ sex + site + s(age_z, k = 10, bs = "cr") + s(sleep_z,
##     k = 5, bs = "cr") + icv + income_scaled + education_scaled +
##     income_scaled:sleep_z + education_scaled:sleep_z
## <environment: 0x55cffc024d68>
##
## Parametric coefficients:
##                          Estimate Std. Error t value Pr(>|t|)
## (Intercept)              672525.8     3822.0 175.963  < 2e-16 ***
```

```
## sexmale                   15577.5      420.3  37.065  < 2e-16 ***
## siteousAvanto            -80433.9     4796.2 -16.770  < 2e-16 ***
## siteousPrisma            -32842.8     4691.2  -7.001 2.59e-12 ***
## siteousSkyra             -49357.9     4096.2 -12.050  < 2e-16 ***
## siteUKB                  -17662.3     3793.4  -4.656 3.24e-06 ***
## siteUOXF                 -49341.6     4273.6 -11.546  < 2e-16 ***
## icv                       46307.6      217.3 213.123  < 2e-16 ***
## income_scaled              2353.3      500.2   4.705 2.55e-06 ***
## education_scaled           1778.9      589.5   3.018  0.00255 **
## income_scaled:sleep_z     -1027.0      498.8  -2.059  0.03949 *
## education_scaled:sleep_z   1006.5      572.4   1.758  0.07870 .
## ---
## Signif. codes:  0 '***' 0.001 '**' 0.01 '*' 0.05 '.' 0.1 ' ' 1
##
## Approximate significance of smooth terms:
##              edf Ref.df      F p-value
## s(age_z)   6.837  6.837 939.60  <2e-16 ***
## s(sleep_z) 3.438  3.438  12.75  <2e-16 ***
## ---
## Signif. codes:  0 '***' 0.001 '**' 0.01 '*' 0.05 '.' 0.1 ' ' 1
##
## R-sq.(adj) =  0.763
## lmer.REML = 7.2587e+05  Scale est. = 1.212e+08  n = 31192
```

We did the same controling for BMI. Below is the model with no covariates but only keeping data with BMI.

```
##
## Family: gaussian
## Link function: identity
##
## Formula:
## value ~ sex + site + s(age_z, k = 10, bs = "cr") + s(sleep_z,
##     k = 5, bs = "cr") + icv
## <environment: 0x55cf415425d0>
##
## Parametric coefficients:
##               Estimate Std. Error t value Pr(>|t|)
## (Intercept)   599356.3     3023.1 198.258  < 2e-16 ***
## sexmale        15038.4      402.3  37.384  < 2e-16 ***
## siteousPrisma  40722.1     3853.5  10.568  < 2e-16 ***
## siteousSkyra   30061.1     2600.7  11.559  < 2e-16 ***
## siteUCAM       23364.4     3137.3   7.447 9.76e-14 ***
## siteUKB        58816.1     3042.4  19.332  < 2e-16 ***
## siteUmU        34980.3     3440.7  10.167  < 2e-16 ***
## icv            46962.7      207.6 226.210  < 2e-16 ***
## ---
## Signif. codes:  0 '***' 0.001 '**' 0.01 '*' 0.05 '.' 0.1 ' ' 1
##
## Approximate significance of smooth terms:
##              edf Ref.df       F p-value
## s(age_z)   7.496  7.496 1222.88  <2e-16 ***
## s(sleep_z) 3.452  3.452   13.17  <2e-16 ***
## ---
## Signif. codes:  0 '***' 0.001 '**' 0.01 '*' 0.05 '.' 0.1 ' ' 1
##
```

```
## R-sq.(adj) =  0.772
## lmer.REML = 7.7733e+05  Scale est. = 1.1321e+08  n = 33447
```

Below is the model output with main effect.

```
##
## Family: gaussian
## Link function: identity
##
## Formula:
## value ~ sex + site + s(age_z, k = 10, bs = "cr") + s(sleep_z,
##     k = 5, bs = "cr") + icv + bmi
## <environment: 0x55cf415425d0>
##
## Parametric coefficients:
##                 Estimate Std. Error t value Pr(>|t|)
## (Intercept)    611000.29    3176.46 192.353  < 2e-16 ***
## sexmale         15404.24     402.57  38.265  < 2e-16 ***
## siteousPrisma   41271.46    3848.60  10.724  < 2e-16 ***
## siteousSkyra    30177.10    2599.45  11.609  < 2e-16 ***
## siteUCAM        23671.95    3133.81   7.554 4.34e-14 ***
## siteUKB         59214.27    3039.21  19.483  < 2e-16 ***
## siteUmU         35513.62    3436.36  10.335  < 2e-16 ***
## icv             47036.96     207.24 226.971  < 2e-16 ***
## bmi              -463.24      39.15 -11.832  < 2e-16 ***
## ---
## Signif. codes:  0 ‘***’ 0.001 ‘**’ 0.01 ‘*’ 0.05 ‘.’ 0.1 ‘ ’ 1
##
## Approximate significance of smooth terms:
##              edf Ref.df       F p-value
## s(age_z)   7.495  7.495 1227.91  <2e-16 ***
## s(sleep_z) 3.348  3.348   11.03  <2e-16 ***
## ---
## Signif. codes:  0 ‘***’ 0.001 ‘**’ 0.01 ‘*’ 0.05 ‘.’ 0.1 ‘ ’ 1
##
## R-sq.(adj) =  0.773
## lmer.REML = 7.7718e+05  Scale est. = 1.1315e+08  n = 33447
```

Next is the model with BMI-sleep interaction.

```
##
## Family: gaussian
## Link function: identity
##
## Formula:
## value ~ sex + site + s(age_z, k = 10, bs = "cr") + s(sleep_z,
##     k = 5, bs = "cr") + icv + bmi + bmi:sleep_z
## <environment: 0x55cf415425d0>
##
## Parametric coefficients:
##                 Estimate Std. Error t value Pr(>|t|)
## (Intercept)    610963.92    3177.79 192.261  < 2e-16 ***
## sexmale         15404.81     402.58  38.266  < 2e-16 ***
## siteousPrisma   41273.74    3848.64  10.724  < 2e-16 ***
## siteousSkyra    30182.38    2599.50  11.611  < 2e-16 ***
## siteUCAM        23686.17    3134.04   7.558 4.21e-14 ***
```

```
## siteUKB        59229.13    3039.45  19.487  < 2e-16 ***
## siteUmU        35526.21    3436.54  10.338  < 2e-16 ***
## icv            47037.65     207.25 226.965  < 2e-16 ***
## bmi             -462.70      39.18 -11.811  < 2e-16 ***
## bmi:sleep_z       14.89      37.22   0.400    0.689
## ---
## Signif. codes:  0 ‘***’ 0.001 ‘**’ 0.01 ‘*’ 0.05 ‘.’ 0.1 ‘ ’ 1
##
## Approximate significance of smooth terms:
##              edf Ref.df        F  p-value
## s(age_z)   7.495  7.495 1227.412  < 2e-16 ***
## s(sleep_z) 3.340  3.340    9.383 2.05e-06 ***
## ---
## Signif. codes:  0 ‘***’ 0.001 ‘**’ 0.01 ‘*’ 0.05 ‘.’ 0.1 ‘ ’ 1
##
## R-sq.(adj) =  0.773
## lmer.REML = 7.7718e+05  Scale est. = 1.1315e+08  n = 33447
```

We did the same controling for depression. Below is the model with no covariates but only keeping data with depression.

```
##
## Family: gaussian
## Link function: identity
##
## Formula:
## value ~ sex + site + s(age_z, k = 10, bs = "cr") + s(sleep_z,
##     k = 5, bs = "cr") + icv
## <environment: 0x55cf3f9d29f0>
##
## Parametric coefficients:
##               Estimate Std. Error t value Pr(>|t|)
## (Intercept)   656906.3     1903.8 345.054  < 2e-16 ***
## sexmale        15204.8      406.5  37.408  < 2e-16 ***
## siteousAvanto -48905.1     4695.3 -10.416  < 2e-16 ***
## siteousPrisma -13980.9     3767.9  -3.711 0.000207 ***
## siteousSkyra  -23131.1     2923.7  -7.912 2.62e-15 ***
## siteUCAM      -34194.1     2621.2 -13.045  < 2e-16 ***
## siteUKB          748.2     1882.2   0.397 0.691005
## siteUmU       -22989.7     2560.8  -8.978  < 2e-16 ***
## icv            46674.0      209.4 222.865  < 2e-16 ***
## ---
## Signif. codes:  0 ‘***’ 0.001 ‘**’ 0.01 ‘*’ 0.05 ‘.’ 0.1 ‘ ’ 1
##
## Approximate significance of smooth terms:
##              edf Ref.df       F p-value
## s(age_z)   7.504  7.504 1057.92  <2e-16 ***
## s(sleep_z) 3.492  3.492   15.14  <2e-16 ***
## ---
## Signif. codes:  0 ‘***’ 0.001 ‘**’ 0.01 ‘*’ 0.05 ‘.’ 0.1 ‘ ’ 1
##
## R-sq.(adj) =  0.769
## lmer.REML = 7.754e+05  Scale est. = 1.0814e+08  n = 33359
```

Below is the model output with main effect.

```
##
## Family: gaussian
## Link function: identity
##
## Formula:
## value ~ sex + site + s(age_z, k = 10, bs = "cr") + s(sleep_z,
##     k = 5, bs = "cr") + icv + depression
## <environment: 0x55cf3f9d29f0>
##
## Parametric coefficients:
##               Estimate Std. Error t value Pr(>|t|)
## (Intercept)   657842.6     1911.4 344.172  < 2e-16 ***
## sexmale        15108.4      406.7  37.149  < 2e-16 ***
## siteousAvanto -48423.6     4694.8 -10.314  < 2e-16 ***
## siteousPrisma -13559.7     3767.2  -3.599 0.000319 ***
## siteousSkyra  -22583.1     2924.3  -7.722 1.17e-14 ***
## siteUCAM      -33980.5     2620.3 -12.968  < 2e-16 ***
## siteUKB          361.0     1882.9   0.192 0.847943
## siteUmU       -20164.5     2615.8  -7.709 1.31e-14 ***
## icv            46688.5      209.4 223.009  < 2e-16 ***
## depression     -7629.3     1455.0  -5.244 1.58e-07 ***
## ---
## Signif. codes:  0 '***' 0.001 '**' 0.01 '*' 0.05 '.' 0.1 ' ' 1
##
## Approximate significance of smooth terms:
##              edf Ref.df       F p-value
## s(age_z)   7.500  7.500 1057.83  <2e-16 ***
## s(sleep_z) 3.421  3.421   12.87  <2e-16 ***
## ---
## Signif. codes:  0 '***' 0.001 '**' 0.01 '*' 0.05 '.' 0.1 ' ' 1
##
## R-sq.(adj) =  0.769
## lmer.REML = 7.7536e+05  Scale est. = 1.081e+08  n = 33359
```

Next is the model with depression-sleep interaction.

```
##
## Family: gaussian
## Link function: identity
##
## Formula:
## value ~ sex + site + s(age_z, k = 10, bs = "cr") + s(sleep_z,
##     k = 5, bs = "cr") + icv + depression + depression:sleep_z
## <environment: 0x55cf3f9d29f0>
##
## Parametric coefficients:
##                    Estimate Std. Error t value Pr(>|t|)
## (Intercept)        657843.8     1911.4 344.170  < 2e-16 ***
## sexmale             15103.8      406.8  37.131  < 2e-16 ***
## siteousAvanto      -48456.0     4695.0 -10.321  < 2e-16 ***
## siteousPrisma      -13594.0     3767.6  -3.608 0.000309 ***
## siteousSkyra       -22614.9     2924.8  -7.732 1.09e-14 ***
## siteUCAM           -34012.0     2620.8 -12.978  < 2e-16 ***
## siteUKB               361.8     1882.9   0.192 0.847631
## siteUmU            -19938.0     2639.9  -7.553 4.38e-14 ***
```

```
## icv                   46688.8      209.4 223.007  < 2e-16 ***
## depression           -7690.4     1458.4  -5.273 1.35e-07 ***
## depression:sleep_z    -690.1     1082.0  -0.638 0.523591
## ---
## Signif. codes:  0 '***' 0.001 '**' 0.01 '*' 0.05 '.' 0.1 ' ' 1
##
## Approximate significance of smooth terms:
##              edf Ref.df       F p-value
## s(age_z)   7.497  7.497 1058.13  <2e-16 ***
## s(sleep_z) 3.426  3.426   11.93  <2e-16 ***
## ---
## Signif. codes:  0 '***' 0.001 '**' 0.01 '*' 0.05 '.' 0.1 ' ' 1
##
## R-sq.(adj) =  0.769
## lmer.REML = 7.7534e+05  Scale est. = 1.081e+08  n = 33359
```

The plot below shows the sleep-volume curve for the original model and for the model with main effects of SES.

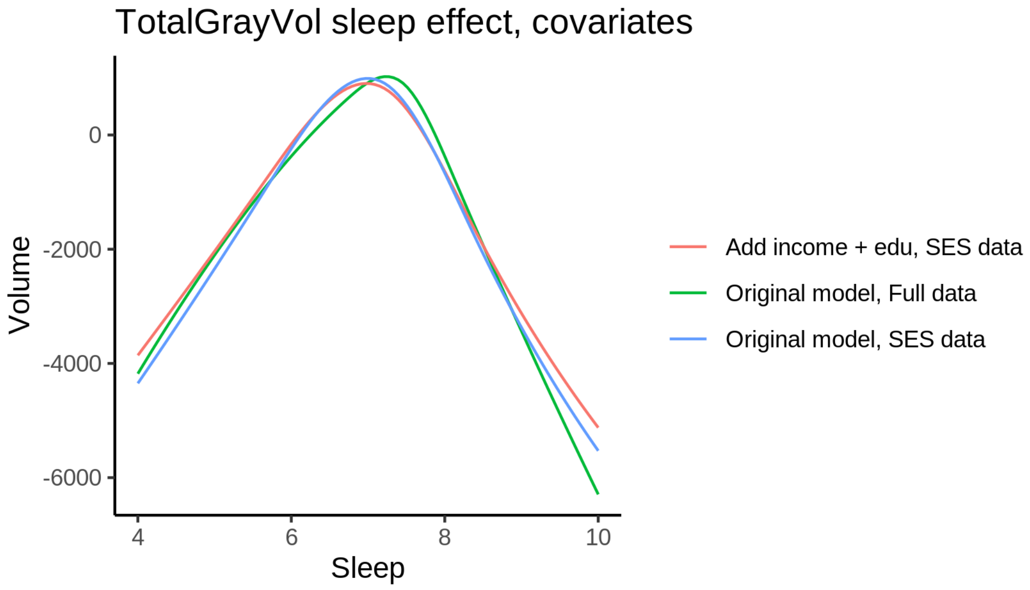


The plot below shows the sleep-volume curve for the original model and for the model with main effects of BMI.

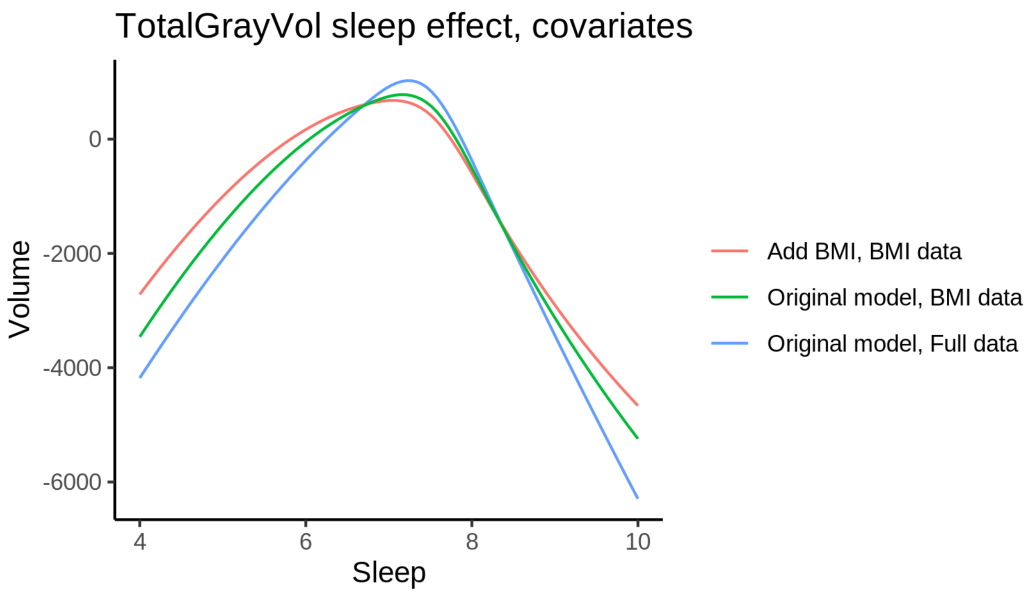


The plot below shows the sleep-volume curve for the original model and for the model with main effects of depression.

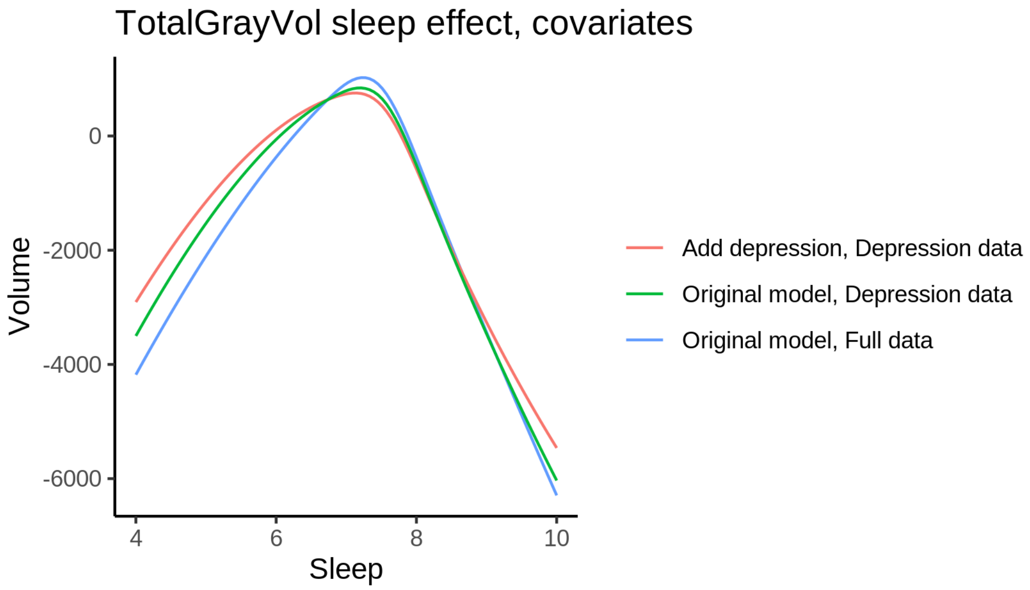


### Ventricles

#### Descriptive statistics

| Study | Observations | Unique IDs | Mean age | Age range |
|---|---|---|---|---|
| HCP | 974 | 974 | 28.8 | 22 - 37 |
| MPIB | 677 | 391 | 63.1 | 24 - 83 |
| UB | 113 | 39 | 70.9 | 64 - 81 |
| UCAM | 884 | 632 | 55.1 | 20 - 88 |

| Study | Observations | Unique IDs | Mean age | Age range |
|---|---|---|---|---|
| UiO | 1460 | 803 | 49.2 | 20 - 89 |
| UKB | 45979 | 43135 | 64.5 | 45 - 83 |
| UmU | 423 | 284 | 62.3 | 25 - 85 |
| UOXF | 769 | 769 | 69.8 | 60 - 85 |

##### Spaghetti plot

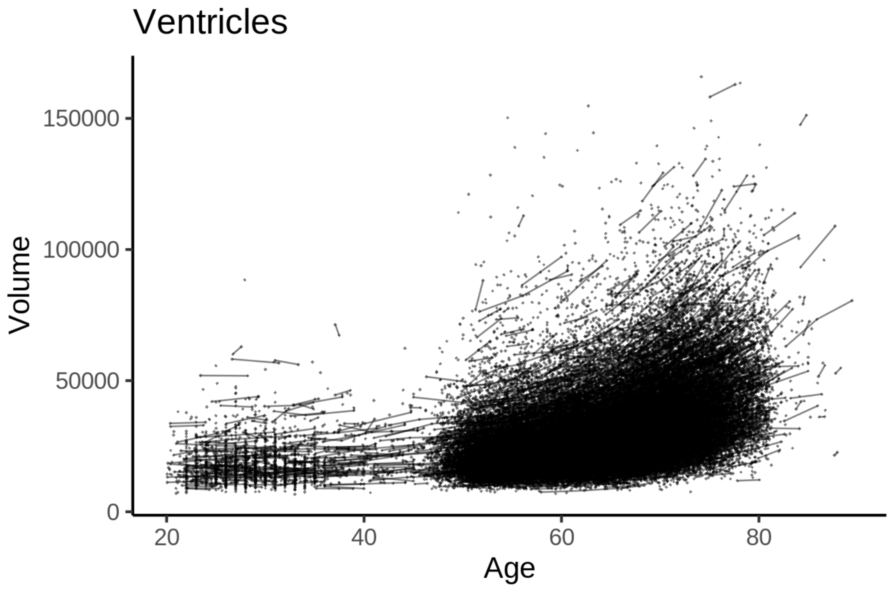


##### Model outputs

Model without sleep term

```
##
## Family: gaussian
## Link function: identity
##
## Formula:
## value ~ sex + site + icv + s(age_z, k = 10, bs = "cr")
## <environment: 0x55cf373050e8>
##
## Parametric coefficients:
##                Estimate Std. Error t value Pr(>|t|)
## (Intercept)    31260.76     600.18  52.085  < 2e-16 ***
## sexmale         1874.58     141.50  13.248  < 2e-16 ***
## siteMPIB       11322.50     846.02  13.383  < 2e-16 ***
## siteousAvanto    822.82     647.71   1.270   0.2040
## siteousPrisma    350.86     970.81   0.361   0.7178
## siteousSkyra    4338.36     644.65   6.730 1.72e-11 ***
## siteUB         -1117.14    2099.51  -0.532   0.5947
## siteUCAM         116.57     724.48   0.161   0.8722
## siteUKB         -590.99     612.01  -0.966   0.3342
## siteUmU         5297.60     941.77   5.625 1.86e-08 ***
```

```
## siteUOXF       1787.82     763.44   2.342   0.0192 *
## icv            6220.00      69.93  88.948  < 2e-16 ***
## ---
## Signif. codes:  0 '***' 0.001 '**' 0.01 '*' 0.05 '.' 0.1 ' ' 1
##
## Approximate significance of smooth terms:
##             edf Ref.df    F p-value
## s(age_z) 8.368  8.368 2516  <2e-16 ***
## ---
## Signif. codes:  0 '***' 0.001 '**' 0.01 '*' 0.05 '.' 0.1 ' ' 1
##
## R-sq.(adj) =  0.392
## lmer.REML = 1.0981e+06  Scale est. = 2.067e+06  n = 51279
```

Model with only main effects of age and sleep

```
##
## Family: gaussian
## Link function: identity
##
## Formula:
## value ~ sex + site + icv + s(age_z, k = 10, bs = "cr") + s(sleep_z,
##     k = 5, bs = "cr")
## <environment: 0x55cf373050e8>
##
## Parametric coefficients:
##                Estimate Std. Error t value Pr(>|t|)
## (Intercept)   31289.51     600.70  52.088  < 2e-16 ***
## sexmale        1877.54     141.48  13.271  < 2e-16 ***
## siteMPIB      11246.69     846.31  13.289  < 2e-16 ***
## siteousAvanto   816.37     647.81   1.260   0.2076
## siteousPrisma   349.41     970.84   0.360   0.7189
## siteousSkyra   4335.93     644.78   6.725 1.78e-11 ***
## siteUB        -1151.01    2099.10  -0.548   0.5835
## siteUCAM         92.60     724.42   0.128   0.8983
## siteUKB        -622.32     612.58  -1.016   0.3097
## siteUmU        5121.63     943.61   5.428 5.73e-08 ***
## siteUOXF       1787.71     763.41   2.342   0.0192 *
## icv            6220.84      69.99  88.888  < 2e-16 ***
## ---
## Signif. codes:  0 '***' 0.001 '**' 0.01 '*' 0.05 '.' 0.1 ' ' 1
##
## Approximate significance of smooth terms:
##               edf Ref.df        F  p-value
## s(age_z)   8.368  8.368 2495.403  < 2e-16 ***
## s(sleep_z) 2.630  2.630    5.651 0.000876 ***
## ---
## Signif. codes:  0 '***' 0.001 '**' 0.01 '*' 0.05 '.' 0.1 ' ' 1
##
## R-sq.(adj) =  0.392
## lmer.REML = 1.0981e+06  Scale est. = 2.0679e+06  n = 51279
```

Model with full interaction between age and sleep

```
##
## Family: gaussian
```

```
## Link function: identity
##
## Formula:
## value ~ sex + site + icv + t2(age_z, sleep_z, k = c(10, 4), bs = "cr")
## <environment: 0x55cf373050e8>
##
## Parametric coefficients:
##                Estimate Std. Error t value Pr(>|t|)
## (Intercept)    31385.34     603.42  52.012  < 2e-16 ***
## sexmale         1893.41     141.73  13.359  < 2e-16 ***
## siteMPIB       11171.74     850.18  13.140  < 2e-16 ***
## siteousAvanto    772.86     651.63   1.186   0.2356
## siteousPrisma    273.58     972.92   0.281   0.7786
## siteousSkyra    4276.86     648.77   6.592 4.37e-11 ***
## siteUB         -1269.23    2100.41  -0.604   0.5457
## siteUCAM          44.34     728.14   0.061   0.9514
## siteUKB         -730.24     615.35  -1.187   0.2353
## siteUmU         5049.92     946.20   5.337 9.49e-08 ***
## siteUOXF        1668.83     766.47   2.177   0.0295 *
## icv             6219.99      69.96  88.905  < 2e-16 ***
## ---
## Signif. codes:  0 '***' 0.001 '**' 0.01 '*' 0.05 '.' 0.1 ' ' 1
##
## Approximate significance of smooth terms:
##                     edf Ref.df    F p-value
## t2(age_z,sleep_z) 15.68  15.68 45.6  <2e-16 ***
## ---
## Signif. codes:  0 '***' 0.001 '**' 0.01 '*' 0.05 '.' 0.1 ' ' 1
##
## R-sq.(adj) =  0.392
## lmer.REML = 1.0981e+06  Scale est. = 2.0575e+06  n = 51279
```

#### Model comparison

`mod_no_sleep` refers to model without sleep term, `mod_no_interaction` refers to model with only main effect of sleep, and `mod_full` refers to model with a full interaction between age and sleep. This is a nested model comparison, and the p-value at a given line refers to comparing the model at the line to the model on the line above. Hence, significance implies that the more complicated model is supported on statistical grounds.

To be even more specific, the p-value on the second row tests whether there is an association between sleep and volume. The p-value on the third row tests whether this association depends on age.

```
## Data: NULL
## Models:
## mod_list$mod_no_sleep$mer: NULL
## mod_list$mod_no_interaction$mer: NULL
## mod_list$mod_full$mer: NULL
##                                 npar     AIC     BIC  logLik deviance  Chisq Df Pr(>Chisq)
## mod_list$mod_no_sleep$mer         16 1098148 1098289 -549058  1098116
## mod_list$mod_no_interaction$mer   18 1098138 1098298 -549051  1098102 13.022  2   0.001487 **
## mod_list$mod_full$mer             20 1098129 1098306 -549044  1098089 13.505  2   0.001168 **
## ---
## Signif. codes:  0 '***' 0.001 '**' 0.01 '*' 0.05 '.' 0.1 ' ' 1
```

We chose the model based on the likelihood ratio test with 5 % significance level, which was `mod_full`.

#### Lifespan brain trajectory

The trajectory shown is from the chosen model `mod_full`.

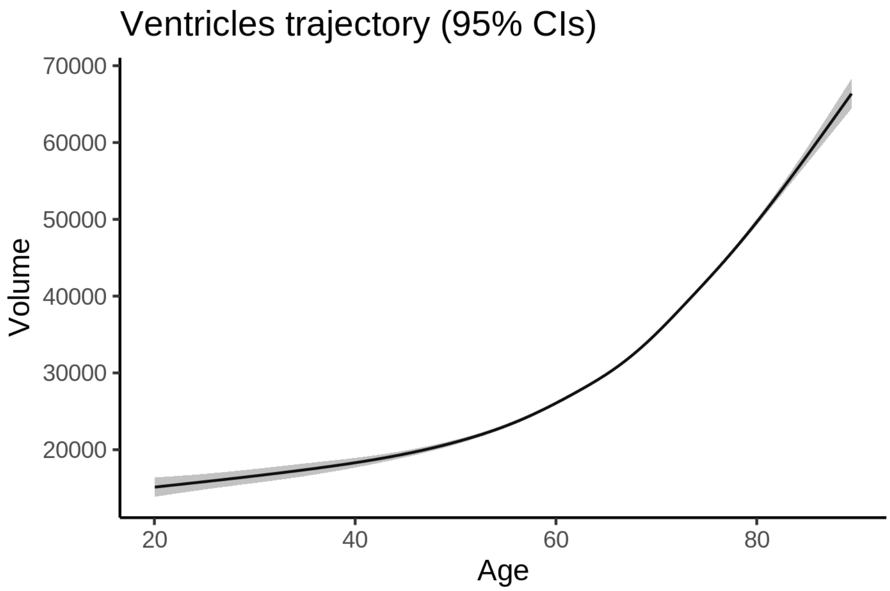


#### Effect of sleep

The chosen model included a full interaction between age and sleep, and the effect of sleep is hence plotted for a set of different ages. For comparability across ages, the sleep-volume curves at each age have been standardized so that they sum to zero.

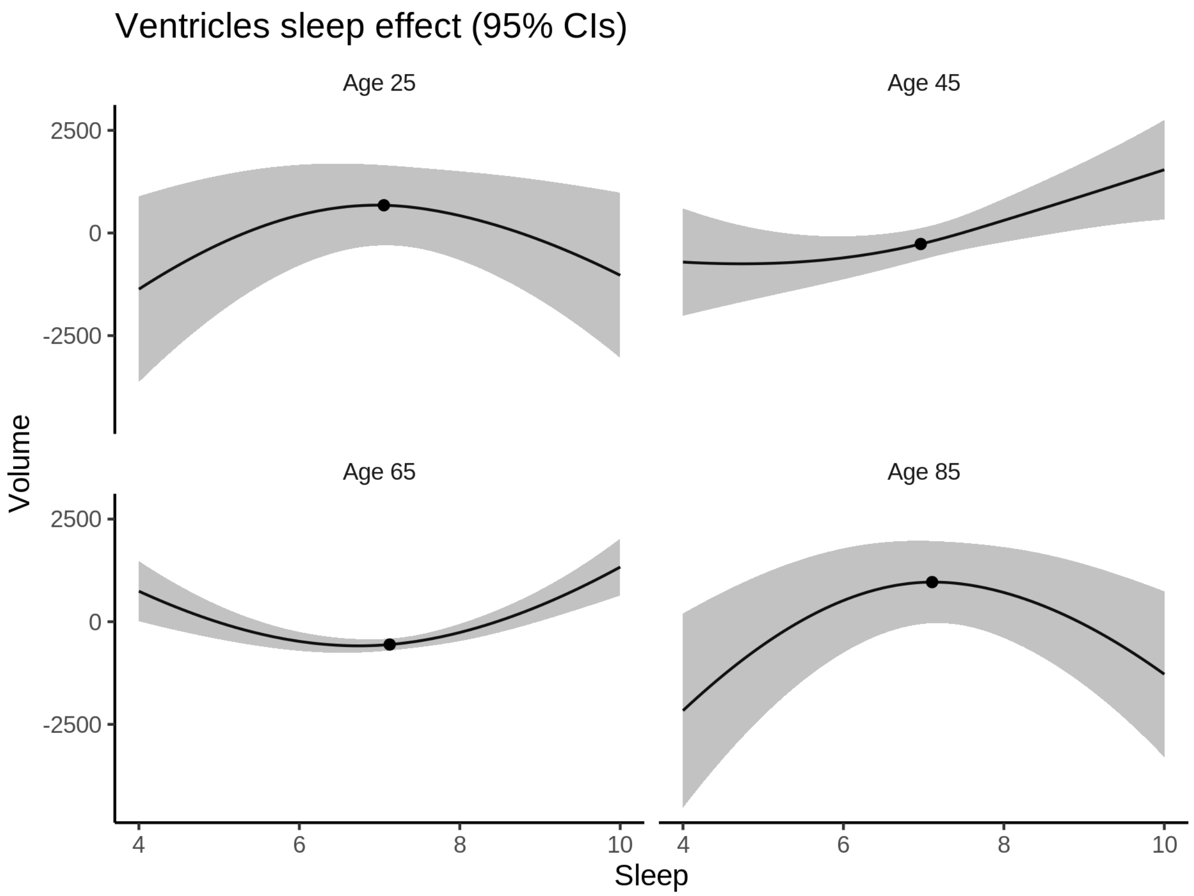


**Deviation from sleep associated with maximal volume**

The table show the sleep associated with maximum volume at chosen ages.

| Age | Sleep at max vol |
|---|---|
| 25 | 7.0 |
| 45 | 10.0 |
| 65 | 10.0 |
| 85 | 7.1 |

Model with sleep-age interaction was chosen, so we show it for four selected ages. The percentage values in the plot are calculated as follows: The maximum at 100 % refers to a person at the given age with a sleep duration associated with maximum volume. The other percentage values show how large the expected volume is for someone with other sleep durations. For example, 99 % implies a 1 % reduction.

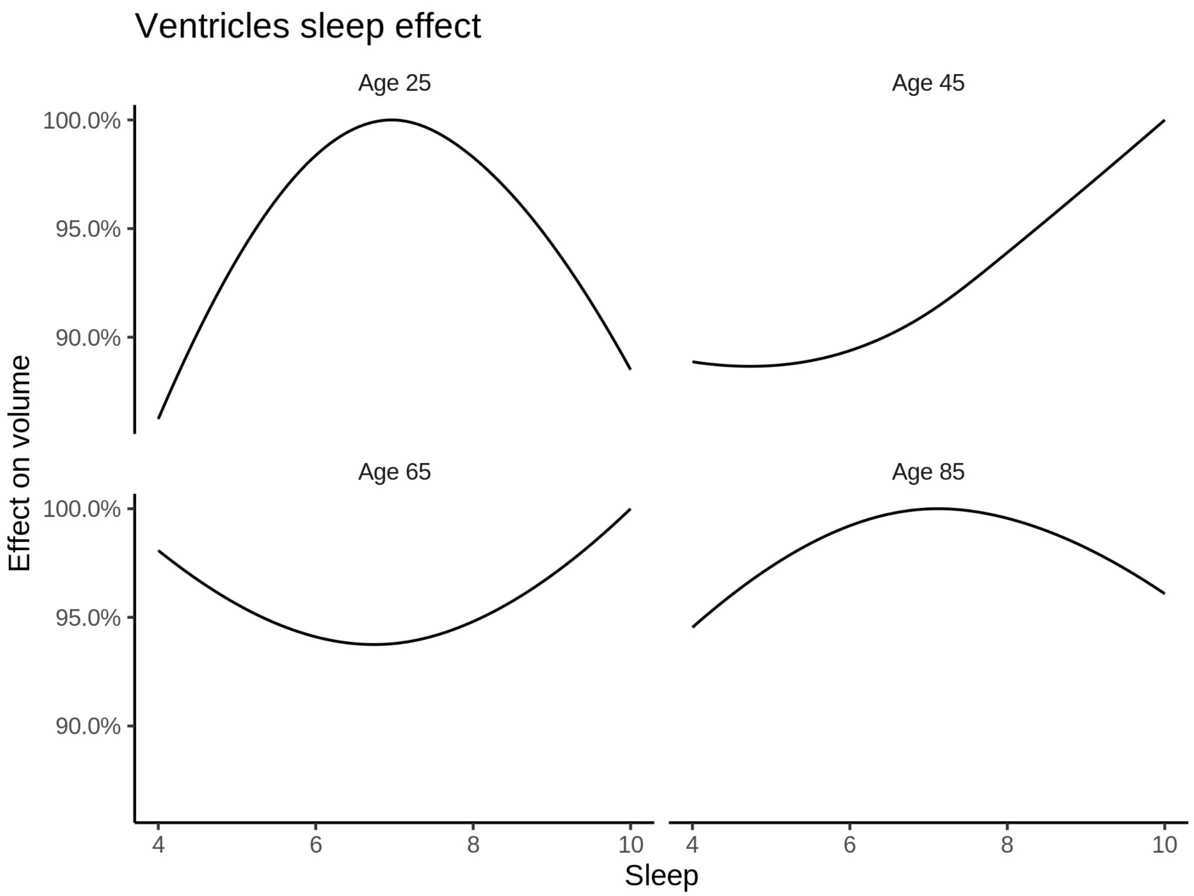


**Comparison of mean sleep and sleep associated with maximum volume**

The plot below compares average sleep to the sleep associated with maximum volume.

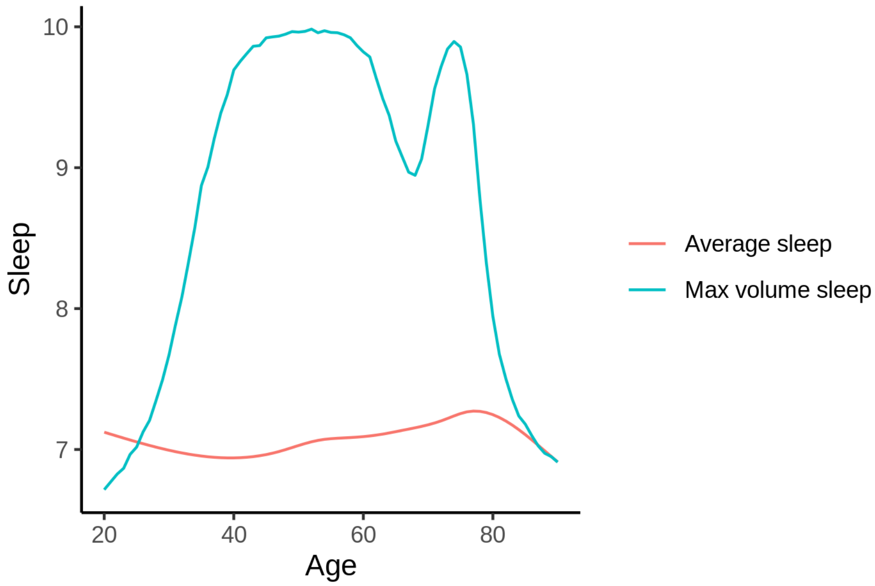

The next plot shows the difference between average sleep and sleep associated with maximum volume. The shaded region is a 95 % confidence interval.

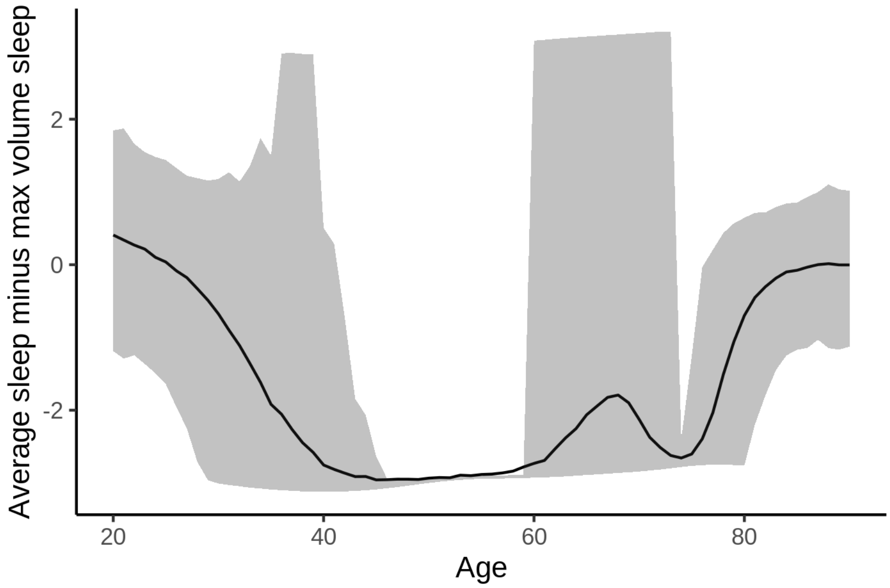


The next plot shows the probability that the sleep duration associated with maximum volume is longer than the average sleep duration, as a function of age. Probility below .05 can be interpreted as evidence that the sleep associated with maximum volume is shorter than the mean sleep, and probability above .95 can be interpreted the opposite way.

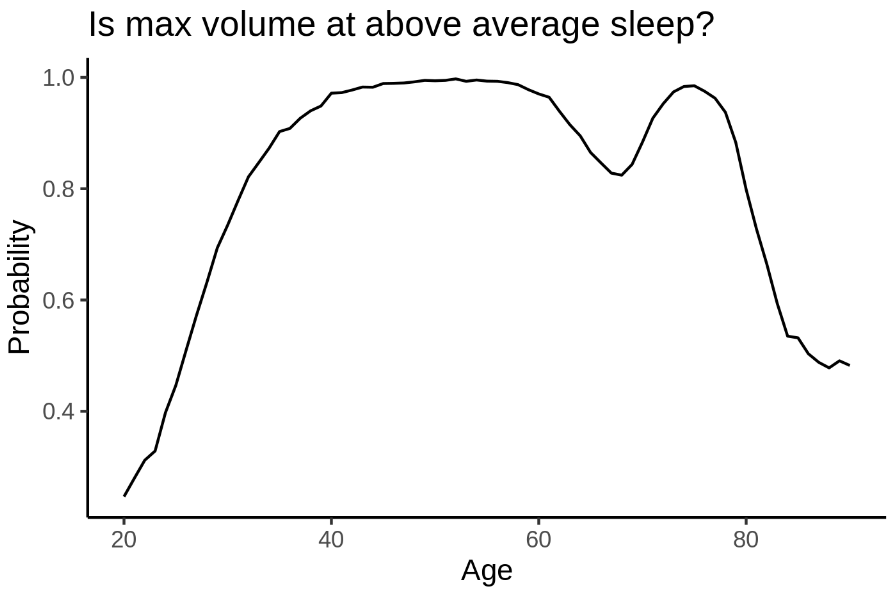


#### Summary

The plot below shows estimated sleep at maximum volume and 95 % confidence intervals for selected regions.

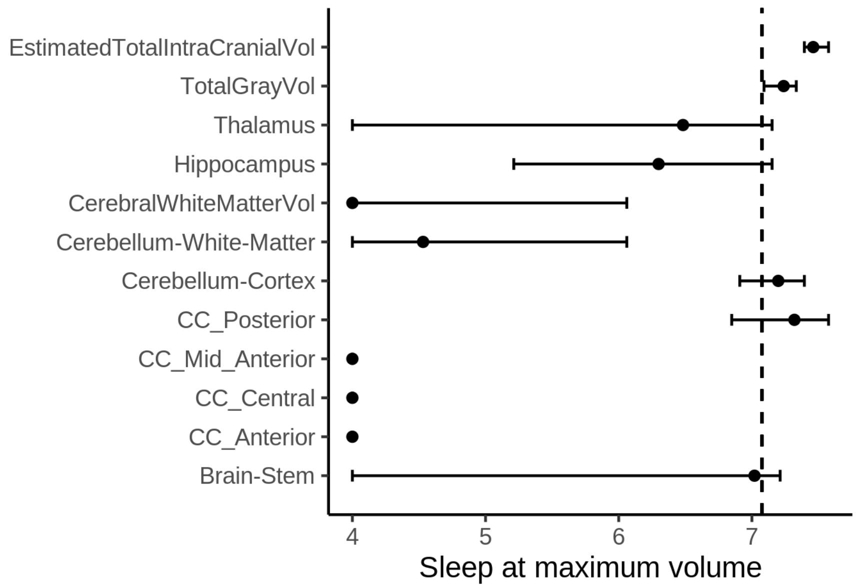


### More details on covariates

#### Raw data

##### UKB

For the UKB, we used the income in US dollars. The distribution of this variable is shown in the histogram below.

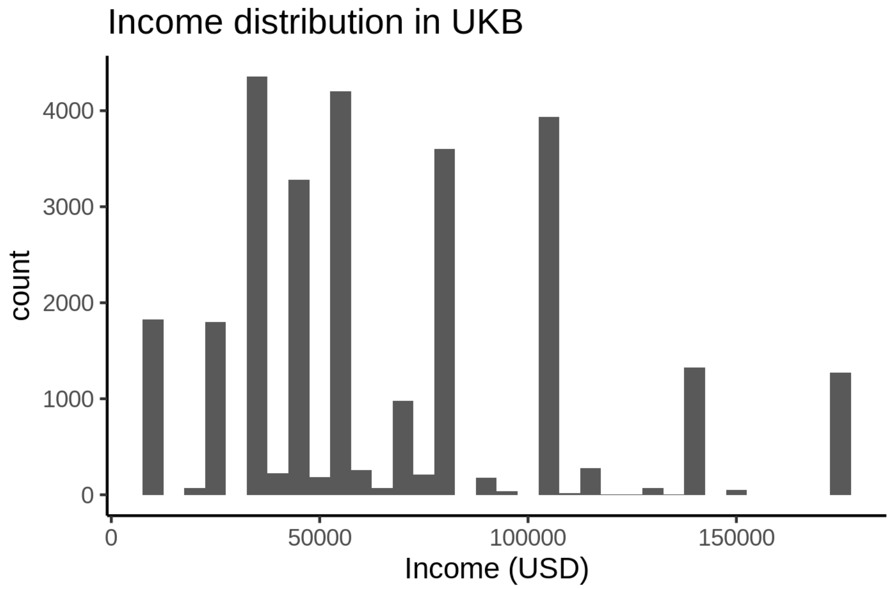


Education took four different categorical values in UKB, shown below.

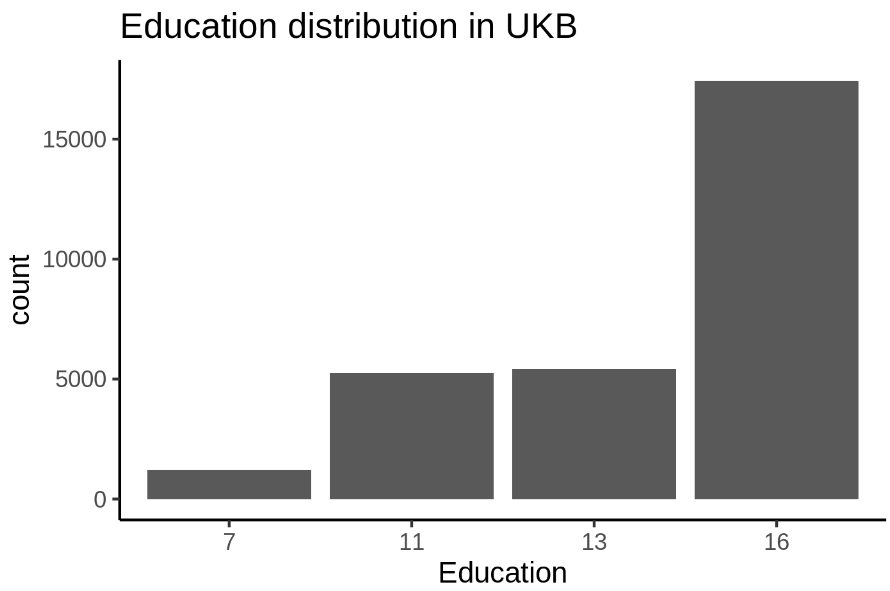


BMI distribution is shown below.

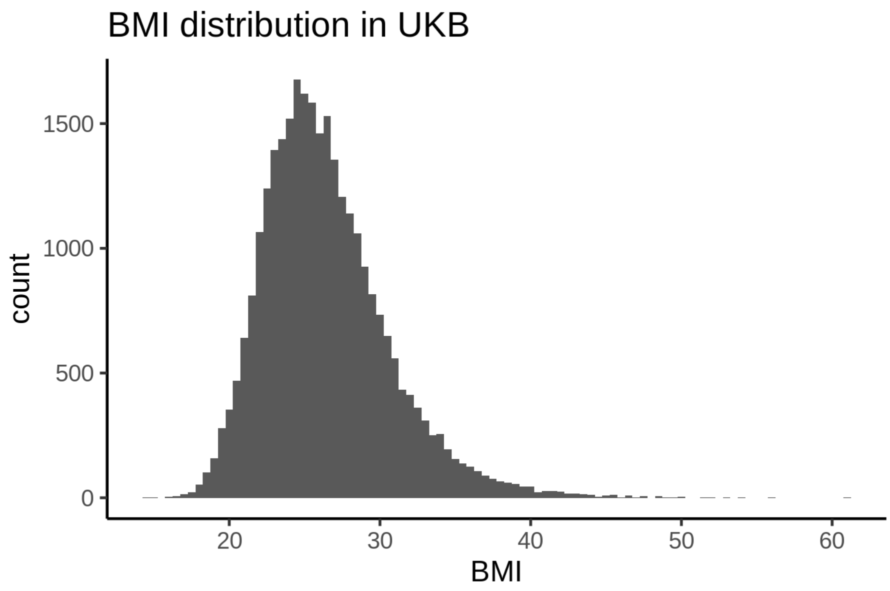


Depression distribution is shown below. This is based on the first principal component of depression covariates.

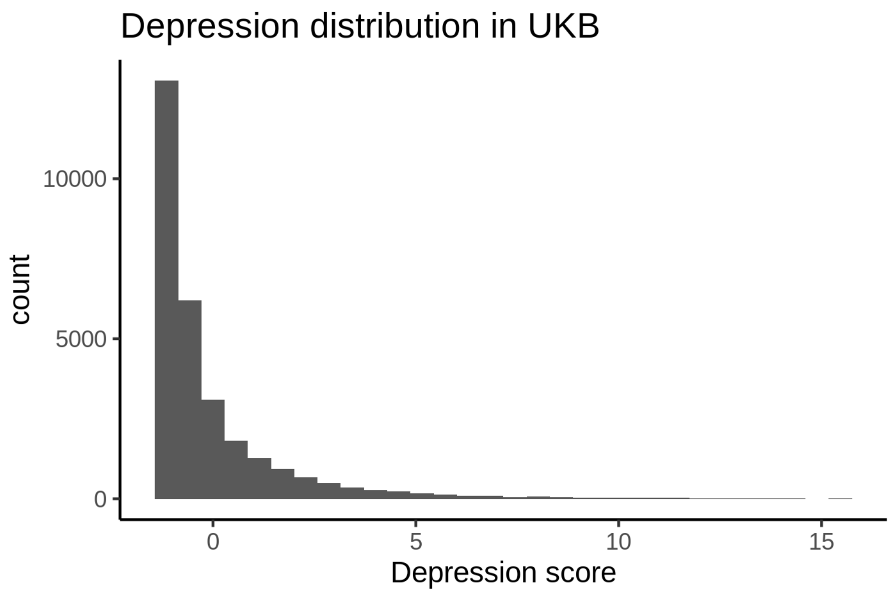


**BASEII**

Income is taken from the variable named “pnett”, which measures monthly income after tax.

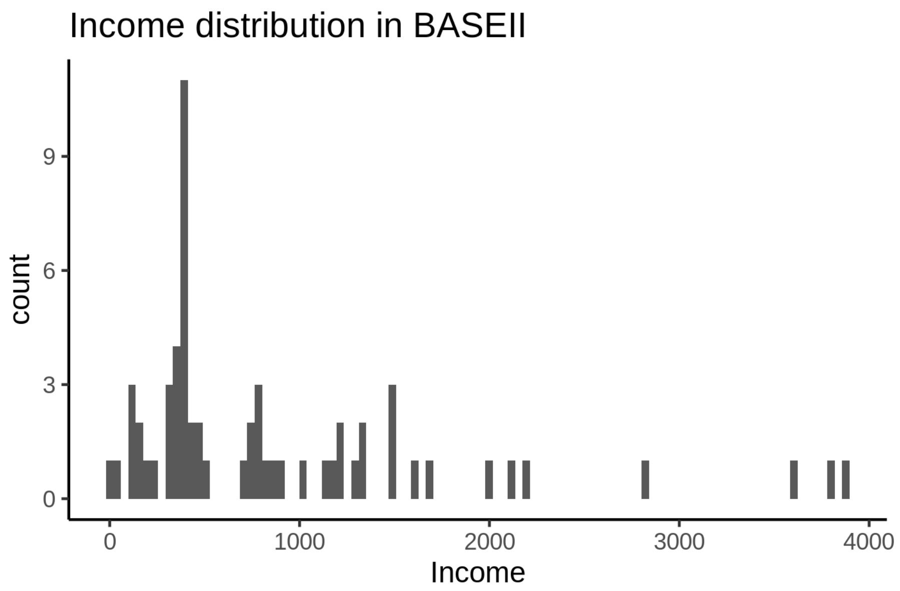

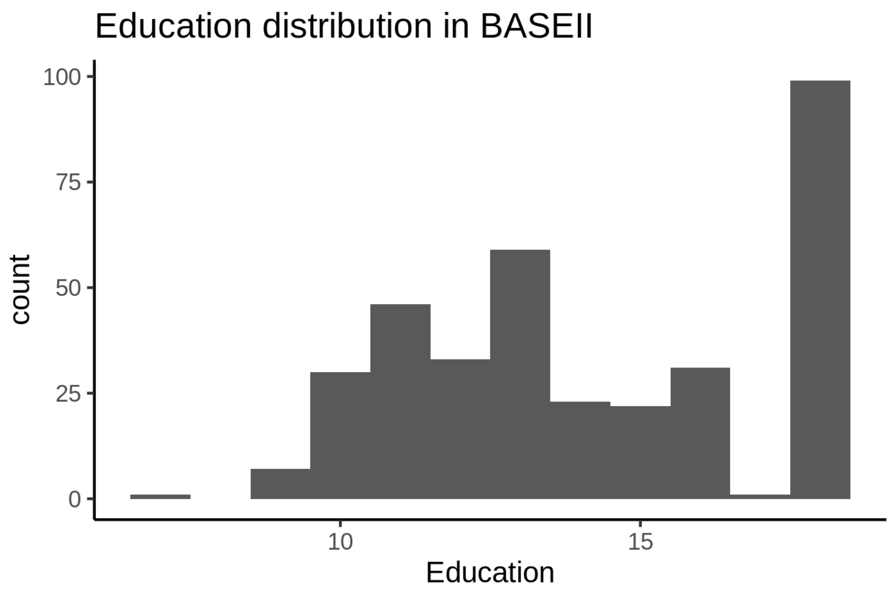


We did not have BMI information in BASEII. Depression scores are shown below.

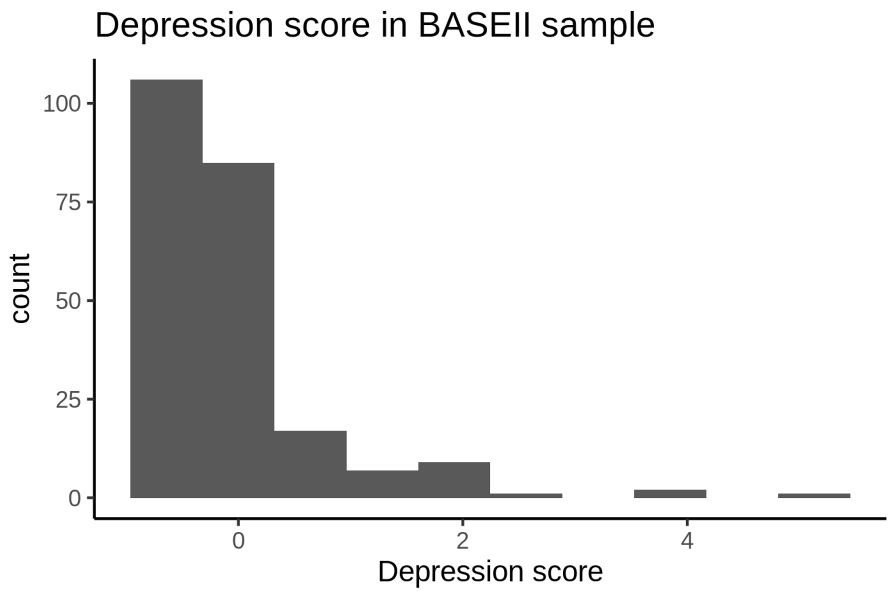


**Betula**

We did not have identifiable education and income variables for Betula, as they were anonymized. We did however have BMI and depression.

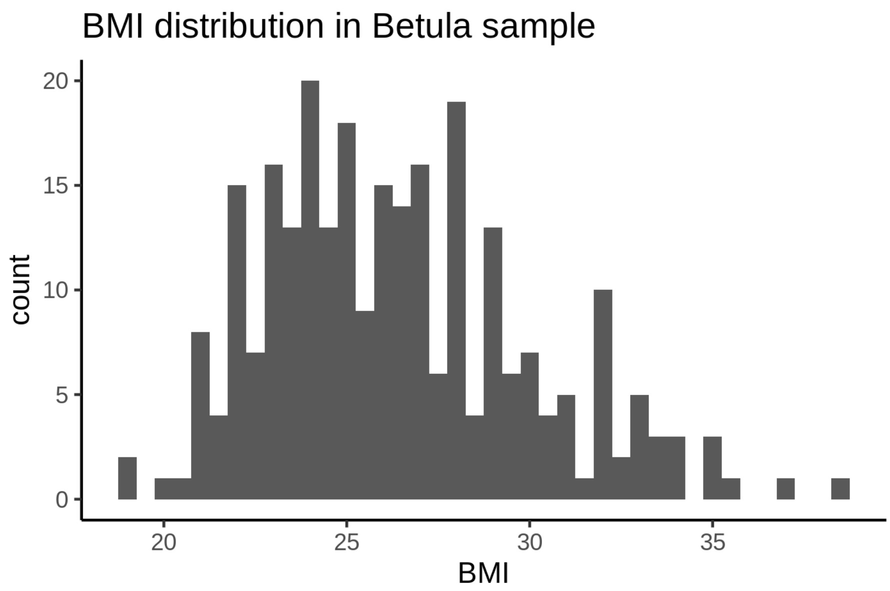


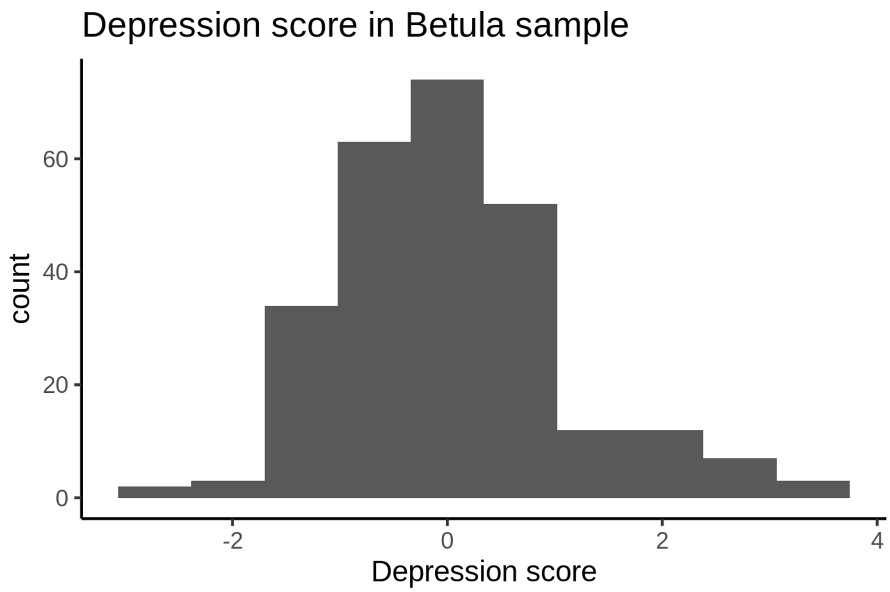


**Cam-CAN**

The income is household income.

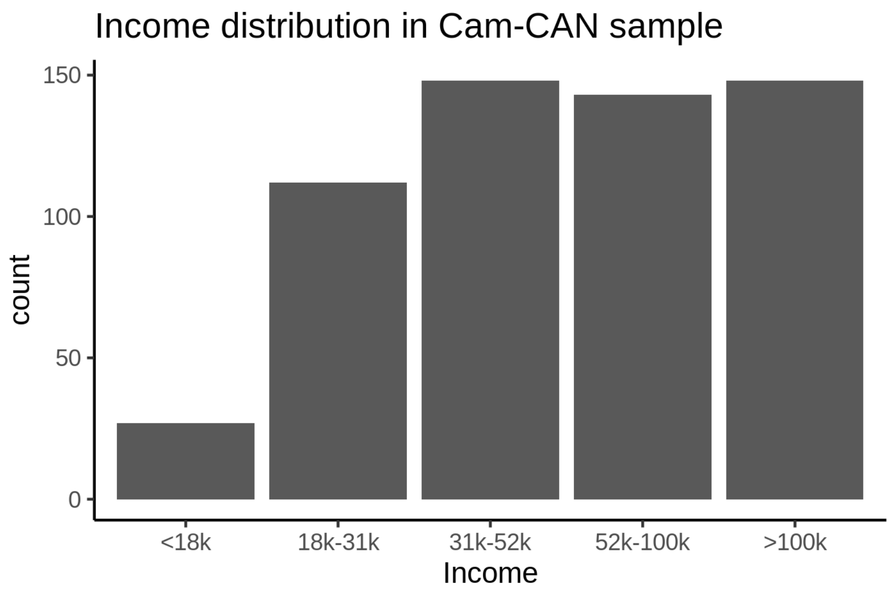


Age at completed education is shown below. The outlier with age above 50 should probably be removed.

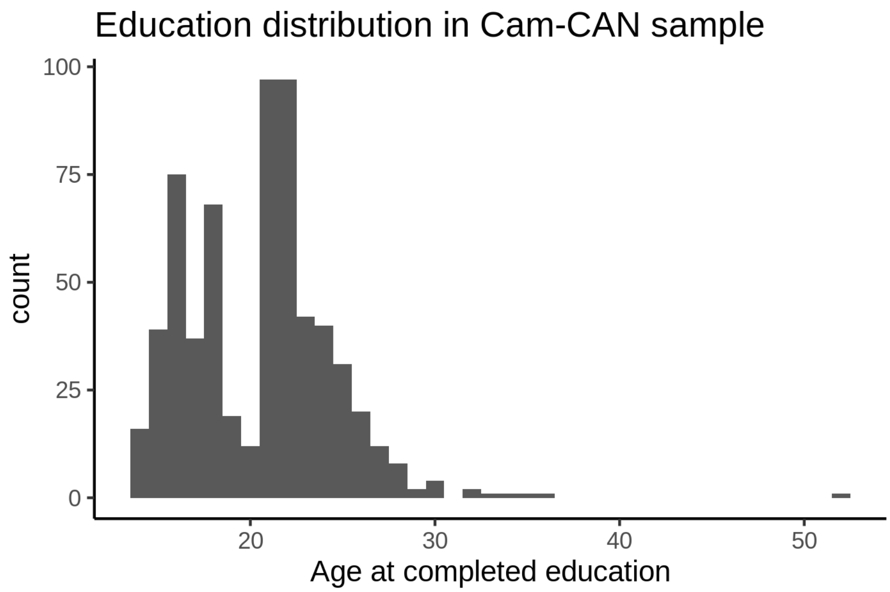


BMI distribution is shown below.

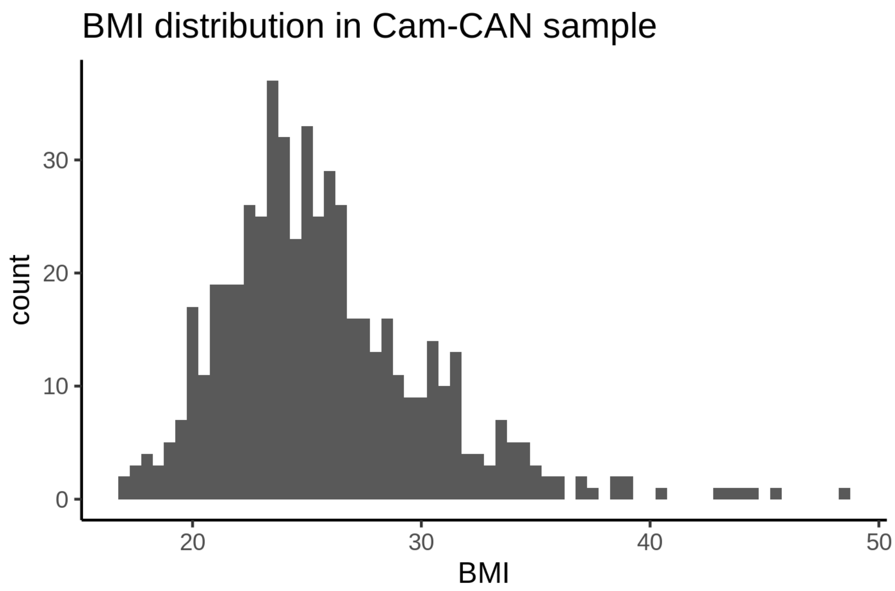
BMI distribution in Cam-CAN sample
count
0
10
20
30
20
30
40
50
BMI

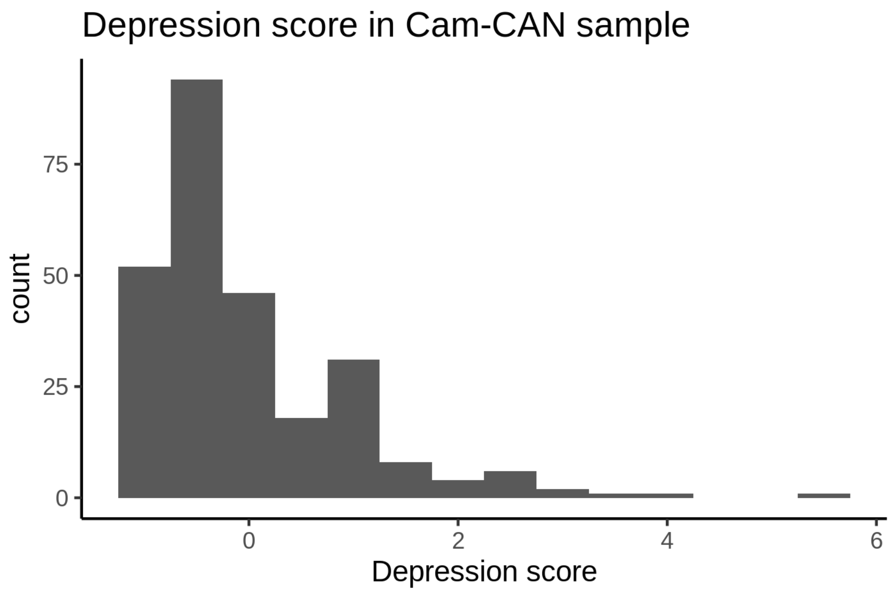
Depression score in Cam-CAN sample
count
0
25
50
75
0
2
4
6
Depression score

**LCBC**

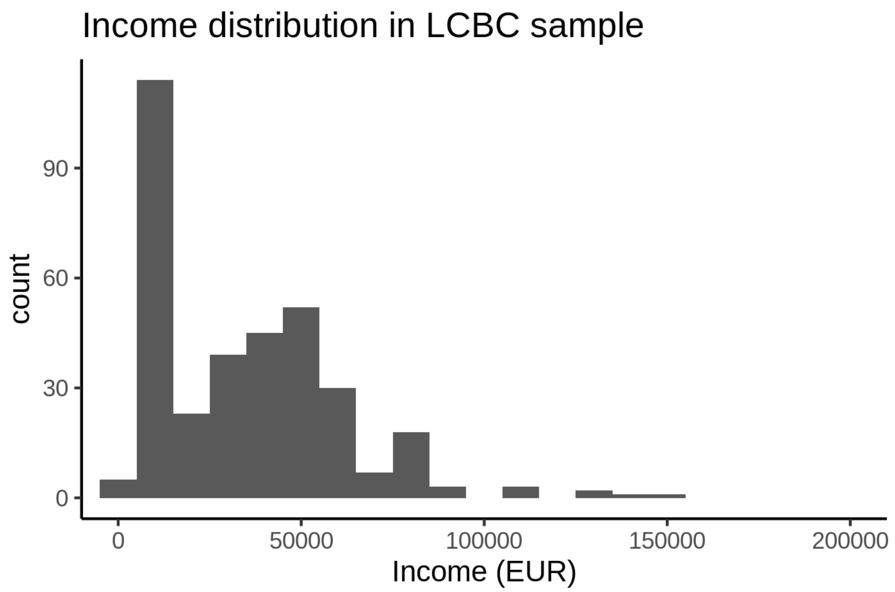
Income distribution in LCBC sample
count
90
60
30
0
0
50000
100000
150000
200000
Income (EUR)

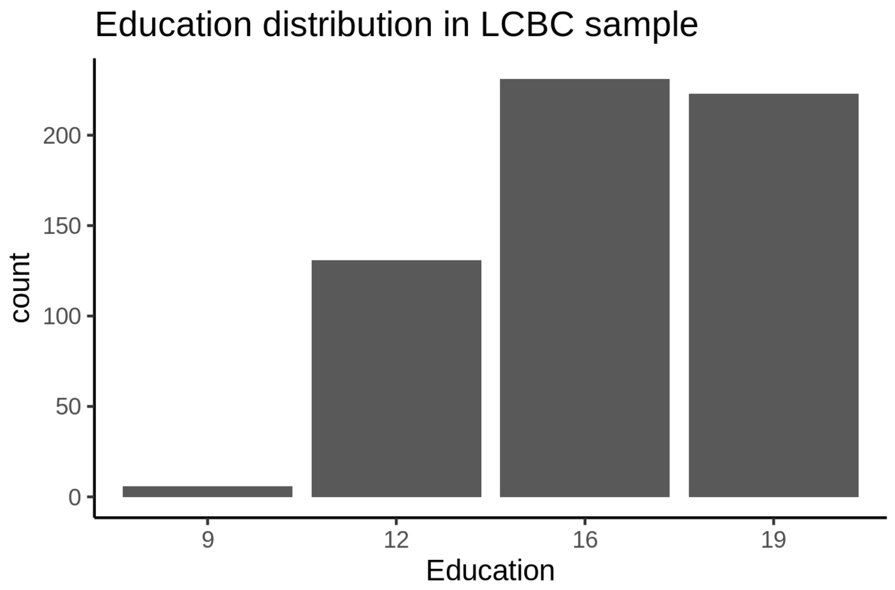
Education distribution in LCBC sample
count
200
150
100
50
0
9
12
16
19
Education

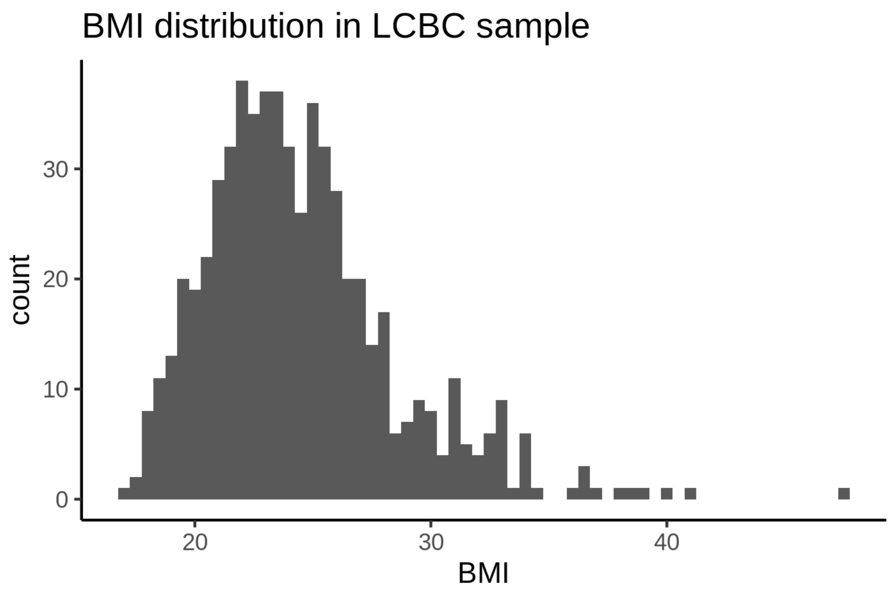


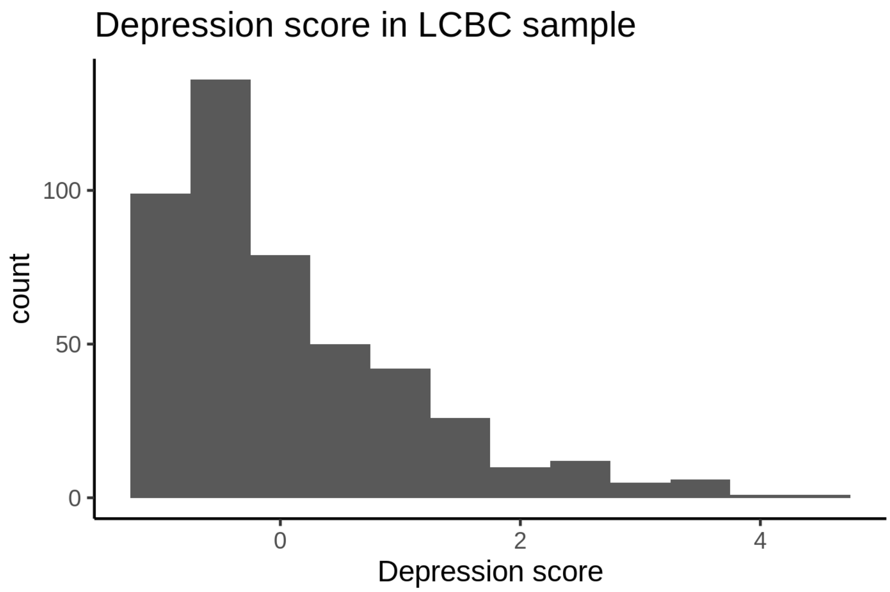


**Oxford**

Household income is on a ten-point categorical scale, cf. supplementary material to SES paper.

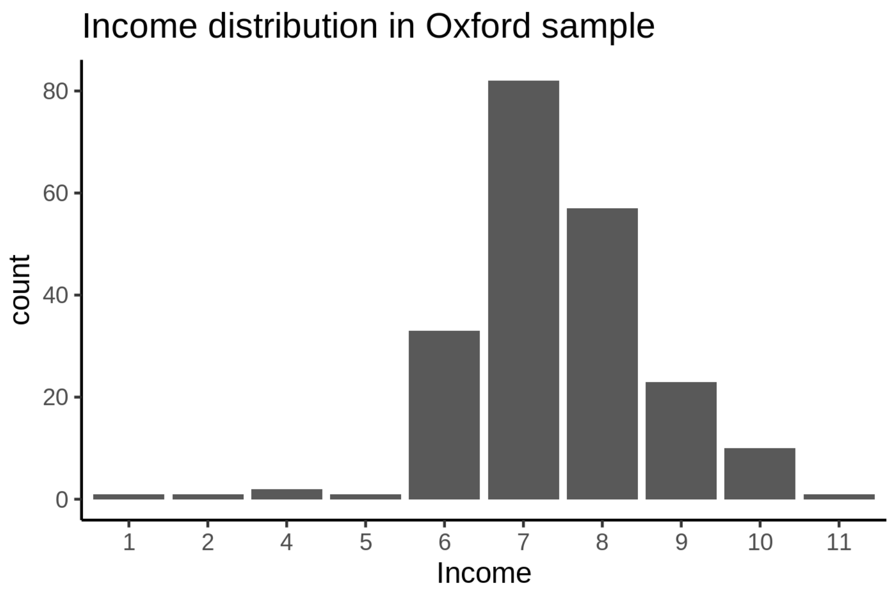


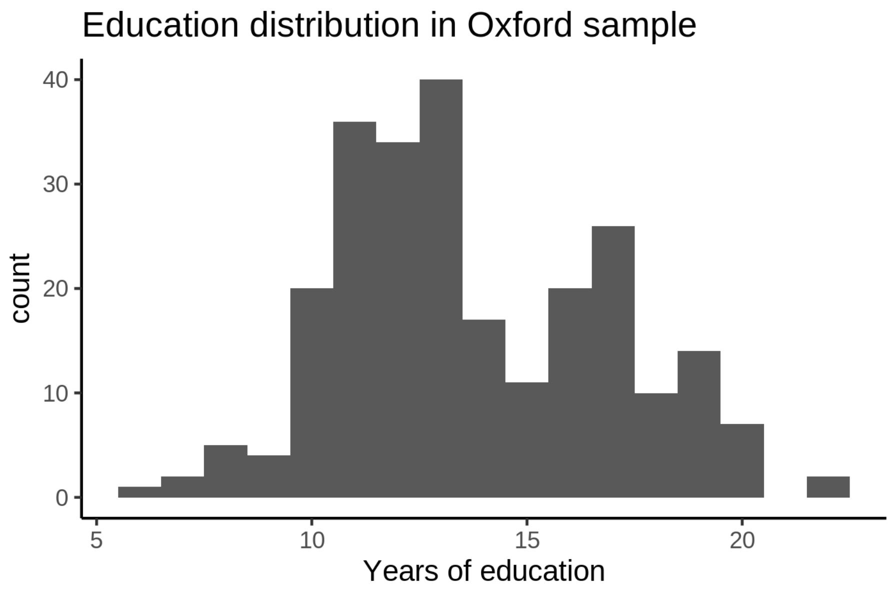


#### Data harmonization

Since the income and education variables at each site are not directly comparable, we had to convert them to something which can be compared across sites. This is described here.

##### Income

The plot below illustrates the data harmonization issue. The horizontal axis shows education levels mapped to the interval $[0, 1]$. While some sites have continuous income with very fat tail to the right (UiO, MPIB), others have predefined categories with very similar numbers in each. Since we are not able to split up the most coarse numbers into finer grained ones, we instead make the fine grained ones coarser.

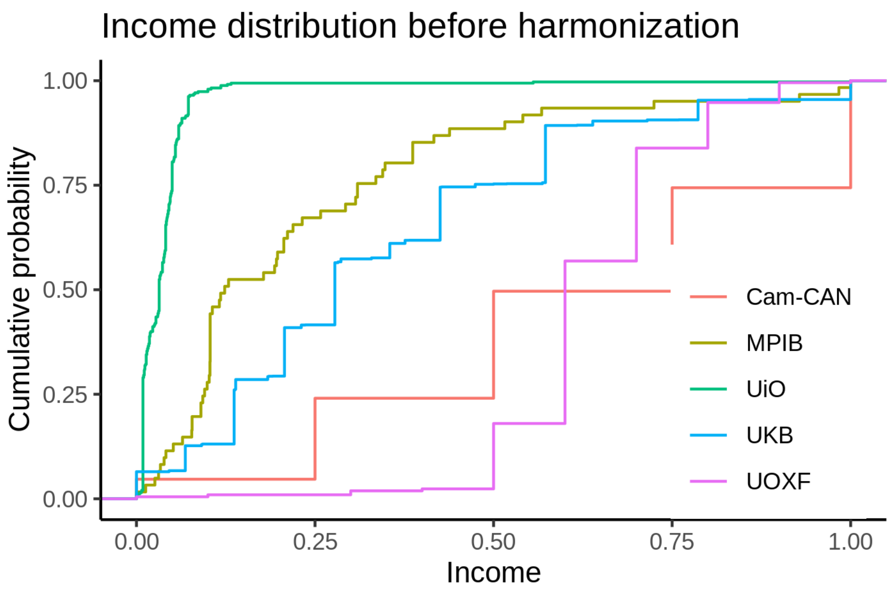


For each study separately, the income values were split into four equally sized bins, except for UOXF and Cam-CAN, for which we only had sufficient data to create three bins. The resulting distribution of values is shown in the plot below, and contains the income values used in subsequent analyses.

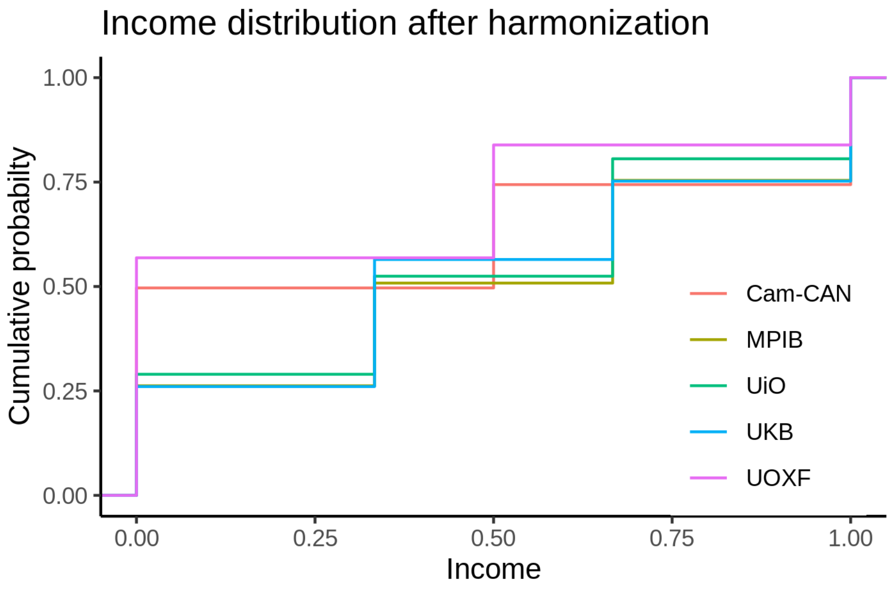


**Education**

A similar strategy was used to harmonize the education data. The plot below shows the original values.

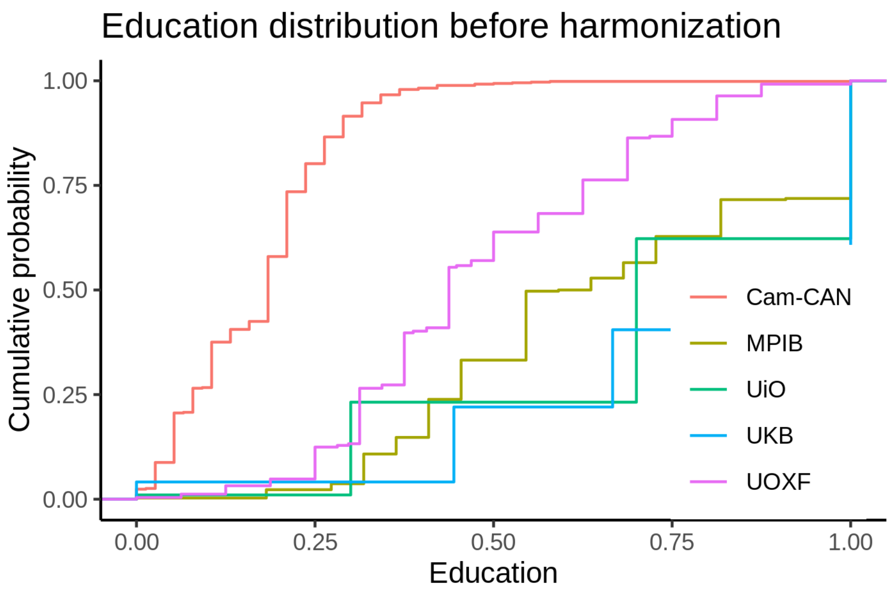


The next plot shows the distribution after harmonization. We are not able to make the distributions very similar, but at least it's an improvement over the raw values.

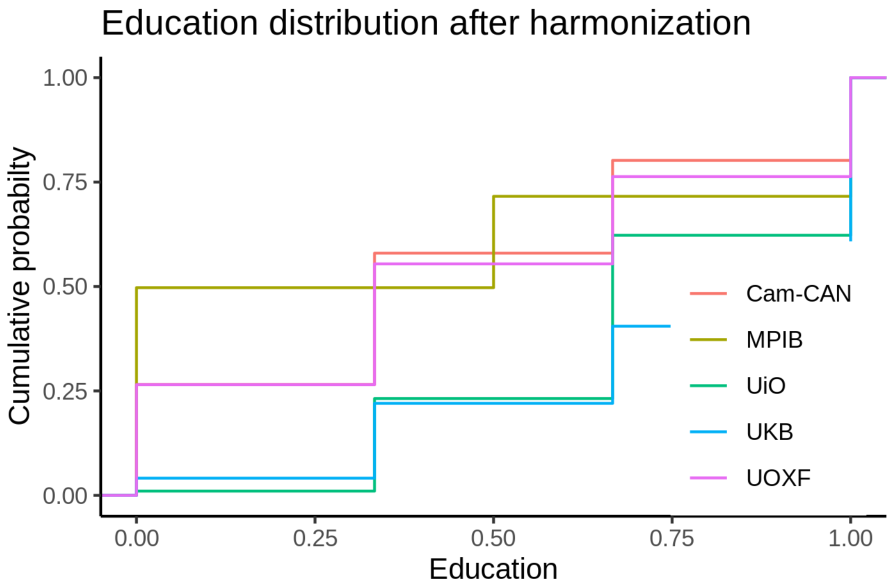


##### BMI

BMI is measured exactly the same way in each sample, so it represents the same thing. We hence use this variable as it is.

##### Depression

Distribution of depression covariates per site is shown below. The data are fairly harmonized.

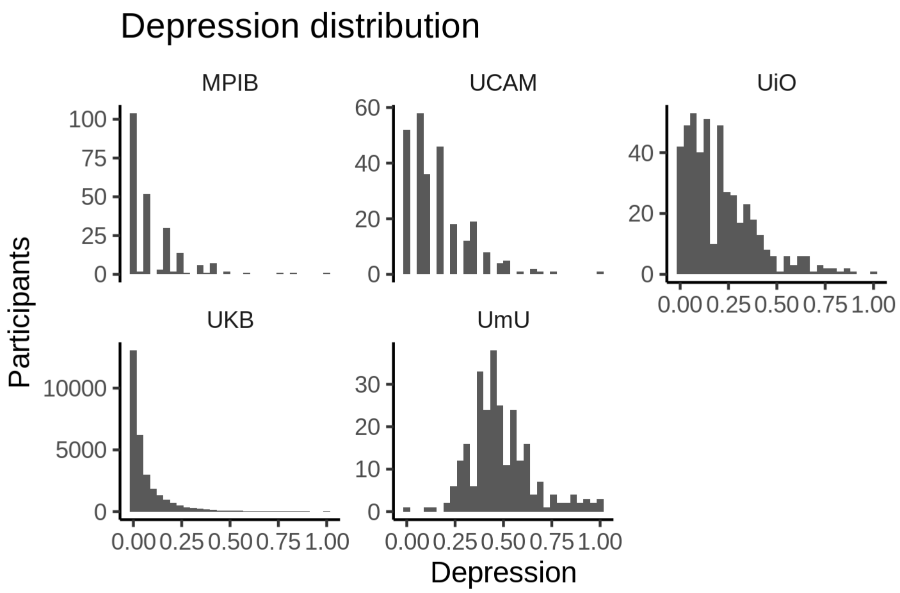


##### Relationship between covariates and sleep

###### Education and sleep

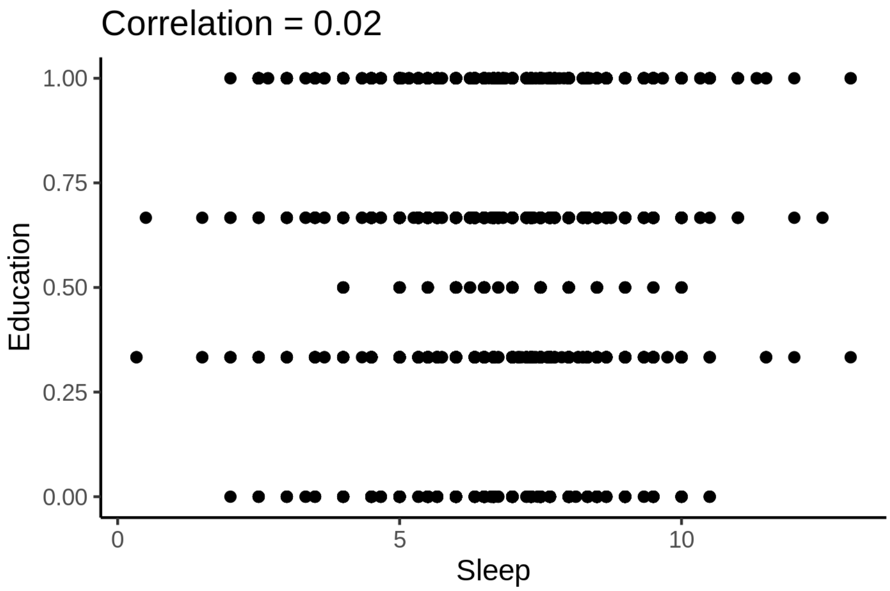

**Income and sleep**

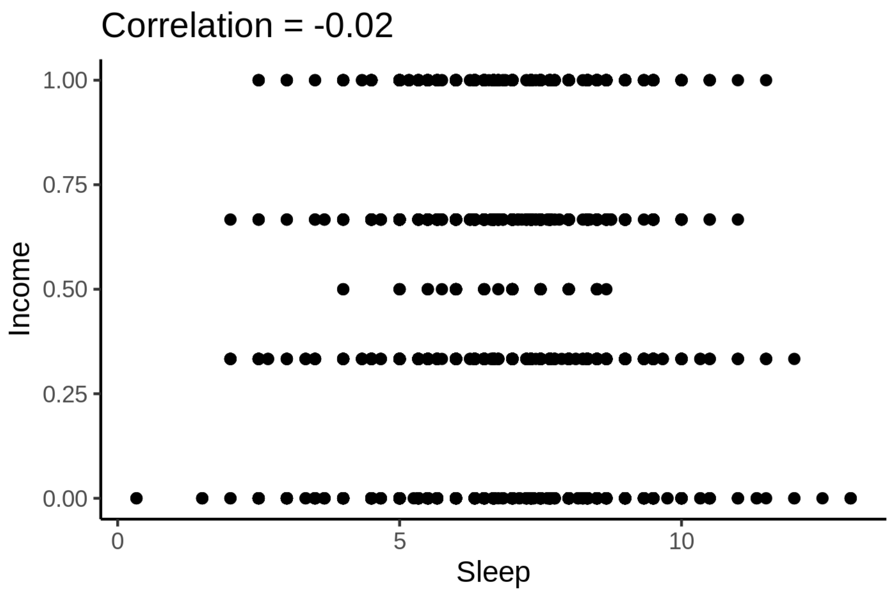


**BMI and sleep**

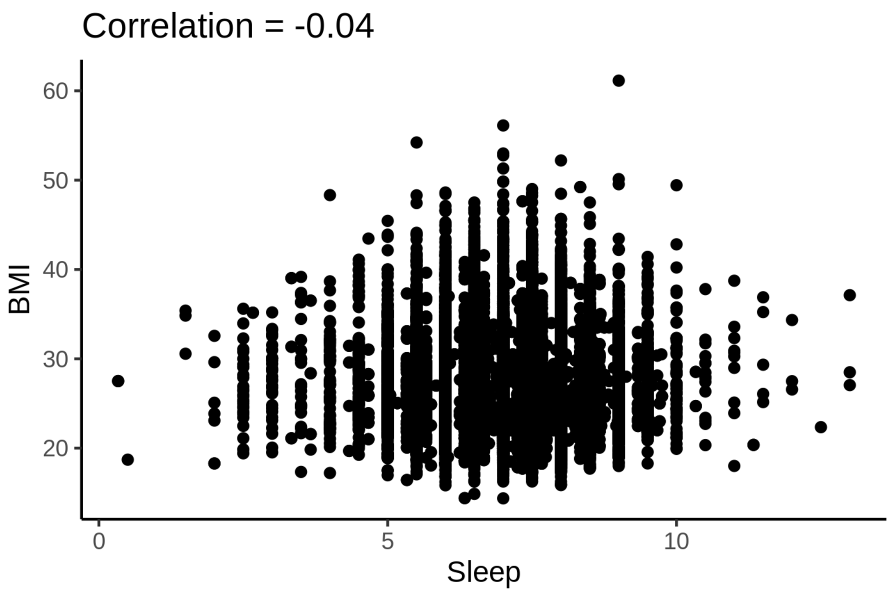

**Depression and sleep**

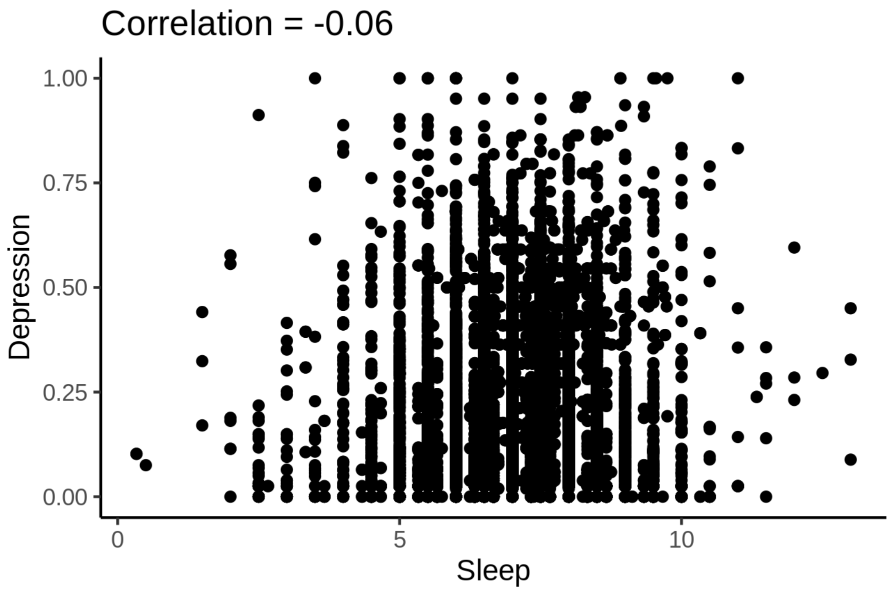
