## Supplementary material for "Sleep duration and brain structure – phenotypic associations and genotypic covariance": SI MRI Methods

**MAGNETIC RESONNANCE IMAGING METHODS**

**Specific information on image acquisition in the different samples**

T1 weighted structural scans were acquired at Siemens, Philips and GE scanners at the various sites.

| **Sample** | **Scanner** | **Field strength (Tesla)** | **Sequence parameters** |
| --- | --- | --- | --- |
| BASE-II | Tim Trio Siemens | 3.0 | TR: 2500 ms, TE: 4.77 ms, TI: 1100 ms, flip angle: 7°, slice thickness: 1.0 mm, FoV 256×256 mm, 176 slices |
| Betula | Discovery GE | 3.0 | TR: 8.19 ms, TE: 3.2 ms, TI: 450 ms, flip angle: 12°, slice thickness: 1 mm, FOV 250×250 mm, 180 slices |
| Cam-CAN | Tim Trio  Siemens | 3.0 | TR: 2250 ms, TE: 2.98 ms, TI: 900 ms, flip angle: 9°, slice thickness 1 mm, FOV 256×240 mm, 192 slices |
| LCBC | Avanto Siemens | 1.5 | TR: 2400 ms, TE: 3.61 ms, TI: 1000 ms, flip angle: 8°, slice thickness: 1.2 mm, FoV: 240×240 m, 160 slices, iPat = 2 |
|  | Avanto Siemens | 1.5 | TR: 2400 ms, TE = 3.79 ms, TI = 1000 ms, flip angle = 8, slice thickness: 1.2 mm, FoV: 240 x 240 mm, 160 slices |
|  | Skyra Siemens | 3.0 | TR: 2300 ms, TE: 2.98 ms, TI: 850 ms, flip angle: 8°, slice thickness: 1 mm, FoV: 256×256 mm, 176 slices |
|  | Prisma Siemens | 3.0 | TR: 2400 ms, TE: 2.22 ms, TI: 1000 ms, flip angle: 8°, slice thickness: 0.8 mm, FoV: 240×256 mm, 208 slices, iPat = 2 |
| UB | Tim Trio Siemens | 3.0 | TR: 2300 ms, TE: 2.98, TI: 900 ms, slice thickness 1 mm, flip angle: 9°, FoV 256×256 mm, 240 slices |
| WH-II | Verio Siemens | 3.0 | TR: 2530 ms, TE: 1.79/3.65/5.51/7.37 ms, TI: 1380 ms, flip angle: 7°, slice thickness: 1.0 mm, FOV: 256×256 mm |
| HCP | Connectome  Skyra  Siemens* | 3.0 | TR: 2400 ms, TE: 2.14 ms, TI: 1000 ms, flip angle: 8°, slice thickness: 0.7 mm, FOV: 224 mm, 256 slices, GRAPPA = 2 |
| UKB | Skyra  Siemens | 3.0 | TR: 2000 ms, TI: 880 ms, slice thickness: 1 mm, FoV: 208×256 mm, 256 slices, iPAT=2 |

**Supplementary Table** MR acquisition parameters

TR: Repetition time, TE: Echo time, TI: Inversion time, FoV: Field of View, iPat: in-plane acceleration, GRAPPA: GRAPPA acceleration factor. *Customized

**Lifebrain MRI scanning and processing**

MRI data originated from 11 different scanners (table above), and were processed with FreeSurfer (<https://surfer.nmr.mgh.harvard.edu/>) (1-4). To avoid introducing possible site-specific biases, quality control measures were imposed and no manual editing was done. We previously have reported that across-scanner consistencies in estimated hippocampal volumes for the scanners used in Lifebrain are high (5).

**HCP MRI scanning and processing**

Imaging data were collected and processed by the human connectome project (https://www.humanconnectome.org/study/hcp-young-adult) as described in (6). Imaging data were collected a customized Siemens 3T “Connectome Skyra” housed at Washington University in St. Louis, using a standard 32-channel Siemens receive head coil and a “body” transmission coil designed by Siemens specifically for the smaller space available using the special gradients of the WU-Minn and MGH-UCLA Connectome scanners. Anatomical T1-weighted magnetization-prepared rapid gradient echo (MPRAGE) images were obtained in the sagittal plane at 0.7mm isotropic resolution, and T2 weighted FLAIR images were acquired at identical resolution in the sagittal plane. Images were processed using a custom combination of tools from FSL and FreeSurfer (7-9).

**UKB MRI scanning and processing**

Imaging data were collected and processed by the UK Biobank (https://www.ukbiobank.ac.uk) as described in (10), using the FreeSurfer 6.0 software package. Imaging data were collected using 3.0 T Siemens Skyra (32-channel head coil). Anatomical T1-weighted magnetization-prepared rapid gradient echo (MPRAGE) images were obtained in the sagittal plane at 1mm isotropic resolution, and T2 weighted FLAIR images were acquired at 1.05x1x1mm resolution in the sagittal plane.
