## Supplementary material for "Sleep duration and brain structure – phenotypic associations and genotypic covariance": SI Sample characteristics

**SAMPLE DESCRIPTIVES**

**BASE II**

*Population, recruitment, inclusion/exclusion criteria and general description of study*

Participants of the Berlin Aging Study II (BASE II) were community-dwelling older adults recruited from the greater Berlin metropolitan area through advertisements in newspapers and public areas (for cohort characteristics and additional details, see [1, 2]). The baseline sample comprised 2200 participants. Of these, 1600 were older adults aged 61–88 years (mean age 71.5, SD 3.89; 793 female), and 600 were younger adults aged 24–40 years (mean age 31.1, SD 3.38; 247 female). Participants were invited to a medical exam consisting of a 2-day protocol, and two cognitive testing sessions scheduled 1 week apart, and were tested in small groups (e.g. about 6 participants per group) on a comprehensive cognitive battery that covers key cognitive abilities measured by 21 tasks. Each session lasted about 3.5 h. 1828 had valid data to be included in the present analyses.

MR Sample: After completion of the cognitive examination of BASE-II, eligible participants were invited to take part in one MRI session within a time window of 2–4 weeks after cognitive testing, consisting of 341 older adults aged 61–82 years (mean age 70.1, SD = 3.89; 131 female) and 103 younger adults (mean age 31.4, SD = 3.7; 39 female). MR scans and cognitive scores were obtained 2012-2013. A subsample of the MR sample was later re-invited for follow-up. The different elements of the study were approved by the ethics committees of the Max Planck Institute for Human Development, the Charité University ethics committee and by the ethics committees of The German Association for Psychology (DGPs). Participants signed written informed consent and received monetary compensation for their participation in BASE-II and the MRI study. All experiments were performed in accordance with relevant guidelines and regulations.

*Inclusion/exclusion criteria/ screening*

Inclusion criteria for taking part in this study were age between 20 and 35 or 60 and 80 years, apparently healthy. Exclusion criteria were untreated diabetes and hypertension; prior stroke, head injuries or brain surgery; psychiatric illness; major depression; dementia with a score < 24 on the Mini-Mental State Examination. To that end, none of the participants took medication that might affect memory function or had a history of head injuries, medical (e.g., heart attack), neurological (e.g., epilepsy), or psychiatric disorders (e.g., depression). All participants reported normal or corrected to normal vision, were right-handed, and scored over 27 on the Mini-Mental Status Examination.

**Betula**

*Population, recruitment and general description of study/ procedures*

In the Betula longitudinal study on aging, memory and dementia, population-based sampling of healthy middle-aged and older adults was used for recruitment. Detailed recruitment procedures are found in [3, 4]. For the current analyses, the MRI subsample of the study is used. Participation in the neuroimaging study was offered to all participants who had remained in the study and completed cognitive testing at the 5th Betula test wave in 2008-2009, and 376 participants from underwent structural and functional MRI in 2009-2010.

*Inclusion/exclusion criteria/ screening*

Exclusion criteria were severe visual or auditory handicaps, intellectual or developmental disabilities, suspected dementia, having a mother tongue other than Swedish, MRI contraindications, neurological disorders, or visual/motor deficits that could interfere with fMRI data collection, MMSE <24, brain or head surgery, and substantial brain anatomical deviations. Eight participants were excluded completely post scanning due to discovered neurological conditions (Schizophrenia, Multiple Sclerosis, Parkinson’s Disease, Hydrocephalus, Alcoholism, and dementia), and an additional two participants were excluded due to MMSE scores below 24. In addition, for 29 participants MRI data only was excluded due to anatomical deviations (subdural hematoma, localized loss of brain tissue, subcortical atrophy, and previous brain or head surgery (n=2)), movement artifacts (n=21), or FreeSurfer processing failures (n=3). Three individuals had missing T1 images due to incomplete acquisition. Age at MRI-scanning, reported with a one decimal precision, was used in the analyses. Testing was performed 1-18 months prior to scanning (mean interval: 9 months).

**Cam-Can**

*Population, recruitment and general description of study/ procedures*

Recruitment was done by invitation letters based on the patient lists of general practitioners within the Cambridge City area. A population-based cohort of 3000 adults aged 18 or above was recruited to Stage 1 of the project, where they completed an interview including health and lifestyle questions, a core cognitive assessment, and a self-completed questionnaire of lifetime experiences and physical activity. Of those interviewed, ~700 participants aged 18-87 (100 per age decile) continued to Stage 2 where they undergo cognitive testing and provide measures of brain structure and function. A subset of ~250 adults returned for longitudinal follow-up data. The study is conducted in compliance with the Helsinki Declaration, and has been approved by the local ethics committee, Cambridgeshire 2 Research Ethics Committee (reference: 10/H0308/50).

*Inclusion/exclusion criteria/ screening*

General exclusion criteria: Term-time residents of colleges and universities, and participants whose Primary Care Physician feel are inappropriate to include. Exclusion criteria for the MRI part of the study: Not cognitively normal (MMSE < 24, memory defect, consent difficulties), communication difficulties (hearing problems [35db at 1000 Hz], insufficient English language, vision difficulties), medical problems by self-report of diagnosis (dementia diagnosis /Alzheimer’s Disease, Parkinson’s Disease, Motor Neurone disease, Multiple sclerosis, cancer, stroke, encephalitis, meningitis, epilepsy, head injury with serious results [coma, unconscious for >2 hours, skull fracture], recently diagnosed or uncontrolled high blood pressure, possible pregnancy, current psychiatric conditions [bipolar disorder, schizophrenia, psychosis]), mobility problems (restricted mobility which could prevent further participation, inability to walk 10 metres), substance abuse (past or current treatment for drug abuse, current drug usage), MRI/ MEG safety and comfort exclusions.

**LCBC**

*Population, recruitment and general description of study/ procedures*

Cognitively healthy, community dwelling participants across the lifespan were drawn from studies coordinated by the Research Group for Lifespan Changes in Brain and Cognition (LCBC [www.oslobrains.no](http://www.oslobrains.no)), approved by a Norwegian Regional Committee for Medical and Health Research Ethics. Written informed consent was obtained from all adult participants and from parents or other legal guardians for participants below age of majority. The samples were recruited by newspaper and web page adds, and part of the developmental sample was recruited through the population registry study MoBa <https://www.fhi.no/en/studies/moba/> Most participants, including all children, were recruited for observational studies, while some adults were recruited to enter into cognitive training studies after baseline assessment.

*Inclusion/exclusion criteria/ screening*

Adult participants were screened using a standardized health interview prior to inclusion in the study. Participants with a history of self- or parent-reported neurological or psychiatric conditions, including clinically significant stroke, serious head injury, untreated hypertension, diabetes, and use of psychoactive drugs within the last two years, were excluded. Further, participants reporting worries concerning their cognitive status, including memory function, were excluded. All participants above 40 years scored ≥24 on the Mini Mental State Examination [5].

**UB**

*Population, recruitment and general description of study/ procedures, inclusion/ exclusion criteria/ screening*

Cohorts from the University of Barcelona (UB) site were collapsed across a number of substudies described below.

*WAHA cohort [6]:* Recruitment and selection of participants took place between May 2012 and May 2014; the trial ended May 31, 2016. Participants were healthy elderly men and women with normal cognitive and visual function. Inclusion criteria were age between 63 and 79 years, apparently healthy, and equally willing to be in either of the two groups. Exclusion criteria included inability to undergo neuropsychological testing; morbid obesity (BMI ≥ 40 kg/m^2^); uncontrolled diabetes (HbA1c > 8%); uncontrolled hypertension (on-treatment blood pressure ≥ 150/100 mmHg); prior stroke, significant head trauma or brain surgery; relevant psychiatric illness; major depression; cognitive deterioration or dementia with a score < 24 on the Mini-Mental State Examination; other neurodegenerative disorders like Parkinson’s disease; advanced AMD or eye-related conditions precluding ophthalmological evaluation; prior chemotherapy; chronic illness with projected shortened lifespan; allergy to walnuts; customary use of fish oil and/or tree nuts (> 2 servings/week) and/or other relevant sources of ALA, such as flaxseed oil or soy lecithin. Eligible participants were recruited via mailing study brochures (LLU) or through the non-profit organization Institute of Aging (BCN), advertisements in the study centers, and word of mouth. Interested individuals attended an informational group meeting, completed a short medical questionnaire and signed the informed consent. Next candidates had a face-to-face interview with the study clinician, who assessed potential compliance, reviewed the medical history, inclusion and exclusion criteria, and recent blood work and use of medications or supplements, and administered the MMSE. Eligible participants were scheduled to have baseline tests (neuropsychological and ophthalmologic evaluations and collection of fasting blood and urine) and were then randomized to either the control or walnut group using a computerized random number table with stratification by center, sex, and age range. Couples entering the study were treated as one number and were randomized into the same group.

*CR and iTBS cohorts [7]:* Healthy volunteers older than 60 were recruited via the Institute of Aging, Barcelona. Individuals willing to participate were gathered in an informal meeting to tell them about the investigation, which included repetitive transcranial magnetic stimulation (TMS). Eligible participants had a normal cognitive profile with MMSE scores ≥ 24 and performances not below 1.5 SD according to normative scores (adjusted for age and education (Peña-Casanova et al., 2009)) on a neuropsychological evaluation.

*GABA cohort [8]:* Participants were recruited from the Institute of Aging (Barcelona) and the University of Experience, an initiative by the University of Barcelona for students aged 55 and older, offering special one and two-year degrees. Older adults, aged ≥ 60 years (age (mean ± *SD*), 68.15 ± 4.6 years; age range: 60–79 years), naive to stimulation, participated in this study after giving informed consent, in accordance with the Declaration of Helsinki (1964, last revision 2013). None of the participants reported a diagnosis of a neurological or psychiatric disorder or any TMS contraindication. Inclusion criteria for the older subjects included a normal cognitive profile with mini-mental state examination (MMSE) scores of ≥ 24 and performance scores not more than 1.5 standard deviation (*SD*) below normative data (adjusted for age and years of education) on any of the administered neuropsychological tests (i.e., they did not fulfil the criteria for mild cognitive impairment (MCI). The neuropsychological battery included (1) a screening test for dementia, using the MMSE, and an evaluation of: (2) premorbid cognition and intelligence quotient (IQ), using the vocabulary subtest of the Wechsler Adult Intelligence Scale-III (WAIS-III) and National Adult Reading Test (NART); (3) verbal memory, using the Free and Cued Selective Reminding Test (SRT); (4) executive functions, using the phonemic fluency task and Trail Making Test B (TMTB); (5) language, using the semantic fluency task and Boston Naming Test (BNT); and (6) speed of processing, using the Symbol Digit Modalities Test (SDMT).

*PD and MSA cohorts [9]:* Healthy volunteers were recruited from the Aging Institute in Barcelona. Inclusion criteria were: scores within normality in a battery of neuropsychological tests described below (adjusted for age and education), IQ within normality as measured by the Vocabulary subtest (WAIS, scalar score >7). Exclusion criteria included: diagnosis of any neurological or psychiatric disease, mild cognitive impairment (MCI), any other condition that might affect cognition (chemotherapy or radiotherapy), diagnosis of fibromyalgia and any incompatibility to undergo MRI. Visuospatial and visuoperceptual functions were assessed with Benton Visual Form Discrimination (VFD) and Judgment of Line Orientation (JLO) tests; executive functions were evaluated with phonemic (words beginning with the letter “p” in 1 minute) and semantic (animals in 1 minute) fluencies; memory through total learning recall (sum of correct responses from trial I to trial V) and delayed recall (total recall after 20 min) through scores on Rey’s Auditory Verbal Learning Test (RAVLT). Attention and WM were assessed with Digit Span Forward and Backward, the Stroop Color-Word Test, Symbol Digits Modalities Tests (SDMT) and the Trail Making Test (in seconds), part A (TMTA) and part B (TMTB); and language was assessed by the total number of correct responses in the short version of the Boston Naming Test (BNT).

**Whitehall**

*Population, recruitment and general description of study/ procedures*

The Whitehall II study, starting in 1985, includes 10.308 British civil servants followed over time, which allows exploring factors hypothesized to affect brain health and cognitive aging. At study start, phase 1 (1985-1988), the population representative cohort was in the age range 35-55 years. MRI was done in Phase 11 (2012-2013) of this study, at which time the total number participants was 6035, and the age range was 60-85 years [10]. A random sample willing and able to give informed consent to participant in the imaging sub-study of Whitehall II was included, and it is this sub-cohort of the Whitehall study which is included in the present. analyses. Ethical approval was granted generically for the “Protocol for non-invasive magnetic resonance investigations in healthy volunteers” (MSD/IDREC/2010/P17.2) by the University of Oxford Central University/ Medical Science Division Interdisciplinary Research Ethics Committee (CUREC/MSD-IDREC), who also approved the specific protocol: “Predicting MRI abnormalities with longitudinal data of the Whitehall II sub-study” (MSD-IDREC-C1-2011-71).

*Inclusion/exclusion criteria/ screening*

A random selection of 800 participants from the Whitehall II phase 11 study was done. Exclusion criteria were MRI contraindications, or being unable to travel to Oxford without assistance. After the exclusion of those with excess motion (N=4) and self-reported history of stroke (N=16). No participant was diagnosed with dementia.

**HCP**

*Population, recruitment and general description of study/ procedures*

The Human Connectome project (HCP) aimed to study and freely share data from 1200 young adults (ages 22-35) from families with twins and non-twin siblings, using a protocol that includes structural and functional magnetic resonance imaging (MRI, fMRI), diffusion tensor imaging (dMRI) at 3 Tesla (3T) and behavioral and genetic testing. The dataset used was the 1200 Subjects Release (<https://www.humanconnectome.org/storage/app/media/documentation/s1200/HCP_S1200_Release_Reference_Manual.pdf>). 243 MZ and DZ twin pairs were identified by genotyping. Age was entered as years without decimals.

*Inclusion/exclusion criteria/ screening*

The participant pool comes from healthy individuals born in Missouri to families that include twins, based on data from the Missouri Department of Health and Senior Services Bureau of Vital Records. Additional recruiting efforts in HCP were used to ensure that participants broadly reflect the ethnic and racial composition of the U.S. population as represented in the 2000 decennial census. ‘Healthy’ was broadly defined, aiming for a pool that is generally representative of the population at large, to capture a wide range of variability in healthy individuals with respect to behavioral, ethnic, and socioeconomic diversity. Sibships with individuals having severe neurodevelopmental disorders (e.g., autism), documented neuropsychiatric disorders (e.g., schizophrenia or depression) or neurologic disorders (e.g., Parkinson's disease) were excluded. Individuals with illnesses such as diabetes or high blood pressure were additionally excluded, as were non-twins (current sample) born prior to 37 weeks gestation. Individuals who were smokers, overweight, or had a history of heavy drinking or recreational drug use without having experienced severe symptoms were included. For a full list of inclusion and exclusion criteria, see [11].

**UKB**

*Population, recruitment and general description of study/ procedures*

UK Biobank (UKB) (<https://www.ukbiobank.ac.uk/about-biobank-uk/>) is a major national and international health resource with the aim of improving the prevention, diagnosis and treatment of a wide range of illnesses. UK Biobank recruited ≈500,000 people aged between 40-69 years in 2006-2010 from across the country to take part in this project [12]. Potential participants were identified through National Health Service (NHS) registers according to being aged 40-69 and living within a reasonable travelling distance of an assessment centre. Assessment centres (22 in total) are located in accessible and convenient locations with a large surrounding population. Participants have undergone measures and provided samples and detailed information about themselves and agreed to have their health followed. Age was calculated from year and month of birth (day of month is missing, and was set to 1 for all subjects) to date of assessment. Age was calculated at the 0_0 timepoint for participants without MRI, and 2_0 timepoint for participants with MRI.
