## Supplementary material for "Sleep duration and brain structure – phenotypic associations and genotypic covariance": SI Cortical clustering

### Clustering of cortical regions

Data preparation, including outlier removal, was done exactly as for the subcortical volumes, and won't be repeated here.

For each measure (area, thickness, volume), the following GAMMs was fitted to each of 33 cortical ROIs. When the measure was thickness, the term `icv` was not included.

```
mod <- gamm4(value ~ sex + site + icv + s(age) + s(sleep),
             random =~(1|id), data = dat)
```

K-means clustering was run for each measure separately, dividing the regions based on the similarity of the sleep-brain curves described by the term `s(age)`. The plots below show how the total within-cluster sum-of-squares depends on the number of clusters. As usual, the plots are not very conclusive, but three clusters were chosen for each measure.

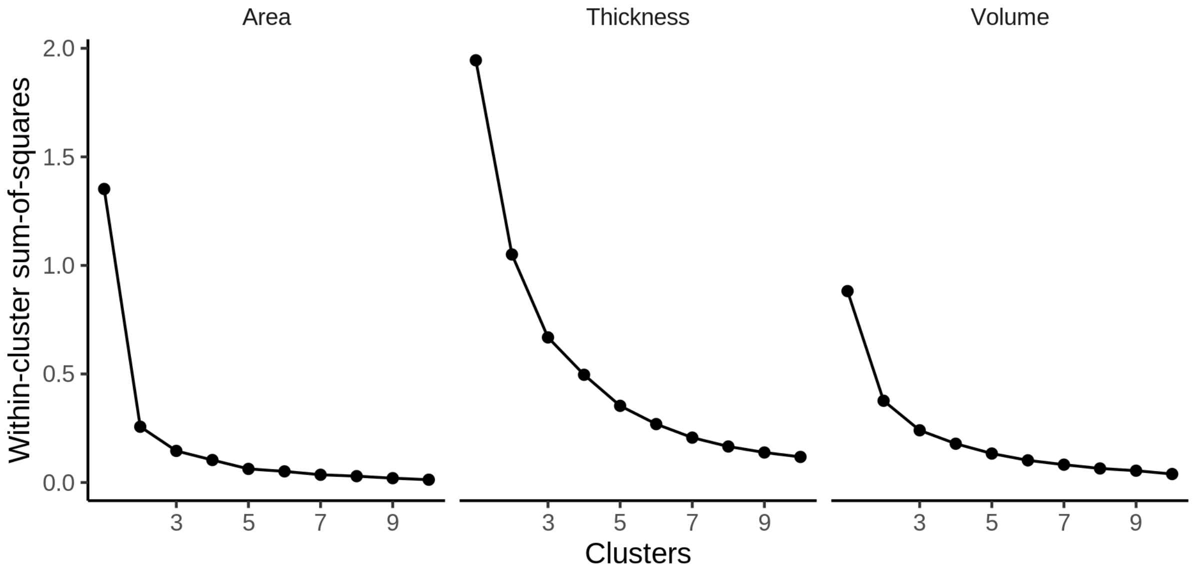


#### Area

The following regions go in each cluster:

- Cluster 1: cuneus, inferiorparietal, lateraloccipital, lingual, pericalcarine, superiorparietal
- Cluster 2: caudalmiddlefrontal, lateralorbitofrontal, middletemporal, parsorbitalis
- Cluster 3: bankssts, caudalanteriorcingulate, entorhinal, frontalpole, fusiform, inferiortemporal, insula, isthmuscingulate, medialorbitofrontal, paracentral, parahippocampal, parsopercularis, parstriangularis, postcentral, posteriorcingulate, precentral, precuneus, rostralanteriorcingulate, rostralmiddlefrontal, superiorfrontal, superiortemporal, supramarginal, transversetemporal

#### Description of clusters

First are spaghetti plots for the three clusters.

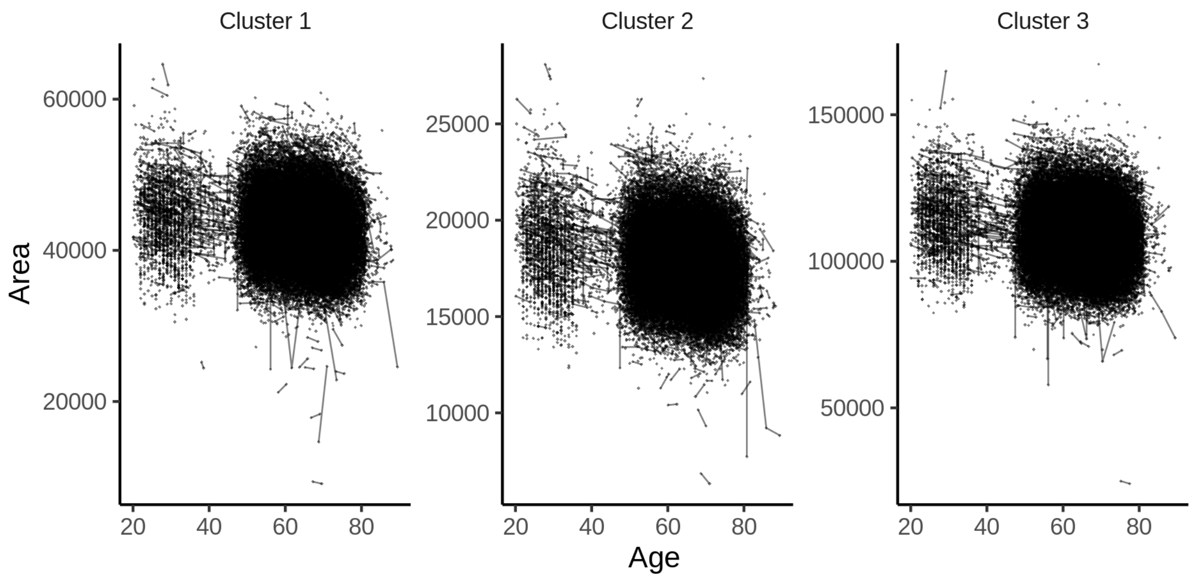


The plot below shows the sleep-volume curves for the three clusters.

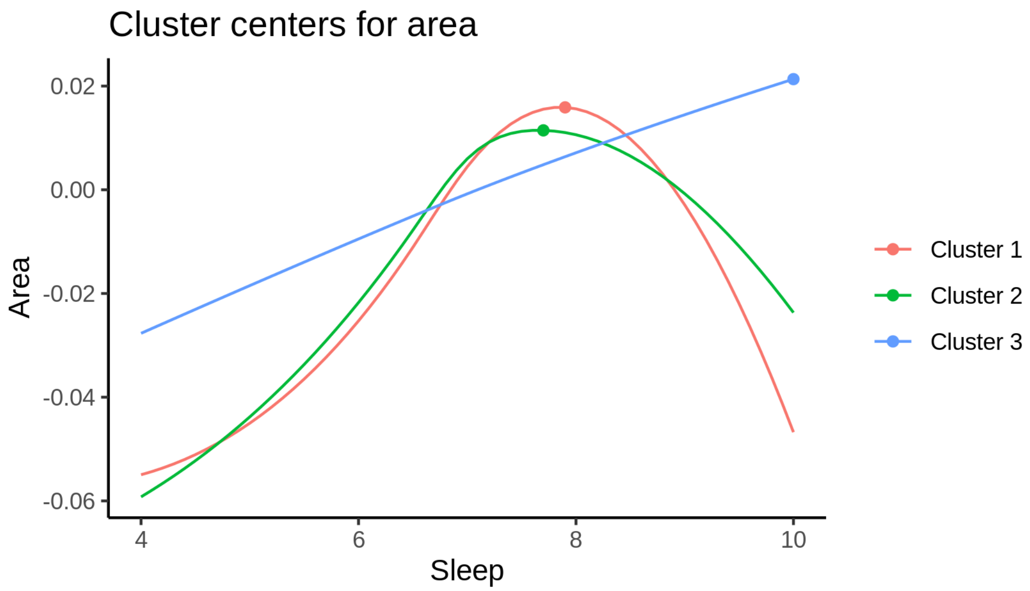


The maximum values for each cluster occur at the following sleep durations:

| Cluster | Sleep |
|---|---|
| Cluster 1 | 7.9 |
| Cluster 2 | 7.7 |
| Cluster 3 | 10.0 |

The next plot shows the effect of varying the sleep duration as a percentage of the maximum value, obtained at the top of the curve. We have to scrutinize these percentages a bit, because actually defining the mean and standard deviations for each cluster turned out to be not that straightforward, but I believe that what I did makes sense. The problem is that the original clustering is done on a Z-transformed scale, since we are interested in differences in shapes of the curves, and not differences in absolute values of area/thickness/volume between the regions. Then I had to define conversion factors by weighting the mean and variances of each region in each cluster, and try to scale it back.

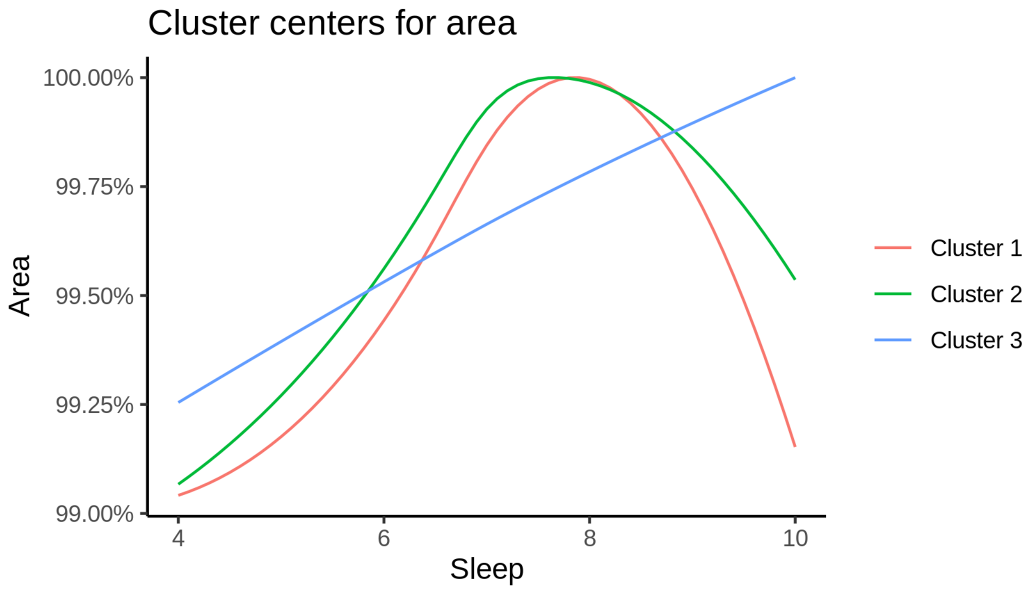


For completeness, we also show the cluster curves upon which the percentage plot above was based. Note that the values differ between clusters, since their sizes are different.

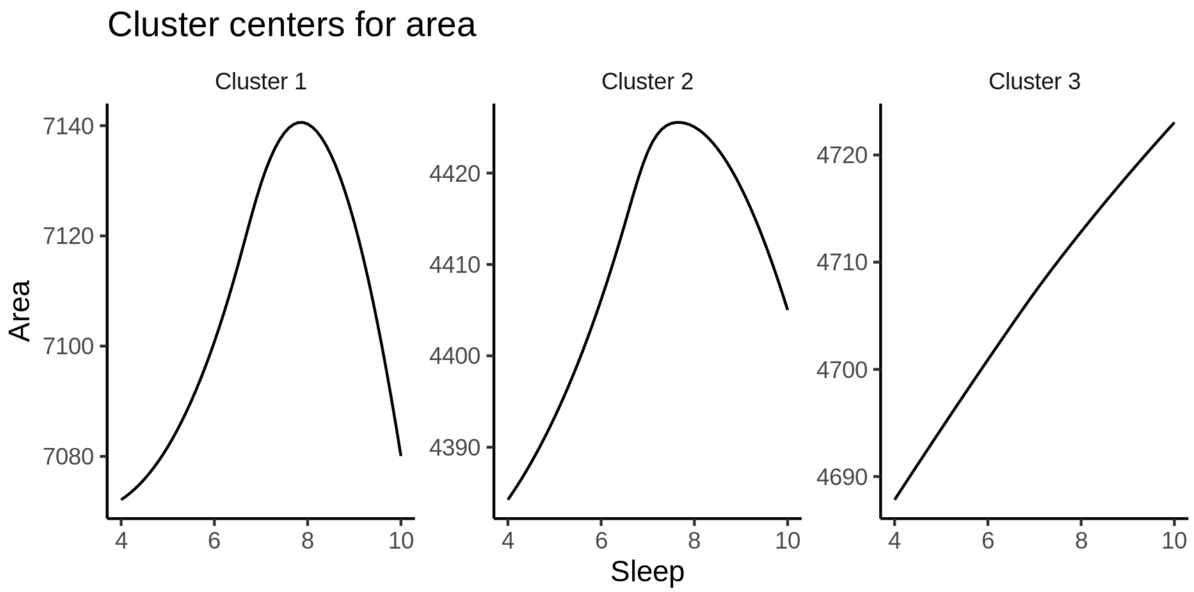


The next plot shows the location of the clusters.

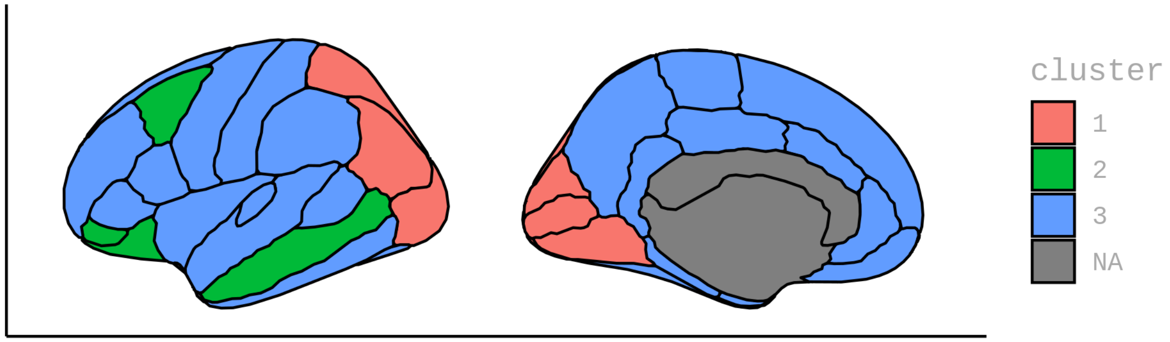


##### GAMM fits in each cluster

The plot below shows the results of fitting GAMMs to the (weighted) average in each cluster.

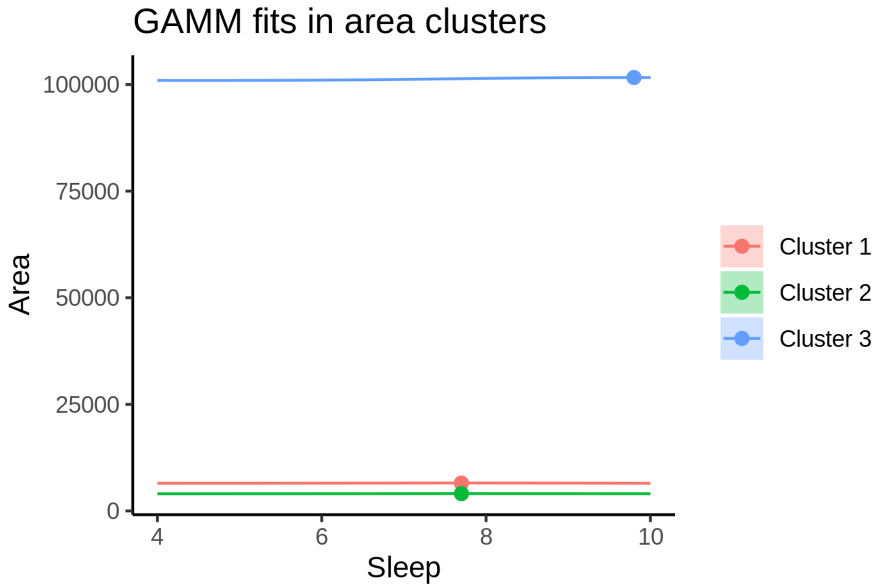


The next plot shows the same thing with percentages of cluster maximum.

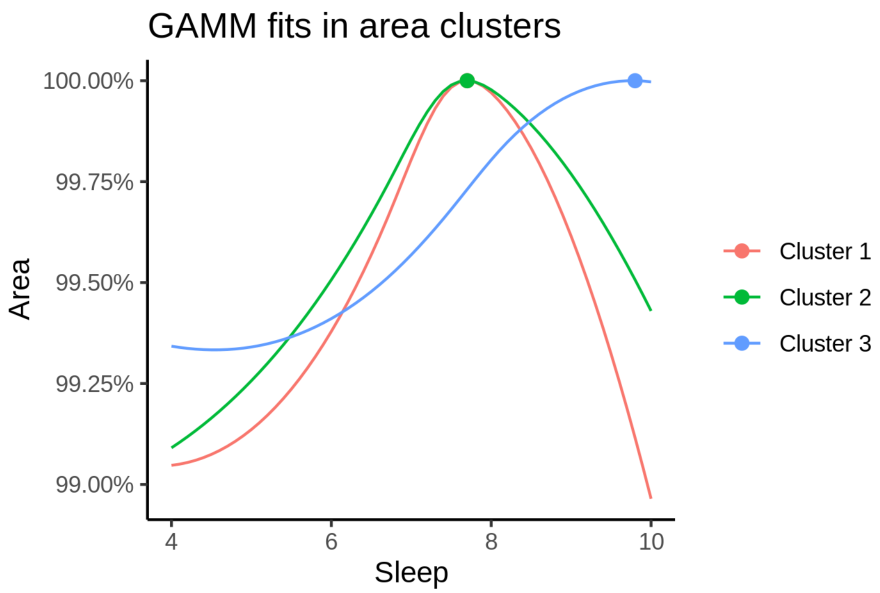


The table below shows the sleep duration associated with maximum value in each cluster, with 95% CIs.

| Sleep at max. (95% CI) |
|---|
| 7.7 (7.5, 8.1) |
| 7.7 (7.5, 8.2) |
| 9.8 (8.7, 10) |

#### Thickness

The following regions go in each cluster:

- Cluster 1: caudalanteriorcingulate, cuneus, isthmuscingulate, lateraloccipital, lingual, parsorbitalis, pericalcarine, precuneus, superiorparietal
- Cluster 2: bankssts, caudalmiddlefrontal, entorhinal, fusiform, inferiorparietal, inferiortemporal, lateralorbitofrontal, middletemporal, parstriangularis, postcentral, precentral, rostralmiddlefrontal, superiorfrontal, superiortemporal, supramarginal, transversetemporal
- Cluster 3: frontalpole, insula, medialorbitofrontal, paracentral, parahippocampal, parsopercularis, posteriorcingulate, rostralanteriorcingulate

##### Description of clusters

First are spaghetti plots for the three clusters.

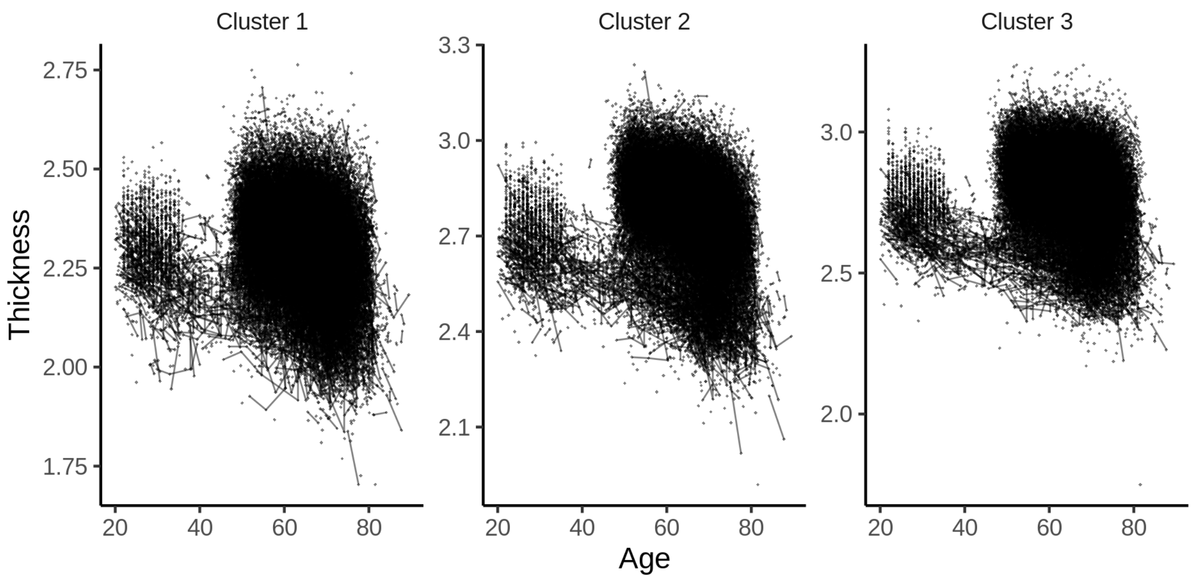


The plot below shows the sleep-volume curves for the three clusters.

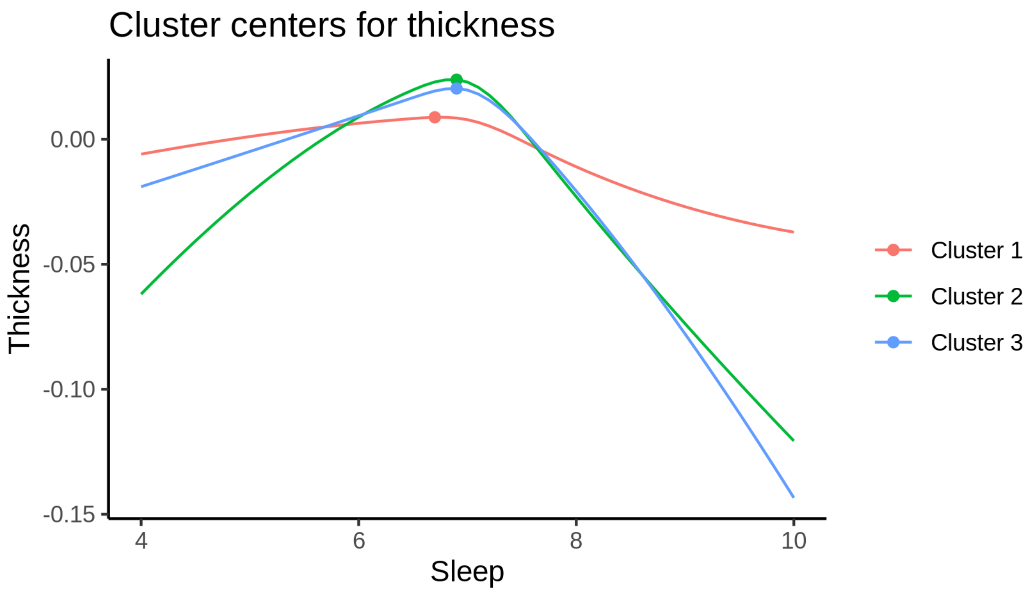


The maximum values for each cluster occur at the following sleep durations:

| Cluster | Sleep |
|---|---|
| Cluster 1 | 6.7 |
| Cluster 2 | 6.9 |
| Cluster 3 | 6.9 |

The next plot shows the effect of varying the sleep duration as a percentage of the maximum value, obtained at the top of the curve. We have to scrutinize these percentages a bit, because actually defining the mean and standard deviations for each cluster turned out to be not that straightforward, but I believe that what I did makes sense. The problem is that the original clustering is done on a Z-transformed scale, since we are interested in differences in shapes of the curves, and not differences in absolute values of area/thickness/volume between the regions. Then I had to define conversion factors by weighting the mean and variances of each region in each cluster, and try to scale it back.

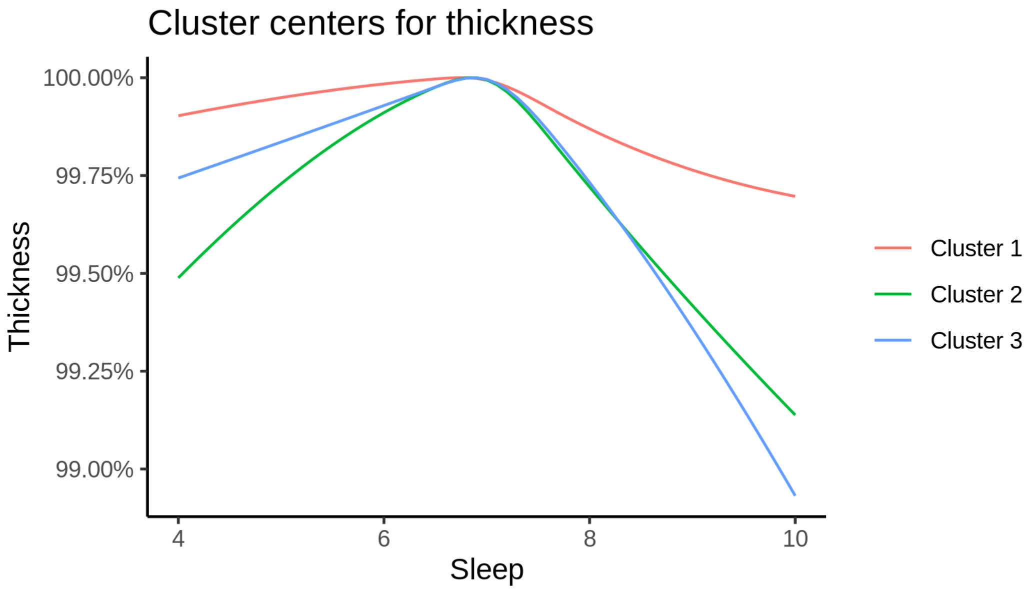


For completeness, we also show the cluster curves upon which the percentage plot above was based.

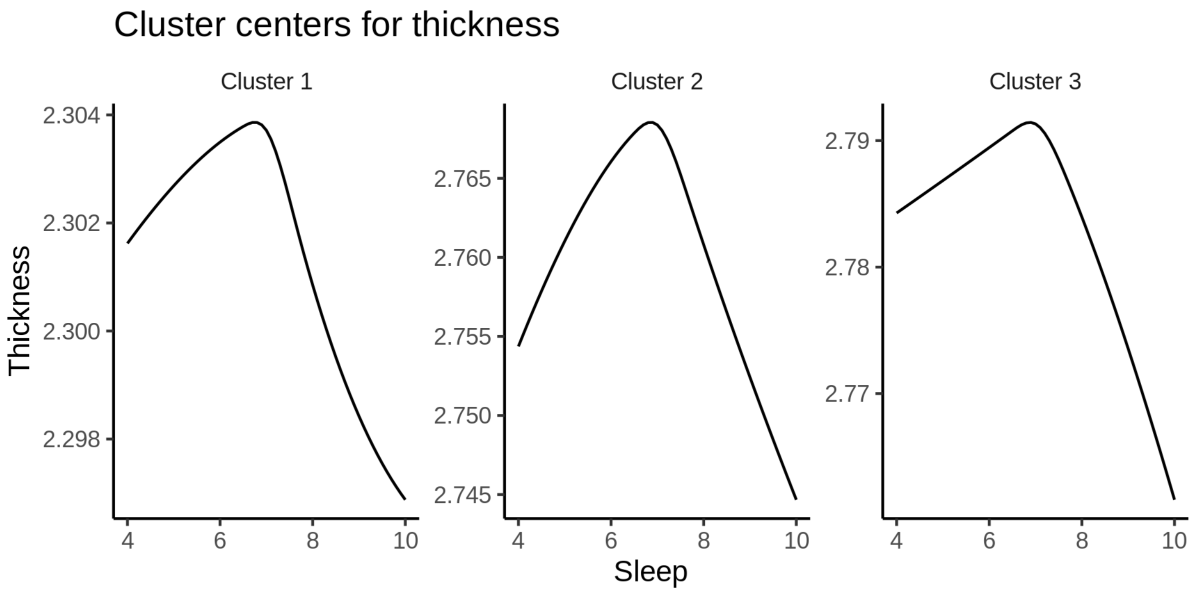


The next plot shows the location of the clusters.

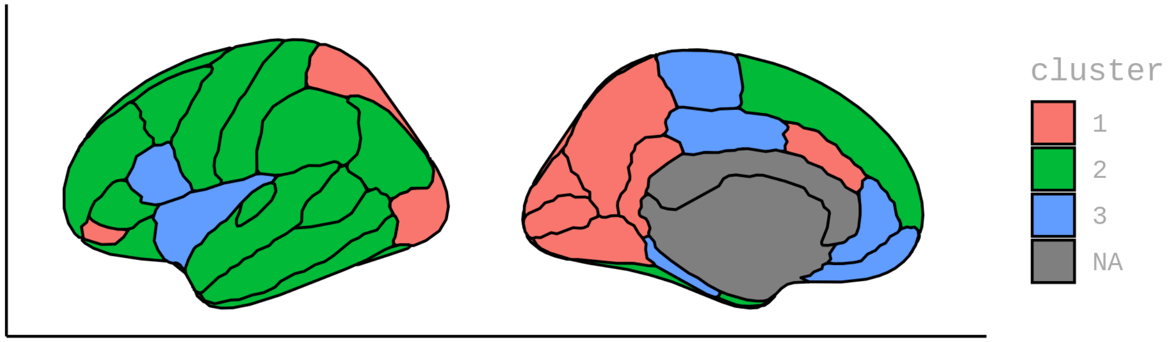


#### GAMM fits in each cluster

The plot below shows the results of fitting GAMMs to the (weighted) average in each cluster.

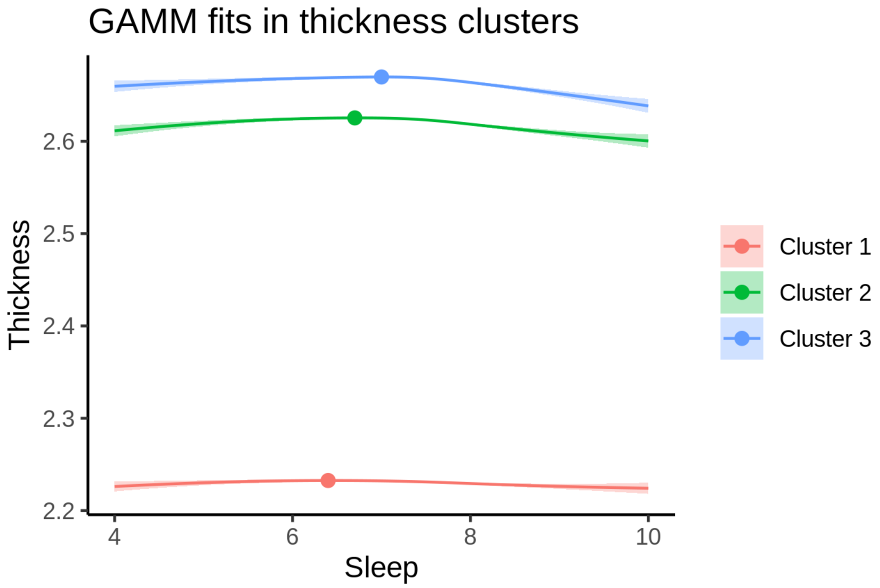


The next plot shows the same thing with percentages of cluster maximum.

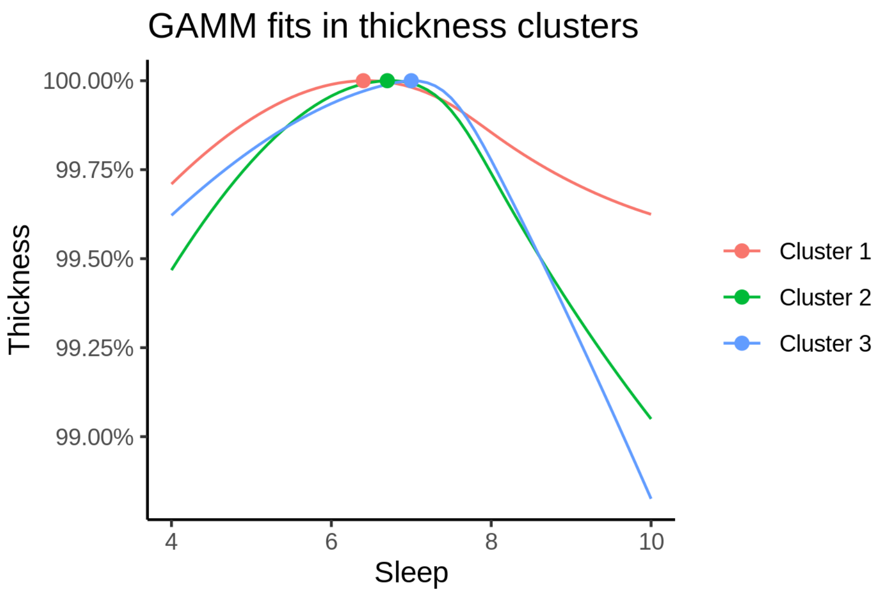


The table below shows the sleep duration associated with maximum value in each cluster, with 95% CIs.

| Sleep at max. (95% CI) |
|---|
| 6.4 (5.7, 7.1) |
| 6.7 (6.3, 7.1) |
| 7 (6.4, 7.2) |

### Volume

The following regions go in each cluster:

- Cluster 1: bankssts, fusiform, inferiortemporal, lateraloccipital, medialorbitofrontal, paracentral, parsopercularis, pericalcarine, postcentral, superiorfrontal, superiorparietal, superiortemporal, supramarginal
- Cluster 2: caudalanteriorcingulate, cuneus, entorhinal, frontalpole, isthmuscingulate, lateralorbitofrontal, lingual, parahippocampal, parsorbitalis, parstriangularis, posteriorcingulate, precuneus, rostralanteriorcingulate, rostralmiddlefrontal, transversetemporal
- Cluster 3: caudalmiddlefrontal, inferiorparietal, insula, middletemporal, precentral

#### Description of clusters

First are spaghetti plots for the three clusters.

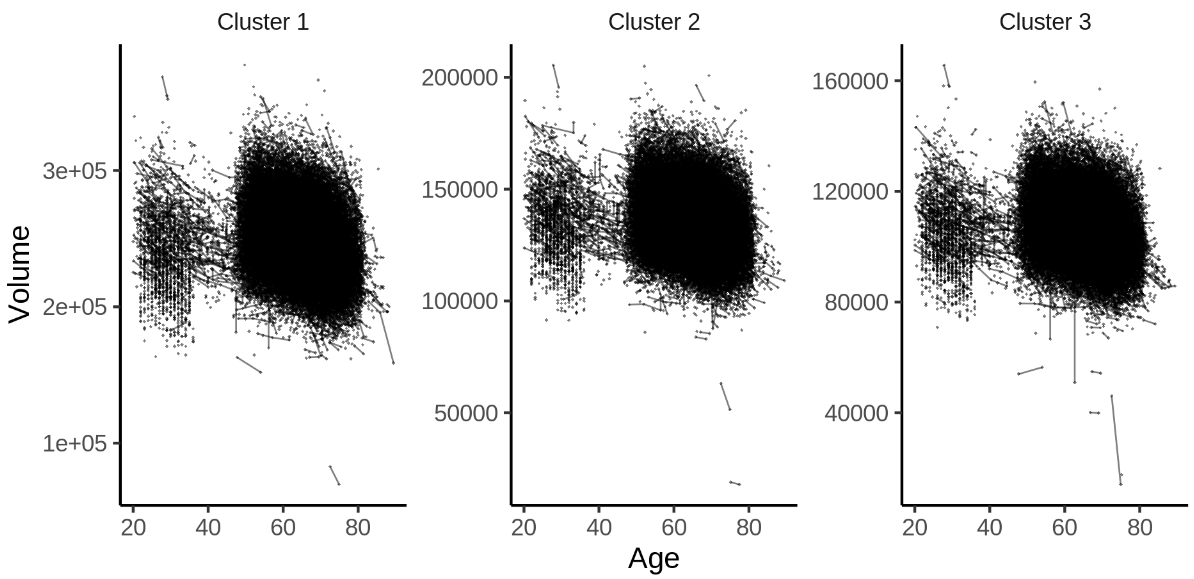


The plot below shows the sleep-volume curves for the three clusters.

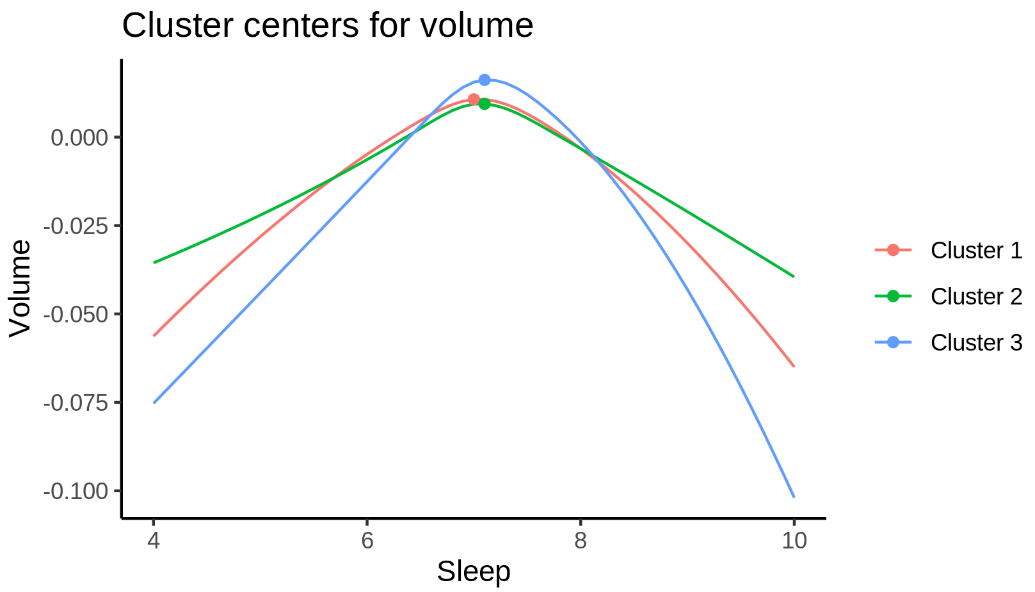


The maximum values for each cluster occur at the following sleep durations:

| Cluster | Sleep |
|---|---|
| Cluster 1 | 7.0 |
| Cluster 2 | 7.1 |
| Cluster 3 | 7.1 |

The next plot shows the effect of varying the sleep duration as a percentage of the maximum value, obtained at the top of the curve. We have to scrutinize these percentages a bit, because actually defining the mean and standard deviations for each cluster turned out to be not that straightforward, but I believe that what I did makes sense. The problem is that the original clustering is done on a Z-transformed scale, since we are interested in differences in shapes of the curves, and not differences in absolute values of area/thickness/volume between the regions. Then I had to define conversion factors by weighting the mean and variances of each region in each cluster, and try to scale it back.

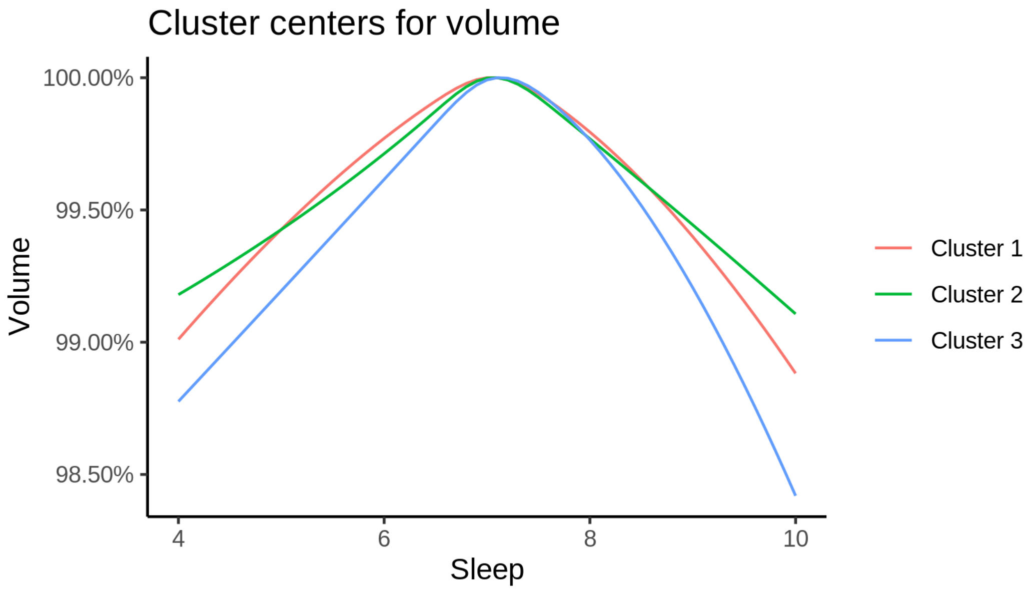


For completeness, we also show the cluster curves upon which the percentage plot above was based. Note that the values differ between clusters, since their sizes are different.

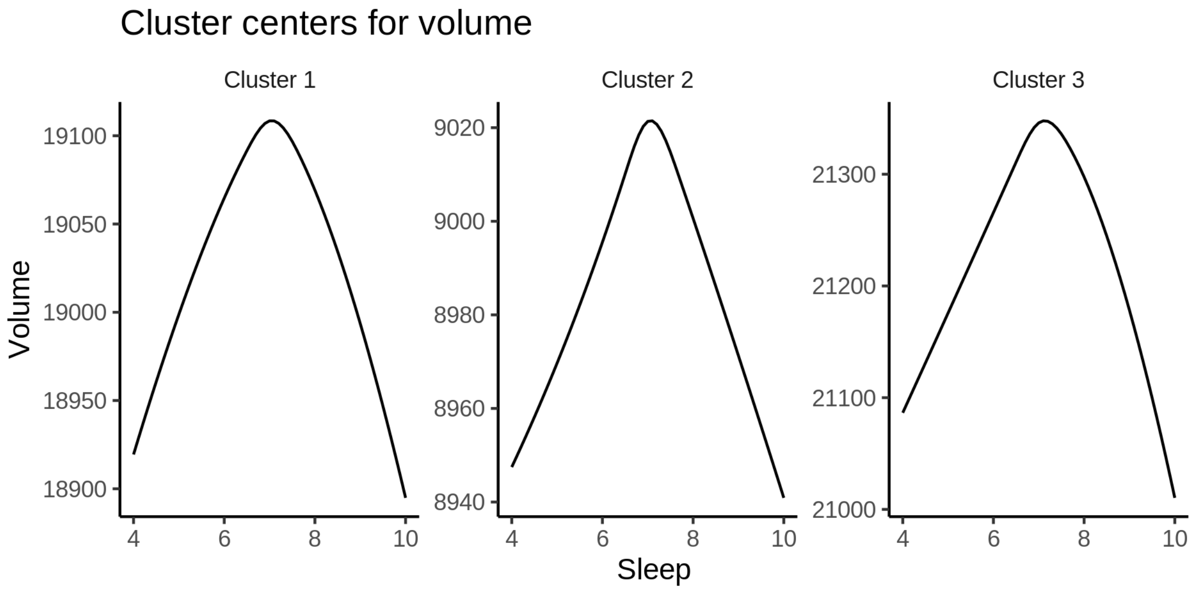


The next plot shows the location of the clusters.

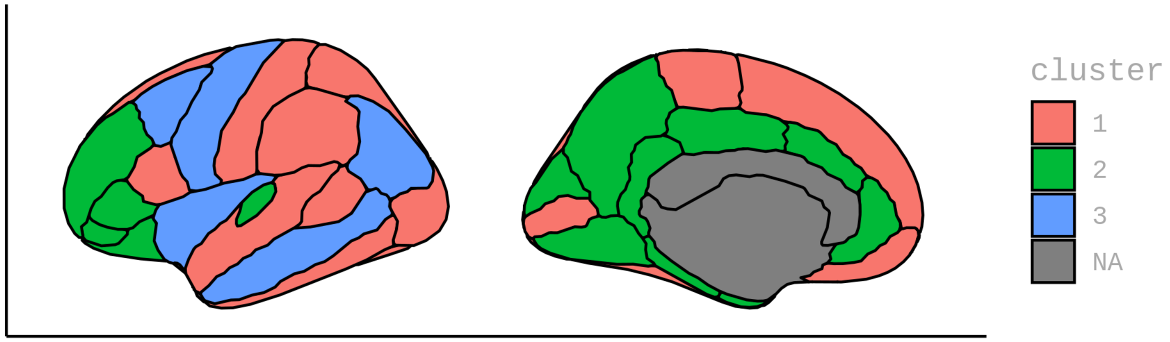


##### GAMM fits in each cluster

The plot below shows the results of fitting GAMMs to the (weighted) average in each cluster.

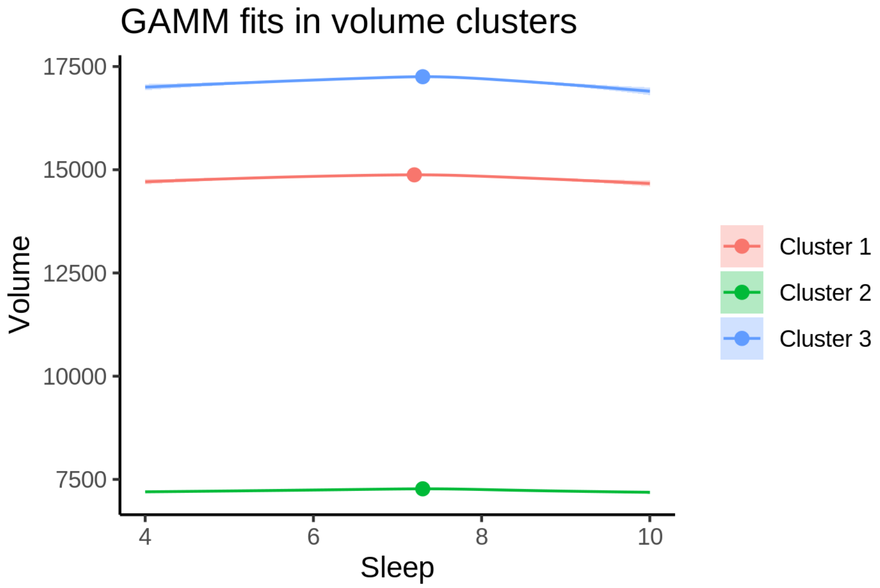


The next plot shows the same thing with percentages of cluster maximum.

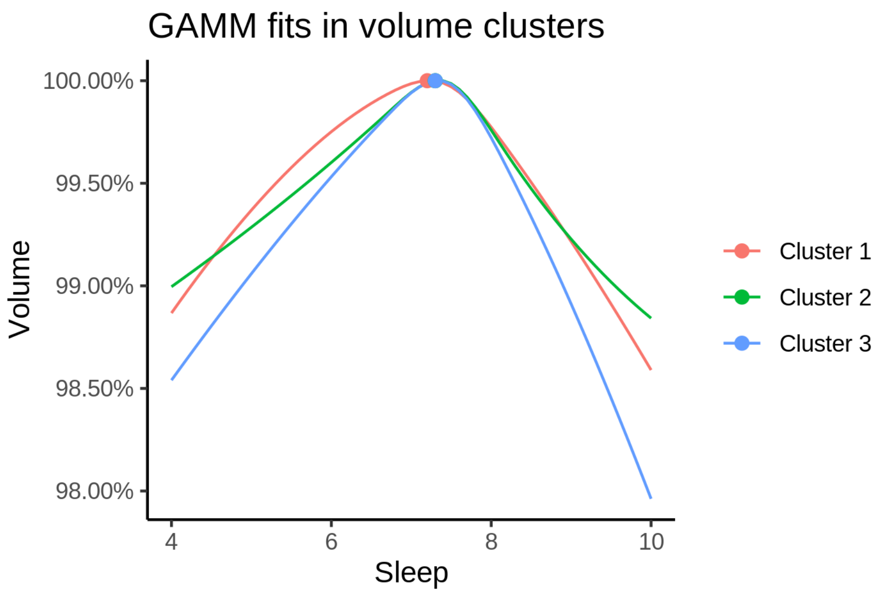


The table below shows the sleep duration associated with maximum value in each cluster, with 95% CIs.

| Sleep at max. (95% CI) |
|---|
| 7.2 (6.9, 7.4) |
| 7.3 (7.2, 7.4) |
| 7.3 (7.2, 7.4) |
