## Supplementary material for "Sleep duration and brain structure – phenotypic associations and genotypic covariance": SI Cortical surface analyses

The table shows the number of observations for each of the groups.

| Age group | Sleep group | n |
|---|---|---|
| Age 20-60 | Sleep above mean | 6421 |
| Age 20-60 | Sleep below mean | 9464 |
| Age above 60 | Sleep above mean | 15053 |
| Age above 60 | Sleep below mean | 17119 |

All plots show p-values for the effect of sleep, corrected with 10,000 Z Monte Carlo simulations. The cluster-forming p-value threshold was 0.05 (arguments to `mri_glmfit-sim` were `--cache 1.3 abs --cwp 0.01`).

#### Area

##### Left hemisphere

###### Age 20-60

**Sleep above mean** Cluster corrected significance plots are shown below.

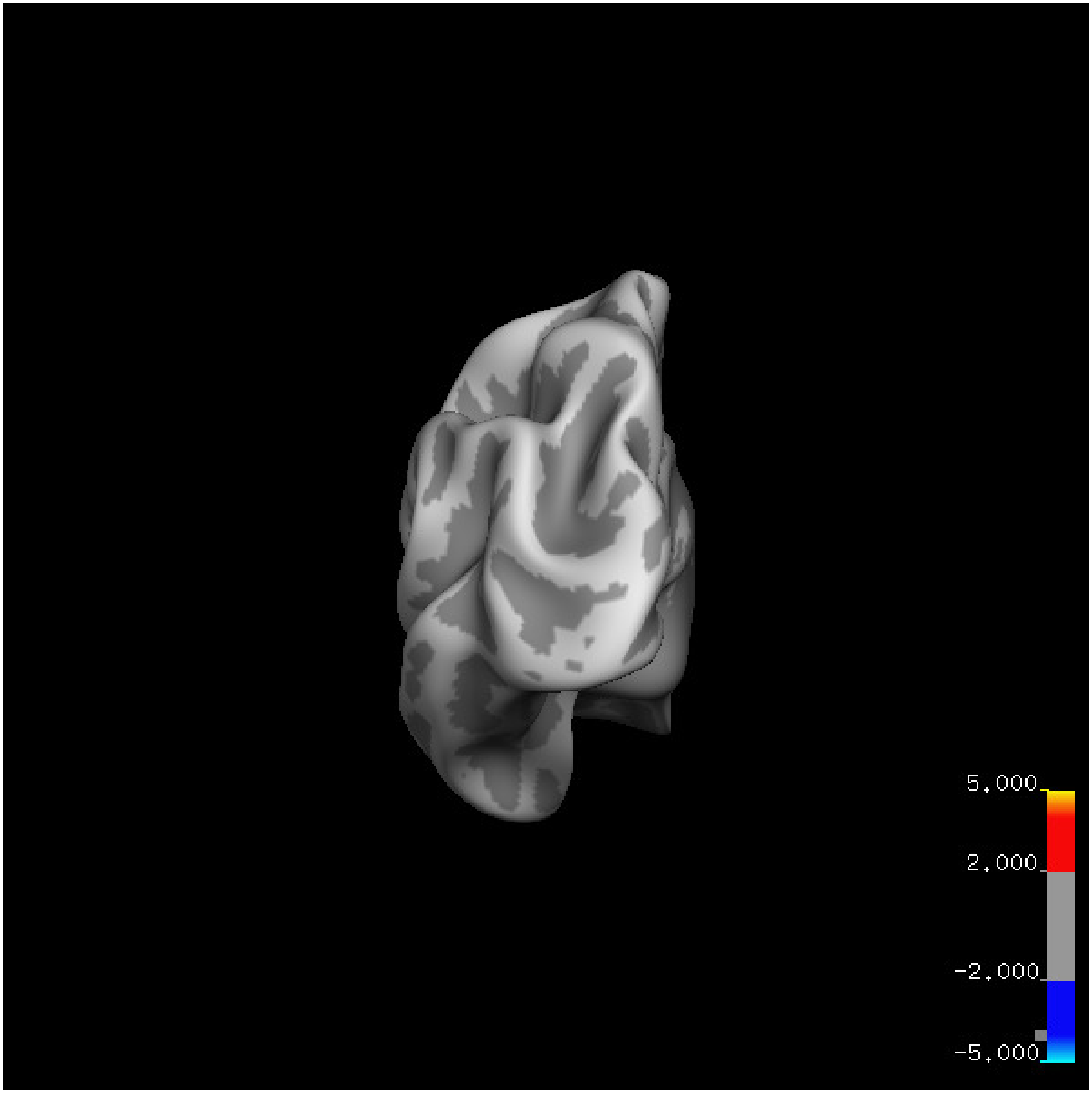
5.000
2.000
-2.000
-5.000

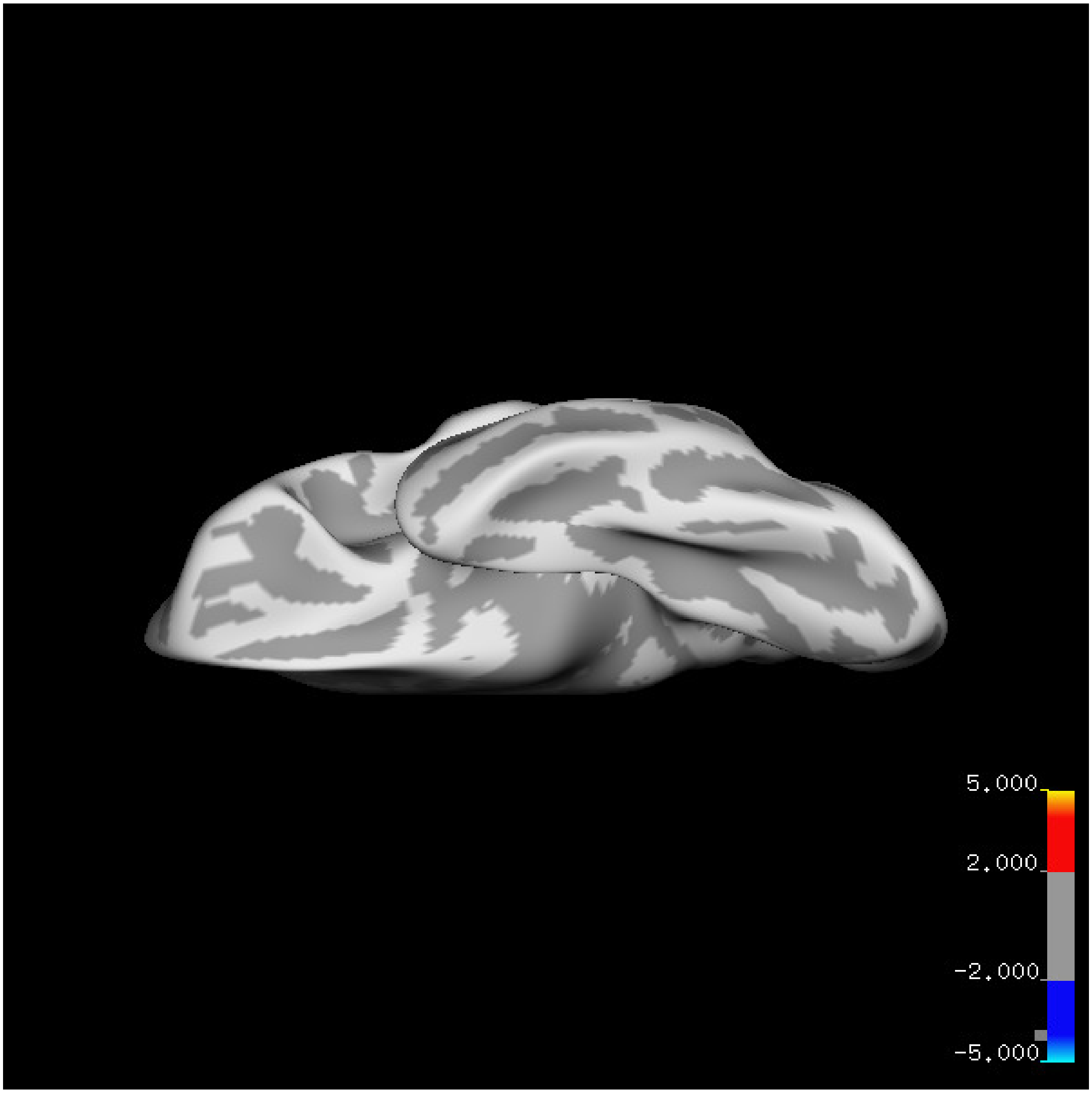
5.000
2.000
-2.000
-5.000

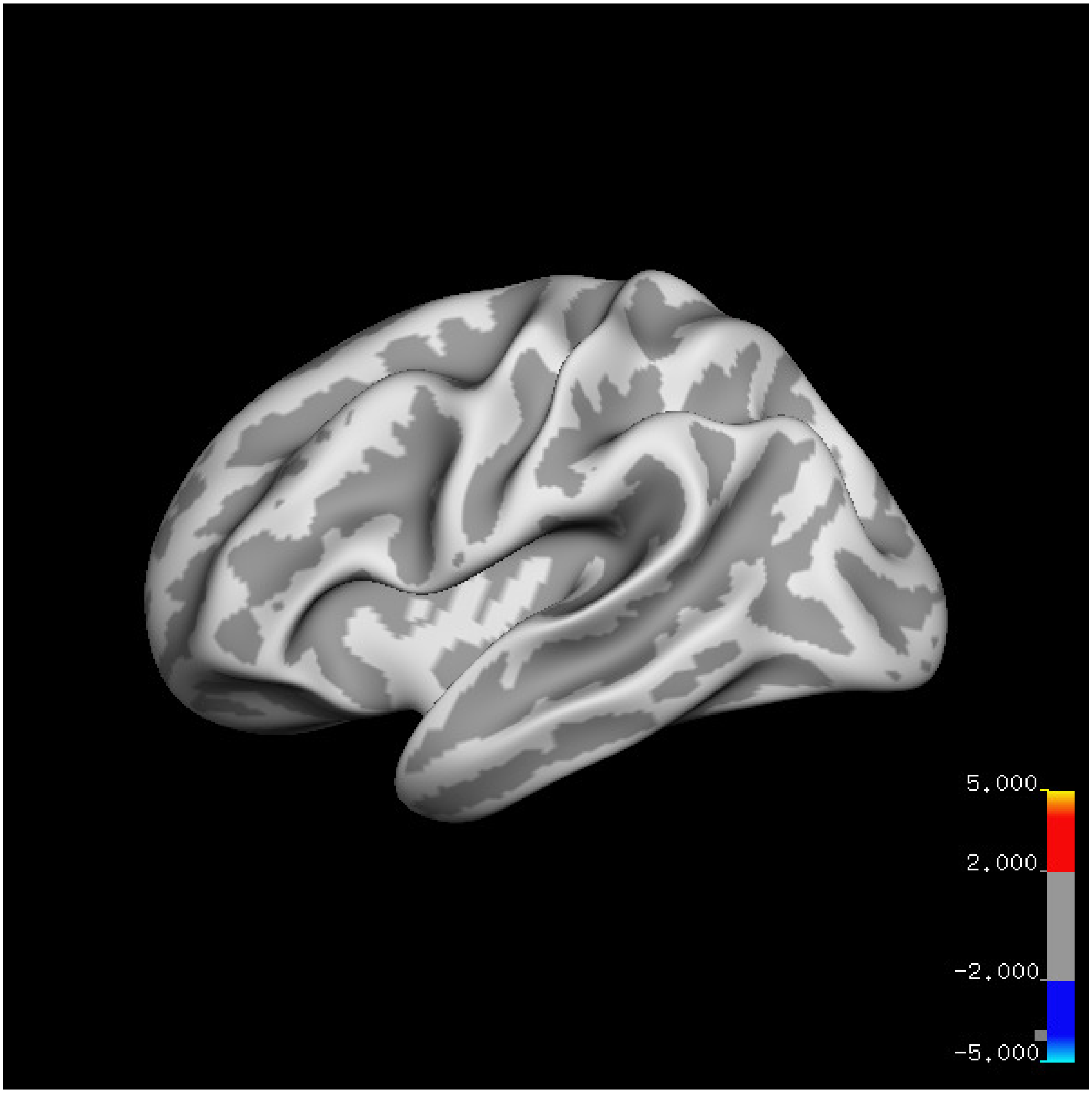
5.000
2.000
-2.000
-5.000

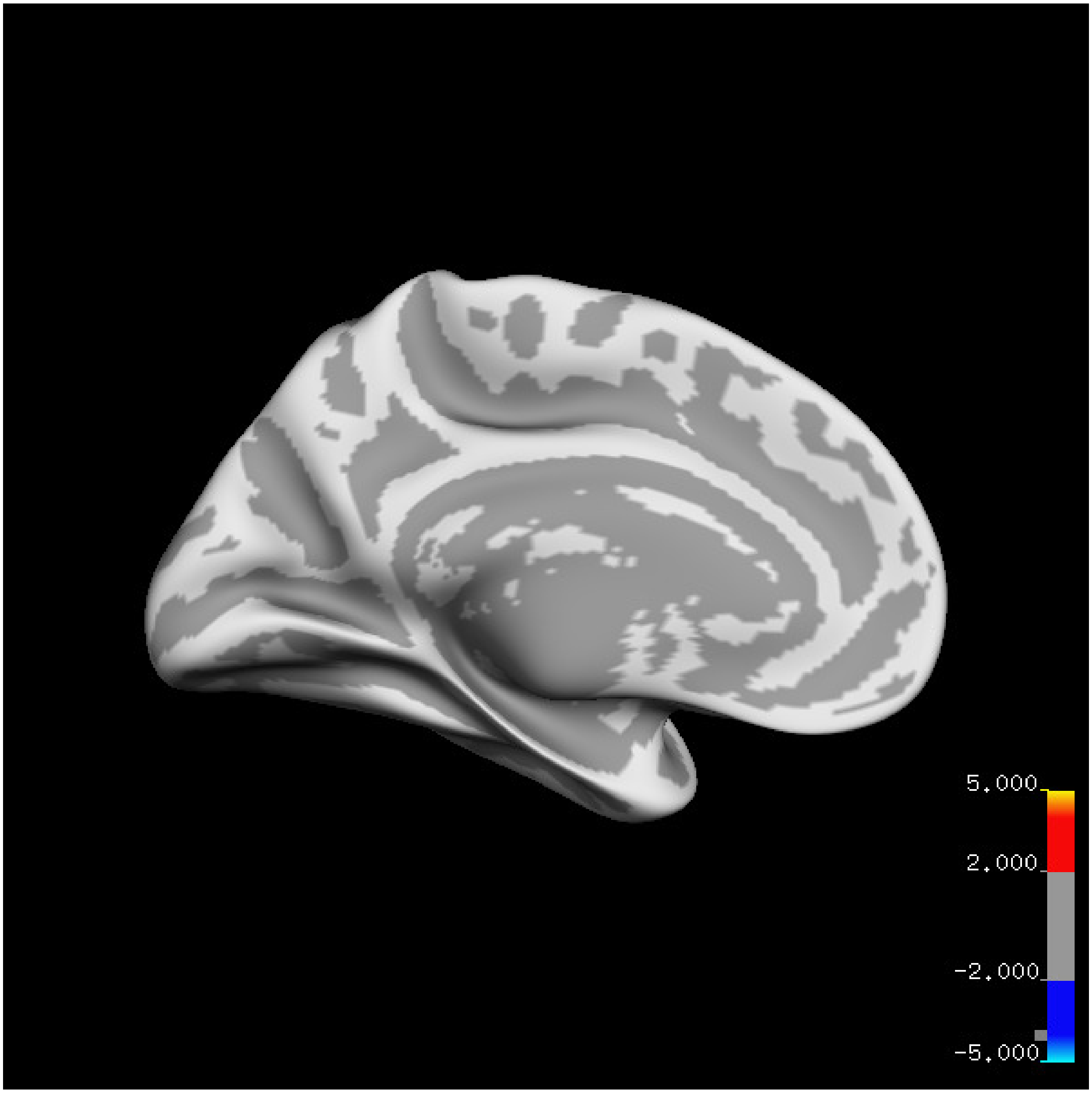

5.000
2.000
-2.000
-5.000

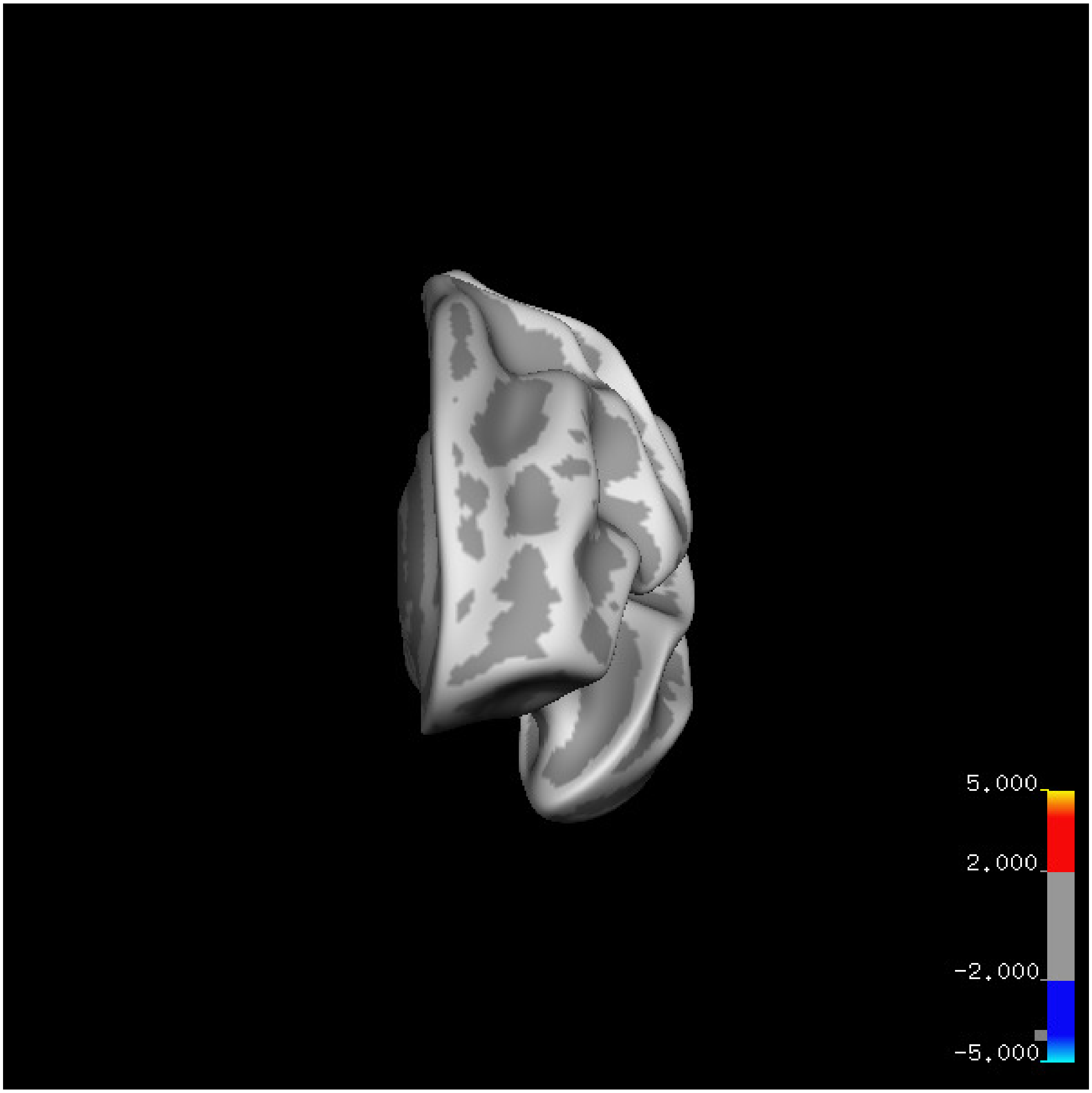
5.000
2.000
-2.000
-5.000

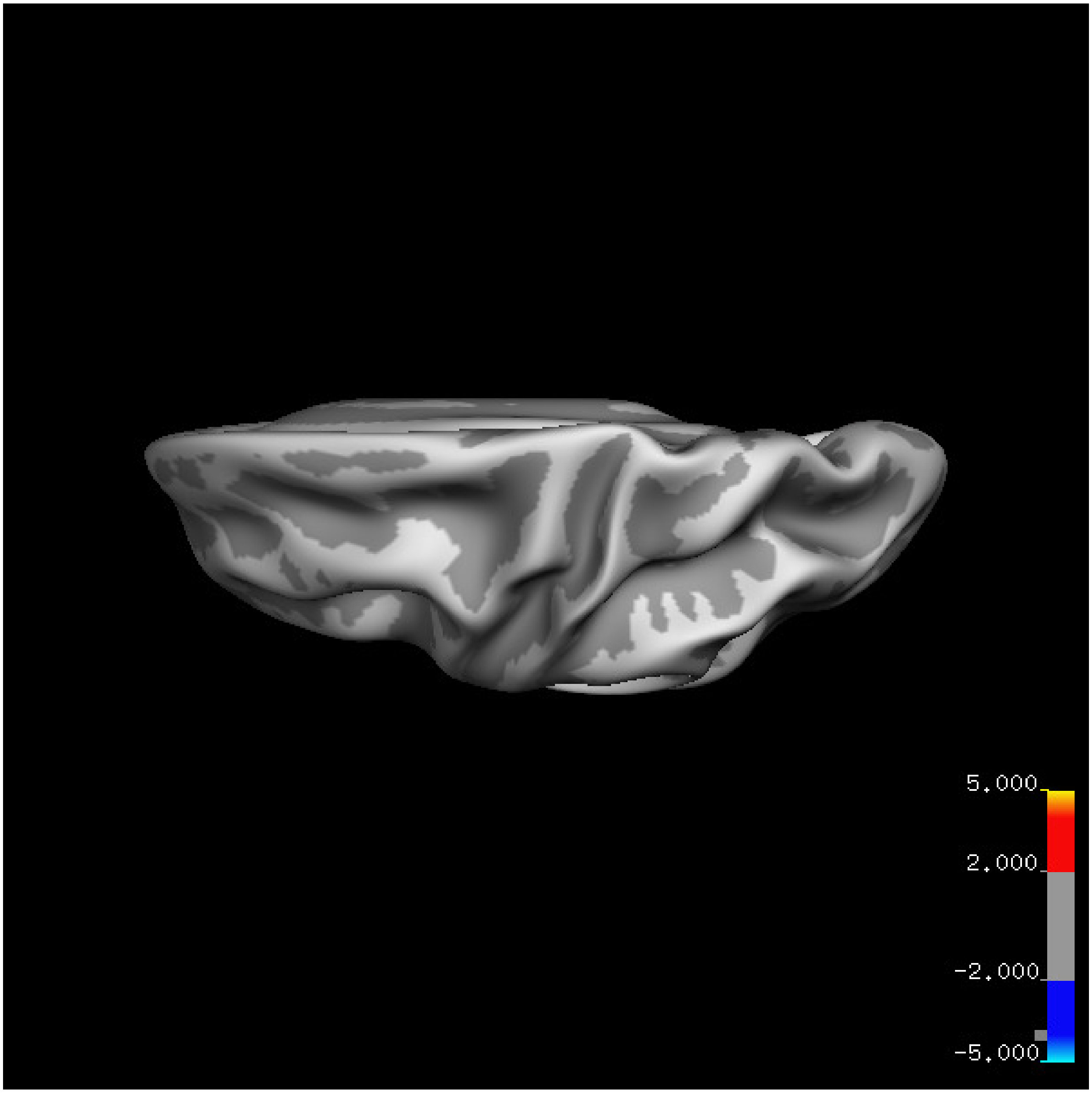
5.000
2.000
-2.000
-5.000

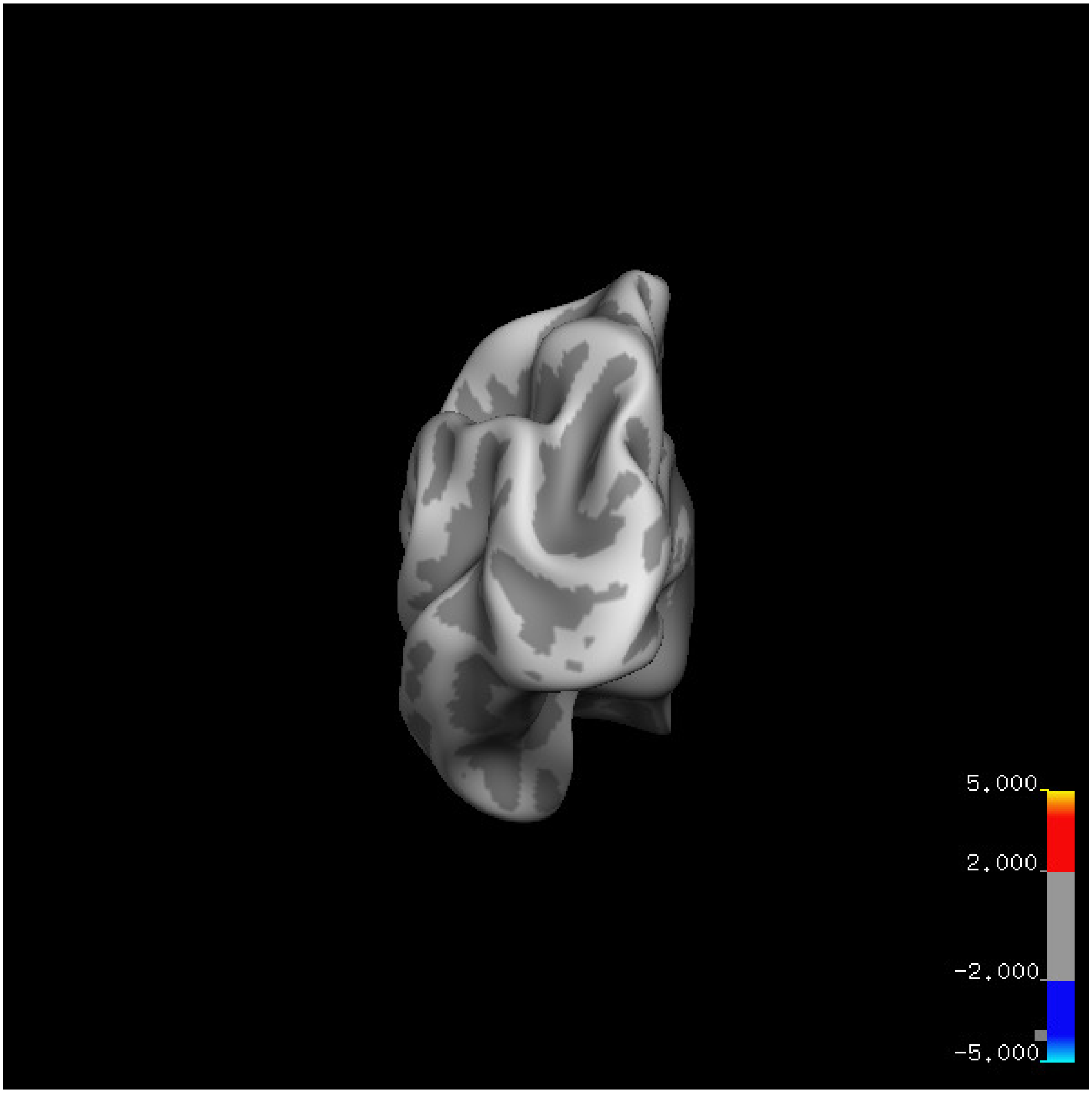

5.000
2.000
-2.000
-5.000

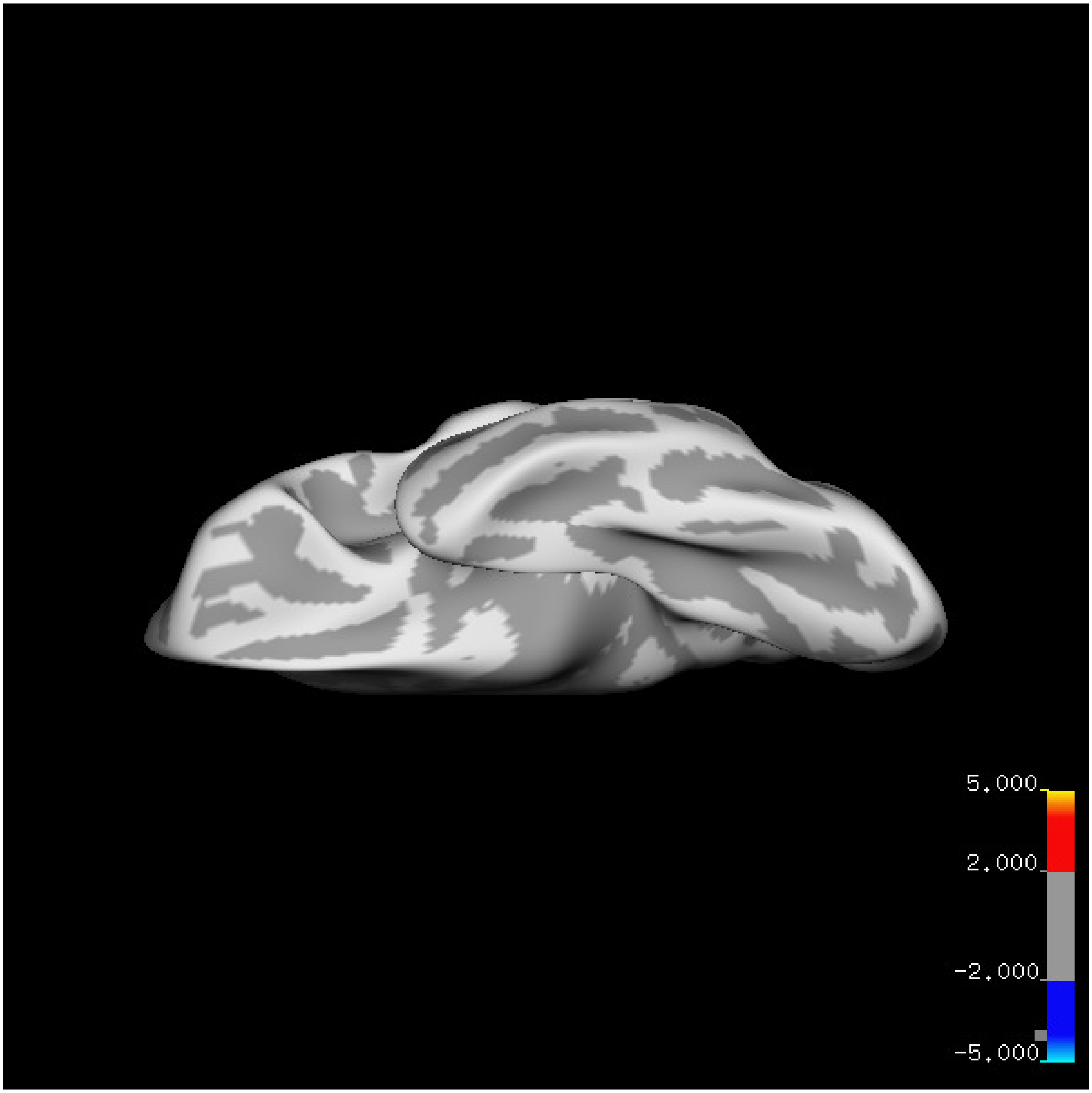
5.000
2.000
-2.000
-5.000

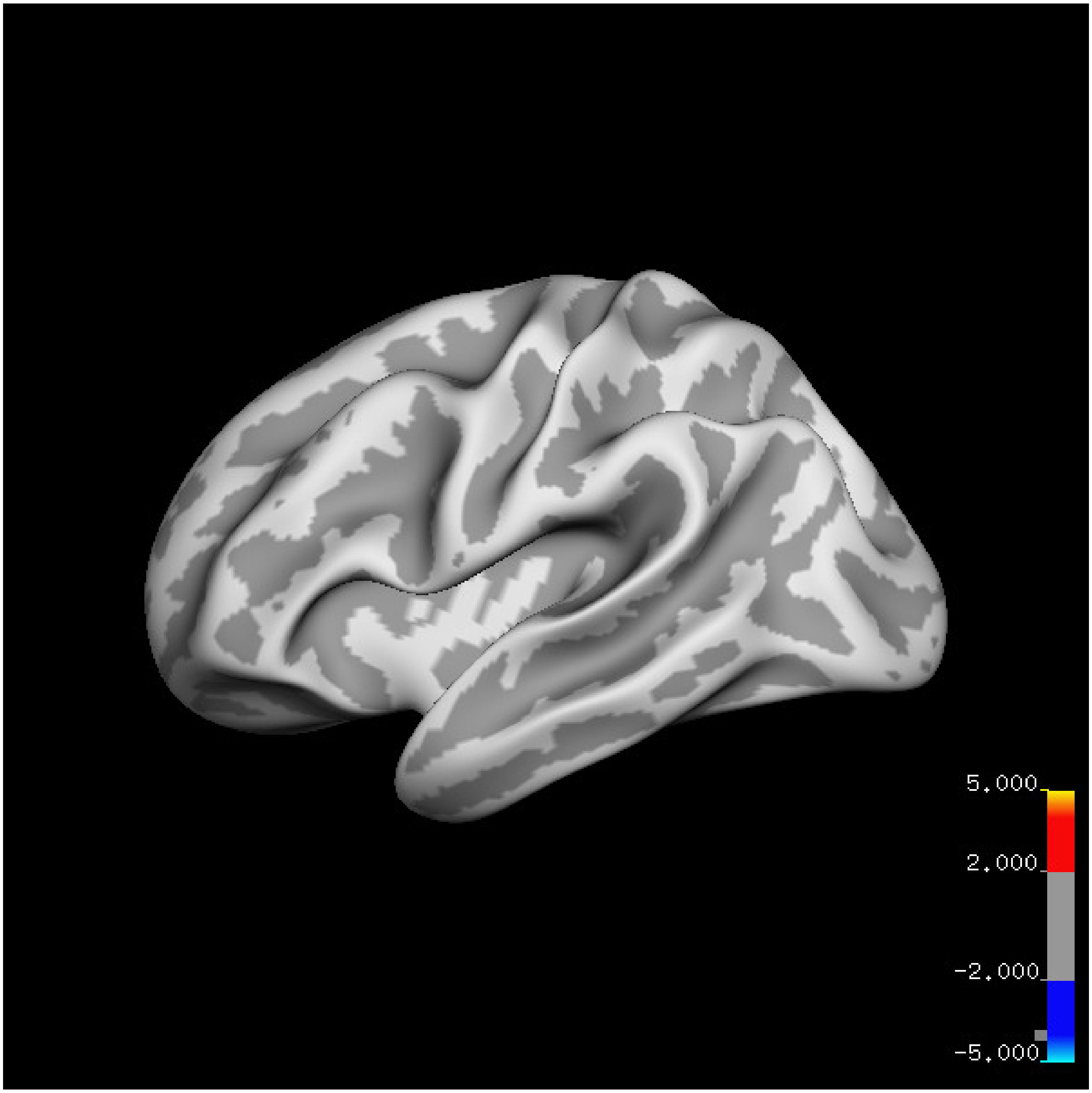
5.000
2.000
-2.000
-5.000

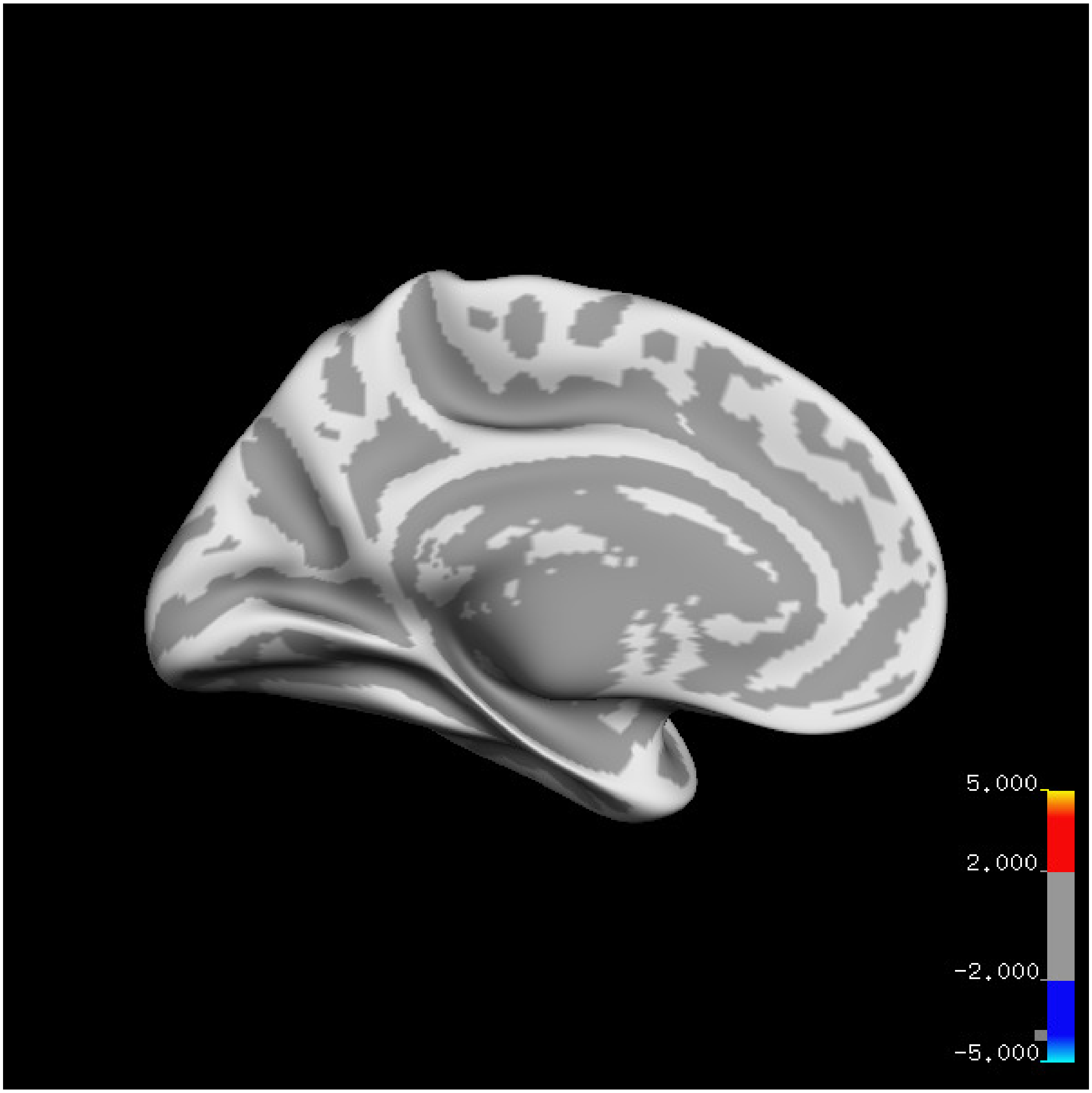
5.000
2.000
-2.000
-5.000

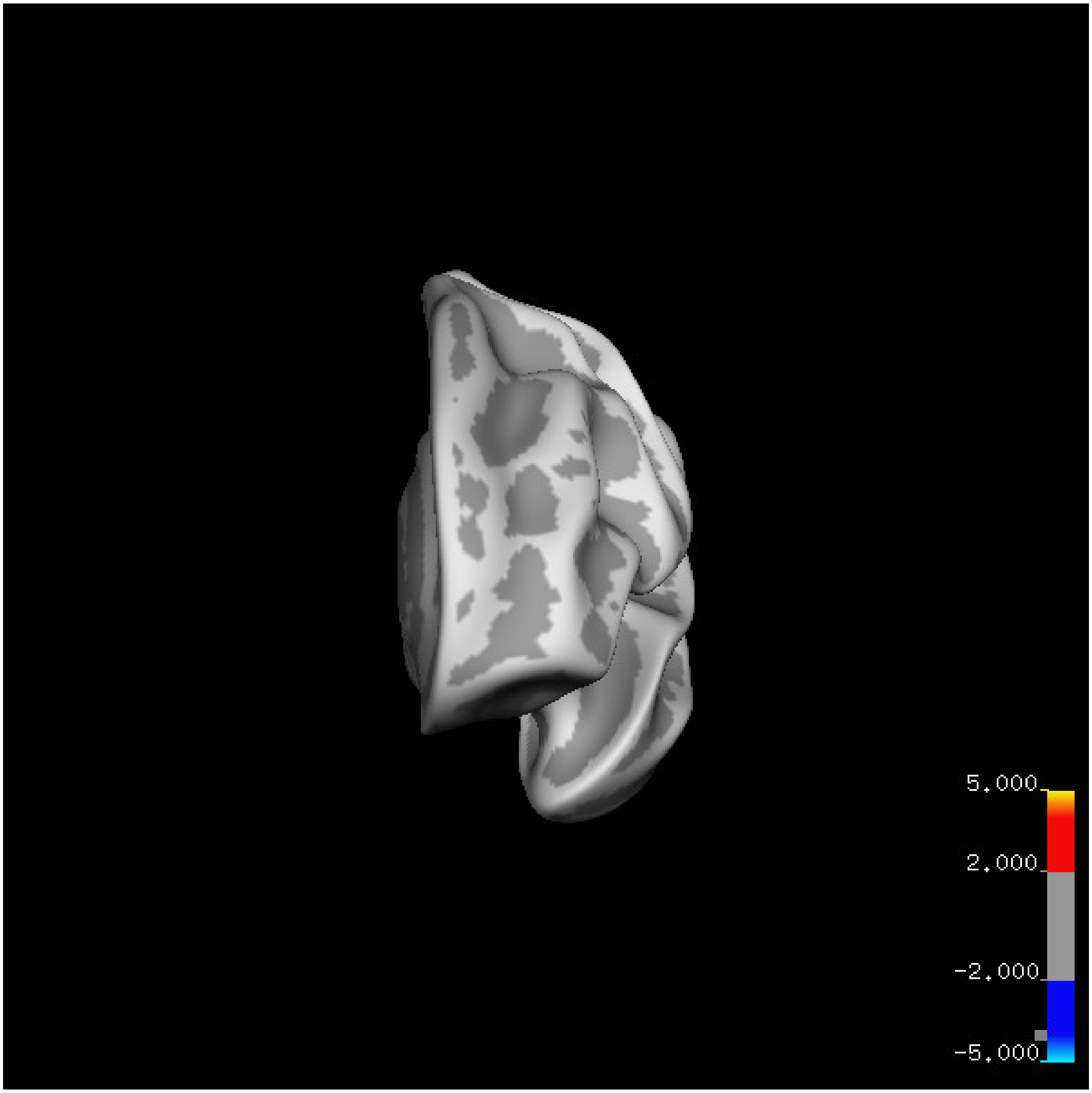
5.000
2.000
-2.000
-5.000

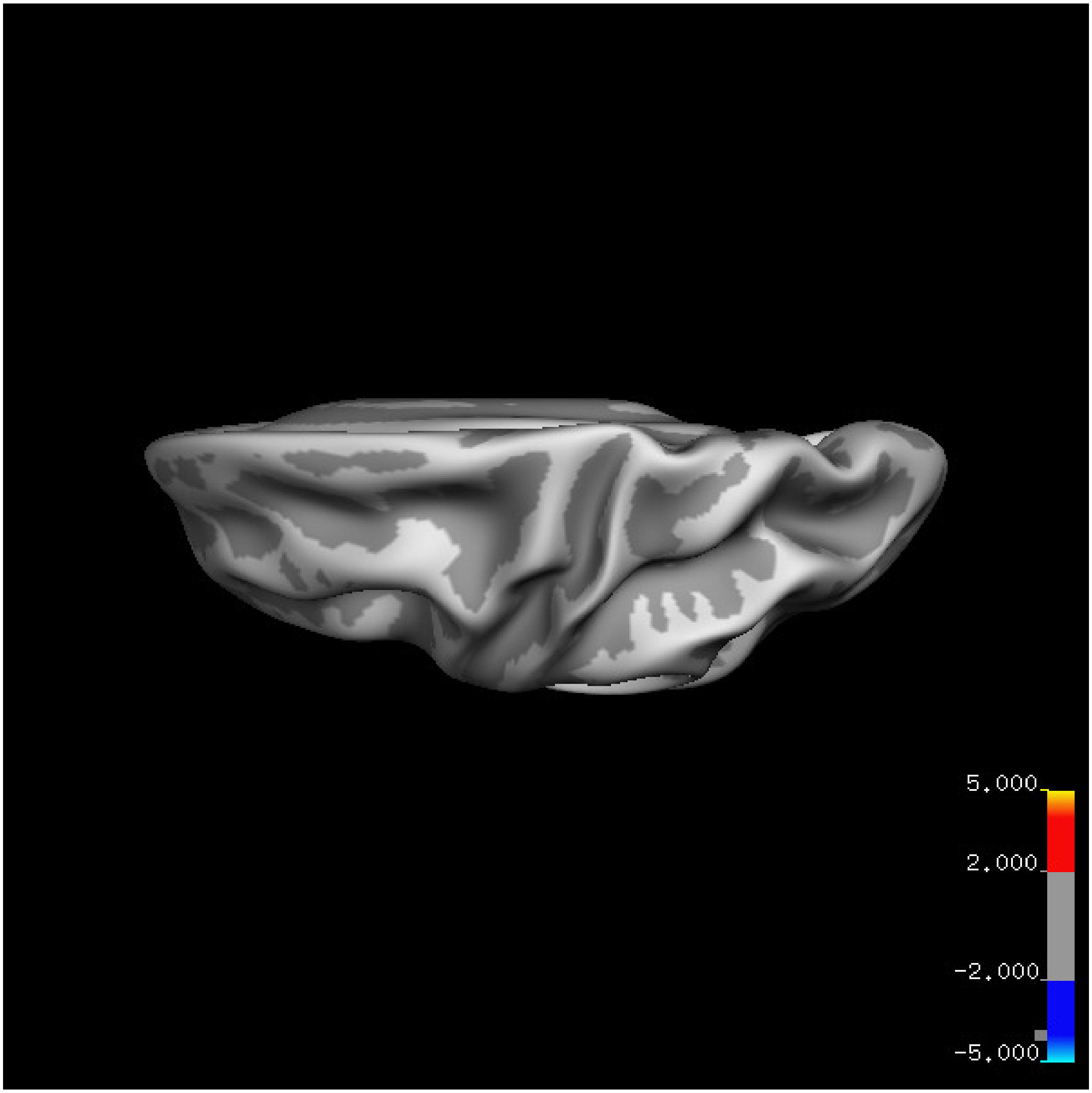


Cluster summary is shown below.

```
## # Cluster Growing Summary (mri_surfcluster)

## # $Id: mri_surfcluster.c,v 1.57.2.3 2016/11/17 18:19:42 zkaufman Exp $

## # $Id: mrisurf.c,v 1.781.2.6 2016/12/27 16:47:14 zkaufman Exp $

## # CreationTime 2021/10/04-10:48:37-GMT

## # cmdline mri_surfcluster.bin --in surface_analyses/results/cluster_comparison/fsaverage6.sm15.lh/ar

## # cwd /gpfs/projects02/p274/cluster/projects/p003-sleep_b

## # sysname  Linux
```

```
## # hostname c1-18

## # machine  x86_64

## # FixVertexAreaFlag 1

## # FixSurfClusterArea 1

## #

## # Input      surface_analyses/results/cluster_comparison/fsaverage6.sm15.lh/area.20_60.above_mean/gl[illegible]

## # Frame Number      0

## # srcsubj fsaverage6

## # hemi lh

## # surface white

## # group_avg_surface_area 84969.3

## # group_avg_vtxarea_loaded 1

## # annot aparc

## # SUBJECTS_DIR /cluster/projects/p274/tools/mri/freesurfer/current/subjects

## # SearchSpace_mm2 79241.3

## # SearchSpace_vtx 37476

## # Bonferroni 0

## # Minimum Threshold 1.3

## # Maximum Threshold infinity

## # Threshold Sign    abs

## # AdjustThreshWhenOneTail 1

## # CW PValue Threshold: 0.01

## # Area Threshold    0 mm^2

## # CSD thresh  1.300000

## # CSD nreps    10000

## # CSD simtype  null-z

## # CSD contrast NA
```

```
## # CSD confint  90.000000

## # Overall max 1.72739 at vertex 15683

## # Overall min -2.54553 at vertex 22175

## # NClusters          0

## # FixMNI = 0

## #

## # ClusterNo  Max   VtxMax   Size(mm^2)  MNIX   MNIY   MNIZ    CWP    CWPLow    CWPHi   NVtxs    Wght
```

**Sleep below mean** Cluster corrected significance plots are shown below.

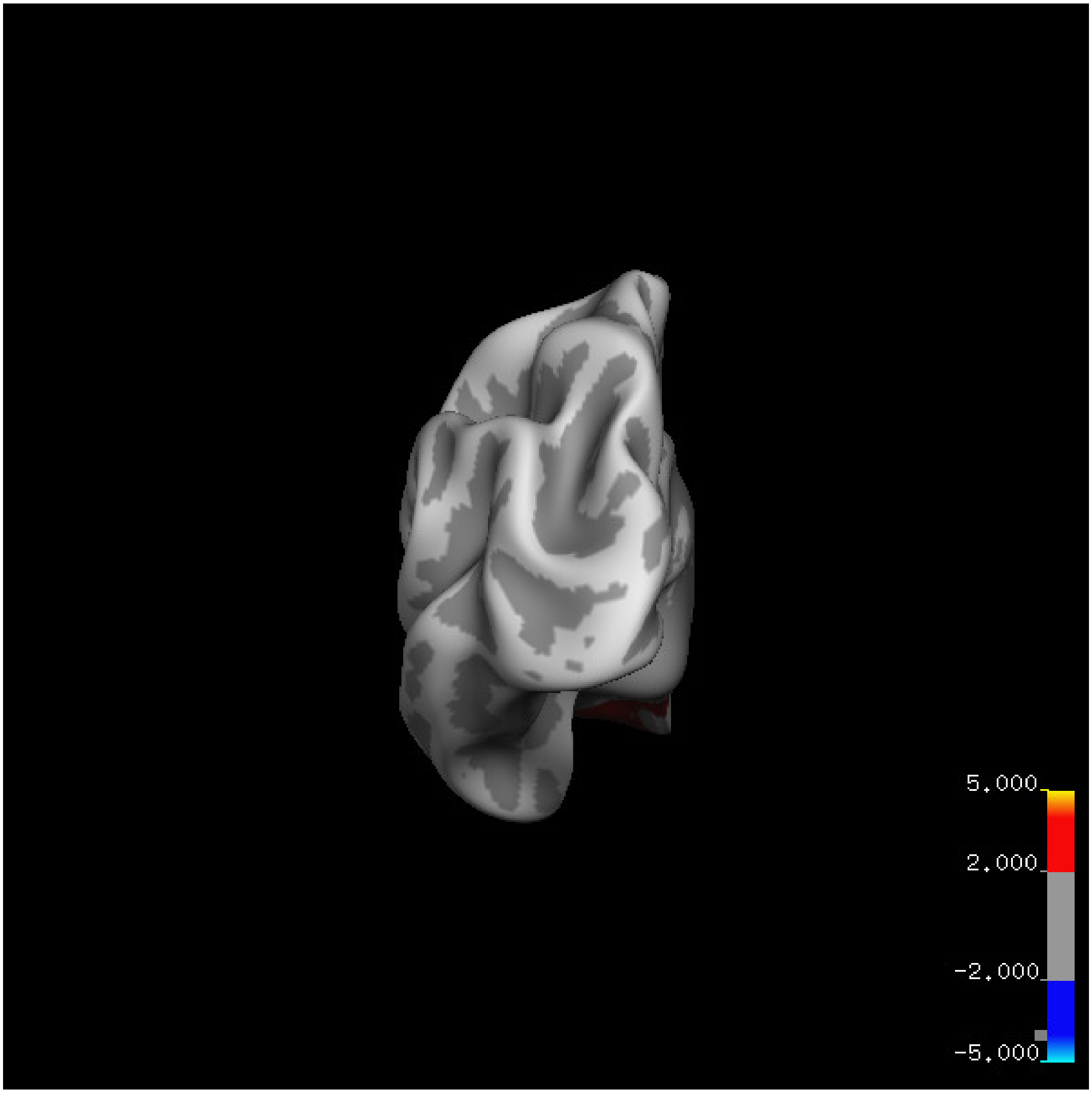
5.000
2.000
-2.000
-5.000

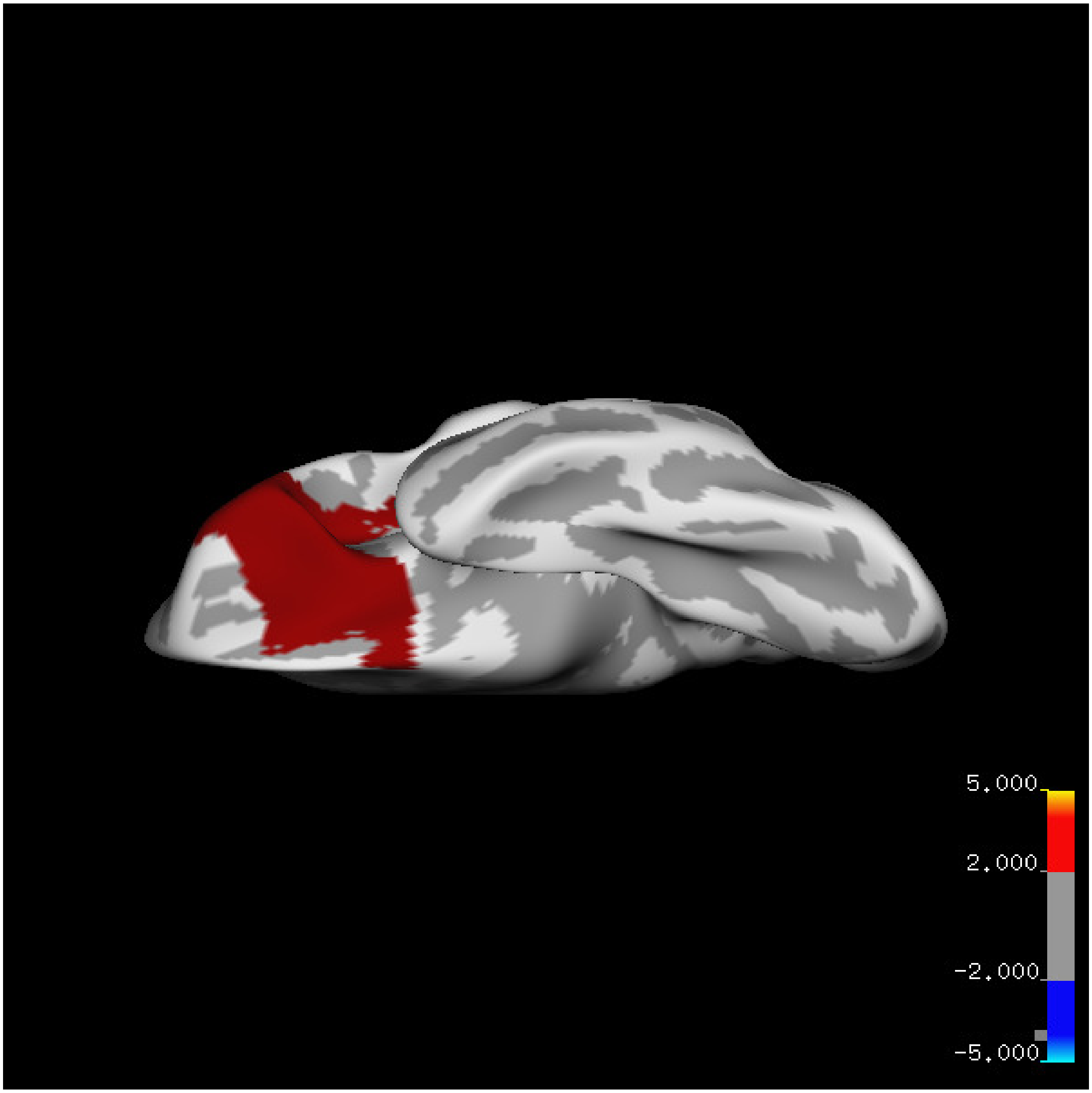
5.000
2.000
-2.000
-5.000

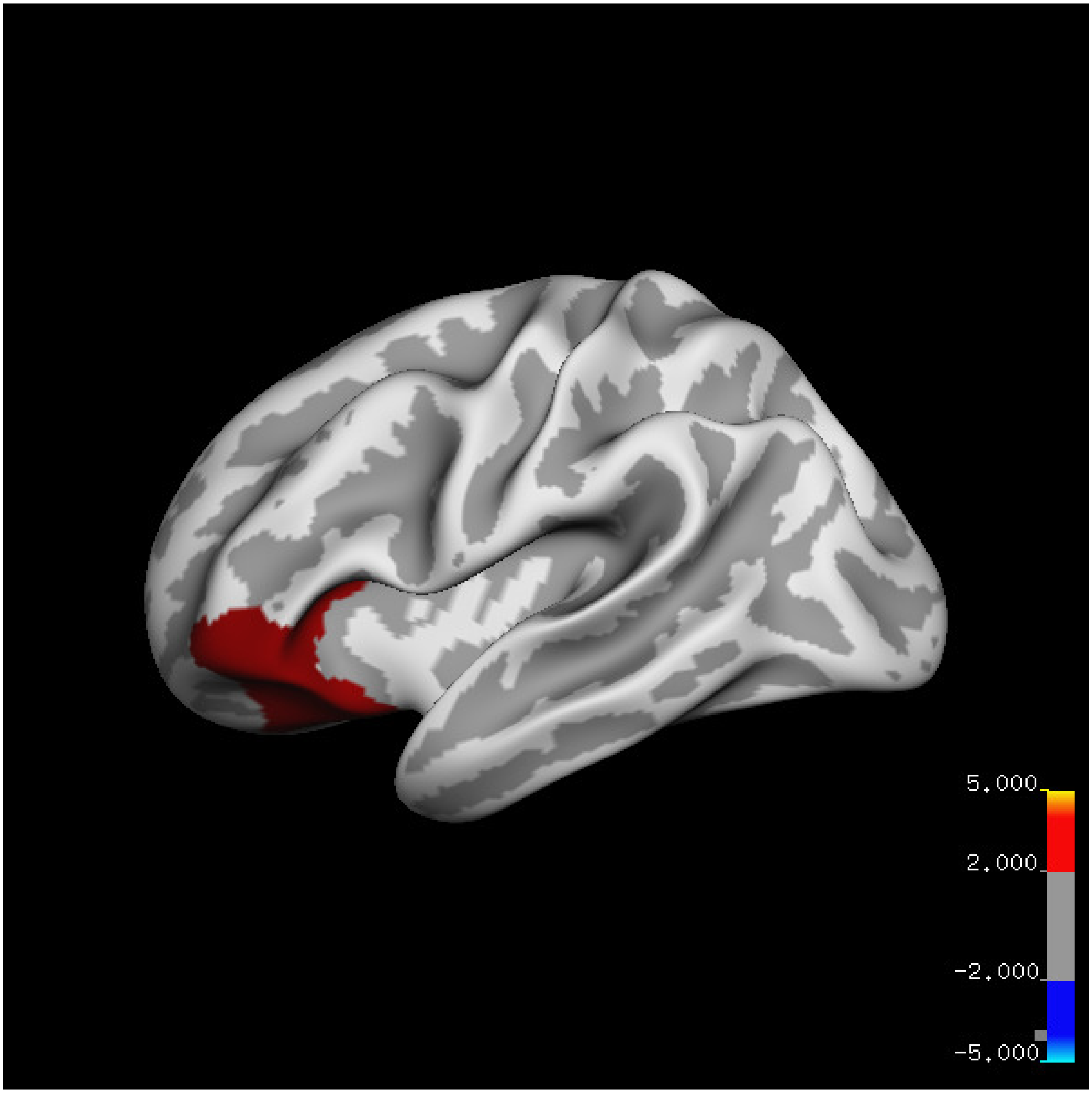
5.000
2.000
-2.000
-5.000

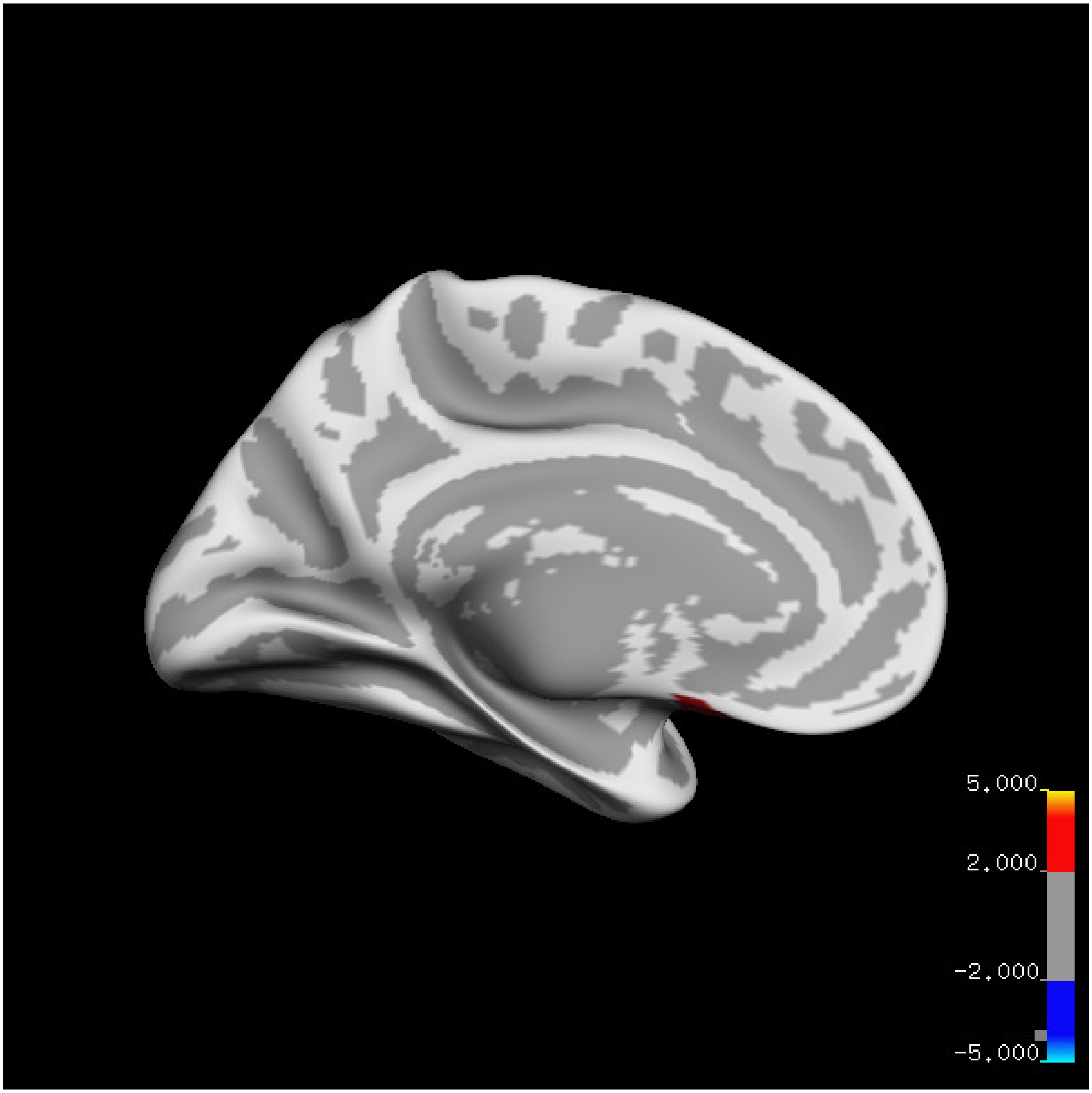
5.000
2.000
-2.000
-5.000

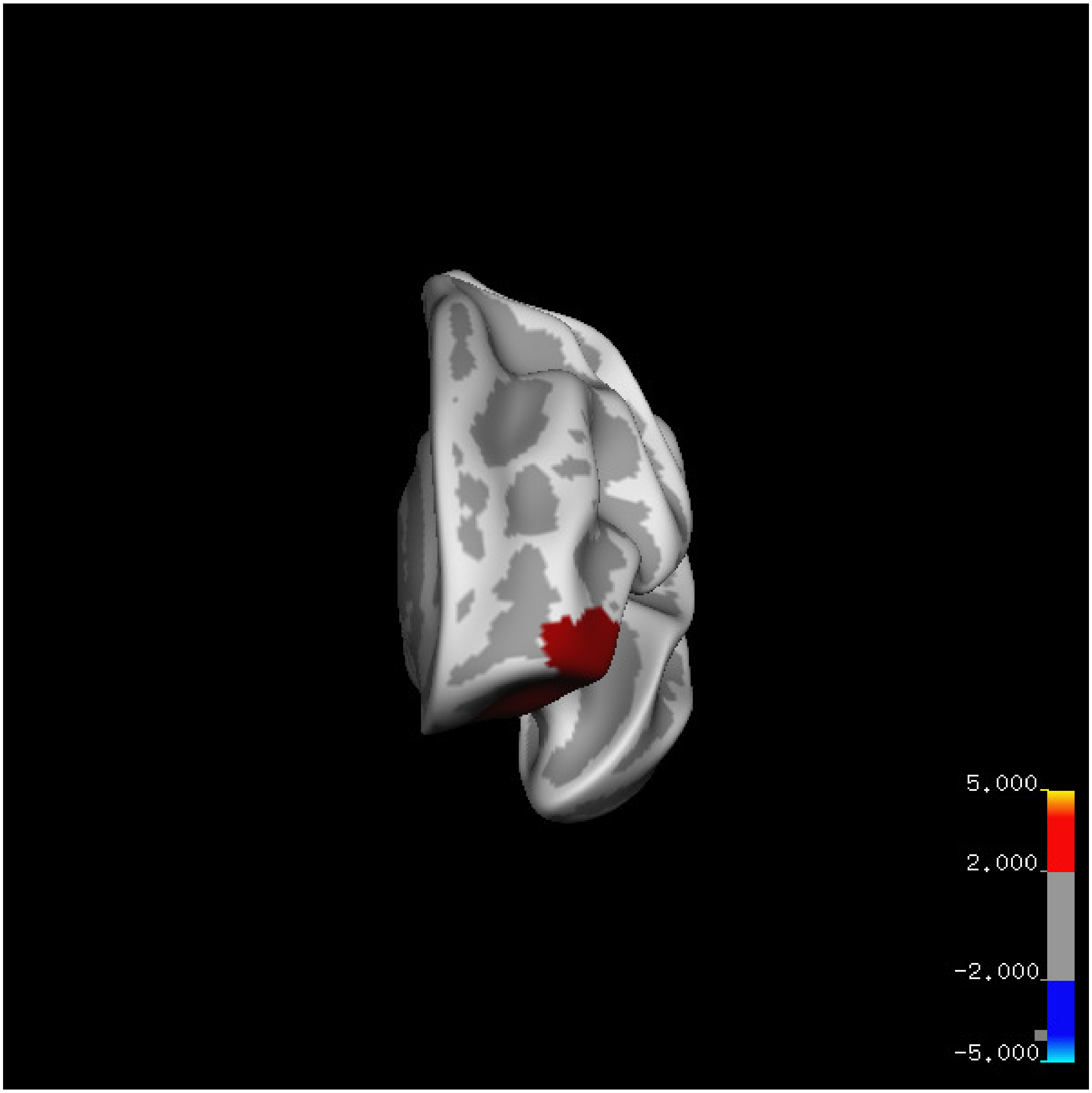
5.000
2.000
-2.000
-5.000

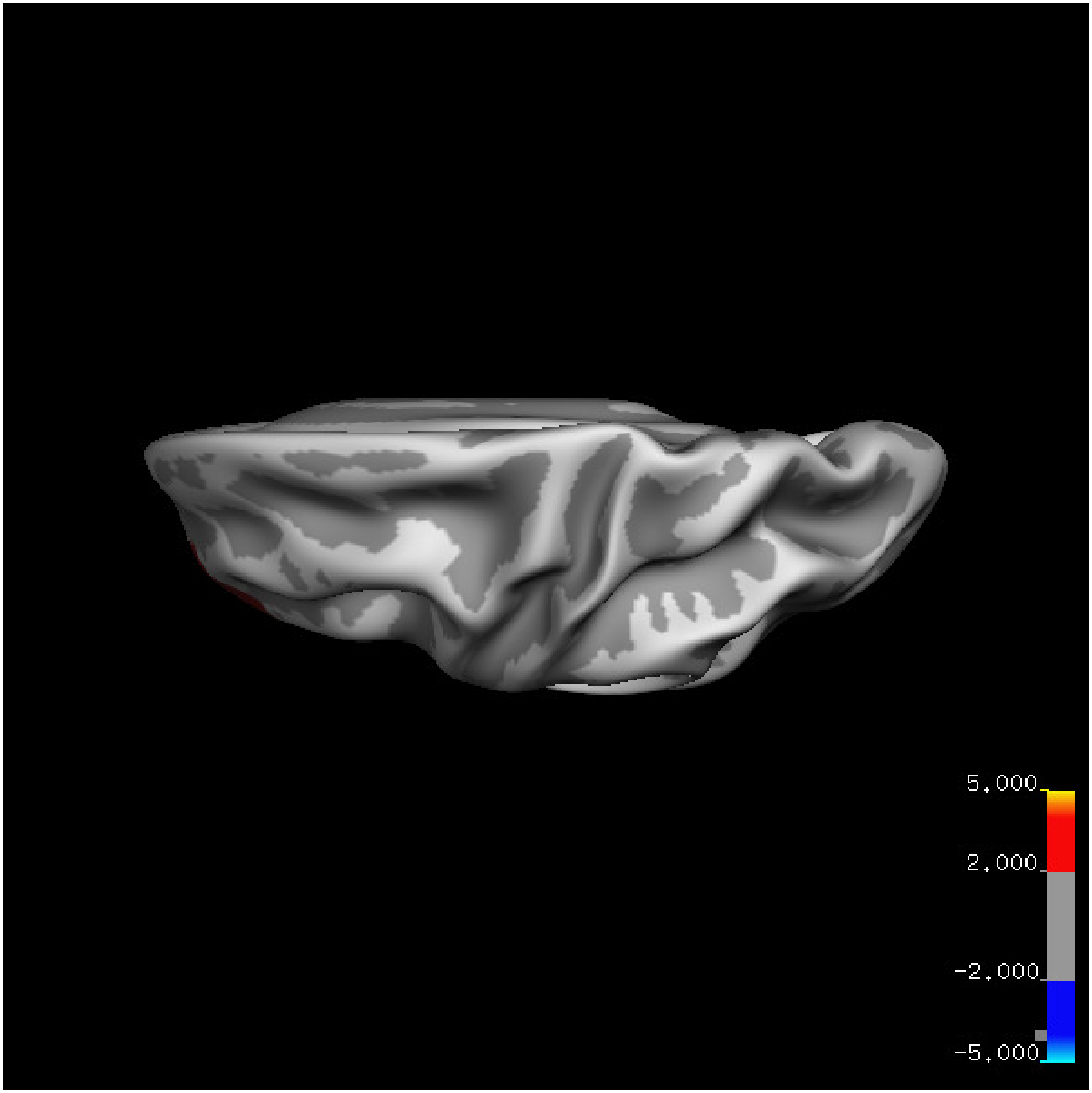
5.000
2.000
-2.000
-5.000

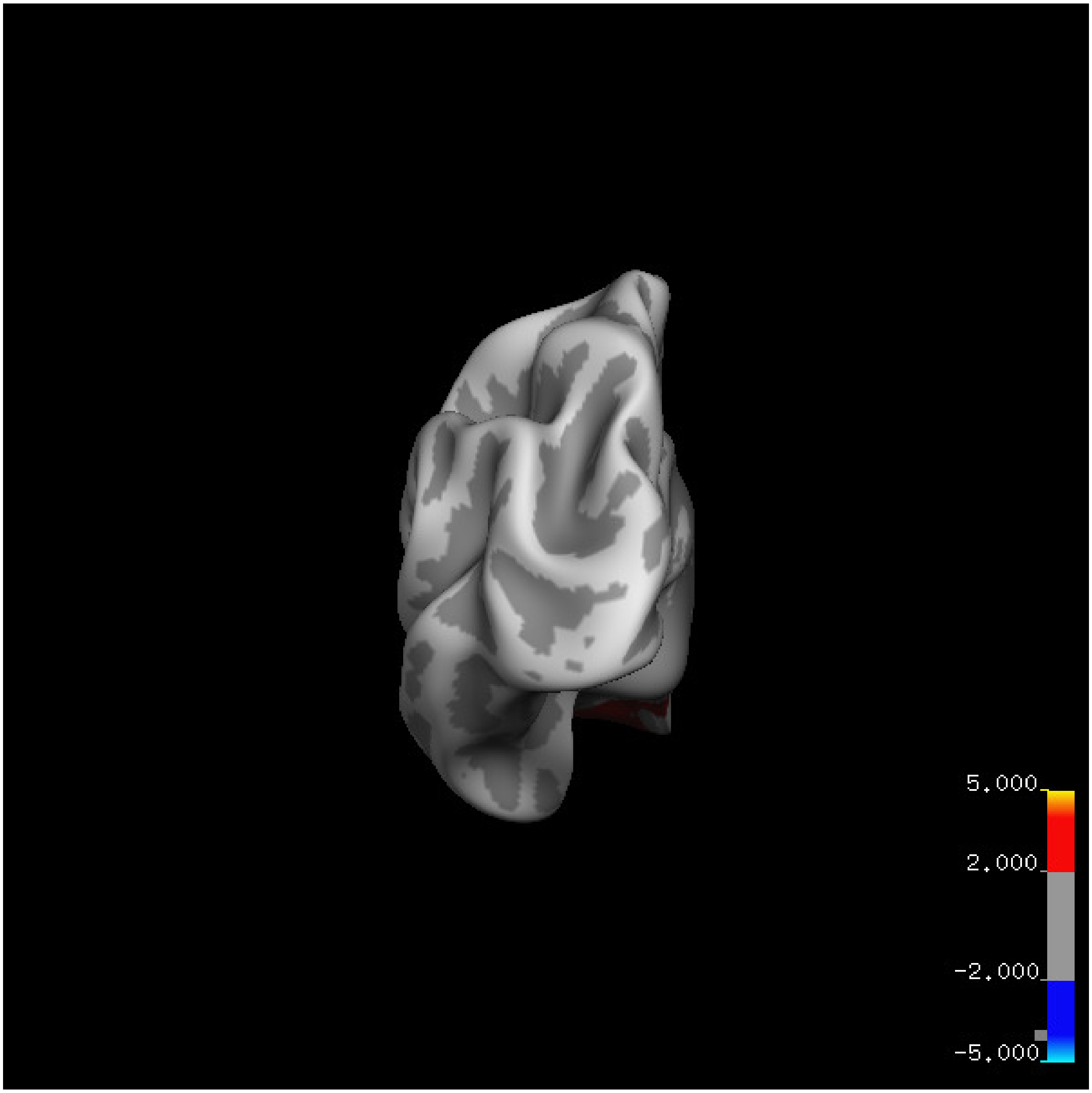
5.000
2.000
-2.000
-5.000

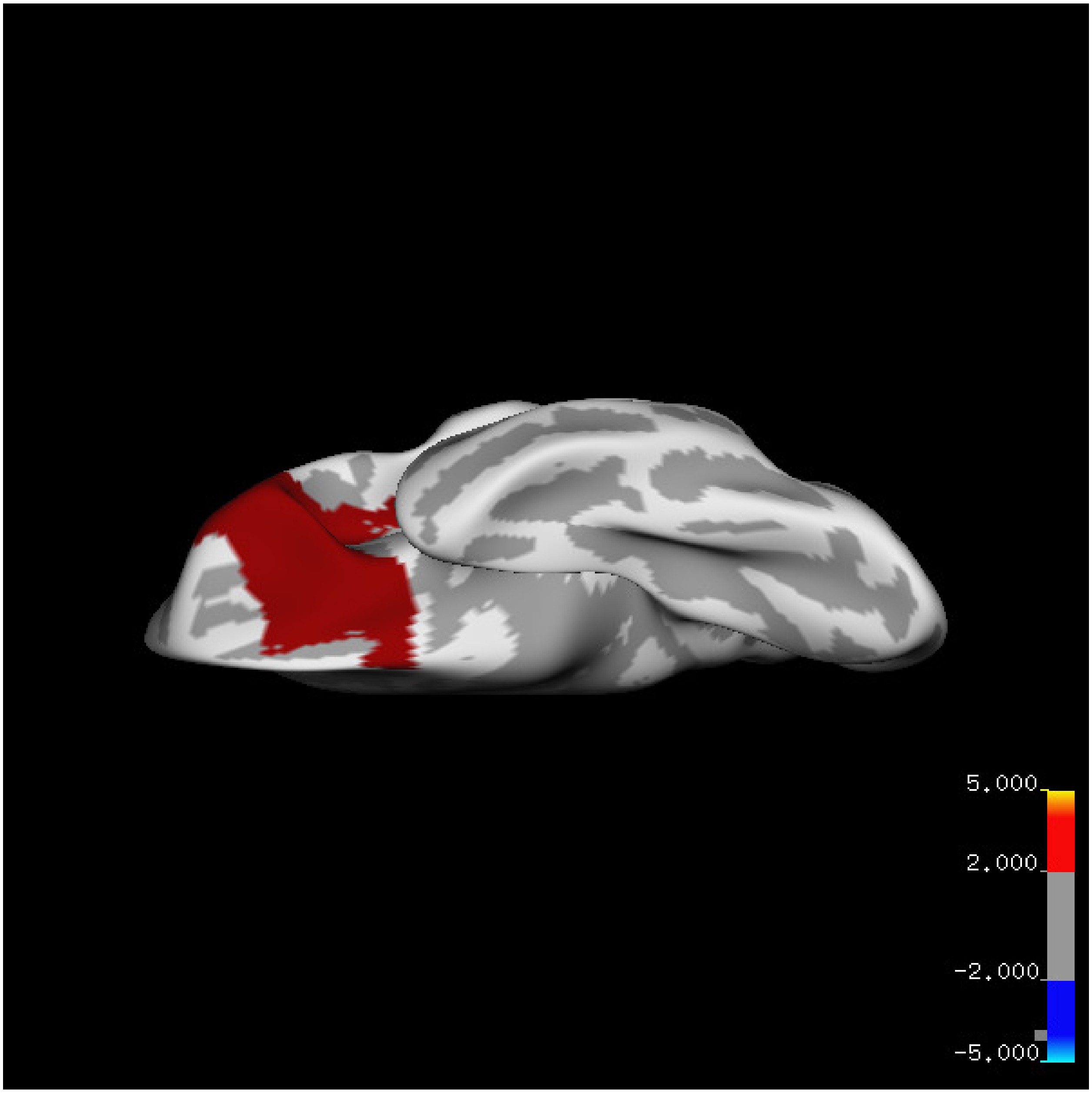
5.000
2.000
-2.000
-5.000

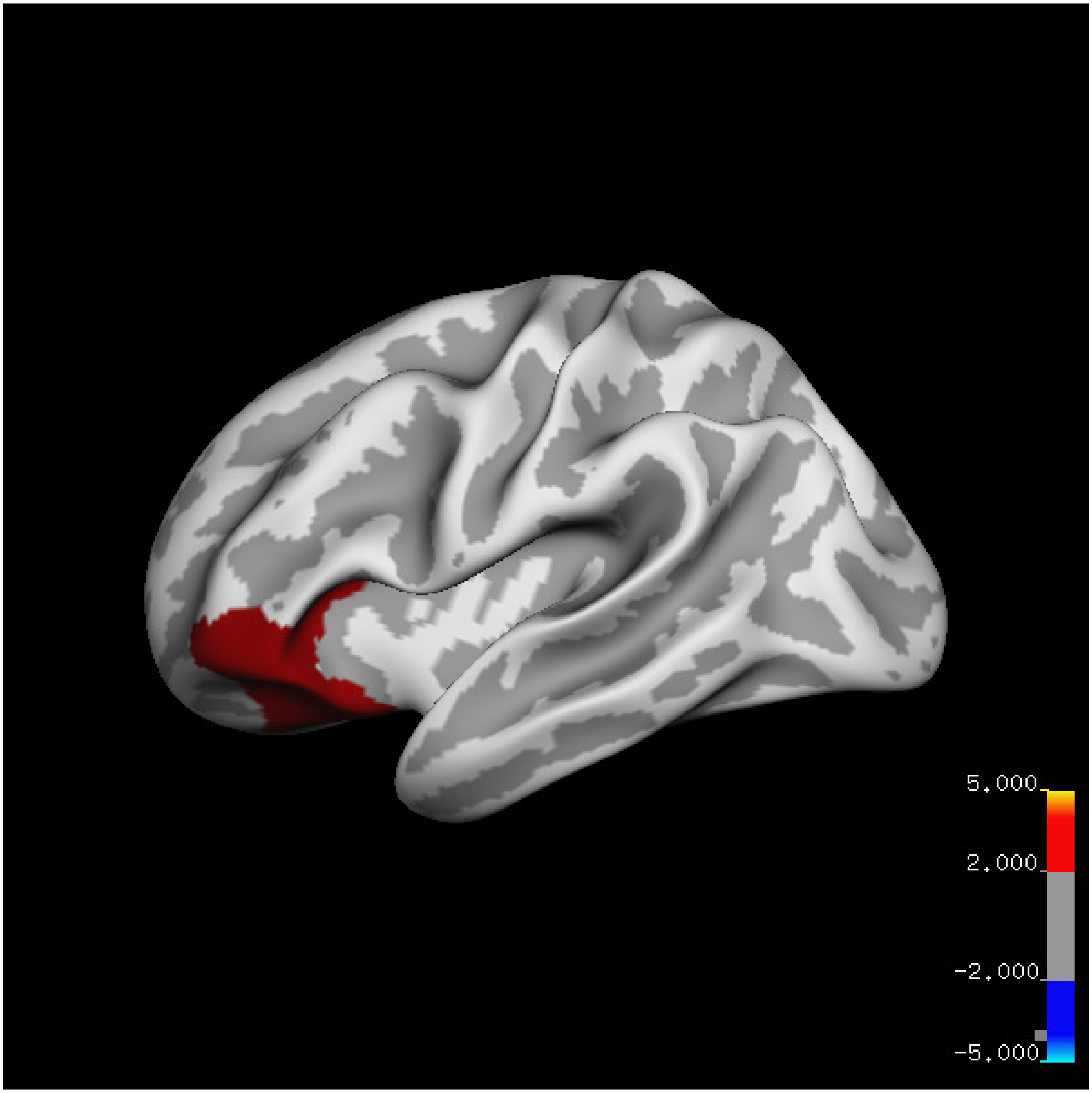
5.000
2.000
-2.000
-5.000

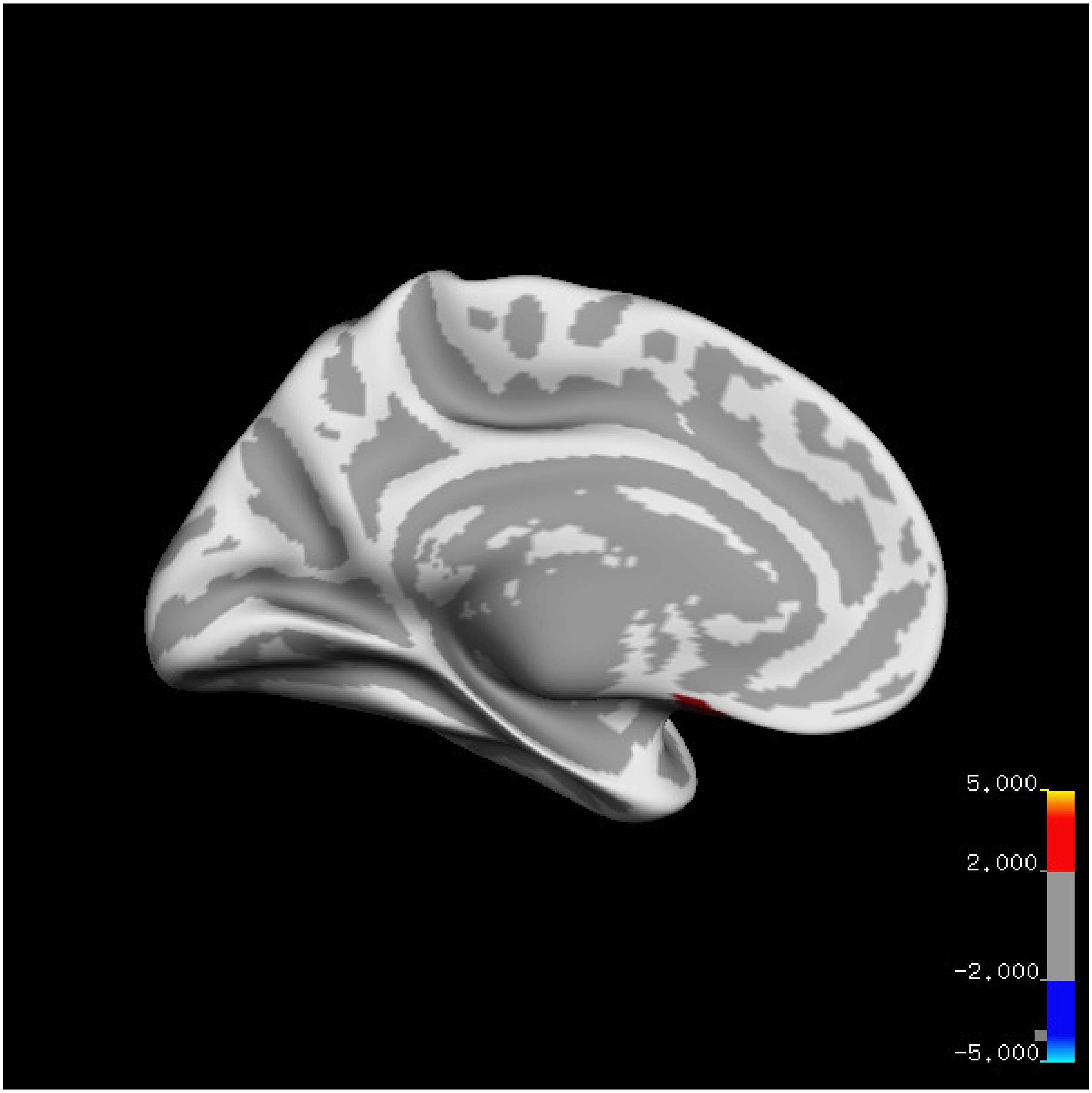
5.000
2.000
-2.000
-5.000

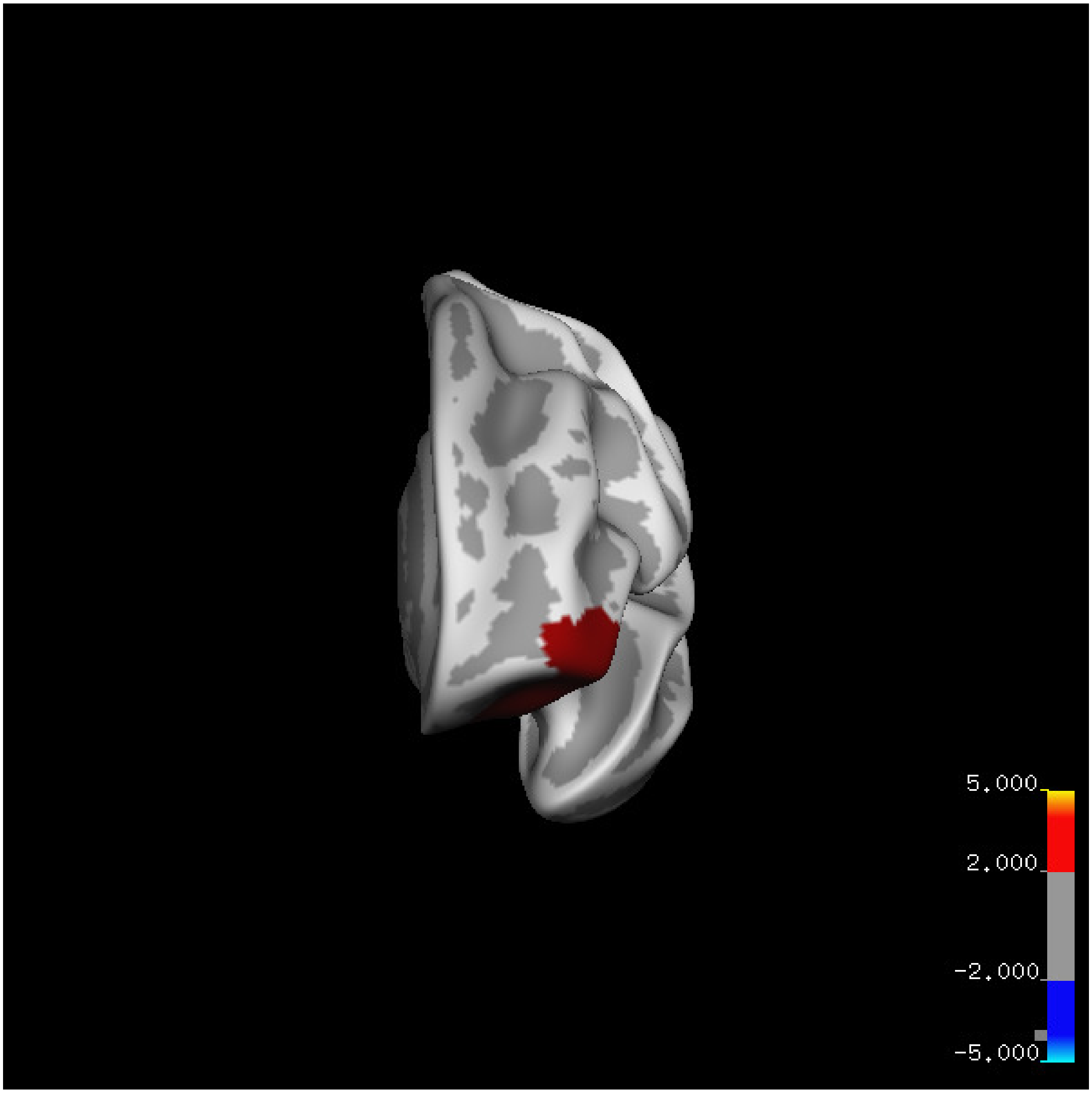
5.000
2.000
-2.000
-5.000

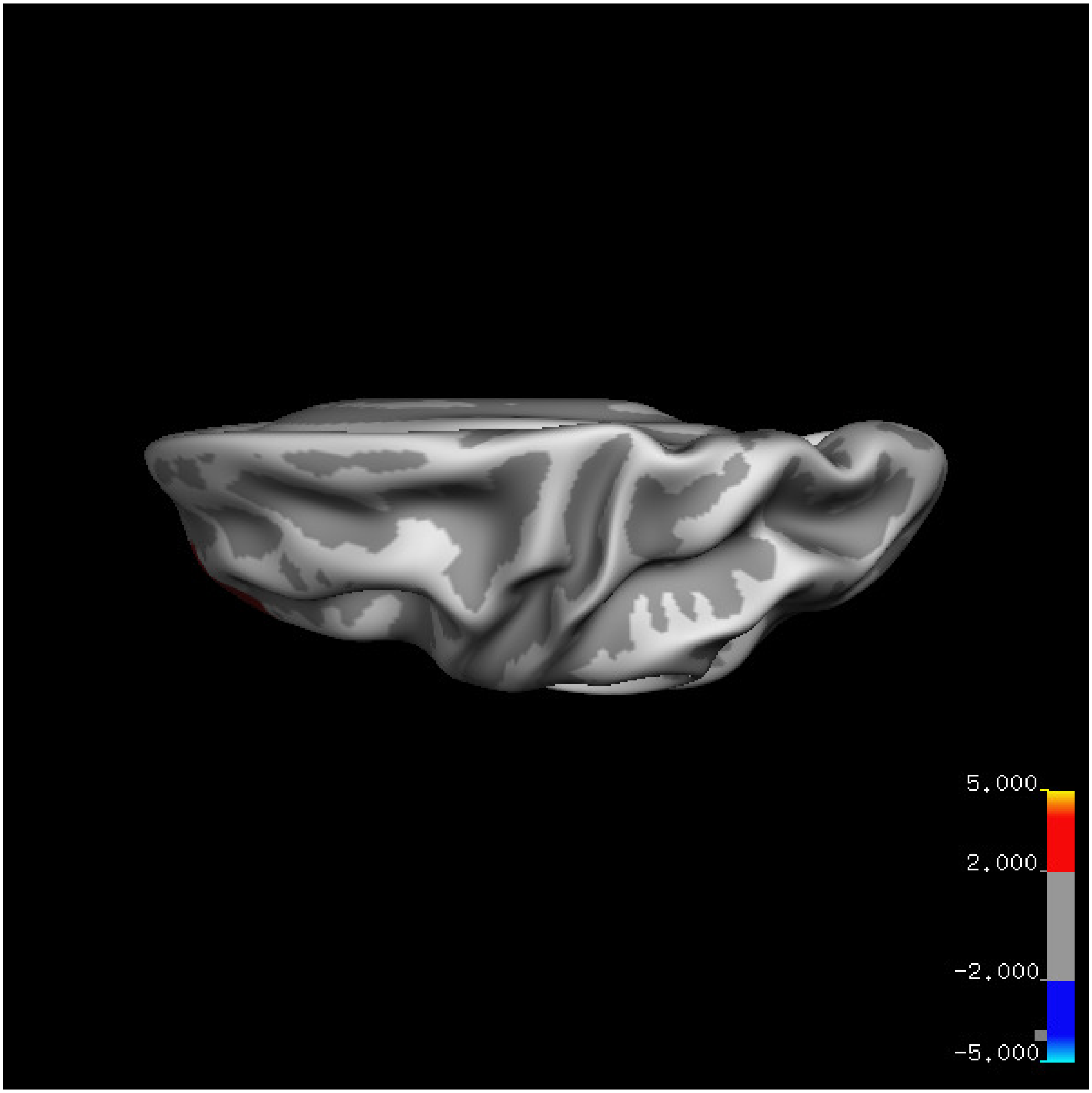


Cluster summary is shown below.

```
## # Cluster Growing Summary (mri_surfcluster)

## # $Id: mri_surfcluster.c,v 1.57.2.3 2016/11/17 18:19:42 zkaufman Exp $

## # $Id: mrisurf.c,v 1.781.2.6 2016/12/27 16:47:14 zkaufman Exp $

## # CreationTime 2021/10/04-11:24:45-GMT

## # cmdline mri_surfcluster.bin --in surface_analyses/results/cluster_comparison/fsaverage6.sm15.lh/ar

## # cwd /gpfs/projects02/p274/cluster/projects/p003-sleep_b

## # sysname  Linux
```

```
## # hostname c1-18
## # machine  x86_64
## # FixVertexAreaFlag 1
## # FixSurfClusterArea 1
## #
## # Input      surface_analyses/results/cluster_comparison/fsaverage6.sm15.lh/area.20_60.below_mean/gl
## # Frame Number      0
## # srcsubj fsaverage6
## # hemi lh
## # surface white
## # group_avg_surface_area 84969.3
## # group_avg_vtxarea_loaded 1
## # annot aparc
## # SUBJECTS_DIR /cluster/projects/p274/tools/mri/freesurfer/current/subjects
## # SearchSpace_mm2 79241.3
## # SearchSpace_vtx 37476
## # Bonferroni 0
## # Minimum Threshold 1.3
## # Maximum Threshold infinity
## # Threshold Sign    abs
## # AdjustThreshWhenOneTail 1
## # CW PValue Threshold: 0.01
## # Area Threshold    0 mm^2
## # CSD thresh  1.300000
## # CSD nreps    10000
## # CSD simtype  null-z
## # CSD contrast NA
```

```
## # CSD confint  90.000000

## # Overall max 5.29361 at vertex 20247

## # Overall min -3.18052 at vertex 38177

## # NClusters          1

## # FixMNI = 0

## #

## # ClusterNo  Max   VtxMax   Size(mm^2)  MNIX   MNIY   MNIZ    CWP    CWPLow    CWPHi   NVtxs    Wght

## 1        5.294   20247   3258.53    -29.8   29.5  -16.1  0.00940  0.00820  0.01060   1546     370
```

**Age above 60**

**Sleep above mean** Cluster corrected significance plots are shown below.

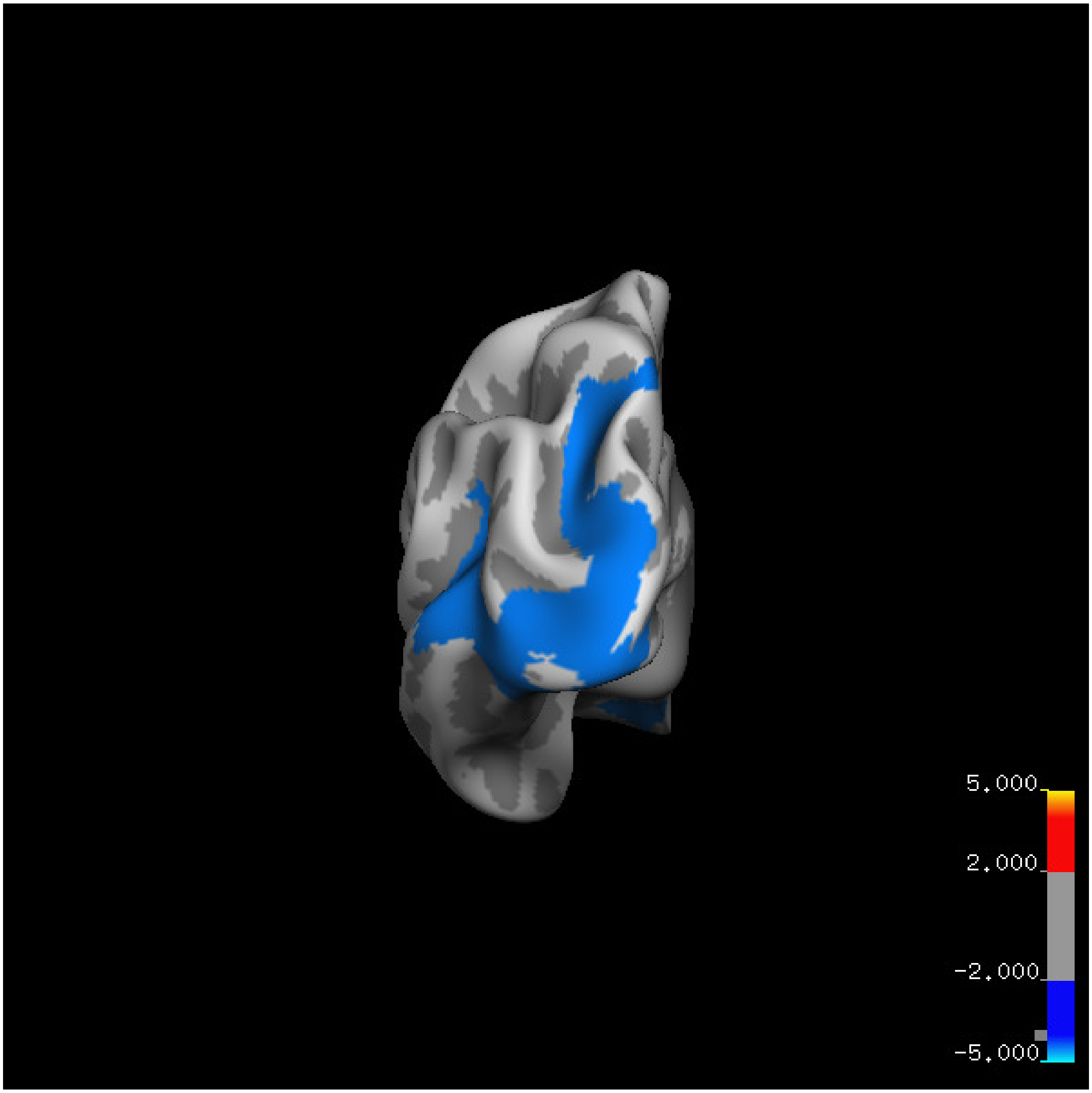
5.000
2.000
-2.000
-5.000

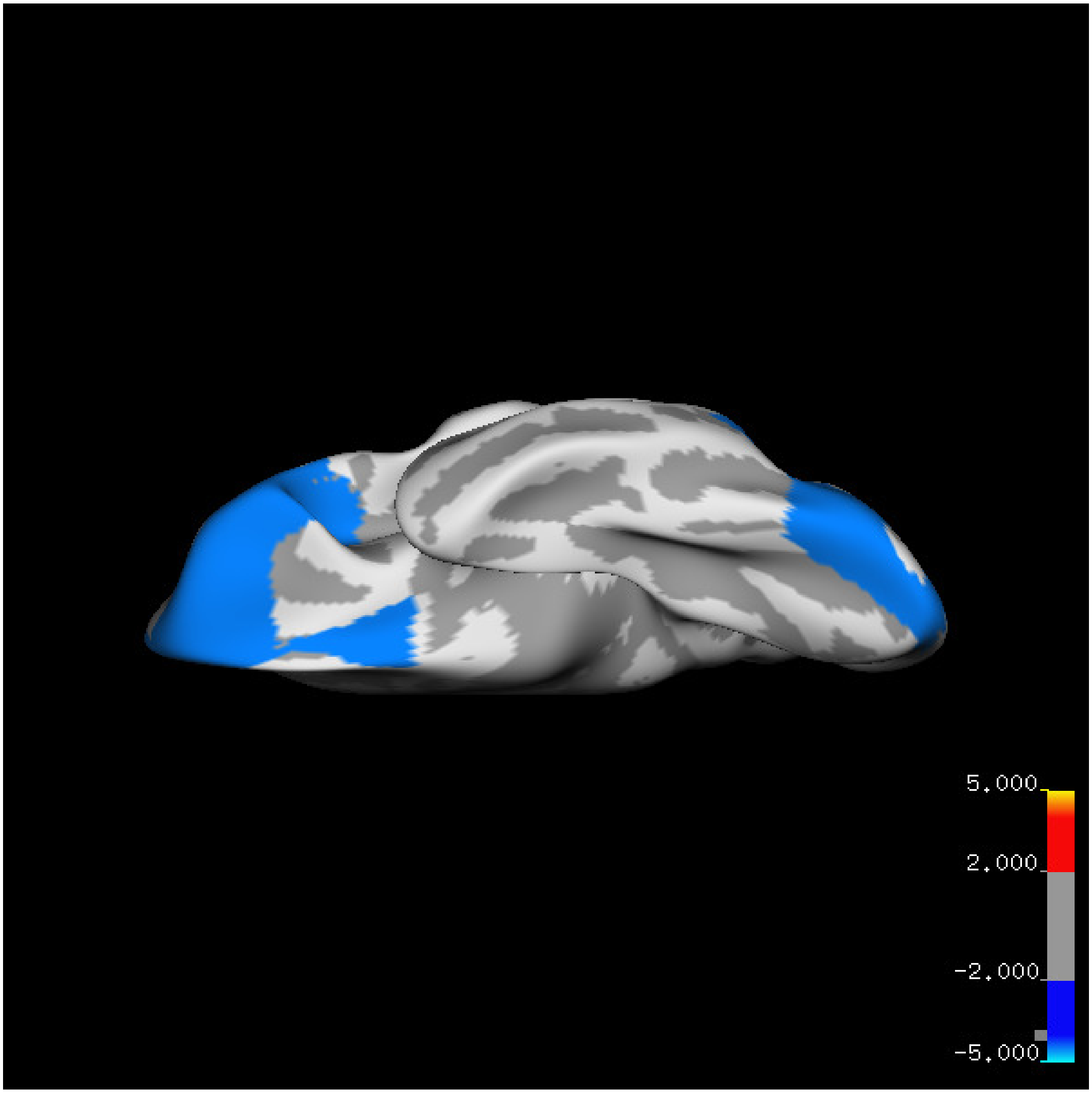

5.000
2.000
-2.000
-5.000

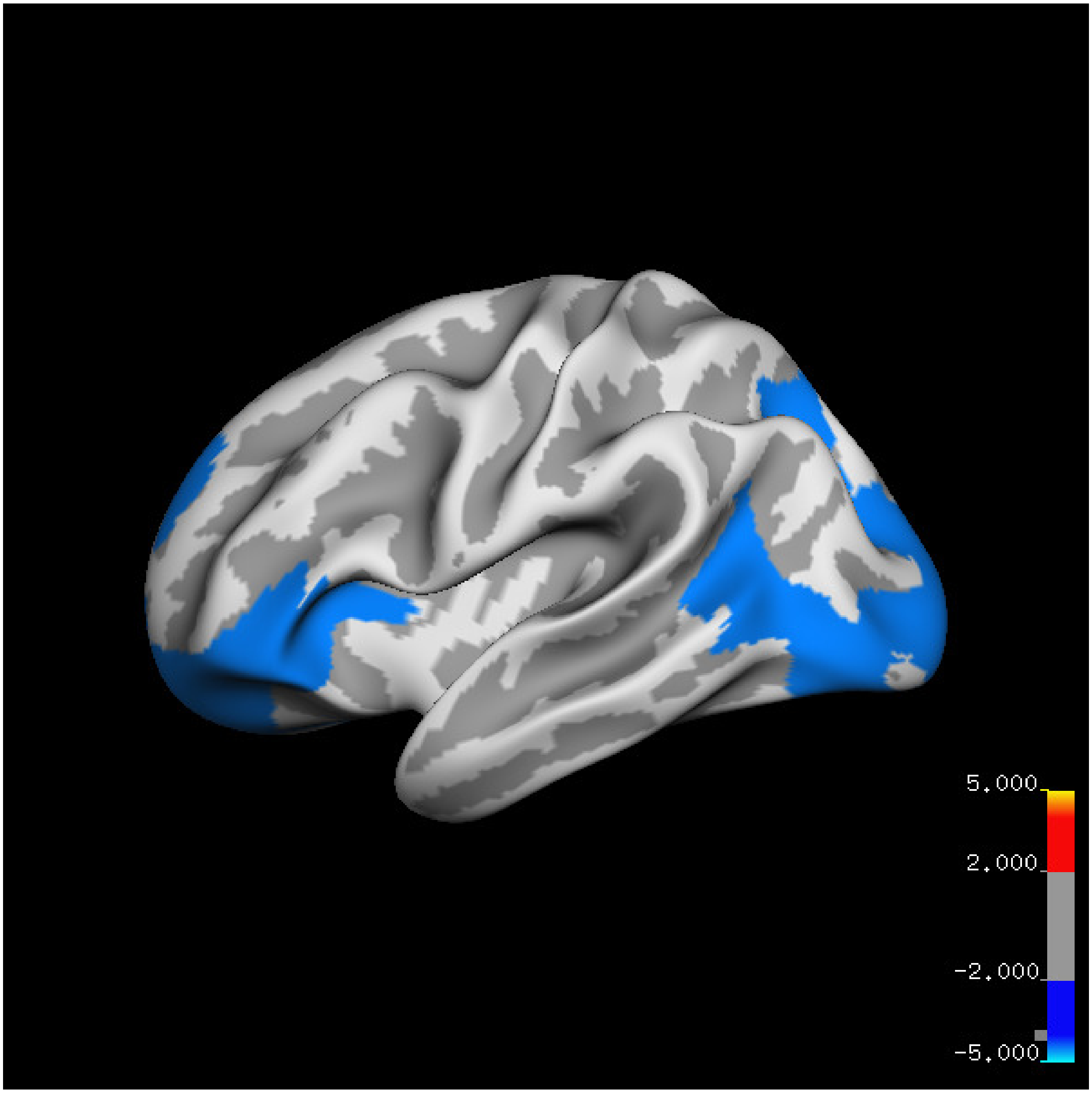
5.000
2.000
-2.000
-5.000

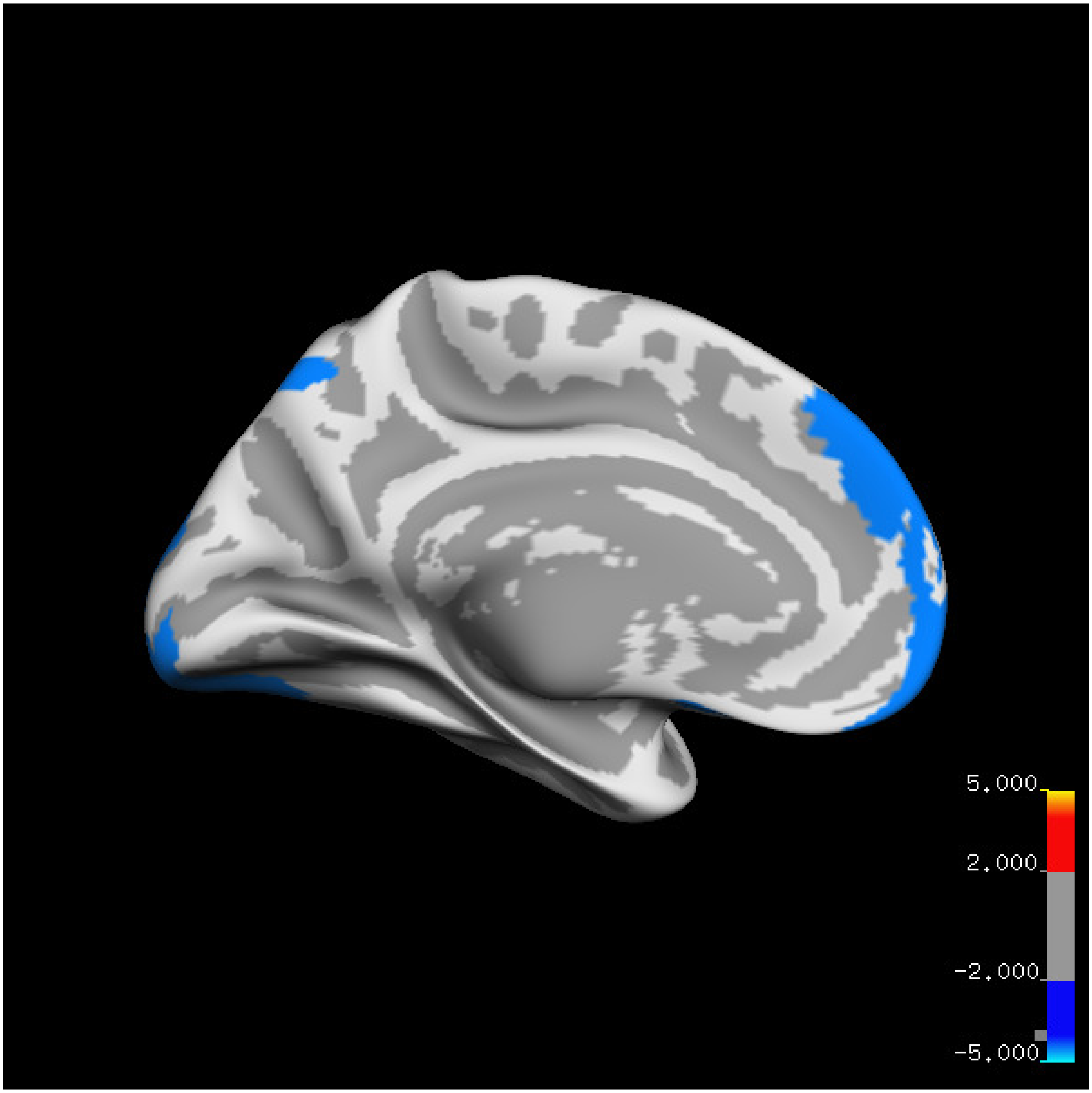

5.000
2.000
-2.000
-5.000

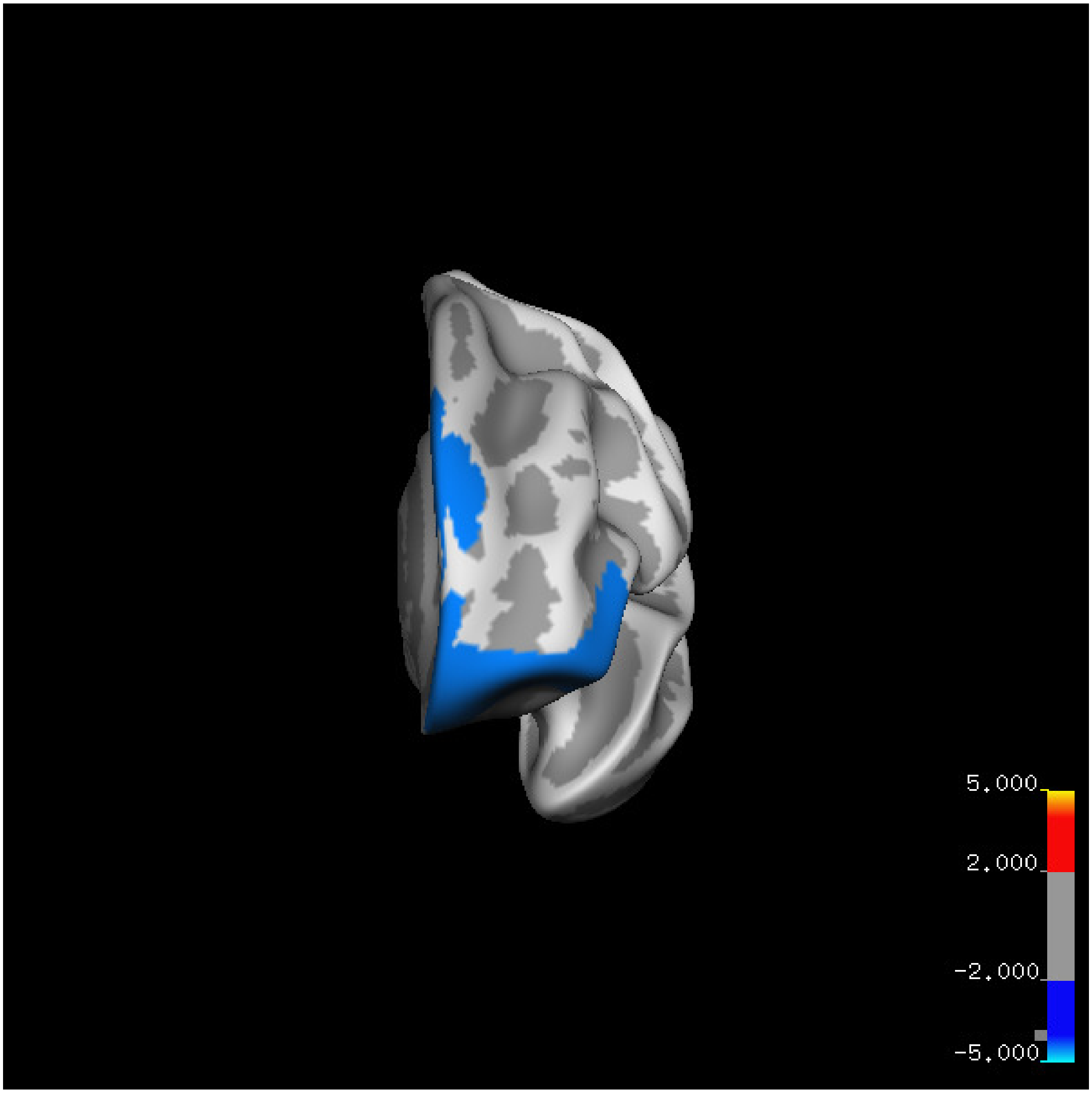
5.000
2.000
-2.000
-5.000

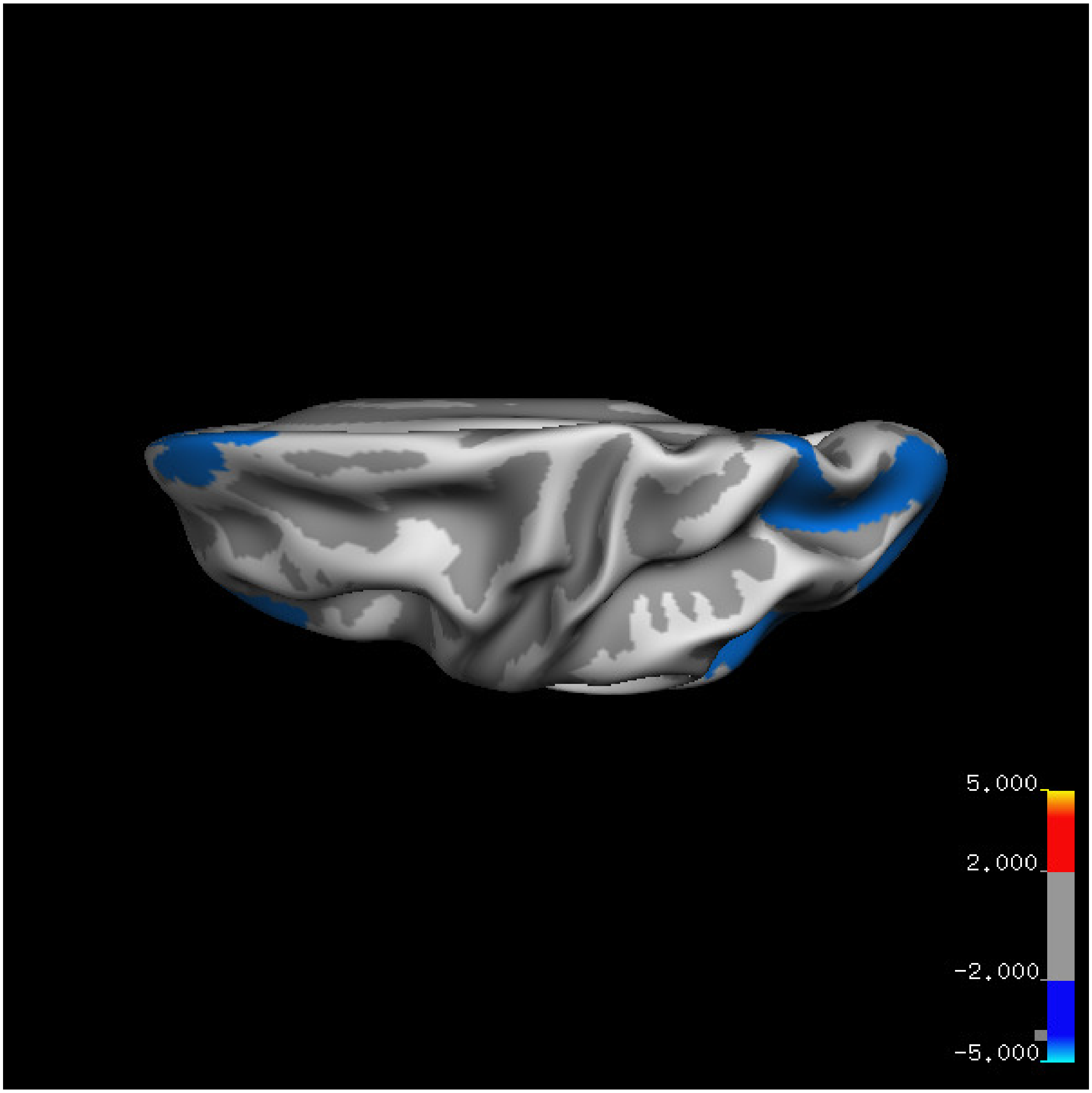
5.000
2.000
-2.000
-5.000

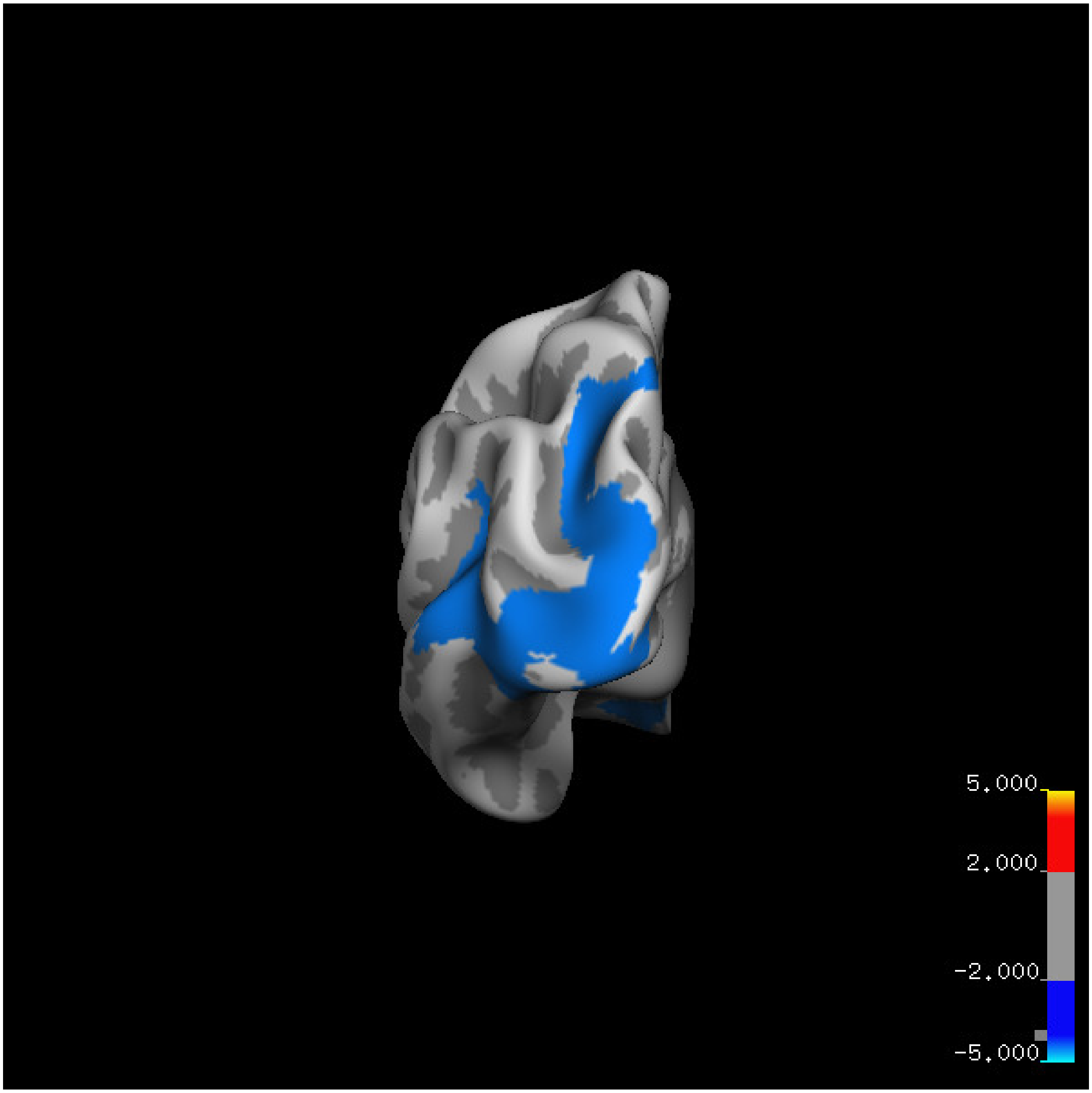
5.000
2.000
-2.000
-5.000

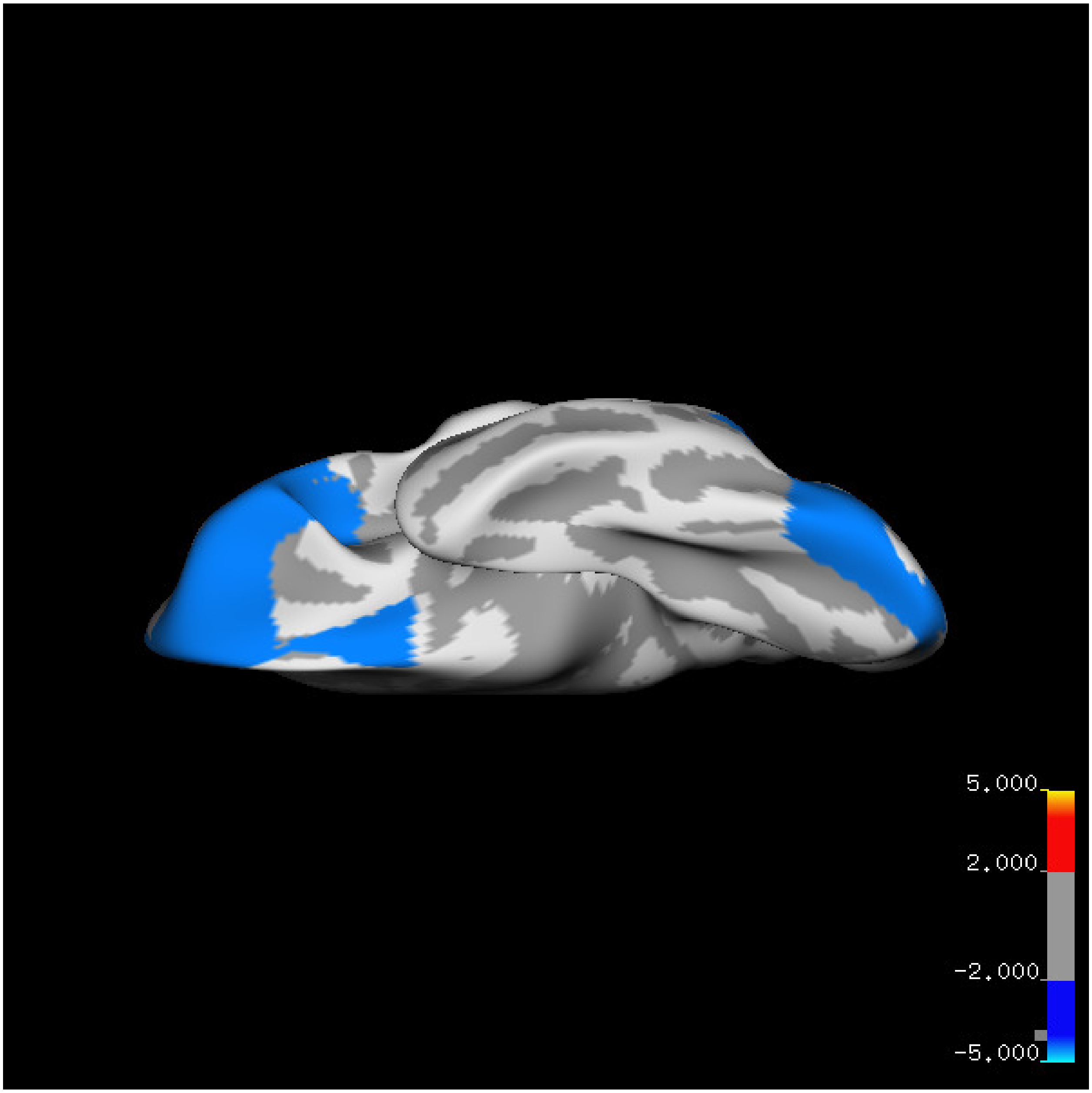
5.000
2.000
-2.000
-5.000

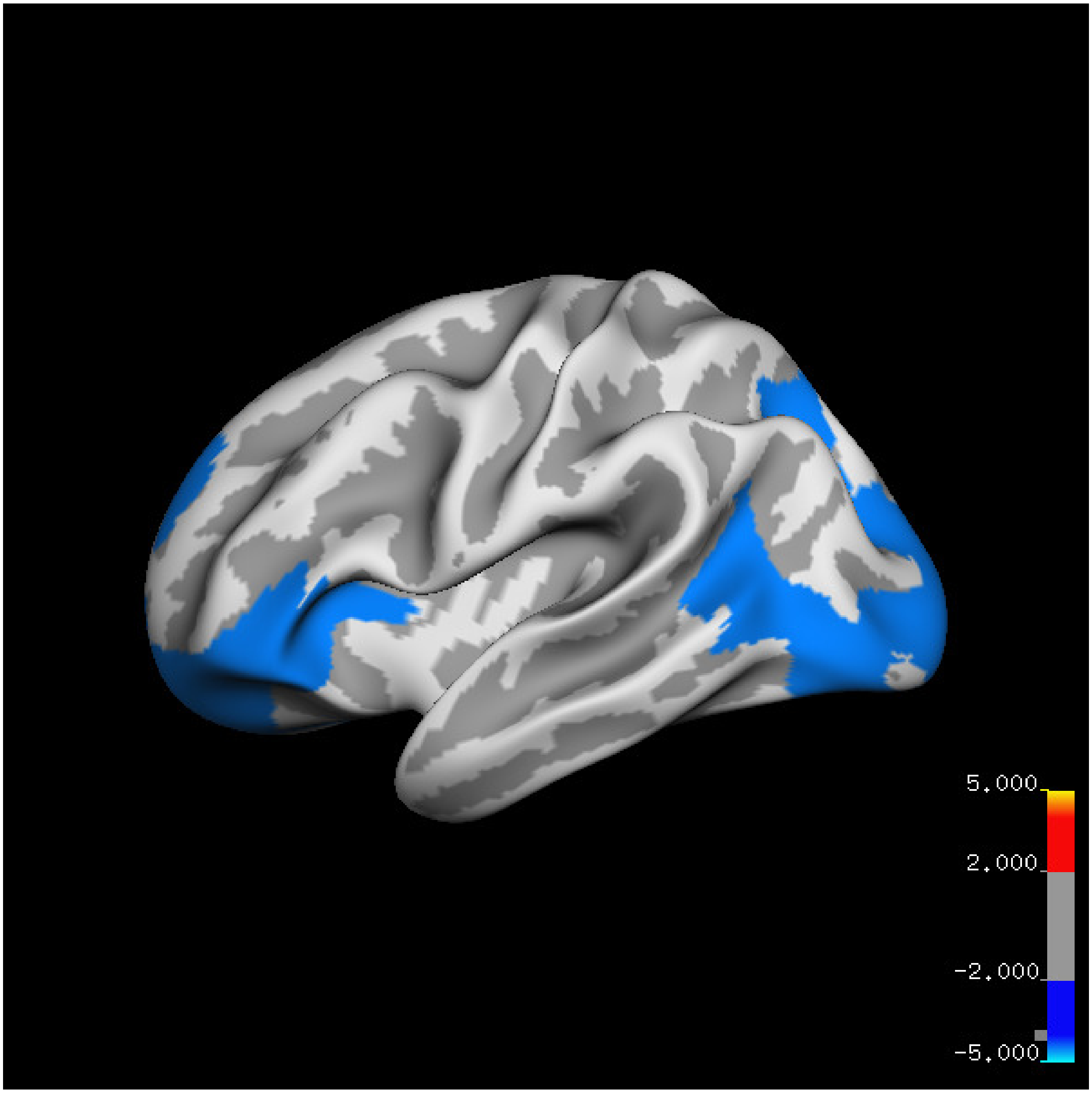
5.000
2.000
-2.000
-5.000

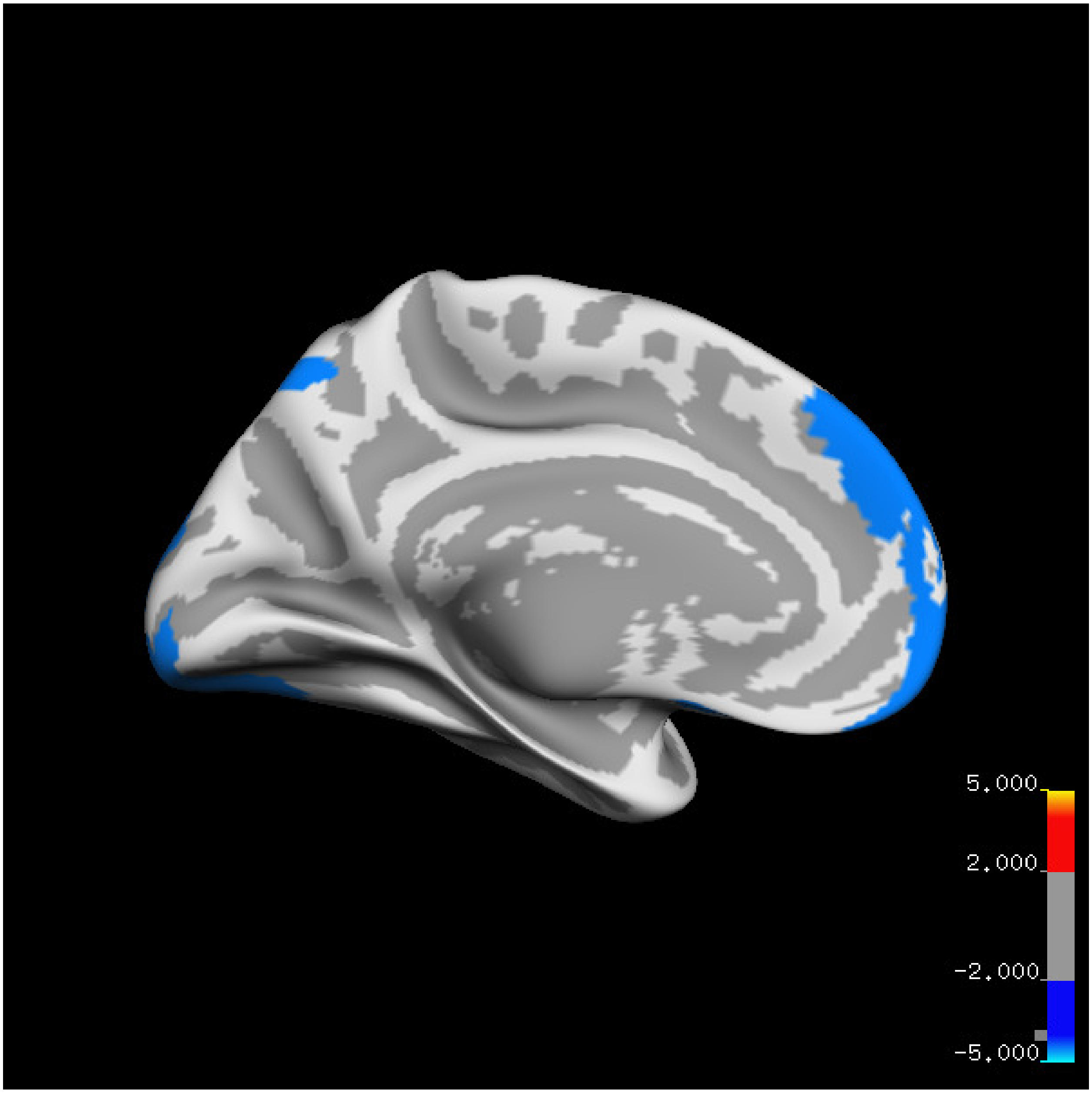
5.000
2.000
-2.000
-5.000

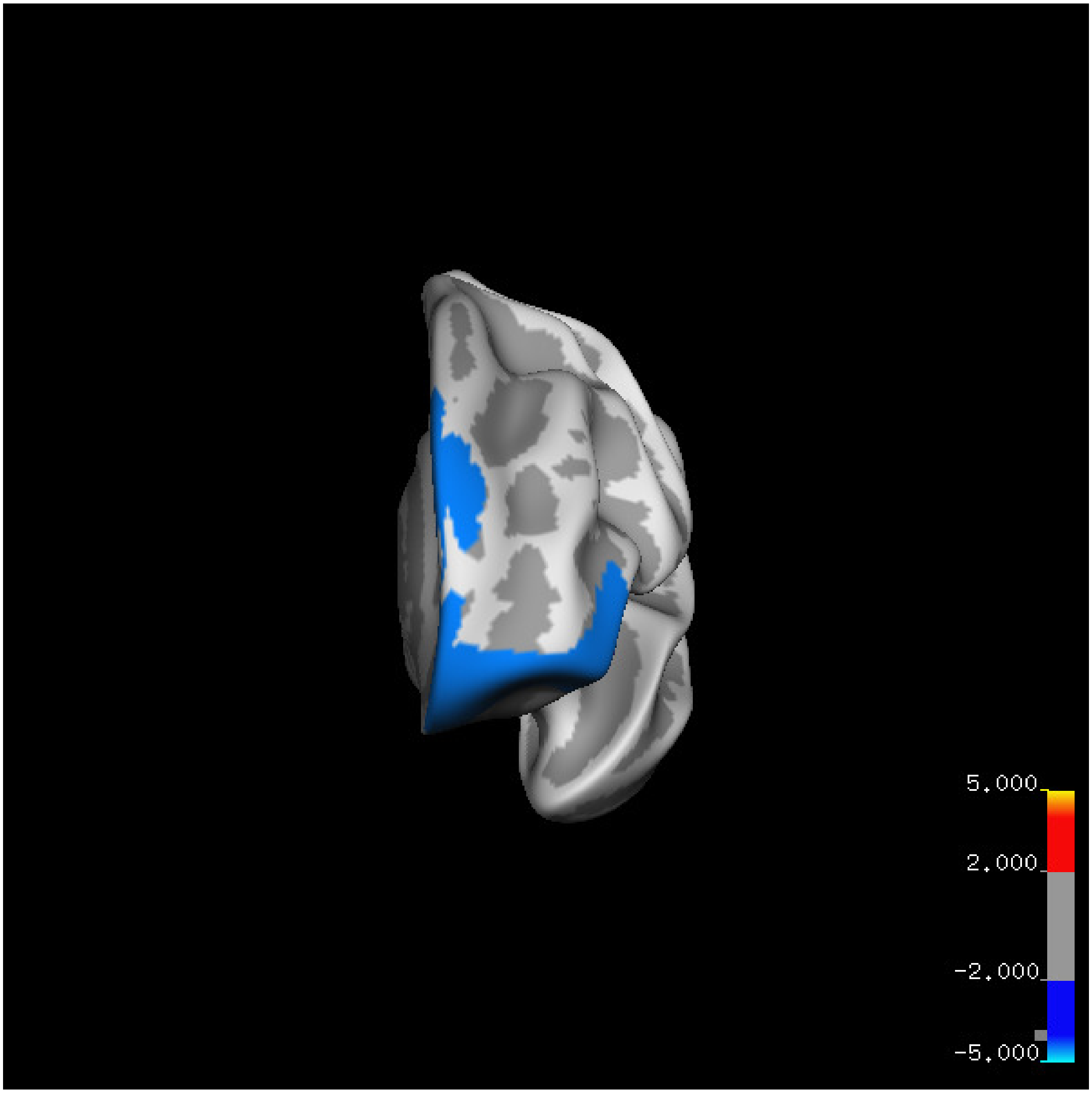
5.000
2.000
-2.000
-5.000

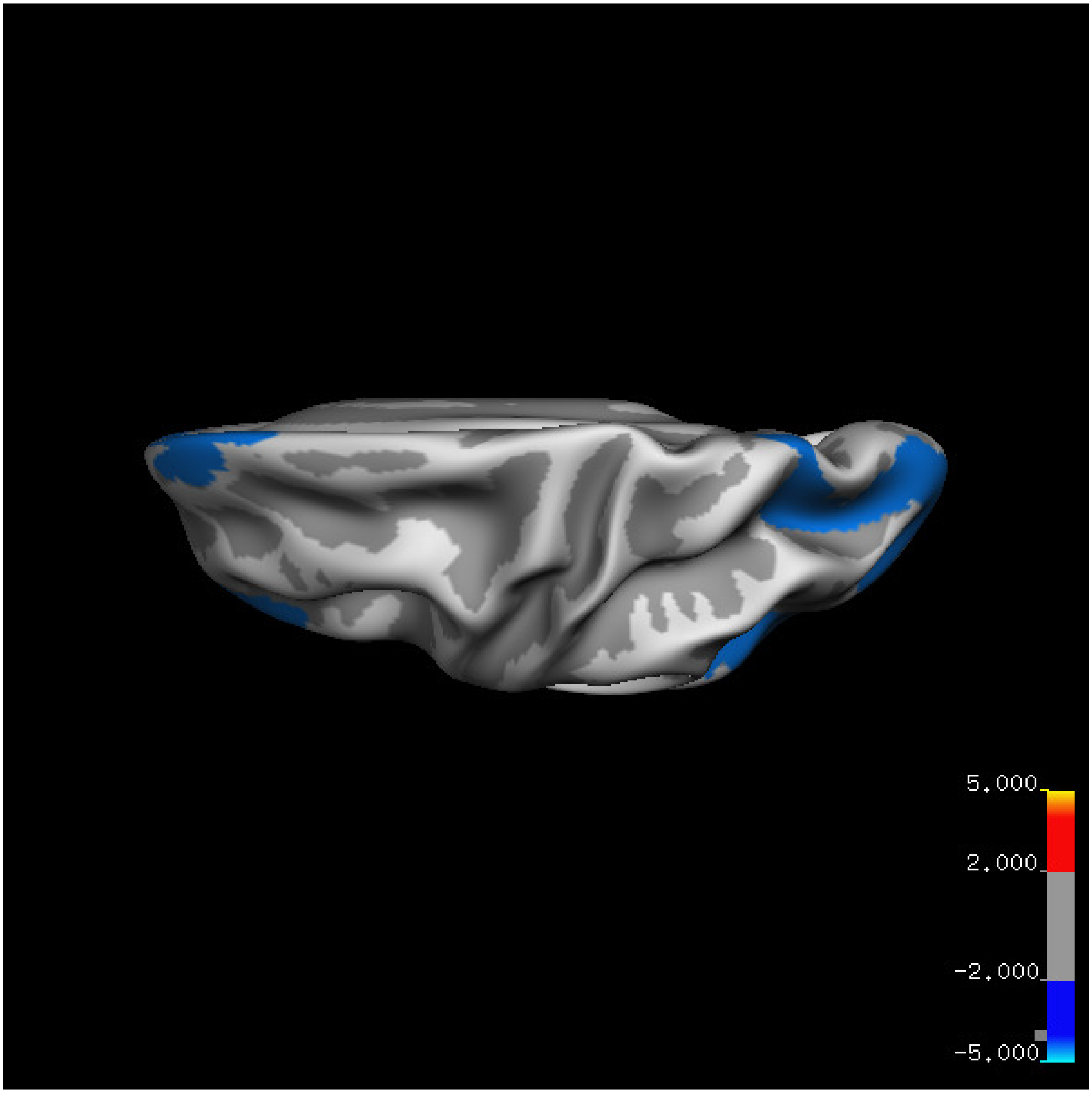


Cluster summary is shown below.

```
## # Cluster Growing Summary (mri_surfcluster)

## # $Id: mri_surfcluster.c,v 1.57.2.3 2016/11/17 18:19:42 zkaufman Exp $

## # $Id: mrisurf.c,v 1.781.2.6 2016/12/27 16:47:14 zkaufman Exp $

## # CreationTime 2021/10/04-13:08:36-GMT

## # cmdline mri_surfcluster.bin --in surface_analyses/results/cluster_comparison/fsaverage6.sm15.lh/ar

## # cwd /gpfs/projects02/p274/cluster/projects/p003-sleep_b

## # sysname  Linux
```

```
## # hostname c1-18
## # machine  x86_64
## # FixVertexAreaFlag 1
## # FixSurfClusterArea 1
## #
## # Input      surface_analyses/results/cluster_comparison/fsaverage6.sm15.lh/area.above60.above_mean/g
## # Frame Number      0
## # srcsubj fsaverage6
## # hemi lh
## # surface white
## # group_avg_surface_area 84969.3
## # group_avg_vtxarea_loaded 1
## # annot aparc
## # SUBJECTS_DIR /cluster/projects/p274/tools/mri/freesurfer/current/subjects
## # SearchSpace_mm2 79241.3
## # SearchSpace_vtx 37476
## # Bonferroni 0
## # Minimum Threshold 1.3
## # Maximum Threshold infinity
## # Threshold Sign    abs
## # AdjustThreshWhenOneTail 1
## # CW PValue Threshold: 0.01
## # Area Threshold    0 mm^2
## # CSD thresh  1.300000
## # CSD nreps    10000
## # CSD simtype  null-z
## # CSD contrast NA
```

```
## # CSD confint  90.000000

## # Overall max 4.47219 at vertex 27406

## # Overall min -3.89879 at vertex 20309

## # NClusters          2

## # FixMNI = 0

## #

## # ClusterNo  Max   VtxMax   Size(mm^2)  MNIX   MNIY   MNIZ    CWP    CWPLow    CWPHi   NVtxs    WghtV

##    1       -2.899    5121   6773.71    -47.6  -59.8   11.8  0.00010  0.00000  0.00020   2556    -4313

##    2       -3.899   20309   5644.96    -15.1   54.8  -15.7  0.00010  0.00000  0.00020   2295    -4655
```

**Sleep below mean** Cluster corrected significance plots are shown below.

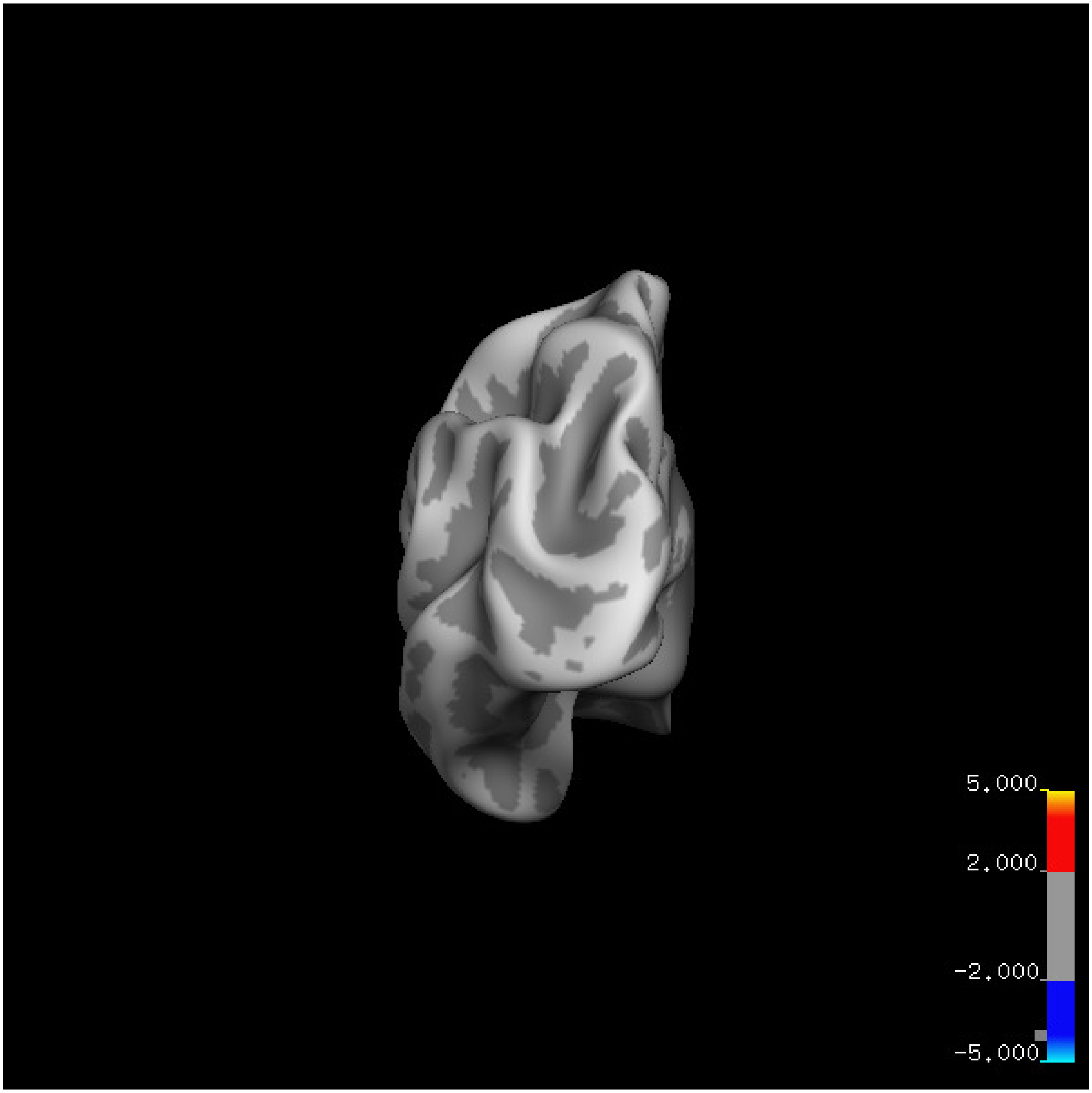
5.000
2.000
-2.000
-5.000

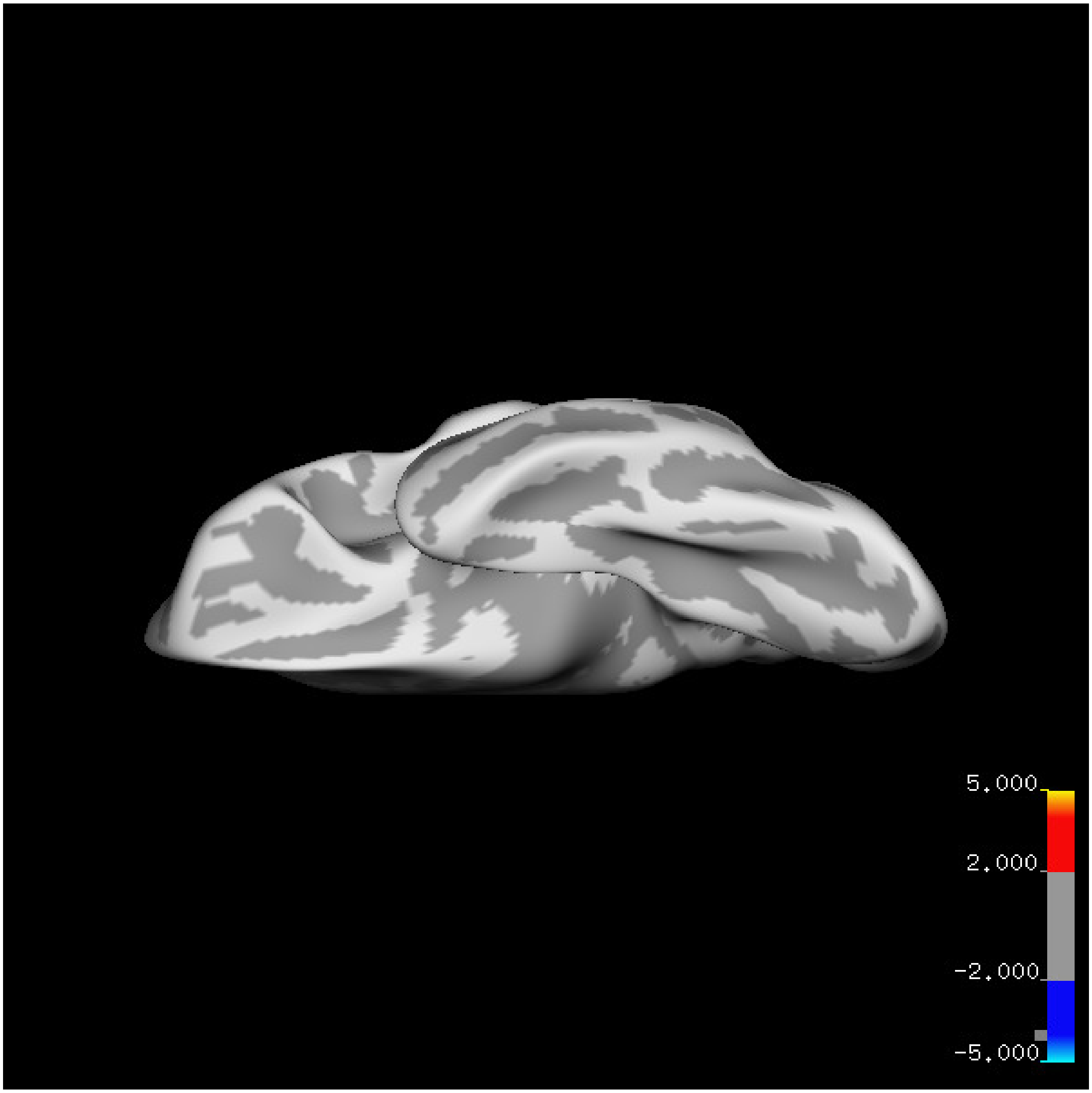
5.000
2.000
-2.000
-5.000

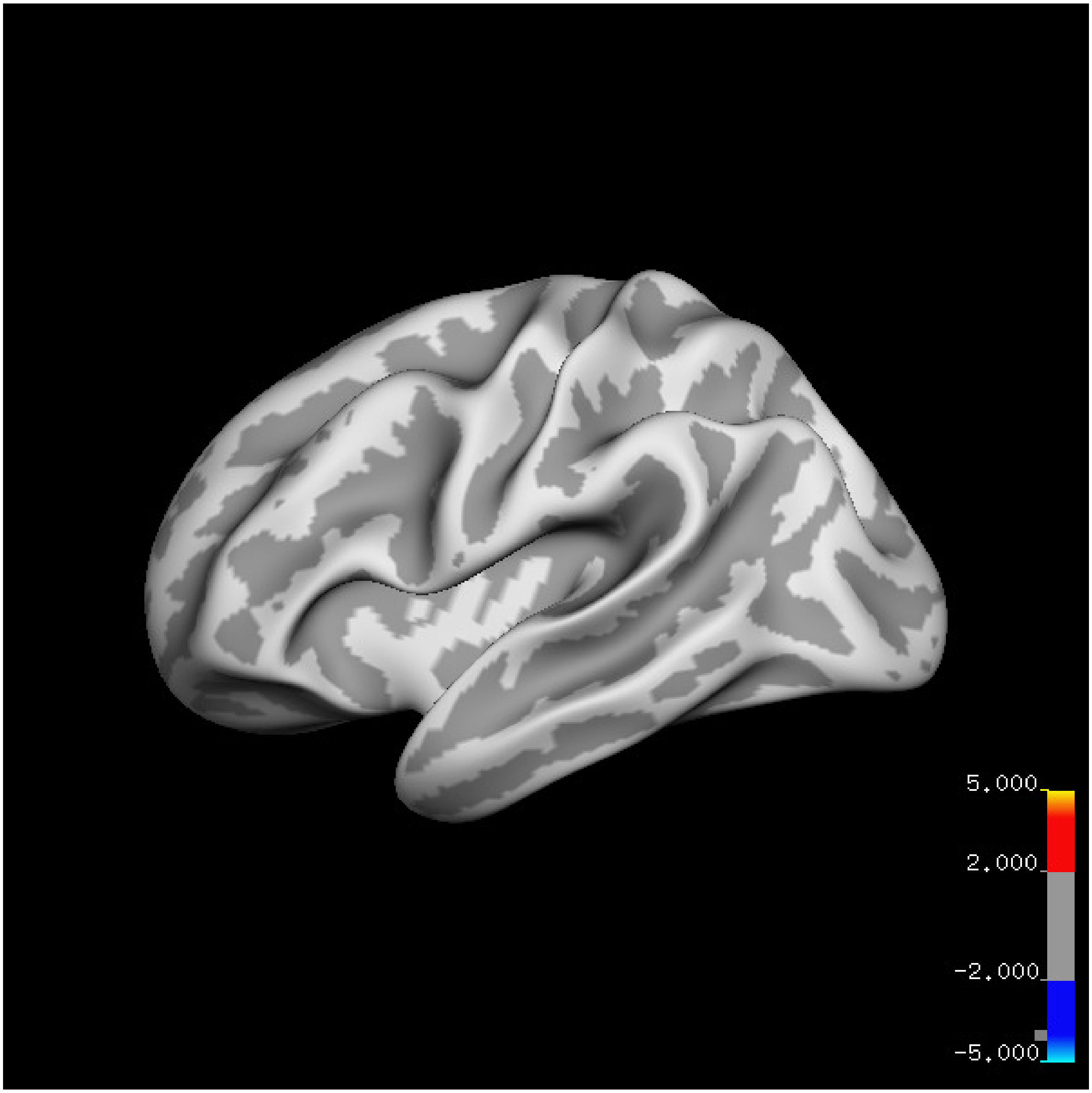
5.000
2.000
-2.000
-5.000

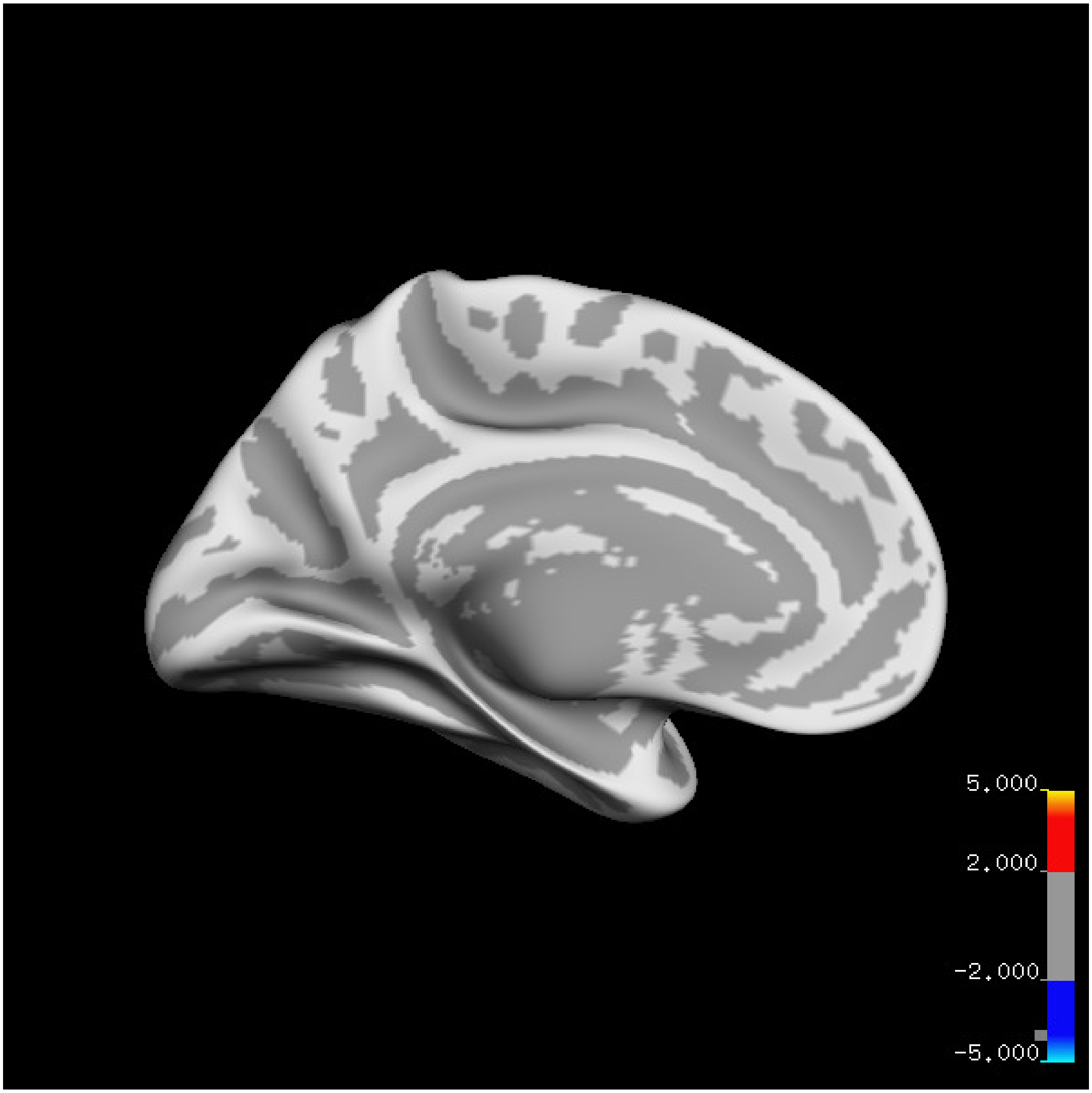

5.000
2.000
-2.000
-5.000

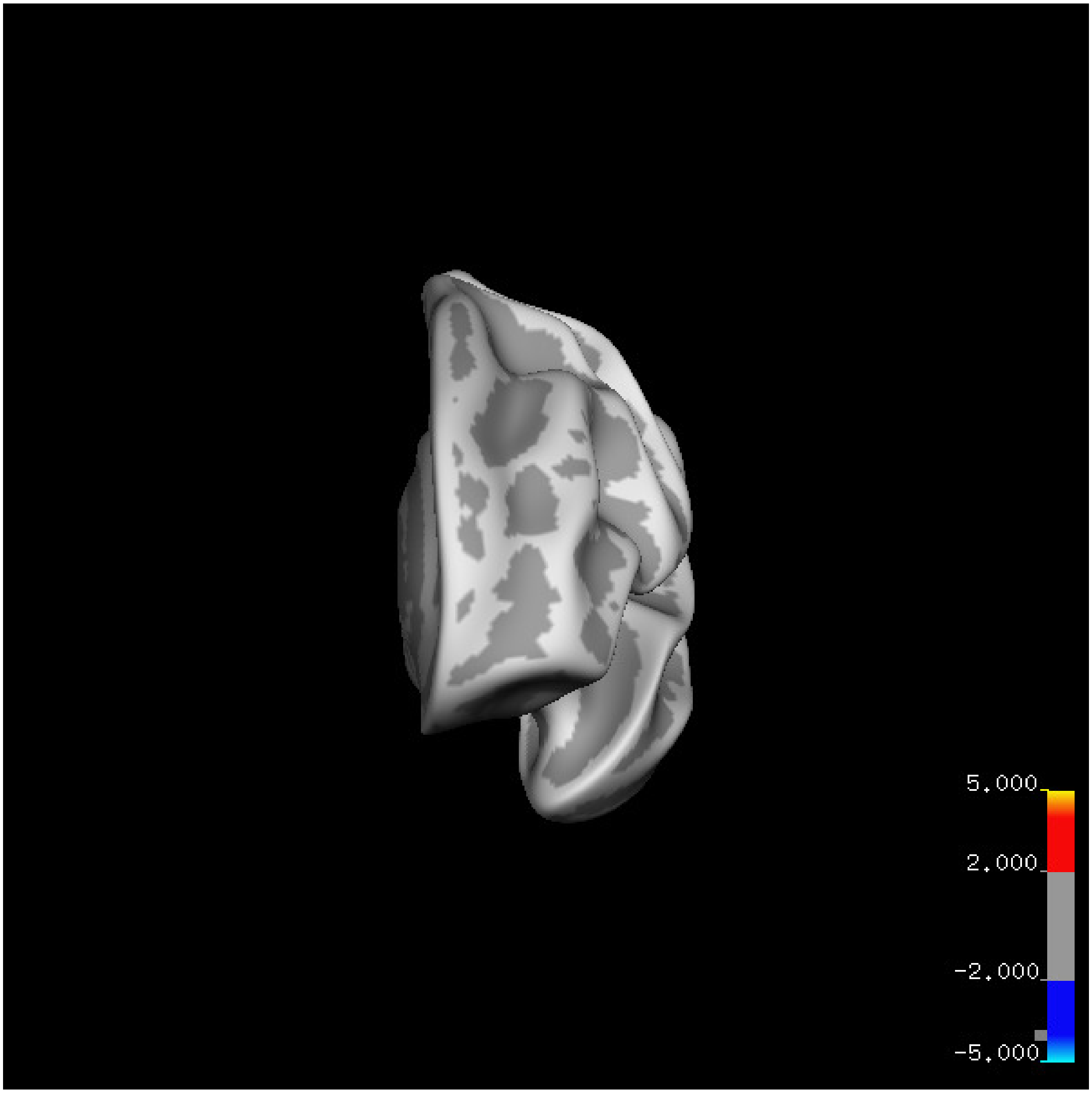

5.000
2.000
-2.000
-5.000

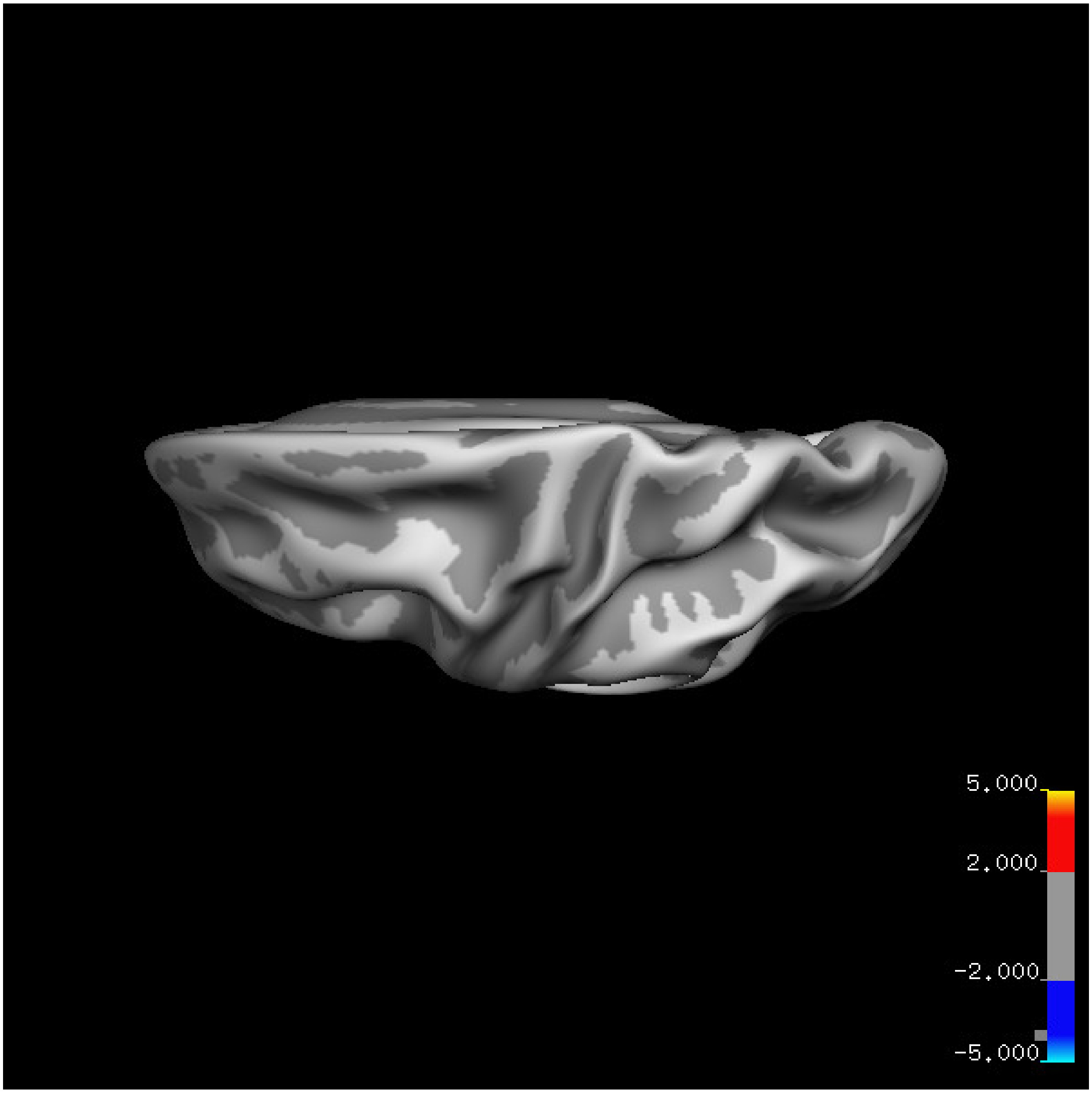

5.000
2.000
-2.000
-5.000

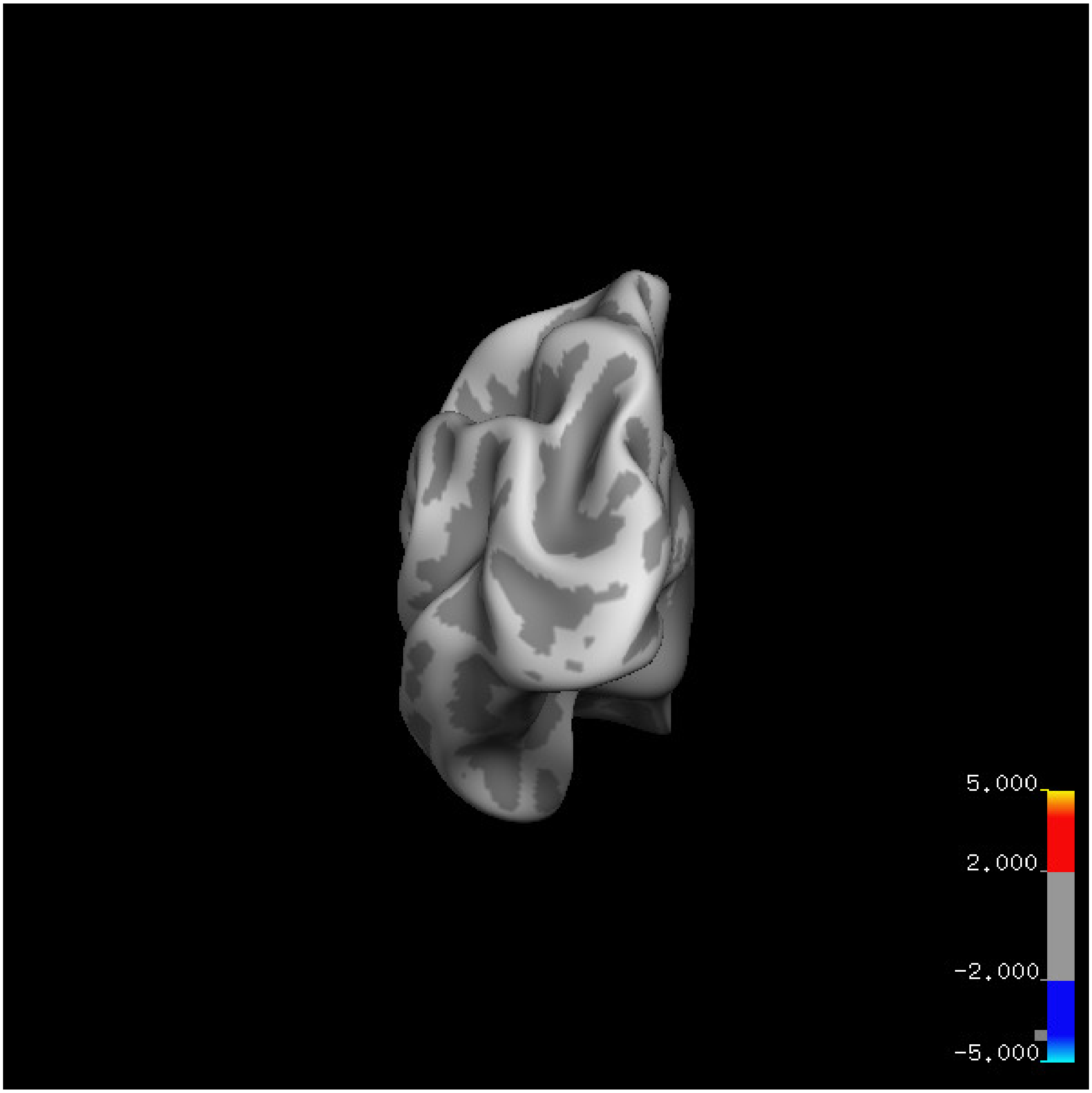
5.000
2.000
-2.000
-5.000

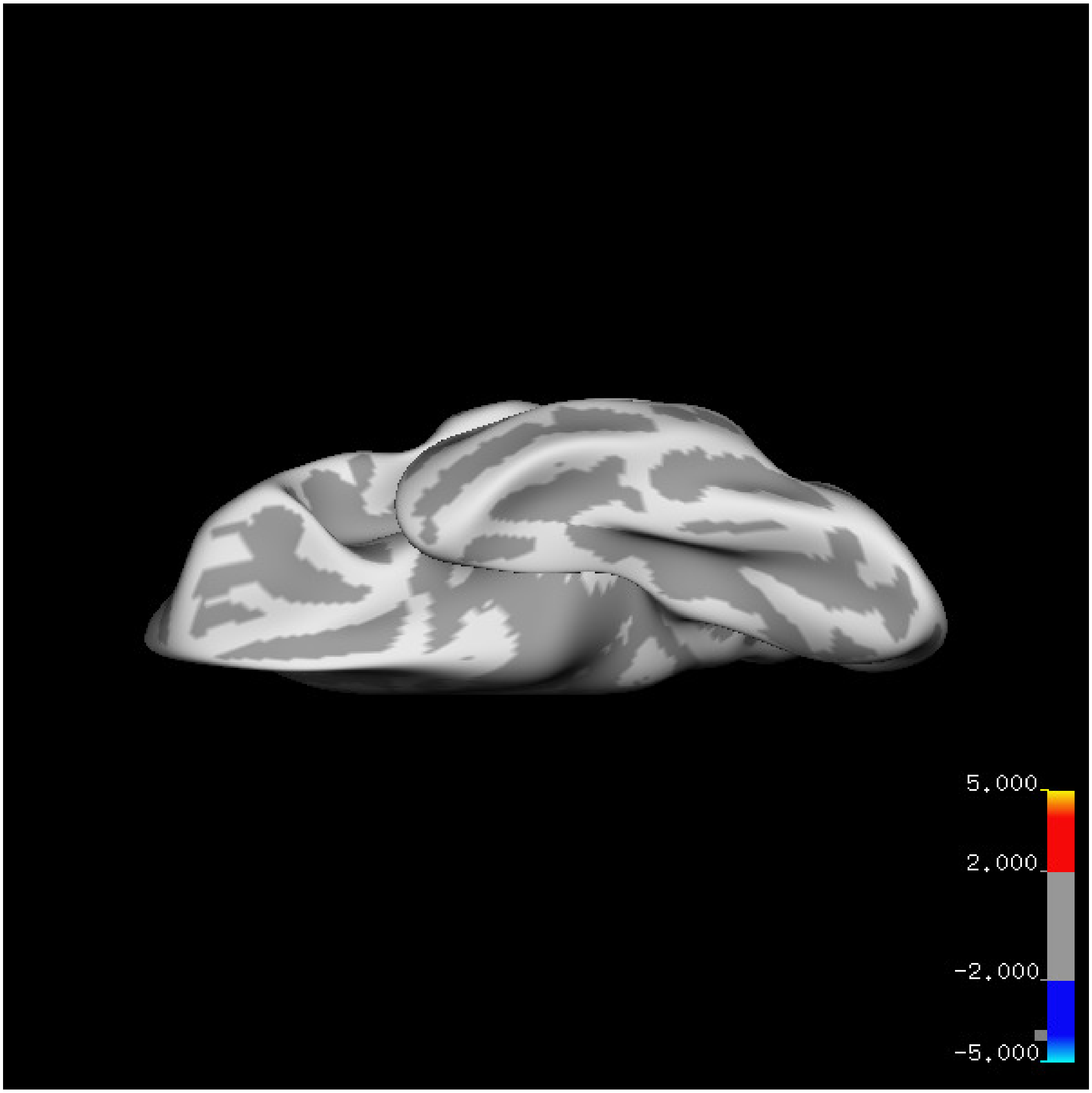
5.000
2.000
-2.000
-5.000

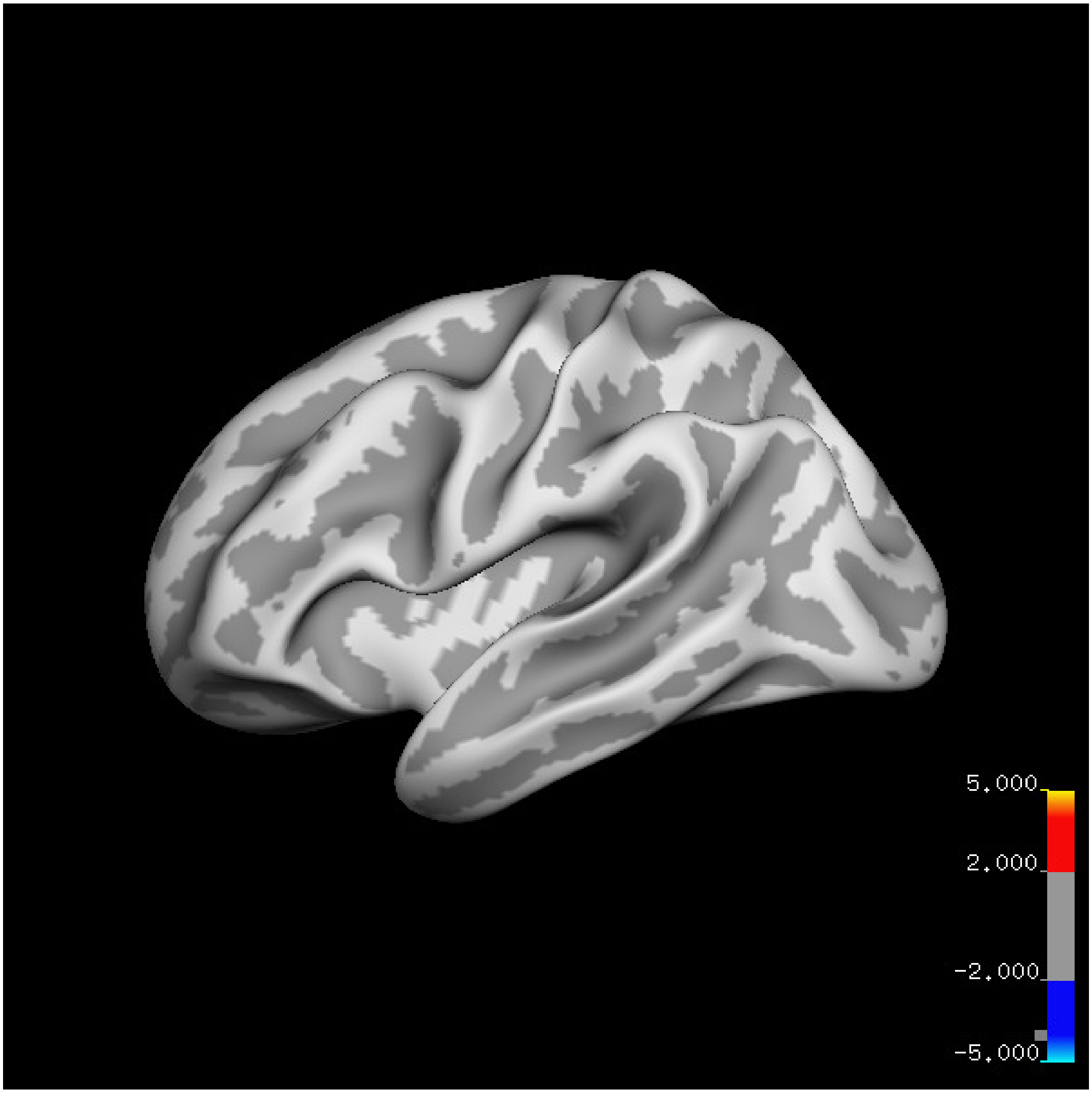
5.000
2.000
-2.000
-5.000

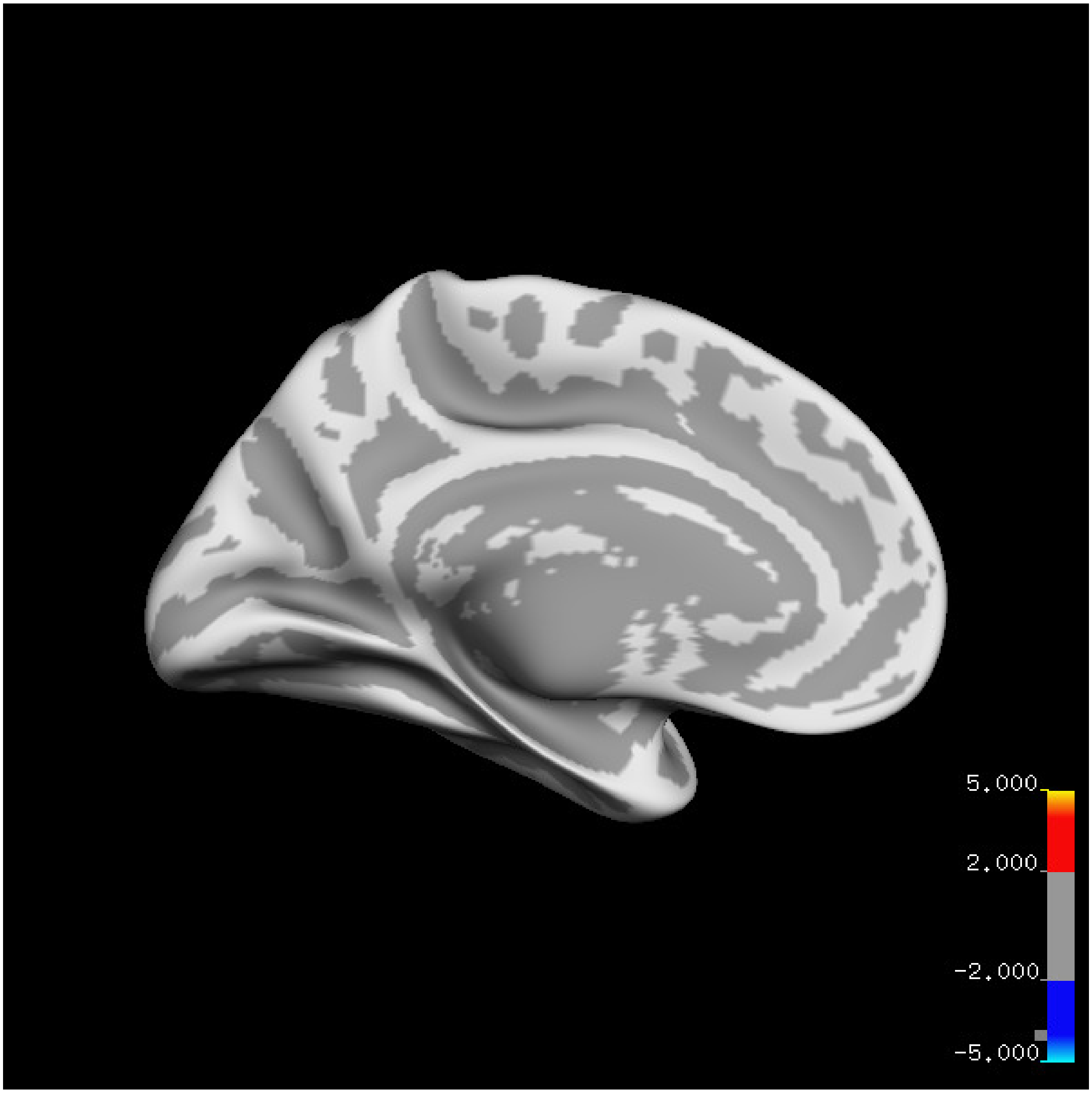
5.000
2.000
-2.000
-5.000

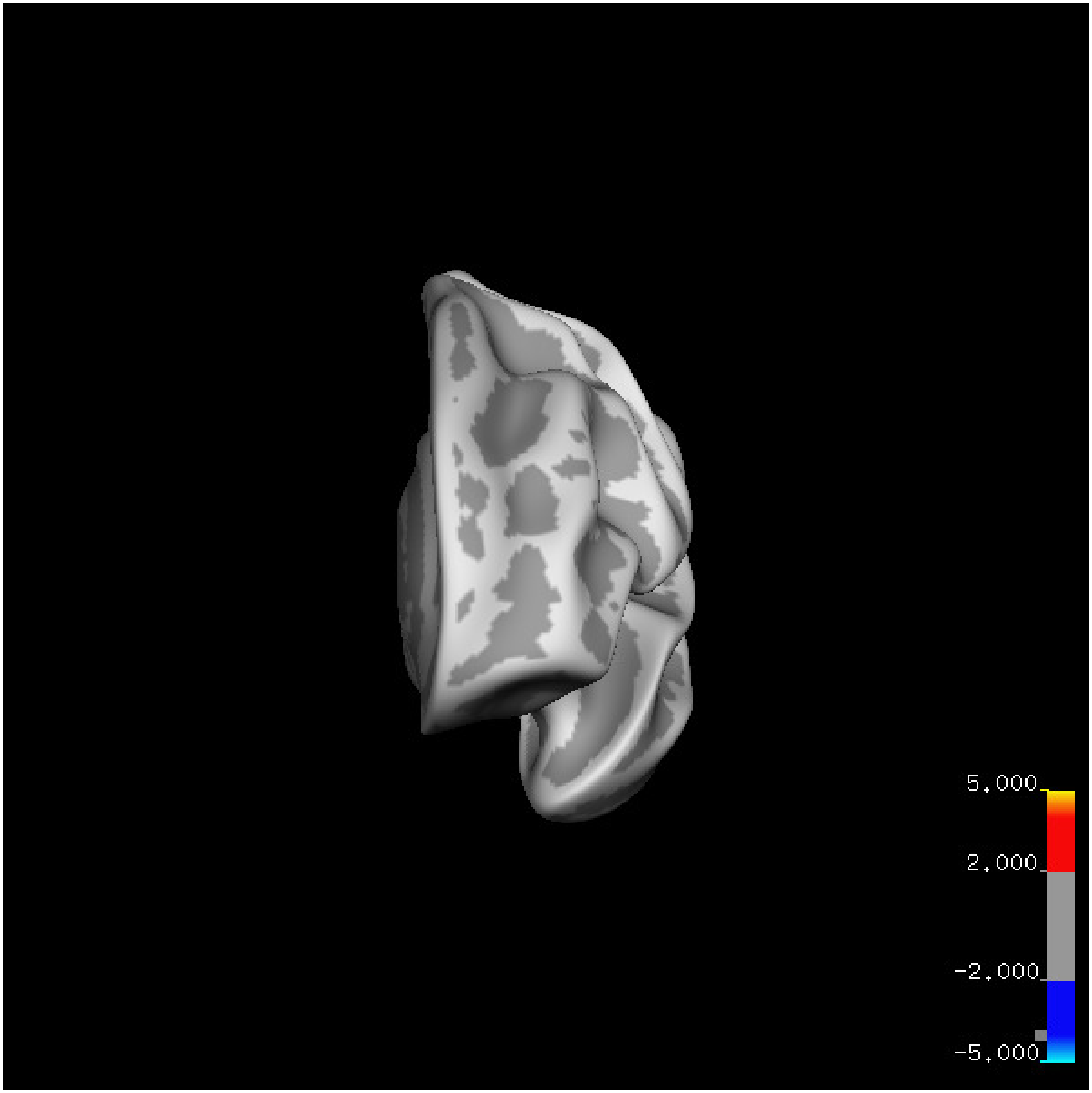
5.000
2.000
-2.000
-5.000

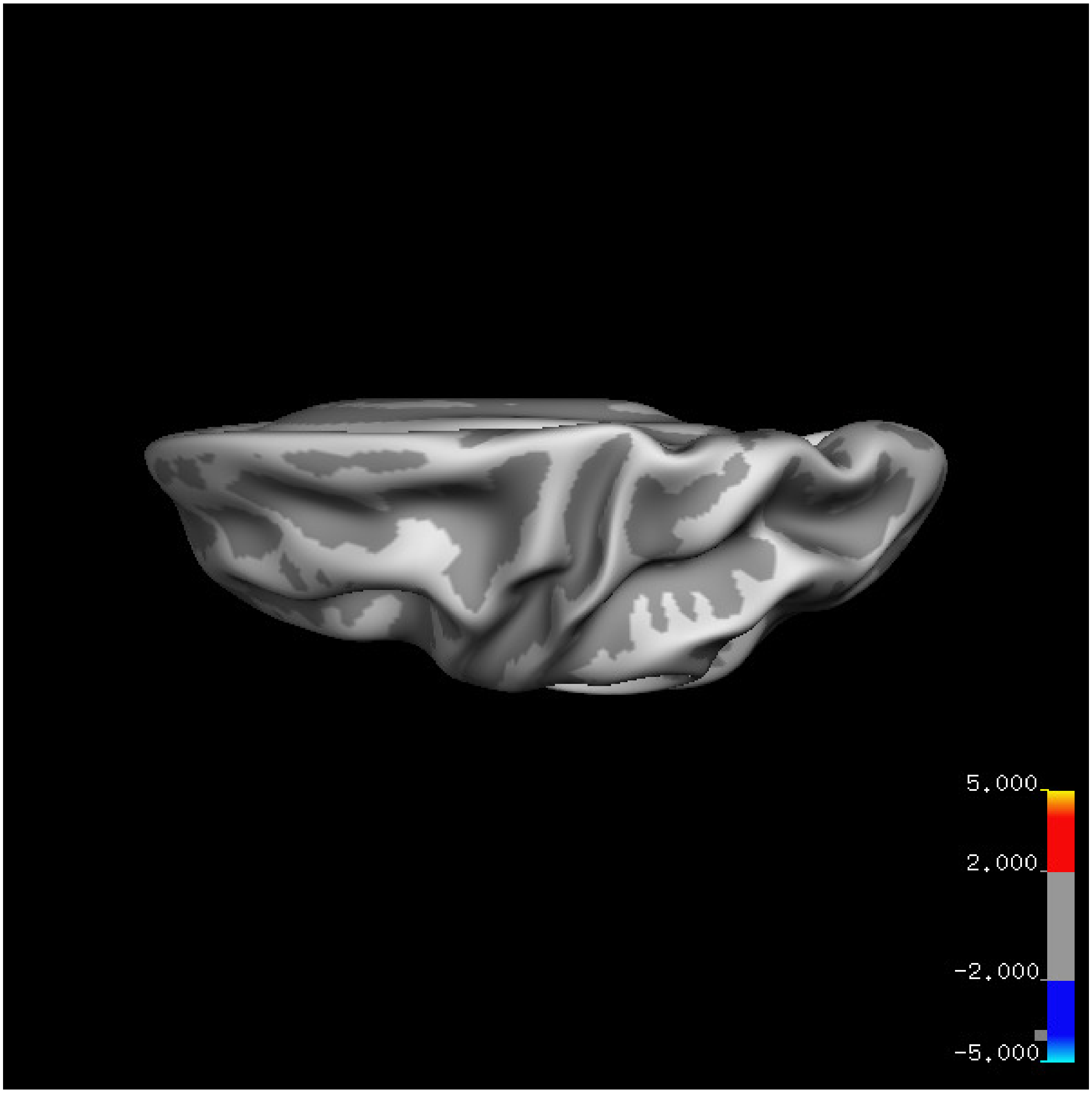


Cluster summary is shown below.

```
## # Cluster Growing Summary (mri_surfcluster)

## # $Id: mri_surfcluster.c,v 1.57.2.3 2016/11/17 18:19:42 zkaufman Exp $

## # $Id: mrisurf.c,v 1.781.2.6 2016/12/27 16:47:14 zkaufman Exp $

## # CreationTime 2021/10/04-13:58:36-GMT

## # cmdline mri_surfcluster.bin --in surface_analyses/results/cluster_comparison/fsaverage6.sm15.lh/ar

## # cwd /gpfs/projects02/p274/cluster/projects/p003-sleep_b

## # sysname  Linux
```

```
## # hostname c1-18
## # machine  x86_64
## # FixVertexAreaFlag 1
## # FixSurfClusterArea 1
## #
## # Input      surface_analyses/results/cluster_comparison/fsaverage6.sm15.lh/area.above60.below_mean/g
## # Frame Number      0
## # srcsubj fsaverage6
## # hemi lh
## # surface white
## # group_avg_surface_area 84969.3
## # group_avg_vtxarea_loaded 1
## # annot aparc
## # SUBJECTS_DIR /cluster/projects/p274/tools/mri/freesurfer/current/subjects
## # SearchSpace_mm2 79241.3
## # SearchSpace_vtx 37476
## # Bonferroni 0
## # Minimum Threshold 1.3
## # Maximum Threshold infinity
## # Threshold Sign    abs
## # AdjustThreshWhenOneTail 1
## # CW PValue Threshold: 0.01
## # Area Threshold    0 mm^2
## # CSD thresh  1.300000
## # CSD nreps    10000
## # CSD simtype  null-z
## # CSD contrast NA
```

```
## # CSD confint  90.000000
## # Overall max 3.52128 at vertex 37618
## # Overall min -3.52666 at vertex 24018
## # NClusters          0
## # FixMNI = 0
## #
## # ClusterNo  Max   VtxMax   Size(mm^2)  MNIX   MNIY   MNIZ    CWP    CWPLow    CWPHi   NVtxs    Wght
```

#### Right hemisphere

##### Age 20-60

**Sleep above mean** Cluster corrected significance plots are shown below.

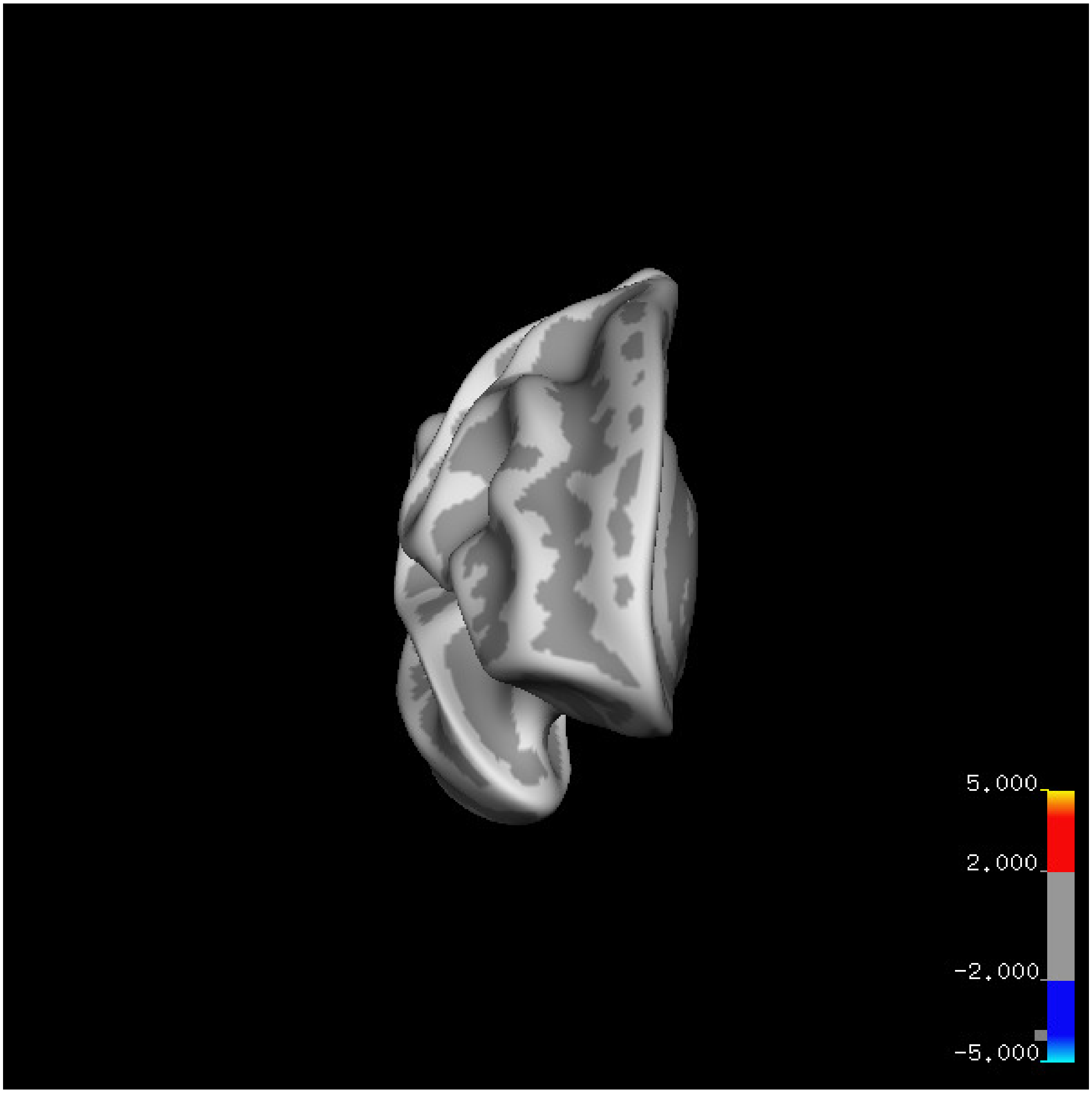

5.000
2.000
-2.000
-5.000

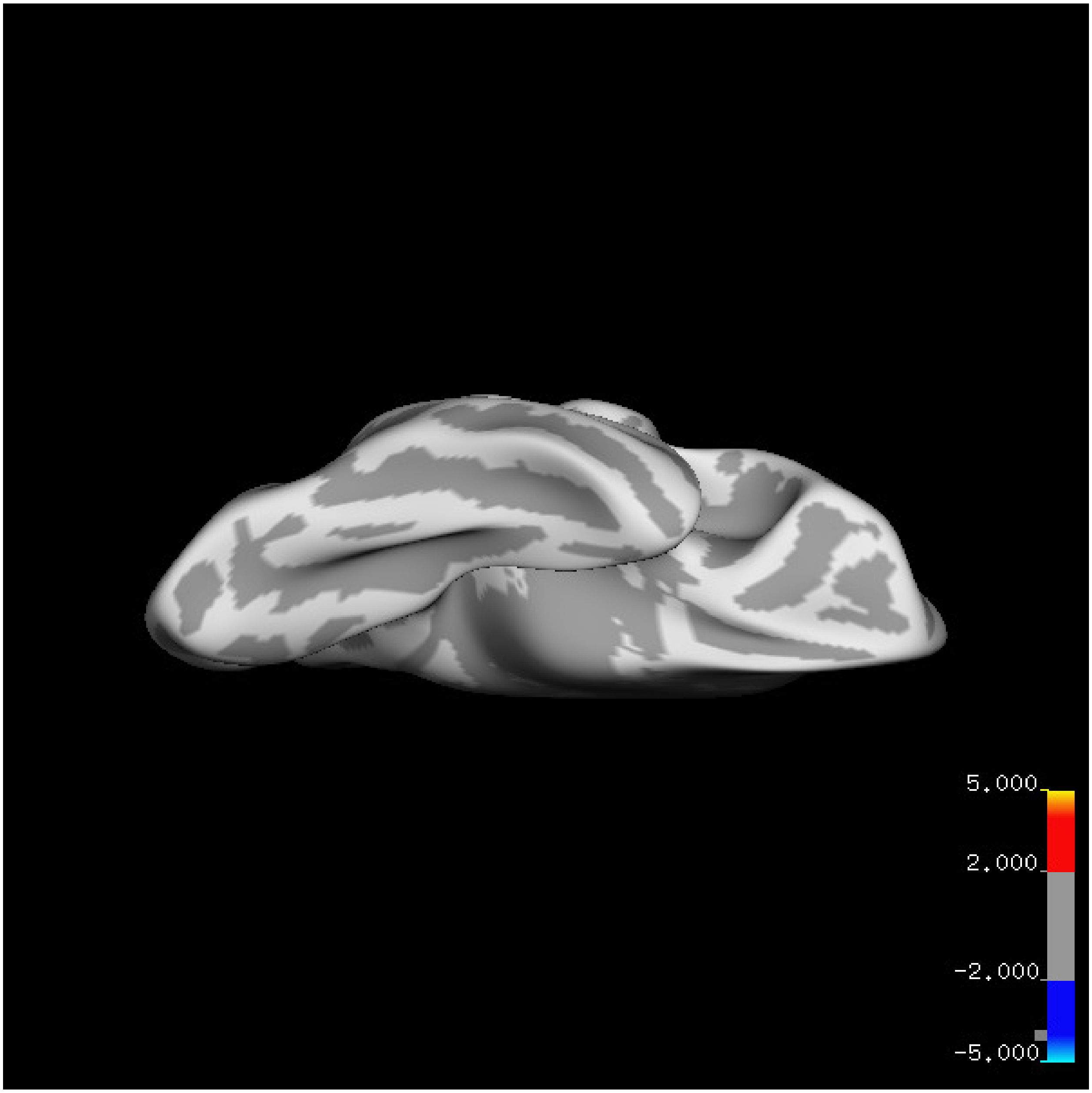
5.000
2.000
-2.000
-5.000

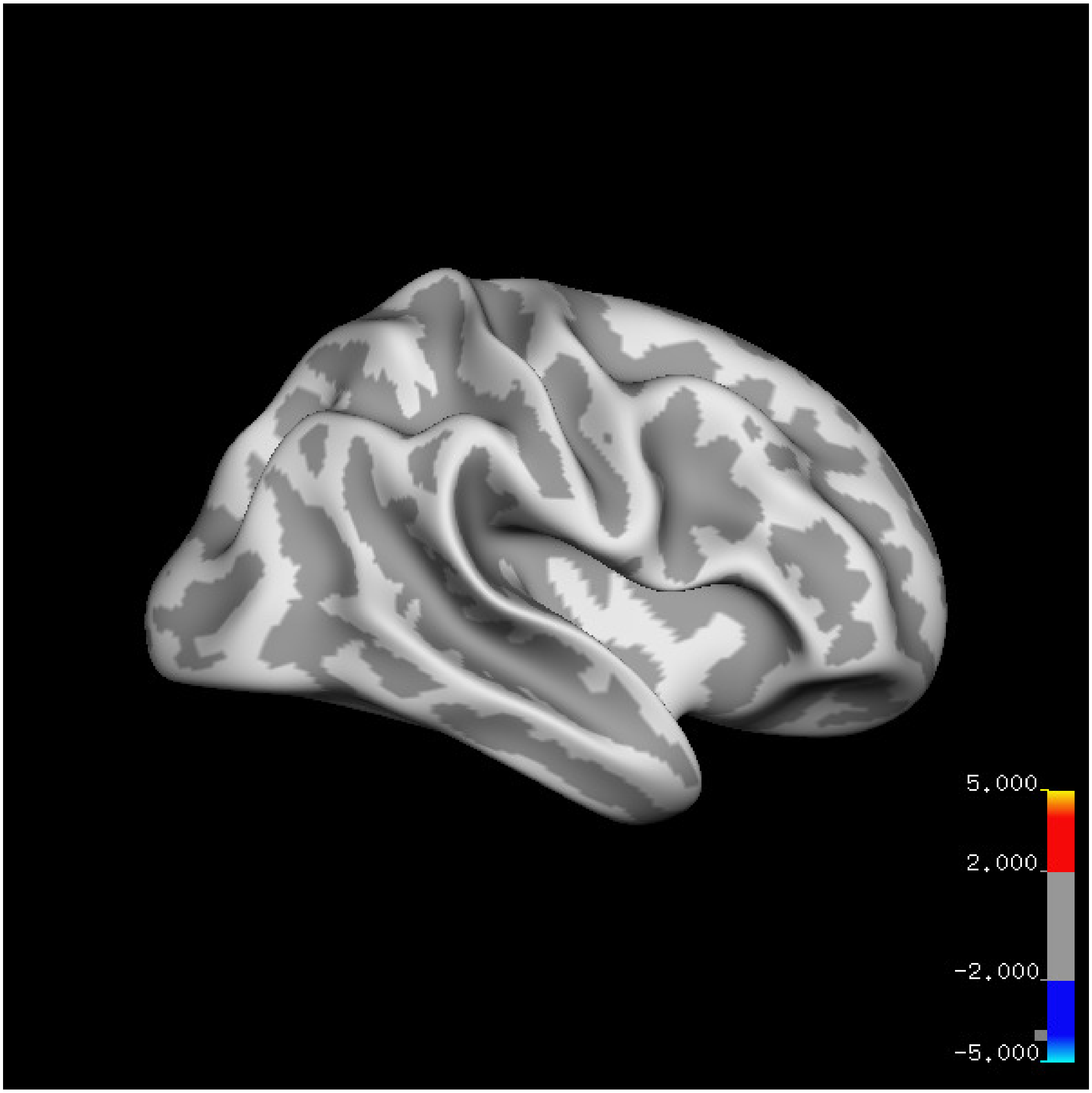
5.000
2.000
-2.000
-5.000

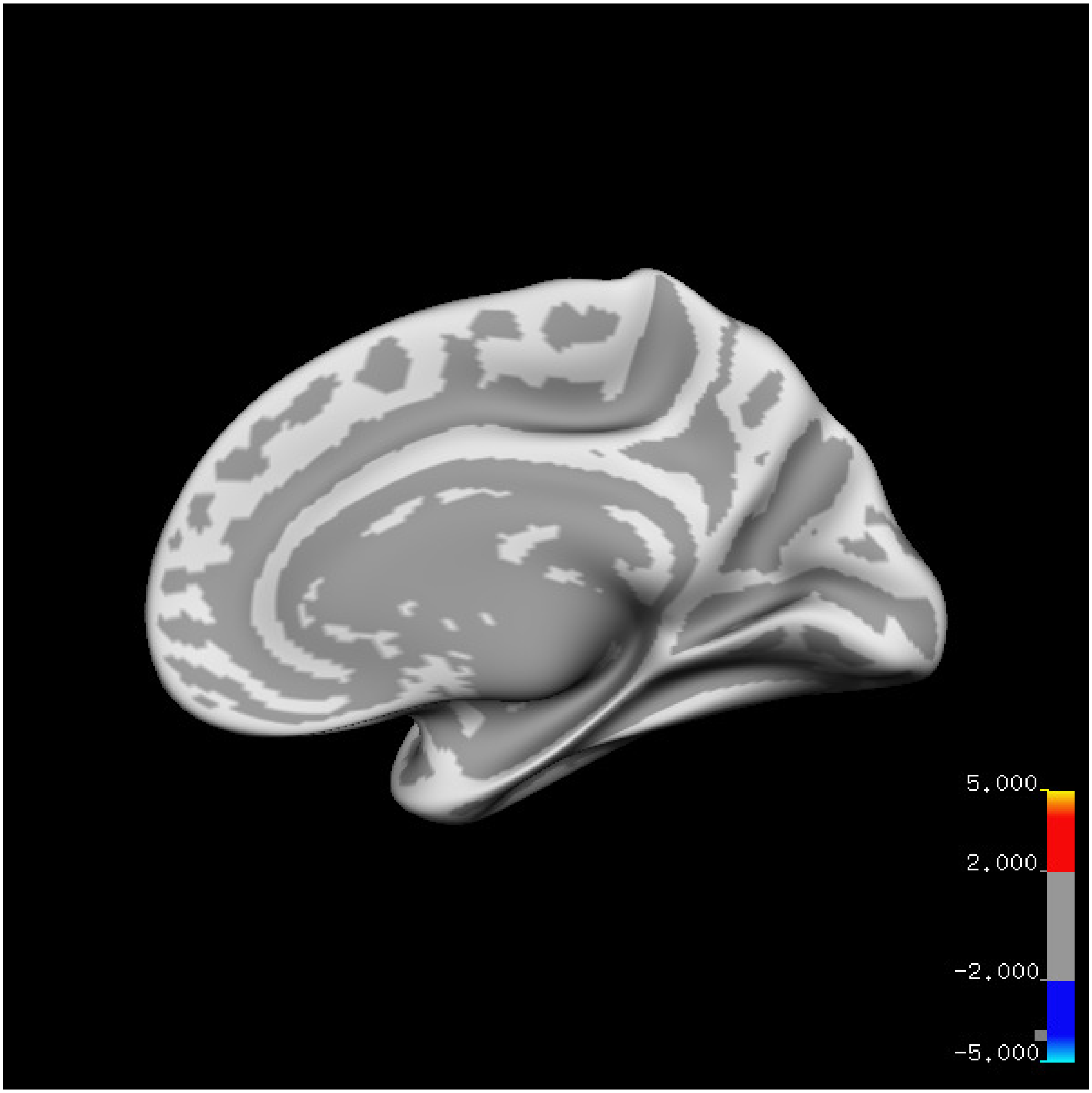
5.000
2.000
-2.000
-5.000

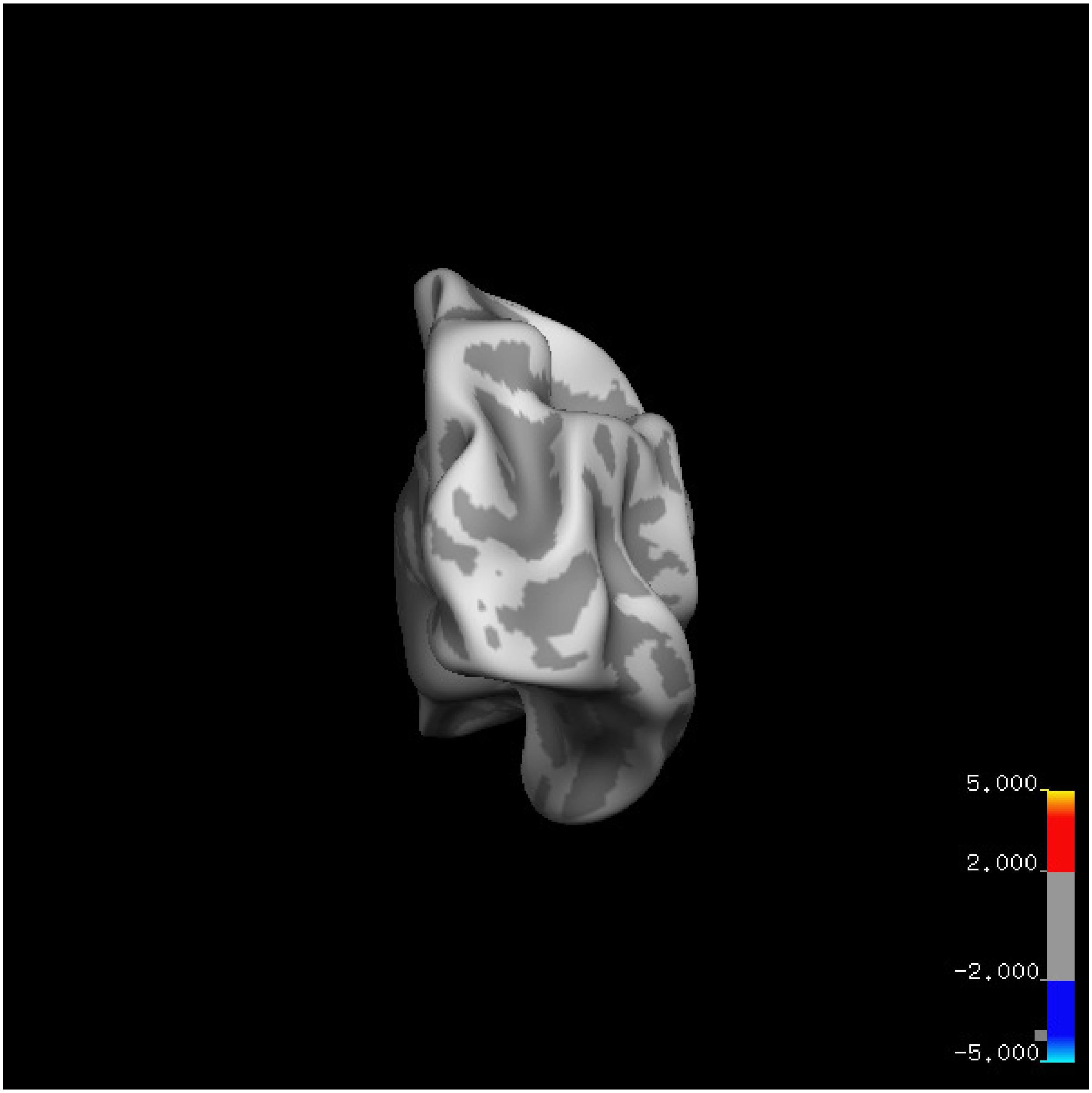
5.000
2.000
-2.000
-5.000

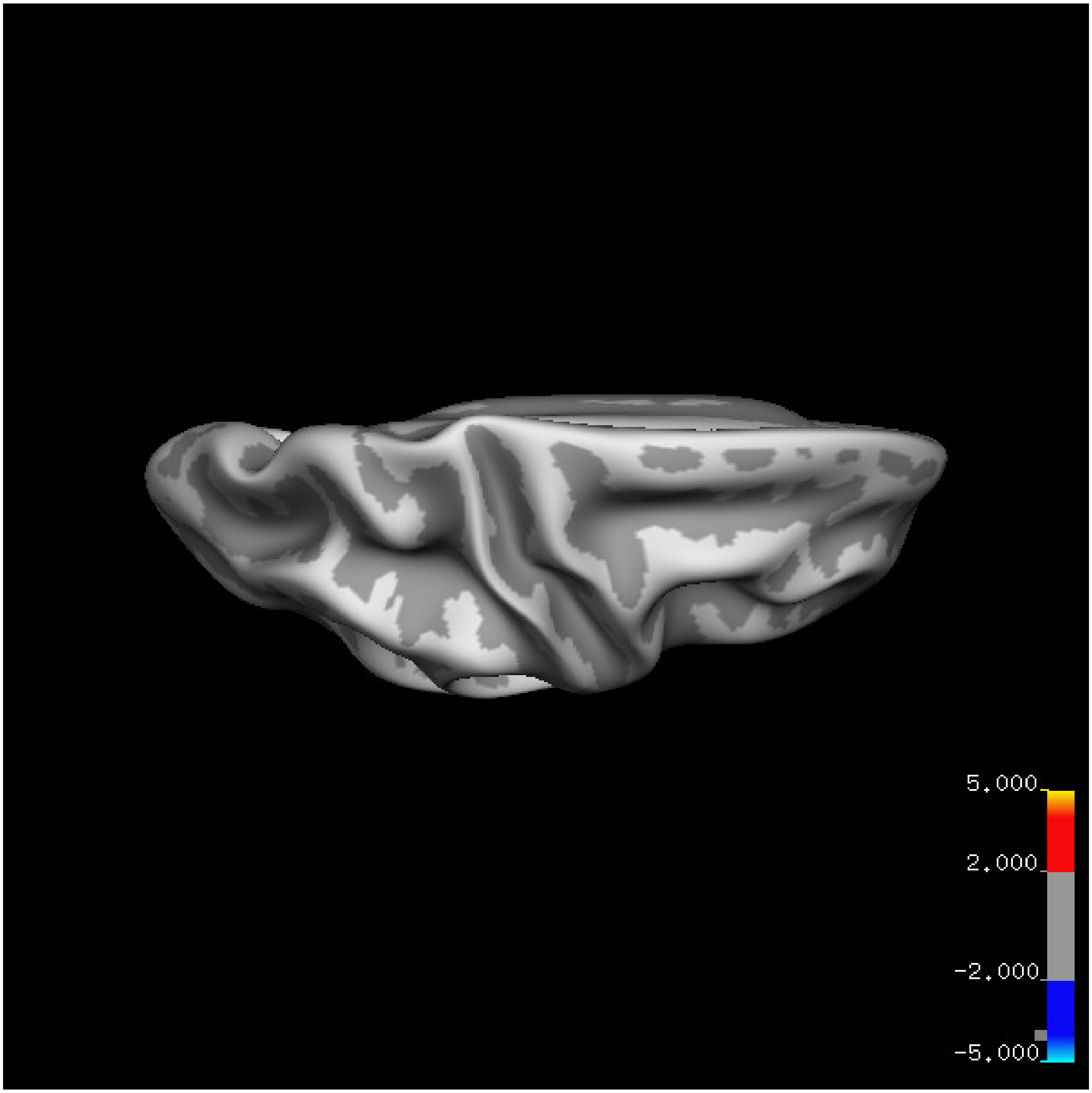
5.000
2.000
-2.000
-5.000

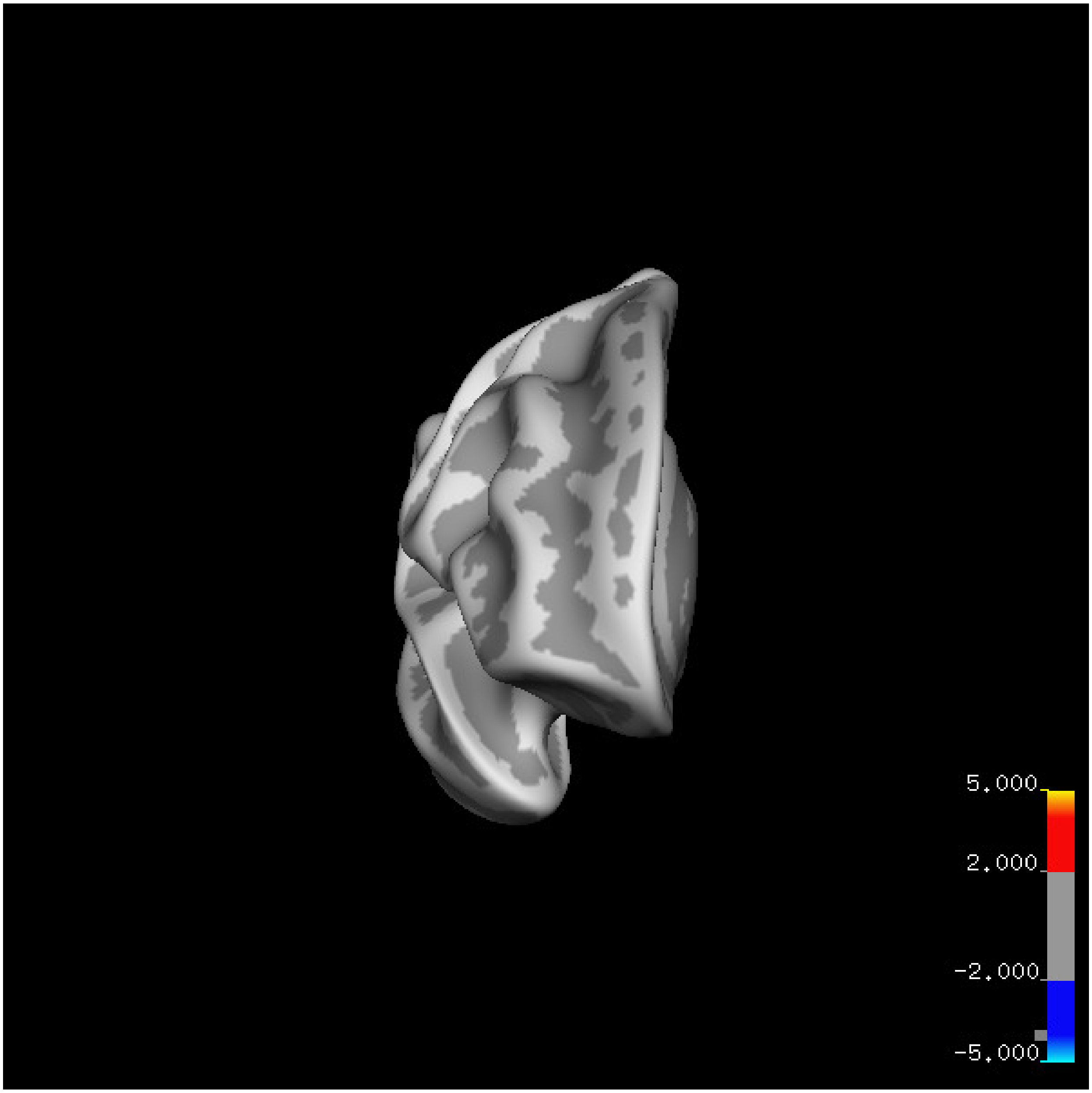
5.000
2.000
-2.000
-5.000

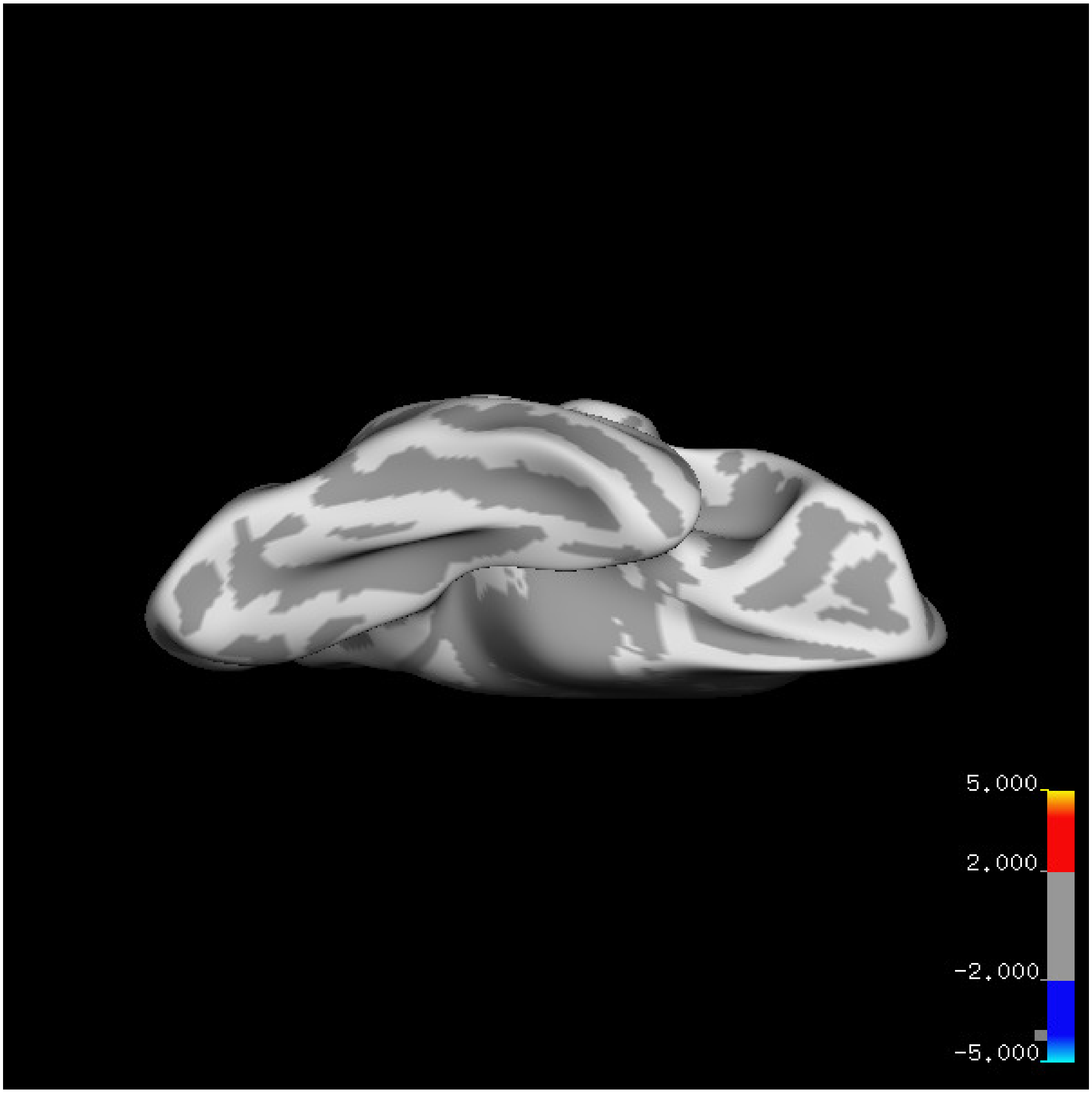
5.000
2.000
-2.000
-5.000

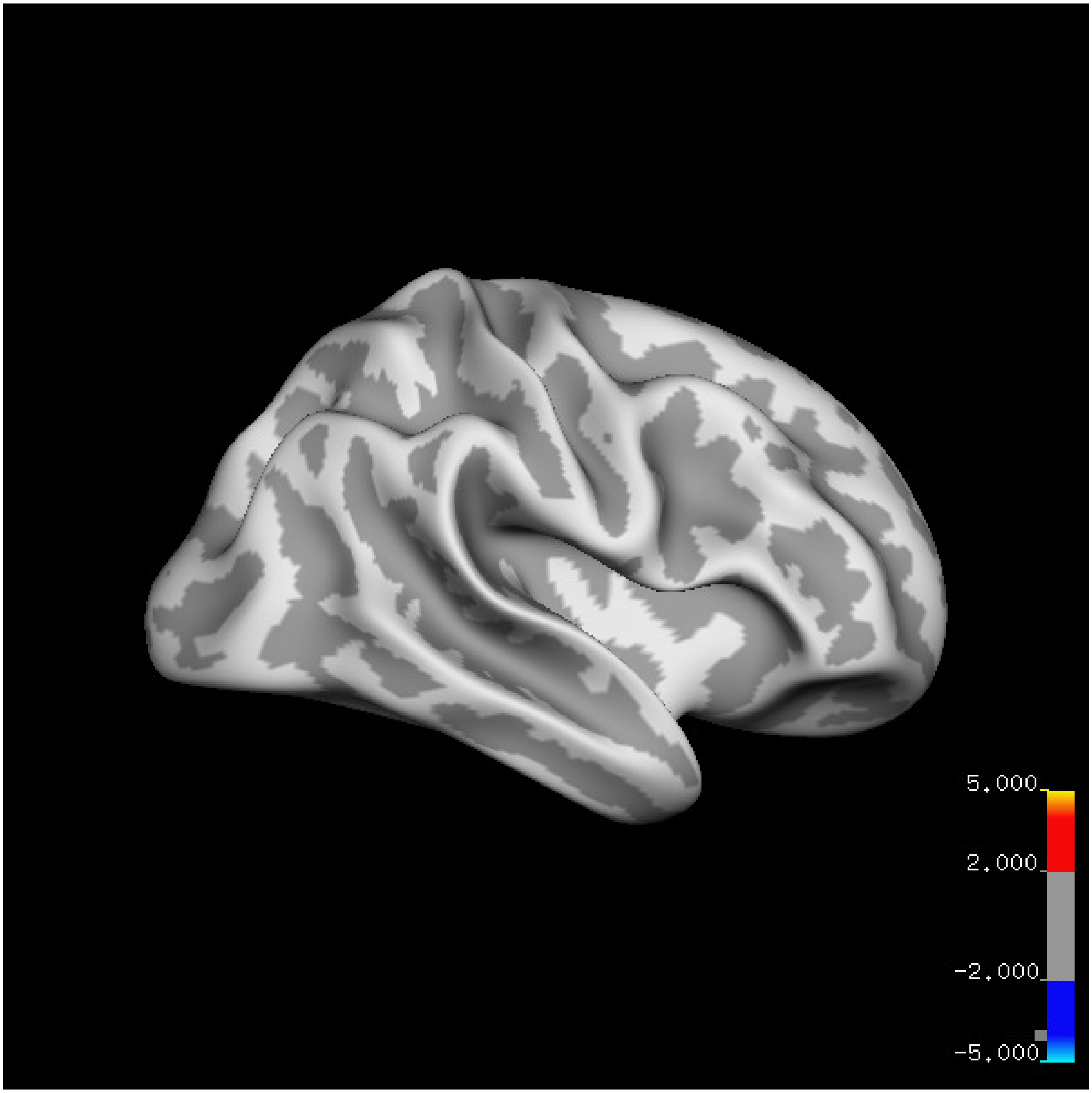
5.000
2.000
-2.000
-5.000

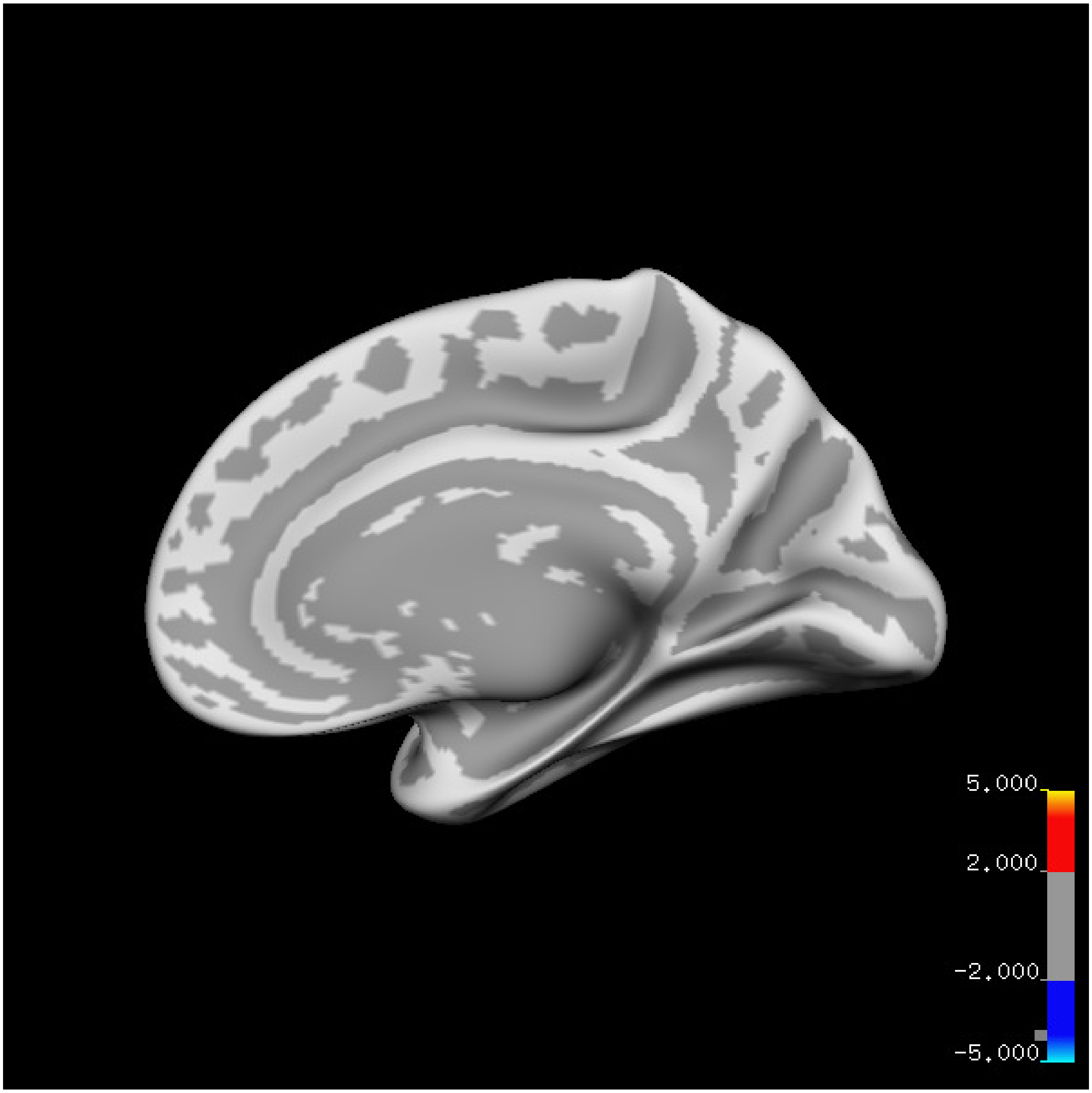

5.000
2.000
-2.000
-5.000

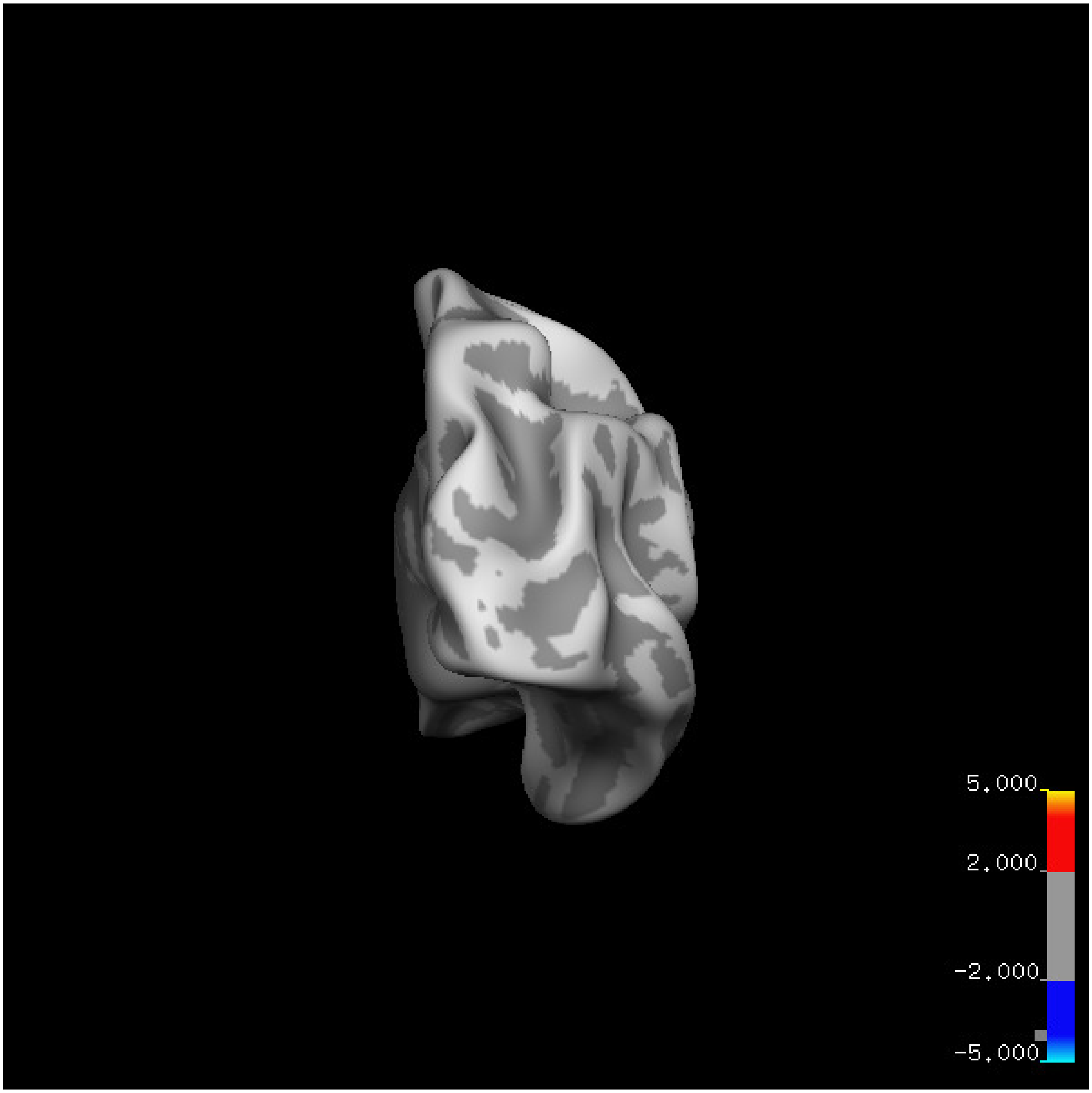
5.000
2.000
-2.000
-5.000

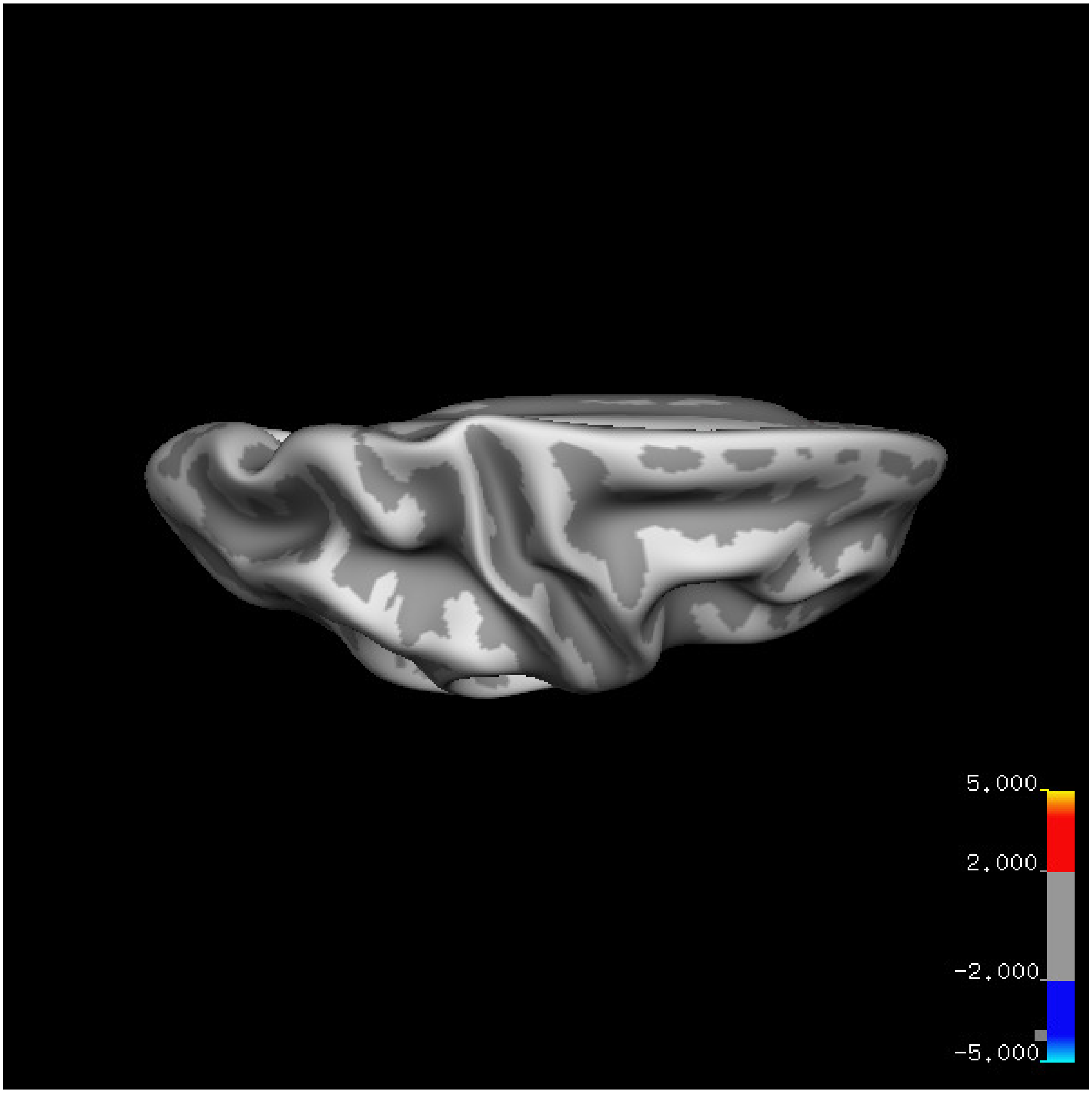


Cluster summary is shown below.

```
## # Cluster Growing Summary (mri_surfcluster)

## # $Id: mri_surfcluster.c,v 1.57.2.3 2016/11/17 18:19:42 zkaufman Exp $

## # $Id: mrisurf.c,v 1.781.2.6 2016/12/27 16:47:14 zkaufman Exp $

## # CreationTime 2021/10/04-10:48:05-GMT

## # cmdline mri_surfcluster.bin --in surface_analyses/results/cluster_comparison/fsaverage6.sm15.rh/ar

## # cwd /gpfs/projects02/p274/cluster/projects/p003-sleep_b

## # sysname  Linux
```

```
## # hostname c1-19
## # machine  x86_64
## # FixVertexAreaFlag 1
## # FixSurfClusterArea 1
## #
## # Input      surface_analyses/results/cluster_comparison/fsaverage6.sm15.rh/area.20_60.above_mean/gl
## # Frame Number      0
## # srcsubj fsaverage6
## # hemi rh
## # surface white
## # group_avg_surface_area 85131.2
## # group_avg_vtxarea_loaded 1
## # annot aparc
## # SUBJECTS_DIR /cluster/projects/p274/tools/mri/freesurfer/current/subjects
## # SearchSpace_mm2 79329.9
## # SearchSpace_vtx 37471
## # Bonferroni 0
## # Minimum Threshold 1.3
## # Maximum Threshold infinity
## # Threshold Sign    abs
## # AdjustThreshWhenOneTail 1
## # CW PValue Threshold: 0.01
## # Area Threshold    0 mm^2
## # CSD thresh  1.300000
## # CSD nreps    10000
## # CSD simtype  null-z
## # CSD contrast NA
```

```
## # CSD confint  90.000000
## # Overall max 2.69234 at vertex 28685
## # Overall min -2.59333 at vertex 5227
## # NClusters          0
## # FixMNI = 0
## #
## # ClusterNo  Max   VtxMax   Size(mm^2)  MNIX   MNIY   MNIZ    CWP    CWPLow    CWPHi   NVtxs    Wght
```

**Sleep below mean** Cluster corrected significance plots are shown below.

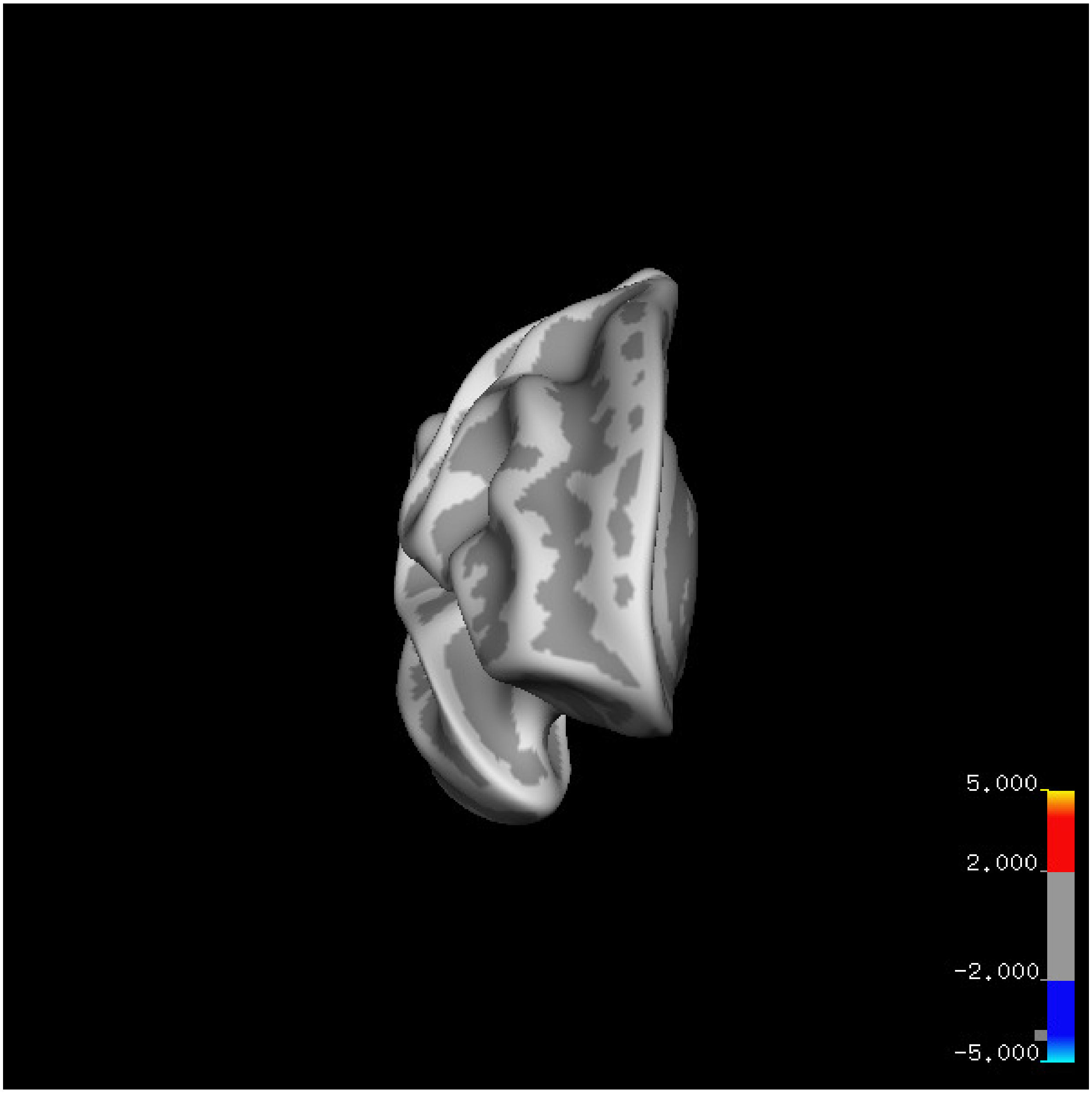

5.000
2.000
-2.000
-5.000

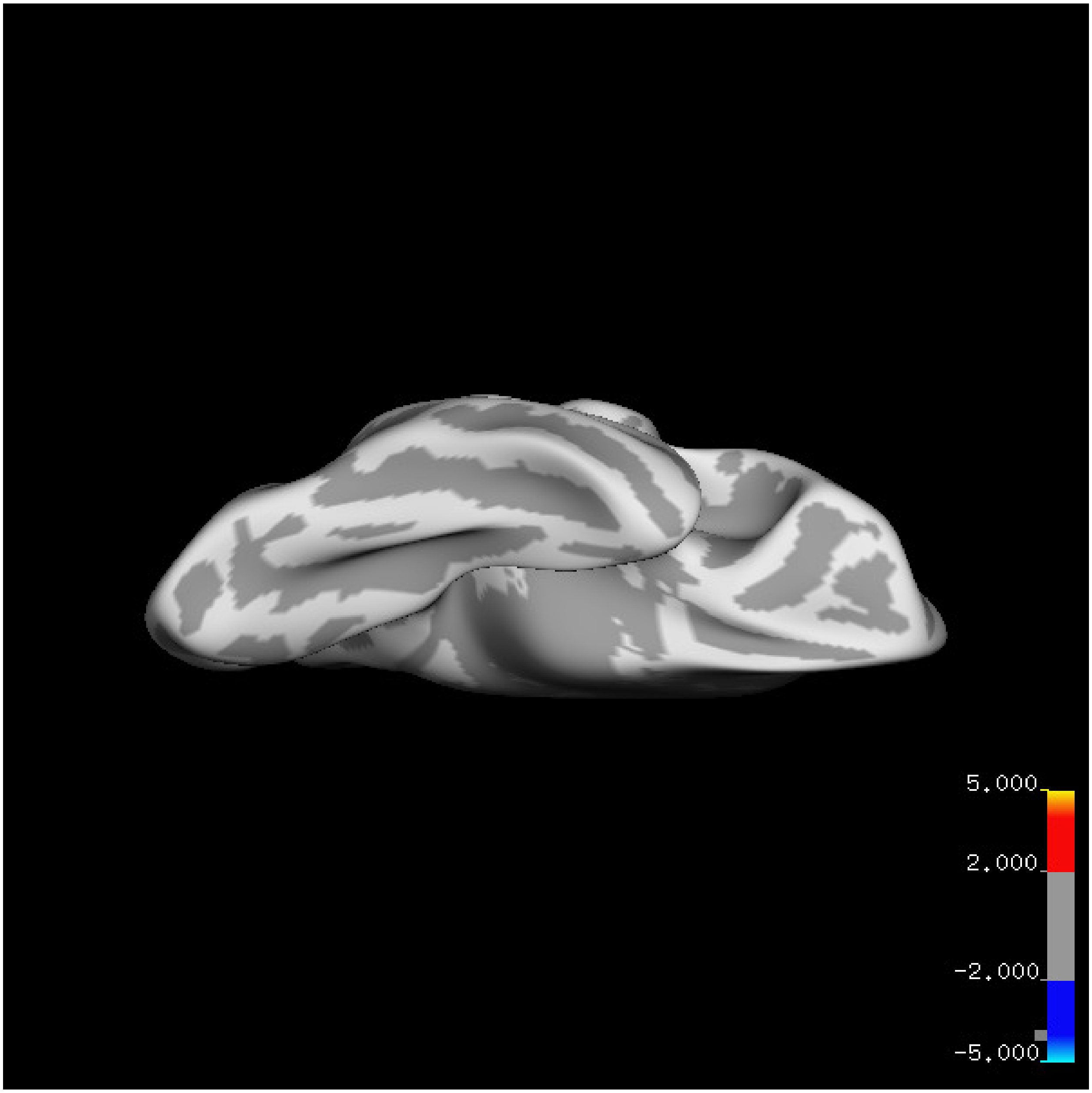
5.000
2.000
-2.000
-5.000

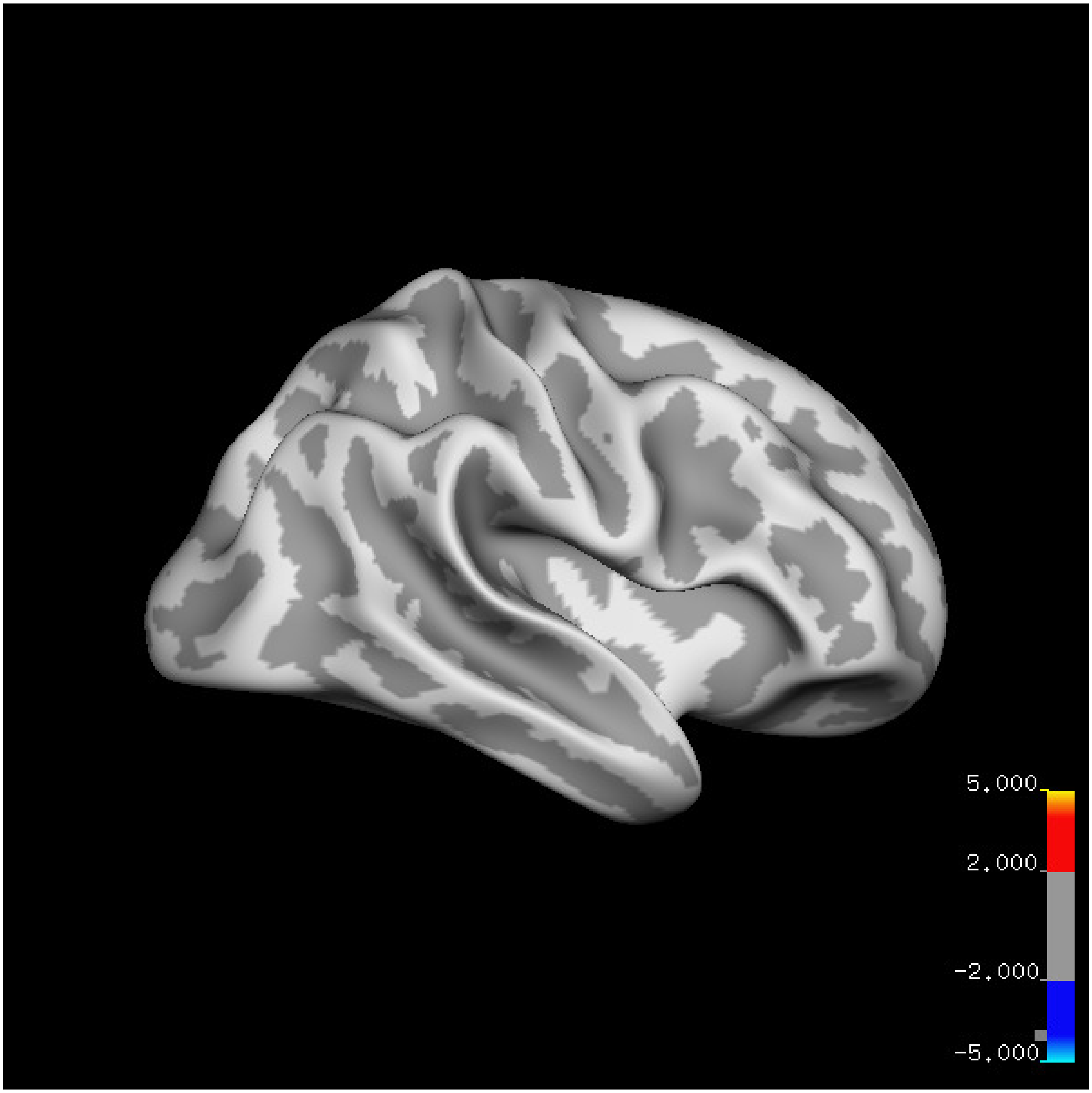
5.000
2.000
-2.000
-5.000

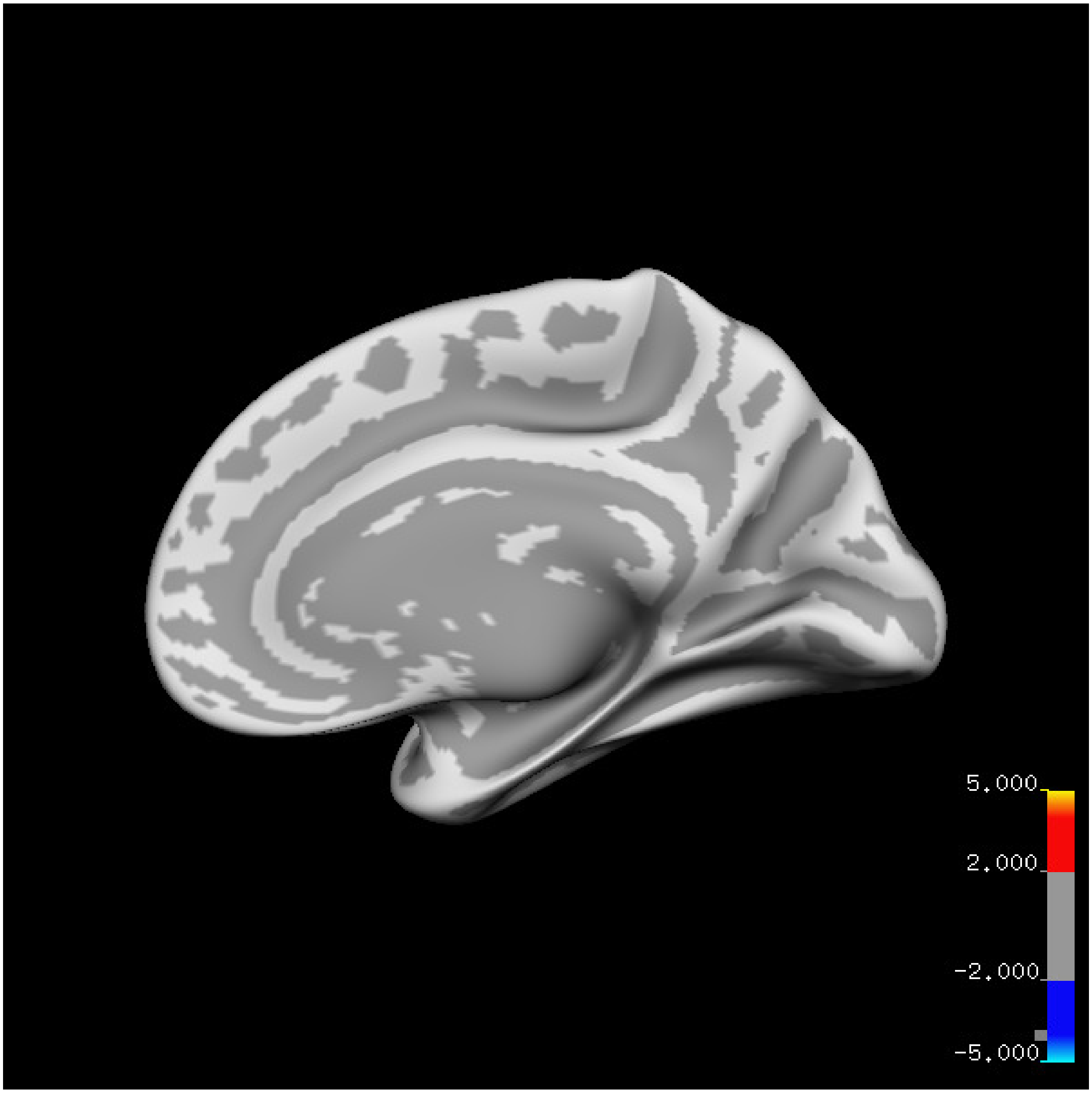
5.000
2.000
-2.000
-5.000

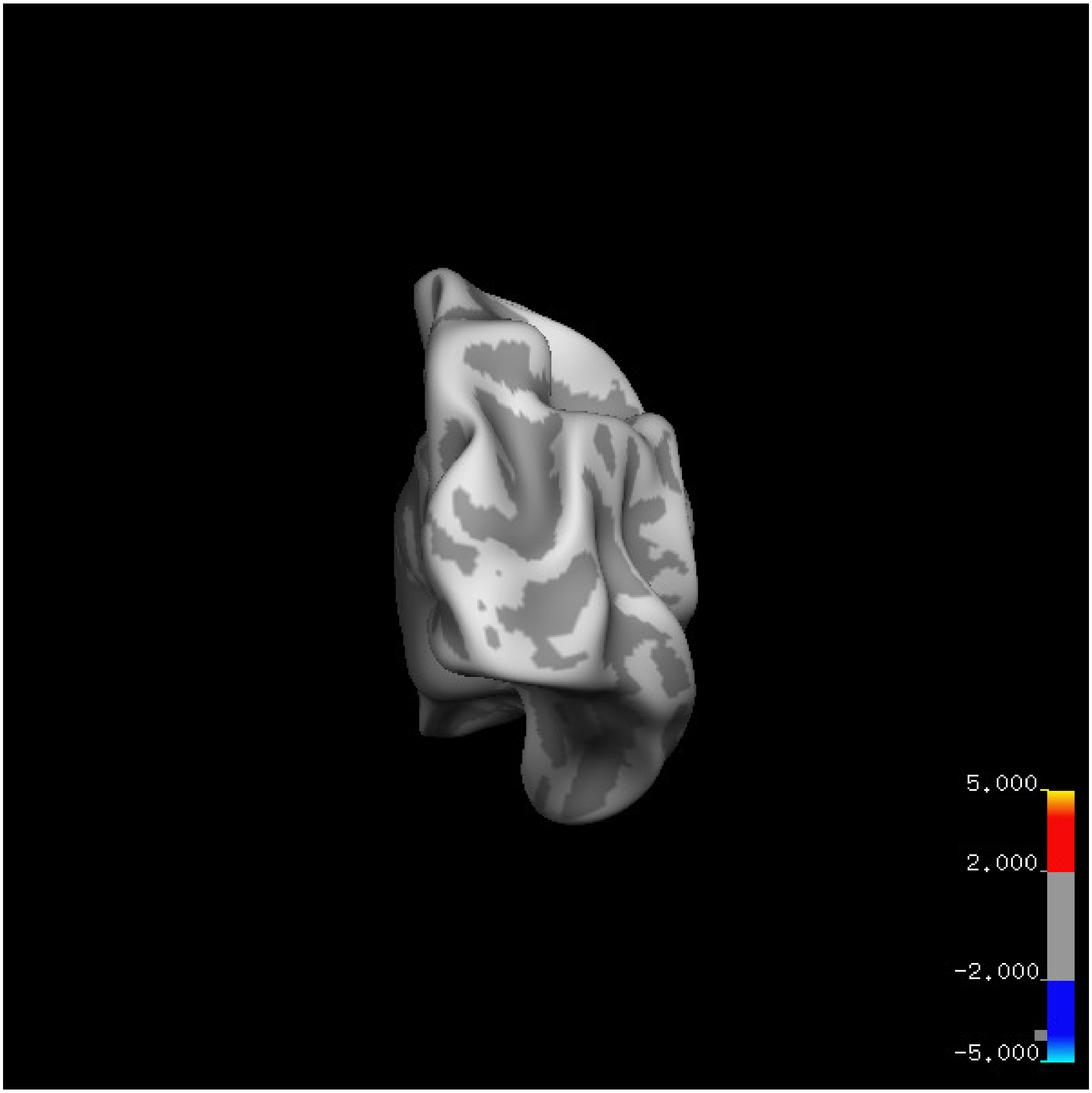
5.000
2.000
-2.000
-5.000

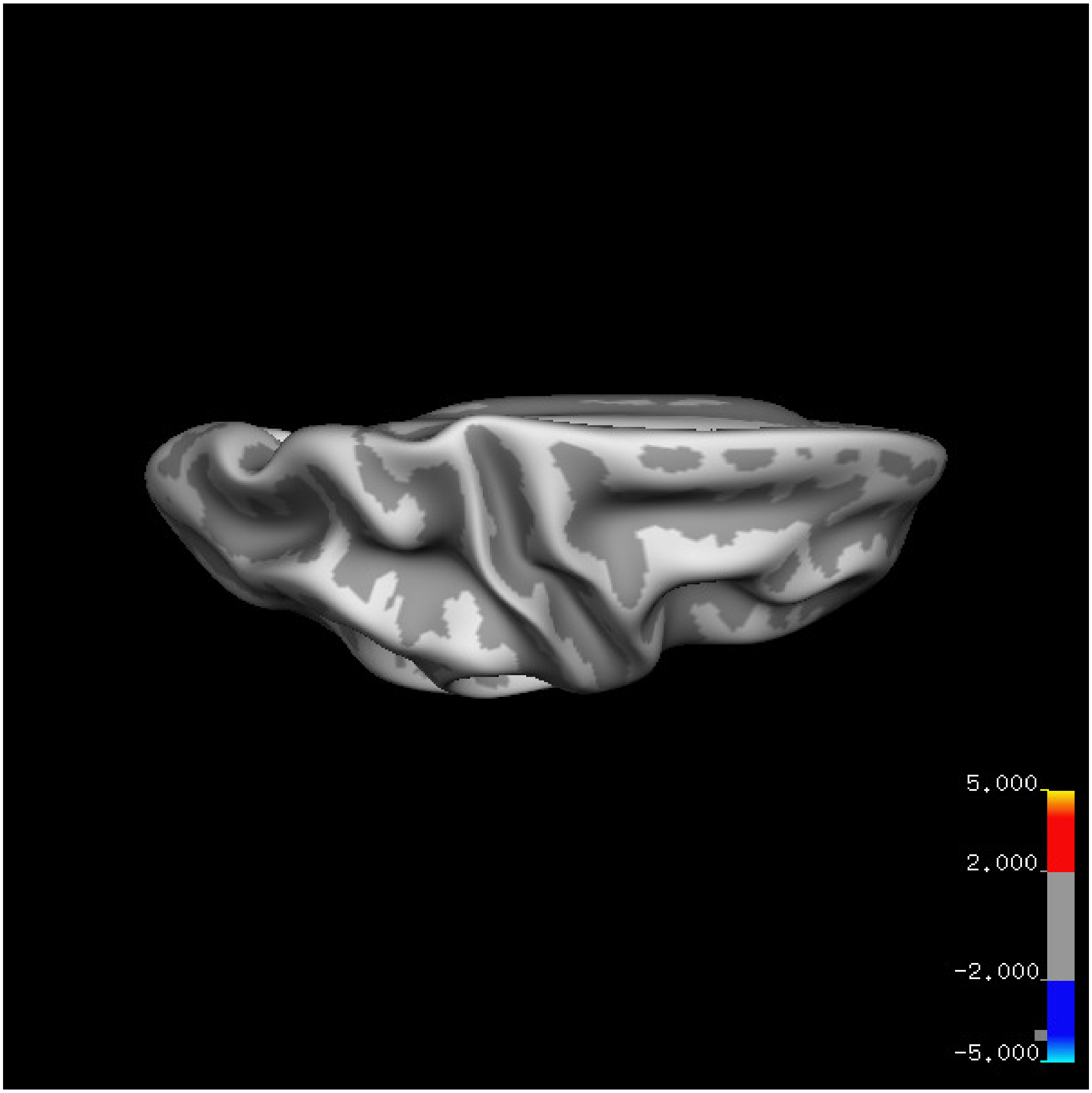
5.000
2.000
-2.000
-5.000

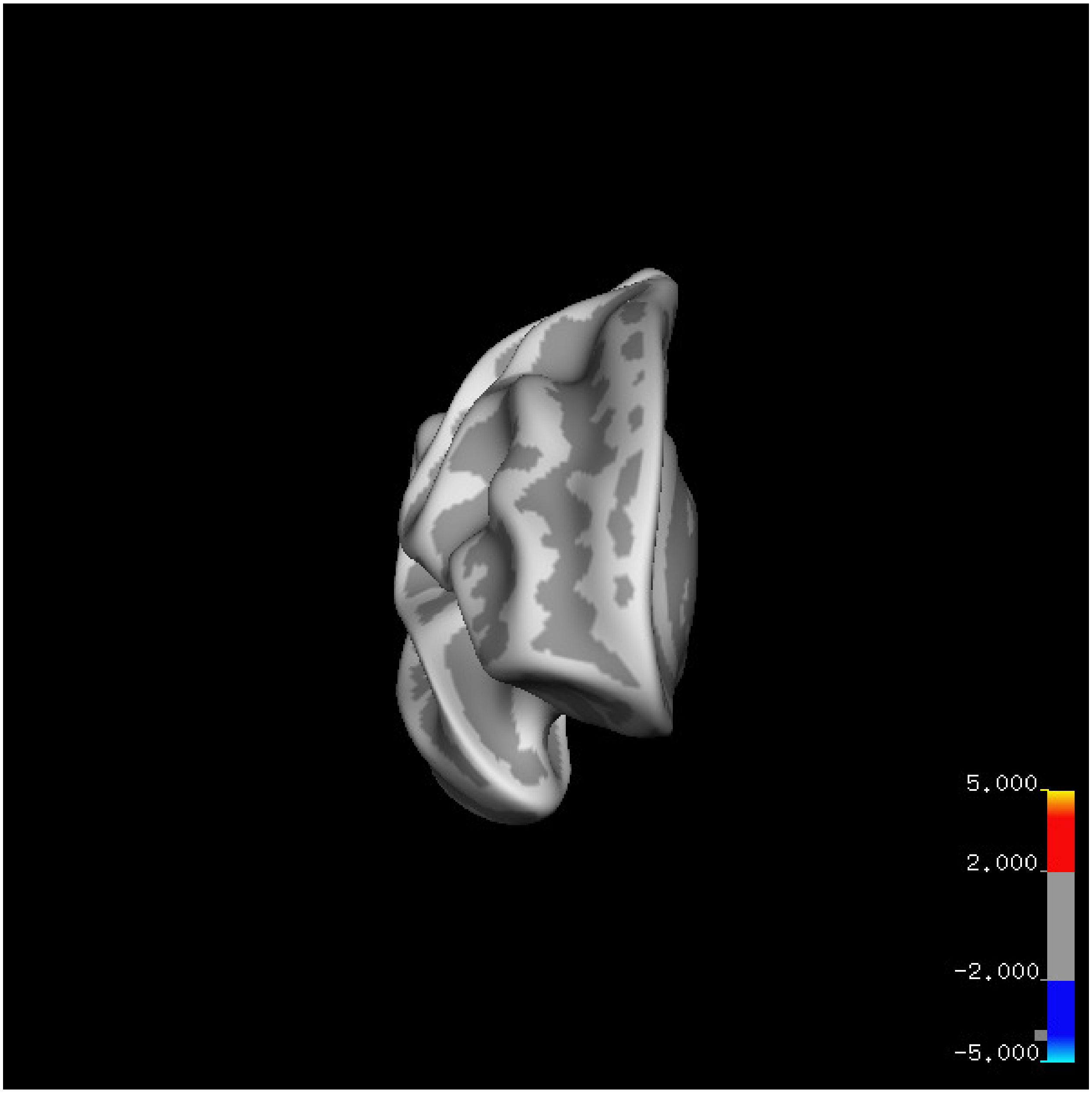
5.000
2.000
-2.000
-5.000

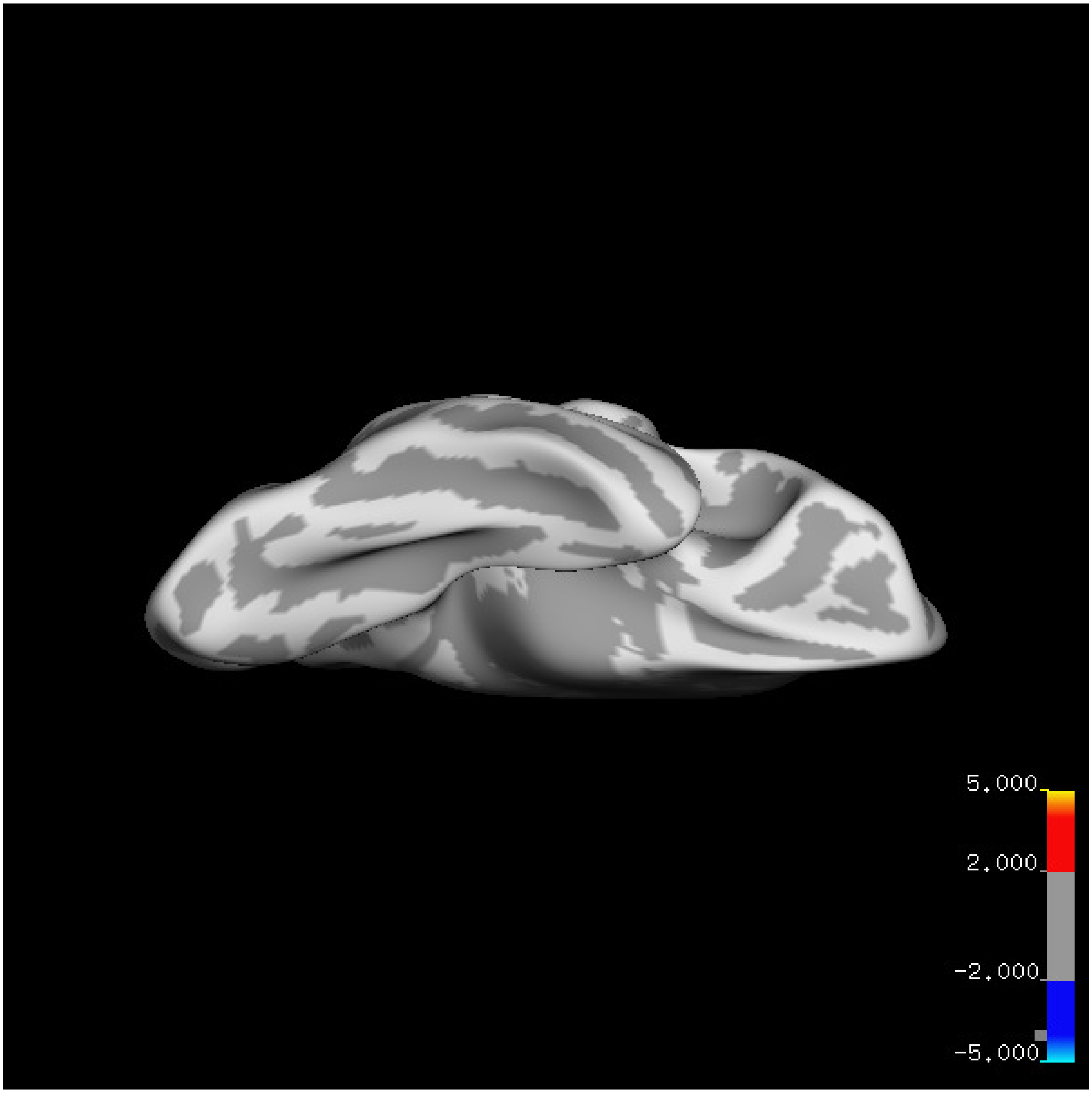

5.000
2.000
-2.000
-5.000

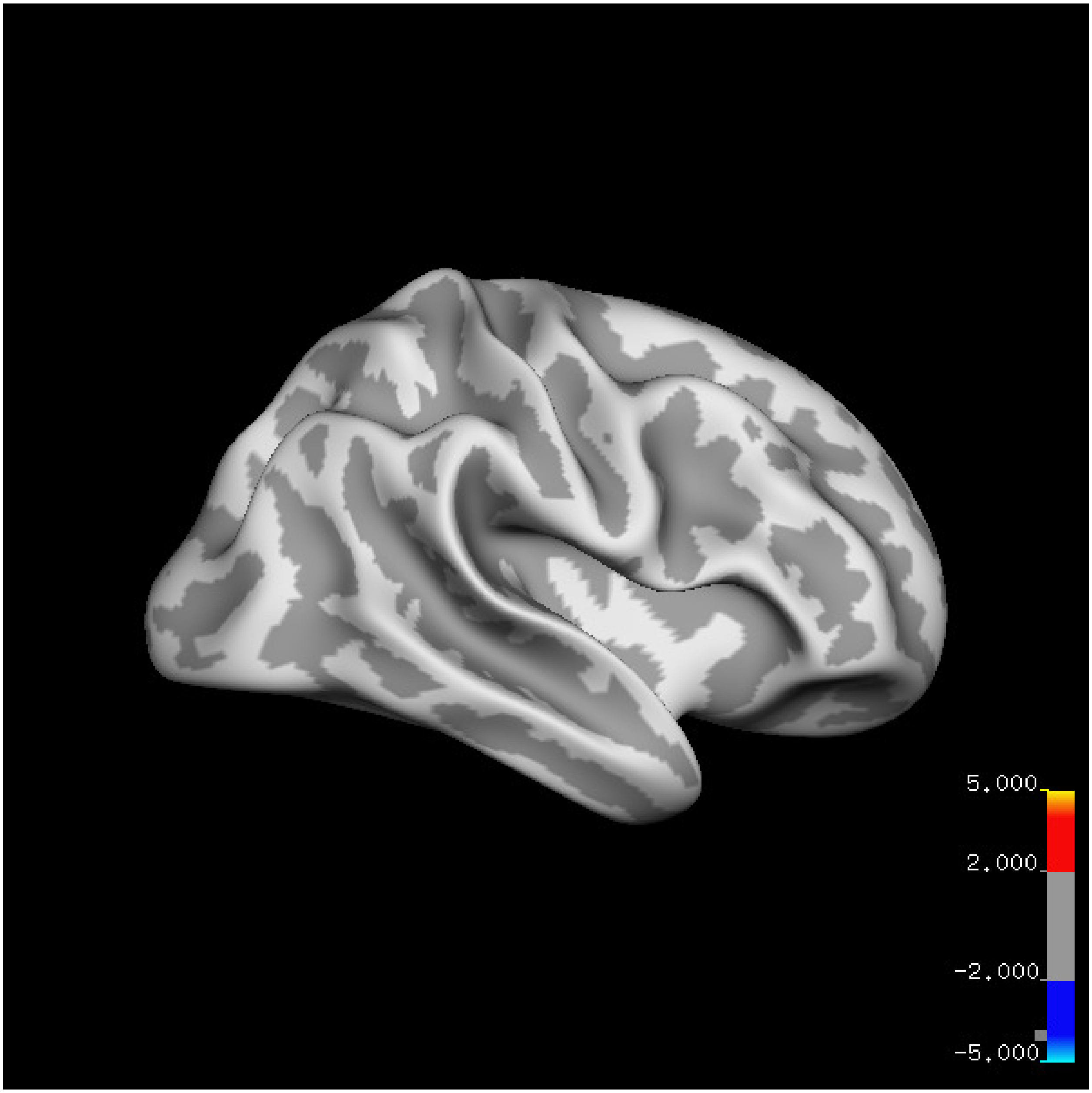
5.000
2.000
-2.000
-5.000

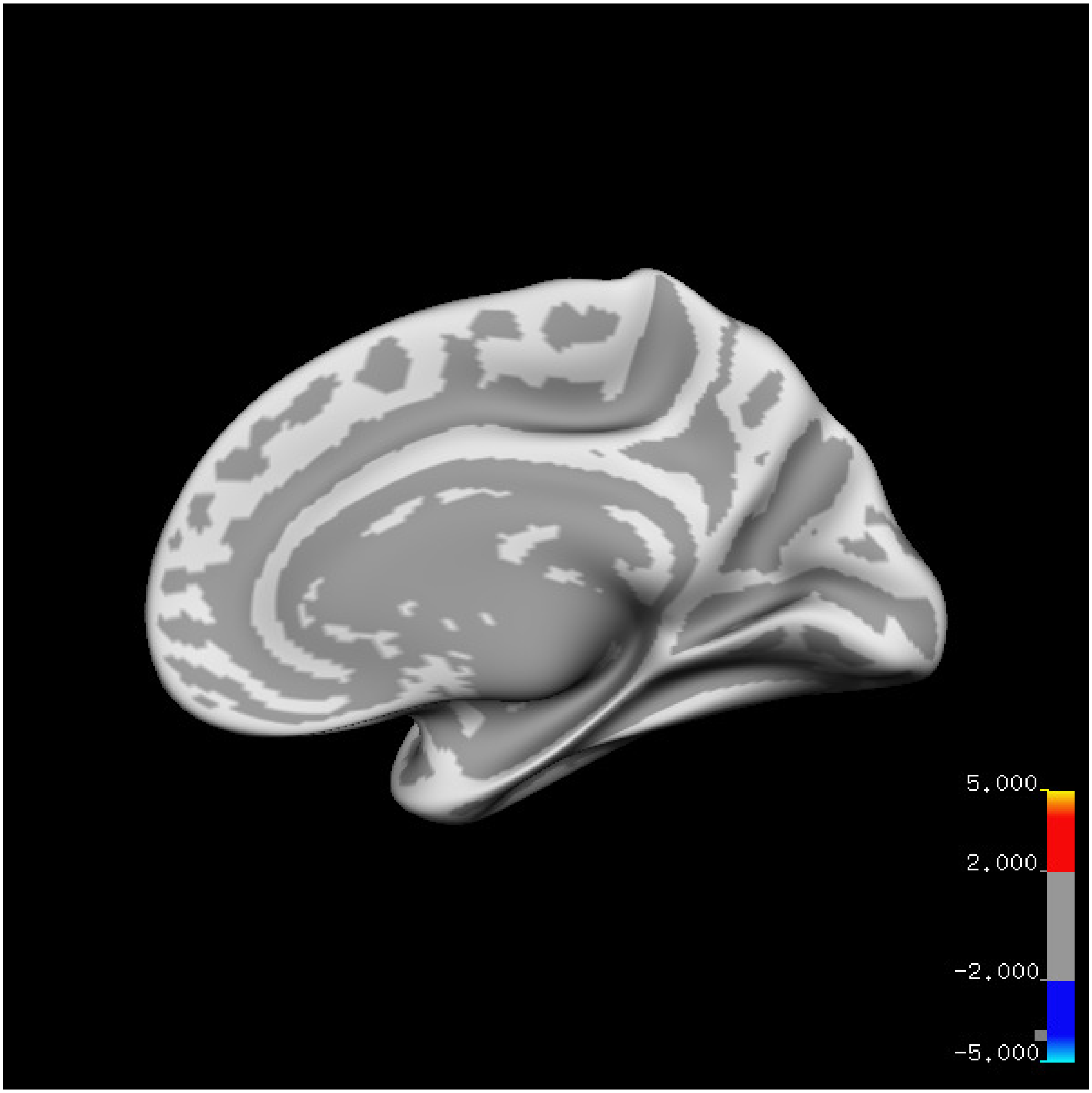
5.000
2.000
-2.000
-5.000

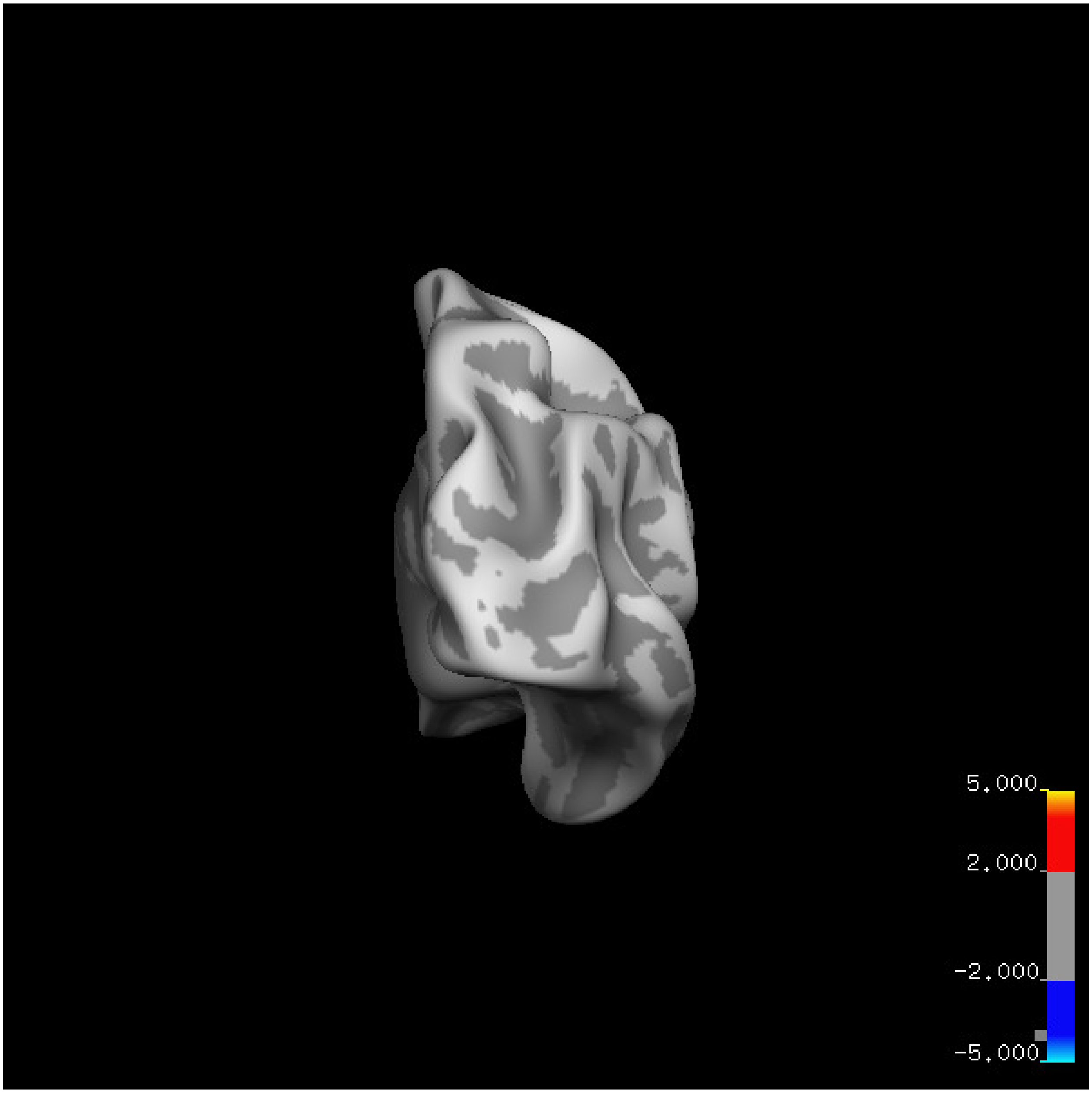
5.000
2.000
-2.000
-5.000

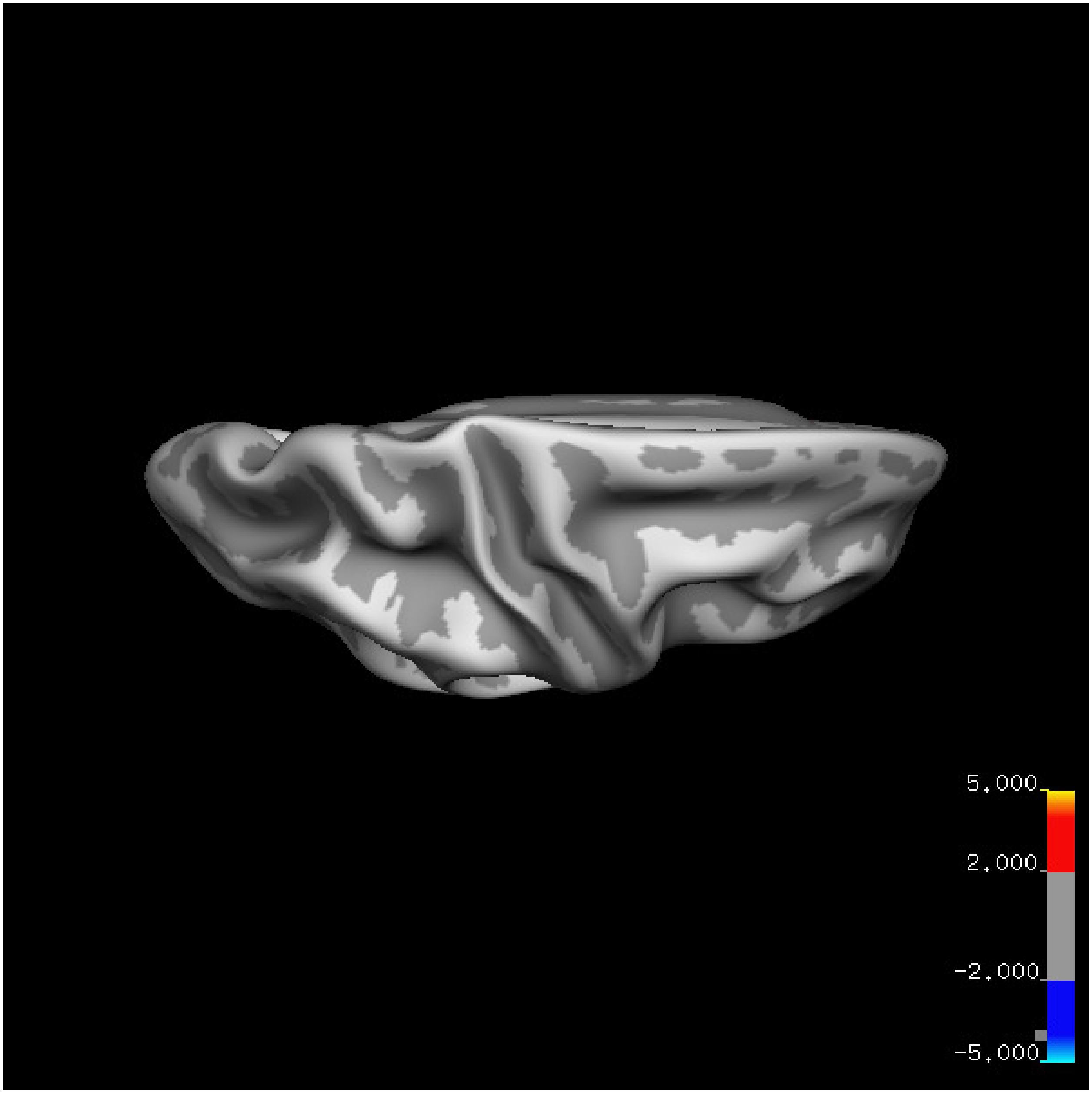


Cluster summary is shown below.

```
## # Cluster Growing Summary (mri_surfcluster)

## # $Id: mri_surfcluster.c,v 1.57.2.3 2016/11/17 18:19:42 zkaufman Exp $

## # $Id: mrisurf.c,v 1.781.2.6 2016/12/27 16:47:14 zkaufman Exp $

## # CreationTime 2021/10/04-11:24:46-GMT

## # cmdline mri_surfcluster.bin --in surface_analyses/results/cluster_comparison/fsaverage6.sm15.rh/ar

## # cwd /gpfs/projects02/p274/cluster/projects/p003-sleep_b

## # sysname  Linux
```

```
## # hostname c1-19
## # machine  x86_64
## # FixVertexAreaFlag 1
## # FixSurfClusterArea 1
## #
## # Input      surface_analyses/results/cluster_comparison/fsaverage6.sm15.rh/area.20_60.below_mean/gl[illegible]
## # Frame Number      0
## # srcsubj fsaverage6
## # hemi rh
## # surface white
## # group_avg_surface_area 85131.2
## # group_avg_vtxarea_loaded 1
## # annot aparc
## # SUBJECTS_DIR /cluster/projects/p274/tools/mri/freesurfer/current/subjects
## # SearchSpace_mm2 79329.9
## # SearchSpace_vtx 37471
## # Bonferroni 0
## # Minimum Threshold 1.3
## # Maximum Threshold infinity
## # Threshold Sign    abs
## # AdjustThreshWhenOneTail 1
## # CW PValue Threshold: 0.01
## # Area Threshold    0 mm^2
## # CSD thresh  1.300000
## # CSD nreps    10000
## # CSD simtype  null-z
## # CSD contrast NA
```

```
## # CSD confint  90.000000

## # Overall max 4.59274 at vertex 38746

## # Overall min -3.02139 at vertex 39545

## # NClusters          0

## # FixMNI = 0

## #

## # ClusterNo  Max   VtxMax   Size(mm^2)  MNIX   MNIY   MNIZ    CWP    CWPLow    CWPHi   NVtxs    Wght
```

**Age above 60**

**Sleep above mean** Cluster corrected significance plots are shown below.

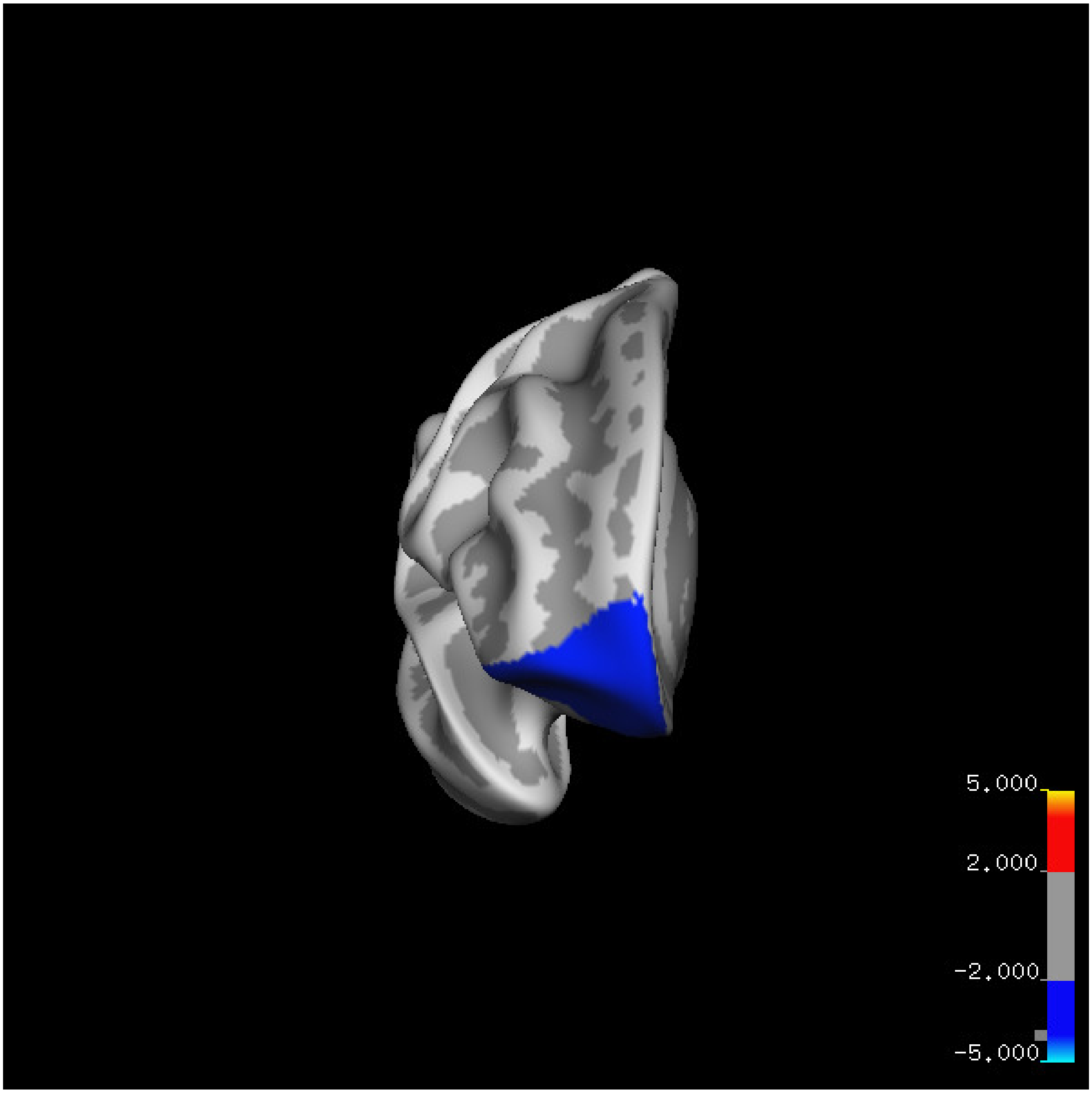
5.000
2.000
-2.000
-5.000

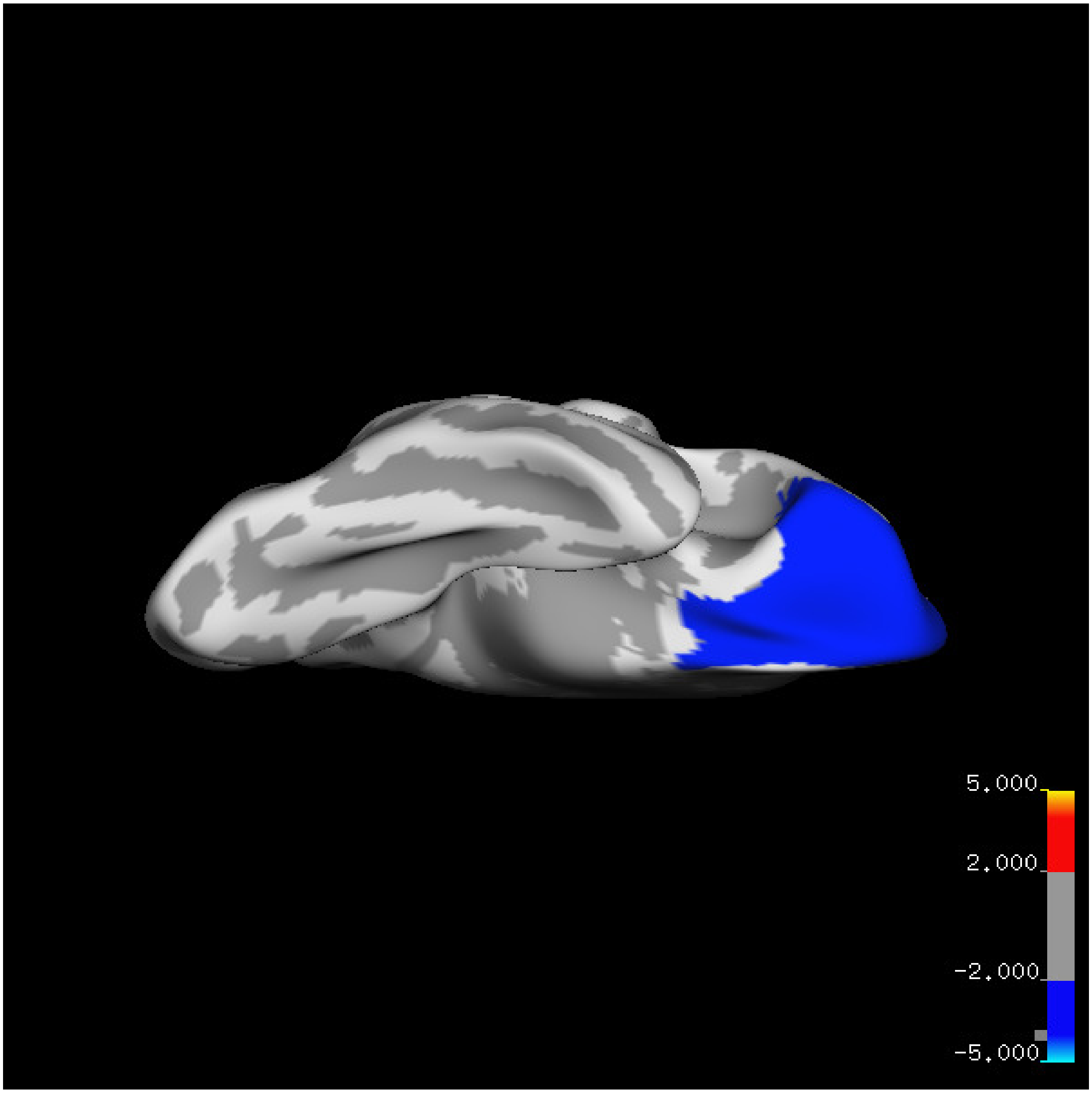
5.000
2.000
-2.000
-5.000

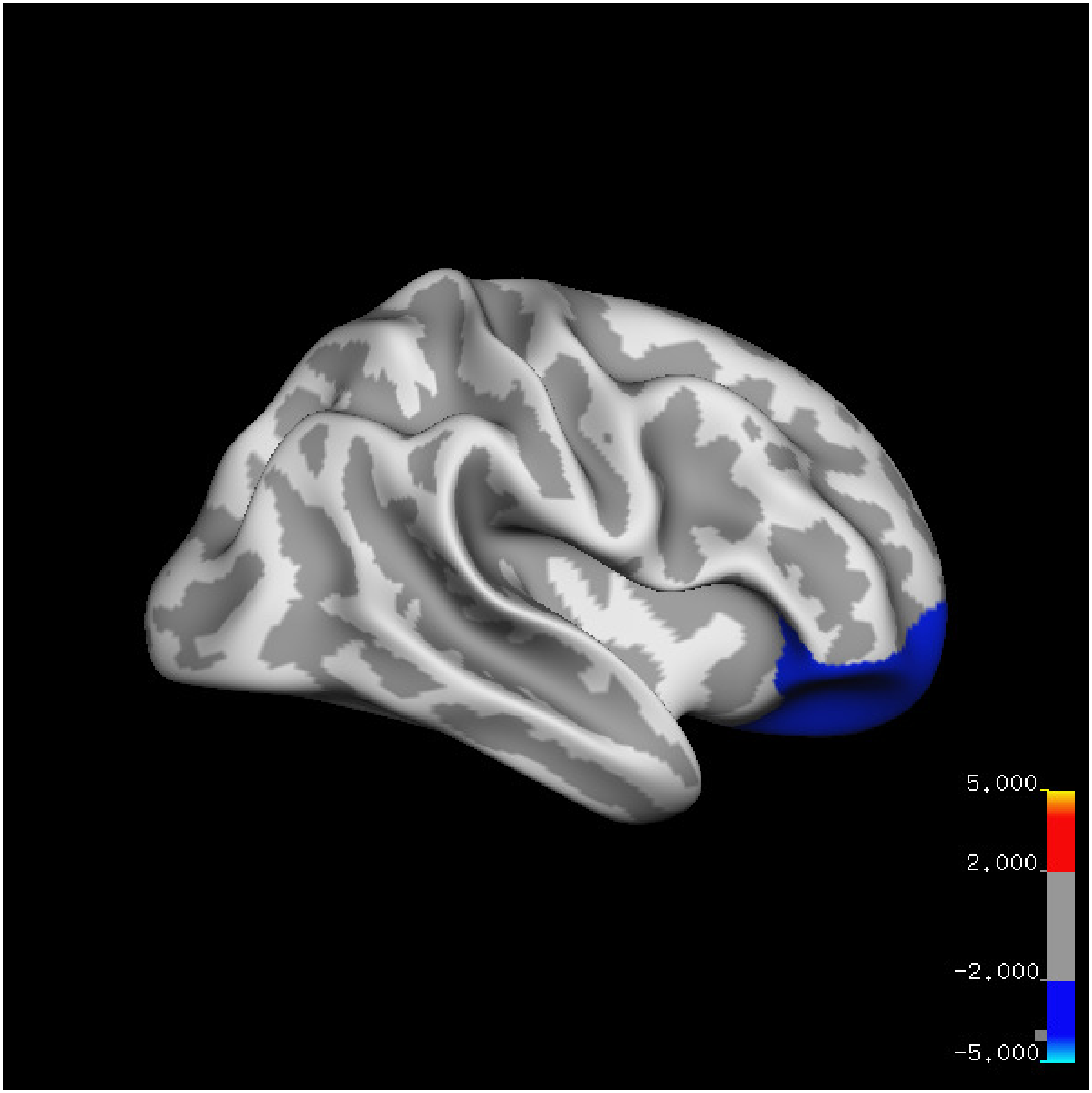
5.000
2.000
-2.000
-5.000

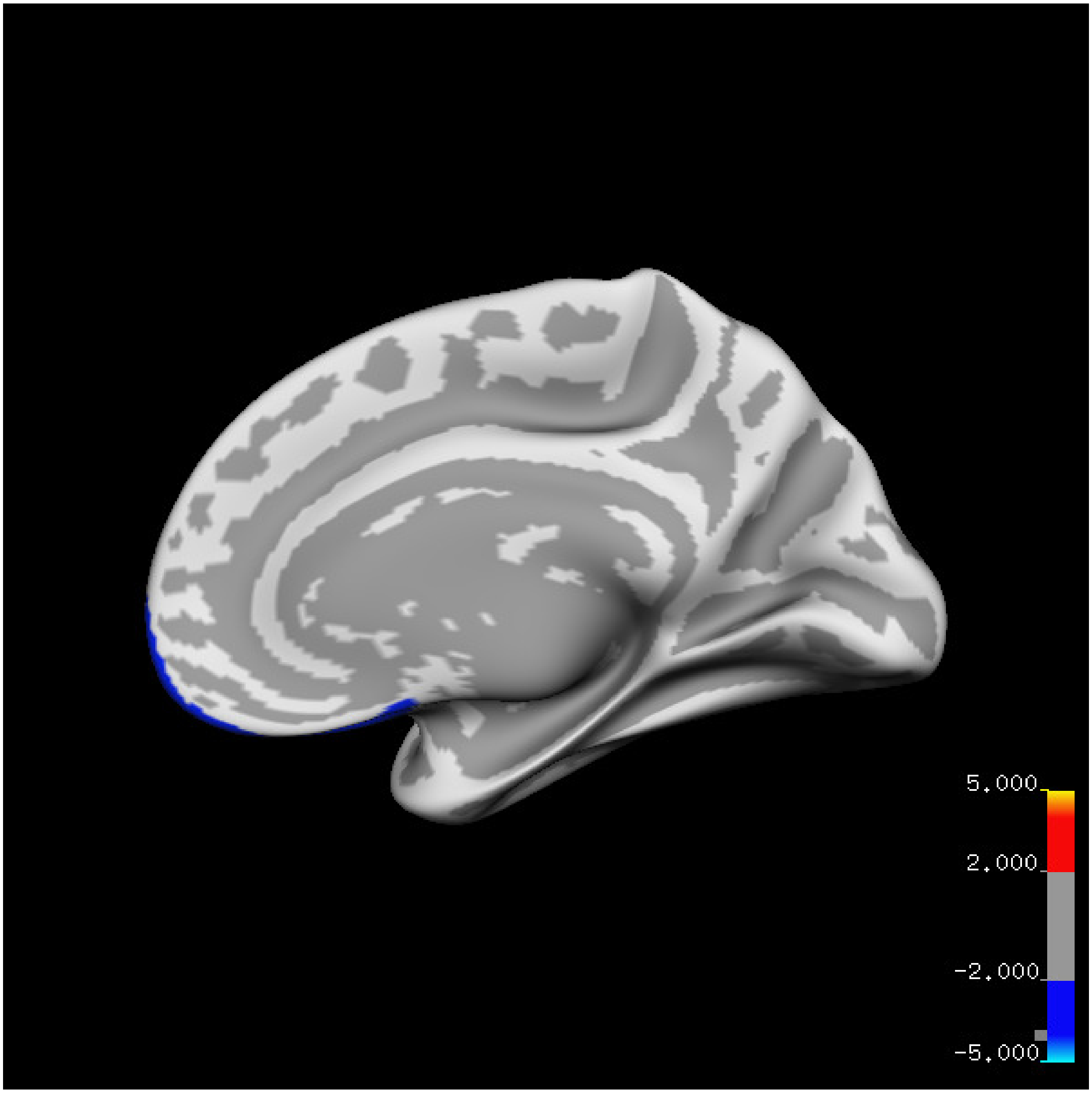
5.000
2.000
-2.000
-5.000

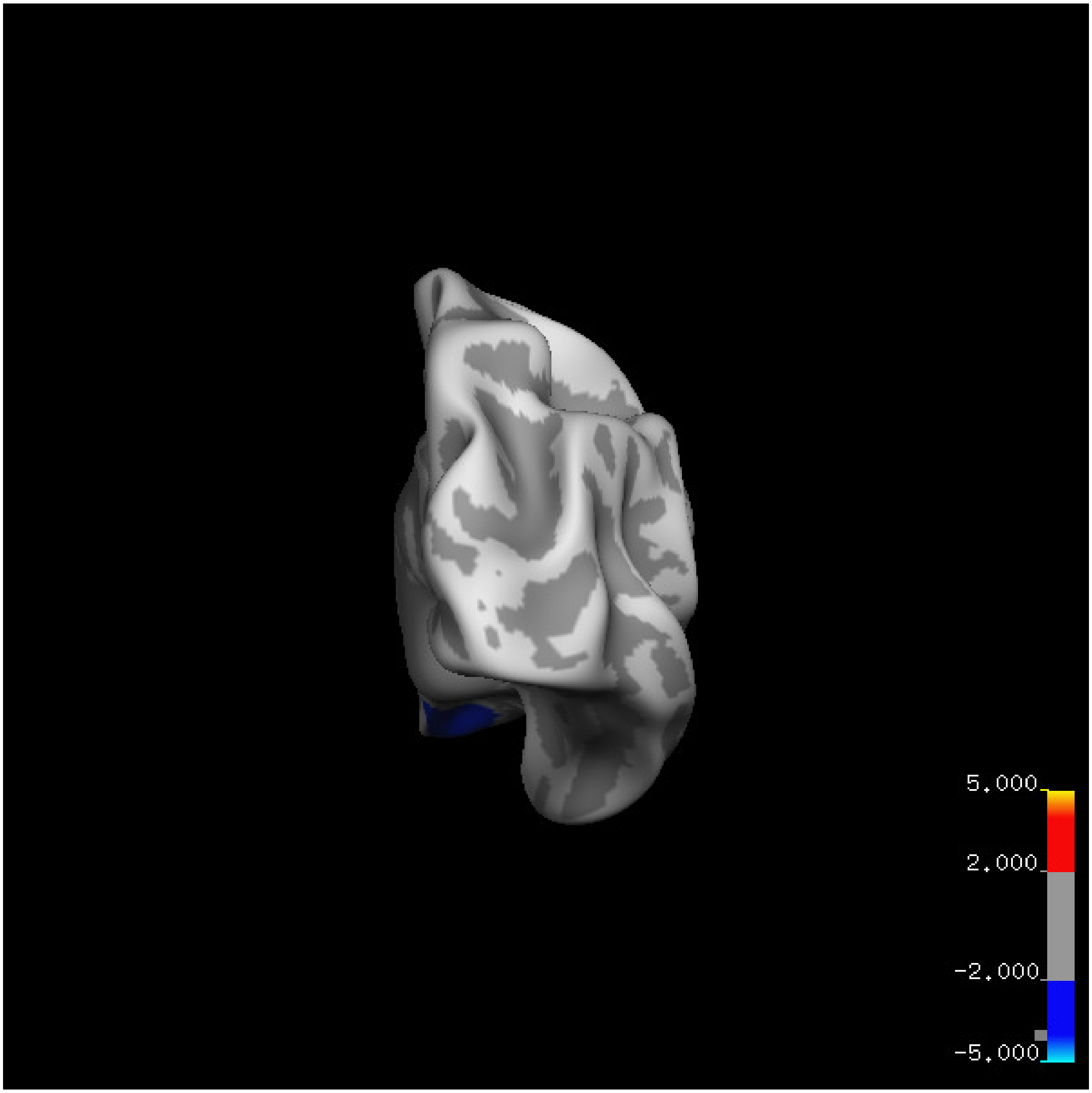
5.000
2.000
-2.000
-5.000

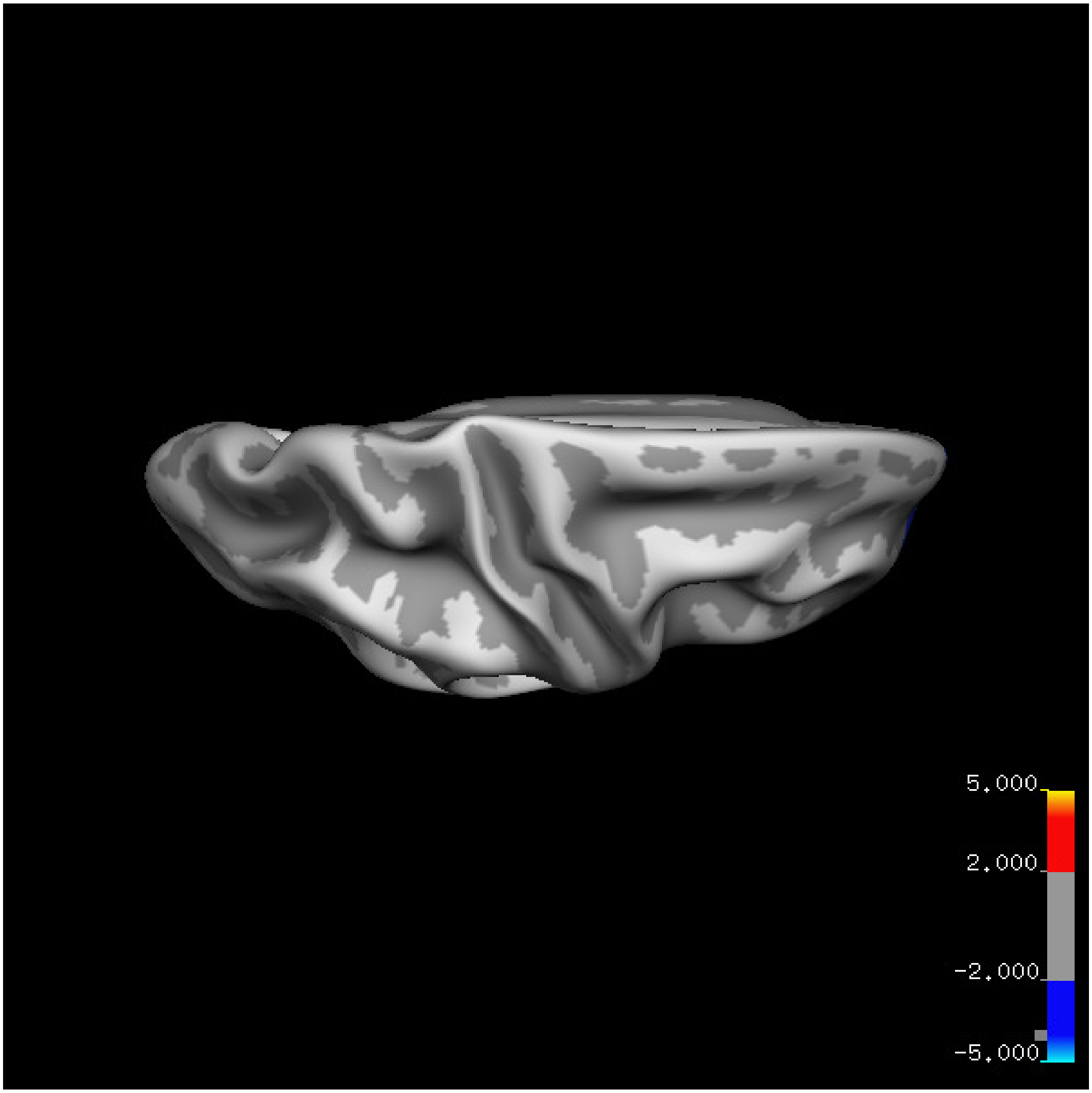
5.000
2.000
-2.000
-5.000

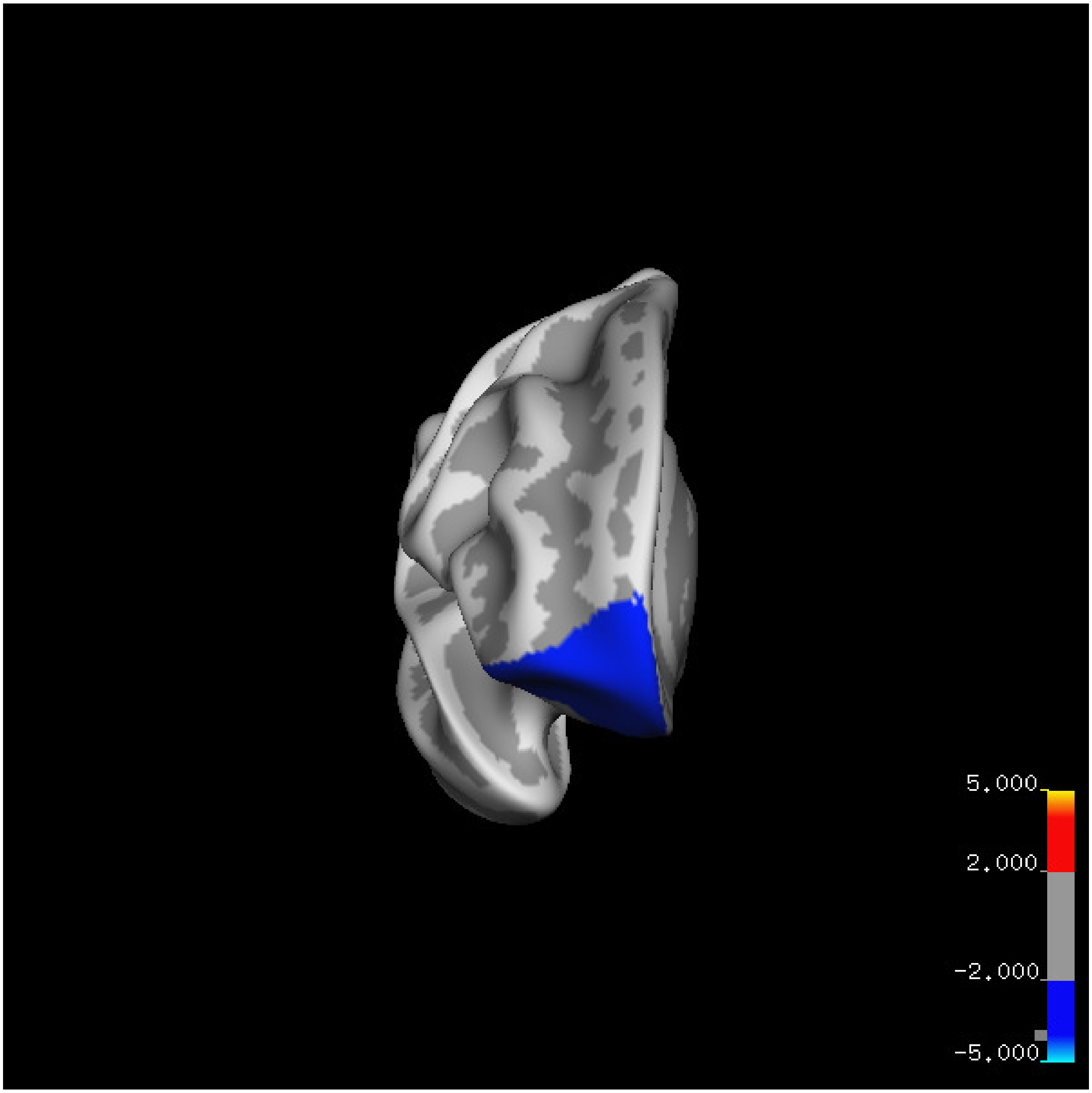
5.000
2.000
-2.000
-5.000

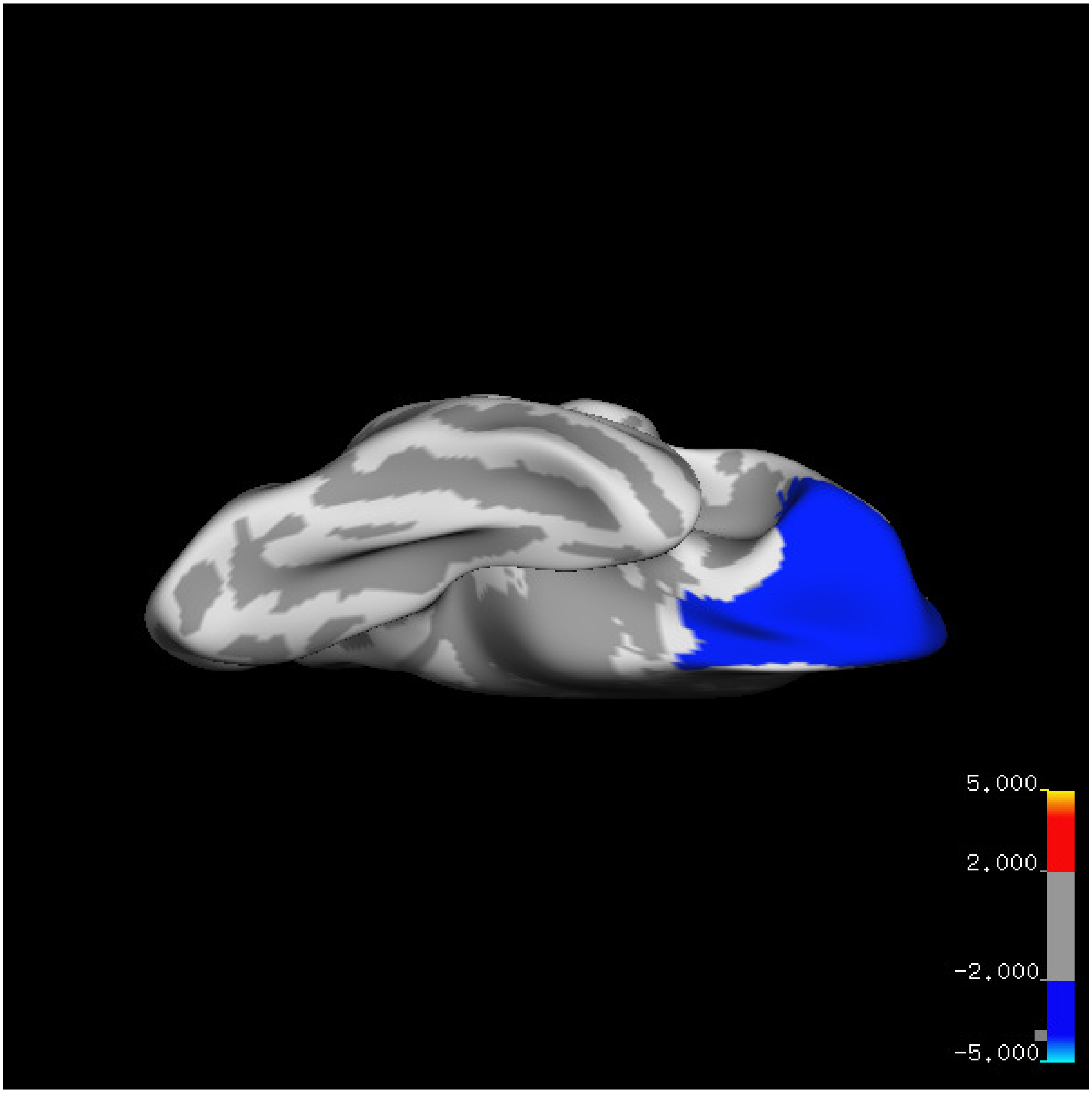
5.000
2.000
-2.000
-5.000

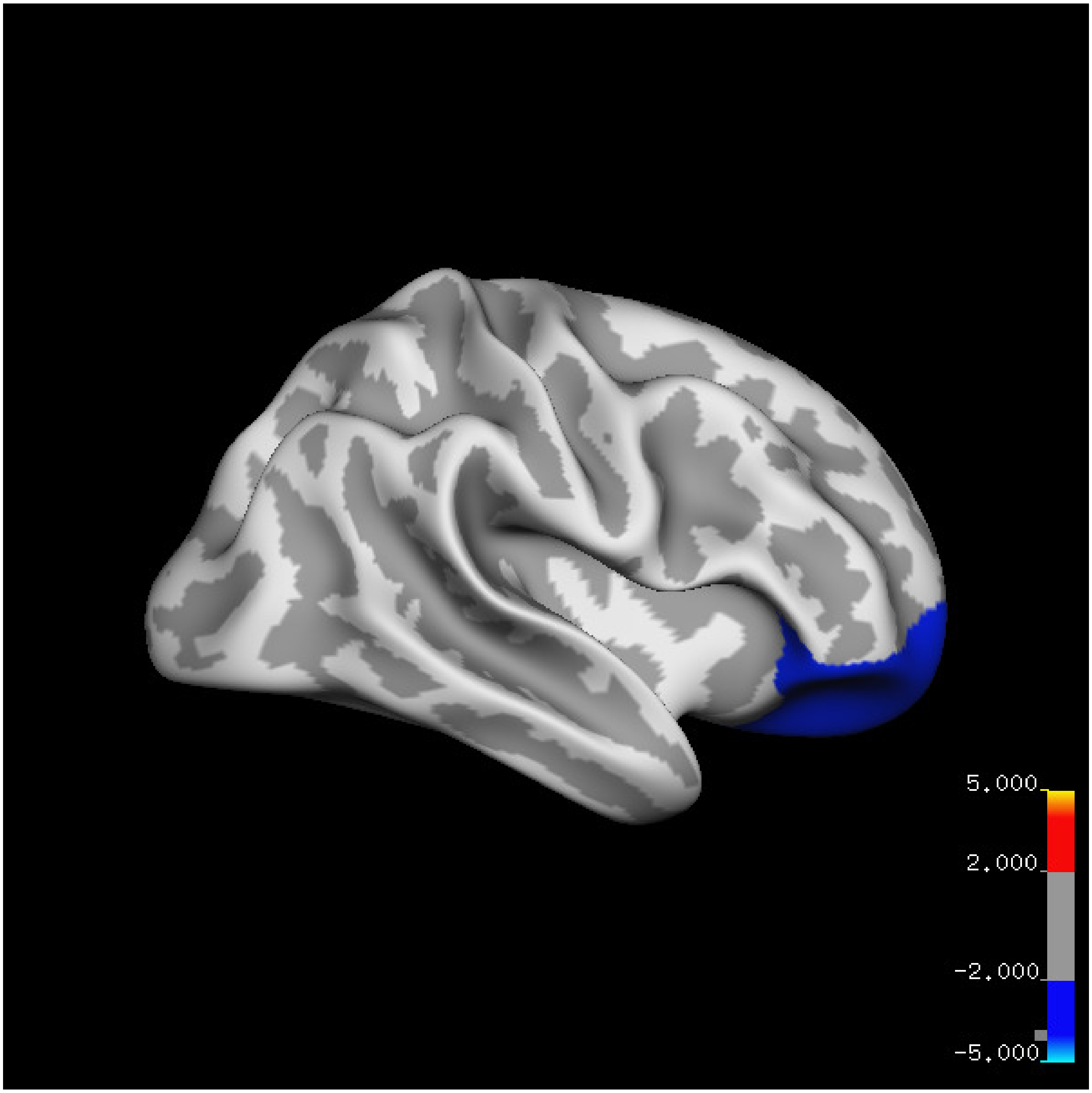
5.000
2.000
-2.000
-5.000

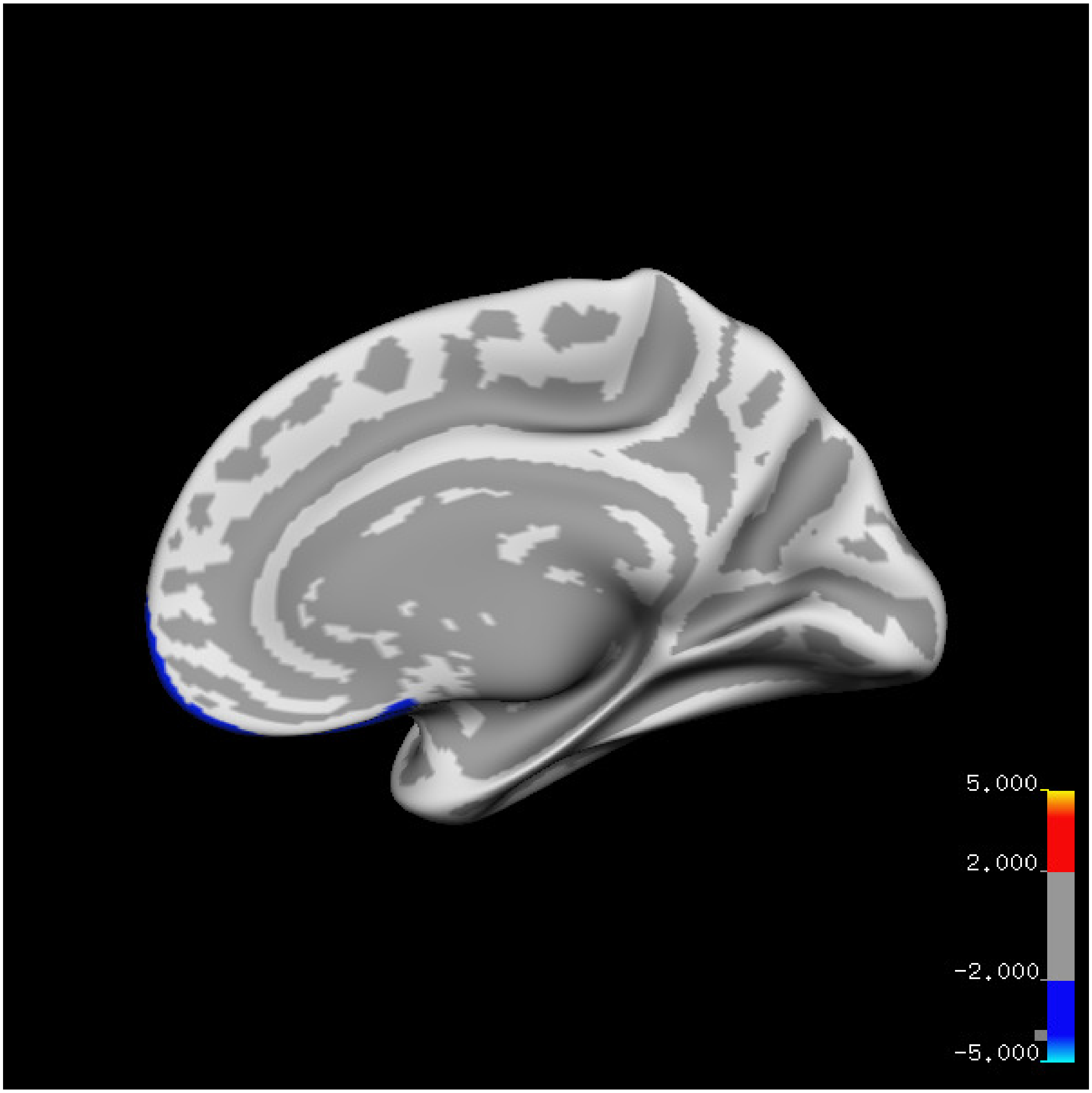
5.000
2.000
-2.000
-5.000

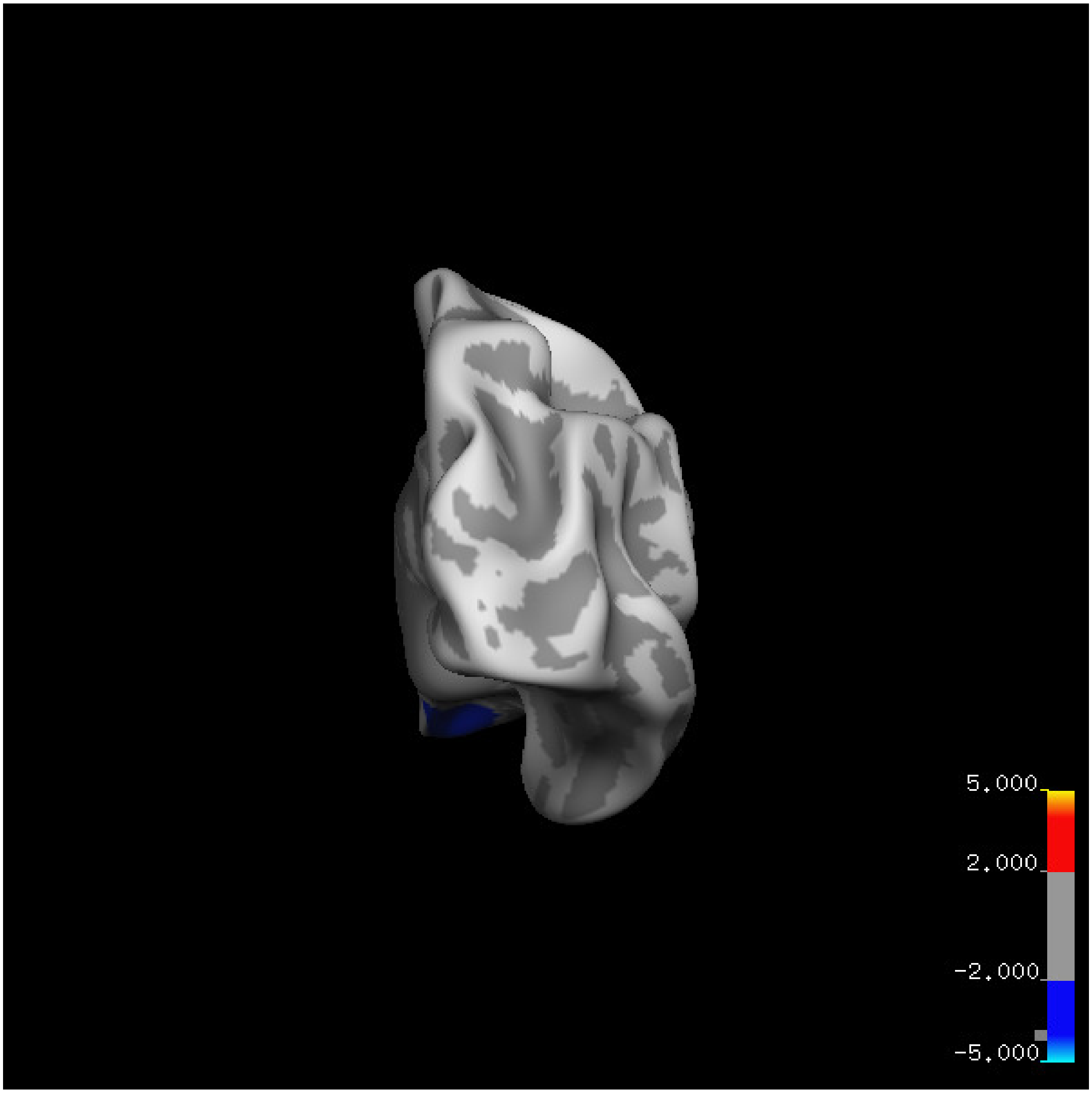
5.000
2.000
-2.000
-5.000

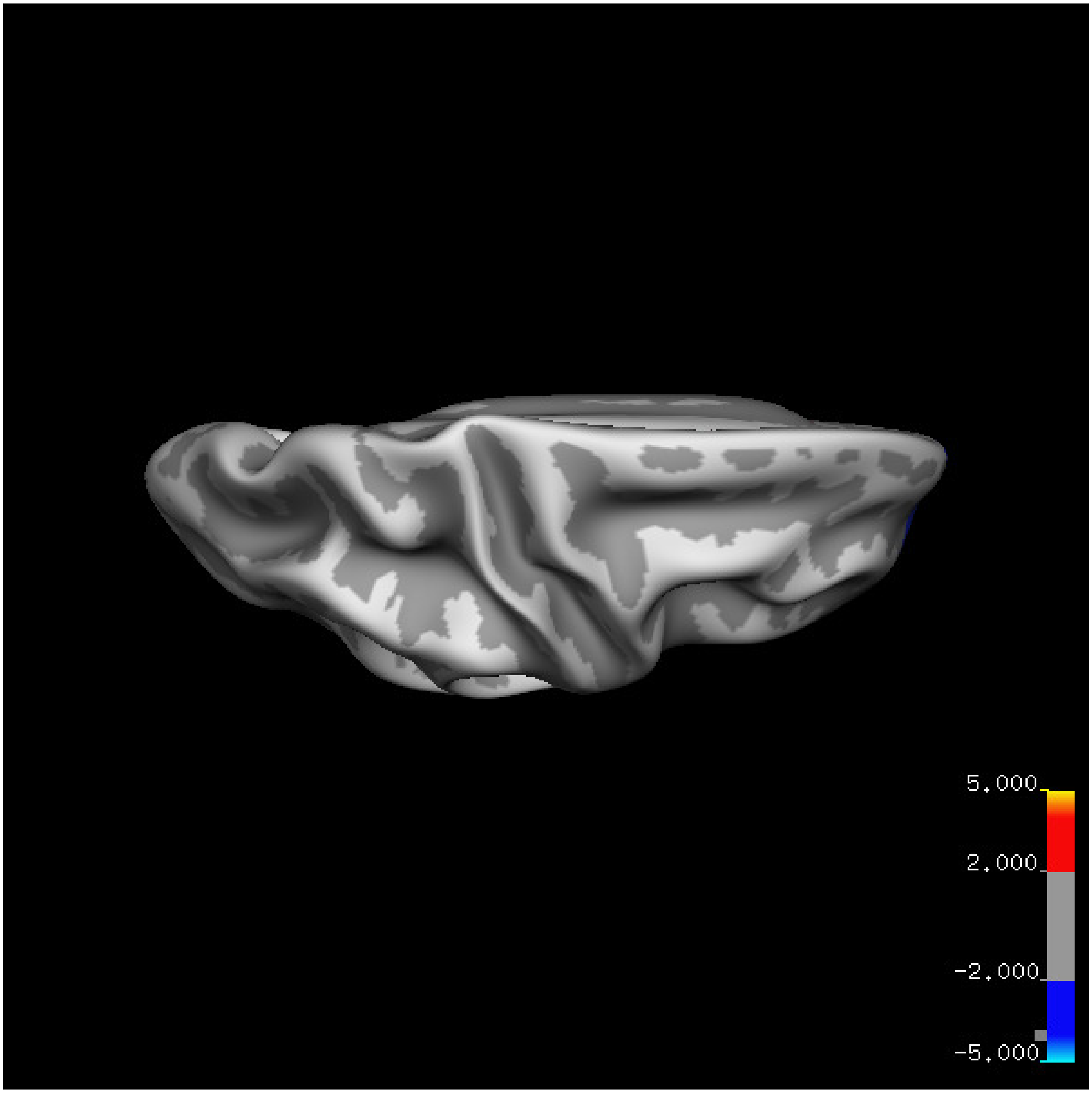


Cluster summary is shown below.

```
## # Cluster Growing Summary (mri_surfcluster)

## # $Id: mri_surfcluster.c,v 1.57.2.3 2016/11/17 18:19:42 zkaufman Exp $

## # $Id: mrisurf.c,v 1.781.2.6 2016/12/27 16:47:14 zkaufman Exp $

## # CreationTime 2021/10/04-13:08:12-GMT

## # cmdline mri_surfcluster.bin --in surface_analyses/results/cluster_comparison/fsaverage6.sm15.rh/ar

## # cwd /gpfs/projects02/p274/cluster/projects/p003-sleep_b

## # sysname  Linux
```

```
## # hostname c1-19
## # machine  x86_64
## # FixVertexAreaFlag 1
## # FixSurfClusterArea 1
## #
## # Input      surface_analyses/results/cluster_comparison/fsaverage6.sm15.rh/area.above60.above_mean/
## # Frame Number      0
## # srcsubj fsaverage6
## # hemi rh
## # surface white
## # group_avg_surface_area 85131.2
## # group_avg_vtxarea_loaded 1
## # annot aparc
## # SUBJECTS_DIR /cluster/projects/p274/tools/mri/freesurfer/current/subjects
## # SearchSpace_mm2 79329.9
## # SearchSpace_vtx 37471
## # Bonferroni 0
## # Minimum Threshold 1.3
## # Maximum Threshold infinity
## # Threshold Sign    abs
## # AdjustThreshWhenOneTail 1
## # CW PValue Threshold: 0.01
## # Area Threshold    0 mm^2
## # CSD thresh  1.300000
## # CSD nreps    10000
## # CSD simtype  null-z
## # CSD contrast NA
```

```
## # CSD confint  90.000000
## # Overall max 3.82101 at vertex 28087
## # Overall min -6.68779 at vertex 5954
## # NClusters          1
## # FixMNI = 0
## #
## # ClusterNo  Max   VtxMax   Size(mm^2)  MNIX   MNIY   MNIZ    CWP    CWPLow    CWPHi   NVtxs    Wght
##    1       -6.688    5954   4006.31     15.1   59.8  -14.9  0.00060  0.00030  0.00090   1593    -438
```

**Sleep below mean** Cluster corrected significance plots are shown below.

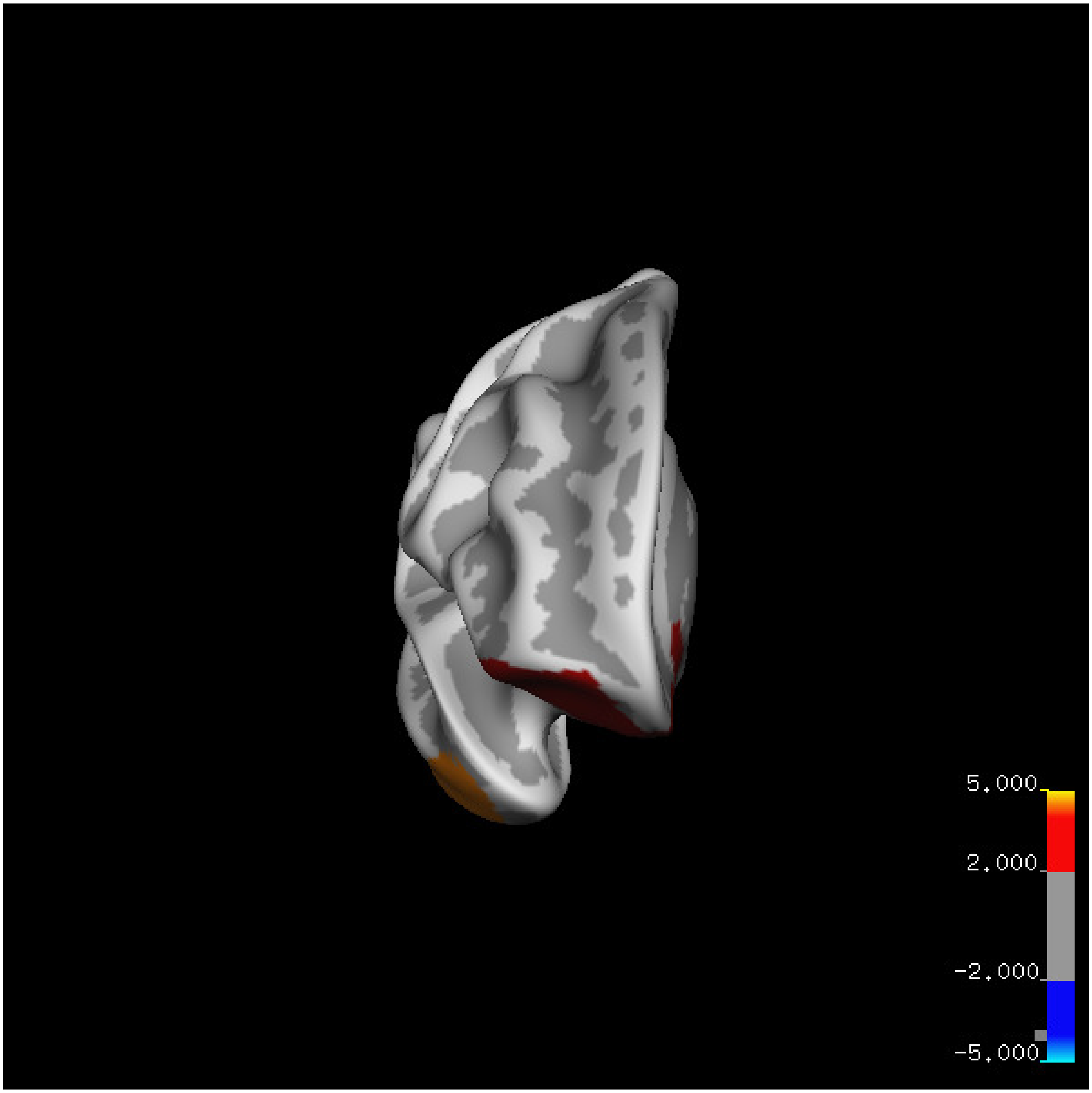
5.000
2.000
-2.000
-5.000

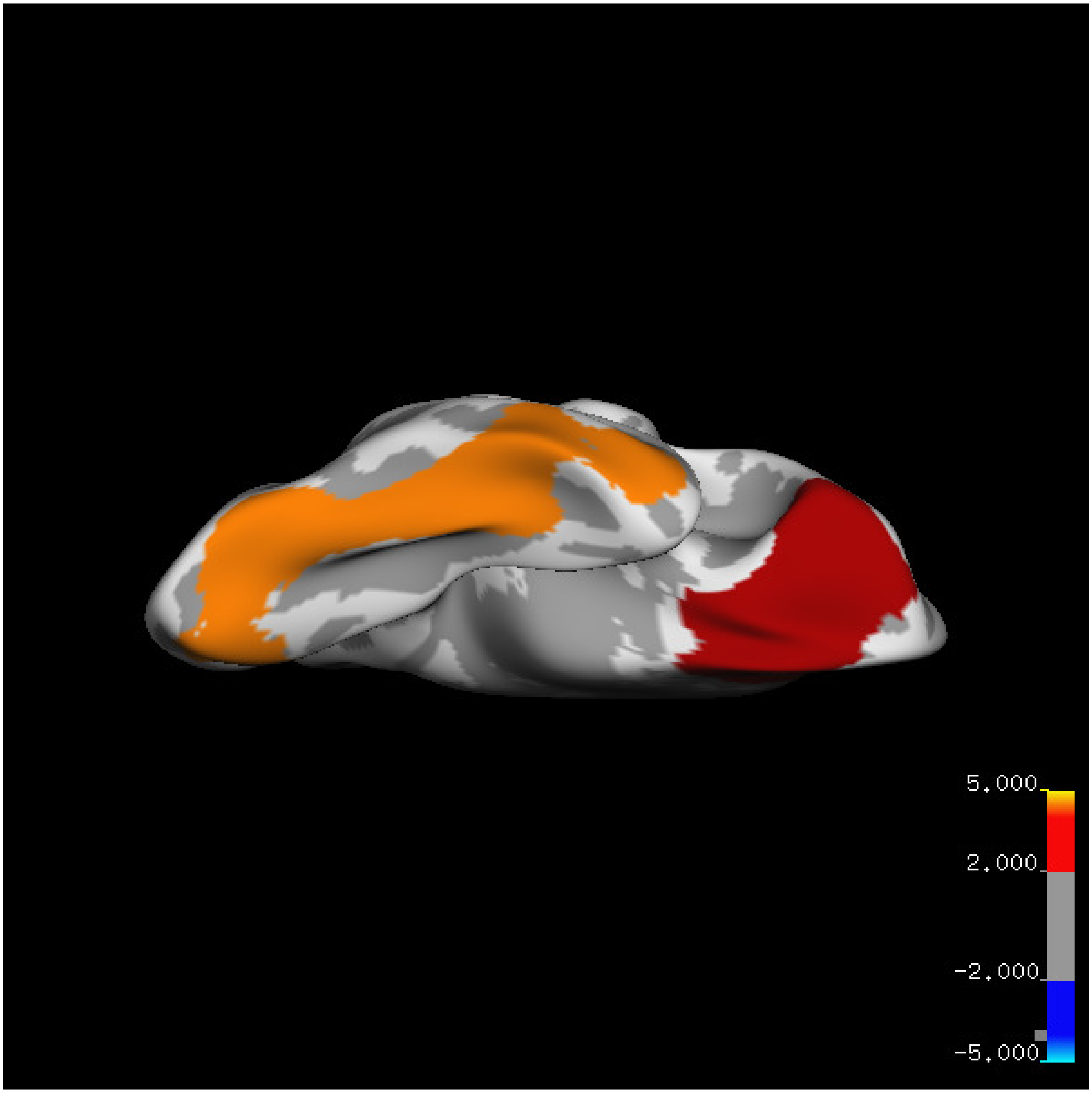
5.000
2.000
-2.000
-5.000

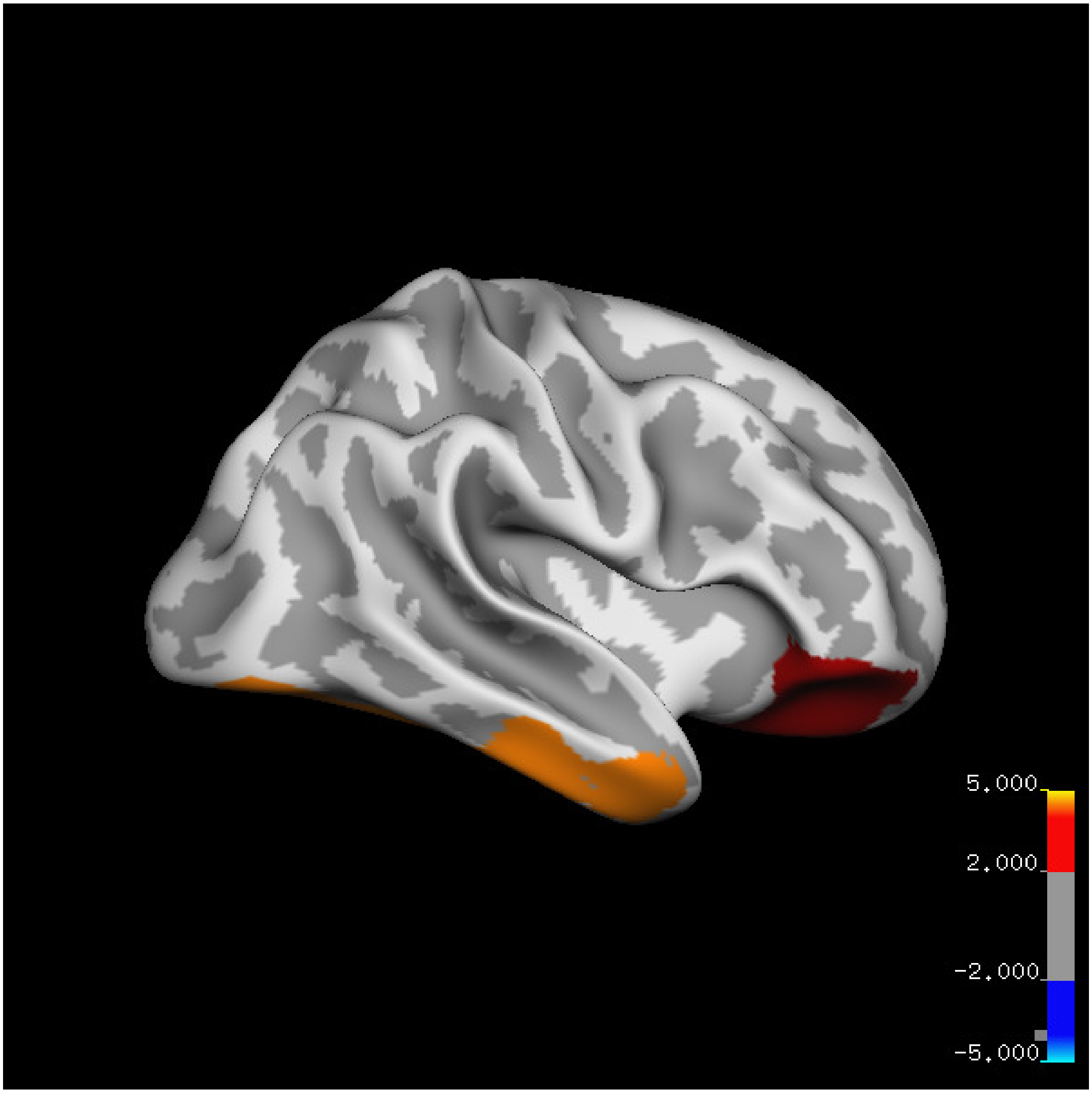
5.000
2.000
-2.000
-5.000

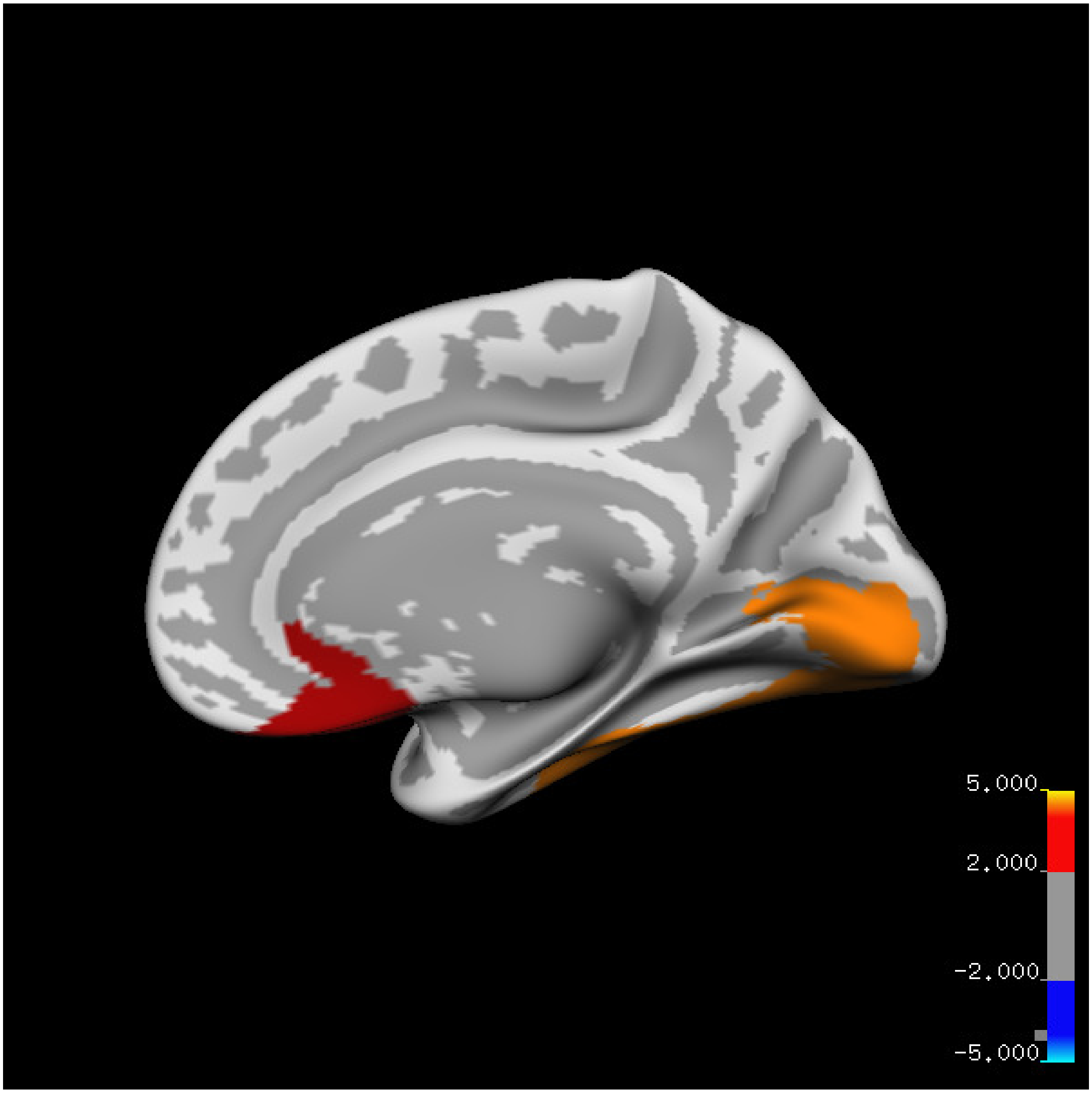
5.000
2.000
-2.000
-5.000

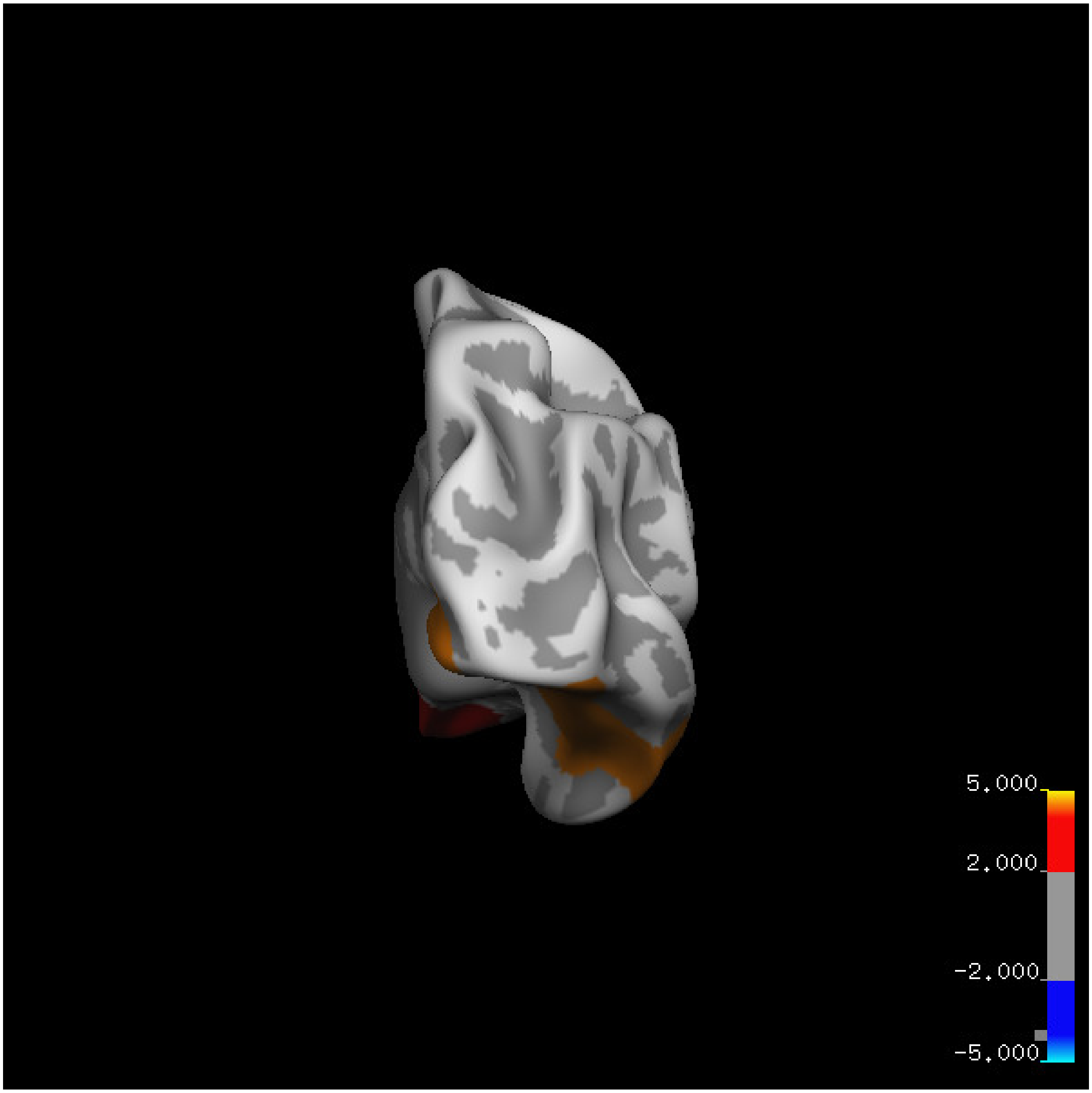
5.000
2.000
-2.000
-5.000

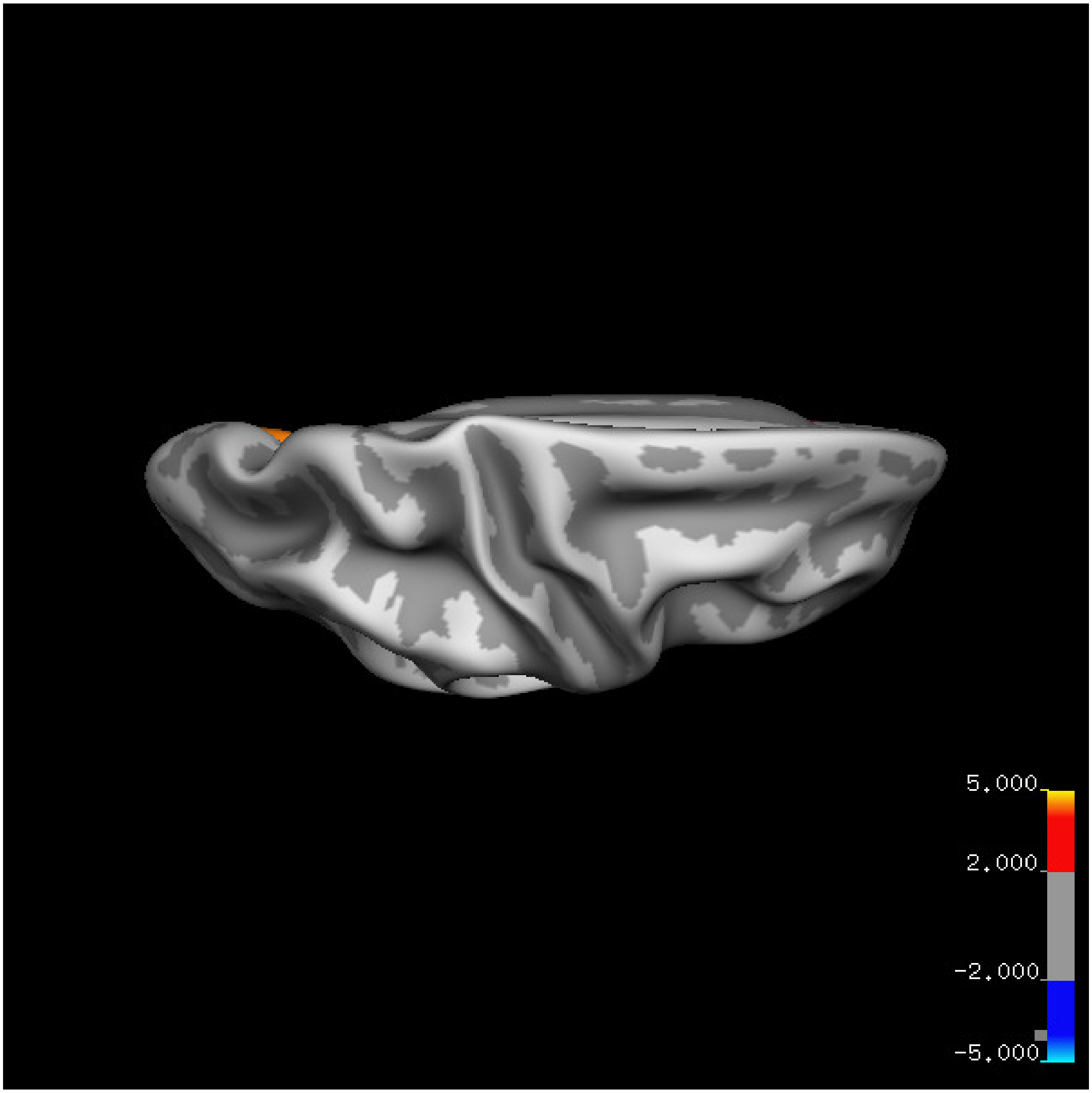

5.000
2.000
-2.000
-5.000

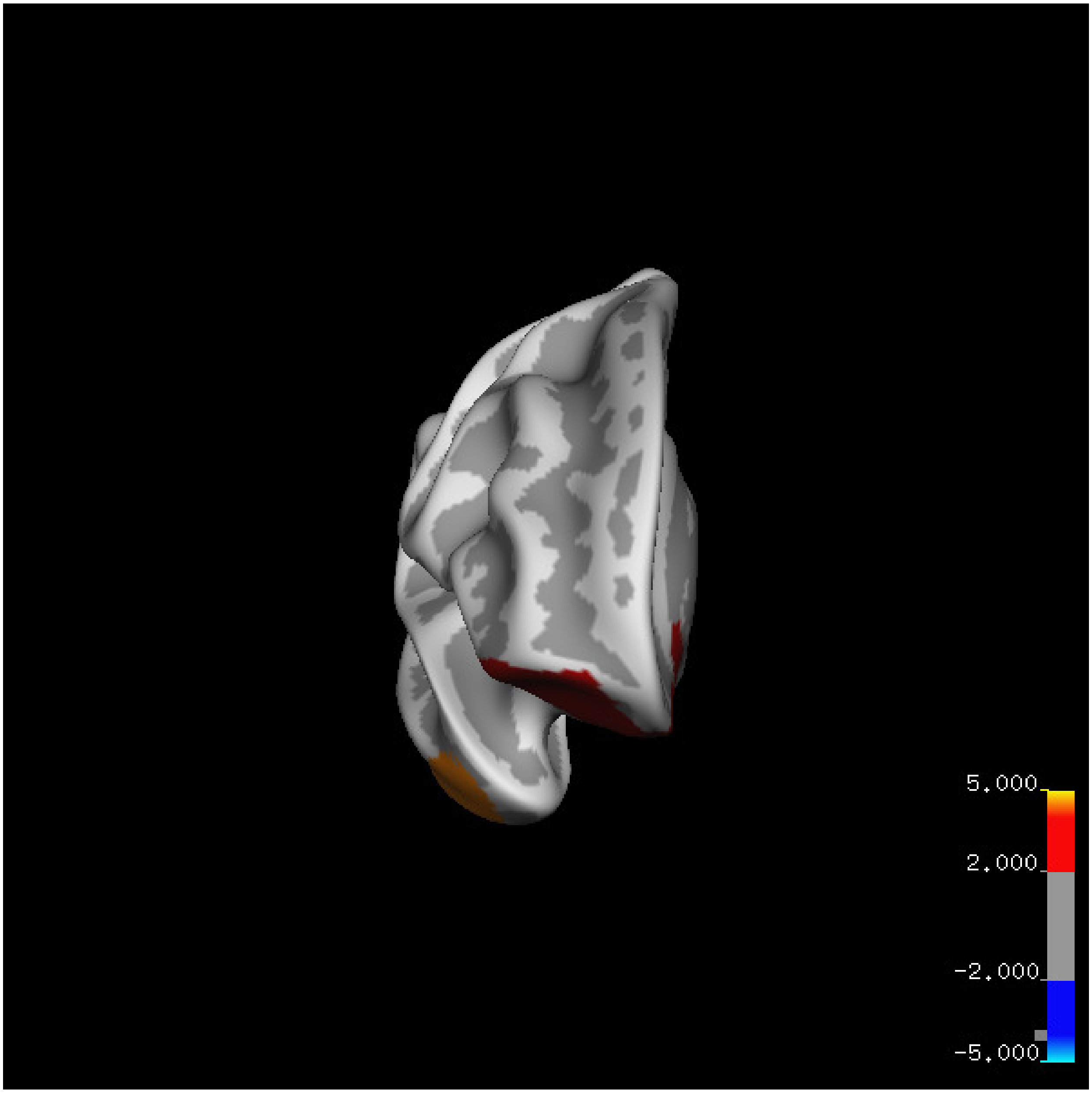
5.000
2.000
-2.000
-5.000

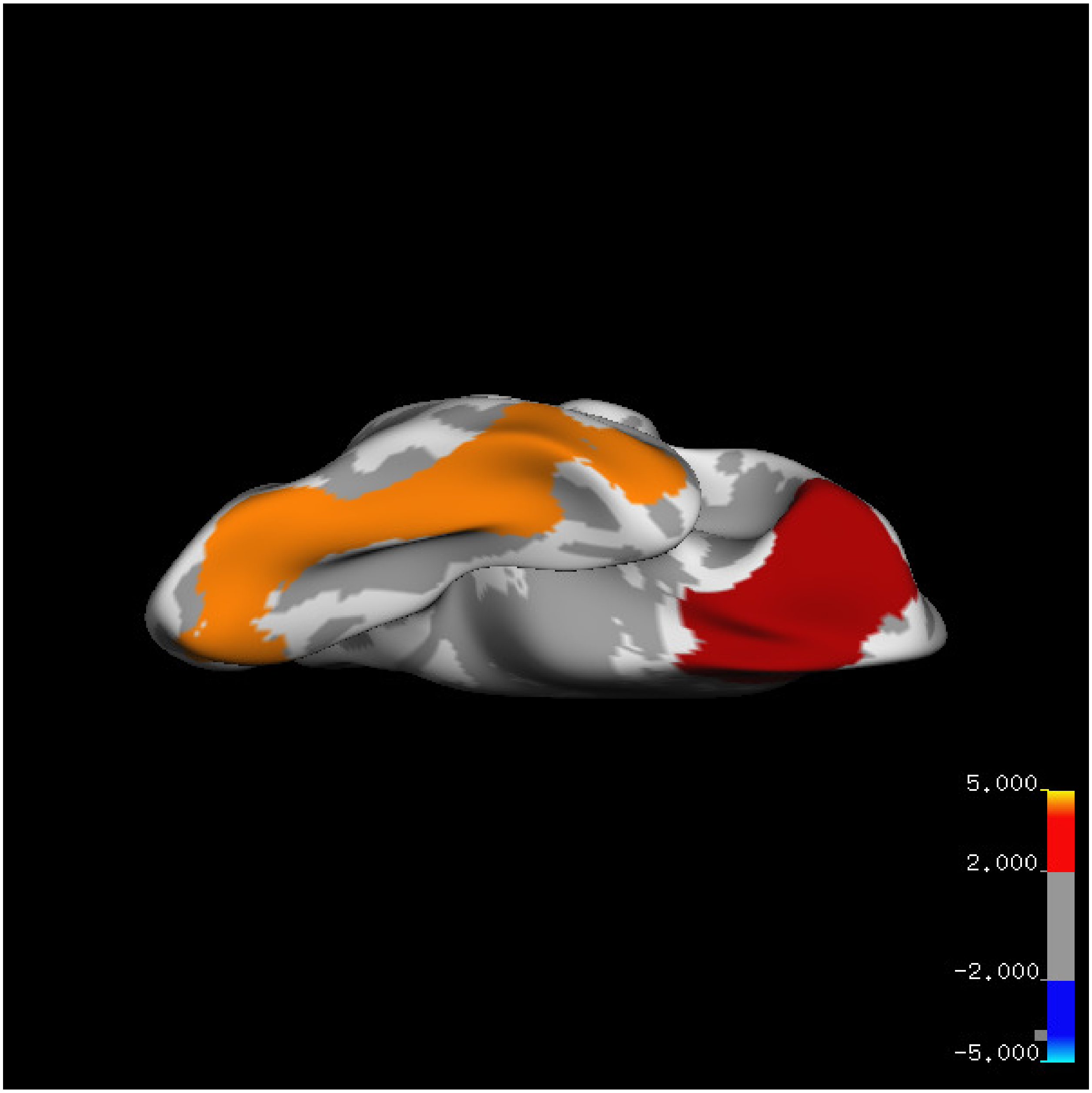
5.000
2.000
-2.000
-5.000

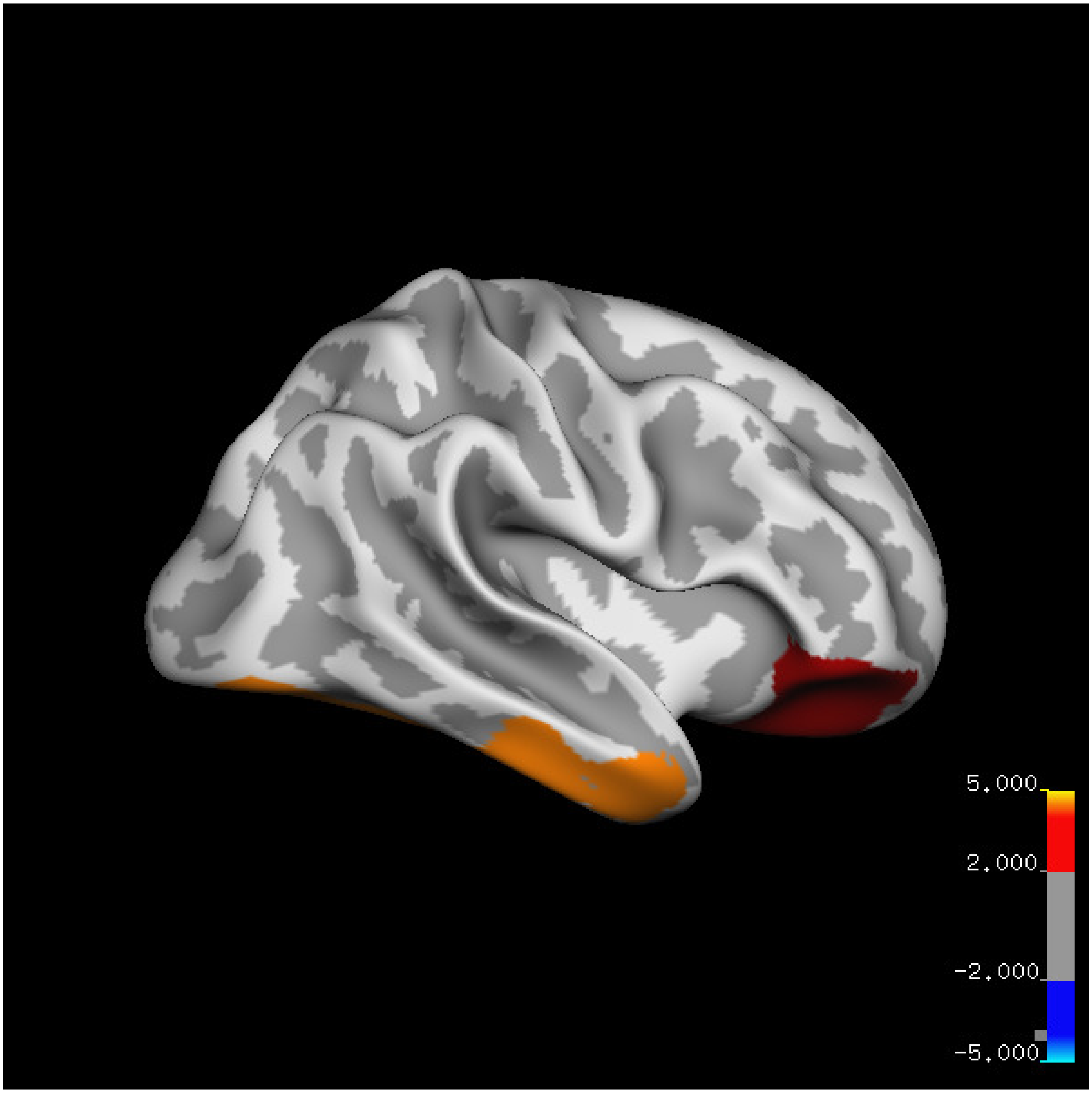
5.000
2.000
-2.000
-5.000

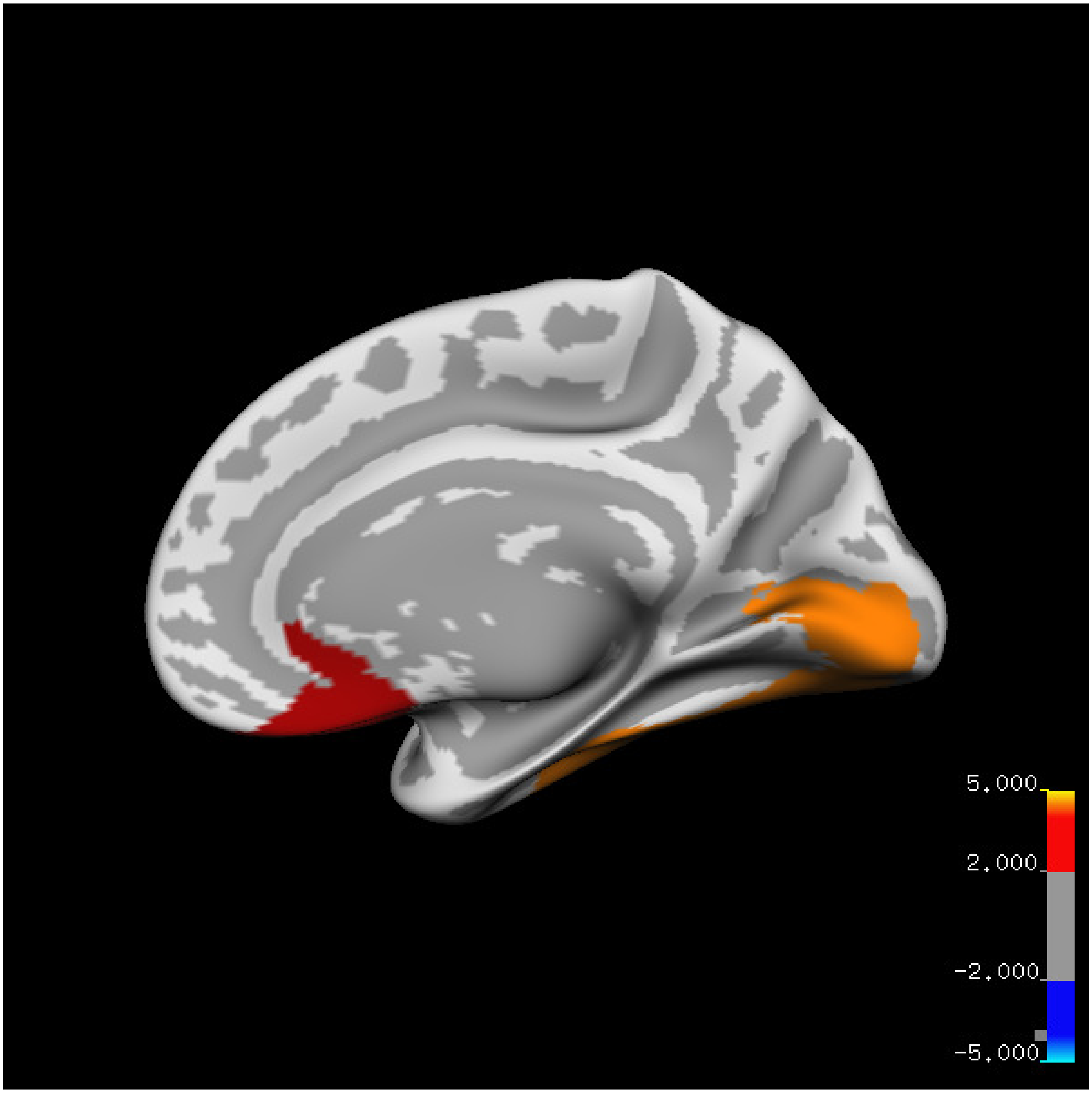

5.000
2.000
-2.000
-5.000

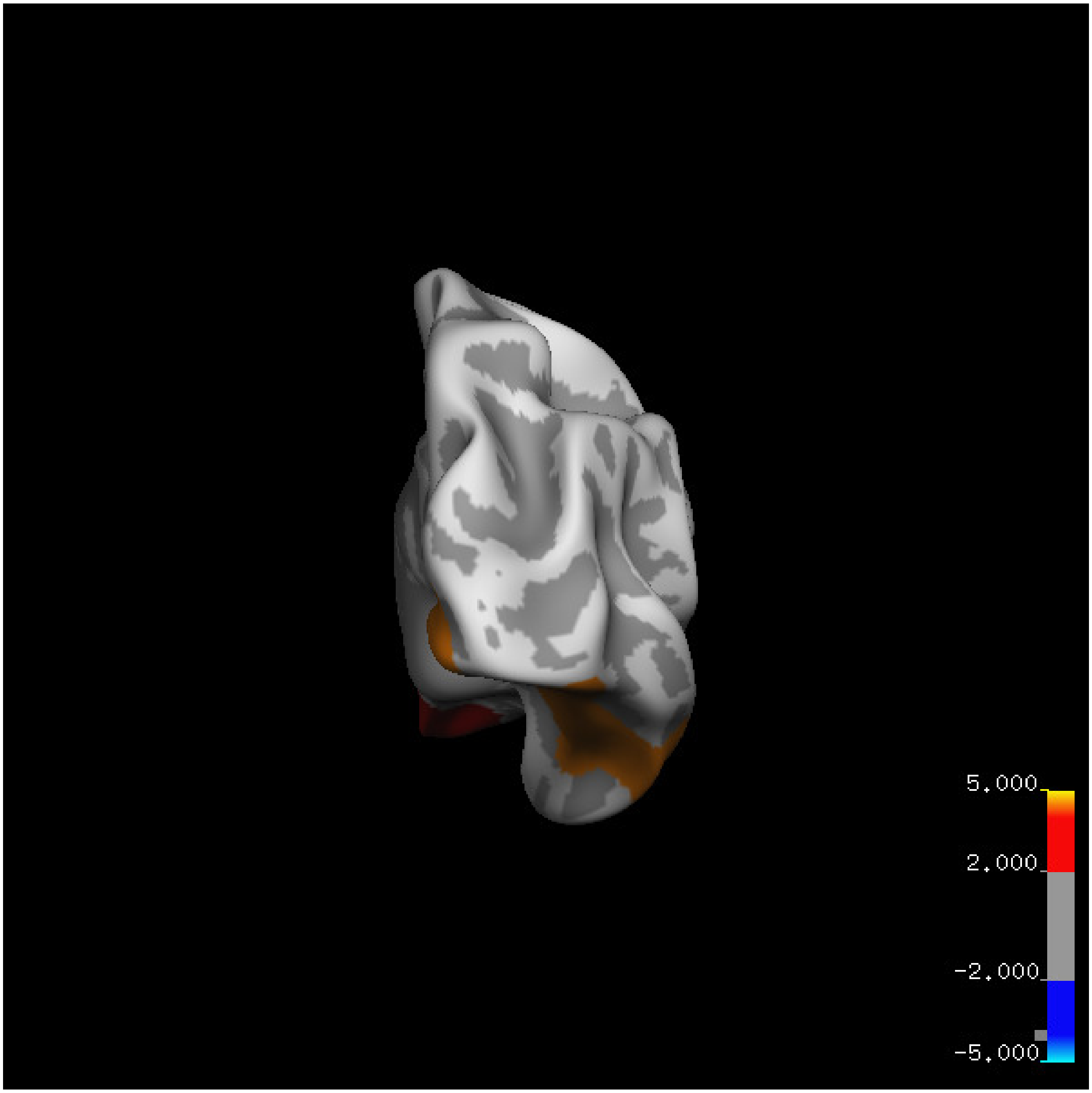

5.000
2.000
-2.000
-5.000

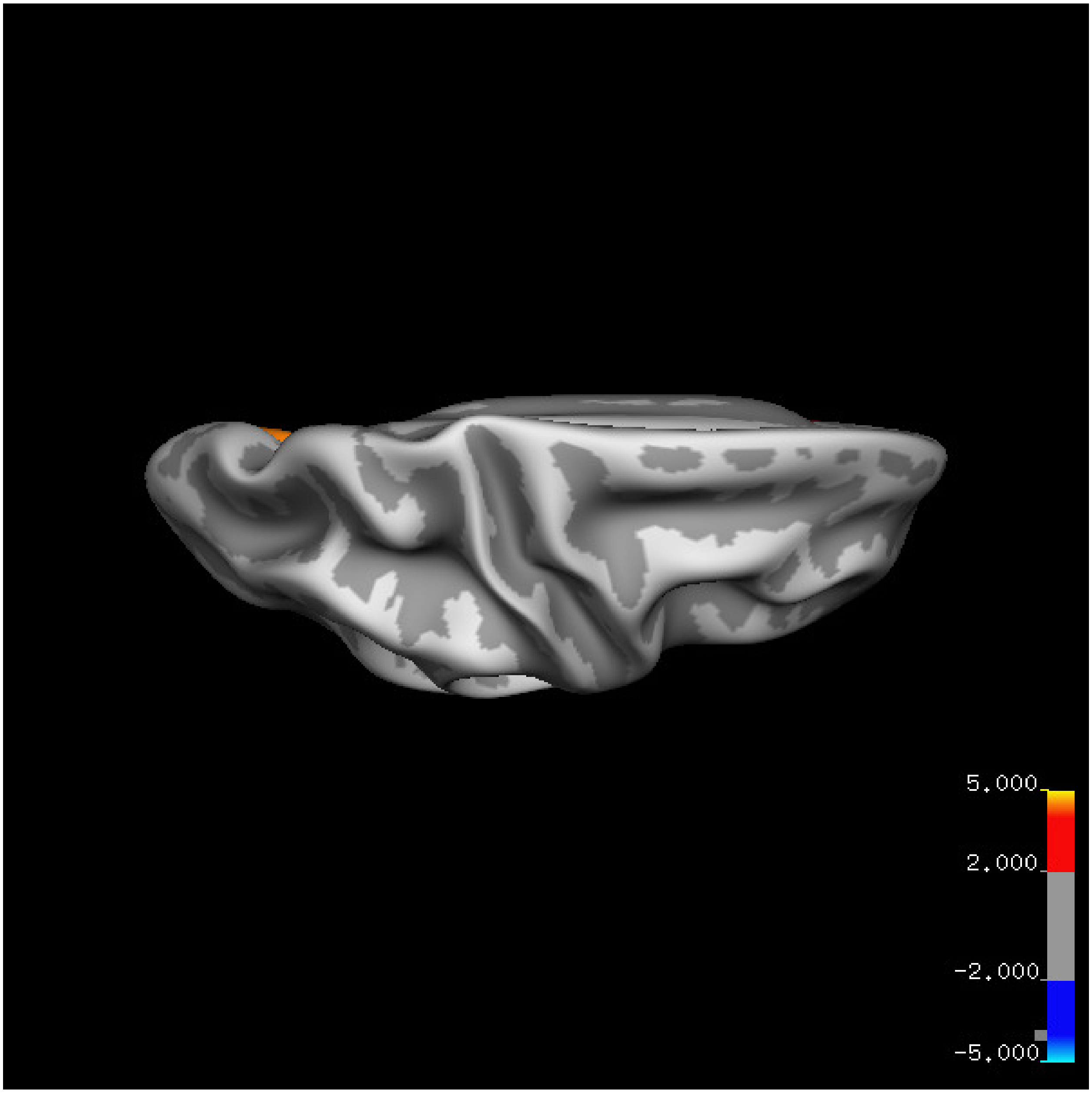


Cluster summary is shown below.

```
## # Cluster Growing Summary (mri_surfcluster)

## # $Id: mri_surfcluster.c,v 1.57.2.3 2016/11/17 18:19:42 zkaufman Exp $

## # $Id: mrisurf.c,v 1.781.2.6 2016/12/27 16:47:14 zkaufman Exp $

## # CreationTime 2021/10/04-13:55:32-GMT

## # cmdline mri_surfcluster.bin --in surface_analyses/results/cluster_comparison/fsaverage6.sm15.rh/ar

## # cwd /gpfs/projects02/p274/cluster/projects/p003-sleep_b

## # sysname  Linux
```

```
## # hostname c1-20
## # machine  x86_64
## # FixVertexAreaFlag 1
## # FixSurfClusterArea 1
## #
## # Input      surface_analyses/results/cluster_comparison/fsaverage6.sm15.rh/area.above60.below_mean/
## # Frame Number      0
## # srcsubj fsaverage6
## # hemi rh
## # surface white
## # group_avg_surface_area 85131.2
## # group_avg_vtxarea_loaded 1
## # annot aparc
## # SUBJECTS_DIR /cluster/projects/p274/tools/mri/freesurfer/current/subjects
## # SearchSpace_mm2 79329.9
## # SearchSpace_vtx 37471
## # Bonferroni 0
## # Minimum Threshold 1.3
## # Maximum Threshold infinity
## # Threshold Sign    abs
## # AdjustThreshWhenOneTail 1
## # CW PValue Threshold: 0.01
## # Area Threshold    0 mm^2
## # CSD thresh  1.300000
## # CSD nreps    10000
## # CSD simtype  null-z
## # CSD contrast NA
```

```
## # CSD confint  90.000000

## # Overall max 5.38019 at vertex 27459

## # Overall min -2.71998 at vertex 23464

## # NClusters          2

## # FixMNI = 0

## #

## # ClusterNo  Max   VtxMax   Size(mm^2)  MNIX   MNIY   MNIZ    CWP    CWPLow    CWPHi   NVtxs    Wght

##    1        3.040   13372   6288.87     45.7  -32.2  -20.9  0.00010  0.00000  0.00020   2252     385

##    2        5.380   27459   3449.59      8.9   24.3  -20.7  0.00620  0.00520  0.00720   1455     407
```

### Thickness

#### Left hemisphere

##### Age 20-60

**Sleep above mean** Cluster corrected significance plots are shown below.

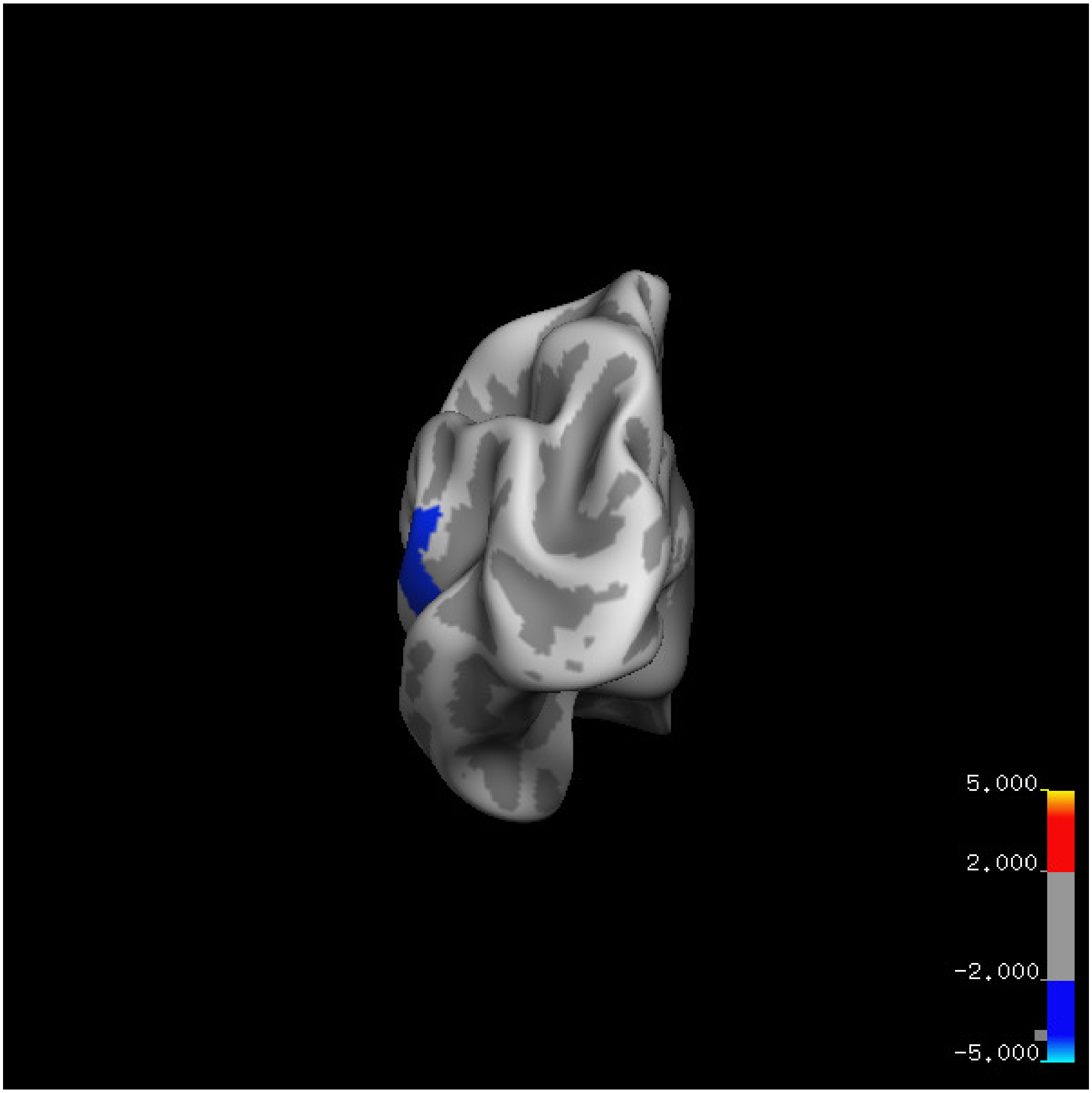

5.000
2.000
-2.000
-5.000

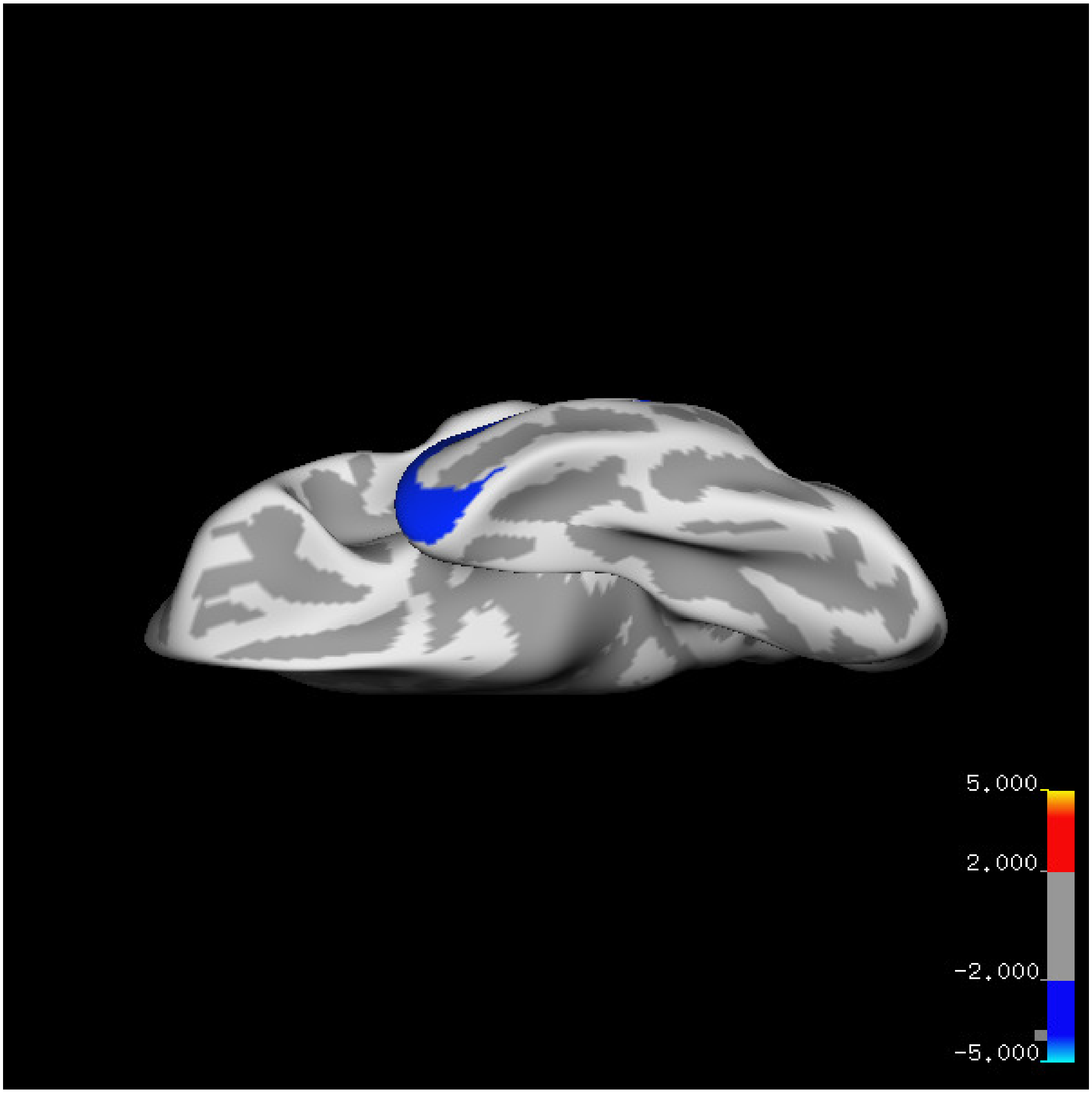
5.000
2.000
-2.000
-5.000

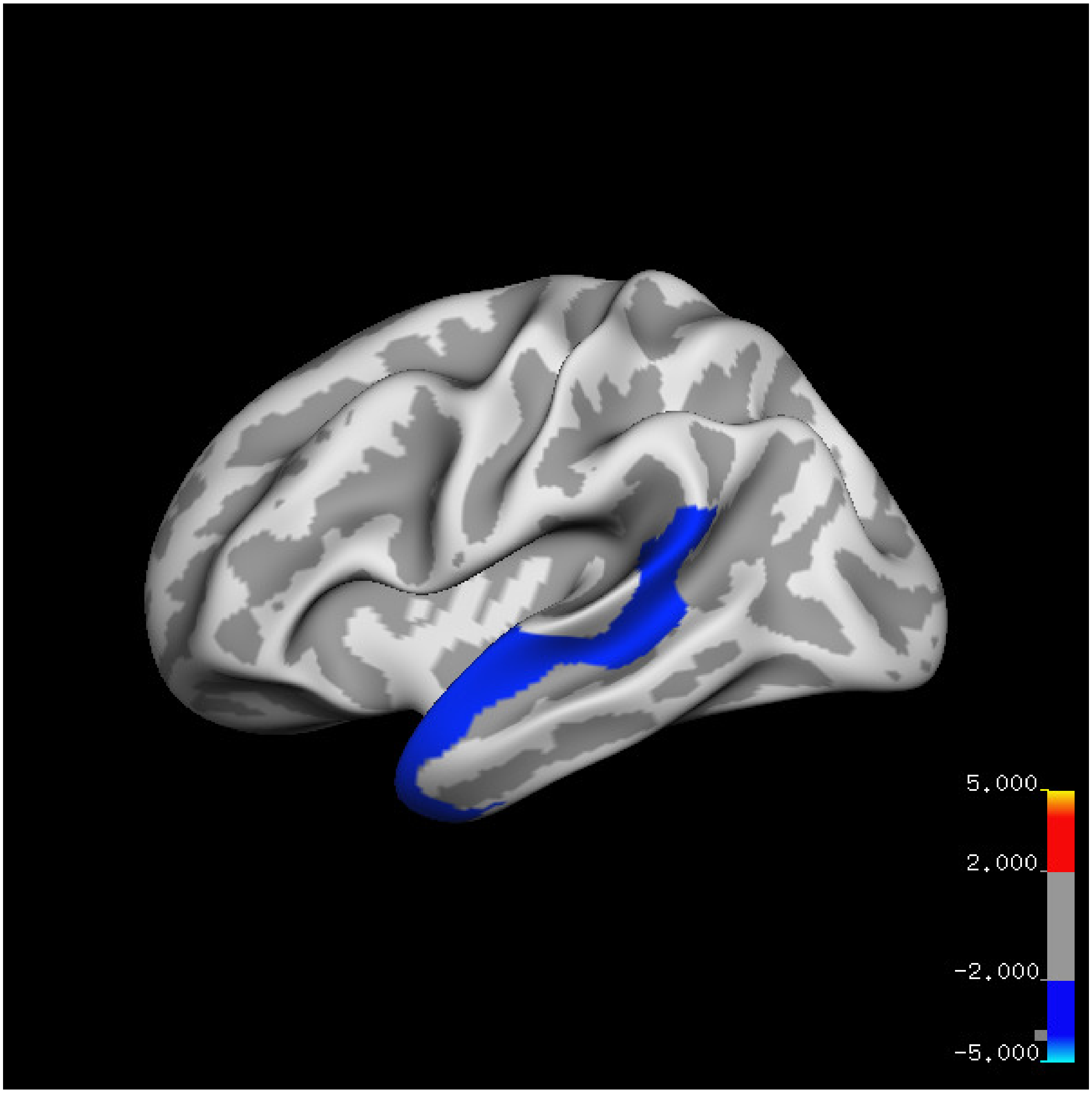

5.000
2.000
-2.000
-5.000

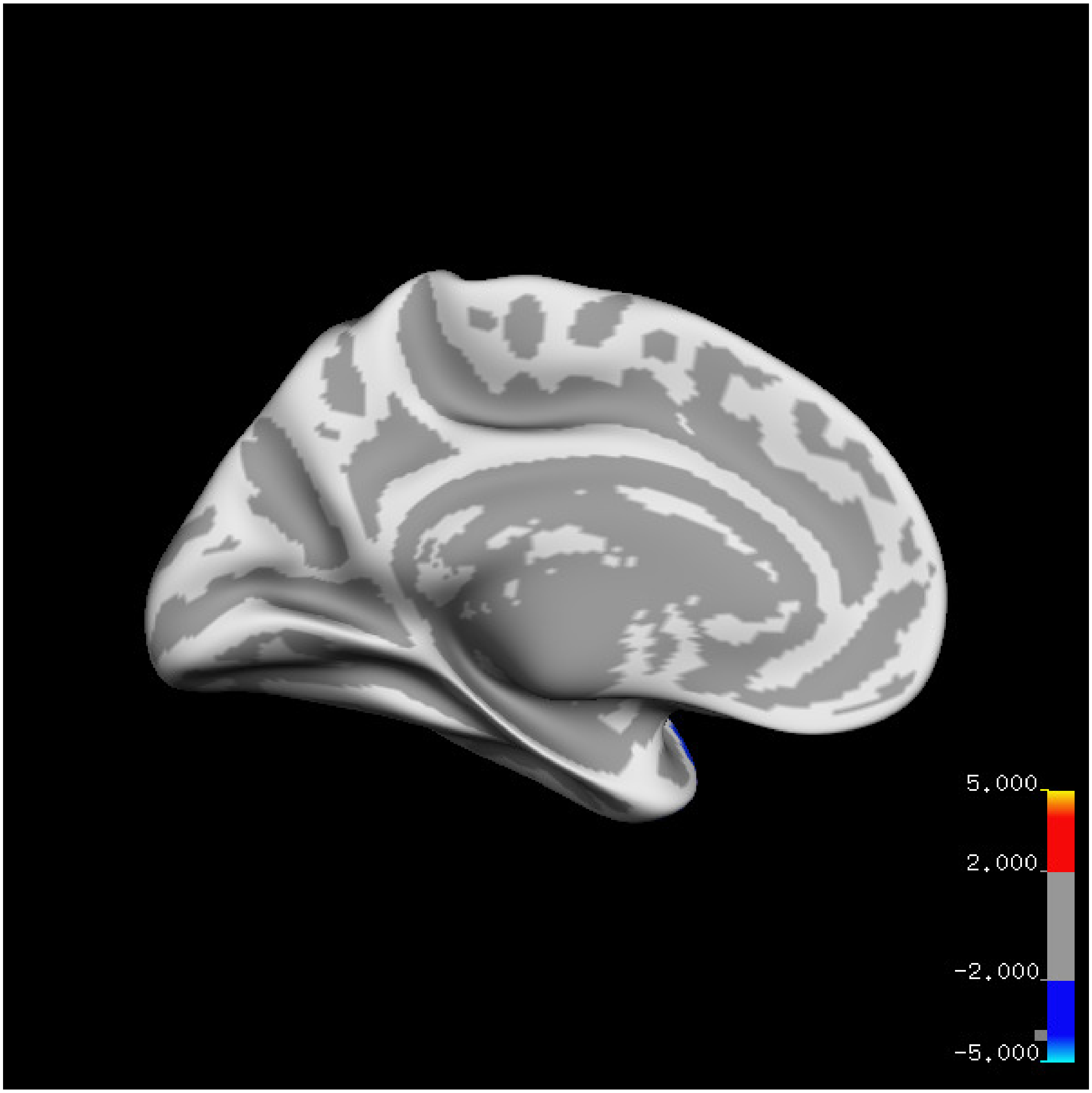

5.000
2.000
-2.000
-5.000

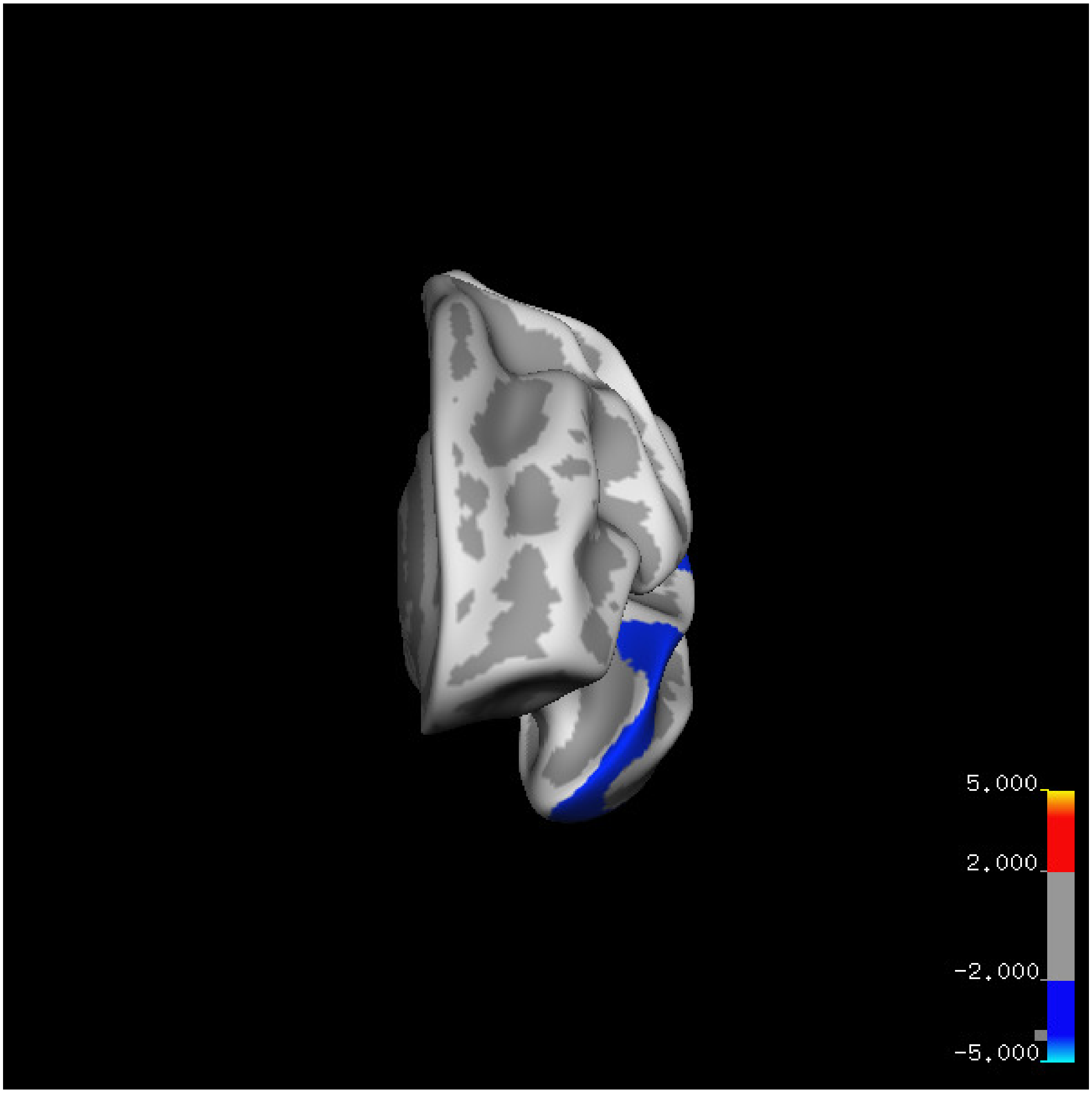
5.000
2.000
-2.000
-5.000

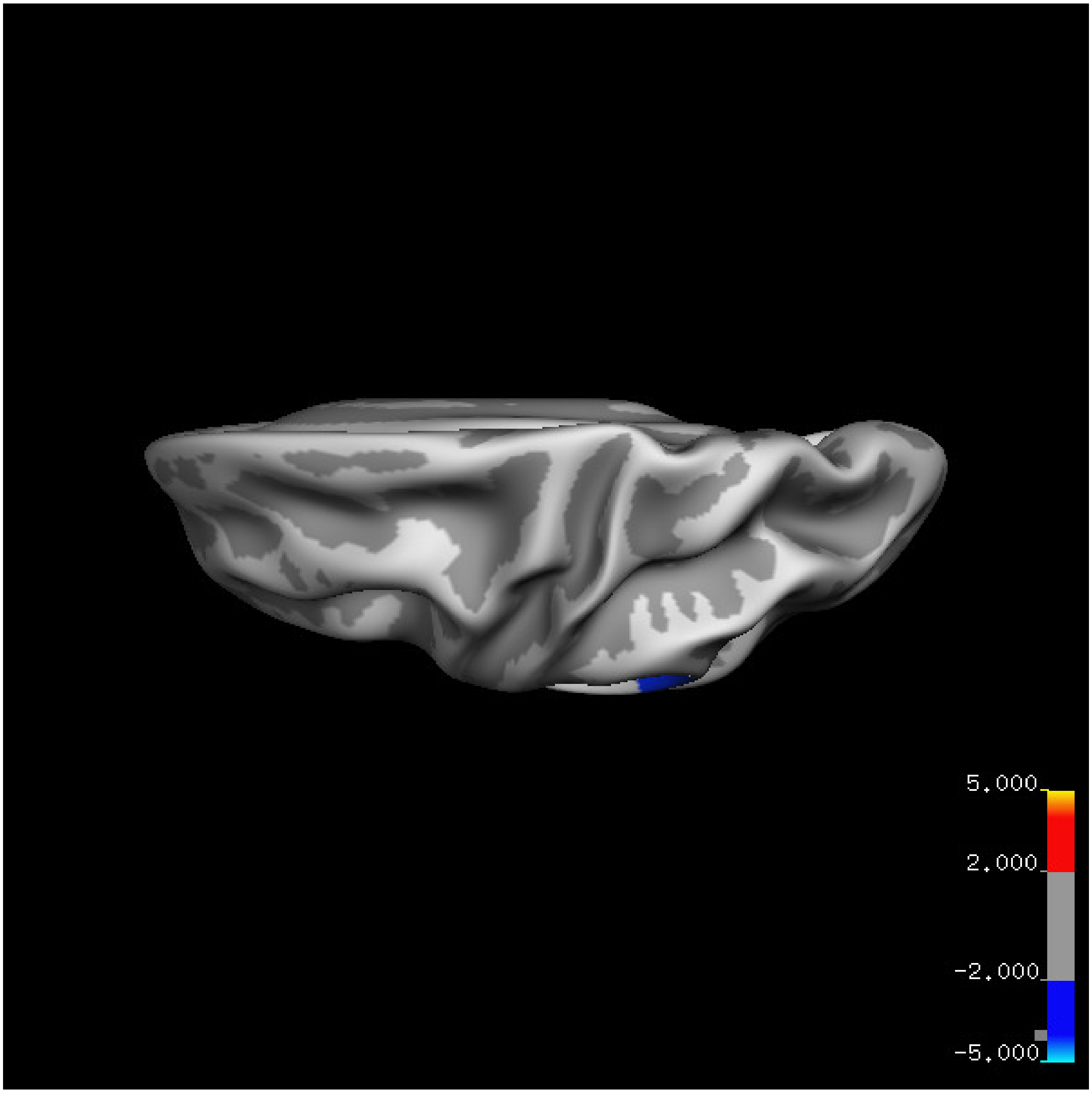
5.000
2.000
-2.000
-5.000

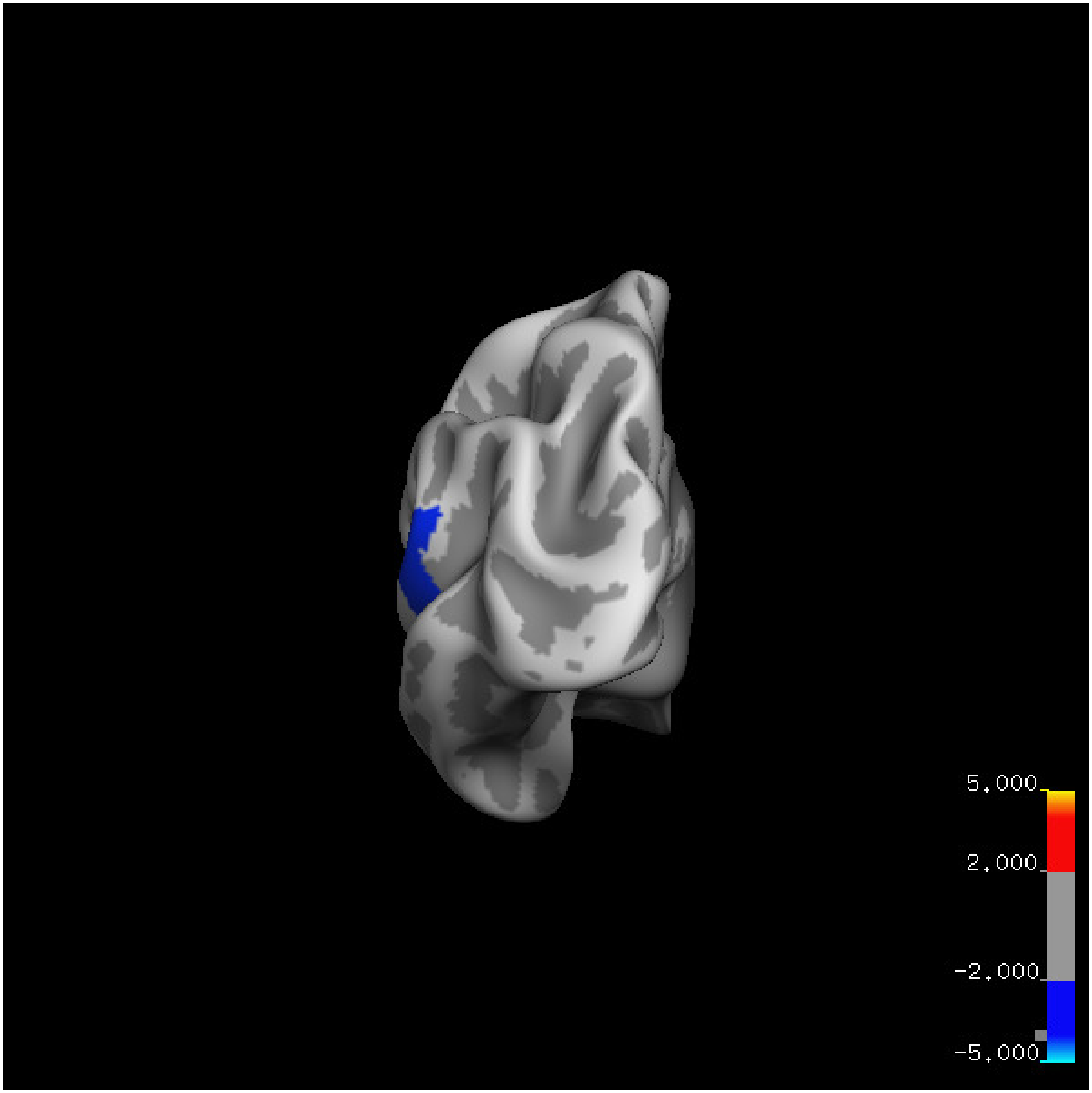
5.000
2.000
-2.000
-5.000

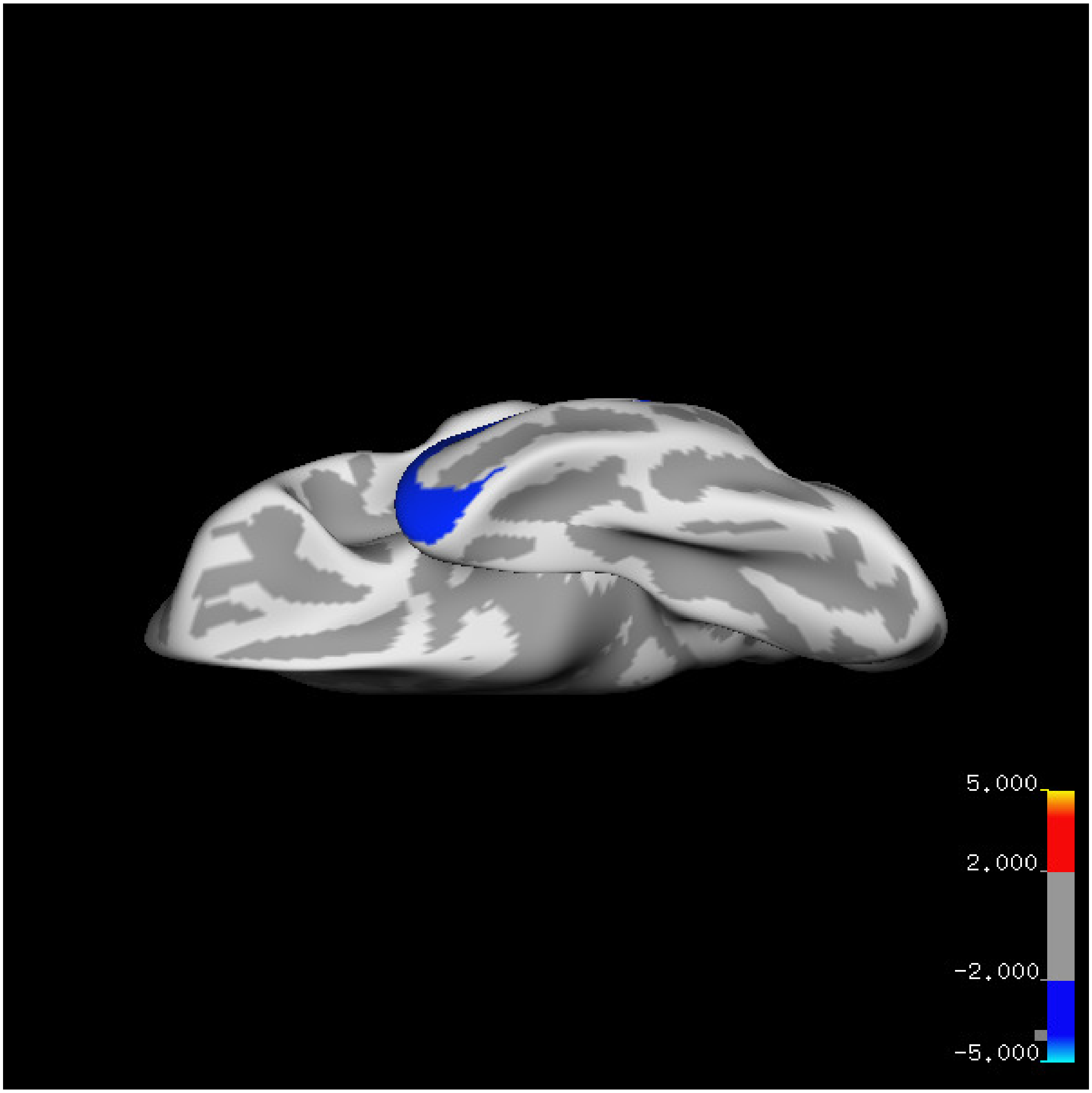
5.000
2.000
-2.000
-5.000

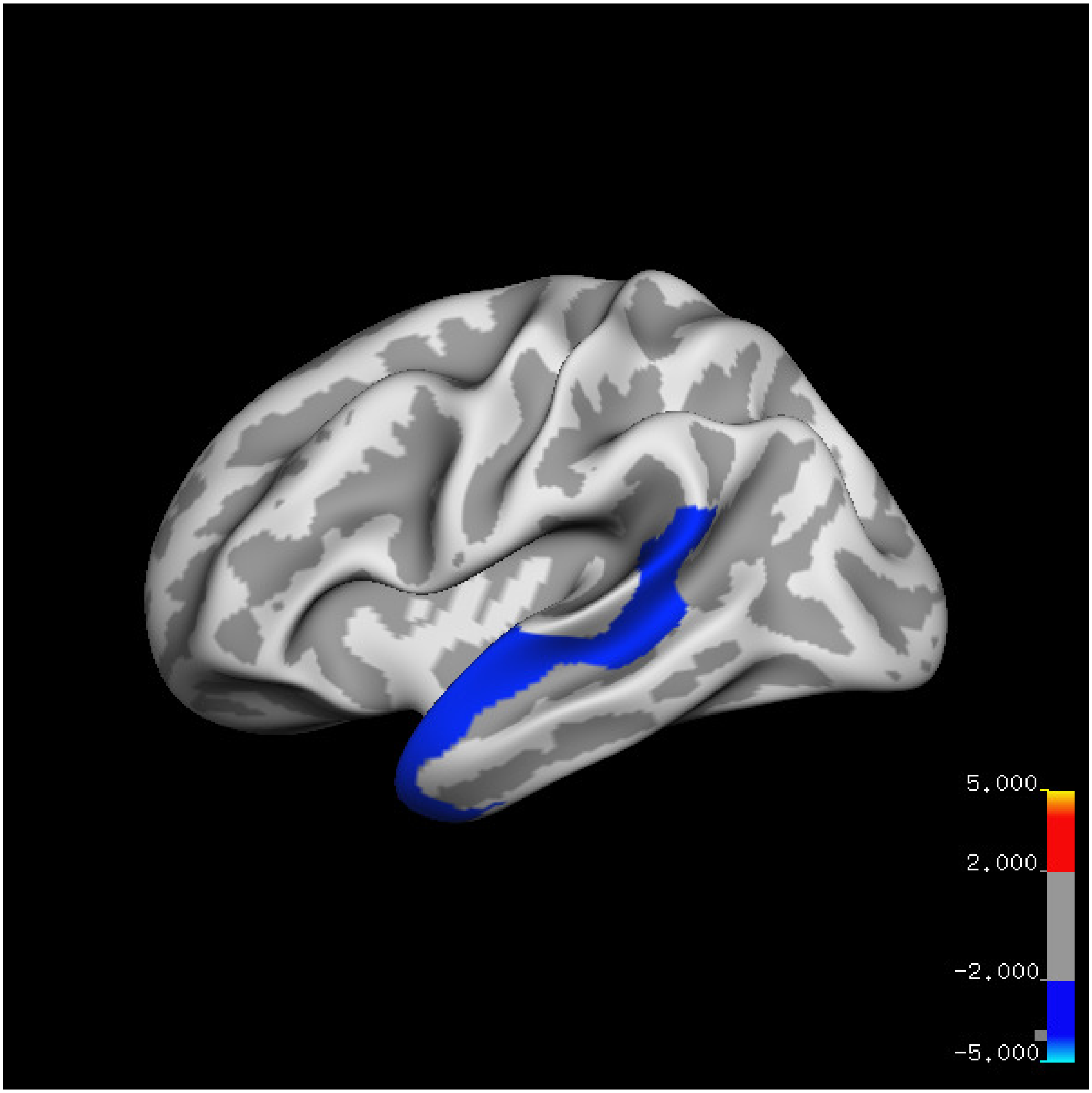
5.000
2.000
-2.000
-5.000

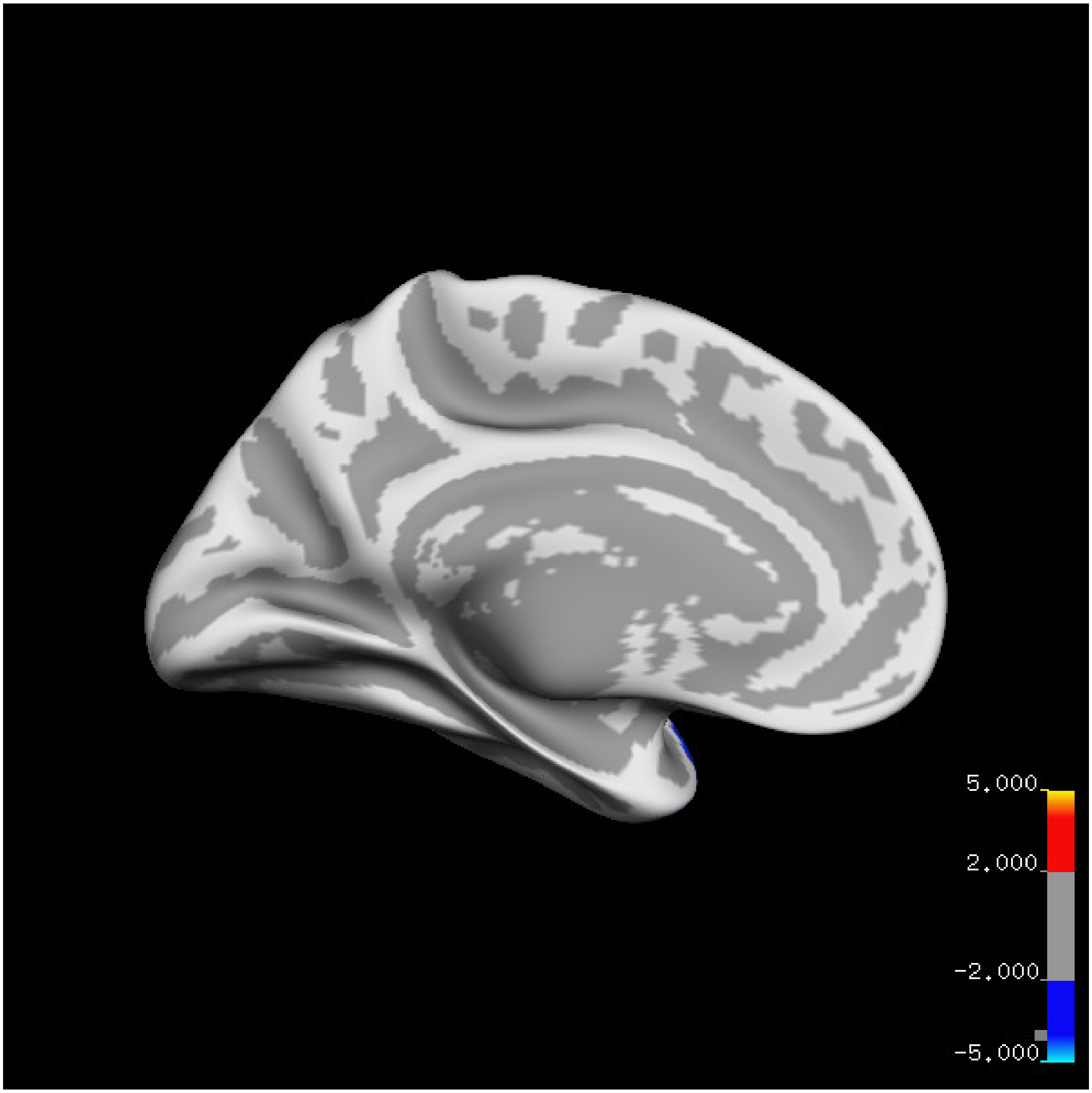
5.000
2.000
-2.000
-5.000

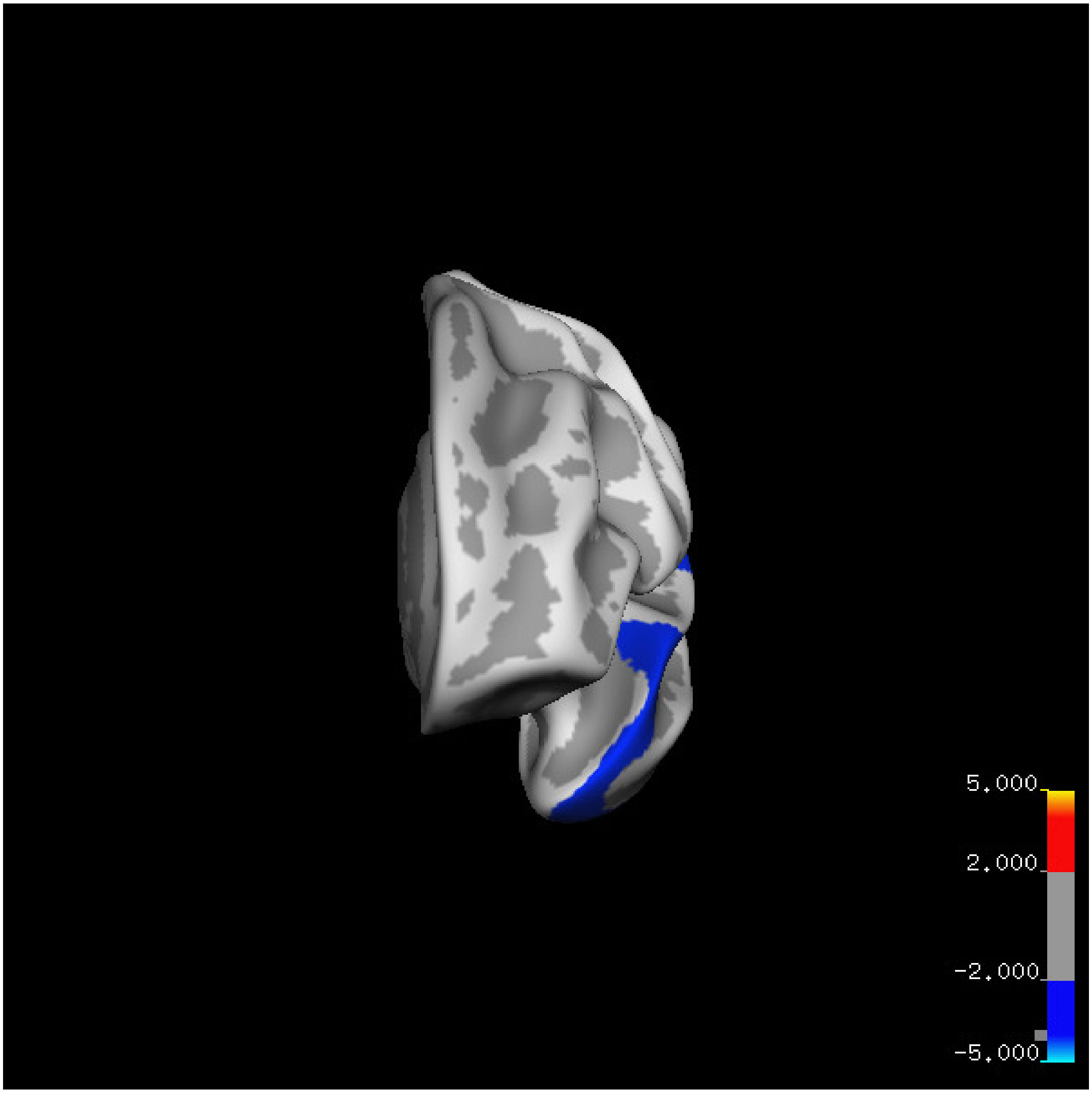
5.000
2.000
-2.000
-5.000

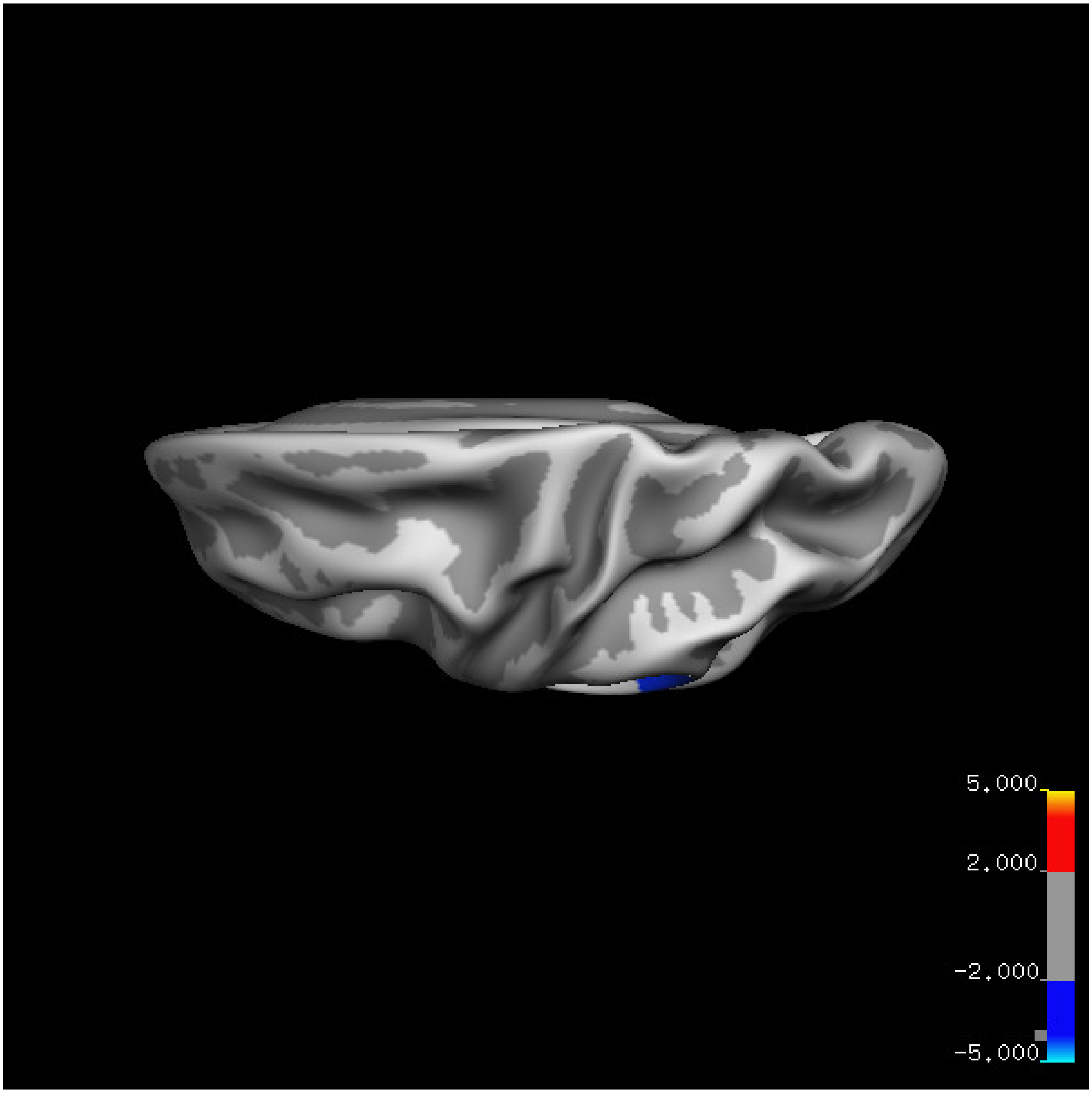


Cluster summary is shown below.

```
## # Cluster Growing Summary (mri_surfcluster)

## # $Id: mri_surfcluster.c,v 1.57.2.3 2016/11/17 18:19:42 zkaufman Exp $

## # $Id: mrisurf.c,v 1.781.2.6 2016/12/27 16:47:14 zkaufman Exp $

## # CreationTime 2021/10/04-10:48:07-GMT

## # cmdline mri_surfcluster.bin --in surface_analyses/results/cluster_comparison/fsaverage6.sm15.lh/th

## # cwd /gpfs/projects02/p274/cluster/projects/p003-sleep_b

## # sysname  Linux
```

```
## # hostname c1-18
## # machine  x86_64
## # FixVertexAreaFlag 1
## # FixSurfClusterArea 1
## #
## # Input      surface_analyses/results/cluster_comparison/fsaverage6.sm15.lh/thickness.20_60.above_mea
## # Frame Number      0
## # srcsubj fsaverage6
## # hemi lh
## # surface white
## # group_avg_surface_area 84969.3
## # group_avg_vtxarea_loaded 1
## # annot aparc
## # SUBJECTS_DIR /cluster/projects/p274/tools/mri/freesurfer/current/subjects
## # SearchSpace_mm2 79241.3
## # SearchSpace_vtx 37476
## # Bonferroni 0
## # Minimum Threshold 1.3
## # Maximum Threshold infinity
## # Threshold Sign    abs
## # AdjustThreshWhenOneTail 1
## # CW PValue Threshold: 0.01
## # Area Threshold    0 mm^2
## # CSD thresh  1.300000
## # CSD nreps    10000
## # CSD simtype  null-z
## # CSD contrast NA
```

```
## # CSD confint  90.000000

## # Overall max 2.4495 at vertex 40575

## # Overall min -3.1882 at vertex 27685

## # NClusters          1

## # FixMNI = 0

## #

## # ClusterNo  Max   VtxMax   Size(mm^2)  MNIX   MNIY   MNIZ    CWP    CWPLow    CWPHi   NVtxs    Wght

##    1       -3.188   27685   3091.96    -50.0  -38.6    1.7  0.00050  0.00020  0.00080   1443    -266
```

**Sleep below mean** Cluster corrected significance plots are shown below.

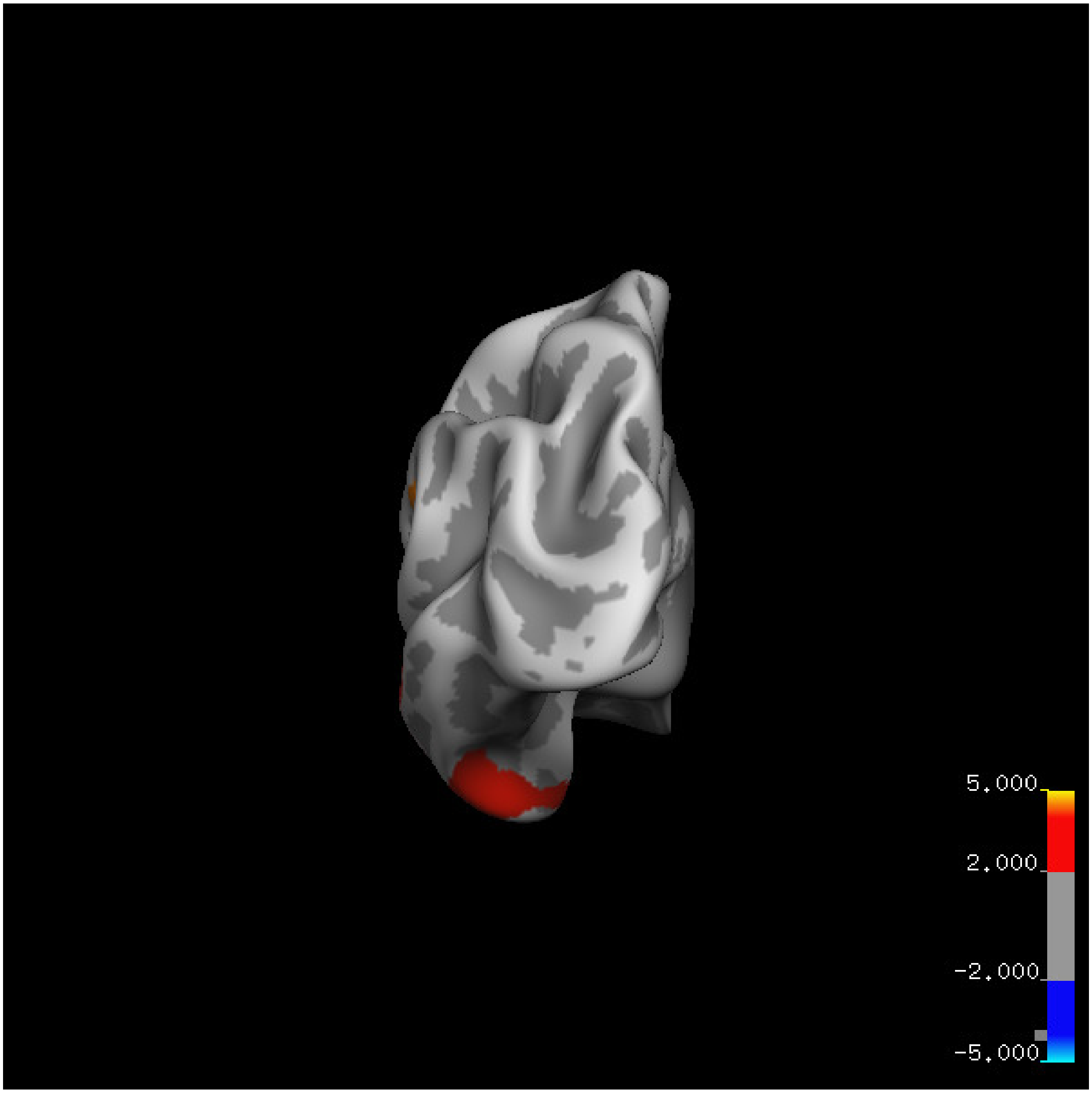
5.000
2.000
-2.000
-5.000

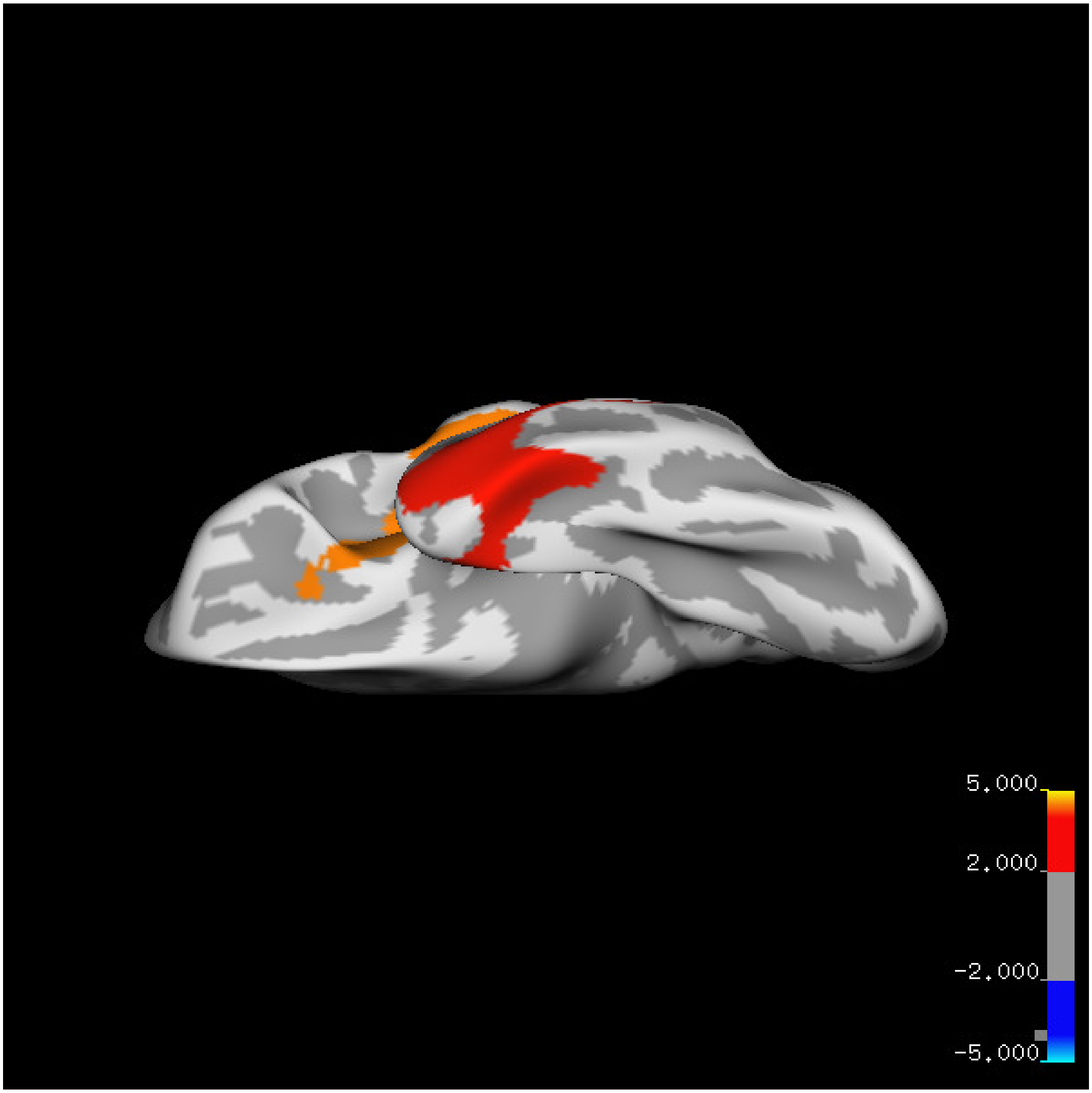
5.000
2.000
-2.000
-5.000

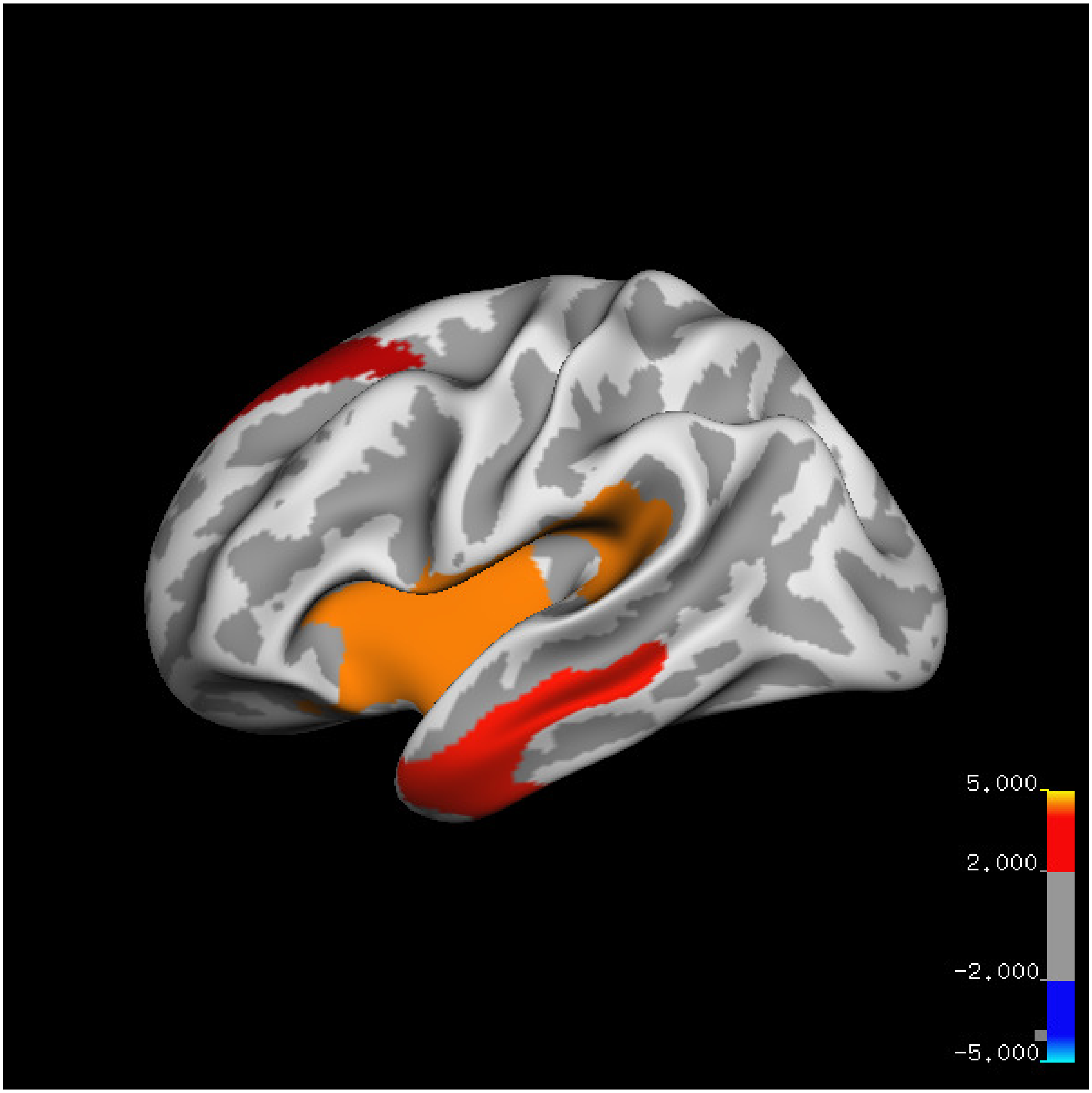
5.000
2.000
-2.000
-5.000

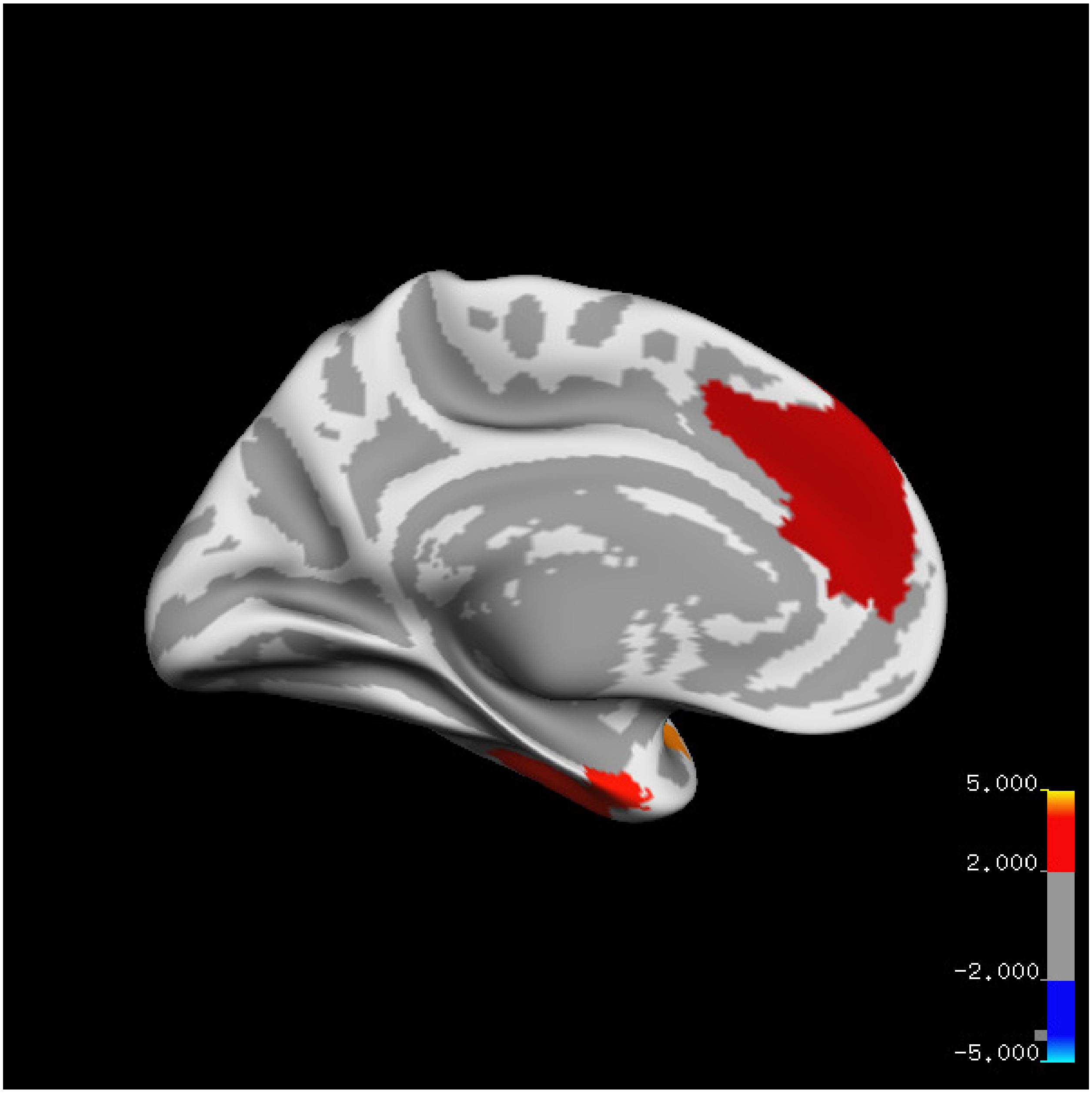
5.000
2.000
-2.000
-5.000

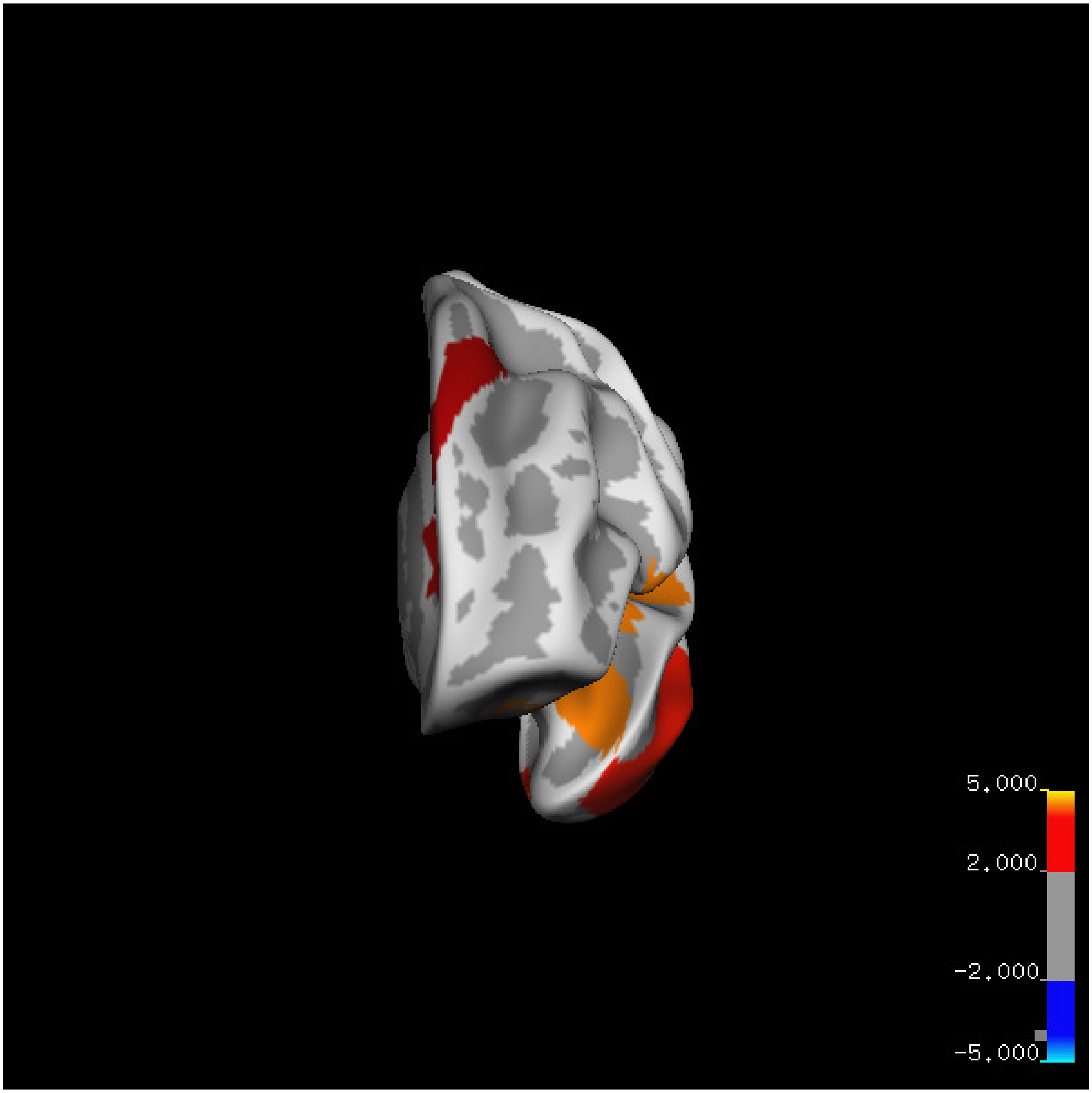
5.000
2.000
-2.000
-5.000

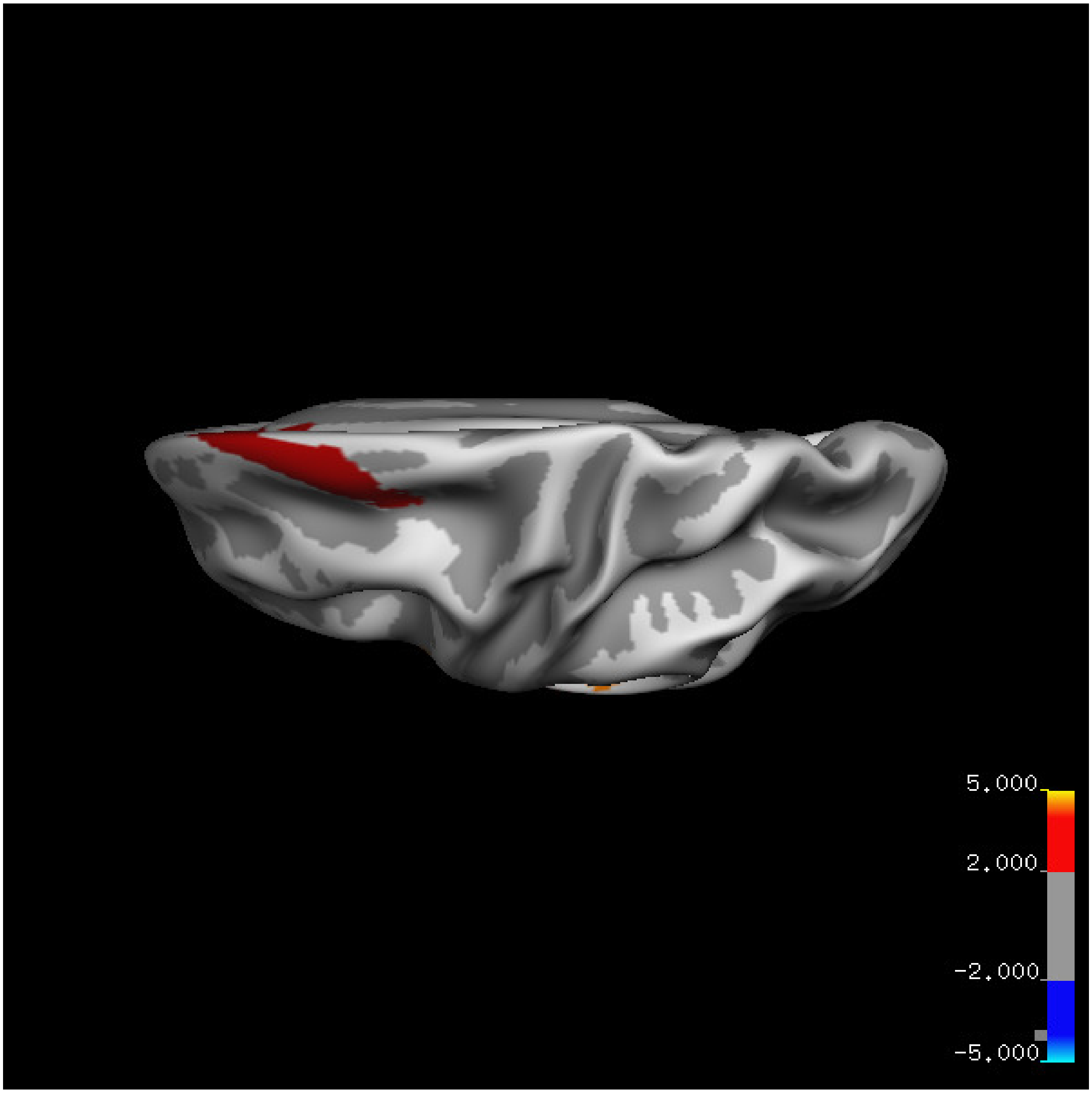
5.000
2.000
-2.000
-5.000

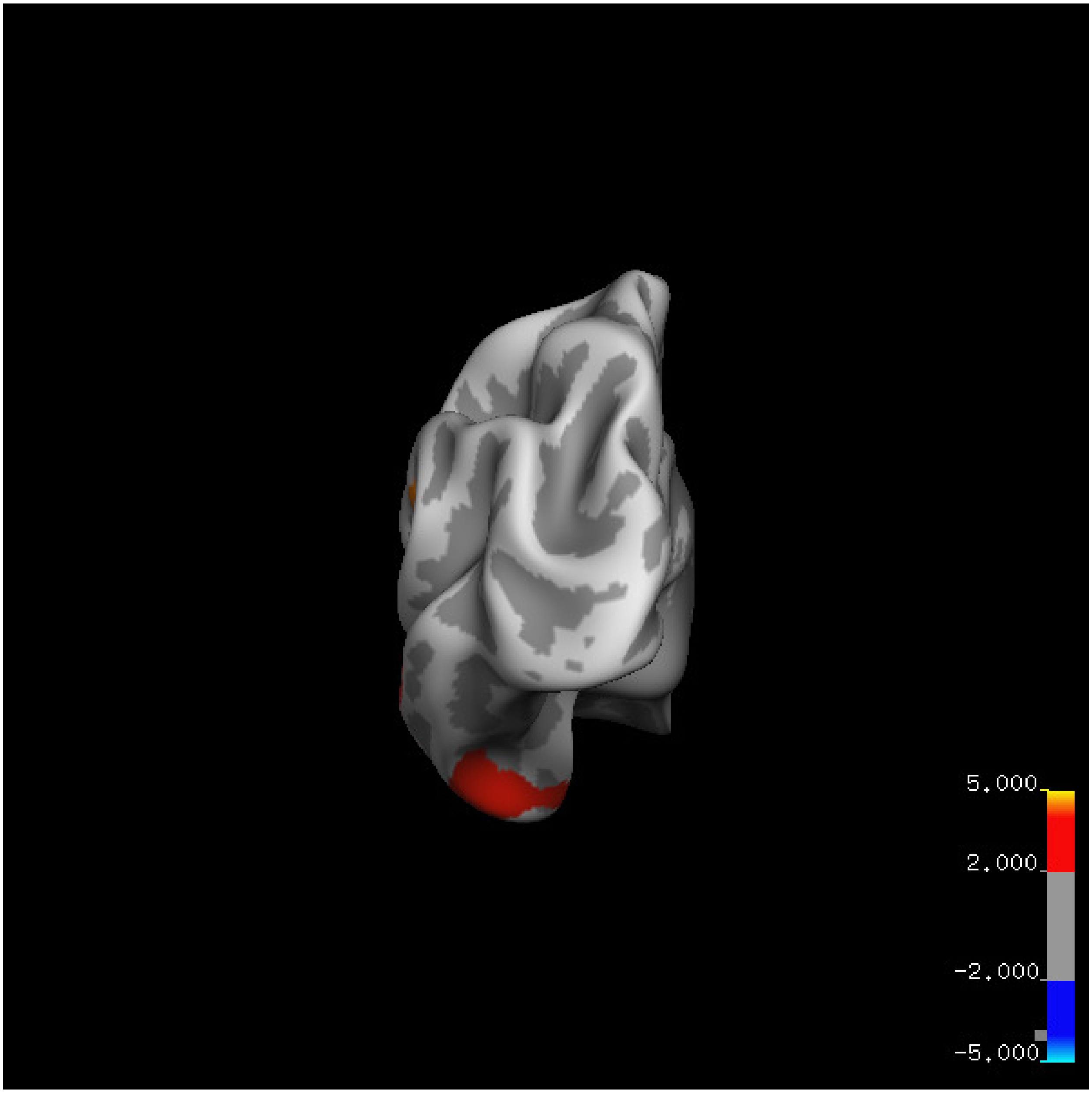
5.000
2.000
-2.000
-5.000

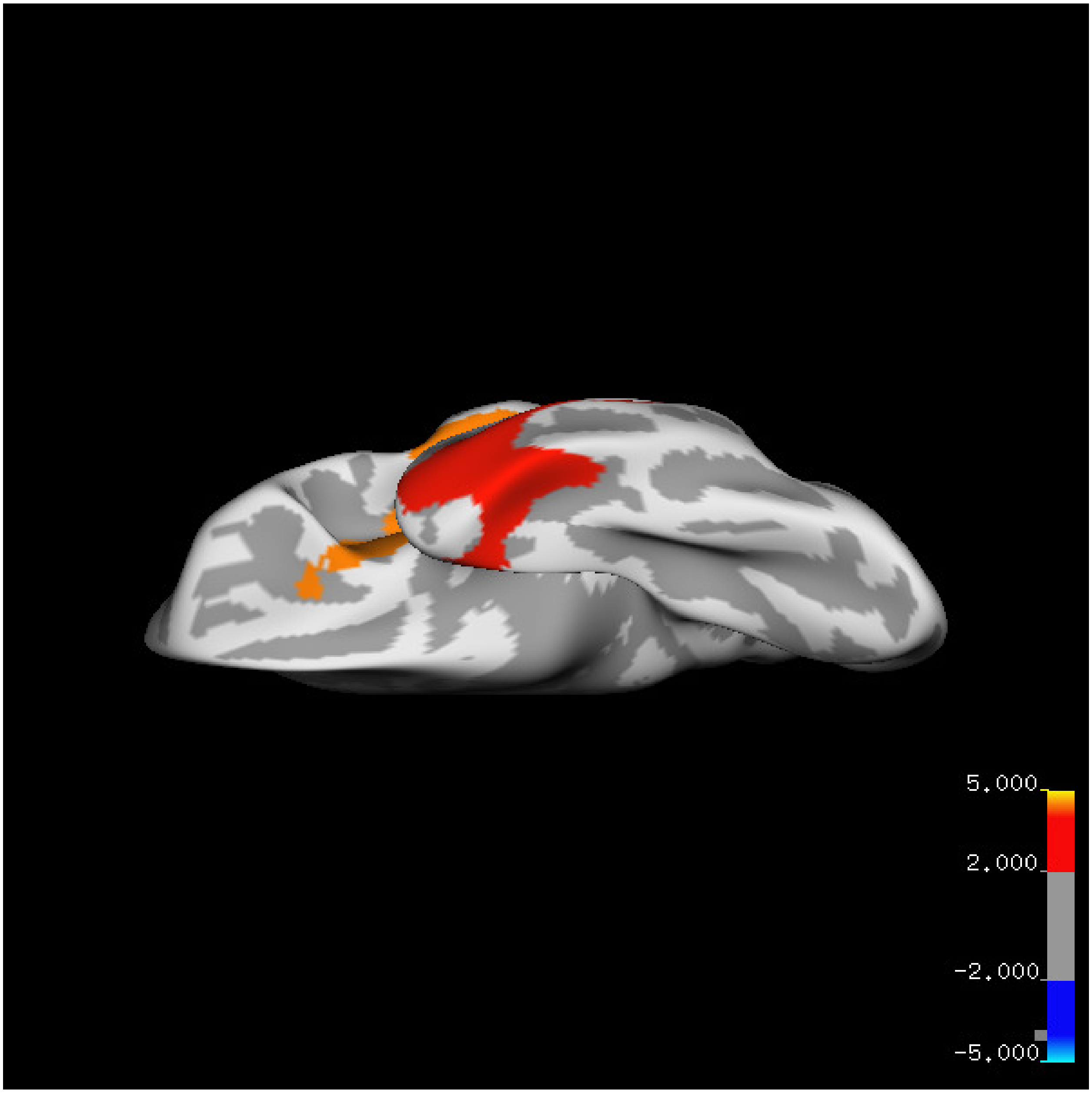
5.000
2.000
-2.000
-5.000

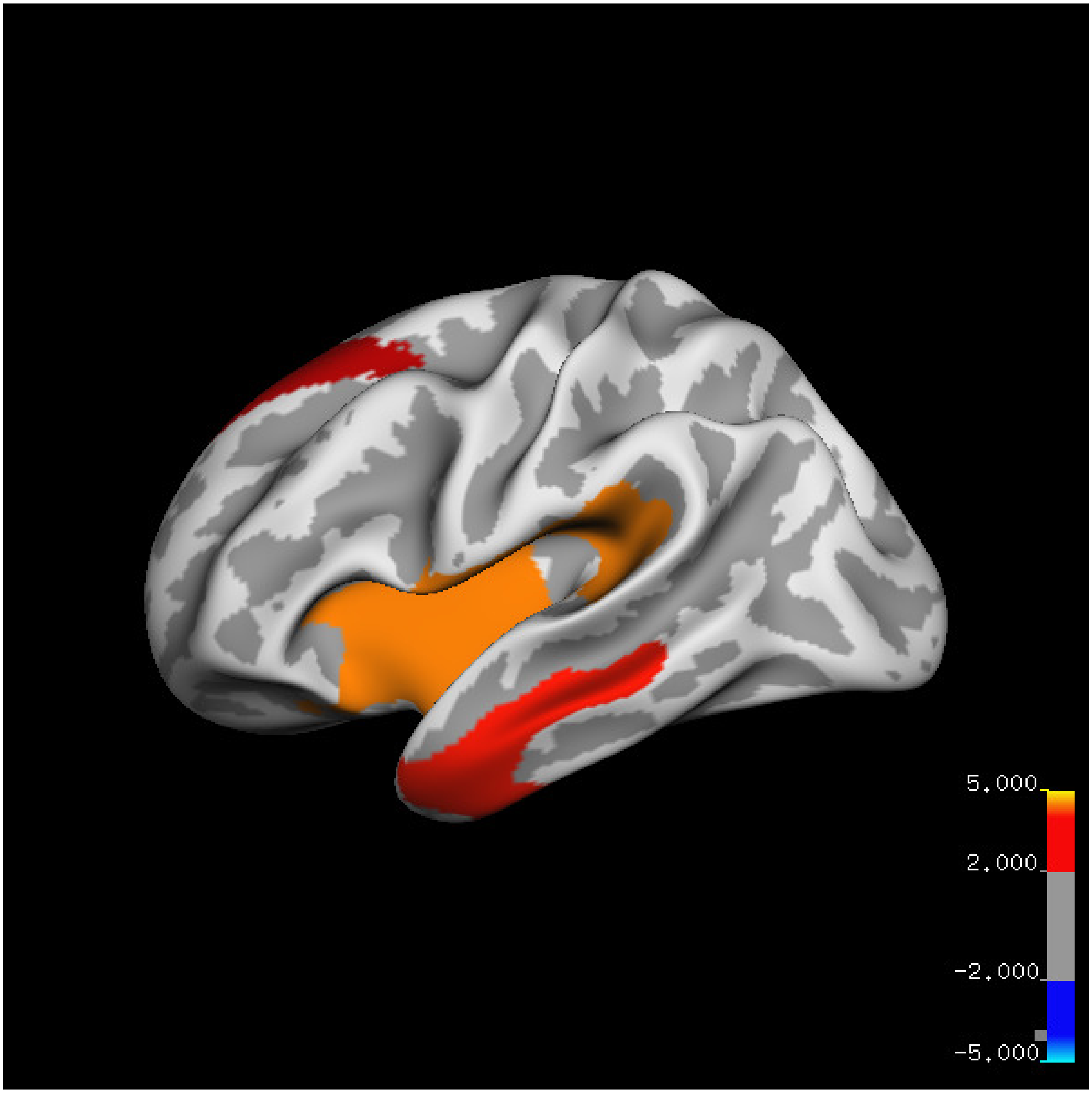
5.000
2.000
-2.000
-5.000

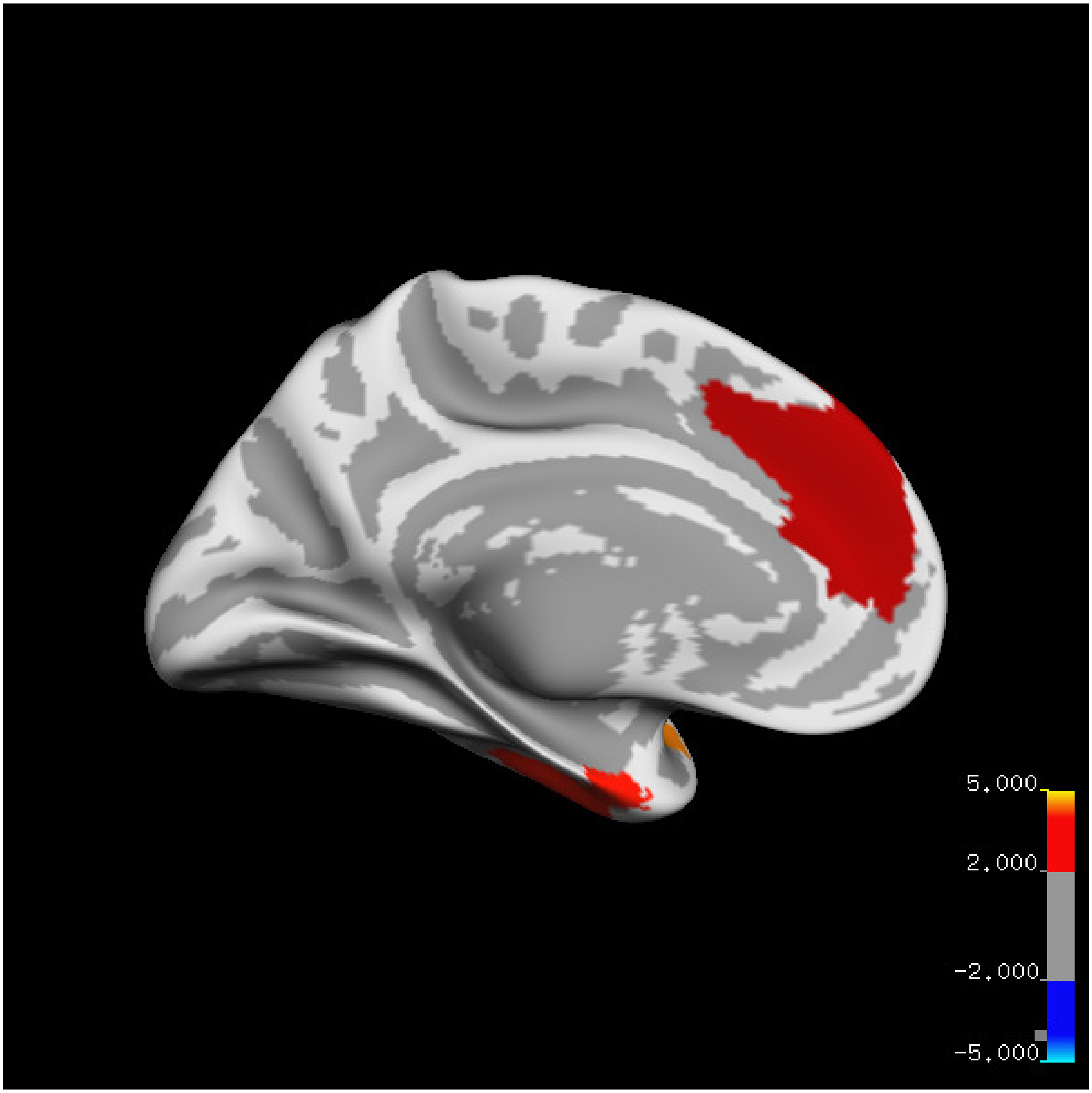
5.000
2.000
-2.000
-5.000

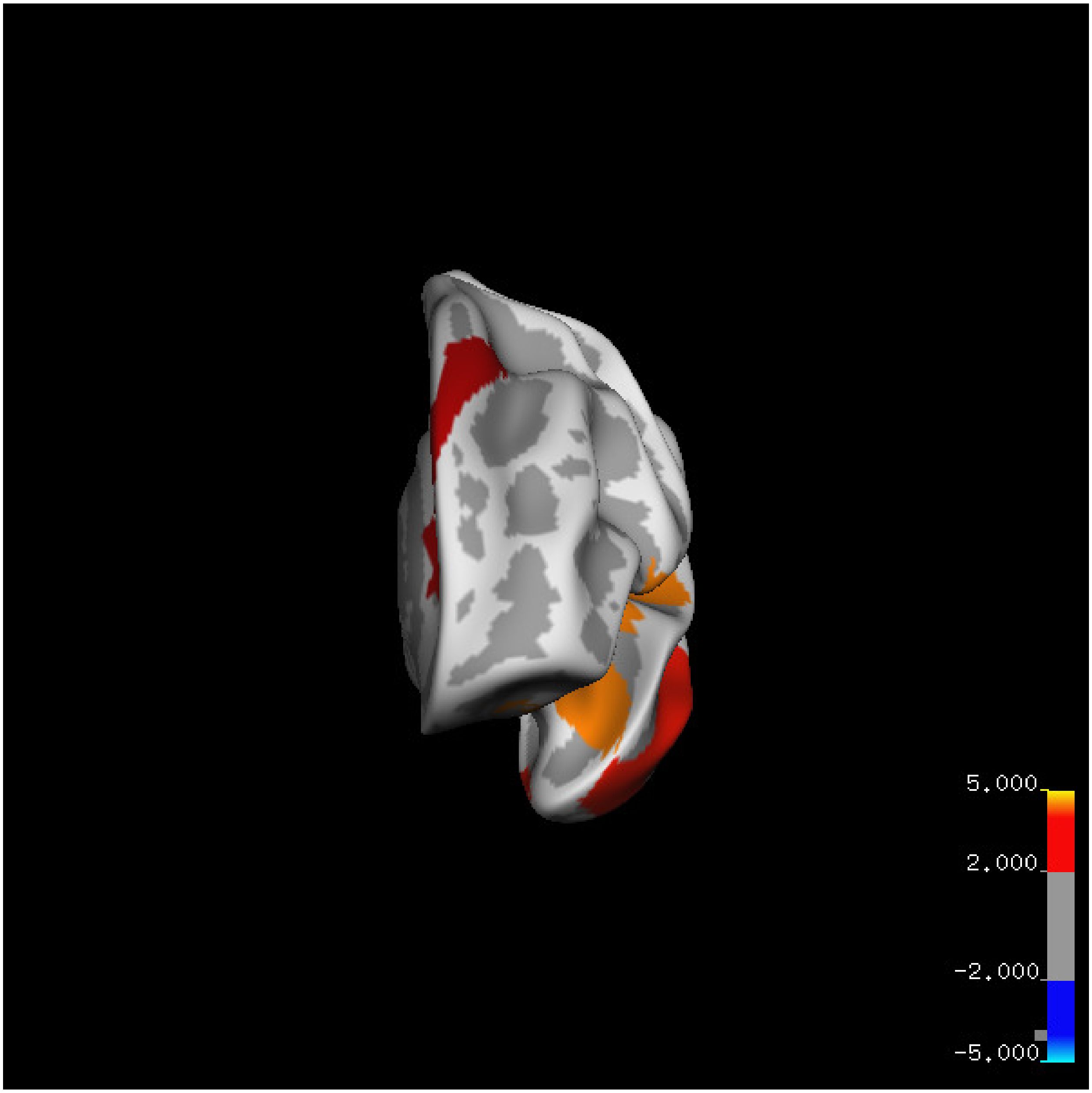
5.000
2.000
-2.000
-5.000

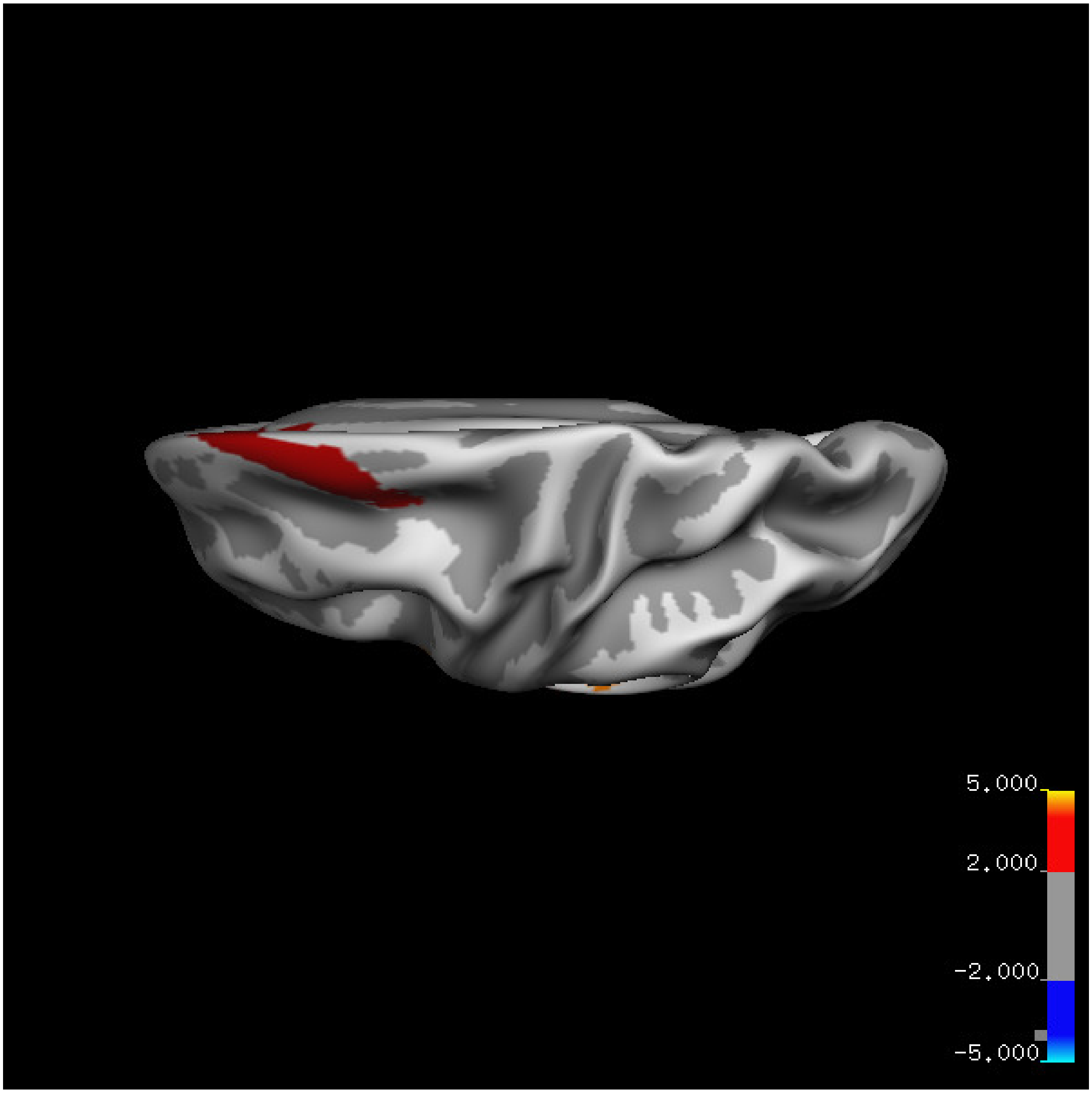


Cluster summary is shown below.

```
## # Cluster Growing Summary (mri_surfcluster)

## # $Id: mri_surfcluster.c,v 1.57.2.3 2016/11/17 18:19:42 zkaufman Exp $

## # $Id: mrisurf.c,v 1.781.2.6 2016/12/27 16:47:14 zkaufman Exp $

## # CreationTime 2021/10/04-11:24:45-GMT

## # cmdline mri_surfcluster.bin --in surface_analyses/results/cluster_comparison/fsaverage6.sm15.lh/th

## # cwd /gpfs/projects02/p274/cluster/projects/p003-sleep_b

## # sysname  Linux
```

```
## # hostname c1-18
## # machine  x86_64
## # FixVertexAreaFlag 1
## # FixSurfClusterArea 1
## #
## # Input      surface_analyses/results/cluster_comparison/fsaverage6.sm15.lh/thickness.20_60.below_mea
## # Frame Number      0
## # srcsubj fsaverage6
## # hemi lh
## # surface white
## # group_avg_surface_area 84969.3
## # group_avg_vtxarea_loaded 1
## # annot aparc
## # SUBJECTS_DIR /cluster/projects/p274/tools/mri/freesurfer/current/subjects
## # SearchSpace_mm2 79241.3
## # SearchSpace_vtx 37476
## # Bonferroni 0
## # Minimum Threshold 1.3
## # Maximum Threshold infinity
## # Threshold Sign    abs
## # AdjustThreshWhenOneTail 1
## # CW PValue Threshold: 0.01
## # Area Threshold    0 mm^2
## # CSD thresh  1.300000
## # CSD nreps    10000
## # CSD simtype  null-z
## # CSD contrast NA
```

```
## # CSD confint  90.000000

## # Overall max 5.67193 at vertex 19748

## # Overall min -0.898067 at vertex 24665

## # NClusters          3

## # FixMNI = 0

## #

## # ClusterNo  Max   VtxMax   Size(mm^2)  MNIX   MNIY   MNIZ    CWP    CWPLow    CWPHi   NVtxs    Wght

##    1        4.495   35252   4843.19    -42.0  -25.8   22.1  0.00010  0.00000  0.00020   2901     628

##    2        3.540   28386   3050.14    -51.7   -0.5  -32.7  0.00070  0.00040  0.00100   1193     251

##    3        3.596    7585   2477.64     -7.4   35.8   32.4  0.00410  0.00330  0.00490    985     198
```

**Age above 60**

**Sleep above mean** Cluster corrected significance plots are shown below.

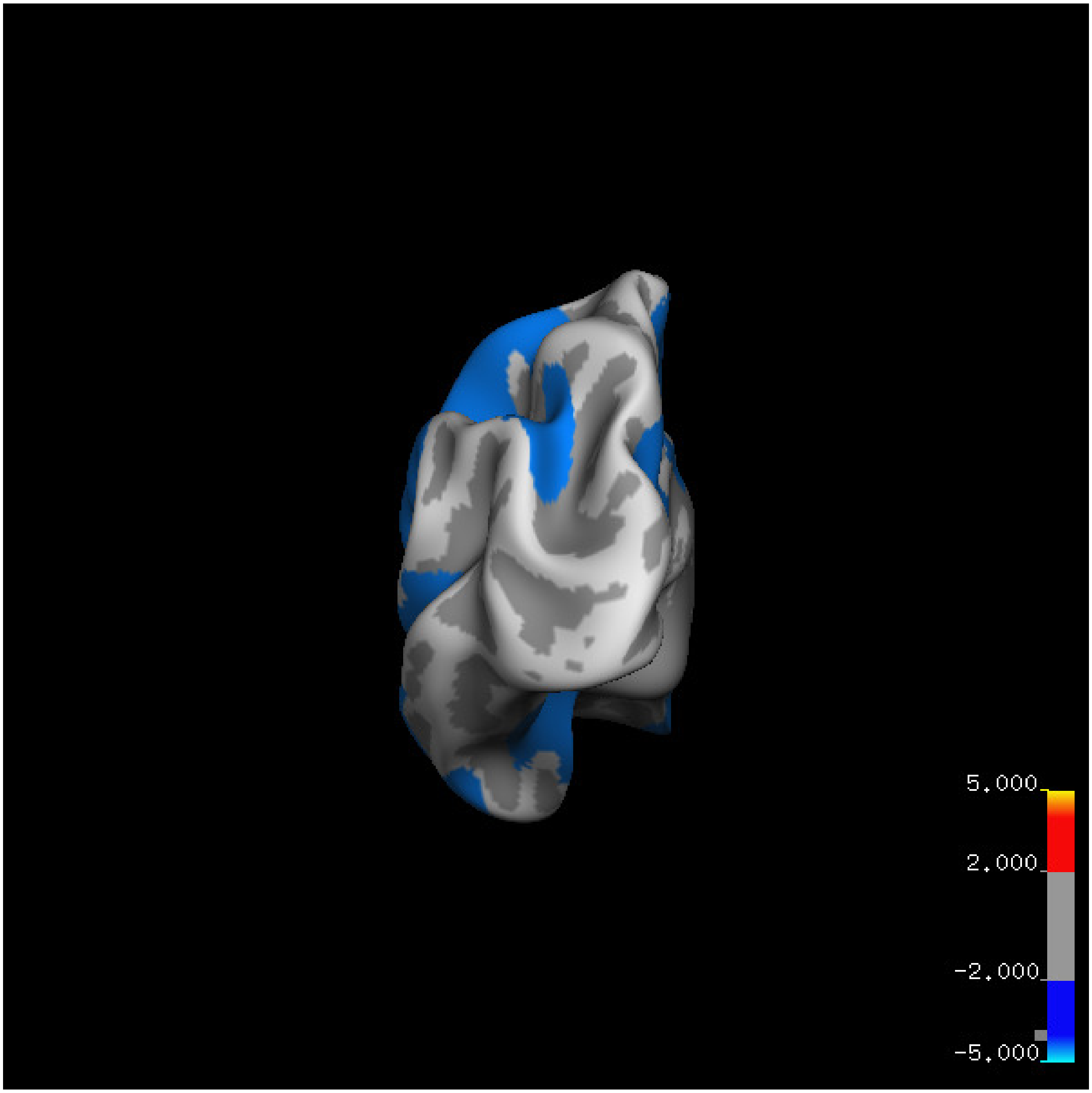
5.000
2.000
-2.000
-5.000

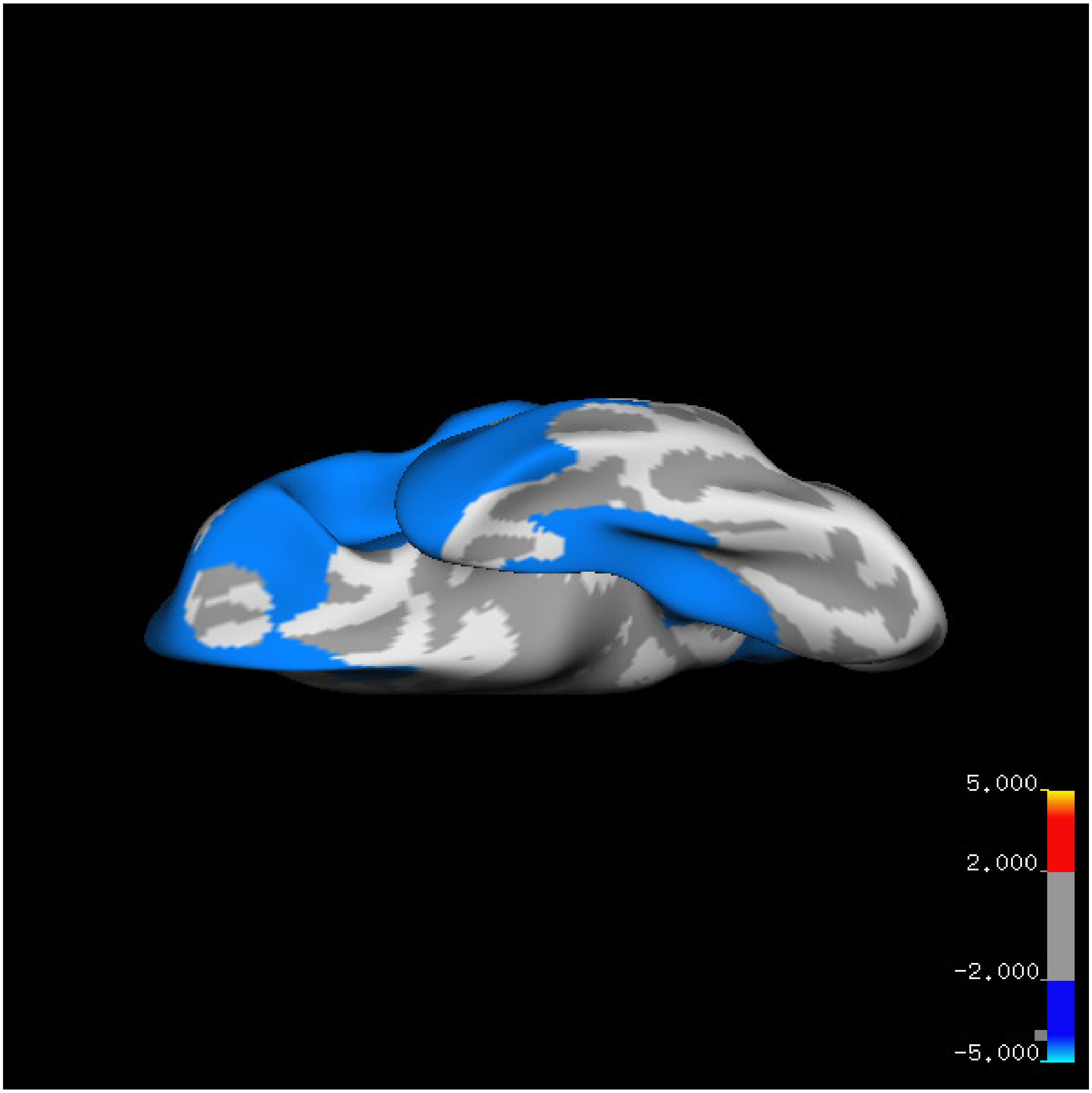
5.000
2.000
-2.000
-5.000

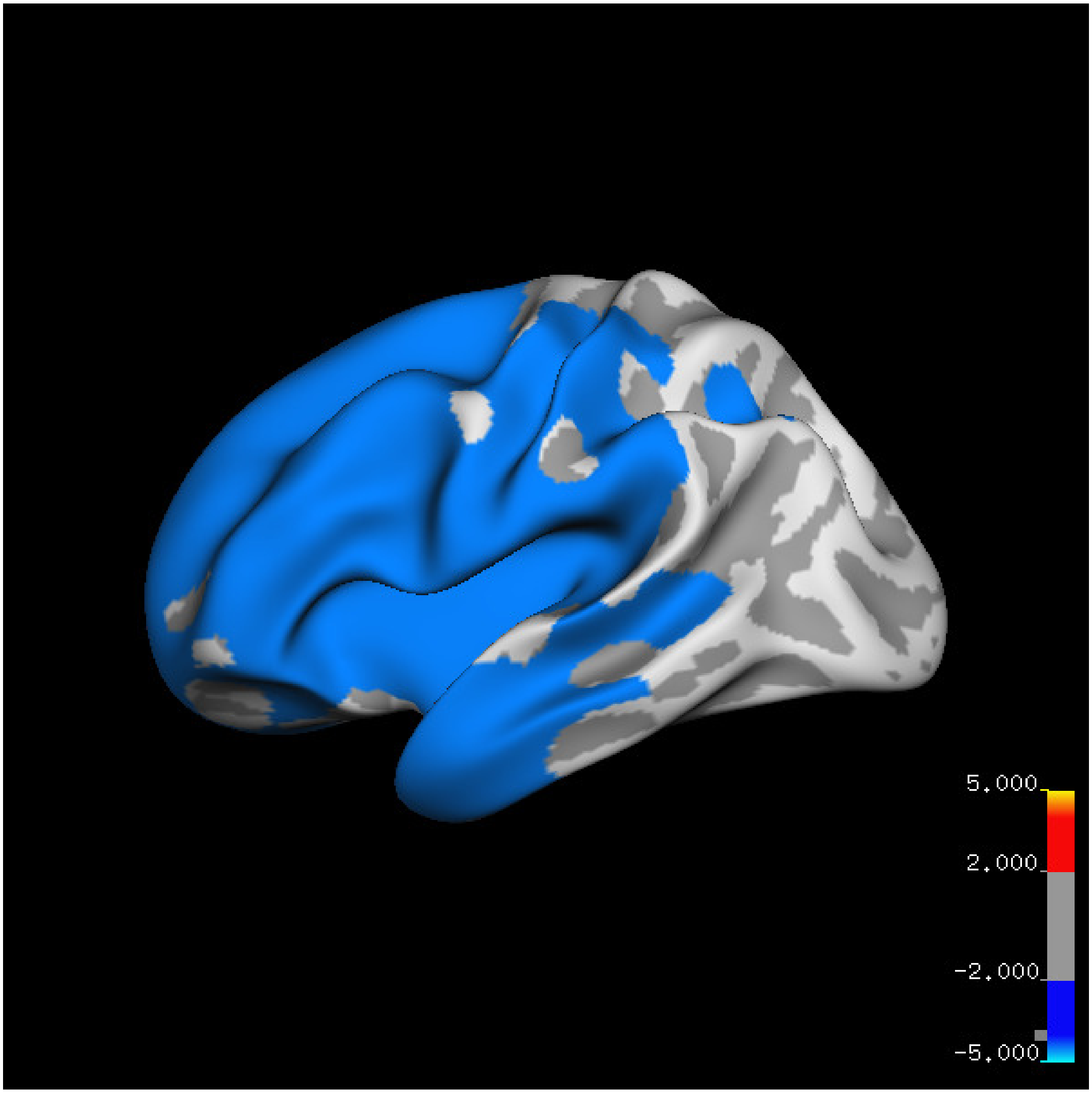
5.000
2.000
-2.000
-5.000

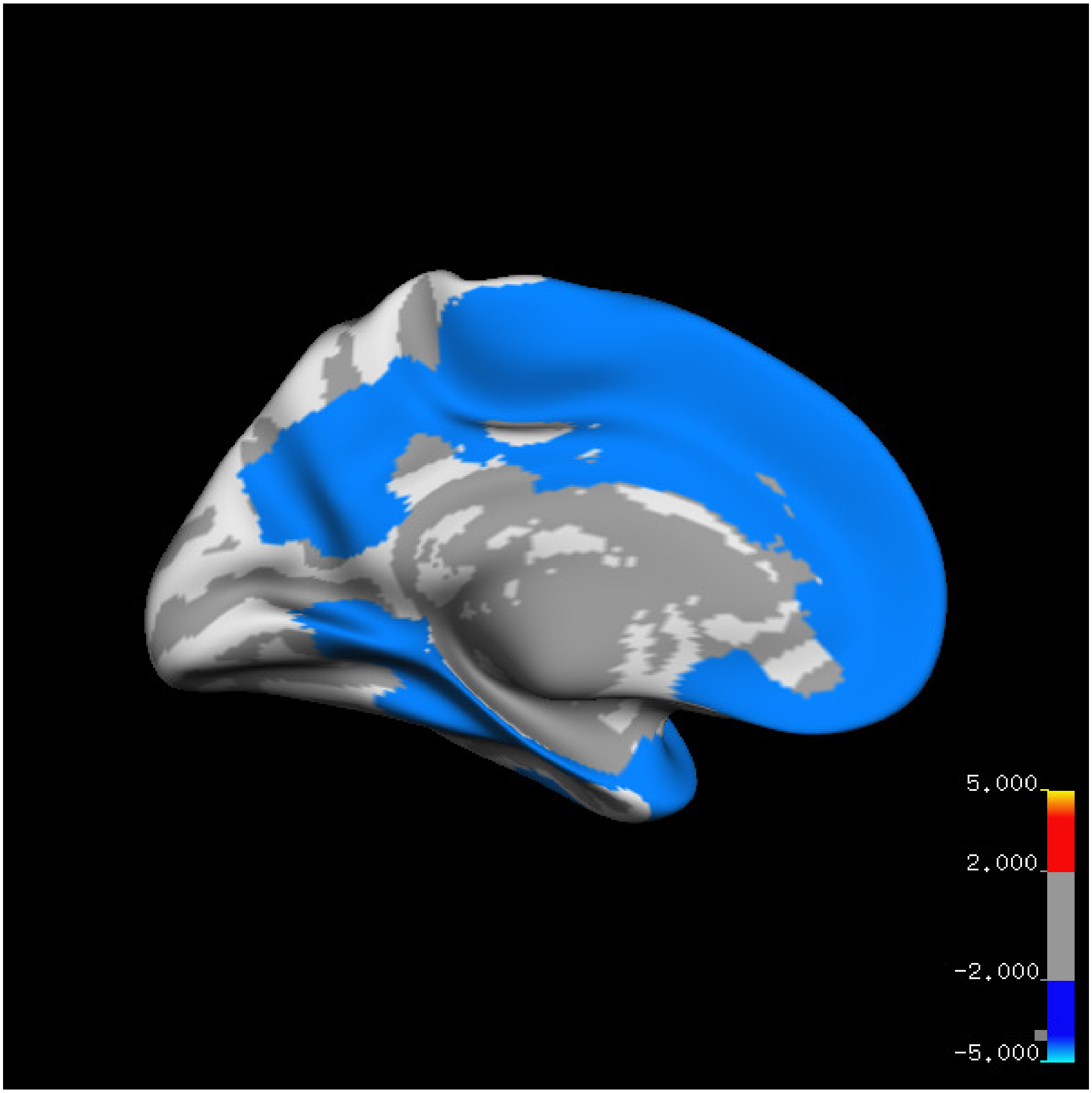

5.000
2.000
-2.000
-5.000

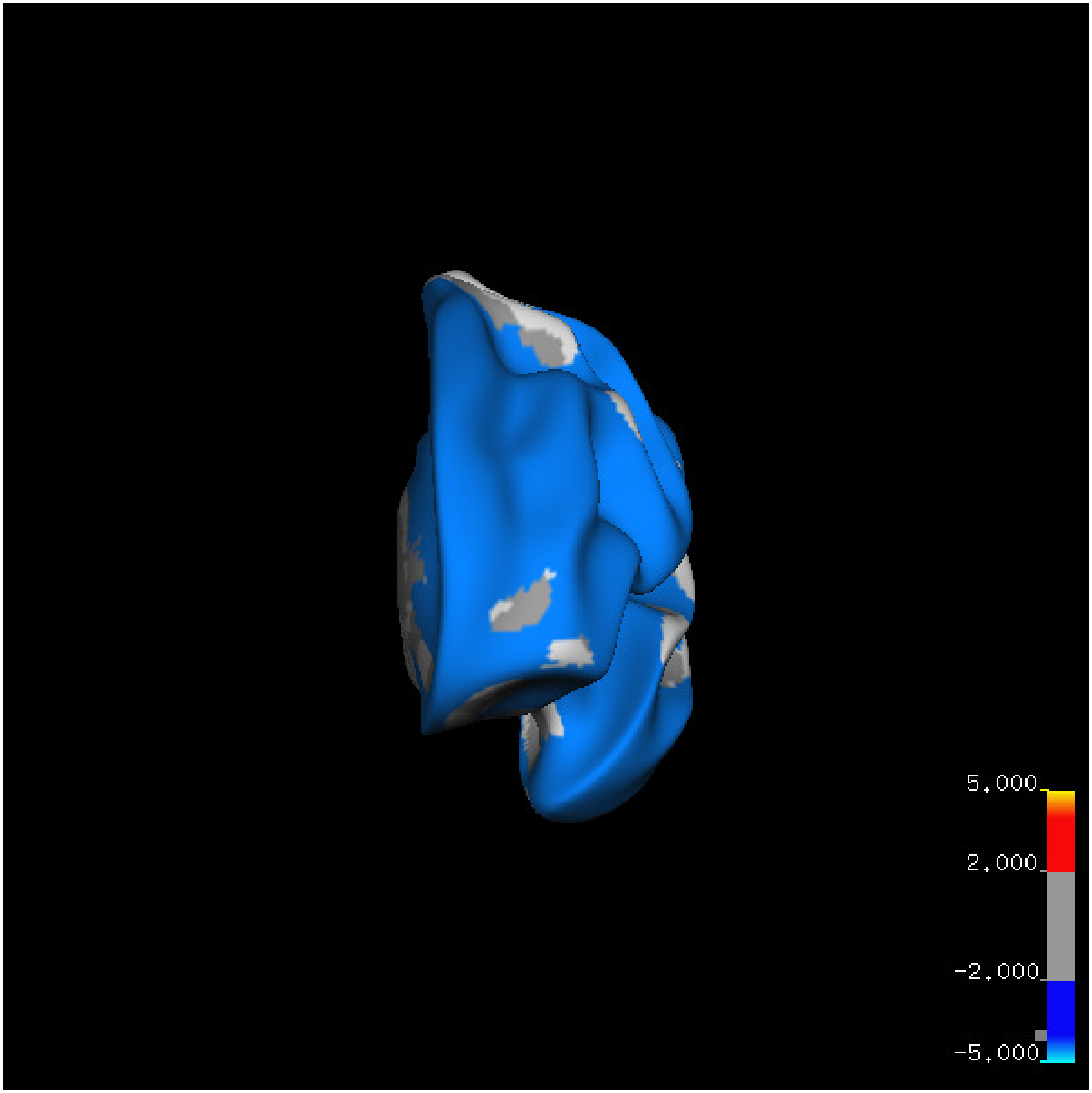
5.000
2.000
-2.000
-5.000

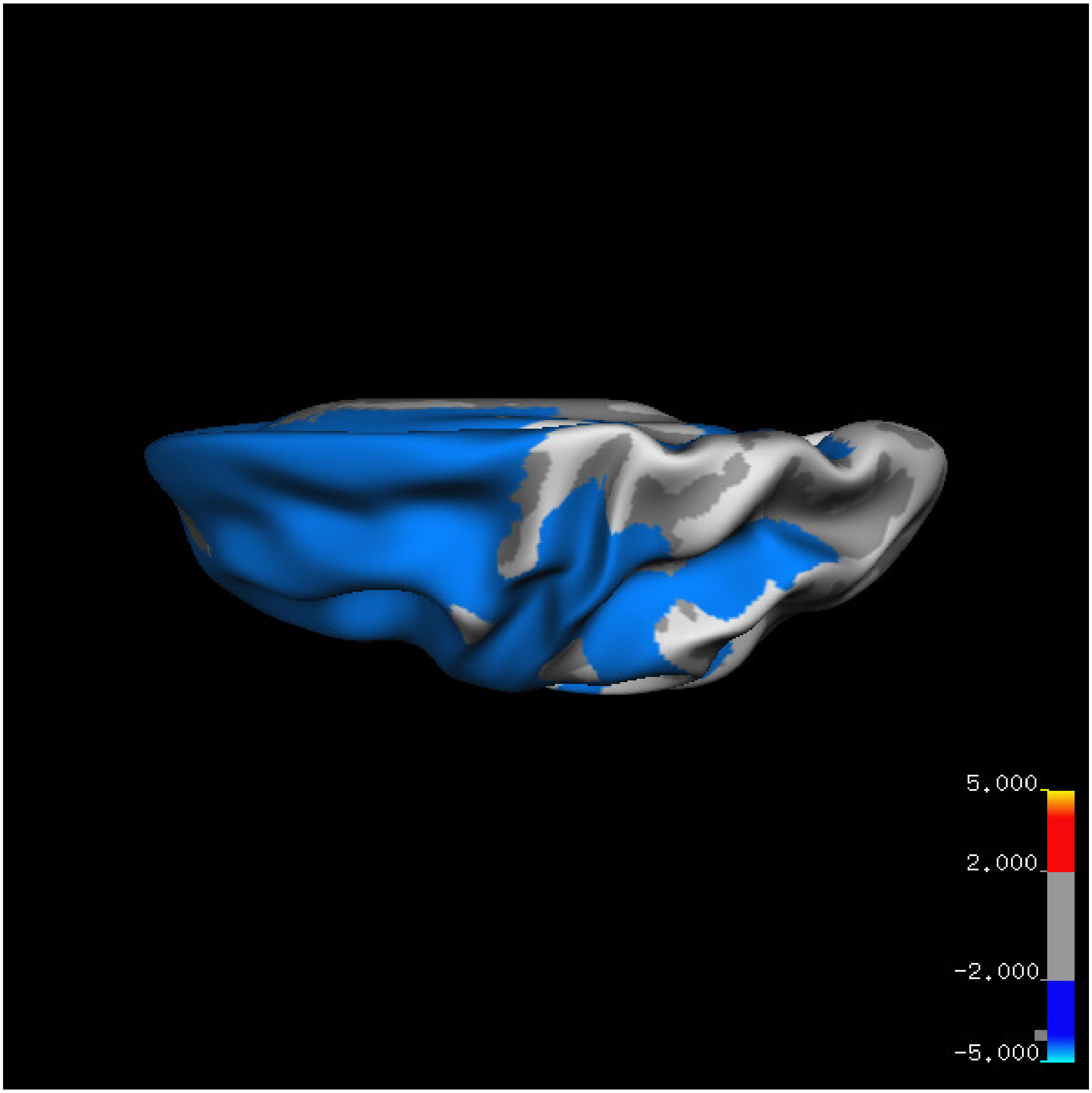
5.000
2.000
-2.000
-5.000

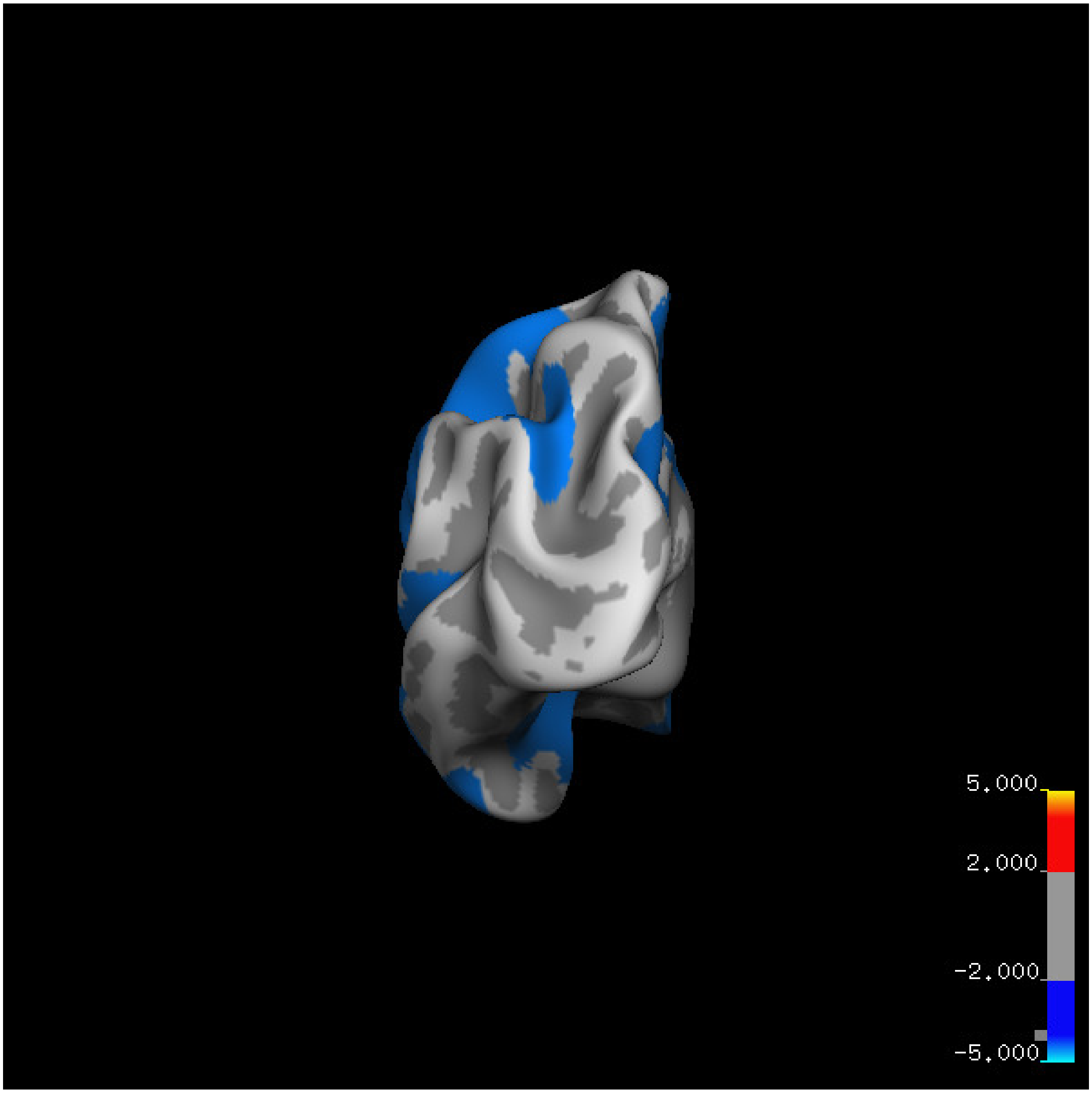
5.000
2.000
-2.000
-5.000

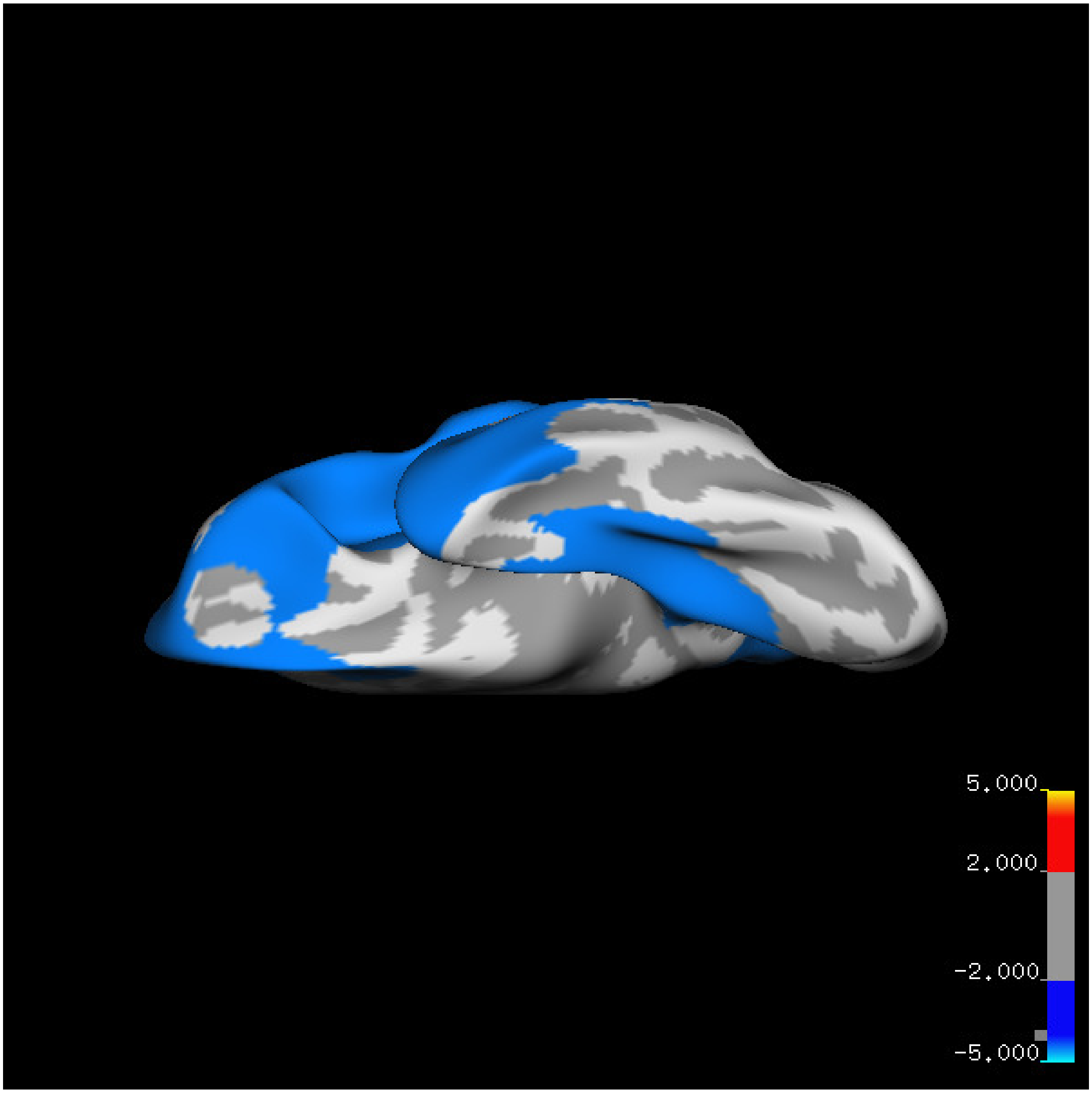
5.000
2.000
-2.000
-5.000

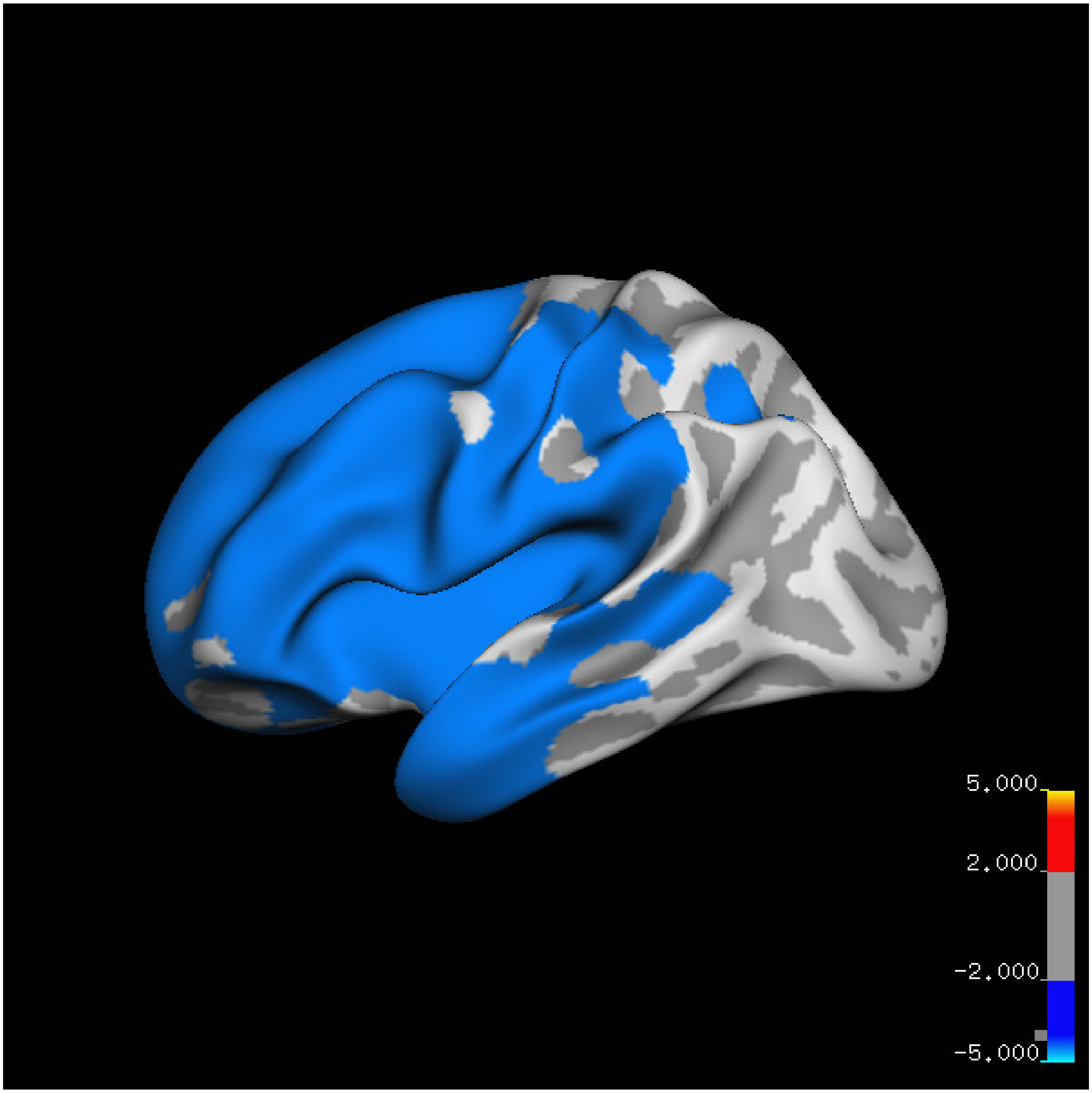
5.000
2.000
-2.000
-5.000

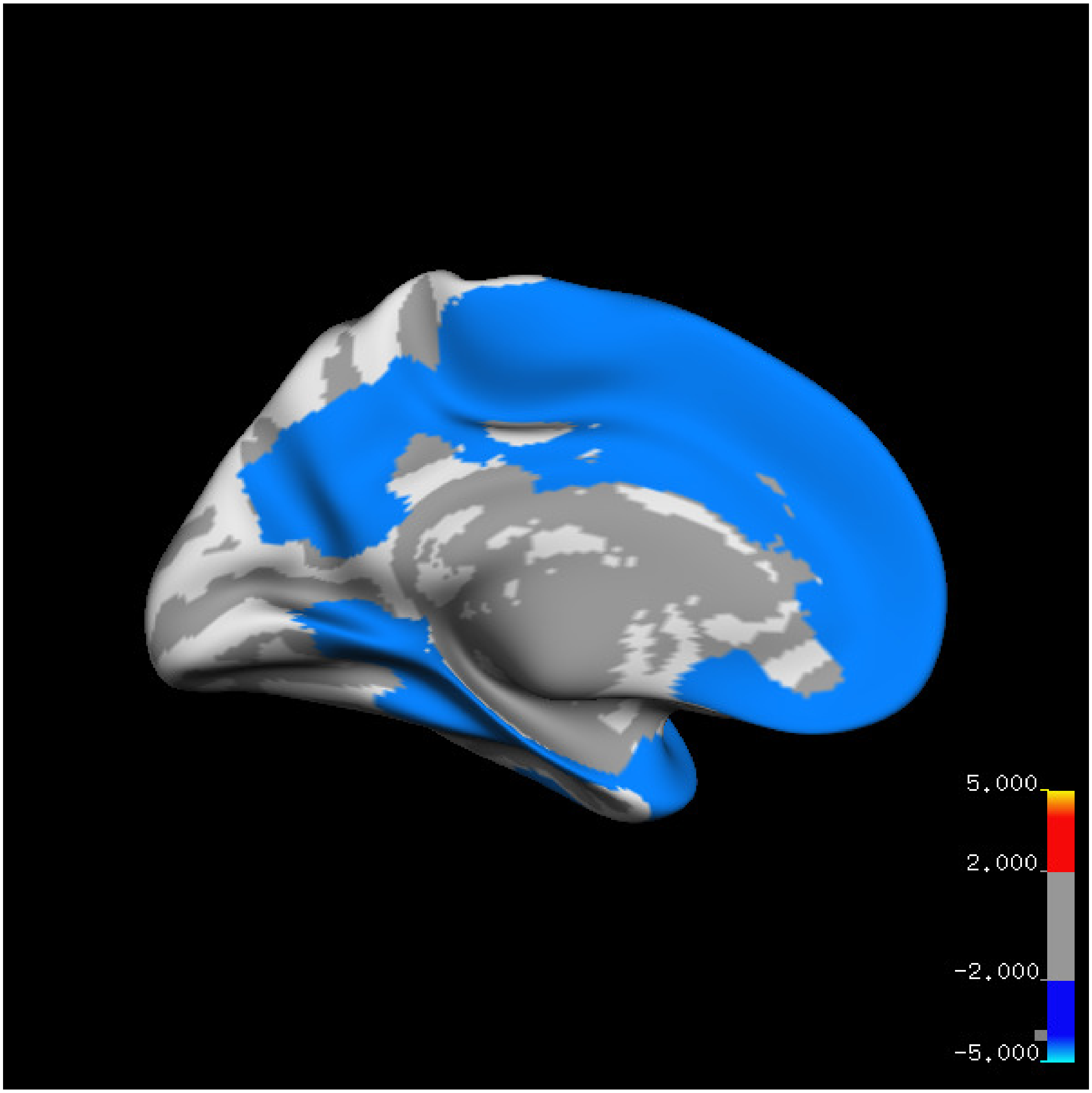
5.000
2.000
-2.000
-5.000

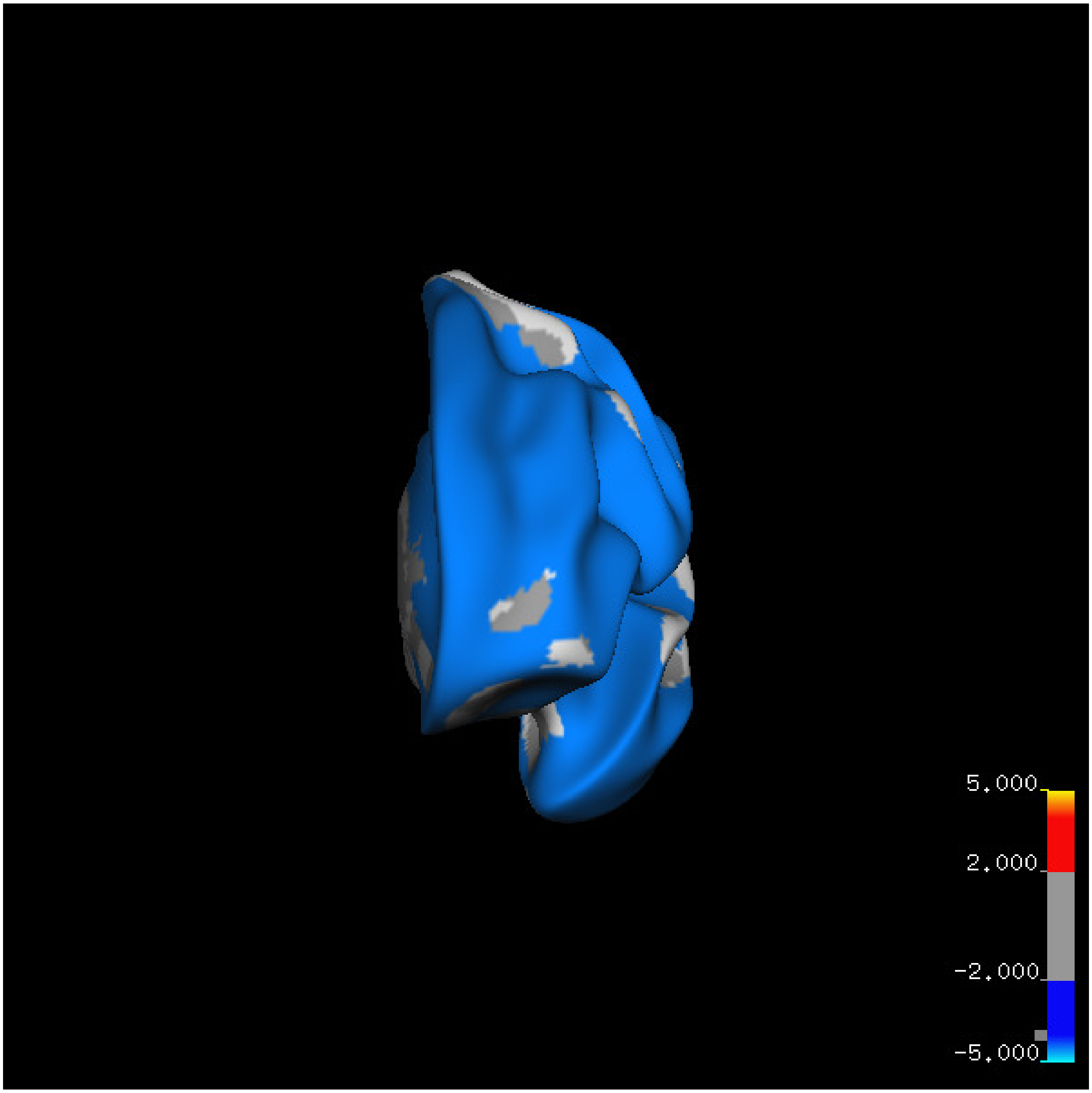
5.000
2.000
-2.000
-5.000

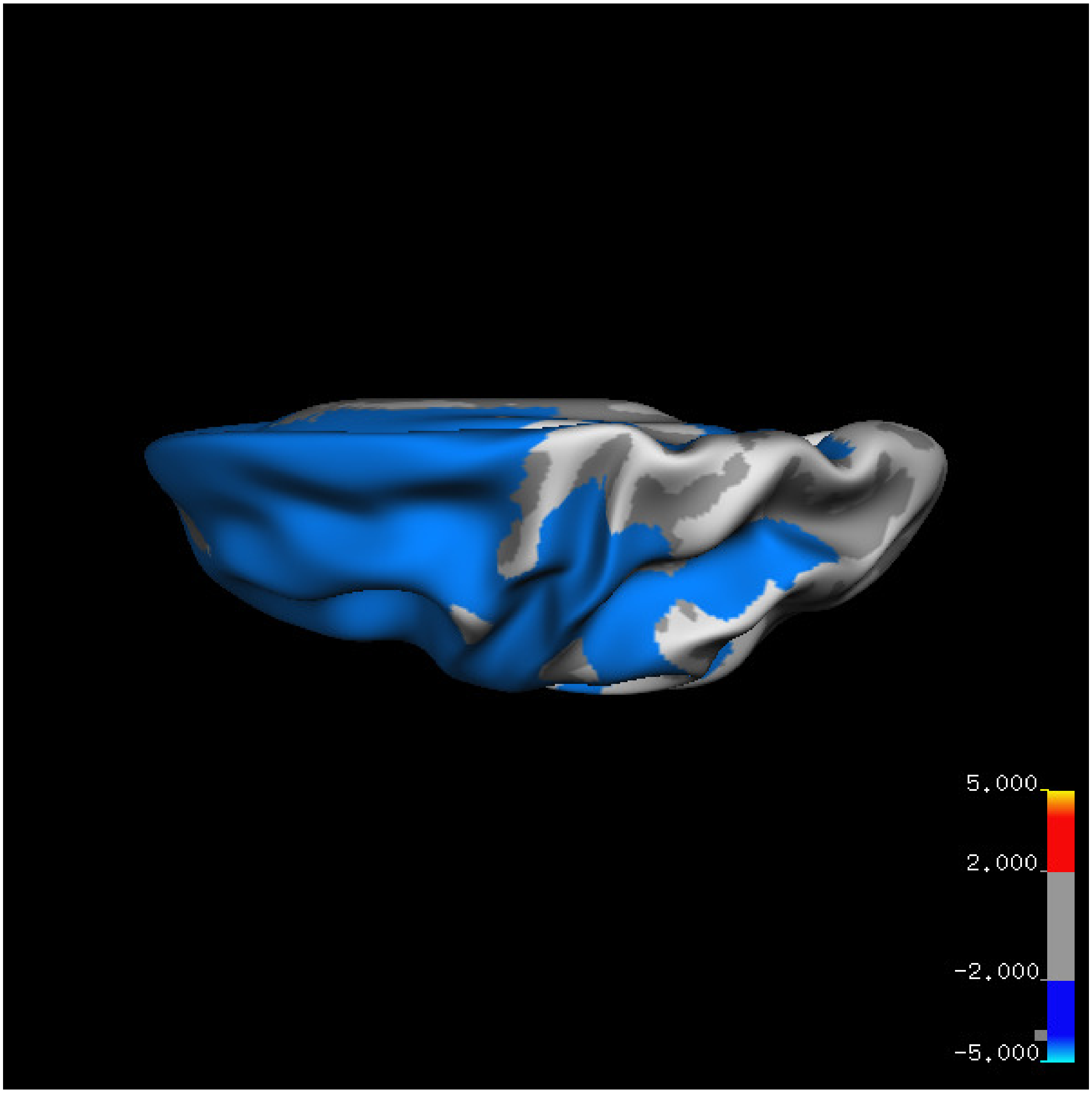


Cluster summary is shown below.

```
## # Cluster Growing Summary (mri_surfcluster)

## # $Id: mri_surfcluster.c,v 1.57.2.3 2016/11/17 18:19:42 zkaufman Exp $

## # $Id: mrisurf.c,v 1.781.2.6 2016/12/27 16:47:14 zkaufman Exp $

## # CreationTime 2021/10/04-13:10:38-GMT

## # cmdline mri_surfcluster.bin --in surface_analyses/results/cluster_comparison/fsaverage6.sm15.lh/thi

## # cwd /gpfs/projects02/p274/cluster/projects/p003-sleep_b

## # sysname  Linux
```

```
## # hostname c1-18
## # machine  x86_64
## # FixVertexAreaFlag 1
## # FixSurfClusterArea 1
## #
## # Input      surface_analyses/results/cluster_comparison/fsaverage6.sm15.lh/thickness.above60.above_
## # Frame Number      0
## # srcsubj fsaverage6
## # hemi lh
## # surface white
## # group_avg_surface_area 84969.3
## # group_avg_vtxarea_loaded 1
## # annot aparc
## # SUBJECTS_DIR /cluster/projects/p274/tools/mri/freesurfer/current/subjects
## # SearchSpace_mm2 79241.3
## # SearchSpace_vtx 37476
## # Bonferroni 0
## # Minimum Threshold 1.3
## # Maximum Threshold infinity
## # Threshold Sign    abs
## # AdjustThreshWhenOneTail 1
## # CW PValue Threshold: 0.01
## # Area Threshold    0 mm^2
## # CSD thresh  1.300000
## # CSD nreps    10000
## # CSD simtype  null-z
## # CSD contrast NA
```

```
## # CSD confint  90.000000
## # Overall max 0.929544 at vertex 39232
## # Overall min -8.75008 at vertex 20083
## # NClusters          1
## # FixMNI = 0
## #
## # ClusterNo  Max   VtxMax   Size(mm^2)  MNIX   MNIY   MNIZ    CWP    CWPLow    CWPHi   NVtxs    Wght
##    1       -8.750   20083  47716.18    -12.8   46.3   10.1  0.00010  0.00000  0.00020  23395   -7115
```

**Sleep below mean** Cluster corrected significance plots are shown below.

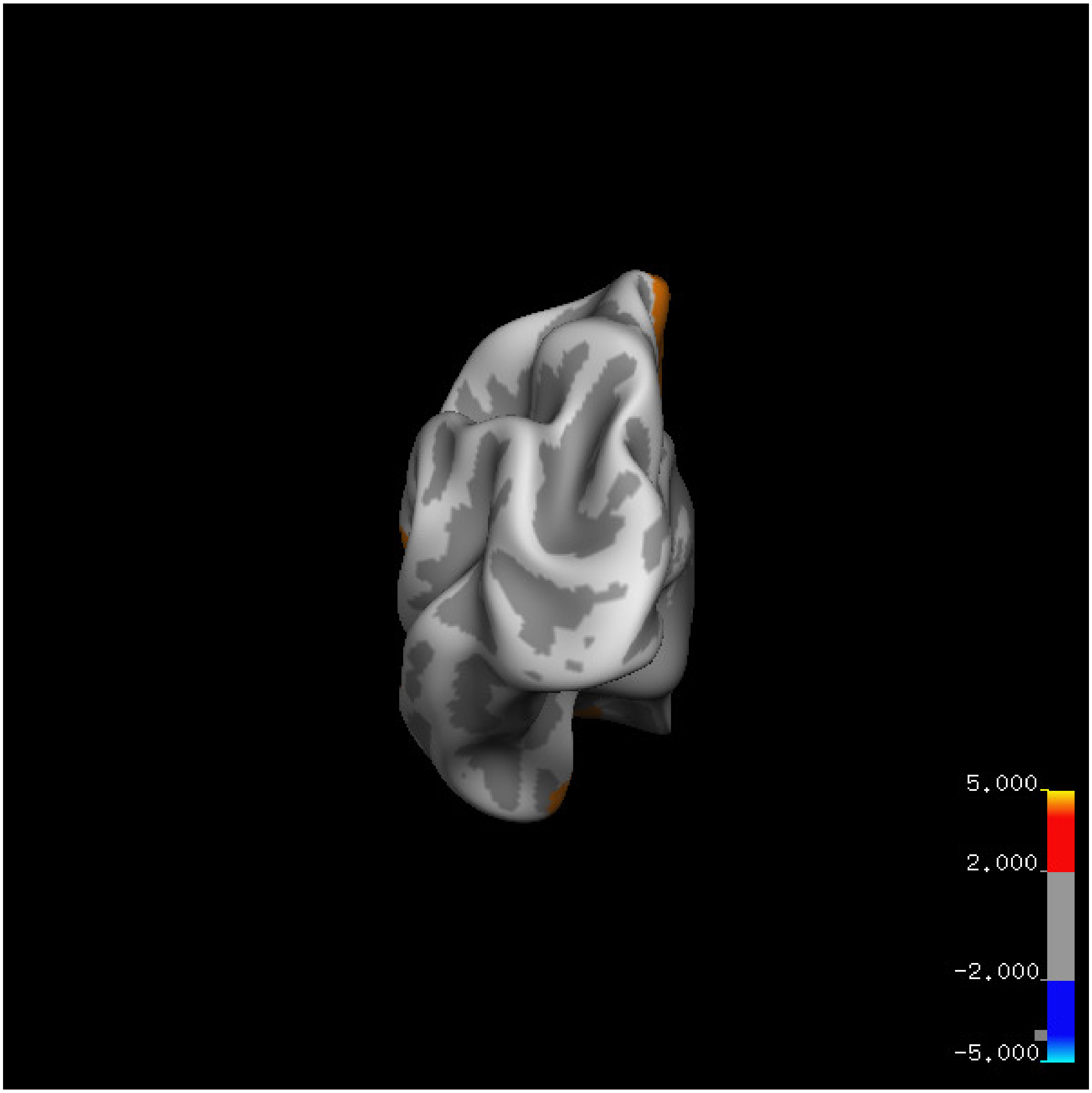

5.000
2.000
-2.000
-5.000

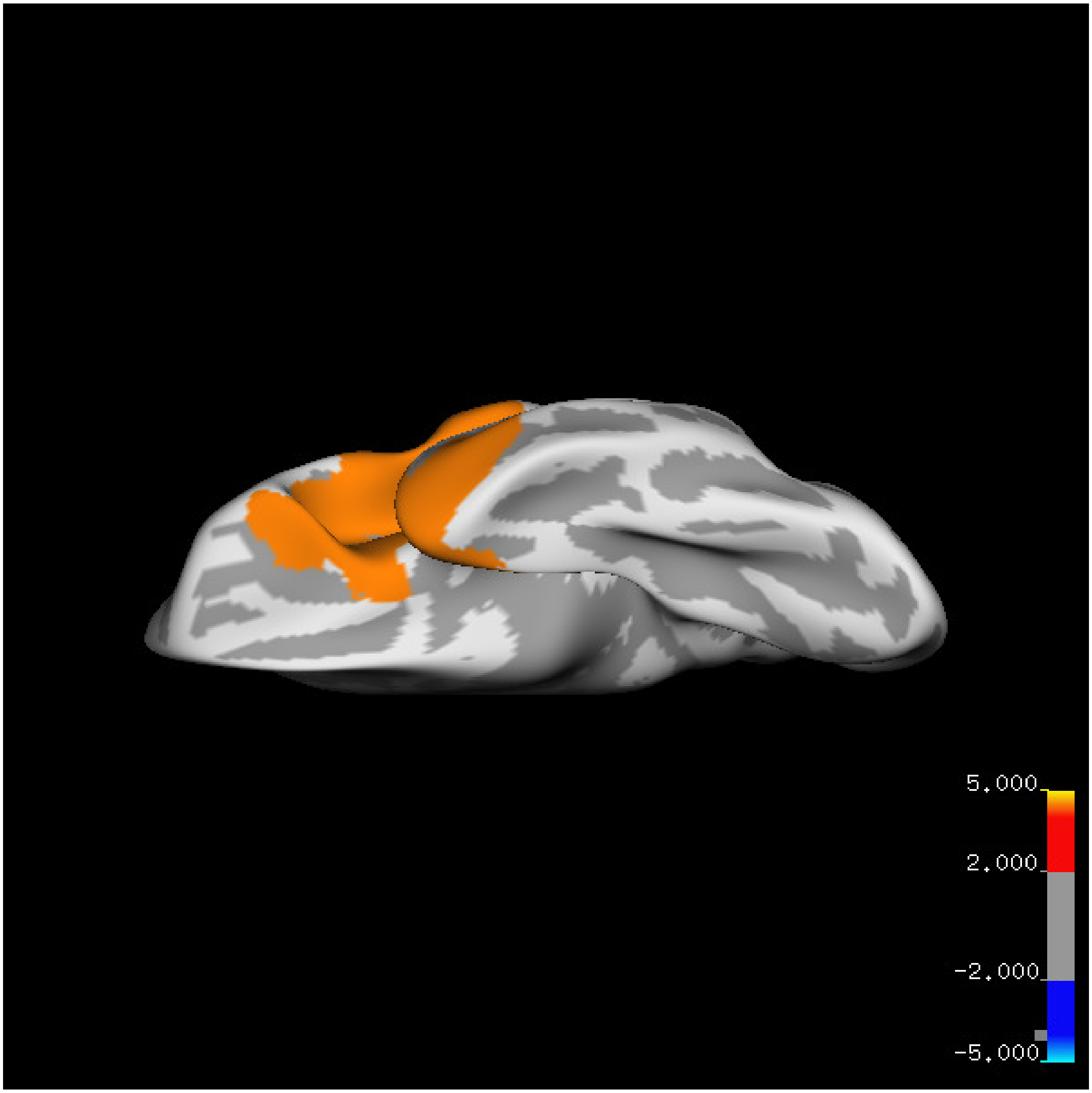
5.000
2.000
-2.000
-5.000

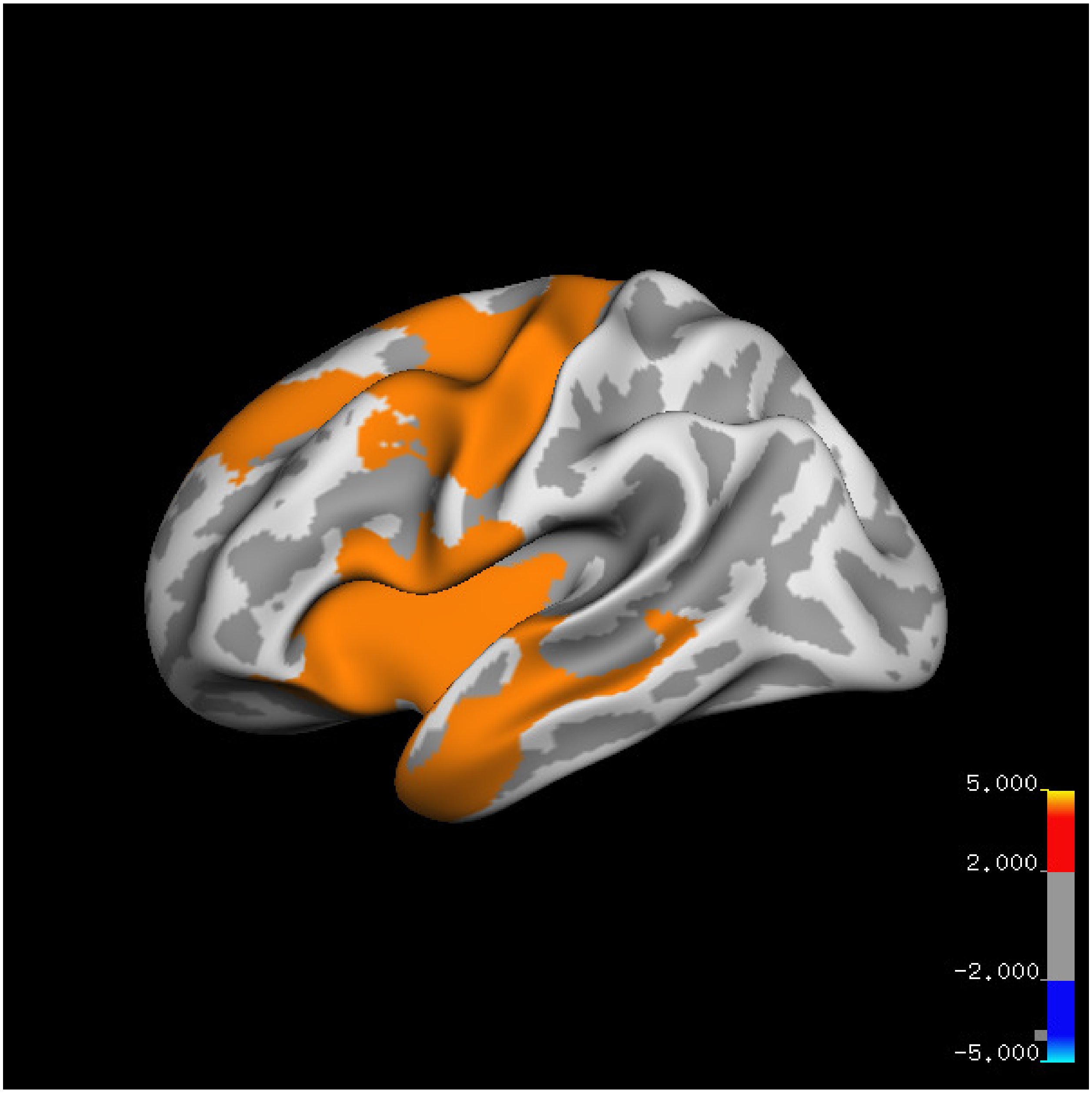
5.000
2.000
-2.000
-5.000

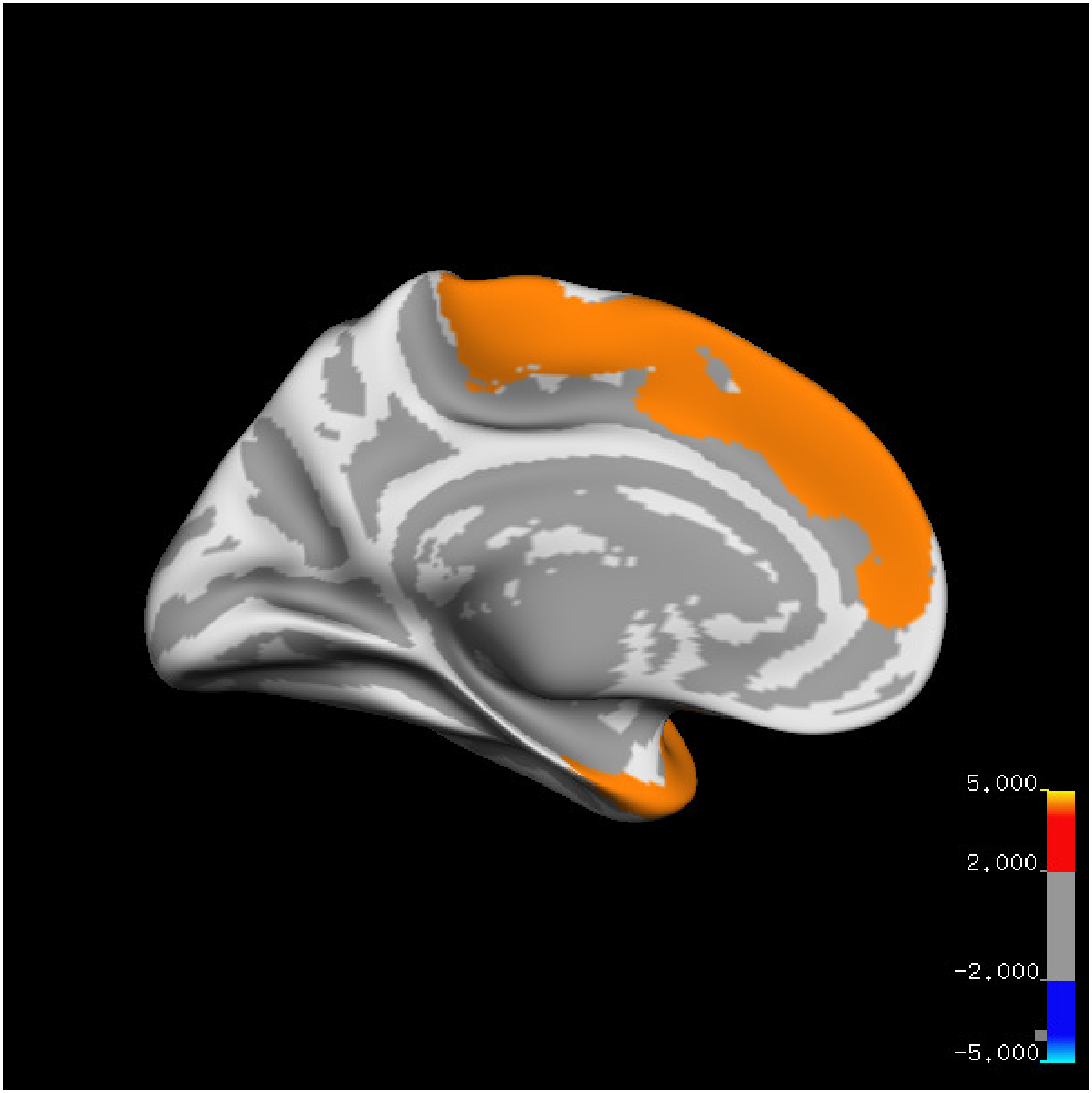
5.000
2.000
-2.000
-5.000

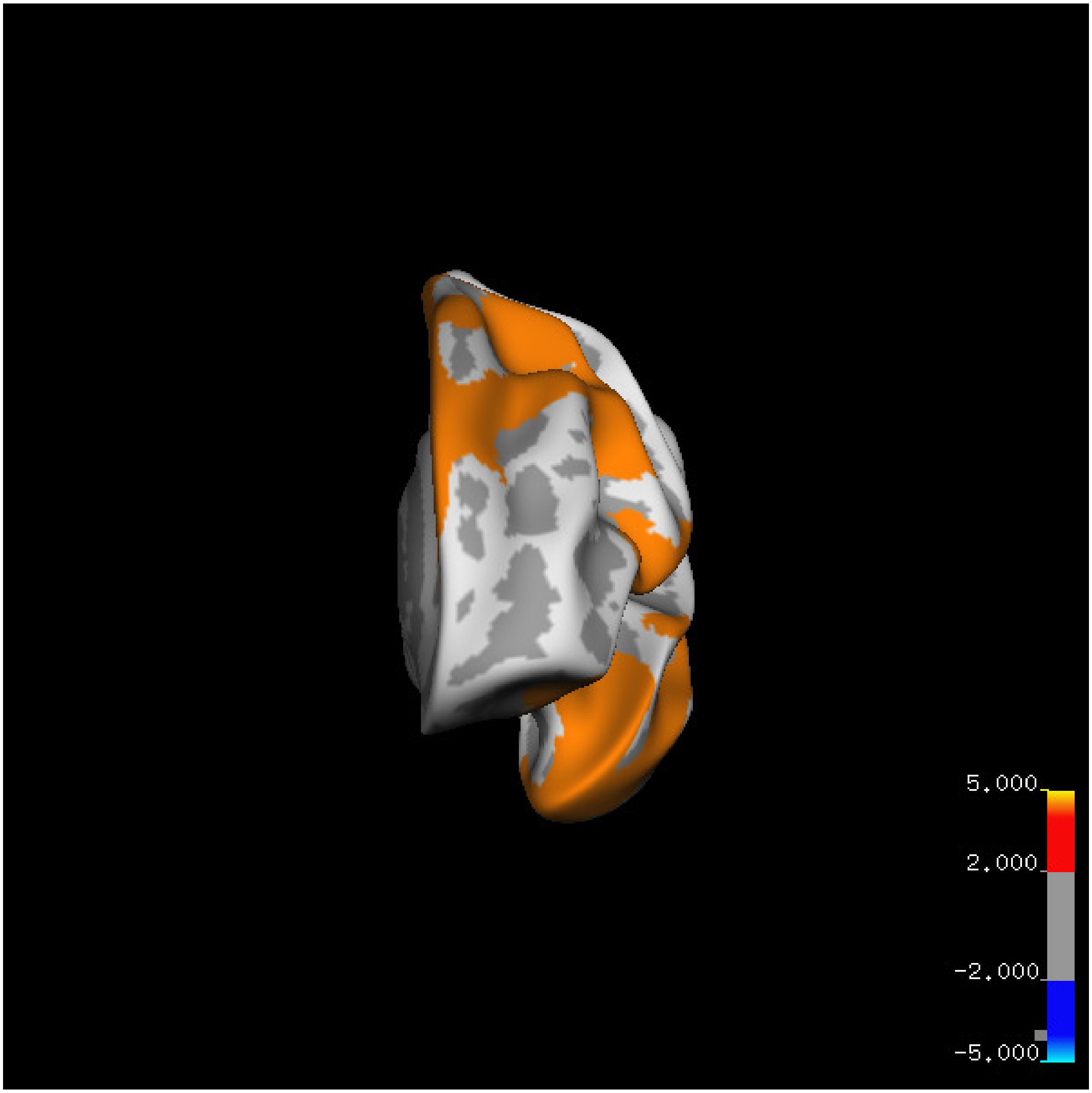
5.000
2.000
-2.000
-5.000

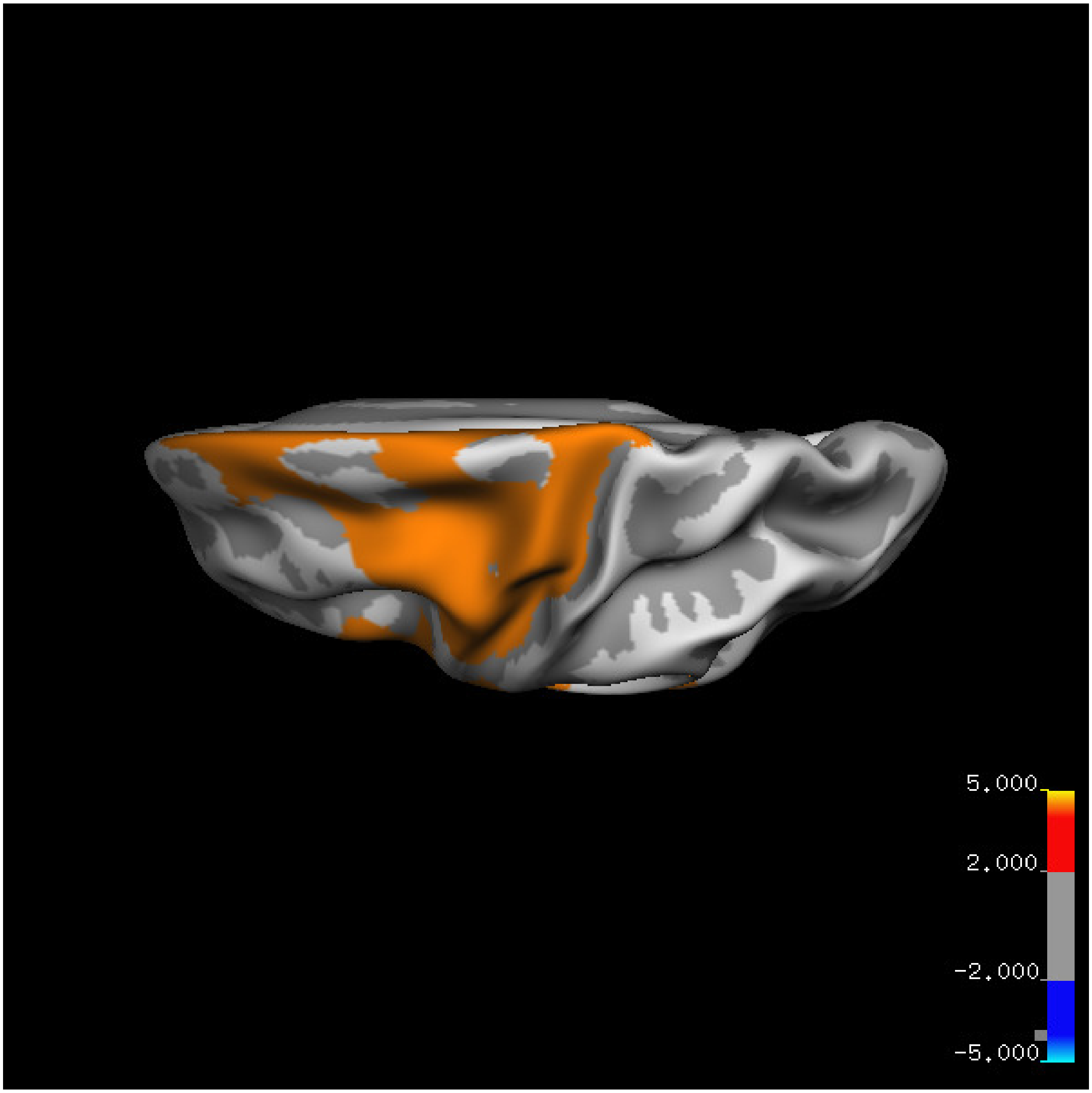
5.000
2.000
-2.000
-5.000

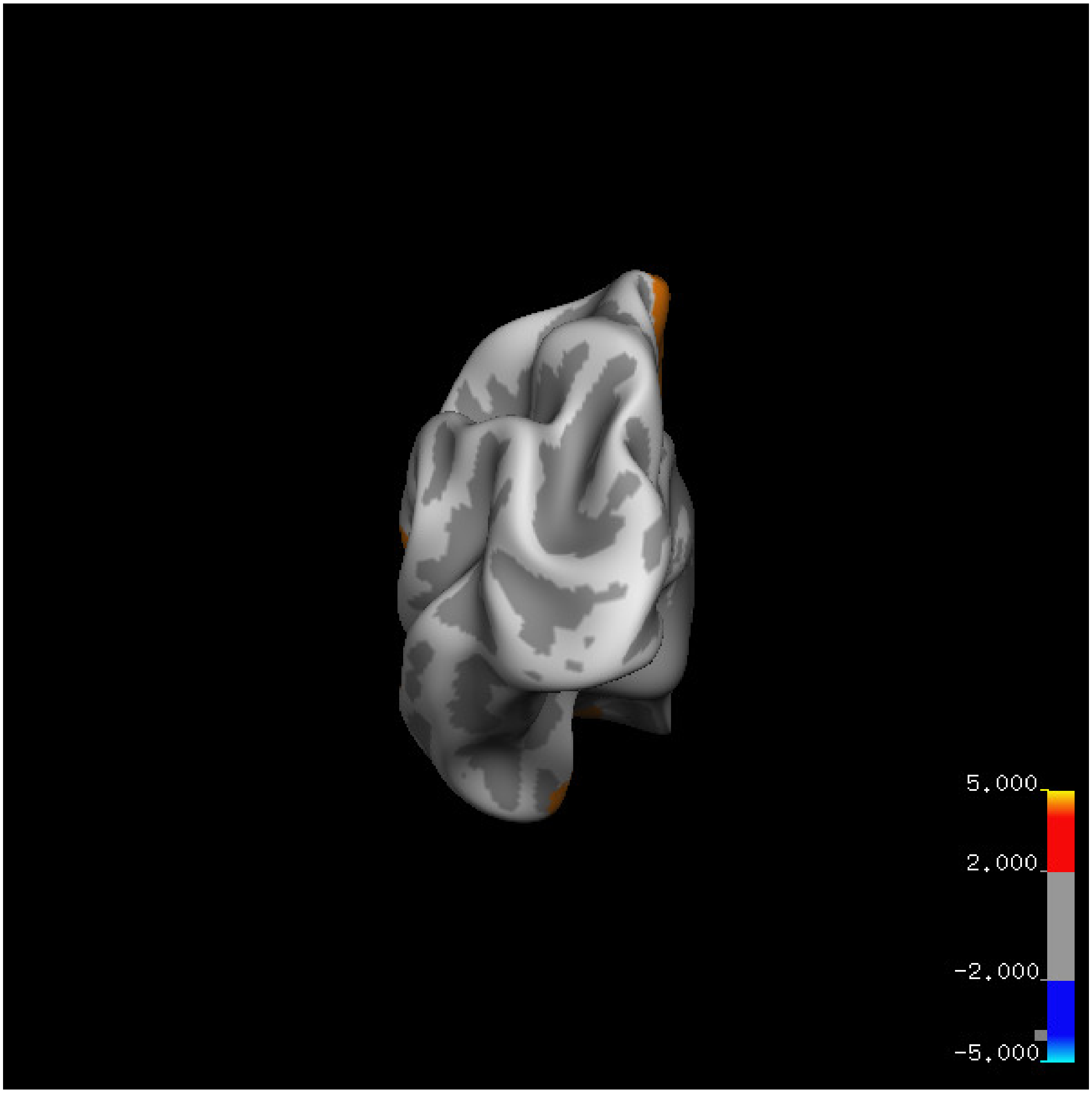
5.000
2.000
-2.000
-5.000

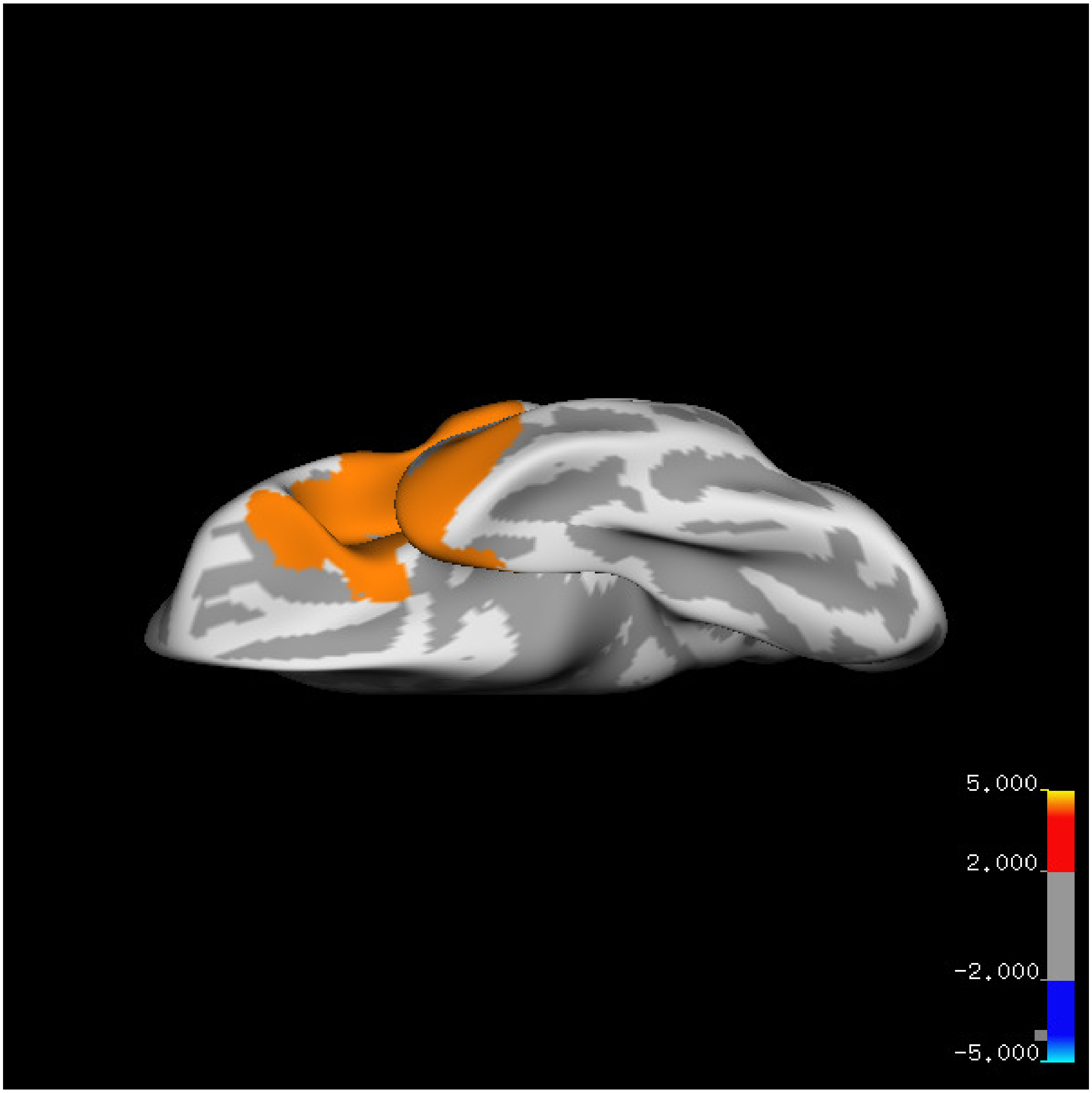
5.000
2.000
-2.000
-5.000

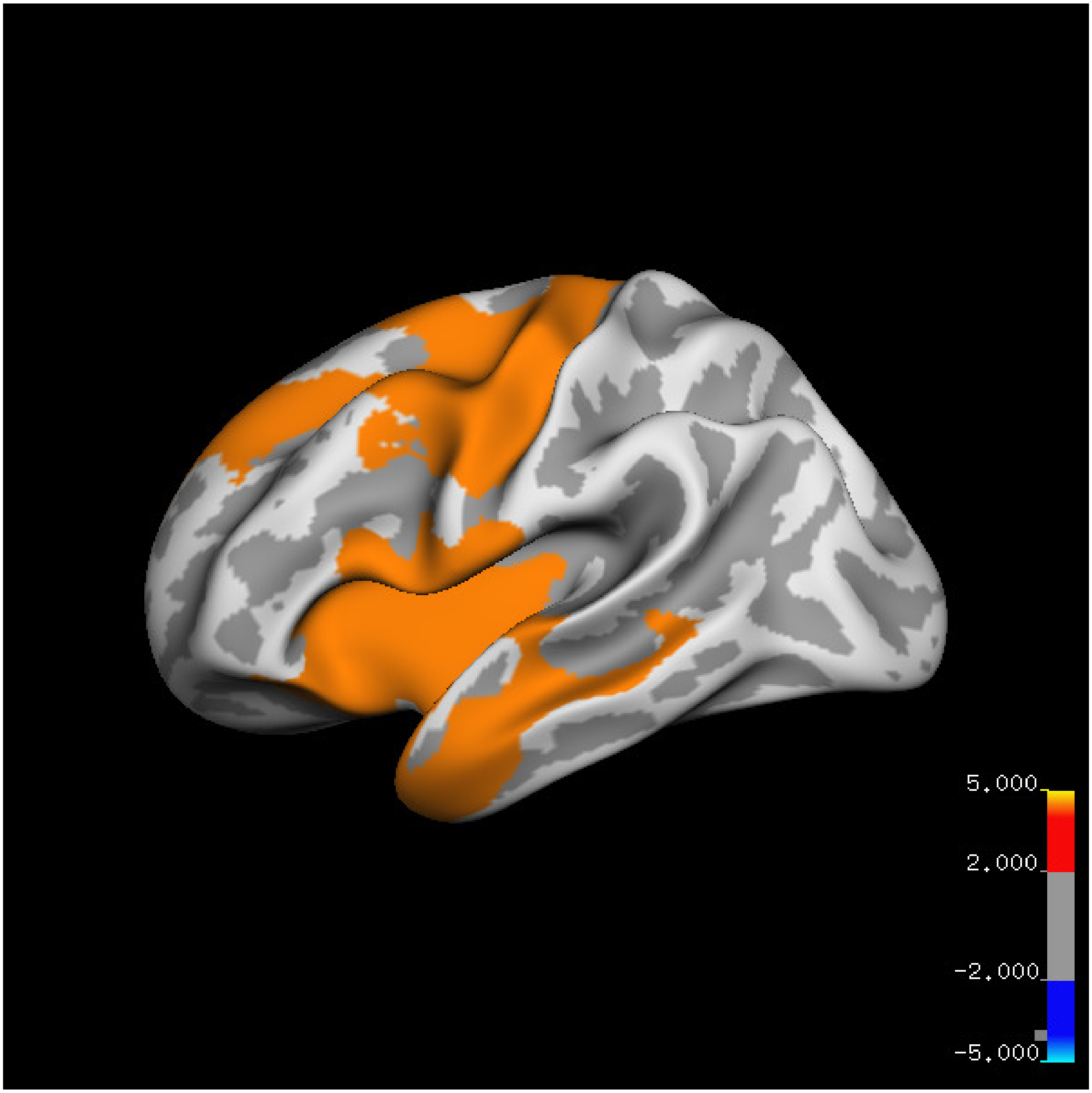
5.000
2.000
-2.000
-5.000

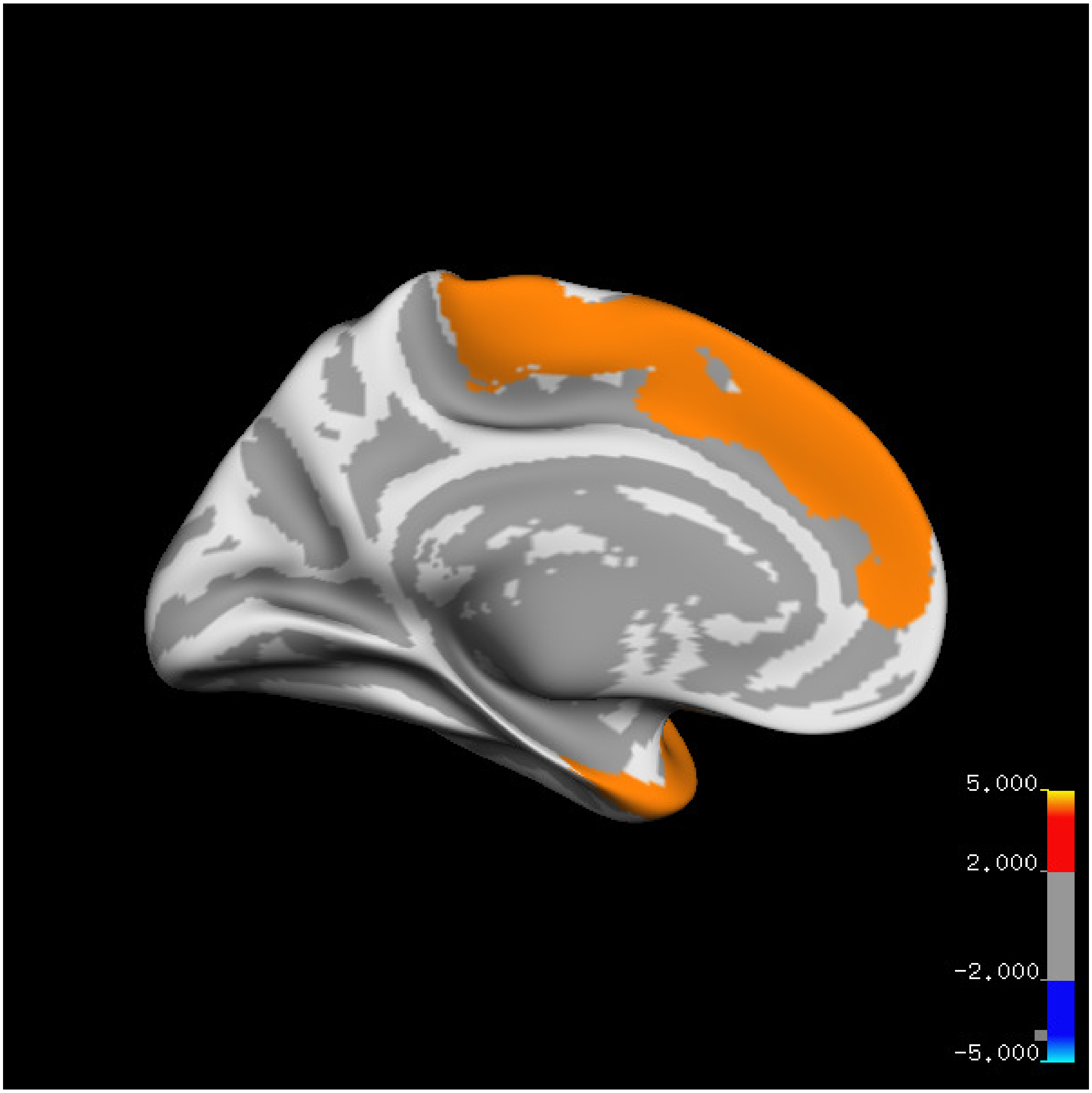
5.000
2.000
-2.000
-5.000

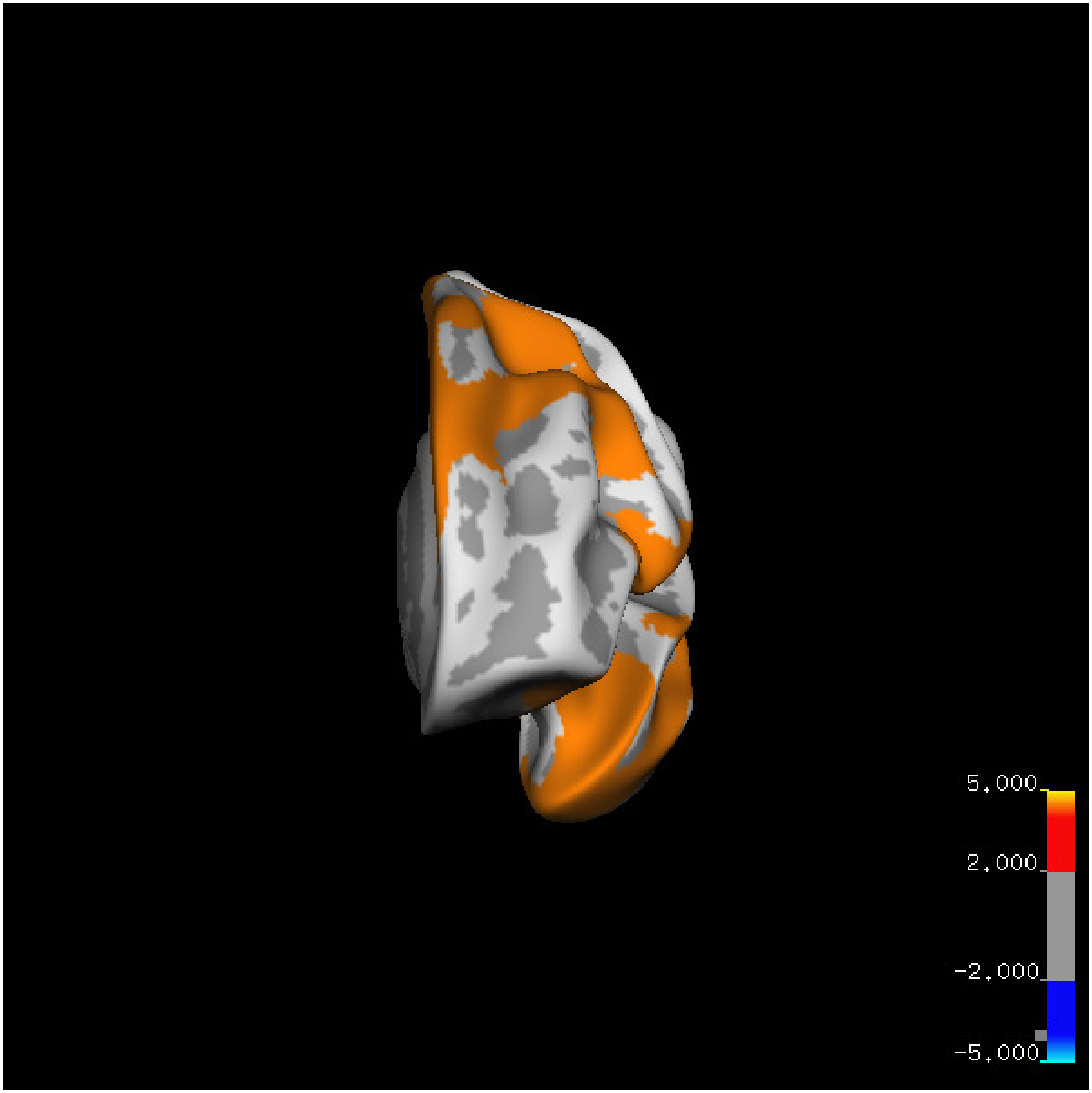

5.000
2.000
-2.000
-5.000

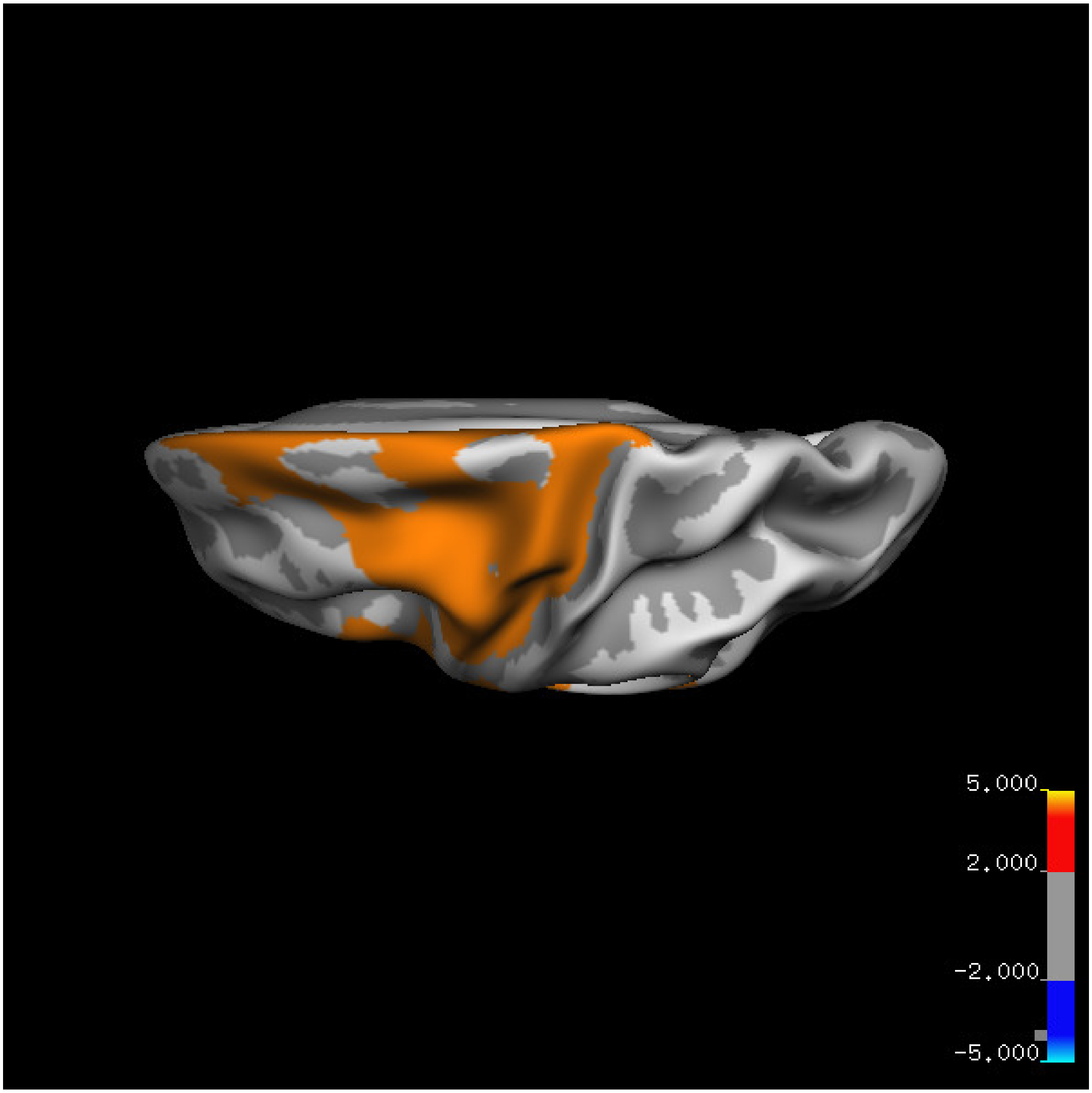


Cluster summary is shown below.

```
## # Cluster Growing Summary (mri_surfcluster)

## # $Id: mri_surfcluster.c,v 1.57.2.3 2016/11/17 18:19:42 zkaufman Exp $

## # $Id: mrisurf.c,v 1.781.2.6 2016/12/27 16:47:14 zkaufman Exp $

## # CreationTime 2021/10/04-13:56:17-GMT

## # cmdline mri_surfcluster.bin --in surface_analyses/results/cluster_comparison/fsaverage6.sm15.lh/th

## # cwd /gpfs/projects02/p274/cluster/projects/p003-sleep_b

## # sysname  Linux
```

```
## # hostname c1-18
## # machine  x86_64
## # FixVertexAreaFlag 1
## # FixSurfClusterArea 1
## #
## # Input      surface_analyses/results/cluster_comparison/fsaverage6.sm15.lh/thickness.above60.below_
## # Frame Number      0
## # srcsubj fsaverage6
## # hemi lh
## # surface white
## # group_avg_surface_area 84969.3
## # group_avg_vtxarea_loaded 1
## # annot aparc
## # SUBJECTS_DIR /cluster/projects/p274/tools/mri/freesurfer/current/subjects
## # SearchSpace_mm2 79241.3
## # SearchSpace_vtx 37476
## # Bonferroni 0
## # Minimum Threshold 1.3
## # Maximum Threshold infinity
## # Threshold Sign    abs
## # AdjustThreshWhenOneTail 1
## # CW PValue Threshold: 0.01
## # Area Threshold    0 mm^2
## # CSD thresh  1.300000
## # CSD nreps    10000
## # CSD simtype  null-z
## # CSD contrast NA
```

```
## # CSD confint  90.000000

## # Overall max 9.75372 at vertex 24708

## # Overall min -2.92715 at vertex 14450

## # NClusters          2

## # FixMNI = 0

## #

## # ClusterNo  Max   VtxMax   Size(mm^2)  MNIX   MNIY   MNIZ    CWP    CWPLow    CWPHi   NVtxs    Wght

##    1        5.857   13451  10371.63    -36.7  -22.4   54.6  0.00010  0.00000  0.00020   5143    1111

##    2        9.754   24708   8867.18    -33.0   11.1  -10.3  0.00010  0.00000  0.00020   4615    1226
```

#### Right hemisphere

##### Age 20-60

**Sleep above mean** Cluster corrected significance plots are shown below.

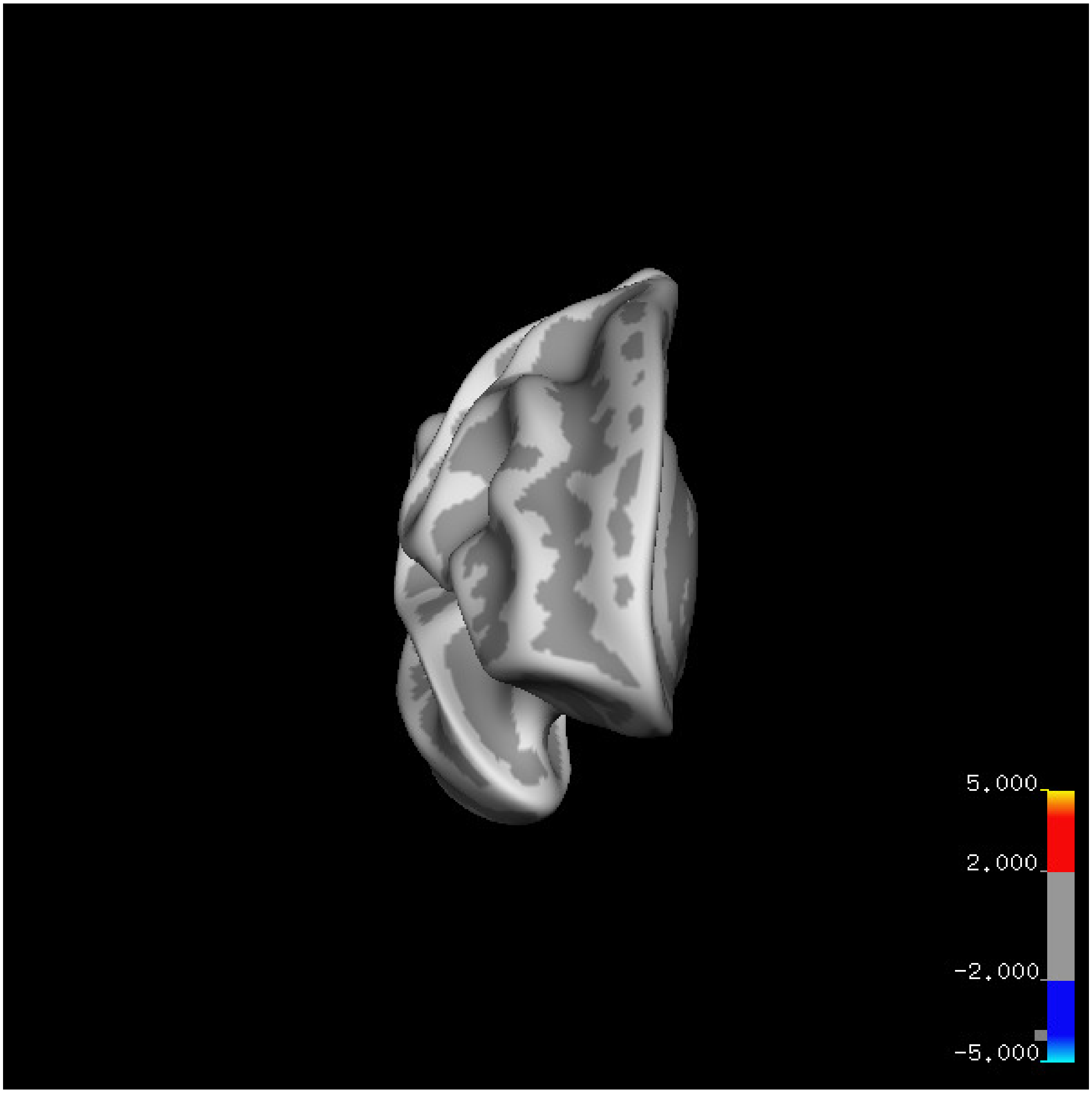
5.000
2.000
-2.000
-5.000

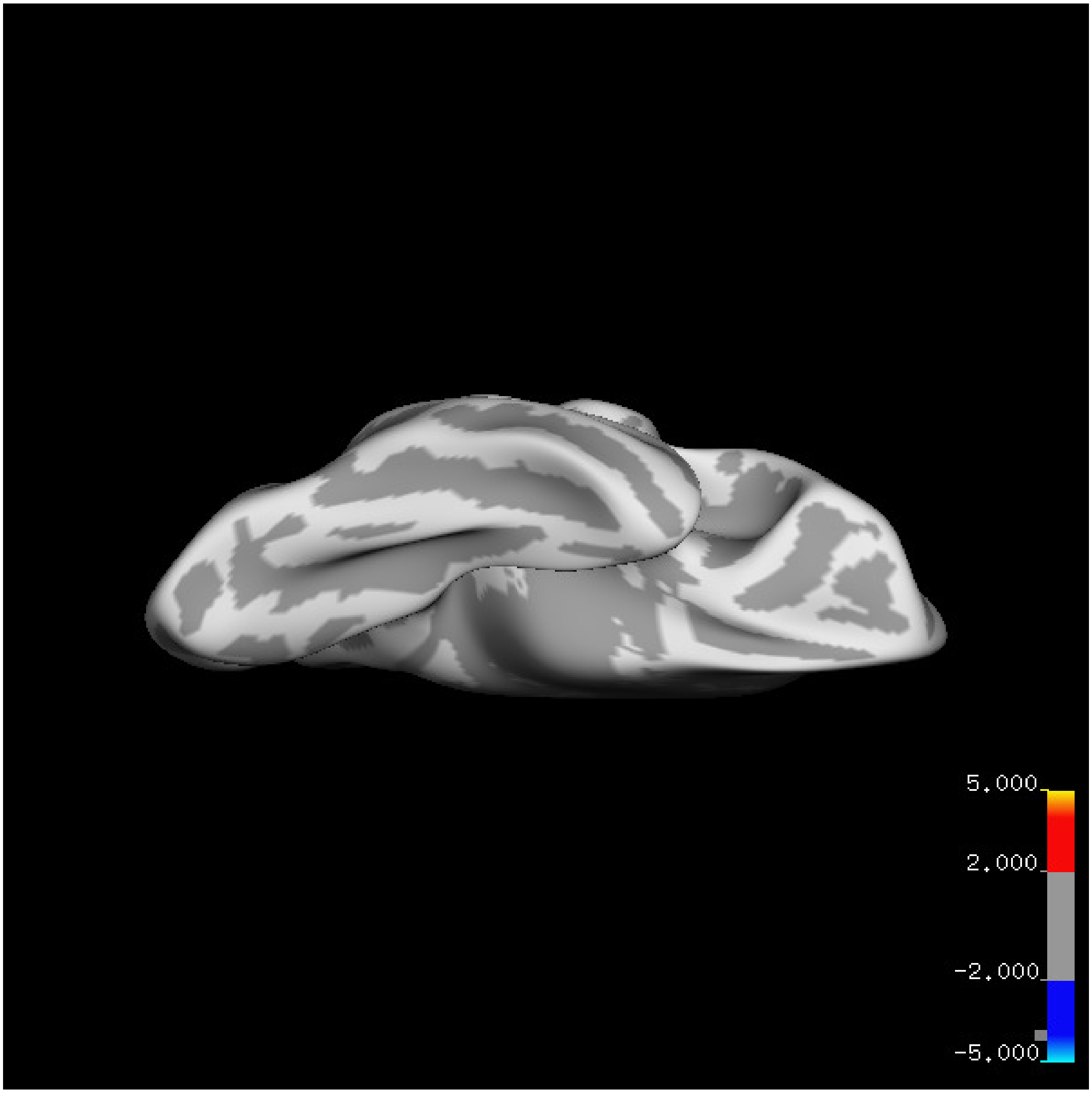
5.000
2.000
-2.000
-5.000

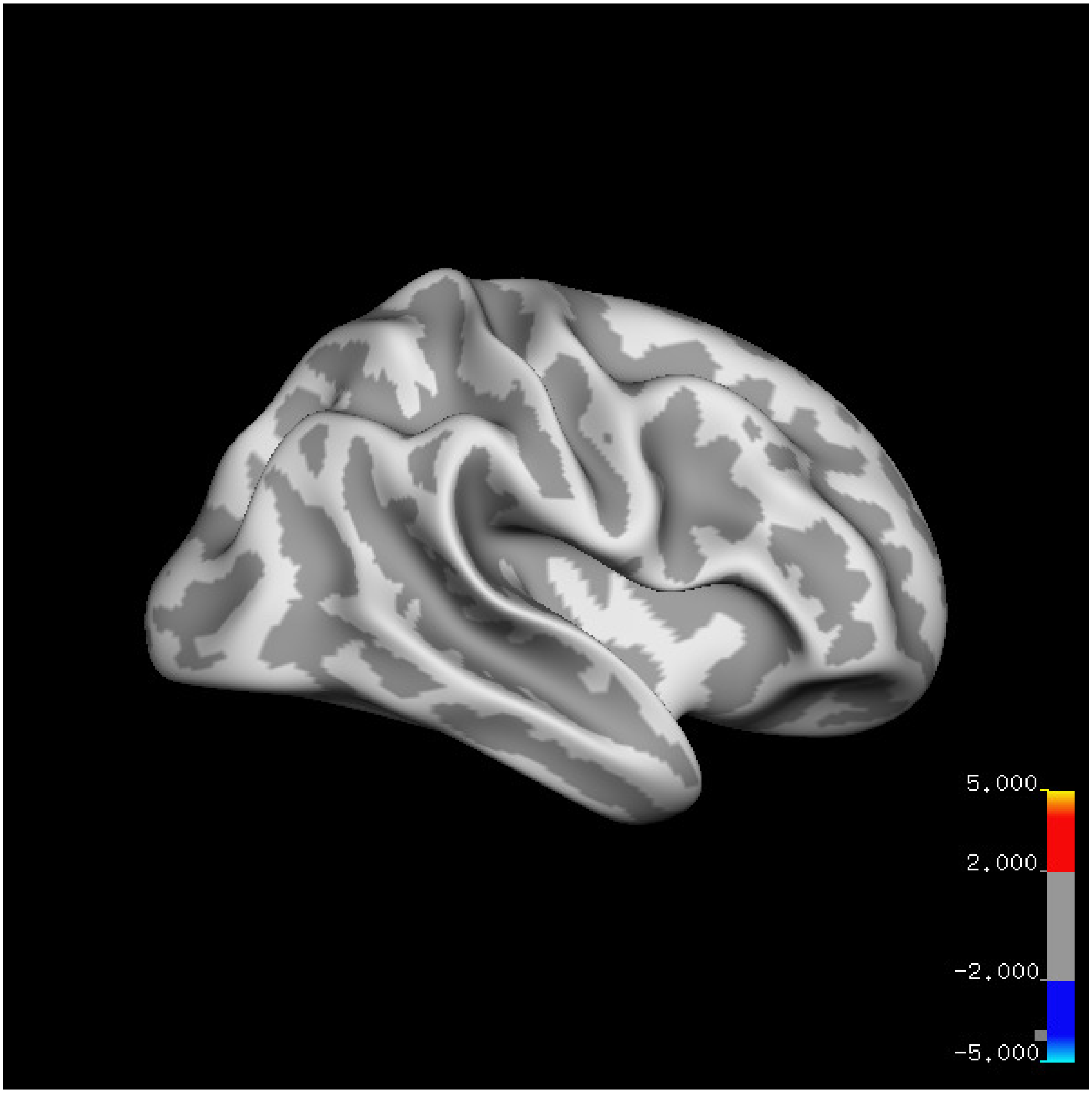
5.000
2.000
-2.000
-5.000

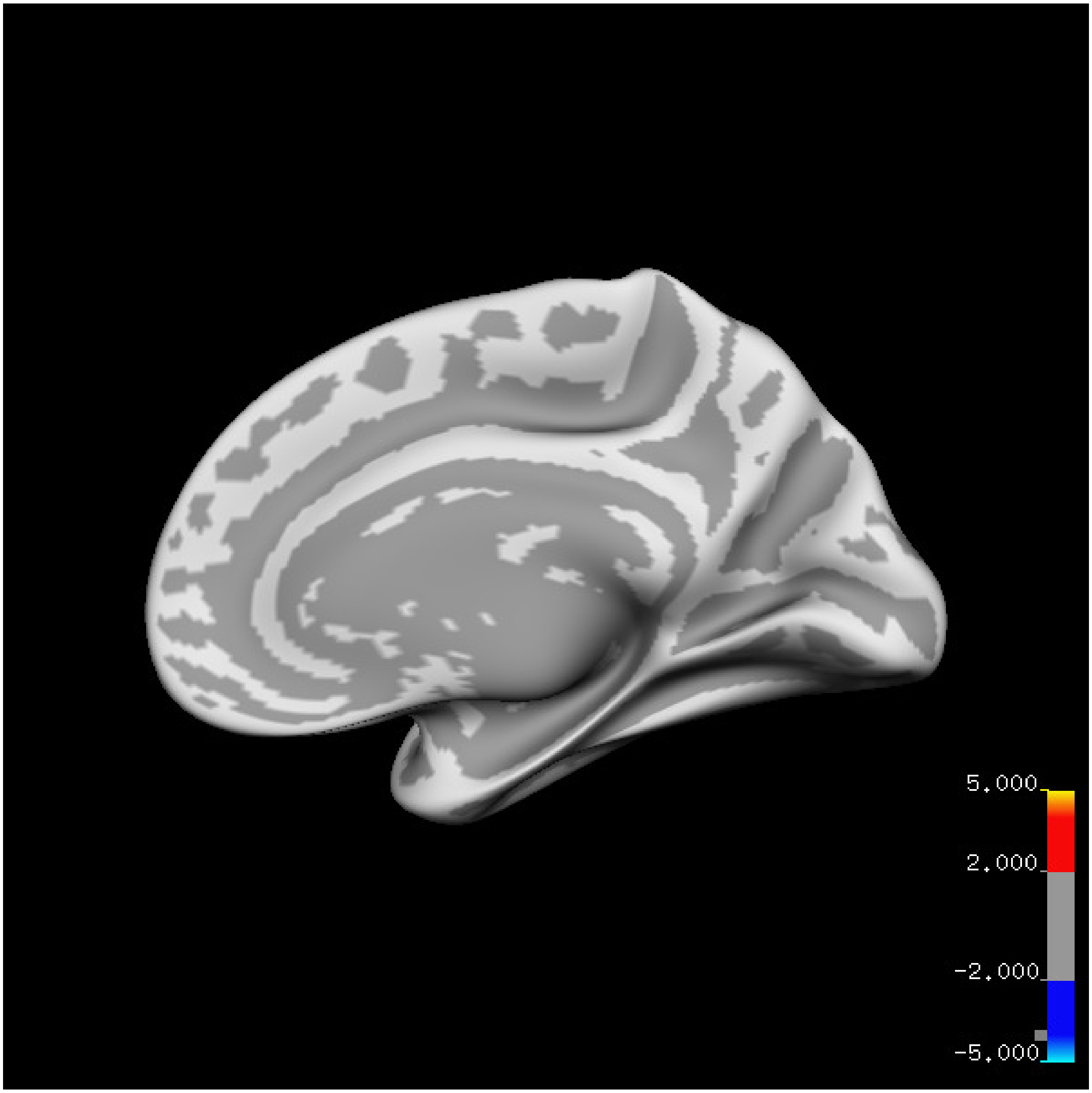
5.000
2.000
-2.000
-5.000

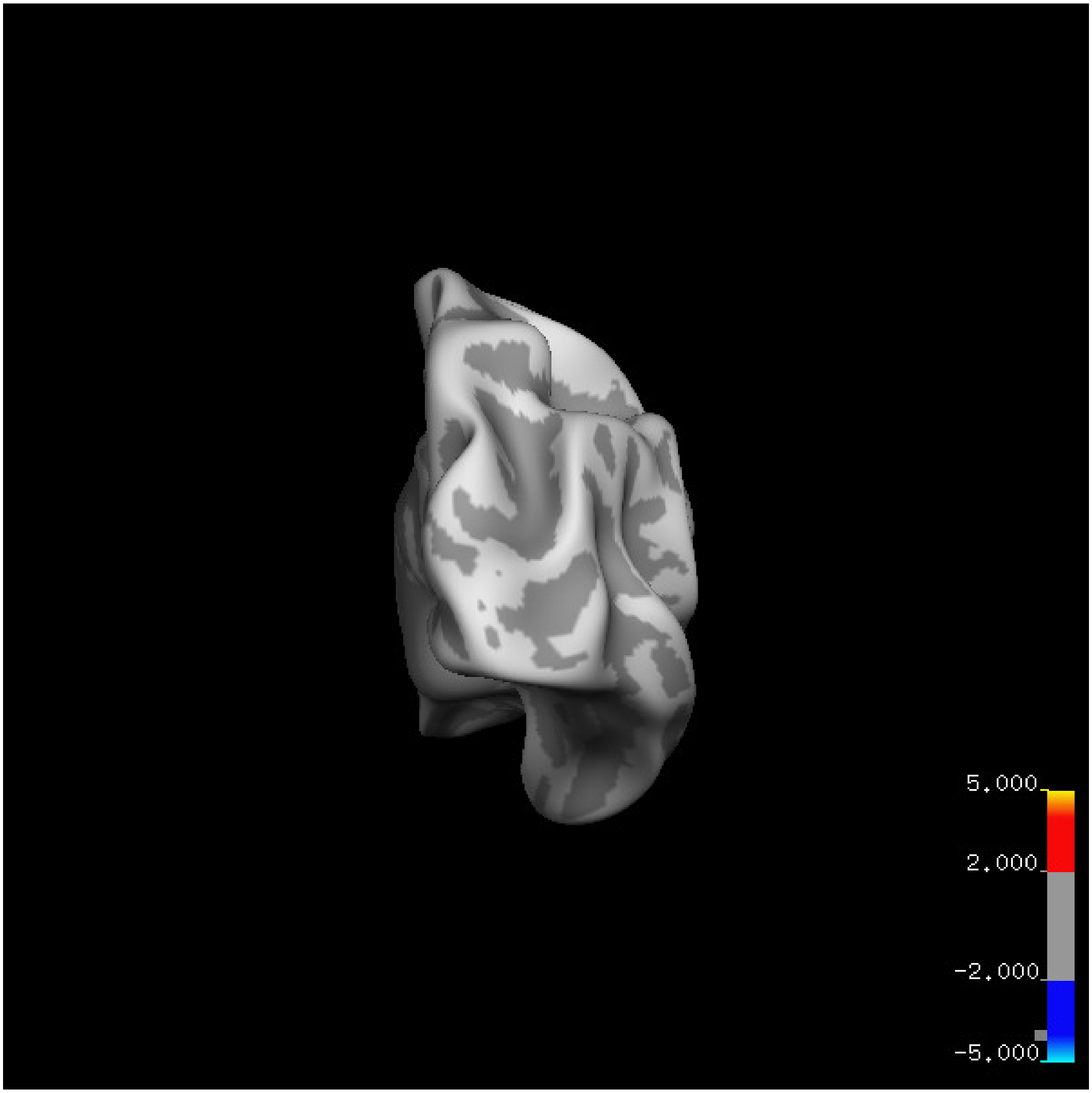
5.000
2.000
-2.000
-5.000

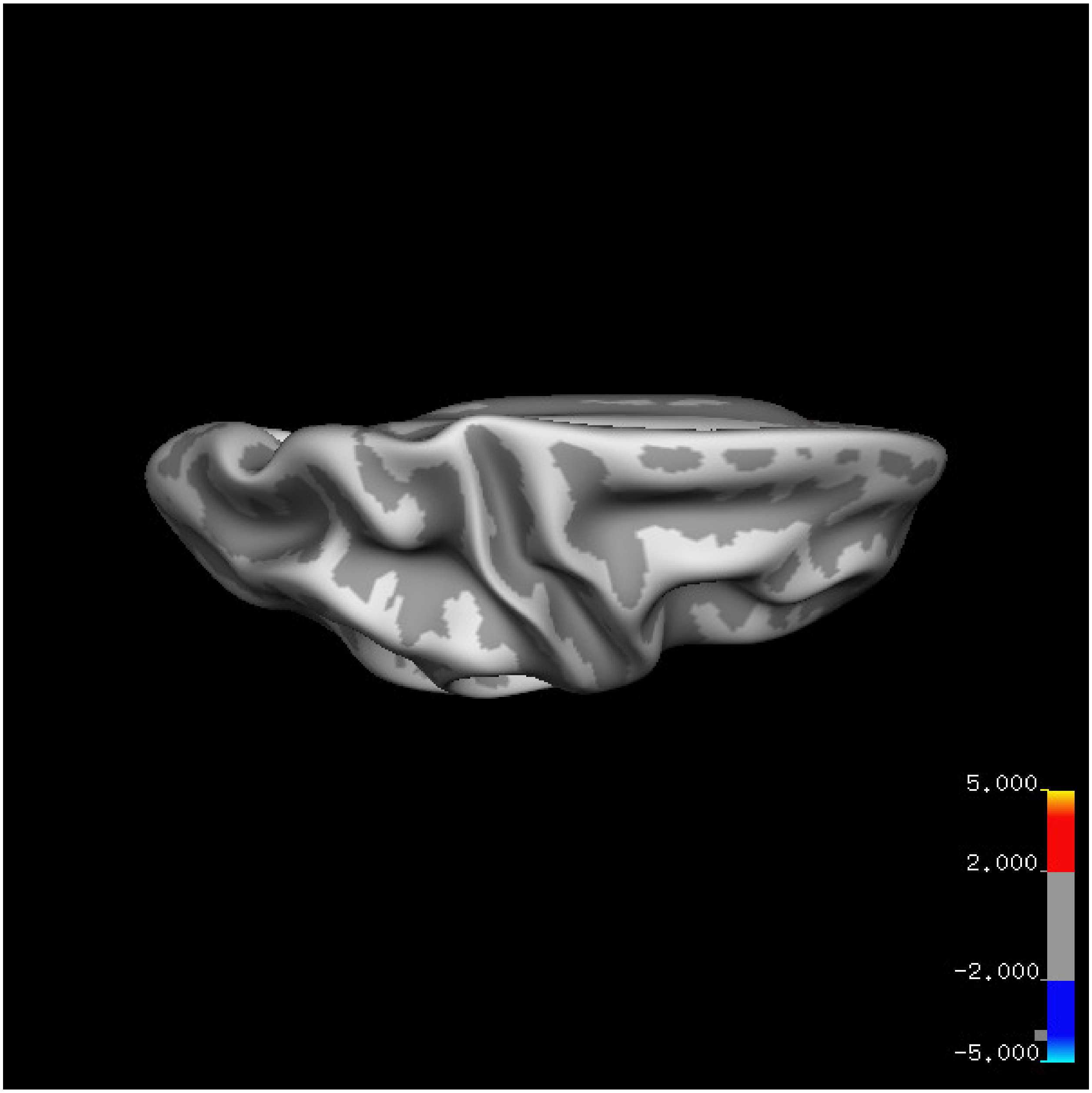

5.000
2.000
-2.000
-5.000

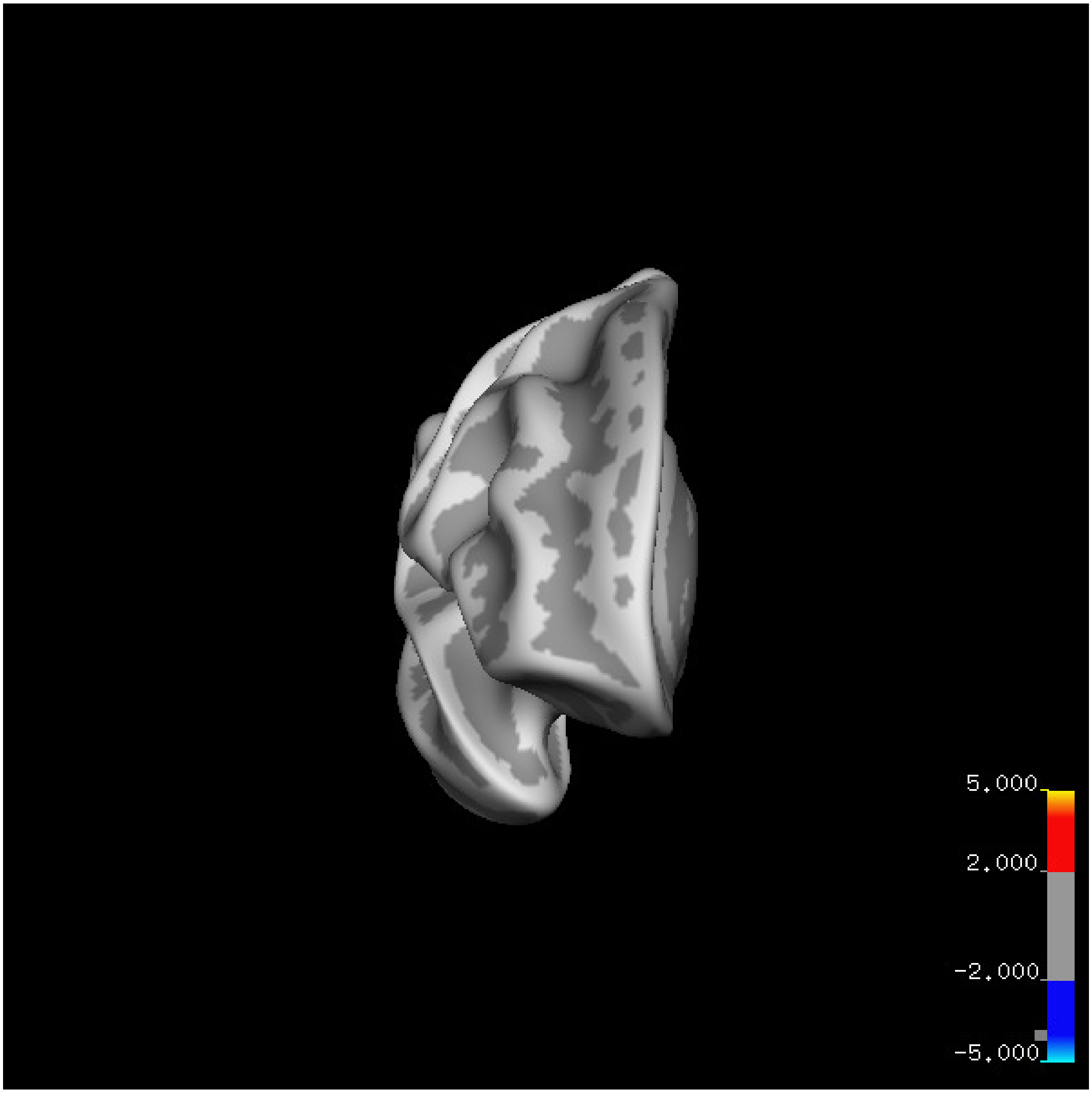
5.000
2.000
-2.000
-5.000

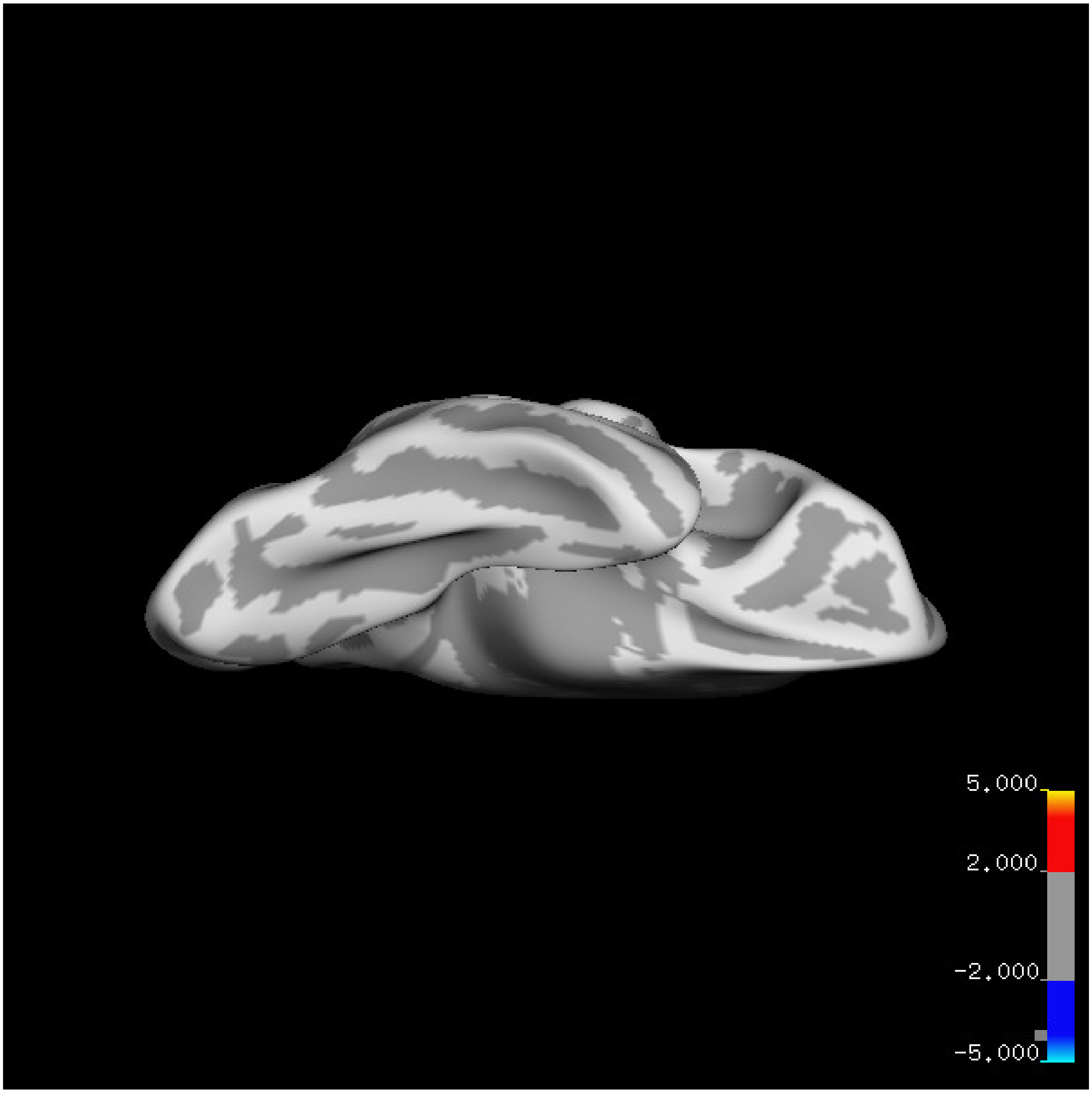
5.000
2.000
-2.000
-5.000

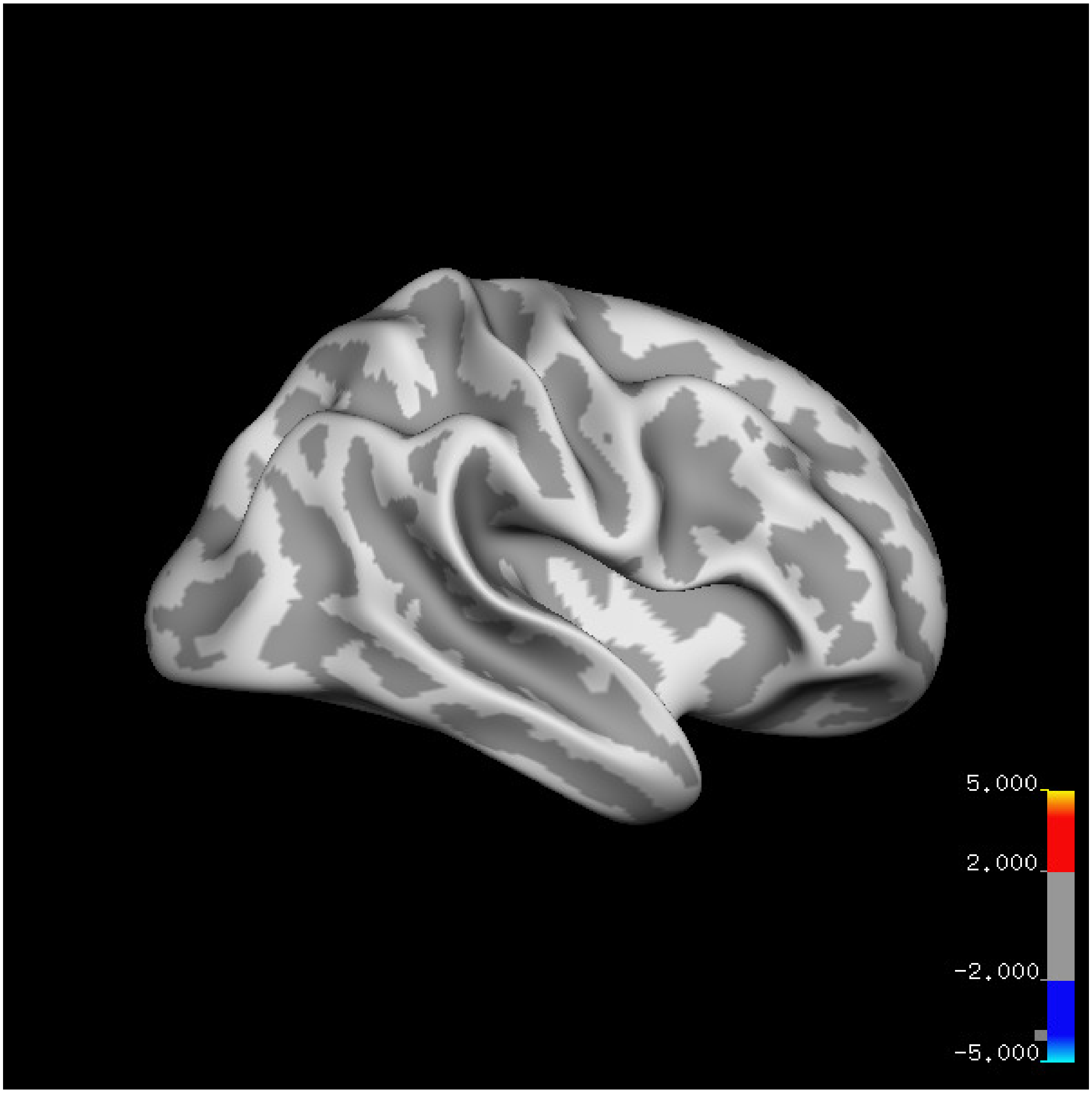
5.000
2.000
-2.000
-5.000

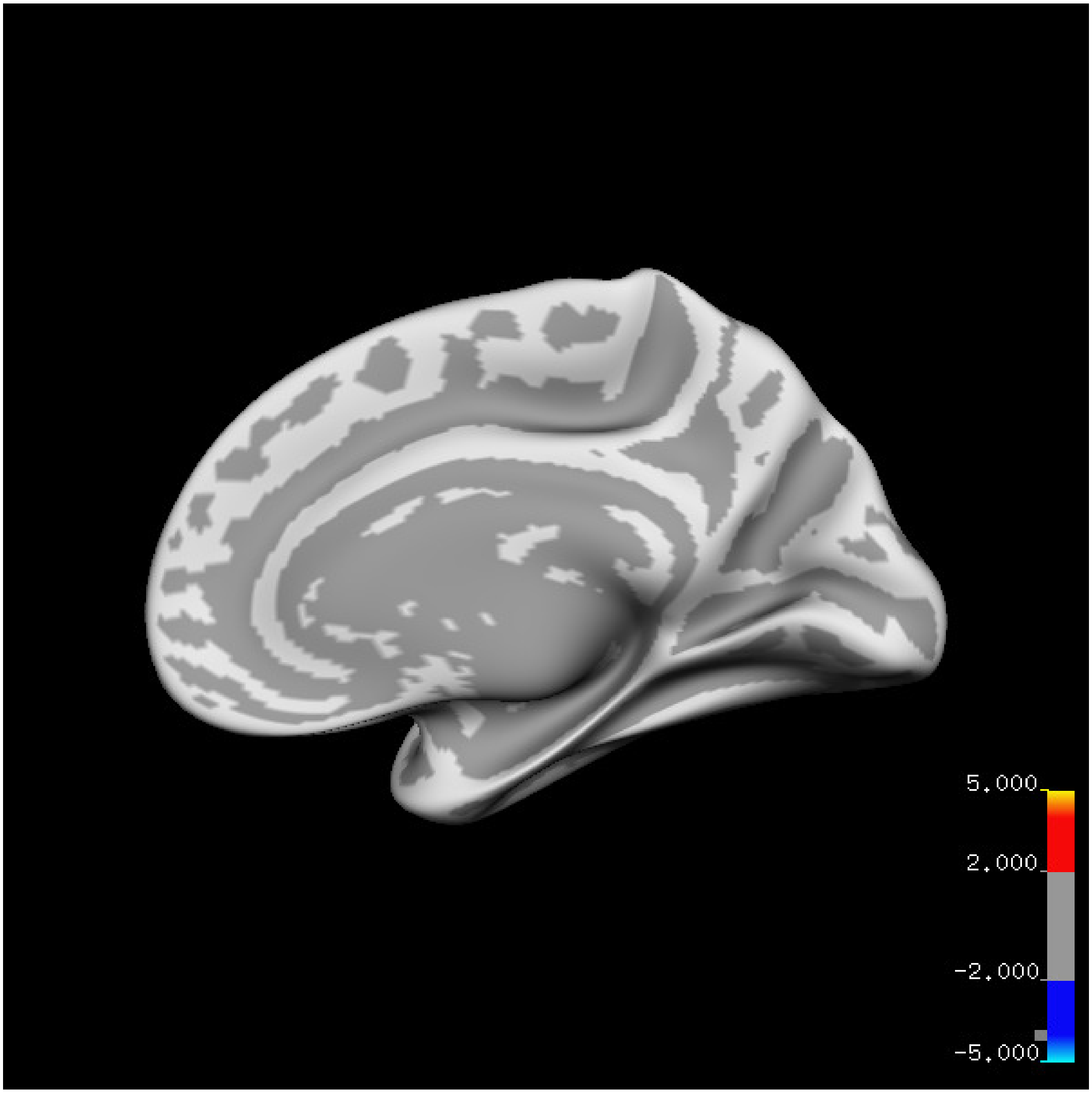
5.000
2.000
-2.000
-5.000

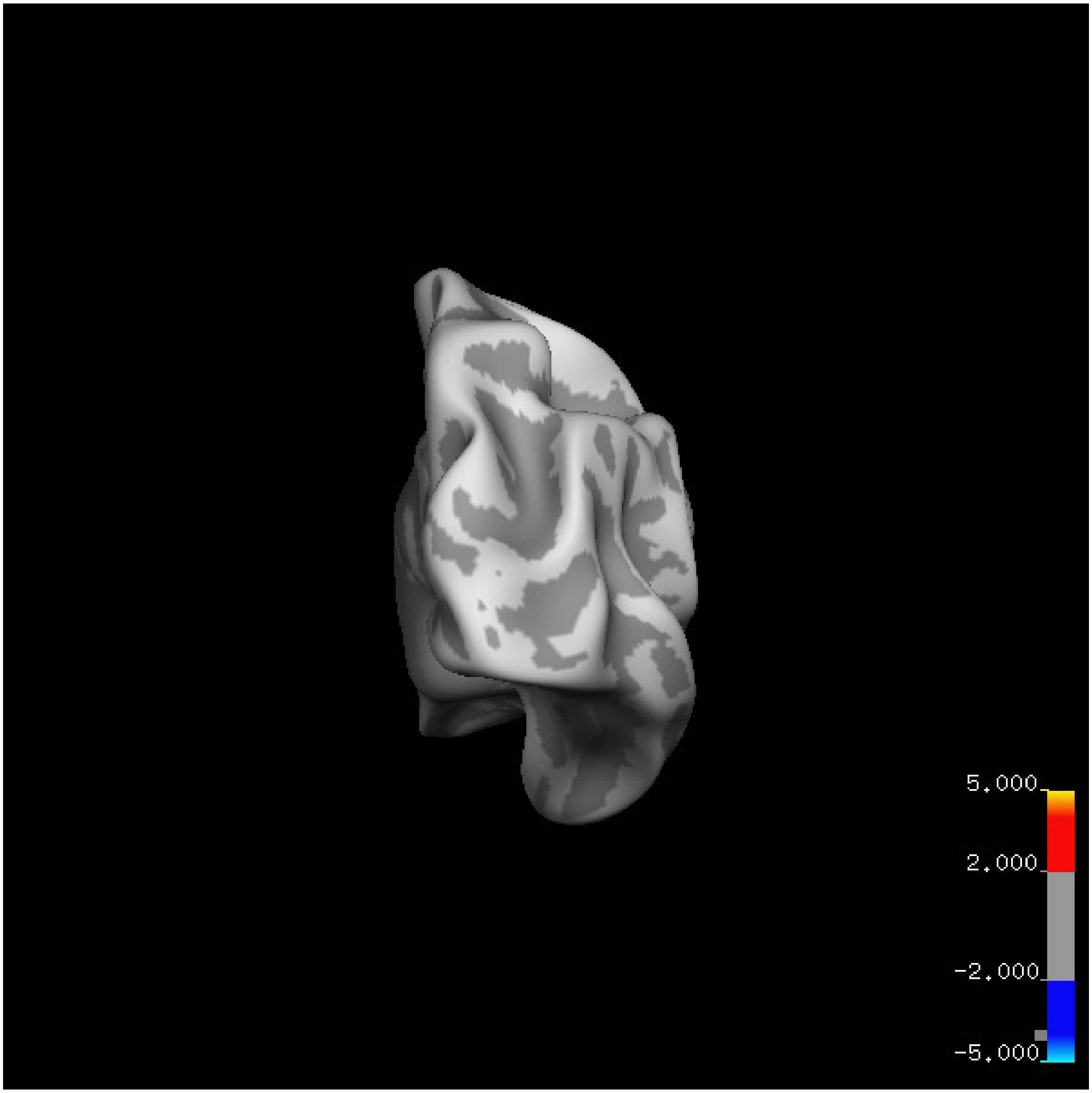
5.000
2.000
-2.000
-5.000

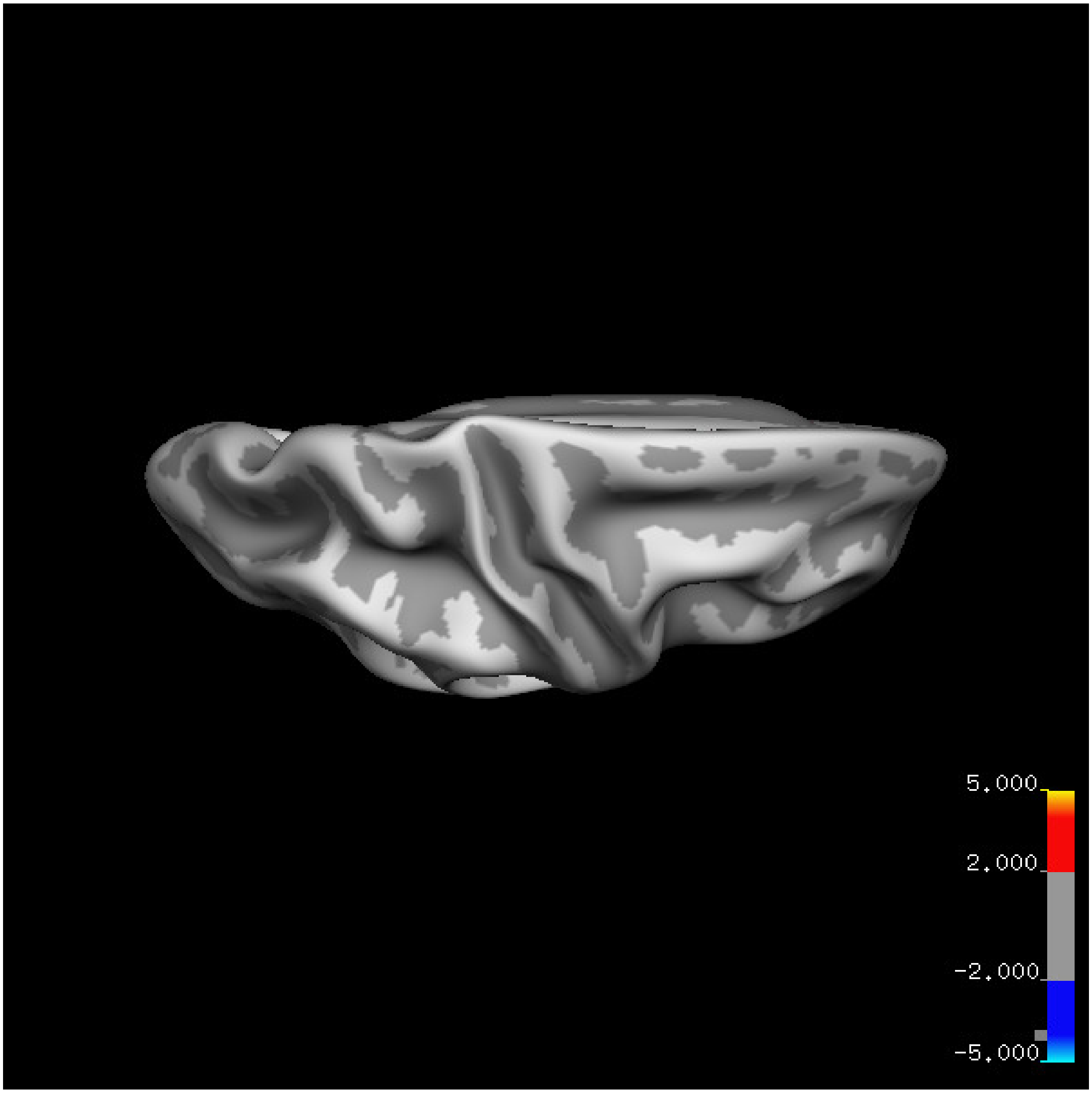


Cluster summary is shown below.

```
## # Cluster Growing Summary (mri_surfcluster)

## # $Id: mri_surfcluster.c,v 1.57.2.3 2016/11/17 18:19:42 zkaufman Exp $

## # $Id: mrisurf.c,v 1.781.2.6 2016/12/27 16:47:14 zkaufman Exp $

## # CreationTime 2021/10/04-10:48:05-GMT

## # cmdline mri_surfcluster.bin --in surface_analyses/results/cluster_comparison/fsaverage6.sm15.rh/thi

## # cwd /gpfs/projects02/p274/cluster/projects/p003-sleep_b

## # sysname  Linux
```

```
## # hostname c1-19
## # machine  x86_64
## # FixVertexAreaFlag 1
## # FixSurfClusterArea 1
## #
## # Input      surface_analyses/results/cluster_comparison/fsaverage6.sm15.rh/thickness.20_60.above_mea
## # Frame Number      0
## # srcsubj fsaverage6
## # hemi rh
## # surface white
## # group_avg_surface_area 85131.2
## # group_avg_vtxarea_loaded 1
## # annot aparc
## # SUBJECTS_DIR /cluster/projects/p274/tools/mri/freesurfer/current/subjects
## # SearchSpace_mm2 79329.9
## # SearchSpace_vtx 37471
## # Bonferroni 0
## # Minimum Threshold 1.3
## # Maximum Threshold infinity
## # Threshold Sign    abs
## # AdjustThreshWhenOneTail 1
## # CW PValue Threshold: 0.01
## # Area Threshold    0 mm^2
## # CSD thresh  1.300000
## # CSD nreps    10000
## # CSD simtype  null-z
## # CSD contrast NA
```

```
## # CSD confint  90.000000

## # Overall max 1.78049 at vertex 21221

## # Overall min -3.48972 at vertex 18719

## # NClusters          0

## # FixMNI = 0

## #

## # ClusterNo  Max   VtxMax   Size(mm^2)  MNIX   MNIY   MNIZ    CWP    CWPLow    CWPHi   NVtxs    Wght
```

**Sleep below mean** Cluster corrected significance plots are shown below.

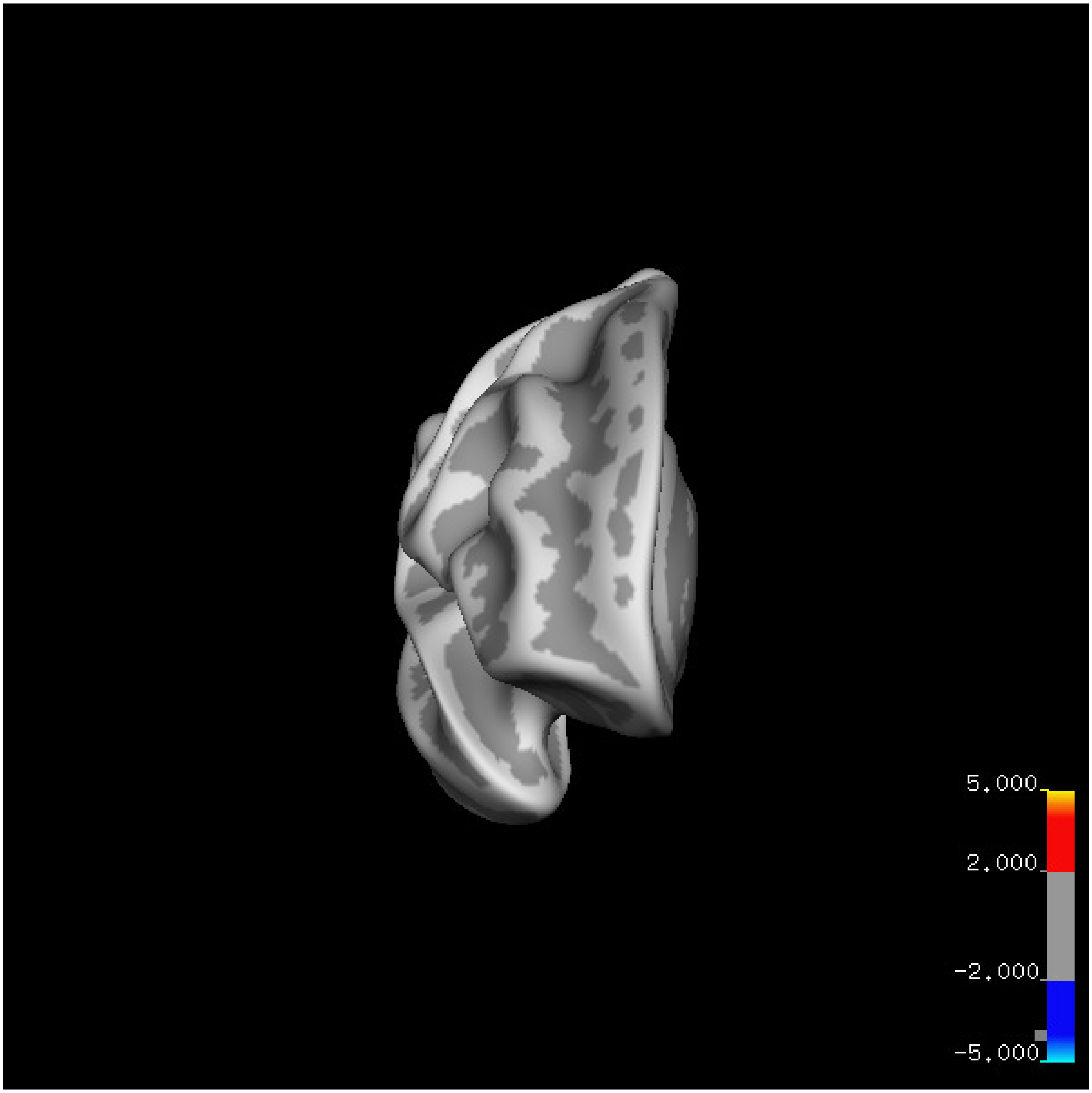
5.000
2.000
-2.000
-5.000

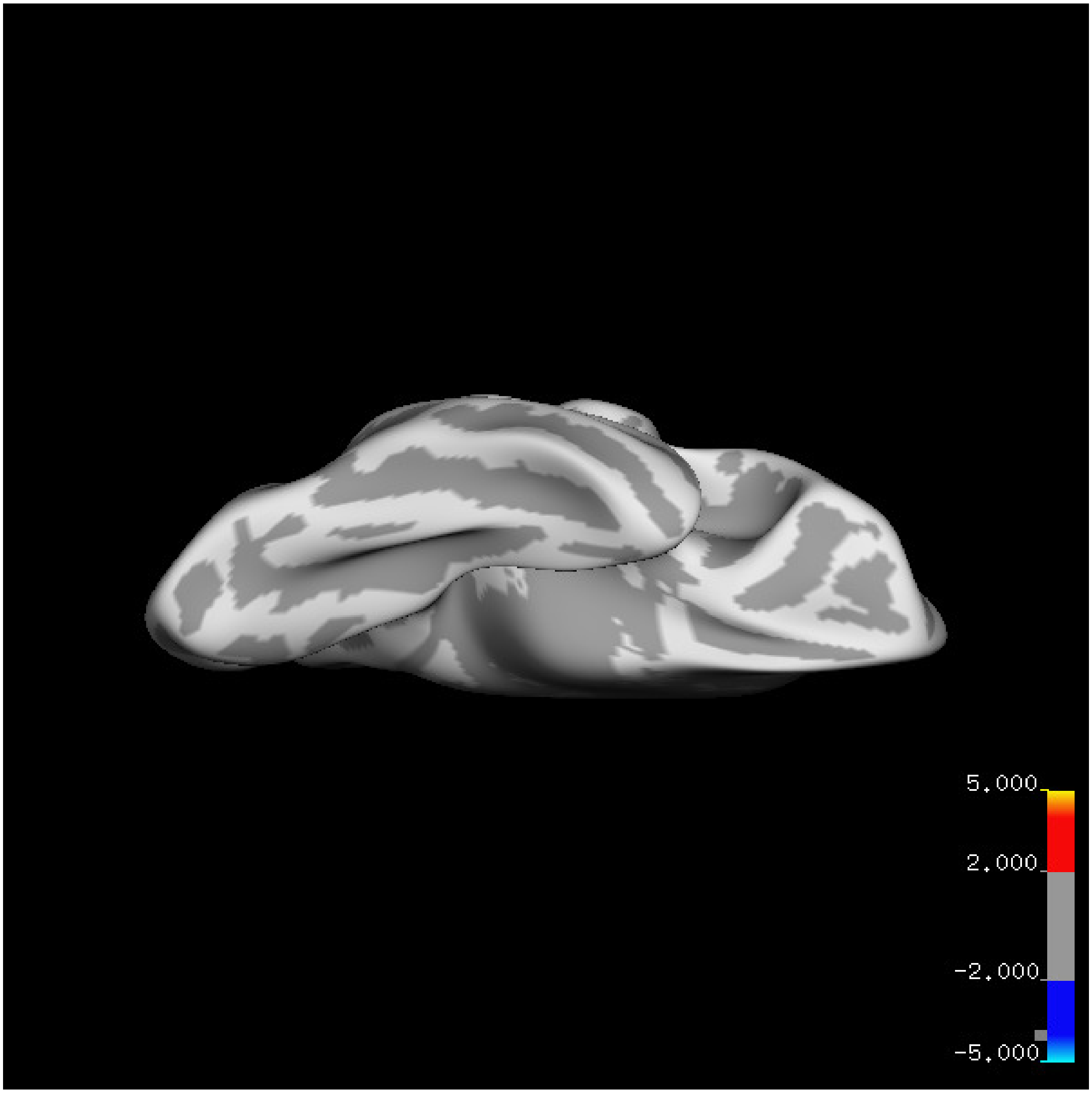
5.000
2.000
-2.000
-5.000

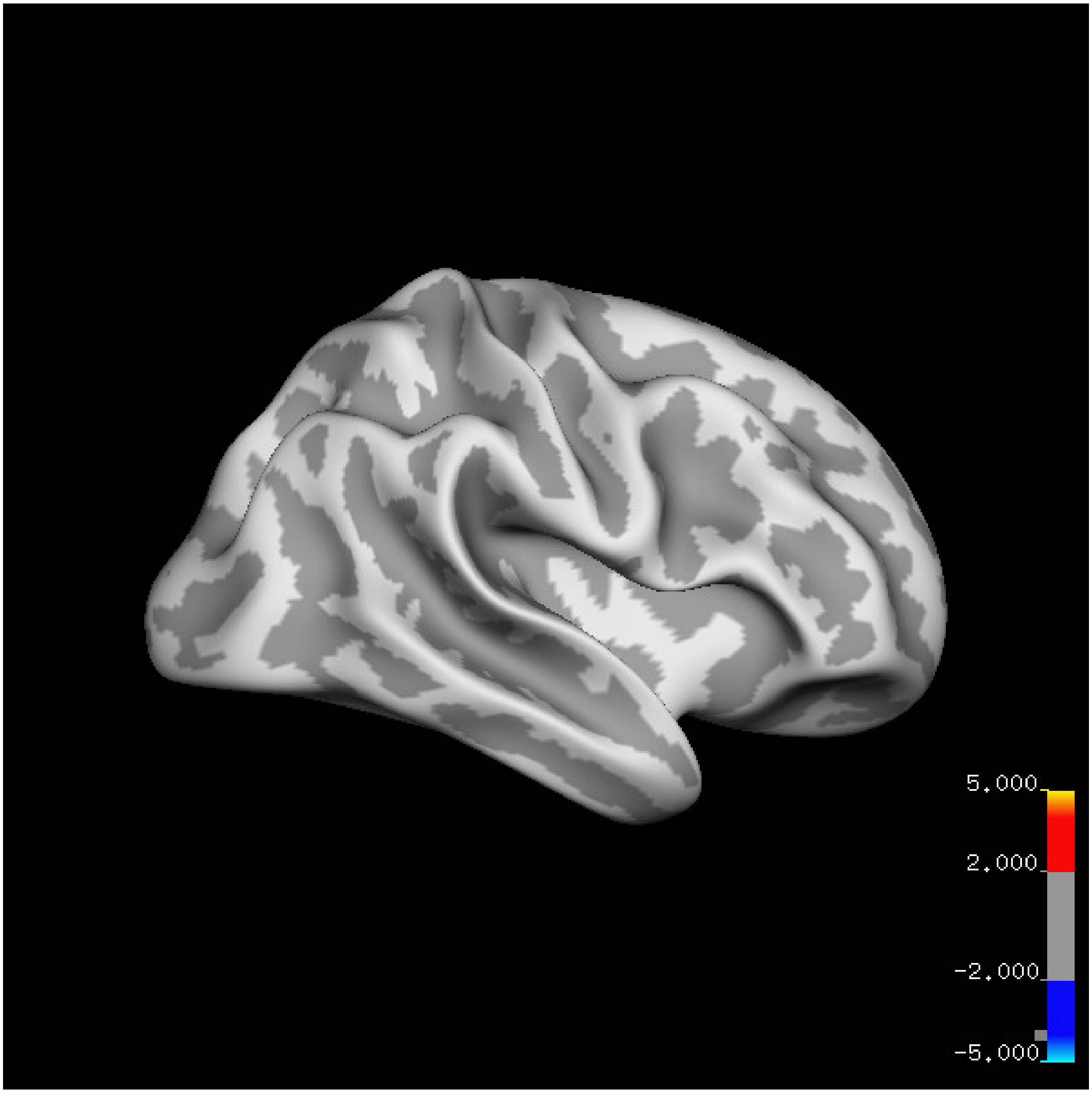
5.000
2.000
-2.000
-5.000

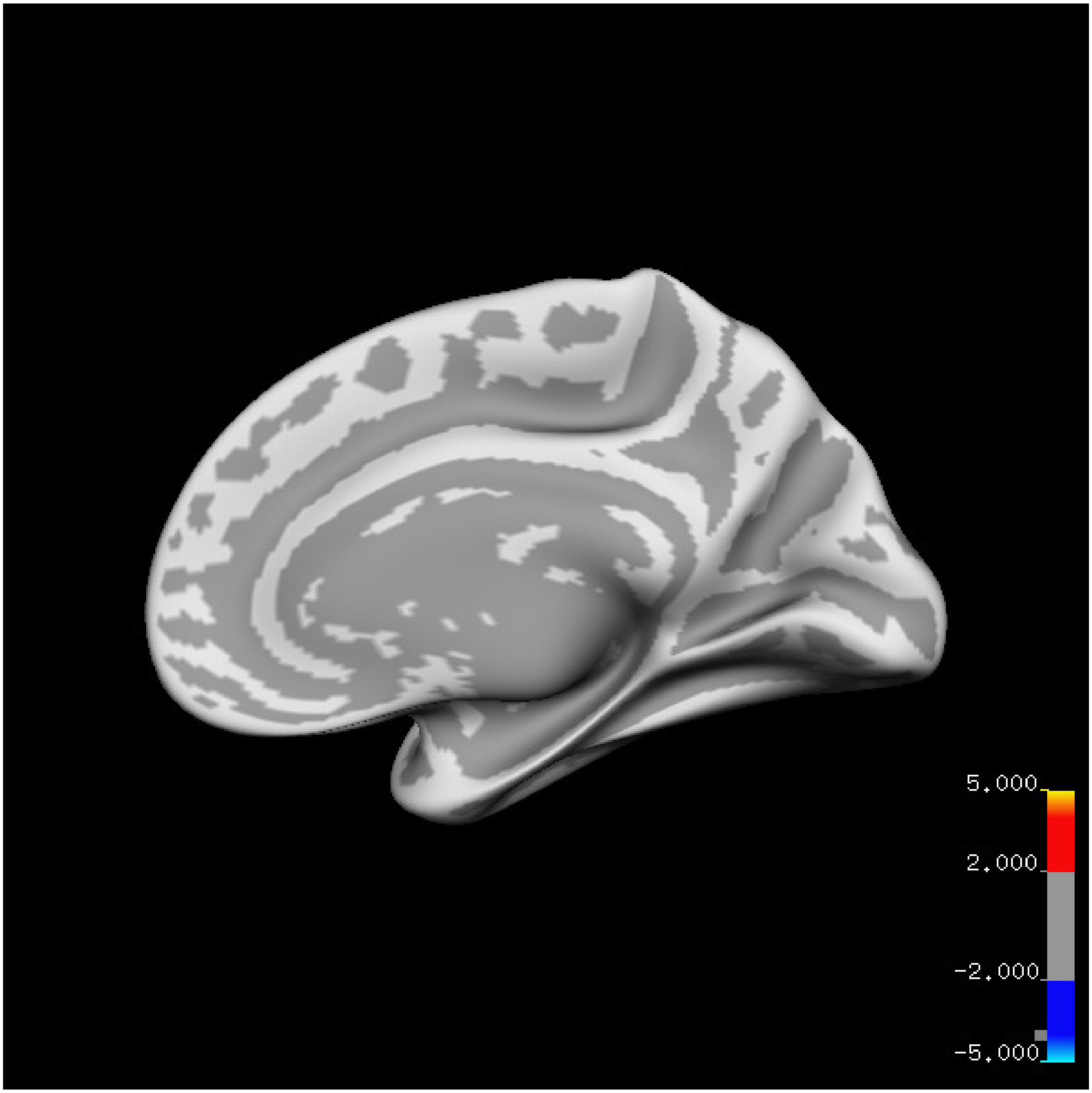
5.000
2.000
-2.000
-5.000

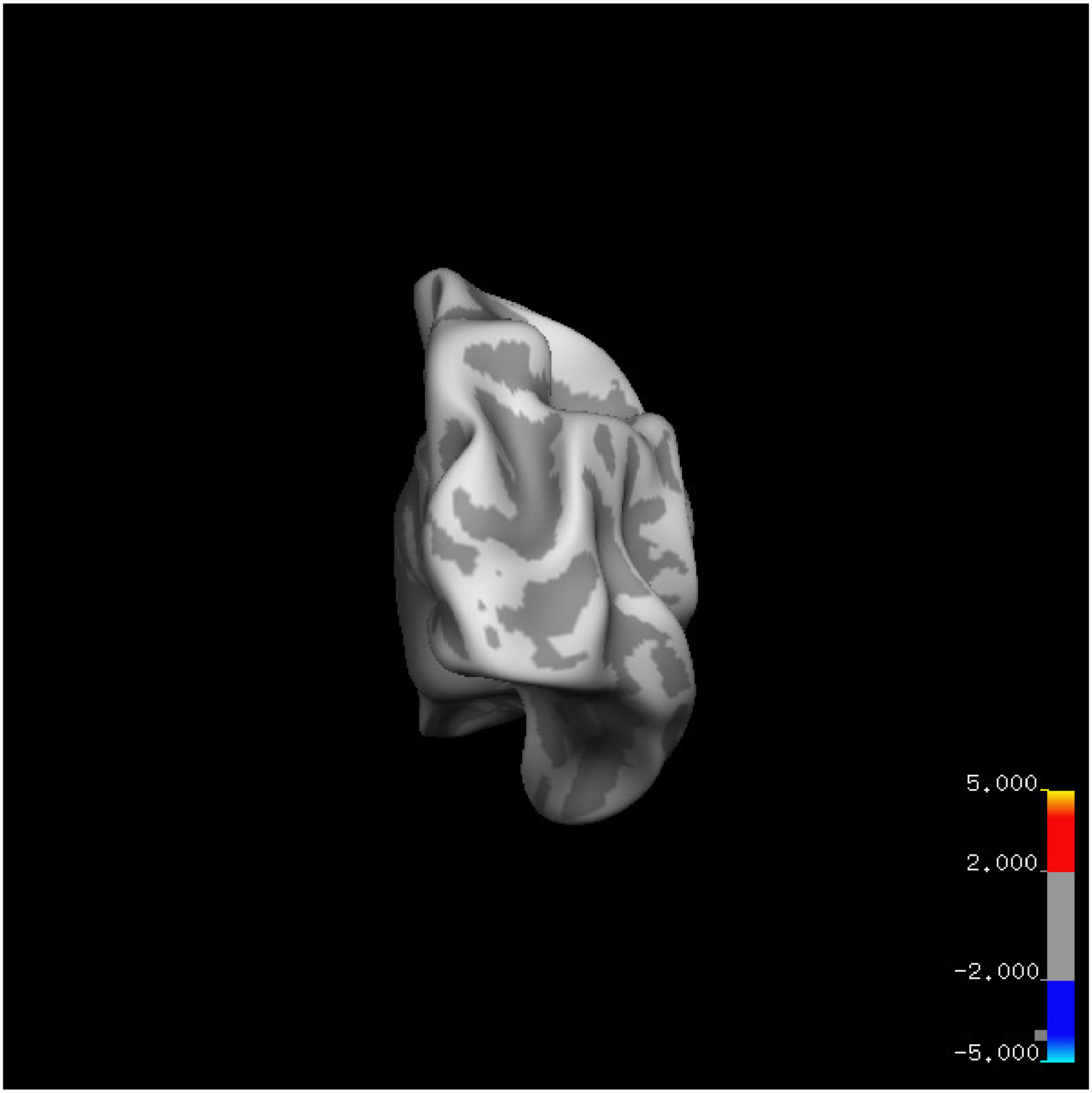
5.000
2.000
-2.000
-5.000

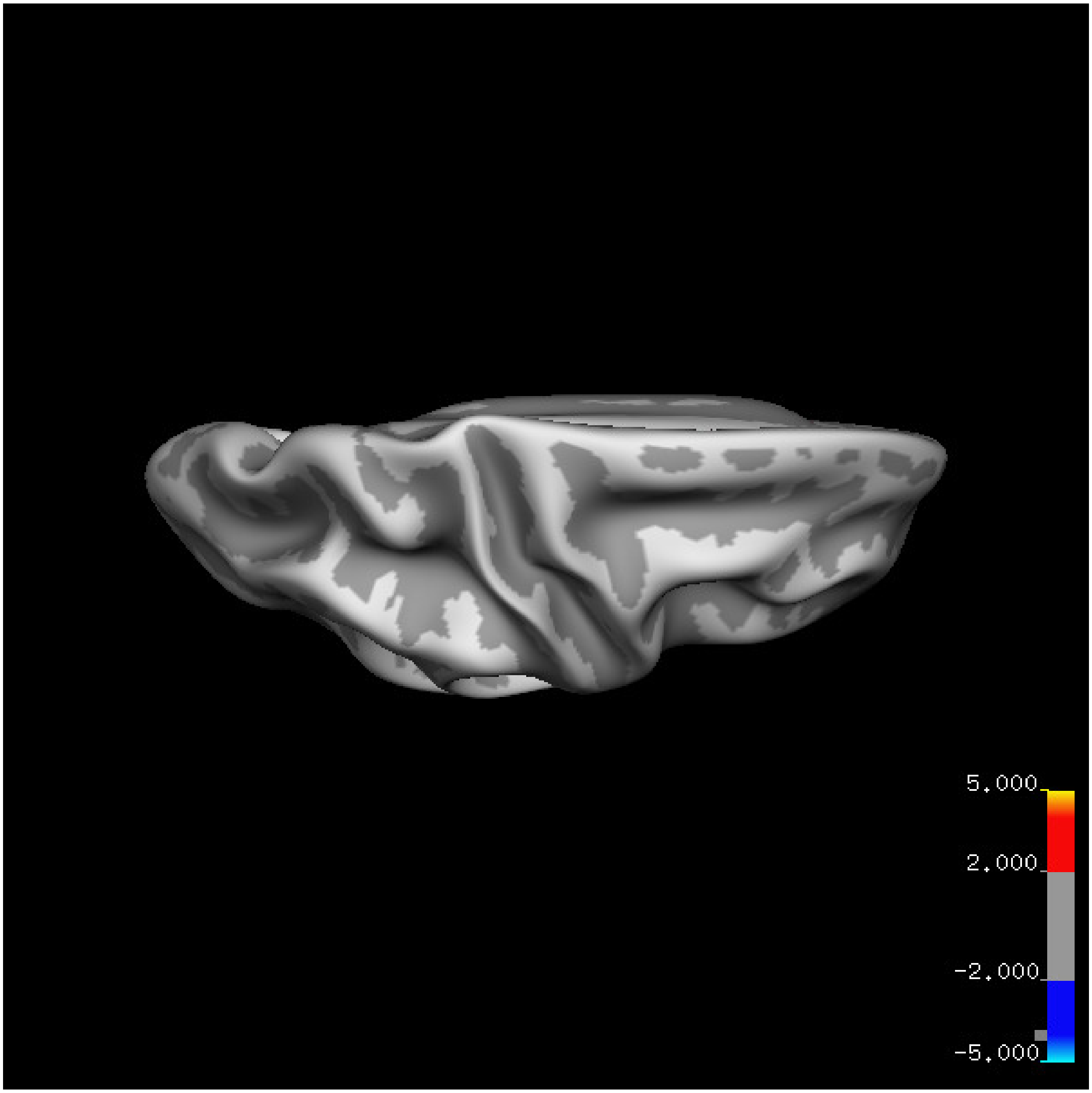
5.000
2.000
-2.000
-5.000

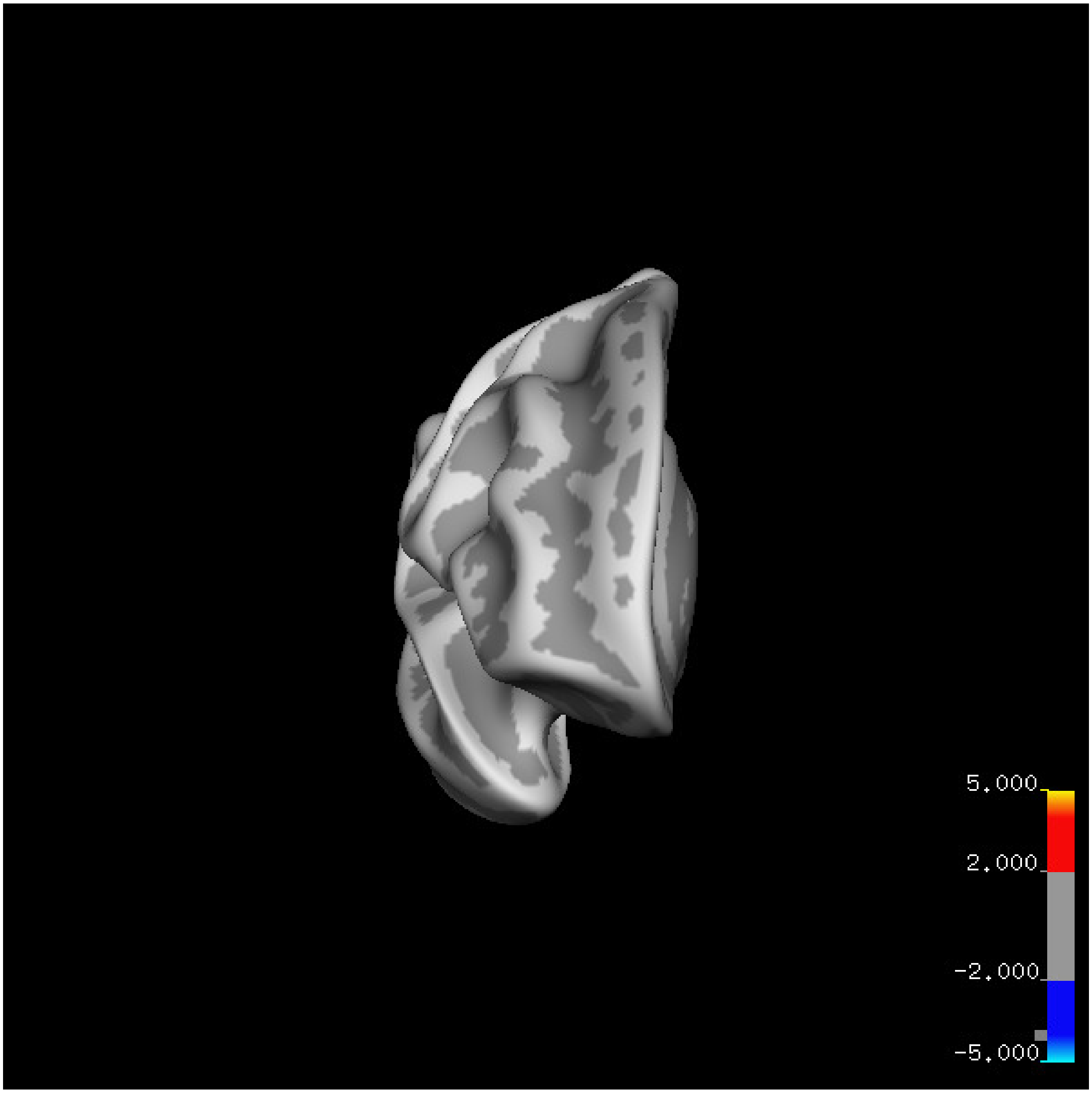
5.000
2.000
-2.000
-5.000

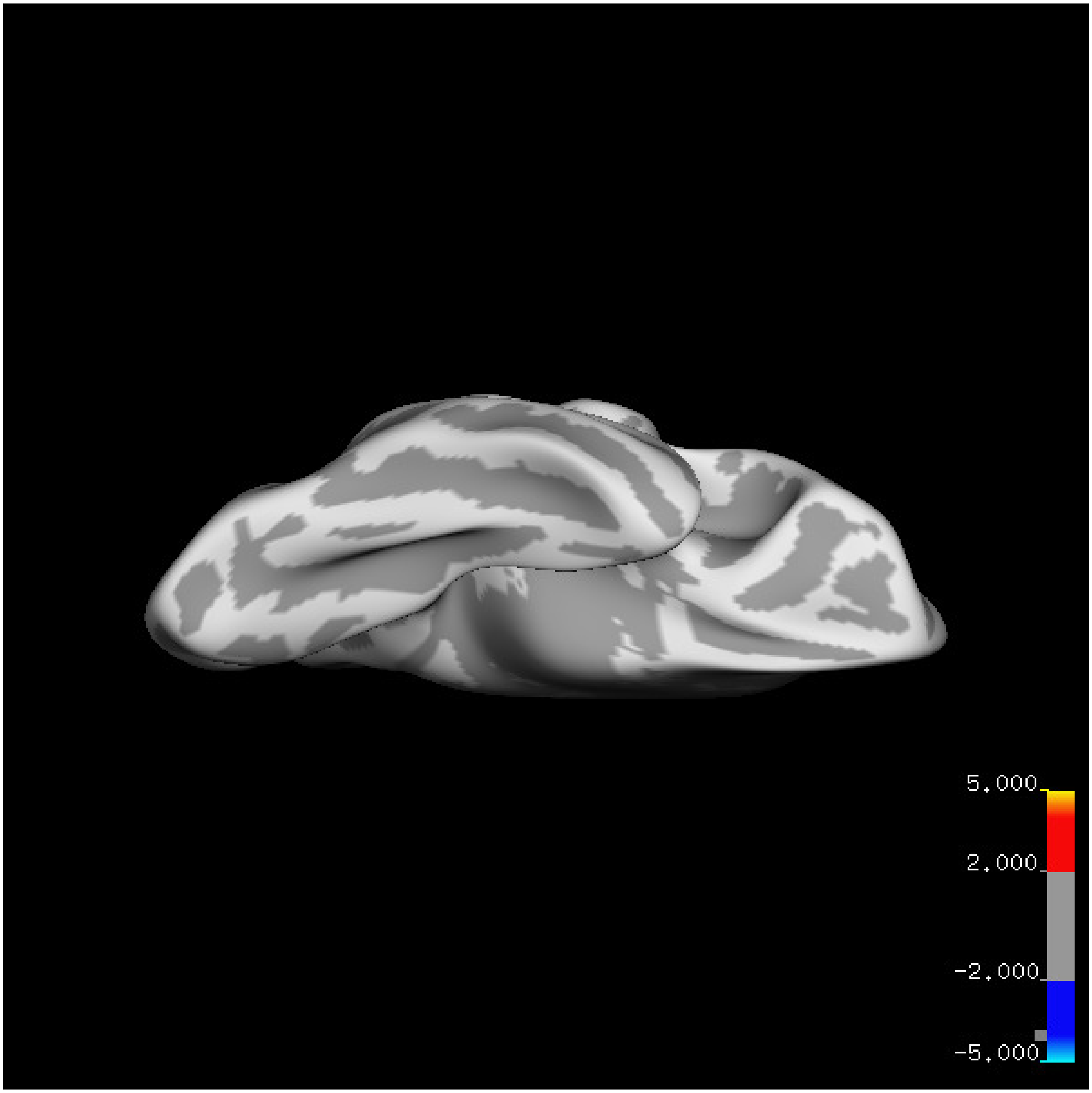

5.000
2.000
-2.000
-5.000

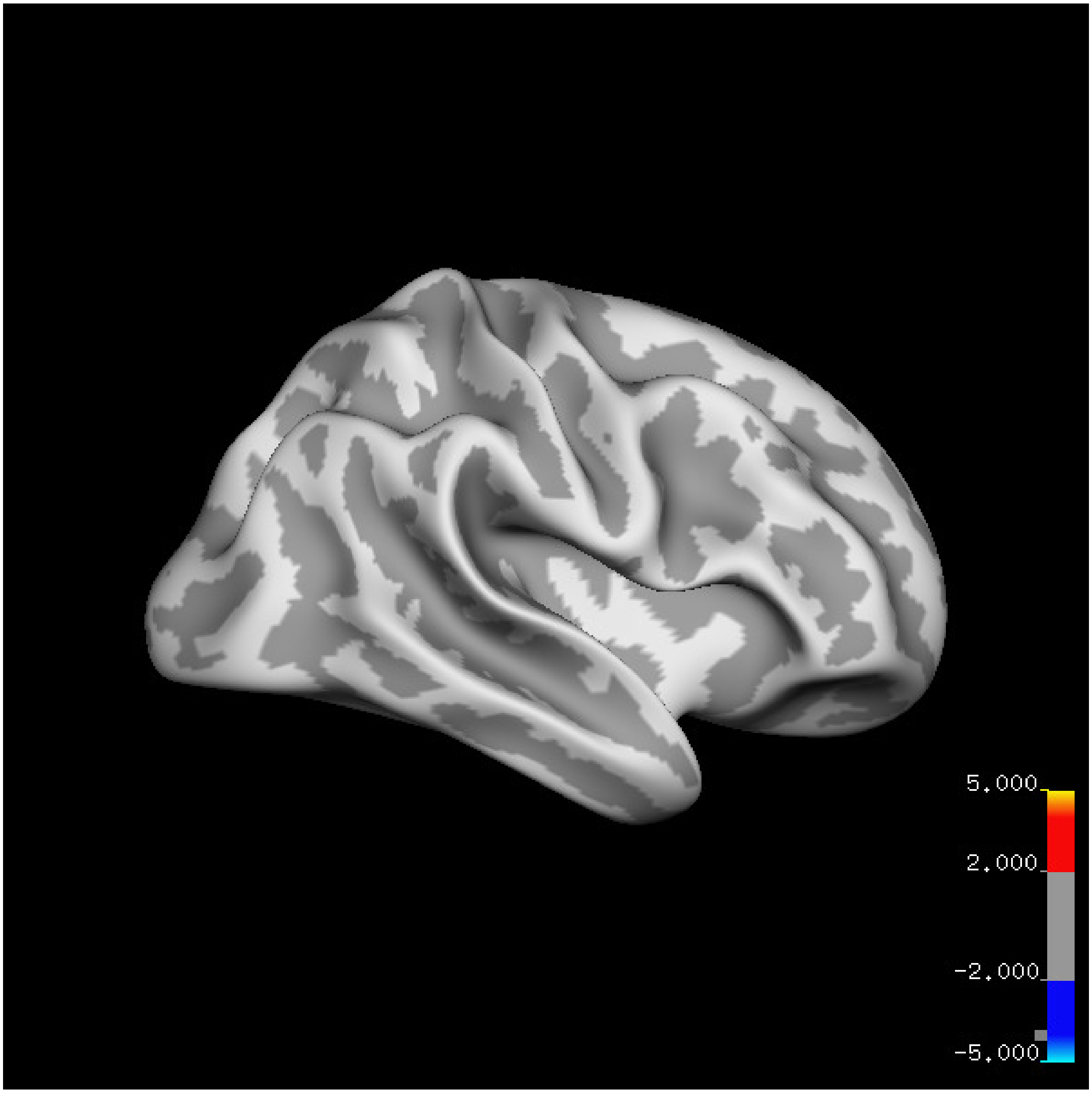

5.000
2.000
-2.000
-5.000

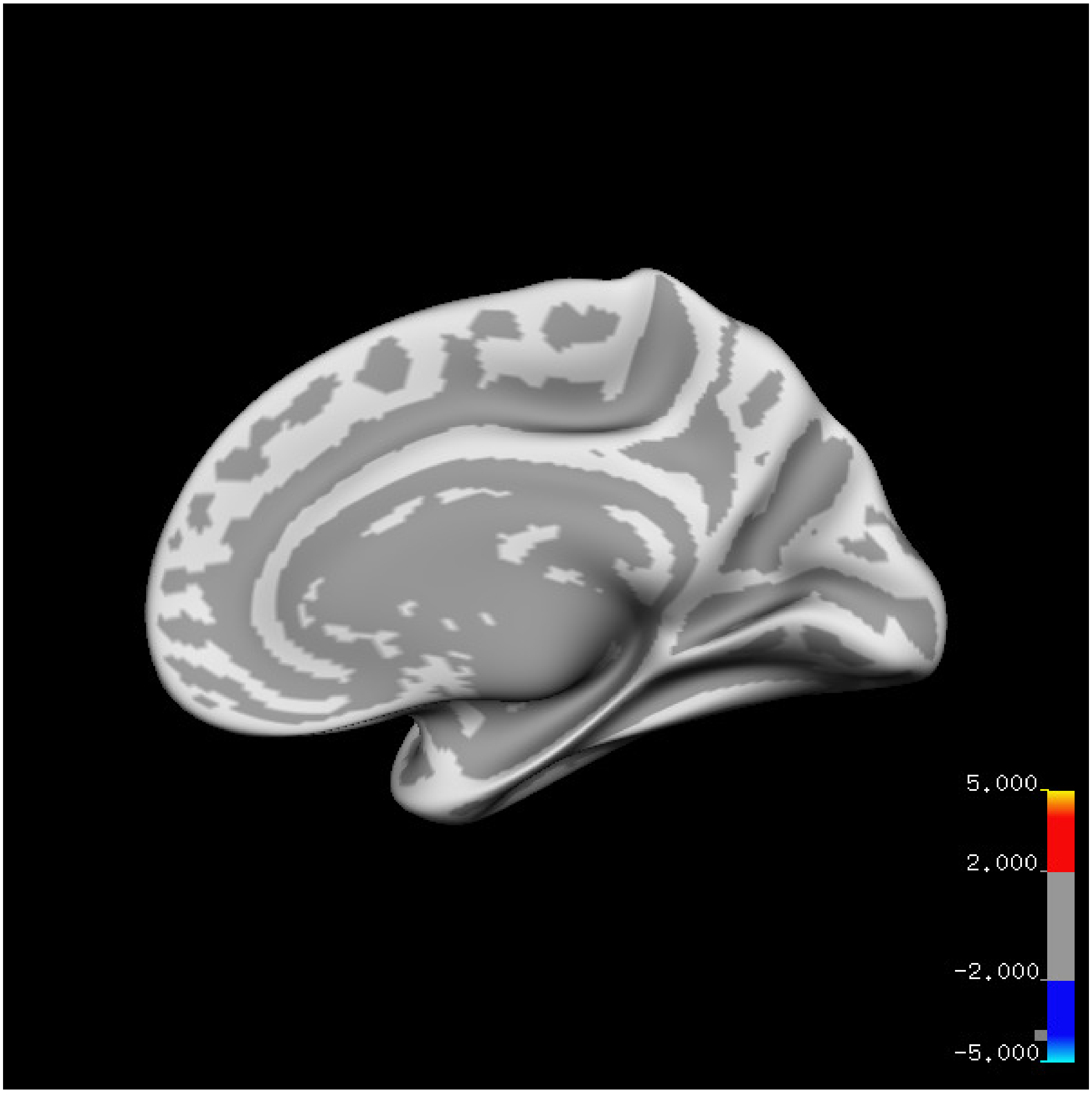
5.000
2.000
-2.000
-5.000

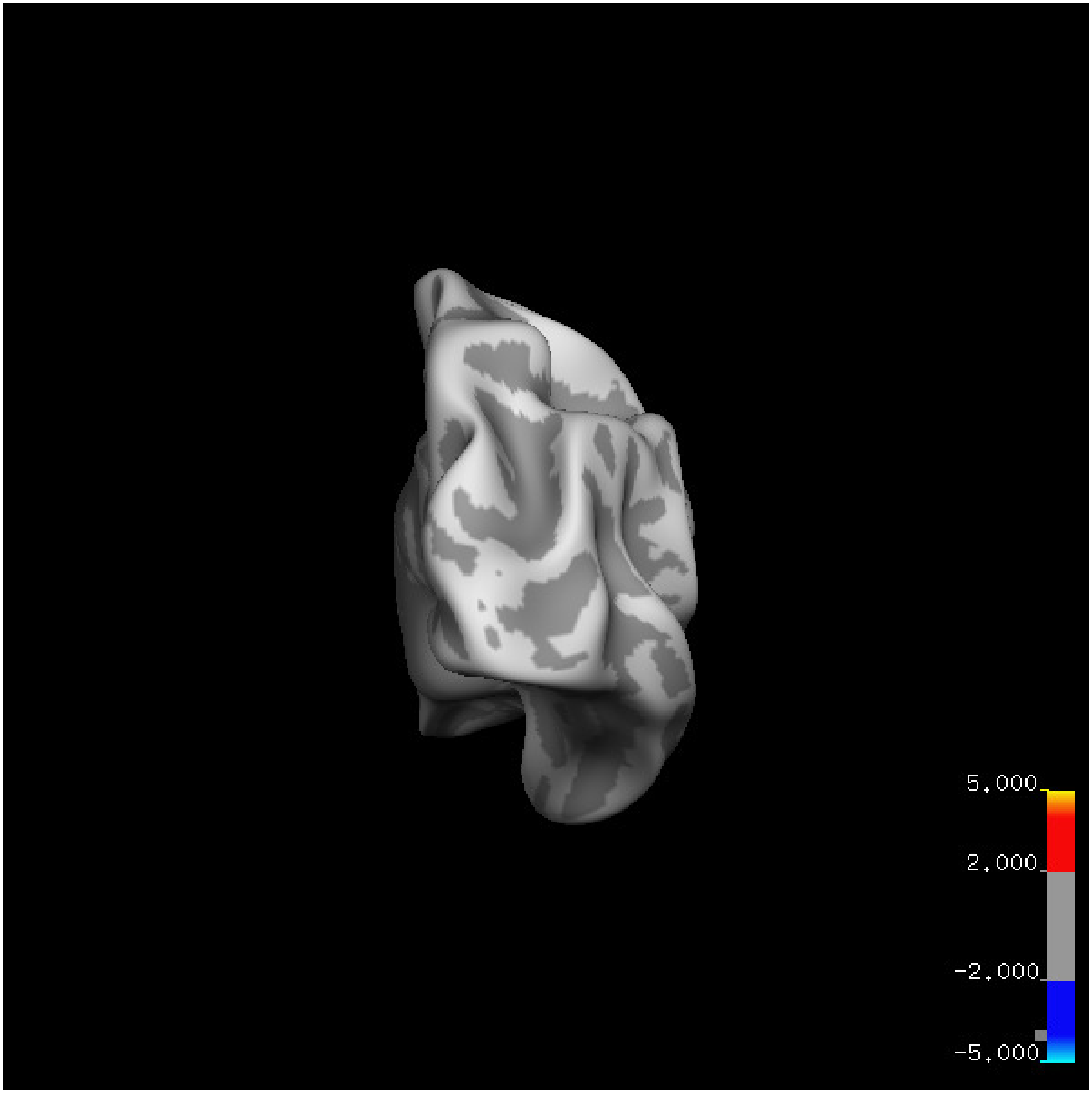
5.000
2.000
-2.000
-5.000

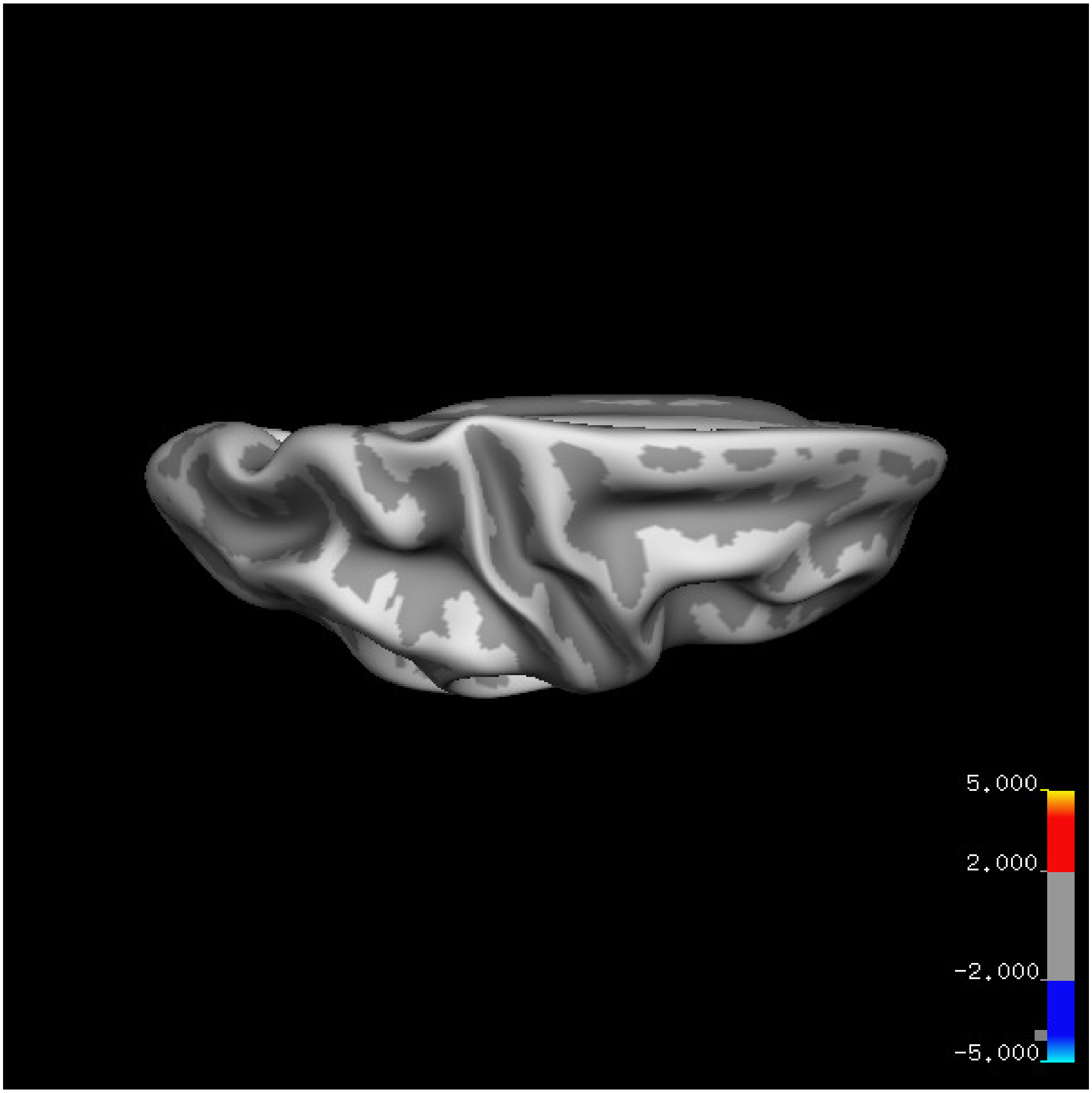


Cluster summary is shown below.

```
## # Cluster Growing Summary (mri_surfcluster)

## # $Id: mri_surfcluster.c,v 1.57.2.3 2016/11/17 18:19:42 zkaufman Exp $

## # $Id: mrisurf.c,v 1.781.2.6 2016/12/27 16:47:14 zkaufman Exp $

## # CreationTime 2021/10/04-11:24:43-GMT

## # cmdline mri_surfcluster.bin --in surface_analyses/results/cluster_comparison/fsaverage6.sm15.rh/th

## # cwd /gpfs/projects02/p274/cluster/projects/p003-sleep_b

## # sysname  Linux
```

```
## # hostname c1-19
## # machine  x86_64
## # FixVertexAreaFlag 1
## # FixSurfClusterArea 1
## #
## # Input      surface_analyses/results/cluster_comparison/fsaverage6.sm15.rh/thickness.20_60.below_mea
## # Frame Number      0
## # srcsubj fsaverage6
## # hemi rh
## # surface white
## # group_avg_surface_area 85131.2
## # group_avg_vtxarea_loaded 1
## # annot aparc
## # SUBJECTS_DIR /cluster/projects/p274/tools/mri/freesurfer/current/subjects
## # SearchSpace_mm2 79329.9
## # SearchSpace_vtx 37471
## # Bonferroni 0
## # Minimum Threshold 1.3
## # Maximum Threshold infinity
## # Threshold Sign    abs
## # AdjustThreshWhenOneTail 1
## # CW PValue Threshold: 0.01
## # Area Threshold    0 mm^2
## # CSD thresh  1.300000
## # CSD nreps    10000
## # CSD simtype  null-z
## # CSD contrast NA
```

```
## # CSD confint  90.000000

## # Overall max 3.62774 at vertex 35320

## # Overall min -3.44113 at vertex 31171

## # NClusters          0

## # FixMNI = 0

## #

## # ClusterNo  Max   VtxMax   Size(mm^2)  MNIX   MNIY   MNIZ    CWP    CWPLow    CWPHi   NVtxs    Wght
```

**Age above 60**

**Sleep above mean** Cluster corrected significance plots are shown below.

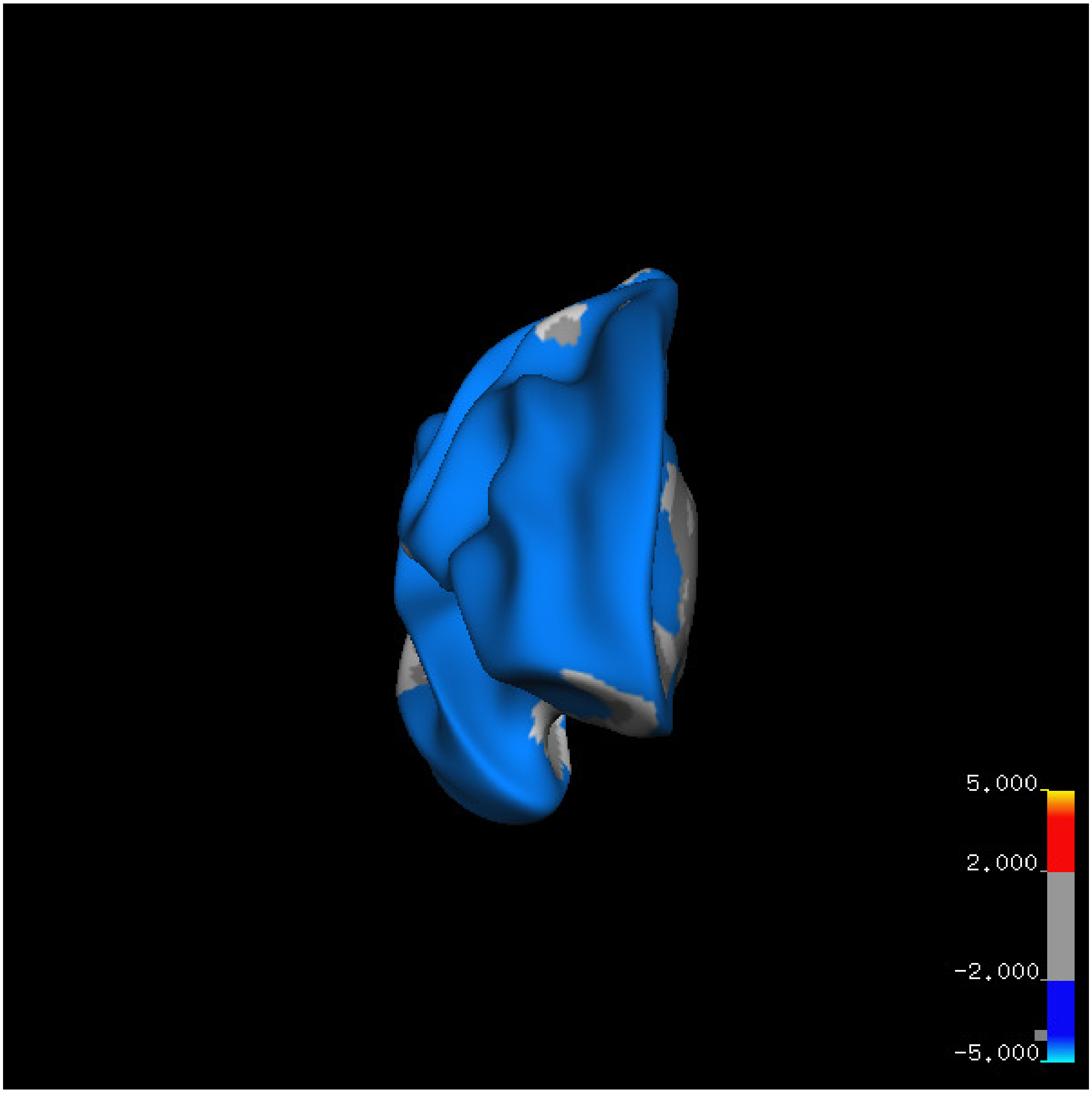
5.000
2.000
-2.000
-5.000

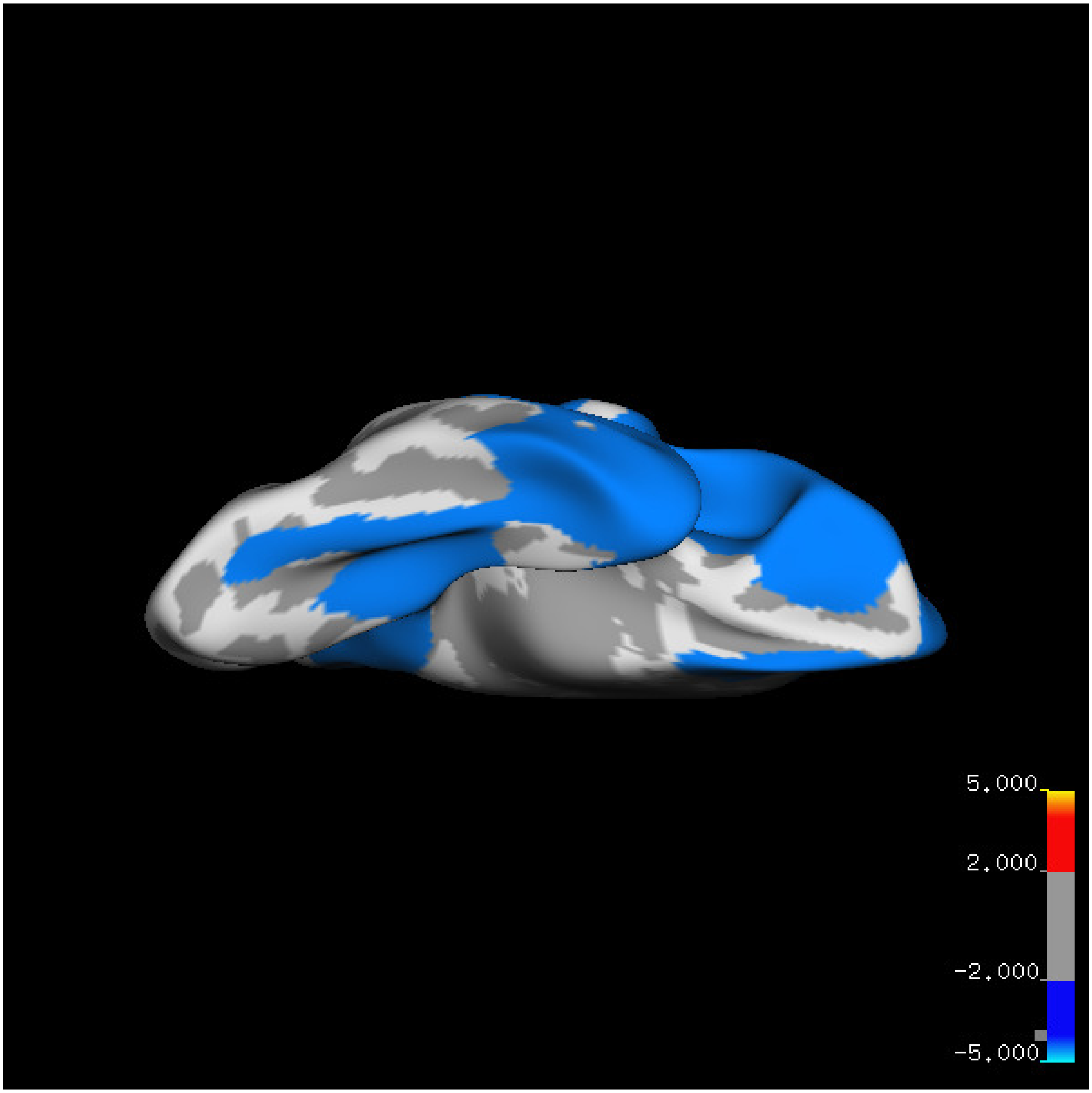
5.000
2.000
-2.000
-5.000

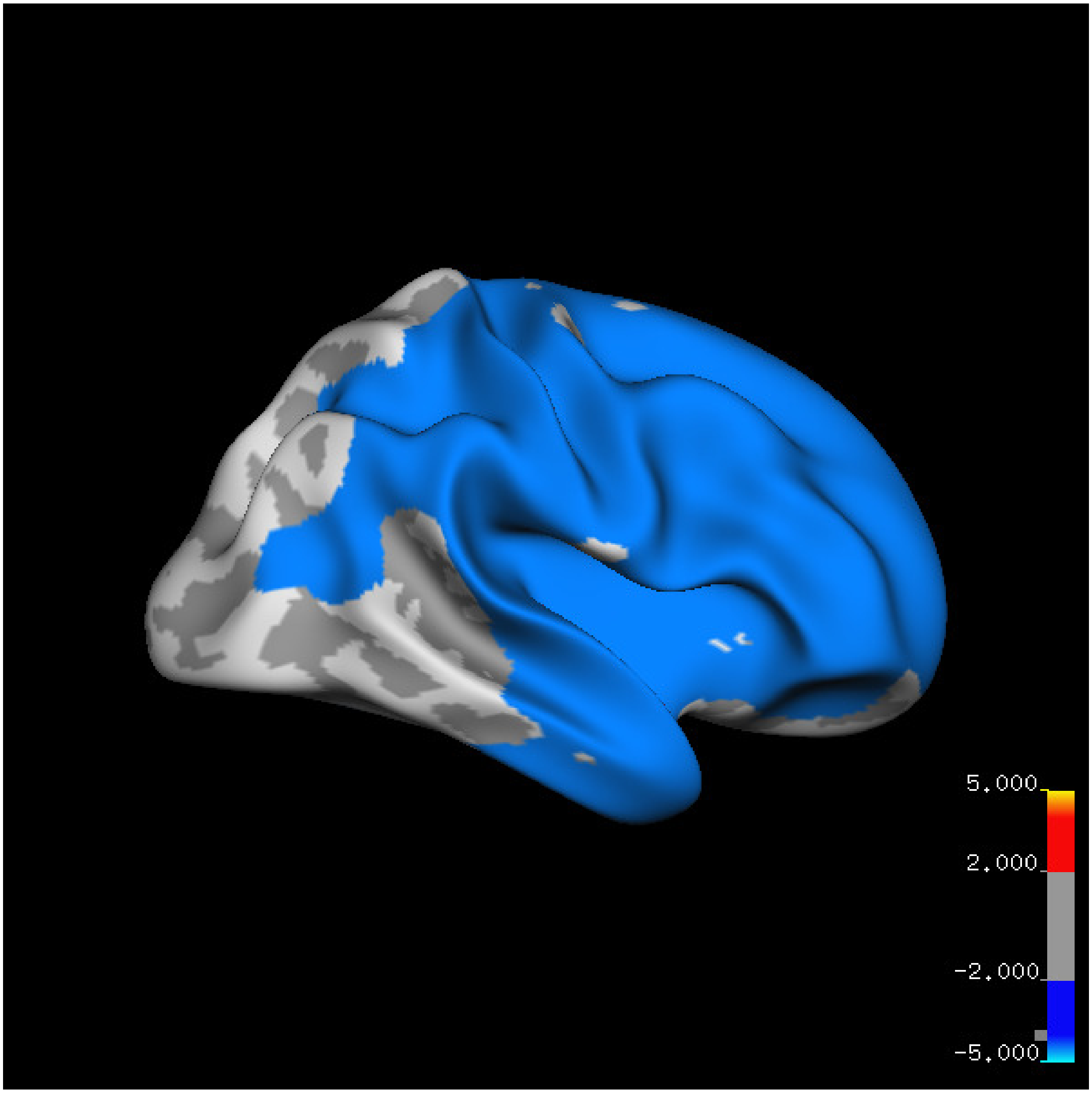
5.000
2.000
-2.000
-5.000

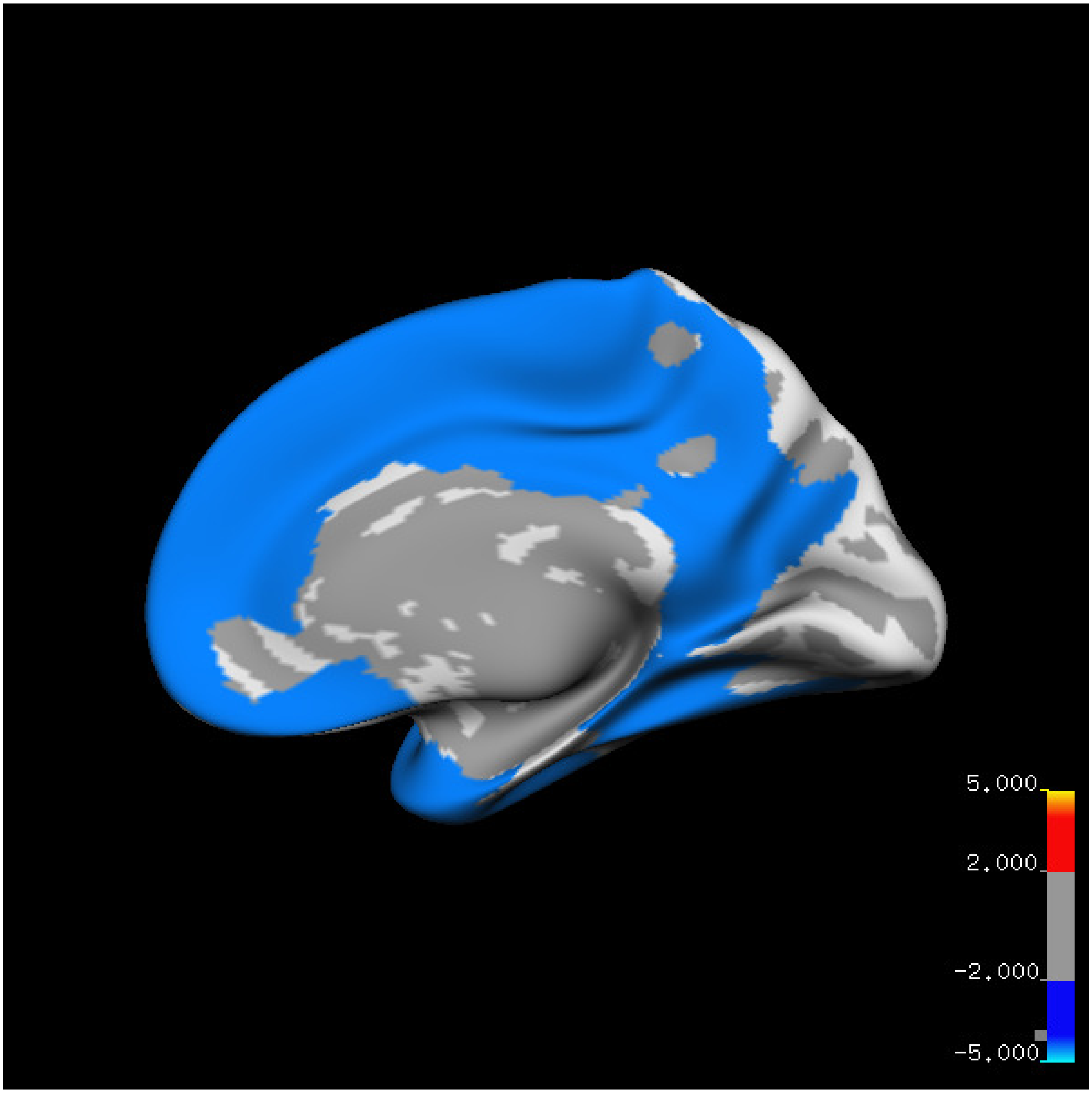
5.000
2.000
-2.000
-5.000

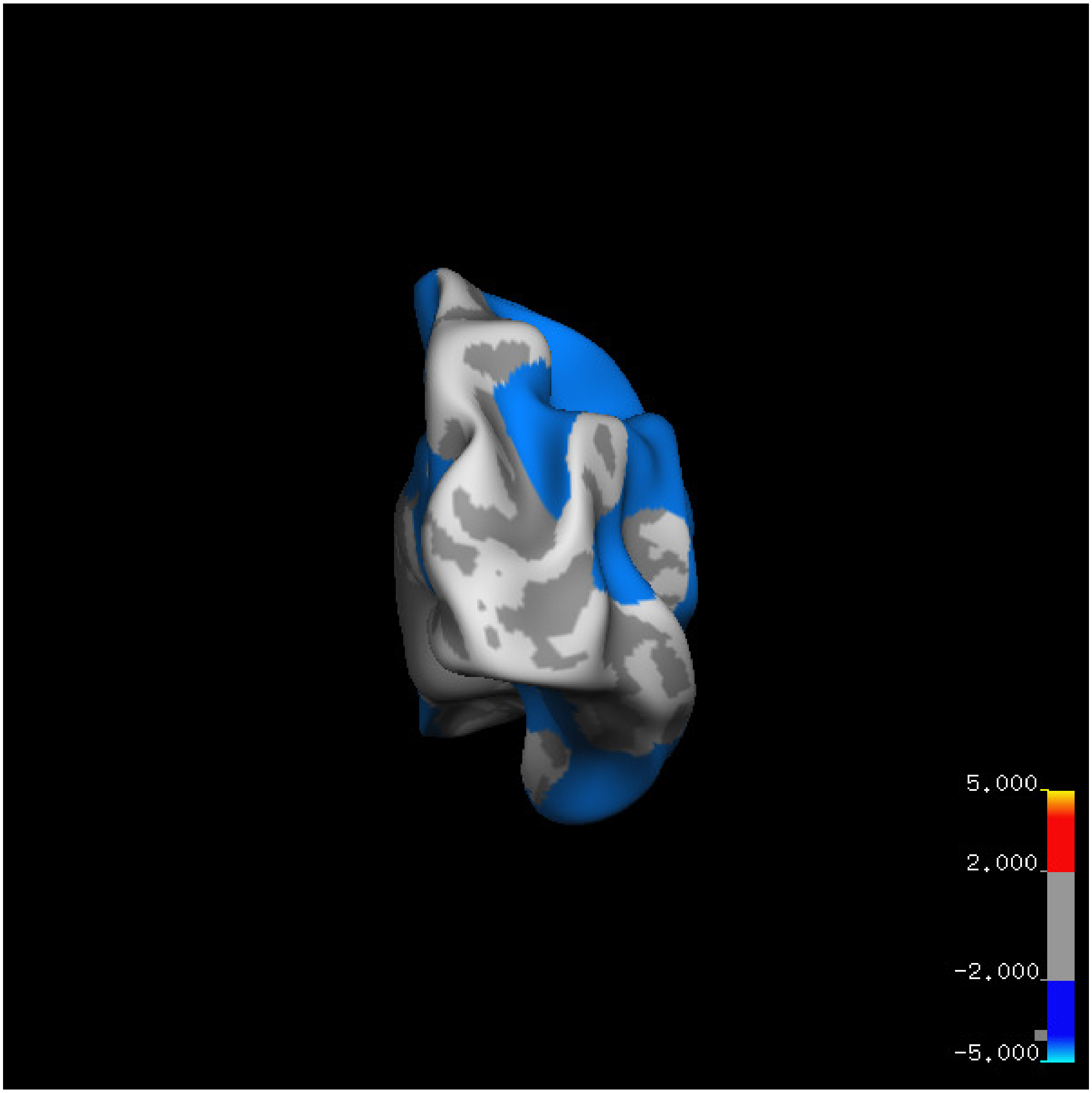
5.000
2.000
-2.000
-5.000

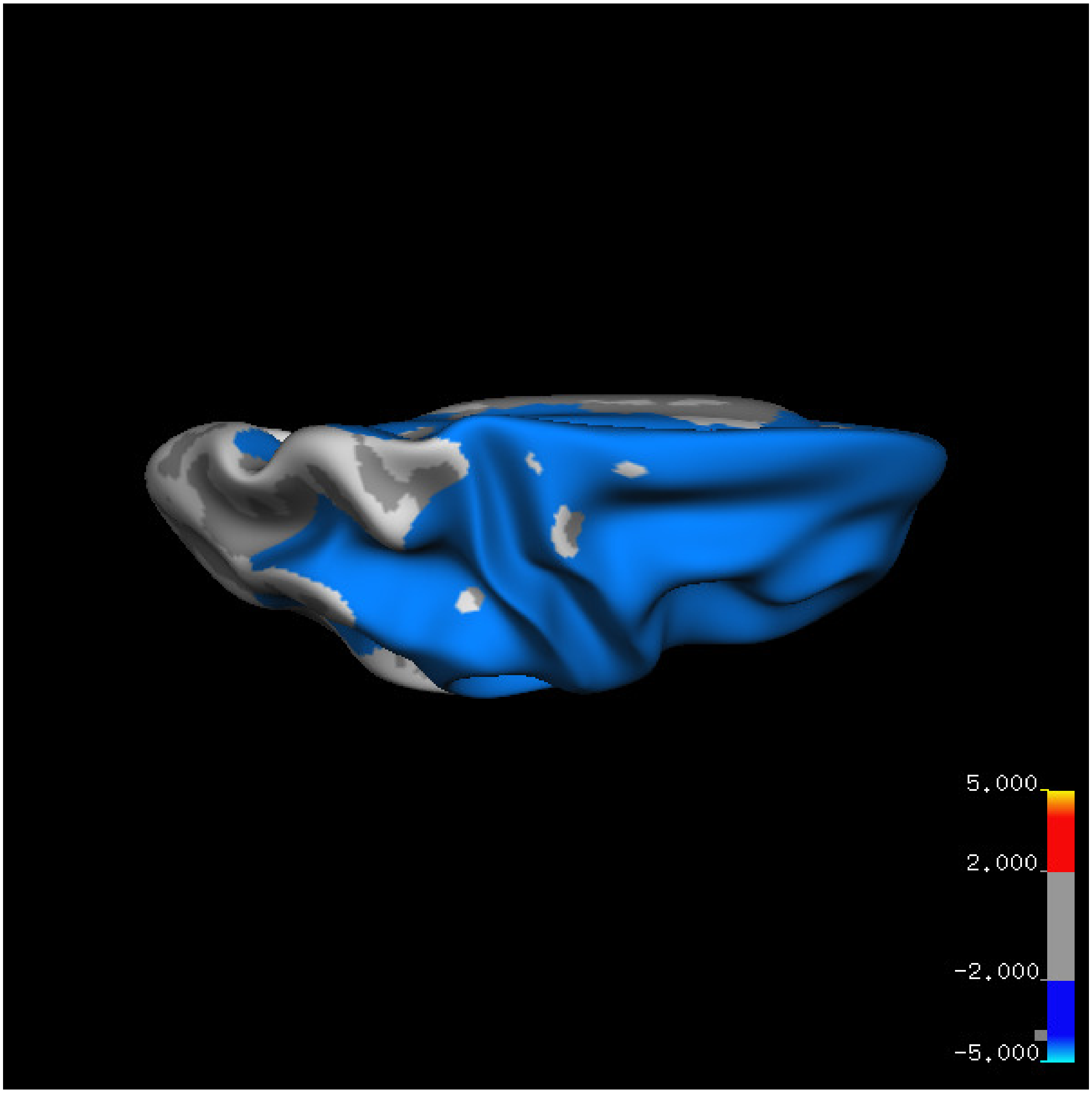
5.000
2.000
-2.000
-5.000

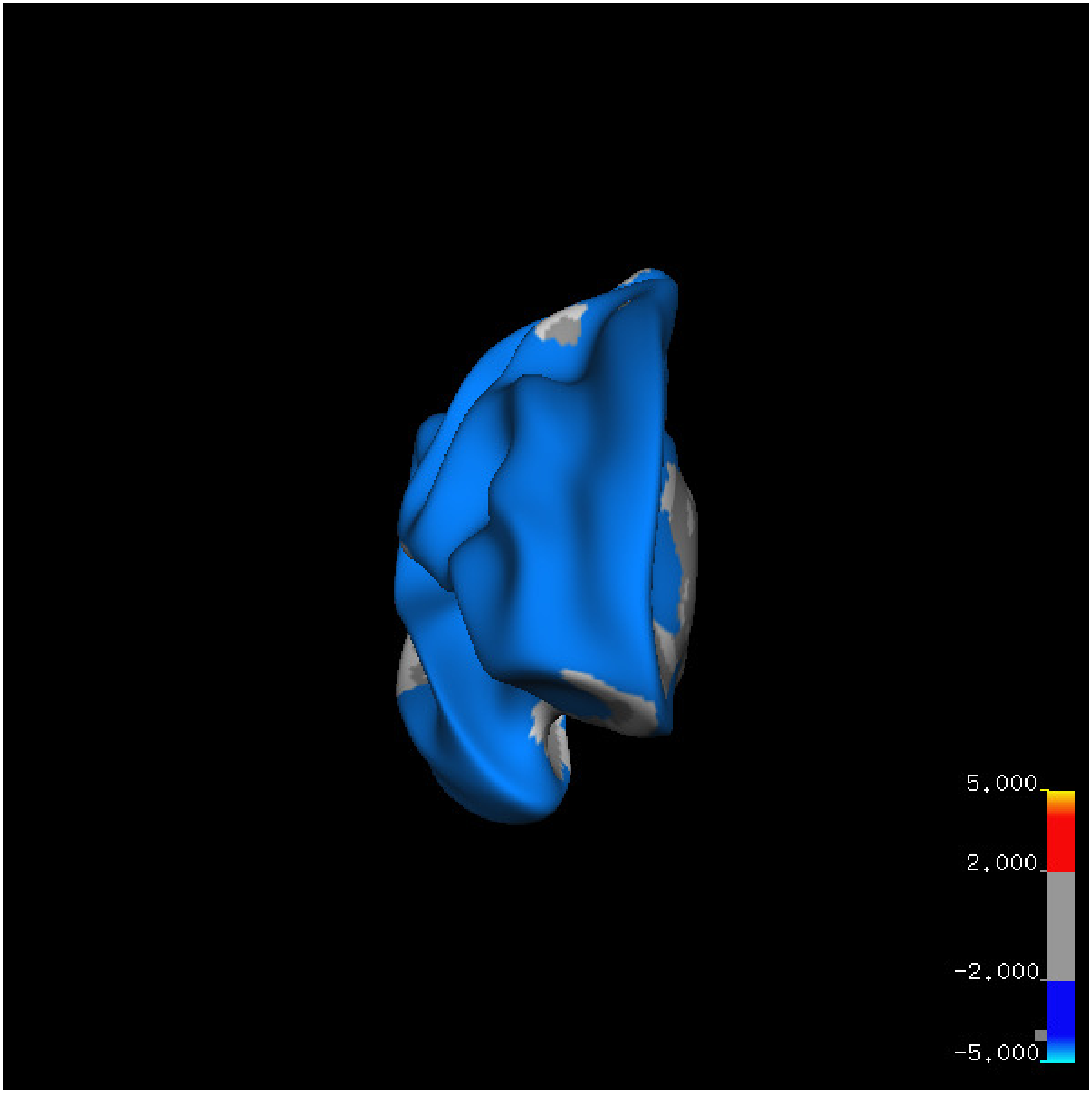
5.000
2.000
-2.000
-5.000

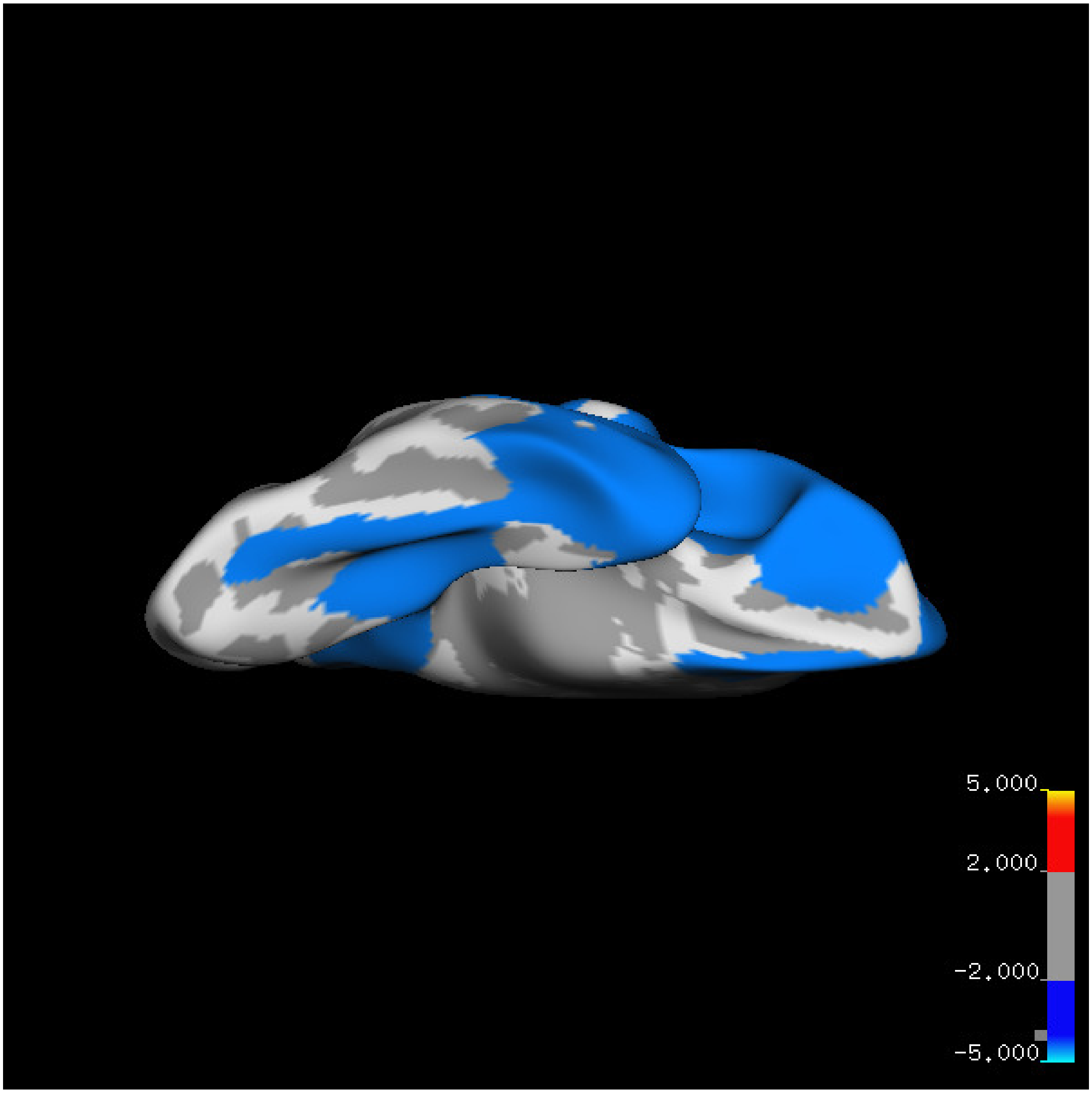
5.000
2.000
-2.000
-5.000

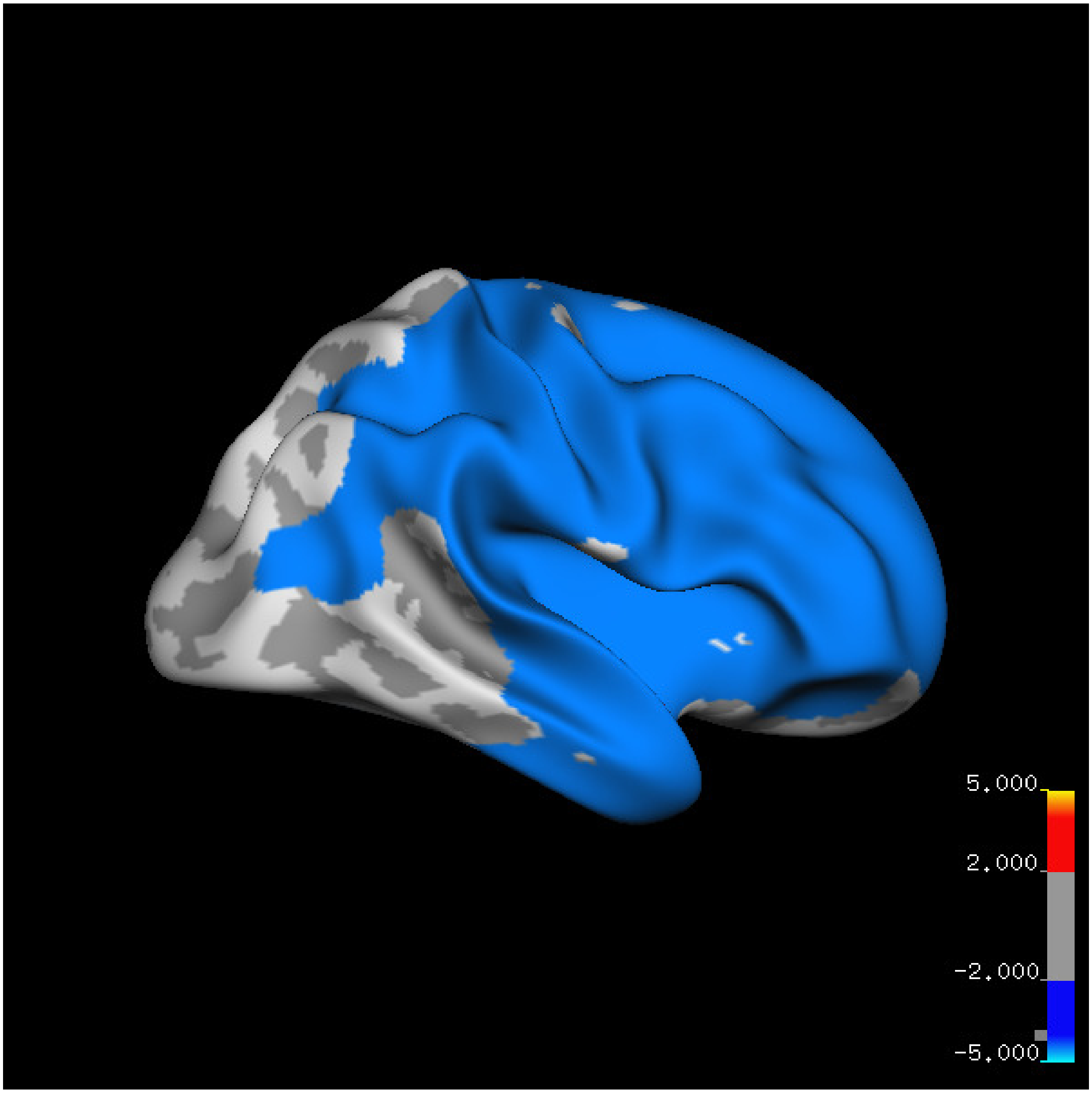

5.000
2.000
-2.000
-5.000

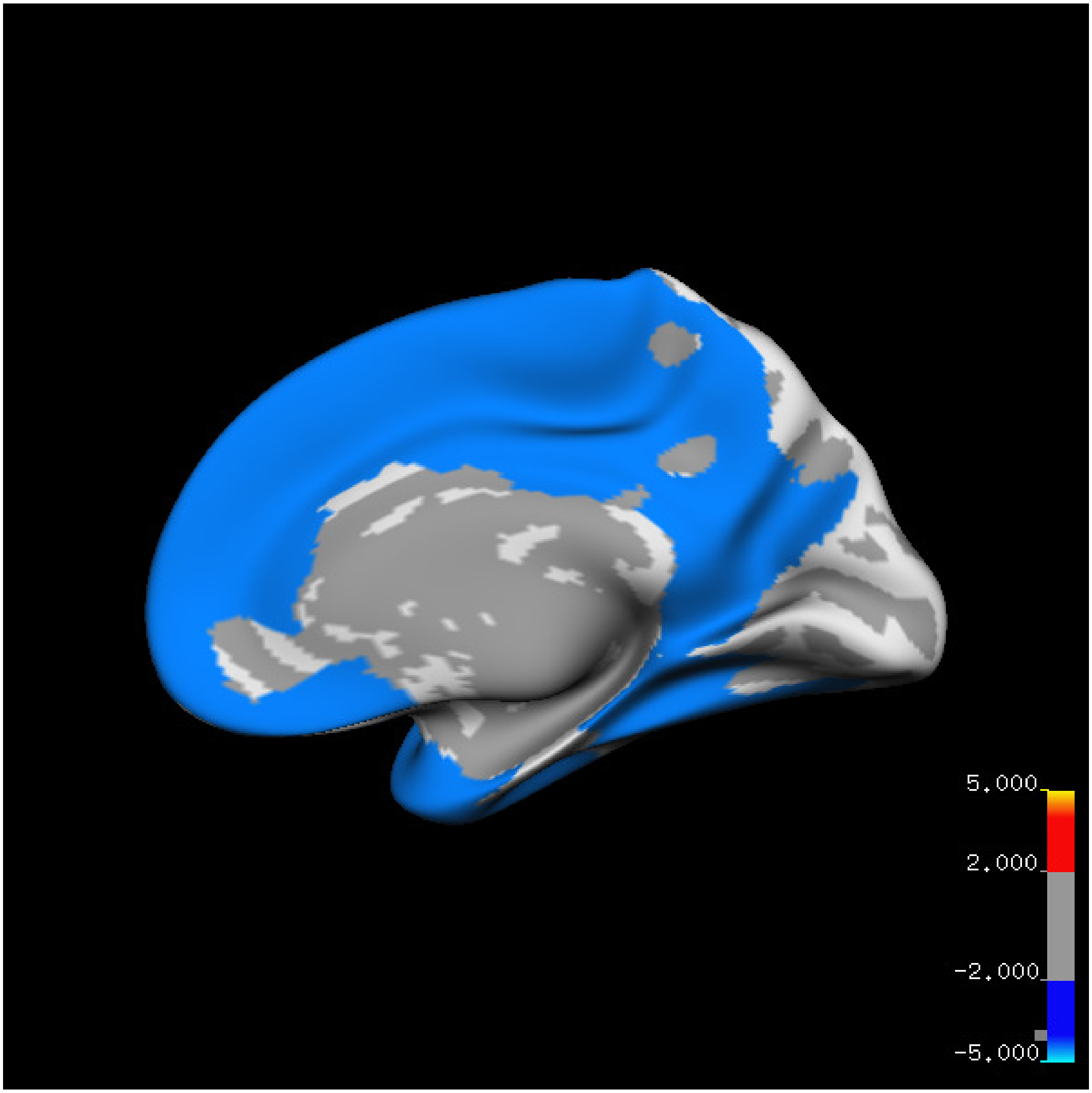

5.000
2.000
-2.000
-5.000

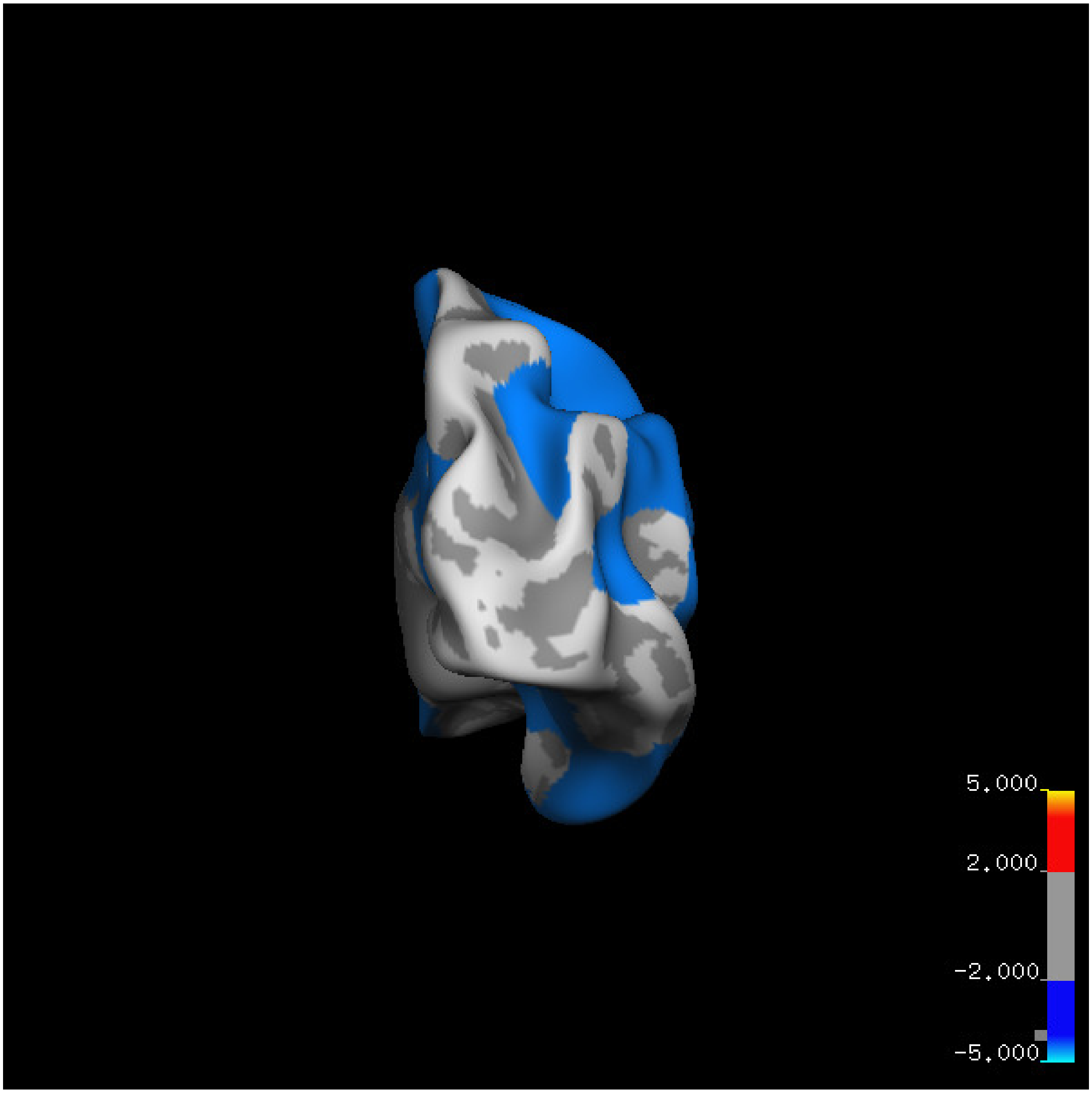
5.000
2.000
-2.000
-5.000

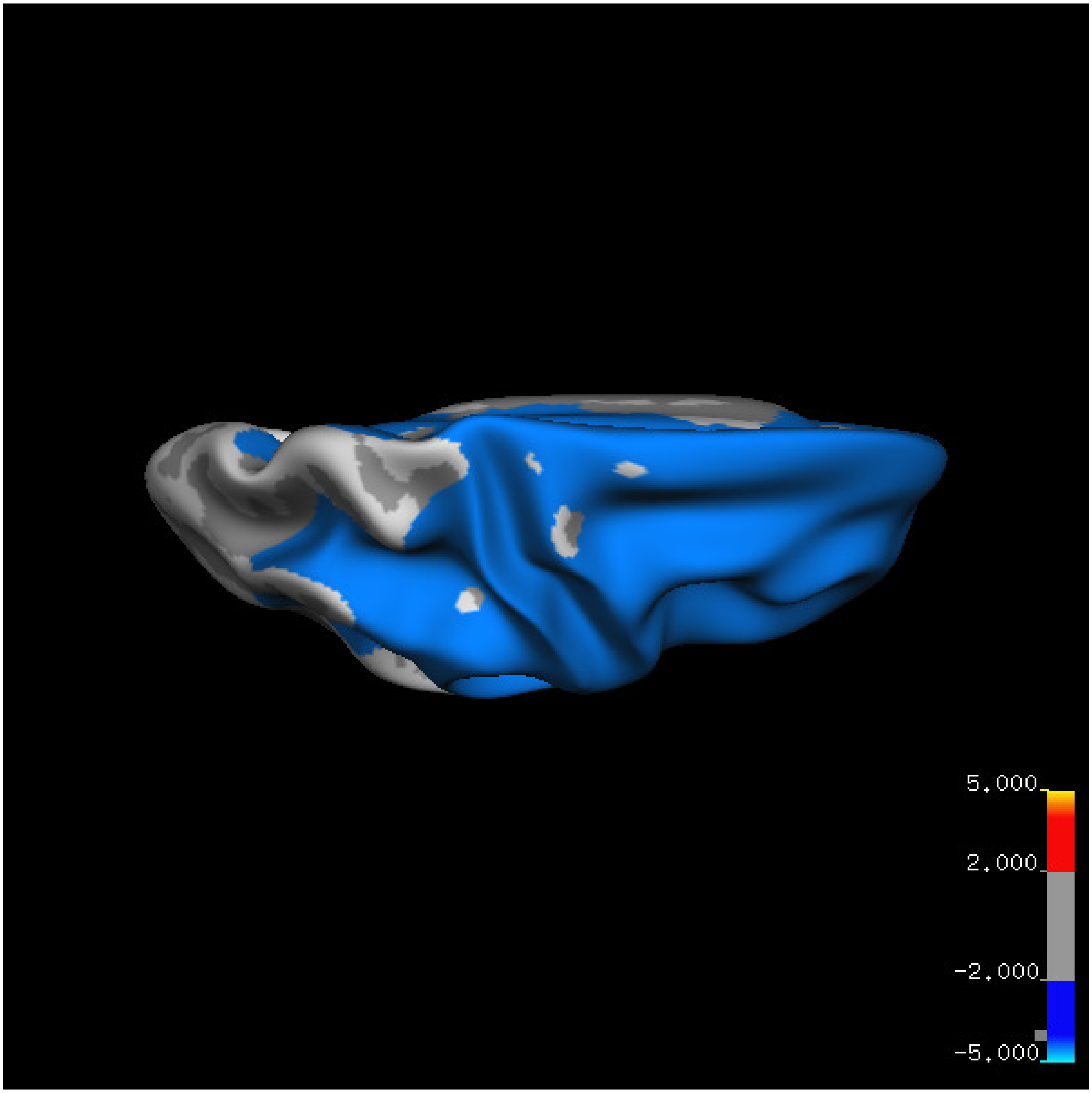


Cluster summary is shown below.

```
## # Cluster Growing Summary (mri_surfcluster)

## # $Id: mri_surfcluster.c,v 1.57.2.3 2016/11/17 18:19:42 zkaufman Exp $

## # $Id: mrisurf.c,v 1.781.2.6 2016/12/27 16:47:14 zkaufman Exp $

## # CreationTime 2021/10/04-13:06:16-GMT

## # cmdline mri_surfcluster.bin --in surface_analyses/results/cluster_comparison/fsaverage6.sm15.rh/thi

## # cwd /gpfs/projects02/p274/cluster/projects/p003-sleep_b

## # sysname  Linux
```

```
## # hostname c1-19
## # machine  x86_64
## # FixVertexAreaFlag 1
## # FixSurfClusterArea 1
## #
## # Input      surface_analyses/results/cluster_comparison/fsaverage6.sm15.rh/thickness.above60.above_
## # Frame Number      0
## # srcsubj fsaverage6
## # hemi rh
## # surface white
## # group_avg_surface_area 85131.2
## # group_avg_vtxarea_loaded 1
## # annot aparc
## # SUBJECTS_DIR /cluster/projects/p274/tools/mri/freesurfer/current/subjects
## # SearchSpace_mm2 79329.9
## # SearchSpace_vtx 37471
## # Bonferroni 0
## # Minimum Threshold 1.3
## # Maximum Threshold infinity
## # Threshold Sign    abs
## # AdjustThreshWhenOneTail 1
## # CW PValue Threshold: 0.01
## # Area Threshold    0 mm^2
## # CSD thresh  1.300000
## # CSD nreps    10000
## # CSD simtype  null-z
## # CSD contrast NA
```

```
## # CSD confint  90.000000
## # Overall max 0.89553 at vertex 9431
## # Overall min -10.8348 at vertex 6130
## # NClusters          1
## # FixMNI = 0
## #
## # ClusterNo  Max   VtxMax   Size(mm^2)  MNIX   MNIY   MNIZ    CWP    CWPLow    CWPHi   NVtxs    Wght
##    1      -10.835    6130  55051.37     22.4    4.6   48.6  0.00010  0.00000  0.00020  27434   -9218
```

**Sleep below mean** Cluster corrected significance plots are shown below.

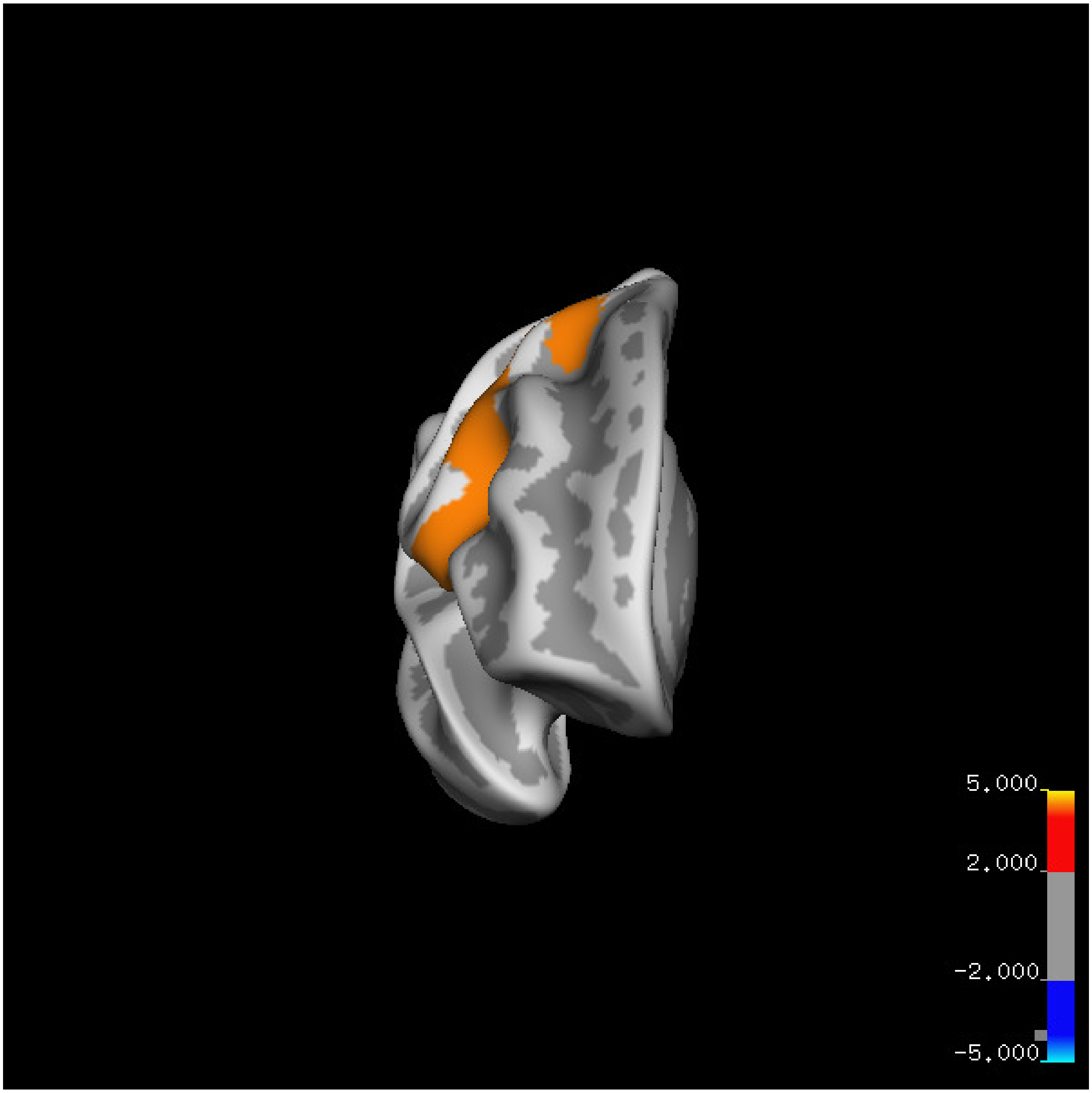

5.000
2.000
-2.000
-5.000

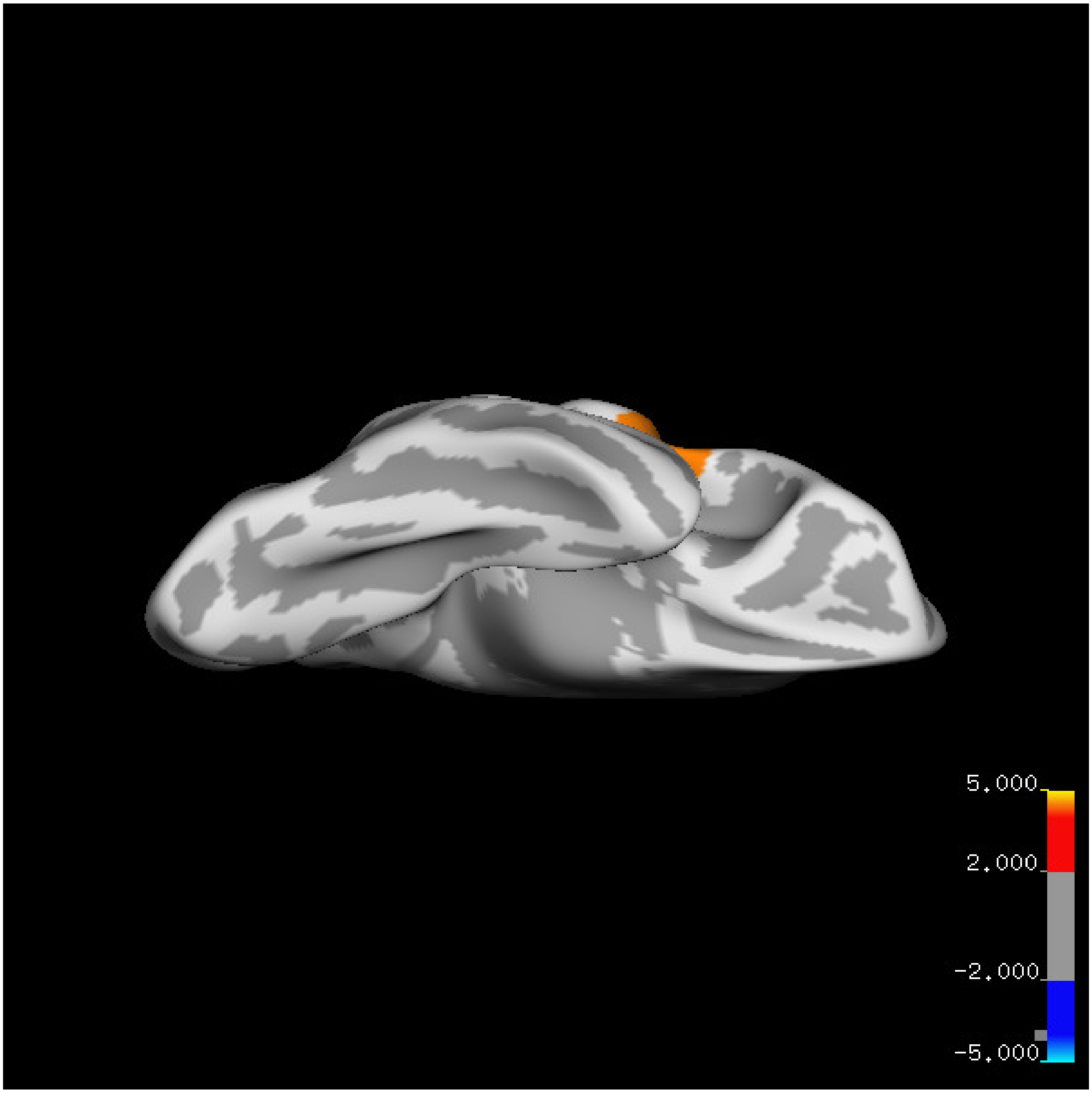

5.000
2.000
-2.000
-5.000

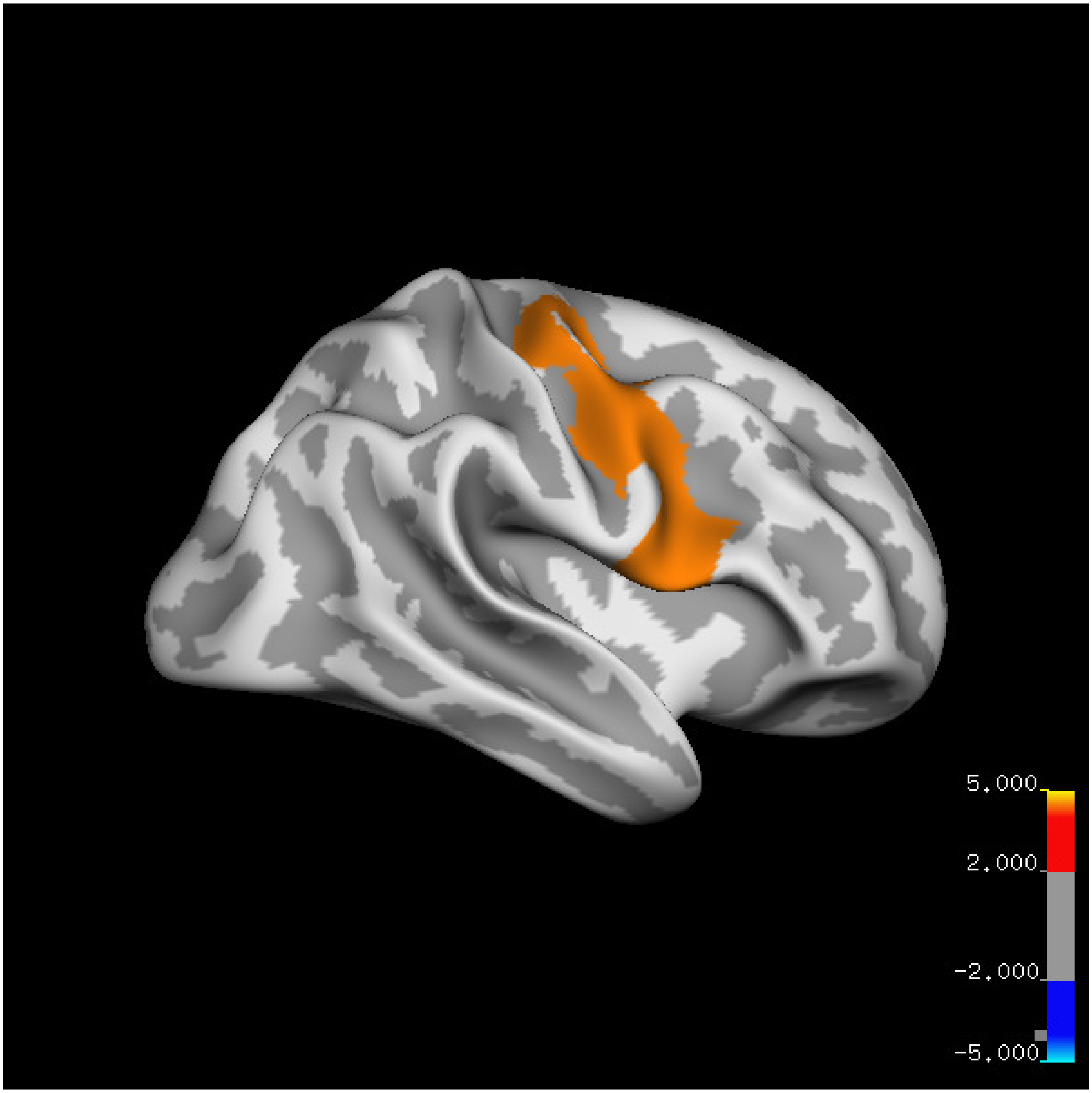
5.000
2.000
-2.000
-5.000

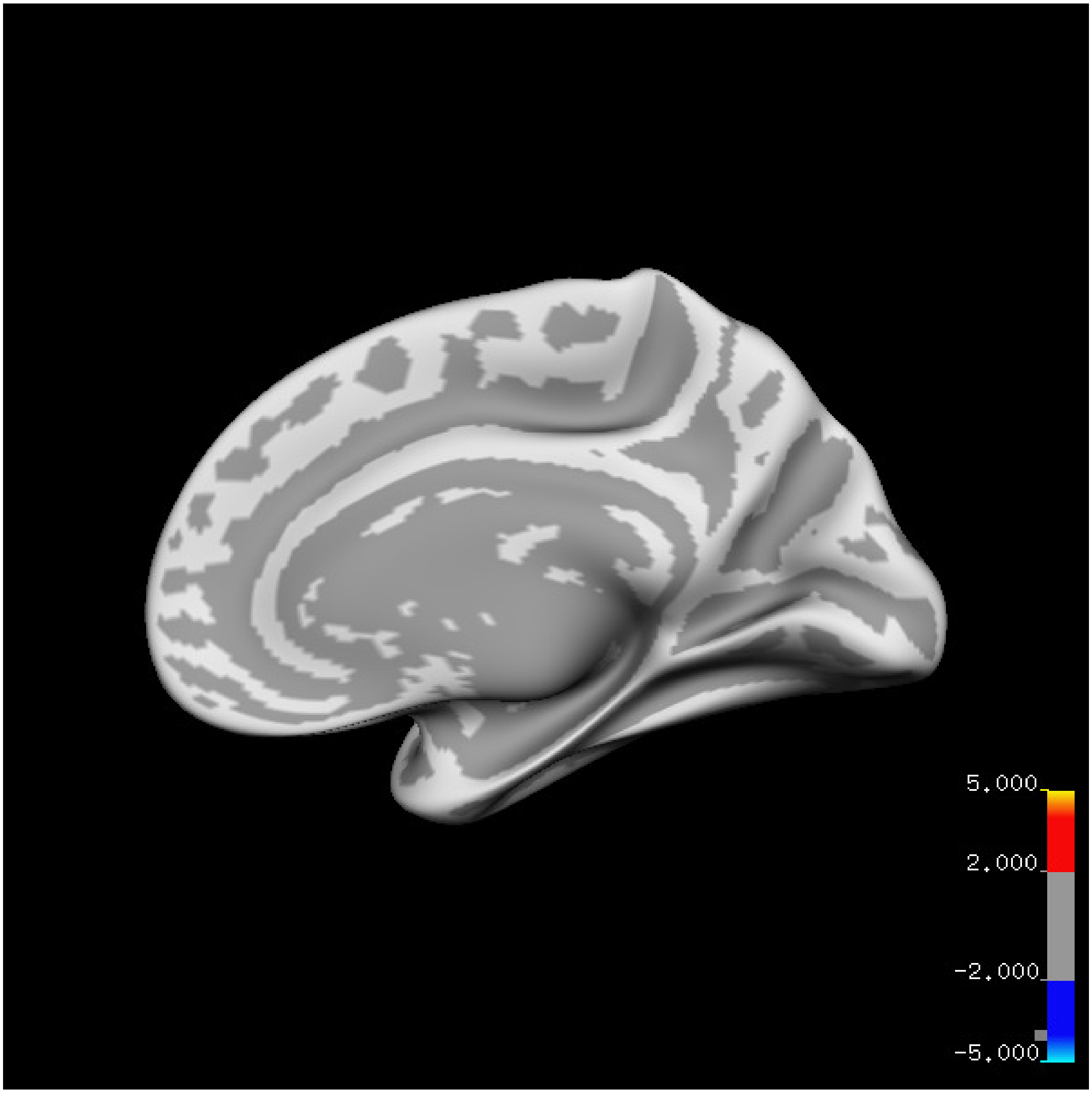
5.000
2.000
-2.000
-5.000

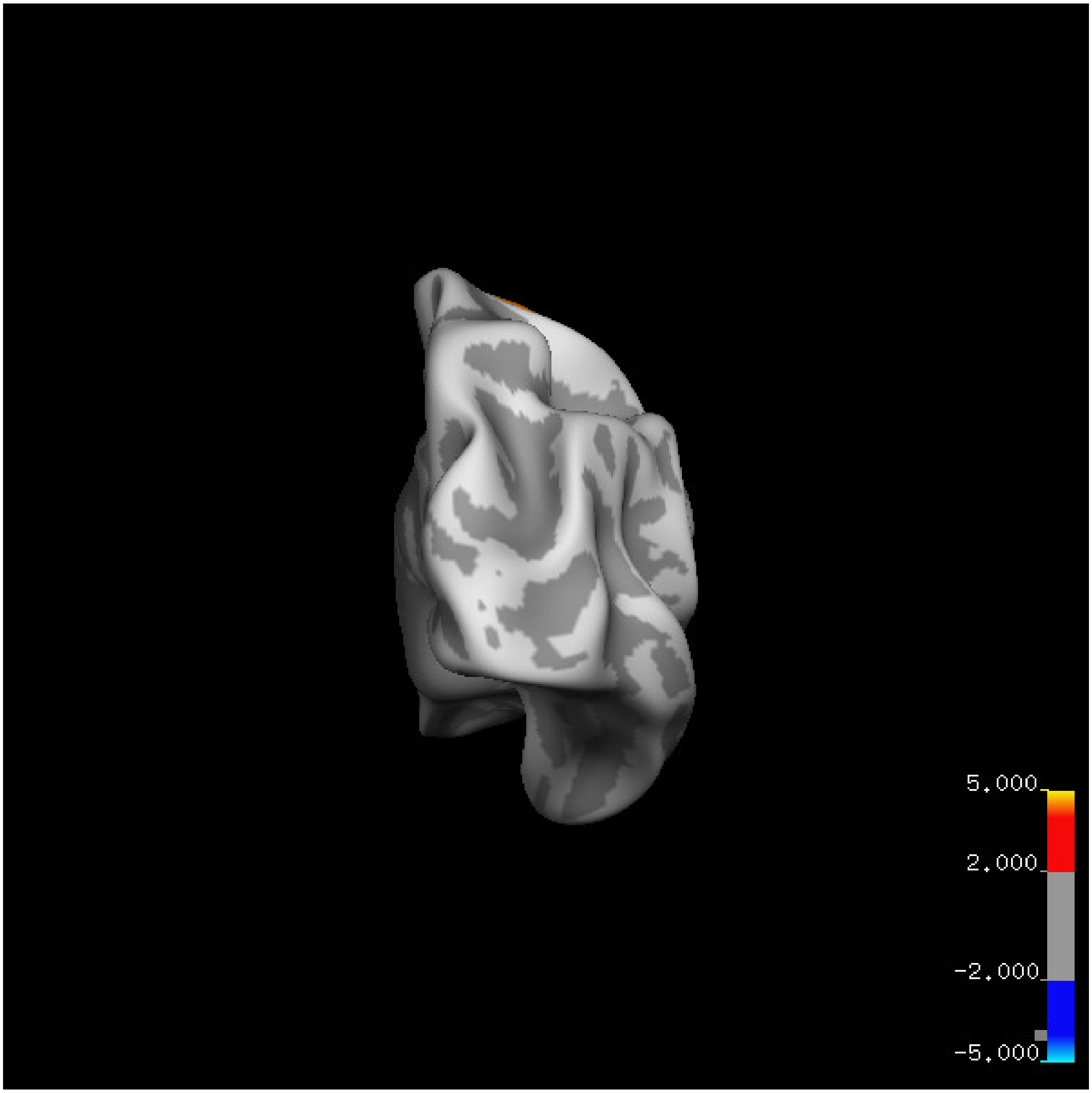
5.000
2.000
-2.000
-5.000

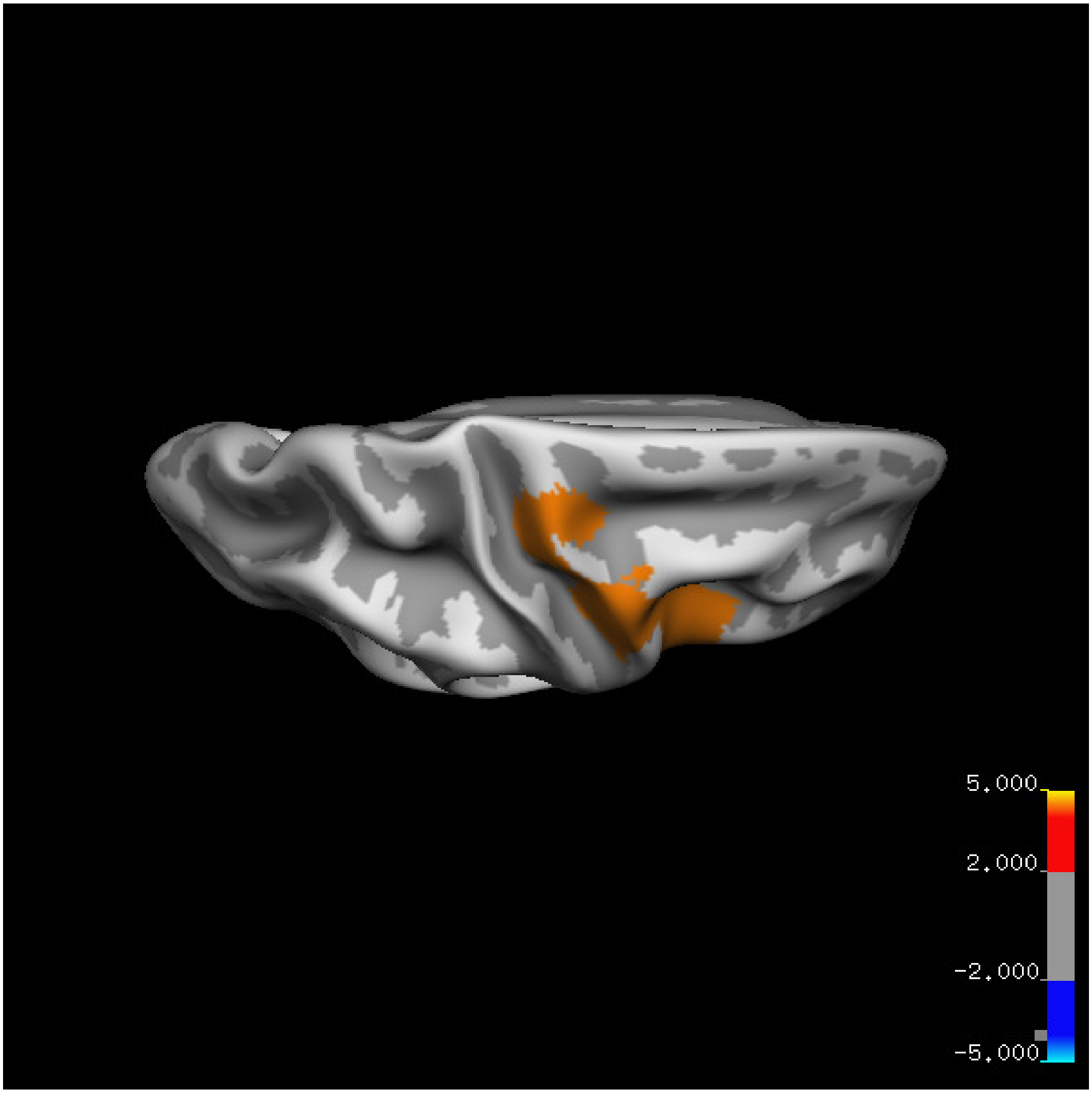

5.000
2.000
-2.000
-5.000

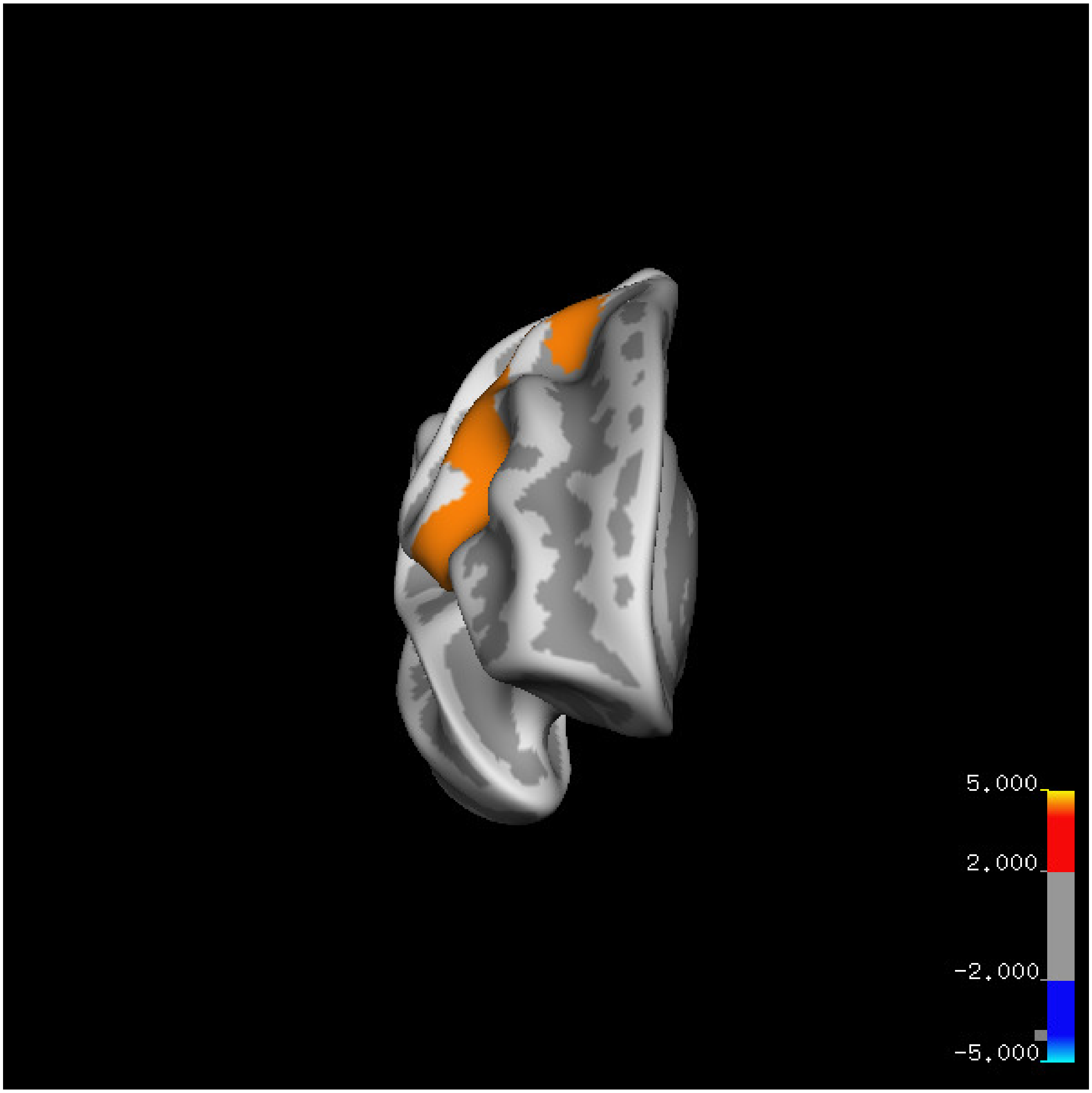
5.000
2.000
-2.000
-5.000

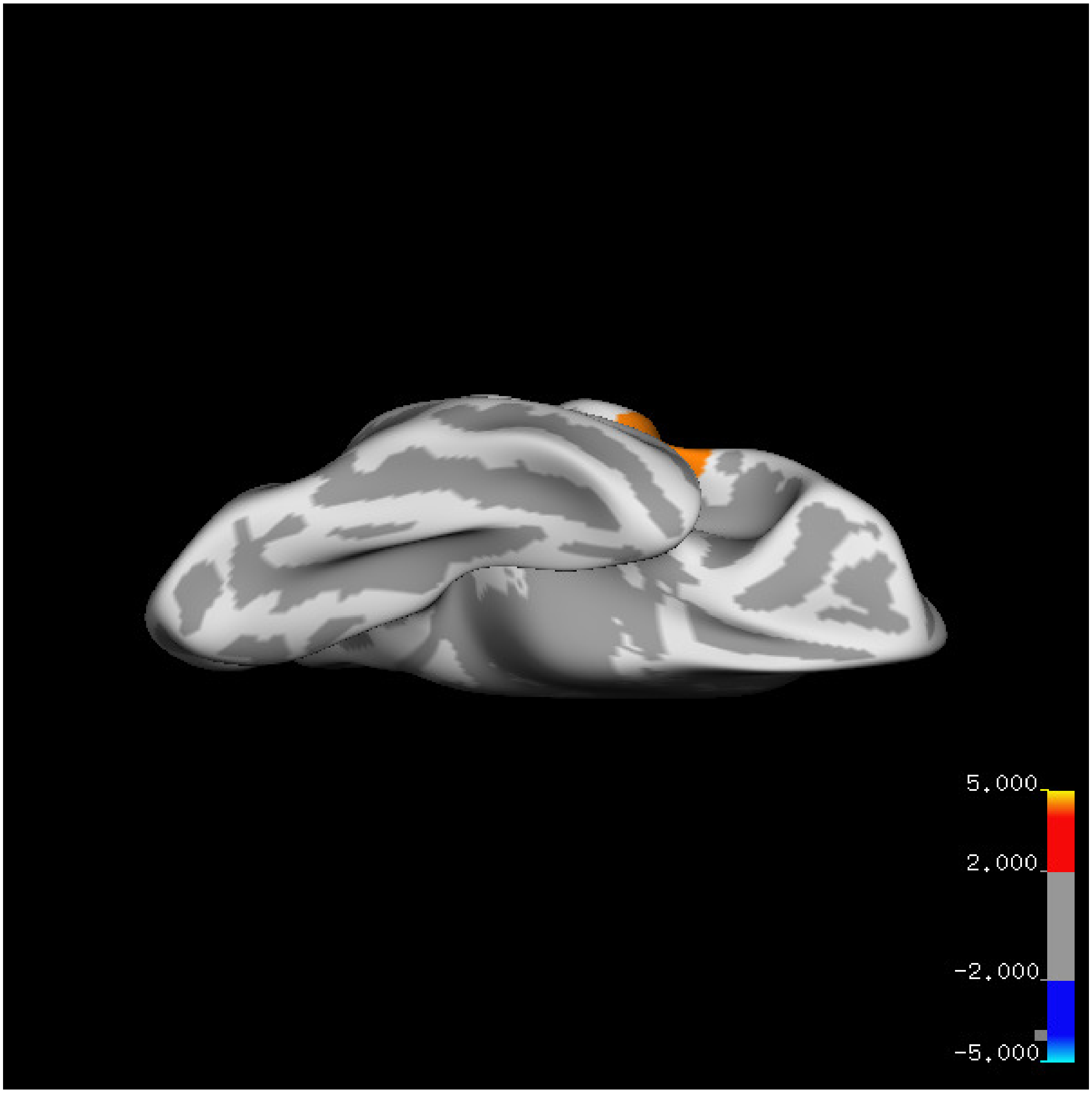
5.000
2.000
-2.000
-5.000

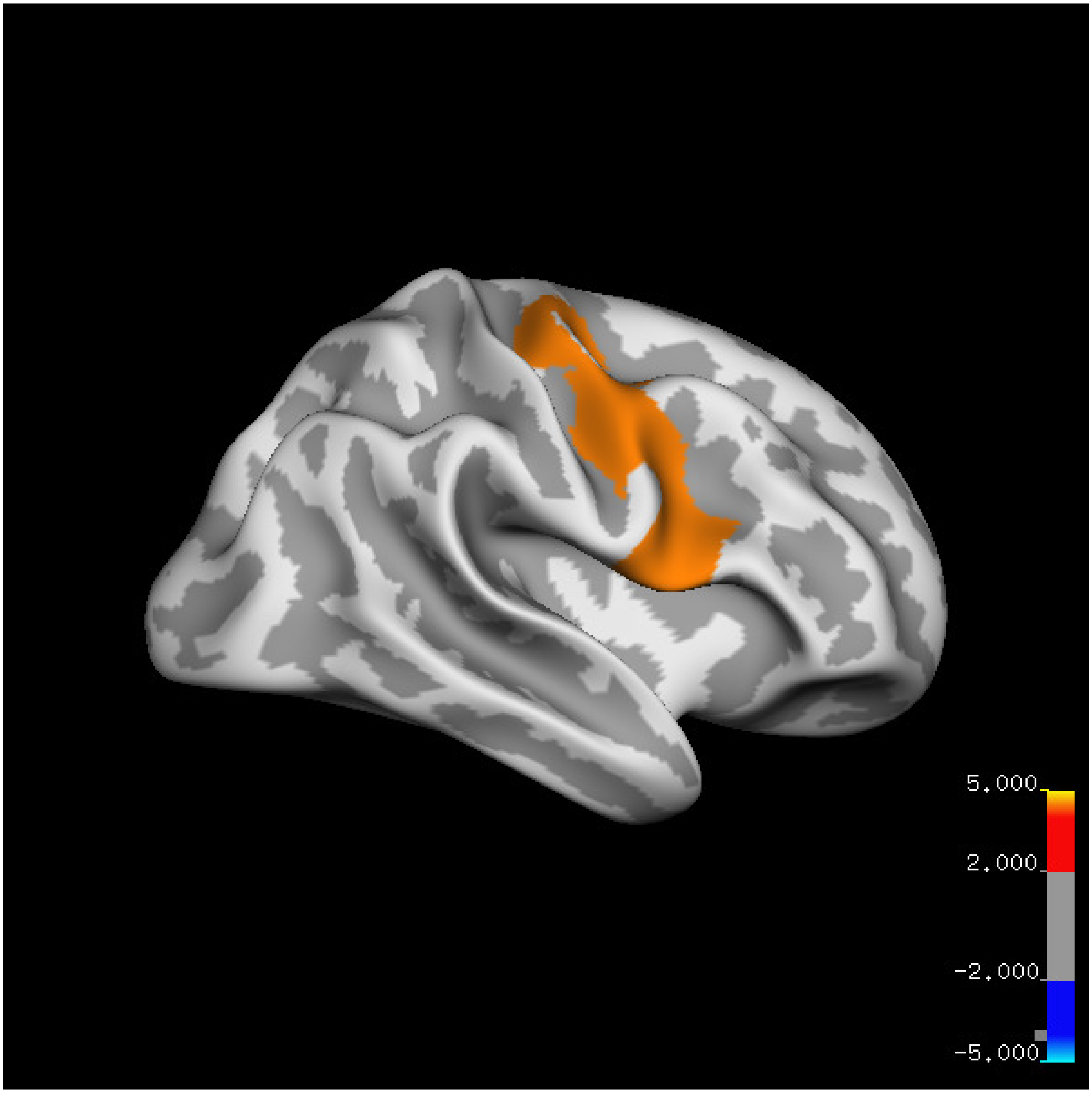
5.000
2.000
-2.000
-5.000

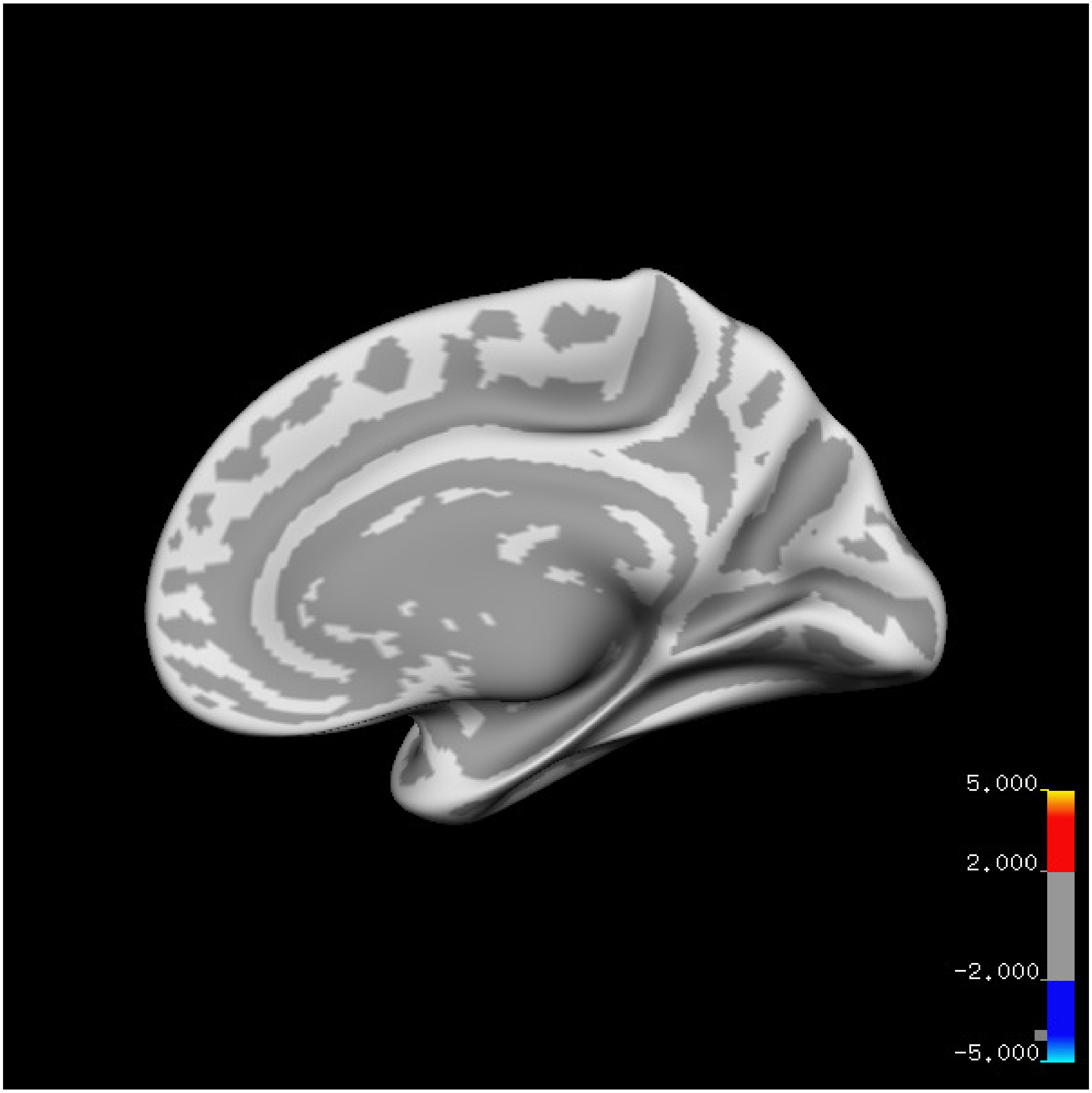
5.000
2.000
-2.000
-5.000

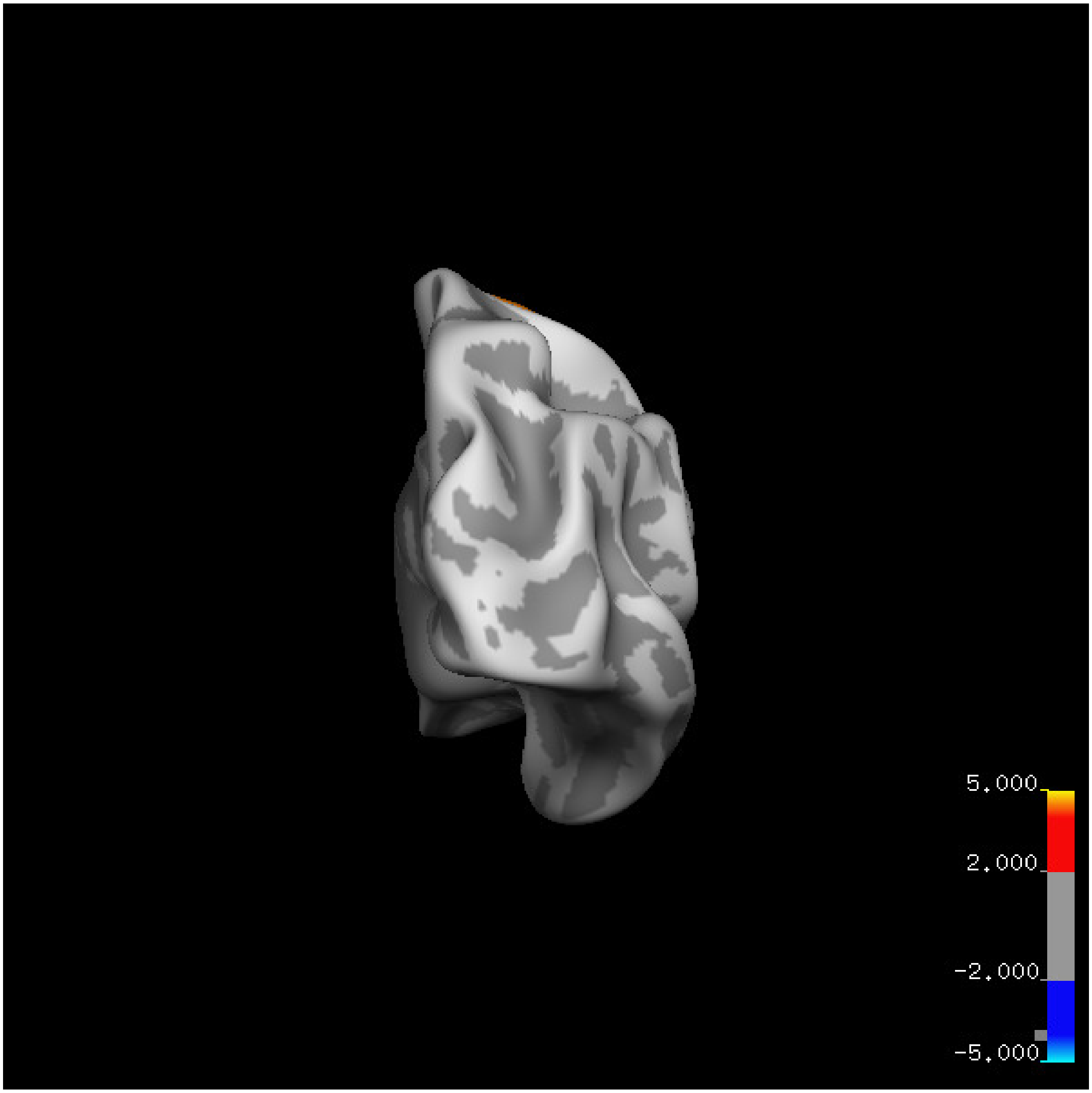
5.000
2.000
-2.000
-5.000

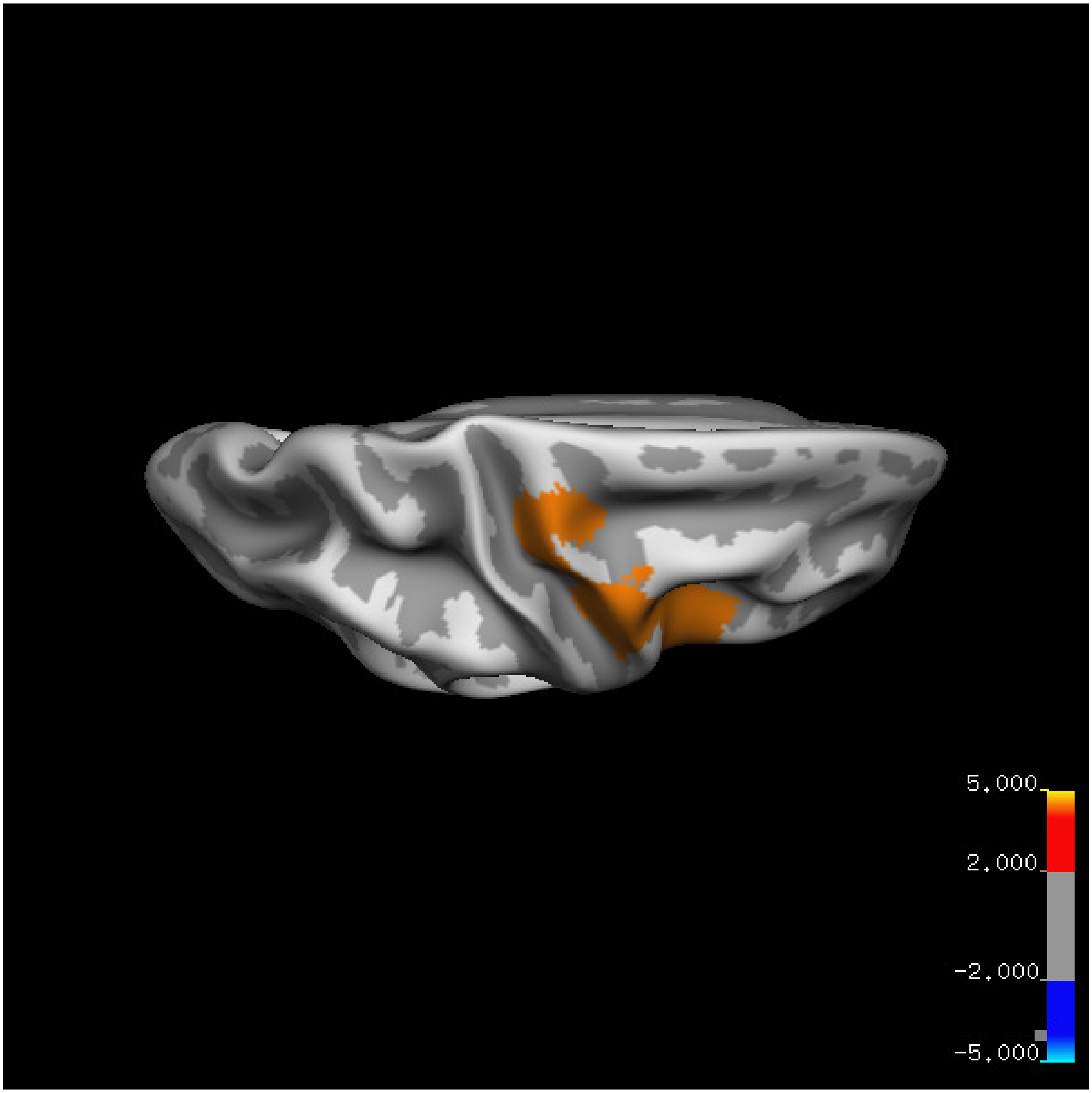


Cluster summary is shown below.

```
## # Cluster Growing Summary (mri_surfcluster)

## # $Id: mri_surfcluster.c,v 1.57.2.3 2016/11/17 18:19:42 zkaufman Exp $

## # $Id: mrisurf.c,v 1.781.2.6 2016/12/27 16:47:14 zkaufman Exp $

## # CreationTime 2021/10/04-13:56:08-GMT

## # cmdline mri_surfcluster.bin --in surface_analyses/results/cluster_comparison/fsaverage6.sm15.rh/th

## # cwd /gpfs/projects02/p274/cluster/projects/p003-sleep_b

## # sysname  Linux
```

```
## # hostname c1-19
## # machine  x86_64
## # FixVertexAreaFlag 1
## # FixSurfClusterArea 1
## #
## # Input      surface_analyses/results/cluster_comparison/fsaverage6.sm15.rh/thickness.above60.below_
## # Frame Number      0
## # srcsubj fsaverage6
## # hemi rh
## # surface white
## # group_avg_surface_area 85131.2
## # group_avg_vtxarea_loaded 1
## # annot aparc
## # SUBJECTS_DIR /cluster/projects/p274/tools/mri/freesurfer/current/subjects
## # SearchSpace_mm2 79329.9
## # SearchSpace_vtx 37471
## # Bonferroni 0
## # Minimum Threshold 1.3
## # Maximum Threshold infinity
## # Threshold Sign    abs
## # AdjustThreshWhenOneTail 1
## # CW PValue Threshold: 0.01
## # Area Threshold    0 mm^2
## # CSD thresh  1.300000
## # CSD nreps    10000
## # CSD simtype  null-z
## # CSD contrast NA
```

```
## # CSD confint  90.000000

## # Overall max 4.32648 at vertex 33127

## # Overall min -2.95686 at vertex 5098

## # NClusters          1

## # FixMNI = 0

## #

## # ClusterNo  Max   VtxMax   Size(mm^2)  MNIX   MNIY   MNIZ    CWP    CWPLow    CWPHi   NVtxs    Wght

##    1        4.326   33127   3524.27     50.7   -0.2   40.6  0.00010  0.00000  0.00020   1797     390
```

### Volume

#### Left hemisphere

##### Age 20-60

**Sleep above mean** Cluster corrected significance plots are shown below.

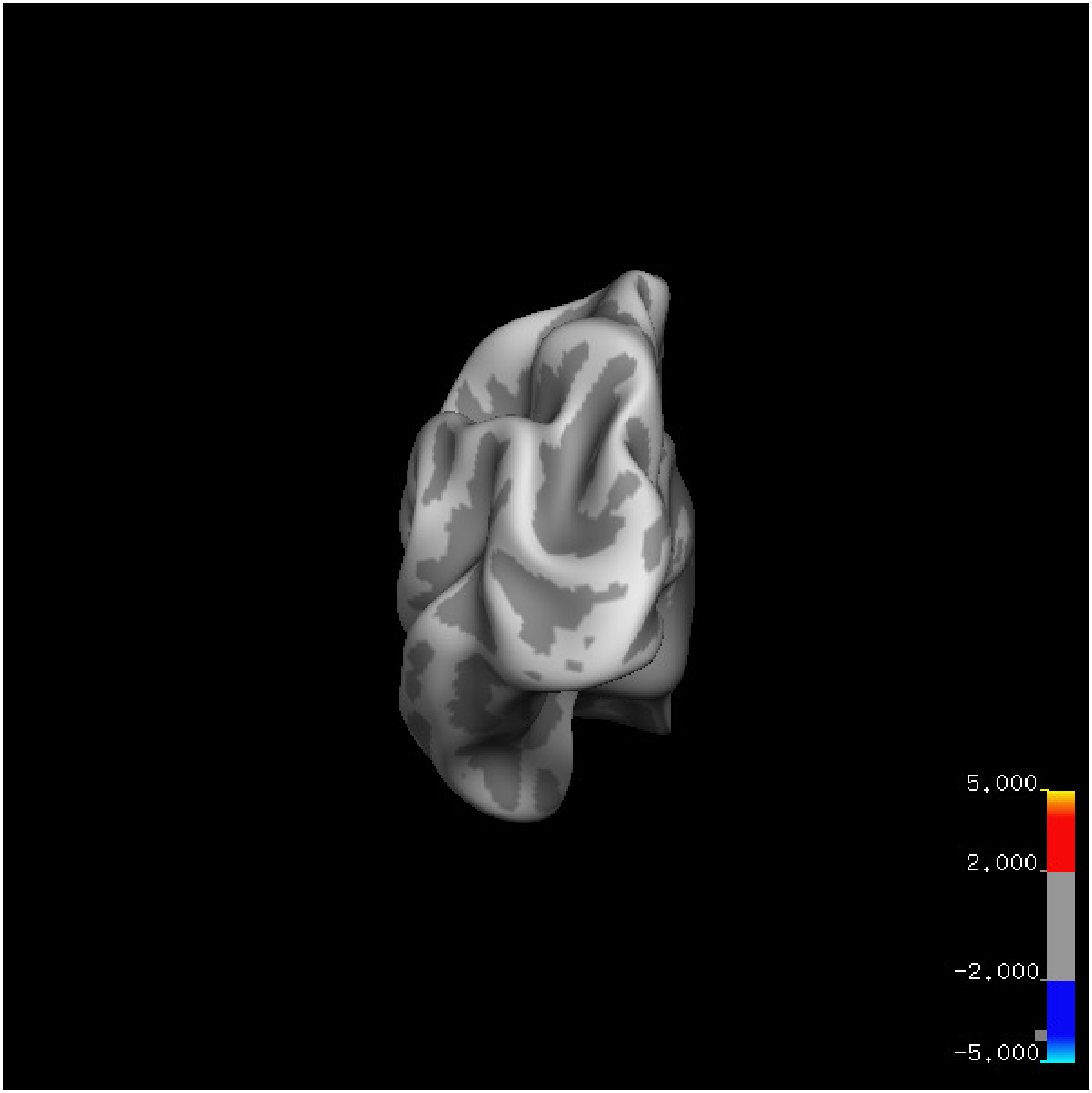
5.000
2.000
-2.000
-5.000

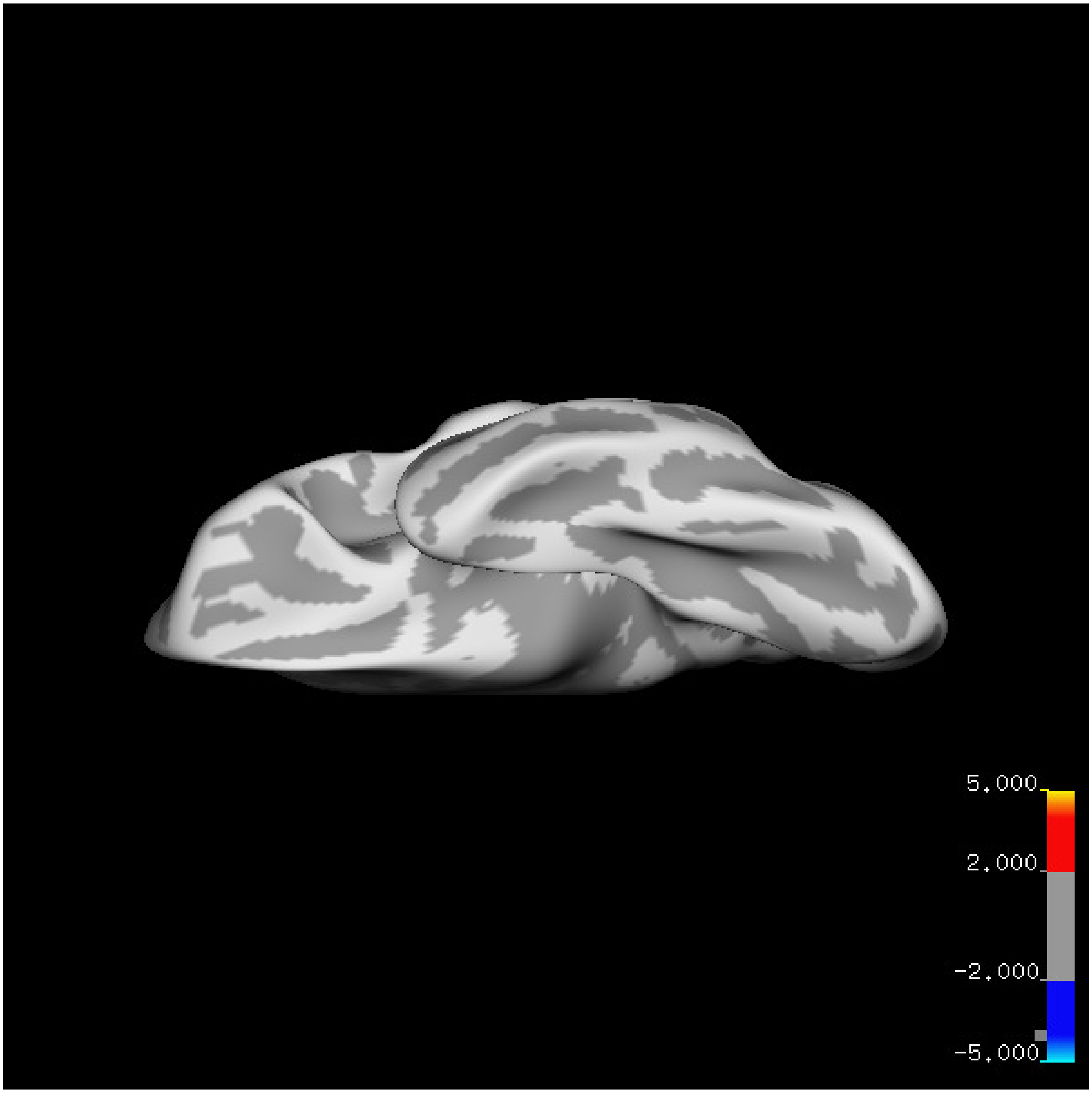
5.000
2.000
-2.000
-5.000

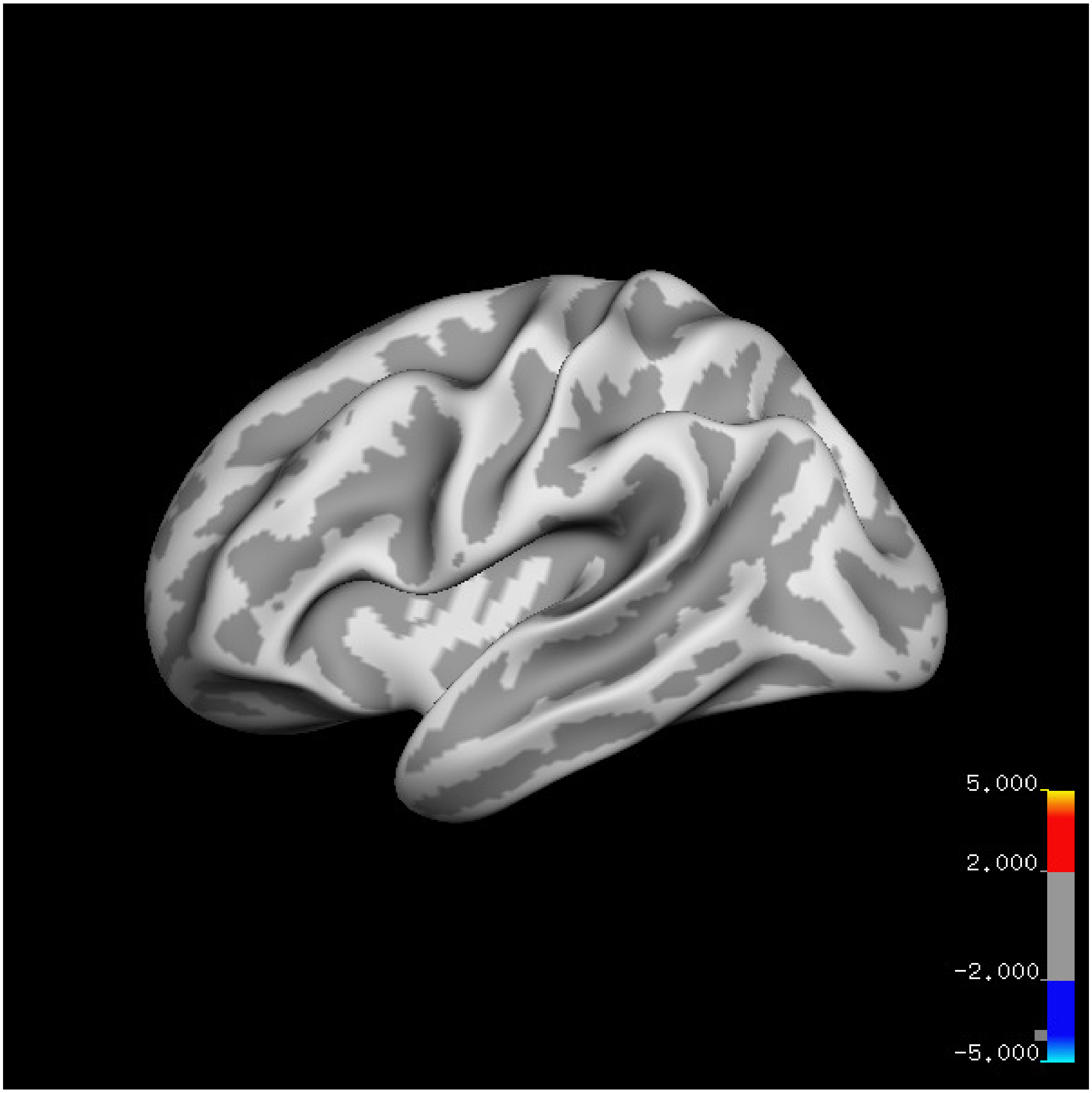
5.000
2.000
-2.000
-5.000

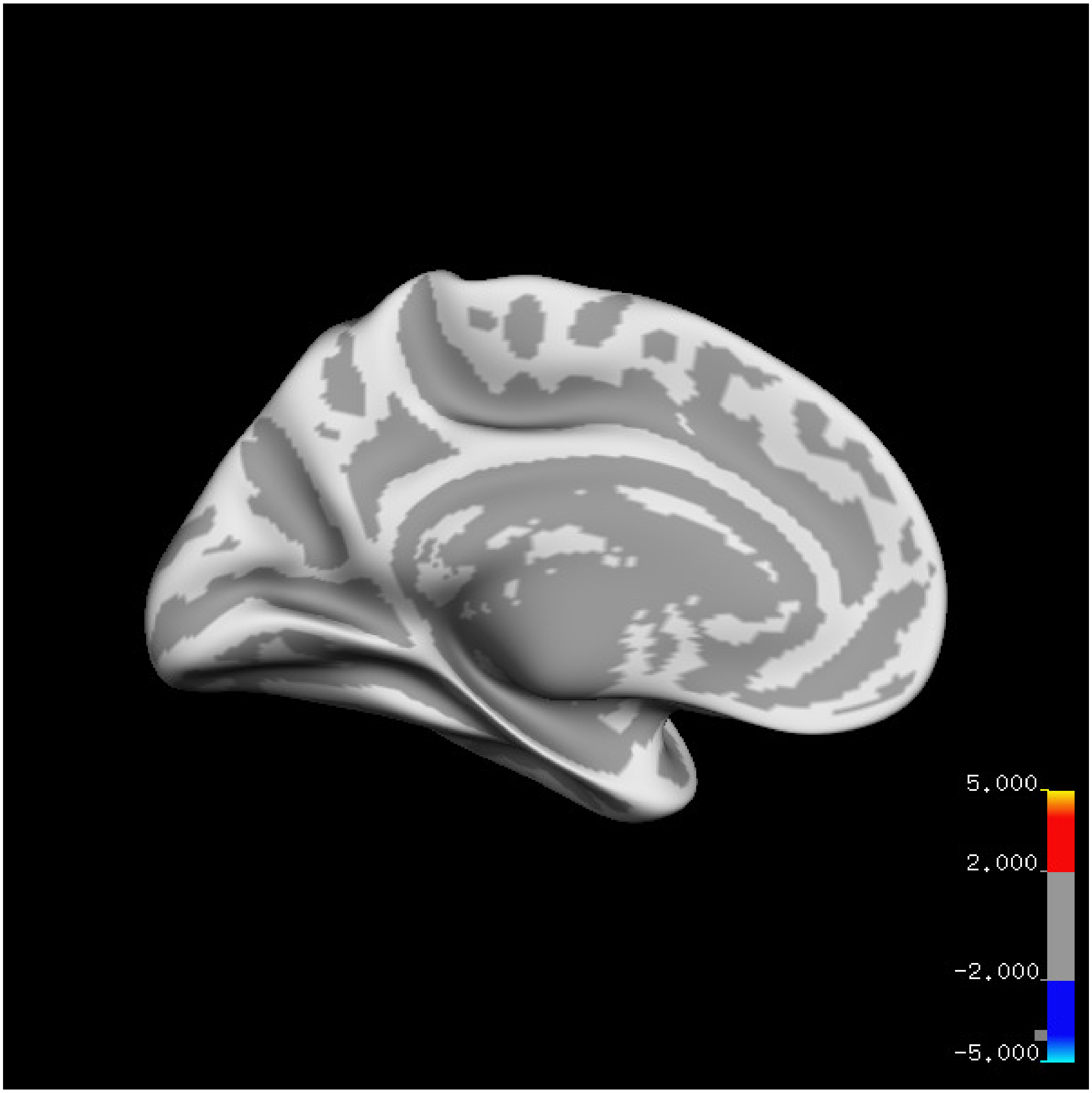

5.000
2.000
-2.000
-5.000

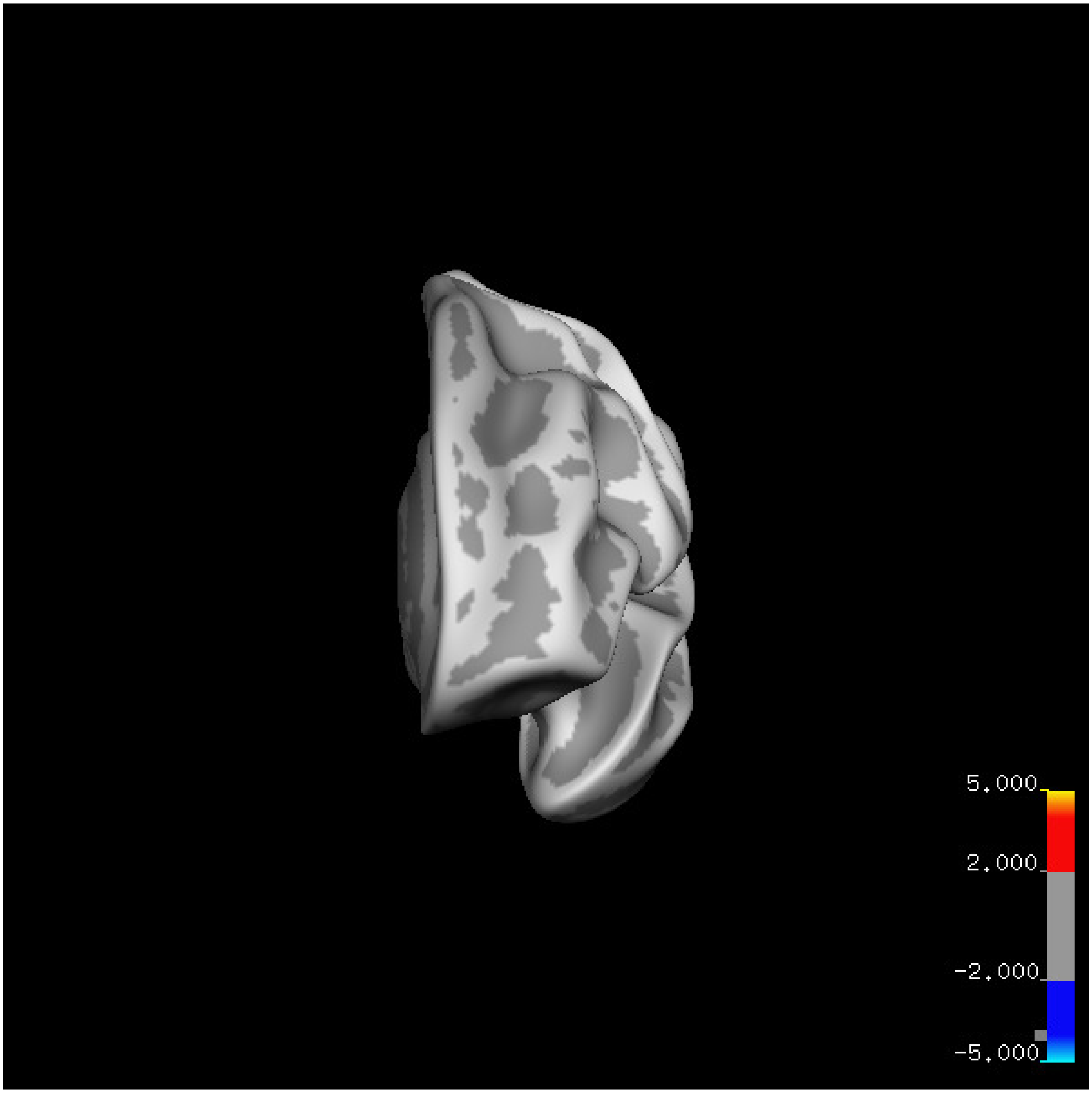
5.000
2.000
-2.000
-5.000

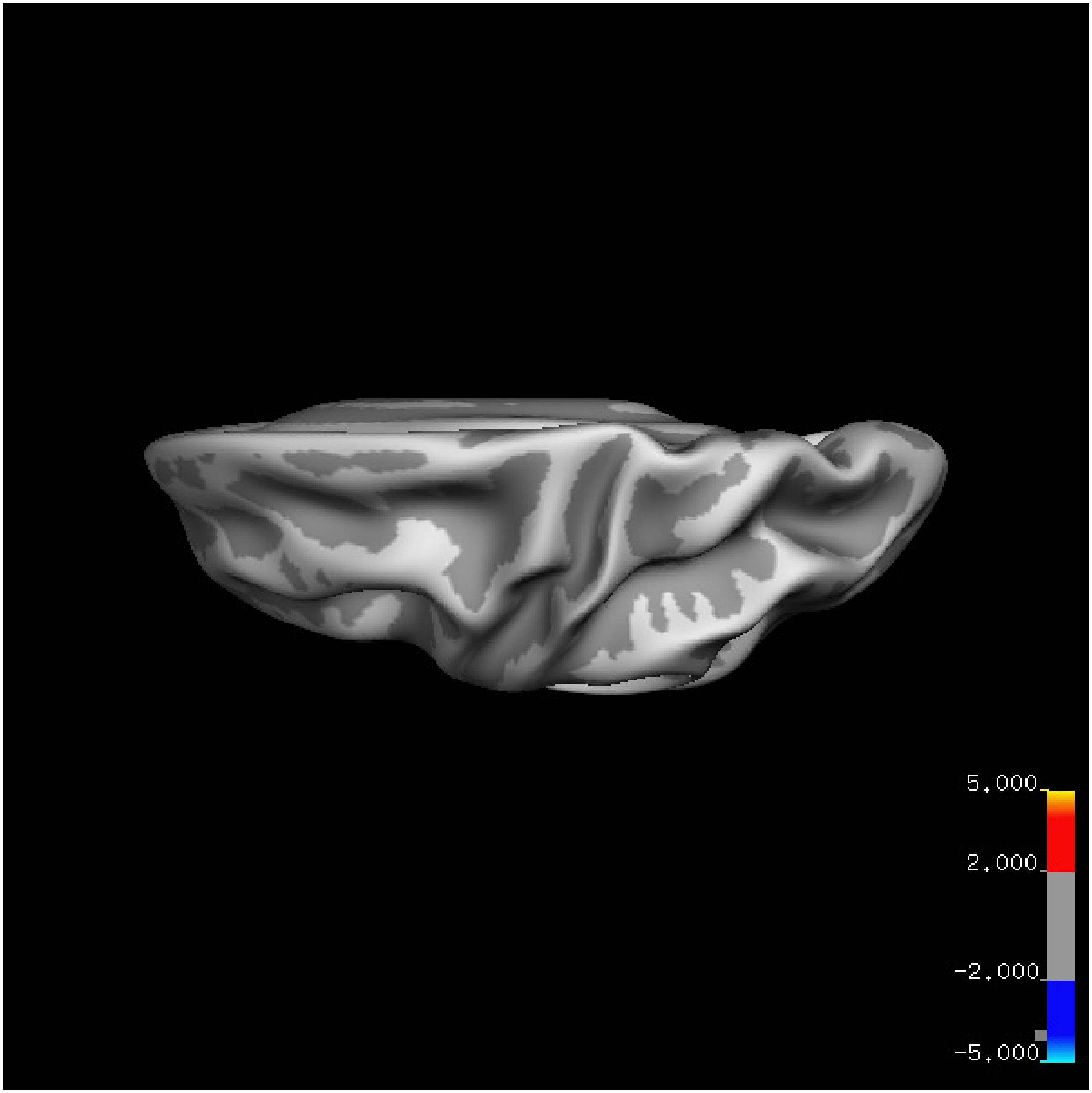
5.000
2.000
-2.000
-5.000

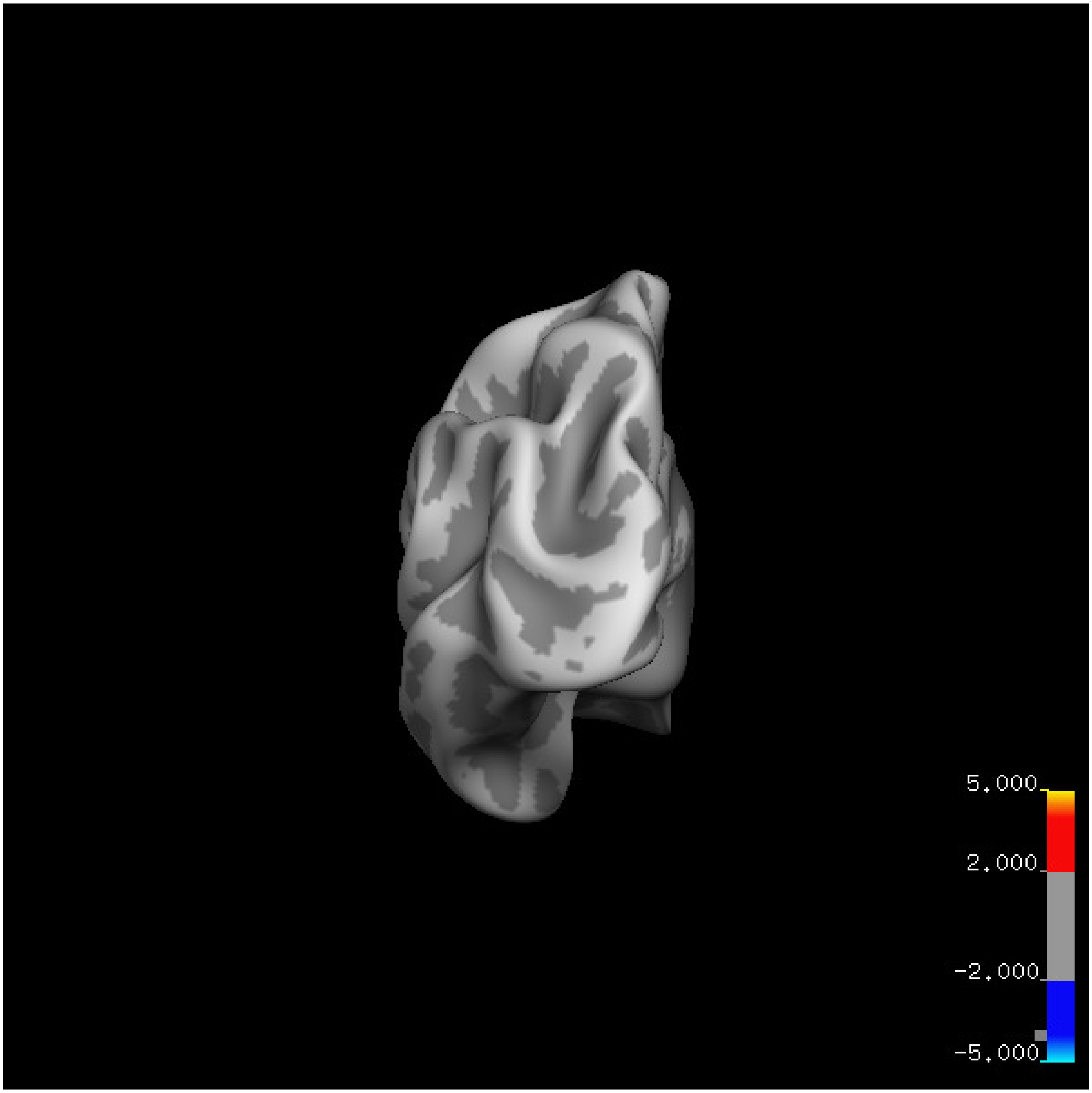
5.000
2.000
-2.000
-5.000

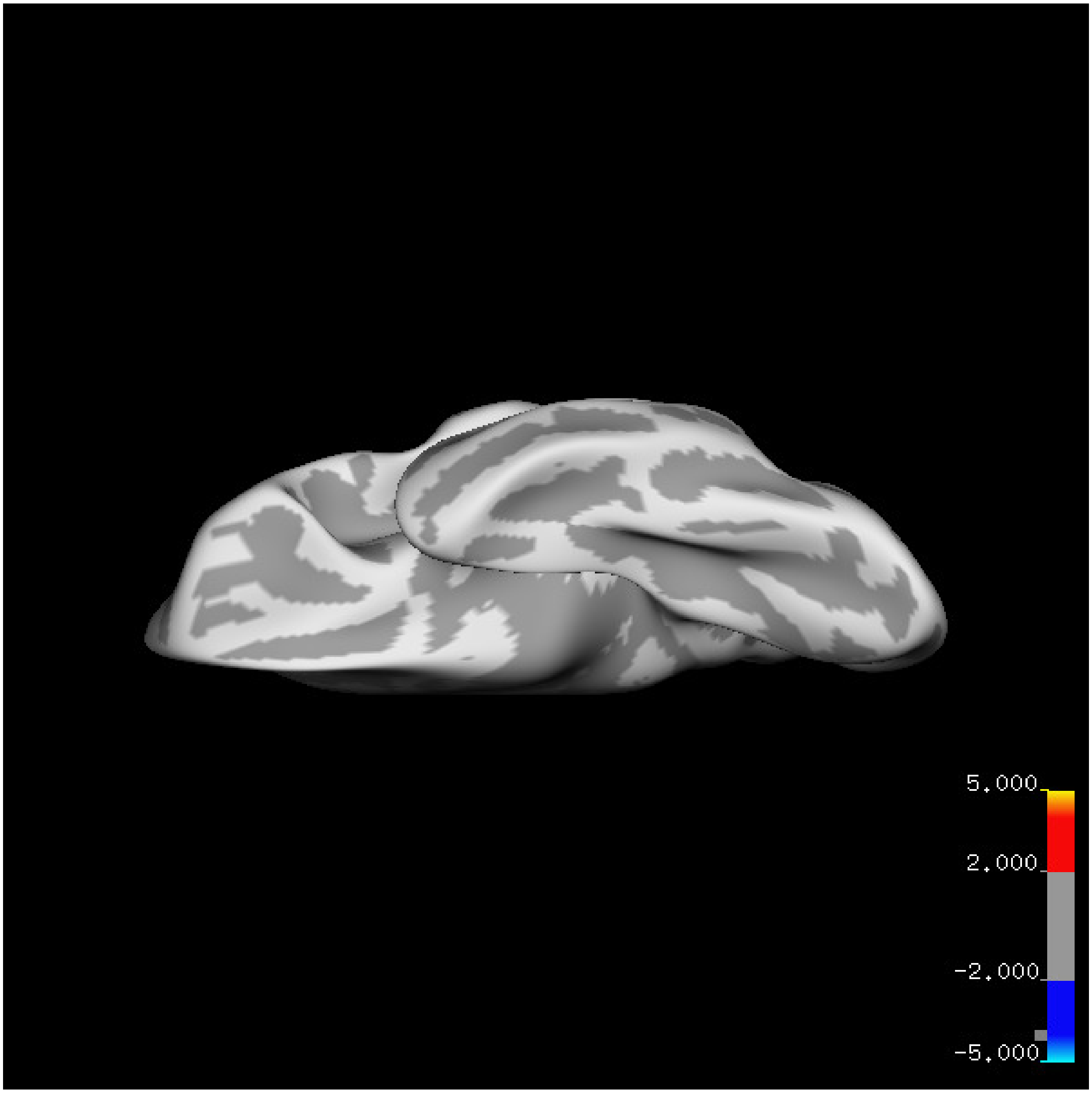
5.000
2.000
-2.000
-5.000

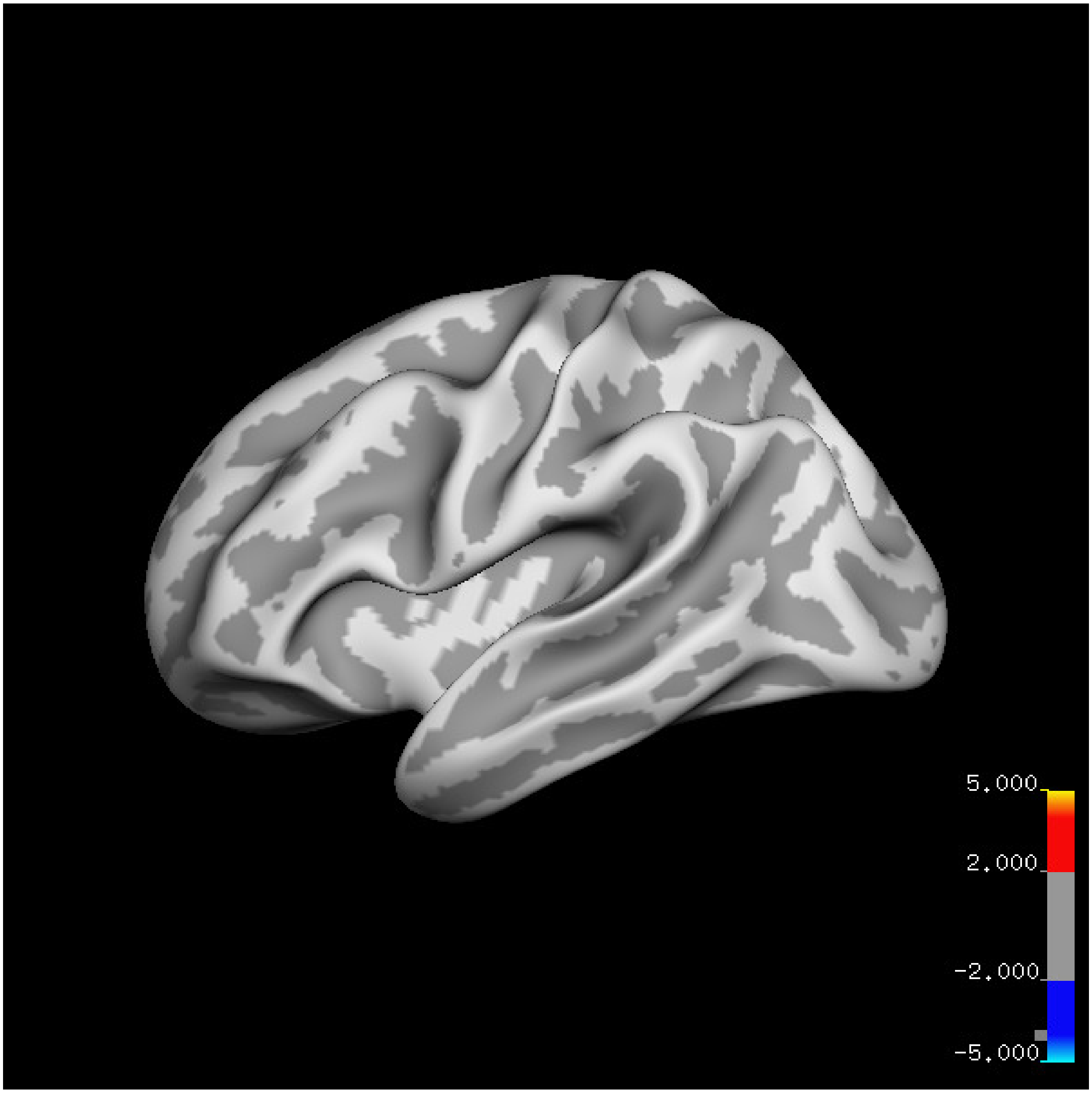
5.000
2.000
-2.000
-5.000

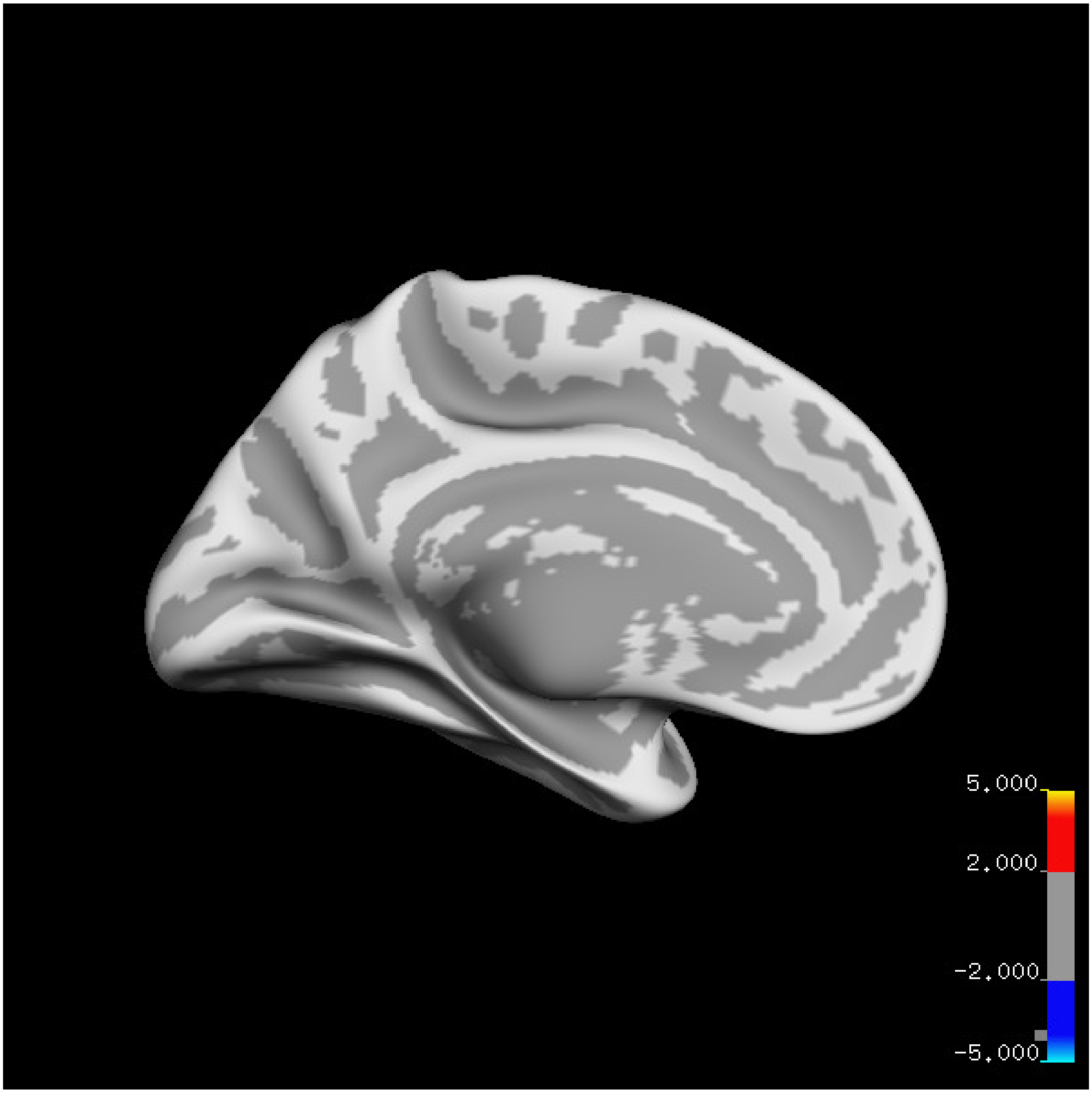
5.000
2.000
-2.000
-5.000

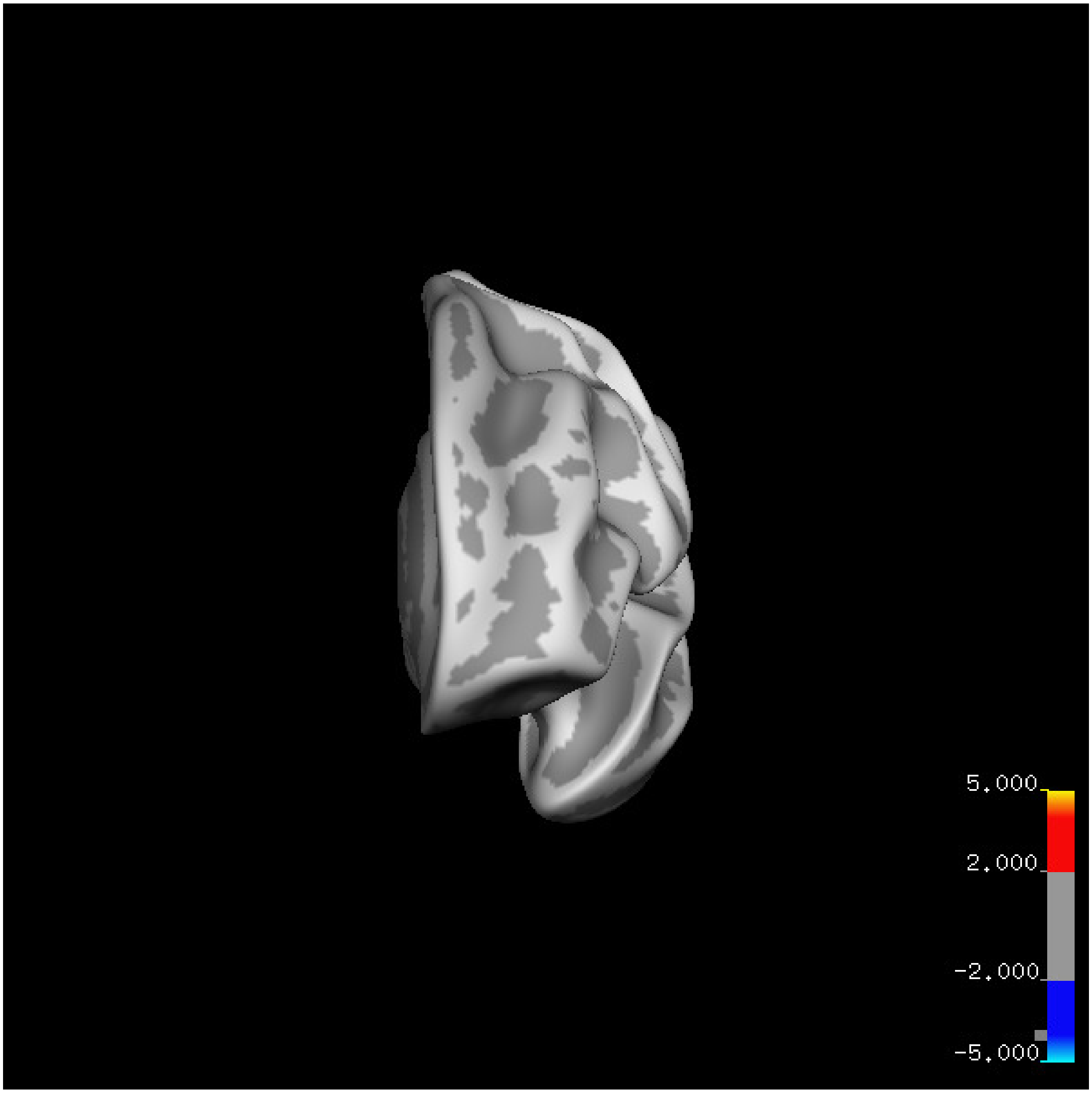
5.000
2.000
-2.000
-5.000

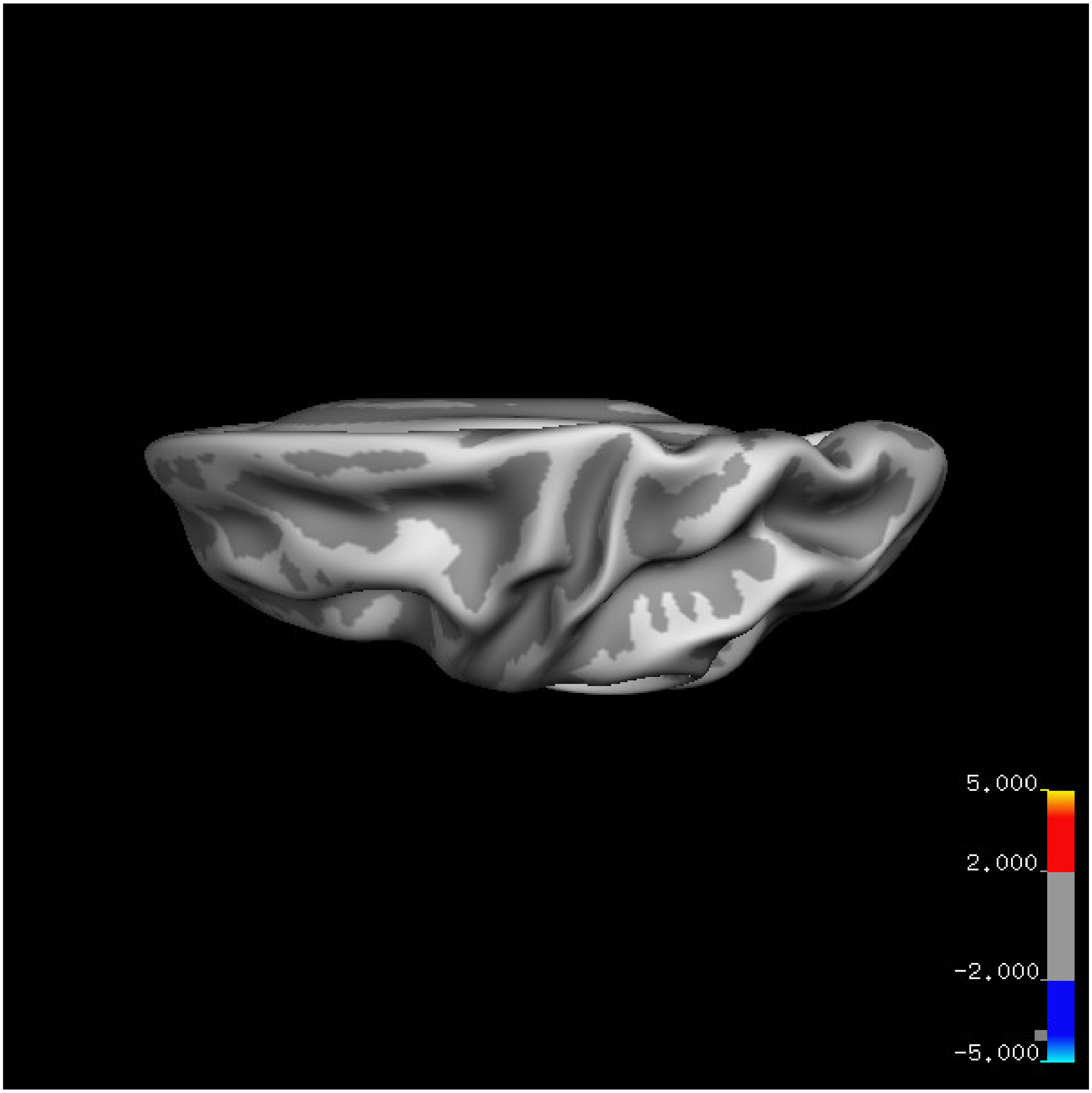


Cluster summary is shown below.

```
## # Cluster Growing Summary (mri_surfcluster)

## # $Id: mri_surfcluster.c,v 1.57.2.3 2016/11/17 18:19:42 zkaufman Exp $

## # $Id: mrisurf.c,v 1.781.2.6 2016/12/27 16:47:14 zkaufman Exp $

## # CreationTime 2021/10/04-10:48:03-GMT

## # cmdline mri_surfcluster.bin --in surface_analyses/results/cluster_comparison/fsaverage6.sm15.lh/vo

## # cwd /gpfs/projects02/p274/cluster/projects/p003-sleep_b

## # sysname  Linux
```

```
## # hostname c1-18
## # machine  x86_64
## # FixVertexAreaFlag 1
## # FixSurfClusterArea 1
## #
## # Input      surface_analyses/results/cluster_comparison/fsaverage6.sm15.lh/volume.20_60.above_mean/g
## # Frame Number      0
## # srcsubj fsaverage6
## # hemi lh
## # surface white
## # group_avg_surface_area 84969.3
## # group_avg_vtxarea_loaded 1
## # annot aparc
## # SUBJECTS_DIR /cluster/projects/p274/tools/mri/freesurfer/current/subjects
## # SearchSpace_mm2 79241.3
## # SearchSpace_vtx 37476
## # Bonferroni 0
## # Minimum Threshold 1.3
## # Maximum Threshold infinity
## # Threshold Sign    abs
## # AdjustThreshWhenOneTail 1
## # CW PValue Threshold: 0.01
## # Area Threshold    0 mm^2
## # CSD thresh  1.300000
## # CSD nreps    10000
## # CSD simtype  null-z
## # CSD contrast NA
```

```
## # CSD confint  90.000000
## # Overall max 1.67798 at vertex 25969
## # Overall min -3.27209 at vertex 7376
## # NClusters          0
## # FixMNI = 0
## #
## # ClusterNo  Max   VtxMax   Size(mm^2)  MNIX   MNIY   MNIZ    CWP    CWPLow    CWPHi   NVtxs    Wght
```

**Sleep below mean** Cluster corrected significance plots are shown below.

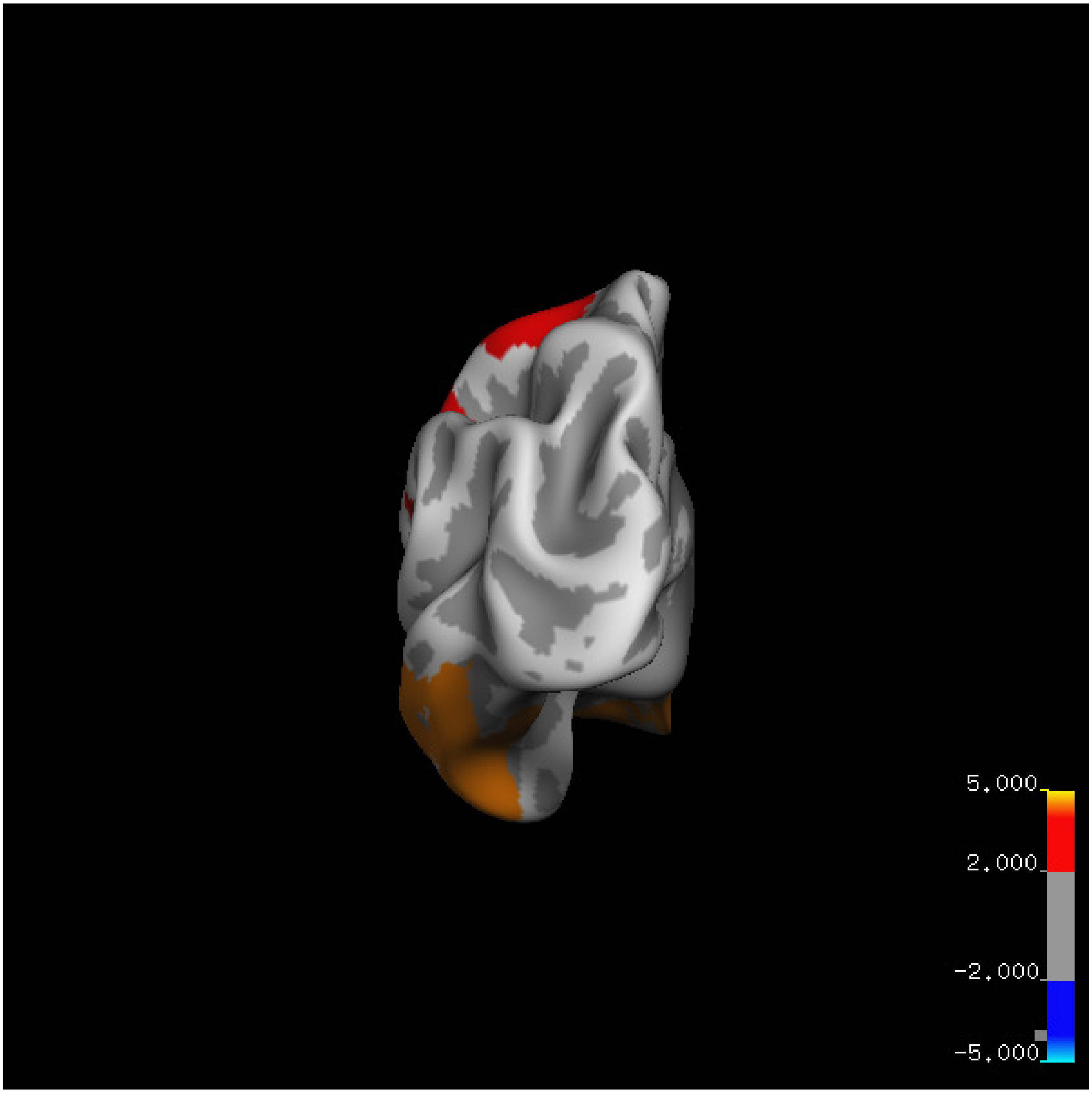
5.000
2.000
-2.000
-5.000

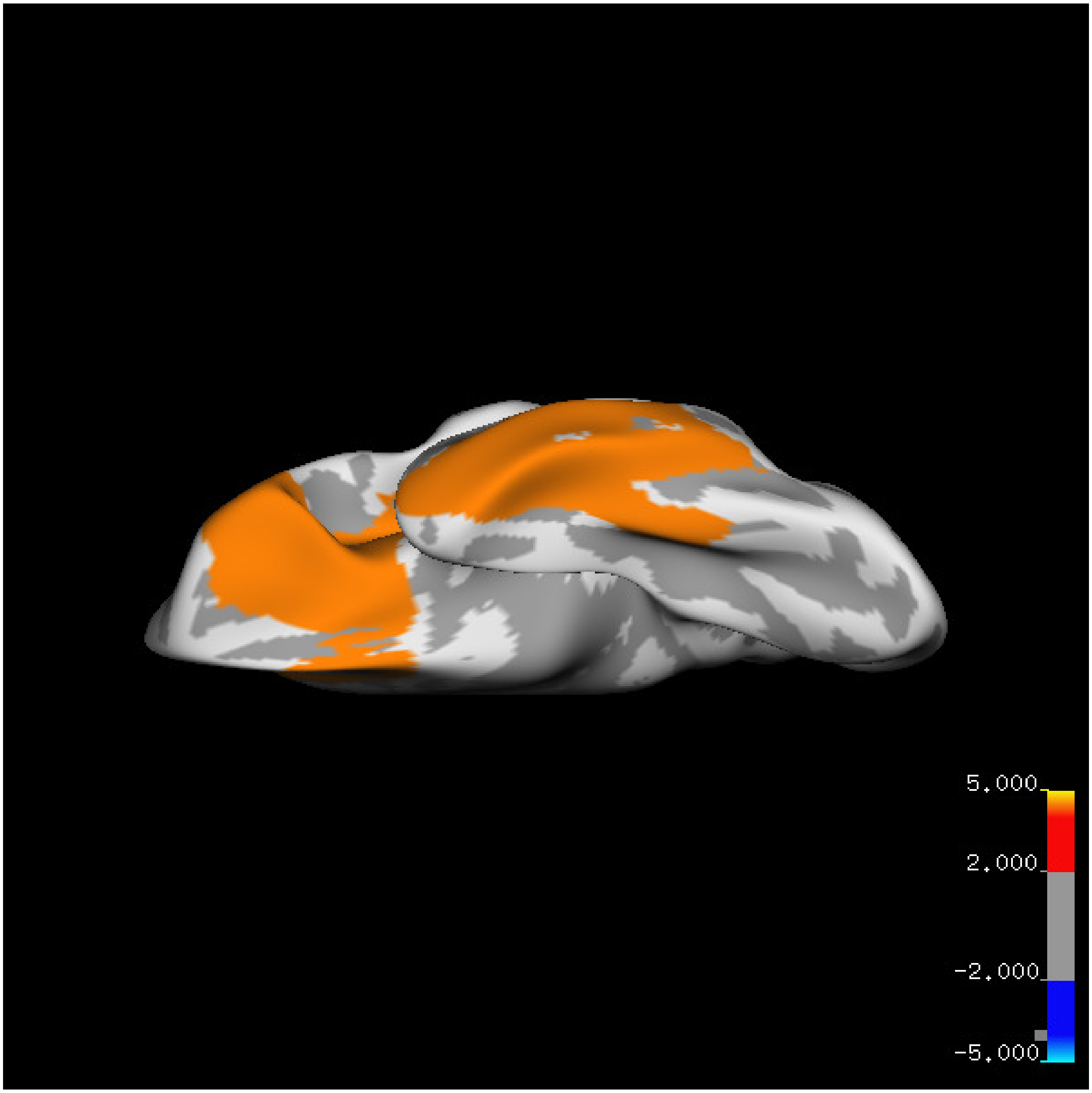
5.000
2.000
-2.000
-5.000

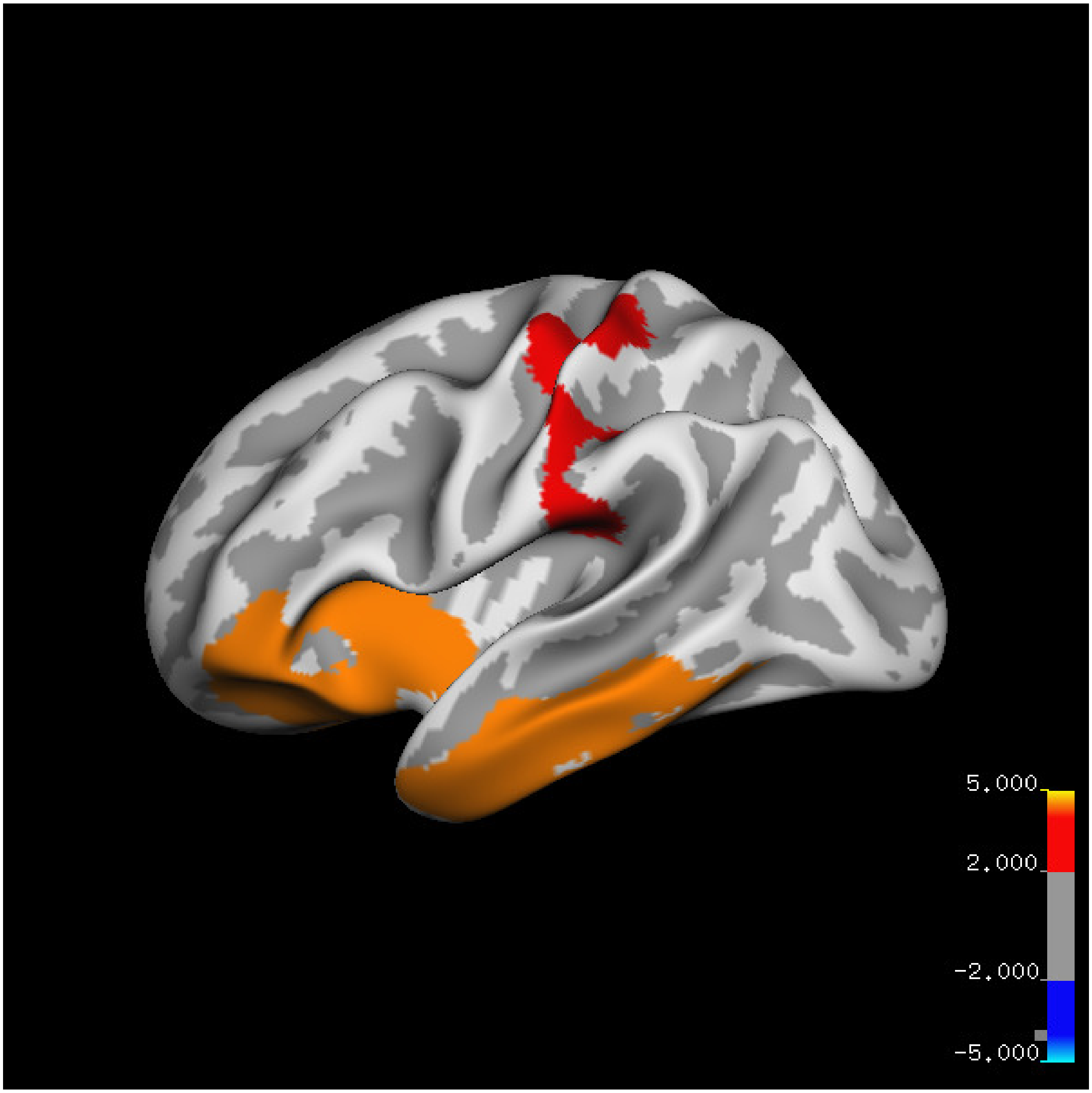
5.000
2.000
-2.000
-5.000

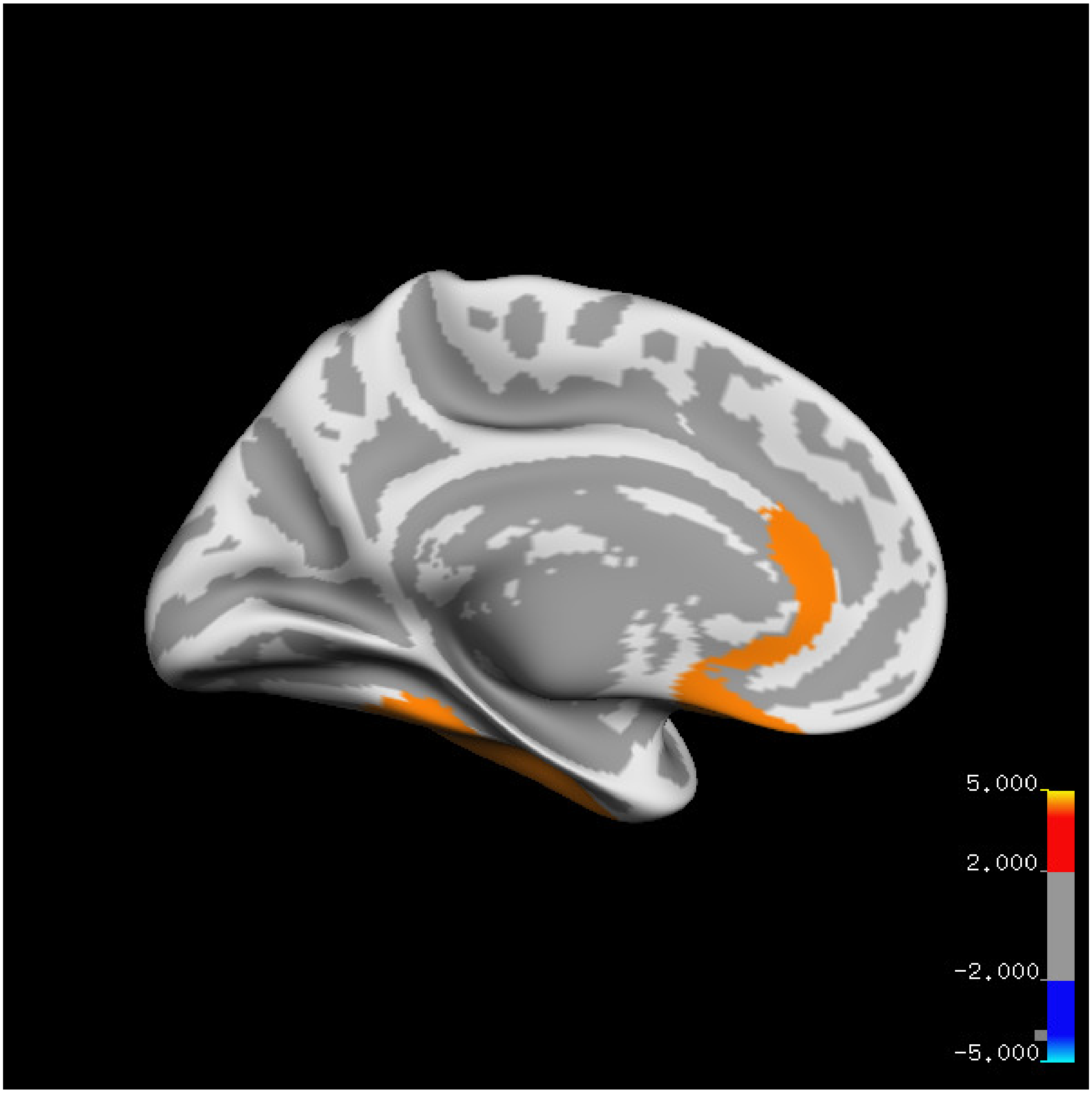
5.000
2.000
-2.000
-5.000

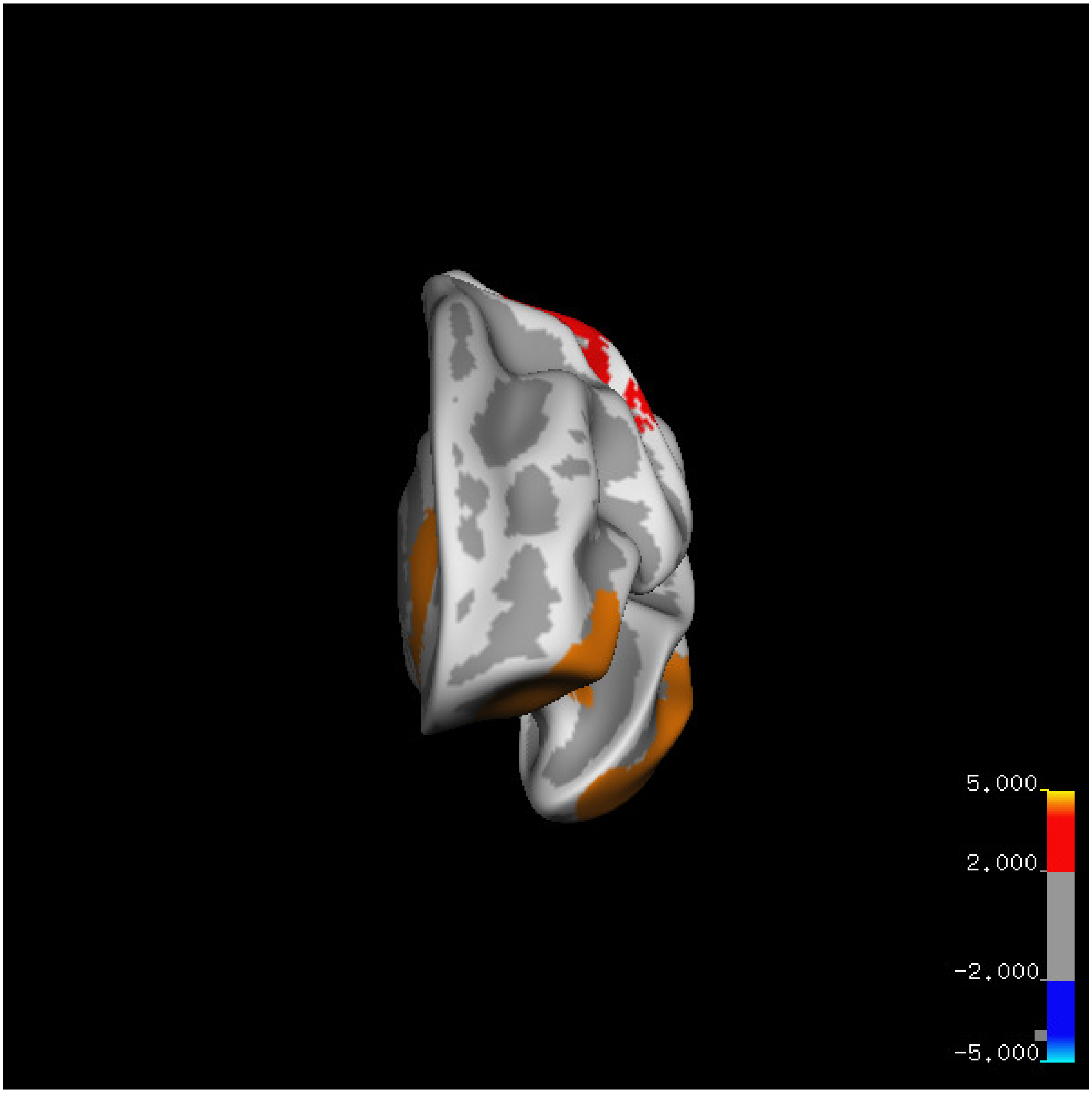
5.000
2.000
-2.000
-5.000

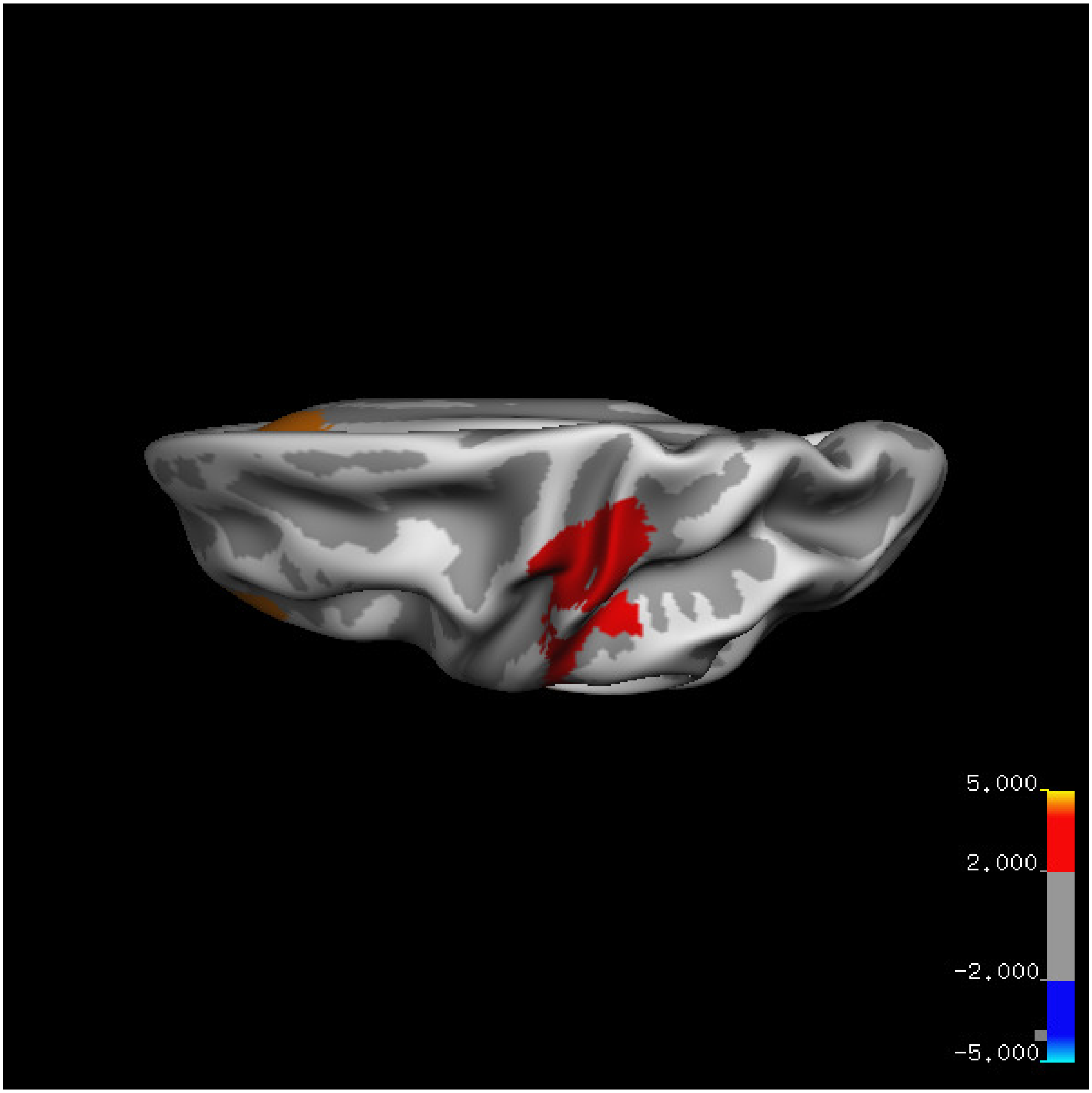

5.000
2.000
-2.000
-5.000

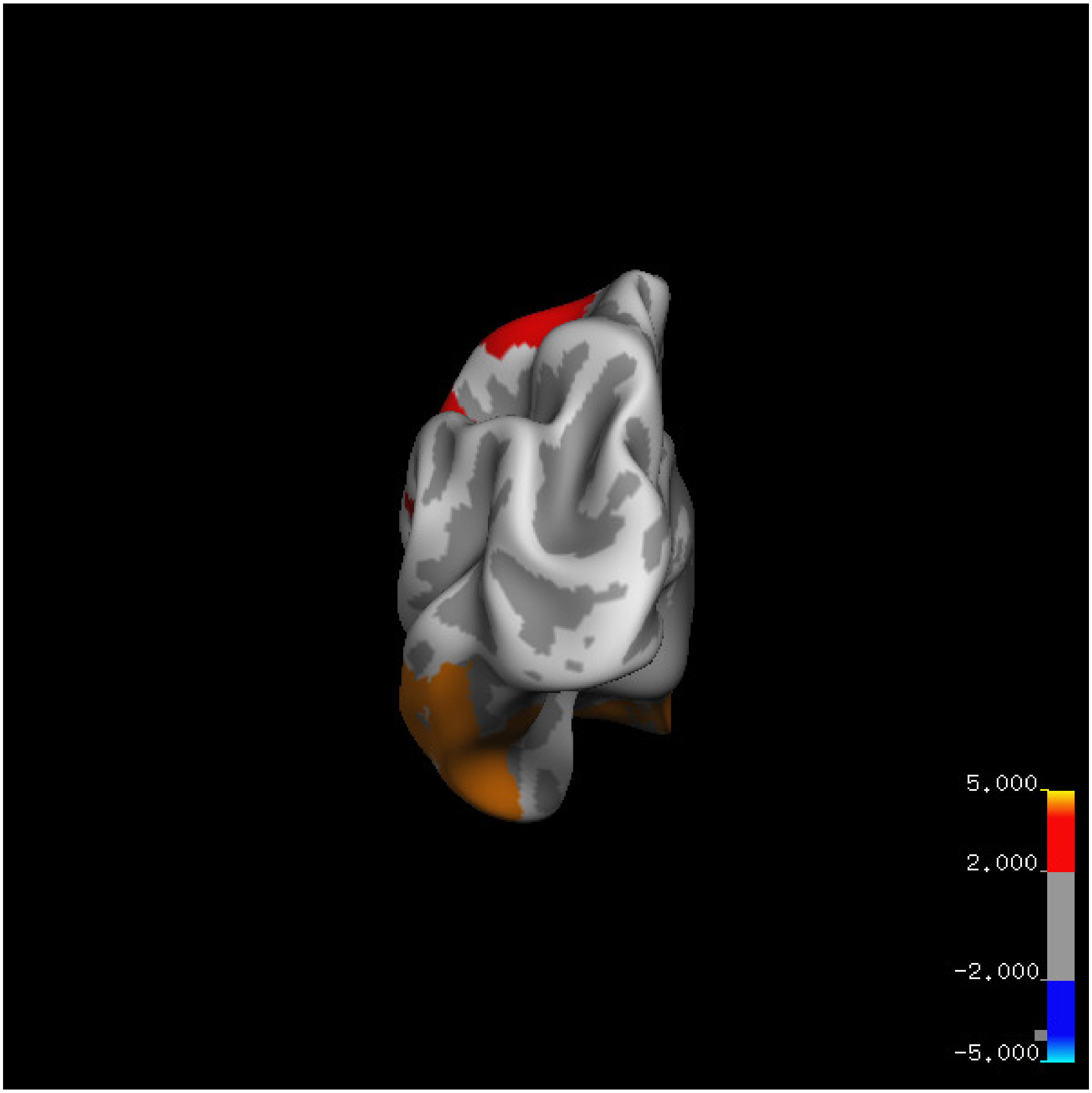

5.000
2.000
-2.000
-5.000

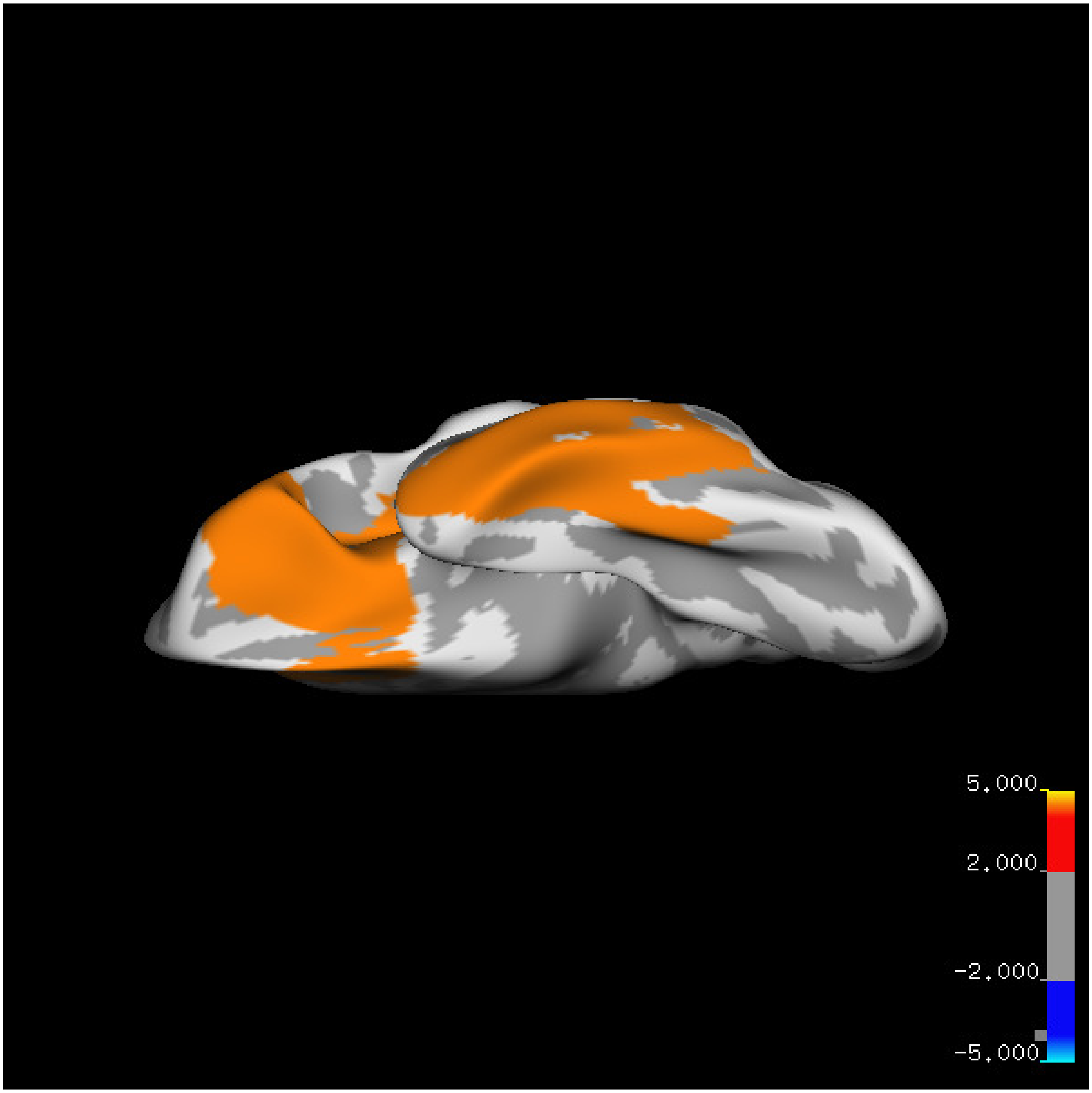
5.000
2.000
-2.000
-5.000

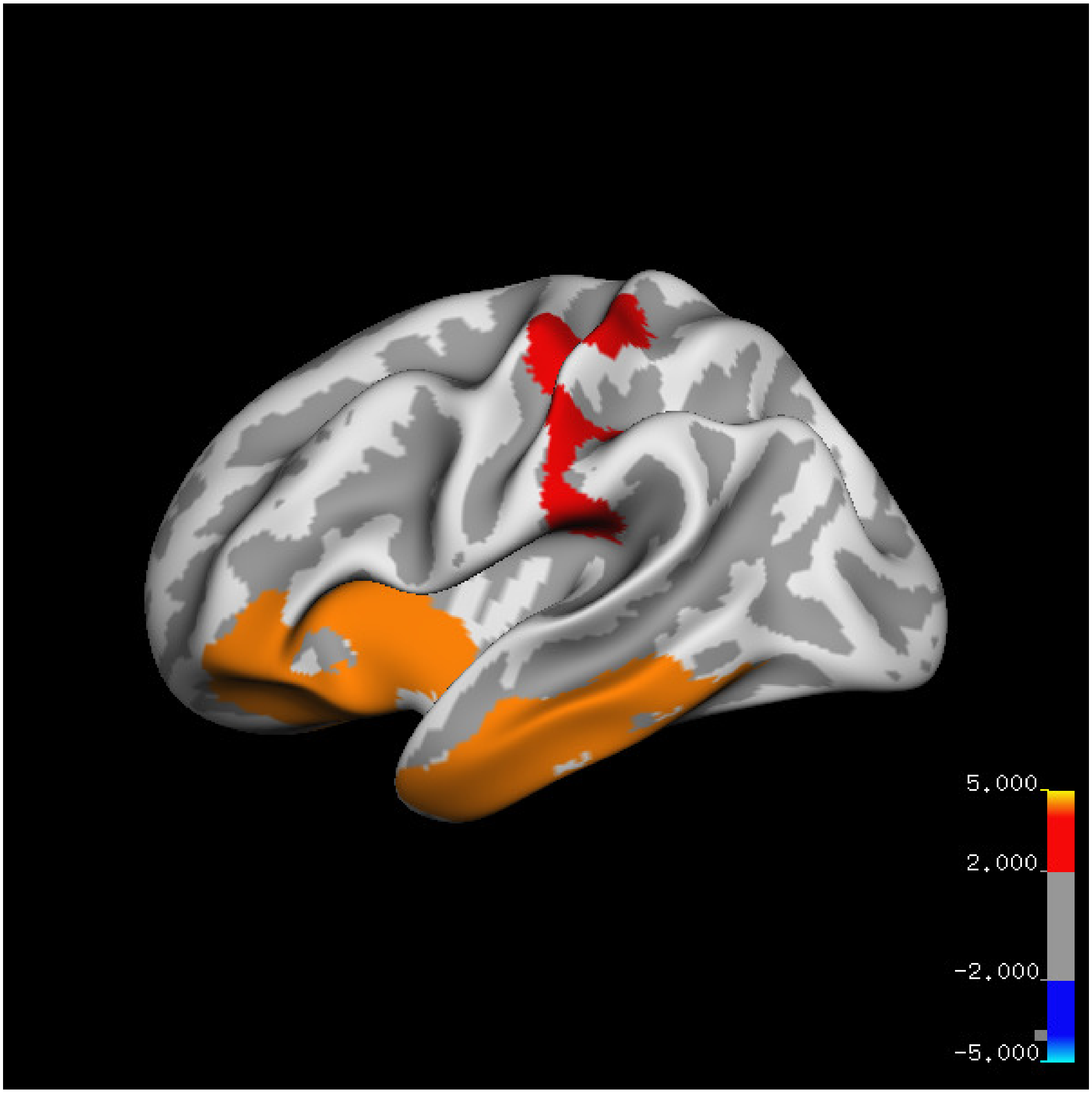
5.000
2.000
-2.000
-5.000

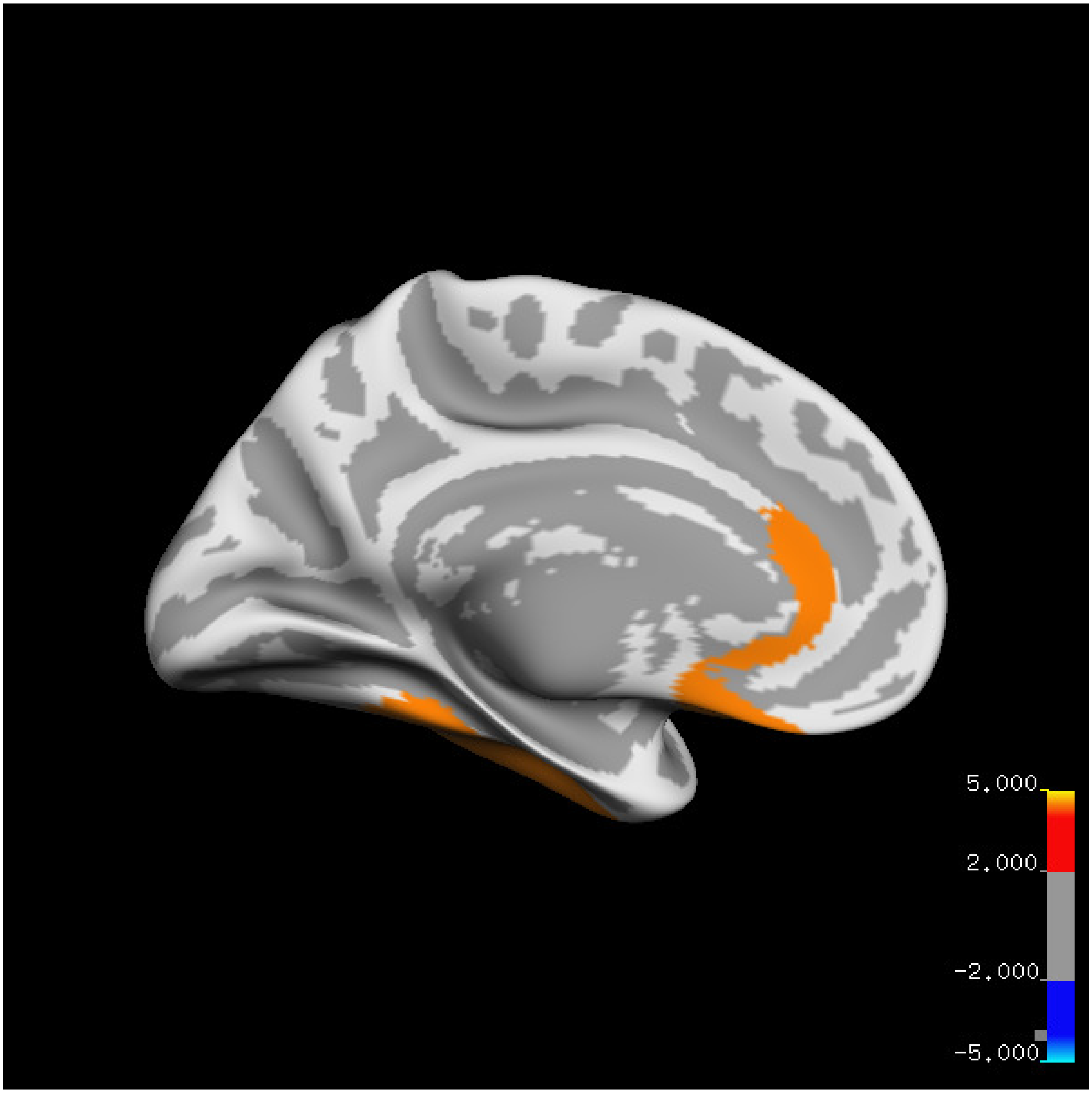
5.000
2.000
-2.000
-5.000

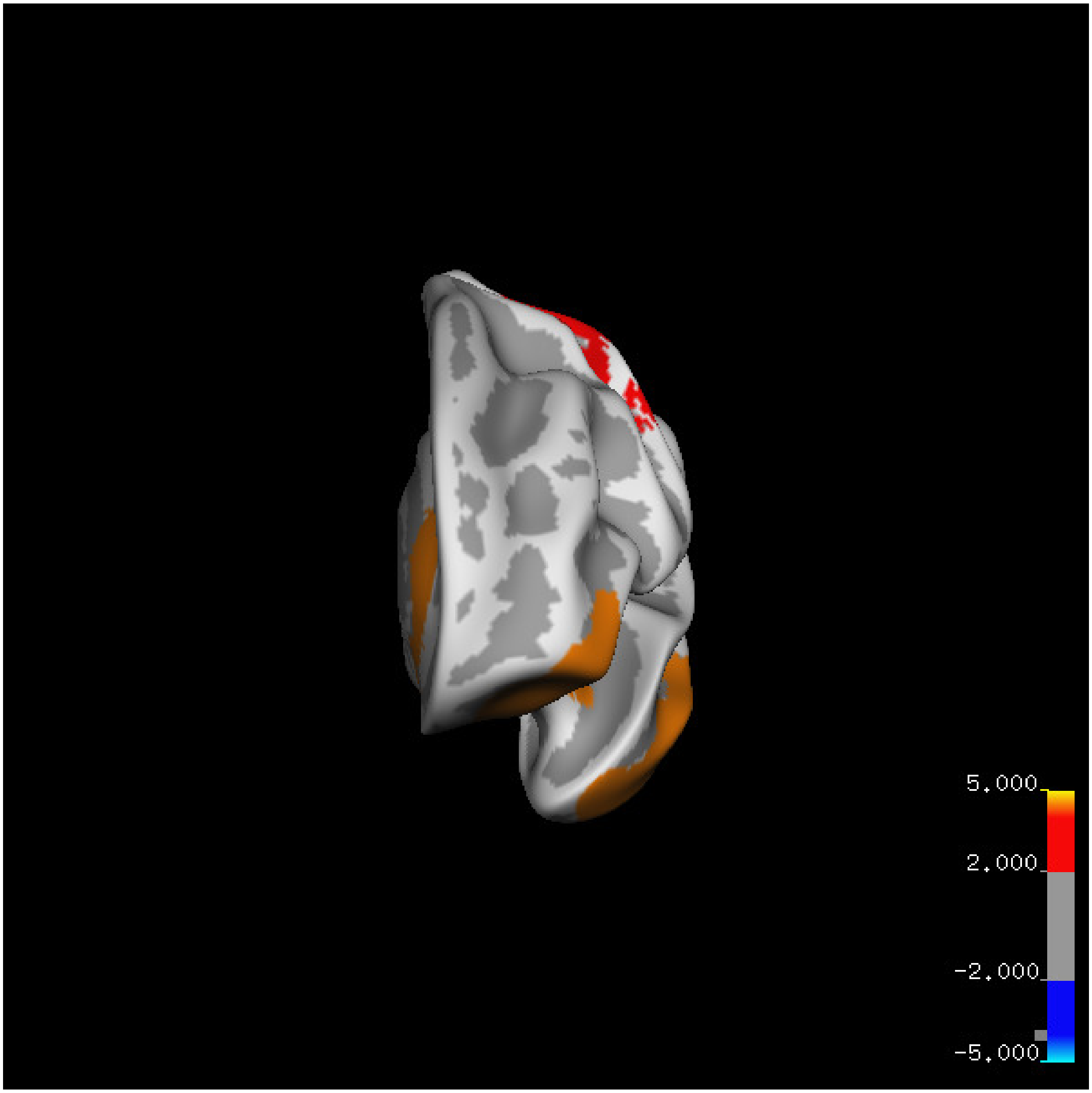
5.000
2.000
-2.000
-5.000

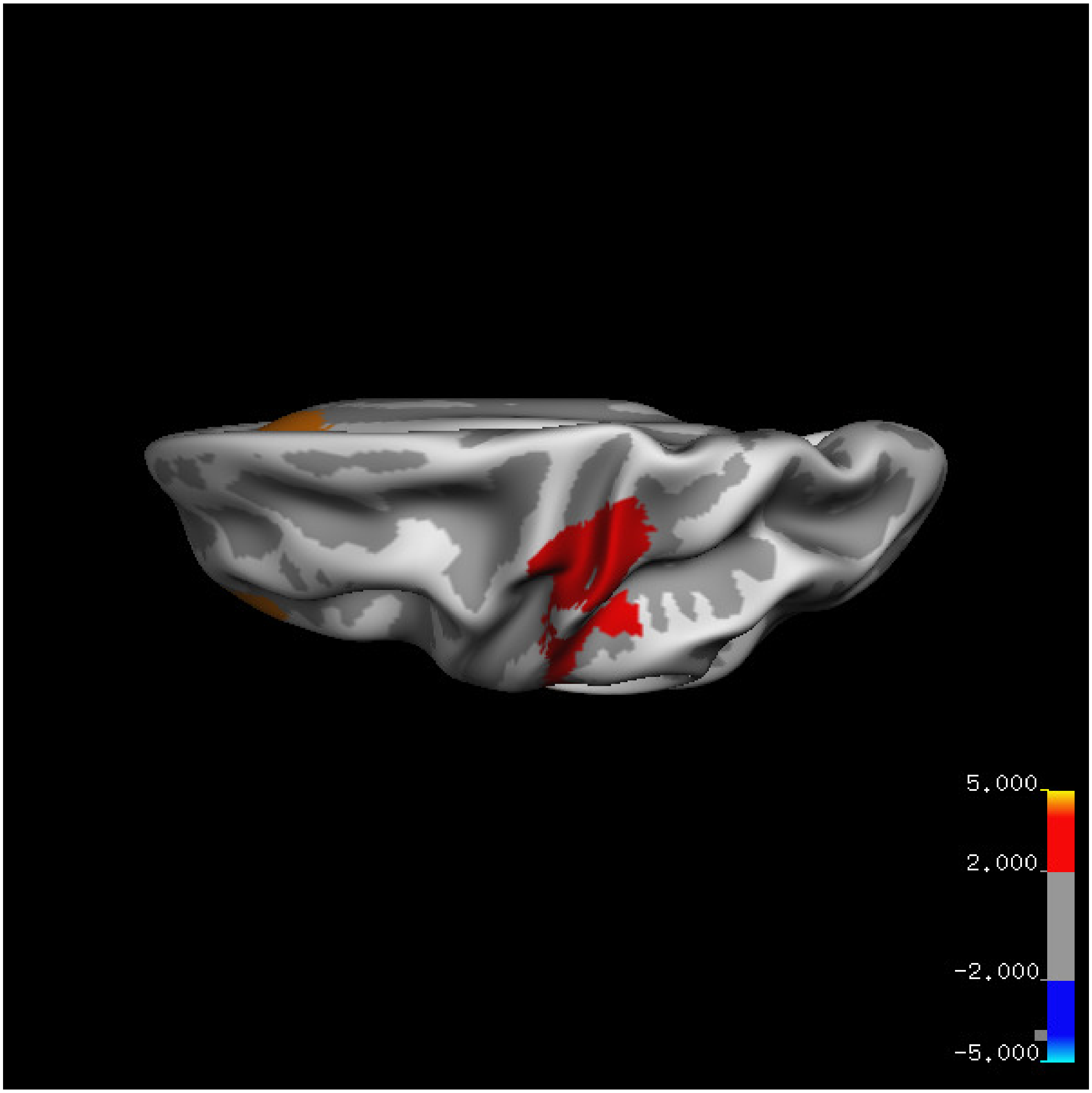


Cluster summary is shown below.

```
## # Cluster Growing Summary (mri_surfcluster)

## # $Id: mri_surfcluster.c,v 1.57.2.3 2016/11/17 18:19:42 zkaufman Exp $

## # $Id: mrisurf.c,v 1.781.2.6 2016/12/27 16:47:14 zkaufman Exp $

## # CreationTime 2021/10/04-11:24:51-GMT

## # cmdline mri_surfcluster.bin --in surface_analyses/results/cluster_comparison/fsaverage6.sm15.lh/vo

## # cwd /gpfs/projects02/p274/cluster/projects/p003-sleep_b

## # sysname  Linux
```

```
## # hostname c1-19
## # machine  x86_64
## # FixVertexAreaFlag 1
## # FixSurfClusterArea 1
## #
## # Input      surface_analyses/results/cluster_comparison/fsaverage6.sm15.lh/volume.20_60.below_mean/g
## # Frame Number      0
## # srcsubj fsaverage6
## # hemi lh
## # surface white
## # group_avg_surface_area 84969.3
## # group_avg_vtxarea_loaded 1
## # annot aparc
## # SUBJECTS_DIR /cluster/projects/p274/tools/mri/freesurfer/current/subjects
## # SearchSpace_mm2 79241.3
## # SearchSpace_vtx 37476
## # Bonferroni 0
## # Minimum Threshold 1.3
## # Maximum Threshold infinity
## # Threshold Sign    abs
## # AdjustThreshWhenOneTail 1
## # CW PValue Threshold: 0.01
## # Area Threshold    0 mm^2
## # CSD thresh  1.300000
## # CSD nreps    10000
## # CSD simtype  null-z
## # CSD contrast NA
```

```
## # CSD confint  90.000000

## # Overall max 8.82261 at vertex 3247

## # Overall min -0.69944 at vertex 8164

## # NClusters          3

## # FixMNI = 0

## #

## # ClusterNo  Max   VtxMax   Size(mm^2)  MNIX   MNIY   MNIZ    CWP    CWPLow    CWPHi   NVtxs    WghtV

##    1        4.814   40059   5109.03    -60.2   -9.9  -21.7  0.00010  0.00000  0.00020   1893     4251

##    2        8.823    3247   4677.96    -32.1   21.8  -20.8  0.00010  0.00000  0.00020   2328     5860

##    3        4.142   15897   2826.67    -31.0  -24.2   53.8  0.00160  0.00110  0.00210   1684     3233
```

**Age above 60**

**Sleep above mean** Cluster corrected significance plots are shown below.

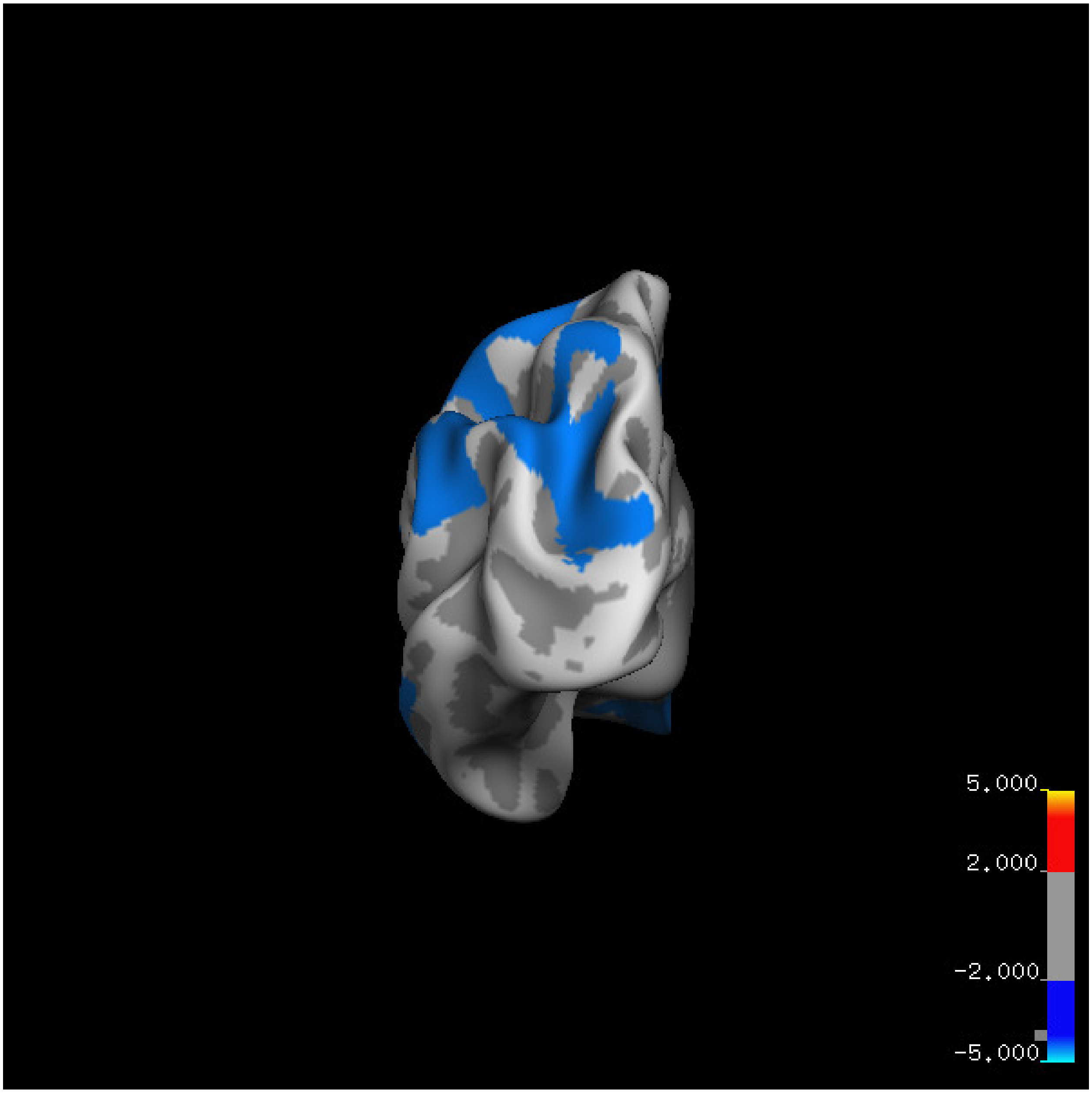
5.000
2.000
-2.000
-5.000

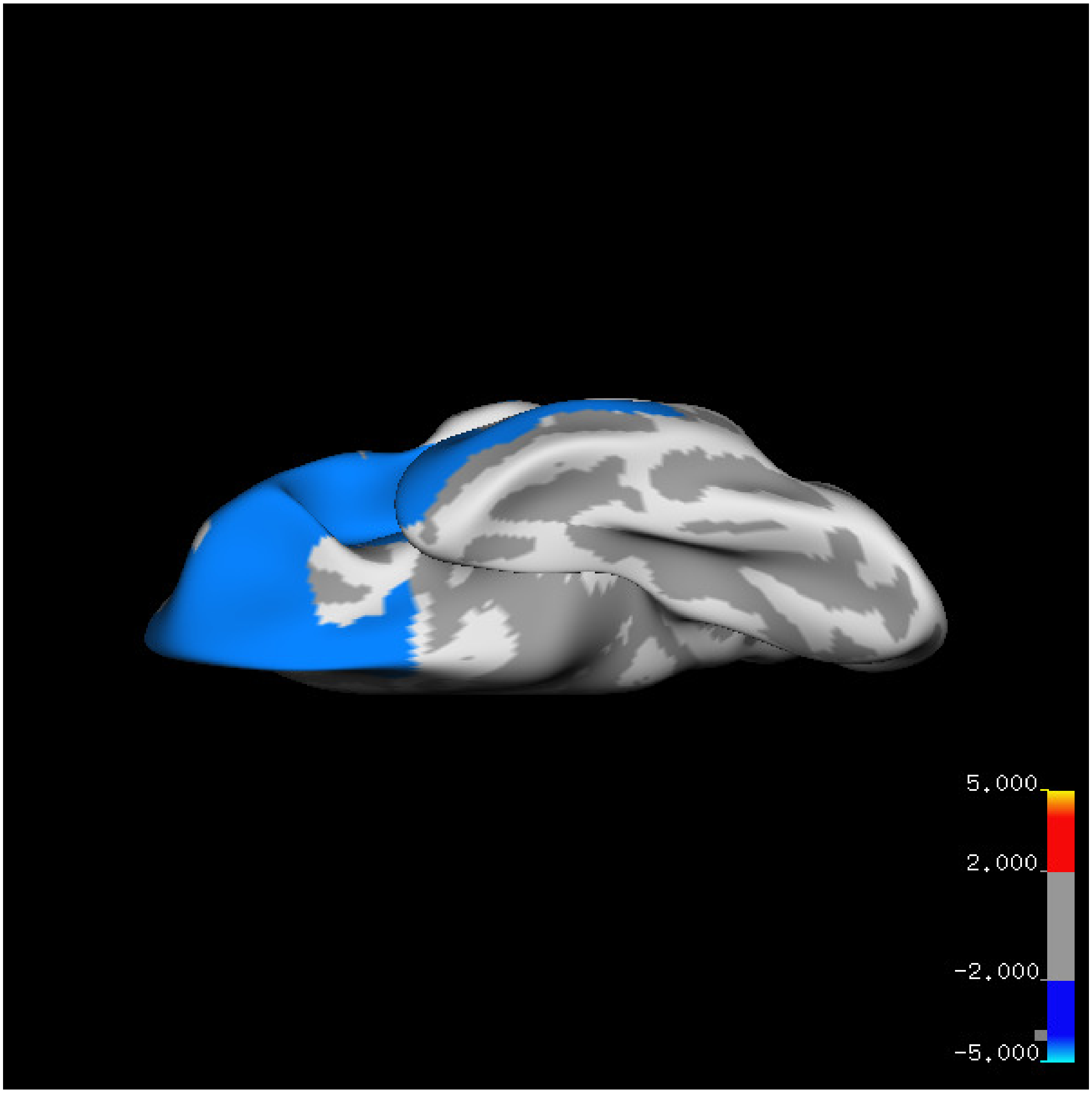
5.000
2.000
-2.000
-5.000

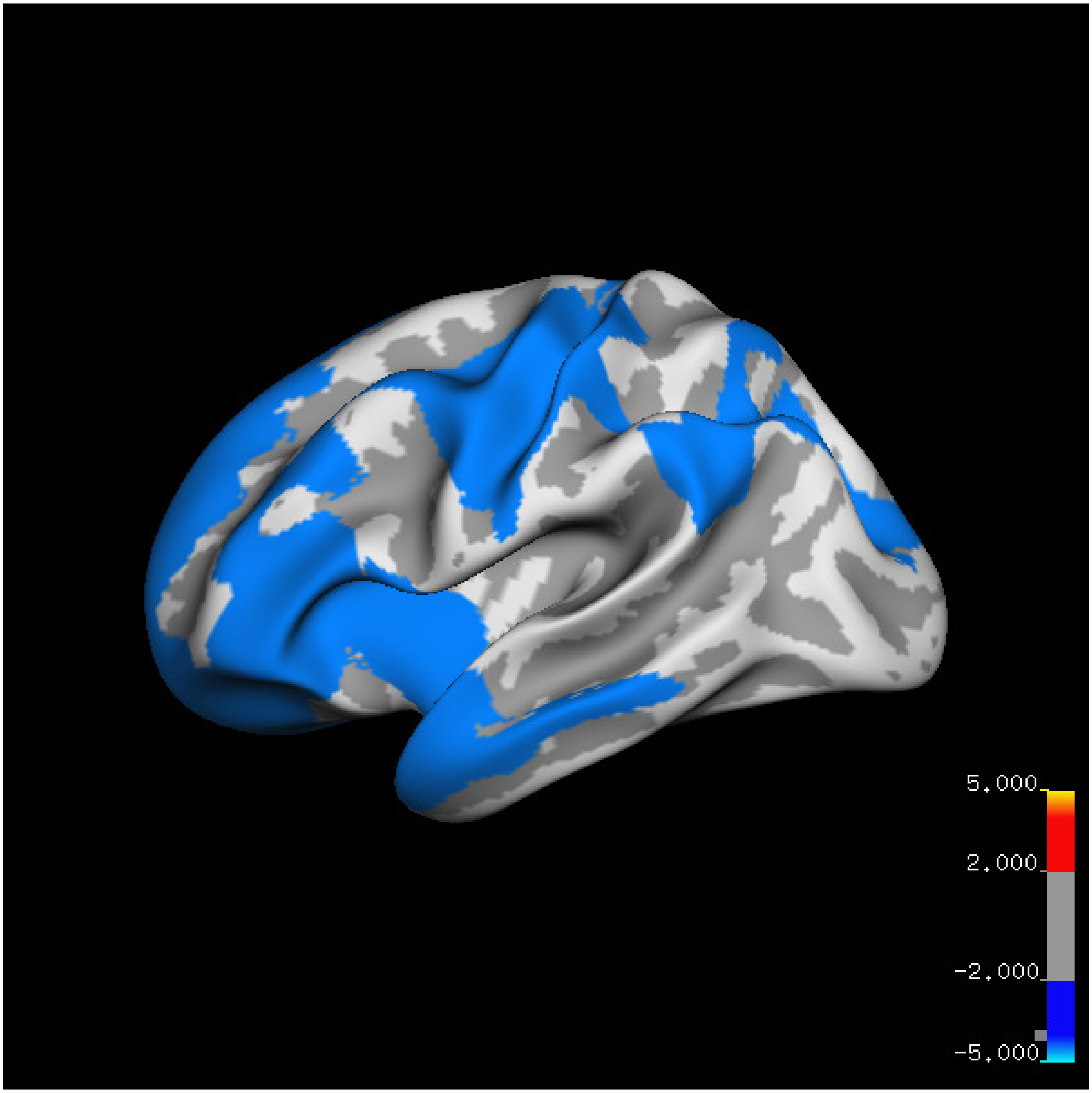
5.000
2.000
-2.000
-5.000

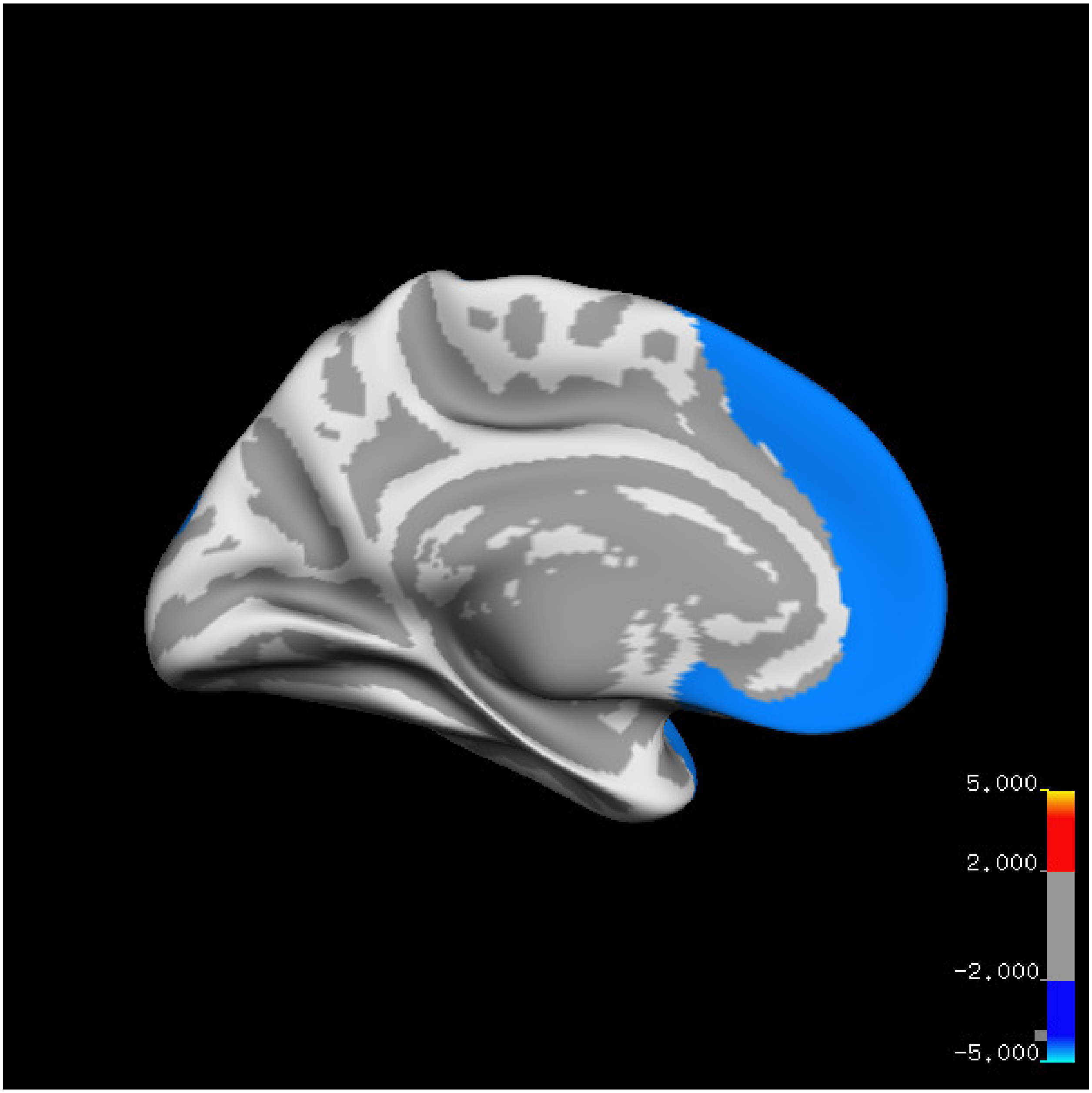
5.000
2.000
-2.000
-5.000

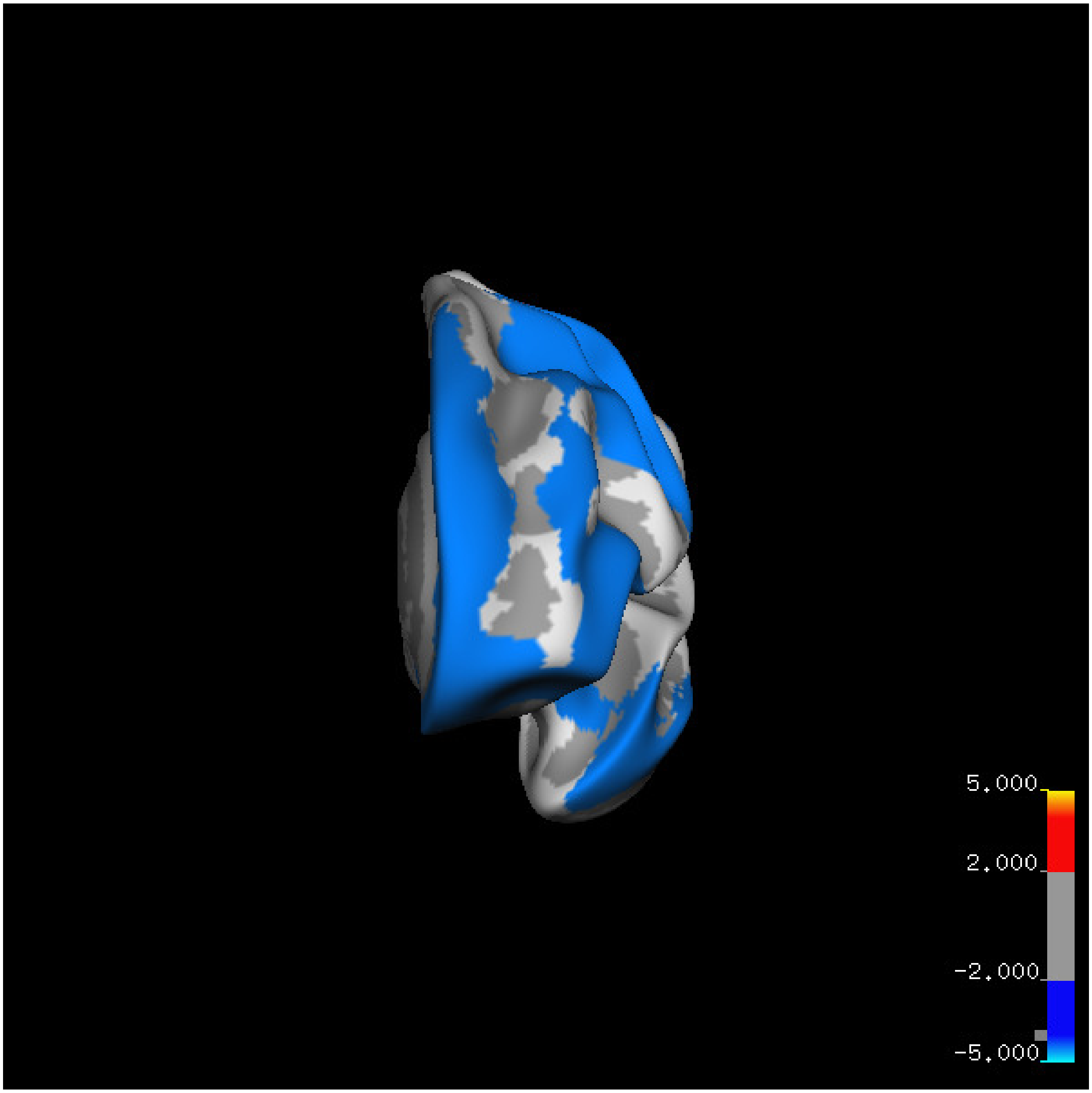
5.000
2.000
-2.000
-5.000

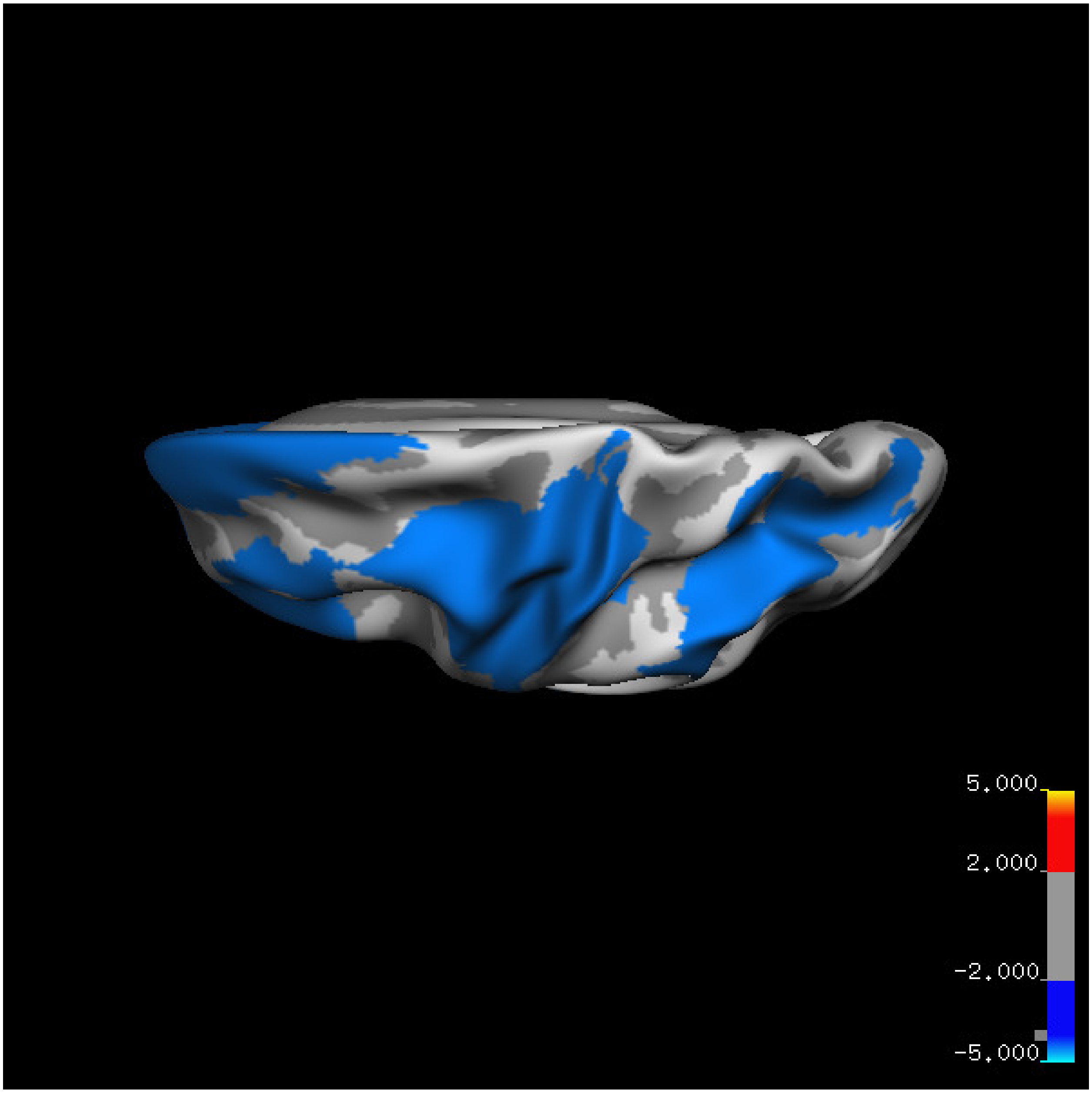
5.000
2.000
-2.000
-5.000

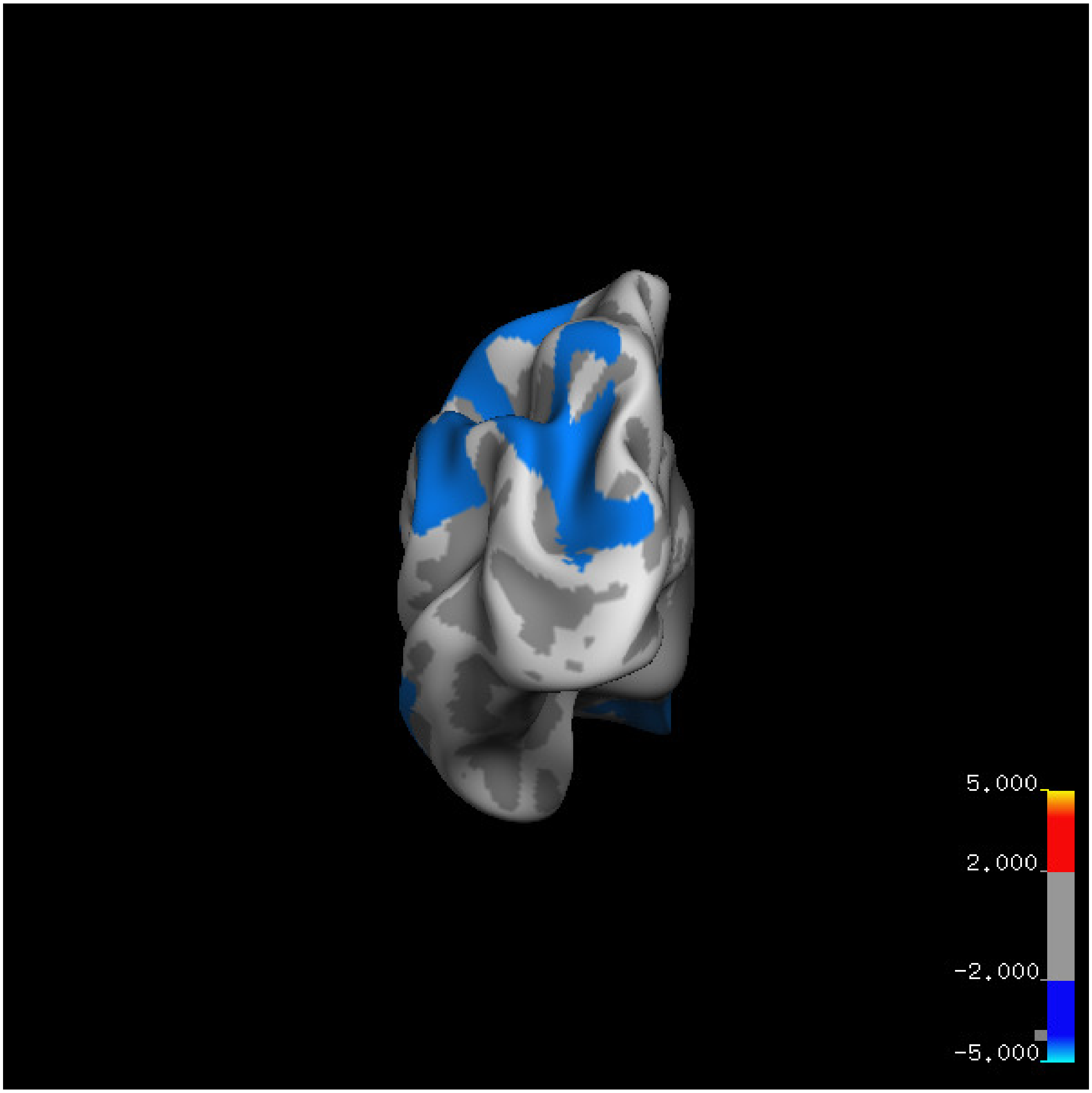

5.000
2.000
-2.000
-5.000

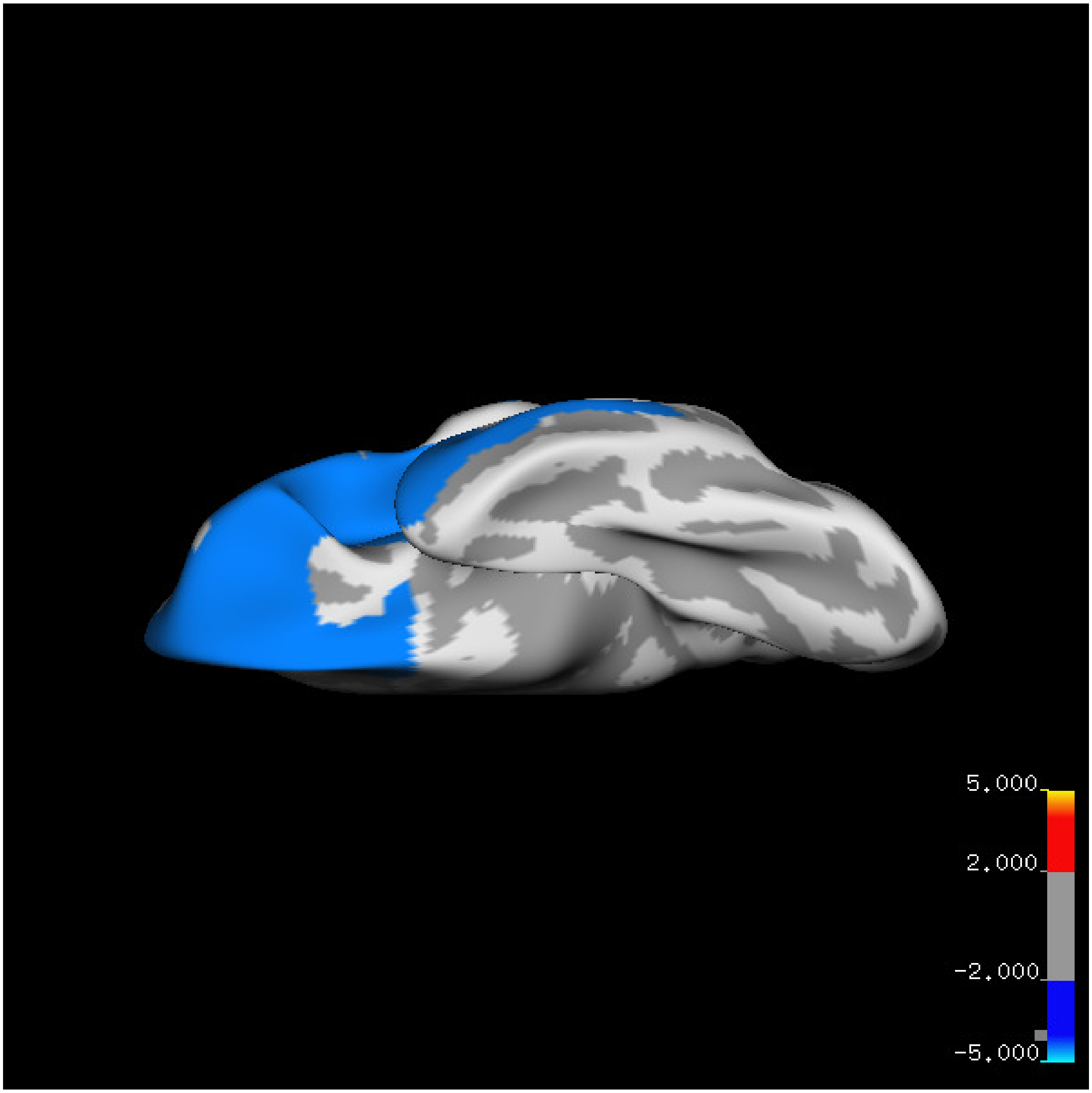
5.000
2.000
-2.000
-5.000

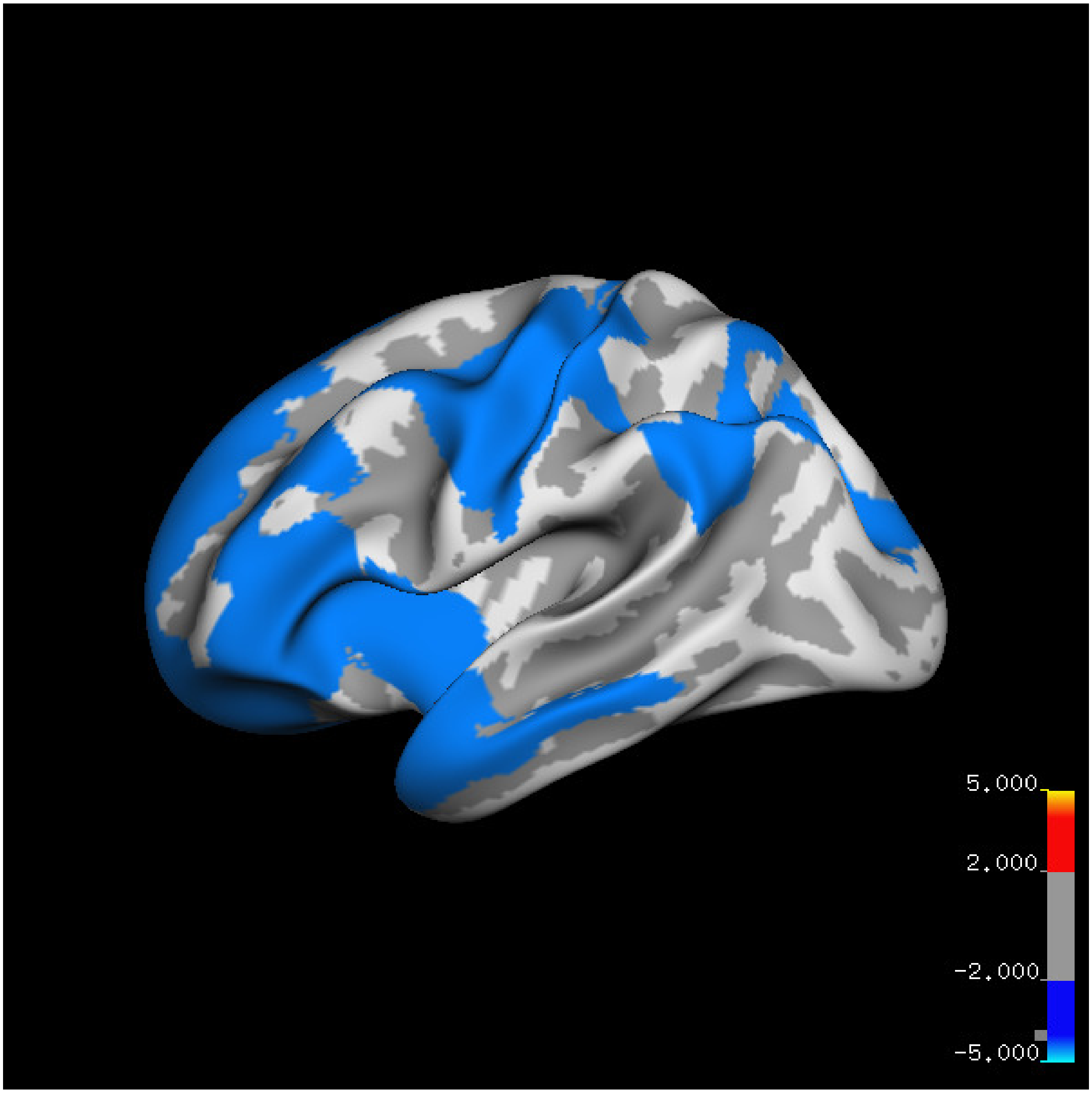
5.000
2.000
-2.000
-5.000

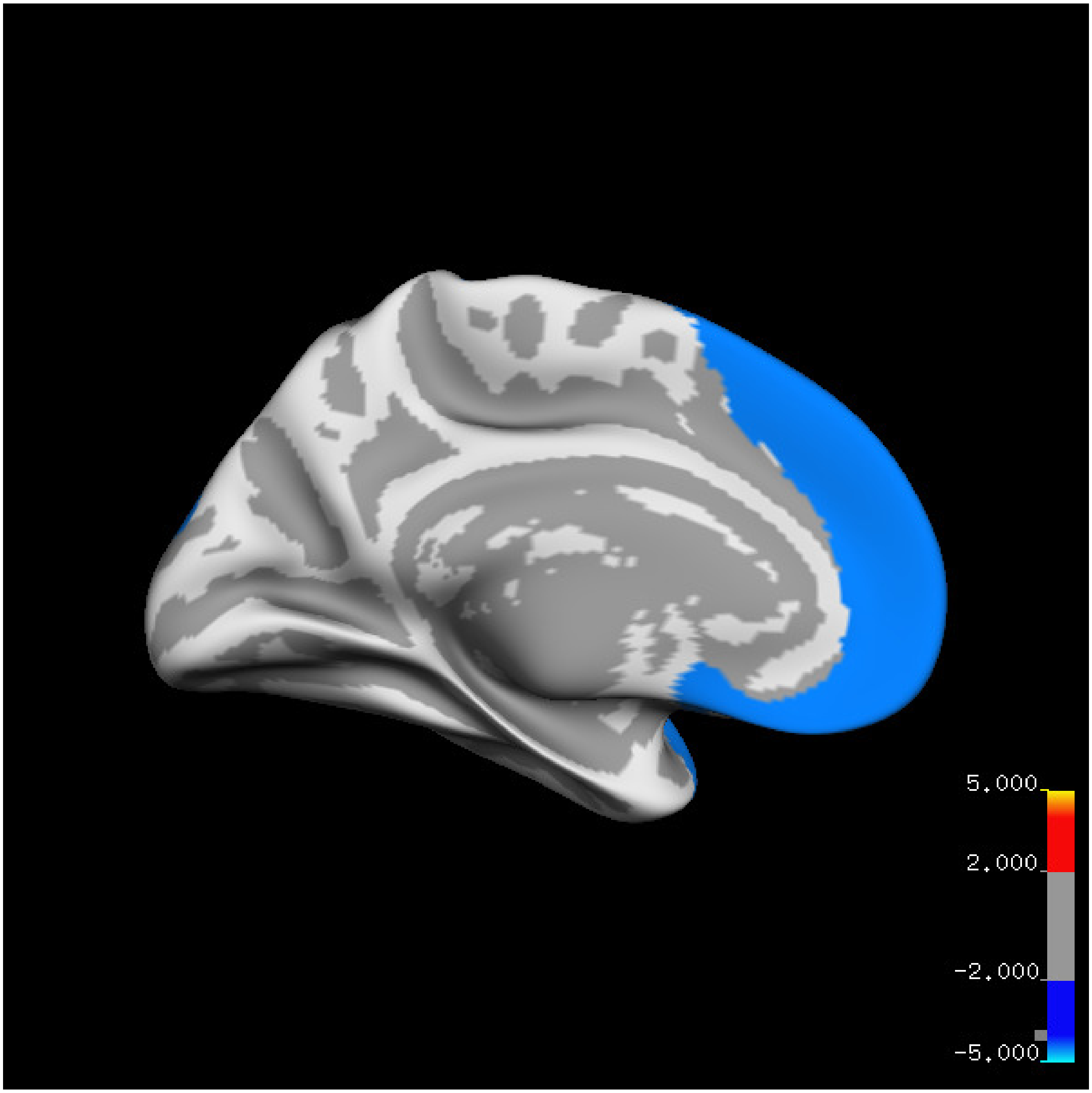

5.000
2.000
-2.000
-5.000

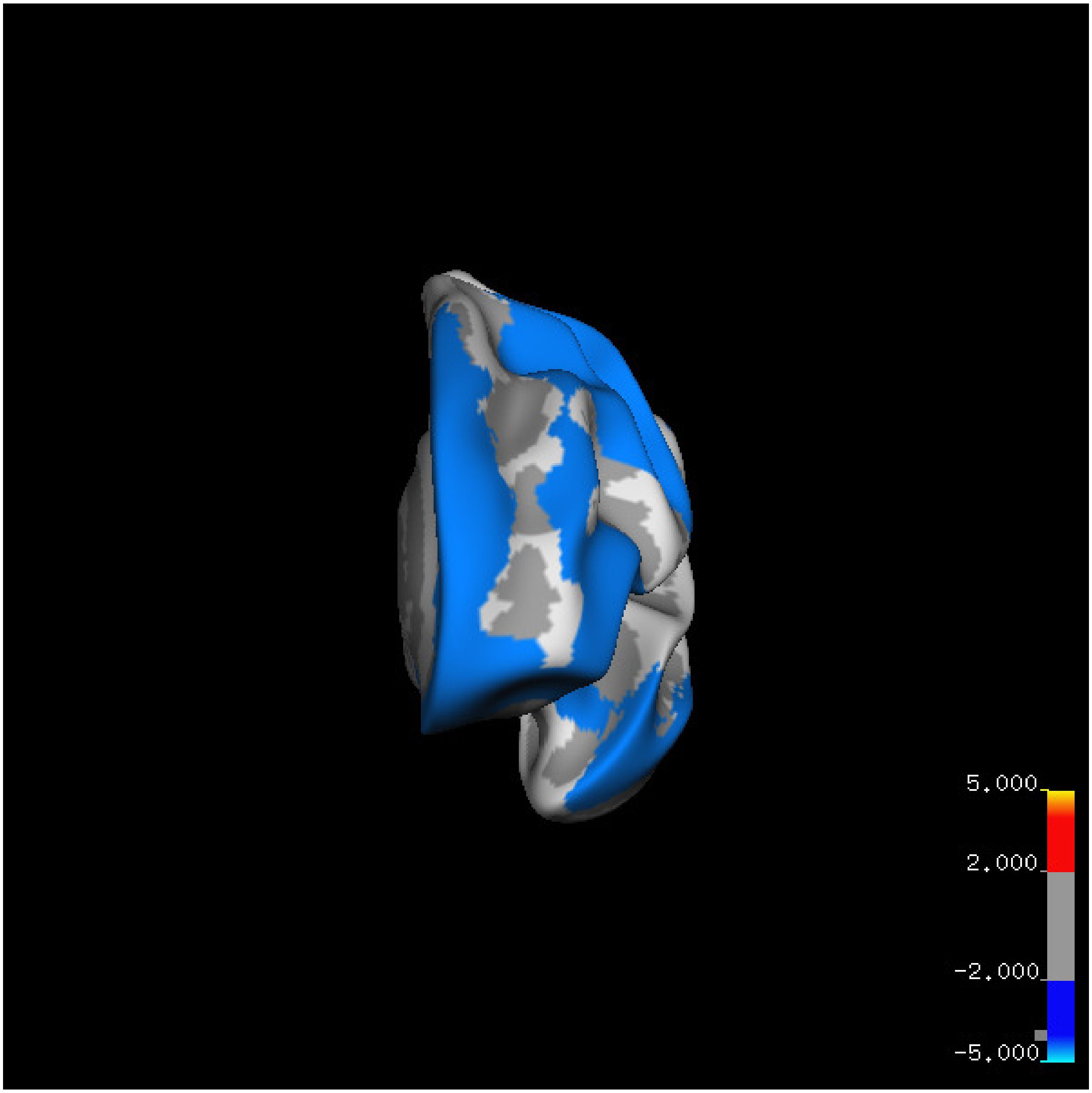
5.000
2.000
-2.000
-5.000

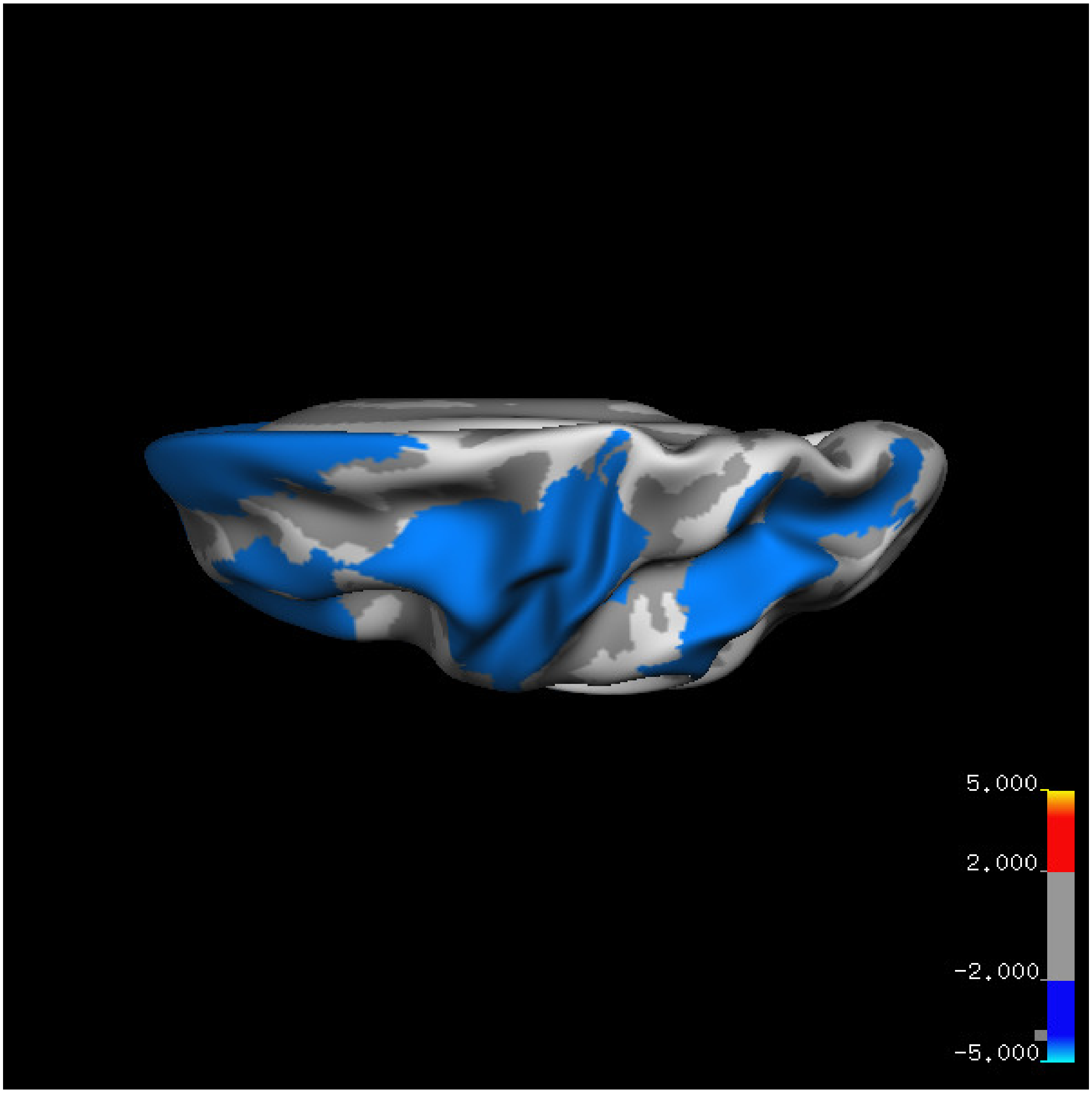


Cluster summary is shown below.

```
## # Cluster Growing Summary (mri_surfcluster)

## # $Id: mri_surfcluster.c,v 1.57.2.3 2016/11/17 18:19:42 zkaufman Exp $

## # $Id: mrisurf.c,v 1.781.2.6 2016/12/27 16:47:14 zkaufman Exp $

## # CreationTime 2021/10/04-13:08:13-GMT

## # cmdline mri_surfcluster.bin --in surface_analyses/results/cluster_comparison/fsaverage6.sm15.lh/vol

## # cwd /gpfs/projects02/p274/cluster/projects/p003-sleep_b

## # sysname  Linux
```

```
## # hostname c1-19
## # machine  x86_64
## # FixVertexAreaFlag 1
## # FixSurfClusterArea 1
## #
## # Input      surface_analyses/results/cluster_comparison/fsaverage6.sm15.lh/volume.above60.above_mea
## # Frame Number      0
## # srcsubj fsaverage6
## # hemi lh
## # surface white
## # group_avg_surface_area 84969.3
## # group_avg_vtxarea_loaded 1
## # annot aparc
## # SUBJECTS_DIR /cluster/projects/p274/tools/mri/freesurfer/current/subjects
## # SearchSpace_mm2 79241.3
## # SearchSpace_vtx 37476
## # Bonferroni 0
## # Minimum Threshold 1.3
## # Maximum Threshold infinity
## # Threshold Sign    abs
## # AdjustThreshWhenOneTail 1
## # CW PValue Threshold: 0.01
## # Area Threshold    0 mm^2
## # CSD thresh  1.300000
## # CSD nreps    10000
## # CSD simtype  null-z
## # CSD contrast NA
```

```
## # CSD confint  90.000000
## # Overall max 0.778486 at vertex 12656
## # Overall min -9.40712 at vertex 9959
## # NClusters          2
## # FixMNI = 0
## #
## # ClusterNo  Max   VtxMax   Size(mm^2)  MNIX   MNIY   MNIZ    CWP    CWPLow    CWPHi   NVtxs    WghtV
##    1       -9.407    9959  22752.42    -43.7    9.5  -30.8  0.00010  0.00000  0.00020  10337   -2974
##    2       -3.579   23551   4601.52    -40.4  -52.4   34.4  0.00010  0.00000  0.00020   2356    -471
```

**Sleep below mean** Cluster corrected significance plots are shown below.

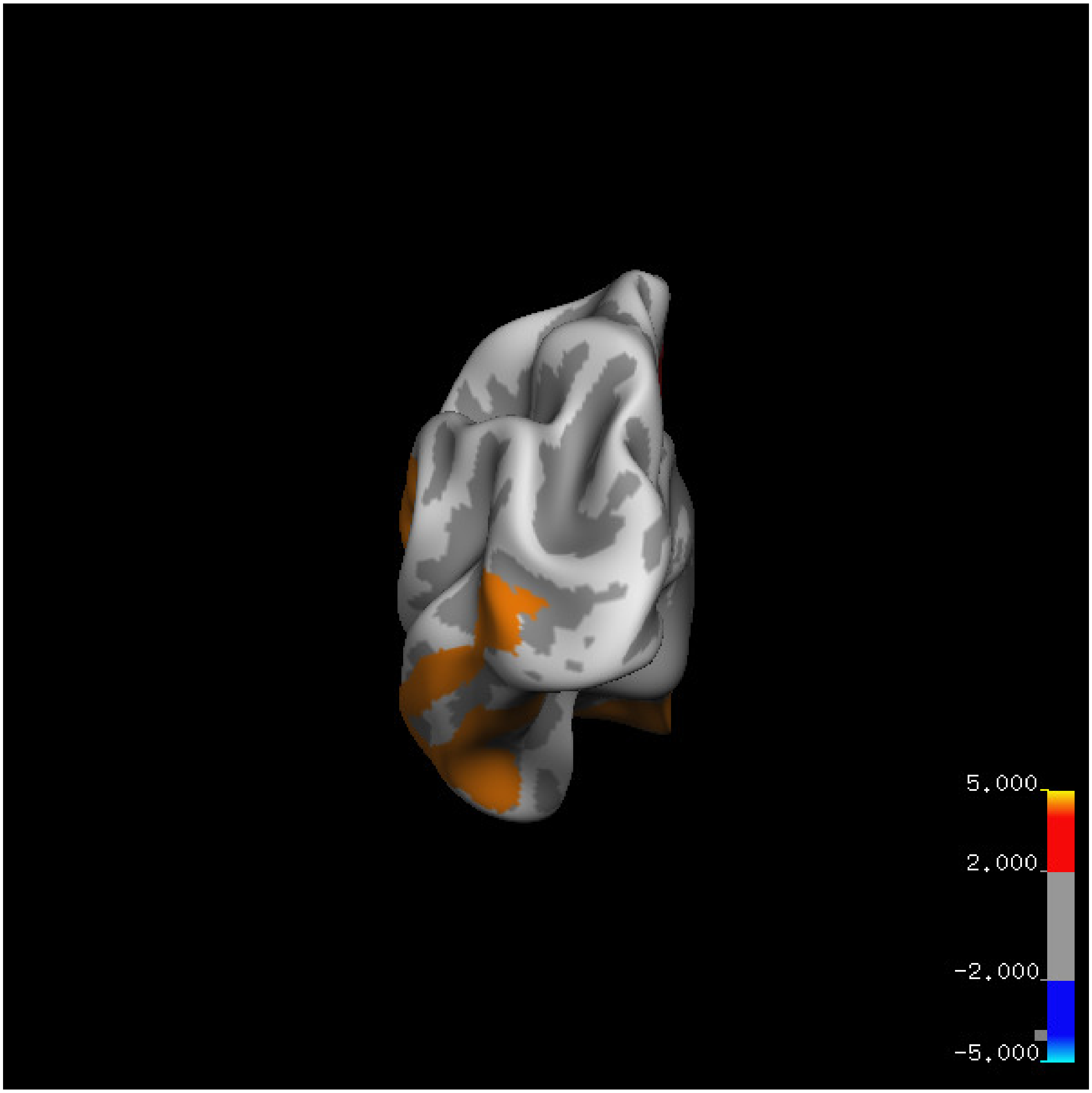
5.000
2.000
-2.000
-5.000

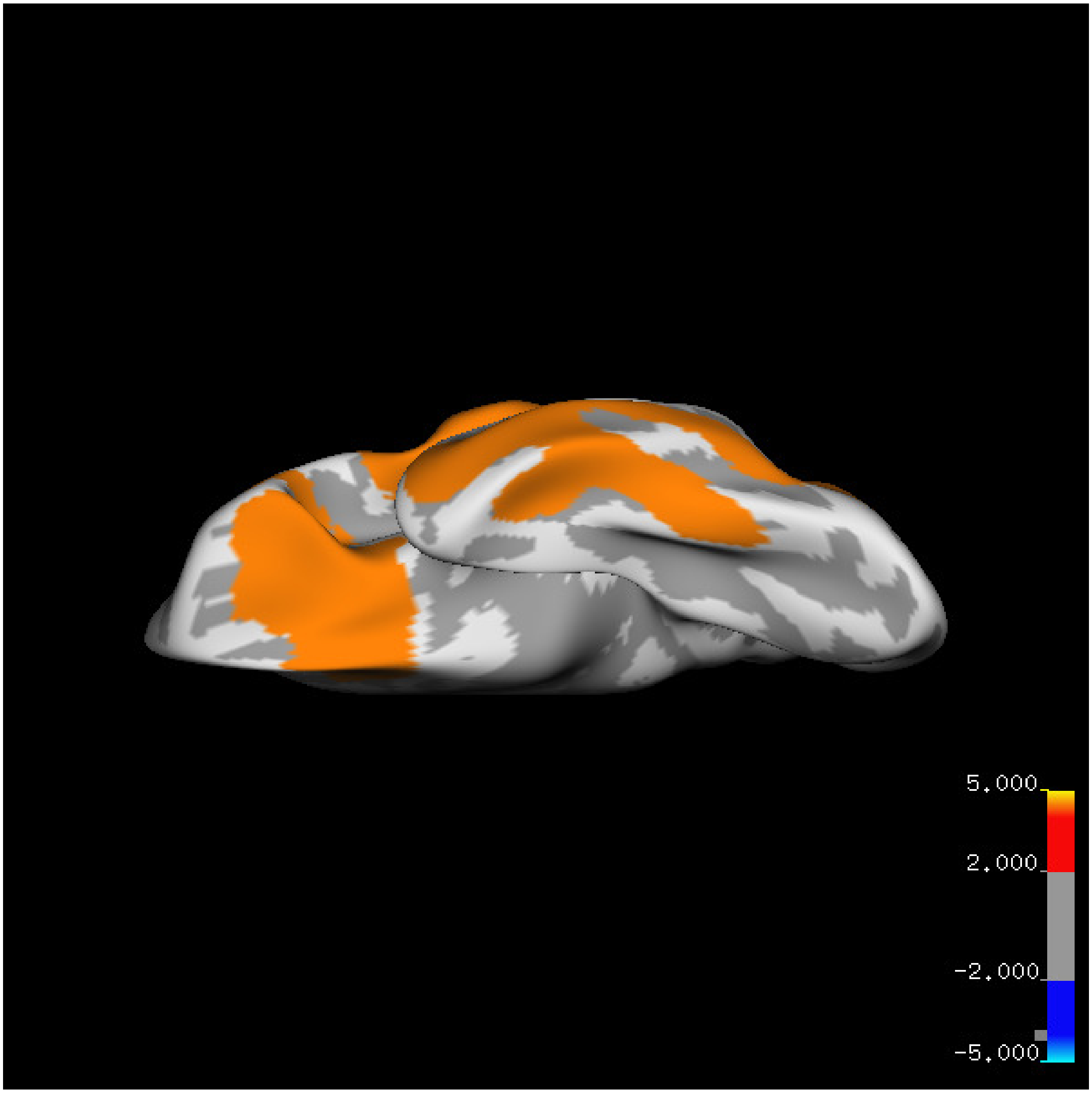
5.000
2.000
-2.000
-5.000

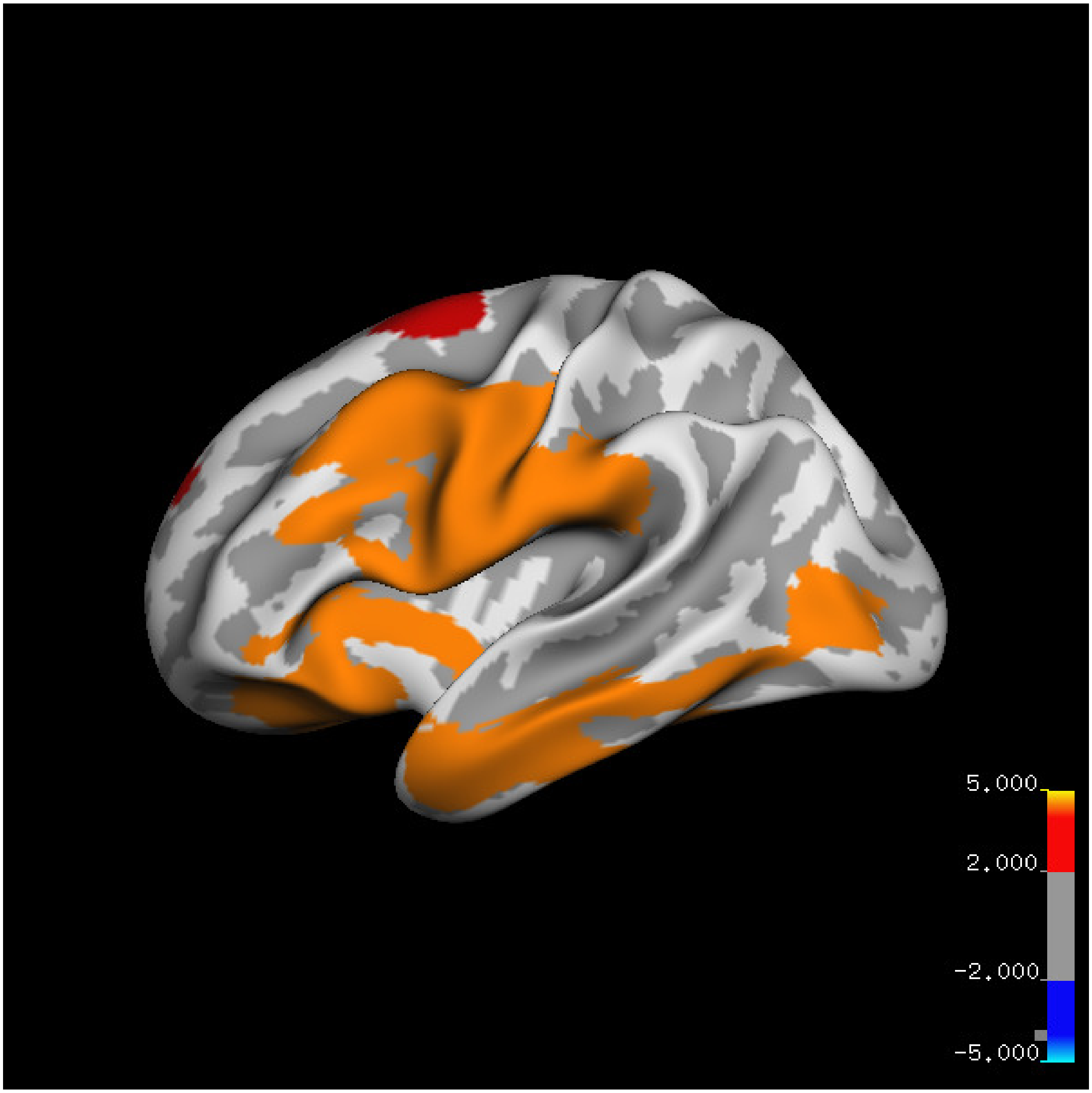
5.000
2.000
-2.000
-5.000

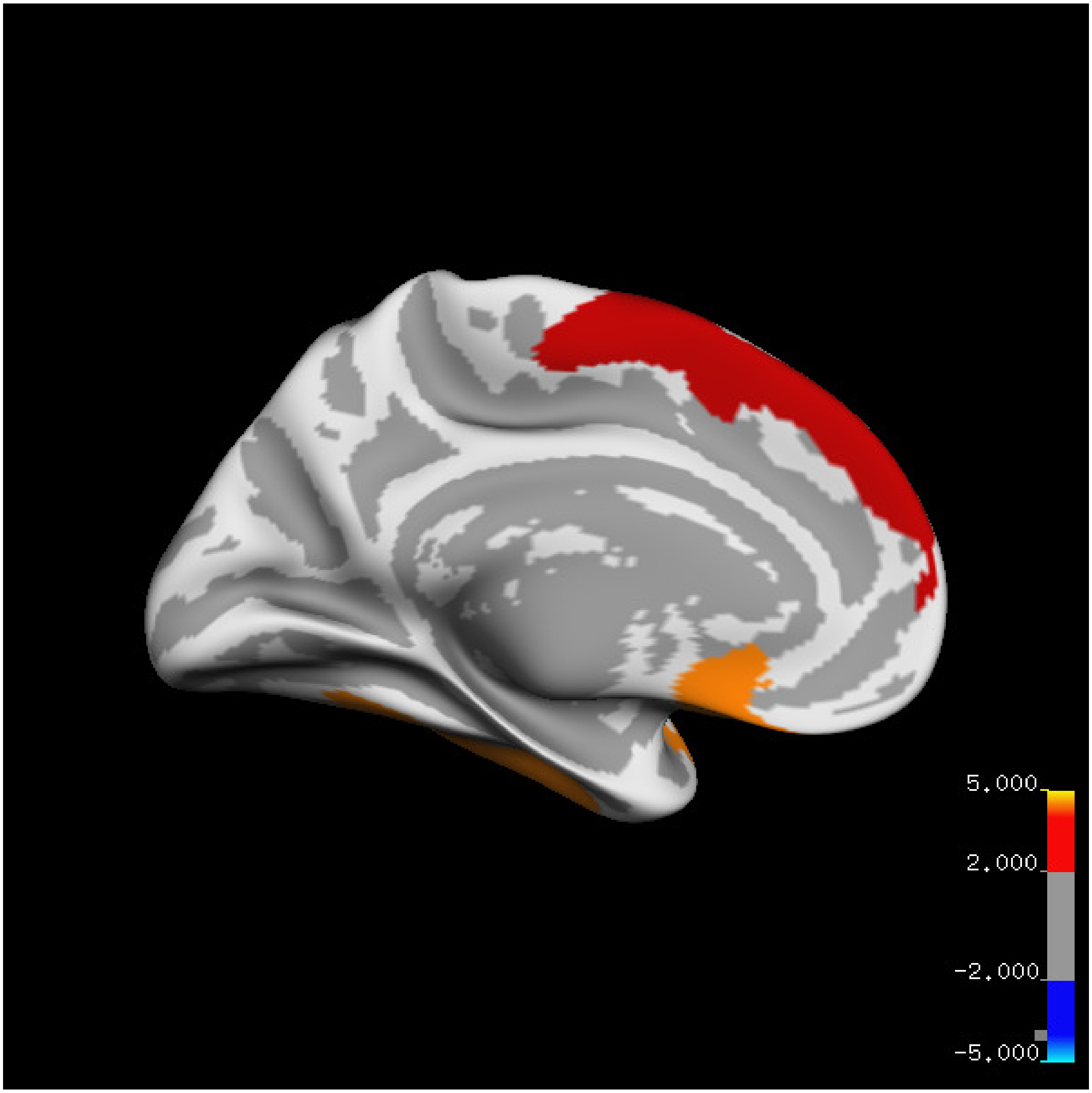

5.000
2.000
-2.000
-5.000

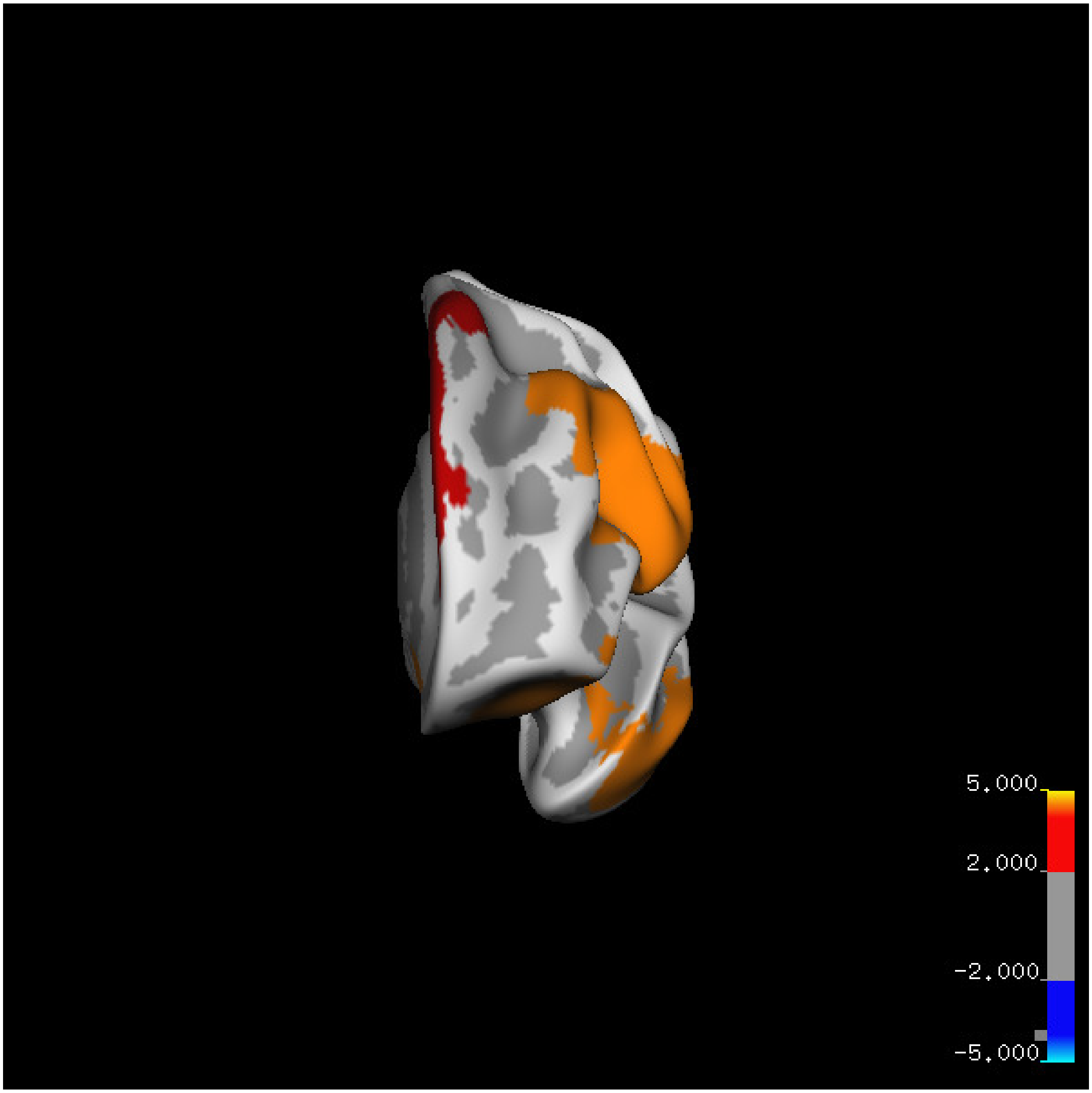
5.000
2.000
-2.000
-5.000

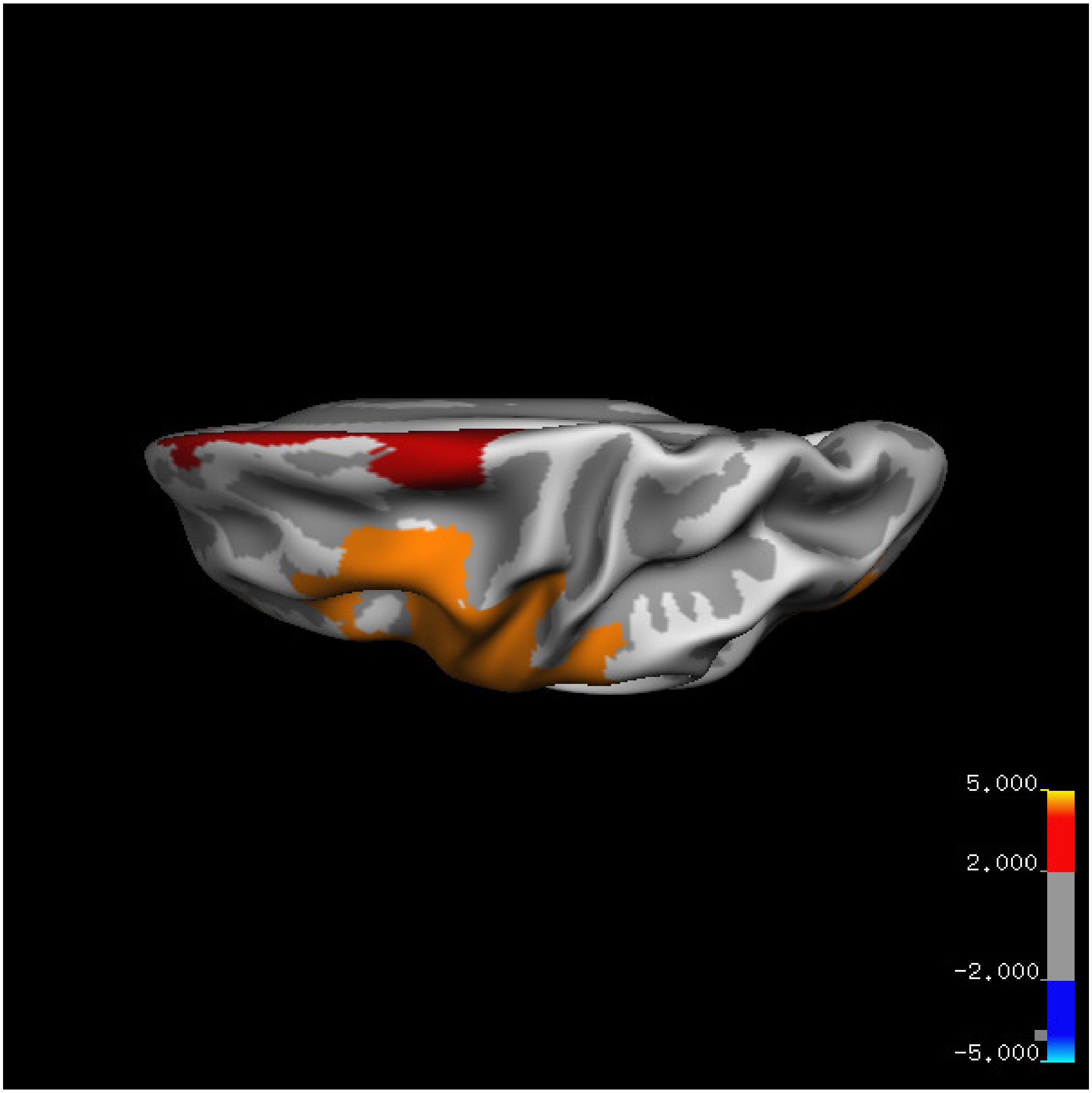
5.000
2.000
-2.000
-5.000

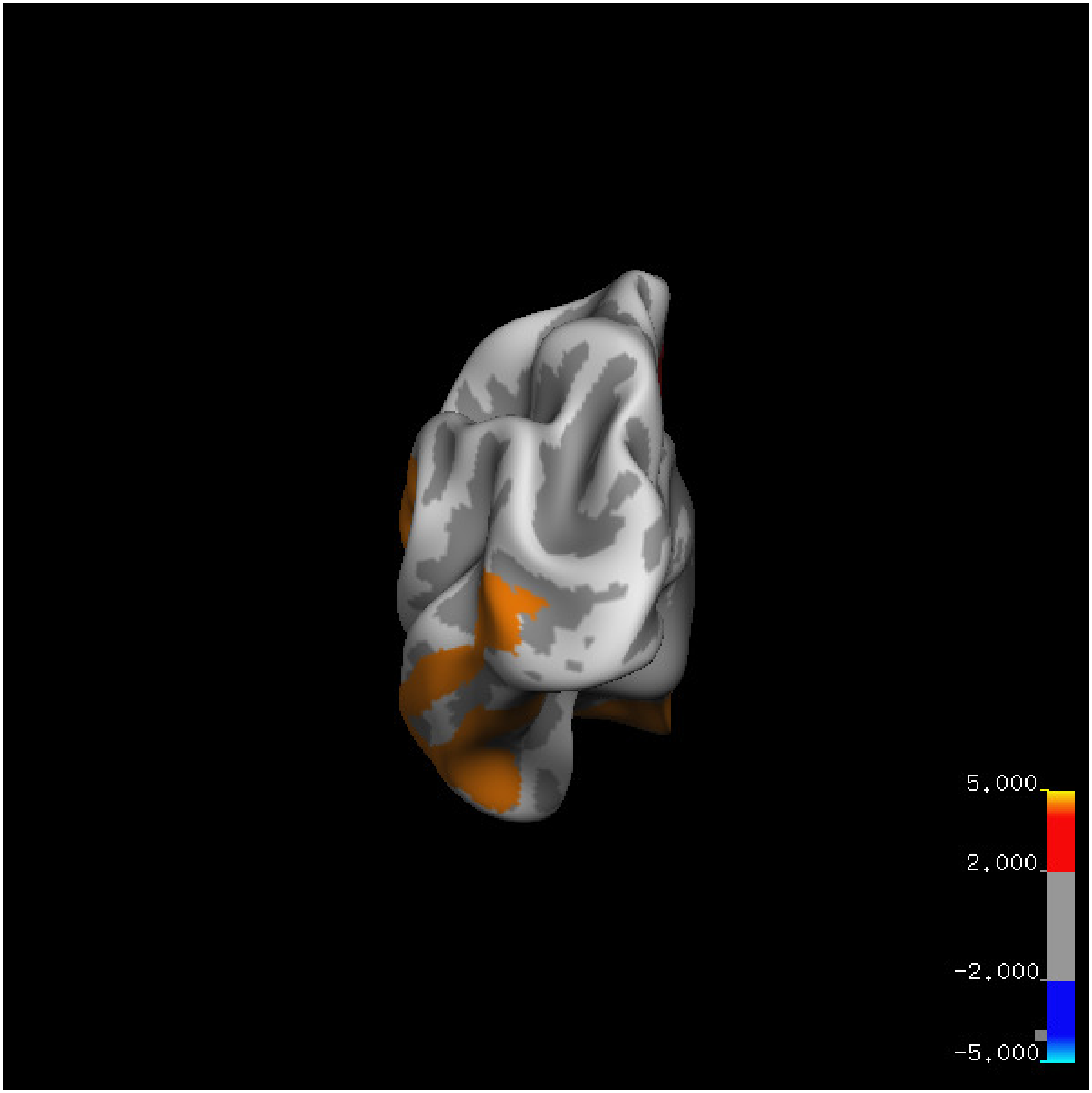
5.000
2.000
-2.000
-5.000

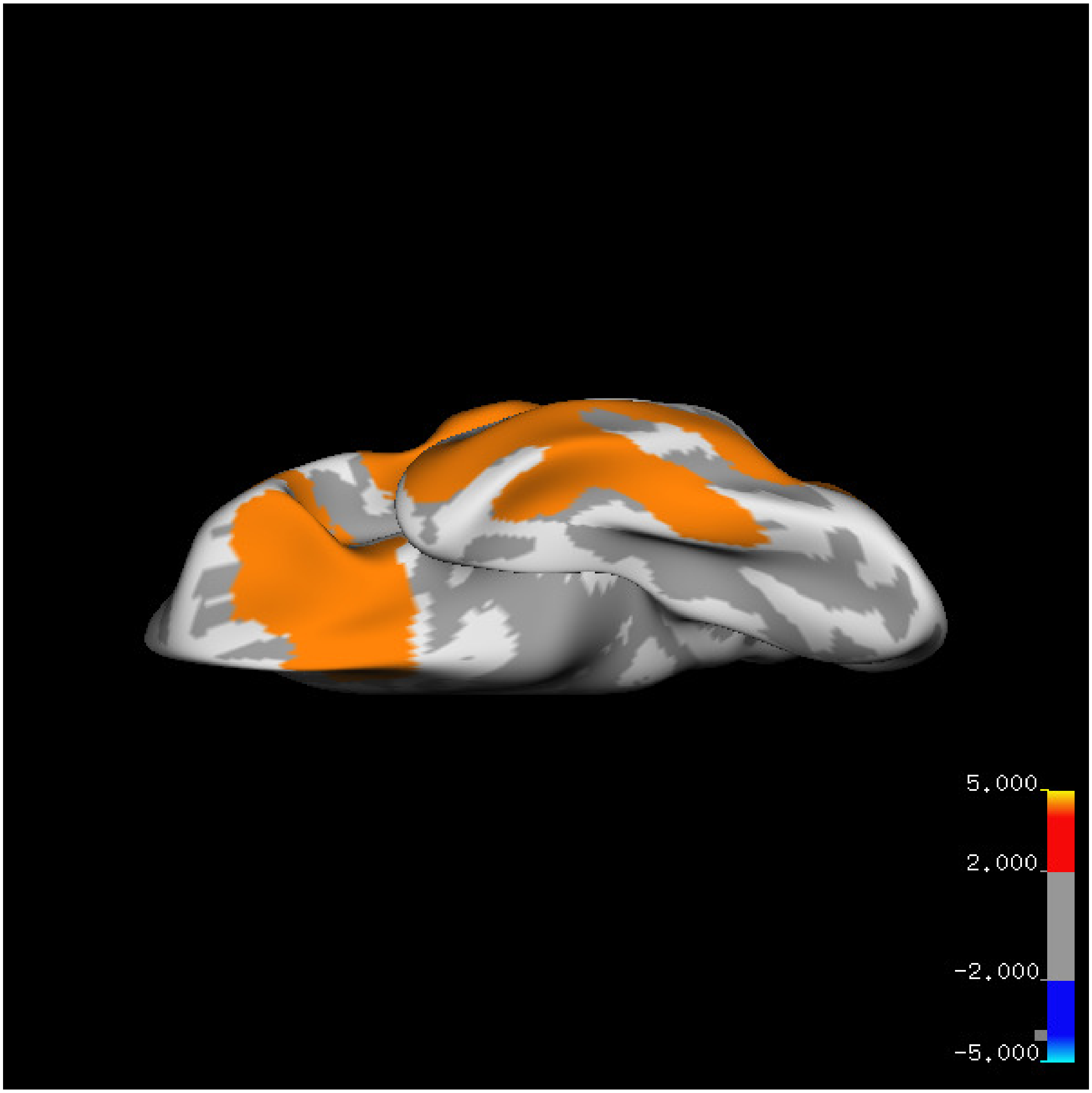
5.000
2.000
-2.000
-5.000

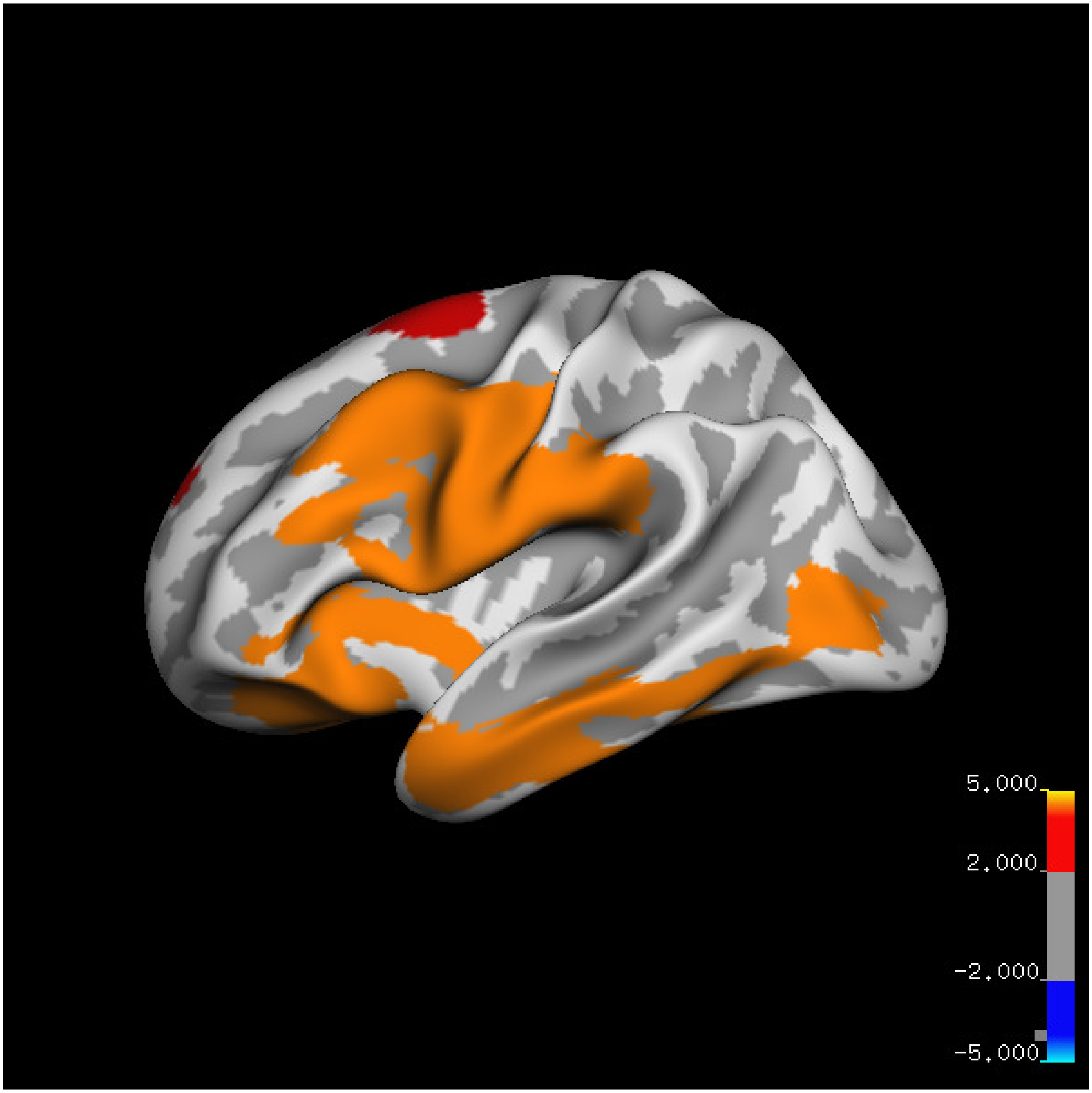
5.000
2.000
-2.000
-5.000

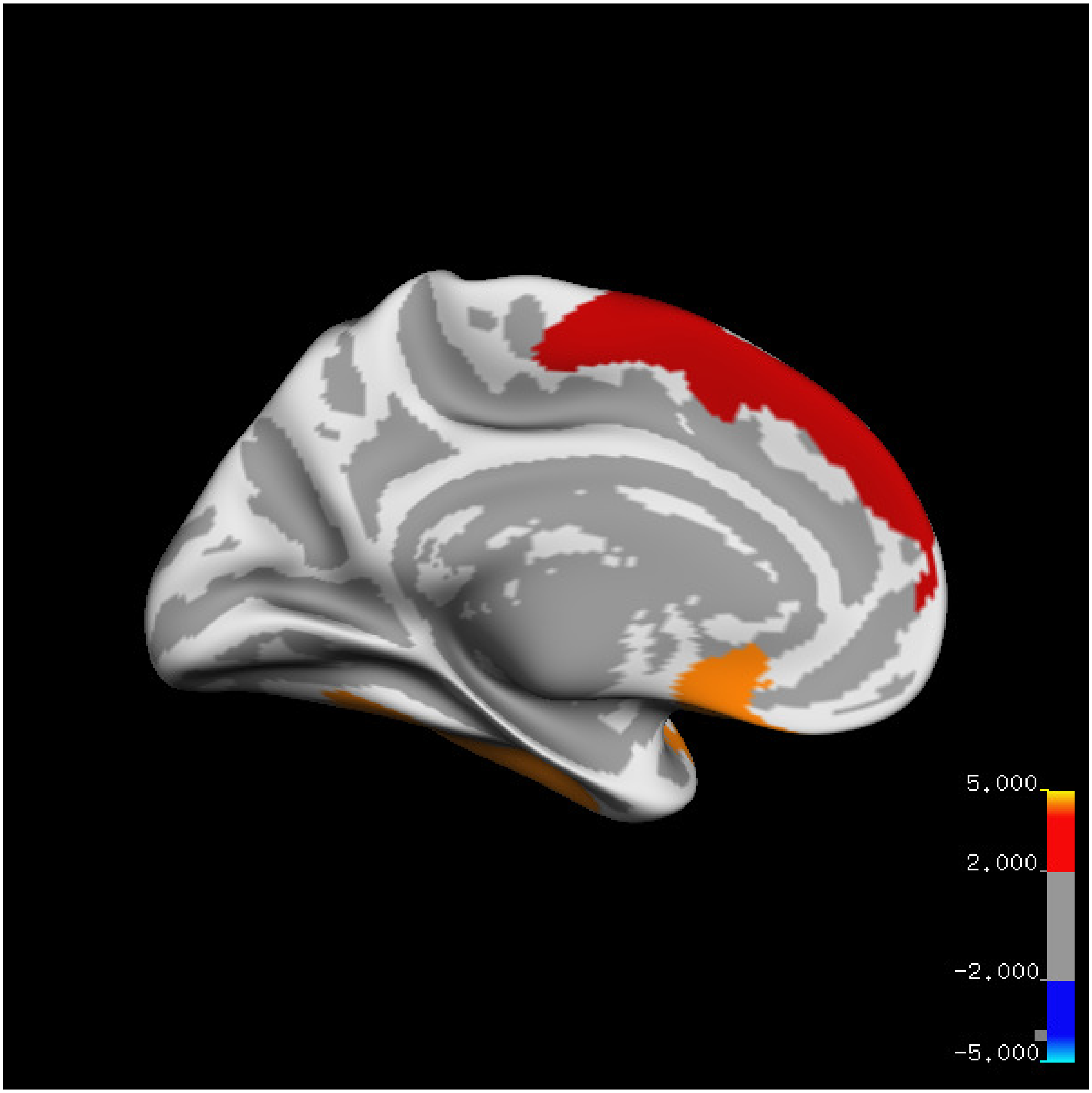
5.000
2.000
-2.000
-5.000

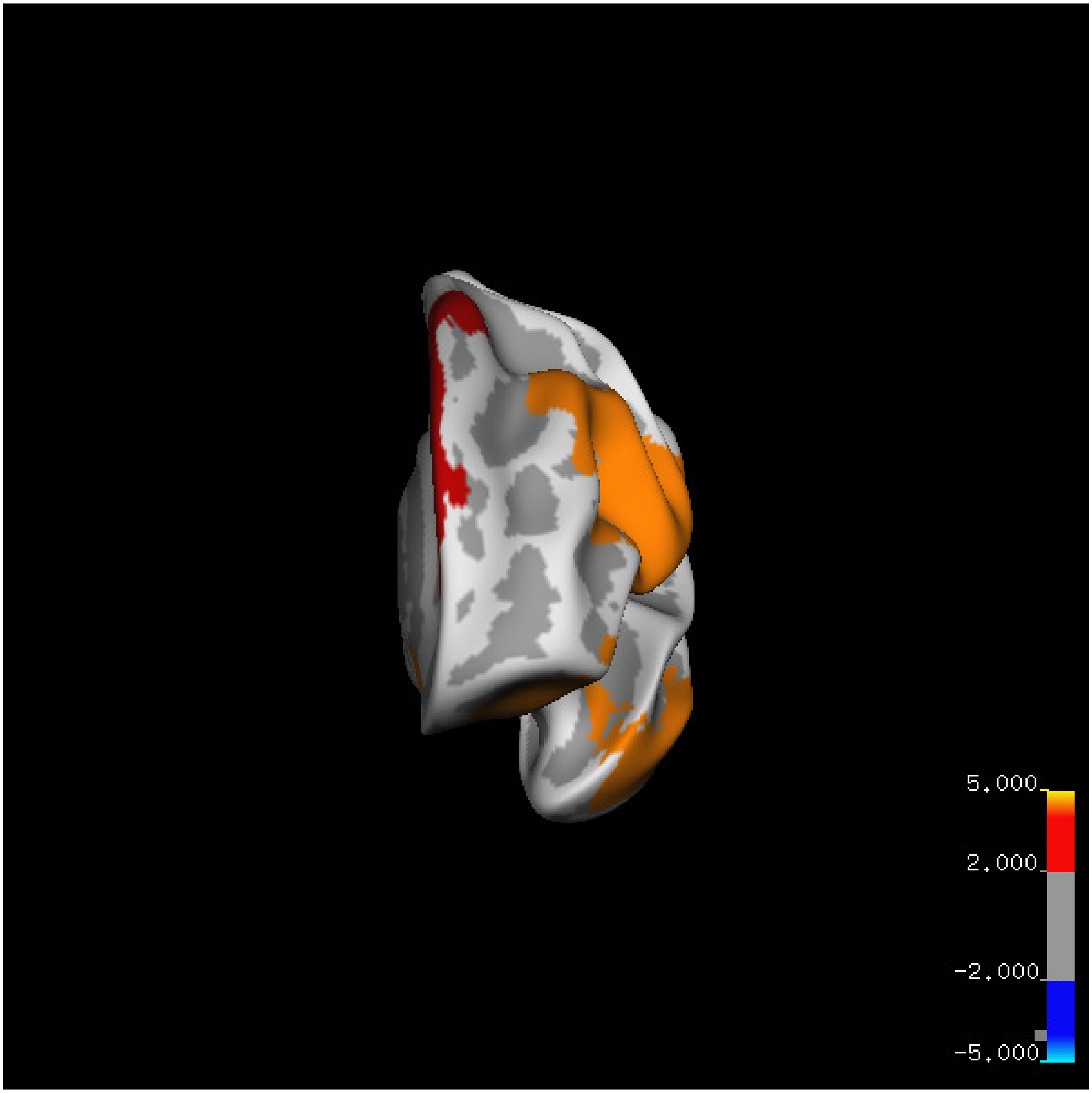
5.000
2.000
-2.000
-5.000

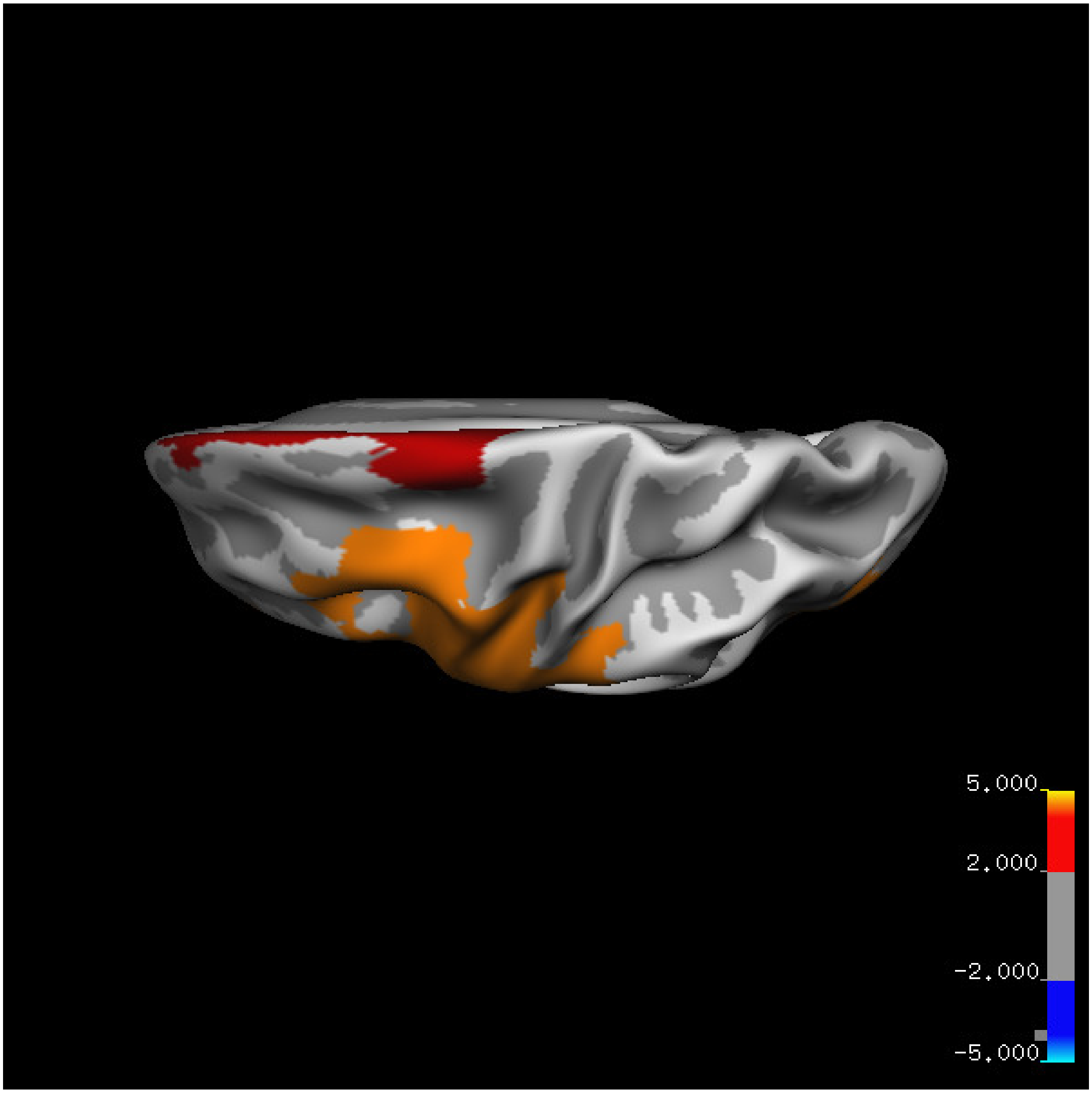


Cluster summary is shown below.

```
## # Cluster Growing Summary (mri_surfcluster)

## # $Id: mri_surfcluster.c,v 1.57.2.3 2016/11/17 18:19:42 zkaufman Exp $

## # $Id: mrisurf.c,v 1.781.2.6 2016/12/27 16:47:14 zkaufman Exp $

## # CreationTime 2021/10/04-13:58:30-GMT

## # cmdline mri_surfcluster.bin --in surface_analyses/results/cluster_comparison/fsaverage6.sm15.lh/vo

## # cwd /gpfs/projects02/p274/cluster/projects/p003-sleep_b

## # sysname  Linux
```

```
## # hostname c1-19
## # machine  x86_64
## # FixVertexAreaFlag 1
## # FixSurfClusterArea 1
## #
## # Input      surface_analyses/results/cluster_comparison/fsaverage6.sm15.lh/volume.above60.below_mea
## # Frame Number      0
## # srcsubj fsaverage6
## # hemi lh
## # surface white
## # group_avg_surface_area 84969.3
## # group_avg_vtxarea_loaded 1
## # annot aparc
## # SUBJECTS_DIR /cluster/projects/p274/tools/mri/freesurfer/current/subjects
## # SearchSpace_mm2 79241.3
## # SearchSpace_vtx 37476
## # Bonferroni 0
## # Minimum Threshold 1.3
## # Maximum Threshold infinity
## # Threshold Sign    abs
## # AdjustThreshWhenOneTail 1
## # CW PValue Threshold: 0.01
## # Area Threshold    0 mm^2
## # CSD thresh  1.300000
## # CSD nreps    10000
## # CSD simtype  null-z
## # CSD contrast NA
```

```
## # CSD confint  90.000000

## # Overall max 6.90806 at vertex 20247

## # Overall min -1.32982 at vertex 6366

## # NClusters          3

## # FixMNI = 0

## #

## # ClusterNo  Max   VtxMax   Size(mm^2)  MNIX   MNIY   MNIZ    CWP    CWPLow    CWPHi   NVtxs    WghtV

##    1        6.908   20247   9043.97    -29.8   29.5  -16.1  0.00010  0.00000  0.00020   3894     9008

##    2        5.249   11686   7737.53    -47.6  -10.2   42.4  0.00010  0.00000  0.00020   3978     9250

##    3        4.352   30728   2549.92     -6.1   -2.4   61.1  0.00360  0.00280  0.00440   1192     2645
```

#### Right hemisphere

##### Age 20-60

**Sleep above mean** Cluster corrected significance plots are shown below.

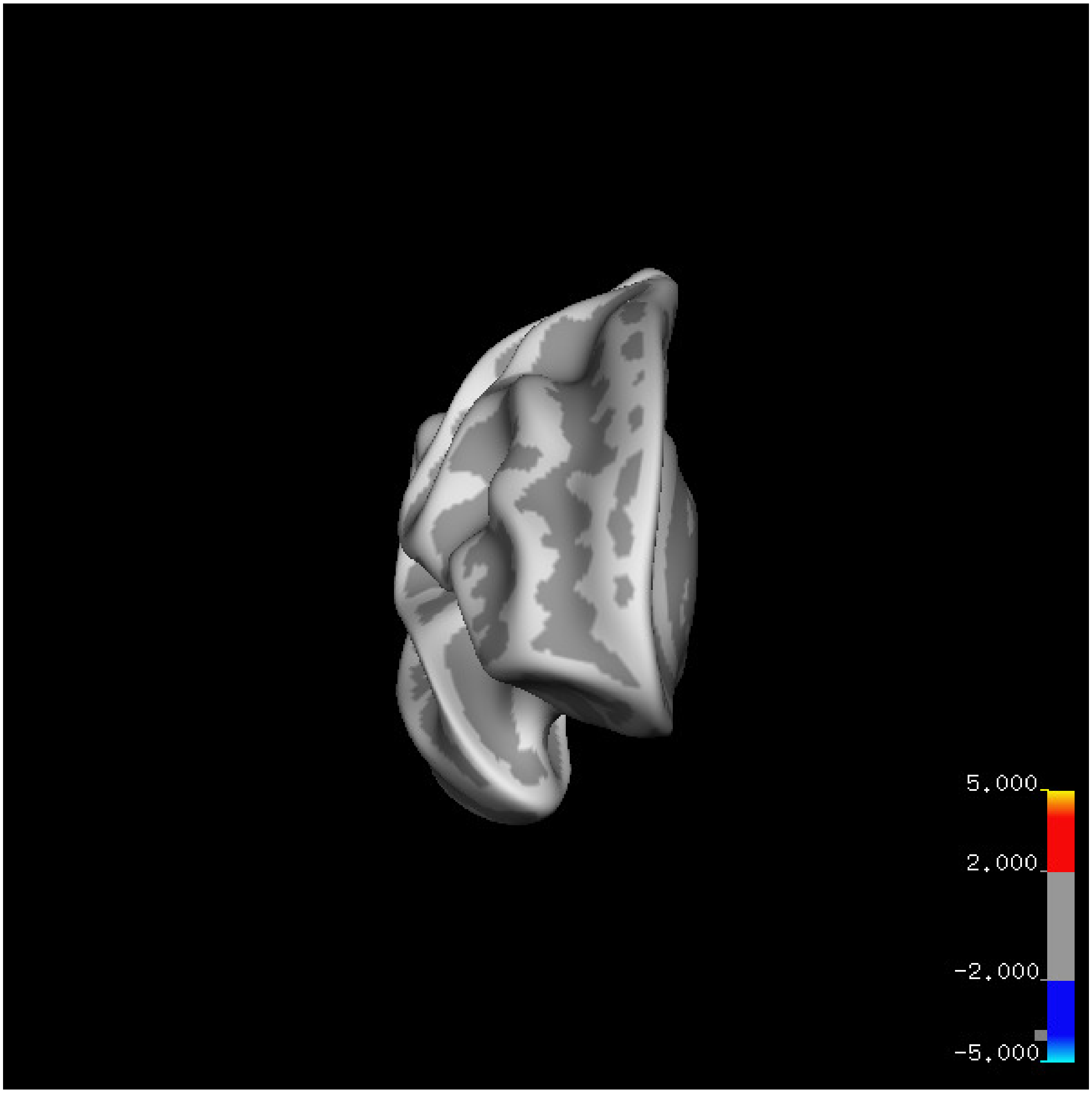
5.000
2.000
-2.000
-5.000

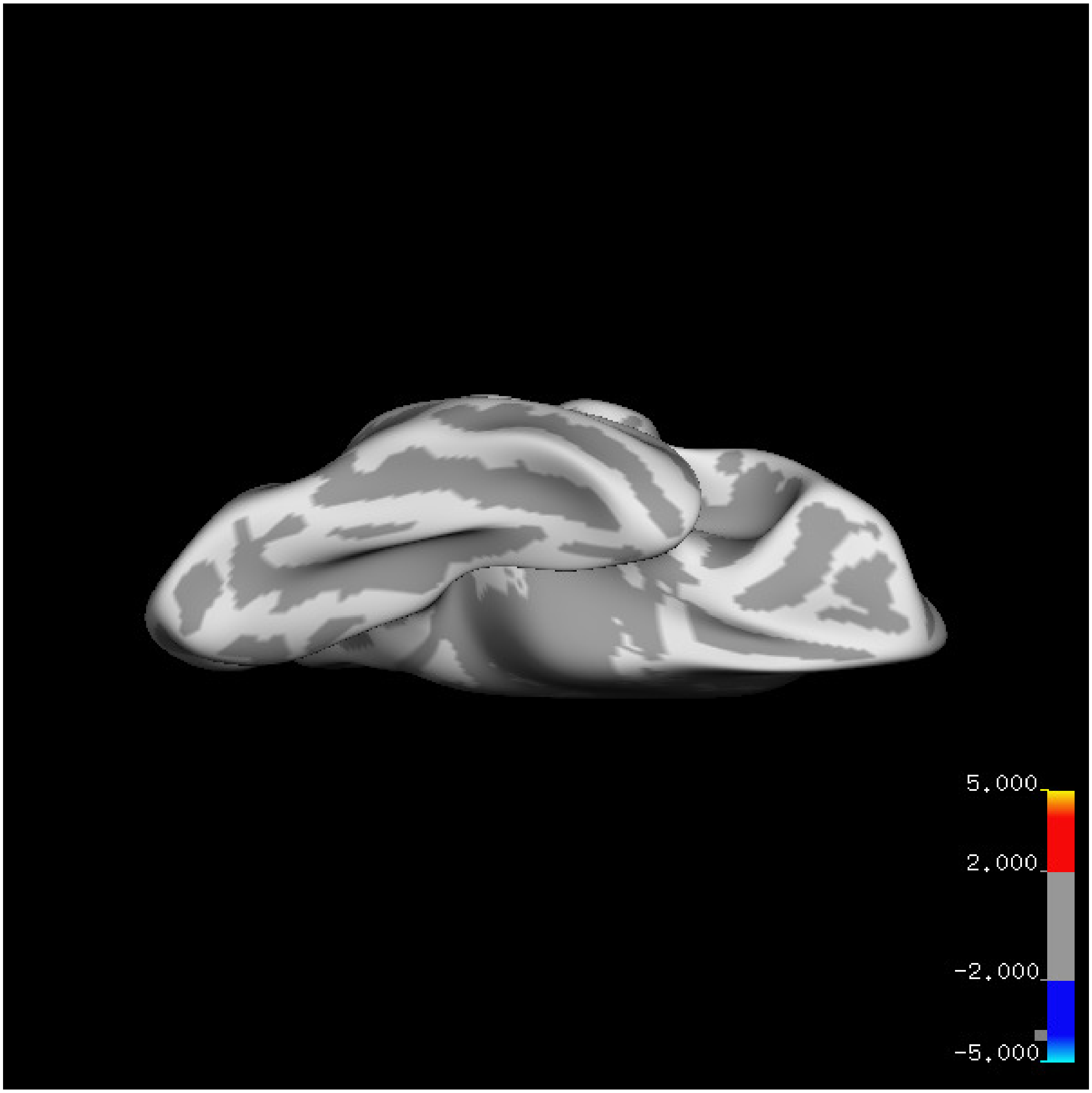
5.000
2.000
-2.000
-5.000

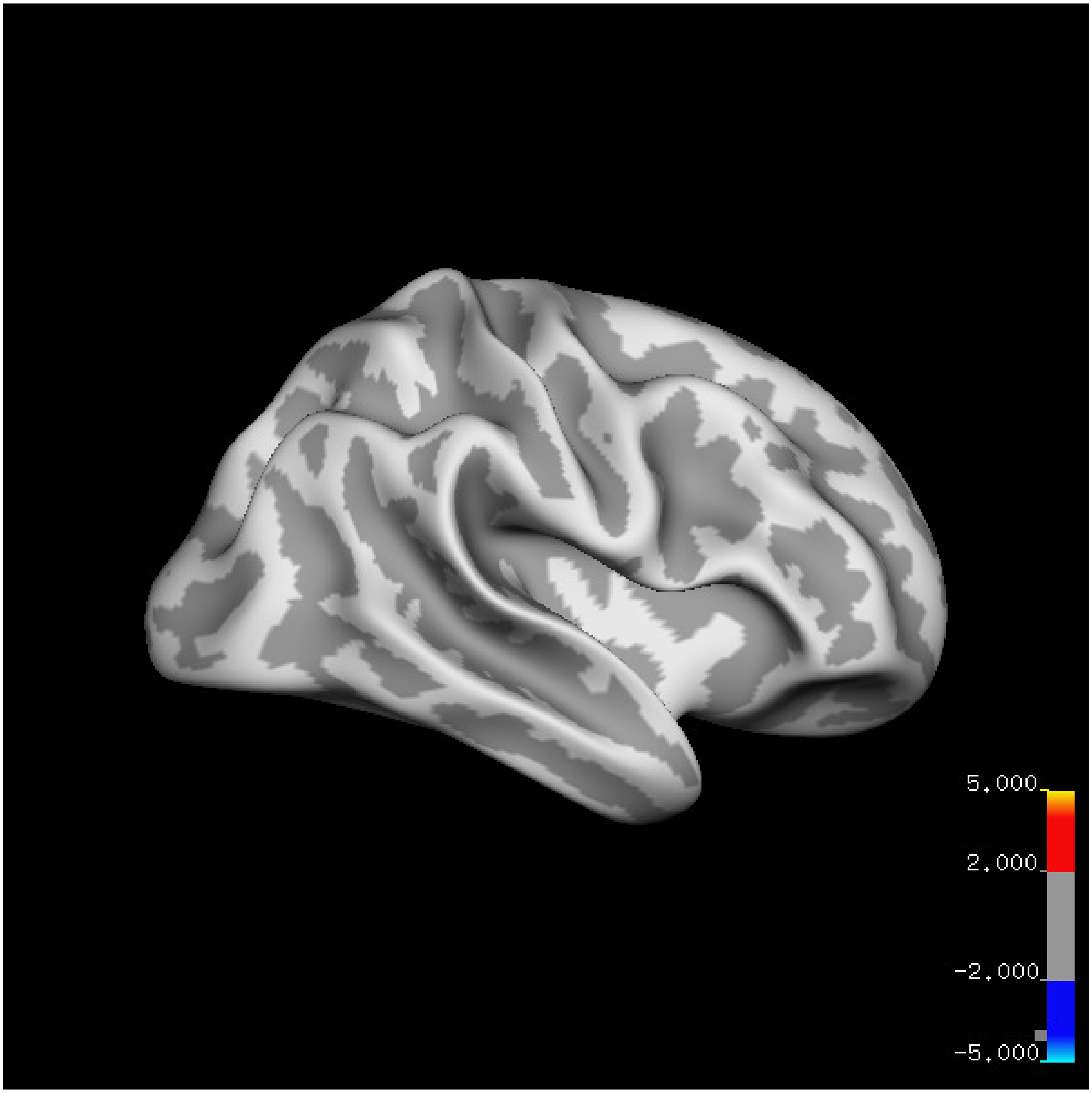

5.000
2.000
-2.000
-5.000

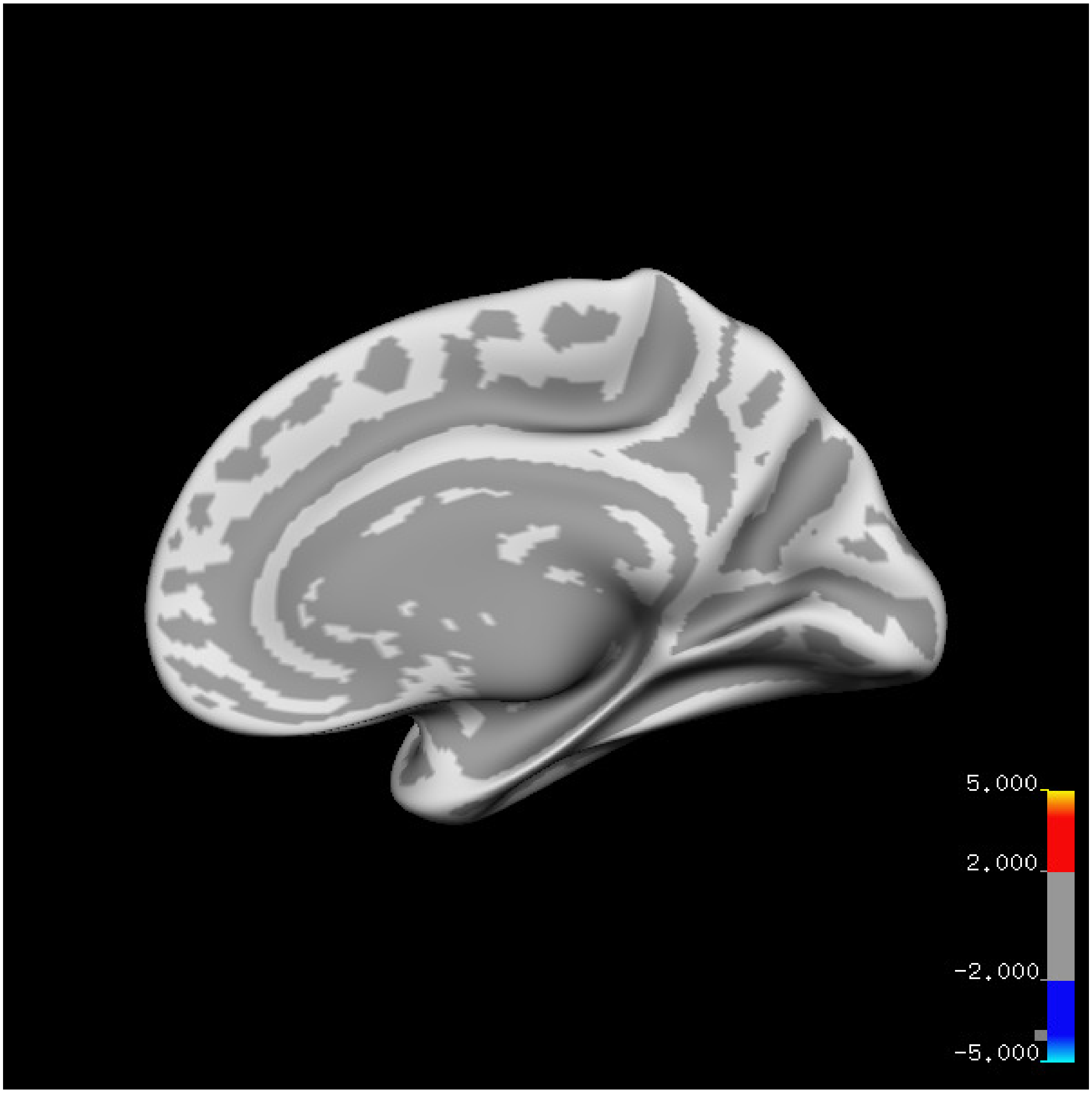
5.000
2.000
-2.000
-5.000

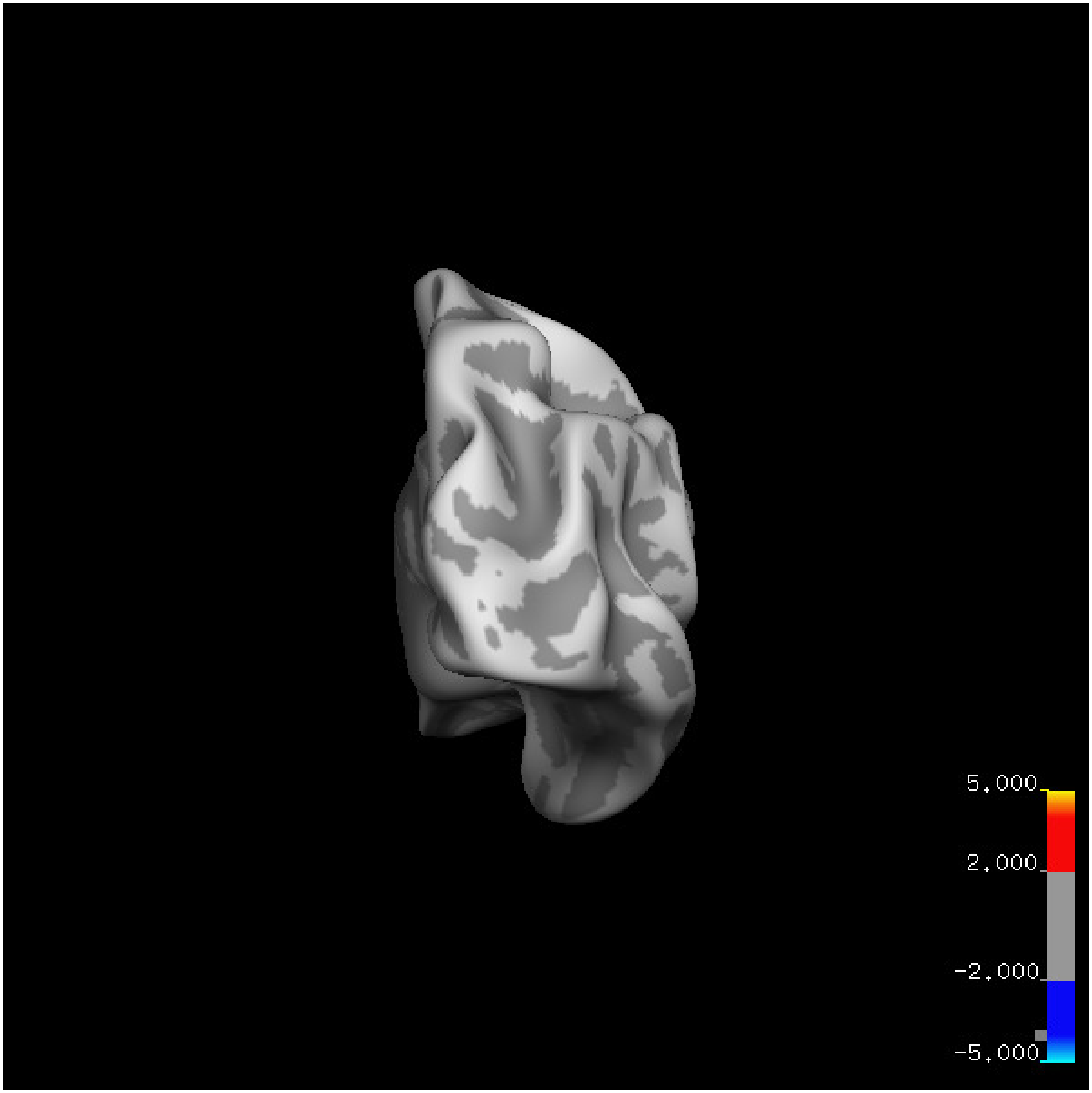

5.000
2.000
-2.000
-5.000

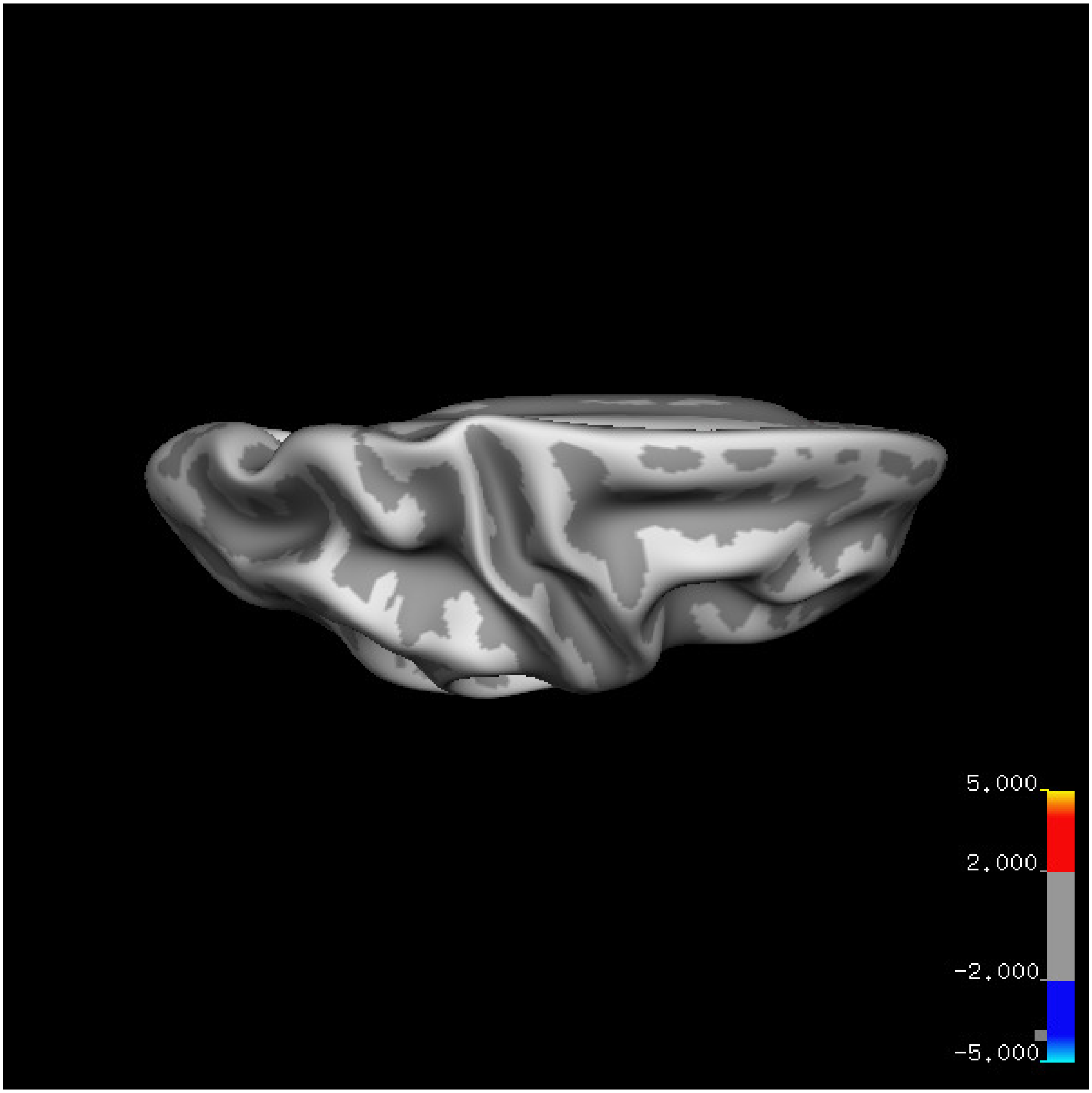
5.000
2.000
-2.000
-5.000

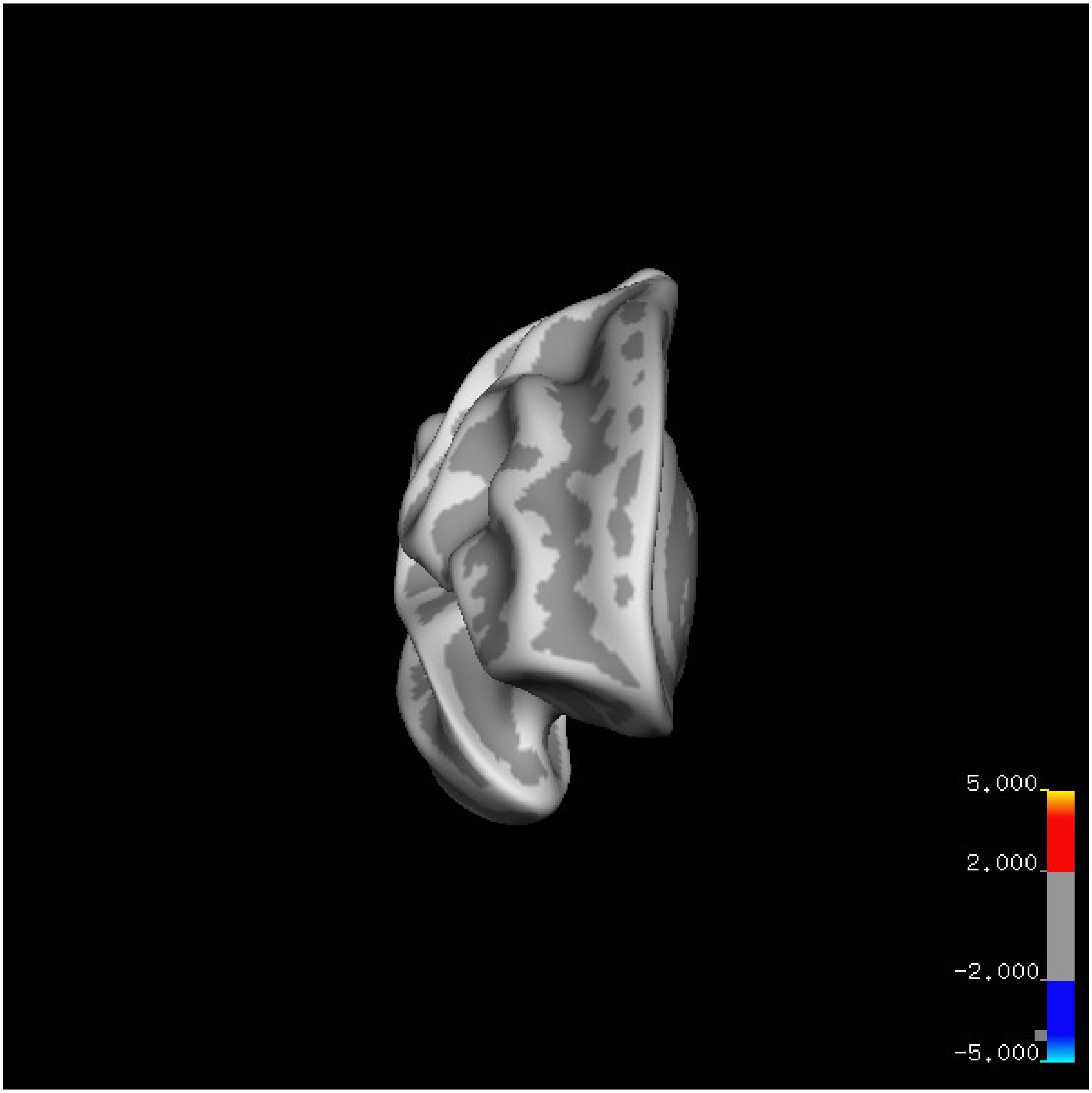
5.000
2.000
-2.000
-5.000

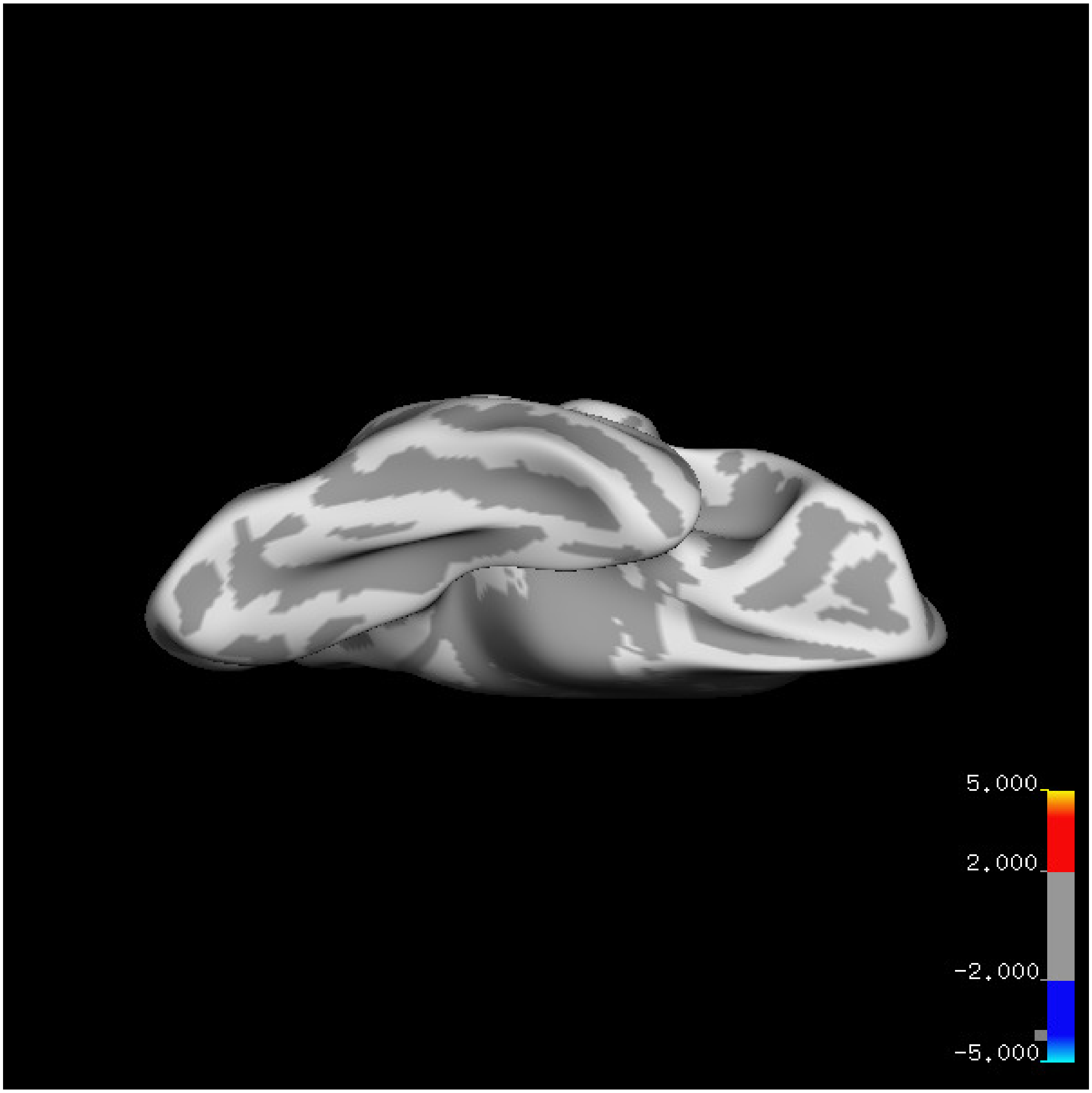
5.000
2.000
-2.000
-5.000

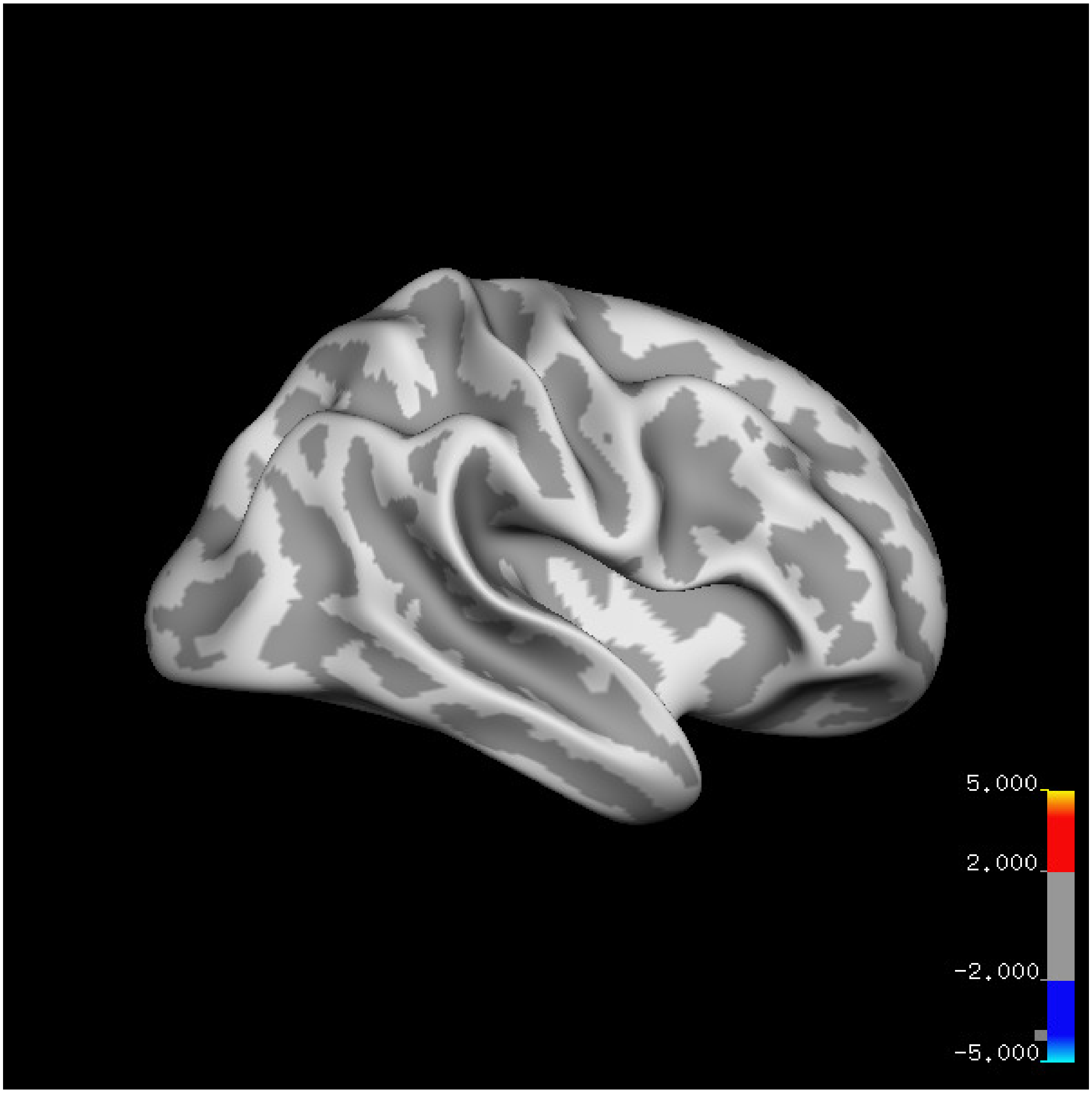
5.000
2.000
-2.000
-5.000

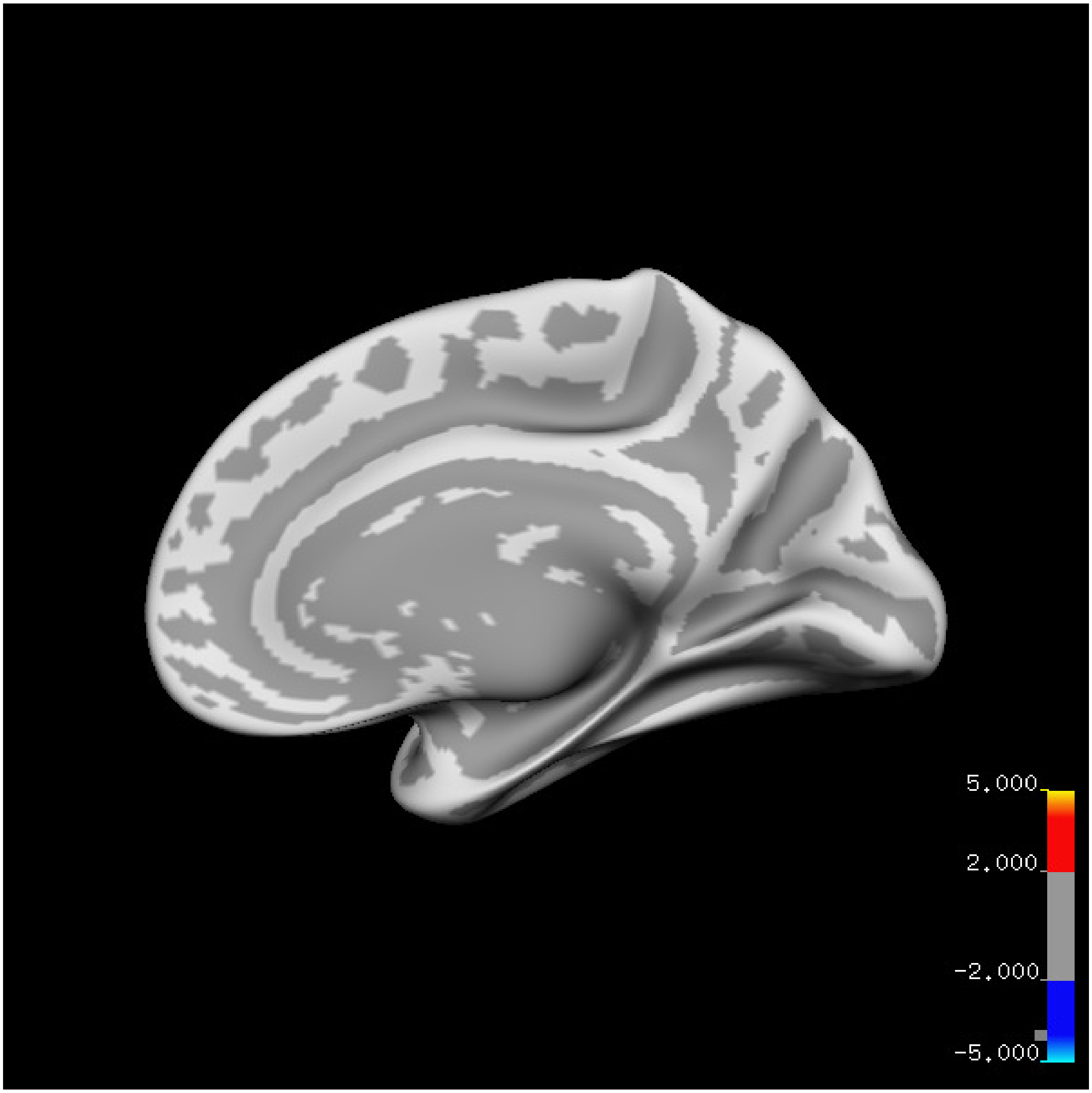
5.000
2.000
-2.000
-5.000

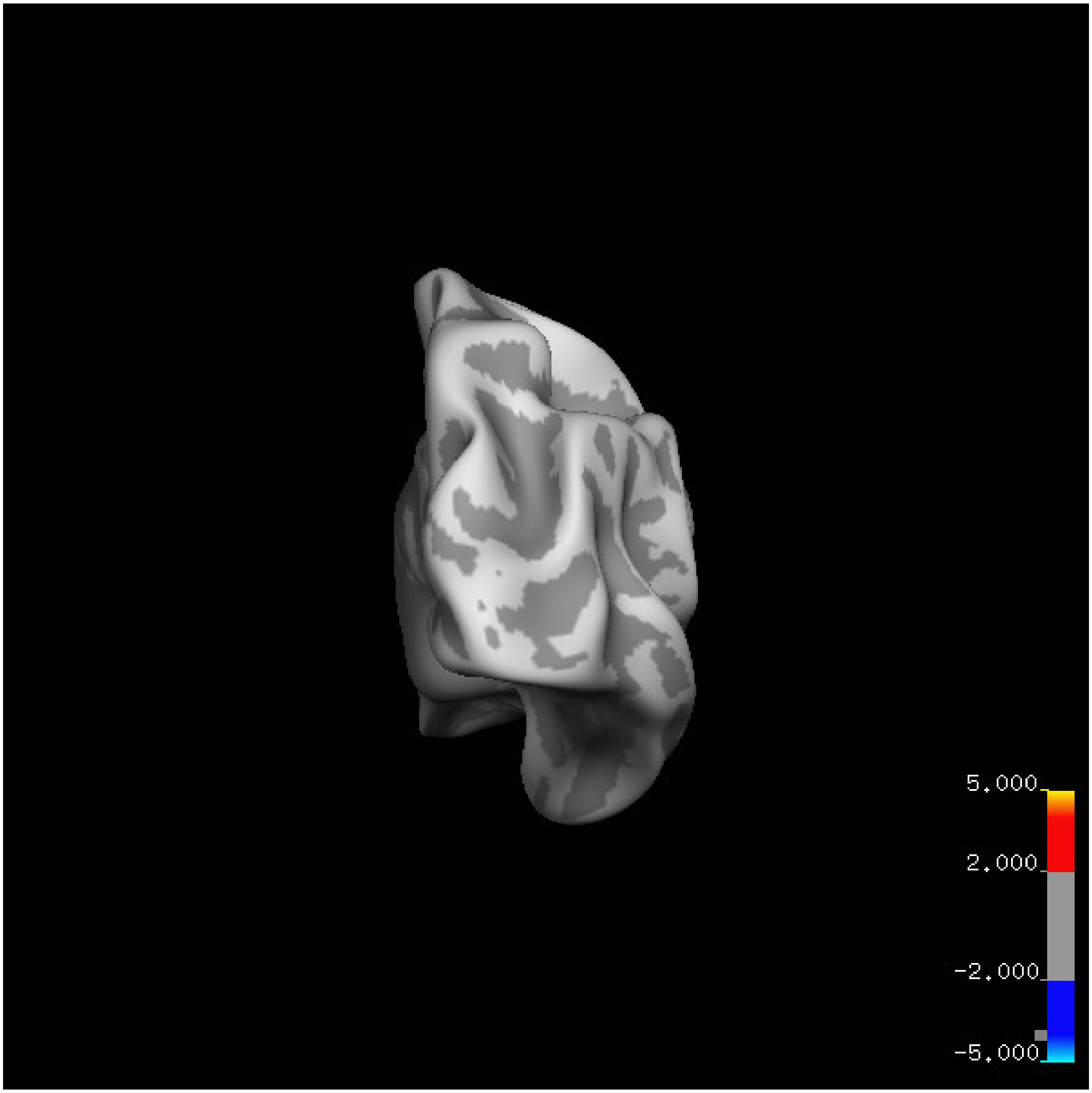
5.000
2.000
-2.000
-5.000

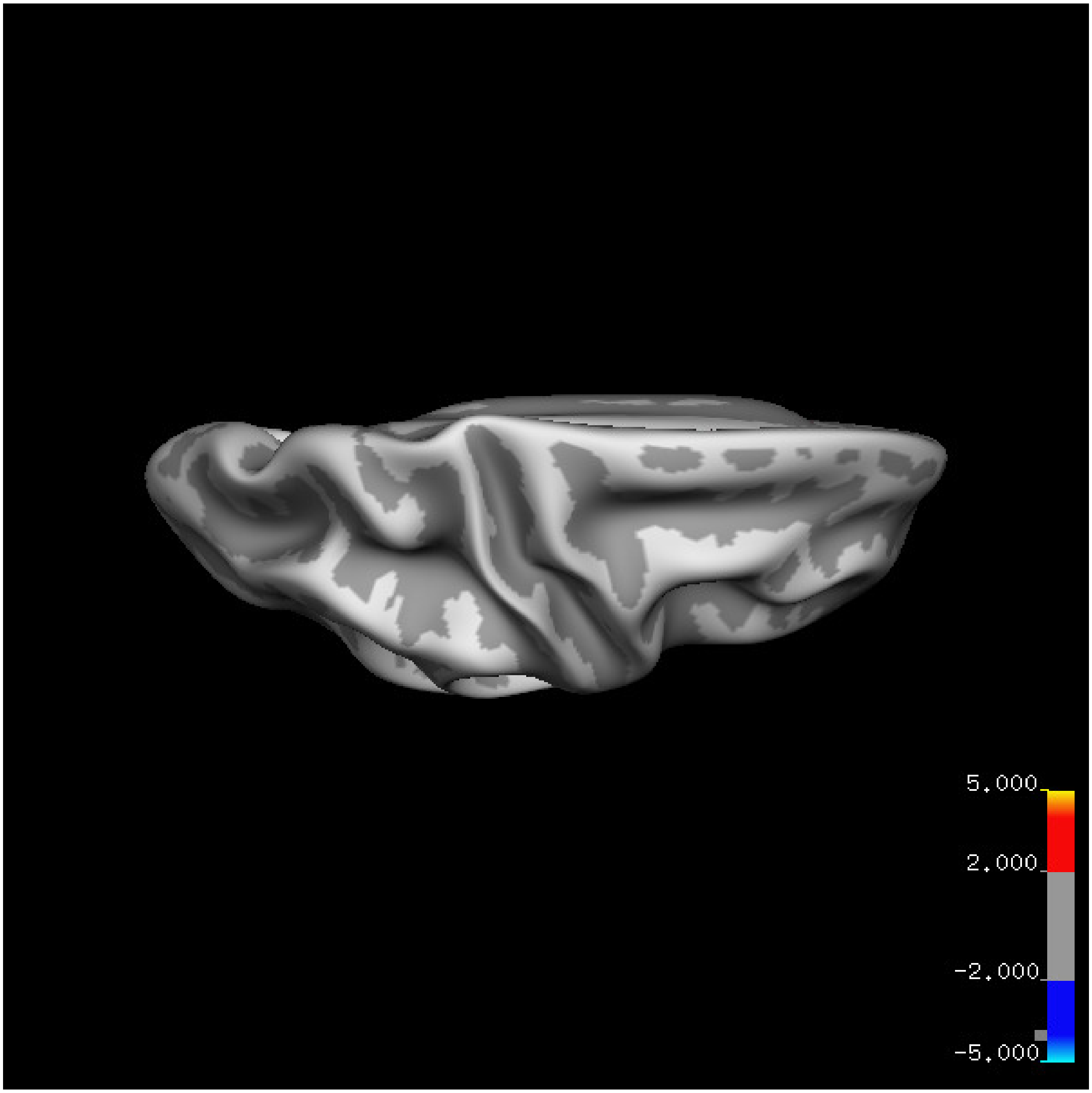


Cluster summary is shown below.

```
## # Cluster Growing Summary (mri_surfcluster)

## # $Id: mri_surfcluster.c,v 1.57.2.3 2016/11/17 18:19:42 zkaufman Exp $

## # $Id: mrisurf.c,v 1.781.2.6 2016/12/27 16:47:14 zkaufman Exp $

## # CreationTime 2021/10/04-10:48:00-GMT

## # cmdline mri_surfcluster.bin --in surface_analyses/results/cluster_comparison/fsaverage6.sm15.rh/vol

## # cwd /gpfs/projects02/p274/cluster/projects/p003-sleep_b

## # sysname  Linux
```

```
## # hostname c1-20
## # machine  x86_64
## # FixVertexAreaFlag 1
## # FixSurfClusterArea 1
## #
## # Input      surface_analyses/results/cluster_comparison/fsaverage6.sm15.rh/volume.20_60.above_mean/g
## # Frame Number      0
## # srcsubj fsaverage6
## # hemi rh
## # surface white
## # group_avg_surface_area 85131.2
## # group_avg_vtxarea_loaded 1
## # annot aparc
## # SUBJECTS_DIR /cluster/projects/p274/tools/mri/freesurfer/current/subjects
## # SearchSpace_mm2 79329.9
## # SearchSpace_vtx 37471
## # Bonferroni 0
## # Minimum Threshold 1.3
## # Maximum Threshold infinity
## # Threshold Sign    abs
## # AdjustThreshWhenOneTail 1
## # CW PValue Threshold: 0.01
## # Area Threshold    0 mm^2
## # CSD thresh  1.300000
## # CSD nreps    10000
## # CSD simtype  null-z
## # CSD contrast NA
```

```
## # CSD confint  90.000000

## # Overall max 1.44439 at vertex 21799

## # Overall min -3.67192 at vertex 5227

## # NClusters          0

## # FixMNI = 0

## #

## # ClusterNo  Max   VtxMax   Size(mm^2)  MNIX   MNIY   MNIZ    CWP    CWPLow    CWPHi   NVtxs    Wght
```

**Sleep below mean** Cluster corrected significance plots are shown below.

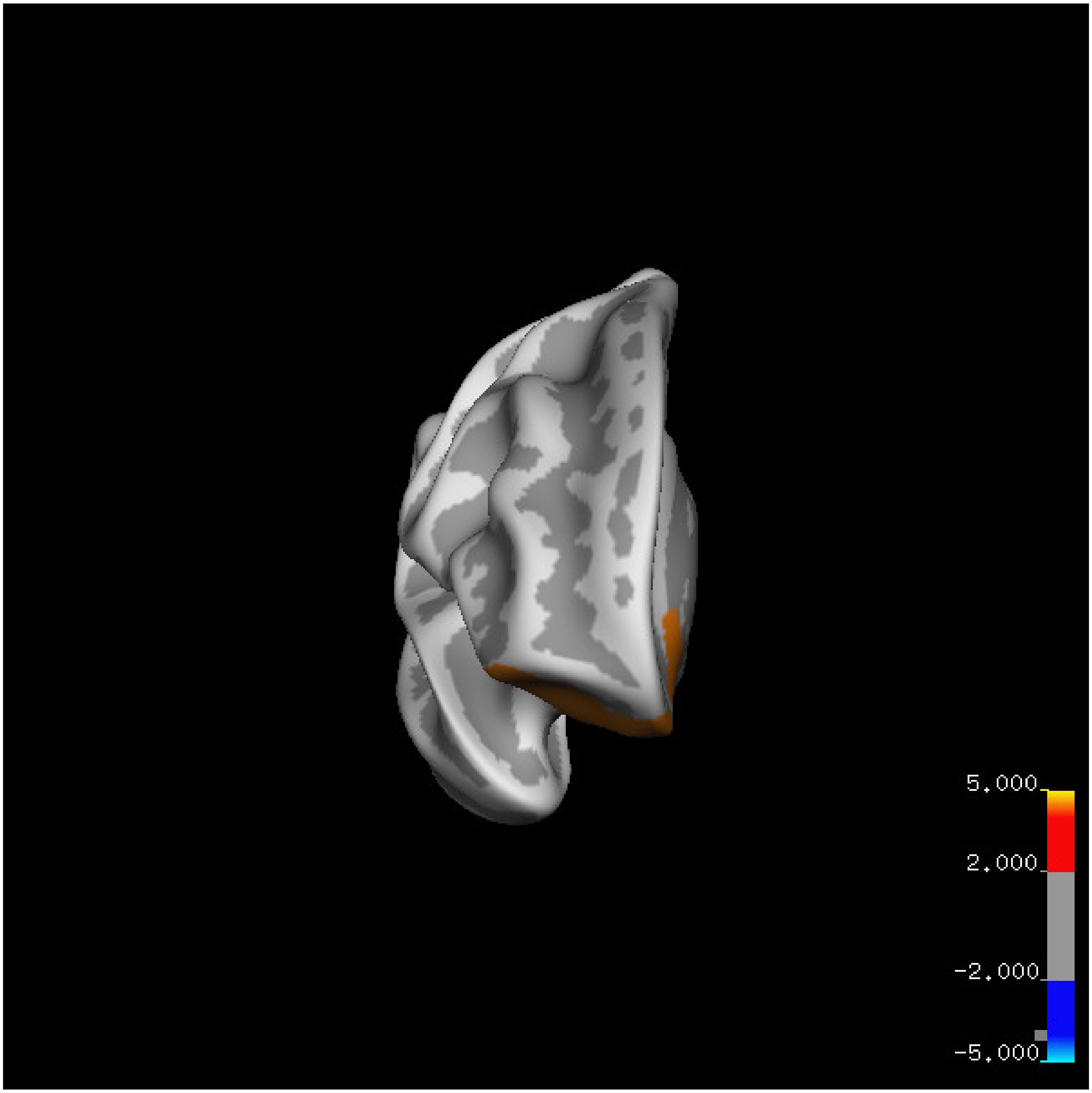
5.000
2.000
-2.000
-5.000

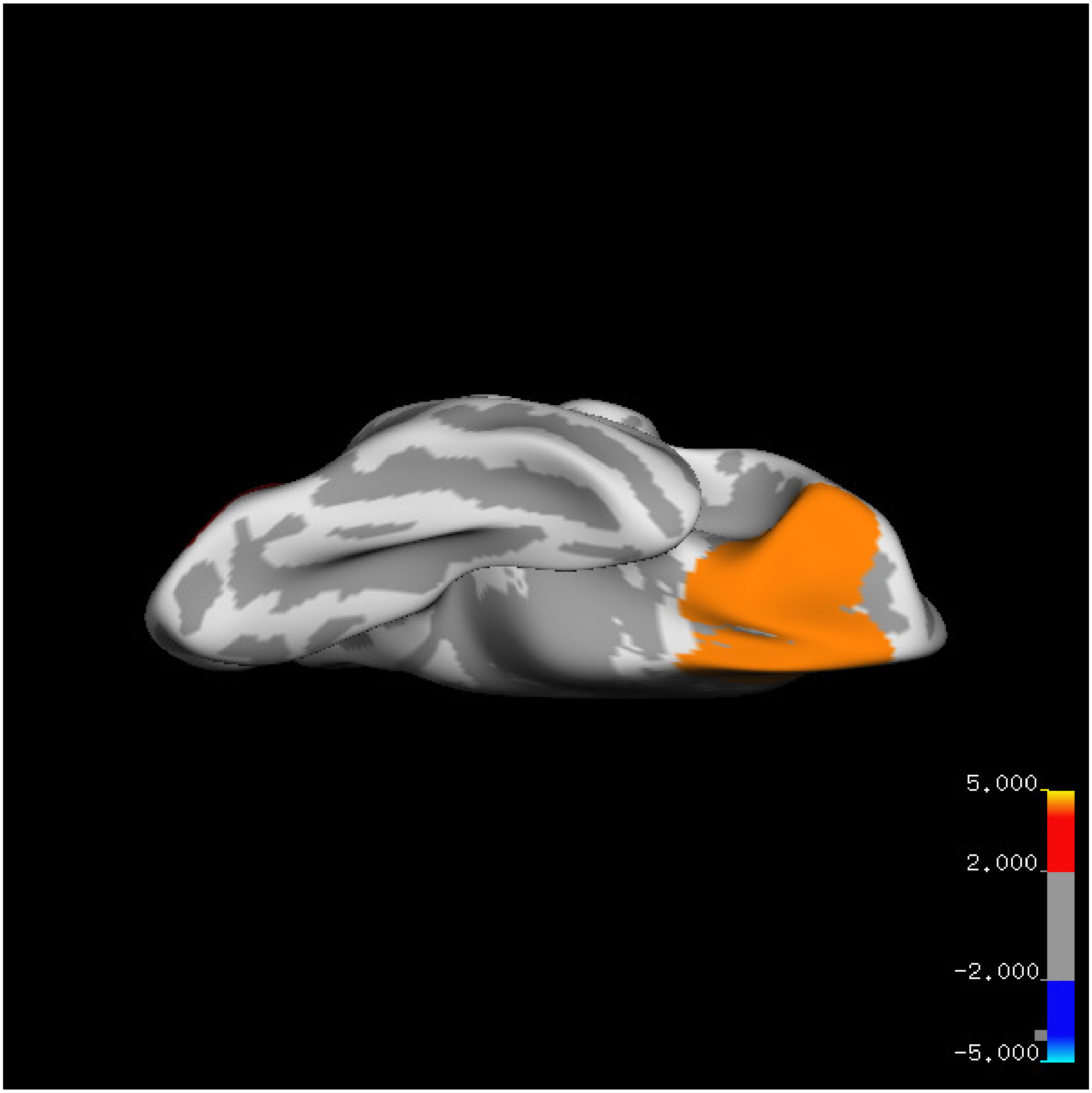
5.000
2.000
-2.000
-5.000

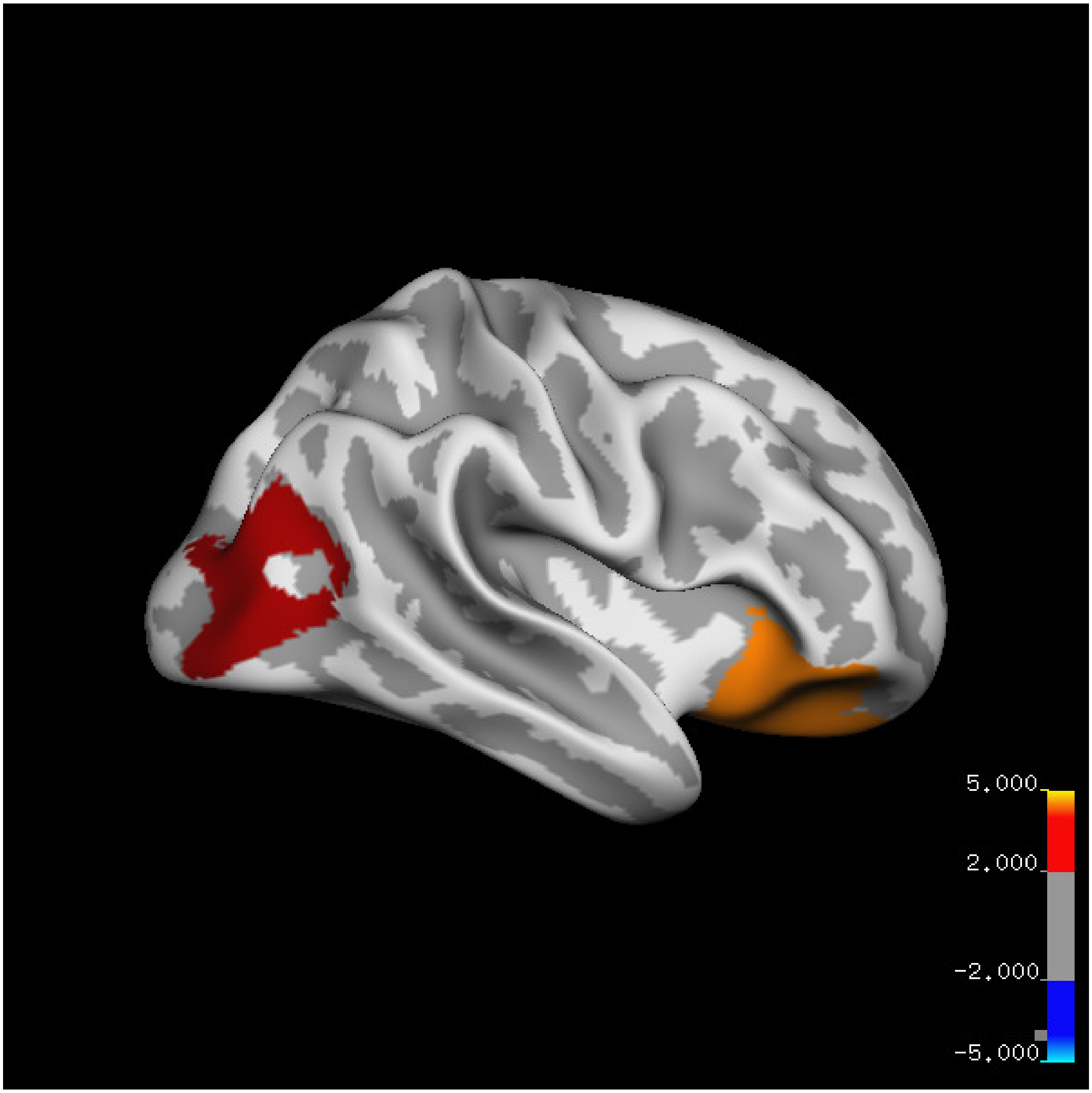
5.000
2.000
-2.000
-5.000

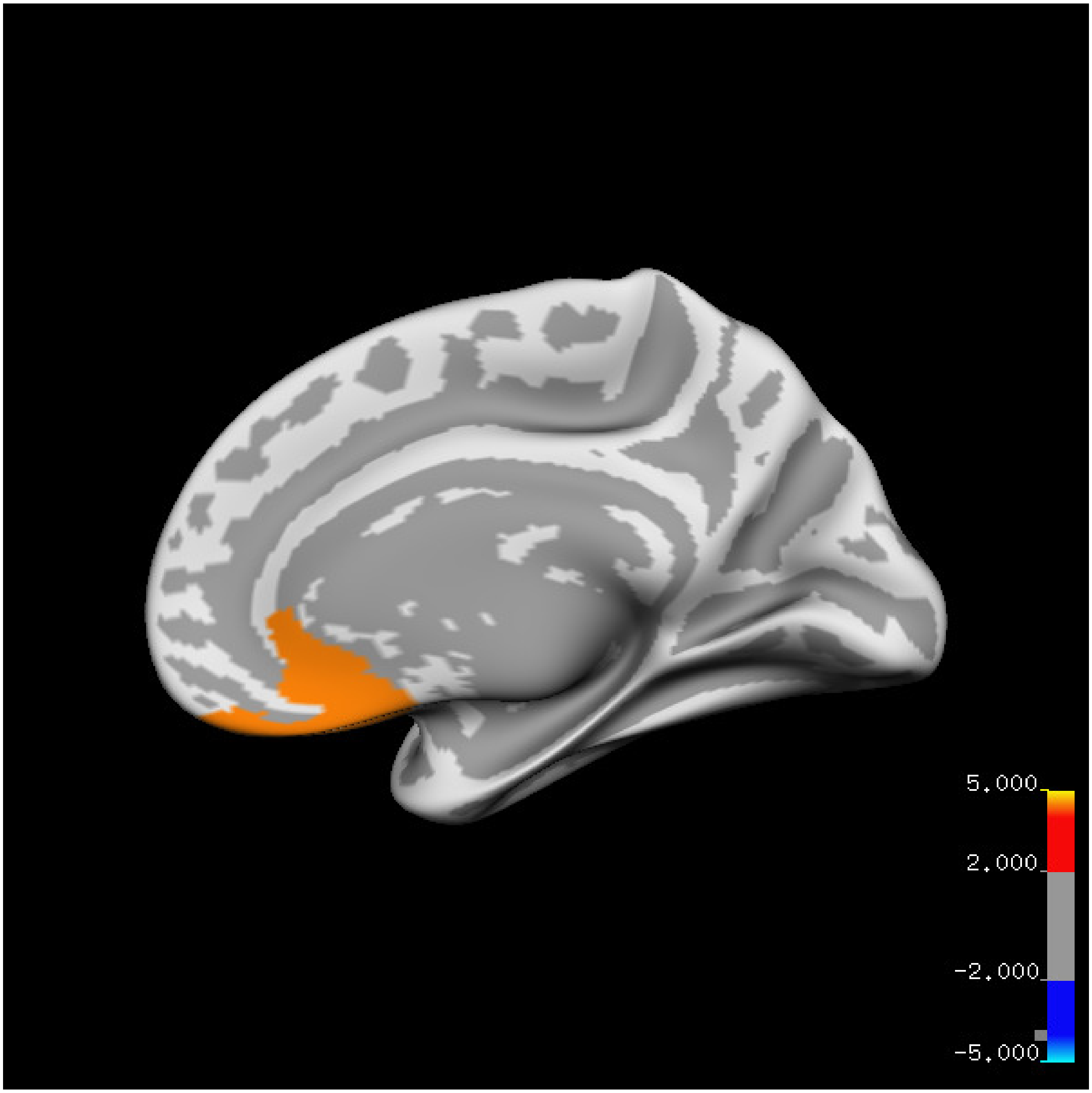
5.000
2.000
-2.000
-5.000

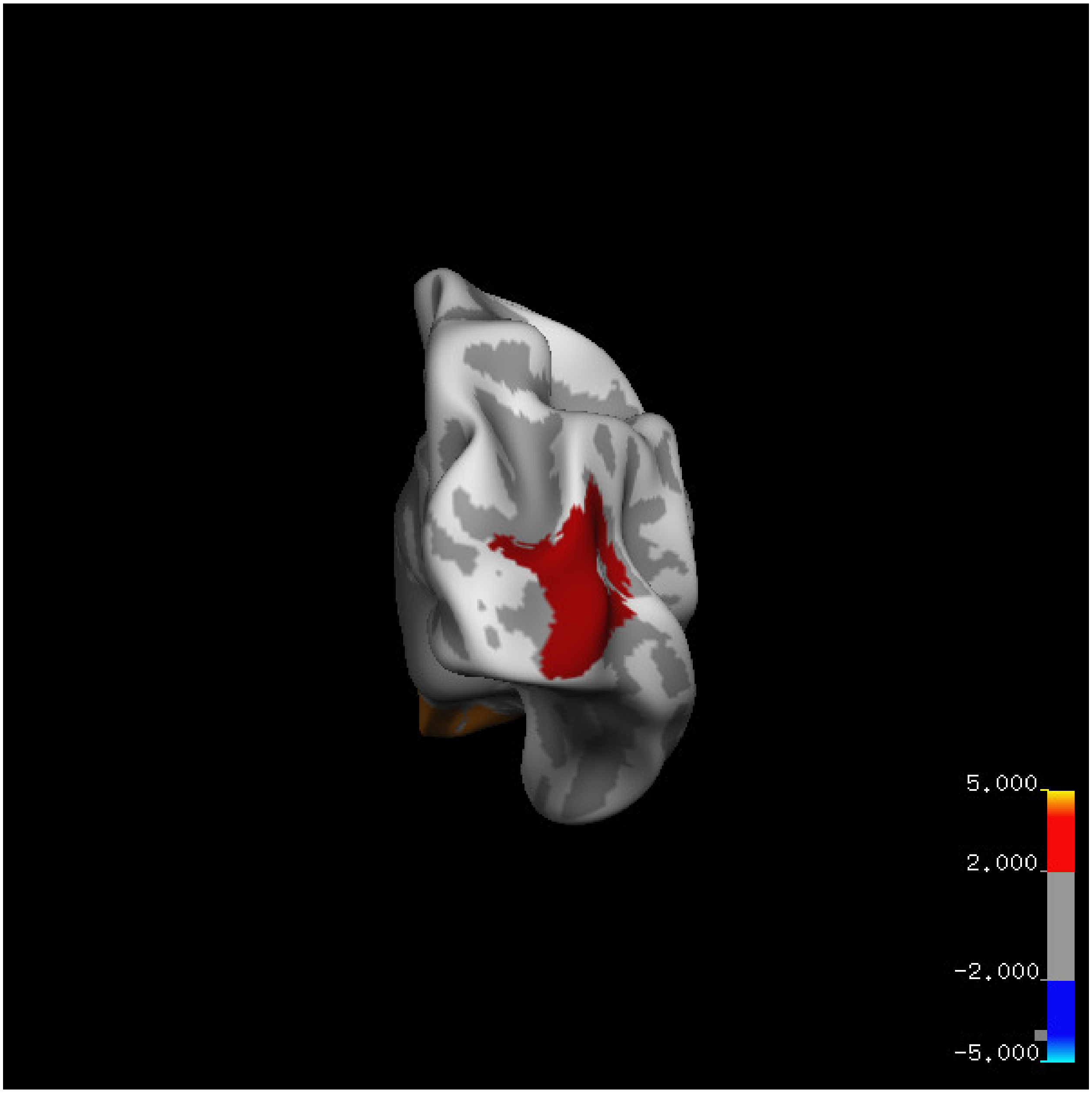
5.000
2.000
-2.000
-5.000

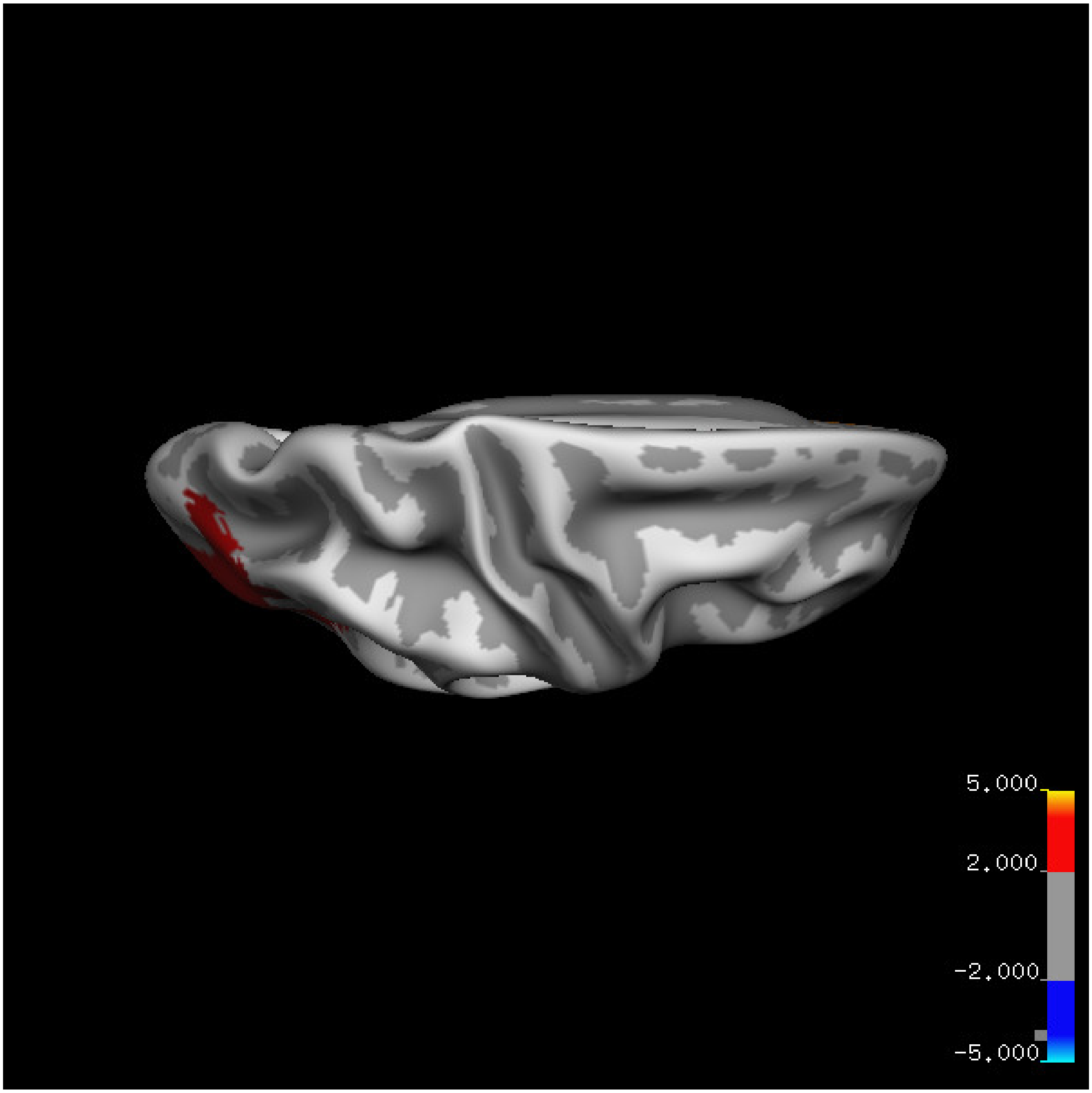
5.000
2.000
-2.000
-5.000

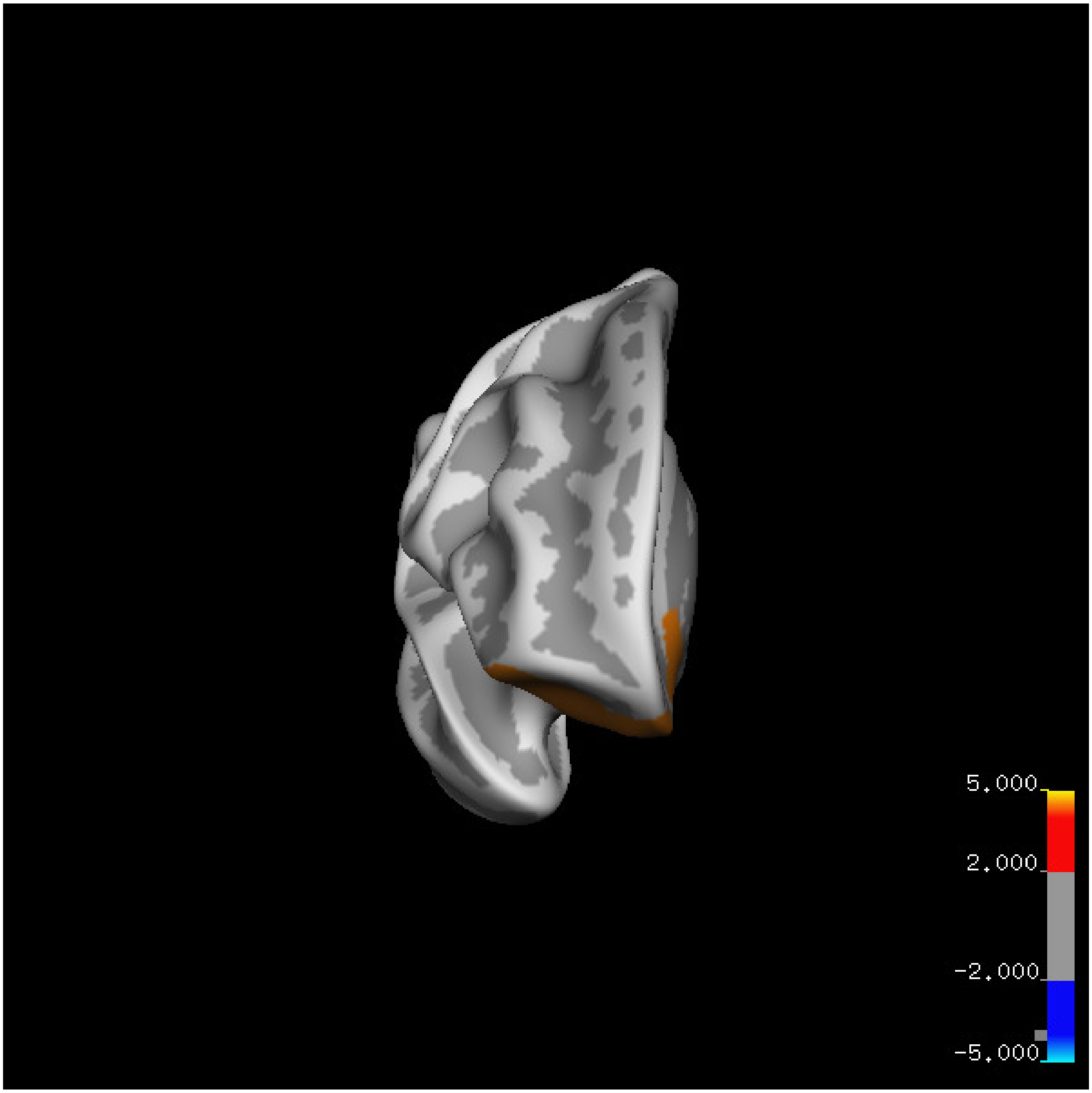
5.000
2.000
-2.000
-5.000

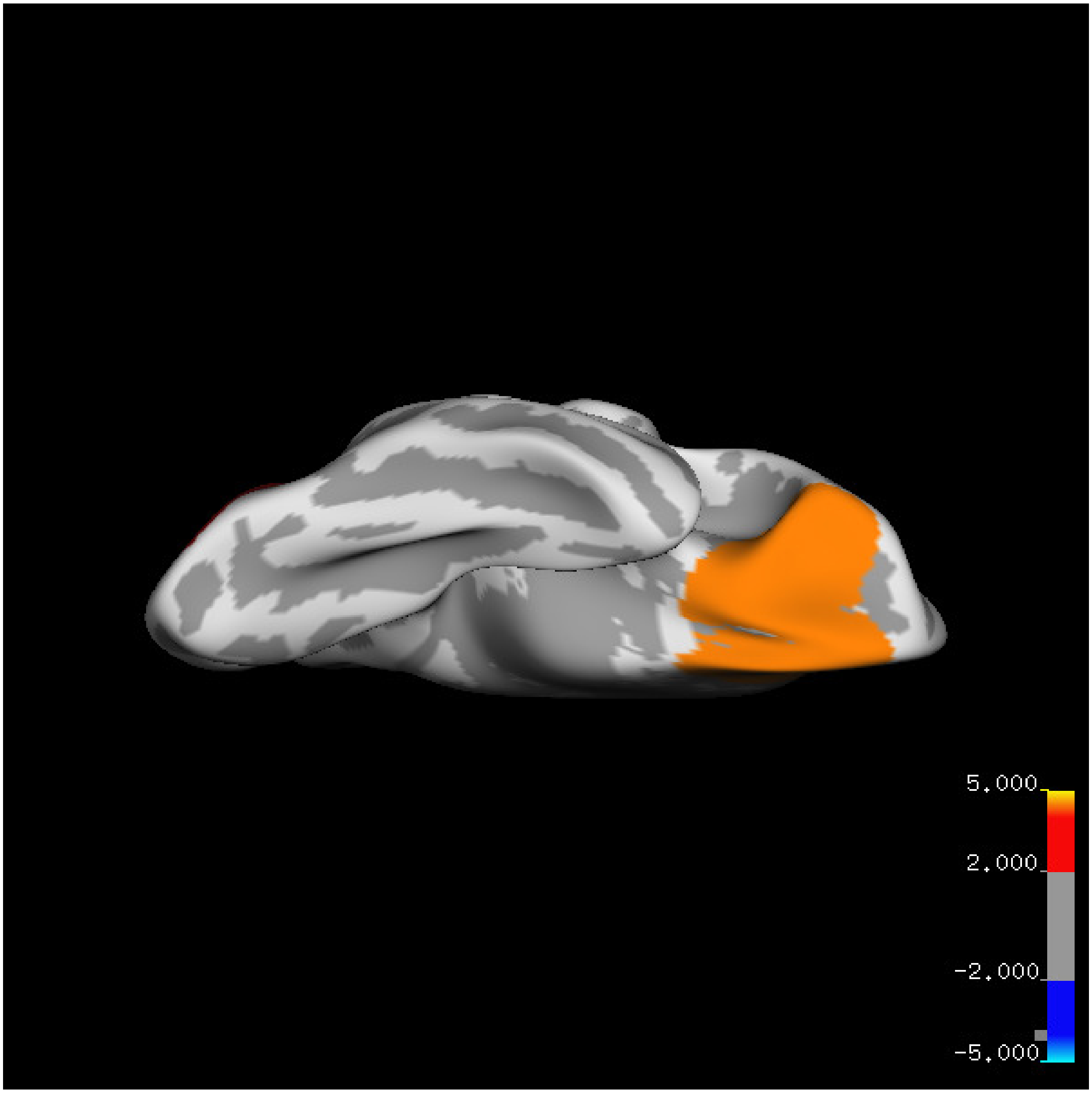
5.000
2.000
-2.000
-5.000

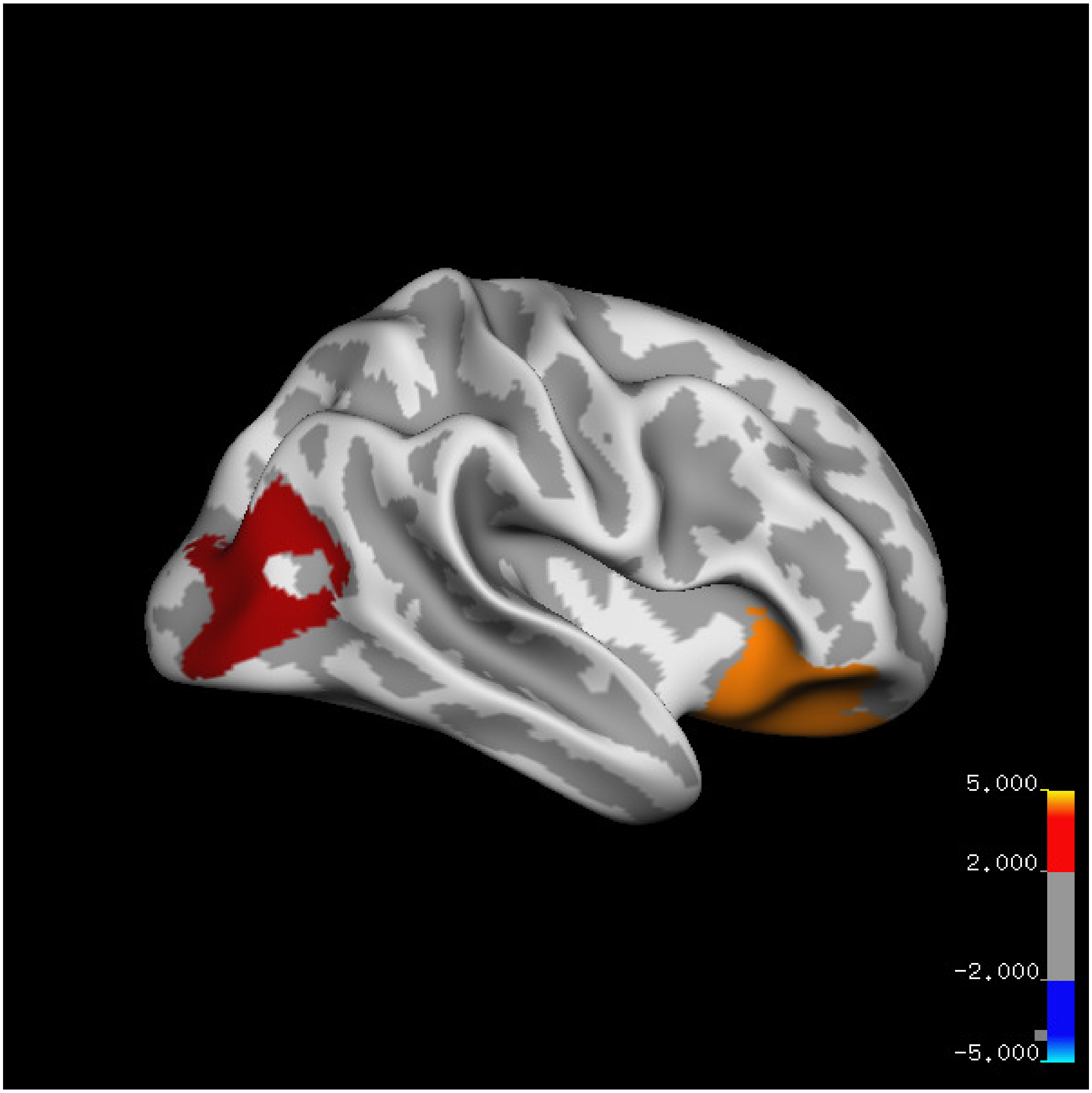
5.000
2.000
-2.000
-5.000

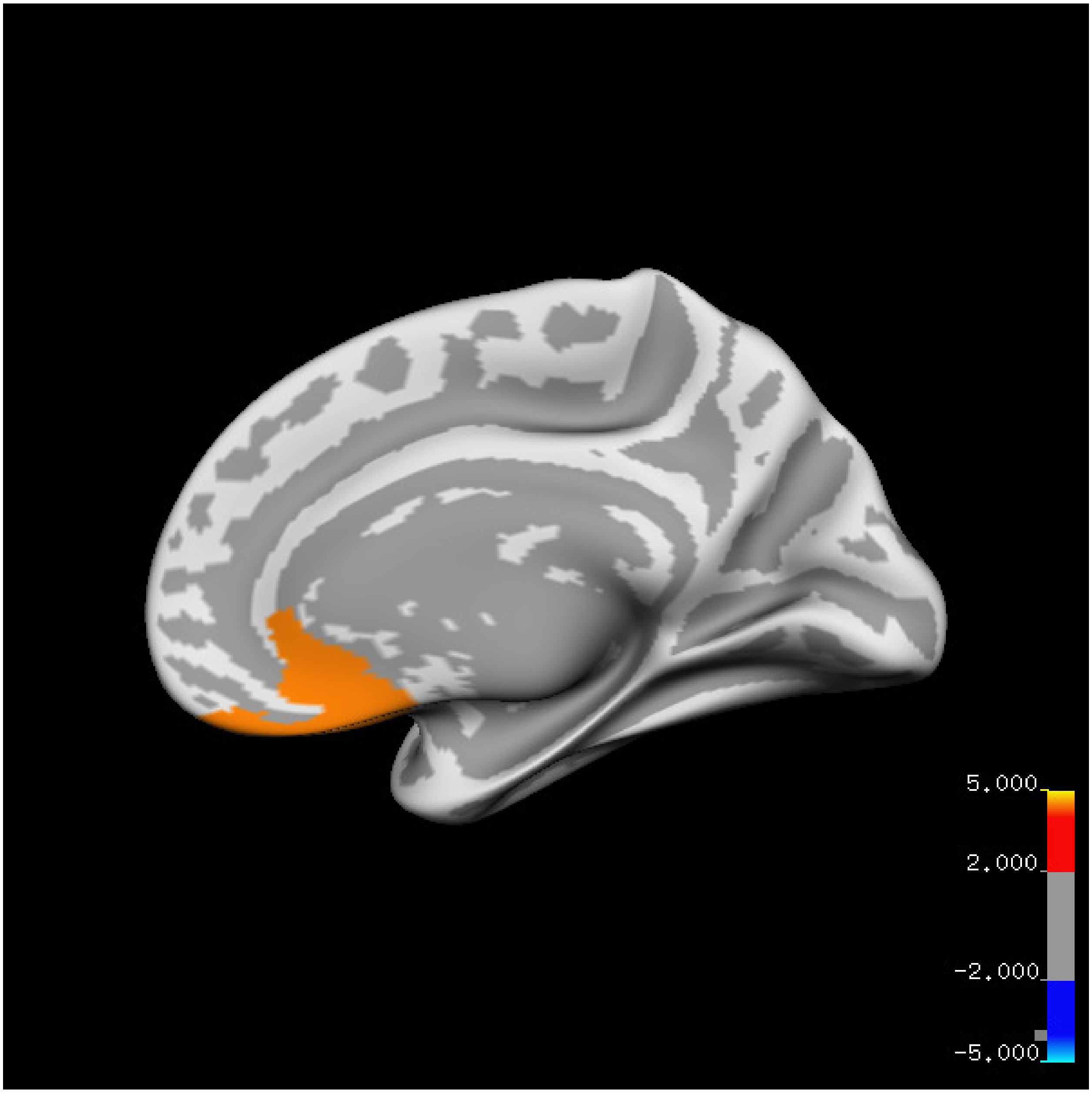
5.000
2.000
-2.000
-5.000

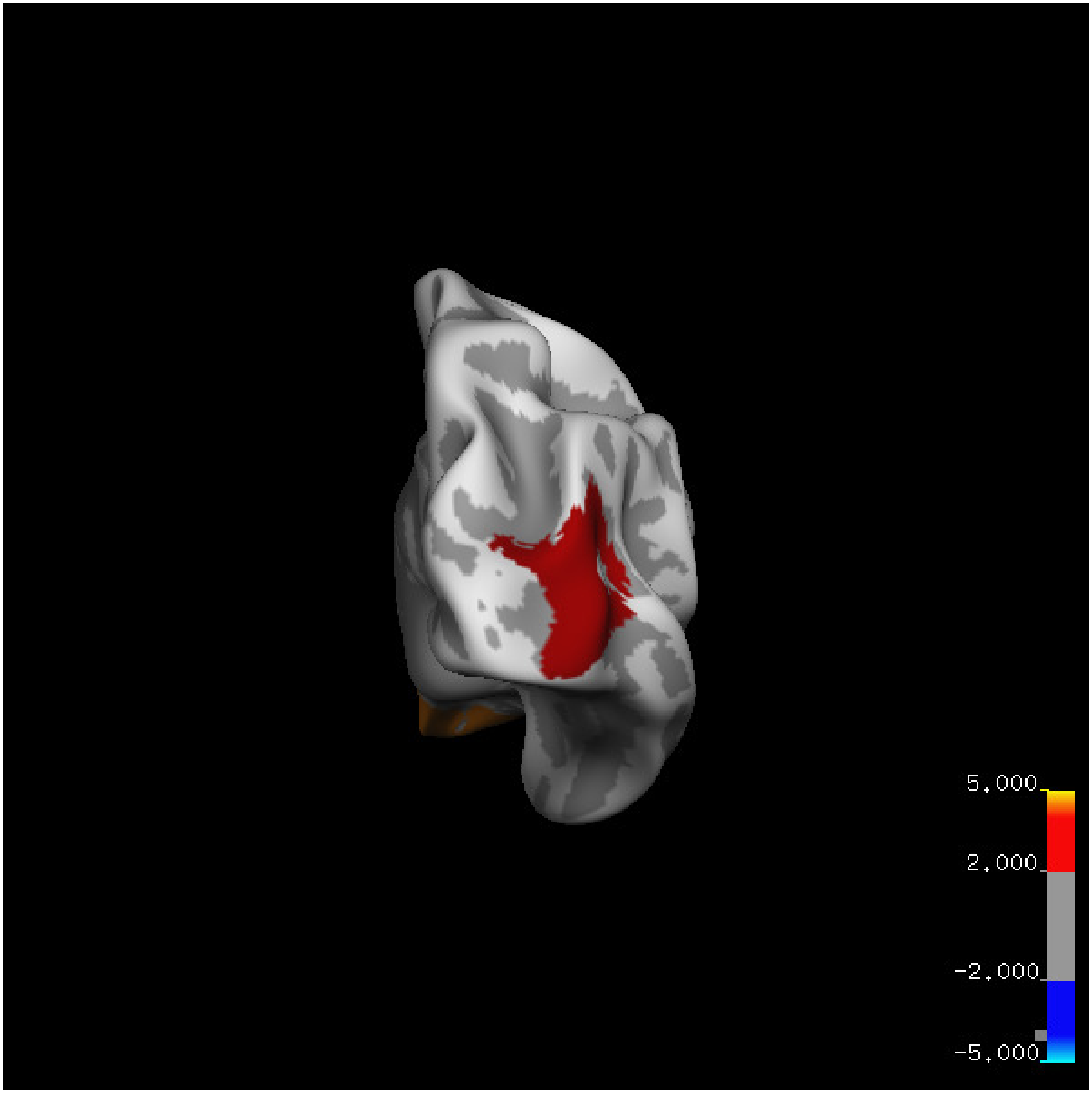
5.000
2.000
-2.000
-5.000

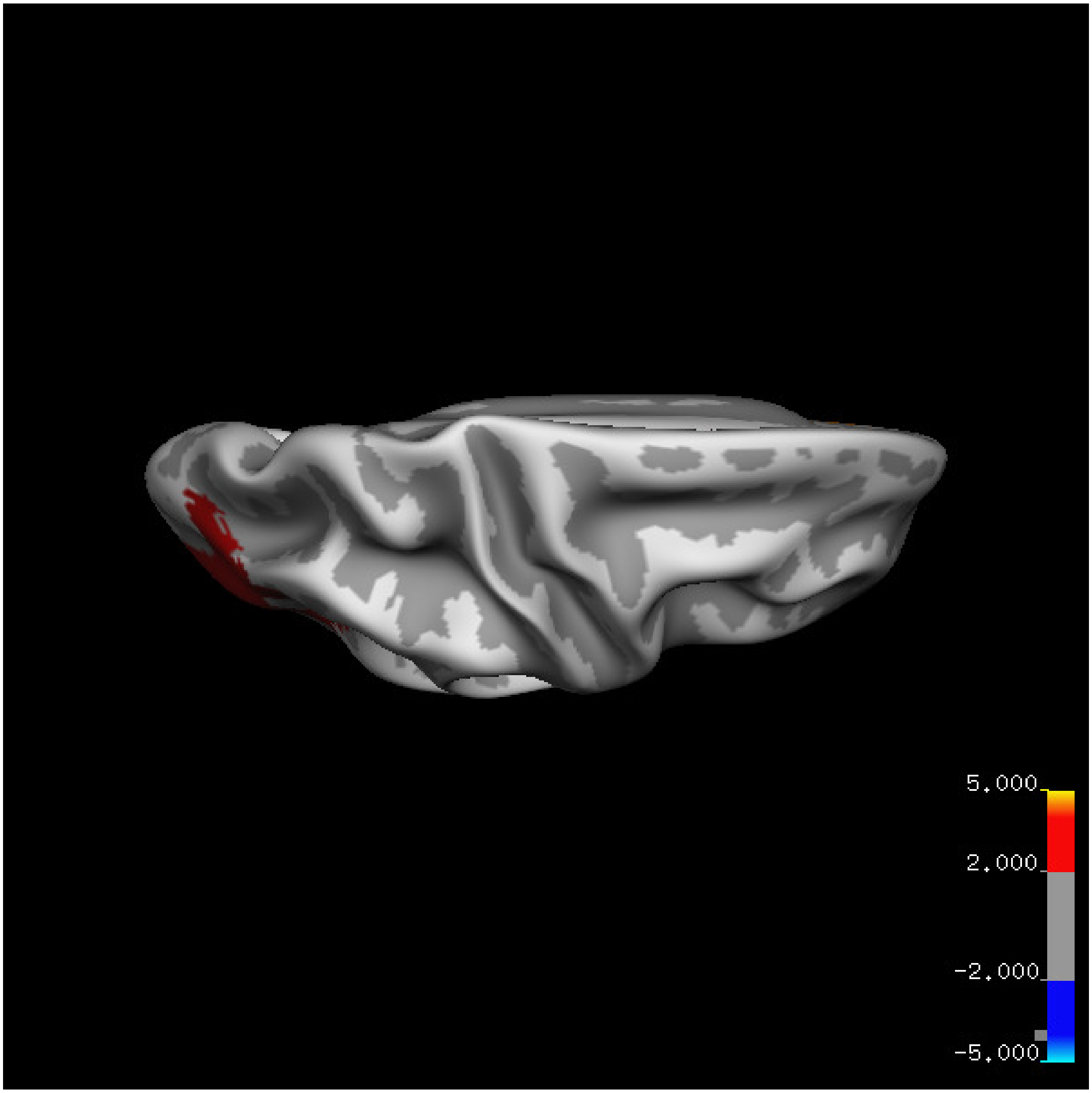


Cluster summary is shown below.

```
## # Cluster Growing Summary (mri_surfcluster)

## # $Id: mri_surfcluster.c,v 1.57.2.3 2016/11/17 18:19:42 zkaufman Exp $

## # $Id: mrisurf.c,v 1.781.2.6 2016/12/27 16:47:14 zkaufman Exp $

## # CreationTime 2021/10/04-11:24:36-GMT

## # cmdline mri_surfcluster.bin --in surface_analyses/results/cluster_comparison/fsaverage6.sm15.rh/vo

## # cwd /gpfs/projects02/p274/cluster/projects/p003-sleep_b

## # sysname  Linux
```

```
## # hostname c1-20
## # machine  x86_64
## # FixVertexAreaFlag 1
## # FixSurfClusterArea 1
## #
## # Input      surface_analyses/results/cluster_comparison/fsaverage6.sm15.rh/volume.20_60.below_mean/g
## # Frame Number      0
## # srcsubj fsaverage6
## # hemi rh
## # surface white
## # group_avg_surface_area 85131.2
## # group_avg_vtxarea_loaded 1
## # annot aparc
## # SUBJECTS_DIR /cluster/projects/p274/tools/mri/freesurfer/current/subjects
## # SearchSpace_mm2 79329.9
## # SearchSpace_vtx 37471
## # Bonferroni 0
## # Minimum Threshold 1.3
## # Maximum Threshold infinity
## # Threshold Sign    abs
## # AdjustThreshWhenOneTail 1
## # CW PValue Threshold: 0.01
## # Area Threshold    0 mm^2
## # CSD thresh  1.300000
## # CSD nreps    10000
## # CSD simtype  null-z
## # CSD contrast NA
```

```
## # CSD confint  90.000000

## # Overall max 5.26514 at vertex 27177

## # Overall min -1.89591 at vertex 4523

## # NClusters          2

## # FixMNI = 0

## #

## # ClusterNo  Max   VtxMax   Size(mm^2)  MNIX   MNIY   MNIZ    CWP    CWPLow    CWPHi   NVtxs    Wght

##    1        5.265   27177   3724.51     34.9   25.5  -10.9  0.00010  0.00000  0.00020   1733     4065

##    2        2.709   35941   2378.35     46.5  -78.6    1.4  0.00690  0.00590  0.00800    899     1599
```

**Age above 60**

**Sleep above mean** Cluster corrected significance plots are shown below.

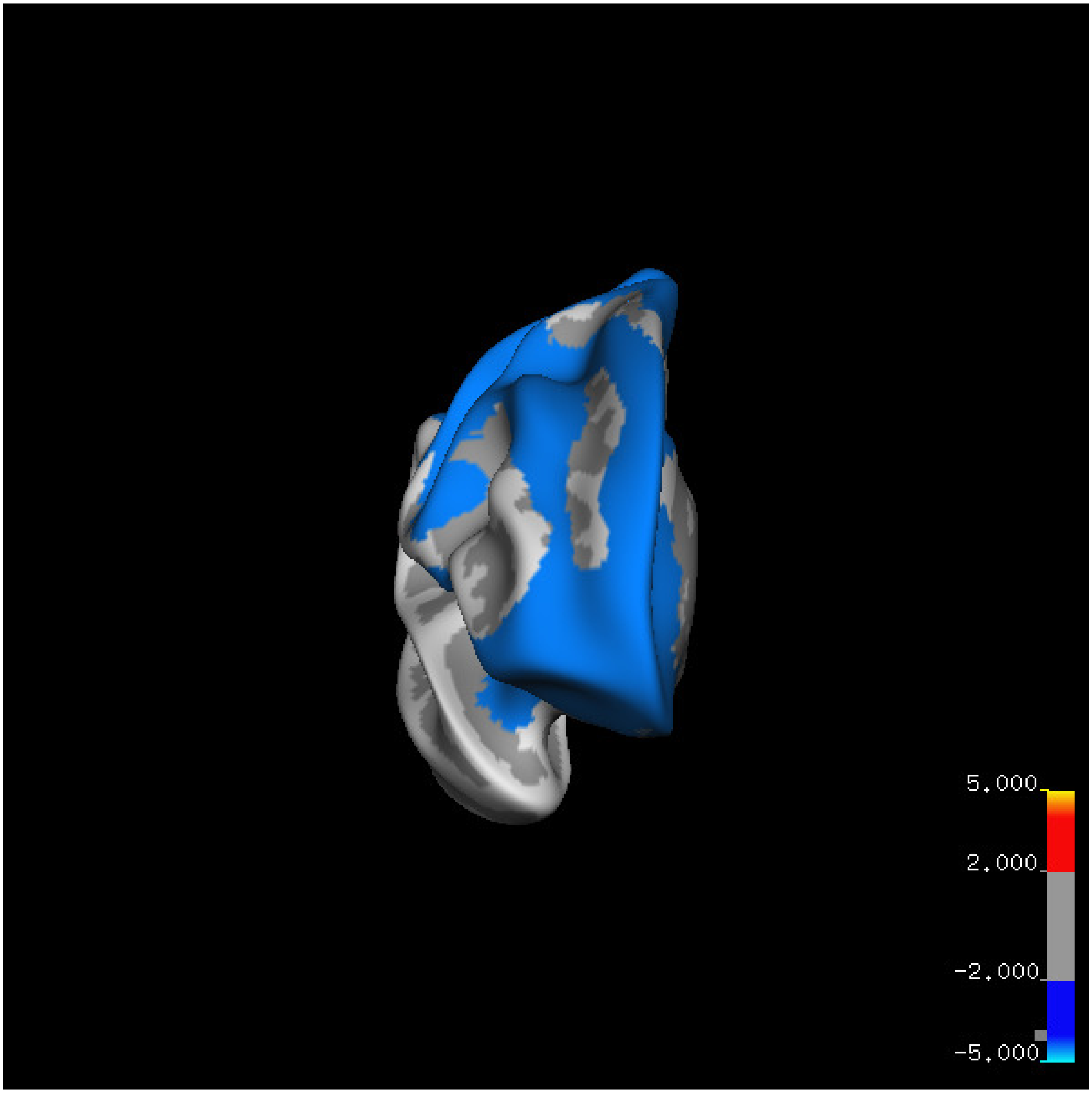
5.000
2.000
-2.000
-5.000

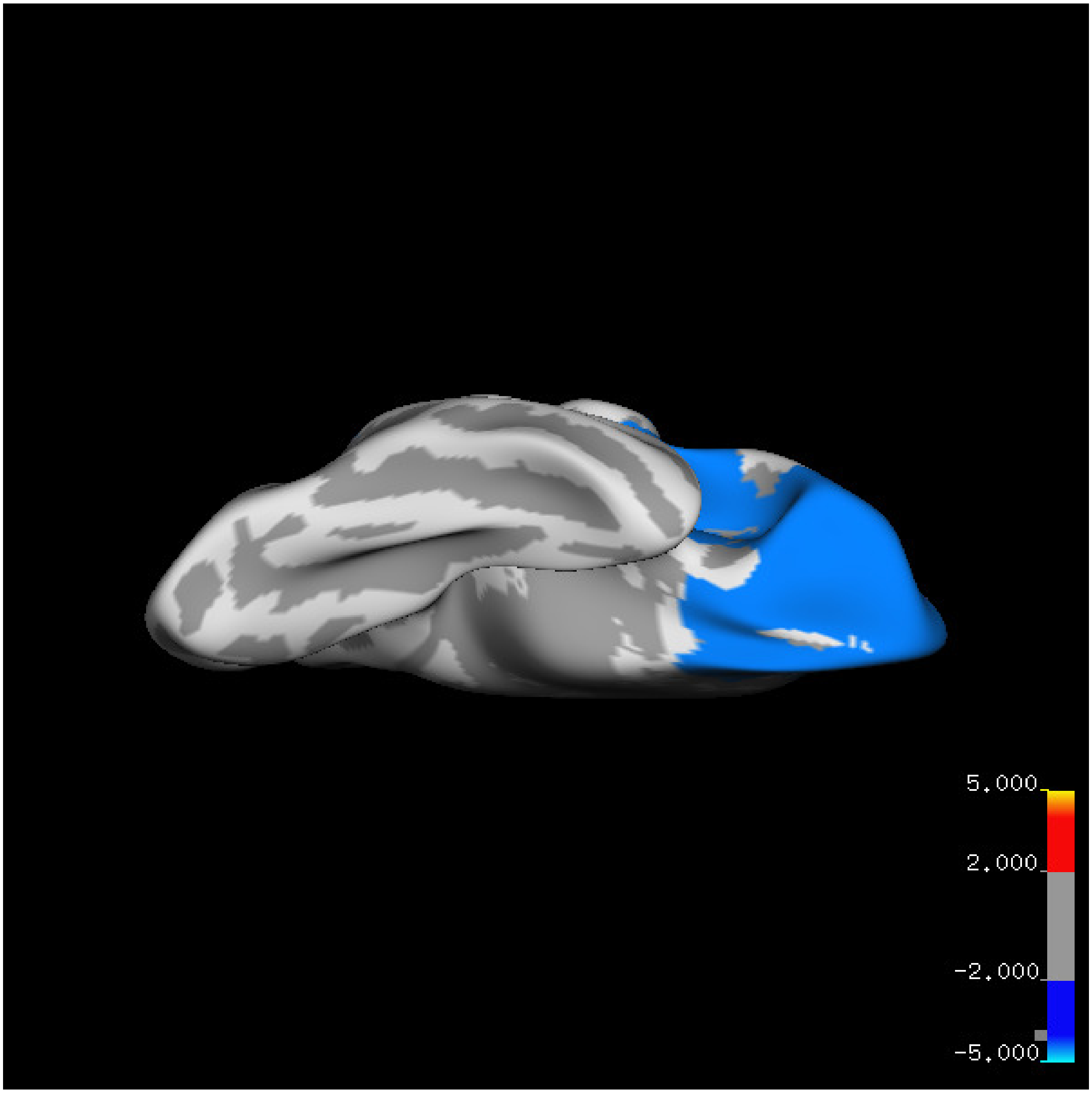
5.000
2.000
-2.000
-5.000

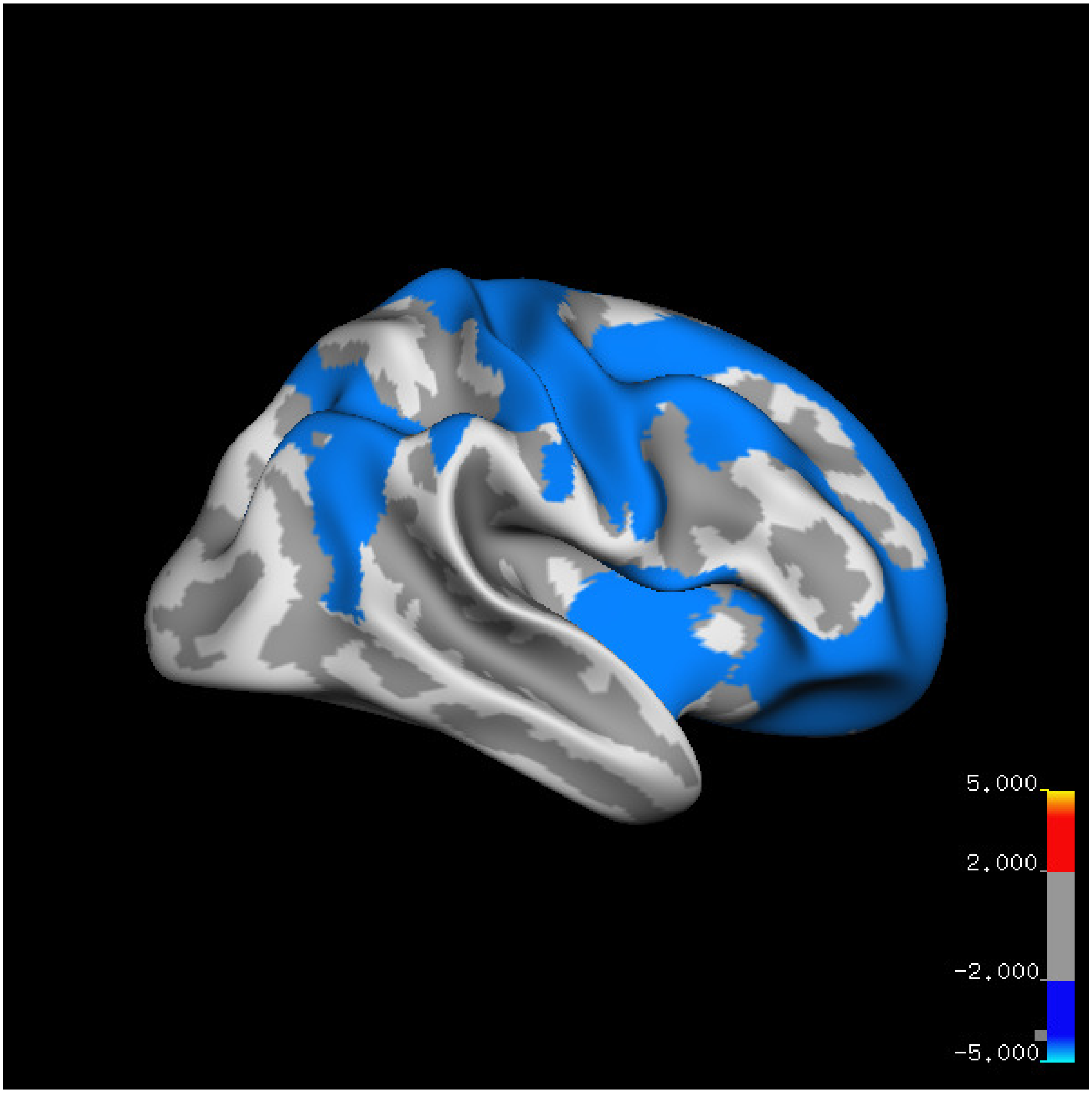

5.000
2.000
-2.000
-5.000

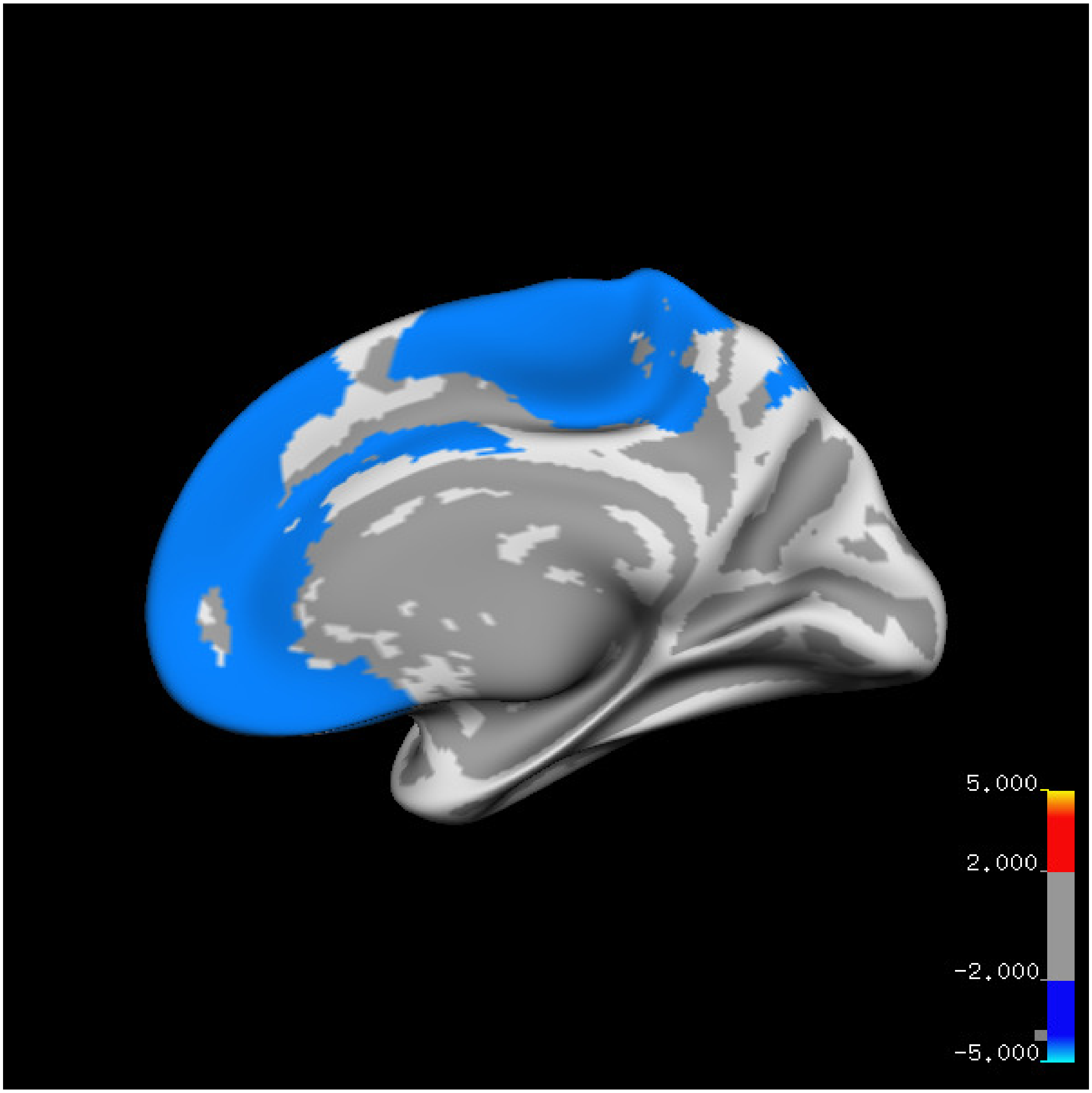
5.000
2.000
-2.000
-5.000

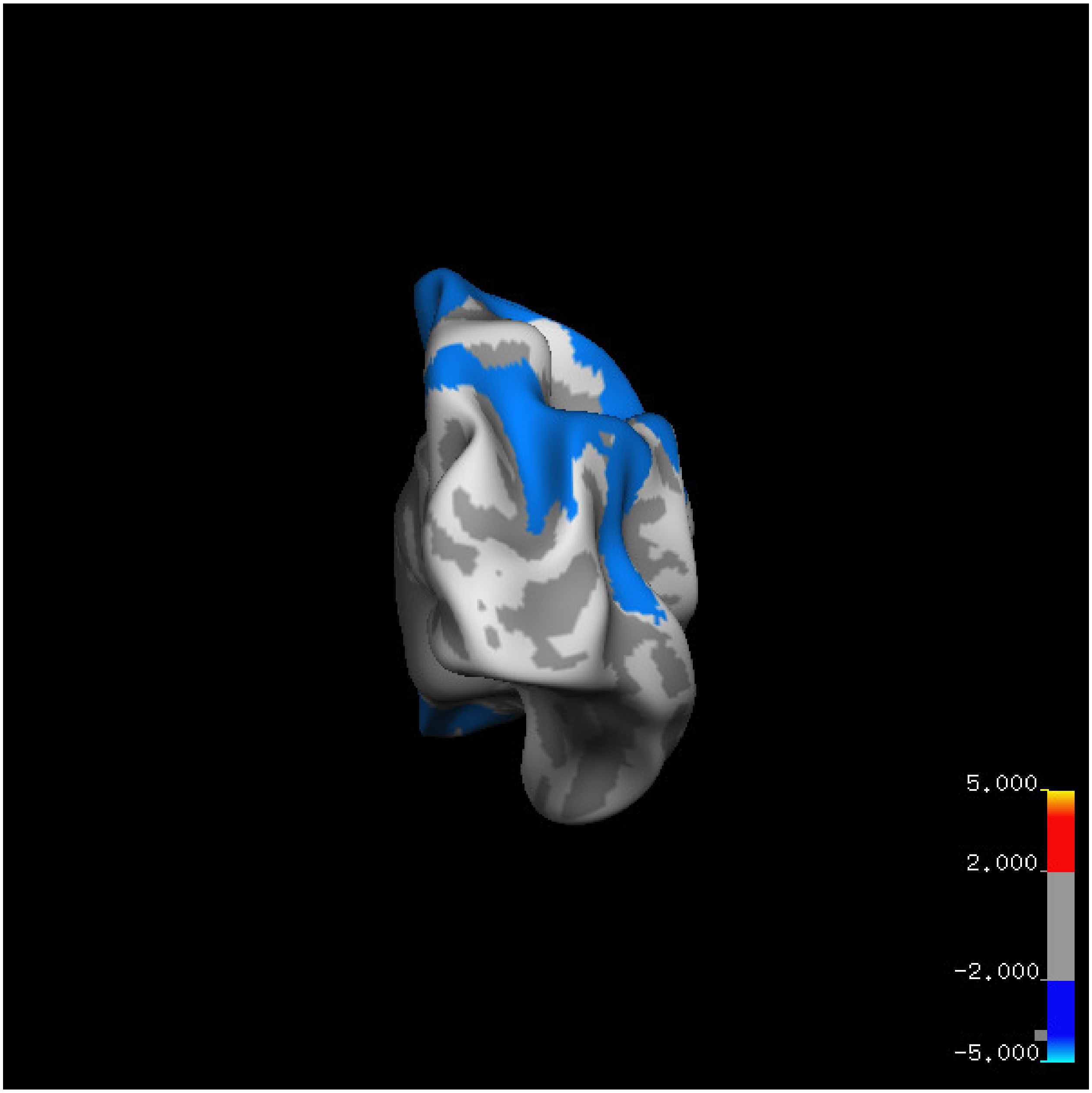
5.000
2.000
-2.000
-5.000

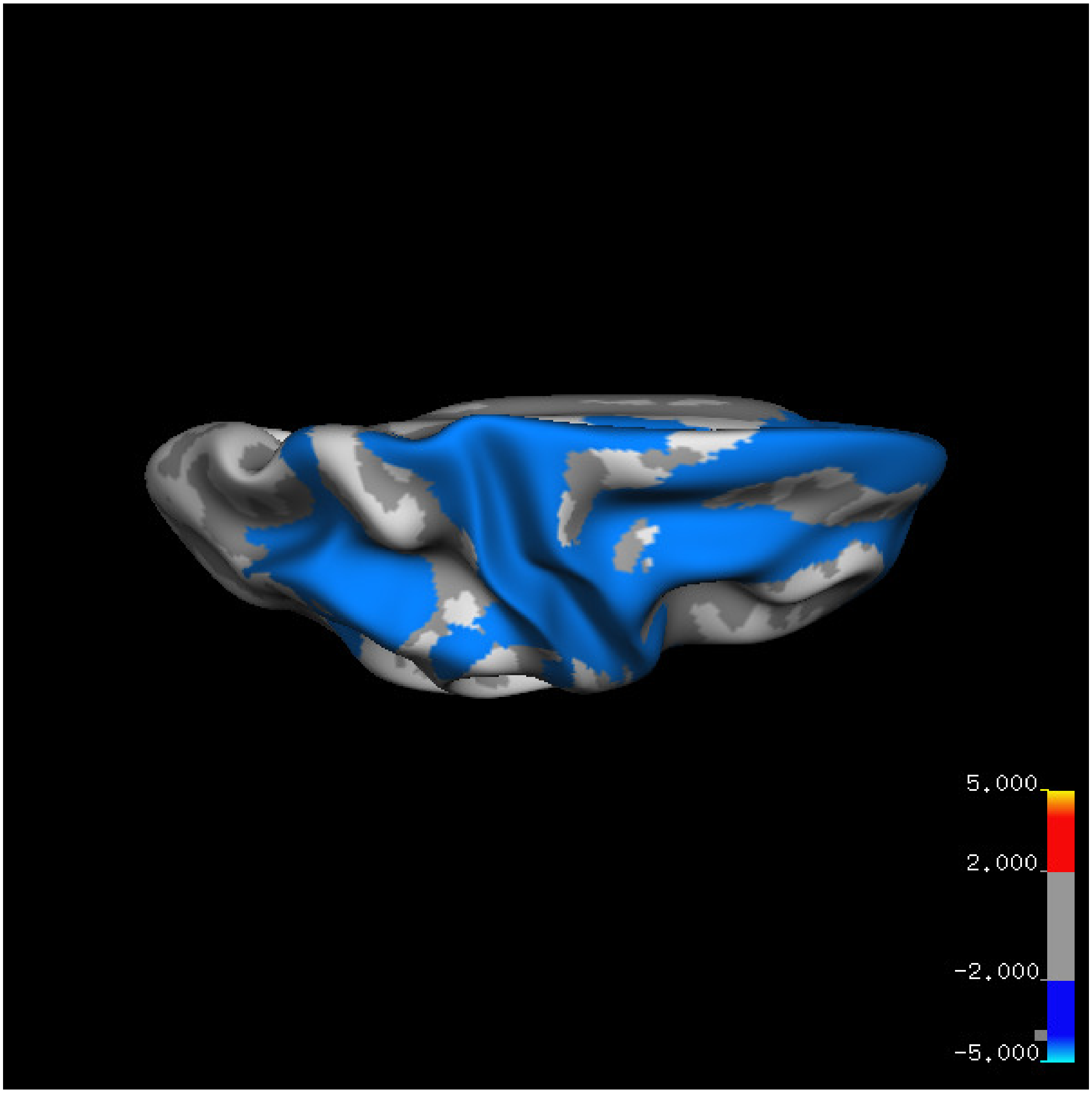
5.000
2.000
-2.000
-5.000

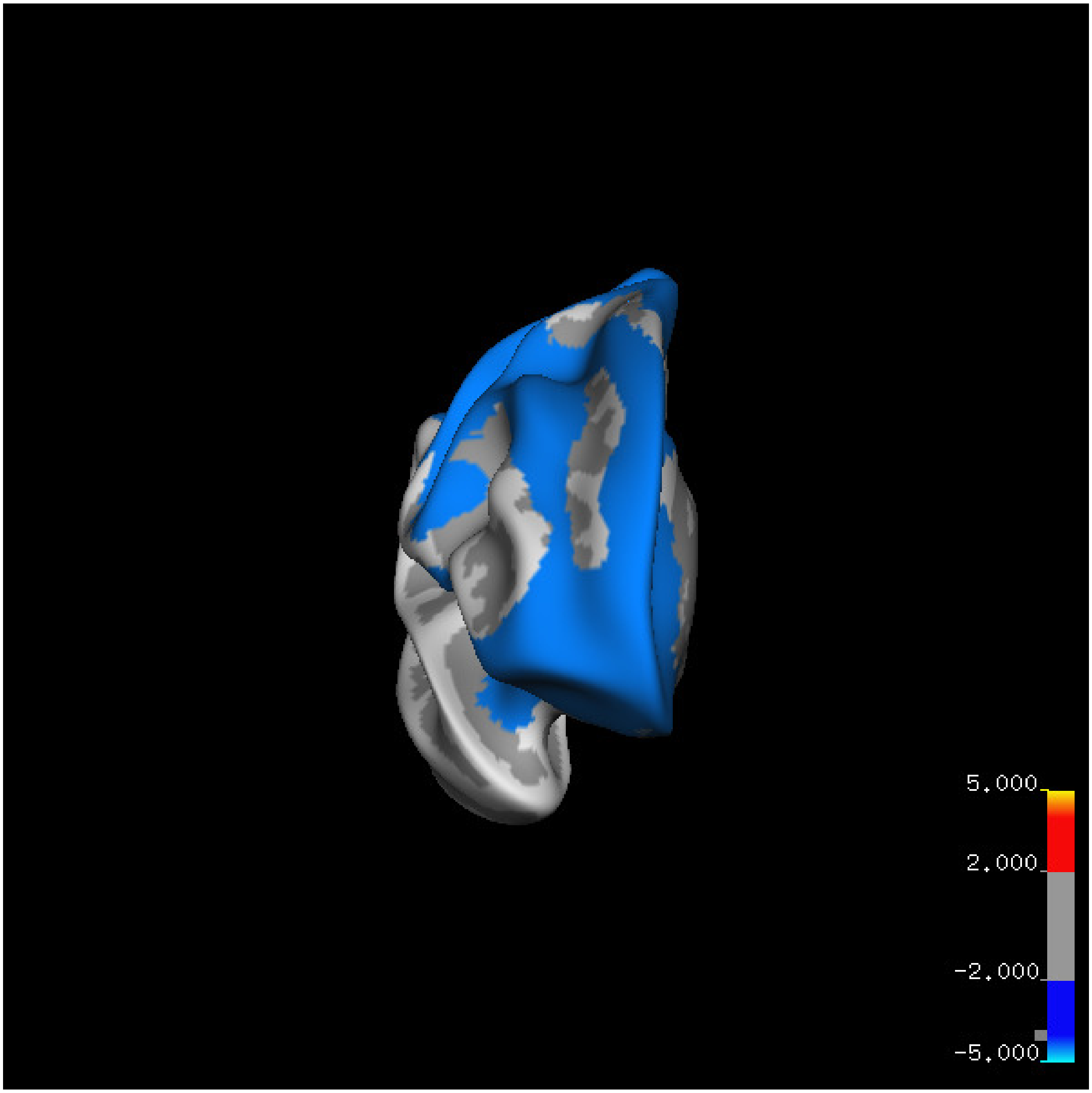
5.000
2.000
-2.000
-5.000

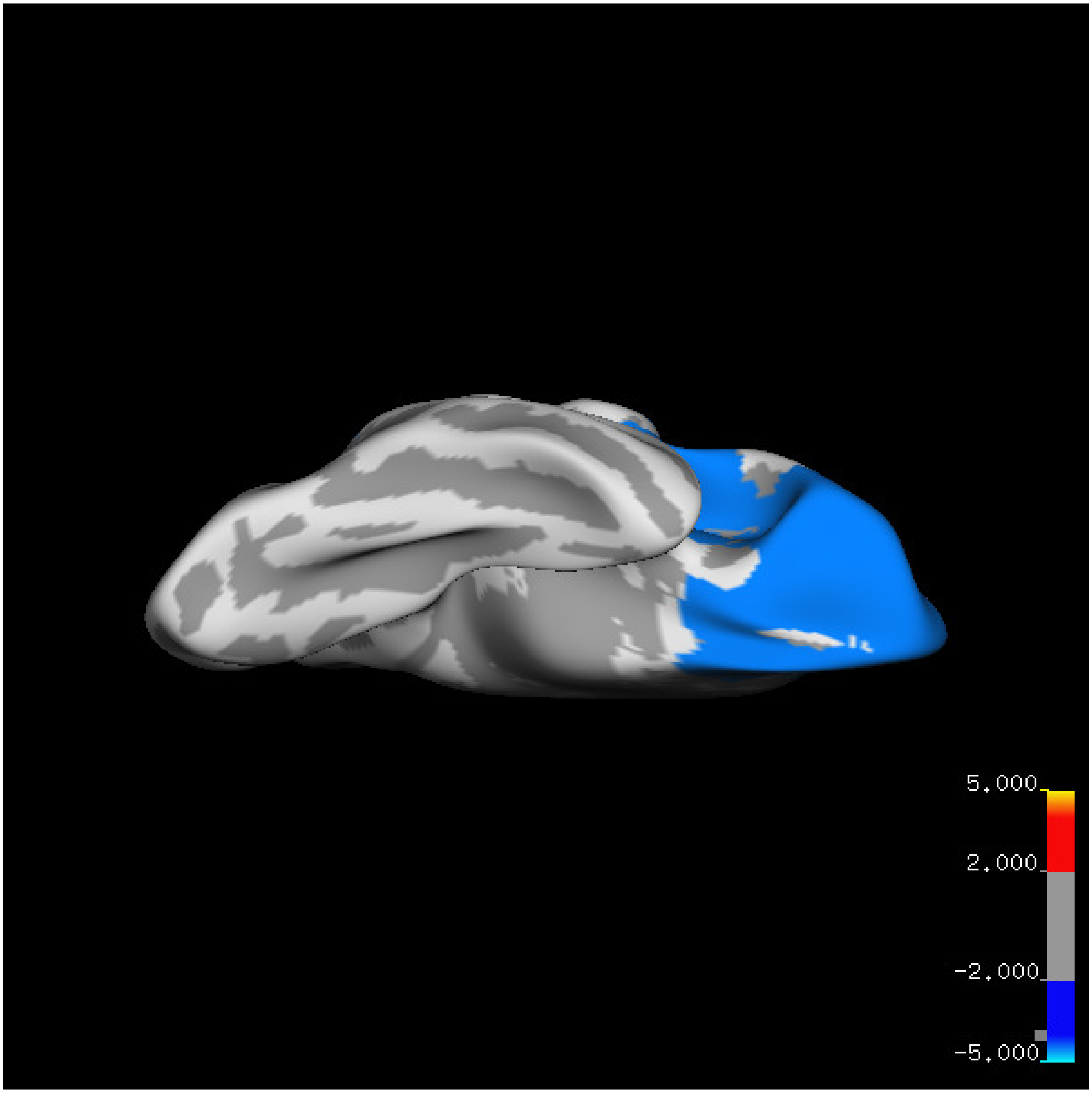

5.000
2.000
-2.000
-5.000

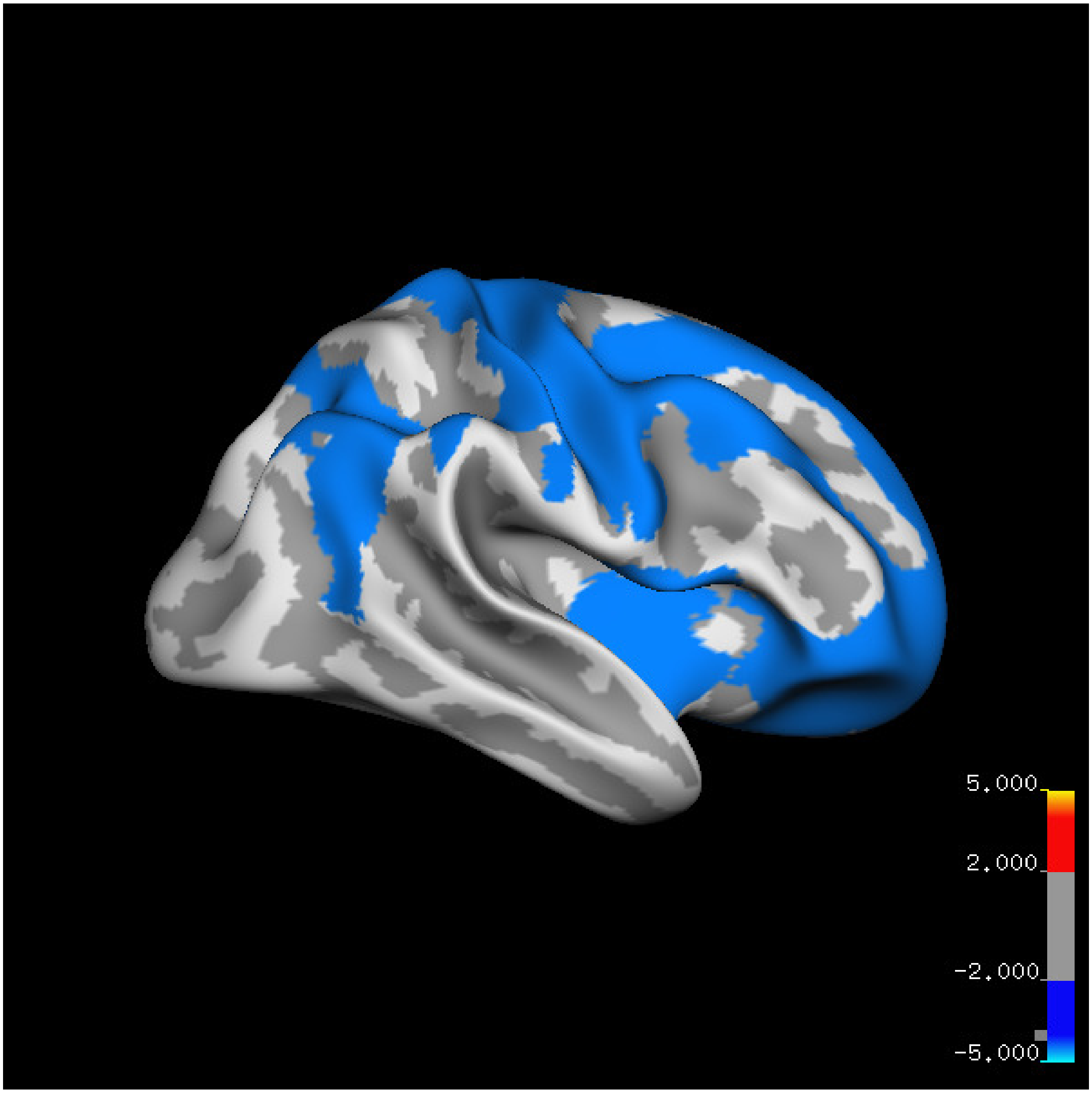
5.000
2.000
-2.000
-5.000

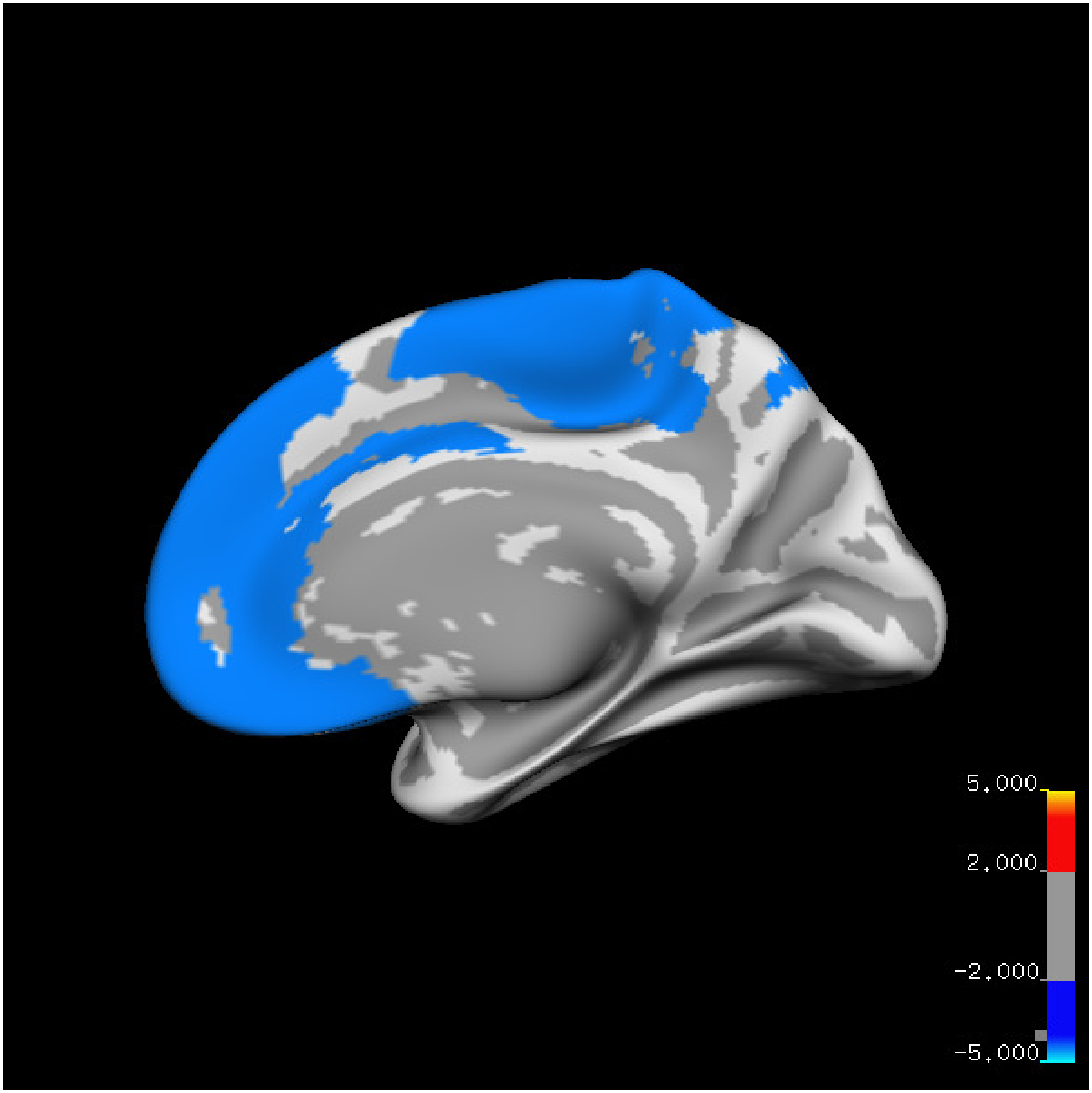
5.000
2.000
-2.000
-5.000

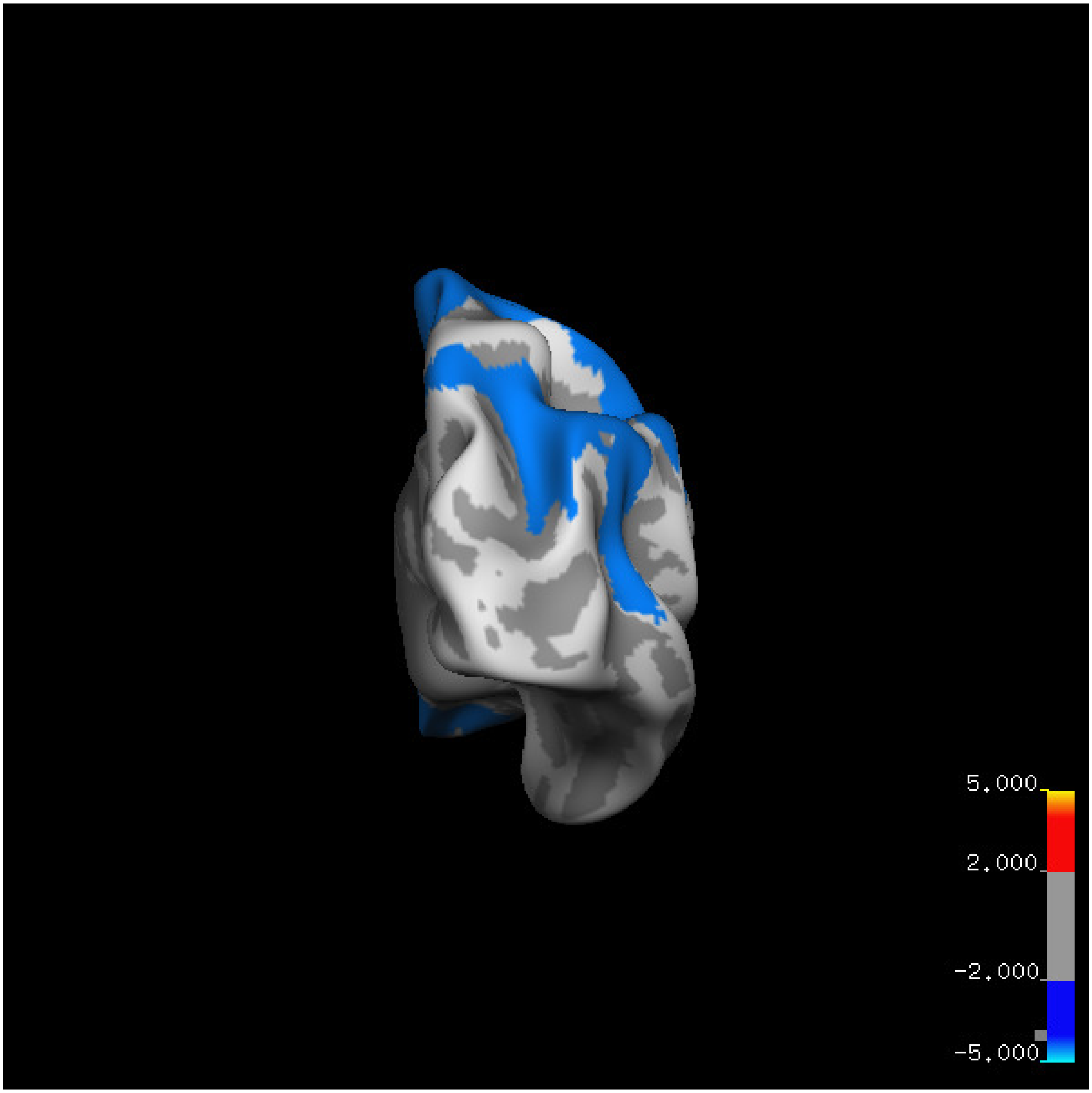
5.000
2.000
-2.000
-5.000

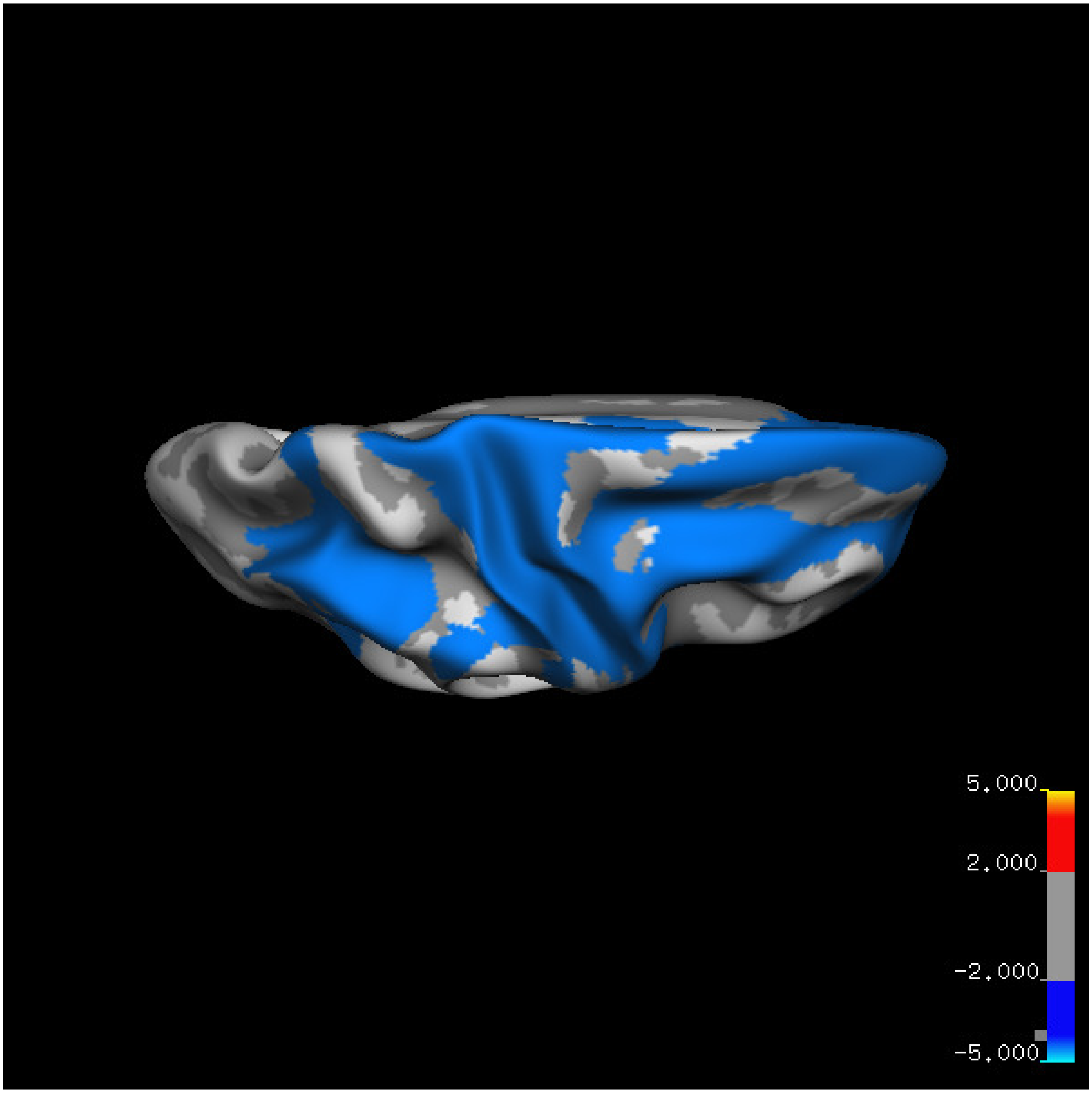


Cluster summary is shown below.

```
## # Cluster Growing Summary (mri_surfcluster)

## # $Id: mri_surfcluster.c,v 1.57.2.3 2016/11/17 18:19:42 zkaufman Exp $

## # $Id: mrisurf.c,v 1.781.2.6 2016/12/27 16:47:14 zkaufman Exp $

## # CreationTime 2021/10/04-13:08:00-GMT

## # cmdline mri_surfcluster.bin --in surface_analyses/results/cluster_comparison/fsaverage6.sm15.rh/vol

## # cwd /gpfs/projects02/p274/cluster/projects/p003-sleep_b

## # sysname  Linux
```

```
## # hostname c1-20
## # machine  x86_64
## # FixVertexAreaFlag 1
## # FixSurfClusterArea 1
## #
## # Input      surface_analyses/results/cluster_comparison/fsaverage6.sm15.rh/volume.above60.above_mear
## # Frame Number      0
## # srcsubj fsaverage6
## # hemi rh
## # surface white
## # group_avg_surface_area 85131.2
## # group_avg_vtxarea_loaded 1
## # annot aparc
## # SUBJECTS_DIR /cluster/projects/p274/tools/mri/freesurfer/current/subjects
## # SearchSpace_mm2 79329.9
## # SearchSpace_vtx 37471
## # Bonferroni 0
## # Minimum Threshold 1.3
## # Maximum Threshold infinity
## # Threshold Sign    abs
## # AdjustThreshWhenOneTail 1
## # CW PValue Threshold: 0.01
## # Area Threshold    0 mm^2
## # CSD thresh  1.300000
## # CSD nreps    10000
## # CSD simtype  null-z
## # CSD contrast NA
```

```
## # CSD confint  90.000000

## # Overall max 1.00354 at vertex 24055

## # Overall min -9.76718 at vertex 27171

## # NClusters          2

## # FixMNI = 0

## #

## # ClusterNo  Max   VtxMax   Size(mm^2)  MNIX   MNIY   MNIZ    CWP    CWPLow    CWPHi   NVtxs    WghtV

##    1       -9.767   27171  26058.29     42.5   26.2  -14.0  0.00010  0.00000  0.00020  12870   -34056

##    2       -5.321    2710   4401.72     52.4  -53.3   37.5  0.00010  0.00000  0.00020   2258    -4901
```

**Sleep below mean** Cluster corrected significance plots are shown below.

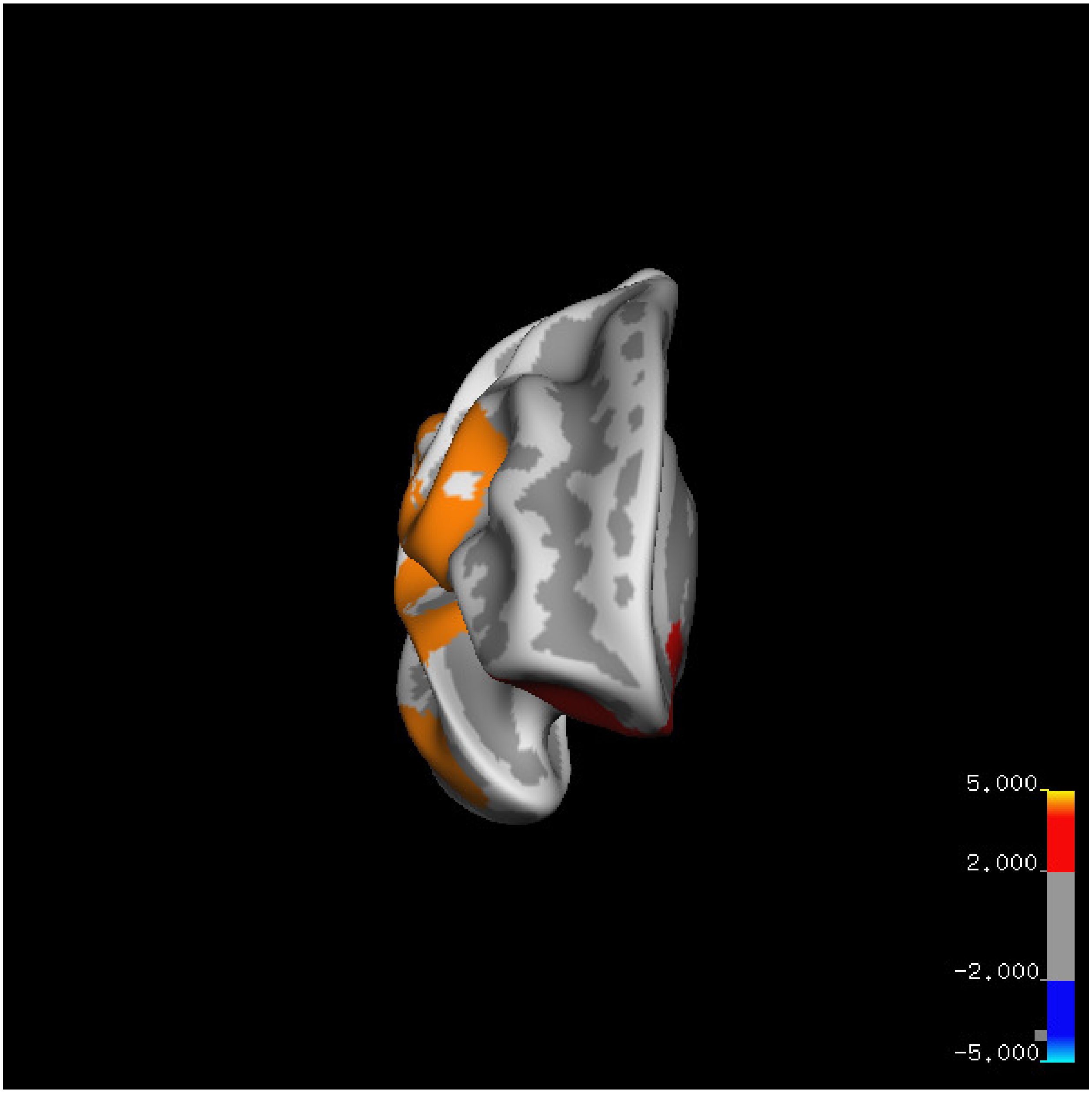
5.000
2.000
-2.000
-5.000

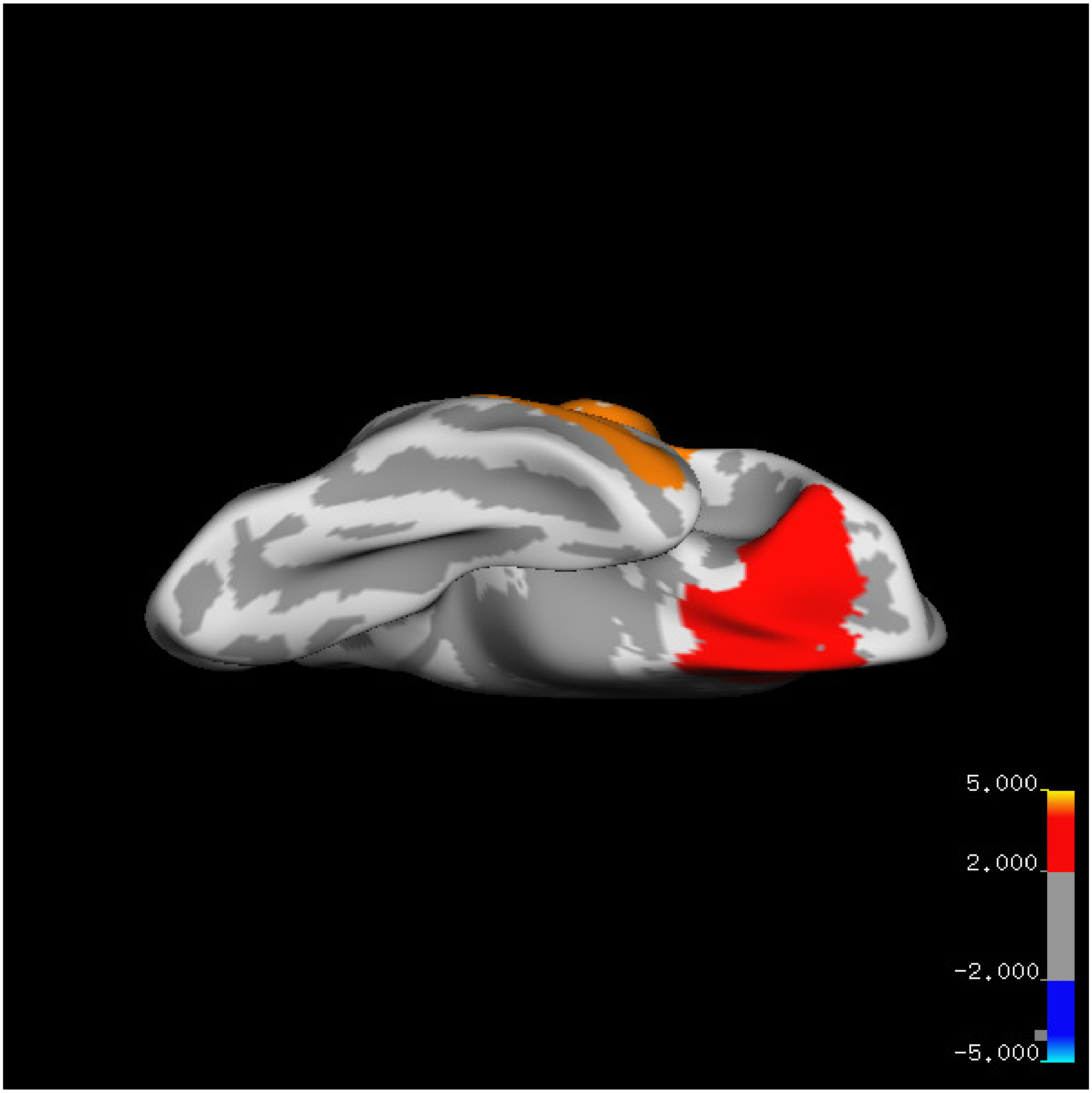
5.000
2.000
-2.000
-5.000

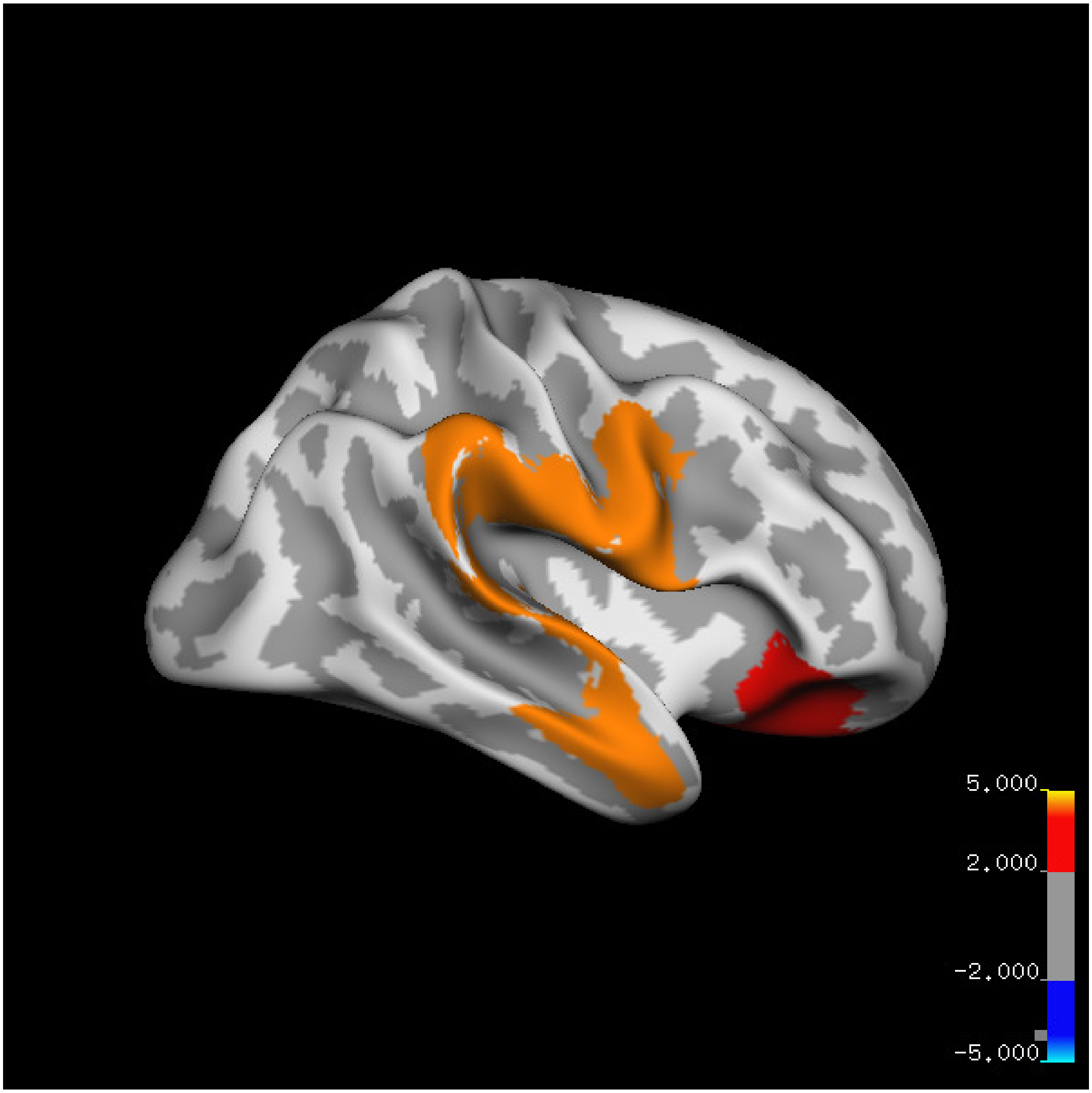

5.000
2.000
-2.000
-5.000

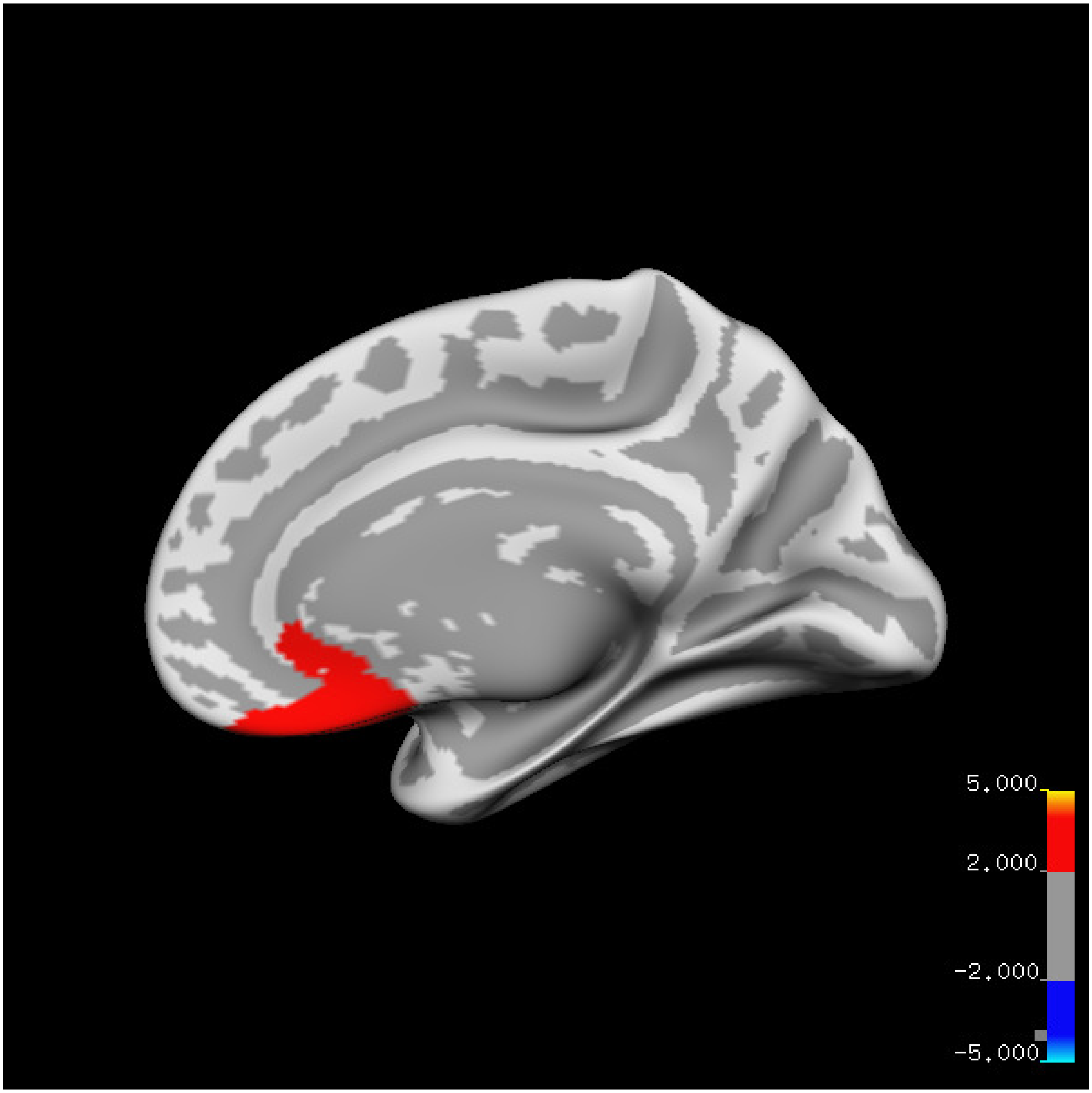
5.000
2.000
-2.000
-5.000

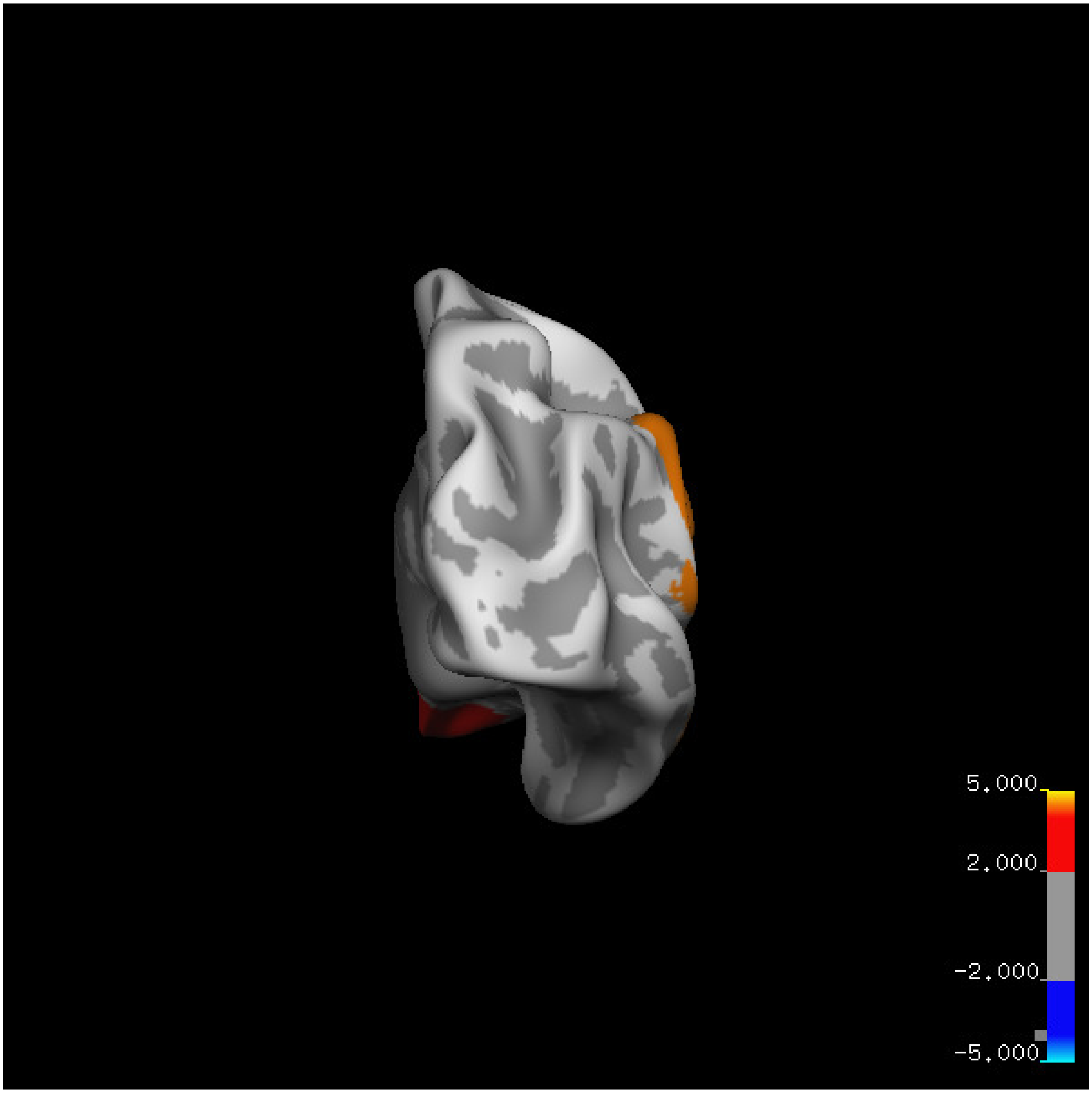
5.000
2.000
-2.000
-5.000

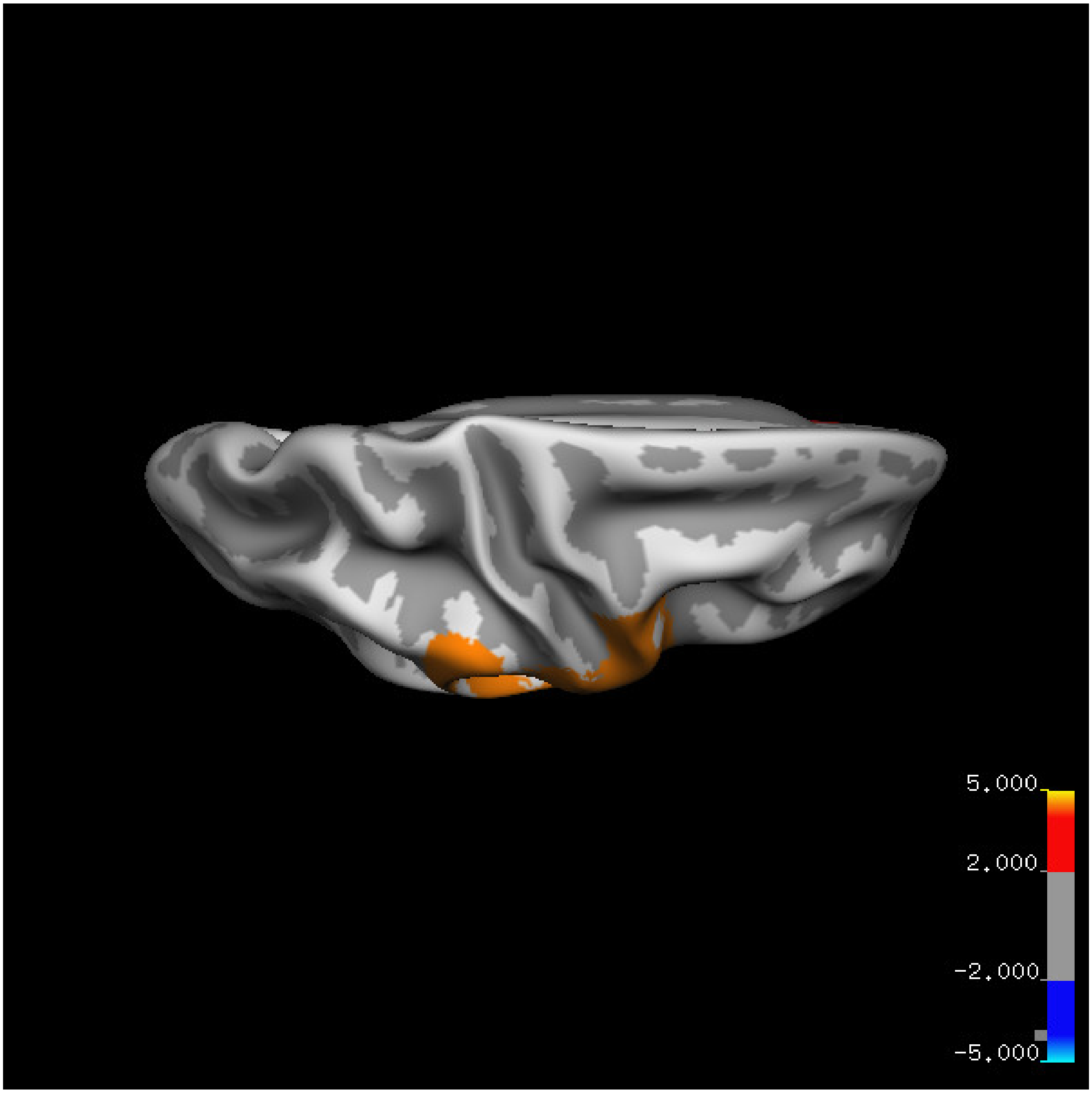

5.000
2.000
-2.000
-5.000

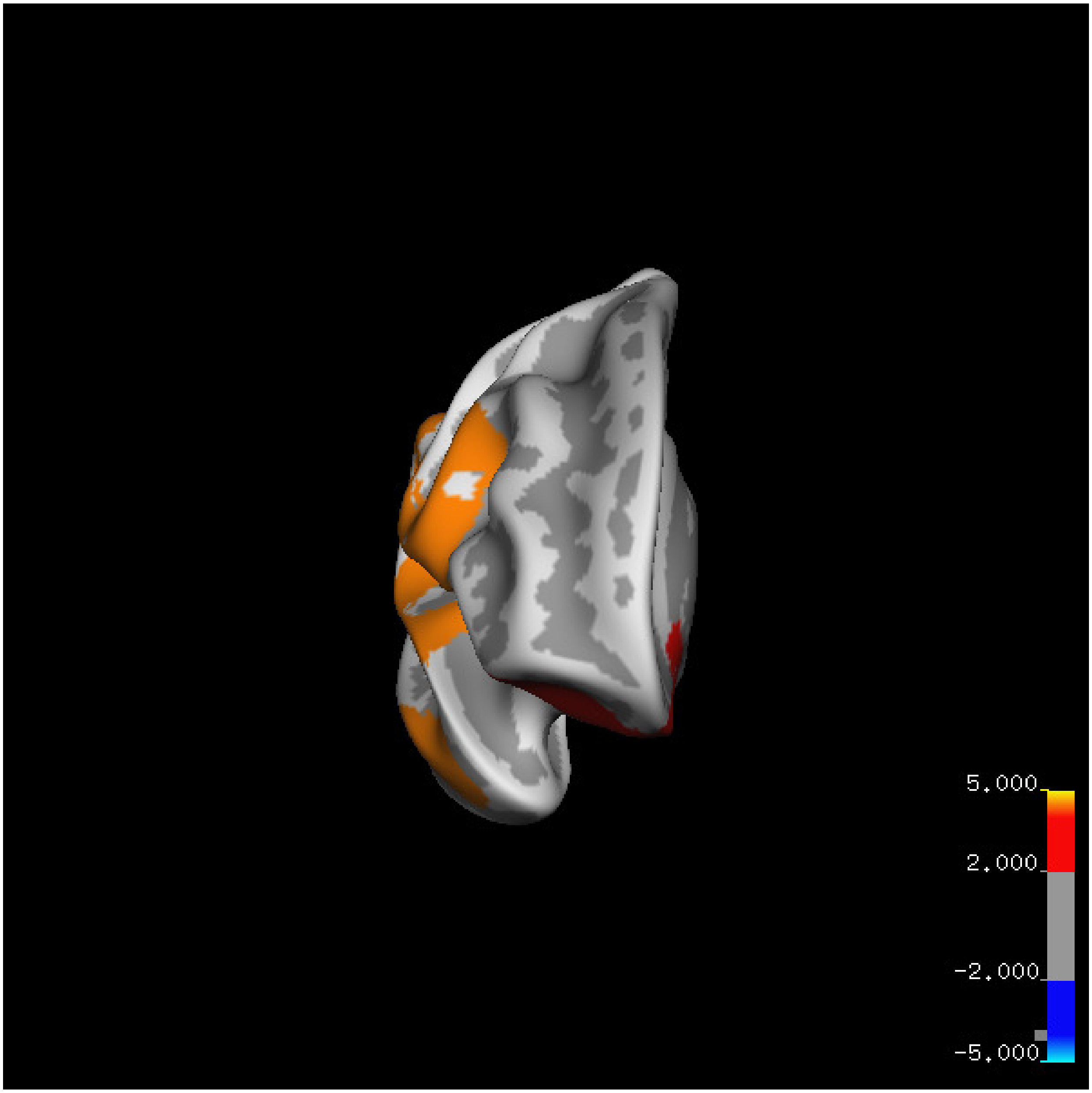
5.000
2.000
-2.000
-5.000

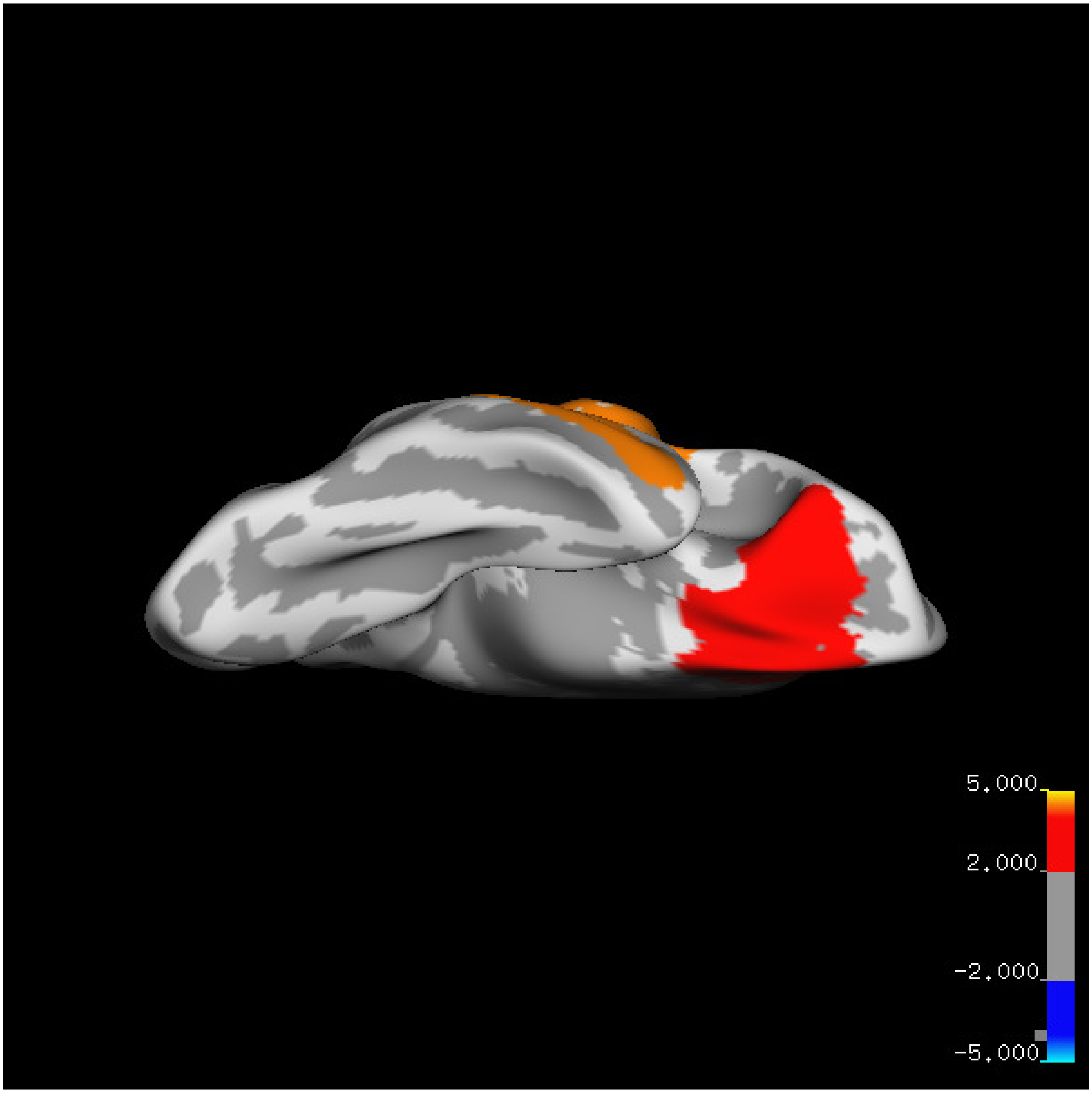
5.000
2.000
-2.000
-5.000

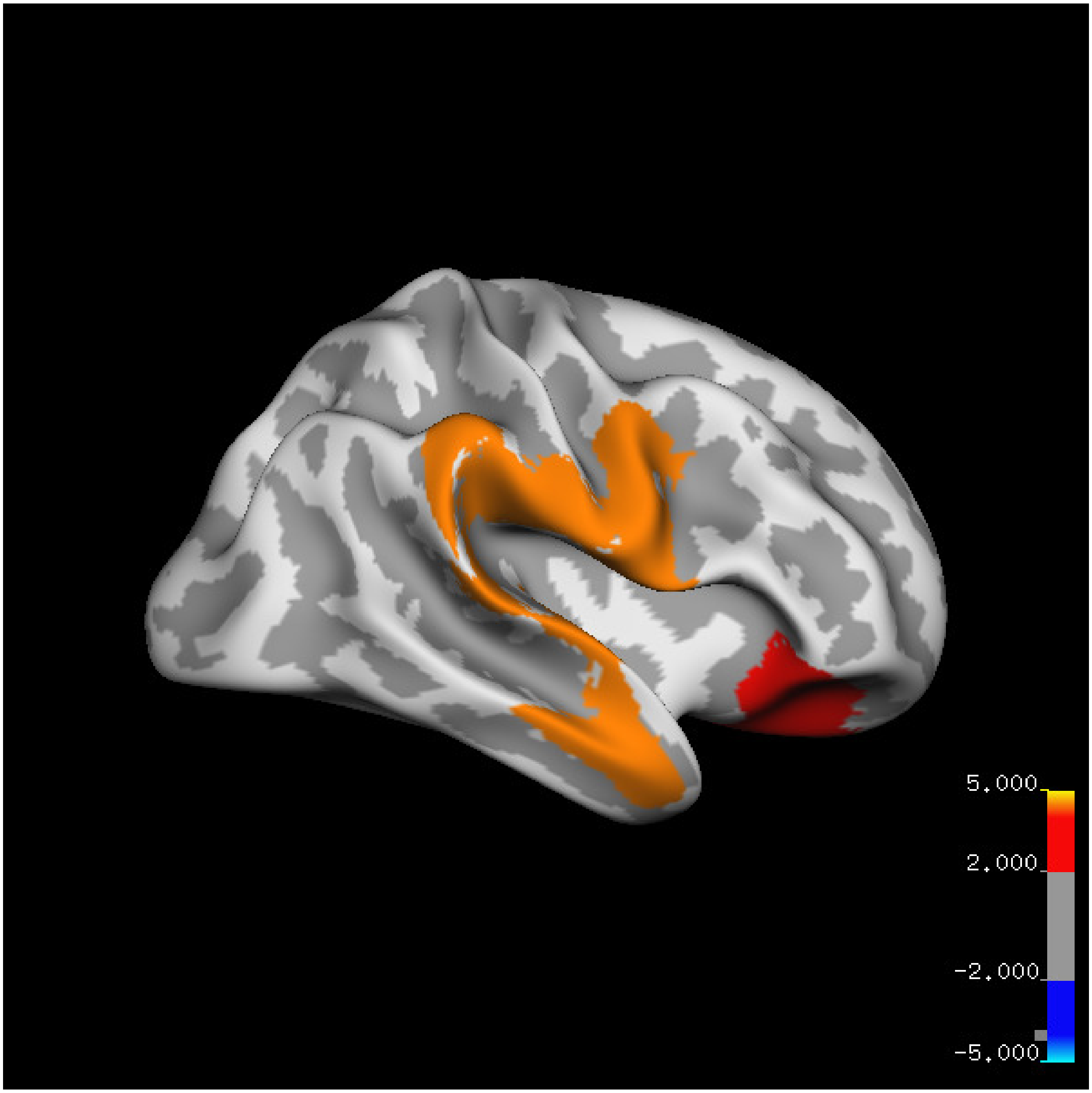

5.000
2.000
-2.000
-5.000

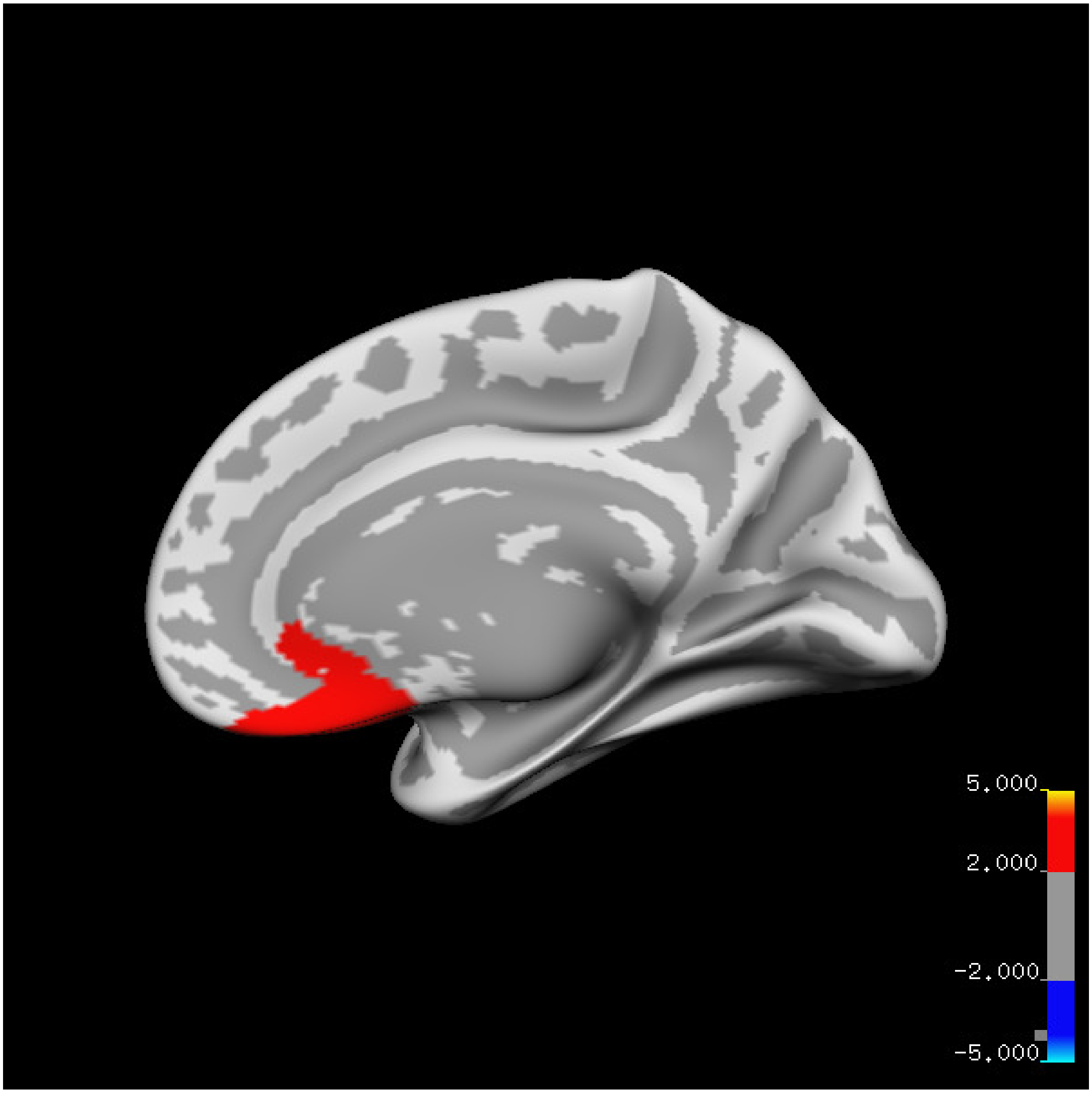
5.000
2.000
-2.000
-5.000

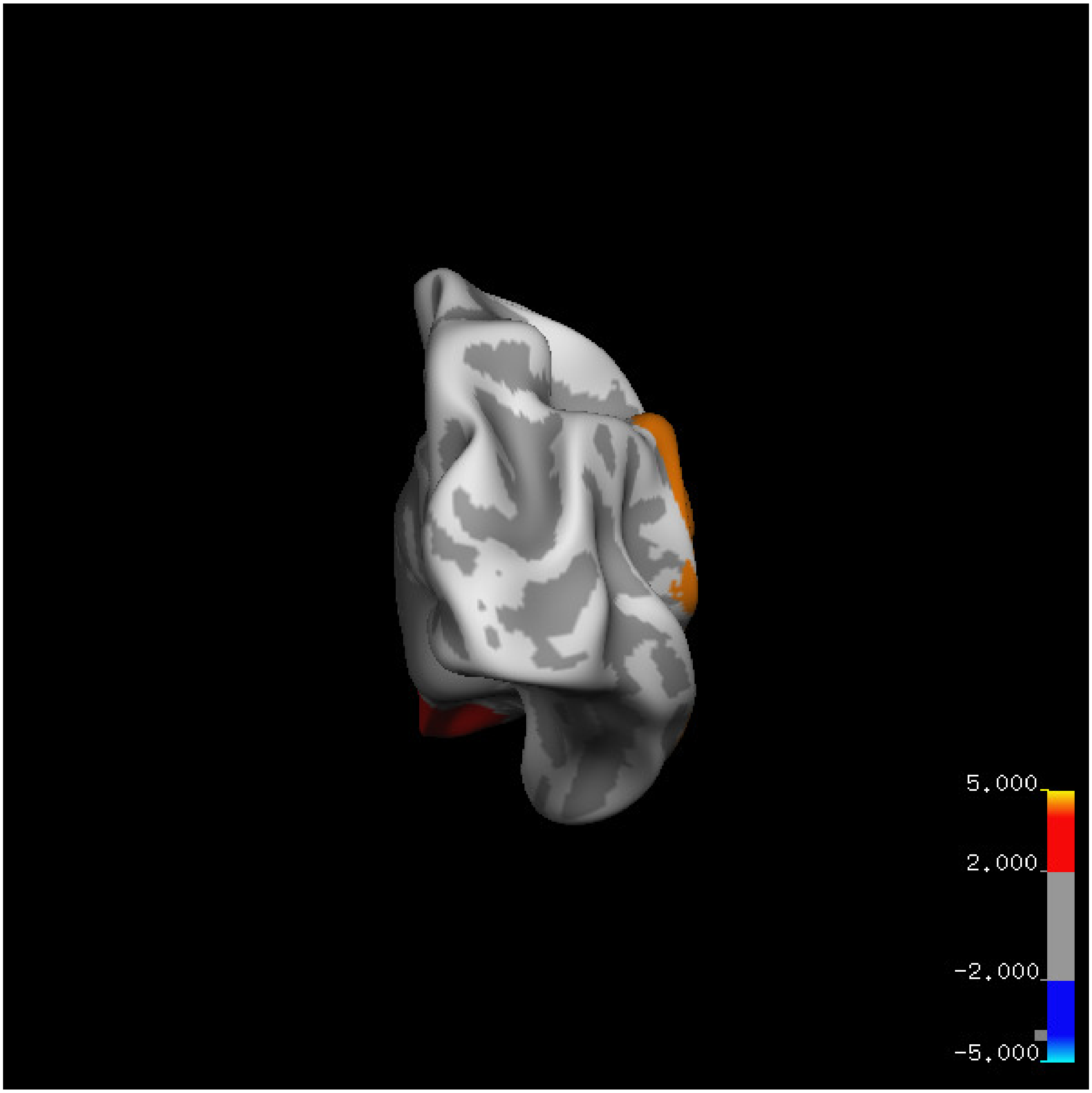
5.000
2.000
-2.000
-5.000

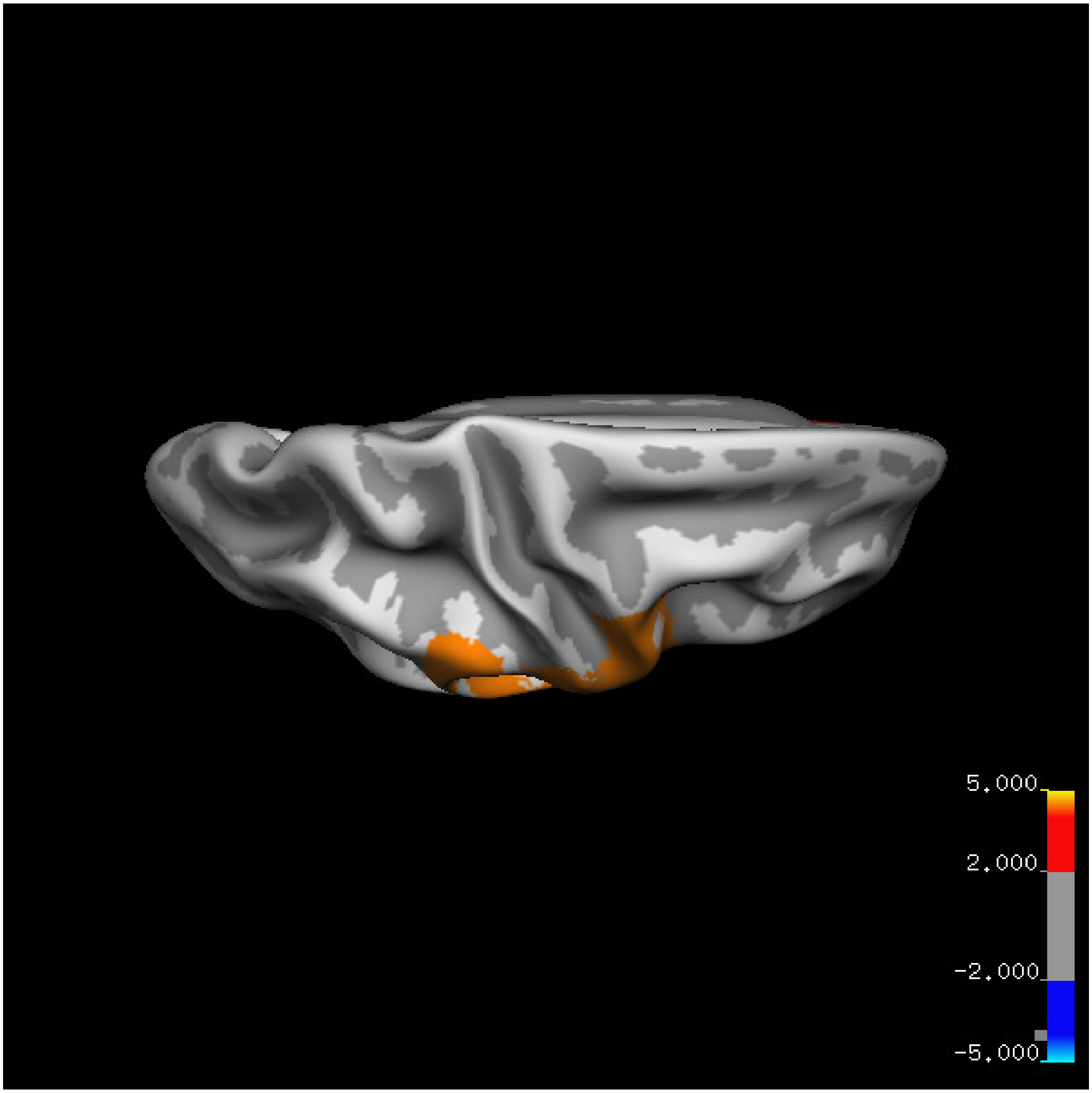


Cluster summary is shown below.

```
## # Cluster Growing Summary (mri_surfcluster)

## # $Id: mri_surfcluster.c,v 1.57.2.3 2016/11/17 18:19:42 zkaufman Exp $

## # $Id: mrisurf.c,v 1.781.2.6 2016/12/27 16:47:14 zkaufman Exp $

## # CreationTime 2021/10/04-13:56:02-GMT

## # cmdline mri_surfcluster.bin --in surface_analyses/results/cluster_comparison/fsaverage6.sm15.rh/vol

## # cwd /gpfs/projects02/p274/cluster/projects/p003-sleep_b

## # sysname  Linux
```

```
## # hostname c1-20
## # machine  x86_64
## # FixVertexAreaFlag 1
## # FixSurfClusterArea 1
## #
## # Input      surface_analyses/results/cluster_comparison/fsaverage6.sm15.rh/volume.above60.below_mea
## # Frame Number      0
## # srcsubj fsaverage6
## # hemi rh
## # surface white
## # group_avg_surface_area 85131.2
## # group_avg_vtxarea_loaded 1
## # annot aparc
## # SUBJECTS_DIR /cluster/projects/p274/tools/mri/freesurfer/current/subjects
## # SearchSpace_mm2 79329.9
## # SearchSpace_vtx 37471
## # Bonferroni 0
## # Minimum Threshold 1.3
## # Maximum Threshold infinity
## # Threshold Sign    abs
## # AdjustThreshWhenOneTail 1
## # CW PValue Threshold: 0.01
## # Area Threshold    0 mm^2
## # CSD thresh  1.300000
## # CSD nreps    10000
## # CSD simtype  null-z
## # CSD contrast NA
```

```
## # CSD confint  90.000000
## # Overall max 6.3661 at vertex 12693
## # Overall min -1.12699 at vertex 2528
## # NClusters          2
## # FixMNI = 0
## #
## # ClusterNo  Max   VtxMax   Size(mm^2)  MNIX   MNIY   MNIZ    CWP    CWPLow    CWPHi   NVtxs    WghtV
##    1        5.608   20164   6356.48     57.3    2.0   11.9  0.00010  0.00000  0.00020   3347     6508
##    2        6.366   12693   3007.20      6.7   26.3  -24.9  0.00090  0.00050  0.00130   1372     3693
```
